# Multi-omics analysis of aspirin treatment response in mice provide molecular insights and targets linked to liver fibrosis regression

**DOI:** 10.1101/2020.07.03.186015

**Authors:** Adil Bhat, Sudrishti Chaudhary, Gaurav Yadav, Anupama prasar, Chhagan Bihari, Jaswinder Singh Maras, Shiv K Sarin

**Author notes:** ***Correspondence:*** Dr. Jaswinder Singh Maras, Assistant Professor, Department of Molecular and Cellular Medicine, Institute of Liver and Biliary Sciences, New Delhi-110070, India, ext-24215 & Dr. Shiv K Sarin, MD, DM, D.Sc (Hony.). Senior Professor, Department of Hepatology, Institute of Liver and Biliary Sciences, New Delhi-110070, India. ***Authors’ contribution:*** AB and JSM conceptualised the work. AB and JSM were responsible for sample processing and experimental work and were helped by SC, AP, and GY, CB. Data analysis was performed by AB and JSM. The manuscript was drafted by AB and JSM. Professor Sarin proof read the manuscript and provided critical review for the manuscript. This manuscript has been seen approved by all authors. ***Abbreviations*** HSC, hepatic stellate cells; TGF-b1, transforming growth factor-b1; PDGF, platelet derived growth factor; PDGFR, platelet derived growth factor receptor; α-SMA, α-smooth muscle actin; ALOX5, arachidonate-5-lipoxygenase; ARG1, arginase-1; KMO, kynurenine-3-monooxygenase; RYR2, ryanodine-receptor-2; Col1α1, collagen type1 alpha 1; ALT, alanine transaminase; AST, aspartate aminotransferase; RT-PCR, real time-polymerase chain reaction.

## Abstract

**Background & Aims:** Aspirin has potent anti-platelet activities and possibly helps regression of fibrosis. We investigated antifibrotic mechanisms of aspirin in the murine CCl_4_ model and in patients with hepatic fibrosis.

**Methods:** Multiomics analysis identified networks and molecular targets regulated by aspirin which were validated in murine model and in patients with liver fibrosis.

**Results:** Biochemical/histopathological changes and hepatic fibrosis were greater in CCl_4_-treated mice compared to CCl_4_-aspirin (CCl_4_+ASA) or control mice (p<0.05). In CCl_4_+ASA mice, integrated proteome-metabolome analysis showed an increase in autophagy, drug metabolism, glutathione and energy metabolism (p<0.05) and decrease in inflammatory pathways, arachidonic acid and butanoate metabolism (p<0.05). Global cross-correlation analysis linked fibrosis markers with protein-metabolite pathways (r2>0.5, p<0.05). Liver proteome enrichment for immune clusters using blood transcription module correlated with histidine and tryptophan metabolism (r2>0.5, p<0.05). Aspirin decreased Ryanodine-receptor-2 (RYR2;oxidative-stress), Arginase-1 (ARG-1;urea cycle), Arachidonate-5-lipoxygenase (ALOX5;leukotriene metabolism), and Kynurenine-3-monooxygenase (KMO;tryptophan metabolism; p<0.05) which correlated with reduction in α-SMA, PDGFR-β and degree of hepatic fibrosis (r2>0.75; p<0.05) in animal and human studies, and, in-vitro analysis. Aspirin modulated intracellular-calcium and oxidative-stress levels by reducing RYR2 expression in activated LX-2 cells. It modulated the liver microbiome and its functions which also correlated with ARG1, ALOX5, RYR2 expression (r2>0.5, p<0.05). Metaproteome analysis showed significant microbiome similarity at phylum level in murine liver tissues and fecal samples. Aspirin increased the abundance of Firmicutes (Ruminococcaceae, Lachnospiraceae, and Clostridiaceae) and their functionality, as assessed by glycerol-3-phosphate dehydrogenase (NAD(P)(+) and dTMP-kinase activity (p<0.05).

**Conclusions:** Aspirin demonstrates broad beneficial effects following oxidative injury, inflammation, and hepatic fibrosis. Aspirin induces distinctive hepatic proteome/metabolome and intrahepatic microbiome changes which are indicative of fibrosis regression and could be further explored as therapeutic targets.

## Introduction

Liver cirrhosis is a life threatening consequence of liver fibrosis (1). Fibrosis is characterized by excessive deposition of extracellular matrix with decreased parenchymal cells and inflammation (2). Development of liver fibrosis involves persistent hepatic inflammation mediated differentiation of quiescent hepatic stellate cells (HSCs) into myofibroblast like cells. These activated HSCs then express many extracellular matrix (ECM) proteins such as collagen type-I, α-smooth muscle actin (α-SMA), transforming growth factor-β (TGF-β1) which mediate fibrosis progression (3). Further, a significant increase in liver fibrosis was seen in mice with higher TGF-β1 levels in their platelets. In addition, platelet derived growth factors mediated activation of hepatic stellate cells and fibrosis, could be amiloriated with aspirin (4).

Aspirin is known to prevent thrombosis and reduced risk and severity of liver disease (5). Activated hepatic stellate cells over express proinflammatory COX-2 enzyme which is reduced by aspirin. In animal model of chronic liver diseases (6) and in large-scale human cohort studies befinit of aspirin as antifibrotic molecule is reported (7). Previous studies have linked effect of aspirin on liver fibrosis to its anti-platelet activity and showed significane of aspirin over other NSAIDs (8). Regular aspirin administration is associated with the improvement of NAFLD features and reduced risk for liver fibrosis progression (9).

Aspirin is shown to enhance autophagy (10). As a calorie restriction mimetic, it is shown to improve the life span by modulating the clock genes, protein acetylation and liver metabolism (11). Regular aspirin use reduces platelet activation, thrombosis and liver fibrosis progression (12). Previous literature shows that aspirin exerts its effects majorly through inhibition of cyclooxygenases (COX), thereby blocking prostaglandin E2 (PGE2) synthesis. However, other observational studies suggest that aspirin may also have effects on non-COX-mediated pathways, such as TNF-α, NF-κB, and PI3K signaling, autophagy, energy metabolism, modulation of antioxidant enzymes and others (13).

Overall understanding of the anti-fibrotic mechanisms of aspirin in the liver is far from clear (14). In liver cirrhosis bacterial over growth and dysbiosis increases significantly. Further gut microbiome also contributes for the synthesis of inflammatory mediators such as prostaglandin (15). Aspirin inhibits PEG2 production and modulates gut microbiome and intestinal permeability (15). This suggests that use of aspirin is beneficial and studies associated to delineate the anti-fibrotic mechanisms of aspirin within the liver are warranted.

To address these issues, the antifibrotic effect of aspirin was studied in a CCl_4_ murine model. This was complemented with global proteome/metaproteome/metabolome analysis of the liver followed by an in-depth analysis of the regulatory networks. The integrated analysis identified promising antifibrotic targets regulated by aspirin and its correlation with immune clusters. Targets identified in our analysis were validated in murine model as well as in patients with liver fibrosis. We also analysed the association of fecal/liver microbiome with hepatic inflammation and fibrosis. Finally we state that using multi-omics approach we were able to identify significant alterations linked to aspirin in mice model of CCl_4_ induce liver fibrosis. This analysis focused on the liver per se, which is a primary responsive organ and likely mediates many of the beneficial effects of aspirin. Our analysis allowed us to dissect the unique differences in response at the metabolite, protein and metaprotein levels between aspirin treated animals and the liver fibrosis induced by repeated doses of CCl_4_-treated animals. Together, these data highlight the significant regulation of the KMO; kynurenine-3-monooxygenase (tryptophan metabolism), ARG-1; arginase-1 (glutathione/energy metabolism) and ALOX-5; arachidonate-5-lipoxygenase, and RyR2; ryanodine-receptor-2 (inflammation) in response to the aspirin intervention, suggesting modulation of these pathways could be useful in therapeutic interventions for regression of liver fibrosis.

## Materials and Methods

### Animals and Sample Collection

Male C57BL/6J mice of 6-8 weeks, weighting 20-25 g were divided into four groups: aspirin (ASA), carbon tetrachloride (CCl_4_), CCl_4_+ASA and control group (n=8 animals/group). After grouping, mice were subjected to twice a week intraperitoneal (i.p.) injections of 10% CCl_4_ (Sigma, 270652) (diluted in olive oil) at a dose of 0.5 µl/g body weight for 12 weeks to induce liver fibrosis (16) or vehicle (olive oil). while the ASA group were treated with aspirin at the same time concominant to CCl_4_ treatment. Low dosage of aspirin treatment were used as described in previous studies (17) whereas dose of CCl_4_ were routinely adjusted based on weight of the animal. Harvested tissues/serum was snap-frozen in liquid nitrogen (N_2_) and kept at −80°C until batch analysis. Animal grouping, model development and statement for ethics are detailed in Supplementary-methods.

### Liver multi-omics analysis

Liver samples from different groups were subjected to proteomics, metabolomics and metaproteomics analysis as detailed in Supplementary-methods.

### Global cross-correlation, clustering and integration analysis

Differentially expressed proteins (DEPs) and metabolites (DEMs) were identified and cross correlated to biochemical/fibrosis indicators and was clustered. Pathway analysis was performed for proteins (enricher/KEGG) and metabolites (metaboanalyst) significantly correlating to markers of fibrosis cluster or biochemical clusters. This was followed by developing a global cross-correlation map between biochemical/fibrosis parameters and pathways linked to the proteins and metabolites using Cytoscape (https://cytoscape.org/) (18). In addition, protein– metabolite joint pathway analysis was performed for the DEPs and DEMs of CCl_4_+ASA group compared to the CCl_4_ group using Metscape-3 (19) which was followed by pathway over representation and visualization of the most altered pathways (20).

### Histopathology and Immunohistochemistry of Liver Sections

Liver tissues were fixed in 10% paraformaldehyde, embedded in paraffin and sectioned at 4 μm thickness which was then stained with haematoxylin-eosin (H&E), Masson’s trichrome (MT) and sirius red staining (21). The extent of hepatic injury and fibrosis were measured through the Ishak index score (22). In addition, expression for α-SMA (Cat.E-AB-34268), PDGFR-β (Cat.E-AB-32531), ARG1 (Cat. M01106-1), RYR2 (Cat. E-AB-64002) and ALOX-5 (Cat. E-AB-10040) was estimated in the membrane and cytoplasmic space of the positively stained cells, counted in consecutive 10 high power fields (40x) and relative quantitation as mean number of cells/10 high power field (40x) was recorded.

### Statistical Analyses

Results are shown as mean and standard deviation unless indicated otherwise. Using Graph Pad Prism v6, statistical analyses were performed and P-values of < 0.05 were considered statistically significant. To compare variables between the two groups, the unpaired (two-tail) Student’s t test, Mann-Whitney U test were performed. For comparison among more than two groups, one-way analysis of variance, Kruskal-Wallis test was performed. All correlations were performed using Spearman correlation analysis (Detailed in Supplementary-methods).

## RESULTS

### Aspirin treatment prevented hepatic injury and reduces fibrosis in CCl_4_ murine model of liver fibrosis

Schema for the development of the CCl_4_ murine model of fibrosis and treatment with aspirin is shown in Figure-1A (Supplementary-Figure-1). The murine model receiving CCl_4_ showed significant increase in liver weight: total body weight and hepatic collagen deposition as compared to vehicle or aspirin (5mg/kg) treated mice (Figure-1B, 1C, p<0.01). At 12-week, ALT and total bilirubin levels were significantly high whereas albumin levels were low in the CCl_4_ model compared to other groups (Figure-1D, p<0.05). Liver fibrosis and liver functions were significantly improved in the aspirin treated group (Supplementary-Table-1). This was complemented with a significantly decreased expression of collagen-IαI, alpha smooth muscle actin (α-SMA) and transforming growth factor β-1 (TGF-β1) in the aspirin treated group (Figure-1E, Supplementary-Table-2). These results suggest that aspirin treatment can reduce liver fibrosis level in the murine model, probably by modulating oxidative stress and inflammation.

**Figure-1:**
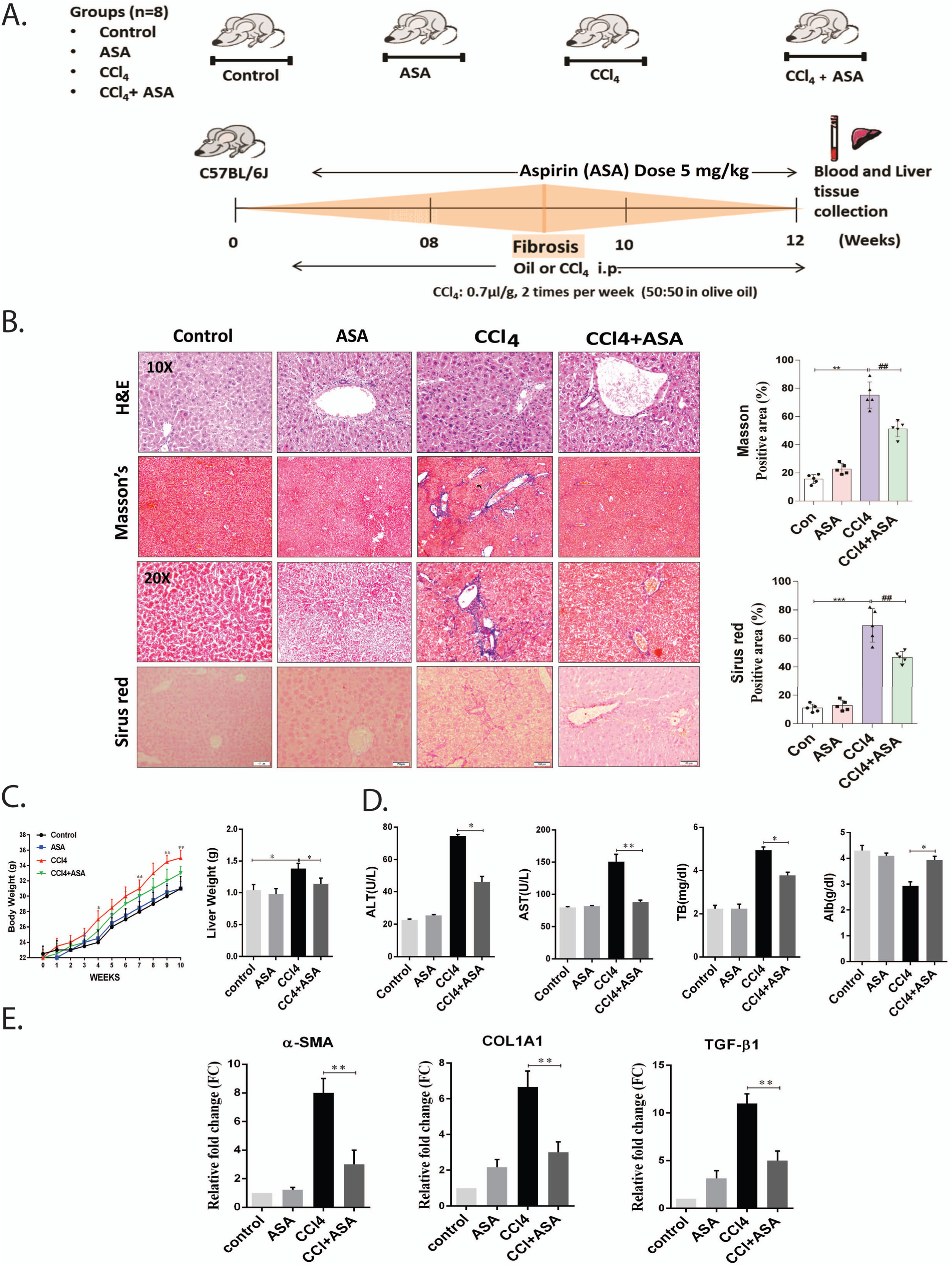
Aspirin administration significantly attenuates hepatic damage and fibrosis in CCl_4_ induced chronic liver injury. A: Schematic flowchart: Mice were given repeated injections of CCl_4_ for 12 weeks to develop liver fibrosis in presence and absence of aspirin (ASA), administered in drinking water. B: Representative histostaining images and quantification showing liver injury as assessed by haematoxylin and eosin (H&E) and fibrosis by using Masson’s trichrome, Sirius red (collagen staining dyes) in healthy control, ASA, CCl_4_ and CCl_4_+ASA mice groups. C: Effect of aspirin on the liver weights of C57BL/6 across mice models during the progression stages and liver/body weight ratio at 12 weeks period. D: Serum amino transferases, total bilirubin and albumin levels are plotted and their levels post aspirin treatment in CCl_4_ model. E: Relative fold expression of α-SMA, COL1A1 and TGF-β1 (liver fibrosis markers) in murine model and other groups. Abbreviations: Con; Control, ASA; Aspirin alone,CCl_4_; carbon tetra chloride group,CCl_4_+ASA; carbon tetra chloride plus aspirin, TB; total bilirubin, %; Percentage. The values are expressed as means ± SEM. n=05 animals per group (*p < 0.05, **p <0.005).

### Aspirin administration decreaed inflammation and liver fibrosis associated proteome

Global proteome analysis can provide an unbiased view of changes in protein/enzyme abundance, which are the key mediators in metabolic /endocrine signaling that may be altered during fibrosis regression. To understand the mechanism linked to aspirin mediated liver fibrosis reduction, total liver tissue proteome was first analysed in murine model of fibrosis with and without aspirin treatment (CCl_4_+ASA versus CCl_4_) (Figure-2A). We also performed unbiased proteome analysis across the four animal groups detailed in Supplementary Figure-2. We identified and quantitated >4,200 proteins across all treatment groups (Supplementary-Table-3). Of the 471 proteins p < 0.05 (5% false discovery rate [FDR]) a total of 161 proteins were upregulated and 310 proteins were downregulated in CCl4+ASA as compared to CCl4 (Figure 2B). Partial least square discriminant analysis (PLS-DA; Figure-2C) and unsupervised clustering analysis showed clear distinction of CCl4+ASA from CCl4 (Figure-2C and 2D). Pathway analysis of the up-regulated proteins specific to the CCl_4_+ASA group were linked to autophagy, drug metabolism, fatty acid degradation, PPAR signalling, histidine metabolism, amino acid metabolism and glutathione metabolism (Figure 2E, Supplementary-Figure-3A). Whereas, pathways linked to downregulated proteins were associated to cholesterol metabolism, EGFR1 signalling, arachidonic acid, ferroptosis, phagosome, and inflammatory signaling pathway (TNF-alpha NFKB signaling pathway; Supplementary-Figure-3B). Finally, to identify fibrosis linked proteins, the identified DEP’s were searched against the OMIM database for liver fibrosis. A total of 33 proteins were identified; of them 21 were up-and 12 were downregulated in CCl_4_ induced liver fibrosis model. Aspirin treatment significantly reduced the expression of acyl-CoA oxidase-2 (Acox2), collagen Type XXV Alpha-1-chain (Col25a1), ryanodine receptor-2 (Ryr2), G-protein subunit beta-1 (Gnb1), protein phosphatase-2 (Ppa2) and others in CCl_4_ model of liver fibrosis (Figure-2F) Together these findings suggest that aspirin reduces fatty acid accumulation, oxidative stress, and inflammation (Supplementary-Figure-4). Further proteins such as Acox2, Col25a1, Ryr2, Gnb1, Ppa2 and others are regulated by aspirin and could be validated for a probable indicator of liver fibrosis (Supplementary-Table-4).

**Figure-2:**
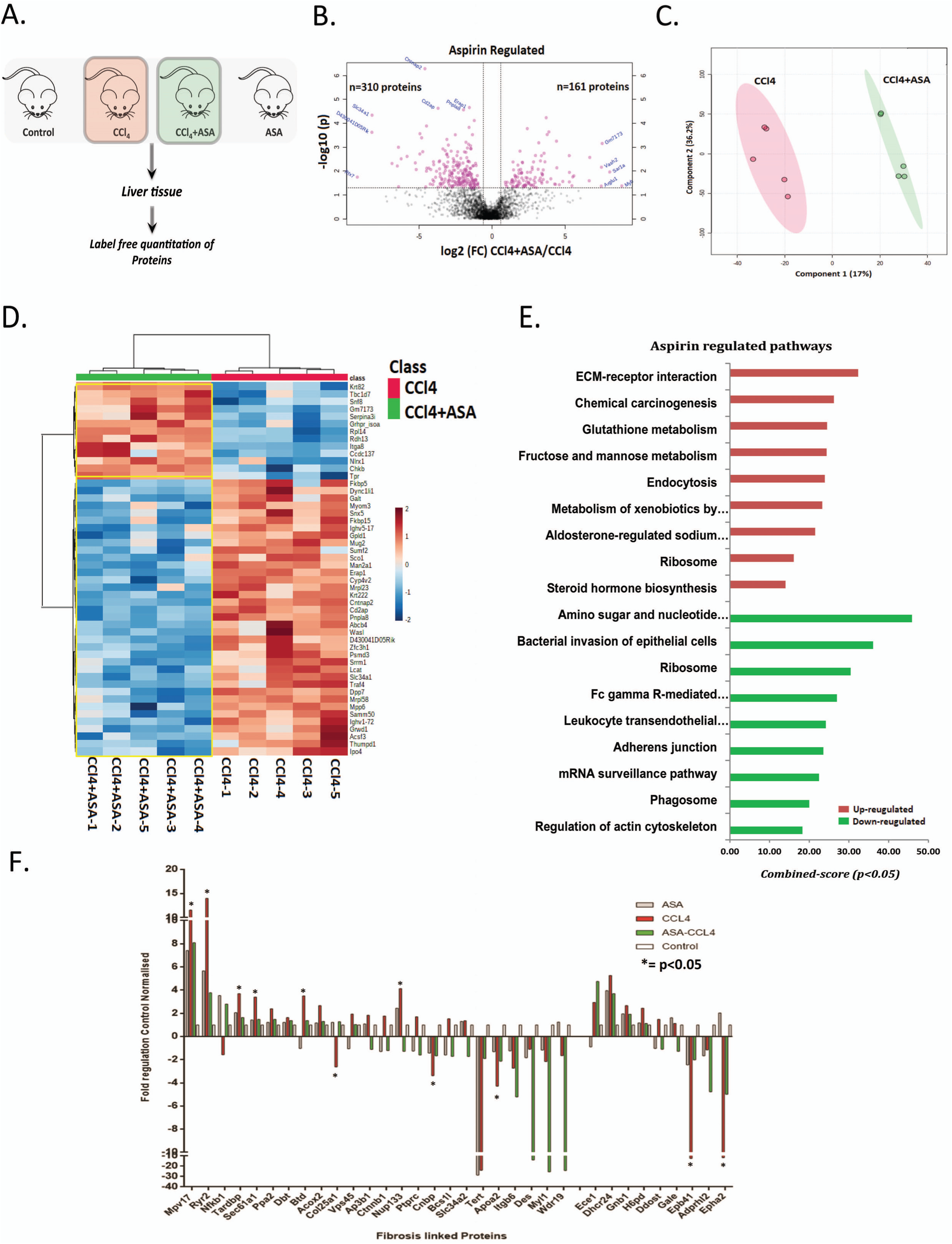
Proteomics analysis of the murine model of liver fibrosis at 12 weeks and comparison to aspirin treatment. A: Workflow for mass spectrometry-based label free proteomic study performed across the mice tissue samples. B: Differentially expressed proteins were visualized by volcano plot and filtered so as to include only statistically significantly changed proteins (P < 0.05, t-test) that were changed by more than 1.5-fold in mice livers. C: PCA plot analysis between the mice groups. D: Heatmap dataset analyzed by 2-way ANOVA, representing the hierarchical cluster of differentially abundant proteins among liver samples from mice groups classified as Con, CCl_4_, CCl_4_+ASA and ASA. E: Bar chart diagram showing pathway associated with up regulated proteins in different mice groups. Red colour = upregulated pathways and Green colour= downregulated pathways. F: Relative fold expression of various proteins linked to liver fibrosis in hepatic tissue. Expression changes are shown in control, CCl_4_, CCl_4_+ASA and ASA mice groups respectively. Interestingly proteins such as (DES, Myl1, Wdr19) were down regulated but were not considered as they lacked to achieve significance. The proteomic analysis was performed using of n=5 animals per group. Abbreviations: Con; Control, ASA; Aspirin alone, CCl4; carbon tetra chloride group, CCl_4_+ASA; carbon tetra chloride plus aspirin. The values are expressed as means ± SEM. n=05 per group (*p < 0.05, **p < 0.005).

### Aspirin treatment enhances metabolic function in murine model of liver fibrosis

Aspirin could modulate liver metabolome, recently, a distinctive plasma metabolomic profile associated with liver fibrosis is also reported (23). Thus reduction of fibrosis post-aspirin treatment could be attributed to a change in the fibrotic liver metabotype. Differentially expressed metabolites (DEMs) were first individually analysed across all the four mice groups detailed in Supplemantary Figure-5 and Supplementary Table-5. We next evaluated the liver metabolome in the CCl_4_ model of fibrosis with and without aspirin treatment (Figure-3A). Of the 164 metabolite differentially expressed a total of 104 metabolite were upregulated and 60 were down regulated in CCl4+ASA group compared to CCl4 (p<0.05; Figure 3B). Partial least square discriminant analysis (PLS-DA) and unsupervised clustering analysis showed a clear distinction between the groups (Figure-3C, 3D). Metabolites upregulated in CCl4+ASA were associated to glutamate metabolism, TCA cycle and pentose phosphate pathway, arginine biosynthesis and cysteine and methionine metabolism (Figure 3E). Metabolites down regulated were linked to arachadonic acid metabolism, lysine degration, tryptophan metabolism and others (Figure 3F). These findings suggest that aspirin induces energy metabolism, amino acid metabolism and reduces inflammatory pahways such as arachadonic acid metabolism and tryptophan metabolism.

**Figure-3:**
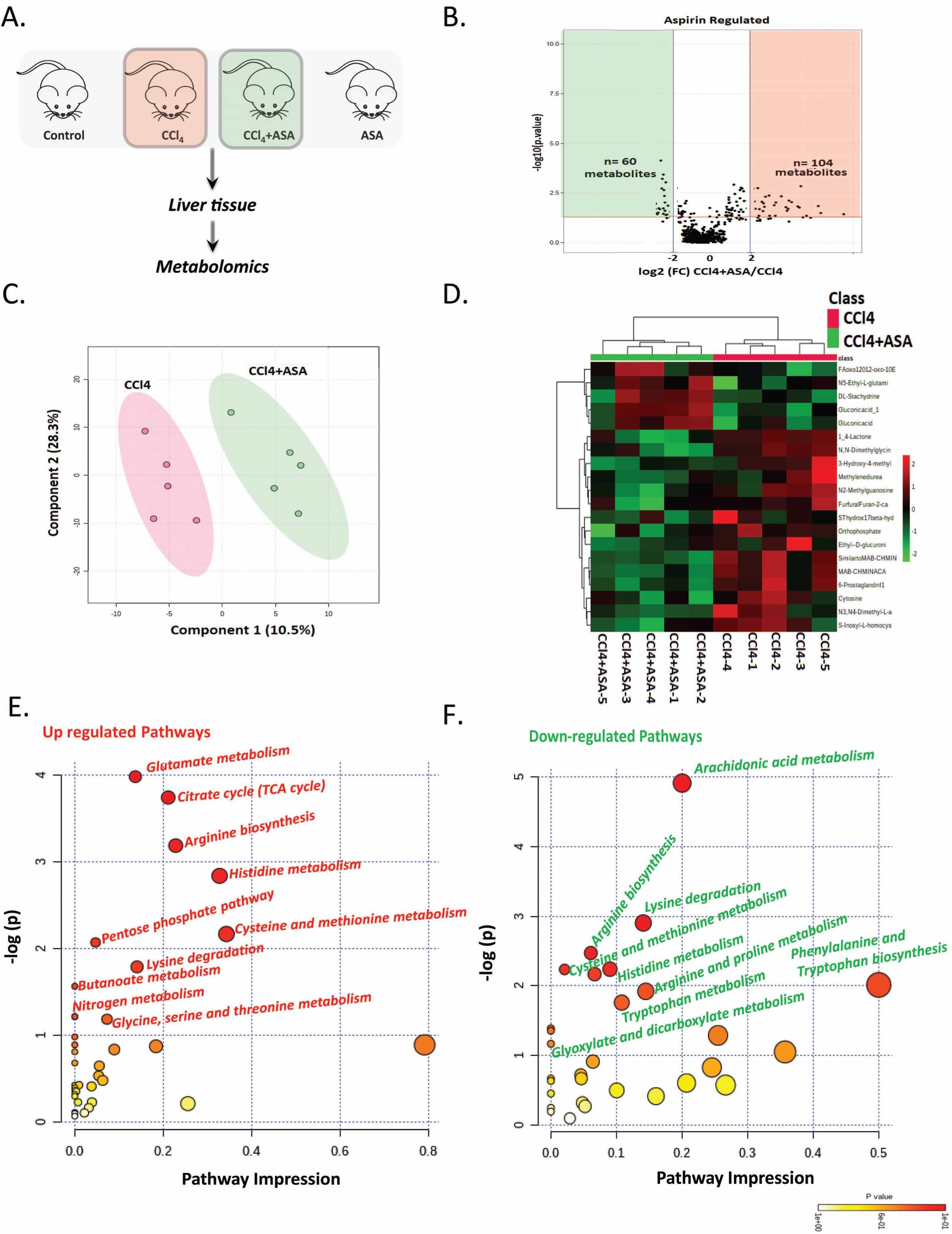
Effect of aspirin treatment on liver metabolome in murine model of liver fibrosis. A: Overview of metabolomic study performed across the mice groups. B: Three dimensional PLS-DA score plots of the aspirin treatment CCl_4_+ASA, and CCl_4_. Each dot represents one liver sample, projected onto first (horizontal axis) and second (vertical axis) PLS-DA variables. Mice groups are shown in different colours (left panel). Three dimensional loading plots of ASA, CCl_4_, CCl_4_+ASA, and con combined analysis both in the negative ion mode (ESI–) and positive mode (ESI+). C: Differentially expressed metabolites were visualized by volcano plot and filtered so as to include only statistically significantly changed proteins (P < 0.05, t-test) that were changed by more than 1.5-fold in mice livers. D: Heatmaps of the differential metabolites in aspirin treated CCl_4_+ASA compared to CCl_4_ mice in the positive ion mode (ESI+). Red and green colours indicate values above and below the mean, respectively. Black indicates are values close to the mean. E: The pathways associated with the upregulated metabolites are represented in bubble plot (log2-fold change >1.5; p-<0.05). F: The pathways associated with the significantly downregulated metabolites are represented in bubble plot, (log2-fold change <1.5; p-<0.05). The analysis was performed using of n=5 animals per group. Abbreviations: CCl4; carbon tetra chloride group, CCl4+ASA; carbon tetra chloride plus aspirin.

**Figure-4:**
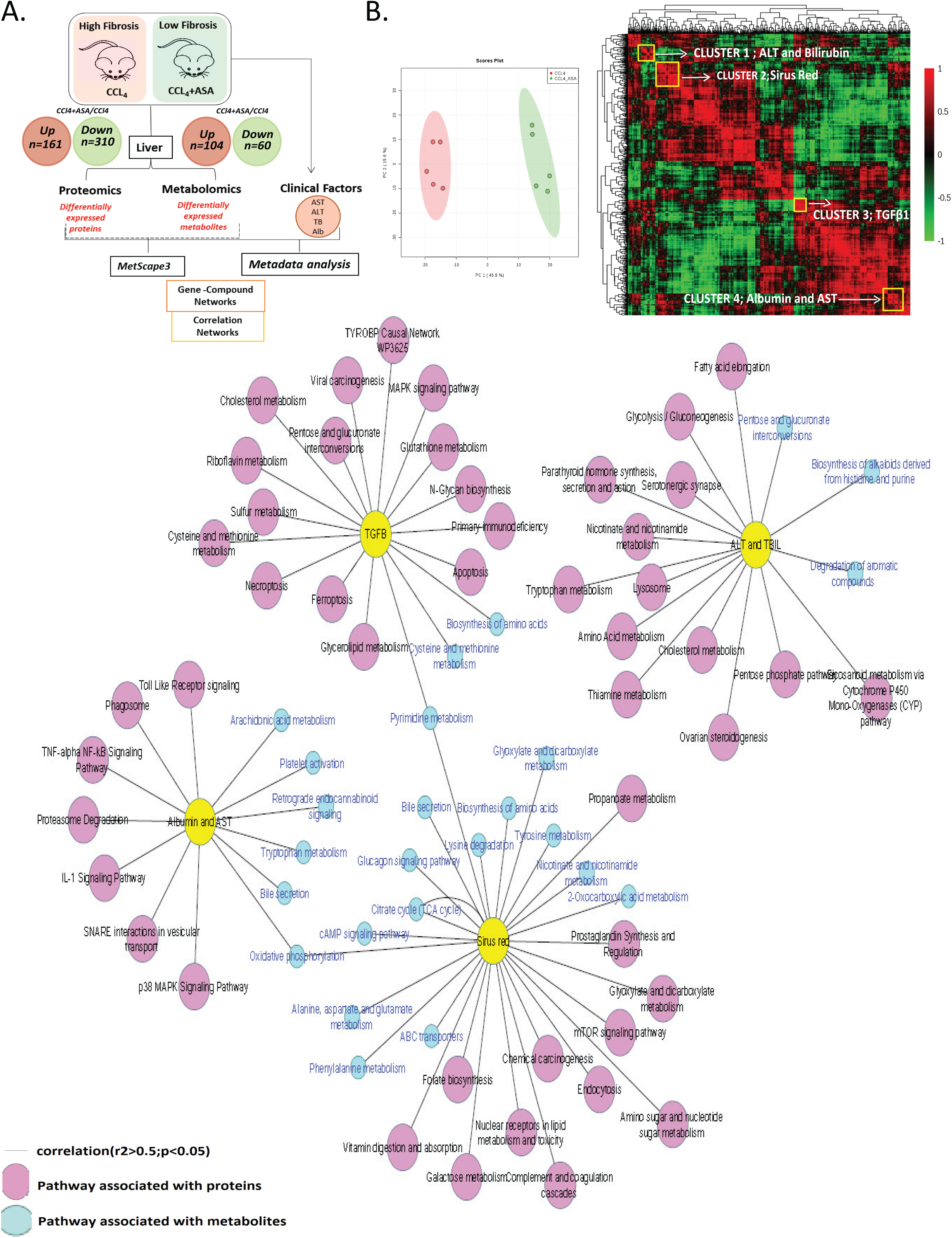
Global cross correlation and hierarchical cluster analysis of hepatic proteome, metabolome with biochemical/fibrosis parameters. A: Schematic representation of the integration analysis performed in this study. Protein-protein/metabolite interaction network was visualized by Cytoscape. B: PCA showing clear distinction between two groups. Global correlation map of the hepatic proteome, metabolome and clinical variables in mice groups. Four clusters were formed containing clinical factors. Correlation analysis with clinical factors and associated proteomic and metabolomics pathways. Pink color indicates the pathway associated with proteins while blue color indicates the pathway associated with metabolites. Yellow color is indicative of clusters containing clinical parameters or liver fibrosis marker.

**Figure-5:**
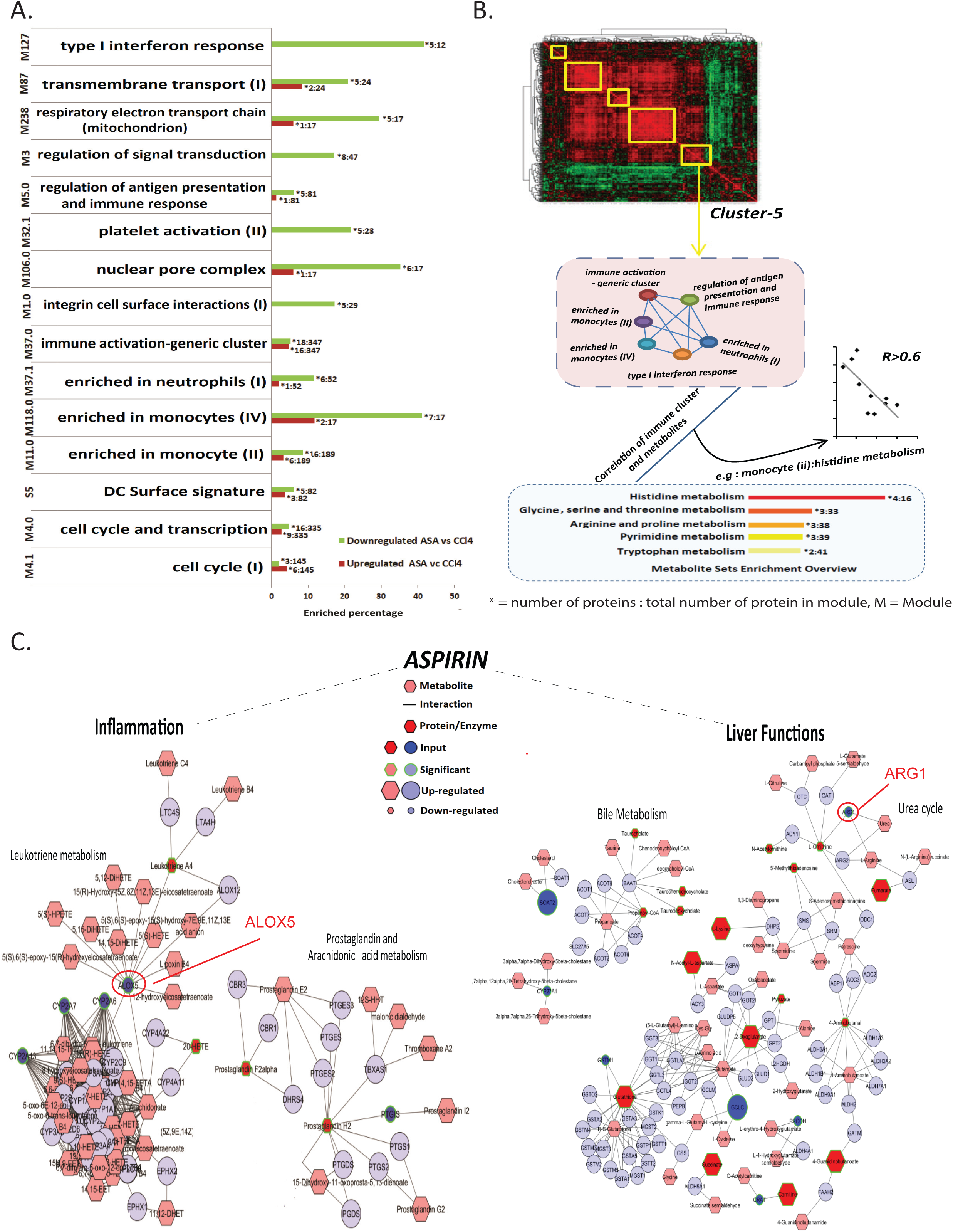
Aspirin decreases inflammation and improves metabolic liver function. A: BTM enrichment analysis of hepatic proteins: Immune clusters are represented as bar diagram showing the number of proteins present in each module, red and green colour bar chart showing the %age of proteins present between the CCl_4_ vs. CCl_4_+ASA mice groups. Also correlation map of immune cluster-5 and its metabolic function is given. B: Whole interactome of metabolites (hexagons), interactions (dashed lines) and proteins/enzymes (circles) associated with aspirin regulated pathways and target molecules in chronic liver disease. Most differentiating fibrosis targets with significant AUC>0.75 under the regulation of aspirin werethen selected for validation.The analysis was performed using of n=5 animals per group. Abbreviations: %age; Percentage, Con; Control, ASA; Aspirin alone, CCl_4_; carbon tetra chloride group, CCl_4_+ASA; carbon tetra chloride plus aspirin.

### Global correlation analysis of hepatic proteome, metabolome and biochemical/fibrosis parameters reveals anti-oxidative state post-aspirin treatment

We hypothesized that the changes in the biochemical/fibrosis parameters should reflect an analogous change in the proteome and metabolome. In particular, this should hold true for liver proteome, metabolome, and liver function/fibrosis parameters. For example, a decrease in fibrosis or biochemical parameters should recapitulate a decrease in proteins or metabolites associated pathways and members of known pathways should cocluster. Integration of the proteome and metabolome could help to impute the class membership of metabolites, proteins, familial enzyme pathways or novel enzyme reaction models associated to change in biochemical or fibrosis indicators. To explore this, global cross-correlation and hierarchal clustering analysis were performed for differentially expressed proteins (798) and metabolites (138) in CCl_4_+ASA (low fibrosis) group as compared to CCl_4_ (high fibrosis) group along with biochemical parameters like ALT, AST, total bilirubin, Albumin, TGF-β1, MT/sirius red score (Figure-4A). Principle component analysis recapitulates the distinction between high (CCl_4_) and low fibrosis (CCl_4_+ASA) groups. Correlation analysis followed by hierarchical clustering led to a global correlation map (r^2^>0.5), where the biochemical parameters/fibrosis markers were clustered with proteins and metabolites. This analysis identified 4 clusters (Figure-4B). Pathway analysis of the proteins and metabolites associated with each cluster showed that alteration in ALT, total bilirubin:cluster-1 was associated with alteration in fatty acid metabolism, tryptophan metabolism, pentose gluconate inter-conversions, cholesterol metabolism and more (r^2^>0.5, p<0.05). Change in fibrosis marker (Sirius red): cluster-2 correlated with mTOR signaling, complement and coagulation, prostaglandin synthesis, cAMP signaling, TCA cycle, and others (r^2^>0.5, p<0.05). Further change in TGF-β1: cluster-3 correlated with MAPK signaling, glutathione metabolism, apoptosis, and others (r^2^>0.5, p<0.05). Similarly a change in albumin and AST levels: cluster-4 was linked to change in TNF-NFκB signalling pathway, IL-1 signaling, p38 MAPK signaling, oxidative phosphorylation, arachidonic acid metabolism and platelet activation (r^2^>0.5, p<0.05; Figure-4B). Over all, these results establish a linear and direct relationship between biochemical/fibrosis markers, proteome, and metabolome of the liver.

### Aspirin treatment reduced inflammation and enhances liver function in murine model of liver fibrosis

Interestingly, our results show that ALT is reduced by aspirin, indicating decrease in hepatic injury. It can be argued that the resulting antifibrotic effects are indirect and not truly anti-fibrotic. To understand this we investigated the role of immune cells, by studying differentially expressed proteins (DEPs) in aspirin treated vs CCl_4_ treated animals using blood transcription module (BTM) (24). The upregulated proteins in CCl4 were enriched for platelet activation-I (>21%), monocytes (>4%), recruitment of neutrophils (>6%) and inflammation modules (>10%). These were significantly downregulated on treatment with aspirin (Figure-5A, Supplementary-Table-6). Under aspirin treatment, the metabolic function of the immune clusters (immune activation – generic cluster, enriched in monocytes (II), enriched in monocytes (IV), enriched in neutrophils, type I interferon response) were decreased and showed direct correlation with the liver metabolome particularly metabolites linked to (histidine, tryptophan metabolism). Correlation of immune clusters with metabolic pathways highlights the anti-inflamatory and anti-oxidative state induced by aspirin treatment in CCl_4_ mice.

Finally, to understand the antifibrotic effect of aspirin, The DEP’s and DEM’s in CCl_4_+ASA as compared to CCl_4_ model was subjected to integrated pathway analysis using Metscape from Cytoscape. We identified a total of 115 pathways significantly perturbed. Proteins/metabolites significantly upregulated in CCl_4_+ASA highlighted increase in 24 (20.7%) pathways such as endocytosis, thermogenesis, protein processing in ER, autophagy, drug metabolism, Rap1 signaling and others (p<0.05) whereas proteins/metabolite downregulated highlighted significant decreases in 53 (45.7%) pathways such as phagosome, MAPK signalling, TNF signaling pathway, fatty acid synthesis, platelet activation and others (p<0.05; Supplementary-Figure-6A). Interestingly, most perturbed 39 pathways (33.6%; which involves both up and down regulated proteins/metabolites) in CCl_4_+ASA group were linked to liver function (urea cycle, bile acid biosynthesis, xenobiotic metabolism), inflammation(arachidonic acid, prostaglandin and leukotriene metabolism), energy metabolism (TCA cycle, pentose phosphate) and amino acid metabolism (tryptophan, tyrosine metabolism)(Supplementary-Figure-6B). Visualisation of key pathways showed that aspirin downregulates inflammation by down regulating central metabolites and proteins of the arachidonic acid pathway such as leukotriene A4, prostaglandin F2alpha, CYP2A6, ALOX5 and others (Figure-5B, left-side). Aspirin regulated bile acid metabolism and urea cycle reflective of liver function by decreasing bile metabolites (propanoyl-CoA, taurocholeate, and others) and urea cycle proteins and metabolites like arginase-1, ornithine, N-acetyl ornithine and others (Figure-5B, right-side). Aspirin treatment to CCl_4_ model increased energy metabolism linked proteins/metabolites (Supplementary-Figure-7A). Finally, aspirin regulates tryptophan metabolism by reduction of kynurenine and kynurenine 3-monooxygenase (KMO) which is a rate limiting enzyme in the kynurenine pathway (Supplementary-Figure-7B).Together these results demonstrate that aspirin reduces inflammation and improves liver function which may have a direct effect on fibrosis regression in murine model.

### Validated in patients RyR2 and other-target proteins expression is reduced in the murine model under aspirin treatment

To identify putative candidates associated with fibrosis regression, AUROC analysis was performed for the DEPs and DEMs in CCl_4_+ASA compared to the CCl_4_ model. A partial list of the top 20 proteins and metabolites along with biochemical parameters and fibrosis indicators is provided as (Supplementary-Table-7). Based on the AUROC levels and p-value significance, we identified RYR2 linked to oxidative stress (25), ALOX-5, KMO linked to inflammation (26), ARG-1 linked to ammonia production (27) significantly associated with liver fibrosis. The expression of these proteins was validated on the representative liver sections of mice, as well as in patients with increasing grades of liver fibrosis. The liver section of the CCl_4_ model showed a significant increase in RYR2, ALOX-5, and ARG-1 expression in the sinusoidal spaces and near portal areas which was decreased when CCl_4_ murine was treated with aspirin (p□<□0.05; Figure-6A). We also observed significant correlation of these proteins with degree of fibrosis, α-SMA and PDGFR-β (r^2^>0.7, p<0.01, Figure-6B). In addition, the identified markers (RYR2, ALOX5, ARG-1 and KMO) showed linear increase in expression (Figure-6C) and correlated significantly with degree of liver fibrosis (F0-F4) in patients with liver fibrosis (p□<□0.05, Figure-6D). Collectively, these results suggest that aspirin may reduce fibrosis by decreasing oxidative stress (RYR2), inflammation (ALOX5, KMO), ammonia production (ARG1),activation of the autophagy (Deptor, VMP1) and energy pathways (IDH3B, MDH1) in liver.

**Figure-6:**
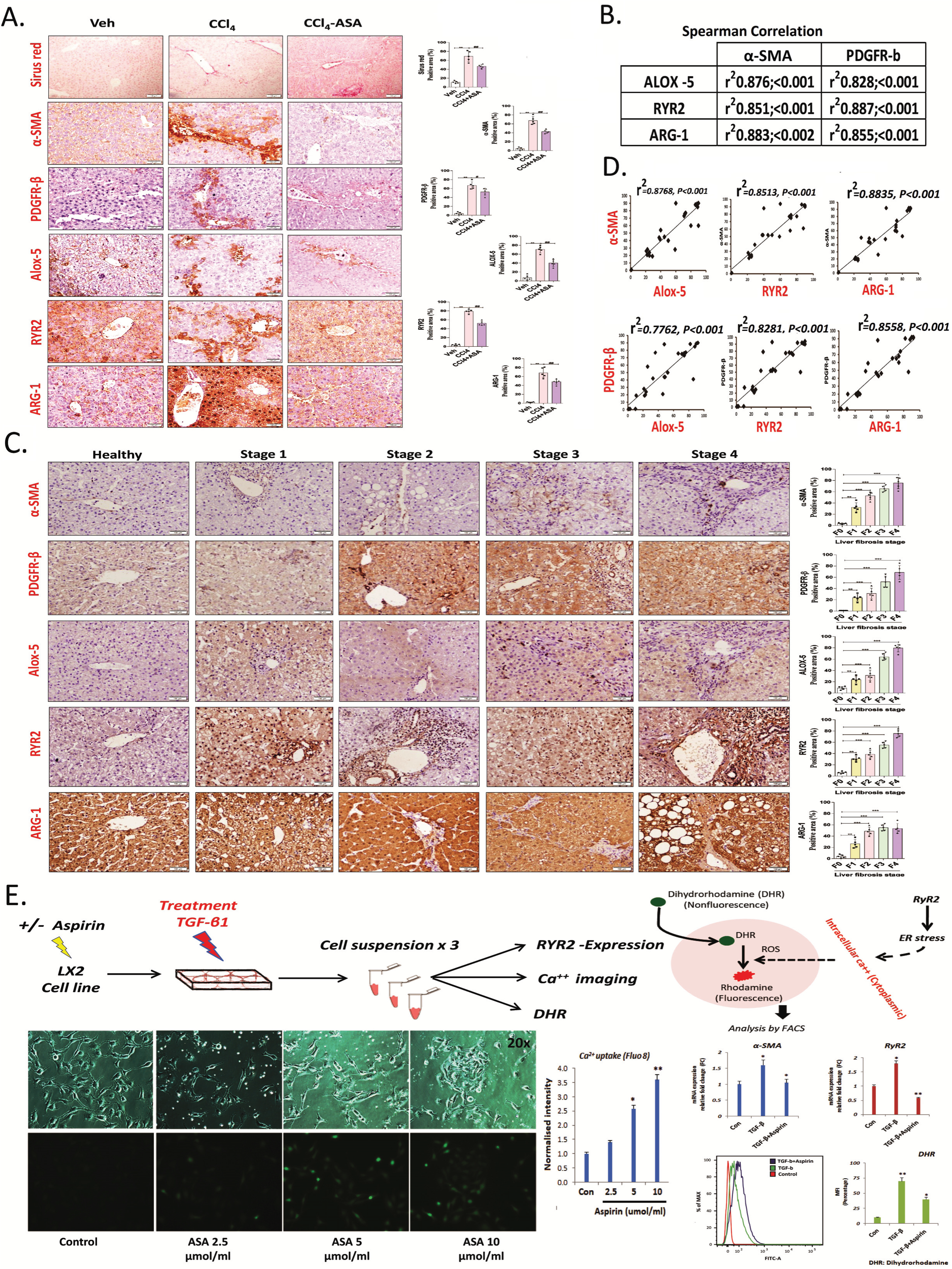
Aspirin decreases the elevated expression of ALOX-5, RYR2 and ARG-1 in liver fibrosis and inhibits oxidative stress. A: IHC staining of target proteins (ALOX-5, RYR2 and ARG-1) were compared and correlated with liver fibrosis markers (α-SMA and PDGFR-β) in model and aspirin treated hepatic tissue.Representativerelative quantification for all IHC analysis of positively stained cells are expressed as percentage of positive cells/10 high power field (40×).The results are given as mean ±SEM; n=5 ** p<0.01 vs. control; ## p<0.01 vs. model. B: IHC staining of target proteins (ALOX-5, RYR2 and ARG-1) were compared and correlated with liver fibrosis markers (α-SMA and PDGFR-β) in human liver (n=5). Degree of liver fibrosis in humans. Results are given aspercentageof positive cells/10 high power field (40×)and are expressed as mean ± SEM; n = 5/stage; *P < 0.05; **P < 0.01; ***P <0.005. C: Expression levels of ALOX-5, RYR2 and ARG-1 in murine model of liver fibrosis showed significant correlation with liver fibrosis markers (α-SMA and PDGFR-β). D: Expression of ALOX-5, RYR2 and ARG-1 with disease stage of liver fibrosis in humans showed significant correlation with liver fibrosis (α-SMA and PDGFR-β).The analysis was performed using n=5 animals per group. Abbreviations: ALOX-5;arachidonate-5-lipoxygenase, RYR2; ryanodine-receptor-2, ARG-1;arginase, α-SMA; alpha-smooth muscle actin, PDGFR-β; platelet derived growth factor receptor-beta. E. Intracellular calcium and ROS generation in TGF-β1-induced LX2 cells was detected by fluorescence microscope and flow cytometer in presence and absence of aspirin. The bar graph represents increase in Ca^++^ levels, ROS and RyR2 expression in activated LX2 and significant decreae in levels after aspirin treatment.

### Aspirin reduces RyR2 expression and inhibit oxidative stress: in-vitro analysis

To evaluate the inhibitory effects of aspirin on LX2-cells (human hepatic stellate cells;HSCs) in terms of calcium release, RyR2 expression and oxidative stress in-vitro. LX2, were treated with TGF-β1 in the presence or absence of aspirin.The results of cell immunofluorescence fluo-8 assay suggest increase in cytoplasmic calcium (Ca^++^) levels upon aspirin treatment with decrease in RyR2 expression (Fig. 4E). Next, LX2 cells were loaded with DHR-123(dihydrorhodamine) to detect the accumulation of intracellular ROS (reactive oxygen species). The addition of aspirin significantly decreased the accumulation of ROS induced by TGF-β1 by over 30%. Consistant with RyR2 mRNA expression, intracellular Ca^++^ and ROS widely distributed in the cytoplasm of TGF-β1-stimulated cells, while coincubation significantly decreased the expression of these markers when compared with the TGF-β1 group (Figure-6E). This indicates that aspirin could decrease the RyR2 expression and inhibit oxidative stress in activated LX2 cells.

### Aspirin ameliorates hepatic fibrosis via modulation of intrahepatic microbiome in murine model of liver fibrosis

Translocation of the pathogenic microorganisms and derived factors from the gut to liver is one of mechanisms linked to the pathogenesis of cirrhosis-associated liver diseases in animals and humans (28). We tested whether livers of mice affected by fibrosis exhibit higher load of pathogenic microorganisms. Our analysis showed higher load of bacterial DNA fragments in CCl_4_ group (Supplementary-Figure-8A). This was complemented with the metaproteome analysis which resulted in the identification of more than 7000 peptide sequences linked to bacterial or host proteins. Based on the peptide sequences the lowest common ancestor (LCA) was identified and visualised (29). An interesting observation was the similarity between the fecal metaproteome and liver metaproteome at the phylum level (Figure-7A). Based on this data, we undertook to study the metaproteome of the liver in CCl_4_ mouse model of cirrhosis in presence or absence of aspirin. Principle component analysis along with unsupervised clustering analysis showed clear segregation of groups based on their metaproteome profiles (Figure-7B). Aspirin treatment in CCl_4_ mice significantly increased bacteria linked to the order Aquifexaeolicus, Chlorobilaes, Tissierellales, Neisserllas, Epsilonproteo bacteria and Spirochaetales (p<0.05) whereas bacteria linked to the order Micrococalae, Prochloraceae and Synechococcaceae were significantly reduced (p<0.05;Figure-7C,D). Bacterial species such as Proteobacteria, Enterobacteriaceae, Fusobacteria and Leptotrichiaceae which are known to be increased in cirrhosis were reduced with aspirin treatment. On the other hand, Firmicutes, Lachnospiraceae and Ruminococcaceaewere enriched with aspirin treatment (Figure-7E). Linear discriminant analysis identified significant increases in bacteria of phylum Actinobacteria:Mycobacteriaceae, Corynebacteriales, Streptomyces;Proteobacteria:Hyphomonas, Anaplasma, Agrobacterium tumefaciens;Firmicutes:Clostridia, Bacilli, Streptococcus; and others:Tenericutes, Spirochaetes, Aquificae. Whereas several bacteria linked to phylum Actinobacteria:Corynebacterium efficiens, Proteobacteria:Alphaproteobacteria, Teredinibacter turnerae; Firmicutes:Bacilli, Clostridium botulinum and others:Thermotogae, Deinococcus-Thermuswere decreased (p<0.05; Figure-7F, Supplementary-Table-8).

**Figure-7:**
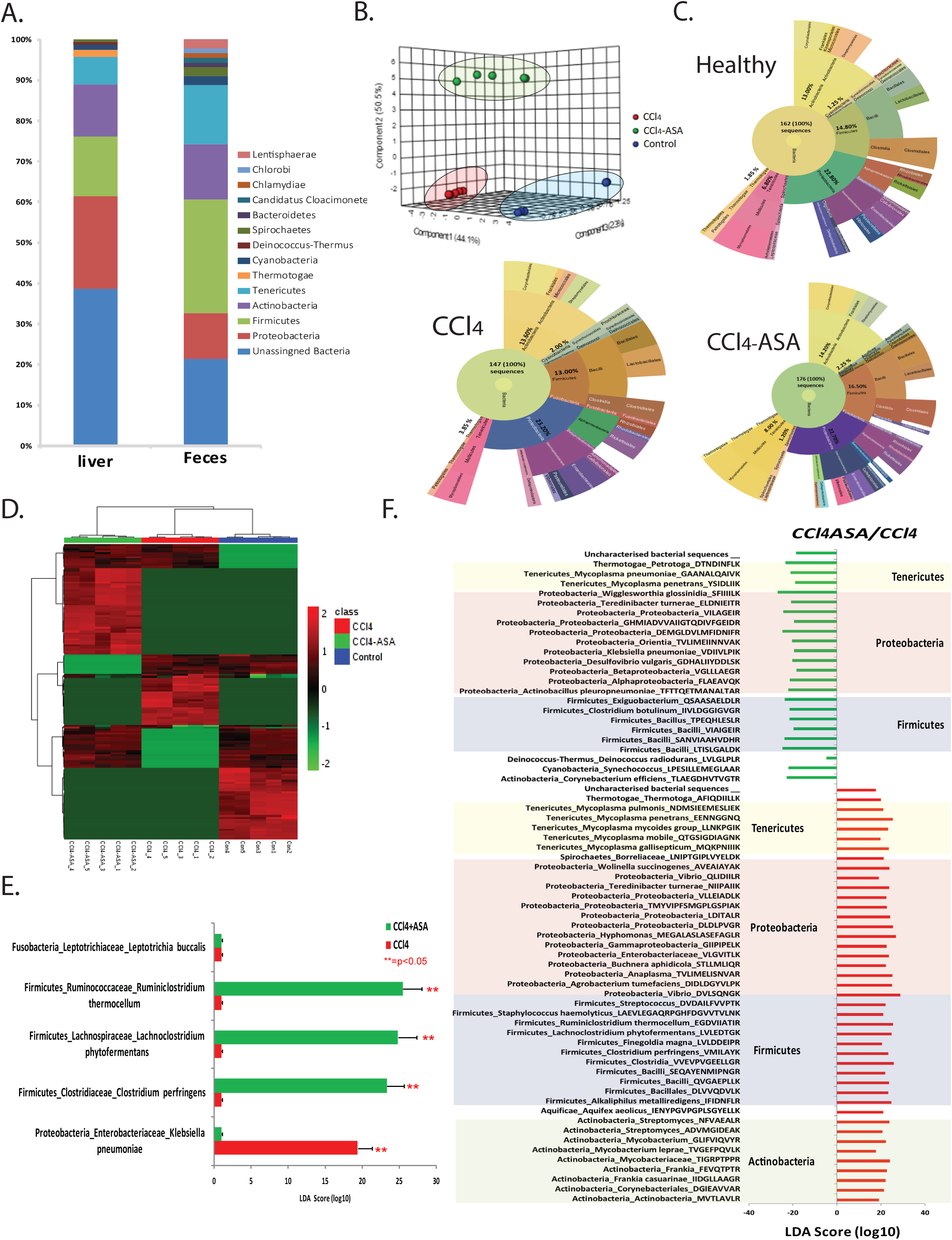
Aspirin adininistartion modulates interahepatic metaproteome in murine model. A: Representative bar charts showing similarity of microbiome alteration derived from liver and fecal samples of mice models. B: Principal component analysis (PCA) of metaproteome data using the Bray-Curtis distance metric C: Sunburst plot representative of microbial population difference across the mice groups. D-E: Metaproteome relative abundance heatmap. Samples are clustered by 1-Pearson correlation and proteins are grouped using KMeans clustering. Relative abundances per protein are colored on a spectrum with red as row maxima and green as row minima. Functional and taxonomic bias within each KMeans cluster is displayed on the right for the significantly altered bacterial species associated with chronic liver disease. F: Linear discriminant analysis reveals taxonomic composition of significant proteins associated with CCl_4_ and CCl_4_+ASA mice group.

Functionality of the hepatic microbiome could also be assessed through metaproteome studies. Aspirin treatment resulted in increased 2-isopropylmalate synthase, Alanine--tRNA ligase activity and reduced DNA-directed RNA polymeraseactivity in in Actinobacteria. Bacteria of the phylum Firmicutes showed higher expression of H(+)-transporting two-sector ATPase and lysidine synthetase and reduced histidine kinase (Figure-8A). This was supplemented with association analysis of the identified metaproteome profile with ALOX5, ARG1 and RYR2 (identified /validated targets associated with hepatic fibrosis). Interestingly, we observed that ARG1 and ALOX5 showed crosscorrelation with many bacterial species and their functional enzymes involved with anti-fibrotic mechanisms (r^2^>0.5, p<0.05, Supplementary-Figure-8B). Together these results suggest that aspirin administration modulated the gut and hepatic microbiome of cirrhotic animals to anti-fibrotic milieu.

**Figure-8:**
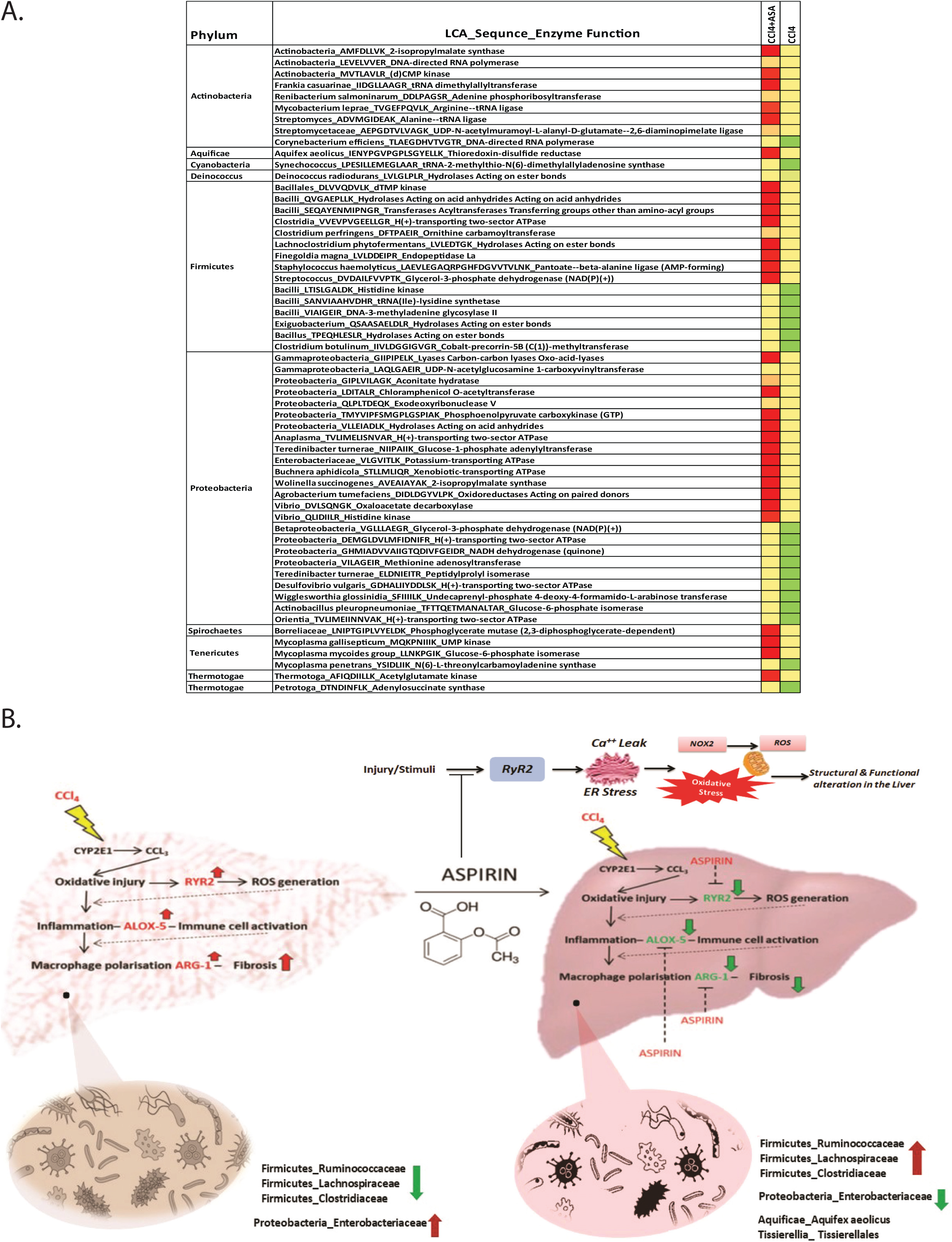
Functionality assesement of hepatic metaproteome. A: Functional analysis of murine liver metaproteome:Taxonomic and functional characterization; Enzyme commission associated with liver metaproteome in high-fibrosis and low-fibrosis (aspirin treated mice group). B: Schematic proposed anti-fibrotic action of aspirin; Chronic liver disease is associated with altered microbiome and bacterial translocation. In this study aspirin treatment was able to reduce bacterial species previously linked to liver disease progression. Aspirin reduces oxidativs stress and improves liver functions by enhancing autophagy and energy metabolic pathways. In brief, expression of key target proteins/metabolites were downregulated linked with oxidative stress proteins (RYR2) (25) which reduces Ca^2+^ leak from ER resulted in decrease in ROS. Simultaneously inflammation (ALOX-5 and KMO) (26, 37) and urea cycle/macrophages polarization (ARG-1) (44) were reduced by aspirin treatment which are otherwise increased in disease proression. Together our finding suggests that during chronic liver injury higher expression of the validated RYR2, ALOX-5, ARG-1 and KMO proteins correlates to increase in liver fibrosis progression and can be modulated by aspirin administration. These results confirm the utility of aspirin as therapeutic in liver fibrosis.

## DISCUSSION

In the present study, the antifibrotic effect of aspirin was studied in the liverofmurine model of CCl_4_. This was complemented with global proteome, metaproteome and metabolome analysis of the liver followed by an in-depth analysis of the regulatory networks. Integrated analysis identified promising antifibrotic targets proteins/metabolites/metaproteins which are under the regulation of aspirin. We validated the expression of RYR2; ryanodine receptor-2, arachidonate-5-lipoxygenase; ALOX-5, arginase-1; ARG-1 in the murine model and showedthattheirexpression correlated with markers of liver fibrosis (α-SMA, PDGR-β levels) and with increase in liver fibrosis. These molecules could be used as a putative candidate for fibrosis monitoring or modulation/therapeutic targets.

Aspirin treatment in murine model of CCl_4_ showed decrease in the hepatic fibrosis. These findings were supported by the liver to body weight ratio, biochemical parameters and by expression of hepatic α-SMA and TGF-β1 expression. These results are consistent with previous studies in rodent model (30).

Proteomics of aspirin treatment in mice with CCl_4_ induced liver fibrosis, showed significant upregulation of drug metabolism, fatty acid degradation, PPAR signalling, and tryptophan metabolism and a significant decrease of cholesterol metabolism, EGFR1 signalling and inflammatory signalling pathways suggesting that aspirin reduces fatty acid accumulation, oxidative stress and inflammation by directly regulating these biological pathways in the liver (31). In total, 33 proteins linked to liver fibrosis were identified and were under the regulation of aspirin. Aspirin treatment significantly reduced the expression of Acox2, Col25a1, Ryr2, Ppa2 and others. Of them ryanodine receptor-2 (RYR2), showed a significant difference.In CCl_4_inducedoxidative injury, the role of RYR2 is well documented (25). Our data suggests that RYR2 modulation could play a role in aspirin mediated fibrosis regression and could be validated as a putative candidate for liver fibrosis.

Concordant to the proteomic analysis, results of the metabolome analysis showed that aspirin induces galactose metabolism, TCA metabolism, mitochondria electron transport chain and reduces butanoate and tryptophan metabolism, taurine/hypotaurine metabolism, phosphatidylethanolamine biosynthesis, urea cycle, leukotriene A4, prostaglandin H2, 20-HETE and others involved in Arachidonic acid metabolism. These results indicate that aspirin improves liver functions and ameliorates liver fibrosis by reduction in inflammatory pathways.

The change in the biochemical/fibrosis parameters reflects an analogous change in the proteome/metabolome and this was documented by global cross-correlation analysis (32). Our analysis showed that biochemical/ fibrosis parameters correlate with proteins/metabolites associated with previously known pathways. For example, change in TGF-β1 correlates to change in MAPK pathways (33), glutathione (34), apoptosis, ferroptosis, and others. Similarly, sirius red score correlated with mTOR signaling (13), prostaglandin synthesis (35) energy pathway and others. Previously our group and others showed that albumin levels or modification is associated with inflammation linked pathways (36). Our results were concordant and showed that change in albumin or AST levels correlate with changes in TNF-alpha, Toll like receptor, IL-1 signaling, and P38-MAP kinase pathways (36). In addition, total bilirubin and ALT was correlated to cholesterol, fatty acid, and tryptophan metabolism linked proteins/metabolites. Overall, these analyses served to validate the association and accuracy of the proteomic/metabolomic measurements.

Analysis of our results highlights an interesting observation pertaining to the utility of aspirin in decreasing the liver damage (ALT levels) which corroborates to the decrease in fibrosis. This observation was validated by identification of immune clusters and its activity followed by its correlation with liver metabolome to determine associated metabolic function. Analysis of our results showed concordant increase in inflammatory clusters with CCl_4_ which got reduced under aspirin treatment, particularly the decrease in immune clusters (5) activity involving “immune activation, enriched in monocytes (II), enriched in monocytes (IV), enriched in neutrophils, type I interferon response” correlated significantly with decrease in histidine and tryptophan metabolism suggesting that aspirin modulates liver micro enviroment by decreasing activation of immune clusters (neutrophils, monocytes, dendritic cells and platelets) and functionality. However. this observation warrants indepth validation.

Integration analysis further validated that aspirin downregulates leukotriene A4, prostaglandin F2 alpha, prostaglandin H2 and arachidonate-5-lipoxygenase (ALOX5) associated to arachidonic acid metabolism in CCl_4_ model. Higher ALOX5 expression is known to be associated with hepatic inflammation and tumurogensis (37). In our analysis aspirin showed modulation of liver function by altering expression of key proteins/metabolites in bile acid metabolism (SOAT2, CYP27A1, taurocholeate) and urea cycle (methylthioadenisine, ornithine, ARG1). Higher expression of ARG-1 is associated with liver steatosis and inhibiting ARG-1 improves hepatic function, reduces lipid accumulation (38). It was interesting to note that, aspirin treatment in the CCl_4_ model of liver fibrosis enhances metabolic energy consumption of the liver by inducing key proteins (IDH3B, MDH) and metabolites (succinate, fumarate, citrate, gluconic acid) in TCA cycle and pentose phosphate pathway.

Recently studies have shown that a significant increase in kynurenine and quinolinic acids areassociated with the pathogenesis of the chronic liver disease (26). Our analysis also showsasignificant reduction of kynurenine and KMO was seen in fibrotic livers post-aspirin treatment. Further our study also showed increase in autophagy related proteins (Deptor, VMP1) which were otherwise decreased in the liver disease (Supplementary-Figure-9) (36).

In our proteomics analysis putative candidate marker RyR2 (oxidative stress protein;(25) was found to be significantly reduced with aspirin. To evaluate the role of RyR2 in liver fibrosis, *in-vitro* model of liver fibrosis using TGF-β as a fibrotic agent in human stellate hepatic cells (LX2) were used and cultured. The treatment with TGF-β leading to increase in intracellular stress and ROS levels contributes to collagen release causing the onset and progression of fibrosis (39). In the present study, we revealed that aspirin inhibited the oxidative stress by reducing intracellular Ca^++^stress regulated by RyR2 expression in activated LX2 cells. These results can be explained by the principle that treatment with TGF-β induces prolonged and sustained ER stress in HSCs and the ER stress-related leak in Ca^++^ resulted in disbalance and oxidative stress (40). This finding was corroborated in our analysis with the cytosolic Ca^++^ levels in activated LX-2 cells which were concordant with RyR2 expression (Figure 5E). In chronic liver disease, perturbations in ER homeostasis leading to ER stress/dysfunction, which promotes calcium release from the ER accompanied by some related apoptosis effectors released from mitochondria (41). Although the exact functional role of Ca^++^ mobilization mediated by RyR2 over expression in liver fibrosis and its modulation by aspirin warrants further elucidation, the possibility that the calcium released from the ER triggers mitochondrial Ca^2+^ uptake, promotes ROS production and apoptosis seems reasonable.

Aspirin administration modulates intra-hepatic microbiome. We noted the higher presence of E. coli in livers of CCl_4_ treated mice compared to aspirin treated group. Previously, bacterial translocation and derived factors from gut to liver has been linked with the major mechanism of pathogenesis of NAFLD (28) and cirrhosis-associated liver diseases in animals and humans (36). In addition, change in the microbiome and aspirin bioavailability isalso interlinked. We reasoned that aspirin could modulate liver microbiome changing the microenvironment and hence could inhibit fibrosis progression. Metaproteome analysis of the stool and liver were similar at the phylum level. Further metaproteins analysis of the liver showed thatFirmicutes; Streptococcaceae, Enterococcaceae, Staphylococcaceae and Proteobacteria; Enterobacteriaceaeweresignificantly reduced with aspirin treatment which were otherwise increased in cirrhotics (42). In addition, the levels of Lachnospiracea and Ruminococcaceae (43); bacteria which are reduced in chronic liver diseases, were increased post-aspirin treatment. Aspirin treatment modified the functional process of the bacteria as evident by changes in several bacterial enzymes, like significant increase in function of glycerol-3-phosphate dehydrogenase (NAD(P)(+) and dTMP kinase and decrease in the function tRNA(Ile)-lysidine synthetase, DNA-3-methyladenine glycosylase II in Firmicutes. A dominant change in enzyme activities of proteobacteria and actinobacteria was also observed. Cross correlation analysis of the validated targets; ARG-1, ALOX5 and RYR2 with the intrahepatic bacteria taxonomy highlighted significant and direct association of many bacterial species and their enzyme activity. These results clearly show that aspirin administration modulates hepatic microbiome and its functionsthus making the microenvironment favourable for fibrosis regression.

Results of the validated targets RYR2; ryanodine-receptor-2 (25), ALOX5; arachidonate-5-lipoxygenase (37), ARG-1:arginase-1 (44) and KMO;kynurenine-3-monooxygenase (45) correlated significantly with the degree of liver fibrosis. These validated targets also documented significant correlation with the liver microbiome. These results clearly indicate that aspirin modulates liver microenvironment by i) altering the microbiome, ii) decreasing RYR2 expression (oxidative stress), iii) reducing ALOX5 and KMO expression (inflammation). Moreover, aspirin significantly induced initiation of autophagy, energy metabolism (TCA and pentose phosphate pathway) and improves liver functions by reducing bile acid metabolites and ARG-1 expression in the murine model of liver fibrosis. Suggesting multi modular effect of aspirin in fibrosis regression.

In conclusion, we present a hepatic multi-omics profiling study in the murine model of liver fibrosis in presence or absence of aspirin treatment. We documented many beneficial effect of aspirin administration which included reduction in oxidative stress, inflammation and hepatic fibrosis. We were able to underline and validate previously known (ALOX5, ARG1) and novel targets of aspirin mediated reduction in hepatic fibrosis. In the study the reduction of RYR2 via aspirin in mice model of liver fibrosis highlights as a new and promising antifibrotic target in the liver which could be further explored for theraputic intervention. Additionally we show that aspirin modulates liver microbiome and improves liver functions and can be used as an attractive therapeutic molecule for preventing hepatic disease progression and possibly fibrosis regression.

## References

1. Rosenbloom J, Macarak E, Piera-Velazquez S, and Jimenez SA. Human Fibrotic Diseases: Current Challenges in Fibrosis Research. Methods in molecular biology. 2017;1627(1–23.

2. Wynn TA, and Ramalingam TR. Mechanisms of fibrosis: therapeutic translation for fibrotic disease. Nature medicine. 2012;18(7):1028–40.

3. Mederacke I, Hsu CC, Troeger JS, Huebener P, Mu X, Dapito DH, Pradere JP, and Schwabe RF. Fate tracing reveals hepatic stellate cells as dominant contributors to liver fibrosis independent of its aetiology. Nature communications. 2013;4(2823.

4. Yoshida S, Ikenaga N, Liu SB, Peng ZW, Chung J, Sverdlov DY, Miyamoto M, Kim YO, Ogawa S, Arch RH, et al. Extrahepatic platelet-derived growth factor-beta, delivered by platelets, promotes activation of hepatic stellate cells and biliary fibrosis in mice. Gastroenterology. 2014;147(6):1378–92.

5. Sahasrabuddhe VV, Gunja MZ, Graubard BI, Trabert B, Schwartz LM, Park Y, Hollenbeck AR, Freedman ND, and McGlynn KA. Nonsteroidal anti-inflammatory drug use, chronic liver disease, and hepatocellular carcinoma. Journal of the National Cancer Institute. 2012;104(23):1808–14.

6. Paik YH, Kim JK, Lee JI, Kang SH, Kim DY, An SH, Lee SJ, Lee DK, Han KH, Chon CY, et al. Celecoxib induces hepatic stellate cell apoptosis through inhibition of Akt activation and suppresses hepatic fibrosis in rats. Gut. 2009;58(11):1517–27.

7. Jiang ZG, Feldbrugge L, Tapper EB, Popov Y, Ghaziani T, Afdhal N, Robson SC, and Mukamal KJ. Aspirin use is associated with lower indices of liver fibrosis among adults in the United States. Alimentary pharmacology & therapeutics. 2016;43(6):734–43.

8. Malehmir M, Pfister D, Gallage S, Szydlowska M, Inverso D, Kotsiliti E, Leone V, Peiseler M, Surewaard BGJ, Rath D, et al. Platelet GPIbalpha is a mediator and potential interventional target for NASH and subsequent liver cancer. Nature medicine. 2019;25(4):641–55.

9. Simon TG, Henson J, Osganian S, Masia R, Chan AT, Chung RT, and Corey KE. Daily Aspirin Use Associated With Reduced Risk For Fibrosis Progression In Patients With Nonalcoholic Fatty Liver Disease. Clin Gastroenterol Hepatol. 2019;17(13):2776–84 e4.

10. Pietrocola F, Castoldi F, Markaki M, Lachkar S, Chen G, Enot DP, Durand S, Bossut N, Tong M, Malik SA, et al. Aspirin Recapitulates Features of Caloric Restriction. Cell Rep. 2018;22(9):2395–407.

11. Sato S, Solanas G, Peixoto FO, Bee L, Symeonidi A, Schmidt MS, Brenner C, Masri S, Benitah SA, and Sassone-Corsi P. Circadian Reprogramming in the Liver Identifies Metabolic Pathways of Aging. Cell. 2017;170(4):664–77 e11.

12. Celikbilek M. Letter: increased platelet activation in chronic liver disease--hit two targets with a single shot. Alimentary pharmacology & therapeutics. 2016;43(9):1023.

13. Din FV, Valanciute A, Houde VP, Zibrova D, Green KA, Sakamoto K, Alessi DR, and Dunlop MG. Aspirin inhibits mTOR signaling, activates AMP-activated protein kinase, and induces autophagy in colorectal cancer cells. Gastroenterology. 2012;142(7):1504–15 e3.

14. Simon TG, Henson J, Osganian S, Masia R, Chan AT, Chung RT, and Corey KE. Daily Aspirin Use Associated With Reduced Risk For Fibrosis Progression In Patients With Nonalcoholic Fatty Liver Disease. Clin Gastroenterol Hepatol. 2019.

15. Kim YG, Udayanga KG, Totsuka N, Weinberg JB, Nunez G, and Shibuya A. Gut dysbiosis promotes M2 macrophage polarization and allergic airway inflammation via fungi-induced PGE(2). Cell host & microbe. 2014;15(1):95–102.

16. Ghiassi-Nejad Z, Hernandez-Gea V, Woodrell C, Lang UE, Dumic K, Kwong A, and Friedman SL. Reduced hepatic stellate cell expression of Kruppel-like factor 6 tumor suppressor isoforms amplifies fibrosis during acute and chronic rodent liver injury. Hepatology. 2013;57(2):786–96.

17. Bulckaen H, Prevost G, Boulanger E, Robitaille G, Roquet V, Gaxatte C, Garcon G, Corman B, Gosset P, Shirali P, et al. Low-dose aspirin prevents age-related endothelial dysfunction in a mouse model of physiological aging. American journal of physiology Heart and circulatory physiology. 2008;294(4):H1562–70.

18. Shannon P, Markiel A, Ozier O, Baliga NS, Wang JT, Ramage D, Amin N, Schwikowski B, and Ideker T. Cytoscape: a software environment for integrated models of biomolecular interaction networks. Genome Res. 2003;13(11):2498–504.

19. Basu S, Duren W, Evans CR, Burant CF, Michailidis G, and Karnovsky A. Sparse network modeling and metscape-based visualization methods for the analysis of large-scale metabolomics data. Bioinformatics. 2017;33(10):1545–53.

20. Song YN, Dong S, Wei B, Liu P, Zhang YY, and Su SB. Metabolomic mechanisms of gypenoside against liver fibrosis in rats: An integrative analysis of proteomics and metabolomics data. PloS one. 2017;12(3):e0173598.

21. Potter JJ, Rennie-Tankesley L, and Mezey E. Influence of leptin in the development of hepatic fibrosis produced in mice by Schistosoma mansoni infection and by chronic carbon tetrachloride administration. Journal of hepatology. 2003;38(3):281–8.

22. Fujii T, Fuchs BC, Yamada S, Lauwers GY, Kulu Y, Goodwin JM, Lanuti M, and Tanabe KK. Mouse model of carbon tetrachloride induced liver fibrosis: Histopathological changes and expression of CD133 and epidermal growth factor. BMC gastroenterology. 2010;10(79.

23. Cano A, Marino Z, Millet O, Martinez-Arranz I, Navasa M, Falcon-Perez JM, Perez-Cormenzana M, Caballeria J, Embade N, Forns X, et al. A Metabolomics Signature Linked To Liver Fibrosis In The Serum Of Transplanted Hepatitis C Patients. Scientific reports. 2017;7(1):10497.

24. Li S, Sullivan NL, Rouphael N, Yu T, Banton S, Maddur MS, McCausland M, Chiu C, Canniff J, Dubey S, et al. Metabolic Phenotypes of Response to Vaccination in Humans. Cell. 2017;169(5):862–77 e17.

25. Liao B, Zhang Y, Sun H, Ma B, and Qian J. Ryanodine Receptor 2 Plays a Critical Role in Spinal Cord Injury via Induction of Oxidative Stress. Cell Physiol Biochem. 2016;38(3):1129–37.

26. Claria J, Moreau R, Fenaille F, Amoros A, Junot C, Gronbaek H, Coenraad MJ, Pruvost A, Ghettas A, Chu-Van E, et al. Orchestration of Tryptophan-Kynurenine Pathway, Acute Decompensation, and Acute-on-Chronic Liver Failure in Cirrhosis. Hepatology. 2019;69(4):1686–701.

27. Sin YY, Ballantyne LL, Mukherjee K, St Amand T, Kyriakopoulou L, Schulze A, and Funk CD. Inducible arginase 1 deficiency in mice leads to hyperargininemia and altered amino acid metabolism. PLoS One. 2013;8(11):e80001.

28. Woodhouse CA, Patel VC, Singanayagam A, and Shawcross DL. Review article: the gut microbiome as a therapeutic target in the pathogenesis and treatment of chronic liver disease. Alimentary pharmacology & therapeutics. 2018;47(2):192–202.

29. Gurdeep Singh R, Tanca A, Palomba A, Van der Jeugt F, Verschaffelt P, Uzzau S, Martens L, Dawyndt P, and Mesuere B. Unipept 4.0: Functional Analysis of Metaproteome Data. Journal of proteome research. 2019;18(2):606–15.

30. Li CJ, Yang ZH, Shi XL, and Liu DL. Effects of aspirin and enoxaparin in a rat model of liver fibrosis. World journal of gastroenterology. 2017;23(35):6412–9.

31. Hu X, Shen H, Wang Y, Zhang L, and Zhao M. Aspirin-triggered resolvin D1 alleviates paraquat-induced acute lung injury in mice. Life Sci. 2019;218(38–46.

32. Niu L, Geyer PE, Wewer Albrechtsen NJ, Gluud LL, Santos A, Doll S, Treit PV, Holst JJ, Knop FK, Vilsboll T, et al. Plasma proteome profiling discovers novel proteins associated with non-alcoholic fatty liver disease. Molecular systems biology. 2019;15(3):e8793.

33. Lee MK, Pardoux C, Hall MC, Lee PS, Warburton D, Qing J, Smith SM, and Derynck R. TGF-beta activates Erk MAP kinase signalling through direct phosphorylation of ShcA. EMBO J. 2007;26(17):3957–67.

34. Liu RM, and Gaston Pravia KA. Oxidative stress and glutathione in TGF-beta-mediated fibrogenesis. Free Radic Biol Med. 2010;48(1):1–15.

35. Boutaud O, Sosa IR, Amin T, Oram D, Adler D, Hwang HS, Crews BC, Milne G, Harris BK, Hoeksema M, et al. Inhibition of the Biosynthesis of Prostaglandin E2 By Low-Dose Aspirin: Implications for Adenocarcinoma Metastasis. Cancer Prev Res (Phila). 2016;9(11):855–65.

36. Starkel P, and Schnabl B. Bidirectional Communication between Liver and Gut during Alcoholic Liver Disease. Seminars in liver disease. 2016;36(4):331–9.

37. Titos E, Ferre N, Lozano JJ, Horrillo R, Lopez-Parra M, Arroyo V, and Claria J. Protection from hepatic lipid accumulation and inflammation by genetic ablation of 5-lipoxygenase. Prostaglandins & other lipid mediators. 2010;92(1-4):54–61.

38. Alisi A, Comparcola D, De Stefanis C, and Nobili V. Arginase 1: a potential marker of a common pattern of liver steatosis in HCV and NAFLD children. Journal of hepatology. 2015;62(5):1207–8.

39. Hernandez-Gea V, Hilscher M, Rozenfeld R, Lim MP, Nieto N, Werner S, Devi LA, and Friedman SL. Endoplasmic reticulum stress induces fibrogenic activity in hepatic stellate cells through autophagy. Journal of hepatology. 2013;59(1):98–104.

40. Koo JH, Lee HJ, Kim W, and Kim SG. Endoplasmic Reticulum Stress in Hepatic Stellate Cells Promotes Liver Fibrosis via PERK-Mediated Degradation of HNRNPA1 and Up-regulation of SMAD2. Gastroenterology. 2016;150(1):181–93 e8.

41. Malhi H, and Kaufman RJ. Endoplasmic reticulum stress in liver disease. Journal of hepatology. 2011;54(4):795–809.

42. De Minicis S, Rychlicki C, Agostinelli L, Saccomanno S, Candelaresi C, Trozzi L, Mingarelli E, Facinelli B, Magi G, Palmieri C, et al. Dysbiosis contributes to fibrogenesis in the course of chronic liver injury in mice. Hepatology. 2014;59(5):1738–49.

43. Bajaj JS, Heuman DM, Hylemon PB, Sanyal AJ, White MB, Monteith P, Noble NA, Unser AB, Daita K, Fisher AR, et al. Altered profile of human gut microbiome is associated with cirrhosis and its complications. Journal of hepatology. 2014;60(5):940–7.

44. Moon J, Do HJ, Cho Y, and Shin MJ. Arginase inhibition ameliorates hepatic metabolic abnormalities in obese mice. PLoS One. 2014;9(7):e103048.

45. Vilar-Gomez E, and Chalasani N. Daily Aspirin Use Reduces Risk of Fibrosis Progression in Patients With Nonalcoholic Fatty Liver Disease, Providing New Uses for an Old Drug. Clin Gastroenterol Hepatol. 2019.

