## supplementary methods and tables for "Multi-omics analysis of aspirin treatment response in mice provide molecular insights and targets linked to liver fibrosis regression"

**Table of contents:**

Supplementary Material and Methods

Supplementary Figure 1
Supplementary Figure 2
Supplementary Figure 3A, 3B
Supplementary Figure 4
Supplementary Figure 5
Supplementary Figure 6A, 6B
Supplementary Figure 7A, 7B
Supplementary Figure 8A, 8B
Supplementary Figure 9

Supplementary Table 1
Supplementary Table 2
Supplementary Table 3
Supplementary Table 4
Supplementary Table 5
Supplementary Table 6
Supplementary Table 7
Supplementary Table 8

#### Supplementary Material and Methods

##### Animals and Sample Collection:

Male C57BL/6J mice of 6-8 weeks, weighting 20-25 g were randomly divided into control group and experimental group, the control group, Aspirin (ASA) group, Carbon tetrachloride ($CCl_4$) group, and $CCl_4$+ASA group (n=08 animals/group). Mice were subjected to twice a week intraperitoneal (i.p.) injections of 10% Carbon tetrachloride ($CCl_4$) (Sigma, 270652 (diluted in olive oil) at a dose of 0.5 µl/g body weight for 12 weeks to induce liver fibrosis (1) or vehicle (olive oil) and concomitant with aspirin treatment. Dose of $CCl_4$ were routinely adjusted based on weight of the animal. Harvested tissues/serum was snap-frozen in liquid nitrogen (N2) and kept at −80°C until batch analysis. Ethics approval for this study was obtained from the institutional review board (IRB-ILBS-2019). Also for animal study approval was taken from the ethics committee, in accordance with CPSEA (committee for the purpose of control and supervision of experiments on animals) guidelines IAEC/ILBS/16/07. Animal grouping, model development is given in Figure 1A and flow chart of experiments performed in Supplementary Figure-1.

**Aspirin treatments:** Low dose of aspirin (50mg/l in drinking water, pH 6.4) was administered in drinking water which was replaced every other day (2). Considering that each animal drinks in average 3 to 4 mL of water per day, this would be equal to 150µg to 200µg aspirin per day for a mouse of 30g weight, (2) which is significantly less than 6 $mg/kg^{-1}$ per day; known dose for induction of platelet Cox I/II inhibition as described (3). As the study required long term administration of aspirin treatment drinking dose of aspirin was preferred over daily oral gavage.

##### Liver Function Enzymes

Blood was collected from mice by the retrobulbor route. The serum was separated by centrifugation 3000g for 10 minutes. Aspartate amino transferase (AST), alanine amino transferase (ALT) levels and others were detected by an Olympus AU5400 automatic analyser at biochemistry laboratory facility at ILBS.

#### Human specimens

The Fibrotic liver samples (fibrosis stage: F1-4) were obtained from patients undergoing routine transjugular liver biopsy liver biopsy (TJLB) collected from institute bio-bank facility ILBS-New Delhi. Normal human liver samples (F0) were collected from 15 patients undergoing hepatic resection for colorectal metastasis (n = 15). Liver biopsy specimens were immediately fixed, paraffin-embedded, and stained with haematoxylin and eosin, Masson's trichrome. All specimens were re-examined and graded by the pathologists. A liver fibrosis score from F0 to F4 were adopted according to the severity and the extent of damage. Grading of liver fibrosis was evaluated according to the METAVIR scoring system. Fibrosis (F) was staged on a scale of 0 to 4: F0, no fibrosis; F1, portal fibrosis without septa; F2, few septa; F3, numerous septa without cirrhosis; F4, cirrhosis (4). All tissues were obtained with donor consent by writing and the prior approval was taken from Institute Ethics Committee-ILBS.

Table: Baseline characteristics.

| Characteristics | F0-F1(n=15) | F2 (n=15) | F3 (n=15) | F4 (n=15) |
|---|---|---|---|---|
| Age(years) | 42±14 | 49±7 | 47±8 | 48±8 |
| Sex(F/M) | 0/15 | 0/15 | 0/15 | 0/15 |
| ALT (U/L) | 80.0±10.0 | 85.0±10.0 | 98.0±6.0** | 89.0±18.0 |
| AST (U/L) | 34.0±6.0 | 45.0±10.0 | 45.0±10.0 | 60.8±10.0 |
| Bilirubin (mg/dL) | 0.24±0.5 | 3.8±1.5 | 6.5±2.5 | 09.5±6.5** |
| Etiology (HCV) | (15/15) | (15/15) | (15/15) | (15/15) |

Values represent mean±SD; *P<0.05 higher when compared to F0-F1, F2, and F3; **P<0.001 higher when compared to F0 or F2.

#### Liver proteome analysis.

Total proteins were isolated from fifty-milligram liver tissues (n=5 samples/group randomly chosen) and was subjected to reduction, alkylation followed by digestion for 16-20 h at 37°C using sequencing-grade modified trypsin: proteins (1ug:20ug w/w). The samples were desalted and subjected to LC-MS/MS analysis. The peptides were eluted by a 3–95% gradient of buffer B (aqueous 80% acetonitrile in 0.1% formic acid) at a flow rate of 300 nL/min for about 150 minutes on a 50-cm analytical C18 column (C18, 3 μm, 100 Å) The peptides were ionized by nano-electrospray and subsequent tandem mass spectrometry (MS/MS) on a Q-ExactiveTM Plus (Thermo Fisher Scientific, San Jose, CA, United States). The peptides were analyzed using a mass spectrometer with the collision-induced dissociation mode with the electrospray voltage was 2.2 kV. Analysis on the orbitrap was performed with full scan MS spectra with a resolution of 70,000 from m/z 350 to 1800. The MS/MS data was analyzed by Proteome Discoverer (version 2.0, Thermo Fisher Scientific, Waltham, MA, United States) using the mice protein database (UP000000589) and mice proteome with Mascot algorithm (Mascot 2.4, Matrix Science). Significant proteins were identified at ($P<0.05$) and q values ($P<0.05$). The threshold of false discovery rate was kept at 0.01. The identified proteins were subjected to standard statistical analysis and network and pathway analysis (2, 5).

**Liver Metabolome analysis**

Metabolites were extracted from the liver lysate using organic phase extraction protocol (6). To brief 25g liver tissue was crushed in liquid nitrogen in 500ul RIPA buffer. To 100 ul of liver lysate 500 ul methanol was added and kept overnight at -20 degree for protein precipitation. The samples were centrifuged at13.3 rpm for 10 min and the supernatant was dried under vacuum. The dried samples were reconstituted with (5:95:5) 5% acetonitrile: 95% water: 5% internal standards at known concentrations, for reverse-phase chromatography by using C18 column (Thermo Scientific™25003102130: 3 µm, 2.1 mm, 100 mm) and at (60 ACN:40 $H_2O$:5 Internal/external standard) for analysis on HILIC analysis (Thermo Scientific™ 26505102130 5 µm, 2.1 mm, 100 mm) using ultra-high performance liquid chromatographic system followed by high-resolution mass spectrometry (MS)(7). Metabolite features were extracted using compound discoverer 3.0 (Thermo) (8). Annotation of the features was performed using mass list searches (7), mzCloud™ (www.mzcloud.org) and mummichog (6). Identified and annotated features were subjected to log normalization and pareto-scaling using metaboanalyst 4.0 (http://metaboanalyst.ca.) server (9) and into SIMCA P12 software (Umetrix, Sweden) for multivariate projection analyses, such as principal component analysis and partial least square discriminant analysis. Pathway enrichment patterns were analyzed using Metaboanalyst 4.0 (9).

**Metaproteome analysis of liver tissue and fecal samples:**

Proteins were isolated form liver tissue and fecal samples of the study groups. The isolated proteins were reduced, alkylated and digested using trypsin followed by mass spectrometry analysis similar to that stated in liver proteome analysis section. The MS/MS data was analyzed by Proteome Discoverer (version 2.0, Thermo Fisher Scientific, Waltham, MA, United States) using the bacterial/fungal sequence (UniprotSwP_20170609, with sequences 467231 and MG_BG__UPSP with sequences 2019194). This was cross validated using Mascot algorithm (Mascot 2.4, Matrix Science) specifically for all possible microbial species. In order to remove the interference of host proteome analysis was based on peptide matches. In brief, significant peptide groups were identified at ($P<0.05$) and q values ($P<0.05$) and the false discovery rate at 0.01. Only rank 1 peptides with Peptide Sequence Match (PSM)>3 were subjected to biodiversity and functional analysis using unipept (10). Only bacterial species associated peptides were segregated and were subjected to statistical, functional and biodiversity analysis (10).

**Cell Culture**

The LX-2 cells were cultured and maintained in medium DMEM low glucose supplemented with 10% FBS initially followed by 2% FBS and 1% antibiotic solution (Penicillin and Streptomycin) incubated at 37 °C in a 5% $CO_2$ humidified atmosphere. For this study, the cells were seeded in 6-well plates and then treated with 5 ng/mL TGF-β to induce their activation (α-SMA) and the collagen release, in the presence or absence of Aspirin 5 μmol/ml.

**Intracellular calcium**

Calcium signaling regulates hepatic stellate cell (HSC) activation and growth (11) intracellular calcium flux in LX2 was measured using Fluo-8 calcium assay kit (Catalogue No. sc-362561). Briefly, the LX2 cells were mixed with Fluo-8 solution (5μM) and incubated at 37°C.After 30 minutes of incubation LX2 cells were washed by $Ca^{+2}/Mg^{+2}$ free PBS buffer and the intracellular calcium flux was measured by fluorescence microscope.

**ROS detection assay**

ROS were detected by the cell-permeable fluorescent probe dihydrorhodamine-123 (DHR-123) (Sigma) as described previously (12). Briefly, LX2 cells were co-incubated with 10 ng/L DHR-123 for 1 h at 37 °C in the dark. Then, the cells were rinsed, and the signal was detected by a flow cytometer (BD-FACSVerse).

**Gene expression analysis by quantitative real-time PCR**

RNA was isolated form the liver tissue using TRIzol™ Reagent (Cat no. 15596026, Thermo) this was followed by cDNA synthesis (Cat no.AB1453A, Verso cDNA kit, Thermo) and real time PCR analysis on ABI 7300 real-time PCR system. Gene expression was calculated using the ΔΔCT method normalized against GAPDH.

**Bacterial DNA detection in liver using quantitative reverse-transcription PCR**.

Mice liver tissue was subjected to DNA isolation with DNeasy Blood & Tissue kits (Qiagen, USA). PCR was performed using Taqman (Life Technologies) gene expression assay probe for Rps18 (Mm02601778_g1) (13), or custom-made probes for E.coli, (Supplementary Table-2). Expression Rps18 was used as the housekeeping gene. Reactions were performed in real time PCR analysis on a 384-well assay format using ABI 7300 real-time PCR instrument**.**

**Statistical Analysis**

Data are expressed as means ± standard errors of the means of at least three independent experiments. Student t test and Pearson correlation were used for statistical analysis. A P-value (*$P < 0.05$, **$P < 0.01$) were considered statistically significant.

**Supplementary figures legends:**

**Supplementary Figure 1:** Flow chart diagram.

**Supplementary Figure 2:** Proteome analysis across the mice groups.

**Supplementary Figure 3A:** Pathway Analysis of upregulated proteins in CCl4 with Aspirin.

**Supplementary Figure 3B:** Pathway Analysis of downregulated proteins in $CCl_4$ with Aspirin**.**

**Supplementary Figure 4:** The mRNA expression analysis of oxidative stress related genes.

**Supplementary Figure 5:** Metabolome analysis across the mice groups.

**Supplementary Figure 6A:** Common up- and downregulated pathways in low and high fibrosis.

**Supplementary Figure 6B:** Most perturbed 39 pathways analysis in low and high fibrosis.

**Supplementary Figure 7A:** Integration analysis of proteins/metabolites (energy metabolism).

**Supplementary Figure 7B:** Integration analysis of proteins/metabolites-tryptophan metabolism.

**Supplementary Figure 8A:** Modulation of hepatic microbiome with aspirin treatment.

**Supplementary Figure 8B:** Correlation analysis of metaproteome with key anti-fibrotic targets

**Supplementary Figure 9:** Modulation of autophagy related proteins with aspirin administration.

**Supplementary table legends:**

**Supplementary Table 1:** Serum biochemical analysis.

**Supplementary Table 2:** Primer list.

**Supplementary Table 3:** Liver proteome analysis.

**Supplementary Table 4:** OMIM database fibrosis associated proteins.

**Supplementary Table 5:** Liver metabolome analysis.

**Supplementary Table 6:** Blood transcription module (BTM) enrichment analysis.

**Supplementary Table 7:** AUROC analysis.

**Supplementary Table 8:** Liver metaproteome analysis.

.

**Supplementary figure 1:** Flow chart diagram of experimental figures.

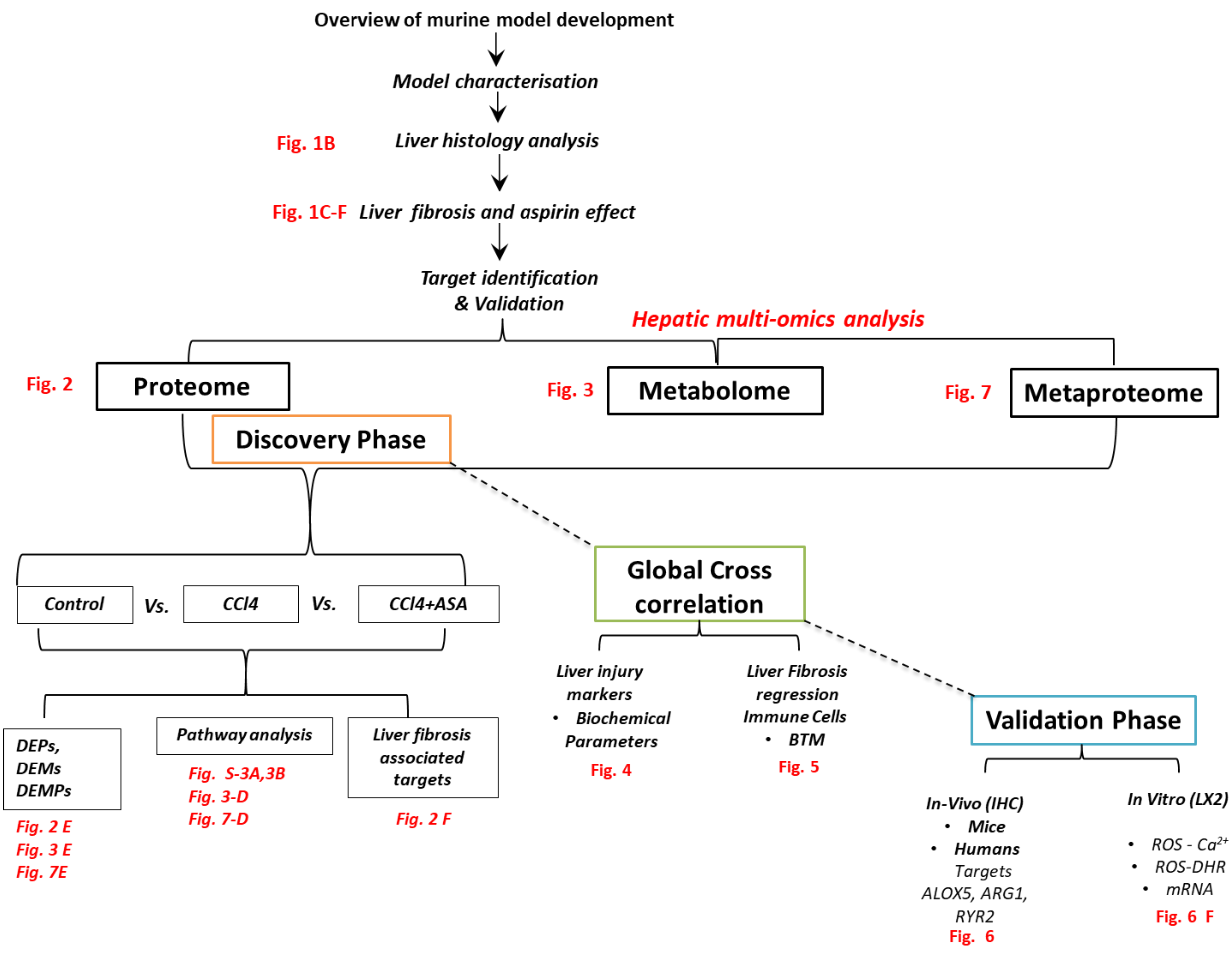

Figure S-1: Workflow and summary of the experiments performed and number of samples used in target determination, and validation experiments including the In-vivo and In-vitro analysis. Respective figures are given in red number for each part of results.

**Explanation for supplementary figure 2:**

**Aspirin administration decreased inflammation and liver fibrosis associated proteome**

Liver proteome was analysed in murine model of fibrosis with and without aspirin treatment across the group (Figure-2A). As compared to controls a total of 176 proteins (96 up- and 80 downregulated) in ASA group, 339 proteins (233 up- and 106 downregulated) in CCl4 and 514 proteins (262 up- and 252 downregulated) in CCl4+ASA groups were differentially expressed (Figure-2B, Supplementary- Table-3). Partial least square discriminant analysis (PLS-DA; Figure-2C) and unsupervised clustering analysis showed clear distinction between the groups (Figure-2D). Of the upregulated (>2-fold) proteins (Figure-2E) a total of 37 (8.1%) proteins were specific to the ASA group, 140 (30.6%) were specific to CCl4, and 165 (36%) specific to CCl4+ASA group (Figure-2E). Proteins specific to the CCl4+ASA group were linked to autophagy, drug metabolism, fatty acid degradation, PPAR signalling, histidine metabolism, amino acid metabolism and glutathione metabolism (Supplementary-Figure-3A).Of the downregulated (<2-fold) proteins, a total of 32 (9.4%) were specific to the ASA group, 48 (14.1%) to CCl4, and 177 (52.1%) to CCl4+ASA group (Figure-2F). Proteins specific to CCl4+ASA were linked to cholesterol metabolism, EGFR1 signalling, arachidonic acid, ferroptosis, phagosome, and inflammatory signaling pathway (TNF-alpha NFKB signaling pathway; Supplementary-Figure-3B).

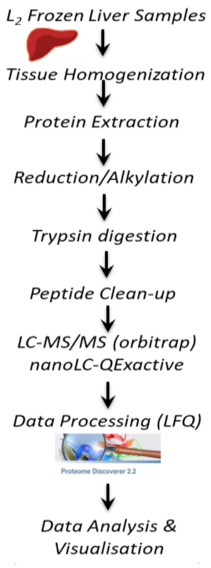

**Supplementary Figure 2:** Proteome analysis across the groups in the $CCl_4$ model of fibrosis with and without aspirin treatment

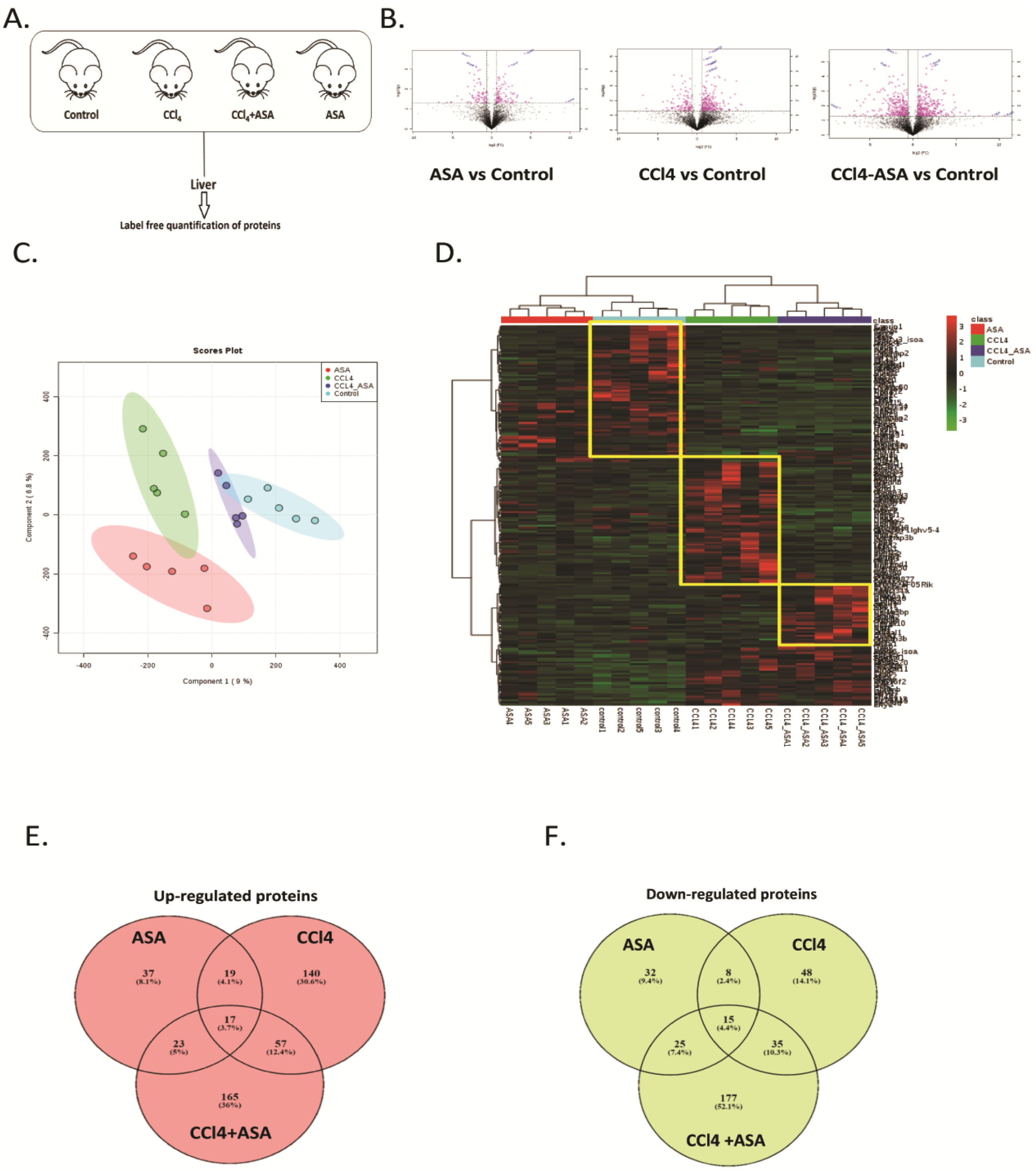

**Figure-S2: Proteomics analysis of the murine model of liver fibrosis at 12 weeks and comparison to aspirin treatment.**

A: Overview of label free proteomic study performed across the mice groups.

B: Differentially expressed proteins were visualized by volcano plot and filtered so as to include only statistically significantly changed proteins ($P < 0.05$, t-test) that were changed by more than 1.5-fold in mice livers.

C: PCA plot analysis between the mice groups.

D: Dendrogram representing the hierarchical cluster of differentially abundant proteins among liver samples from mice groups classified as Con, $CCl_4$, $CCl_4$+ASA and ASA.

E, F: Venn diagram showing differentially regulated proteins in different mice groups. Red venn = upregulated proteins and Green venn= downregulated proteins.

**Supplementary Figure 3A:** Pathway Analysis of upregulated proteins in $CCl_4$ with Aspirin.

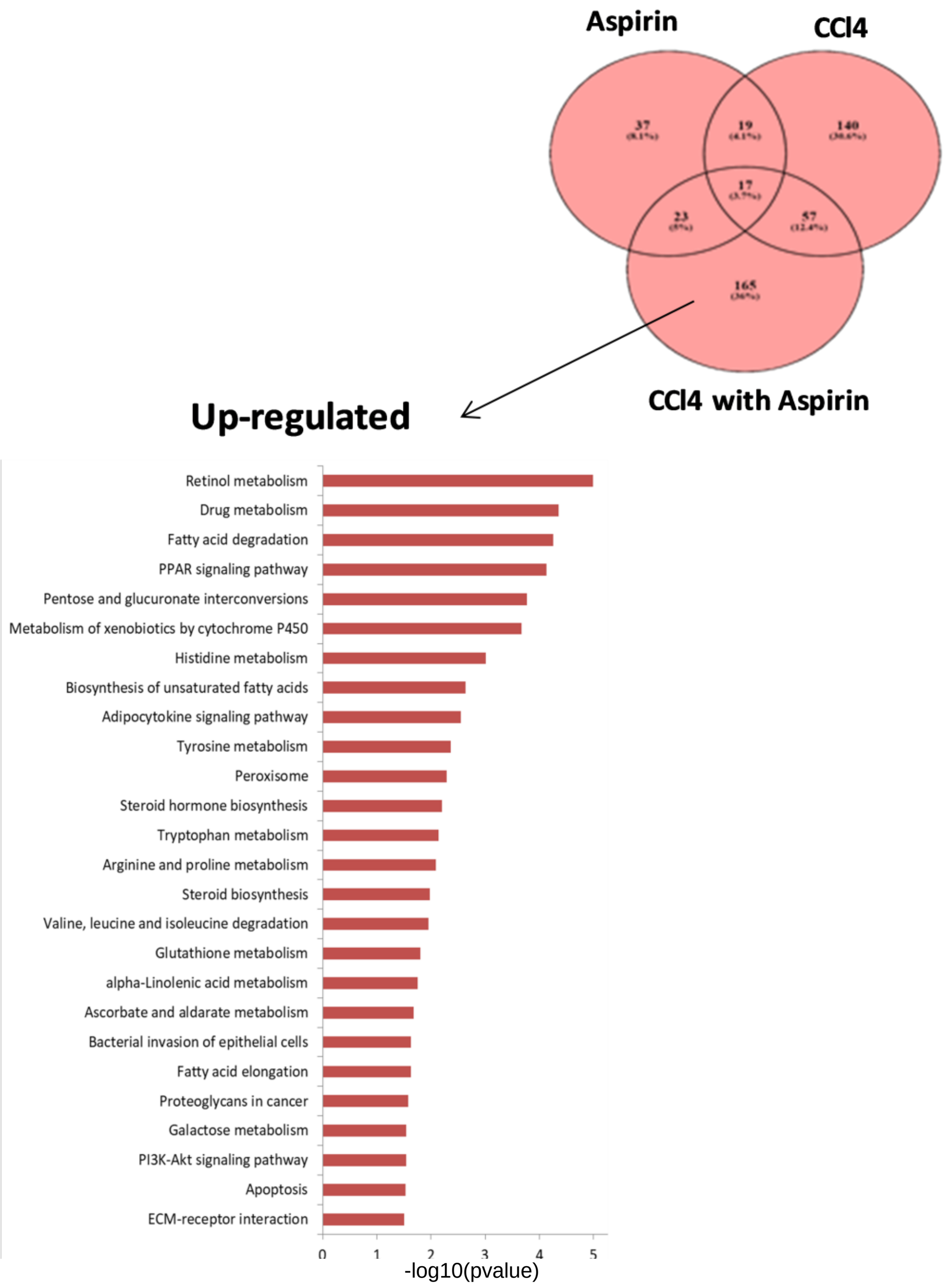

**Figure S-3A:** A total of 165 up-regulated proteins in CCl4+Aspirin were associated to pathway linked to drug metabolism, fatty acid degradation, PPAR signalling, histidine metabolism and amino acid metabolism.

**Supplementary Figure 3B:** Pathway Analysis of downregulated proteins in $CCl_4$ with Aspirin**.**

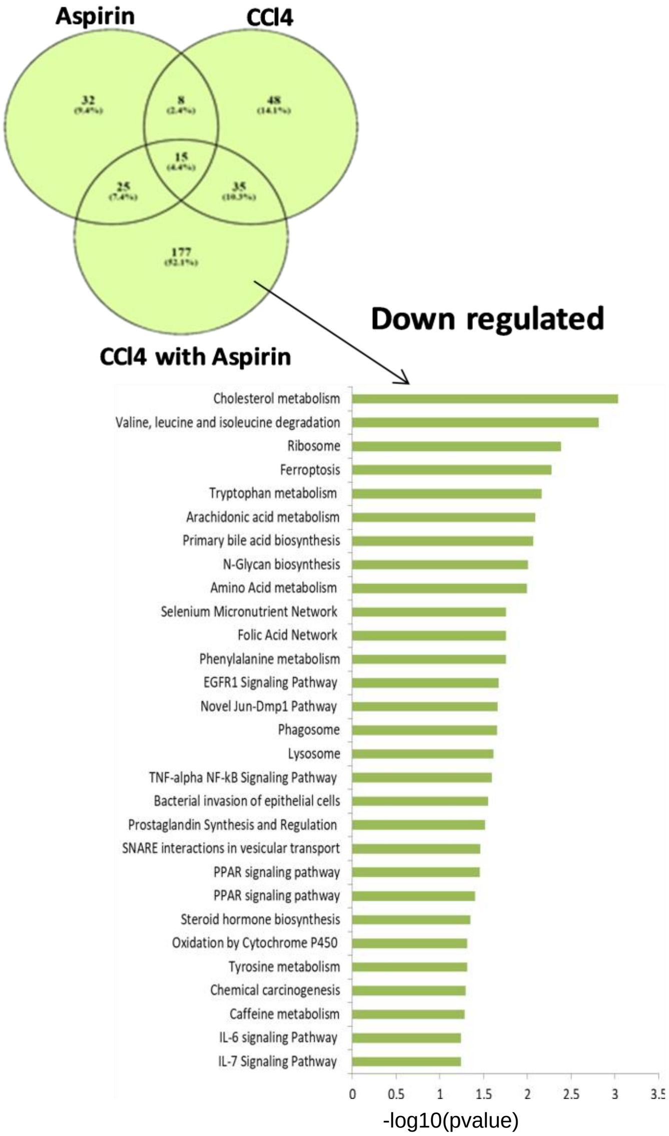

**Figure S-3B:** A total of 177 downregulated proteins in CCl4+Aspirin were associated to pathway linked to cholesterol metabolism, TNF-α signalling, arachidonic acid and tyrosine metabolism.

**Supplementary Figure 4:** RT-PCR expression analysis of oxidative stress related genes.

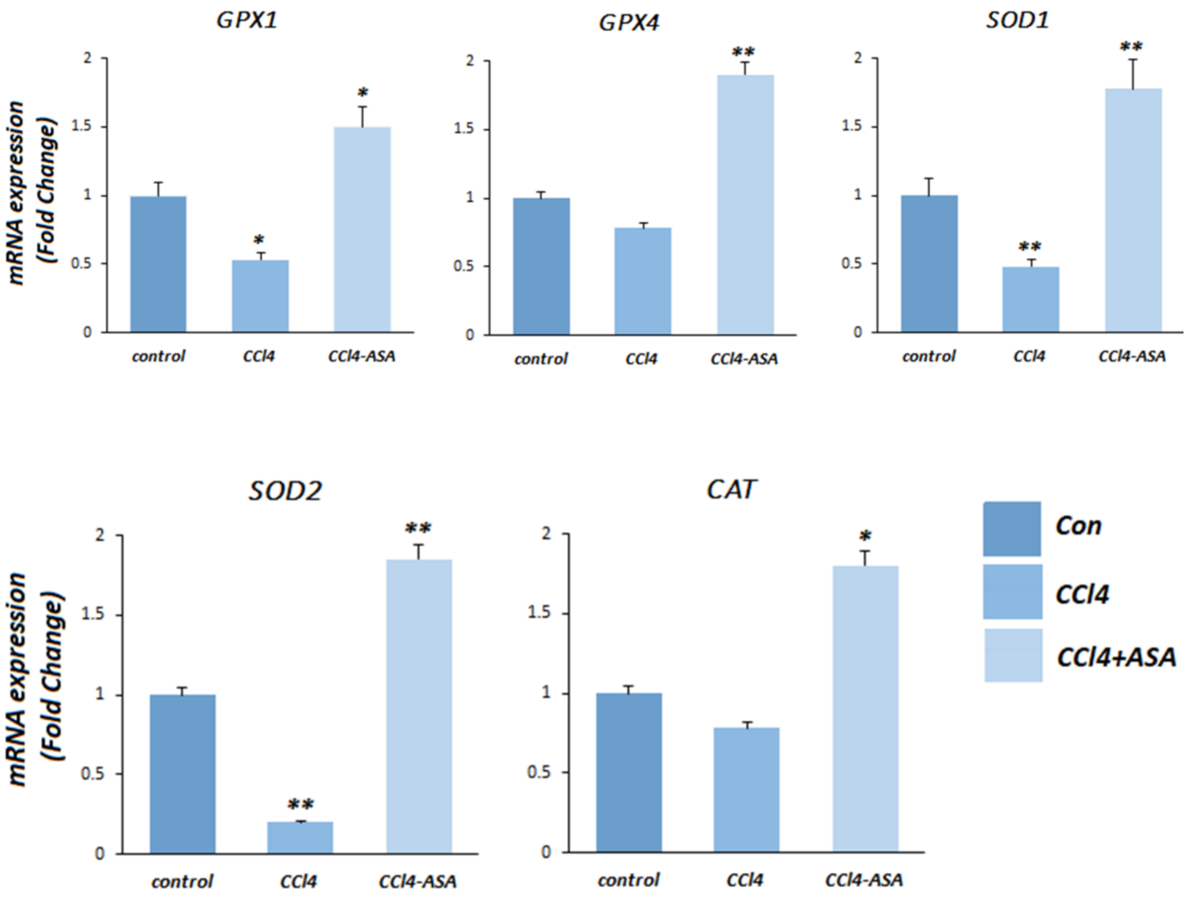

**Figure S-4:** The mRNA expression changes (qPCR analysis) showed a downregulation of the classical genes that protect against oxidative stress (Gpx1, Gpx4, Sod1, Sod2 and Cat) in aspirin treated as compared to control.Abbreviations: Con; Control, ASA; Aspirin alone, $CCl_4$; carbon tetra chloride group, $CCl_4$+ASA; carbon tetra chloride plus aspirinAsterisks denote statistical significance as:(*$p < 0.05$, **$p < 0.005$).

**Explanation for supplementary figure 5:**

**Aspirin treatment enhances metabolic function in murine model of liver fibrosis**

The reduction of fibrosis post-aspirin treatment could be attributed to a change in the fibrotic liver metabotype. We evaluated the liver metabolome across the groups in the $CCl_4$ model of fibrosis with and without aspirin treatment (Figure-3A). Metabolomic analysis of the liver identified differentially expressed metabolites (DEMs) in ASA (26 metabolites up- and 25 downregulated), $CCl_4$ (32 metabolites up- and 32 downregulated) and $CCl_4$+ASA groups (50 metabolites up- and 42 downregulated) as compared to controls (Figure-3B, Supplementary-Table-5). Partial least square discriminant analysis (PLS-DA) and unsupervised clustering analysis showed a clear distinction between the groups (Figure-3C, 3D). In addition, venn analysis of all the upregulated metabolites identified 10 (13.9%) metabolites unique to aspirin group, 7 (9.7%) metabolites unique to $CCl_4$ group and 25 (34.7%) metabolites unique to $CCl_4$+ASA group (Figure-3E). Metabolites unique to $CCl_4$+ASA group were linked to galactose metabolism, mitochondria electron transport chain, ubiquinone biosynthesis, pyrimidine metabolism, tyrosine metabolism, urea cycle, and others. Similarly, venn analysis of the down regulated metabolites identified 10 (15.2%) metabolites unique in the aspirin treatment ($CCl_4$+ASA) group, 14 (21.2%) metabolites unique to $CCl_4$ group and 19 (28.8%) unique to aspirin and $CCl_4$ treatment group. Metabolites unique to aspirin and $CCl_4$ group highlighted significant down regulation of butanoate metabolism, taurine/hypotaurine metabolism, phosphatidylethanolamine biosynthesis pathways and others (Figure-3F).These findings suggest that aspirin induces energy metabolism, amino acid metabolism and reduces butanoate metabolism and phosphatidylethanolamine biosynthesis linked to microbiome metabolism.

**Supplementary Figure 5:** Metabolome analysis across the groups in the $CCl_4$ model of fibrosis with and without aspirin treatment

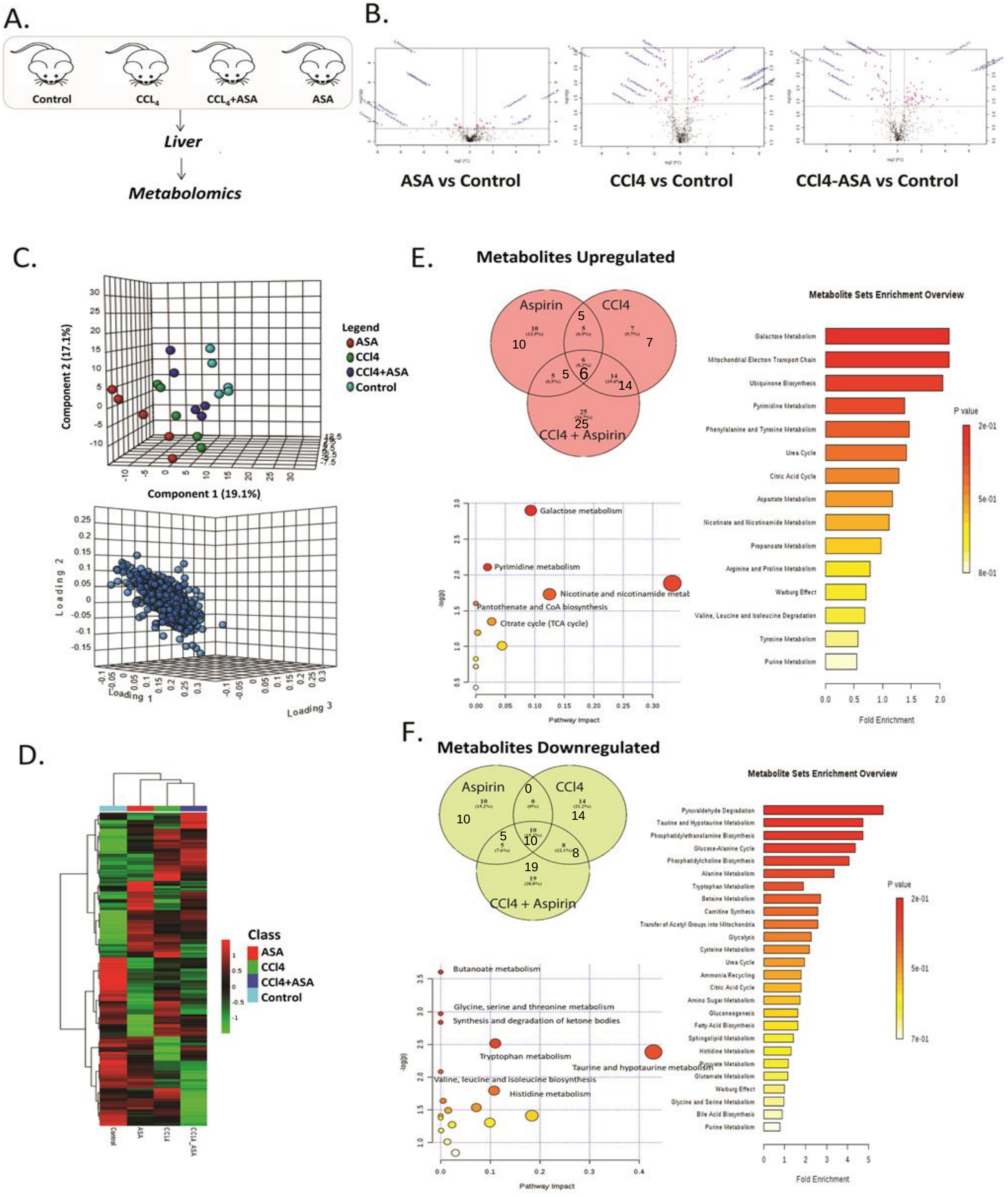

**Figure-S5: Liver metabolome in murine models.** A: Overview of metabolomic study performed across the mice groups. B: Differentially expressed metabolites were visualized by volcano plots C: Three dimensional PLS-DA score plots of the aspirin (ASA), $CCl_4$, $CCl_4$+ASA, and control, each dot represents one liver sample, projected onto first (horizontal axis) and second (vertical axis) PLS-DA variables. Mice groups are shown in different colours (left panel). Three dimensional loading plots of ASA, $CCl_4$, $CCl_4$+ASA, and con combined analysis both in the negative ion mode (ESI–) and positive mode (ESI+). D: Heatmap analysis of the differential metabolites in different groups . Red and green colours indicate values above and below the mean, respectively. Black indicates are values close to the mean. Venn diagram showing the common and unique metabolites between the ASA alone and $CCl_4$ model vs. $CCl_4$+ASA. E: Significantly upregulated metabolites are highlighted in red venny, (log2-fold change >1.5; p-<0.05). F: Significantly downregulated metabolites are highlighted in green venny, (log2-fold change <1.5; p- <0.05). The pathways associated with the up and down-regulated metabolites are represented in bar chart.

**Supplementary Figure 6:** Analysis of common up- and downregulated pathways in low and high fibrosis mice group.

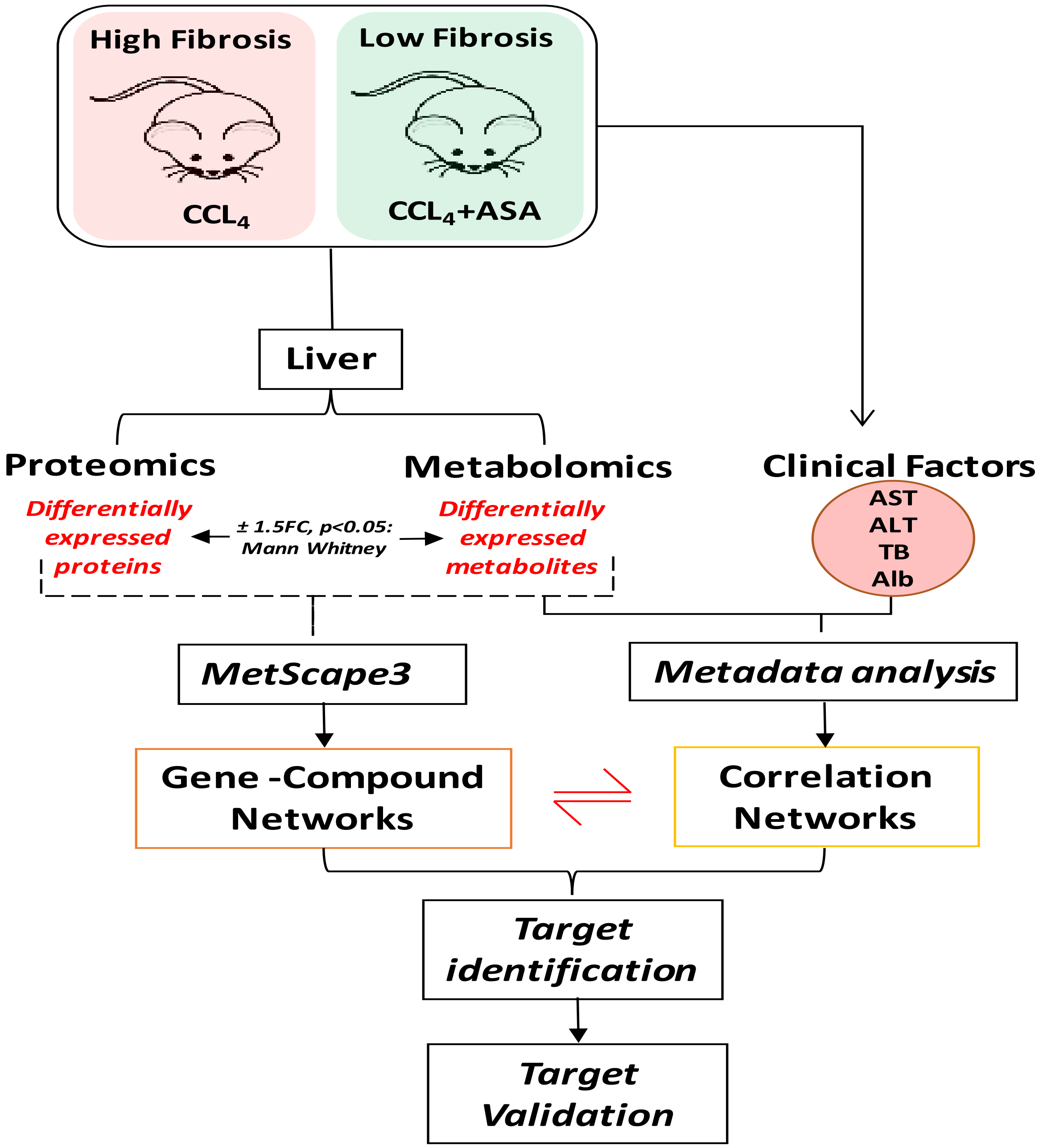

.

**Figure S-6A**: The pathways linked to up-regulated proteins/metabolites in $CCl_4$-ASA

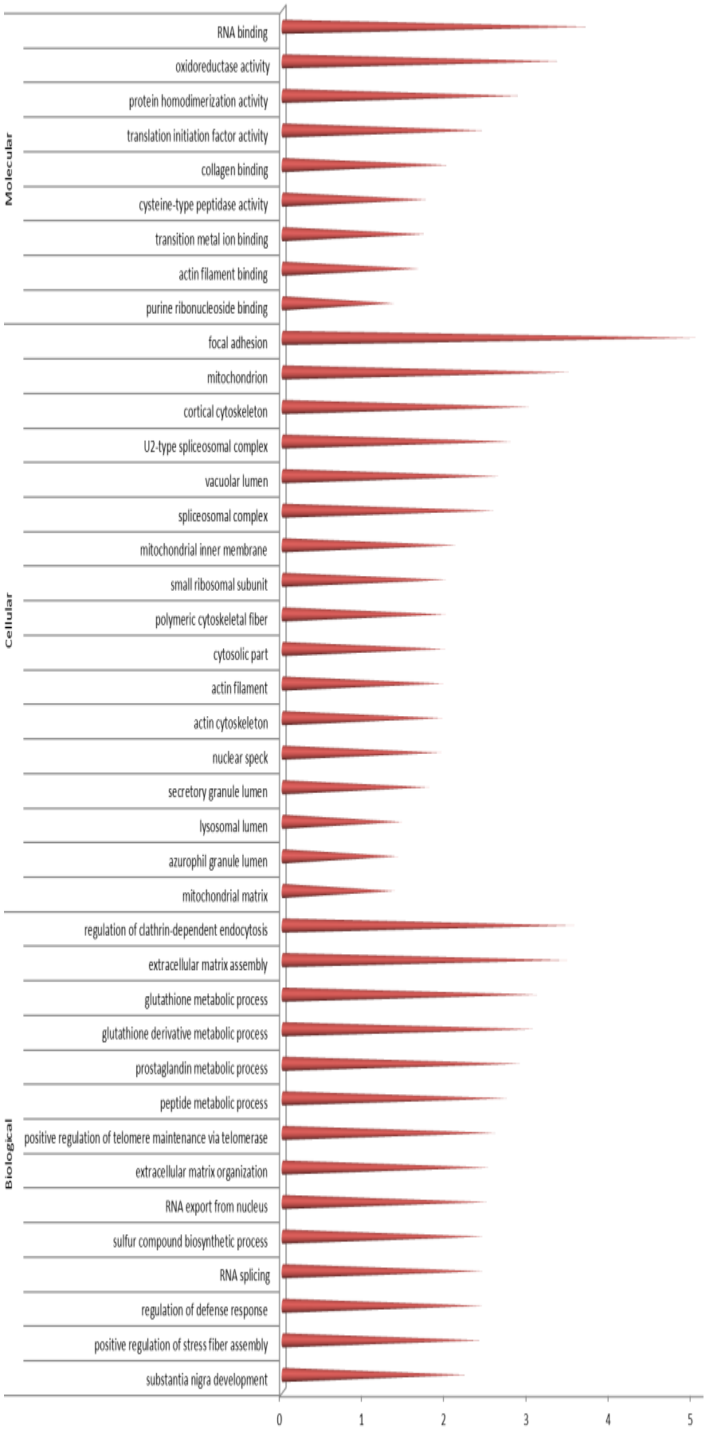

**Figure S-6A**: The up-regulated proteins/metabolites pathways in CCl4-ASA group were associated to liver function such xenobiotic metabolism, TCA cycle, pentose phosphate and amino acid metabolism (energy metabolism) whereas downregulated pathways were associated to urea cycle, bile acid biosynthesis, arachidonic acid, prostaglandin and leukotriene metabolism (inflammation)

**Figure S-6A**: The pathways linked to down-regulated proteins/metabolites in $CCl_4$-ASA

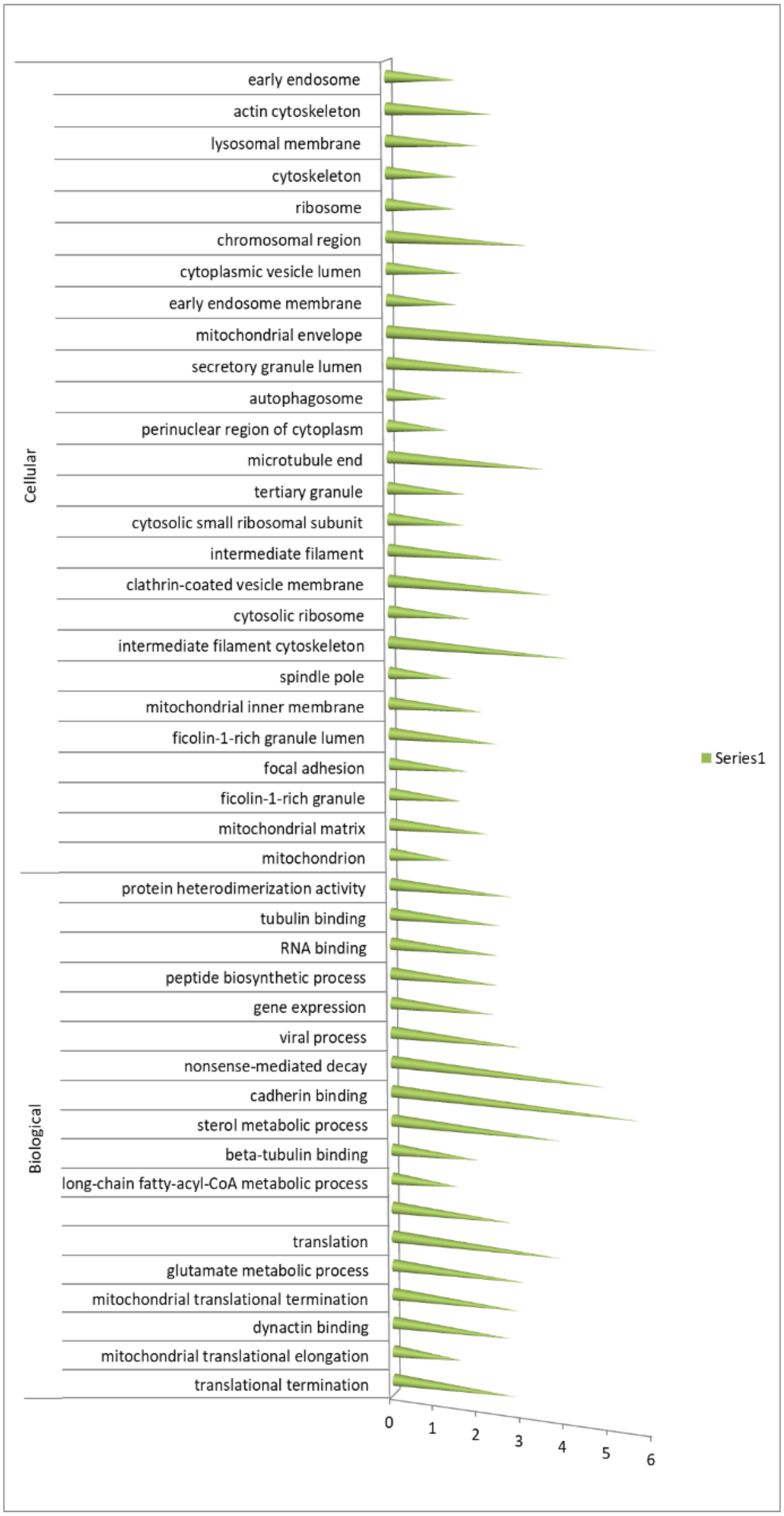

**Supplementary Figure 6B:** Most perturbed 39 pathways analysis in low and high fibrosis.

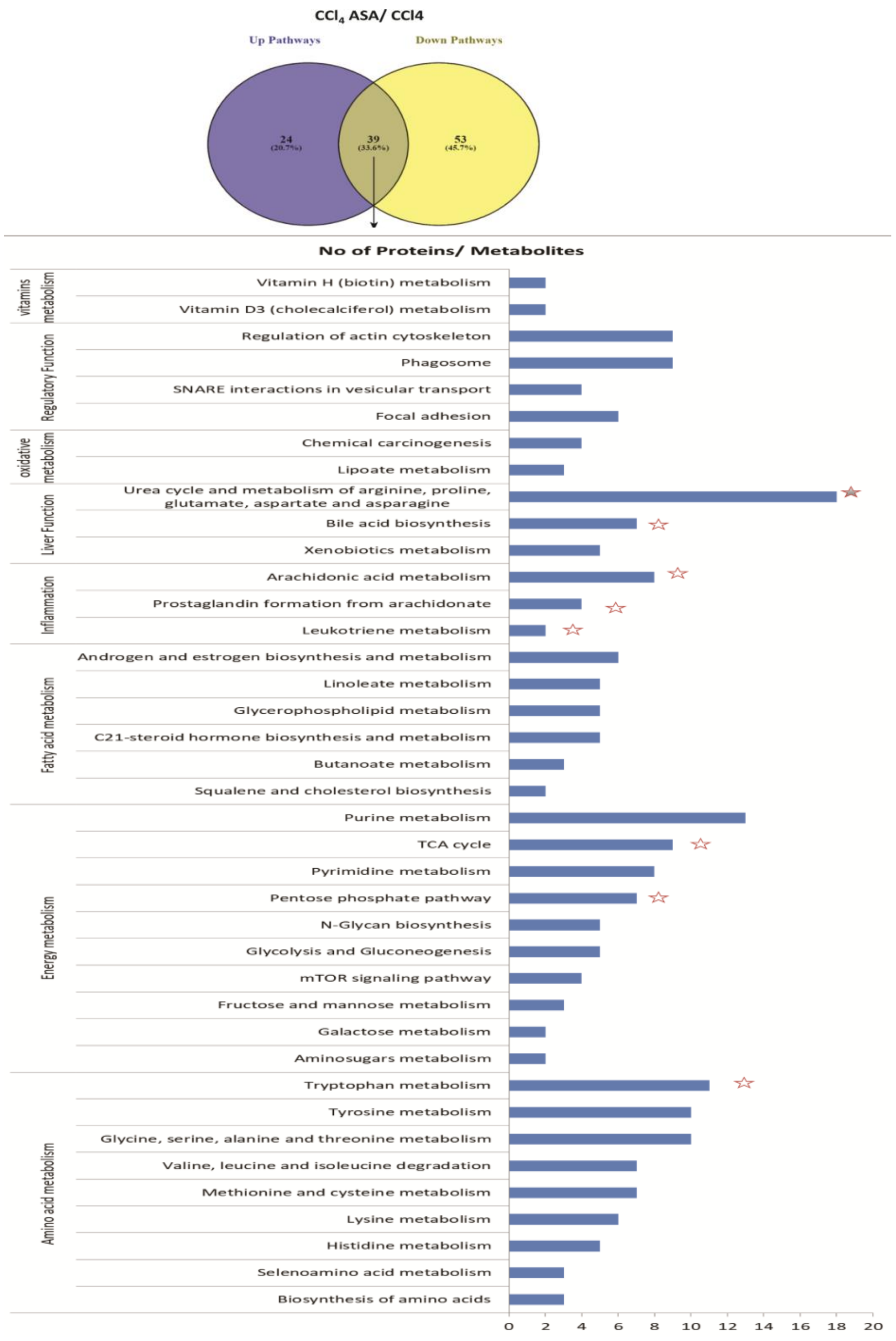

**Figure S-6B**: The most perturbed 39 pathways (33.6%; which involves both up and down-regulated proteins/metabolites) in CCl4-ASA group were associated to liver function (urea cycle, bile acid biosynthesis, xenobiotic metabolism), arachidonic acid, prostaglandin and leukotriene metabolism (inflammation), TCA cycle, pentose phosphate and amino acid metabolism (energy metabolism).

**Supplementary Figure 7A:** Integration analysis of proteins/metabolites associated with energy metabolism.

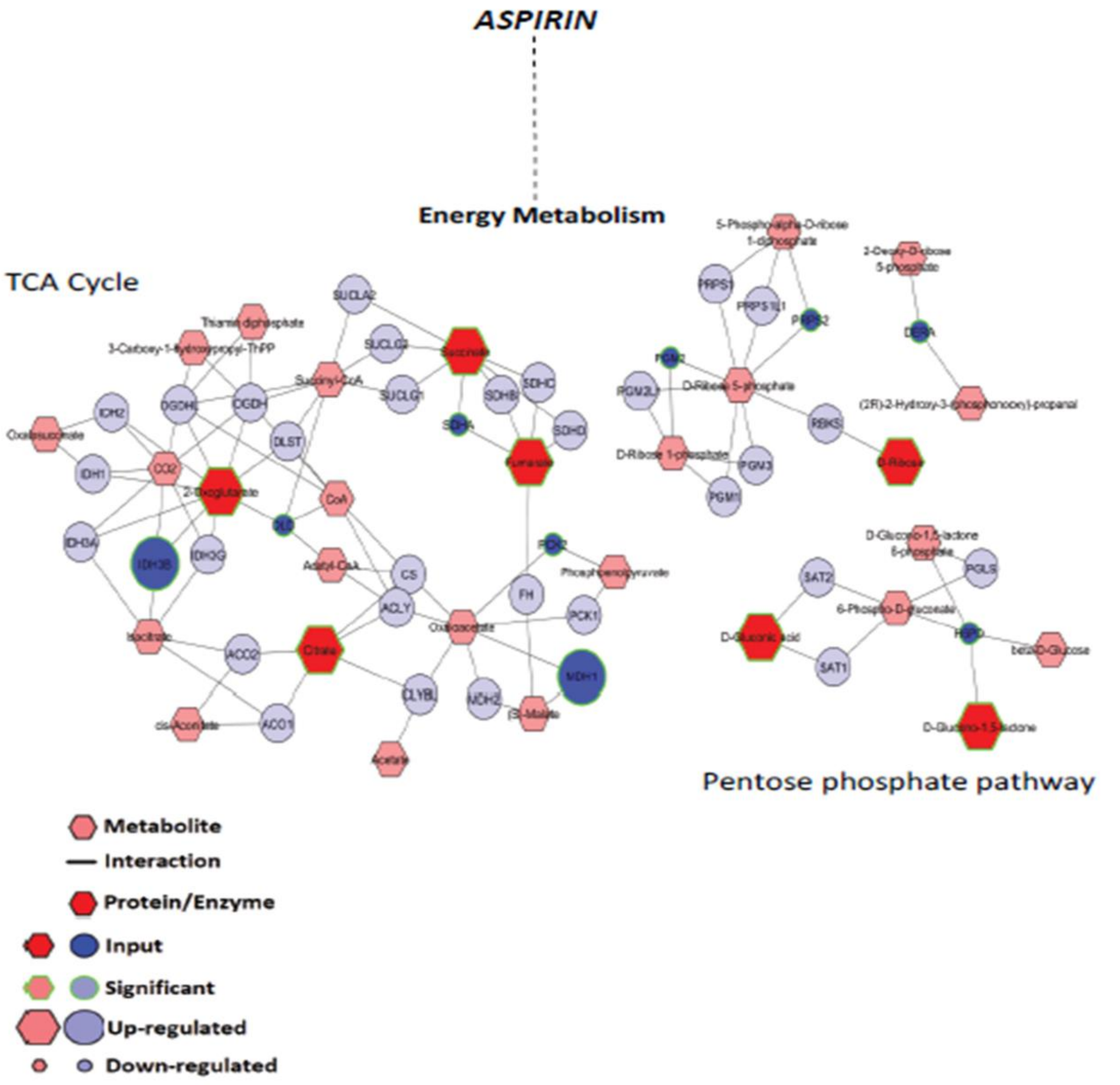

**Figure S-7A:** Interactome of metabolites (hexagons), interactions (dashed lines) and proteins/enzymes (circles) associated with tryptophan metabolism. Analysis shows a significant increase in metabolites associated with energy metabolism in fibrotic livers post-aspirin treatment.

**Supplementary Figure 7B:** Integration analysis of proteins/metabolites associated with tryptophan metabolism.

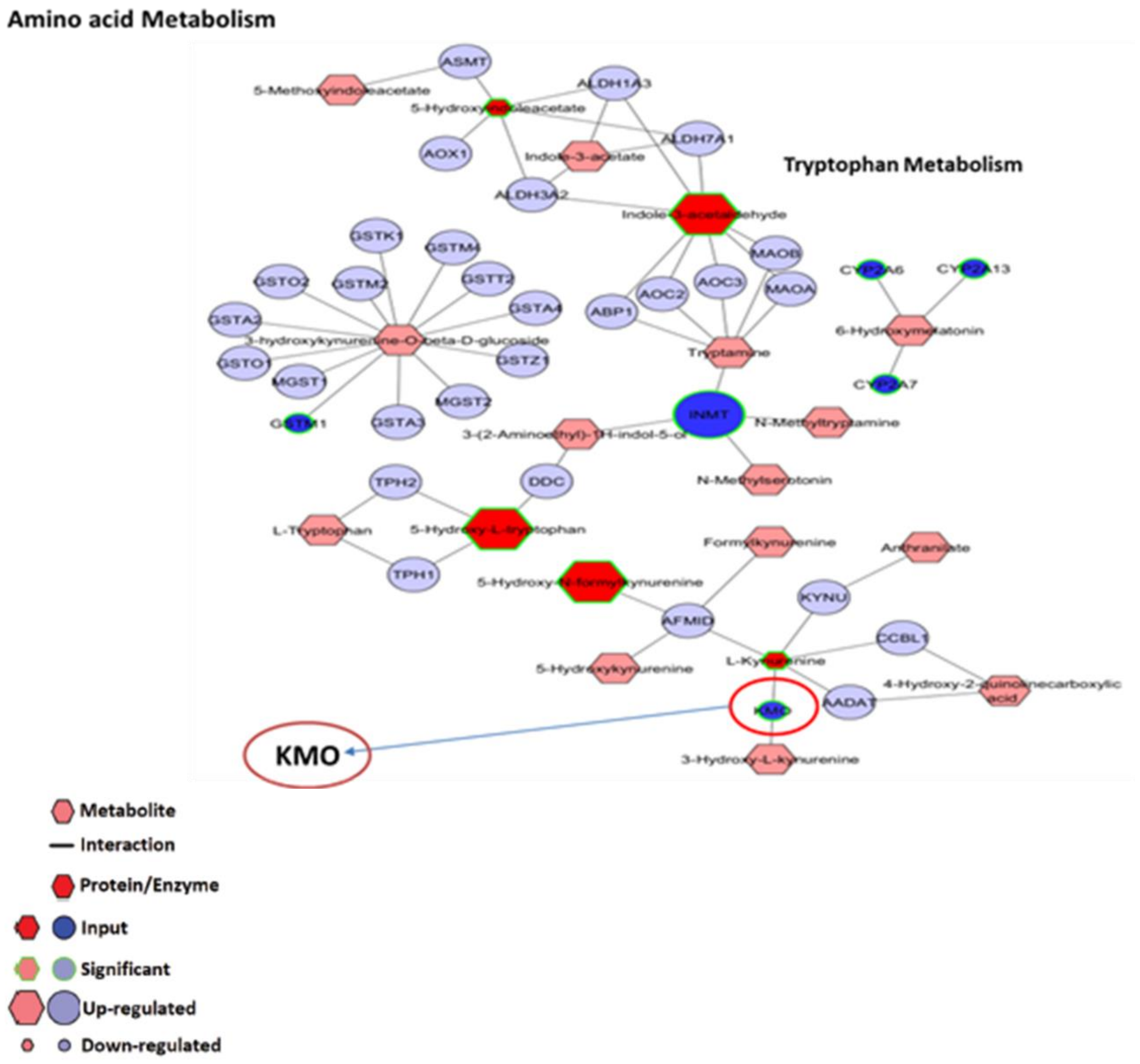

**Figure S-7B:** Interactome of metabolites (hexagons), interactions (dashed lines) and proteins/enzymes (circles) associated with tryptophan metabolism. Analysis shows a significant reduction of kynurenine-3-monoxygenase (KMO) was seen in fibrotic livers post-aspirin treatment.

**Supplementary Figure 8A:** Modulation of hepatic microbiome with aspirin treatment.

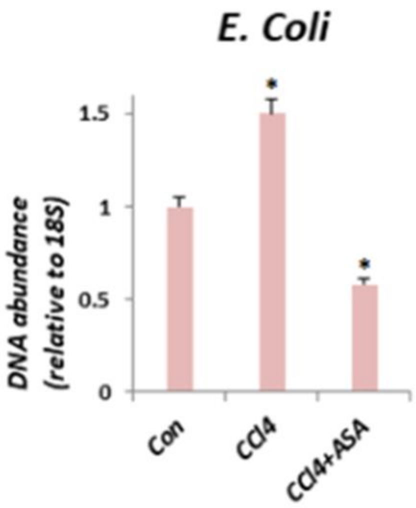

**Figure S-8A:** Pathogenic microorganism analysis in the liver in $CCl_4$- and $CCl_4$-ASA mouse model. Presence of E.coli DNA in the liver tissue was assessed by qPCR. Scatter plots show the individual values (dots), group medians (line)and standard deviations (whiskers). Abbreviations: Con; Control, ASA; Aspirin alone, $CCl_4$; carbon tetra chloride group, $CCl_4$+ASA; carbon tetra chloride plus aspirinAsterisks denote statistical significance as: (*$p < 0.05$, **$p < 0.005$).

**Supplementary Figure 8B:** Correlation of hepatic microbiome with key target proteins post aspirin treatment.

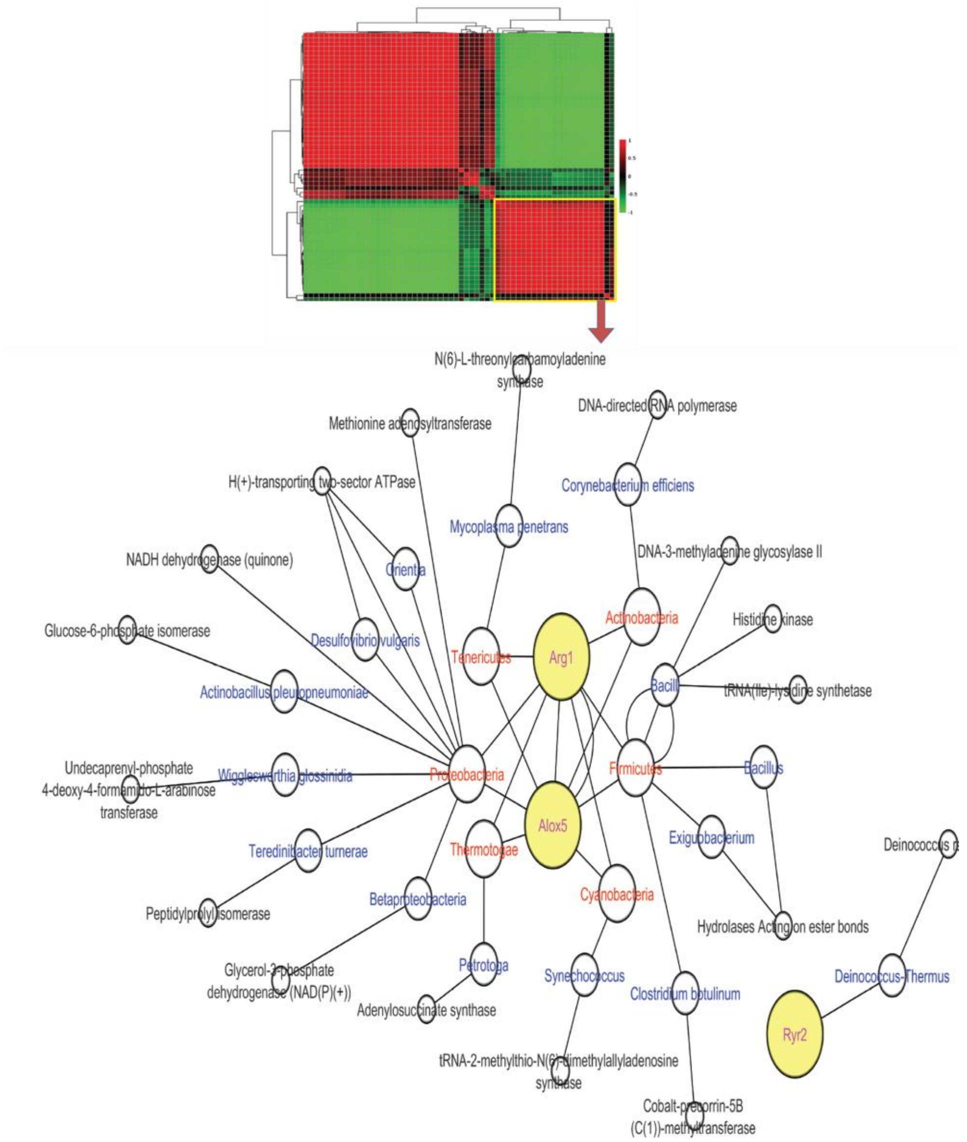

**Figure S-8B:** Cross correlation analysis between identified aspirin regulated key target protein and taxonomic features with associated enzymatic functionality. Target proteins ARG1, ALOX5 and RYR2 (encircled yellow), black lines (correlations, $r^2>0.7; p<0.05$), bacterial species (red circle), sub phylum bacterial species (blue circle) and enzyme function (outer black circle).

**Supplementary Figure 9:** Modulation of autophagy related proteins with aspirin administration.

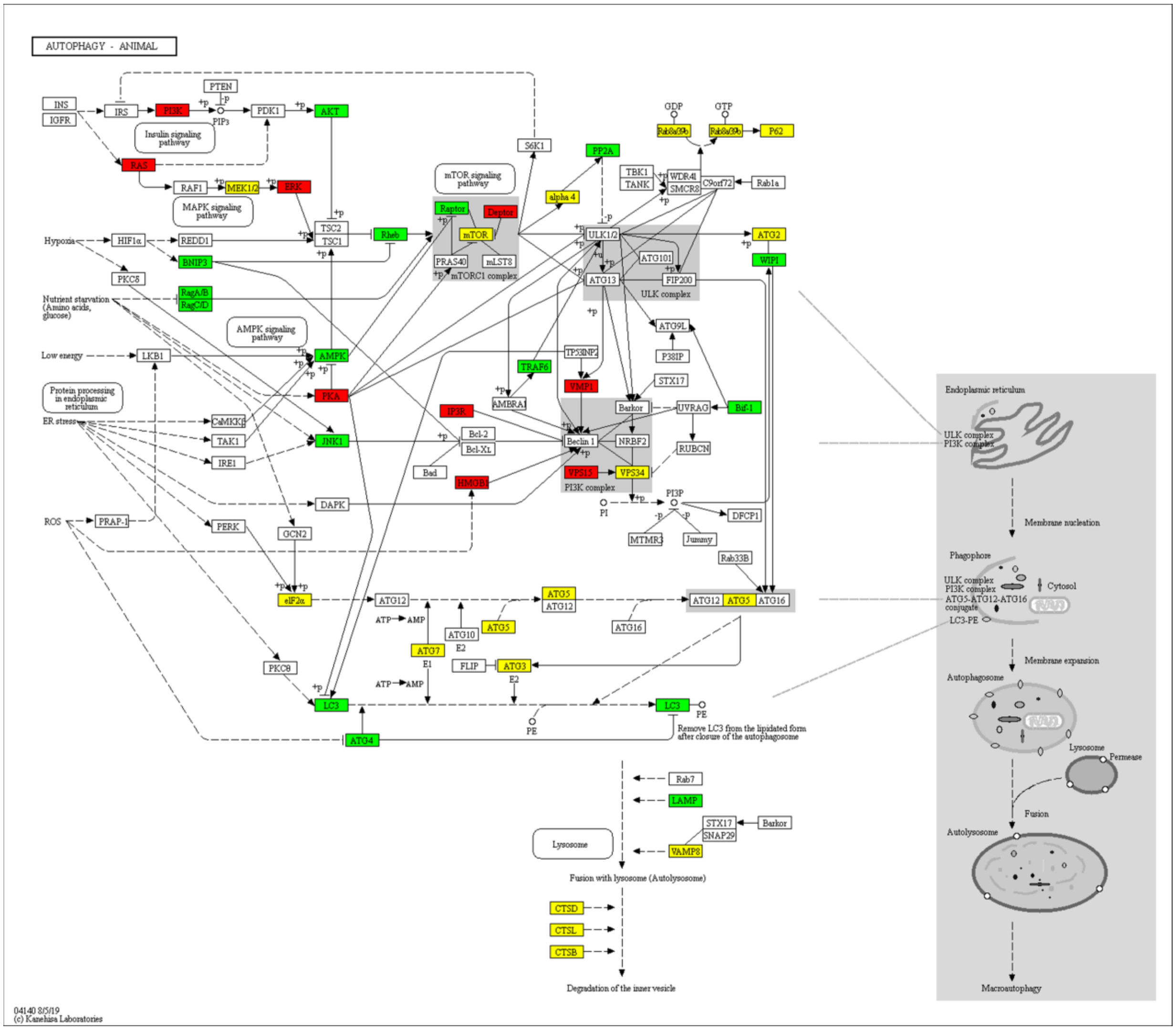

**Figure S-9:** KEGG analysis of autophagy pathway: Proteins up-regulated (Red), proteins identified but not significant (Yellow) and proteins down-regulated (Green) associated with autophagy. Analysis shows a significant increase in autophagy related proteins were observed in fibrotic livers post-aspirin treatment.

**References:**

1. Ghiassi-Nejad Z, Hernandez-Gea V, Woodrell C, Lang UE, Dumic K, Kwong A, Friedman SL. Reduced hepatic stellate cell expression of Kruppel-like factor 6 tumor suppressor isoforms amplifies fibrosis during acute and chronic rodent liver injury. Hepatology 2013;57:786-796.
2. Zhou Y, Peng H, Liu Z, Zhang KK, Jendrusch C, Drake M, Hao Y, et al. Sex-associated preventive effects of low-dose aspirin on obesity and non-alcoholic fatty liver disease in mouse offspring with over-nutrition in utero. Lab Invest 2019;99:244-259.
3. Tancevski I, Wehinger A, Schgoer W, Eller P, Cuzzocrea S, Foeger B, Patsch JR, et al. Aspirin regulates expression and function of scavenger receptor-BI in macrophages: studies in primary human macrophages and in mice. FASEB J 2006;20:1328-1335.
4. Bedossa P, Poynard T. An algorithm for the grading of activity in chronic hepatitis C. The METAVIR Cooperative Study Group. Hepatology 1996;24:289-293.
5. Malehmir M, Pfister D, Gallage S, Szydlowska M, Inverso D, Kotsiliti E, Leone V, et al. Platelet GPIbalpha is a mediator and potential interventional target for NASH and subsequent liver cancer. Nat Med 2019;25:641-655.
6. Song YN, Dong S, Wei B, Liu P, Zhang YY, Su SB. Metabolomic mechanisms of gypenoside against liver fibrosis in rats: An integrative analysis of proteomics and metabolomics data. PLoS One 2017;12:e0173598.
7. Starkel P, Leclercq S, de Timary P, Schnabl B. Intestinal dysbiosis and permeability: the yin and yang in alcohol dependence and alcoholic liver disease. Clin Sci (Lond) 2018;132:199-212.
8. Basu S, Duren W, Evans CR, Burant CF, Michailidis G, Karnovsky A. Sparse network modeling and metscape-based visualization methods for the analysis of large-scale metabolomics data. Bioinformatics 2017;33:1545-1553.
9. Chong J, Soufan O, Li C, Caraus I, Li S, Bourque G, Wishart DS, et al. MetaboAnalyst 4.0: towards more transparent and integrative metabolomics analysis. Nucleic Acids Res 2018;46:W486-W494.
10. Zhang X, Deeke SA, Ning Z, Starr AE, Butcher J, Li J, Mayne J, et al. Metaproteomics reveals associations between microbiome and intestinal extracellular vesicle proteins in pediatric inflammatory bowel disease. Nat Commun 2018;9:2873.
11. Soliman EM, Rodrigues MA, Gomes DA, Sheung N, Yu J, Amaya MJ, Nathanson MH, et al. Intracellular calcium signals regulate growth of hepatic stellate cells via specific effects on cell cycle progression. Cell Calcium 2009;45:284-292.
12. Langer DA, Das A, Semela D, Kang-Decker N, Hendrickson H, Bronk SF, Katusic ZS, et al. Nitric oxide promotes caspase-independent hepatic stellate cell apoptosis through the generation of reactive oxygen species. Hepatology 2008;47:1983-1993.
13. Furuya S, Argemi J, Uehara T, Katou Y, Fouts DE, Schnabl B, Dubuquoy L, et al. A Novel Mouse Model of Acute-on-Chronic Cholestatic Alcoholic Liver Disease: A Systems Biology Comparison With Human Alcoholic Hepatitis. Alcohol Clin Exp Res 2020;44:87-101.

**Supplementary Table -1: Liver biochemical parameters in control and experimental mice groups.**

| Table 1: Liver biochemical parameters in control and experimental mice groups. | | | | |
|---|---|---|---|---|
| **Treatment** | **ALT(U/L)** | **AST(U/L)** | **Alb(g/dl)** | **TB(mg/dl)** |
| Negative Control | 22.6±1.2 | 79.5±1.4 | 4.3±0.2 | 2.2±0.3 |
| CCl4 | 74.2±2.1** | 150.7±11.5** | 2.9±0.2** | 4.9±0.4** |
| $CCl_4$+ASA (5mg/kg) | 46.0±4.3## | 88.1±2.4## | 3.9±0.2## | 3.7±0.2## |
| The result are showed as mean ±SD; ** p<0.01 vs. negative control ; ## p<0.01 vs. model control | | | | |

**Supplementary Table-2: Primers used in quantitative real-time PCR (mouse).**

| Table 2: Primers used in quantitative real-time PCR (mouse) | | | | |
|---|---|---|---|---|
| **S. no** | **Genes** | **(5'-3')** | **Sequence** | **Refrences** |
| **1** | **α-SMA** | **Forward** | TCCAGCTATGTGTGAAGAGGA | |
| | | **Reverse** | ACGATGGATGGGAAAACAGC | |
| **2** | **PDGFR-β** | **Forward** | GGATGCTTTTGGAGTGAGGA | |
| | | **Reverse** | AAACGGACAGCAGTAGAAGG | |
| **3** | **TGF-β1** | **Forward** | ATCGACATGGAGCTGGTGAA | |
| | | **Reverse** | TGTTGTACAAAGCGAGCACC | |
| **4** | **Col1a** | **Forward** | GCTGCCTTTTCTGCTCCTTT | |
| | | **Reverse** | TGGGACATTTCAGCATTGCC | |
| **5** | **Tgfbr2** | **Forward** | ACATTCAAGCCCCACTCTGA | |
| | | **Reverse** | TGGGGAACGGTCTTCTTTGA | |
| **6** | **TNF-α** | **Forward** | TCAGAAACACAAGATGCTGGG | |
| | | **Reverse** | ACCACTCTCCCTTTGCAGAA | |
| **7** | **IL10** | **Forward** | AGAAACTCATGGGTCTTGGGA | |
| | | **Reverse** | AGTTTTCAGGGATGAAGCGG | |
| **8** | **Gapdh** | **Forward** | GTGGACCTCATGGCCTACAT | |
| | | **Reverse** | TGTGAGGGAGATGCTCAGTG | |
| **9** | **Escherichia coli** | **Forward** | GTTAATACCTTTGCTCATTGA | Malinen et al., 2005 |
| | | **Reverse** | ACCAGGGTATCTAATCCTGTT | |

**Reference:**

01. Malinen E, Rinttila T, Kajander K, Matto J, Kassinen A, Krogius L, Saarela M, Korpela R, Palva A (2005) Analysis of the fecal microbiota of irritable bowel syndrome patients and healthy controls with real-time PCR. Am J Gastroenterol 100(2):373-82.

### Supplementary Table-3. List of identified protein groups across all mice liver samples and their quantification (normalized Log2 intensities)

**Supplementary Table 3. List of identified protein groups across all mice liver samples and their quantification (normalized Log2 intensities)**

| Sr. no | Gene Symbol | UP/Down all groups vs. control | Accession | # Peptides | # PSMs | # Unique Peptides | # AAs | MW [kDa] | calc. pI | Coverage [%] (by Search Engine): Sequest HT | FC ASA/CONTROL | P-value | FC CCl4/CONTROL | P-value | FC ASA-CCl4/CONTROL | P-value | FC CCl4/ ASA-CCl4 | P-value |
|---|---|---|---|---|---|---|---|---|---|---|---|---|---|---|---|---|---|---|
| 1 | **Nrf1** | Unchanged | A0A0J9YU17 | 1 | 1 | 1 | 38 | 4.3 | 5.45 | 100 | 1.000 | 0.600 | 1.300 | 0.800 | 1.100 | 0.900 | 1.300 | 0.612 |
| 2 | **Fabp1** | Unchanged | P12710 | 31 | 1917 | 31 | 127 | 14.2 | 8.56 | 98 | 1.127 | 0.524 | 1.252 | 0.547 | 0.872 | 0.315 | 1.436 | 0.349 |
| 3 | **Rplp2** | Unchanged | P99027 | 12 | 369 | 12 | 115 | 11.6 | 4.54 | 97 | 0.962 | 0.816 | 0.998 | 0.992 | 0.901 | 0.606 | 1.107 | 0.731 |
| 4 | **Acaa2** | Unchanged | Q8BWT1 | 42 | 1607 | 42 | 397 | 41.8 | 8.09 | 97 | 1.129 | 0.346 | 0.934 | 0.680 | 1.098 | 0.585 | 0.851 | 0.435 |
| 5 | **Urah** | Unchanged | Q9CRB3 | 10 | 113 | 6 | 118 | 13.6 | 7.87 | 96 | 0.818 | 0.285 | 0.673 | 0.111 | 1.199 | 0.417 | 0.561 | 0.053 |
| 6 | **Arg1** | Unchanged | Q61176 | 33 | 1329 | 33 | 323 | 34.8 | 7.01 | 96 | 0.885 | 0.576 | 1.032 | 0.869 | 0.812 | 0.309 | 1.271 | 0.061 |
| 7 | **Gchfr** | Unchanged | P99025 | 6 | 162 | 6 | 84 | 9.6 | 6.27 | 83 | 1.154 | 0.558 | 1.212 | 0.528 | 1.419 | 0.182 | 0.854 | 0.556 |
| 8 | **Rida** | Unchanged | P52760 | 15 | 517 | 15 | 135 | 14.2 | 8.68 | 95 | 1.237 | 0.085 | 1.111 | 0.637 | 1.246 | 0.323 | 0.892 | 0.644 |
| 9 | **Aldh2** | Unchanged | A0A0G2JFQ0 | 21 | 288 | 1 | 229 | 25.5 | 8.18 | 94 | 0.568 | 0.379 | 0.555 | 0.364 | 0.572 | 0.394 | 0.969 | 0.959 |
| 10 | **Ddt** | Unchanged | O35215 | 12 | 268 | 12 | 118 | 13.1 | 6.54 | 94 | 1.049 | 0.862 | 1.098 | 0.816 | 1.074 | 0.784 | 1.023 | 0.945 |
| 11 | **Mup14** | Unchanged | B8JI96 | 18 | 431 | 3 | 150 | 17.4 | 4.77 | 94 | 0.562 | 0.298 | 0.450 | 0.223 | 0.461 | 0.212 | 0.977 | 0.959 |
| 12 | **Actb** | Unchanged | P60710 | 41 | 1596 | 13 | 375 | 41.7 | 5.48 | 94 | 0.975 | 0.794 | 0.783 | 0.027 | 1.195 | 0.200 | 0.655 | 0.012 |
| 13 | **Sod1** | Unchanged | P08228 | 11 | 361 | 11 | 154 | 15.9 | 6.51 | 93 | 0.825 | 0.486 | 0.973 | 0.947 | 0.625 | 0.184 | 1.556 | 0.368 |
| 14 | **Cyp2d9** | Unchanged | P11714 | 32 | 512 | 15 | 504 | 56.9 | 6.21 | 93 | 1.235 | 0.600 | 1.351 | 0.570 | 1.451 | 0.247 | 0.931 | 0.883 |
| 15 | **Ephx2** | Unchanged | P34914 | 45 | 1476 | 45 | 554 | 62.5 | 6.19 | 92 | 1.210 | 0.272 | 0.922 | 0.759 | 1.185 | 0.301 | 0.778 | 0.333 |
| 16 | **Fah** | Unchanged | D3Z2R9 | 11 | 167 | 1 | 141 | 15.6 | 9.98 | 92 | 0.108 | 0.083 | 0.004 | 0.058 | 0.010 | 0.059 | 0.403 | 0.085 |
| 17 | **Prdx1** | Unchanged | P35700 | 28 | 878 | 26 | 199 | 22.2 | 8.12 | 92 | 0.820 | 0.398 | 1.109 | 0.705 | 1.069 | 0.799 | 1.037 | 0.905 |
| 18 | **Ahcy** | Unchanged | P50247 | 38 | 1539 | 36 | 432 | 47.7 | 6.54 | 92 | 1.123 | 0.562 | 0.718 | 0.164 | 0.923 | 0.680 | 0.778 | 0.112 |
| 19 | **Fbp1** | UP | Q9QXD6 | 35 | 1432 | 35 | 338 | 36.9 | 6.57 | 91 | 1.312 | 0.101 | 1.523 | 0.007 | 1.751 | 0.005 | 0.870 | 0.255 |
| 20 | **Nqo2** | Unchanged | Q9JI75 | 21 | 162 | 21 | 231 | 26.2 | 7.01 | 91 | 0.818 | 0.222 | 0.821 | 0.169 | 0.931 | 0.630 | 0.882 | 0.378 |
| 21 | **Gapdh** | Unchanged | P16858 | 32 | 1599 | 6 | 333 | 35.8 | 8.25 | 91 | 0.782 | 0.220 | 0.725 | 0.179 | 1.027 | 0.863 | 0.706 | 0.177 |
| 22 | **Hbb-bs** | Down | A8DUK4 | 27 | 804 | 4 | 147 | 15.7 | 7.69 | 91 | 0.658 | 0.122 | 0.544 | 0.045 | 0.606 | 0.105 | 0.898 | 0.674 |
| 23 | **Aldh1l1** | Unchanged | Q8R0Y6 | 84 | 2309 | 78 | 902 | 98.6 | 5.91 | 91 | 1.043 | 0.706 | 0.708 | 0.096 | 0.901 | 0.328 | 0.786 | 0.250 |
| 24 | **Gstp1** | Unchanged | P19157 | 16 | 1350 | 15 | 210 | 23.6 | 7.87 | 91 | 0.875 | 0.492 | 1.032 | 0.933 | 1.093 | 0.583 | 0.944 | 0.874 |
| 25 | **Actg1** | Unchanged | F8WGM8 | 11 | 275 | 1 | 105 | 11.9 | 7.21 | 90 | 1.391 | 0.647 | 1.471 | 0.571 | 1.811 | 0.426 | 0.812 | 0.761 |
| 26 | **Rdh7** | Unchanged | O88451 | 29 | 425 | 16 | 316 | 35.6 | 8.09 | 90 | 0.983 | 0.905 | 0.772 | 0.343 | 0.930 | 0.660 | 0.830 | 0.526 |
| 27 | **Ldha** | Unchanged | P06151 | 34 | 1489 | 32 | 332 | 36.5 | 7.74 | 90 | 0.700 | 0.032 | 1.125 | 0.365 | 1.026 | 0.858 | 1.096 | 0.438 |
| 28 | **Prdx2** | Unchanged | Q61171 | 17 | 300 | 17 | 198 | 21.8 | 5.41 | 89 | 0.963 | 0.676 | 1.087 | 0.385 | 1.035 | 0.768 | 1.050 | 0.692 |
| 29 | **Gm4952** | Unchanged | Q5FW57 | 23 | 379 | 22 | 296 | 34.1 | 7.8 | 89 | 1.295 | 0.082 | 1.096 | 0.704 | 1.123 | 0.415 | 0.976 | 0.917 |
| 30 | **Hsd17b10** | Unchanged | Q99N15 | 21 | 283 | 21 | 261 | 27.3 | 8.76 | 89 | 1.000 | 1.000 | 0.625 | 0.094 | 0.636 | 0.074 | 0.982 | 0.948 |

| | | | | | | | | | | | | | | | | | | |
|---|---|---|---|---|---|---|---|---|---|---|---|---|---|---|---|---|---|---|
| 31 | **Ca3** | Unchanged | P16015 | 23 | 1117 | 23 | 260 | 29.3 | 7.37 | 89 | 1.522 | 0.055 | 0.782 | 0.467 | 0.884 | 0.567 | 0.885 | 0.670 |
| 32 | **Acox1** | UP | Q9R0H0 | 49 | 1458 | 12 | 661 | 74.6 | 8.48 | 89 | 1.172 | 0.277 | 1.784 | 0.030 | 1.591 | 0.003 | 1.121 | 0.540 |
| 33 | **Rgn** | Unchanged | Q64374 | 29 | 1035 | 29 | 299 | 33.4 | 5.34 | 89 | 1.034 | 0.876 | 0.841 | 0.556 | 0.946 | 0.793 | 0.889 | 0.631 |
| 34 | **Adh1** | Unchanged | P00329 | 29 | 1420 | 29 | 375 | 39.7 | 8.1 | 89 | 1.120 | 0.746 | 1.327 | 0.608 | 1.242 | 0.617 | 1.069 | 0.910 |
| 35 | **Gstm1** | Unchanged | P10649 | 33 | 1054 | 11 | 218 | 26 | 7.94 | 81 | 1.094 | 0.787 | 1.671 | 0.231 | 1.621 | 0.365 | 1.031 | 0.951 |
| 36 | **Glyat** | Unchanged | Q91XE0 | 22 | 496 | 21 | 296 | 34.1 | 8.28 | 89 | 0.885 | 0.429 | 1.034 | 0.867 | 0.916 | 0.432 | 1.128 | 0.533 |
| 37 | **Nit2** | Unchanged | Q9JHW2 | 23 | 403 | 23 | 276 | 30.5 | 6.9 | 88 | 1.144 | 0.360 | 1.101 | 0.682 | 1.084 | 0.490 | 1.016 | 0.946 |
| 38 | **Qdpr** | Down | Q8BVI4 | 19 | 479 | 19 | 241 | 25.6 | 7.81 | 88 | 0.841 | 0.475 | 0.792 | 0.274 | 0.655 | 0.043 | 1.209 | 0.295 |
| 39 | **Anxa5** | UP | P48036 | 31 | 480 | 31 | 319 | 35.7 | 4.96 | 88 | 1.021 | 0.923 | 2.134 | 0.034 | 1.828 | 0.018 | 1.167 | 0.540 |
| 40 | **Rack1** | Down | P68040 | 22 | 381 | 22 | 317 | 35.1 | 7.69 | 88 | 0.799 | 0.037 | 0.553 | 0.014 | 0.630 | 0.002 | 0.878 | 0.588 |
| 41 | **Gm3839** | Unchanged | S4R1W1 | 27 | 853 | 1 | 333 | 35.8 | 8.03 | 88 | 0.785 | 0.717 | 1.548 | 0.525 | 0.435 | 0.335 | 3.561 | 0.170 |
| 42 | **Sult2a8** | Unchanged | Q8BGL3 | 25 | 243 | 25 | 282 | 33.3 | 7.02 | 88 | 1.075 | 0.698 | 1.033 | 0.867 | 0.844 | 0.412 | 1.224 | 0.360 |
| 43 | **Selenbp2** | Down | Q63836 | 44 | 1199 | 8 | 472 | 52.6 | 6.18 | 88 | 0.702 | 0.164 | 0.571 | 0.029 | 0.487 | 0.005 | 1.171 | 0.417 |
| 44 | **Hbb-bsiso** | Unchanged | E9Q223 | 16 | 461 | 1 | 103 | 11.1 | 6.37 | 87 | #DIV/0! | #DIV/0! | #DIV/0! | #DIV/0! | #DIV/0! | #DIV/0! | #DIV/0! | #DIV/0! |
| 45 | **Hpd** | Down | P49429 | 36 | 896 | 36 | 393 | 45 | 7.06 | 87 | 1.032 | 0.812 | 0.649 | 0.025 | 0.883 | 0.325 | 0.735 | 0.151 |
| 46 | **Cyp2d10** | Unchanged | P24456 | 33 | 649 | 23 | 504 | 57.2 | 6.61 | 87 | 0.908 | 0.508 | 1.279 | 0.162 | 0.848 | 0.329 | 1.509 | 0.030 |
| 47 | **Selenbp1** | Unchanged | P17563 | 41 | 1197 | 6 | 472 | 52.5 | 6.29 | 87 | 0.758 | 0.496 | 1.639 | 0.211 | 1.182 | 0.614 | 1.386 | 0.245 |
| 48 | **Acat1** | Unchanged | Q8QZT1 | 32 | 826 | 32 | 424 | 44.8 | 8.51 | 87 | 1.295 | 0.244 | 0.894 | 0.448 | 1.031 | 0.763 | 0.867 | 0.337 |
| 49 | **Inmt** | Unchanged | P40936 | 20 | 446 | 20 | 264 | 29.4 | 6.39 | 87 | 1.367 | 0.031 | 0.995 | 0.983 | 1.488 | 0.032 | 0.669 | 0.066 |
| 50 | **Cyp2c29** | Unchanged | Q64458 | 42 | 630 | 27 | 490 | 55.7 | 8.29 | 87 | 1.172 | 0.471 | 0.753 | 0.263 | 0.641 | 0.179 | 1.175 | 0.625 |
| 51 | **Ndufa4** | Unchanged | Q62425 | 10 | 94 | 10 | 82 | 9.3 | 9.52 | 87 | 1.357 | 0.398 | 1.708 | 0.075 | 0.969 | 0.930 | 1.764 | 0.124 |
| 52 | **Bhmt** | Down | O35490 | 40 | 2415 | 36 | 407 | 45 | 7.9 | 86 | 0.991 | 0.964 | 0.519 | 0.028 | 0.809 | 0.385 | 0.642 | 0.164 |
| 53 | **Cyp2d26** | UP | Q8CIM7 | 37 | 571 | 2 | 500 | 56.9 | 6.64 | 80 | 1.980 | 0.028 | 4.588 | 0.110 | 4.663 | 0.166 | 0.984 | 0.981 |
| 54 | **Me1** | Unchanged | P06801 | 43 | 819 | 43 | 572 | 63.9 | 7.44 | 86 | 1.107 | 0.775 | 1.629 | 0.380 | 1.544 | 0.268 | 1.055 | 0.916 |
| 55 | **Nme2** | Unchanged | Q01768 | 14 | 448 | 9 | 152 | 17.4 | 7.5 | 80 | 0.839 | 0.737 | 1.032 | 0.944 | 1.044 | 0.920 | 0.988 | 0.976 |
| 56 | **Sult1a1** | UP | E9QNL5 | 24 | 191 | 24 | 291 | 33.9 | 8.1 | 86 | 1.423 | 0.163 | 2.172 | 0.015 | 1.221 | 0.443 | 1.779 | 0.059 |
| 57 | **Eif4a1** | Unchanged | P60843 | 34 | 498 | 20 | 406 | 46.1 | 5.48 | 86 | 0.868 | 0.441 | 0.781 | 0.245 | 1.050 | 0.687 | 0.744 | 0.132 |
| 58 | **Gnmt** | Unchanged | Q9QXF8 | 25 | 858 | 25 | 293 | 32.7 | 7.44 | 86 | 1.324 | 0.136 | 1.019 | 0.899 | 1.138 | 0.334 | 0.896 | 0.409 |
| 59 | **Gstz1** | Unchanged | Q9WVL0 | 18 | 710 | 18 | 216 | 24.3 | 7.85 | 86 | 1.079 | 0.528 | 0.856 | 0.481 | 1.050 | 0.686 | 0.815 | 0.371 |
| 60 | **Tkfc** | Unchanged | Q8VC30 | 40 | 1182 | 40 | 578 | 59.7 | 6.92 | 86 | 1.226 | 0.207 | 1.568 | 0.150 | 1.356 | 0.123 | 1.156 | 0.591 |
| 61 | **Blvrb** | Unchanged | Q923D2 | 13 | 336 | 13 | 206 | 22.2 | 7.01 | 85 | 1.164 | 0.292 | 0.895 | 0.444 | 1.158 | 0.243 | 0.773 | 0.134 |
| 62 | **Hadh** | Down | Q61425 | 24 | 640 | 24 | 314 | 34.4 | 8.65 | 85 | 0.902 | 0.254 | 0.760 | 0.043 | 0.608 | 0.001 | 1.250 | 0.178 |
| 63 | **Pgm1** | Unchanged | Q9D0F9 | 40 | 930 | 40 | 562 | 61.4 | 6.57 | 85 | 0.999 | 0.993 | 1.057 | 0.788 | 1.001 | 0.993 | 1.056 | 0.809 |
| 64 | **Hba-a1** | Unchanged | Q91VB8 | 11 | 479 | 2 | 142 | 15.1 | 8.22 | 85 | 0.915 | 0.465 | 1.194 | 0.561 | 0.930 | 0.602 | 1.284 | 0.454 |
| 65 | **Cat** | Unchanged | P24270 | 49 | 1643 | 49 | 527 | 59.8 | 7.88 | 85 | 0.917 | 0.717 | 1.126 | 0.715 | 1.013 | 0.957 | 1.111 | 0.764 |
| 66 | **Dmgdh** | Unchanged | Q9DBT9 | 61 | 1304 | 61 | 869 | 97.2 | 7.88 | 85 | 1.109 | 0.490 | 0.919 | 0.557 | 0.739 | 0.075 | 1.243 | 0.244 |
| 67 | **Hspa8** | Unchanged | P63017 | 49 | 1405 | 43 | 646 | 70.8 | 5.52 | 85 | 0.728 | 0.003 | 0.900 | 0.630 | 0.959 | 0.769 | 0.938 | 0.801 |
| 68 | **Abat** | Unchanged | P61922 | 37 | 583 | 37 | 500 | 56.4 | 8.09 | 85 | 1.235 | 0.261 | 1.236 | 0.269 | 1.158 | 0.295 | 1.068 | 0.719 |
| 69 | **Atp5md** | Unchanged | Q78IK2 | 5 | 32 | 5 | 58 | 6.4 | 9.83 | 59 | 0.773 | 0.399 | 0.908 | 0.779 | 0.549 | 0.144 | 1.655 | 0.260 |
| 70 | **Ppa1** | Unchanged | Q9D819 | 28 | 633 | 28 | 289 | 32.6 | 5.6 | 84 | 0.837 | 0.164 | 0.703 | 0.067 | 0.931 | 0.640 | 0.755 | 0.161 |
| 71 | **Esd** | Unchanged | Q9R0P3 | 20 | 465 | 20 | 282 | 31.3 | 7.12 | 84 | 1.201 | 0.272 | 1.332 | 0.274 | 1.186 | 0.233 | 1.124 | 0.589 |

| | | | | | | | | | | | | | | | | | | |
|---|---|---|---|---|---|---|---|---|---|---|---|---|---|---|---|---|---|---|
| 72 | **Acat3** | Unchanged | Q80X81 | 24 | 289 | 7 | 397 | 41.4 | 7.94 | 84 | 1.026 | 0.837 | 1.101 | 0.523 | 1.079 | 0.550 | 1.021 | 0.869 |
| 73 | **Prdx6** | Unchanged | O08709 | 24 | 780 | 2 | 224 | 24.9 | 6.01 | 75 | 0.998 | 0.994 | 0.791 | 0.612 | 0.978 | 0.957 | 0.809 | 0.598 |
| 74 | **Tpt1** | Unchanged | P63028 | 9 | 157 | 9 | 172 | 19.5 | 4.86 | 84 | 1.152 | 0.386 | 1.257 | 0.306 | 1.875 | 0.057 | 0.670 | 0.172 |
| 75 | **Idh1** | Unchanged | A0A087WNN7 | 2 | 6 | 2 | 19 | 2.1 | 4.75 | 84 | 0.098 | 0.224 | 0.953 | 0.961 | 0.512 | 0.505 | 1.863 | 0.511 |
| 76 | **Otc** | Unchanged | P11725 | 30 | 1013 | 30 | 354 | 39.7 | 8.73 | 84 | 1.005 | 0.979 | 0.835 | 0.339 | 1.101 | 0.510 | 0.759 | 0.195 |
| 77 | **Akr1c12** | Unchanged | Q9JLI0 | 23 | 257 | 9 | 323 | 37 | 6.64 | 84 | 1.407 | 0.156 | 1.216 | 0.588 | 1.521 | 0.107 | 0.800 | 0.401 |
| 78 | **Pah** | Unchanged | P16331 | 35 | 623 | 32 | 453 | 51.9 | 6.3 | 84 | 1.000 | 0.999 | 0.731 | 0.221 | 0.750 | 0.241 | 0.975 | 0.943 |
| 79 | **Ephx1** | Unchanged | Q9D379 | 35 | 463 | 35 | 455 | 52.5 | 8.35 | 84 | 1.154 | 0.458 | 1.643 | 0.088 | 1.696 | 0.054 | 0.968 | 0.902 |
| 80 | **Cyp2d26iso** | Unchanged | A0A0R4IZY2 | 37 | 384 | 2 | 500 | 56.9 | 6.64 | 84 | 0.991 | 0.939 | 1.067 | 0.717 | 0.998 | 0.988 | 1.070 | 0.730 |
| 81 | **Hgd** | Unchanged | O09173 | 27 | 701 | 27 | 445 | 49.9 | 7.24 | 84 | 1.164 | 0.185 | 1.315 | 0.208 | 1.147 | 0.176 | 1.146 | 0.495 |
| 82 | **Iah1** | Unchanged | Q9DB29 | 16 | 223 | 16 | 249 | 28 | 5.52 | 84 | 0.944 | 0.788 | 0.961 | 0.839 | 1.091 | 0.650 | 0.881 | 0.514 |
| 83 | **Pgrmc1** | Unchanged | O55022 | 15 | 333 | 14 | 195 | 21.7 | 4.7 | 61 | 1.070 | 0.826 | 0.841 | 0.462 | 0.950 | 0.849 | 0.885 | 0.513 |
| 84 | **Akr1c13** | Unchanged | Q8VC28 | 25 | 452 | 10 | 323 | 37 | 7.08 | 83 | 0.925 | 0.527 | 0.993 | 0.974 | 1.044 | 0.726 | 0.952 | 0.795 |
| 85 | **Adh5** | Unchanged | P28474 | 24 | 575 | 22 | 374 | 39.5 | 7.25 | 83 | 0.911 | 0.500 | 0.684 | 0.031 | 0.837 | 0.158 | 0.817 | 0.265 |
| 86 | **Haao** | Unchanged | Q78JT3 | 20 | 508 | 20 | 286 | 32.8 | 6.54 | 83 | 0.878 | 0.483 | 0.739 | 0.189 | 0.783 | 0.227 | 0.943 | 0.827 |
| 87 | **Aldh2 iso** | Unchanged | P47738 | 46 | 1426 | 16 | 519 | 56.5 | 7.62 | 83 | 1.100 | 0.672 | 1.129 | 0.668 | 1.124 | 0.637 | 1.004 | 0.988 |
| 88 | **Ftcd** | Unchanged | Q91XD4 | 39 | 895 | 39 | 541 | 58.9 | 6.11 | 83 | 1.106 | 0.753 | 1.274 | 0.222 | 1.349 | 0.161 | 0.944 | 0.683 |
| 89 | **Akr1a1** | Unchanged | Q9JII6 | 25 | 479 | 24 | 325 | 36.6 | 7.39 | 79 | 1.015 | 0.869 | 0.833 | 0.240 | 0.932 | 0.687 | 0.894 | 0.641 |
| 90 | **Ass1** | Unchanged | P16460 | 41 | 2481 | 41 | 412 | 46.6 | 8.22 | 83 | 0.983 | 0.918 | 0.647 | 0.132 | 0.919 | 0.618 | 0.703 | 0.297 |
| 91 | **Alb** | Unchanged | P07724 | 60 | 2163 | 60 | 608 | 68.6 | 6.07 | 83 | 1.052 | 0.784 | 1.326 | 0.219 | 1.397 | 0.120 | 0.950 | 0.806 |
| 92 | **Cyp2d9iso** | Unchanged | A0A2R8VHS2 | 13 | 138 | 1 | 206 | 23.1 | 5.22 | 83 | 0.504 | 0.557 | 0.167 | 0.305 | 0.828 | 0.842 | 0.202 | 0.094 |
| 93 | **Aldob** | Unchanged | Q91Y97 | 35 | 2381 | 34 | 364 | 39.5 | 8.27 | 82 | 1.028 | 0.904 | 0.790 | 0.188 | 1.008 | 0.967 | 0.783 | 0.285 |
| 94 | **Tuba1c** | Unchanged | P68373 | 29 | 540 | 3 | 449 | 49.9 | 5.1 | 82 | 1.077 | 0.821 | 0.937 | 0.853 | 0.883 | 0.487 | 1.062 | 0.871 |
| 95 | **Hspd1** | Unchanged | P63038 | 61 | 2515 | 61 | 573 | 60.9 | 6.18 | 82 | 1.098 | 0.539 | 1.116 | 0.549 | 1.175 | 0.156 | 0.950 | 0.777 |
| 96 | **Atp5f1b** | Unchanged | P56480 | 36 | 2069 | 36 | 529 | 56.3 | 5.34 | 82 | 1.165 | 0.374 | 0.965 | 0.904 | 1.029 | 0.893 | 0.938 | 0.843 |
| 97 | **Hebp1** | Unchanged | A0A140T8J4 | 13 | 147 | 3 | 190 | 21 | 5.26 | 82 | 1.062 | 0.824 | 1.273 | 0.493 | 1.047 | 0.855 | 1.215 | 0.593 |
| 98 | **Gstt1** | UP | Q64471 | 18 | 311 | 18 | 240 | 27.4 | 7.27 | 76 | 1.827 | 0.021 | 2.743 | 0.039 | 2.221 | 0.005 | 1.235 | 0.500 |
| 99 | **Cps1** | Unchanged | Q8C196 | 147 | 6730 | 144 | 1500 | 164.5 | 6.92 | 82 | 0.955 | 0.803 | 0.774 | 0.233 | 0.953 | 0.774 | 0.812 | 0.386 |
| 100 | **Akr1c6** | Unchanged | P70694 | 32 | 985 | 23 | 323 | 37 | 8.28 | 82 | 1.034 | 0.923 | 0.731 | 0.462 | 0.830 | 0.600 | 0.882 | 0.799 |
| 101 | **Aldh1a1** | Unchanged | P24549 | 42 | 1727 | 29 | 501 | 54.4 | 7.8 | 82 | 1.173 | 0.386 | 1.081 | 0.737 | 1.138 | 0.418 | 0.950 | 0.814 |
| 102 | **Dynll2** | Unchanged | Q9D0M5 | 6 | 86 | 4 | 89 | 10.3 | 7.37 | 82 | 1.406 | 0.257 | 0.780 | 0.556 | 1.369 | 0.250 | 0.570 | 0.097 |
| 103 | **Mgst1** | Unchanged | Q91VS7 | 13 | 688 | 13 | 155 | 17.5 | 9.66 | 82 | 1.029 | 0.954 | 0.665 | 0.396 | 0.838 | 0.745 | 0.794 | 0.667 |
| 104 | **Mup11** | Unchanged | P04938 | 21 | 949 | 0 | 181 | 20.7 | 4.94 | 82 | 3.049 | 0.359 | 3.404 | 0.268 | 0.680 | 0.562 | 5.004 | 0.210 |
| 105 | **Arf5** | Unchanged | P84084 | 13 | 182 | 6 | 180 | 20.5 | 6.79 | 82 | 0.873 | 0.747 | 1.345 | 0.443 | 0.697 | 0.233 | 1.930 | 0.128 |
| 106 | **Gstm2** | UP | P15626 | 24 | 455 | 12 | 218 | 25.7 | 7.39 | 77 | 1.032 | 0.909 | 2.074 | 0.026 | 0.897 | 0.725 | 2.312 | 0.036 |
| 107 | **Aldh2 isob** | Unchanged | A0A0G2JF60 | 19 | 285 | 1 | 196 | 21.4 | 8.03 | 82 | 0.595 | 0.515 | 0.768 | 0.733 | 0.477 | 0.410 | 1.612 | 0.563 |
| 108 | **Acox1 iso** | Unchanged | A2A848 | 39 | 642 | 2 | 625 | 70 | 7.75 | 82 | 2.056 | 0.220 | 3.018 | 0.059 | 2.891 | 0.016 | 1.044 | 0.885 |
| 109 | **Eno1** | Unchanged | P17182 | 40 | 1685 | 34 | 434 | 47.1 | 6.8 | 82 | 0.961 | 0.465 | 1.060 | 0.662 | 1.128 | 0.244 | 0.940 | 0.689 |
| 110 | **Sardh** | Unchanged | Q99LB7 | 49 | 1083 | 49 | 919 | 101.6 | 6.74 | 81 | 0.999 | 0.994 | 0.674 | 0.035 | 0.684 | 0.017 | 0.984 | 0.924 |
| 111 | **Acat2** | Unchanged | Q8CAY6 | 22 | 503 | 5 | 397 | 41.3 | 7.5 | 81 | 0.817 | 0.399 | 0.937 | 0.880 | 1.053 | 0.874 | 0.890 | 0.823 |
| 112 | **Gstt3** | Unchanged | Q99L20 | 15 | 146 | 15 | 241 | 27.4 | 7.74 | 76 | 1.065 | 0.781 | 0.684 | 0.196 | 0.702 | 0.217 | 0.973 | 0.872 |

| | | | | | | | | | | | | | | | | | | |
|---|---|---|---|---|---|---|---|---|---|---|---|---|---|---|---|---|---|---|
| 113 | **Pgk1** | Unchanged | P09411 | 35 | 929 | 35 | 417 | 44.5 | 7.9 | 81 | 0.956 | 0.779 | 0.948 | 0.804 | 1.183 | 0.159 | 0.802 | 0.263 |
| 114 | **Arf4** | Unchanged | P61750 | 12 | 180 | 6 | 180 | 20.4 | 7.14 | 81 | 1.853 | 0.237 | 3.395 | 0.175 | 4.404 | 0.096 | 0.771 | 0.684 |
| 115 | **Hbb-b1** | Unchanged | P02088 | 19 | 1164 | 2 | 147 | 15.8 | 7.65 | 76 | 8.795 | 0.385 | 0.932 | 0.944 | 0.189 | 0.179 | 4.919 | 0.377 |
| 116 | **P4hb** | Unchanged | P09103 | 56 | 1492 | 56 | 509 | 57 | 4.88 | 81 | 0.846 | 0.341 | 0.853 | 0.390 | 1.054 | 0.819 | 0.809 | 0.448 |
| 117 | **Aldh8a1** | Unchanged | Q8BH00 | 34 | 788 | 34 | 487 | 53.6 | 7.55 | 81 | 1.127 | 0.239 | 1.128 | 0.465 | 1.116 | 0.304 | 1.011 | 0.948 |
| 118 | **Rps3** | Unchanged | P62908 | 23 | 291 | 23 | 243 | 26.7 | 9.66 | 81 | 0.773 | 0.508 | 1.170 | 0.764 | 0.845 | 0.681 | 1.384 | 0.566 |
| 119 | **Acta1** | Unchanged | P68134 | 28 | 622 | 2 | 377 | 42 | 5.39 | 81 | 0.244 | 0.388 | 0.527 | 0.590 | 0.114 | 0.309 | 4.618 | 0.099 |
| 120 | **Actc1** | Unchanged | P68033 | 30 | 646 | 1 | 377 | 42 | 5.39 | 81 | 0.746 | 0.126 | 0.894 | 0.660 | 1.235 | 0.224 | 0.724 | 0.153 |
| 121 | **Mup13** | Unchanged | A2CEK6 | 20 | 393 | 4 | 180 | 20.6 | 5.2 | 81 | 0.700 | 0.611 | 0.464 | 0.292 | 0.131 | 0.081 | 3.538 | 0.155 |
| 122 | **Spr** | Unchanged | Q91XH5 | 15 | 144 | 15 | 262 | 27.9 | 6.21 | 81 | 0.843 | 0.682 | 0.761 | 0.529 | 0.793 | 0.547 | 0.960 | 0.908 |
| 123 | **Uqcrq** | Unchanged | Q9CQ69 | 7 | 65 | 7 | 82 | 9.8 | 10.26 | 70 | 1.234 | 0.734 | 0.989 | 0.985 | 0.512 | 0.363 | 1.931 | 0.200 |
| 124 | **Gsta3** | Unchanged | A0A087WQI6 | 11 | 416 | 1 | 133 | 15.4 | 5.55 | 80 | 3.594 | 0.207 | 2.061 | 0.337 | 3.779 | 0.148 | 0.545 | 0.366 |
| 125 | **4931406C07Rik** | Unchanged | Q91V76 | 19 | 298 | 19 | 315 | 35 | 6.29 | 80 | 1.179 | 0.273 | 1.102 | 0.617 | 1.018 | 0.914 | 1.083 | 0.701 |
| 126 | **Nme1** | Unchanged | P15532 | 14 | 326 | 9 | 152 | 17.2 | 7.37 | 80 | 0.677 | 0.054 | 0.660 | 0.056 | 0.785 | 0.180 | 0.841 | 0.272 |
| 127 | **Pahiso** | Unchanged | A0A1W2P781 | 4 | 23 | 1 | 81 | 8.7 | 5.01 | 80 | 0.786 | 0.713 | 0.319 | 0.188 | 0.055 | 0.063 | 5.840 | 0.197 |
| 128 | **Akr1c19** | Unchanged | G3X9Y6 | 19 | 177 | 15 | 323 | 37 | 7.23 | 80 | 1.115 | 0.621 | 1.031 | 0.893 | 1.584 | 0.057 | 0.651 | 0.050 |
| 129 | **Gpx1** | Unchanged | P11352 | 13 | 345 | 13 | 201 | 22.3 | 7.21 | 80 | 1.036 | 0.791 | 0.990 | 0.954 | 1.157 | 0.270 | 0.855 | 0.336 |
| 130 | **Lap3** | Unchanged | Q9CPY7 | 36 | 579 | 36 | 519 | 56.1 | 7.72 | 80 | 0.826 | 0.516 | 0.812 | 0.539 | 0.792 | 0.471 | 1.026 | 0.935 |
| 131 | **Cyb5a** | Unchanged | P56395 | 8 | 285 | 4 | 134 | 15.2 | 5.07 | 80 | 1.198 | 0.528 | 0.823 | 0.562 | 0.798 | 0.266 | 1.031 | 0.941 |
| 132 | **Sec14l2** | Unchanged | Q99J08 | 29 | 700 | 29 | 403 | 46.3 | 7.11 | 80 | 1.213 | 0.179 | 1.245 | 0.334 | 1.251 | 0.021 | 0.995 | 0.981 |
| 133 | **Atp5pd** | Unchanged | Q9DCX2 | 13 | 327 | 13 | 161 | 18.7 | 5.69 | 80 | 1.241 | 0.176 | 0.782 | 0.301 | 1.084 | 0.719 | 0.722 | 0.249 |
| 134 | **Gdi2** | Unchanged | Q61598 | 36 | 499 | 30 | 445 | 50.5 | 6.25 | 79 | 1.125 | 0.062 | 1.220 | 0.400 | 1.379 | 0.003 | 0.885 | 0.552 |
| 135 | **Calm2** | Unchanged | P0DP27 | 11 | 360 | 6 | 149 | 16.8 | 4.22 | 79 | 0.692 | 0.108 | 1.106 | 0.706 | 0.838 | 0.404 | 1.320 | 0.363 |
| 136 | **Aldh9a1** | Unchanged | Q9JLJ2 | 33 | 552 | 33 | 494 | 53.5 | 6.98 | 79 | 1.132 | 0.673 | 0.873 | 0.649 | 1.195 | 0.426 | 0.731 | 0.144 |
| 137 | **Eif4a2** | Unchanged | P10630 | 26 | 329 | 12 | 407 | 46.4 | 5.48 | 79 | 0.975 | 0.851 | 0.711 | 0.033 | 1.384 | 0.372 | 0.514 | 0.134 |
| 138 | **Gpd1** | Unchanged | P13707 | 26 | 470 | 25 | 349 | 37.5 | 7.17 | 79 | 1.193 | 0.268 | 1.150 | 0.642 | 0.900 | 0.615 | 1.279 | 0.440 |
| 139 | **Mup22** | Down | Q4FZE8 | 21 | 533 | 1 | 181 | 20.8 | 5.02 | 79 | 0.856 | 0.671 | 0.587 | 0.279 | 0.287 | 0.023 | 2.046 | 0.318 |
| 140 | **Idh1 iso** | Unchanged | O88844 | 35 | 1027 | 34 | 414 | 46.6 | 7.17 | 79 | 0.938 | 0.690 | 0.980 | 0.932 | 1.677 | 0.053 | 0.585 | 0.078 |
| 141 | **Ndufb1** | Unchanged | P0DN34 | 4 | 16 | 4 | 57 | 7 | 8.21 | 79 | 0.893 | 0.851 | 0.384 | 0.139 | 0.863 | 0.826 | 0.445 | 0.406 |
| 142 | **Gstm7** | Unchanged | Q80W21 | 22 | 421 | 12 | 218 | 25.7 | 6.8 | 74 | 0.971 | 0.915 | 1.094 | 0.751 | 1.074 | 0.806 | 1.019 | 0.934 |
| 143 | **Park7** | Unchanged | Q99LX0 | 14 | 403 | 14 | 189 | 20 | 6.77 | 79 | 1.022 | 0.924 | 0.752 | 0.199 | 1.041 | 0.810 | 0.722 | 0.013 |
| 144 | **Acsl1** | Unchanged | P41216 | 47 | 1058 | 44 | 699 | 77.9 | 7.15 | 79 | 0.950 | 0.827 | 0.834 | 0.588 | 0.822 | 0.514 | 1.014 | 0.973 |
| 145 | **Pbld2** | Unchanged | Q9CXN7 | 22 | 396 | 15 | 288 | 32 | 5.34 | 79 | 1.182 | 0.106 | 1.171 | 0.326 | 1.309 | 0.110 | 0.895 | 0.559 |
| 146 | **Acaa1b** | UP | Q8VCH0 | 28 | 993 | 12 | 424 | 44 | 8.51 | 79 | 1.443 | 0.227 | 1.308 | 0.561 | 1.937 | 0.014 | 0.675 | 0.206 |
| 147 | **Asl** | Unchanged | Q91YI0 | 36 | 996 | 36 | 464 | 51.7 | 6.96 | 79 | 0.945 | 0.655 | 0.925 | 0.423 | 0.911 | 0.340 | 1.015 | 0.886 |
| 148 | **Pebp1** | Unchanged | P70296 | 14 | 265 | 14 | 187 | 20.8 | 5.4 | 79 | 0.888 | 0.222 | 1.195 | 0.482 | 1.032 | 0.808 | 1.158 | 0.565 |
| 149 | **Hint1** | Unchanged | P70349 | 8 | 209 | 8 | 126 | 13.8 | 6.87 | 79 | 0.874 | 0.590 | 0.786 | 0.406 | 0.867 | 0.653 | 0.907 | 0.761 |
| 150 | **Igkc** | Unchanged | P01837 | 7 | 57 | 7 | 107 | 11.9 | 5.9 | 79 | 0.903 | 0.793 | 1.707 | 0.323 | 0.613 | 0.285 | 2.785 | 0.128 |
| 151 | **Glrx** | Unchanged | Q9QUH0 | 6 | 125 | 6 | 107 | 11.9 | 8.37 | 79 | 0.680 | 0.007 | 0.868 | 0.445 | 0.676 | 0.028 | 1.285 | 0.365 |
| 152 | **Ugdh** | Unchanged | O70475 | 32 | 591 | 32 | 493 | 54.8 | 7.56 | 78 | 0.902 | 0.291 | 1.233 | 0.203 | 1.475 | 0.011 | 0.836 | 0.281 |
| 153 | **Kynu** | Unchanged | Q9CXF0 | 27 | 232 | 27 | 464 | 52.3 | 6.57 | 78 | 1.367 | 0.285 | 1.556 | 0.076 | 1.490 | 0.070 | 1.044 | 0.769 |

| | | | | | | | | | | | | | | | | | | |
|---|---|---|---|---|---|---|---|---|---|---|---|---|---|---|---|---|---|---|
| 154 | **Grhpr** | Unchanged | D6REG4 | 15 | 216 | 1 | 197 | 21.2 | 7.8 | 78 | 1.087 | 0.890 | 0.208 | 0.135 | 4.423 | 0.087 | 0.047 | 0.038 |
| 155 | **Cyp2c50** | Down | Q91X77 | 34 | 456 | 7 | 490 | 55.7 | 7.88 | 78 | 0.900 | 0.671 | 0.685 | 0.082 | 0.249 | 0.001 | 2.751 | 0.005 |
| 156 | **Hagh** | Unchanged | A0A0R4J052 | 13 | 147 | 1 | 260 | 28.9 | 7.12 | 78 | 4.998 | 0.320 | 8.900 | 0.232 | 1.102 | 0.843 | 8.073 | 0.238 |
| 157 | **Sh3bgrl** | Unchanged | Q9JJU8 | 6 | 48 | 6 | 114 | 12.8 | 4.92 | 78 | 0.706 | 0.425 | 1.286 | 0.494 | 0.730 | 0.423 | 1.762 | 0.056 |
| 158 | **Gstm3** | UP | P19639 | 22 | 376 | 9 | 218 | 25.7 | 7.77 | 73 | 0.978 | 0.942 | 5.316 | 0.050 | 4.201 | 0.025 | 1.266 | 0.626 |
| 159 | **Uqcrc2** | Unchanged | Q9DB77 | 24 | 415 | 24 | 453 | 48.2 | 9.25 | 78 | 1.179 | 0.502 | 0.821 | 0.393 | 0.867 | 0.528 | 0.947 | 0.816 |
| 160 | **Oat** | Unchanged | P29758 | 24 | 442 | 24 | 439 | 48.3 | 6.62 | 78 | 1.437 | 0.118 | 0.740 | 0.155 | 0.833 | 0.355 | 0.889 | 0.575 |
| 161 | **Tuba1a** | Unchanged | P68369 | 27 | 496 | 1 | 451 | 50.1 | 5.06 | 78 | 0.843 | 0.620 | 0.915 | 0.864 | 0.627 | 0.193 | 1.459 | 0.559 |
| 162 | **Mat1a** | Unchanged | Q91X83 | 25 | 593 | 21 | 396 | 43.5 | 5.82 | 78 | 0.977 | 0.833 | 0.680 | 0.061 | 0.898 | 0.415 | 0.757 | 0.234 |
| 163 | **Sar1b** | Unchanged | Q9CQC9 | 14 | 207 | 11 | 198 | 22.4 | 6.11 | 78 | 1.106 | 0.593 | 0.958 | 0.850 | 0.870 | 0.557 | 1.101 | 0.650 |
| 164 | **Amdhd1** | UP | Q9DBA8 | 21 | 415 | 21 | 426 | 46.5 | 6.95 | 78 | 1.289 | 0.036 | 1.482 | 0.055 | 1.753 | 0.004 | 0.845 | 0.302 |
| 165 | **Isoc2a** | Unchanged | P85094 | 8 | 235 | 8 | 206 | 22.4 | 8.02 | 78 | 1.053 | 0.899 | 1.851 | 0.200 | 1.325 | 0.548 | 1.397 | 0.426 |
| 166 | **Eef1a1** | Unchanged | D3YZ68 | 26 | 632 | 2 | 249 | 27.6 | 8.65 | 78 | 2.253 | 0.341 | 3.284 | 0.285 | 2.077 | 0.342 | 1.582 | 0.605 |
| 167 | **Dhrs1** | Unchanged | Q99L04 | 17 | 223 | 17 | 313 | 34 | 8.35 | 77 | 1.019 | 0.943 | 0.937 | 0.844 | 0.706 | 0.264 | 1.326 | 0.408 |
| 168 | **Cyb5aiso** | Unchanged | E0CY88 | 5 | 102 | 1 | 110 | 12.3 | 5.45 | 77 | #DIV/0! | #DIV/0! | #DIV/0! | #DIV/0! | #DIV/0! | #DIV/0! | #DIV/0! | #DIV/0! |
| 169 | **Etfb** | Unchanged | Q9DCW4 | 21 | 499 | 21 | 255 | 27.6 | 8.1 | 77 | 0.881 | 0.141 | 1.001 | 0.996 | 0.940 | 0.572 | 1.064 | 0.666 |
| 170 | **Slc25a13** | Unchanged | Q9QXX4 | 35 | 646 | 31 | 676 | 74.4 | 8.6 | 77 | 1.073 | 0.509 | 1.078 | 0.828 | 1.197 | 0.345 | 0.900 | 0.767 |
| 171 | **Tkt** | Unchanged | P40142 | 37 | 1073 | 37 | 623 | 67.6 | 7.5 | 77 | 0.960 | 0.785 | 1.139 | 0.432 | 1.196 | 0.185 | 0.952 | 0.714 |
| 172 | **Pfn1** | Unchanged | P62962 | 12 | 344 | 12 | 140 | 14.9 | 8.28 | 77 | 0.775 | 0.167 | 0.875 | 0.590 | 0.856 | 0.376 | 1.022 | 0.924 |
| 173 | **Pdia3** | Unchanged | P27773 | 45 | 900 | 45 | 505 | 56.6 | 6.21 | 77 | 0.876 | 0.504 | 0.892 | 0.573 | 0.930 | 0.729 | 0.959 | 0.873 |
| 174 | **Akr1c14** | Unchanged | Q91WT7 | 21 | 240 | 19 | 323 | 37.2 | 7.2 | 77 | 1.183 | 0.599 | 1.444 | 0.383 | 1.662 | 0.122 | 0.869 | 0.714 |
| 175 | **Fdps** | Unchanged | Q920E5 | 21 | 360 | 21 | 353 | 40.6 | 5.66 | 77 | 0.739 | 0.080 | 1.065 | 0.817 | 1.006 | 0.952 | 1.058 | 0.829 |
| 176 | **Got1** | Unchanged | P05201 | 27 | 517 | 27 | 413 | 46.2 | 7.14 | 72 | 0.748 | 0.242 | 0.806 | 0.443 | 0.714 | 0.180 | 1.128 | 0.570 |
| 177 | **Acaa1a** | Unchanged | Q921H8 | 27 | 892 | 9 | 424 | 43.9 | 8.44 | 77 | 1.017 | 0.897 | 1.029 | 0.842 | 1.267 | 0.073 | 0.812 | 0.190 |
| 178 | **Mup7** | Unchanged | L7MUC7 | 17 | 400 | 2 | 171 | 19.5 | 4.91 | 77 | 0.014 | 0.057 | 0.013 | 0.056 | 0.011 | 0.056 | 1.119 | 0.538 |
| 179 | **Tuba4a** | Unchanged | P68368 | 25 | 558 | 9 | 448 | 49.9 | 5.06 | 77 | 0.900 | 0.764 | 0.846 | 0.689 | 0.657 | 0.219 | 1.287 | 0.638 |
| 180 | **Slc25a5** | Unchanged | P51881 | 28 | 436 | 17 | 298 | 32.9 | 9.73 | 77 | 0.780 | 0.638 | 0.709 | 0.543 | 0.526 | 0.286 | 1.348 | 0.679 |
| 181 | **Hibadh** | Unchanged | Q99L13 | 19 | 350 | 19 | 335 | 35.4 | 8.13 | 76 | 1.302 | 0.183 | 1.388 | 0.121 | 1.119 | 0.500 | 1.241 | 0.248 |
| 182 | **Pbld1** | Unchanged | Q9DCG6 | 18 | 378 | 11 | 288 | 32 | 7.03 | 76 | 1.049 | 0.842 | 0.986 | 0.950 | 0.851 | 0.477 | 1.159 | 0.311 |
| 183 | **Pgam1** | Unchanged | Q9DBJ1 | 17 | 503 | 14 | 254 | 28.8 | 7.18 | 76 | 1.077 | 0.375 | 1.100 | 0.409 | 1.202 | 0.021 | 0.915 | 0.391 |
| 184 | **Gclc** | UP | P97494 | 35 | 444 | 35 | 637 | 72.5 | 5.9 | 74 | 1.564 | 0.005 | 1.796 | 0.000 | 2.262 | 0.000 | 0.794 | 0.012 |
| 185 | **Ptges3** | Down | Q9R0Q7 | 10 | 117 | 10 | 160 | 18.7 | 4.55 | 76 | 0.805 | 0.177 | 0.826 | 0.314 | 0.456 | 0.003 | 1.812 | 0.015 |
| 186 | **Uox** | Unchanged | P25688 | 25 | 512 | 25 | 303 | 35 | 8.35 | 76 | 1.655 | 0.214 | 1.353 | 0.491 | 2.583 | 0.070 | 0.524 | 0.181 |
| 187 | **Dbi** | Unchanged | D3Z563 | 7 | 80 | 1 | 63 | 7.2 | 9.57 | 76 | 3.612 | 0.286 | 0.083 | 0.138 | 3.653 | 0.248 | 0.023 | 0.121 |
| 188 | **Rhoa** | Unchanged | Q9QUI0 | 12 | 171 | 4 | 193 | 21.8 | 6.1 | 76 | 0.929 | 0.700 | 1.044 | 0.857 | 1.370 | 0.225 | 0.762 | 0.302 |
| 189 | **Rnh1** | UP | Q91VI7 | 22 | 394 | 22 | 456 | 49.8 | 4.78 | 76 | 1.414 | 0.140 | 1.800 | 0.042 | 1.338 | 0.264 | 1.345 | 0.312 |
| 190 | **Rdh16f1** | Unchanged | E9Q9P8 | 21 | 192 | 3 | 317 | 35.8 | 8.81 | 76 | 1.014 | 0.948 | 2.160 | 0.207 | 1.974 | 0.067 | 1.094 | 0.850 |
| 191 | **Dhrs4** | UP | Q99LB2 | 16 | 188 | 16 | 279 | 29.9 | 9.38 | 76 | 1.432 | 0.118 | 1.509 | 0.040 | 1.975 | 0.018 | 0.764 | 0.202 |
| 192 | **Cyp2f2** | Unchanged | P33267 | 38 | 580 | 36 | 491 | 55.9 | 7.9 | 76 | 0.802 | 0.334 | 0.932 | 0.879 | 0.885 | 0.531 | 1.053 | 0.917 |
| 193 | **Comt** | Unchanged | O88587 | 23 | 543 | 23 | 265 | 29.5 | 5.83 | 76 | 0.981 | 0.914 | 1.500 | 0.294 | 1.027 | 0.926 | 1.460 | 0.372 |
| 194 | **Ak2** | Unchanged | Q9WTP6 | 16 | 434 | 13 | 239 | 26.5 | 7.39 | 70 | 1.121 | 0.557 | 1.303 | 0.176 | 1.417 | 0.082 | 0.919 | 0.640 |

| | | | | | | | | | | | | | | | | | |
|---|---|---|---|---|---|---|---|---|---|---|---|---|---|---|---|---|---|
| 195 | **Baat** | Unchanged | Q91X34 | 20 | 270 | 20 | 420 | 46.5 | 6.68 | 76 | 1.092 | 0.814 | 1.471 | 0.451 | 1.154 | 0.770 | 1.275 | 0.682 |
| 196 | **Arf1** | UP | P84078 | 14 | 266 | 2 | 181 | 20.7 | 6.8 | 76 | 1.245 | 0.441 | 1.533 | 0.036 | 1.103 | 0.653 | 1.390 | 0.092 |
| 197 | **Arf3** | Unchanged | P61205 | 13 | 245 | 1 | 181 | 20.6 | 7.43 | 76 | 1.632 | 0.136 | 4.205 | 0.190 | 1.041 | 0.896 | 4.039 | 0.195 |
| 198 | **Keg1** | Unchanged | Q9DCY0 | 16 | 229 | 16 | 295 | 33.7 | 8.43 | 76 | 0.953 | 0.926 | 1.353 | 0.638 | 1.045 | 0.932 | 1.295 | 0.694 |
| 199 | **Tpi1** | Unchanged | P17751 | 19 | 640 | 19 | 299 | 32.2 | 5.74 | 76 | 0.905 | 0.545 | 1.114 | 0.567 | 1.039 | 0.851 | 1.072 | 0.740 |
| 200 | **Pck1** | Unchanged | Q9Z2V4 | 36 | 426 | 36 | 622 | 69.3 | 6.6 | 76 | 0.667 | 0.217 | 0.691 | 0.242 | 0.841 | 0.615 | 0.822 | 0.508 |
| 201 | **Cyp2c67** | Unchanged | Q569X9 | 26 | 136 | 10 | 491 | 56.1 | 7.36 | 76 | 0.745 | 0.116 | 0.817 | 0.360 | 0.670 | 0.048 | 1.219 | 0.499 |
| 202 | **Vcp** | Unchanged | Q01853 | 64 | 1226 | 64 | 806 | 89.3 | 5.26 | 76 | 1.131 | 0.545 | 1.108 | 0.561 | 1.217 | 0.244 | 0.910 | 0.512 |
| 203 | **Glud1** | Unchanged | P26443 | 49 | 1814 | 49 | 558 | 61.3 | 8 | 75 | 1.314 | 0.121 | 1.207 | 0.155 | 1.477 | 0.041 | 0.817 | 0.236 |
| 204 | **Tmem14c** | Unchanged | Q9CQN6 | 5 | 30 | 5 | 114 | 11.6 | 9.7 | 75 | 1.013 | 0.967 | 1.675 | 0.057 | 0.854 | 0.657 | 1.963 | 0.030 |
| 205 | **Gstt2** | UP | Q61133 | 12 | 112 | 12 | 244 | 27.6 | 7.55 | 70 | 1.836 | 0.048 | 1.407 | 0.129 | 1.610 | 0.004 | 0.874 | 0.334 |
| 206 | **Ywhag** | Unchanged | P61982 | 17 | 315 | 11 | 247 | 28.3 | 4.89 | 75 | 1.018 | 0.958 | 1.073 | 0.830 | 1.035 | 0.911 | 1.037 | 0.894 |
| 207 | **Ces1d** | Unchanged | Q8VCT4 | 39 | 1209 | 35 | 565 | 61.7 | 6.61 | 75 | 1.116 | 0.369 | 1.141 | 0.548 | 1.311 | 0.044 | 0.870 | 0.460 |
| 208 | **Got2** | Unchanged | P05202 | 32 | 911 | 32 | 430 | 47.4 | 9 | 75 | 0.936 | 0.844 | 0.684 | 0.346 | 0.723 | 0.390 | 0.945 | 0.892 |
| 209 | **Phb** | Unchanged | P67778 | 14 | 223 | 14 | 272 | 29.8 | 5.76 | 75 | 1.059 | 0.851 | 1.266 | 0.562 | 1.036 | 0.908 | 1.222 | 0.668 |
| 210 | **Prdx6iso** | Unchanged | Q6GT24 | 25 | 438 | 3 | 224 | 24.8 | 6.37 | 75 | 0.948 | 0.651 | 1.028 | 0.842 | 0.941 | 0.647 | 1.092 | 0.432 |
| 211 | **Calr** | Unchanged | P14211 | 29 | 1023 | 29 | 416 | 48 | 4.49 | 75 | 0.866 | 0.667 | 0.735 | 0.357 | 0.803 | 0.506 | 0.916 | 0.792 |
| 212 | **Psmd13** | UP | Q9WVJ2 | 21 | 171 | 21 | 376 | 42.8 | 5.71 | 75 | 1.751 | 0.013 | 1.474 | 0.001 | 1.612 | 0.088 | 0.915 | 0.675 |
| 213 | **Fahiso** | Unchanged | P35505 | 23 | 843 | 13 | 419 | 46.1 | 7.18 | 75 | 0.933 | 0.542 | 1.001 | 0.994 | 1.067 | 0.668 | 0.939 | 0.766 |
| 214 | **Acadvl** | Unchanged | P50544 | 45 | 873 | 45 | 656 | 70.8 | 8.75 | 75 | 0.960 | 0.777 | 1.296 | 0.442 | 0.813 | 0.056 | 1.594 | 0.229 |
| 215 | **Lypla1** | Unchanged | P97823 | 9 | 124 | 9 | 230 | 24.7 | 6.62 | 75 | 2.043 | 0.257 | 3.243 | 0.083 | 1.385 | 0.428 | 2.341 | 0.149 |
| 216 | **Cyb5r3** | Unchanged | Q9DCN2 | 22 | 432 | 22 | 301 | 34.1 | 8.38 | 75 | 0.743 | 0.001 | 0.959 | 0.771 | 0.823 | 0.167 | 1.165 | 0.453 |
| 217 | **Grhpriso** | Unchanged | Q91Z53 | 21 | 575 | 7 | 328 | 35.3 | 7.65 | 75 | 0.899 | 0.387 | 0.724 | 0.049 | 0.950 | 0.584 | 0.761 | 0.081 |
| 218 | **Shmt2** | Unchanged | Q9CZN7 | 30 | 253 | 29 | 504 | 55.7 | 8.47 | 75 | 0.783 | 0.082 | 0.832 | 0.305 | 0.766 | 0.056 | 1.085 | 0.695 |
| 219 | **S100a11** | UP | P50543 | 4 | 16 | 4 | 98 | 11.1 | 5.45 | 74 | 1.287 | 0.556 | 3.815 | 0.211 | 3.602 | 0.044 | 1.059 | 0.929 |
| 220 | **Pcx** | Unchanged | E9QPD7 | 67 | 801 | 67 | 1179 | 129.7 | 6.71 | 74 | 0.983 | 0.896 | 1.081 | 0.612 | 0.864 | 0.432 | 1.252 | 0.291 |
| 221 | **Ppia** | Unchanged | P17742 | 12 | 405 | 12 | 164 | 18 | 7.9 | 63 | 0.878 | 0.358 | 1.048 | 0.823 | 1.314 | 0.126 | 0.798 | 0.351 |
| 222 | **Gc** | Unchanged | P21614 | 30 | 434 | 30 | 476 | 53.6 | 5.5 | 74 | 0.888 | 0.536 | 1.075 | 0.796 | 1.360 | 0.246 | 0.790 | 0.448 |
| 223 | **Amacr** | Unchanged | O09174 | 15 | 228 | 15 | 381 | 41.7 | 7.4 | 74 | 1.047 | 0.921 | 0.819 | 0.685 | 0.768 | 0.558 | 1.067 | 0.817 |
| 224 | **Acads** | Unchanged | Q07417 | 24 | 504 | 24 | 412 | 44.9 | 8.47 | 74 | 0.847 | 0.351 | 0.832 | 0.498 | 0.808 | 0.160 | 1.030 | 0.921 |
| 225 | **Pklr** | Unchanged | G3X925 | 39 | 530 | 38 | 574 | 62.2 | 7.09 | 74 | 1.332 | 0.141 | 1.328 | 0.102 | 1.460 | 0.072 | 0.910 | 0.598 |
| 226 | **Aldh6a1** | Unchanged | Q9EQ20 | 34 | 932 | 34 | 535 | 57.9 | 8.07 | 74 | 1.053 | 0.550 | 0.907 | 0.503 | 1.006 | 0.960 | 0.902 | 0.497 |
| 227 | **Uba1** | Unchanged | Q02053 | 53 | 731 | 53 | 1058 | 117.7 | 5.66 | 74 | 1.101 | 0.369 | 1.190 | 0.170 | 1.058 | 0.578 | 1.125 | 0.362 |
| 228 | **Etfa** | Unchanged | Q99LC5 | 26 | 721 | 26 | 333 | 35 | 8.38 | 74 | 0.986 | 0.894 | 1.162 | 0.367 | 1.154 | 0.366 | 1.008 | 0.971 |
| 229 | **Krt8** | Down | P11679 | 37 | 545 | 31 | 490 | 54.5 | 5.82 | 74 | 0.659 | 0.088 | 1.137 | 0.797 | 0.405 | 0.008 | 2.808 | 0.179 |
| 230 | **Eif5a** | Unchanged | P63242 | 19 | 455 | 19 | 154 | 16.8 | 5.24 | 74 | 1.001 | 0.998 | 1.177 | 0.525 | 1.186 | 0.531 | 0.993 | 0.979 |
| 231 | **Ak3** | Unchanged | Q9WTP7 | 18 | 292 | 18 | 227 | 25.4 | 8.84 | 74 | 0.994 | 0.973 | 1.141 | 0.362 | 0.914 | 0.514 | 1.248 | 0.075 |
| 232 | **Pygl** | Unchanged | Q9ET01 | 61 | 1345 | 55 | 850 | 97.4 | 7.09 | 74 | 0.968 | 0.820 | 0.832 | 0.197 | 0.782 | 0.150 | 1.064 | 0.657 |
| 233 | **Krt18** | Down | P05784 | 34 | 662 | 33 | 423 | 47.5 | 5.33 | 74 | 0.788 | 0.136 | 0.655 | 0.074 | 0.550 | 0.005 | 1.192 | 0.508 |
| 234 | **Ces1f** | Unchanged | Q91WU0 | 36 | 993 | 32 | 561 | 61.6 | 6.48 | 74 | 1.068 | 0.568 | 0.899 | 0.554 | 1.140 | 0.305 | 0.788 | 0.173 |
| 235 | **Dynlrb1** | Unchanged | P62627 | 4 | 42 | 4 | 96 | 11 | 7.25 | 74 | 0.613 | 0.189 | 0.926 | 0.839 | 0.745 | 0.370 | 1.244 | 0.492 |

| | | | | | | | | | | | | | | | | | | |
|---|---|---|---|---|---|---|---|---|---|---|---|---|---|---|---|---|---|---|
| 236 | **Cltc** | Unchanged | Q68FD5 | 103 | 1408 | 103 | 1675 | 191.4 | 5.69 | 74 | 1.060 | 0.525 | 1.328 | 0.142 | 0.971 | 0.852 | 1.367 | 0.171 |
| 237 | **Selenbp2iso** | Unchanged | G3UZT2 | 10 | 86 | 3 | 134 | 14.6 | 7.74 | 74 | 0.720 | 0.333 | 0.403 | 0.016 | 0.214 | 0.001 | 1.886 | 0.144 |
| 238 | **Cyp2a12** | Unchanged | P56593 | 36 | 441 | 33 | 492 | 56.1 | 9.19 | 74 | 0.698 | 0.195 | 0.910 | 0.729 | 0.655 | 0.116 | 1.388 | 0.311 |
| 239 | **Psmc4** | Unchanged | P54775 | 18 | 234 | 17 | 418 | 47.4 | 5.21 | 74 | 1.560 | 0.217 | 1.998 | 0.092 | 1.662 | 0.093 | 1.202 | 0.584 |
| 240 | **Aldh1b1** | UP | Q9CZS1 | 34 | 480 | 32 | 519 | 57.5 | 7.02 | 74 | 1.231 | 0.167 | 2.004 | 0.012 | 4.105 | 0.048 | 0.488 | 0.161 |
| 241 | **Ehhadh** | UP | Q9DBM2 | 45 | 815 | 45 | 718 | 78.3 | 9.13 | 74 | 1.175 | 0.636 | 1.394 | 0.411 | 2.093 | 0.043 | 0.666 | 0.215 |
| 242 | **Fabp5** | Unchanged | Q05816 | 11 | 85 | 11 | 135 | 15.1 | 6.54 | 73 | 0.267 | 0.063 | 0.435 | 0.154 | 0.589 | 0.335 | 0.739 | 0.555 |
| 243 | **Gamt** | Unchanged | O35969 | 10 | 227 | 10 | 236 | 26.3 | 5.71 | 73 | 1.161 | 0.465 | 1.139 | 0.757 | 1.567 | 0.089 | 0.727 | 0.374 |
| 244 | **Pccb** | Unchanged | Q99MN9 | 25 | 416 | 25 | 541 | 58.4 | 7.66 | 73 | 1.221 | 0.364 | 1.483 | 0.202 | 1.256 | 0.252 | 1.181 | 0.539 |
| 245 | **Scp2** | Unchanged | P32020 | 36 | 1131 | 36 | 547 | 59.1 | 7.44 | 73 | 0.883 | 0.426 | 0.736 | 0.189 | 0.892 | 0.433 | 0.825 | 0.362 |
| 246 | **Cth** | Unchanged | Q8VCN5 | 22 | 505 | 22 | 398 | 43.5 | 7.65 | 73 | 1.135 | 0.393 | 1.153 | 0.021 | 1.043 | 0.637 | 1.106 | 0.213 |
| 247 | **Rps15a** | Down | P62245 | 12 | 89 | 12 | 130 | 14.8 | 10.13 | 73 | 0.832 | 0.347 | 0.591 | 0.007 | 0.943 | 0.699 | 0.627 | 0.022 |
| 248 | **S100a8** | UP | P27005 | 3 | 24 | 3 | 89 | 10.3 | 5.68 | 73 | 0.378 | 0.167 | 0.723 | 0.497 | 14.694 | 0.049 | 0.049 | 0.045 |
| 249 | **Mup20** | Unchanged | Q5FW60 | 14 | 246 | 11 | 181 | 20.9 | 4.89 | 73 | 1.114 | 0.618 | 1.893 | 0.115 | 1.378 | 0.233 | 1.374 | 0.386 |
| 250 | **Agxt2** | Unchanged | Q3UEG6 | 30 | 390 | 30 | 513 | 57.1 | 7.93 | 73 | 1.096 | 0.592 | 0.843 | 0.543 | 0.883 | 0.539 | 0.955 | 0.879 |
| 251 | **Uqcrc1** | Unchanged | A0A0A6YW82 | 10 | 95 | 1 | 191 | 20.8 | 5.39 | 73 | 3.563 | 0.265 | 3.019 | 0.497 | 0.618 | 0.381 | 4.883 | 0.421 |
| 252 | **Pgls** | Unchanged | Q9CQ60 | 11 | 183 | 11 | 257 | 27.2 | 5.85 | 73 | 1.423 | 0.052 | 1.371 | 0.004 | 1.322 | 0.061 | 1.037 | 0.780 |
| 253 | **Apoa1** | Unchanged | Q00623 | 22 | 319 | 22 | 264 | 30.6 | 5.73 | 73 | 0.591 | 0.243 | 0.628 | 0.287 | 0.737 | 0.485 | 0.851 | 0.595 |
| 254 | **Uqcrfs1** | Down | Q9CR68 | 15 | 240 | 15 | 274 | 29.3 | 8.7 | 73 | 0.809 | 0.057 | 0.658 | 0.004 | 0.598 | 0.001 | 1.101 | 0.528 |
| 255 | **Hadha** | Unchanged | Q8BMS1 | 49 | 1484 | 49 | 763 | 82.6 | 9.14 | 73 | 1.174 | 0.325 | 1.258 | 0.482 | 1.186 | 0.323 | 1.060 | 0.850 |
| 256 | **Hsd11b1** | Unchanged | P50172 | 19 | 293 | 19 | 292 | 32.3 | 8.46 | 73 | 0.869 | 0.221 | 1.118 | 0.499 | 0.841 | 0.279 | 1.329 | 0.185 |
| 257 | **Sult1d1** | UP | Q3UZZ6 | 15 | 181 | 15 | 295 | 35.1 | 5.76 | 73 | 1.402 | 0.218 | 1.179 | 0.586 | 1.728 | 0.016 | 0.682 | 0.087 |
| 258 | **Msra** | Unchanged | Q9D6Y7 | 14 | 216 | 14 | 233 | 26 | 8.41 | 73 | 1.199 | 0.194 | 0.999 | 0.993 | 1.132 | 0.329 | 0.882 | 0.253 |
| 259 | **Aldoa** | Unchanged | P05064 | 17 | 180 | 6 | 364 | 39.3 | 8.09 | 73 | 0.656 | 0.305 | 1.346 | 0.449 | 1.143 | 0.708 | 1.178 | 0.610 |
| 260 | **Gsto1** | Unchanged | O09131 | 14 | 121 | 14 | 240 | 27.5 | 7.36 | 73 | 0.972 | 0.896 | 1.209 | 0.534 | 1.440 | 0.339 | 0.840 | 0.657 |
| 261 | **Pdxk** | Unchanged | Q8K183 | 15 | 210 | 15 | 312 | 35 | 6.29 | 72 | 1.445 | 0.037 | 1.579 | 0.099 | 0.997 | 0.980 | 1.585 | 0.089 |
| 262 | **Hao1** | Unchanged | Q9WU19 | 22 | 334 | 22 | 370 | 41 | 7.72 | 72 | 0.945 | 0.814 | 1.371 | 0.085 | 0.927 | 0.696 | 1.478 | 0.005 |
| 263 | **Tubb5** | Unchanged | P99024 | 28 | 496 | 5 | 444 | 49.6 | 4.89 | 72 | 0.651 | 0.303 | 0.628 | 0.329 | 0.670 | 0.359 | 0.936 | 0.847 |
| 264 | **Aldh1a7** | UP | O35945 | 34 | 903 | 21 | 501 | 54.6 | 7.71 | 72 | 1.374 | 0.043 | 1.560 | 0.031 | 1.696 | 0.015 | 0.920 | 0.608 |
| 265 | **Psma1** | Unchanged | Q9R1P4 | 16 | 251 | 16 | 263 | 29.5 | 6.46 | 72 | 0.742 | 0.425 | 0.821 | 0.572 | 0.786 | 0.430 | 1.044 | 0.866 |
| 266 | **Tubb4b** | Unchanged | P68372 | 28 | 570 | 3 | 445 | 49.8 | 4.89 | 72 | 0.976 | 0.919 | 0.948 | 0.854 | 0.930 | 0.794 | 1.019 | 0.957 |
| 267 | **Tubb2a** | UP | Q7TMM9 | 28 | 543 | 6 | 445 | 49.9 | 4.89 | 72 | 1.469 | 0.265 | 2.486 | 0.017 | 1.538 | 0.153 | 1.617 | 0.090 |
| 268 | **Atic** | Unchanged | Q9CWJ9 | 34 | 356 | 34 | 592 | 64.2 | 6.76 | 70 | 0.920 | 0.575 | 1.165 | 0.559 | 0.994 | 0.980 | 1.172 | 0.616 |
| 269 | **Fasn** | Unchanged | P19096 | 134 | 3026 | 134 | 2504 | 272.3 | 6.58 | 72 | 0.855 | 0.295 | 0.957 | 0.780 | 0.902 | 0.570 | 1.062 | 0.779 |
| 270 | **Ugt2b5** | Unchanged | Q8K169 | 31 | 349 | 27 | 530 | 60.8 | 7.87 | 72 | 0.862 | 0.350 | 0.906 | 0.771 | 1.065 | 0.783 | 0.850 | 0.668 |
| 271 | **Akr1d1** | Unchanged | Q8VCX1 | 24 | 303 | 22 | 325 | 37.3 | 6.9 | 72 | 1.405 | 0.210 | 1.047 | 0.859 | 1.148 | 0.646 | 0.912 | 0.786 |
| 272 | **Psmd7** | Unchanged | P26516 | 16 | 168 | 16 | 321 | 36.5 | 6.77 | 70 | 0.806 | 0.565 | 0.815 | 0.556 | 0.717 | 0.433 | 1.137 | 0.587 |
| 273 | **Gsta3iso** | Unchanged | P30115 | 25 | 1566 | 12 | 221 | 25.3 | 8.73 | 72 | 1.210 | 0.220 | 0.738 | 0.211 | 1.443 | 0.203 | 0.512 | 0.074 |
| 274 | **Ndrg2** | Unchanged | A0A2I3BQM5 | 5 | 55 | 1 | 114 | 12.7 | 5 | 72 | 3.587 | 0.292 | 3.260 | 0.112 | 1.569 | 0.610 | 2.077 | 0.223 |
| 275 | **Sds** | Unchanged | Q8VBT2 | 14 | 257 | 14 | 327 | 34.6 | 7.12 | 72 | 1.013 | 0.958 | 0.994 | 0.987 | 0.927 | 0.758 | 1.072 | 0.869 |
| 276 | **Hba** | Unchanged | P01942 | 10 | 548 | 1 | 142 | 15.1 | 8.22 | 64 | 1.369 | 0.697 | 0.648 | 0.516 | 0.640 | 0.400 | 1.013 | 0.987 |

| | | | | | | | | | | | | | | | | | | |
|---|---|---|---|---|---|---|---|---|---|---|---|---|---|---|---|---|---|---|
| 277 | **Fh** | Unchanged | P97807 | 26 | 614 | 26 | 507 | 54.3 | 9.04 | 72 | 1.016 | 0.905 | 1.161 | 0.431 | 0.942 | 0.663 | 1.233 | 0.204 |
| 278 | **Acsf2** | Unchanged | Q8VCW8 | 40 | 710 | 40 | 615 | 67.9 | 8.18 | 72 | 1.280 | 0.090 | 0.992 | 0.980 | 1.106 | 0.512 | 0.897 | 0.735 |
| 279 | **Ttpa** | Unchanged | Q8BWP5 | 16 | 235 | 16 | 278 | 32 | 8.37 | 72 | 1.096 | 0.361 | 1.102 | 0.531 | 0.917 | 0.497 | 1.201 | 0.337 |
| 280 | **Gpt** | Unchanged | Q8QZR5 | 28 | 512 | 27 | 496 | 55.1 | 6.64 | 72 | 0.482 | 0.211 | 0.514 | 0.242 | 0.561 | 0.281 | 0.916 | 0.487 |
| 281 | **Rps14** | Unchanged | P62264 | 12 | 171 | 12 | 151 | 16.3 | 10.05 | 72 | 0.648 | 0.219 | 0.864 | 0.703 | 0.767 | 0.401 | 1.128 | 0.740 |
| 282 | **Acox2** | UP | Q9QXD1 | 37 | 397 | 37 | 681 | 76.8 | 7.46 | 72 | 1.174 | 0.197 | 2.667 | 0.043 | 1.299 | 0.075 | 2.053 | 0.087 |
| 283 | **Sdha** | Unchanged | Q8K2B3 | 33 | 564 | 33 | 664 | 72.5 | 7.37 | 71 | 0.828 | 0.080 | 1.293 | 0.245 | 0.684 | 0.011 | 1.890 | 0.038 |
| 284 | **Ech1** | Unchanged | O35459 | 19 | 241 | 19 | 327 | 36.1 | 7.71 | 71 | 1.136 | 0.405 | 1.011 | 0.955 | 0.979 | 0.869 | 1.033 | 0.859 |
| 285 | **Mthfd1** | Unchanged | Q922D8 | 48 | 717 | 48 | 935 | 101.1 | 7.14 | 71 | 1.060 | 0.312 | 0.823 | 0.001 | 0.907 | 0.196 | 0.908 | 0.212 |
| 286 | **Agmat** | Unchanged | A2AS89 | 13 | 177 | 13 | 358 | 38.2 | 7.87 | 71 | 1.052 | 0.927 | 1.068 | 0.920 | 1.111 | 0.818 | 0.961 | 0.945 |
| 287 | **Ca2** | Unchanged | P00920 | 11 | 179 | 11 | 260 | 29 | 7.01 | 71 | 1.200 | 0.489 | 1.078 | 0.789 | 0.942 | 0.794 | 1.145 | 0.552 |
| 288 | **Cox6b1** | Unchanged | P56391 | 8 | 86 | 8 | 86 | 10.1 | 8.72 | 71 | 1.332 | 0.592 | 1.247 | 0.733 | 1.405 | 0.498 | 0.887 | 0.825 |
| 289 | **Hdhd3** | Unchanged | Q9CYW4 | 10 | 80 | 10 | 251 | 28 | 6.8 | 71 | 1.126 | 0.739 | 1.212 | 0.565 | 1.263 | 0.410 | 0.960 | 0.843 |
| 290 | **Nit1** | Unchanged | Q8VDK1 | 16 | 200 | 16 | 323 | 35.7 | 7.94 | 71 | 1.312 | 0.130 | 1.066 | 0.814 | 0.917 | 0.520 | 1.163 | 0.615 |
| 291 | **Ddah1** | UP | Q9CWS0 | 14 | 133 | 13 | 285 | 31.4 | 5.97 | 71 | 2.172 | 0.034 | 2.855 | 0.206 | 6.411 | 0.016 | 0.445 | 0.148 |
| 292 | **Hsp90ab1** | Unchanged | P11499 | 57 | 1221 | 35 | 724 | 83.2 | 5.03 | 65 | 1.065 | 0.674 | 1.270 | 0.293 | 0.956 | 0.679 | 1.329 | 0.203 |
| 293 | **Ak2 iso** | Unchanged | F7BP55 | 4 | 53 | 1 | 72 | 7.4 | 8.94 | 71 | 0.830 | 0.632 | 0.912 | 0.844 | 0.959 | 0.902 | 0.951 | 0.899 |
| 294 | **Khk** | Unchanged | A0A0J9YU79 | 19 | 248 | 3 | 298 | 32.7 | 6.16 | 71 | 1.200 | 0.797 | 1.546 | 0.425 | 0.890 | 0.886 | 1.738 | 0.256 |
| 295 | **Acsm1** | Unchanged | Q91VA0 | 34 | 788 | 34 | 573 | 64.7 | 7.17 | 69 | 0.946 | 0.718 | 1.086 | 0.730 | 1.021 | 0.888 | 1.063 | 0.768 |
| 296 | **Cct2** | Unchanged | P80314 | 28 | 375 | 28 | 535 | 57.4 | 6.4 | 71 | 1.182 | 0.501 | 1.184 | 0.482 | 1.455 | 0.048 | 0.813 | 0.305 |
| 297 | **Hacl1** | Unchanged | Q9QXE0 | 29 | 616 | 29 | 581 | 63.6 | 6.27 | 67 | 0.844 | 0.389 | 1.093 | 0.704 | 1.026 | 0.891 | 1.065 | 0.782 |
| 298 | **Hal** | Unchanged | P35492 | 35 | 524 | 35 | 657 | 72.2 | 6.34 | 70 | 1.326 | 0.389 | 1.652 | 0.259 | 1.232 | 0.538 | 1.341 | 0.485 |
| 299 | **Cyp2c37** | Down | P56654 | 28 | 392 | 5 | 490 | 55.6 | 7.11 | 70 | 1.052 | 0.830 | 0.442 | 0.005 | 0.250 | 0.000 | 1.770 | 0.023 |
| 300 | **Khkiso** | Unchanged | P97328 | 20 | 435 | 2 | 298 | 32.7 | 6.16 | 67 | 0.707 | 0.142 | 0.987 | 0.965 | 0.808 | 0.320 | 1.222 | 0.588 |
| 301 | **Shmt1** | Unchanged | P50431 | 29 | 447 | 28 | 478 | 52.6 | 6.95 | 70 | 1.031 | 0.915 | 0.873 | 0.704 | 1.016 | 0.952 | 0.859 | 0.656 |
| 302 | **Gulo** | Unchanged | P58710 | 27 | 418 | 27 | 440 | 50.4 | 8.24 | 70 | 0.948 | 0.897 | 0.965 | 0.942 | 0.542 | 0.209 | 1.779 | 0.276 |
| 303 | **Dhdh** | Unchanged | Q9DBB8 | 17 | 255 | 17 | 333 | 36.3 | 6.44 | 70 | 1.297 | 0.241 | 0.896 | 0.686 | 1.087 | 0.697 | 0.824 | 0.340 |
| 304 | **Pnp** | Unchanged | P23492 | 17 | 280 | 17 | 289 | 32.3 | 6.16 | 70 | 0.806 | 0.326 | 1.213 | 0.468 | 1.193 | 0.342 | 1.017 | 0.943 |
| 305 | **Actn4** | Unchanged | P57780 | 55 | 759 | 29 | 912 | 104.9 | 5.41 | 70 | 0.959 | 0.907 | 0.889 | 0.763 | 1.283 | 0.496 | 0.693 | 0.385 |
| 306 | **Ndufa5** | Unchanged | Q9CPP6 | 8 | 99 | 8 | 116 | 13.4 | 8.1 | 70 | 1.288 | 0.568 | 1.222 | 0.215 | 1.088 | 0.713 | 1.124 | 0.484 |
| 307 | **Eef1b** | Unchanged | O70251 | 11 | 227 | 11 | 225 | 24.7 | 4.69 | 67 | 1.053 | 0.876 | 0.931 | 0.795 | 1.030 | 0.926 | 0.904 | 0.627 |
| 308 | **Hprt1** | Unchanged | P00493 | 10 | 101 | 10 | 218 | 24.6 | 6.68 | 70 | 0.924 | 0.778 | 1.499 | 0.194 | 0.659 | 0.083 | 2.275 | 0.029 |
| 309 | **Eef2** | Unchanged | P58252 | 60 | 1305 | 59 | 858 | 95.3 | 6.83 | 69 | 0.859 | 0.217 | 0.965 | 0.842 | 1.051 | 0.696 | 0.918 | 0.663 |
| 310 | **Acly** | Unchanged | Q3V117 | 59 | 602 | 59 | 1101 | 120.7 | 7.44 | 69 | 0.946 | 0.783 | 1.040 | 0.754 | 1.124 | 0.465 | 0.926 | 0.506 |
| 311 | **Aldh7a1** | Unchanged | Q9DBF1 | 28 | 721 | 28 | 539 | 58.8 | 7.47 | 69 | 0.897 | 0.223 | 0.899 | 0.534 | 0.908 | 0.102 | 0.989 | 0.954 |
| 312 | **Eef1g** | Unchanged | Q9D8N0 | 25 | 379 | 25 | 437 | 50 | 6.74 | 69 | 0.768 | 0.219 | 0.776 | 0.350 | 1.138 | 0.560 | 0.682 | 0.149 |
| 313 | **Cbr1** | Unchanged | P48758 | 18 | 302 | 11 | 277 | 30.6 | 8.31 | 69 | 1.554 | 0.492 | 1.301 | 0.561 | 1.713 | 0.071 | 0.759 | 0.445 |
| 314 | **Hspa9** | Unchanged | P38647 | 44 | 1052 | 43 | 679 | 73.4 | 6.07 | 69 | 1.093 | 0.497 | 1.072 | 0.730 | 1.222 | 0.180 | 0.877 | 0.536 |
| 315 | **Lcp1** | Unchanged | Q61233 | 38 | 439 | 31 | 627 | 70.1 | 5.33 | 69 | 0.919 | 0.457 | 1.214 | 0.339 | 1.285 | 0.005 | 0.944 | 0.746 |
| 316 | **Eci1** | Unchanged | P42125 | 17 | 296 | 17 | 289 | 32.2 | 8.98 | 69 | 1.310 | 0.021 | 1.103 | 0.487 | 0.920 | 0.484 | 1.199 | 0.202 |
| 317 | **Agl** | Unchanged | F8VPN4 | 70 | 435 | 70 | 1532 | 174.2 | 6.74 | 69 | 1.006 | 0.970 | 0.764 | 0.164 | 0.752 | 0.161 | 1.017 | 0.908 |

| | | | | | | | | | | | | | | | | | |
|---|---|---|---|---|---|---|---|---|---|---|---|---|---|---|---|---|---|
| 318 | **Paics** | Unchanged | Q9DCL9 | 20 | 199 | 20 | 425 | 47 | 7.23 | 69 | 1.049 | 0.785 | 1.321 | 0.173 | 1.309 | 0.079 | 1.010 | 0.941 |
| 319 | **Slc25a20** | UP | Q9Z2Z6 | 16 | 172 | 16 | 301 | 33 | 9.11 | 69 | 2.898 | 0.006 | 2.860 | 0.016 | 2.889 | 0.001 | 0.990 | 0.961 |
| 320 | **Ces2a** | Unchanged | Q8QZR3 | 23 | 403 | 18 | 558 | 61.9 | 6.16 | 69 | 1.022 | 0.944 | 1.077 | 0.877 | 1.211 | 0.626 | 0.889 | 0.836 |
| 321 | **Ndufa12** | Unchanged | Q7TMF3 | 7 | 93 | 7 | 145 | 17.1 | 9.36 | 69 | 1.726 | 0.297 | 1.773 | 0.086 | 1.948 | 0.173 | 0.910 | 0.815 |
| 322 | **Mcfd2** | Unchanged | Q8K5B2 | 5 | 119 | 5 | 145 | 16.2 | 4.72 | 69 | 0.743 | 0.613 | 0.698 | 0.529 | 0.805 | 0.724 | 0.867 | 0.809 |
| 323 | **Hebp1iso** | Unchanged | Q9R257 | 12 | 222 | 1 | 190 | 21.1 | 5.26 | 69 | 0.777 | 0.797 | 5.013 | 0.149 | 8.848 | 0.062 | 0.567 | 0.403 |
| 324 | **Fabp4** | Unchanged | P04117 | 10 | 45 | 10 | 132 | 14.6 | 8.4 | 69 | 0.703 | 0.242 | 1.548 | 0.453 | 1.687 | 0.115 | 0.917 | 0.857 |
| 325 | **Hmgcs2** | Unchanged | P54869 | 37 | 1890 | 35 | 508 | 56.8 | 8.41 | 69 | 1.255 | 0.593 | 1.067 | 0.868 | 1.015 | 0.971 | 1.052 | 0.894 |
| 326 | **Hsp90aa1** | Unchanged | P07901 | 57 | 849 | 37 | 733 | 84.7 | 5.01 | 63 | 0.944 | 0.698 | 0.808 | 0.068 | 1.014 | 0.924 | 0.797 | 0.186 |
| 327 | **Por** | Unchanged | P37040 | 36 | 500 | 36 | 678 | 77 | 5.53 | 69 | 0.963 | 0.885 | 1.147 | 0.668 | 1.173 | 0.524 | 0.978 | 0.938 |
| 328 | **Adk** | Unchanged | P55264 | 29 | 752 | 29 | 361 | 40.1 | 6.21 | 69 | 1.254 | 0.018 | 1.062 | 0.572 | 0.956 | 0.666 | 1.111 | 0.326 |
| 329 | **Hspe1** | UP | Q64433 | 9 | 165 | 9 | 102 | 11 | 8.35 | 69 | 1.657 | 0.133 | 1.558 | 0.247 | 2.004 | 0.033 | 0.777 | 0.453 |
| 330 | **Atp5f1a** | Unchanged | Q03265 | 37 | 1322 | 37 | 553 | 59.7 | 9.19 | 69 | 1.054 | 0.456 | 0.872 | 0.384 | 0.938 | 0.626 | 0.930 | 0.700 |
| 331 | **Fmo5** | Unchanged | P97872 | 29 | 511 | 29 | 533 | 60 | 8.73 | 68 | 1.068 | 0.642 | 1.172 | 0.605 | 1.321 | 0.217 | 0.887 | 0.694 |
| 332 | **Mup3** | Down | Q3KQQ2 | 13 | 97 | 11 | 184 | 21.4 | 4.75 | 68 | 0.648 | 0.102 | 0.815 | 0.399 | 0.442 | 0.004 | 1.846 | 0.052 |
| 333 | **Haghiso** | Unchanged | Q99KB8 | 13 | 280 | 1 | 309 | 34.1 | 7.75 | 68 | 1.035 | 0.835 | 1.285 | 0.485 | 1.039 | 0.881 | 1.237 | 0.594 |
| 334 | **Sec14l4** | Unchanged | Q8R0F9 | 25 | 296 | 25 | 403 | 46 | 7.12 | 68 | 1.242 | 0.221 | 1.799 | 0.178 | 1.072 | 0.749 | 1.677 | 0.239 |
| 335 | **Txndc5** | Unchanged | Q91W90 | 20 | 260 | 20 | 417 | 46.4 | 5.78 | 68 | 1.198 | 0.156 | 1.105 | 0.749 | 1.123 | 0.435 | 0.984 | 0.956 |
| 336 | **Sord** | Unchanged | Q64442 | 27 | 992 | 27 | 357 | 38.2 | 7.02 | 68 | 0.944 | 0.516 | 0.790 | 0.018 | 0.721 | 0.027 | 1.097 | 0.547 |
| 337 | **Hsd17b8** | Unchanged | P50171 | 11 | 132 | 11 | 259 | 26.6 | 6.54 | 68 | 1.109 | 0.748 | 0.340 | 0.076 | 0.791 | 0.529 | 0.430 | 0.219 |
| 338 | **Glo1** | Unchanged | Q9CPU0 | 14 | 292 | 14 | 184 | 20.8 | 5.47 | 68 | 1.101 | 0.612 | 1.368 | 0.228 | 1.208 | 0.298 | 1.133 | 0.616 |
| 339 | **Uroc1** | UP | Q8VC12 | 33 | 583 | 33 | 676 | 74.5 | 7.53 | 68 | 1.297 | 0.062 | 1.966 | 0.050 | 1.597 | 0.001 | 1.231 | 0.418 |
| 340 | **Hadhb** | Unchanged | Q99JY0 | 31 | 739 | 31 | 475 | 51.4 | 9.38 | 68 | 1.320 | 0.060 | 1.228 | 0.203 | 1.553 | 0.168 | 0.791 | 0.405 |
| 341 | **Psme2** | UP | P97372 | 14 | 203 | 14 | 239 | 27 | 5.76 | 68 | 1.169 | 0.552 | 1.712 | 0.033 | 1.416 | 0.164 | 1.210 | 0.251 |
| 342 | **Pls3** | Unchanged | Q99K51 | 37 | 370 | 29 | 630 | 70.7 | 5.62 | 68 | 1.148 | 0.310 | 1.315 | 0.021 | 1.233 | 0.128 | 1.066 | 0.594 |
| 343 | **Hsd17b4** | Unchanged | P51660 | 37 | 665 | 37 | 735 | 79.4 | 8.57 | 68 | 1.242 | 0.180 | 1.230 | 0.212 | 1.155 | 0.335 | 1.065 | 0.670 |
| 344 | **Sec61b** | Unchanged | Q9CQS8 | 5 | 16 | 5 | 96 | 10 | 11.56 | 52 | 0.908 | 0.882 | 0.192 | 0.098 | 0.487 | 0.329 | 0.394 | 0.307 |
| 345 | **Mettl7b** | Unchanged | Q9DD20 | 14 | 171 | 14 | 244 | 28 | 8.59 | 68 | 1.352 | 0.383 | 1.465 | 0.484 | 0.971 | 0.948 | 1.508 | 0.430 |
| 346 | **Ldhd** | Down | Q7TNG8 | 19 | 276 | 19 | 484 | 51.8 | 6.62 | 68 | 0.784 | 0.303 | 0.941 | 0.762 | 0.625 | 0.039 | 1.506 | 0.074 |
| 347 | **Bpnt1** | Unchanged | Q9Z0S1 | 14 | 164 | 14 | 308 | 33.2 | 5.82 | 68 | 0.875 | 0.522 | 1.346 | 0.398 | 0.777 | 0.219 | 1.733 | 0.148 |
| 348 | **Eef1a1iso** | Unchanged | P10126 | 33 | 1487 | 6 | 462 | 50.1 | 9.01 | 68 | 0.866 | 0.274 | 0.855 | 0.238 | 0.939 | 0.585 | 0.911 | 0.425 |
| 349 | **Slc25a11** | Unchanged | Q9CR62 | 15 | 135 | 15 | 314 | 34.1 | 9.94 | 68 | 1.543 | 0.271 | 1.197 | 0.591 | 1.767 | 0.388 | 0.677 | 0.520 |
| 350 | **Dynll1** | Unchanged | P63168 | 3 | 33 | 1 | 89 | 10.4 | 7.4 | 67 | 0.682 | 0.649 | 1.261 | 0.742 | 1.094 | 0.881 | 1.153 | 0.773 |
| 351 | **Atp5po** | Unchanged | Q9DB20 | 14 | 261 | 14 | 213 | 23.3 | 9.99 | 67 | 1.007 | 0.983 | 0.998 | 0.997 | 1.040 | 0.920 | 0.960 | 0.942 |
| 352 | **Ppib** | Unchanged | P24369 | 20 | 329 | 20 | 216 | 23.7 | 9.55 | 67 | 0.656 | 0.070 | 0.900 | 0.667 | 0.994 | 0.980 | 0.905 | 0.728 |
| 353 | **Slc25a15** | Unchanged | Q9WVD5 | 15 | 149 | 15 | 301 | 32.8 | 8.85 | 67 | 0.783 | 0.434 | 1.106 | 0.709 | 0.752 | 0.392 | 1.471 | 0.265 |
| 354 | **Ndufa9** | Unchanged | Q9DC69 | 20 | 215 | 20 | 377 | 42.5 | 9.74 | 67 | 0.993 | 0.976 | 1.121 | 0.715 | 0.816 | 0.421 | 1.373 | 0.287 |
| 355 | **Aldh4a1** | Unchanged | Q8CHT0 | 31 | 797 | 31 | 562 | 61.8 | 8.24 | 67 | 1.142 | 0.417 | 0.822 | 0.343 | 0.839 | 0.265 | 0.981 | 0.938 |
| 356 | **Scrn2** | Unchanged | Q8VCA8 | 16 | 205 | 16 | 425 | 46.6 | 5.71 | 67 | 1.696 | 0.440 | 3.521 | 0.201 | 0.688 | 0.196 | 5.117 | 0.154 |
| 357 | **Atp5mg** | Unchanged | Q9CPQ8 | 6 | 139 | 6 | 103 | 11.4 | 9.74 | 67 | 1.142 | 0.443 | 0.866 | 0.525 | 1.181 | 0.343 | 0.733 | 0.164 |
| 358 | **Ugt2b36** | UP | Q3UEP4 | 25 | 287 | 22 | 530 | 61 | 8.25 | 67 | 1.067 | 0.698 | 1.342 | 0.407 | 1.546 | 0.023 | 0.868 | 0.614 |

| | | | | | | | | | | | | | | | | | | |
|---|---|---|---|---|---|---|---|---|---|---|---|---|---|---|---|---|---|---|
| 359 | **Acy3** | Unchanged | Q91XE4 | 12 | 185 | 12 | 318 | 35.3 | 5.5 | 67 | 1.158 | 0.276 | 0.977 | 0.923 | 1.515 | 0.166 | 0.645 | 0.175 |
| 360 | **Cyp27a1** | Down | Q9DBG1 | 28 | 283 | 28 | 533 | 60.7 | 9.14 | 67 | 0.872 | 0.592 | 0.961 | 0.796 | 0.578 | 0.014 | 1.664 | 0.023 |
| 361 | **Rps4x** | Unchanged | P62702 | 23 | 306 | 23 | 263 | 29.6 | 10.15 | 67 | 0.904 | 0.430 | 1.034 | 0.820 | 1.376 | 0.238 | 0.752 | 0.324 |
| 362 | **Cndp2** | Unchanged | Q9D1A2 | 22 | 236 | 22 | 475 | 52.7 | 5.66 | 67 | 0.832 | 0.464 | 0.876 | 0.534 | 1.145 | 0.577 | 0.765 | 0.296 |
| 363 | **Csad** | Unchanged | Q9DBE0 | 28 | 485 | 28 | 493 | 55.1 | 6.61 | 67 | 0.993 | 0.980 | 1.492 | 0.351 | 1.693 | 0.189 | 0.881 | 0.762 |
| 364 | **Tpmt** | Unchanged | O55060 | 14 | 148 | 14 | 240 | 27.6 | 6.44 | 67 | 1.097 | 0.773 | 1.764 | 0.176 | 1.034 | 0.913 | 1.706 | 0.179 |
| 365 | **Cbs** | Unchanged | Q91WT9 | 27 | 465 | 27 | 561 | 61.5 | 6.48 | 62 | 1.092 | 0.592 | 1.083 | 0.615 | 1.433 | 0.247 | 0.756 | 0.359 |
| 366 | **Rbp1** | UP | Q00915 | 8 | 52 | 8 | 135 | 15.8 | 5.25 | 60 | 1.418 | 0.559 | 3.019 | 0.031 | 2.326 | 0.040 | 1.298 | 0.443 |
| 367 | **Rplp1** | Unchanged | P47955 | 3 | 134 | 3 | 114 | 11.5 | 4.32 | 67 | 0.957 | 0.846 | 1.191 | 0.670 | 0.829 | 0.510 | 1.437 | 0.444 |
| 368 | **Gpi** | Unchanged | P06745 | 31 | 472 | 31 | 558 | 62.7 | 8.13 | 67 | 0.925 | 0.646 | 1.792 | 0.055 | 1.468 | 0.088 | 1.221 | 0.412 |
| 369 | **Mdh2** | Unchanged | P08249 | 21 | 504 | 21 | 338 | 35.6 | 8.68 | 67 | 0.827 | 0.673 | 0.994 | 0.989 | 0.890 | 0.787 | 1.117 | 0.795 |
| 370 | **Ndufv1** | Unchanged | Q91YT0 | 23 | 250 | 23 | 464 | 50.8 | 8.21 | 66 | 0.940 | 0.696 | 1.106 | 0.539 | 1.031 | 0.876 | 1.074 | 0.702 |
| 371 | **Ces3a** | Unchanged | Q63880 | 24 | 792 | 13 | 571 | 63.3 | 6.14 | 66 | 0.818 | 0.495 | 0.951 | 0.916 | 0.887 | 0.743 | 1.072 | 0.903 |
| 372 | **Mecr** | Unchanged | Q9DCS3 | 14 | 187 | 14 | 373 | 40.3 | 9.07 | 66 | 1.058 | 0.863 | 1.277 | 0.490 | 1.143 | 0.623 | 1.118 | 0.758 |
| 373 | **Atp5me** | Unchanged | Q06185 | 6 | 91 | 6 | 71 | 8.2 | 9.35 | 66 | 2.162 | 0.092 | 1.661 | 0.267 | 2.022 | 0.081 | 0.822 | 0.638 |
| 374 | **Rab1b** | Unchanged | Q9D1G1 | 11 | 117 | 5 | 201 | 22.2 | 5.73 | 66 | 1.120 | 0.729 | 1.358 | 0.410 | 1.551 | 0.372 | 0.876 | 0.775 |
| 375 | **Hist2h2aa1** | Unchanged | Q6GSS7 | 9 | 114 | 5 | 130 | 14.1 | 10.9 | 66 | 3.278 | 0.218 | 5.353 | 0.204 | 43.840 | 0.113 | 0.122 | 0.151 |
| 376 | **Hist1h2ae** | Unchanged | C0HKE4 | 9 | 128 | 5 | 130 | 14.1 | 11.05 | 66 | 1.259 | 0.355 | 1.113 | 0.557 | 20.240 | 0.205 | 0.055 | 0.207 |
| 377 | **Anxa6** | Unchanged | P14824 | 41 | 747 | 2 | 673 | 75.8 | 5.5 | 66 | 0.936 | 0.352 | 1.228 | 0.004 | 1.193 | 0.034 | 1.029 | 0.607 |
| 378 | **Aadac** | Unchanged | Q99PG0 | 26 | 465 | 26 | 398 | 45.2 | 7.49 | 66 | 1.191 | 0.121 | 1.247 | 0.251 | 1.288 | 0.055 | 0.968 | 0.853 |
| 379 | **Tf** | Unchanged | Q921I1 | 54 | 1016 | 2 | 697 | 76.7 | 7.18 | 66 | 0.829 | 0.756 | 1.454 | 0.606 | 2.632 | 0.264 | 0.552 | 0.437 |
| 380 | **Cnpy2** | Unchanged | Q9QXT0 | 10 | 120 | 10 | 182 | 20.8 | 5.07 | 66 | 0.630 | 0.053 | 0.650 | 0.099 | 0.911 | 0.707 | 0.713 | 0.240 |
| 381 | **Rplp0** | Unchanged | P14869 | 20 | 344 | 20 | 317 | 34.2 | 6.25 | 66 | 1.044 | 0.882 | 1.010 | 0.968 | 1.259 | 0.425 | 0.802 | 0.509 |
| 382 | **Gstk1** | Unchanged | Q9DCM2 | 14 | 275 | 14 | 226 | 25.7 | 8.88 | 66 | 1.174 | 0.049 | 1.463 | 0.434 | 0.866 | 0.060 | 1.690 | 0.317 |
| 383 | **Adi1** | Unchanged | Q99JT9 | 11 | 53 | 11 | 179 | 21.5 | 5.5 | 58 | 1.072 | 0.867 | 1.293 | 0.608 | 1.163 | 0.792 | 1.112 | 0.861 |
| 384 | **Hsd17b13** | UP | A8Y5N4 | 16 | 139 | 2 | 264 | 29.4 | 9.58 | 66 | 2.026 | 0.003 | 3.214 | 0.099 | 1.863 | 0.079 | 1.725 | 0.307 |
| 385 | **Pipox** | UP | Q9D826 | 19 | 336 | 19 | 390 | 43.8 | 7.55 | 66 | 1.184 | 0.300 | 1.739 | 0.034 | 1.724 | 0.007 | 1.009 | 0.960 |
| 386 | **Cyp1a2** | Unchanged | P00186 | 28 | 302 | 28 | 513 | 58.1 | 8.76 | 66 | 1.037 | 0.852 | 1.303 | 0.159 | 0.917 | 0.578 | 1.422 | 0.075 |
| 387 | **Acadl** | UP | P51174 | 26 | 449 | 26 | 430 | 47.9 | 8.31 | 66 | 1.580 | 0.010 | 1.212 | 0.292 | 1.425 | 0.021 | 0.850 | 0.307 |
| 388 | **Ube2n** | Unchanged | P61089 | 6 | 98 | 6 | 152 | 17.1 | 6.57 | 66 | 0.297 | 0.180 | 0.298 | 0.184 | 0.378 | 0.231 | 0.789 | 0.379 |
| 389 | **Ethe1** | UP | Q9DCM0 | 10 | 137 | 10 | 254 | 27.7 | 7.23 | 66 | 1.105 | 0.424 | 0.994 | 0.972 | 1.556 | 0.000 | 0.639 | 0.008 |
| 390 | **Ywhaz** | Unchanged | P63101 | 16 | 364 | 14 | 245 | 27.8 | 4.79 | 66 | 0.906 | 0.546 | 1.228 | 0.663 | 1.417 | 0.340 | 0.867 | 0.771 |
| 391 | **Rpl9** | Unchanged | P51410 | 11 | 222 | 11 | 192 | 21.9 | 9.95 | 65 | 1.018 | 0.968 | 0.910 | 0.837 | 0.759 | 0.553 | 1.199 | 0.671 |
| 392 | **Clic4** | Unchanged | Q9QYB1 | 12 | 115 | 12 | 253 | 28.7 | 5.59 | 66 | 1.327 | 0.179 | 1.750 | 0.155 | 1.100 | 0.608 | 1.592 | 0.222 |
| 393 | **Fmo1** | UP | P50285 | 25 | 247 | 25 | 532 | 59.9 | 8.47 | 66 | 1.342 | 0.302 | 1.813 | 0.035 | 1.593 | 0.027 | 1.138 | 0.474 |
| 394 | **Hmgcl** | Unchanged | P38060 | 18 | 349 | 18 | 325 | 34.2 | 8.41 | 66 | 1.158 | 0.350 | 0.790 | 0.180 | 1.120 | 0.260 | 0.705 | 0.053 |
| 395 | **Actn4 iso** | Unchanged | D3Z0L8 | 12 | 139 | 1 | 206 | 23.3 | 7.97 | 66 | 3.006 | 0.314 | 1.202 | 0.584 | 2.989 | 0.139 | 0.402 | 0.169 |
| 396 | **Psma4** | Unchanged | Q9R1P0 | 12 | 130 | 12 | 261 | 29.5 | 7.72 | 66 | 0.805 | 0.445 | 0.918 | 0.772 | 0.578 | 0.108 | 1.588 | 0.086 |
| 397 | **Dbiiso** | Unchanged | P31786 | 10 | 323 | 4 | 87 | 10 | 8.82 | 66 | 0.751 | 0.033 | 1.083 | 0.832 | 1.103 | 0.534 | 0.982 | 0.962 |
| 398 | **Vapa** | Unchanged | Q9WV55 | 11 | 142 | 10 | 249 | 27.8 | 8.4 | 65 | 1.532 | 0.324 | 1.999 | 0.113 | 1.111 | 0.614 | 1.800 | 0.164 |
| 399 | **Aox3** | Unchanged | G3X982 | 59 | 996 | 58 | 1335 | 146.8 | 6.46 | 65 | 0.767 | 0.092 | 0.784 | 0.182 | 0.726 | 0.078 | 1.079 | 0.619 |

| | | | | | | | | | | | | | | | | | | |
|---|---|---|---|---|---|---|---|---|---|---|---|---|---|---|---|---|---|---|
| 400 | **Cmbl** | Unchanged | Q8R1G2 | 14 | 205 | 14 | 245 | 27.9 | 7.18 | 65 | 0.856 | 0.382 | 0.811 | 0.230 | 0.789 | 0.396 | 1.028 | 0.926 |
| 401 | **Adh4** | Unchanged | Q9QYY9 | 16 | 139 | 16 | 377 | 40.2 | 7.83 | 65 | 1.185 | 0.565 | 1.408 | 0.356 | 1.034 | 0.913 | 1.361 | 0.394 |
| 402 | **Atp5pb** | Unchanged | Q9CQQ7 | 16 | 243 | 16 | 256 | 28.9 | 9.06 | 62 | 1.283 | 0.608 | 1.912 | 0.249 | 1.447 | 0.404 | 1.322 | 0.569 |
| 403 | **Hspa5** | Unchanged | P20029 | 43 | 1864 | 41 | 655 | 72.4 | 5.16 | 65 | 0.777 | 0.158 | 0.899 | 0.592 | 0.946 | 0.788 | 0.951 | 0.846 |
| 404 | **Serpina3k** | Down | A0A0R4J0I1 | 27 | 378 | 12 | 418 | 46.6 | 5.16 | 65 | 0.622 | 0.050 | 0.733 | 0.136 | 0.593 | 0.035 | 1.236 | 0.006 |
| 405 | **Txnl1** | Unchanged | Q8CDN6 | 15 | 109 | 15 | 289 | 32.2 | 4.96 | 57 | 1.095 | 0.805 | 1.125 | 0.691 | 0.960 | 0.899 | 1.172 | 0.636 |
| 406 | **Ywhab** | Unchanged | Q9CQV8 | 17 | 335 | 12 | 246 | 28.1 | 4.83 | 65 | 0.734 | 0.242 | 0.722 | 0.299 | 0.926 | 0.707 | 0.779 | 0.265 |
| 407 | **Ube2l3** | Unchanged | P68037 | 8 | 125 | 8 | 154 | 17.9 | 8.51 | 65 | 1.108 | 0.682 | 0.986 | 0.971 | 1.557 | 0.055 | 0.633 | 0.130 |
| 408 | **Anxa6 iso** | Unchanged | F8WIT2 | 40 | 395 | 1 | 667 | 75.2 | 5.55 | 65 | 1.281 | 0.727 | 1.430 | 0.589 | 1.230 | 0.762 | 1.162 | 0.818 |
| 409 | **Ivd** | Unchanged | Q9JHI5 | 19 | 311 | 19 | 424 | 46.3 | 8.29 | 65 | 1.022 | 0.902 | 0.790 | 0.421 | 0.742 | 0.063 | 1.065 | 0.853 |
| 410 | **Alad** | Unchanged | P10518 | 14 | 387 | 14 | 330 | 36 | 6.79 | 65 | 1.091 | 0.665 | 1.470 | 0.141 | 1.188 | 0.263 | 1.238 | 0.378 |
| 411 | **Gcdh** | Unchanged | Q60759 | 20 | 553 | 20 | 438 | 48.6 | 8.73 | 65 | 1.161 | 0.190 | 1.049 | 0.715 | 1.033 | 0.744 | 1.015 | 0.923 |
| 412 | **Ces3b** | Unchanged | Q8VCU1 | 21 | 517 | 11 | 571 | 63.3 | 5.92 | 65 | 0.826 | 0.462 | 1.121 | 0.818 | 0.818 | 0.560 | 1.370 | 0.598 |
| 413 | **Prodh** | Unchanged | Q9WU79 | 30 | 271 | 27 | 599 | 68 | 8.24 | 65 | 1.166 | 0.651 | 1.240 | 0.445 | 0.684 | 0.312 | 1.813 | 0.012 |
| 414 | **Cyp4a12a** | UP | Q91WL5 | 26 | 184 | 8 | 508 | 58.3 | 9.13 | 65 | 2.992 | 0.061 | 4.400 | 0.066 | 1.897 | 0.045 | 2.320 | 0.164 |
| 415 | **Slc27a5** | Unchanged | Q4LDG0 | 30 | 557 | 30 | 689 | 76.2 | 8.38 | 65 | 0.833 | 0.202 | 0.997 | 0.991 | 0.869 | 0.517 | 1.148 | 0.633 |
| 416 | **Mdh1** | Down | P14152 | 18 | 485 | 18 | 334 | 36.5 | 6.58 | 65 | 0.828 | 0.133 | 0.545 | 0.000 | 0.954 | 0.353 | 0.571 | 0.002 |
| 417 | **Psme1** | Unchanged | P97371 | 17 | 302 | 17 | 249 | 28.7 | 5.97 | 65 | 0.896 | 0.278 | 1.016 | 0.891 | 1.188 | 0.345 | 0.855 | 0.413 |
| 418 | **Ndufs6** | Unchanged | P52503 | 7 | 82 | 7 | 116 | 13 | 8.65 | 65 | 0.531 | 0.170 | 0.571 | 0.081 | 0.551 | 0.120 | 1.035 | 0.928 |
| 419 | **Ugt2b1** | Unchanged | Q8R084 | 25 | 262 | 22 | 529 | 60.1 | 8.37 | 65 | 0.827 | 0.538 | 0.916 | 0.845 | 0.690 | 0.309 | 1.327 | 0.626 |
| 420 | **Anxa2** | Unchanged | P07356 | 20 | 166 | 20 | 339 | 38.7 | 7.69 | 65 | 1.023 | 0.941 | 4.818 | 0.059 | 4.814 | 0.077 | 1.001 | 0.999 |
| 421 | **Psma7** | Unchanged | Q9Z2U0 | 15 | 242 | 15 | 248 | 27.8 | 8.46 | 65 | 1.072 | 0.695 | 1.462 | 0.248 | 1.244 | 0.412 | 1.175 | 0.631 |
| 422 | **Hsbp1** | Unchanged | Q9CQZ1 | 3 | 24 | 3 | 76 | 8.6 | 4.31 | 64 | 1.427 | 0.573 | 2.860 | 0.094 | 3.341 | 0.124 | 0.856 | 0.760 |
| 423 | **Hsp90b1** | Unchanged | P08113 | 64 | 1396 | 62 | 802 | 92.4 | 4.82 | 64 | 0.806 | 0.304 | 0.788 | 0.139 | 0.742 | 0.073 | 1.063 | 0.714 |
| 424 | **Aprt** | Unchanged | P08030 | 8 | 53 | 8 | 180 | 19.7 | 6.79 | 64 | 0.855 | 0.673 | 1.726 | 0.373 | 3.018 | 0.147 | 0.572 | 0.403 |
| 425 | **Nipsnap1** | Unchanged | O55125 | 17 | 316 | 17 | 284 | 33.3 | 9.44 | 64 | 1.473 | 0.076 | 1.465 | 0.189 | 1.325 | 0.131 | 1.105 | 0.667 |
| 426 | **Pcbd1** | Unchanged | P61458 | 7 | 88 | 7 | 104 | 12 | 6.8 | 64 | 0.824 | 0.537 | 0.801 | 0.591 | 0.995 | 0.986 | 0.805 | 0.633 |
| 427 | **Pkm** | Unchanged | P52480 | 27 | 310 | 26 | 531 | 57.8 | 7.47 | 64 | 0.807 | 0.282 | 2.121 | 0.185 | 1.401 | 0.290 | 1.514 | 0.414 |
| 428 | **Iqgap2** | Unchanged | Q3UQ44 | 77 | 798 | 74 | 1575 | 180.4 | 5.64 | 64 | 1.030 | 0.892 | 1.376 | 0.217 | 0.894 | 0.648 | 1.540 | 0.074 |
| 429 | **Pcca** | Down | Q91ZA3 | 42 | 481 | 42 | 724 | 79.9 | 7.25 | 64 | 0.672 | 0.003 | 0.756 | 0.042 | 0.432 | 0.000 | 1.751 | 0.004 |
| 430 | **Actn1** | Unchanged | Q7TPR4 | 44 | 403 | 26 | 892 | 103 | 5.38 | 64 | 0.898 | 0.525 | 1.288 | 0.397 | 1.135 | 0.663 | 1.135 | 0.712 |
| 431 | **Psmc3** | Unchanged | O88685 | 21 | 274 | 21 | 442 | 49.5 | 5.19 | 64 | 1.000 | 1.000 | 0.947 | 0.720 | 0.687 | 0.045 | 1.378 | 0.011 |
| 432 | **Rab7a** | Unchanged | P51150 | 12 | 140 | 12 | 207 | 23.5 | 6.7 | 64 | 0.935 | 0.836 | 1.033 | 0.909 | 0.907 | 0.756 | 1.139 | 0.592 |
| 433 | **Cyp2d22** | Unchanged | Q9JKY7 | 26 | 187 | 21 | 500 | 56.4 | 6.98 | 64 | 0.998 | 0.990 | 1.005 | 0.981 | 0.659 | 0.080 | 1.525 | 0.116 |
| 434 | **Rab14** | Unchanged | Q91V41 | 14 | 164 | 12 | 215 | 23.9 | 6.21 | 64 | 0.952 | 0.745 | 1.267 | 0.405 | 0.821 | 0.354 | 1.545 | 0.222 |
| 435 | **Marc1;Mosc1** | Unchanged | Q9CW42 | 20 | 300 | 16 | 340 | 38 | 8.59 | 58 | 0.781 | 0.638 | 1.261 | 0.692 | 0.873 | 0.798 | 1.444 | 0.557 |
| 436 | **Mpst** | Unchanged | Q99J99 | 14 | 134 | 13 | 297 | 33.1 | 6.47 | 64 | 0.916 | 0.738 | 1.306 | 0.532 | 1.151 | 0.579 | 1.134 | 0.739 |
| 437 | **Anxa4** | UP | P97429 | 20 | 179 | 19 | 319 | 35.9 | 5.57 | 64 | 0.901 | 0.684 | 1.632 | 0.014 | 1.337 | 0.170 | 1.221 | 0.190 |
| 438 | **Gcat** | UP | O88986 | 15 | 277 | 15 | 416 | 44.9 | 7.33 | 64 | 1.414 | 0.082 | 1.545 | 0.010 | 1.535 | 0.009 | 1.006 | 0.957 |
| 439 | **Cyp2e1** | Unchanged | Q05421 | 33 | 658 | 33 | 493 | 56.8 | 8.4 | 64 | 1.121 | 0.499 | 0.754 | 0.238 | 0.710 | 0.117 | 1.063 | 0.835 |
| 440 | **Ganab** | Unchanged | Q8BHN3 | 42 | 581 | 42 | 944 | 106.8 | 6.06 | 64 | 0.815 | 0.132 | 0.957 | 0.882 | 0.963 | 0.805 | 0.993 | 0.983 |

| | | | | | | | | | | | | | | | | | | |
|---|---|---|---|---|---|---|---|---|---|---|---|---|---|---|---|---|---|---|
| 441 | **Ugt1a9** | Unchanged | Q62452 | 23 | 411 | 12 | 528 | 60 | 8.35 | 64 | 1.048 | 0.909 | 1.357 | 0.588 | 1.598 | 0.169 | 0.849 | 0.647 |
| 442 | **Glul** | Unchanged | P15105 | 20 | 490 | 20 | 373 | 42.1 | 7.08 | 64 | 1.051 | 0.874 | 0.504 | 0.121 | 0.744 | 0.467 | 0.678 | 0.409 |
| 443 | **Tmem126a** | Unchanged | Q9D8Y1 | 6 | 24 | 6 | 196 | 21.5 | 9.41 | 64 | 0.531 | 0.127 | 1.084 | 0.883 | 0.631 | 0.160 | 1.719 | 0.419 |
| 444 | **Naprt** | Unchanged | Q8CC86 | 22 | 340 | 22 | 538 | 58.2 | 6.46 | 64 | 1.004 | 0.986 | 0.973 | 0.921 | 1.157 | 0.461 | 0.841 | 0.496 |
| 445 | **C3** | Unchanged | H3BL60 | 7 | 27 | 1 | 102 | 11.3 | 6.07 | 64 | 1.189 | 0.813 | 1.551 | 0.619 | 0.298 | 0.290 | 5.200 | 0.216 |
| 446 | **Aco2** | Unchanged | Q99KI0 | 42 | 862 | 42 | 780 | 85.4 | 7.93 | 64 | 1.027 | 0.740 | 0.889 | 0.193 | 1.052 | 0.490 | 0.845 | 0.071 |
| 447 | **Eif1** | Unchanged | P48024 | 5 | 47 | 5 | 113 | 12.7 | 7.44 | 64 | 1.950 | 0.256 | 2.324 | 0.376 | 2.334 | 0.158 | 0.996 | 0.995 |
| 448 | **Acsm5** | Unchanged | Q8BGA8 | 24 | 202 | 24 | 578 | 64.3 | 7.72 | 64 | 1.190 | 0.010 | 1.322 | 0.291 | 1.385 | 0.018 | 0.955 | 0.842 |
| 449 | **Acot13** | Unchanged | Q9CQR4 | 6 | 80 | 6 | 140 | 15.2 | 8.82 | 64 | 0.967 | 0.951 | 1.241 | 0.676 | 1.091 | 0.868 | 1.138 | 0.713 |
| 450 | **Mttp** | Unchanged | O08601 | 52 | 882 | 52 | 894 | 99 | 7.62 | 64 | 0.818 | 0.054 | 0.970 | 0.768 | 1.264 | 0.084 | 0.768 | 0.047 |
| 451 | **Ndufb7** | Unchanged | Q9CR61 | 7 | 61 | 7 | 137 | 16.3 | 8.18 | 64 | 1.557 | 0.673 | 11.183 | 0.227 | 0.628 | 0.437 | 17.811 | 0.212 |
| 452 | **Cyp2c54** | Down | Q6XVG2 | 28 | 334 | 7 | 490 | 55.8 | 7.47 | 63 | 1.291 | 0.144 | 0.357 | 0.002 | 0.663 | 0.082 | 0.539 | 0.017 |
| 453 | **Rab1A** | Unchanged | P62821 | 11 | 127 | 5 | 205 | 22.7 | 6.21 | 63 | 0.768 | 0.557 | 0.739 | 0.578 | 0.655 | 0.418 | 1.128 | 0.828 |
| 454 | **Ndufs1** | Unchanged | Q91VD9 | 31 | 509 | 31 | 727 | 79.7 | 5.72 | 63 | 1.043 | 0.834 | 1.108 | 0.626 | 0.899 | 0.557 | 1.232 | 0.356 |
| 455 | **Aass** | Unchanged | Q99K67 | 45 | 662 | 45 | 926 | 102.9 | 6.87 | 63 | 1.280 | 0.288 | 1.333 | 0.420 | 0.889 | 0.663 | 1.499 | 0.336 |
| 456 | **Suox** | Unchanged | Q8R086 | 19 | 352 | 19 | 546 | 60.7 | 6.54 | 63 | 1.112 | 0.661 | 0.934 | 0.759 | 1.161 | 0.542 | 0.804 | 0.431 |
| 457 | **Gstm6** | Unchanged | O35660 | 15 | 156 | 11 | 218 | 25.6 | 6.24 | 59 | 1.181 | 0.626 | 1.522 | 0.062 | 0.881 | 0.539 | 1.728 | 0.013 |
| 458 | **Snd1** | Unchanged | Q78PY7 | 47 | 681 | 47 | 910 | 102 | 7.43 | 63 | 0.979 | 0.903 | 0.993 | 0.969 | 1.021 | 0.886 | 0.972 | 0.861 |
| 459 | **Btf3l4** | Unchanged | Q9CQH7 | 5 | 38 | 5 | 158 | 17.3 | 6.35 | 63 | 0.652 | 0.456 | 0.727 | 0.553 | 0.936 | 0.902 | 0.777 | 0.608 |
| 460 | **Cap1** | UP | P40124 | 19 | 313 | 19 | 474 | 51.5 | 7.52 | 63 | 0.895 | 0.639 | 1.193 | 0.355 | 1.662 | 0.015 | 0.718 | 0.083 |
| 461 | **Cct5** | UP | P80316 | 23 | 322 | 22 | 541 | 59.6 | 6.02 | 63 | 1.413 | 0.035 | 1.290 | 0.102 | 1.534 | 0.010 | 0.841 | 0.218 |
| 462 | **Ywhae** | Unchanged | P62259 | 20 | 383 | 18 | 255 | 29.2 | 4.74 | 63 | 0.779 | 0.218 | 1.277 | 0.455 | 0.980 | 0.921 | 1.304 | 0.416 |
| 463 | **Lsm7** | UP | Q9CQQ8 | 5 | 41 | 5 | 103 | 11.6 | 5.27 | 63 | 1.129 | 0.558 | 1.639 | 0.035 | 1.211 | 0.210 | 1.353 | 0.114 |
| 464 | **Psmd2** | Unchanged | Q8VDM4 | 41 | 465 | 41 | 908 | 100.1 | 5.17 | 63 | 0.889 | 0.617 | 1.146 | 0.562 | 0.980 | 0.902 | 1.169 | 0.506 |
| 465 | **Ndrg2iso** | Unchanged | Q9QYG0 | 15 | 295 | 11 | 371 | 40.8 | 5.4 | 63 | 0.828 | 0.359 | 0.882 | 0.575 | 0.730 | 0.179 | 1.207 | 0.583 |
| 466 | **Ehd3** | Unchanged | Q9QXY6 | 26 | 170 | 14 | 535 | 60.8 | 6.46 | 63 | 1.341 | 0.125 | 1.485 | 0.041 | 1.223 | 0.357 | 1.214 | 0.372 |
| 467 | **Glyctk** | Unchanged | A0A0R4J1H2 | 16 | 163 | 16 | 523 | 55.3 | 6.67 | 63 | 1.045 | 0.859 | 1.255 | 0.630 | 1.083 | 0.746 | 1.159 | 0.757 |
| 468 | **Ywhaq** | Unchanged | P68254 | 16 | 220 | 13 | 245 | 27.8 | 4.78 | 63 | 0.605 | 0.431 | 0.938 | 0.905 | 0.905 | 0.848 | 1.036 | 0.919 |
| 469 | **Vdac1** | Unchanged | Q60932 | 14 | 292 | 13 | 296 | 32.3 | 8.43 | 63 | 1.085 | 0.742 | 0.938 | 0.621 | 0.822 | 0.231 | 1.141 | 0.363 |
| 470 | **Idh2** | Unchanged | P54071 | 31 | 603 | 30 | 452 | 50.9 | 8.69 | 63 | 1.192 | 0.378 | 2.010 | 0.114 | 1.610 | 0.056 | 1.248 | 0.509 |
| 471 | **Gale** | Unchanged | Q8R059 | 12 | 117 | 12 | 347 | 38.2 | 6.74 | 63 | 1.622 | 0.147 | 1.135 | 0.724 | 0.777 | 0.401 | 1.460 | 0.326 |
| 472 | **Adhfe1** | Unchanged | Q8R0N6 | 16 | 116 | 16 | 465 | 49.9 | 7.58 | 63 | 1.089 | 0.730 | 1.006 | 0.982 | 0.756 | 0.255 | 1.330 | 0.233 |
| 473 | **Sec22b** | Unchanged | O08547 | 8 | 74 | 8 | 215 | 24.7 | 8.51 | 63 | 1.122 | 0.788 | 0.994 | 0.989 | 0.422 | 0.199 | 2.354 | 0.038 |
| 474 | **Sec23a** | Unchanged | Q01405 | 30 | 345 | 26 | 765 | 86.1 | 7.08 | 63 | 1.077 | 0.662 | 1.370 | 0.192 | 1.159 | 0.521 | 1.182 | 0.510 |
| 475 | **Slc27a2** | Unchanged | O35488 | 42 | 646 | 42 | 620 | 70.4 | 8.9 | 59 | 1.505 | 0.065 | 1.403 | 0.072 | 1.468 | 0.159 | 0.956 | 0.840 |
| 476 | **Slc25a10** | Unchanged | Q9QZD8 | 12 | 125 | 12 | 287 | 31.7 | 9.32 | 63 | 1.276 | 0.682 | 2.686 | 0.174 | 0.709 | 0.588 | 3.787 | 0.087 |
| 477 | **Thrsp** | Unchanged | Q62264 | 6 | 95 | 6 | 150 | 17.1 | 4.98 | 63 | 1.374 | 0.539 | 1.436 | 0.381 | 1.187 | 0.772 | 1.210 | 0.619 |
| 478 | **Cope** | Unchanged | O89079 | 14 | 186 | 14 | 308 | 34.5 | 5.06 | 56 | 1.252 | 0.474 | 1.375 | 0.221 | 1.193 | 0.614 | 1.152 | 0.609 |
| 479 | **Aldh3a2** | UP | P47740 | 25 | 545 | 24 | 484 | 53.9 | 8.35 | 63 | 1.622 | 0.039 | 1.462 | 0.167 | 1.823 | 0.012 | 0.802 | 0.256 |
| 480 | **Ugp2** | Unchanged | Q91ZJ5 | 35 | 628 | 35 | 508 | 56.9 | 7.61 | 63 | 1.192 | 0.343 | 1.089 | 0.574 | 1.145 | 0.451 | 0.951 | 0.641 |
| 481 | **Cct3** | UP | P80318 | 26 | 260 | 26 | 545 | 60.6 | 6.7 | 63 | 1.127 | 0.425 | 1.159 | 0.428 | 1.512 | 0.000 | 0.766 | 0.121 |

| | | | | | | | | | | | | | | | | | | |
|---|---|---|---|---|---|---|---|---|---|---|---|---|---|---|---|---|---|---|
| 482 | **Bckdhb** | UP | Q6P3A8 | 14 | 126 | 14 | 390 | 42.9 | 6.7 | 63 | 1.186 | 0.434 | 2.141 | 0.007 | 1.178 | 0.459 | 1.819 | 0.028 |
| 483 | **Stard10** | Unchanged | Q9JMD3 | 18 | 221 | 18 | 291 | 32.9 | 7.12 | 63 | 1.253 | 0.511 | 1.273 | 0.444 | 1.402 | 0.126 | 0.908 | 0.631 |
| 484 | **Ca1** | Unchanged | P13634 | 8 | 49 | 8 | 261 | 28.3 | 6.96 | 62 | 1.612 | 0.154 | 1.406 | 0.338 | 1.032 | 0.902 | 1.362 | 0.320 |
| 485 | **Pon3** | Unchanged | Q62087 | 11 | 114 | 11 | 354 | 39.3 | 5.74 | 62 | 0.789 | 0.258 | 0.775 | 0.341 | 1.243 | 0.417 | 0.623 | 0.124 |
| 486 | **Gstm1iso** | Unchanged | F6WHQ7 | 20 | 285 | 1 | 194 | 22.6 | 9.07 | 62 | 1.315 | 0.579 | 1.215 | 0.683 | 2.214 | 0.140 | 0.549 | 0.235 |
| 487 | **Fgb** | Unchanged | Q8K0E8 | 26 | 262 | 25 | 481 | 54.7 | 7.08 | 62 | 0.567 | 0.058 | 0.606 | 0.104 | 0.566 | 0.059 | 1.071 | 0.814 |
| 488 | **Aspdh** | Down | Q9DCQ2 | 13 | 183 | 13 | 287 | 30.3 | 6.95 | 62 | 0.340 | 0.023 | 0.397 | 0.052 | 1.133 | 0.778 | 0.351 | 0.115 |
| 489 | **Maob** | Unchanged | Q8BW75 | 25 | 401 | 24 | 520 | 58.5 | 8.29 | 62 | 1.209 | 0.081 | 0.976 | 0.868 | 1.115 | 0.440 | 0.876 | 0.433 |
| 490 | **Kmo** | Unchanged | Q91WN4 | 22 | 250 | 22 | 479 | 54.5 | 8.88 | 62 | 0.978 | 0.891 | 1.134 | 0.485 | 0.739 | 0.015 | 1.535 | 0.045 |
| 491 | **Sod2** | Unchanged | P09671 | 10 | 201 | 10 | 222 | 24.6 | 8.62 | 62 | 0.811 | 0.160 | 0.829 | 0.471 | 1.240 | 0.260 | 0.669 | 0.153 |
| 492 | **Cox6a1** | Unchanged | P43024 | 4 | 102 | 4 | 111 | 12.3 | 9.98 | 62 | 1.299 | 0.610 | 1.576 | 0.271 | 1.165 | 0.733 | 1.352 | 0.207 |
| 493 | **Dera** | Down | Q91YP3 | 16 | 88 | 16 | 318 | 35 | 8.72 | 62 | 1.025 | 0.936 | 1.366 | 0.441 | 0.390 | 0.044 | 3.500 | 0.040 |
| 494 | **Abhd14b** | Unchanged | E9QN99 | 10 | 144 | 10 | 210 | 22.4 | 6.52 | 62 | 1.047 | 0.885 | 0.844 | 0.655 | 1.058 | 0.873 | 0.798 | 0.620 |
| 495 | **Phb2** | Unchanged | O35129 | 15 | 222 | 15 | 299 | 33.3 | 9.83 | 62 | 0.919 | 0.634 | 0.713 | 0.183 | 0.839 | 0.592 | 0.850 | 0.675 |
| 496 | **Sfxn1** | Unchanged | Q99JR1 | 12 | 159 | 11 | 322 | 35.6 | 9.23 | 62 | 0.805 | 0.314 | 0.894 | 0.646 | 0.669 | 0.117 | 1.337 | 0.225 |
| 497 | **Canx** | Unchanged | P35564 | 30 | 393 | 30 | 591 | 67.2 | 4.64 | 62 | 0.764 | 0.501 | 1.067 | 0.845 | 0.799 | 0.533 | 1.336 | 0.180 |
| 498 | **Ube2k** | Unchanged | P61087 | 8 | 52 | 8 | 200 | 22.4 | 5.44 | 62 | 0.979 | 0.925 | 0.959 | 0.862 | 0.970 | 0.906 | 0.989 | 0.970 |
| 499 | **Crot** | UP | Q9DC50 | 32 | 523 | 32 | 612 | 70.2 | 6.73 | 61 | 1.808 | 0.032 | 2.470 | 0.021 | 2.936 | 0.000 | 0.841 | 0.411 |
| 500 | **Glod4** | Unchanged | Q9CPV4 | 14 | 159 | 14 | 298 | 33.3 | 5.47 | 61 | 0.860 | 0.318 | 0.765 | 0.185 | 1.141 | 0.064 | 0.671 | 0.046 |
| 501 | **Cct7** | Unchanged | P80313 | 25 | 287 | 25 | 544 | 59.6 | 7.84 | 61 | 0.342 | 0.089 | 0.588 | 0.300 | 0.347 | 0.089 | 1.696 | 0.206 |
| 502 | **Pcyt2** | Unchanged | Q922E4 | 16 | 147 | 16 | 404 | 45.2 | 6.58 | 61 | 1.180 | 0.729 | 1.363 | 0.547 | 1.271 | 0.584 | 1.072 | 0.889 |
| 503 | **C3 iso** | Unchanged | P01027 | 74 | 782 | 68 | 1663 | 186.4 | 6.73 | 61 | 0.699 | 0.176 | 1.656 | 0.344 | 1.271 | 0.356 | 1.303 | 0.572 |
| 504 | **Gpt2** | Unchanged | Q8BGT5 | 22 | 281 | 21 | 522 | 57.9 | 8 | 61 | 1.429 | 0.260 | 0.706 | 0.136 | 1.022 | 0.936 | 0.691 | 0.276 |
| 505 | **C1qbp** | Unchanged | Q8R5L1 | 9 | 124 | 9 | 279 | 31 | 4.87 | 61 | 0.969 | 0.931 | 1.007 | 0.986 | 1.486 | 0.288 | 0.678 | 0.270 |
| 506 | **Aco1** | Unchanged | P28271 | 45 | 864 | 45 | 889 | 98.1 | 7.5 | 60 | 1.183 | 0.331 | 1.168 | 0.456 | 1.291 | 0.067 | 0.905 | 0.572 |
| 507 | **Rpn2** | Unchanged | Q9DBG6 | 26 | 472 | 26 | 631 | 69 | 5.81 | 61 | 1.162 | 0.621 | 1.012 | 0.971 | 1.183 | 0.604 | 0.856 | 0.658 |
| 508 | **Hnrnpk** | Unchanged | P61979 | 23 | 391 | 23 | 463 | 50.9 | 5.54 | 61 | 0.892 | 0.492 | 1.055 | 0.831 | 0.968 | 0.864 | 1.090 | 0.727 |
| 509 | **Cpt2** | Unchanged | P52825 | 30 | 612 | 30 | 658 | 73.9 | 8.37 | 61 | 1.079 | 0.661 | 1.087 | 0.691 | 1.261 | 0.162 | 0.862 | 0.416 |
| 510 | **Decr1** | Unchanged | Q9CQ62 | 18 | 412 | 18 | 335 | 36.2 | 8.95 | 61 | 1.204 | 0.272 | 1.538 | 0.102 | 1.280 | 0.291 | 1.201 | 0.498 |
| 511 | **Rdh16** | UP | O54909 | 16 | 143 | 4 | 317 | 35.7 | 9.11 | 61 | 2.191 | 0.110 | 3.421 | 0.069 | 1.935 | 0.030 | 1.768 | 0.247 |
| 512 | **Hsd17b13 iso** | Unchanged | Q8VCR2 | 18 | 264 | 4 | 304 | 33.4 | 8.88 | 61 | 1.084 | 0.657 | 1.151 | 0.482 | 1.318 | 0.027 | 0.873 | 0.482 |
| 513 | **Idi1** | Unchanged | P58044 | 10 | 111 | 10 | 227 | 26.3 | 6.16 | 56 | 0.503 | 0.073 | 0.911 | 0.801 | 0.657 | 0.283 | 1.387 | 0.439 |
| 514 | **Skp1** | Unchanged | Q9WTX5 | 8 | 112 | 8 | 163 | 18.7 | 4.54 | 61 | 0.976 | 0.923 | 0.645 | 0.110 | 0.718 | 0.272 | 0.899 | 0.775 |
| 515 | **Pdia5** | Unchanged | Q921X9 | 20 | 274 | 20 | 517 | 59.2 | 7.5 | 61 | 1.858 | 0.416 | 1.720 | 0.309 | 1.517 | 0.341 | 1.134 | 0.793 |
| 516 | **Cryz** | Unchanged | P47199 | 15 | 350 | 15 | 331 | 35.2 | 8.07 | 61 | 1.231 | 0.197 | 0.712 | 0.082 | 0.826 | 0.250 | 0.862 | 0.367 |
| 517 | **Mpc2** | Unchanged | Q9D023 | 8 | 122 | 8 | 127 | 14.3 | 10.61 | 61 | 1.828 | 0.140 | 1.560 | 0.436 | 1.322 | 0.531 | 1.180 | 0.758 |
| 518 | **Clic1** | Unchanged | Q9Z1Q5 | 7 | 55 | 7 | 241 | 27 | 5.17 | 61 | 0.372 | 0.172 | 0.276 | 0.121 | 0.692 | 0.556 | 0.398 | 0.223 |
| 519 | **Mccc2** | Down | Q3ULD5 | 23 | 264 | 23 | 563 | 61.3 | 8 | 61 | 0.825 | 0.569 | 0.647 | 0.147 | 0.518 | 0.044 | 1.248 | 0.400 |
| 520 | **Urod** | UP | P70697 | 14 | 144 | 14 | 367 | 40.7 | 6.65 | 60 | 1.494 | 0.116 | 2.126 | 0.034 | 1.559 | 0.092 | 1.364 | 0.251 |
| 521 | **Mmut** | Unchanged | P16332 | 26 | 411 | 26 | 748 | 82.8 | 6.89 | 60 | 1.037 | 0.707 | 1.077 | 0.706 | 0.959 | 0.738 | 1.123 | 0.582 |
| 522 | **Mb** | Unchanged | P04247 | 7 | 29 | 7 | 154 | 17.1 | 7.62 | 50 | 0.494 | 0.454 | 0.513 | 0.471 | 0.363 | 0.353 | 1.412 | 0.503 |

| | | | | | | | | | | | | | | | | | | |
|---|---|---|---|---|---|---|---|---|---|---|---|---|---|---|---|---|---|---|
| 523 | **Rps5** | Unchanged | Q91V55 | 14 | 138 | 14 | 204 | 22.9 | 9.72 | 60 | 1.560 | 0.051 | 3.247 | 0.223 | 1.367 | 0.267 | 2.375 | 0.301 |
| 524 | **Actr3** | Unchanged | Q99JY9 | 16 | 225 | 13 | 418 | 47.3 | 5.88 | 60 | 1.035 | 0.815 | 1.187 | 0.493 | 1.057 | 0.717 | 1.123 | 0.625 |
| 525 | **Cyp2c68** | Unchanged | K7N6C2 | 22 | 113 | 5 | 491 | 56 | 7.36 | 60 | 1.033 | 0.946 | 2.743 | 0.229 | 1.124 | 0.803 | 2.440 | 0.272 |
| 526 | **Cox7a2** | Unchanged | P48771 | 5 | 81 | 5 | 83 | 9.3 | 10.27 | 60 | 0.652 | 0.066 | 0.689 | 0.240 | 0.685 | 0.036 | 1.005 | 0.988 |
| 527 | **Hspa4** | Unchanged | Q3U2G2 | 43 | 246 | 40 | 842 | 94.1 | 5.21 | 60 | 0.672 | 0.017 | 1.012 | 0.883 | 0.754 | 0.031 | 1.342 | 0.020 |
| 528 | **Elob** | Unchanged | P62869 | 8 | 39 | 8 | 118 | 13.2 | 5.01 | 60 | 1.482 | 0.485 | 1.469 | 0.515 | 1.299 | 0.500 | 1.130 | 0.811 |
| 529 | **Ftl1** | Unchanged | Q9CPX4 | 10 | 115 | 10 | 183 | 20.8 | 6 | 60 | 0.984 | 0.963 | 0.679 | 0.383 | 1.294 | 0.214 | 0.525 | 0.054 |
| 530 | **Rps8** | Unchanged | P62242 | 13 | 190 | 13 | 208 | 24.2 | 10.32 | 60 | 0.805 | 0.272 | 0.689 | 0.063 | 1.011 | 0.946 | 0.682 | 0.010 |
| 531 | **Gsta1** | UP | P13745 | 12 | 645 | 2 | 223 | 25.6 | 8.98 | 60 | 2.780 | 0.092 | 2.583 | 0.021 | 5.907 | 0.007 | 0.437 | 0.054 |
| 532 | **Cycs** | Unchanged | P62897 | 13 | 342 | 13 | 105 | 11.6 | 9.58 | 60 | 0.785 | 0.139 | 1.186 | 0.554 | 0.980 | 0.896 | 1.209 | 0.534 |
| 533 | **Cyp3a13** | Unchanged | Q64464 | 28 | 197 | 27 | 503 | 57.5 | 8.57 | 60 | 0.792 | 0.422 | 0.920 | 0.779 | 0.638 | 0.202 | 1.441 | 0.217 |
| 534 | **Gpd2** | Unchanged | Q64521 | 31 | 254 | 31 | 727 | 80.9 | 6.61 | 60 | 1.144 | 0.487 | 1.316 | 0.116 | 0.922 | 0.642 | 1.427 | 0.109 |
| 535 | **Ptgr2** | UP | Q8VDQ1 | 12 | 144 | 12 | 351 | 38 | 5.41 | 60 | 1.460 | 0.115 | 1.832 | 0.249 | 1.631 | 0.038 | 1.123 | 0.769 |
| 536 | **Ndufs2** | Unchanged | Q91WD5 | 19 | 217 | 19 | 463 | 52.6 | 6.99 | 60 | 1.184 | 0.561 | 1.079 | 0.811 | 0.767 | 0.459 | 1.407 | 0.227 |
| 537 | **Acot2** | UP | Q9QYR9 | 17 | 188 | 9 | 453 | 49.6 | 7.36 | 60 | 3.763 | 0.214 | 7.829 | 0.068 | 4.544 | 0.000 | 1.723 | 0.338 |
| 538 | **Rpl7a** | Unchanged | P12970 | 20 | 345 | 20 | 266 | 30 | 10.56 | 60 | 1.341 | 0.187 | 1.353 | 0.462 | 1.569 | 0.175 | 0.863 | 0.718 |
| 539 | **Clybl** | Down | Q8R4N0 | 16 | 151 | 16 | 338 | 37.5 | 8.54 | 60 | 0.577 | 0.023 | 0.625 | 0.065 | 0.498 | 0.014 | 1.254 | 0.292 |
| 540 | **Psmd14** | Unchanged | O35593 | 9 | 88 | 9 | 310 | 34.6 | 6.52 | 60 | 0.947 | 0.747 | 1.324 | 0.189 | 1.364 | 0.011 | 0.970 | 0.866 |
| 541 | **Arf2** | Unchanged | Q8BSL7 | 10 | 207 | 1 | 181 | 20.7 | 6.58 | 60 | 0.691 | 0.578 | 2.931 | 0.134 | 1.574 | 0.317 | 1.863 | 0.265 |
| 542 | **Vim** | Unchanged | P20152 | 23 | 115 | 21 | 466 | 53.7 | 5.12 | 60 | 1.015 | 0.961 | 0.922 | 0.807 | 0.946 | 0.848 | 0.974 | 0.915 |
| 543 | **Ces2g** | Unchanged | E9PV38 | 19 | 146 | 13 | 560 | 62.7 | 6.04 | 60 | 1.123 | 0.721 | 1.449 | 0.246 | 1.407 | 0.216 | 1.030 | 0.900 |
| 544 | **Cfl1** | UP | P18760 | 11 | 262 | 10 | 166 | 18.5 | 8.09 | 60 | 1.150 | 0.691 | 1.765 | 0.023 | 1.868 | 0.061 | 0.945 | 0.798 |
| 545 | **Gk** | Unchanged | Q64516 | 23 | 228 | 23 | 559 | 61.2 | 5.87 | 60 | 1.044 | 0.885 | 1.036 | 0.905 | 0.869 | 0.623 | 1.191 | 0.432 |
| 546 | **Dpyd** | Unchanged | Q8CHR6 | 43 | 645 | 43 | 1025 | 111.2 | 7.27 | 60 | 0.844 | 0.235 | 1.078 | 0.486 | 1.083 | 0.382 | 0.996 | 0.967 |
| 547 | **Papss2** | UP | O88428 | 26 | 260 | 25 | 621 | 70.3 | 7.58 | 59 | 1.128 | 0.477 | 2.271 | 0.003 | 1.549 | 0.002 | 1.466 | 0.037 |
| 548 | **Ppa2** | UP | Q91VM9 | 15 | 142 | 15 | 330 | 38.1 | 6.98 | 59 | 1.220 | 0.101 | 2.388 | 0.006 | 1.484 | 0.036 | 1.609 | 0.056 |
| 549 | **Snu13** | UP | Q9D0T1 | 5 | 19 | 5 | 128 | 14.2 | 8.46 | 59 | 0.971 | 0.965 | 1.691 | 0.478 | 2.941 | 0.042 | 0.575 | 0.243 |
| 550 | **Ugt1a6b** | Unchanged | K9J7B2 | 22 | 283 | 6 | 531 | 60.5 | 8.69 | 59 | 1.077 | 0.680 | 1.463 | 0.153 | 0.949 | 0.768 | 1.541 | 0.080 |
| 551 | **Entpd5** | Unchanged | Q9WUZ9 | 16 | 286 | 16 | 427 | 47.1 | 5.29 | 59 | 1.191 | 0.409 | 1.428 | 0.046 | 1.167 | 0.365 | 1.224 | 0.104 |
| 552 | **Hist1h4a** | Unchanged | P62806 | 14 | 180 | 14 | 103 | 11.4 | 11.36 | 59 | 1.814 | 0.391 | 3.901 | 0.237 | 2.716 | 0.209 | 1.436 | 0.644 |
| 553 | **Psmb2** | UP | Q9R1P3 | 11 | 152 | 11 | 201 | 22.9 | 7.02 | 59 | 1.012 | 0.941 | 1.793 | 0.008 | 1.205 | 0.246 | 1.488 | 0.012 |
| 554 | **Acy1** | Unchanged | A0A0R4J050 | 14 | 96 | 14 | 408 | 45.8 | 6.21 | 59 | 1.195 | 0.588 | 1.054 | 0.868 | 0.864 | 0.577 | 1.221 | 0.627 |
| 555 | **Gda** | UP | Q9R111 | 16 | 126 | 16 | 454 | 51 | 5.53 | 59 | 3.310 | 0.186 | 3.834 | 0.051 | 5.479 | 0.005 | 0.700 | 0.355 |
| 556 | **Srm** | Unchanged | Q64674 | 10 | 78 | 10 | 302 | 34 | 5.5 | 59 | 0.915 | 0.830 | 1.256 | 0.643 | 0.588 | 0.160 | 2.138 | 0.213 |
| 557 | **Acadm** | Unchanged | P45952 | 25 | 508 | 25 | 421 | 46.5 | 8.37 | 59 | 1.078 | 0.518 | 1.362 | 0.218 | 1.444 | 0.025 | 0.944 | 0.774 |
| 558 | **Mvp** | Unchanged | E9Q3X0 | 33 | 258 | 33 | 870 | 96.8 | 5.64 | 59 | 1.207 | 0.679 | 1.376 | 0.269 | 0.998 | 0.994 | 1.379 | 0.220 |
| 559 | **Dhfr** | Unchanged | P00375 | 9 | 101 | 9 | 187 | 21.6 | 8.53 | 49 | 1.126 | 0.324 | 1.145 | 0.079 | 1.366 | 0.185 | 0.838 | 0.419 |
| 560 | **Atox1** | Unchanged | O08997 | 4 | 107 | 4 | 68 | 7.3 | 6.51 | 59 | 0.948 | 0.821 | 0.769 | 0.394 | 0.999 | 0.997 | 0.769 | 0.423 |
| 561 | **Ufm1** | Unchanged | P61961 | 3 | 39 | 3 | 85 | 9.1 | 9.31 | 59 | 1.295 | 0.454 | 1.167 | 0.657 | 0.696 | 0.319 | 1.677 | 0.180 |
| 562 | **Rpl12** | Unchanged | P35979 | 8 | 217 | 8 | 165 | 17.8 | 9.42 | 59 | 0.903 | 0.775 | 0.618 | 0.254 | 1.120 | 0.717 | 0.552 | 0.075 |
| 563 | **Ociad1** | Unchanged | Q9CRD0 | 9 | 58 | 9 | 247 | 27.6 | 7.81 | 59 | 5.063 | 0.367 | 1.857 | 0.256 | 1.976 | 0.251 | 0.939 | 0.908 |

| | | | | | | | | | | | | | | | | | | |
|---|---|---|---|---|---|---|---|---|---|---|---|---|---|---|---|---|---|---|
| 564 | **Suclg2** | Unchanged | Q9Z2I8 | 22 | 470 | 22 | 433 | 46.8 | 7.02 | 59 | 1.116 | 0.268 | 1.132 | 0.211 | 1.074 | 0.434 | 1.054 | 0.628 |
| 565 | **Rpsa** | Unchanged | P14206 | 17 | 314 | 17 | 295 | 32.8 | 4.87 | 59 | 1.388 | 0.115 | 1.331 | 0.152 | 1.407 | 0.013 | 0.946 | 0.742 |
| 566 | **Cdc42** | Unchanged | P60766 | 9 | 115 | 9 | 191 | 21.2 | 6.55 | 59 | 1.269 | 0.503 | 1.643 | 0.297 | 1.119 | 0.700 | 1.468 | 0.331 |
| 567 | **Cs** | Unchanged | Q9CZU6 | 22 | 355 | 22 | 464 | 51.7 | 8.57 | 59 | 0.974 | 0.890 | 1.012 | 0.954 | 0.997 | 0.988 | 1.015 | 0.934 |
| 568 | **Pdia4** | Down | P08003 | 45 | 709 | 45 | 638 | 71.9 | 5.31 | 59 | 0.726 | 0.047 | 0.554 | 0.002 | 0.695 | 0.055 | 0.796 | 0.217 |
| 569 | **Psmd6** | Unchanged | Q99JI4 | 21 | 125 | 21 | 389 | 45.5 | 5.52 | 54 | 1.518 | 0.150 | 0.843 | 0.228 | 2.124 | 0.106 | 0.397 | 0.069 |
| 570 | **Lonp1** | UP | Q8CGK3 | 38 | 490 | 38 | 949 | 105.8 | 6.57 | 59 | 1.169 | 0.585 | 2.342 | 0.049 | 1.281 | 0.362 | 1.829 | 0.112 |
| 571 | **Tst** | Unchanged | P52196 | 18 | 440 | 17 | 297 | 33.4 | 7.85 | 59 | 1.017 | 0.899 | 1.202 | 0.468 | 0.976 | 0.812 | 1.231 | 0.403 |
| 572 | **Ugt1a6** | Unchanged | Q64435 | 22 | 451 | 6 | 531 | 60.4 | 8.62 | 59 | 1.309 | 0.062 | 1.937 | 0.059 | 1.345 | 0.225 | 1.440 | 0.260 |
| 573 | **Hoga1** | Unchanged | Q9DCU9 | 16 | 226 | 16 | 321 | 34.6 | 7.72 | 59 | 0.888 | 0.545 | 1.214 | 0.433 | 0.795 | 0.290 | 1.526 | 0.092 |
| 574 | **Rhoc** | Unchanged | Q62159 | 9 | 96 | 1 | 193 | 22 | 6.58 | 59 | 0.887 | 0.730 | 1.134 | 0.713 | 0.653 | 0.299 | 1.736 | 0.129 |
| 575 | **Nudt2** | Unchanged | P56380 | 5 | 14 | 5 | 147 | 17 | 6.28 | 59 | 1.386 | 0.353 | 1.244 | 0.308 | 1.156 | 0.389 | 1.076 | 0.723 |
| 576 | **Nudt7** | Unchanged | Q99P30 | 13 | 176 | 13 | 236 | 26.8 | 6.34 | 58 | 0.705 | 0.253 | 0.654 | 0.163 | 0.853 | 0.616 | 0.766 | 0.407 |
| 577 | **Pa2g4** | Unchanged | P50580 | 20 | 208 | 20 | 394 | 43.7 | 6.86 | 58 | 0.832 | 0.166 | 0.923 | 0.719 | 0.750 | 0.120 | 1.230 | 0.441 |
| 578 | **Atp5f1d** | Unchanged | Q9D3D9 | 4 | 76 | 4 | 168 | 17.6 | 5.08 | 58 | 1.191 | 0.603 | 0.450 | 0.204 | 0.635 | 0.427 | 0.708 | 0.653 |
| 579 | **Uqcrc1iso** | Unchanged | Q9CZ13 | 24 | 516 | 14 | 480 | 52.8 | 6.21 | 58 | 1.059 | 0.698 | 0.752 | 0.048 | 1.041 | 0.792 | 0.722 | 0.066 |
| 580 | **Ces2e** | Unchanged | Q8BK48 | 25 | 461 | 20 | 559 | 62.3 | 6.16 | 58 | 1.275 | 0.223 | 1.286 | 0.304 | 1.366 | 0.049 | 0.941 | 0.737 |
| 581 | **Nipsnap3b** | UP | Q9CQE1 | 9 | 103 | 9 | 247 | 28.3 | 9.48 | 58 | 1.602 | 0.018 | 2.118 | 0.002 | 1.603 | 0.008 | 1.322 | 0.069 |
| 582 | **Myl6** | UP | Q60605 | 8 | 125 | 7 | 151 | 16.9 | 4.65 | 58 | 1.516 | 0.153 | 1.165 | 0.547 | 2.237 | 0.025 | 0.521 | 0.052 |
| 583 | **Nudt19** | Unchanged | P11930 | 14 | 94 | 14 | 357 | 40.3 | 6.68 | 58 | 0.931 | 0.890 | 0.647 | 0.421 | 0.759 | 0.546 | 0.852 | 0.711 |
| 584 | **Ywhah** | UP | P68510 | 15 | 170 | 11 | 246 | 28.2 | 4.89 | 58 | 1.437 | 0.339 | 1.440 | 0.198 | 2.675 | 0.016 | 0.538 | 0.075 |
| 585 | **Bckdha** | Unchanged | Q3U3J1 | 20 | 119 | 20 | 446 | 50.7 | 8.06 | 58 | 1.505 | 0.166 | 2.321 | 0.209 | 1.149 | 0.485 | 2.020 | 0.268 |
| 586 | **Pgd** | Unchanged | Q9DCD0 | 25 | 307 | 25 | 483 | 53.2 | 7.23 | 58 | 1.063 | 0.617 | 0.964 | 0.897 | 1.091 | 0.491 | 0.884 | 0.649 |
| 587 | **Pdha1** | Unchanged | P35486 | 21 | 216 | 20 | 390 | 43.2 | 8.19 | 58 | 1.391 | 0.186 | 0.803 | 0.375 | 0.788 | 0.187 | 1.019 | 0.944 |
| 588 | **Ttr** | Unchanged | P07309 | 5 | 77 | 5 | 147 | 15.8 | 6.16 | 58 | 2.011 | 0.202 | 1.133 | 0.842 | 2.041 | 0.051 | 0.555 | 0.143 |
| 589 | **Rpl22** | Unchanged | P67984 | 7 | 89 | 7 | 128 | 14.8 | 9.19 | 58 | 0.753 | 0.391 | 0.867 | 0.687 | 0.640 | 0.277 | 1.355 | 0.574 |
| 590 | **Gls2** | Down | Q571F8 | 25 | 268 | 25 | 602 | 66.3 | 7.66 | 58 | 0.877 | 0.652 | 0.850 | 0.277 | 0.657 | 0.020 | 1.293 | 0.234 |
| 591 | **Sdhb** | Unchanged | Q9CQA3 | 16 | 198 | 16 | 282 | 31.8 | 8.68 | 58 | 1.288 | 0.162 | 0.782 | 0.302 | 0.754 | 0.290 | 1.037 | 0.887 |
| 592 | **Tagln2** | UP | Q9WVA4 | 11 | 187 | 11 | 199 | 22.4 | 8.24 | 58 | 0.991 | 0.964 | 2.213 | 0.000 | 2.275 | 0.017 | 0.973 | 0.893 |
| 593 | **Slc25a12** | Unchanged | Q8BH59 | 22 | 182 | 18 | 677 | 74.5 | 8.25 | 58 | 0.940 | 0.903 | 1.205 | 0.692 | 0.667 | 0.489 | 1.807 | 0.056 |
| 594 | **Ndufa10** | Unchanged | Q99LC3 | 14 | 163 | 14 | 355 | 40.6 | 7.78 | 58 | 1.348 | 0.385 | 1.419 | 0.488 | 1.381 | 0.437 | 1.028 | 0.953 |
| 595 | **Rps27a** | Unchanged | P62983 | 9 | 180 | 9 | 156 | 17.9 | 9.64 | 58 | 1.018 | 0.965 | 1.090 | 0.849 | 0.965 | 0.929 | 1.130 | 0.787 |
| 596 | **Gldc** | UP | Q91W43 | 31 | 305 | 31 | 1025 | 113.2 | 7.56 | 58 | 1.746 | 0.003 | 1.910 | 0.007 | 1.162 | 0.366 | 1.643 | 0.021 |
| 597 | **Ces1e** | Unchanged | Q64176 | 23 | 384 | 19 | 562 | 61.5 | 6.15 | 58 | 1.110 | 0.780 | 1.755 | 0.158 | 0.815 | 0.526 | 2.152 | 0.057 |
| 598 | **Galm** | Unchanged | Q8K157 | 11 | 122 | 11 | 342 | 37.8 | 6.74 | 58 | 1.120 | 0.509 | 1.338 | 0.304 | 1.287 | 0.269 | 1.040 | 0.883 |
| 599 | **Ndufa7** | Unchanged | Q9Z1P6 | 6 | 47 | 6 | 113 | 12.6 | 10.17 | 58 | 1.134 | 0.811 | 1.857 | 0.301 | 0.620 | 0.414 | 2.995 | 0.123 |
| 600 | **Pnpo** | Unchanged | Q91XF0 | 9 | 196 | 9 | 261 | 30.1 | 8.22 | 57 | 0.771 | 0.132 | 0.695 | 0.135 | 0.879 | 0.421 | 0.791 | 0.289 |
| 601 | **Gsr** | Down | P47791 | 16 | 148 | 16 | 500 | 53.6 | 7.99 | 57 | 0.785 | 0.308 | 1.242 | 0.510 | 0.544 | 0.030 | 2.282 | 0.073 |
| 602 | **Slc25a4** | Unchanged | P48962 | 18 | 236 | 7 | 298 | 32.9 | 9.72 | 57 | 0.137 | 0.249 | 0.430 | 0.442 | 0.256 | 0.317 | 1.680 | 0.248 |
| 603 | **Actr3 iso** | Unchanged | A0A087WS98 | 4 | 52 | 1 | 129 | 14.7 | 4.56 | 57 | 1.239 | 0.833 | 12.657 | 0.088 | 1.882 | 0.479 | 6.726 | 0.111 |
| 604 | **Copg1** | Unchanged | Q9QZE5 | 32 | 363 | 30 | 874 | 97.5 | 5.35 | 57 | 1.151 | 0.509 | 1.297 | 0.097 | 0.918 | 0.704 | 1.413 | 0.060 |

| | | | | | | | | | | | | | | | | | | |
|---|---|---|---|---|---|---|---|---|---|---|---|---|---|---|---|---|---|---|
| 605 | Uqcrh | Unchanged | P99028 | 4 | 82 | 4 | 89 | 10.4 | 4.87 | 57 | 1.374 | 0.083 | 1.176 | 0.534 | 1.258 | 0.252 | 0.935 | 0.782 |
| 606 | Hnrnpc | Unchanged | A0A2I3BRM6 | 6 | 28 | 1 | 117 | 12.8 | 9.5 | 57 | 0.832 | 0.728 | 0.456 | 0.217 | 0.285 | 0.115 | 1.600 | 0.273 |
| 607 | Dpys | Unchanged | Q9EQF5 | 23 | 540 | 23 | 519 | 56.7 | 7.2 | 57 | 1.276 | 0.234 | 1.125 | 0.402 | 1.416 | 0.025 | 0.794 | 0.037 |
| 608 | Txn | Unchanged | P10639 | 7 | 178 | 7 | 105 | 11.7 | 4.92 | 57 | 0.959 | 0.746 | 1.160 | 0.702 | 0.882 | 0.211 | 1.316 | 0.516 |
| 609 | Prdx5 | Unchanged | P99029 | 12 | 367 | 12 | 210 | 21.9 | 8.85 | 57 | 1.079 | 0.388 | 1.364 | 0.021 | 1.086 | 0.175 | 1.257 | 0.063 |
| 610 | Gnpnat1 | Unchanged | Q9JK38 | 6 | 78 | 6 | 184 | 20.8 | 7.69 | 57 | 1.272 | 0.539 | 0.673 | 0.354 | 1.386 | 0.481 | 0.485 | 0.181 |
| 611 | Rap1b | Unchanged | Q99JI6 | 10 | 102 | 4 | 184 | 20.8 | 5.78 | 57 | 0.878 | 0.707 | 1.181 | 0.727 | 0.909 | 0.811 | 1.299 | 0.643 |
| 612 | Rap1a | Unchanged | P62835 | 10 | 112 | 4 | 184 | 21 | 6.67 | 57 | 0.974 | 0.919 | 1.196 | 0.413 | 0.993 | 0.972 | 1.205 | 0.221 |
| 613 | Dpp3 | Unchanged | Q99KK7 | 26 | 233 | 26 | 738 | 82.8 | 5.38 | 57 | 1.917 | 0.248 | 3.531 | 0.139 | 1.552 | 0.293 | 2.275 | 0.246 |
| 614 | Ndufa8 | Unchanged | Q9DCJ5 | 8 | 80 | 8 | 172 | 20 | 8.46 | 47 | 0.532 | 0.100 | 1.319 | 0.554 | 0.724 | 0.326 | 1.823 | 0.243 |
| 615 | Dstn | Unchanged | Q9R0P5 | 8 | 113 | 8 | 165 | 18.5 | 7.97 | 57 | 1.123 | 0.628 | 0.659 | 0.179 | 1.390 | 0.360 | 0.474 | 0.129 |
| 616 | Cct4 | Unchanged | P80315 | 23 | 310 | 22 | 539 | 58 | 8.02 | 57 | 0.942 | 0.713 | 1.269 | 0.154 | 1.231 | 0.168 | 1.031 | 0.825 |
| 617 | Pla2g12b | Unchanged | Q8VC81 | 6 | 33 | 6 | 195 | 21.7 | 6.15 | 57 | 1.170 | 0.707 | 1.058 | 0.919 | 0.845 | 0.694 | 1.251 | 0.620 |
| 618 | Ddost | Unchanged | O54734 | 15 | 215 | 15 | 441 | 49 | 5.83 | 57 | 0.966 | 0.814 | 1.480 | 0.063 | 0.902 | 0.624 | 1.640 | 0.033 |
| 619 | Pecr | Unchanged | Q99MZ7 | 12 | 228 | 12 | 303 | 32.4 | 8.29 | 57 | 1.062 | 0.834 | 1.200 | 0.593 | 0.831 | 0.549 | 1.443 | 0.357 |
| 620 | Eif6 | Unchanged | O55135 | 8 | 104 | 8 | 245 | 26.5 | 4.74 | 57 | 0.951 | 0.876 | 1.073 | 0.866 | 1.152 | 0.620 | 0.931 | 0.860 |
| 621 | Lhpp | Unchanged | Q9D7I5 | 6 | 65 | 6 | 270 | 29.1 | 5.1 | 57 | 1.069 | 0.892 | 1.122 | 0.842 | 0.574 | 0.346 | 1.954 | 0.233 |
| 622 | Faah | Unchanged | O08914 | 20 | 263 | 20 | 579 | 63.2 | 7.87 | 57 | 1.170 | 0.741 | 2.102 | 0.290 | 1.499 | 0.468 | 1.403 | 0.589 |
| 623 | Ugt1a1 | Unchanged | Q63886 | 24 | 748 | 14 | 535 | 60 | 8.62 | 57 | 1.042 | 0.775 | 1.329 | 0.288 | 1.100 | 0.637 | 1.208 | 0.494 |
| 624 | Cmpk1 | UP | Q9DBP5 | 11 | 193 | 11 | 196 | 22.2 | 5.83 | 57 | 1.123 | 0.665 | 1.807 | 0.048 | 1.492 | 0.098 | 1.211 | 0.394 |
| 625 | Galnt11 | Unchanged | D3YUC2 | 1 | 1 | 1 | 46 | 5.3 | 5.21 | 57 | #DIV/0! | #DIV/0! | #DIV/0! | #DIV/0! | #DIV/0! | #DIV/0! | #DIV/0! | #DIV/0! |
| 626 | Tcp1 | Unchanged | P11983 | 21 | 283 | 21 | 556 | 60.4 | 6.16 | 56 | 1.191 | 0.252 | 1.175 | 0.476 | 1.402 | 0.069 | 0.838 | 0.431 |
| 627 | Lrpprc | Unchanged | Q6PB66 | 58 | 522 | 58 | 1392 | 156.5 | 6.83 | 56 | 1.171 | 0.153 | 1.271 | 0.083 | 1.089 | 0.520 | 1.167 | 0.286 |
| 628 | Eci2 | UP | E9PUY9 | 11 | 75 | 1 | 202 | 22.2 | 7.88 | 56 | 0.758 | 0.612 | 1.943 | 0.368 | 0.593 | 0.349 | 3.276 | 0.187 |
| 629 | Psmd11 | Down | Q8BG32 | 20 | 170 | 20 | 422 | 47.4 | 6.48 | 56 | 0.905 | 0.675 | 1.011 | 0.964 | 0.622 | 0.037 | 1.625 | 0.093 |
| 630 | Taldo1 | Down | Q93092 | 19 | 237 | 19 | 337 | 37.4 | 7.03 | 56 | 0.642 | 0.013 | 0.819 | 0.219 | 1.230 | 0.462 | 0.666 | 0.206 |
| 631 | Tomm7 | Unchanged | Q9D173 | 2 | 5 | 2 | 55 | 6.2 | 10.17 | 56 | 5.374 | 0.175 | 24.426 | 0.283 | 7.961 | 0.094 | 3.068 | 0.449 |
| 632 | Bphl | Unchanged | Q8R164 | 15 | 279 | 15 | 291 | 32.8 | 8.94 | 56 | 1.072 | 0.668 | 0.950 | 0.864 | 0.981 | 0.937 | 0.969 | 0.930 |
| 633 | Ppp2cb | Unchanged | P62715 | 10 | 94 | 1 | 309 | 35.6 | 5.43 | 56 | 0.908 | 0.474 | 0.894 | 0.595 | 0.918 | 0.666 | 0.973 | 0.917 |
| 634 | Ppp2ca | Unchanged | P63330 | 10 | 107 | 1 | 309 | 35.6 | 5.54 | 56 | 1.160 | 0.858 | 1.443 | 0.671 | 0.120 | 0.129 | 12.000 | 0.163 |
| 635 | Wdr1 | UP | O88342 | 17 | 182 | 17 | 606 | 66.4 | 6.6 | 56 | 1.283 | 0.239 | 1.863 | 0.221 | 1.728 | 0.038 | 1.078 | 0.853 |
| 636 | Marc2;Mosc2 | Unchanged | Q922Q1 | 22 | 343 | 18 | 338 | 38.2 | 8.68 | 56 | 1.035 | 0.937 | 1.704 | 0.235 | 1.144 | 0.774 | 1.489 | 0.376 |
| 637 | Psmc5 | Unchanged | P62196 | 19 | 218 | 17 | 406 | 45.6 | 7.55 | 56 | 0.601 | 0.065 | 0.881 | 0.613 | 0.881 | 0.536 | 1.000 | 1.000 |
| 638 | Cpt1a | Unchanged | P97742 | 29 | 318 | 28 | 773 | 88.2 | 8.62 | 56 | 1.127 | 0.516 | 0.831 | 0.526 | 0.826 | 0.470 | 1.006 | 0.984 |
| 639 | Gm20425 | Unchanged | E9Q035 | 60 | 584 | 8 | 978 | 107.7 | 7.77 | 56 | 0.771 | 0.304 | 1.245 | 0.385 | 1.488 | 0.218 | 0.837 | 0.529 |
| 640 | Ttc39c | Unchanged | G3X8X1 | 23 | 92 | 23 | 580 | 65.4 | 6.87 | 56 | 2.724 | 0.264 | 1.464 | 0.399 | 0.717 | 0.281 | 2.042 | 0.209 |
| 641 | Mpc1 | Unchanged | P63030 | 5 | 42 | 5 | 109 | 12.4 | 9.61 | 56 | 1.437 | 0.637 | 3.180 | 0.180 | 1.994 | 0.275 | 1.595 | 0.481 |
| 642 | Rbbp9 | Unchanged | O88851 | 8 | 24 | 8 | 186 | 20.9 | 5.97 | 56 | 0.908 | 0.784 | 0.903 | 0.808 | 0.654 | 0.292 | 1.381 | 0.525 |
| 643 | Bbox1 | Unchanged | Q924Y0 | 15 | 144 | 15 | 387 | 44.7 | 6.92 | 56 | 0.821 | 0.391 | 1.220 | 0.429 | 0.762 | 0.274 | 1.602 | 0.068 |
| 644 | Arl1 | Unchanged | P61211 | 7 | 73 | 7 | 181 | 20.4 | 5.72 | 56 | 0.662 | 0.416 | 1.245 | 0.619 | 0.761 | 0.565 | 1.636 | 0.149 |
| 645 | Aars | Unchanged | Q8BGQ7 | 31 | 410 | 31 | 968 | 106.8 | 5.67 | 56 | 0.995 | 0.980 | 0.940 | 0.842 | 0.963 | 0.888 | 0.976 | 0.946 |

| | | | | | | | | | | | | | | | | | | |
|---|---|---|---|---|---|---|---|---|---|---|---|---|---|---|---|---|---|---|
| 646 | **Hyou1** | Unchanged | Q9JKR6 | 47 | 699 | 47 | 999 | 111.1 | 5.19 | 56 | 0.940 | 0.688 | 0.966 | 0.847 | 0.782 | 0.193 | 1.236 | 0.226 |
| 647 | **Sucla2** | Down | Q9Z2I9 | 26 | 342 | 26 | 463 | 50.1 | 7.01 | 56 | 1.022 | 0.872 | 0.634 | 0.005 | 0.487 | 0.000 | 1.300 | 0.235 |
| 648 | **Cyp2j5** | Unchanged | O54749 | 20 | 196 | 19 | 501 | 57.7 | 8.73 | 56 | 1.058 | 0.701 | 1.119 | 0.388 | 0.793 | 0.133 | 1.411 | 0.008 |
| 649 | **Cox4i1** | Unchanged | P19783 | 11 | 162 | 11 | 169 | 19.5 | 9.23 | 56 | 0.972 | 0.906 | 0.810 | 0.418 | 0.779 | 0.351 | 1.040 | 0.807 |
| 650 | **Itpa** | Unchanged | Q9D892 | 7 | 100 | 7 | 198 | 21.9 | 5.87 | 56 | 1.233 | 0.556 | 1.107 | 0.734 | 1.628 | 0.076 | 0.680 | 0.113 |
| 651 | **Tbca** | Unchanged | P48428 | 8 | 38 | 8 | 108 | 12.8 | 5.27 | 56 | 1.276 | 0.618 | 1.961 | 0.194 | 1.817 | 0.146 | 1.079 | 0.866 |
| 652 | **Atp5pf** | Unchanged | P97450 | 9 | 217 | 9 | 108 | 12.5 | 9.36 | 56 | 0.978 | 0.793 | 0.917 | 0.570 | 0.762 | 0.012 | 1.203 | 0.294 |
| 653 | **Qprt** | Unchanged | Q91X91 | 12 | 291 | 12 | 299 | 31.5 | 6.71 | 56 | 0.923 | 0.536 | 0.927 | 0.751 | 0.803 | 0.178 | 1.154 | 0.593 |
| 654 | **Echs1** | Unchanged | Q8BH95 | 14 | 315 | 14 | 290 | 31.5 | 8.48 | 56 | 0.869 | 0.716 | 0.578 | 0.199 | 0.716 | 0.397 | 0.806 | 0.341 |
| 655 | **Pm20d1** | Unchanged | Q8C165 | 17 | 169 | 17 | 503 | 55.6 | 6.43 | 55 | 0.963 | 0.900 | 1.426 | 0.230 | 0.797 | 0.403 | 1.789 | 0.033 |
| 656 | **Aldh5a1** | Down | Q8BWF0 | 20 | 307 | 20 | 523 | 55.9 | 8.25 | 55 | 0.982 | 0.897 | 0.551 | 0.031 | 0.882 | 0.711 | 0.625 | 0.339 |
| 657 | **Zadh2** | Unchanged | Q8BGC4 | 14 | 145 | 14 | 377 | 40.5 | 7.42 | 55 | 1.117 | 0.687 | 1.750 | 0.073 | 1.574 | 0.226 | 1.112 | 0.718 |
| 658 | **Cyp3a11** | UP | Q64459 | 26 | 291 | 25 | 504 | 57.8 | 8.82 | 55 | 1.693 | 0.020 | 3.000 | 0.015 | 3.130 | 0.001 | 0.958 | 0.861 |
| 659 | **Psmd5** | Unchanged | Q8BJY1 | 17 | 123 | 17 | 504 | 55.9 | 5.21 | 55 | 0.865 | 0.562 | 1.086 | 0.725 | 0.970 | 0.897 | 1.120 | 0.437 |
| 660 | **Dpysl2** | Unchanged | O08553 | 17 | 166 | 16 | 572 | 62.2 | 6.38 | 55 | 0.806 | 0.318 | 0.822 | 0.483 | 0.913 | 0.683 | 0.900 | 0.719 |
| 661 | **Mrpl12** | Unchanged | Q9DB15 | 9 | 122 | 9 | 201 | 21.7 | 9.29 | 55 | 0.831 | 0.719 | 1.068 | 0.904 | 0.774 | 0.608 | 1.380 | 0.566 |
| 662 | **Hint2** | Unchanged | Q9D0S9 | 6 | 154 | 6 | 163 | 17.3 | 9.82 | 55 | 1.380 | 0.251 | 0.854 | 0.678 | 1.059 | 0.812 | 0.807 | 0.449 |
| 663 | **Rps15** | Unchanged | P62843 | 4 | 165 | 4 | 145 | 17 | 10.39 | 55 | 1.080 | 0.792 | 0.809 | 0.642 | 0.651 | 0.237 | 1.243 | 0.621 |
| 664 | **Hsd17b12** | Unchanged | O70503 | 15 | 147 | 15 | 312 | 34.7 | 9.52 | 55 | 0.727 | 0.255 | 1.270 | 0.562 | 1.089 | 0.719 | 1.166 | 0.688 |
| 665 | **Akr7a2** | Unchanged | Q8CG76 | 16 | 175 | 16 | 367 | 40.6 | 8.12 | 55 | 1.290 | 0.115 | 1.067 | 0.878 | 0.988 | 0.958 | 1.080 | 0.862 |
| 666 | **Cyp2c70** | Down | Q91W64 | 19 | 208 | 19 | 489 | 56 | 8.09 | 55 | 0.814 | 0.415 | 1.300 | 0.161 | 0.647 | 0.033 | 2.010 | 0.005 |
| 667 | **Rps3a** | Unchanged | P97351 | 17 | 261 | 17 | 264 | 29.9 | 9.73 | 55 | 0.447 | 0.091 | 0.678 | 0.294 | 0.631 | 0.209 | 1.075 | 0.687 |
| 668 | **Napa** | Unchanged | Q9DB05 | 11 | 72 | 10 | 295 | 33.2 | 5.45 | 55 | 1.612 | 0.463 | 1.346 | 0.591 | 1.282 | 0.691 | 1.050 | 0.939 |
| 669 | **Tsta3** | Unchanged | P23591 | 11 | 135 | 11 | 321 | 35.9 | 6.74 | 55 | 0.850 | 0.765 | 0.515 | 0.268 | 0.345 | 0.127 | 1.494 | 0.255 |
| 670 | **Prodhiso** | Unchanged | F6XMC0 | 4 | 20 | 1 | 115 | 13.4 | 6.95 | 55 | 0.850 | 0.790 | 0.871 | 0.878 | 0.949 | 0.933 | 0.918 | 0.928 |
| 671 | **Psma5** | Unchanged | Q9Z2U1 | 11 | 196 | 11 | 241 | 26.4 | 4.79 | 55 | 0.860 | 0.227 | 1.066 | 0.702 | 0.923 | 0.452 | 1.155 | 0.425 |
| 672 | **Acad8** | Unchanged | Q9D7B6 | 18 | 149 | 18 | 413 | 45 | 8.13 | 55 | 0.972 | 0.906 | 0.831 | 0.647 | 1.269 | 0.202 | 0.655 | 0.264 |
| 673 | **Apoe** | UP | P08226 | 15 | 351 | 14 | 311 | 35.8 | 5.68 | 55 | 0.972 | 0.955 | 0.923 | 0.870 | 0.979 | 0.960 | 0.943 | 0.899 |
| 674 | **Cyp2c23** | Unchanged | E9Q5K4 | 21 | 116 | 20 | 494 | 56.4 | 8.47 | 55 | 0.389 | 0.109 | 0.668 | 0.432 | 0.970 | 0.946 | 0.689 | 0.439 |
| 675 | **Capzb** | Unchanged | A2AMW0 | 12 | 92 | 12 | 260 | 29.3 | 6.92 | 55 | 1.052 | 0.865 | 1.267 | 0.564 | 0.953 | 0.884 | 1.330 | 0.521 |
| 676 | **RTRAF** | UP | Q9CQE8 | 11 | 88 | 11 | 244 | 28.1 | 6.89 | 55 | 1.461 | 0.249 | 1.813 | 0.280 | 1.659 | 0.020 | 1.093 | 0.836 |
| 677 | **Mettl7a1** | Unchanged | Q8C6B0 | 9 | 59 | 9 | 244 | 28.1 | 8.31 | 55 | 0.835 | 0.715 | 1.573 | 0.326 | 1.091 | 0.871 | 1.442 | 0.435 |
| 678 | **Prpsap1** | UP | Q9D0M1 | 13 | 115 | 12 | 356 | 39.4 | 7.2 | 54 | 1.407 | 0.152 | 1.869 | 0.028 | 1.789 | 0.019 | 1.045 | 0.848 |
| 679 | **Pcbp1** | Unchanged | P60335 | 12 | 184 | 9 | 356 | 37.5 | 7.09 | 54 | 0.783 | 0.661 | 1.090 | 0.839 | 0.862 | 0.761 | 1.265 | 0.314 |
| 680 | **Kpnb1** | UP | P70168 | 27 | 244 | 27 | 876 | 97.1 | 4.78 | 54 | 1.567 | 0.133 | 3.643 | 0.019 | 1.346 | 0.323 | 2.707 | 0.036 |
| 681 | **Gbe1** | Unchanged | Q9D6Y9 | 29 | 403 | 29 | 702 | 80.3 | 6.43 | 54 | 1.228 | 0.142 | 1.173 | 0.302 | 1.297 | 0.046 | 0.904 | 0.366 |
| 682 | **Ttc38** | Unchanged | A3KMP2 | 18 | 201 | 18 | 465 | 52.2 | 6.3 | 54 | 1.182 | 0.722 | 0.915 | 0.838 | 0.945 | 0.892 | 0.969 | 0.936 |
| 683 | **Prdx4** | Unchanged | O08807 | 12 | 209 | 10 | 274 | 31 | 7.15 | 54 | 0.777 | 0.247 | 1.221 | 0.325 | 1.185 | 0.480 | 1.030 | 0.877 |
| 684 | **Sephs1** | Unchanged | Q8BH69 | 11 | 104 | 11 | 392 | 42.9 | 5.97 | 54 | 0.967 | 0.913 | 1.398 | 0.272 | 1.236 | 0.491 | 1.131 | 0.596 |
| 685 | **Fkbp2** | Unchanged | P45878 | 7 | 105 | 7 | 140 | 15.3 | 8.88 | 54 | 0.805 | 0.522 | 1.333 | 0.241 | 0.923 | 0.832 | 1.444 | 0.230 |
| 686 | **Psmc2** | Unchanged | P46471 | 24 | 214 | 24 | 433 | 48.6 | 5.95 | 54 | 1.066 | 0.675 | 1.227 | 0.194 | 0.810 | 0.012 | 1.514 | 0.037 |

| | | | | | | | | | | | | | | | | | | |
|---|---|---|---|---|---|---|---|---|---|---|---|---|---|---|---|---|---|---|
| 687 | **Mdp1** | Unchanged | Q9D967 | 5 | 45 | 5 | 164 | 18.6 | 6.8 | 54 | 0.381 | 0.089 | 0.804 | 0.642 | 1.422 | 0.484 | 0.566 | 0.304 |
| 688 | **Psmb1** | Unchanged | O09061 | 12 | 230 | 12 | 240 | 26.4 | 7.81 | 54 | 1.271 | 0.442 | 1.567 | 0.472 | 1.110 | 0.668 | 1.411 | 0.566 |
| 689 | **Ighv1-42** | Unchanged | A0A075B5V6 | 3 | 29 | 1 | 98 | 10.8 | 8.73 | 54 | 1.667 | 0.149 | 90.095 | 0.068 | 97.733 | 0.085 | 0.922 | 0.909 |
| 690 | **Sec31a** | UP | S4R192 | 12 | 60 | 1 | 248 | 27.8 | 5.57 | 54 | 1.281 | 0.692 | 0.761 | 0.772 | 1.595 | 0.516 | 0.477 | 0.446 |
| 691 | **Copa** | Unchanged | Q8CIE6 | 46 | 365 | 46 | 1224 | 138.3 | 7.65 | 54 | 0.732 | 0.196 | 1.061 | 0.755 | 0.968 | 0.879 | 1.097 | 0.393 |
| 692 | **Tufm** | Down | Q8BFR5 | 18 | 181 | 18 | 452 | 49.5 | 7.56 | 54 | 0.619 | 0.050 | 1.169 | 0.615 | 0.493 | 0.011 | 2.370 | 0.047 |
| 693 | **Bdh1** | Unchanged | Q80XN0 | 18 | 336 | 18 | 343 | 38.3 | 9.01 | 54 | 1.165 | 0.653 | 1.489 | 0.404 | 1.101 | 0.827 | 1.352 | 0.579 |
| 694 | **Tln1** | Unchanged | P26039 | 80 | 792 | 74 | 2541 | 269.7 | 6.18 | 54 | 0.995 | 0.971 | 1.302 | 0.243 | 1.036 | 0.710 | 1.257 | 0.284 |
| 695 | **Pxmp2** | Unchanged | Q5D073 | 8 | 53 | 8 | 193 | 22.1 | 9.99 | 54 | 0.821 | 0.622 | 1.331 | 0.531 | 0.895 | 0.794 | 1.487 | 0.386 |
| 696 | **Ptbp1** | Unchanged | Q922I7 | 18 | 116 | 15 | 555 | 59.3 | 9.16 | 54 | 0.898 | 0.320 | 1.204 | 0.478 | 0.689 | 0.078 | 1.747 | 0.133 |
| 697 | **Memo1** | Unchanged | Q91VH6 | 11 | 64 | 11 | 297 | 33.7 | 7.14 | 54 | 1.350 | 0.341 | 2.002 | 0.081 | 1.628 | 0.089 | 1.230 | 0.408 |
| 698 | **Kyat3** | Unchanged | Q71RI9 | 18 | 185 | 18 | 455 | 51.1 | 8.37 | 54 | 0.854 | 0.582 | 1.054 | 0.874 | 0.948 | 0.882 | 1.111 | 0.792 |
| 699 | **Thnsl2** | Unchanged | Q80W22 | 18 | 177 | 18 | 483 | 54.2 | 6.39 | 54 | 0.911 | 0.735 | 2.540 | 0.134 | 1.850 | 0.284 | 1.373 | 0.562 |
| 700 | **Gss** | Unchanged | P51855 | 18 | 212 | 18 | 474 | 52.2 | 5.8 | 54 | 1.320 | 0.304 | 1.633 | 0.286 | 1.161 | 0.485 | 1.407 | 0.410 |
| 701 | **Ces1c** | Unchanged | P23953 | 21 | 265 | 15 | 554 | 61 | 5.06 | 54 | 0.807 | 0.275 | 1.673 | 0.056 | 0.969 | 0.862 | 1.726 | 0.030 |
| 702 | **Hibch** | Unchanged | Q8QZS1 | 19 | 258 | 19 | 385 | 43 | 8.06 | 54 | 0.917 | 0.712 | 0.838 | 0.542 | 0.715 | 0.038 | 1.172 | 0.621 |
| 703 | **Serpina1b** | Unchanged | P22599 | 15 | 271 | 5 | 413 | 45.9 | 5.54 | 54 | 1.215 | 0.358 | 1.210 | 0.272 | 1.240 | 0.329 | 0.976 | 0.916 |
| 704 | **Psmd12** | Down | Q9D8W5 | 20 | 179 | 20 | 456 | 52.9 | 7.06 | 54 | 0.941 | 0.797 | 1.112 | 0.766 | 0.582 | 0.042 | 1.912 | 0.174 |
| 705 | **Gdi1** | Unchanged | P50396 | 20 | 236 | 14 | 447 | 50.5 | 5.08 | 54 | 1.246 | 0.261 | 1.452 | 0.265 | 1.651 | 0.053 | 0.880 | 0.673 |
| 706 | **Hsdl2** | Unchanged | Q2TPA8 | 16 | 146 | 16 | 490 | 54.2 | 6.74 | 54 | 1.097 | 0.497 | 1.012 | 0.962 | 1.018 | 0.898 | 0.994 | 0.981 |
| 707 | **Rps7** | Unchanged | P62082 | 10 | 219 | 10 | 194 | 22.1 | 10.1 | 54 | 0.865 | 0.658 | 1.125 | 0.707 | 0.976 | 0.929 | 1.153 | 0.514 |
| 708 | **Cyc1** | Unchanged | Q9D0M3 | 13 | 273 | 13 | 325 | 35.3 | 9.16 | 54 | 0.786 | 0.202 | 0.985 | 0.956 | 0.812 | 0.309 | 1.213 | 0.593 |
| 709 | **Isoc1** | UP | Q91V64 | 12 | 156 | 12 | 297 | 32 | 7.39 | 54 | 1.140 | 0.449 | 1.622 | 0.014 | 1.570 | 0.018 | 1.033 | 0.798 |
| 710 | **Ppp1cb** | Unchanged | P62141 | 12 | 149 | 4 | 327 | 37.2 | 6.19 | 54 | 1.275 | 0.642 | 1.969 | 0.254 | 0.945 | 0.926 | 2.083 | 0.274 |
| 711 | **Ddx39b** | Unchanged | Q9Z1N5 | 17 | 154 | 10 | 428 | 49 | 5.67 | 54 | 1.127 | 0.463 | 1.132 | 0.681 | 1.373 | 0.147 | 0.825 | 0.506 |
| 712 | **Akr1e2** | Unchanged | Q9DCT1 | 11 | 83 | 10 | 301 | 34.4 | 7.33 | 53 | 1.008 | 0.987 | 0.988 | 0.982 | 0.985 | 0.972 | 1.003 | 0.995 |
| 713 | **Rdh16f2** | UP | Q8K3M1 | 11 | 58 | 8 | 318 | 35.9 | 8.91 | 53 | 2.948 | 0.016 | 6.529 | 0.000 | 5.888 | 0.014 | 1.109 | 0.720 |
| 714 | **Abcd3** | UP | P55096 | 31 | 463 | 31 | 659 | 75.4 | 9.26 | 53 | 1.105 | 0.653 | 1.615 | 0.019 | 1.409 | 0.044 | 1.146 | 0.328 |
| 715 | **Rps2** | Unchanged | P25444 | 19 | 215 | 19 | 293 | 31.2 | 10.24 | 53 | 0.978 | 0.927 | 0.908 | 0.724 | 0.705 | 0.109 | 1.288 | 0.452 |
| 716 | **Psmc6** | Unchanged | P62334 | 15 | 187 | 15 | 389 | 44.1 | 7.49 | 53 | 1.225 | 0.499 | 0.813 | 0.545 | 0.885 | 0.617 | 0.918 | 0.773 |
| 717 | **Igkv3-4** | Unchanged | P01665 | 3 | 36 | 3 | 111 | 12 | 4.61 | 53 | 0.492 | 0.120 | 0.678 | 0.328 | 0.934 | 0.798 | 0.726 | 0.414 |
| 718 | **Gys2** | UP | Q8VCB3 | 28 | 208 | 28 | 704 | 80.8 | 6.89 | 53 | 1.514 | 0.123 | 1.642 | 0.030 | 1.565 | 0.184 | 1.049 | 0.821 |
| 719 | **Naxd** | Unchanged | Q9CZ42 | 13 | 96 | 13 | 343 | 36.7 | 7.77 | 53 | 0.959 | 0.836 | 1.234 | 0.310 | 0.984 | 0.919 | 1.254 | 0.202 |
| 720 | **Cyp4f14** | Unchanged | Q9EP75 | 17 | 116 | 9 | 524 | 59.8 | 6.87 | 53 | 0.972 | 0.933 | 1.487 | 0.433 | 1.203 | 0.571 | 1.236 | 0.658 |
| 721 | **Apeh** | Unchanged | Q8R146 | 21 | 187 | 21 | 732 | 81.5 | 5.59 | 53 | 0.774 | 0.408 | 0.797 | 0.464 | 0.977 | 0.938 | 0.815 | 0.370 |
| 722 | **Psmd1** | Unchanged | Q3TXS7 | 32 | 328 | 32 | 953 | 105.7 | 5.39 | 53 | 1.313 | 0.290 | 1.148 | 0.140 | 1.179 | 0.465 | 0.974 | 0.895 |
| 723 | **Acot1** | UP | O55137 | 17 | 180 | 5 | 419 | 46.1 | 6.58 | 53 | 1.644 | 0.025 | 1.770 | 0.085 | 1.900 | 0.003 | 0.932 | 0.732 |
| 724 | **Cct8** | UP | P42932 | 29 | 238 | 29 | 548 | 59.5 | 5.62 | 53 | 1.845 | 0.026 | 1.301 | 0.339 | 0.948 | 0.817 | 1.373 | 0.323 |
| 725 | **Tgm2** | Unchanged | P21981 | 24 | 262 | 24 | 686 | 77 | 5.1 | 53 | 1.245 | 0.612 | 0.806 | 0.561 | 1.179 | 0.631 | 0.684 | 0.069 |
| 726 | **Rpl38** | Unchanged | Q9JJI8 | 7 | 109 | 7 | 70 | 8.2 | 10.1 | 53 | 0.999 | 0.998 | 0.983 | 0.974 | 1.891 | 0.071 | 0.520 | 0.169 |
| 727 | **Capns1** | Unchanged | O88456 | 8 | 53 | 8 | 269 | 28.4 | 5.63 | 53 | 0.620 | 0.098 | 0.744 | 0.307 | 0.535 | 0.050 | 1.390 | 0.226 |

| | | | | | | | | | | | | | | | | | | |
|---|---|---|---|---|---|---|---|---|---|---|---|---|---|---|---|---|---|---|
| 728 | **Cisd1** | Unchanged | Q91WS0 | 7 | 82 | 7 | 108 | 12.1 | 9.06 | 53 | 1.130 | 0.491 | 0.747 | 0.318 | 0.845 | 0.339 | 0.884 | 0.686 |
| 729 | **Ces2c** | Down | Q91WG0 | 20 | 235 | 13 | 561 | 62.4 | 5.69 | 53 | 0.330 | 0.041 | 0.337 | 0.041 | 0.267 | 0.028 | 1.260 | 0.126 |
| 730 | **Ckm** | UP | P07310 | 17 | 61 | 17 | 381 | 43 | 7.06 | 53 | 0.775 | 0.703 | 0.763 | 0.676 | 9.373 | 0.030 | 0.081 | 0.026 |
| 731 | **Farsa** | Unchanged | Q8C0C7 | 17 | 167 | 17 | 508 | 57.6 | 8.28 | 53 | 1.175 | 0.354 | 1.743 | 0.085 | 1.018 | 0.899 | 1.713 | 0.086 |
| 732 | **Pdia6** | Unchanged | Q922R8 | 24 | 605 | 2 | 440 | 48.1 | 5.14 | 53 | 1.184 | 0.698 | 18.241 | 0.275 | 20.474 | 0.154 | 0.891 | 0.910 |
| 733 | **Acsl5** | Unchanged | Q8JZR0 | 29 | 376 | 26 | 683 | 76.2 | 7.09 | 53 | 0.965 | 0.859 | 0.812 | 0.249 | 0.730 | 0.107 | 1.112 | 0.674 |
| 734 | **Apip** | UP | Q9WVQ5 | 9 | 42 | 9 | 241 | 26.9 | 6.9 | 53 | 0.858 | 0.627 | 1.958 | 0.042 | 0.932 | 0.802 | 2.101 | 0.036 |
| 735 | **Pon1** | Unchanged | P52430 | 14 | 157 | 14 | 355 | 39.5 | 5.22 | 51 | 0.731 | 0.274 | 1.066 | 0.821 | 0.876 | 0.660 | 1.216 | 0.471 |
| 736 | **Hpx** | Unchanged | Q91X72 | 25 | 280 | 25 | 460 | 51.3 | 7.8 | 53 | 0.553 | 0.163 | 0.628 | 0.235 | 0.532 | 0.119 | 1.181 | 0.513 |
| 737 | **Rps17** | UP | P63276 | 7 | 183 | 7 | 135 | 15.5 | 9.85 | 53 | 1.205 | 0.215 | 1.129 | 0.430 | 1.788 | 0.002 | 0.631 | 0.003 |
| 738 | **Hdlbp** | Unchanged | Q8VDJ3 | 54 | 652 | 54 | 1268 | 141.7 | 6.87 | 53 | 0.881 | 0.571 | 0.886 | 0.692 | 1.084 | 0.735 | 0.817 | 0.547 |
| 739 | **Mettl26** | Unchanged | Q9DCS2 | 8 | 191 | 8 | 204 | 22.7 | 6.52 | 52 | 1.231 | 0.286 | 1.072 | 0.792 | 0.727 | 0.142 | 1.474 | 0.178 |
| 740 | **Nadk2** | Unchanged | Q8C5H8 | 19 | 166 | 19 | 452 | 50.8 | 8.07 | 52 | 0.870 | 0.340 | 1.131 | 0.424 | 1.114 | 0.288 | 1.015 | 0.902 |
| 741 | **Ppp1ca** | Unchanged | P62137 | 13 | 146 | 5 | 330 | 37.5 | 6.33 | 52 | 1.004 | 0.991 | 1.273 | 0.350 | 1.120 | 0.683 | 1.137 | 0.477 |
| 742 | **Slc25a3** | Unchanged | Q8VEM8 | 16 | 173 | 16 | 357 | 39.6 | 9.26 | 52 | 0.882 | 0.689 | 0.679 | 0.215 | 0.681 | 0.245 | 0.997 | 0.994 |
| 743 | **Psmb7** | Unchanged | P70195 | 12 | 101 | 12 | 277 | 29.9 | 7.99 | 52 | 1.245 | 0.722 | 1.617 | 0.343 | 0.741 | 0.648 | 2.181 | 0.044 |
| 744 | **Gckr** | Unchanged | A0A0J9YUI8 | 21 | 122 | 21 | 623 | 68.7 | 7.18 | 52 | 0.894 | 0.732 | 0.896 | 0.715 | 0.688 | 0.237 | 1.302 | 0.296 |
| 745 | **Arpc5** | Unchanged | Q9CPW4 | 4 | 45 | 4 | 151 | 16.3 | 5.67 | 52 | 0.862 | 0.152 | 0.916 | 0.729 | 1.092 | 0.614 | 0.838 | 0.551 |
| 746 | **Hsd3b3** | Down | P26150 | 15 | 129 | 12 | 373 | 42 | 7.97 | 48 | 0.954 | 0.775 | 0.960 | 0.924 | 0.604 | 0.029 | 1.591 | 0.396 |
| 747 | **Rtn4ip1** | Unchanged | Q924D0 | 14 | 73 | 14 | 396 | 43.3 | 9.2 | 52 | 2.538 | 0.211 | 0.796 | 0.660 | 1.010 | 0.981 | 0.788 | 0.448 |
| 748 | **Gatd3a** | Unchanged | Q9D172 | 11 | 151 | 11 | 266 | 28.1 | 8.78 | 52 | 0.823 | 0.527 | 1.490 | 0.310 | 0.870 | 0.627 | 1.714 | 0.208 |
| 749 | **Tmed7** | Unchanged | D3YZZ5 | 8 | 25 | 8 | 224 | 25.2 | 6.89 | 52 | 0.830 | 0.421 | 1.176 | 0.641 | 0.560 | 0.100 | 2.099 | 0.147 |
| 750 | **Ap1b1** | Unchanged | O35643 | 35 | 345 | 18 | 943 | 103.9 | 5.17 | 52 | 1.091 | 0.665 | 1.101 | 0.631 | 0.968 | 0.898 | 1.138 | 0.619 |
| 751 | **Snrpe** | UP | P62305 | 3 | 29 | 3 | 92 | 10.8 | 9.44 | 52 | 1.923 | 0.032 | 3.295 | 0.008 | 2.906 | 0.003 | 1.134 | 0.586 |
| 752 | **Cyp4a12b** | Unchanged | A2A974 | 21 | 79 | 3 | 508 | 58.3 | 8.9 | 52 | 0.987 | 0.961 | 2.275 | 0.240 | 0.699 | 0.086 | 3.254 | 0.156 |
| 753 | **Cpox** | Unchanged | P36552 | 21 | 250 | 21 | 443 | 49.7 | 8.53 | 52 | 1.152 | 0.247 | 1.031 | 0.902 | 1.495 | 0.009 | 0.689 | 0.095 |
| 754 | **Rpl8** | Unchanged | P62918 | 13 | 137 | 13 | 257 | 28 | 11.03 | 52 | 1.098 | 0.785 | 1.027 | 0.960 | 0.638 | 0.287 | 1.610 | 0.484 |
| 755 | **Pdia6iso** | Unchanged | Q3TML0 | 24 | 340 | 2 | 445 | 48.7 | 5.19 | 52 | 0.889 | 0.518 | 1.026 | 0.835 | 1.000 | 1.000 | 1.026 | 0.878 |
| 756 | **Coq7** | UP | P97478 | 7 | 35 | 7 | 217 | 24 | 7.17 | 52 | 3.228 | 0.066 | 4.602 | 0.000 | 3.470 | 0.011 | 1.326 | 0.262 |
| 757 | **Ehd1** | Unchanged | Q9WVK4 | 21 | 119 | 8 | 534 | 60.6 | 6.83 | 52 | 1.530 | 0.178 | 1.743 | 0.070 | 1.418 | 0.061 | 1.229 | 0.414 |
| 758 | **Galk1** | Unchanged | Q9R0N0 | 14 | 191 | 14 | 392 | 42.3 | 5.26 | 52 | 1.444 | 0.415 | 1.278 | 0.428 | 1.132 | 0.489 | 1.129 | 0.700 |
| 759 | **Ndufv2** | Unchanged | Q9D6J6 | 10 | 228 | 10 | 248 | 27.3 | 7.4 | 52 | 1.143 | 0.524 | 1.407 | 0.053 | 0.951 | 0.750 | 1.480 | 0.026 |
| 760 | **Arhgdia** | Unchanged | Q99PT1 | 9 | 203 | 9 | 204 | 23.4 | 5.2 | 52 | 0.787 | 0.300 | 1.323 | 0.473 | 1.141 | 0.631 | 1.159 | 0.712 |
| 761 | **Psmb5** | Unchanged | O55234 | 12 | 93 | 12 | 264 | 28.5 | 7.02 | 52 | 1.023 | 0.964 | 0.617 | 0.377 | 0.614 | 0.352 | 1.004 | 0.986 |
| 762 | **Fgg** | Unchanged | Q8VCM7 | 17 | 175 | 17 | 436 | 49.4 | 5.86 | 52 | 1.053 | 0.843 | 1.247 | 0.397 | 1.232 | 0.310 | 1.013 | 0.959 |
| 763 | **Mccc1** | Unchanged | Q99MR8 | 25 | 348 | 25 | 717 | 79.3 | 7.83 | 52 | 0.926 | 0.884 | 1.111 | 0.861 | 0.485 | 0.213 | 2.289 | 0.237 |
| 764 | **Fuom** | Unchanged | Q8R2K1 | 5 | 53 | 5 | 153 | 16.8 | 5.36 | 52 | 1.415 | 0.567 | 3.456 | 0.151 | 2.271 | 0.054 | 1.522 | 0.482 |
| 765 | **Rdx** | Unchanged | P26043 | 31 | 528 | 21 | 583 | 68.5 | 6.2 | 52 | 1.134 | 0.595 | 0.967 | 0.859 | 1.136 | 0.353 | 0.852 | 0.390 |
| 766 | **Farsb** | Unchanged | Q9WUA2 | 21 | 166 | 21 | 589 | 65.7 | 7.12 | 52 | 1.132 | 0.493 | 1.367 | 0.220 | 0.810 | 0.404 | 1.689 | 0.117 |
| 767 | **Actr1b** | Unchanged | Q8R5C5 | 12 | 99 | 5 | 376 | 42.3 | 6.4 | 52 | 1.888 | 0.412 | 1.568 | 0.365 | 0.675 | 0.148 | 2.324 | 0.159 |
| 768 | **Aimp2** | Unchanged | Q8R010 | 7 | 77 | 7 | 320 | 35.4 | 7.83 | 52 | 1.004 | 0.988 | 1.091 | 0.752 | 0.955 | 0.824 | 1.142 | 0.632 |

| | | | | | | | | | | | | | | | | | | |
|---|---|---|---|---|---|---|---|---|---|---|---|---|---|---|---|---|---|---|
| 769 | **Hnrnpa2b1** | Unchanged | O88569 | 18 | 311 | 16 | 353 | 37.4 | 8.95 | 52 | 0.794 | 0.265 | 0.751 | 0.229 | 0.874 | 0.505 | 0.860 | 0.394 |
| 770 | **Ubxn1** | Unchanged | Q922Y1 | 7 | 20 | 7 | 297 | 33.6 | 5.26 | 52 | 0.640 | 0.372 | 1.827 | 0.262 | 2.796 | 0.096 | 0.653 | 0.405 |
| 771 | **Dars** | Unchanged | Q922B2 | 23 | 243 | 23 | 501 | 57.1 | 6.49 | 51 | 0.883 | 0.540 | 0.870 | 0.382 | 1.056 | 0.834 | 0.824 | 0.430 |
| 772 | **Naxe** | Unchanged | Q8K4Z3 | 10 | 133 | 10 | 282 | 31 | 7.69 | 51 | 0.911 | 0.480 | 1.136 | 0.374 | 0.930 | 0.592 | 1.222 | 0.092 |
| 773 | **Ndufb9** | UP | Q9CQJ8 | 6 | 85 | 6 | 179 | 22 | 7.8 | 51 | 0.956 | 0.793 | 1.950 | 0.021 | 1.301 | 0.265 | 1.499 | 0.117 |
| 774 | **Stip1** | Down | Q60864 | 29 | 270 | 29 | 543 | 62.5 | 6.8 | 51 | 0.819 | 0.370 | 0.782 | 0.279 | 0.609 | 0.022 | 1.284 | 0.261 |
| 775 | **Cox5a** | Unchanged | P12787 | 8 | 134 | 8 | 146 | 16.1 | 6.54 | 51 | 1.173 | 0.626 | 1.259 | 0.549 | 1.294 | 0.251 | 0.973 | 0.941 |
| 776 | **Rps12** | Unchanged | A0A1W2P7A1 | 7 | 68 | 7 | 148 | 16 | 8.16 | 51 | 1.005 | 0.993 | 0.674 | 0.523 | 0.628 | 0.402 | 1.074 | 0.903 |
| 777 | **H6pd** | UP | Q8CFX1 | 28 | 328 | 28 | 789 | 88.9 | 6.93 | 51 | 1.163 | 0.477 | 2.438 | 0.006 | 1.132 | 0.261 | 2.154 | 0.009 |
| 778 | **Actr1a** | UP | P61164 | 11 | 98 | 4 | 376 | 42.6 | 6.64 | 51 | 1.937 | 0.002 | 0.882 | 0.741 | 1.764 | 0.028 | 0.500 | 0.050 |
| 779 | **Yars** | Down | Q91WQ3 | 21 | 106 | 21 | 528 | 59.1 | 7.01 | 51 | 0.770 | 0.171 | 0.567 | 0.020 | 0.505 | 0.007 | 1.122 | 0.685 |
| 780 | **Rpl30** | Unchanged | P62889 | 5 | 93 | 5 | 115 | 12.8 | 9.63 | 51 | 1.033 | 0.924 | 1.260 | 0.571 | 0.912 | 0.774 | 1.382 | 0.446 |
| 781 | **Usp5** | Unchanged | Q3U4W8 | 25 | 150 | 25 | 835 | 93.3 | 5.06 | 51 | 0.557 | 0.122 | 0.790 | 0.512 | 0.483 | 0.075 | 1.633 | 0.132 |
| 782 | **Tsfm** | UP | Q9CZR8 | 10 | 35 | 10 | 324 | 35.3 | 7.06 | 51 | 1.656 | 0.070 | 2.380 | 0.031 | 1.576 | 0.146 | 1.510 | 0.222 |
| 783 | **Mthfsl** | UP | L7N466 | 7 | 35 | 2 | 203 | 23.2 | 6.7 | 51 | 1.578 | 0.132 | 1.419 | 0.425 | 1.701 | 0.037 | 0.834 | 0.532 |
| 784 | **Xylb** | Unchanged | Q3TNA1 | 19 | 255 | 19 | 551 | 59.5 | 6.9 | 51 | 0.869 | 0.623 | 1.730 | 0.288 | 0.781 | 0.272 | 2.214 | 0.165 |
| 785 | **Suclg1** | Unchanged | Q9WUM5 | 15 | 280 | 15 | 346 | 36.1 | 9.39 | 51 | 0.924 | 0.807 | 0.841 | 0.524 | 0.858 | 0.615 | 0.980 | 0.938 |
| 786 | **Rpl10a** | Unchanged | Q5XJF6 | 12 | 137 | 12 | 217 | 24.8 | 9.94 | 51 | 0.960 | 0.888 | 0.598 | 0.141 | 1.288 | 0.473 | 0.464 | 0.104 |
| 787 | **Ugt2a3** | Unchanged | Q8BWQ1 | 22 | 306 | 19 | 534 | 61.1 | 7.93 | 50 | 0.655 | 0.325 | 0.834 | 0.743 | 0.670 | 0.338 | 1.245 | 0.743 |
| 788 | **Lta4h** | Unchanged | P24527 | 21 | 244 | 21 | 611 | 69 | 6.42 | 48 | 1.250 | 0.422 | 1.282 | 0.333 | 1.445 | 0.313 | 0.888 | 0.734 |
| 789 | **Eif3b** | Unchanged | Q8JZQ9 | 30 | 185 | 30 | 803 | 91.3 | 5.02 | 51 | 0.484 | 0.110 | 1.483 | 0.326 | 0.654 | 0.342 | 2.266 | 0.077 |
| 790 | **Arpc2** | Unchanged | Q9CVB6 | 14 | 166 | 14 | 300 | 34.3 | 7.36 | 51 | 1.161 | 0.718 | 0.853 | 0.634 | 0.752 | 0.517 | 1.134 | 0.791 |
| 791 | **Stoml2** | Unchanged | Q99JB2 | 11 | 191 | 4 | 353 | 38.4 | 8.87 | 51 | 1.793 | 0.214 | 1.356 | 0.187 | 0.953 | 0.854 | 1.422 | 0.150 |
| 792 | **Mri1** | Unchanged | Q9CQT1 | 10 | 63 | 10 | 369 | 39.4 | 5.91 | 51 | 0.936 | 0.823 | 0.837 | 0.499 | 0.776 | 0.478 | 1.078 | 0.790 |
| 793 | **Oplah** | Unchanged | Q8K010 | 42 | 432 | 42 | 1288 | 137.5 | 6.28 | 51 | 0.991 | 0.963 | 1.035 | 0.848 | 0.920 | 0.579 | 1.125 | 0.523 |
| 794 | **Otub1** | Unchanged | Q7TQI3 | 9 | 92 | 9 | 271 | 31.3 | 4.94 | 51 | 1.137 | 0.618 | 1.132 | 0.802 | 1.198 | 0.650 | 0.945 | 0.916 |
| 795 | **Mtpn** | Unchanged | P62774 | 5 | 72 | 5 | 118 | 12.9 | 5.52 | 51 | 1.061 | 0.843 | 1.049 | 0.853 | 1.014 | 0.961 | 1.034 | 0.871 |
| 796 | **Cyp2a5** | UP | P20852 | 21 | 151 | 18 | 494 | 56.7 | 9.2 | 51 | 1.532 | 0.056 | 1.804 | 0.016 | 0.888 | 0.563 | 2.031 | 0.016 |
| 797 | **Tpm3** | Unchanged | E9Q7Q3 | 22 | 141 | 7 | 248 | 28.7 | 4.72 | 51 | 0.809 | 0.168 | 1.166 | 0.687 | 2.654 | 0.078 | 0.439 | 0.136 |
| 798 | **Eprs** | Unchanged | Q8CGC7 | 59 | 539 | 59 | 1512 | 170 | 7.66 | 51 | 0.916 | 0.672 | 0.940 | 0.772 | 0.787 | 0.231 | 1.195 | 0.383 |
| 799 | **Ap2a2** | Unchanged | P17427 | 33 | 316 | 27 | 938 | 104 | 6.93 | 51 | 0.802 | 0.175 | 0.866 | 0.577 | 0.728 | 0.072 | 1.189 | 0.602 |
| 800 | **Tomm22** | Unchanged | Q9CPQ3 | 4 | 16 | 4 | 142 | 15.5 | 4.34 | 51 | 0.944 | 0.783 | 0.977 | 0.916 | 0.793 | 0.418 | 1.233 | 0.519 |
| 801 | **Cyb5b** | Unchanged | Q9CQX2 | 4 | 117 | 4 | 146 | 16.3 | 4.89 | 51 | 1.123 | 0.717 | 1.089 | 0.798 | 0.859 | 0.506 | 1.268 | 0.431 |
| 802 | **Echdc3** | Unchanged | Q9D7J9 | 12 | 136 | 12 | 300 | 32.4 | 8.34 | 51 | 1.219 | 0.063 | 1.275 | 0.061 | 1.220 | 0.083 | 1.045 | 0.699 |
| 803 | **Igha** | UP | A0A0A6YXW6 | 11 | 80 | 11 | 389 | 42 | 4.94 | 51 | 13.054 | 0.070 | 2.517 | 0.042 | 1.461 | 0.320 | 1.723 | 0.121 |
| 804 | **Acp1** | UP | Q561M1 | 6 | 25 | 1 | 158 | 17.9 | 6.54 | 51 | 1.050 | 0.877 | 0.911 | 0.795 | 0.925 | 0.798 | 0.984 | 0.972 |
| 805 | **Coasy** | Unchanged | Q9DBL7 | 15 | 164 | 15 | 563 | 62 | 7.11 | 51 | 1.032 | 0.907 | 1.113 | 0.810 | 0.973 | 0.899 | 1.145 | 0.778 |
| 806 | **Selenof** | Down | Q9ERR7 | 6 | 31 | 6 | 162 | 17.8 | 5.35 | 51 | 0.874 | 0.742 | 1.118 | 0.824 | 0.399 | 0.042 | 2.804 | 0.194 |
| 807 | **Myl12a** | UP | Q6ZWQ9 | 7 | 38 | 4 | 172 | 19.9 | 4.81 | 51 | 1.561 | 0.199 | 1.502 | 0.140 | 1.588 | 0.037 | 0.946 | 0.771 |
| 808 | **Mpv17** | UP | P19258 | 4 | 17 | 4 | 176 | 19.7 | 9.44 | 51 | 7.426 | 0.016 | 11.566 | 0.000 | 8.098 | 0.007 | 1.428 | 0.229 |
| 809 | **Trap1** | Unchanged | Q9CQN1 | 28 | 386 | 27 | 706 | 80.2 | 6.68 | 51 | 1.179 | 0.490 | 1.373 | 0.409 | 1.465 | 0.113 | 0.938 | 0.859 |

| | | | | | | | | | | | | | | | | | | |
|---|---|---|---|---|---|---|---|---|---|---|---|---|---|---|---|---|---|---|
| 810 | **Vdac3** | Down | Q60931 | 11 | 137 | 10 | 283 | 30.7 | 8.79 | 51 | 0.969 | 0.820 | 1.165 | 0.338 | 0.596 | 0.008 | 1.954 | 0.004 |
| 811 | **Apoc3** | Down | P33622 | 4 | 100 | 4 | 99 | 11 | 4.75 | 51 | 1.079 | 0.740 | 0.881 | 0.749 | 0.533 | 0.041 | 1.654 | 0.320 |
| 812 | **Fahd2a** | UP | A0A0R4J094 | 9 | 52 | 9 | 313 | 34.7 | 8.16 | 50 | 2.063 | 0.010 | 1.068 | 0.794 | 1.375 | 0.208 | 0.777 | 0.345 |
| 813 | **Calu** | Unchanged | O35887 | 12 | 129 | 2 | 315 | 37 | 4.67 | 50 | 1.727 | 0.419 | 2.520 | 0.206 | 2.425 | 0.187 | 1.039 | 0.949 |
| 814 | **Ugt1a5** | UP | B2RT14 | 19 | 196 | 9 | 529 | 60 | 8.09 | 50 | 1.276 | 0.289 | 1.805 | 0.009 | 1.194 | 0.445 | 1.512 | 0.015 |
| 815 | **Dbt** | Unchanged | P53395 | 17 | 235 | 17 | 482 | 53.2 | 8.6 | 50 | 1.212 | 0.549 | 1.631 | 0.261 | 1.380 | 0.172 | 1.182 | 0.665 |
| 816 | **Bola1** | Unchanged | Q9D8S9 | 5 | 24 | 5 | 137 | 14.4 | 8.76 | 50 | 0.829 | 0.598 | 3.162 | 0.079 | 0.543 | 0.225 | 5.821 | 0.040 |
| 817 | **Rps19** | Unchanged | Q9CZX8 | 10 | 167 | 10 | 145 | 16.1 | 10.4 | 50 | 0.757 | 0.399 | 0.525 | 0.072 | 0.823 | 0.480 | 0.638 | 0.189 |
| 818 | **Cand1** | Unchanged | Q6ZQ38 | 40 | 375 | 38 | 1230 | 136.2 | 5.78 | 50 | 1.224 | 0.551 | 1.420 | 0.289 | 0.704 | 0.221 | 2.018 | 0.063 |
| 819 | **Dld** | Unchanged | O08749 | 18 | 320 | 18 | 509 | 54.2 | 7.9 | 50 | 0.934 | 0.609 | 1.291 | 0.110 | 0.876 | 0.372 | 1.475 | 0.033 |
| 820 | **Uggt1** | Unchanged | Q6P5E4 | 46 | 381 | 46 | 1551 | 176.3 | 5.62 | 50 | 1.020 | 0.940 | 0.913 | 0.674 | 0.926 | 0.726 | 0.985 | 0.954 |
| 821 | **Strap** | Unchanged | Q9Z1Z2 | 11 | 88 | 11 | 350 | 38.4 | 5.12 | 50 | 1.185 | 0.500 | 1.314 | 0.363 | 1.113 | 0.643 | 1.180 | 0.449 |
| 822 | **Nsdhl** | Unchanged | Q9R1J0 | 11 | 84 | 11 | 362 | 40.7 | 7.85 | 50 | 1.375 | 0.427 | 1.309 | 0.578 | 0.917 | 0.772 | 1.428 | 0.489 |
| 823 | **Ola1** | Unchanged | Q9CZ30 | 14 | 108 | 14 | 396 | 44.7 | 7.81 | 50 | 1.110 | 0.777 | 1.159 | 0.662 | 1.087 | 0.825 | 1.066 | 0.719 |
| 824 | **AU022252** | Unchanged | Q5EBG8 | 5 | 68 | 5 | 199 | 21.8 | 5.53 | 50 | 1.389 | 0.327 | 1.313 | 0.041 | 1.072 | 0.831 | 1.225 | 0.498 |
| 825 | **Psmb3** | Unchanged | Q9R1P1 | 10 | 136 | 10 | 205 | 22.9 | 6.55 | 50 | 1.023 | 0.946 | 1.061 | 0.798 | 1.090 | 0.727 | 0.973 | 0.838 |
| 826 | **Psmc1** | Unchanged | P62192 | 18 | 168 | 16 | 440 | 49.2 | 6.21 | 50 | 0.934 | 0.835 | 1.037 | 0.918 | 0.499 | 0.062 | 2.077 | 0.084 |
| 827 | **Fahd1** | UP | Q8R0F8 | 6 | 66 | 6 | 227 | 25.2 | 7.69 | 50 | 0.757 | 0.265 | 0.920 | 0.822 | 1.708 | 0.031 | 0.539 | 0.089 |
| 828 | **Eif3f** | UP | Q9DCH4 | 10 | 137 | 10 | 361 | 38 | 5.58 | 50 | 1.509 | 0.055 | 1.566 | 0.026 | 1.361 | 0.115 | 1.151 | 0.345 |
| 829 | **Cotl1** | UP | Q9CQI6 | 7 | 60 | 7 | 142 | 15.9 | 5.4 | 50 | 2.072 | 0.120 | 3.313 | 0.006 | 2.931 | 0.039 | 1.130 | 0.711 |
| 830 | **Ndufb10** | Unchanged | Q9DCS9 | 9 | 89 | 9 | 176 | 21 | 8.03 | 47 | 1.553 | 0.149 | 1.773 | 0.092 | 1.625 | 0.166 | 1.091 | 0.792 |
| 831 | **Pzp** | Unchanged | Q61838 | 45 | 588 | 44 | 1495 | 165.7 | 6.68 | 50 | 0.915 | 0.577 | 1.131 | 0.451 | 0.907 | 0.553 | 1.246 | 0.197 |
| 832 | **Upb1** | Unchanged | Q8VC97 | 14 | 106 | 14 | 393 | 43.9 | 6.79 | 50 | 1.001 | 0.998 | 0.808 | 0.684 | 0.922 | 0.860 | 0.877 | 0.627 |
| 833 | **Vdac2** | Unchanged | Q60930 | 10 | 141 | 10 | 295 | 31.7 | 7.49 | 50 | 0.996 | 0.990 | 1.221 | 0.508 | 0.907 | 0.798 | 1.345 | 0.106 |
| 834 | **Fggy** | UP | A2AJL3 | 18 | 225 | 18 | 552 | 60.3 | 6.42 | 50 | 1.383 | 0.048 | 1.911 | 0.001 | 1.102 | 0.444 | 1.734 | 0.003 |
| 835 | **Ak4** | Unchanged | Q9WUR9 | 7 | 58 | 7 | 223 | 25 | 7.53 | 50 | 0.927 | 0.793 | 0.809 | 0.582 | 0.946 | 0.806 | 0.855 | 0.670 |
| 836 | **Ap2b1** | UP | Q9DBG3 | 35 | 325 | 18 | 937 | 104.5 | 5.38 | 50 | 0.976 | 0.874 | 2.778 | 0.040 | 1.629 | 0.001 | 1.705 | 0.156 |
| 837 | **Acaca** | Unchanged | Q5SWU9 | 71 | 577 | 64 | 2345 | 265.1 | 6.39 | 50 | 0.917 | 0.481 | 1.273 | 0.198 | 1.024 | 0.852 | 1.243 | 0.259 |
| 838 | **Atp5f1c** | Unchanged | Q91VR2 | 13 | 252 | 1 | 298 | 32.9 | 9.01 | 50 | 0.877 | 0.801 | 0.816 | 0.683 | 0.492 | 0.226 | 1.658 | 0.179 |
| 839 | **Capza2** | Unchanged | P47754 | 8 | 123 | 7 | 286 | 32.9 | 5.85 | 50 | 1.000 | 1.000 | 1.052 | 0.842 | 1.356 | 0.054 | 0.776 | 0.179 |
| 840 | **Micos13** | Unchanged | Q8R404 | 3 | 31 | 3 | 119 | 13.4 | 8.63 | 50 | 1.214 | 0.679 | 1.472 | 0.595 | 0.638 | 0.299 | 2.306 | 0.330 |
| 841 | **Eif2s1** | Unchanged | Q6ZWX6 | 15 | 102 | 15 | 315 | 36.1 | 5.08 | 50 | 0.642 | 0.215 | 0.721 | 0.417 | 0.627 | 0.159 | 1.150 | 0.691 |
| 842 | **Atp5c1** | Down | A2AKU9 | 13 | 122 | 1 | 297 | 32.8 | 9.13 | 49 | 0.329 | 0.023 | 0.308 | 0.014 | 0.195 | 0.006 | 1.576 | 0.254 |
| 843 | **Timm8a1** | Down | Q9WVA2 | 4 | 14 | 4 | 97 | 11 | 5.16 | 49 | 0.555 | 0.137 | 0.513 | 0.105 | 0.242 | 0.006 | 2.117 | 0.214 |
| 844 | **Ndufb8** | Unchanged | Q9D6J5 | 7 | 112 | 7 | 186 | 21.9 | 6.64 | 49 | 1.144 | 0.579 | 1.026 | 0.928 | 1.295 | 0.281 | 0.792 | 0.439 |
| 845 | **Gphn** | Unchanged | Q8BUV3 | 20 | 113 | 20 | 769 | 83.2 | 5.6 | 49 | 0.976 | 0.794 | 0.920 | 0.515 | 1.037 | 0.735 | 0.886 | 0.436 |
| 846 | **Rps11** | Unchanged | P62281 | 7 | 104 | 7 | 158 | 18.4 | 10.3 | 44 | 0.922 | 0.862 | 0.853 | 0.740 | 0.810 | 0.662 | 1.053 | 0.813 |
| 847 | **Etfdh** | Unchanged | Q921G7 | 26 | 356 | 26 | 616 | 68 | 7.58 | 49 | 1.279 | 0.136 | 1.126 | 0.514 | 0.884 | 0.600 | 1.273 | 0.228 |
| 848 | **Hsd17b2** | Unchanged | P51658 | 15 | 187 | 15 | 381 | 41.8 | 8.57 | 49 | 0.787 | 0.520 | 0.805 | 0.540 | 0.847 | 0.681 | 0.950 | 0.836 |
| 849 | **Rps18** | Down | P62270 | 10 | 87 | 10 | 152 | 17.7 | 10.99 | 49 | 0.925 | 0.875 | 0.141 | 0.031 | 0.184 | 0.038 | 0.765 | 0.455 |
| 850 | **Vars** | Unchanged | Q9Z1Q9 | 42 | 466 | 42 | 1263 | 140.1 | 7.77 | 49 | 0.774 | 0.327 | 0.858 | 0.526 | 0.678 | 0.163 | 1.266 | 0.188 |

| | | | | | | | | | | | | | | | | | | |
|---|---|---|---|---|---|---|---|---|---|---|---|---|---|---|---|---|---|---|
| 851 | **Coq8a** | Unchanged | Q60936 | 18 | 143 | 18 | 645 | 71.7 | 6.54 | 49 | 1.305 | 0.643 | 4.989 | 0.189 | 0.608 | 0.365 | 8.205 | 0.150 |
| 852 | **Psma6** | Unchanged | Q9QUM9 | 13 | 160 | 13 | 246 | 27.4 | 6.76 | 49 | 0.794 | 0.167 | 1.317 | 0.077 | 1.296 | 0.242 | 1.016 | 0.935 |
| 853 | **Psmb6** | Unchanged | Q60692 | 7 | 68 | 7 | 238 | 25.4 | 5.11 | 49 | 0.872 | 0.554 | 0.830 | 0.417 | 0.822 | 0.423 | 1.010 | 0.964 |
| 854 | **Mcat** | Unchanged | A0A2R8W722 | 3 | 3 | 1 | 116 | 12.9 | 6.89 | 49 | 0.005 | 0.090 | 1.024 | 0.974 | 0.643 | 0.558 | 1.591 | 0.499 |
| 855 | **Rps10** | Unchanged | P63325 | 9 | 119 | 9 | 165 | 18.9 | 10.15 | 49 | 0.981 | 0.973 | 1.841 | 0.299 | 0.994 | 0.991 | 1.853 | 0.311 |
| 856 | **Rab11a** | Unchanged | P62492 | 8 | 104 | 1 | 216 | 24.4 | 6.57 | 49 | 0.761 | 0.727 | 0.717 | 0.673 | 0.807 | 0.766 | 0.888 | 0.734 |
| 857 | **Isoc2b** | Unchanged | Q9DCC7 | 4 | 25 | 4 | 210 | 23.1 | 8.18 | 49 | 0.459 | 0.212 | 2.146 | 0.054 | 0.535 | 0.329 | 4.011 | 0.006 |
| 858 | **Aifm1** | Unchanged | Q9Z0X1 | 24 | 346 | 24 | 612 | 66.7 | 9.17 | 49 | 0.476 | 0.435 | 0.601 | 0.549 | 0.394 | 0.368 | 1.525 | 0.023 |
| 859 | **Btf3** | Unchanged | Q64152 | 7 | 56 | 7 | 204 | 22 | 9.52 | 49 | 1.074 | 0.806 | 1.217 | 0.434 | 1.084 | 0.822 | 1.123 | 0.640 |
| 860 | **C8g** | Unchanged | Q8VCG4 | 6 | 49 | 6 | 202 | 22.5 | 9.25 | 49 | 1.001 | 0.999 | 0.542 | 0.100 | 1.084 | 0.825 | 0.500 | 0.125 |
| 861 | **Rpn1** | Unchanged | Q91YQ5 | 24 | 315 | 24 | 608 | 68.5 | 6.46 | 48 | 0.959 | 0.929 | 1.141 | 0.745 | 0.849 | 0.691 | 1.344 | 0.476 |
| 862 | **Rsu1** | Unchanged | Q9D031 | 7 | 32 | 7 | 277 | 31.4 | 8.63 | 49 | 1.696 | 0.329 | 2.275 | 0.407 | 1.520 | 0.388 | 1.496 | 0.632 |
| 863 | **Ugt2b34** | Unchanged | Q8K154 | 18 | 171 | 16 | 532 | 60.8 | 7.88 | 49 | 1.131 | 0.571 | 1.436 | 0.359 | 0.970 | 0.931 | 1.481 | 0.409 |
| 864 | **Hmgcs1** | Unchanged | Q8JZK9 | 18 | 161 | 16 | 520 | 57.5 | 5.99 | 46 | 0.908 | 0.656 | 1.383 | 0.389 | 1.170 | 0.497 | 1.182 | 0.655 |
| 865 | **Prdx3** | Unchanged | P20108 | 8 | 209 | 8 | 257 | 28.1 | 7.58 | 49 | 0.940 | 0.645 | 1.132 | 0.582 | 0.877 | 0.481 | 1.291 | 0.348 |
| 866 | **Rab11b** | Unchanged | P46638 | 9 | 115 | 2 | 218 | 24.5 | 5.94 | 49 | 0.751 | 0.309 | 0.782 | 0.488 | 0.689 | 0.249 | 1.135 | 0.762 |
| 867 | **Gstm4** | Down | Q8R5I6 | 11 | 134 | 5 | 218 | 25.5 | 6.93 | 49 | 0.946 | 0.835 | 0.387 | 0.029 | 0.337 | 0.028 | 1.148 | 0.830 |
| 868 | **Eif3i** | Unchanged | Q9QZD9 | 10 | 95 | 10 | 325 | 36.4 | 5.64 | 49 | 0.769 | 0.423 | 0.588 | 0.173 | 0.596 | 0.127 | 0.987 | 0.963 |
| 869 | **Srp54** | Unchanged | P14576 | 16 | 102 | 16 | 504 | 55.7 | 8.75 | 49 | 1.128 | 0.540 | 1.002 | 0.992 | 0.900 | 0.671 | 1.114 | 0.707 |
| 870 | **Rpl7** | Unchanged | P14148 | 19 | 267 | 19 | 270 | 31.4 | 10.89 | 49 | 0.998 | 0.984 | 0.930 | 0.582 | 0.684 | 0.053 | 1.360 | 0.107 |
| 871 | **Usp14** | Unchanged | Q9JMA1 | 18 | 95 | 18 | 493 | 56 | 5.24 | 47 | 2.661 | 0.245 | 2.521 | 0.057 | 1.208 | 0.410 | 2.087 | 0.098 |
| 872 | **Xdh** | UP | Q00519 | 41 | 511 | 40 | 1335 | 146.5 | 7.56 | 48 | 1.341 | 0.135 | 1.686 | 0.003 | 2.153 | 0.015 | 0.783 | 0.224 |
| 873 | **Rab5a** | Down | Q9CQD1 | 7 | 49 | 5 | 215 | 23.6 | 8.15 | 48 | 1.146 | 0.687 | 0.807 | 0.260 | 0.456 | 0.004 | 1.772 | 0.101 |
| 874 | **Eci2iso** | Unchanged | Q9WUR2 | 15 | 251 | 3 | 391 | 43.2 | 8.92 | 48 | 1.566 | 0.050 | 1.783 | 0.036 | 1.580 | 0.039 | 1.128 | 0.602 |
| 875 | **Mthfs** | Unchanged | Q9D110 | 7 | 46 | 2 | 203 | 23.2 | 7.02 | 48 | 1.051 | 0.842 | 1.268 | 0.519 | 1.264 | 0.572 | 1.003 | 0.995 |
| 876 | **Flnb** | Unchanged | Q80X90 | 75 | 672 | 69 | 2602 | 277.7 | 5.71 | 48 | 0.538 | 0.135 | 0.559 | 0.151 | 0.529 | 0.128 | 1.056 | 0.798 |
| 877 | **Purb** | Unchanged | O35295 | 7 | 48 | 6 | 324 | 33.9 | 5.43 | 48 | 1.168 | 0.739 | 5.741 | 0.180 | 1.477 | 0.428 | 3.886 | 0.226 |
| 878 | **Tardbp** | Unchanged | Q921F2 | 10 | 94 | 10 | 414 | 44.5 | 6.7 | 48 | 2.052 | 0.339 | 3.705 | 0.200 | 1.637 | 0.354 | 2.264 | 0.319 |
| 879 | **Echdc2** | Unchanged | Q3TLP5 | 10 | 235 | 10 | 296 | 31.8 | 8.72 | 48 | 1.078 | 0.718 | 1.524 | 0.052 | 1.270 | 0.121 | 1.199 | 0.297 |
| 880 | **Actr2** | Unchanged | P61161 | 13 | 159 | 13 | 394 | 44.7 | 6.74 | 48 | 1.267 | 0.462 | 1.334 | 0.530 | 1.610 | 0.176 | 0.828 | 0.656 |
| 881 | **Arcn1** | Unchanged | Q5XJY5 | 22 | 154 | 22 | 511 | 57.2 | 6.21 | 46 | 1.110 | 0.788 | 1.245 | 0.509 | 1.041 | 0.921 | 1.196 | 0.671 |
| 882 | **Slc25a45** | UP | Q8CFJ7 | 7 | 34 | 7 | 288 | 31.7 | 9.1 | 39 | 1.108 | 0.805 | 2.387 | 0.008 | 1.450 | 0.355 | 1.646 | 0.026 |
| 883 | **Pdhb** | Unchanged | Q9D051 | 15 | 272 | 15 | 359 | 38.9 | 6.87 | 48 | 0.874 | 0.171 | 1.058 | 0.755 | 0.730 | 0.030 | 1.448 | 0.098 |
| 884 | **Rpl23** | Unchanged | P62830 | 8 | 118 | 8 | 140 | 14.9 | 10.51 | 48 | 0.711 | 0.190 | 0.865 | 0.565 | 0.685 | 0.199 | 1.263 | 0.405 |
| 885 | **Fech** | Unchanged | P22315 | 13 | 140 | 13 | 420 | 47.1 | 8.91 | 48 | 0.646 | 0.292 | 0.812 | 0.637 | 0.636 | 0.295 | 1.277 | 0.595 |
| 886 | **Tsn** | Unchanged | Q62348 | 8 | 74 | 8 | 228 | 26.2 | 6.44 | 48 | 1.557 | 0.372 | 2.034 | 0.155 | 2.217 | 0.057 | 0.917 | 0.810 |
| 887 | **Hist1h3b** | UP | P84228 | 5 | 47 | 1 | 136 | 15.4 | 11.27 | 48 | 0.606 | 0.572 | 1.057 | 0.946 | 4.350 | 0.042 | 0.243 | 0.042 |
| 888 | **H3f3c** | Unchanged | P02301 | 5 | 38 | 1 | 136 | 15.3 | 11.14 | 48 | 1.001 | 0.999 | 1.841 | 0.379 | 1.944 | 0.355 | 0.947 | 0.938 |
| 889 | **Hist1h3a** | Unchanged | P68433 | 5 | 27 | 1 | 136 | 15.4 | 11.12 | 48 | 0.691 | 0.464 | 0.474 | 0.173 | 3.155 | 0.357 | 0.150 | 0.255 |
| 890 | **Urahiso** | Unchanged | A0A1B0GSD0 | 6 | 36 | 1 | 153 | 17.1 | 6.4 | 48 | #DIV/0! | #DIV/0! | #DIV/0! | #DIV/0! | #DIV/0! | #DIV/0! | #DIV/0! | #DIV/0! |
| 891 | **Tars** | Unchanged | Q9D0R2 | 32 | 312 | 30 | 722 | 83.3 | 7.36 | 48 | 1.665 | 0.128 | 2.057 | 0.068 | 1.357 | 0.242 | 1.516 | 0.171 |

| | | | | | | | | | | | | | | | | | |
|---|---|---|---|---|---|---|---|---|---|---|---|---|---|---|---|---|---|
| 892 | **Rab2a** | UP | P53994 | 8 | 97 | 8 | 212 | 23.5 | 6.54 | 48 | 1.227 | 0.088 | 1.577 | 0.009 | 0.967 | 0.782 | 1.630 | 0.003 |
| 893 | **Caluiso** | Unchanged | Q6XLQ8 | 13 | 64 | 3 | 315 | 37.1 | 4.59 | 48 | 0.966 | 0.959 | 1.146 | 0.877 | 1.306 | 0.683 | 0.878 | 0.883 |
| 894 | **Sephs2** | Unchanged | P97364 | 14 | 212 | 14 | 452 | 47.8 | 6.06 | 48 | 1.130 | 0.496 | 0.730 | 0.343 | 1.217 | 0.248 | 0.600 | 0.065 |
| 895 | **Hars** | Unchanged | Q61035 | 19 | 137 | 16 | 509 | 57.4 | 6 | 46 | 1.187 | 0.619 | 1.075 | 0.763 | 1.203 | 0.306 | 0.894 | 0.637 |
| 896 | **Echdc1** | Unchanged | Q9D9V3 | 11 | 151 | 11 | 322 | 35.4 | 7.01 | 48 | 1.198 | 0.468 | 1.126 | 0.545 | 1.742 | 0.059 | 0.646 | 0.106 |
| 897 | **Mcts1** | Unchanged | Q9DB27 | 5 | 46 | 5 | 181 | 20.5 | 8.82 | 48 | 1.315 | 0.494 | 1.826 | 0.163 | 1.236 | 0.550 | 1.477 | 0.343 |
| 898 | **Fermt2** | Unchanged | Q8CIB5 | 19 | 143 | 19 | 680 | 77.8 | 6.7 | 48 | 1.599 | 0.135 | 1.852 | 0.213 | 1.222 | 0.421 | 1.516 | 0.370 |
| 899 | **Ndufa2** | Unchanged | Q9CQ75 | 6 | 65 | 5 | 99 | 10.9 | 9.99 | 47 | 1.316 | 0.362 | 0.855 | 0.643 | 0.560 | 0.115 | 1.528 | 0.321 |
| 900 | **Serpina1d** | Unchanged | Q00897 | 14 | 256 | 3 | 413 | 46 | 5.44 | 47 | 1.018 | 0.939 | 0.899 | 0.821 | 1.132 | 0.768 | 0.794 | 0.678 |
| 901 | **Sptan1** | Unchanged | P16546 | 80 | 691 | 80 | 2472 | 284.4 | 5.33 | 47 | 1.041 | 0.911 | 1.161 | 0.693 | 1.447 | 0.278 | 0.803 | 0.486 |
| 902 | **Immt** | Unchanged | Q8CAQ8 | 29 | 282 | 12 | 757 | 83.8 | 6.61 | 47 | 1.107 | 0.716 | 1.358 | 0.312 | 0.837 | 0.456 | 1.623 | 0.112 |
| 903 | **Igkv5-39** | Unchanged | A0A075B5M7 | 3 | 10 | 3 | 95 | 10.3 | 6.25 | 47 | 0.297 | 0.253 | 0.317 | 0.259 | 0.724 | 0.651 | 0.438 | 0.067 |
| 904 | **Vps35** | Unchanged | Q9EQH3 | 29 | 190 | 29 | 796 | 91.7 | 5.44 | 47 | 0.742 | 0.141 | 1.246 | 0.397 | 0.854 | 0.382 | 1.459 | 0.163 |
| 905 | **Ndufb4** | Unchanged | Q9CQC7 | 6 | 74 | 6 | 129 | 15.1 | 9.89 | 47 | 1.084 | 0.714 | 0.804 | 0.362 | 0.973 | 0.898 | 0.827 | 0.330 |
| 906 | **Lyplal1** | Unchanged | E9QLB2 | 7 | 19 | 7 | 239 | 26.3 | 7.88 | 47 | 1.065 | 0.806 | 1.187 | 0.688 | 0.872 | 0.487 | 1.361 | 0.491 |
| 907 | **Hnrnpl** | UP | Q8R081 | 16 | 184 | 16 | 586 | 63.9 | 8.1 | 47 | 13.970 | 0.089 | 17.019 | 0.042 | 1.856 | 0.001 | 9.169 | 0.051 |
| 908 | **Lactb2** | Unchanged | Q99KR3 | 12 | 150 | 12 | 288 | 32.7 | 6.33 | 47 | 1.032 | 0.943 | 1.150 | 0.697 | 0.864 | 0.749 | 1.331 | 0.468 |
| 909 | **Ogdh** | Unchanged | Q60597 | 39 | 455 | 39 | 1023 | 116.4 | 6.83 | 47 | 1.167 | 0.483 | 1.347 | 0.261 | 0.981 | 0.912 | 1.373 | 0.176 |
| 910 | **Rpl31** | Unchanged | P62900 | 7 | 32 | 7 | 125 | 14.5 | 10.54 | 47 | 0.598 | 0.297 | 0.792 | 0.602 | 0.375 | 0.109 | 2.112 | 0.042 |
| 911 | **Timm9** | Unchanged | Q9WV98 | 3 | 12 | 3 | 89 | 10.3 | 7.21 | 47 | 0.731 | 0.662 | 7.128 | 0.114 | 0.427 | 0.280 | 16.706 | 0.086 |
| 912 | **Dnpep** | Unchanged | Q9Z2W0 | 15 | 176 | 15 | 473 | 52.2 | 7.25 | 47 | 1.061 | 0.828 | 1.698 | 0.083 | 1.050 | 0.794 | 1.617 | 0.075 |
| 913 | **Idh3b** | Unchanged | Q91VA7 | 15 | 55 | 15 | 384 | 42.2 | 8.6 | 47 | 0.969 | 0.936 | 0.870 | 0.611 | 1.472 | 0.241 | 0.591 | 0.069 |
| 914 | **Dcxr** | Unchanged | Q91X52 | 9 | 144 | 9 | 244 | 25.7 | 7.31 | 47 | 1.092 | 0.806 | 0.589 | 0.173 | 0.645 | 0.209 | 0.912 | 0.592 |
| 915 | **Dhtkd1** | Unchanged | A2ATU0 | 22 | 245 | 22 | 921 | 102.7 | 6.9 | 47 | 1.285 | 0.306 | 1.418 | 0.193 | 0.911 | 0.713 | 1.556 | 0.072 |
| 916 | **Gnb2** | Unchanged | P62880 | 11 | 85 | 7 | 340 | 37.3 | 6 | 47 | 1.310 | 0.387 | 2.050 | 0.055 | 1.570 | 0.069 | 1.306 | 0.348 |
| 917 | **Plin2** | UP | P43883 | 11 | 72 | 11 | 425 | 46.6 | 6.87 | 47 | 1.673 | 0.131 | 2.522 | 0.326 | 2.536 | 0.040 | 0.994 | 0.993 |
| 918 | **Mug1** | Unchanged | P28665 | 53 | 686 | 35 | 1476 | 165.2 | 6.42 | 47 | 1.101 | 0.504 | 1.477 | 0.231 | 1.433 | 0.270 | 1.031 | 0.932 |
| 919 | **Mtap** | UP | Q9CQ65 | 9 | 120 | 9 | 283 | 31 | 7.14 | 47 | 1.104 | 0.683 | 1.567 | 0.020 | 1.720 | 0.002 | 0.911 | 0.504 |
| 920 | **Ggact** | Unchanged | Q923B0 | 4 | 37 | 4 | 149 | 17.1 | 5.29 | 47 | 2.218 | 0.150 | 3.708 | 0.138 | 2.768 | 0.143 | 1.340 | 0.630 |
| 921 | **Ido2** | Unchanged | Q8R0V5 | 11 | 65 | 11 | 405 | 45.2 | 6.68 | 47 | 1.084 | 0.806 | 1.771 | 0.210 | 1.191 | 0.624 | 1.487 | 0.402 |
| 922 | **Abhd11** | Unchanged | Q8K4F5 | 6 | 47 | 6 | 307 | 33.5 | 9.61 | 47 | 0.918 | 0.886 | 2.182 | 0.338 | 0.880 | 0.838 | 2.480 | 0.306 |
| 923 | **Fkbp3** | Unchanged | Q62446 | 9 | 54 | 9 | 224 | 25.1 | 9.28 | 47 | 0.642 | 0.290 | 1.301 | 0.374 | 1.086 | 0.851 | 1.198 | 0.586 |
| 924 | **Acp1 iso** | Unchanged | Q9D358 | 6 | 31 | 1 | 158 | 18.2 | 6.74 | 47 | 1.329 | 0.557 | 2.527 | 0.042 | 2.201 | 0.189 | 1.148 | 0.727 |
| 925 | **Ppp1r7** | Unchanged | Q3UM45 | 9 | 66 | 9 | 361 | 41.3 | 4.92 | 47 | 0.323 | 0.228 | 0.348 | 0.242 | 0.306 | 0.215 | 1.137 | 0.627 |
| 926 | **Vcl** | Unchanged | Q64727 | 34 | 410 | 34 | 1066 | 116.6 | 6 | 47 | 0.745 | 0.277 | 0.878 | 0.678 | 1.036 | 0.875 | 0.848 | 0.525 |
| 927 | **Acadsb** | Down | Q9DBL1 | 15 | 184 | 15 | 432 | 47.8 | 7.87 | 47 | 0.620 | 0.029 | 0.699 | 0.101 | 0.743 | 0.034 | 0.941 | 0.765 |
| 928 | **Iigp1** | Unchanged | Q9QZ85 | 17 | 246 | 14 | 413 | 47.5 | 6.33 | 47 | 0.881 | 0.373 | 1.016 | 0.943 | 0.859 | 0.304 | 1.183 | 0.438 |
| 929 | **Rpl10** | Unchanged | Q6ZWV3 | 12 | 192 | 12 | 214 | 24.6 | 10.08 | 47 | 0.820 | 0.439 | 0.733 | 0.183 | 0.608 | 0.073 | 1.205 | 0.594 |
| 930 | **Tymp** | Down | Q99N42 | 13 | 97 | 13 | 471 | 49.3 | 7.03 | 47 | 1.239 | 0.537 | 0.471 | 0.028 | 0.287 | 0.006 | 1.639 | 0.094 |
| 931 | **Pter** | Unchanged | Q60866 | 15 | 137 | 15 | 349 | 39.2 | 6.67 | 47 | 0.613 | 0.134 | 0.921 | 0.757 | 0.946 | 0.825 | 0.973 | 0.901 |
| 932 | **Lgals1** | Unchanged | P16045 | 6 | 75 | 6 | 135 | 14.9 | 5.49 | 47 | 1.016 | 0.975 | 0.711 | 0.546 | 0.989 | 0.981 | 0.719 | 0.341 |

| | | | | | | | | | | | | | | | | | | |
|---|---|---|---|---|---|---|---|---|---|---|---|---|---|---|---|---|---|---|
| 933 | **Atp6v1g1** | Unchanged | Q9CR51 | 2 | 5 | 2 | 118 | 13.7 | 7.97 | 47 | 0.829 | 0.634 | 6.514 | 0.131 | 10.563 | 0.142 | 0.617 | 0.564 |
| 934 | **Apoa4** | Unchanged | P06728 | 14 | 59 | 14 | 395 | 45 | 5.47 | 47 | 0.875 | 0.681 | 1.753 | 0.240 | 0.546 | 0.068 | 3.212 | 0.067 |
| 935 | **Lrrc59** | Down | Q922Q8 | 12 | 175 | 12 | 307 | 34.9 | 9.52 | 47 | 1.028 | 0.891 | 0.762 | 0.130 | 0.604 | 0.009 | 1.262 | 0.104 |
| 936 | **Ighg1** | Unchanged | A0A0A6YWR2 | 10 | 69 | 10 | 393 | 43.4 | 6.44 | 47 | 0.420 | 0.104 | 1.564 | 0.460 | 0.758 | 0.493 | 2.062 | 0.267 |
| 937 | **Gnb1** | UP | P62874 | 10 | 47 | 6 | 340 | 37.4 | 6 | 46 | 1.947 | 0.024 | 2.668 | 0.125 | 1.929 | 0.116 | 1.383 | 0.498 |
| 938 | **Rps29** | Unchanged | P62274 | 3 | 16 | 3 | 56 | 6.7 | 10.13 | 27 | 1.306 | 0.518 | 1.327 | 0.426 | 0.599 | 0.227 | 2.213 | 0.076 |
| 939 | **Pcbp2** | Unchanged | Q61990 | 10 | 135 | 7 | 362 | 38.2 | 6.79 | 46 | 0.995 | 0.993 | 0.750 | 0.679 | 0.909 | 0.891 | 0.825 | 0.774 |
| 940 | **Ces1** | Unchanged | Q8VCC2 | 23 | 456 | 19 | 565 | 62.6 | 6 | 46 | 1.490 | 0.081 | 1.188 | 0.500 | 1.470 | 0.146 | 0.809 | 0.333 |
| 941 | **Pitpnb** | Unchanged | Q8JZZ5 | 10 | 22 | 9 | 272 | 31.6 | 6.37 | 46 | 0.648 | 0.128 | 1.011 | 0.968 | 0.650 | 0.102 | 1.555 | 0.131 |
| 942 | **Arf6** | Unchanged | P62331 | 6 | 63 | 6 | 175 | 20.1 | 8.95 | 46 | 2.193 | 0.400 | 1.483 | 0.411 | 3.007 | 0.184 | 0.493 | 0.318 |
| 943 | **Mtatp8** | UP | P03930 | 4 | 27 | 4 | 67 | 7.8 | 9.88 | 46 | 1.870 | 0.069 | 2.453 | 0.231 | 1.935 | 0.009 | 1.268 | 0.662 |
| 944 | **Serpina1a** | Unchanged | P07758 | 14 | 191 | 4 | 413 | 46 | 5.72 | 46 | 2.255 | 0.510 | 2.782 | 0.255 | 2.654 | 0.304 | 1.048 | 0.950 |
| 945 | **Pex19** | Unchanged | Q8VCI5 | 8 | 109 | 8 | 299 | 32.7 | 4.34 | 46 | 1.021 | 0.952 | 1.334 | 0.502 | 1.137 | 0.662 | 1.173 | 0.712 |
| 946 | **Psma2** | Unchanged | P49722 | 9 | 169 | 9 | 234 | 25.9 | 7.43 | 46 | 0.931 | 0.766 | 1.071 | 0.859 | 1.050 | 0.835 | 1.019 | 0.960 |
| 947 | **Stoml2iso** | Unchanged | A2AG39 | 8 | 64 | 1 | 286 | 30.9 | 7.28 | 46 | 1.052 | 0.934 | 1.424 | 0.568 | 1.284 | 0.701 | 1.108 | 0.864 |
| 948 | **Cyp4v2** | Down | Q9DBW0 | 19 | 123 | 19 | 525 | 60.9 | 8.47 | 46 | 0.840 | 0.481 | 1.314 | 0.054 | 0.441 | 0.002 | 2.979 | 0.000 |
| 949 | **Tdo2** | Unchanged | P48776 | 14 | 99 | 14 | 406 | 47.7 | 6.81 | 46 | 0.983 | 0.971 | 0.808 | 0.687 | 0.274 | 0.075 | 2.947 | 0.109 |
| 950 | **Kyat1** | Unchanged | Q8BTY1 | 14 | 206 | 14 | 424 | 47.5 | 6.95 | 46 | 0.959 | 0.768 | 1.061 | 0.772 | 1.688 | 0.057 | 0.629 | 0.117 |
| 951 | **Asna1** | Unchanged | O54984 | 9 | 44 | 9 | 348 | 38.8 | 4.91 | 46 | 0.569 | 0.260 | 0.846 | 0.678 | 0.989 | 0.977 | 0.855 | 0.493 |
| 952 | **Srp19** | Unchanged | Q9D104 | 4 | 26 | 4 | 144 | 16.1 | 9.88 | 46 | 1.204 | 0.780 | 2.587 | 0.258 | 1.781 | 0.064 | 1.452 | 0.554 |
| 953 | **Pctp** | Unchanged | Q5SV41 | 7 | 24 | 7 | 214 | 24.6 | 6.24 | 46 | 1.301 | 0.384 | 1.028 | 0.946 | 0.854 | 0.627 | 1.204 | 0.644 |
| 954 | **Cfl2** | Unchanged | P45591 | 7 | 127 | 6 | 166 | 18.7 | 7.88 | 46 | 1.049 | 0.828 | 0.931 | 0.844 | 1.166 | 0.479 | 0.799 | 0.401 |
| 955 | **Msn** | Unchanged | P26041 | 26 | 255 | 16 | 577 | 67.7 | 6.6 | 44 | 0.867 | 0.618 | 0.812 | 0.472 | 1.092 | 0.706 | 0.744 | 0.318 |
| 956 | **Ctsz** | Unchanged | Q9WUU7 | 9 | 121 | 9 | 306 | 34 | 6.6 | 46 | 0.807 | 0.416 | 1.357 | 0.301 | 1.054 | 0.843 | 1.287 | 0.330 |
| 957 | **Rtcb** | Unchanged | Q99LF4 | 18 | 135 | 18 | 505 | 55.2 | 7.23 | 46 | 1.131 | 0.805 | 1.744 | 0.371 | 0.417 | 0.112 | 4.184 | 0.104 |
| 958 | **Cryzl2** | UP | Q3UNZ8 | 7 | 74 | 7 | 350 | 37.8 | 8.9 | 46 | 1.600 | 0.056 | 1.800 | 0.081 | 1.524 | 0.030 | 1.181 | 0.509 |
| 959 | **Arl8b** | Unchanged | Q9CQW2 | 7 | 58 | 2 | 186 | 21.5 | 8.43 | 46 | 0.866 | 0.577 | 1.203 | 0.535 | 0.562 | 0.099 | 2.141 | 0.025 |
| 960 | **Rps13** | Unchanged | P62301 | 11 | 188 | 11 | 151 | 17.2 | 10.54 | 46 | 1.122 | 0.439 | 0.861 | 0.500 | 1.394 | 0.060 | 0.618 | 0.047 |
| 961 | **Rad23b** | Unchanged | P54728 | 12 | 160 | 11 | 416 | 43.5 | 4.83 | 46 | 0.830 | 0.144 | 0.684 | 0.072 | 0.931 | 0.765 | 0.735 | 0.327 |
| 962 | **Rpl3** | Unchanged | P27659 | 16 | 245 | 16 | 403 | 46.1 | 10.21 | 46 | 1.105 | 0.619 | 0.809 | 0.211 | 0.916 | 0.276 | 0.883 | 0.442 |
| 963 | **Hsd17b11** | Unchanged | Q9EQ06 | 11 | 176 | 9 | 298 | 32.9 | 8.66 | 46 | 1.090 | 0.774 | 0.845 | 0.620 | 1.454 | 0.380 | 0.581 | 0.214 |
| 964 | **Eif3e** | Unchanged | P60229 | 14 | 129 | 14 | 445 | 52.2 | 6.04 | 46 | 1.112 | 0.597 | 1.408 | 0.114 | 0.823 | 0.483 | 1.712 | 0.021 |
| 965 | **Nars** | Unchanged | Q8BP47 | 24 | 330 | 24 | 559 | 64.2 | 5.86 | 46 | 1.045 | 0.779 | 1.215 | 0.288 | 1.137 | 0.510 | 1.069 | 0.777 |
| 966 | **Lgals9** | Down | O08573 | 13 | 141 | 13 | 353 | 40 | 9.31 | 46 | 1.154 | 0.660 | 1.440 | 0.502 | 0.353 | 0.046 | 4.083 | 0.099 |
| 967 | **Ube2i** | Unchanged | P63280 | 5 | 25 | 5 | 158 | 18 | 8.66 | 46 | 1.041 | 0.923 | 1.554 | 0.321 | 0.718 | 0.401 | 2.163 | 0.091 |
| 968 | **Atp6v1a** | UP | P50516 | 20 | 176 | 20 | 617 | 68.3 | 5.58 | 46 | 2.132 | 0.091 | 2.814 | 0.028 | 7.361 | 0.032 | 0.382 | 0.110 |
| 969 | **Nat2** | Down | P50295 | 9 | 65 | 9 | 290 | 33.7 | 5.88 | 46 | 1.405 | 0.057 | 1.175 | 0.291 | 0.634 | 0.014 | 1.854 | 0.014 |
| 970 | **Slc25a22** | Unchanged | Q9D6M3 | 10 | 81 | 10 | 323 | 34.6 | 9.09 | 46 | 0.825 | 0.777 | 1.024 | 0.975 | 0.537 | 0.420 | 1.906 | 0.384 |
| 971 | **Psma3** | Unchanged | O70435 | 11 | 74 | 11 | 255 | 28.4 | 5.44 | 45 | 1.056 | 0.799 | 1.140 | 0.696 | 1.127 | 0.732 | 1.012 | 0.977 |
| 972 | **Psmb4** | Unchanged | P99026 | 7 | 171 | 7 | 264 | 29.1 | 5.64 | 45 | 0.964 | 0.799 | 0.875 | 0.525 | 1.145 | 0.402 | 0.764 | 0.218 |
| 973 | **Eif2s2** | Unchanged | Q99L45 | 14 | 114 | 14 | 331 | 38.1 | 5.8 | 45 | 1.158 | 0.601 | 1.555 | 0.229 | 1.015 | 0.952 | 1.532 | 0.211 |

| | | | | | | | | | | | | | | | | | | |
|---|---|---|---|---|---|---|---|---|---|---|---|---|---|---|---|---|---|---|
| 974 | **Cox5b** | Down | P19536 | 7 | 110 | 7 | 128 | 13.8 | 8.38 | 45 | 0.703 | 0.066 | 0.589 | 0.034 | 0.651 | 0.035 | 0.906 | 0.674 |
| 975 | **Tpm3iso** | Unchanged | E9Q5J9 | 19 | 122 | 4 | 285 | 33.3 | 4.77 | 45 | 0.984 | 0.966 | 0.926 | 0.837 | 2.462 | 0.122 | 0.376 | 0.096 |
| 976 | **Cct6a** | Unchanged | P80317 | 18 | 347 | 18 | 531 | 58 | 7.08 | 45 | 1.003 | 0.982 | 1.207 | 0.360 | 1.089 | 0.512 | 1.108 | 0.617 |
| 977 | **Asah1** | Unchanged | Q9WV54 | 15 | 116 | 15 | 394 | 44.6 | 8.46 | 45 | 1.434 | 0.491 | 2.741 | 0.123 | 0.923 | 0.830 | 2.968 | 0.108 |
| 978 | **Ctsd** | UP | P18242 | 17 | 380 | 17 | 410 | 44.9 | 7.15 | 45 | 1.189 | 0.299 | 2.215 | 0.001 | 1.903 | 0.000 | 1.164 | 0.309 |
| 979 | **Mlycd** | Unchanged | Q99J39 | 15 | 103 | 15 | 492 | 54.7 | 8.94 | 45 | 0.657 | 0.173 | 1.097 | 0.725 | 1.033 | 0.927 | 1.063 | 0.848 |
| 980 | **Ndufb11** | Down | O09111 | 5 | 69 | 5 | 151 | 17.4 | 5.22 | 45 | 0.790 | 0.052 | 0.651 | 0.143 | 0.624 | 0.013 | 1.044 | 0.906 |
| 981 | **Prep** | Unchanged | Q9QUR6 | 22 | 208 | 22 | 710 | 80.7 | 5.73 | 45 | 1.051 | 0.836 | 1.261 | 0.626 | 1.083 | 0.511 | 1.165 | 0.734 |
| 982 | **Scly** | Unchanged | A0A0R4J069 | 12 | 59 | 12 | 432 | 47.2 | 6.67 | 45 | 0.993 | 0.954 | 1.167 | 0.690 | 1.001 | 0.995 | 1.166 | 0.710 |
| 983 | **Eif3h** | UP | Q91WK2 | 11 | 58 | 11 | 352 | 39.8 | 6.67 | 45 | 0.754 | 0.293 | 1.337 | 0.429 | 1.651 | 0.045 | 0.810 | 0.485 |
| 984 | **Nutf2** | UP | P61971 | 4 | 30 | 4 | 127 | 14.5 | 5.38 | 45 | 1.614 | 0.136 | 3.774 | 0.032 | 1.739 | 0.311 | 2.170 | 0.130 |
| 985 | **Immt iso** | Unchanged | A0A0U1RP81 | 19 | 103 | 2 | 488 | 53.7 | 8.66 | 45 | 8.774 | 0.118 | 2.314 | 0.314 | 4.875 | 0.078 | 0.475 | 0.289 |
| 986 | **Rpl27** | Down | P61358 | 6 | 99 | 6 | 136 | 15.8 | 10.56 | 45 | 0.936 | 0.773 | 0.458 | 0.003 | 1.241 | 0.287 | 0.369 | 0.010 |
| 987 | **Copb1** | Unchanged | Q9JIF7 | 32 | 340 | 32 | 953 | 107 | 6 | 45 | 1.034 | 0.837 | 1.109 | 0.497 | 0.903 | 0.433 | 1.228 | 0.254 |
| 988 | **Acaa1a iso** | Unchanged | H3BJG8 | 4 | 50 | 1 | 114 | 12.2 | 6.55 | 45 | 3.547 | 0.201 | 0.720 | 0.674 | 4.984 | 0.096 | 0.145 | 0.077 |
| 989 | **Atl3** | UP | Q91YH5 | 18 | 143 | 17 | 541 | 60.5 | 6.1 | 45 | 1.410 | 0.062 | 2.230 | 0.014 | 1.493 | 0.038 | 1.494 | 0.110 |
| 990 | **Gmppb** | Unchanged | Q8BTZ7 | 10 | 88 | 10 | 360 | 39.9 | 6.74 | 45 | 0.663 | 0.241 | 1.120 | 0.732 | 1.956 | 0.165 | 0.573 | 0.219 |
| 991 | **Acad9** | UP | Q8JZN5 | 17 | 134 | 17 | 625 | 68.7 | 7.46 | 45 | 1.868 | 0.004 | 2.712 | 0.013 | 1.732 | 0.151 | 1.566 | 0.200 |
| 992 | **Afmid** | Unchanged | Q8K4H1 | 7 | 42 | 7 | 305 | 34.2 | 6.11 | 45 | 1.018 | 0.954 | 1.552 | 0.075 | 1.479 | 0.140 | 1.050 | 0.780 |
| 993 | **Erp44** | Unchanged | Q9D1Q6 | 14 | 171 | 14 | 406 | 46.8 | 5.27 | 45 | 1.648 | 0.425 | 0.629 | 0.339 | 1.851 | 0.252 | 0.340 | 0.078 |
| 994 | **Eif3d** | UP | O70194 | 14 | 162 | 14 | 548 | 63.9 | 6.05 | 45 | 1.701 | 0.015 | 1.944 | 0.039 | 1.161 | 0.460 | 1.675 | 0.066 |
| 995 | **Pdcd6** | Unchanged | P12815 | 6 | 19 | 6 | 191 | 21.9 | 5.4 | 45 | 1.486 | 0.316 | 3.789 | 0.058 | 1.180 | 0.631 | 3.211 | 0.068 |
| 996 | **Dglucy** | Unchanged | Q8BH86 | 16 | 112 | 16 | 617 | 66.3 | 7.09 | 44 | 1.321 | 0.318 | 1.188 | 0.347 | 0.803 | 0.313 | 1.481 | 0.106 |
| 997 | **Etf1** | Down | Q8BWY3 | 13 | 56 | 13 | 437 | 49 | 5.71 | 44 | 1.250 | 0.554 | 0.404 | 0.022 | 0.651 | 0.158 | 0.621 | 0.134 |
| 998 | **Ckmt2** | Unchanged | Q6P8J7 | 11 | 32 | 9 | 419 | 47.4 | 8.4 | 44 | 0.150 | 0.350 | 0.154 | 0.352 | 0.721 | 0.799 | 0.214 | 0.391 |
| 999 | **Dcps** | Unchanged | Q9DAR7 | 10 | 59 | 10 | 338 | 39 | 6.48 | 44 | 0.934 | 0.827 | 0.801 | 0.198 | 0.959 | 0.853 | 0.835 | 0.381 |
| 1000 | **Dnm1l** | Unchanged | Q8K1M6 | 21 | 215 | 21 | 742 | 82.6 | 7.05 | 44 | 1.110 | 0.606 | 1.542 | 0.069 | 1.141 | 0.201 | 1.351 | 0.137 |
| 1001 | **Sars** | Unchanged | P26638 | 17 | 176 | 17 | 512 | 58.4 | 6.3 | 44 | 1.070 | 0.828 | 1.730 | 0.098 | 1.351 | 0.294 | 1.280 | 0.366 |
| 1002 | **Rpl18a** | Unchanged | P62717 | 11 | 99 | 11 | 176 | 20.7 | 10.71 | 44 | 0.709 | 0.396 | 0.644 | 0.291 | 0.569 | 0.194 | 1.131 | 0.521 |
| 1003 | **Mocs2** | Unchanged | Q9Z224 | 2 | 6 | 2 | 88 | 9.7 | 4.61 | 44 | 0.930 | 0.900 | 1.837 | 0.354 | 1.213 | 0.643 | 1.515 | 0.444 |
| 1004 | **Pdzk1** | Unchanged | Q9JIL4 | 22 | 209 | 22 | 519 | 56.5 | 5.43 | 44 | 1.205 | 0.628 | 1.190 | 0.682 | 1.089 | 0.819 | 1.093 | 0.848 |
| 1005 | **Ctnna1** | Unchanged | P26231 | 24 | 263 | 20 | 906 | 100 | 6.23 | 44 | 1.137 | 0.571 | 1.100 | 0.758 | 1.083 | 0.754 | 1.016 | 0.961 |
| 1006 | **Rpl23a** | Unchanged | P62751 | 8 | 79 | 8 | 156 | 17.7 | 10.45 | 44 | 0.380 | 0.057 | 0.508 | 0.177 | 0.619 | 0.243 | 0.820 | 0.690 |
| 1007 | **Apex1** | Unchanged | P28352 | 8 | 99 | 8 | 317 | 35.5 | 7.91 | 44 | 0.941 | 0.737 | 1.240 | 0.383 | 1.062 | 0.650 | 1.168 | 0.534 |
| 1008 | **Gclm** | Unchanged | O09172 | 8 | 95 | 8 | 274 | 30.5 | 5.52 | 44 | 0.561 | 0.471 | 0.622 | 0.553 | 0.543 | 0.445 | 1.146 | 0.749 |
| 1009 | **Mrps36** | Unchanged | Q9CQX8 | 2 | 36 | 2 | 102 | 11.1 | 9.99 | 44 | 1.101 | 0.766 | 2.131 | 0.166 | 1.770 | 0.208 | 1.204 | 0.695 |
| 1010 | **Prkar2a** | Unchanged | Q8K1M3 | 14 | 78 | 14 | 402 | 45.6 | 4.93 | 44 | 0.978 | 0.880 | 1.003 | 0.981 | 1.089 | 0.534 | 0.922 | 0.406 |
| 1011 | **Cops8** | Unchanged | Q8VBV7 | 4 | 29 | 4 | 209 | 23.2 | 5.2 | 44 | 0.973 | 0.929 | 0.997 | 0.995 | 1.195 | 0.522 | 0.834 | 0.605 |
| 1012 | **Arhgdib** | Unchanged | Q61599 | 4 | 55 | 4 | 200 | 22.8 | 5.11 | 44 | 1.133 | 0.839 | 1.283 | 0.621 | 1.572 | 0.236 | 0.816 | 0.616 |
| 1013 | **Sar1a** | Unchanged | Q99JZ4 | 7 | 54 | 2 | 198 | 22.4 | 6.68 | 44 | 1.193 | 0.537 | 1.037 | 0.844 | 0.920 | 0.739 | 1.127 | 0.556 |
| 1014 | **Txnrd1** | Unchanged | Q9JMH6 | 18 | 166 | 17 | 613 | 67 | 7.44 | 44 | 1.733 | 0.168 | 2.283 | 0.154 | 1.707 | 0.077 | 1.338 | 0.522 |

| | | | | | | | | | | | | | | | | | | |
|---|---|---|---|---|---|---|---|---|---|---|---|---|---|---|---|---|---|---|
| 1015 | **Prpf19** | Unchanged | Q99KP6 | 11 | 79 | 11 | 504 | 55.2 | 6.61 | 44 | 1.051 | 0.881 | 1.240 | 0.589 | 1.102 | 0.767 | 1.125 | 0.772 |
| 1016 | **Ndufb6** | Unchanged | Q3UIU2 | 4 | 65 | 4 | 128 | 15.5 | 9.79 | 44 | 1.098 | 0.701 | 2.103 | 0.272 | 1.589 | 0.093 | 1.324 | 0.603 |
| 1017 | **Rac1** | Unchanged | P63001 | 7 | 35 | 5 | 192 | 21.4 | 8.5 | 44 | 0.858 | 0.714 | 1.849 | 0.188 | 2.347 | 0.077 | 0.788 | 0.543 |
| 1018 | **Gsta2** | UP | P10648 | 9 | 619 | 1 | 222 | 25.5 | 8.81 | 44 | 1.508 | 0.344 | 1.949 | 0.287 | 8.910 | 0.022 | 0.219 | 0.041 |
| 1019 | **Eef1e1** | Unchanged | Q9D1M4 | 5 | 9 | 5 | 174 | 19.8 | 8.59 | 44 | 0.515 | 0.412 | 0.907 | 0.891 | 0.489 | 0.356 | 1.856 | 0.356 |
| 1020 | **Chdh** | Unchanged | Q8BJ64 | 18 | 180 | 18 | 596 | 66.4 | 8.51 | 44 | 1.419 | 0.309 | 1.203 | 0.566 | 0.676 | 0.345 | 1.780 | 0.009 |
| 1021 | **Serpina3kiso** | Unchanged | P07759 | 16 | 424 | 1 | 418 | 46.9 | 5.16 | 44 | 0.984 | 0.973 | 0.982 | 0.966 | 2.076 | 0.345 | 0.473 | 0.354 |
| 1022 | **Erp29** | Unchanged | P57759 | 9 | 61 | 9 | 262 | 28.8 | 6.15 | 44 | 0.701 | 0.503 | 1.351 | 0.545 | 0.556 | 0.294 | 2.430 | 0.085 |
| 1023 | **Rmdn2** | Unchanged | Q8BSE0 | 12 | 115 | 12 | 410 | 47 | 7.56 | 43 | 1.459 | 0.020 | 1.957 | 0.054 | 1.513 | 0.252 | 1.293 | 0.458 |
| 1024 | **Ddx1** | Unchanged | Q91VR5 | 25 | 166 | 25 | 740 | 82.4 | 7.21 | 43 | 2.437 | 0.317 | 1.287 | 0.447 | 1.831 | 0.226 | 0.703 | 0.466 |
| 1025 | **Kars** | Unchanged | Q99MN1 | 18 | 129 | 18 | 595 | 67.8 | 5.94 | 43 | 0.837 | 0.400 | 1.201 | 0.246 | 1.095 | 0.593 | 1.097 | 0.569 |
| 1026 | **Rps23** | UP | P62267 | 6 | 61 | 6 | 143 | 15.8 | 10.49 | 43 | 1.294 | 0.288 | 2.235 | 0.039 | 1.664 | 0.195 | 1.343 | 0.409 |
| 1027 | **Rars** | Unchanged | Q9D0I9 | 21 | 200 | 21 | 660 | 75.6 | 7.55 | 43 | 0.834 | 0.587 | 1.140 | 0.752 | 1.078 | 0.760 | 1.058 | 0.894 |
| 1028 | **Iba57** | Unchanged | Q8CAK1 | 7 | 21 | 7 | 358 | 38.4 | 9.01 | 43 | 1.096 | 0.894 | 0.425 | 0.333 | 0.352 | 0.260 | 1.206 | 0.738 |
| 1029 | **Aadat** | Unchanged | Q9WVM8 | 11 | 111 | 11 | 425 | 47.6 | 8.25 | 43 | 0.657 | 0.434 | 0.731 | 0.540 | 0.485 | 0.187 | 1.507 | 0.482 |
| 1030 | **Dctn2** | Unchanged | Q99KJ8 | 10 | 83 | 10 | 402 | 44.1 | 5.26 | 43 | 1.276 | 0.365 | 0.929 | 0.764 | 1.248 | 0.369 | 0.744 | 0.320 |
| 1031 | **Mcfd2iso** | Unchanged | D3YVL4 | 1 | 4 | 1 | 74 | 8.2 | 5.53 | 43 | 3.833 | 0.197 | 6.862 | 0.276 | 0.397 | 0.295 | 17.274 | 0.231 |
| 1032 | **Rpl6** | Down | P47911 | 17 | 249 | 17 | 296 | 33.5 | 10.7 | 43 | 0.504 | 0.010 | 0.976 | 0.945 | 0.657 | 0.097 | 1.487 | 0.359 |
| 1033 | **Hdhd2** | Unchanged | Q3UGR5 | 6 | 49 | 6 | 259 | 28.7 | 6.05 | 43 | 1.038 | 0.878 | 2.326 | 0.229 | 1.775 | 0.109 | 1.310 | 0.616 |
| 1034 | **Rnpep** | Unchanged | Q8VCT3 | 18 | 152 | 18 | 650 | 72.4 | 5.35 | 43 | 0.502 | 0.172 | 0.697 | 0.395 | 0.775 | 0.543 | 0.899 | 0.675 |
| 1035 | **Cbr4** | Unchanged | Q91VT4 | 7 | 91 | 7 | 236 | 25.4 | 9.69 | 43 | 0.912 | 0.748 | 1.002 | 0.996 | 0.573 | 0.087 | 1.749 | 0.049 |
| 1036 | **Myh9** | UP | Q8VDD5 | 79 | 760 | 62 | 1960 | 226.2 | 5.66 | 43 | 2.023 | 0.025 | 1.845 | 0.013 | 3.977 | 0.022 | 0.464 | 0.081 |
| 1037 | **Rabggta** | Unchanged | Q9JHK4 | 14 | 47 | 14 | 567 | 64.9 | 5.8 | 43 | 1.530 | 0.207 | 1.874 | 0.128 | 0.798 | 0.632 | 2.348 | 0.043 |
| 1038 | **Ppp2r1a** | UP | Q76MZ3 | 20 | 292 | 14 | 589 | 65.3 | 5.11 | 43 | 1.485 | 0.205 | 2.601 | 0.041 | 1.792 | 0.186 | 1.451 | 0.354 |
| 1039 | **Ndufa13** | Unchanged | Q9ERS2 | 7 | 48 | 7 | 144 | 16.8 | 9.48 | 43 | 0.590 | 0.129 | 1.596 | 0.163 | 1.290 | 0.653 | 1.238 | 0.661 |
| 1040 | **Uchl3** | UP | Q9JKB1 | 6 | 64 | 6 | 230 | 26.1 | 5.05 | 43 | 1.510 | 0.300 | 3.155 | 0.029 | 2.234 | 0.048 | 1.412 | 0.312 |
| 1041 | **Psmd3** | Unchanged | P14685 | 17 | 140 | 17 | 530 | 60.7 | 8.44 | 43 | 0.855 | 0.685 | 1.266 | 0.482 | 0.458 | 0.109 | 2.763 | 0.006 |
| 1042 | **Manf** | Unchanged | Q3TMX5 | 9 | 60 | 9 | 179 | 20.4 | 8.25 | 43 | 0.592 | 0.116 | 1.754 | 0.348 | 0.610 | 0.129 | 2.873 | 0.159 |
| 1043 | **Capza1** | Unchanged | P47753 | 7 | 53 | 6 | 286 | 32.9 | 5.55 | 43 | 1.128 | 0.704 | 0.895 | 0.685 | 1.317 | 0.102 | 0.679 | 0.120 |
| 1044 | **H2afz** | Unchanged | P0C0S6 | 4 | 53 | 2 | 128 | 13.5 | 10.58 | 43 | 1.250 | 0.739 | 2.846 | 0.223 | 2.309 | 0.163 | 1.233 | 0.736 |
| 1045 | **Ndufs3** | Unchanged | Q9DCT2 | 13 | 132 | 13 | 263 | 30.1 | 7.17 | 43 | 1.167 | 0.392 | 1.241 | 0.442 | 1.139 | 0.496 | 1.090 | 0.770 |
| 1046 | **Aarsd1** | Unchanged | Q3THG9 | 9 | 47 | 9 | 412 | 44.9 | 6.42 | 43 | 1.023 | 0.969 | 0.315 | 0.118 | 0.280 | 0.099 | 1.126 | 0.728 |
| 1047 | **Gmfb** | Unchanged | Q9CQI3 | 4 | 43 | 4 | 142 | 16.7 | 5.16 | 43 | 1.357 | 0.547 | 2.160 | 0.102 | 1.574 | 0.330 | 1.372 | 0.357 |
| 1048 | **Fth1** | Down | P09528 | 8 | 42 | 8 | 182 | 21.1 | 5.88 | 43 | 0.945 | 0.903 | 0.491 | 0.187 | 0.217 | 0.032 | 2.264 | 0.206 |
| 1049 | **Ociad2** | Unchanged | Q9D8W7 | 5 | 48 | 5 | 154 | 16.9 | 9.41 | 43 | 1.254 | 0.770 | 2.212 | 0.239 | 1.155 | 0.805 | 1.915 | 0.271 |
| 1050 | **Akr1c20** | UP | Q8VC77 | 10 | 68 | 5 | 323 | 36.8 | 7.84 | 43 | 3.920 | 0.064 | 2.793 | 0.005 | 2.209 | 0.030 | 1.264 | 0.226 |
| 1051 | **Amt** | Unchanged | Q8CFA2 | 13 | 105 | 13 | 403 | 44 | 8.7 | 43 | 1.206 | 0.348 | 1.250 | 0.570 | 1.034 | 0.874 | 1.210 | 0.620 |
| 1052 | **Phyh** | Unchanged | O35386 | 11 | 131 | 11 | 338 | 38.6 | 7.53 | 43 | 0.708 | 0.047 | 0.705 | 0.240 | 0.677 | 0.043 | 1.041 | 0.915 |
| 1053 | **Tspo** | UP | P50637 | 3 | 21 | 3 | 169 | 18.8 | 9.52 | 27 | 7.155 | 0.068 | 10.117 | 0.010 | 8.682 | 0.008 | 1.165 | 0.691 |
| 1054 | **Rab5c** | Unchanged | P35278 | 6 | 56 | 4 | 216 | 23.4 | 8.41 | 43 | 0.867 | 0.594 | 1.200 | 0.708 | 0.568 | 0.103 | 2.114 | 0.226 |
| 1055 | **Gck** | Unchanged | P52792 | 14 | 132 | 14 | 465 | 52.1 | 5.29 | 43 | 1.133 | 0.148 | 0.953 | 0.793 | 1.099 | 0.671 | 0.867 | 0.598 |

| | | | | | | | | | | | | | | | | | | |
|---|---|---|---|---|---|---|---|---|---|---|---|---|---|---|---|---|---|---|
| 1056 | **Copb2** | Unchanged | O55029 | 25 | 244 | 25 | 905 | 102.4 | 5.3 | 43 | 0.847 | 0.179 | 0.739 | 0.009 | 0.814 | 0.058 | 0.908 | 0.480 |
| 1057 | **Lsm6** | Unchanged | P62313 | 2 | 4 | 2 | 80 | 9.1 | 9.58 | 43 | 0.623 | 0.398 | 0.368 | 0.131 | 0.475 | 0.195 | 0.775 | 0.715 |
| 1058 | **Pdcd6ip** | Unchanged | Q9WU78 | 24 | 237 | 24 | 869 | 96 | 6.52 | 42 | 1.297 | 0.353 | 1.082 | 0.747 | 1.034 | 0.855 | 1.047 | 0.865 |
| 1059 | **Agpat2** | Unchanged | Q8K3K7 | 9 | 62 | 9 | 278 | 31 | 8.88 | 42 | 1.355 | 0.426 | 1.321 | 0.287 | 0.691 | 0.098 | 1.910 | 0.079 |
| 1060 | **Slc25a1** | Unchanged | Q8JZU2 | 10 | 108 | 10 | 311 | 33.9 | 9.89 | 42 | 0.890 | 0.617 | 0.808 | 0.516 | 1.011 | 0.934 | 0.799 | 0.513 |
| 1061 | **Gcsh** | Unchanged | Q91WK5 | 4 | 65 | 4 | 170 | 18.6 | 4.75 | 42 | 1.037 | 0.887 | 1.252 | 0.442 | 0.952 | 0.877 | 1.314 | 0.437 |
| 1062 | **Uqcrb** | Unchanged | Q9D855 | 6 | 108 | 6 | 111 | 13.5 | 9.11 | 42 | 2.185 | 0.122 | 1.116 | 0.824 | 0.316 | 0.065 | 3.532 | 0.093 |
| 1063 | **Ilf2** | Unchanged | Q9CXY6 | 10 | 37 | 10 | 390 | 43 | 5.26 | 42 | 1.253 | 0.509 | 1.227 | 0.637 | 1.366 | 0.274 | 0.898 | 0.754 |
| 1064 | **Ppp6c** | UP | Q9CQR6 | 8 | 50 | 8 | 305 | 35.1 | 5.69 | 38 | 1.584 | 0.318 | 2.814 | 0.027 | 0.860 | 0.769 | 3.274 | 0.010 |
| 1065 | **Npc2** | Unchanged | Q9Z0J0 | 6 | 34 | 6 | 149 | 16.4 | 7.68 | 42 | 0.973 | 0.924 | 1.245 | 0.522 | 1.441 | 0.415 | 0.864 | 0.754 |
| 1066 | **Gnai2** | Unchanged | P08752 | 10 | 65 | 8 | 355 | 40.5 | 5.45 | 42 | 1.306 | 0.296 | 2.217 | 0.143 | 0.873 | 0.640 | 2.541 | 0.096 |
| 1067 | **Lamtor1** | Unchanged | Q9CQ22 | 3 | 6 | 3 | 161 | 17.7 | 5.15 | 42 | 1.555 | 0.566 | 2.439 | 0.200 | 1.918 | 0.077 | 1.272 | 0.617 |
| 1068 | **Cisd2** | Unchanged | Q9CQB5 | 5 | 26 | 5 | 135 | 15.2 | 9.51 | 42 | 1.134 | 0.552 | 0.853 | 0.585 | 1.612 | 0.295 | 0.529 | 0.230 |
| 1069 | **Cyp7b1** | Down | Q60991 | 19 | 175 | 19 | 507 | 58.4 | 9.09 | 42 | 0.513 | 0.061 | 0.503 | 0.089 | 0.173 | 0.003 | 2.906 | 0.083 |
| 1070 | **Eif3j2** | Unchanged | Q66JS6 | 9 | 48 | 9 | 263 | 29.5 | 4.81 | 42 | 0.616 | 0.260 | 0.848 | 0.727 | 0.909 | 0.820 | 0.932 | 0.879 |
| 1071 | **Pcyox1** | Unchanged | Q9CQF9 | 13 | 145 | 13 | 505 | 56.5 | 6.92 | 42 | 0.832 | 0.283 | 0.979 | 0.933 | 0.823 | 0.293 | 1.190 | 0.474 |
| 1072 | **Nampt** | Unchanged | Q99KQ4 | 16 | 119 | 16 | 491 | 55.4 | 7.15 | 42 | 1.266 | 0.327 | 1.417 | 0.309 | 1.430 | 0.192 | 0.991 | 0.980 |
| 1073 | **Rpl15** | Unchanged | Q9CZM2 | 10 | 80 | 10 | 204 | 24.1 | 11.62 | 42 | 1.129 | 0.873 | 2.226 | 0.324 | 0.408 | 0.173 | 5.450 | 0.139 |
| 1074 | **Micos10** | Unchanged | Q7TNS2 | 1 | 3 | 1 | 76 | 8.6 | 8.94 | 42 | 1.380 | 0.379 | 19.745 | 0.125 | 1.794 | 0.142 | 11.005 | 0.139 |
| 1075 | **Rpl5** | Down | P47962 | 14 | 276 | 14 | 297 | 34.4 | 9.77 | 42 | 0.746 | 0.286 | 0.451 | 0.036 | 0.968 | 0.942 | 0.466 | 0.207 |
| 1076 | **Nat8f2** | Unchanged | Q8CHQ9 | 8 | 97 | 7 | 238 | 26.4 | 9.58 | 42 | 0.855 | 0.543 | 0.839 | 0.506 | 0.740 | 0.366 | 1.134 | 0.647 |
| 1077 | **Igkv8-27** | Unchanged | A0A075B5N4 | 3 | 4 | 2 | 100 | 11 | 7.96 | 42 | 0.820 | 0.701 | 4.459 | 0.241 | 0.713 | 0.590 | 6.252 | 0.210 |
| 1078 | **Eno3** | Unchanged | P21550 | 12 | 430 | 6 | 434 | 47 | 7.18 | 42 | 0.177 | 0.363 | 0.213 | 0.384 | 0.152 | 0.348 | 1.405 | 0.467 |
| 1079 | **Pafah1b2** | Unchanged | Q61206 | 5 | 42 | 5 | 229 | 25.6 | 5.92 | 42 | 1.995 | 0.380 | 2.441 | 0.342 | 1.475 | 0.490 | 1.655 | 0.525 |
| 1080 | **Rpl17** | Unchanged | Q9CPR4 | 10 | 189 | 10 | 184 | 21.4 | 10.18 | 42 | 0.977 | 0.956 | 1.075 | 0.858 | 0.777 | 0.520 | 1.384 | 0.419 |
| 1081 | **Cstb** | Unchanged | Q62426 | 3 | 38 | 3 | 98 | 11 | 7.39 | 42 | 0.842 | 0.530 | 1.078 | 0.758 | 1.469 | 0.139 | 0.734 | 0.169 |
| 1082 | **Rps16** | Unchanged | P14131 | 8 | 109 | 8 | 146 | 16.4 | 10.21 | 40 | 1.164 | 0.592 | 1.564 | 0.467 | 1.515 | 0.348 | 1.032 | 0.956 |
| 1083 | **Fam71d** | Unchanged | A0A1W2P6P5 | 1 | 1 | 1 | 91 | 10.4 | 9.79 | 42 | 3.181 | 0.246 | 4.276 | 0.137 | 1.469 | 0.668 | 2.911 | 0.221 |
| 1084 | **Hykk** | Unchanged | Q5U5V2 | 12 | 57 | 12 | 376 | 42.3 | 6.02 | 42 | 1.093 | 0.731 | 0.830 | 0.328 | 1.104 | 0.698 | 0.752 | 0.294 |
| 1085 | **Cyp4f15** | Unchanged | Q8VCA4 | 15 | 50 | 7 | 527 | 60.5 | 6.86 | 42 | 1.558 | 0.515 | 1.603 | 0.467 | 0.628 | 0.425 | 2.552 | 0.228 |
| 1086 | **Mif** | Unchanged | P34884 | 4 | 40 | 4 | 115 | 12.5 | 7.34 | 32 | 0.980 | 0.936 | 1.266 | 0.490 | 1.415 | 0.154 | 0.895 | 0.678 |
| 1087 | **Ndufa3** | Unchanged | Q9CQ91 | 2 | 23 | 2 | 84 | 9.3 | 8.47 | 42 | 1.650 | 0.318 | 1.423 | 0.496 | 1.218 | 0.668 | 1.168 | 0.743 |
| 1088 | **Gng12** | Unchanged | Q9DAS9 | 2 | 35 | 2 | 72 | 8 | 8.97 | 42 | 1.152 | 0.740 | 1.502 | 0.442 | 1.591 | 0.291 | 0.944 | 0.916 |
| 1089 | **Dnajc3** | Unchanged | Q91YW3 | 14 | 150 | 14 | 504 | 57.4 | 5.85 | 42 | 1.464 | 0.383 | 1.589 | 0.121 | 0.757 | 0.158 | 2.099 | 0.027 |
| 1090 | **Hsd17b6** | UP | Q9R092 | 12 | 174 | 4 | 317 | 36.1 | 8.4 | 42 | 1.385 | 0.369 | 3.270 | 0.090 | 3.105 | 0.050 | 1.053 | 0.912 |
| 1091 | **Ddi2** | Unchanged | A2ADY9 | 9 | 147 | 9 | 399 | 44.6 | 5.05 | 42 | 0.787 | 0.364 | 0.759 | 0.306 | 0.760 | 0.286 | 0.999 | 0.994 |
| 1092 | **Ctsb** | Unchanged | P10605 | 14 | 292 | 14 | 339 | 37.3 | 5.91 | 42 | 1.030 | 0.888 | 1.571 | 0.307 | 1.121 | 0.567 | 1.401 | 0.417 |
| 1093 | **Mtch2** | Unchanged | Q791V5 | 10 | 136 | 10 | 303 | 33.5 | 8.25 | 42 | 1.287 | 0.457 | 2.188 | 0.217 | 1.215 | 0.262 | 1.800 | 0.302 |
| 1094 | **Ndufs7** | Unchanged | Q9DC70 | 7 | 72 | 7 | 224 | 24.7 | 9.92 | 42 | 0.628 | 0.295 | 0.923 | 0.782 | 0.671 | 0.237 | 1.377 | 0.181 |
| 1095 | **Nt5c** | Unchanged | Q9JM14 | 4 | 44 | 4 | 200 | 23.1 | 5.49 | 42 | 1.138 | 0.790 | 1.174 | 0.738 | 0.850 | 0.757 | 1.381 | 0.408 |
| 1096 | **Rmdn3** | Unchanged | Q3UJU9 | 17 | 217 | 17 | 470 | 52 | 5.21 | 41 | 0.797 | 0.112 | 1.322 | 0.395 | 0.673 | 0.024 | 1.966 | 0.100 |

| | | | | | | | | | | | | | | | | | | |
|---|---|---|---|---|---|---|---|---|---|---|---|---|---|---|---|---|---|---|
| 1097 | **Hsd3b7** | UP | Q9EQC1 | 9 | 56 | 9 | 369 | 41.1 | 8.05 | 41 | 1.866 | 0.017 | 3.736 | 0.093 | 1.773 | 0.245 | 2.106 | 0.240 |
| 1098 | **Hnrnph2** | Unchanged | P70333 | 11 | 119 | 6 | 449 | 49.2 | 6.3 | 41 | 0.783 | 0.256 | 0.748 | 0.225 | 0.792 | 0.218 | 0.944 | 0.829 |
| 1099 | **Lman1** | Unchanged | Q9D0F3 | 15 | 183 | 14 | 517 | 57.8 | 6.34 | 41 | 1.057 | 0.767 | 0.900 | 0.665 | 1.121 | 0.773 | 0.803 | 0.638 |
| 1100 | **Rpl37a** | Unchanged | P61514 | 3 | 52 | 3 | 92 | 10.3 | 10.43 | 41 | 0.756 | 0.603 | 0.289 | 0.141 | 0.892 | 0.834 | 0.324 | 0.098 |
| 1101 | **Plg** | Unchanged | P20918 | 20 | 115 | 20 | 812 | 90.7 | 6.6 | 41 | 0.805 | 0.192 | 1.077 | 0.674 | 0.923 | 0.680 | 1.166 | 0.421 |
| 1102 | **Glrx3** | UP | Q9CQM9 | 10 | 58 | 10 | 337 | 37.8 | 5.59 | 41 | 19.549 | 0.095 | 2.905 | 0.009 | 2.087 | 0.030 | 1.392 | 0.200 |
| 1103 | **Rps9** | Down | Q6ZWN5 | 12 | 144 | 12 | 194 | 22.6 | 10.65 | 41 | 1.366 | 0.550 | 1.014 | 0.979 | 0.148 | 0.008 | 6.860 | 0.097 |
| 1104 | **Prodh2** | Down | Q8VCZ9 | 12 | 119 | 12 | 456 | 50.7 | 8.19 | 41 | 0.986 | 0.963 | 1.833 | 0.375 | 0.410 | 0.019 | 4.473 | 0.140 |
| 1105 | **Ahsg** | Unchanged | P29699 | 8 | 118 | 8 | 345 | 37.3 | 6.51 | 41 | 0.745 | 0.325 | 1.386 | 0.359 | 0.953 | 0.870 | 1.453 | 0.282 |
| 1106 | **Ndufs4** | Unchanged | Q9CXZ1 | 7 | 83 | 7 | 175 | 19.8 | 9.99 | 41 | 0.720 | 0.493 | 0.853 | 0.713 | 0.811 | 0.629 | 1.052 | 0.806 |
| 1107 | **Atp6v1b2** | Unchanged | P62814 | 15 | 92 | 15 | 511 | 56.5 | 5.81 | 41 | 0.773 | 0.461 | 5.192 | 0.122 | 1.608 | 0.481 | 3.229 | 0.196 |
| 1108 | **Rpl4** | Unchanged | Q9D8E6 | 20 | 311 | 20 | 419 | 47.1 | 11 | 41 | 0.920 | 0.576 | 1.192 | 0.449 | 0.910 | 0.515 | 1.310 | 0.248 |
| 1109 | **Npepl1** | Unchanged | Q6NSR8 | 12 | 82 | 12 | 524 | 55.9 | 6.84 | 41 | 0.754 | 0.327 | 0.985 | 0.954 | 1.054 | 0.826 | 0.934 | 0.633 |
| 1110 | **Mcee** | Unchanged | Q9D1I5 | 4 | 46 | 4 | 178 | 19 | 9.09 | 41 | 0.896 | 0.739 | 1.202 | 0.684 | 0.932 | 0.832 | 1.290 | 0.539 |
| 1111 | **Capn2** | UP | O08529 | 20 | 84 | 20 | 700 | 79.8 | 4.96 | 41 | 0.842 | 0.393 | 0.736 | 0.346 | 1.520 | 0.046 | 0.484 | 0.024 |
| 1112 | **Vps26a** | Unchanged | P40336 | 8 | 53 | 8 | 327 | 38.1 | 6.57 | 41 | 0.768 | 0.341 | 1.008 | 0.978 | 0.929 | 0.824 | 1.085 | 0.830 |
| 1113 | **Acad11** | UP | Q80XL6 | 22 | 172 | 22 | 779 | 87.3 | 8.43 | 41 | 1.022 | 0.940 | 2.346 | 0.043 | 0.797 | 0.500 | 2.945 | 0.025 |
| 1114 | **Gfer** | Unchanged | P56213 | 5 | 48 | 5 | 198 | 22.9 | 7.93 | 41 | 0.857 | 0.126 | 1.082 | 0.690 | 0.863 | 0.346 | 1.253 | 0.351 |
| 1115 | **Igkv12-46** | Unchanged | A0A140T8P6 | 3 | 4 | 3 | 115 | 12.6 | 5.91 | 41 | 0.415 | 0.071 | 0.638 | 0.286 | 0.505 | 0.141 | 1.262 | 0.596 |
| 1116 | **Apoe iso** | Unchanged | G3UWW2 | 2 | 2 | 1 | 71 | 8.3 | 4.03 | 41 | 3.076 | 0.080 | 9.125 | 0.040 | 1.769 | 0.479 | 5.159 | 0.064 |
| 1117 | **Ca5a** | Unchanged | P23589 | 8 | 71 | 8 | 299 | 34.1 | 8 | 41 | 0.806 | 0.512 | 0.478 | 0.095 | 1.127 | 0.809 | 0.424 | 0.199 |
| 1118 | **Rab18** | UP | P35293 | 6 | 65 | 6 | 206 | 23 | 5.36 | 41 | 1.408 | 0.213 | 1.548 | 0.251 | 1.526 | 0.016 | 1.015 | 0.962 |
| 1119 | **Rpl13** | Unchanged | P47963 | 10 | 102 | 10 | 211 | 24.3 | 11.55 | 41 | 1.072 | 0.756 | 0.727 | 0.284 | 0.825 | 0.454 | 0.881 | 0.726 |
| 1120 | **Nedd8** | Unchanged | P29595 | 4 | 41 | 4 | 81 | 9 | 7.25 | 41 | 1.044 | 0.913 | 1.598 | 0.362 | 1.616 | 0.057 | 0.989 | 0.976 |
| 1121 | **Sept2** | Unchanged | P42208 | 9 | 112 | 9 | 361 | 41.5 | 6.55 | 41 | 1.073 | 0.709 | 1.497 | 0.033 | 1.079 | 0.699 | 1.388 | 0.034 |
| 1122 | **Gne** | Unchanged | Q91WG8 | 19 | 116 | 19 | 722 | 79.1 | 6.81 | 41 | 0.751 | 0.187 | 0.680 | 0.109 | 0.634 | 0.064 | 1.072 | 0.782 |
| 1123 | **Mtx2** | Unchanged | O88441 | 5 | 46 | 5 | 263 | 29.7 | 5.63 | 41 | 2.228 | 0.209 | 2.208 | 0.340 | 1.287 | 0.580 | 1.716 | 0.479 |
| 1124 | **IghmAC3 8.205.12** | Unchanged | P06330 | 3 | 48 | 1 | 118 | 12.9 | 7.11 | 41 | 0.466 | 0.055 | 0.909 | 0.831 | 0.814 | 0.539 | 1.116 | 0.816 |
| 1125 | **H2-K1** | Unchanged | P01901 | 11 | 98 | 7 | 369 | 41.3 | 6.39 | 41 | 0.925 | 0.793 | 1.269 | 0.498 | 1.115 | 0.753 | 1.138 | 0.731 |
| 1126 | **Psmb8** | Unchanged | P28063 | 8 | 50 | 8 | 276 | 30.2 | 6.68 | 36 | 0.948 | 0.875 | 1.083 | 0.779 | 1.205 | 0.647 | 0.899 | 0.750 |
| 1127 | **Rps28** | Unchanged | P62858 | 4 | 47 | 4 | 69 | 7.8 | 10.7 | 41 | 0.954 | 0.805 | 1.166 | 0.453 | 0.617 | 0.150 | 1.891 | 0.034 |
| 1128 | **Dusp3** | Unchanged | Q9D7X3 | 5 | 42 | 5 | 185 | 20.5 | 6.54 | 41 | 1.046 | 0.788 | 1.379 | 0.219 | 0.906 | 0.504 | 1.523 | 0.137 |
| 1129 | **Mtco2** | Unchanged | P00405 | 5 | 136 | 5 | 227 | 26 | 4.73 | 41 | 1.568 | 0.412 | 2.922 | 0.293 | 1.703 | 0.503 | 1.716 | 0.547 |
| 1130 | **Ighv4-1** | Unchanged | A0A075B5R5 | 3 | 6 | 3 | 116 | 12.9 | 8.9 | 41 | 0.340 | 0.409 | 1.759 | 0.476 | 0.360 | 0.421 | 4.887 | 0.078 |
| 1131 | **Hsd3b5** | Down | Q61694 | 12 | 119 | 7 | 373 | 41.9 | 9.01 | 36 | 0.742 | 0.337 | 0.390 | 0.002 | 0.353 | 0.004 | 1.106 | 0.724 |
| 1132 | **Rps27l** | Unchanged | Q6ZWY3 | 4 | 51 | 2 | 84 | 9.5 | 9.45 | 40 | 1.494 | 0.318 | 1.698 | 0.312 | 1.136 | 0.736 | 1.494 | 0.440 |
| 1133 | **Rps27** | Unchanged | Q6ZWU9 | 4 | 17 | 2 | 84 | 9.5 | 9.45 | 40 | 0.941 | 0.922 | 0.708 | 0.635 | 0.422 | 0.320 | 1.675 | 0.500 |
| 1134 | **Cbx3** | Unchanged | Q9DCC5 | 7 | 20 | 2 | 183 | 20.8 | 5.33 | 40 | 0.656 | 0.533 | 0.628 | 0.487 | 0.233 | 0.120 | 2.692 | 0.183 |
| 1135 | **Ide** | Unchanged | F6RPJ9 | 25 | 79 | 25 | 987 | 114.2 | 6.27 | 40 | 0.871 | 0.269 | 0.974 | 0.879 | 0.804 | 0.297 | 1.211 | 0.461 |
| 1136 | **Decr2** | Unchanged | Q9WV68 | 10 | 131 | 10 | 292 | 31.3 | 8.81 | 40 | 1.367 | 0.128 | 1.324 | 0.244 | 1.265 | 0.163 | 1.046 | 0.831 |
| 1137 | **Tbcel** | Unchanged | Q8C5W3 | 8 | 49 | 8 | 424 | 48 | 5.53 | 40 | 1.320 | 0.473 | 1.864 | 0.187 | 1.521 | 0.101 | 1.225 | 0.586 |

| | | | | | | | | | | | | | | | | | |
|---|---|---|---|---|---|---|---|---|---|---|---|---|---|---|---|---|---|
| 1138 | **Nucb1** | Unchanged | Q02819 | 16 | 132 | 16 | 459 | 53.4 | 5.07 | 40 | 1.614 | 0.490 | 0.889 | 0.622 | 1.267 | 0.343 | 0.701 | 0.102 |
| 1139 | **Lipa** | Unchanged | Q9Z0M5 | 14 | 174 | 14 | 397 | 45.3 | 8.07 | 40 | 1.349 | 0.366 | 0.874 | 0.323 | 0.736 | 0.042 | 1.187 | 0.260 |
| 1140 | **Eif1a** | Unchanged | Q60872 | 4 | 32 | 3 | 144 | 16.5 | 5.24 | 40 | 0.953 | 0.838 | 0.635 | 0.417 | 1.001 | 0.997 | 0.634 | 0.413 |
| 1141 | **Calml3** | Unchanged | Q9D6P8 | 6 | 129 | 1 | 149 | 16.7 | 4.31 | 40 | 3.140 | 0.383 | 4.039 | 0.067 | 5.064 | 0.062 | 0.798 | 0.675 |
| 1142 | **Ighg2b** | Unchanged | A0A0A6YVP0 | 13 | 83 | 2 | 405 | 44.3 | 6.52 | 40 | 0.481 | 0.163 | 0.827 | 0.694 | 0.749 | 0.546 | 1.104 | 0.825 |
| 1143 | **Hnrnpf** | Unchanged | Q9Z2X1 | 10 | 132 | 9 | 415 | 45.7 | 5.49 | 40 | 1.255 | 0.304 | 1.181 | 0.500 | 1.798 | 0.138 | 0.657 | 0.278 |
| 1144 | **Ccdc47** | UP | Q9D024 | 11 | 78 | 11 | 483 | 55.8 | 4.84 | 40 | 1.548 | 0.082 | 2.088 | 0.002 | 1.815 | 0.101 | 1.151 | 0.565 |
| 1145 | **Copz1** | Unchanged | P61924 | 5 | 82 | 5 | 177 | 20.2 | 4.81 | 40 | 0.885 | 0.542 | 1.062 | 0.765 | 0.910 | 0.662 | 1.168 | 0.336 |
| 1146 | **Gsta4** | UP | P24472 | 11 | 150 | 8 | 222 | 25.5 | 7.39 | 40 | 1.429 | 0.250 | 0.986 | 0.969 | 2.389 | 0.035 | 0.413 | 0.045 |
| 1147 | **Hdgf** | Unchanged | P51859 | 7 | 74 | 7 | 237 | 26.3 | 4.83 | 40 | 1.143 | 0.783 | 2.414 | 0.135 | 1.724 | 0.175 | 1.400 | 0.464 |
| 1148 | **Tmsb4x** | Unchanged | P20065 | 4 | 15 | 4 | 50 | 5.7 | 5.06 | 40 | 0.800 | 0.760 | 2.628 | 0.420 | 1.805 | 0.411 | 1.456 | 0.685 |
| 1149 | **Ndufb2** | Unchanged | Q9CPU2 | 2 | 19 | 2 | 105 | 12 | 6.52 | 40 | 1.193 | 0.576 | 0.742 | 0.348 | 1.124 | 0.690 | 0.660 | 0.247 |
| 1150 | **Ugt3a2** | Unchanged | Q8JZZ0 | 15 | 200 | 12 | 523 | 59.6 | 7.53 | 40 | 0.871 | 0.380 | 0.812 | 0.296 | 0.753 | 0.030 | 1.079 | 0.697 |
| 1151 | **Vat1** | UP | Q62465 | 11 | 74 | 11 | 406 | 43.1 | 6.37 | 40 | 1.072 | 0.803 | 1.301 | 0.210 | 1.722 | 0.002 | 0.755 | 0.090 |
| 1152 | **Rpl11** | Unchanged | Q9CXW4 | 7 | 96 | 7 | 178 | 20.2 | 9.6 | 40 | 0.790 | 0.729 | 0.484 | 0.357 | 0.307 | 0.208 | 1.575 | 0.284 |
| 1153 | **Fdxr** | Down | Q61578 | 11 | 54 | 11 | 494 | 54.2 | 8.66 | 40 | 1.119 | 0.833 | 1.027 | 0.958 | 0.154 | 0.038 | 6.689 | 0.049 |
| 1154 | **Ppt1** | Unchanged | O88531 | 9 | 121 | 9 | 306 | 34.5 | 8 | 40 | 1.485 | 0.043 | 1.717 | 0.147 | 1.407 | 0.043 | 1.221 | 0.512 |
| 1155 | **Hnrnpa3** | Unchanged | Q8BG05 | 15 | 259 | 15 | 379 | 39.6 | 9.01 | 40 | 0.206 | 0.071 | 0.196 | 0.068 | 0.379 | 0.149 | 0.519 | 0.121 |
| 1156 | **Atp2a1** | Down | Q8R429 | 32 | 129 | 23 | 994 | 109.4 | 5.22 | 40 | 0.292 | 0.045 | 0.335 | 0.054 | 0.367 | 0.063 | 0.914 | 0.826 |
| 1157 | **Arl8a** | Unchanged | Q8VEH3 | 6 | 41 | 1 | 186 | 21.4 | 7.77 | 40 | #DIV/0! | #DIV/0! | #DIV/0! | #DIV/0! | #DIV/0! | #DIV/0! | #DIV/0! | #DIV/0! |
| 1158 | **Prrc1** | Unchanged | Q3UPH1 | 10 | 69 | 10 | 443 | 46.3 | 5.95 | 40 | 1.239 | 0.174 | 1.385 | 0.044 | 0.882 | 0.568 | 1.572 | 0.036 |
| 1159 | **C4b** | Unchanged | P01029 | 41 | 341 | 41 | 1738 | 192.8 | 7.53 | 40 | 0.814 | 0.288 | 1.208 | 0.527 | 0.785 | 0.168 | 1.538 | 0.202 |
| 1160 | **Pgrmc2** | Unchanged | Q80UU9 | 6 | 40 | 5 | 217 | 23.3 | 5.15 | 40 | 0.821 | 0.748 | 0.948 | 0.921 | 0.574 | 0.366 | 1.650 | 0.271 |
| 1161 | **Pabpc1** | Unchanged | P29341 | 25 | 302 | 17 | 636 | 70.6 | 9.5 | 40 | 0.890 | 0.355 | 0.929 | 0.657 | 1.068 | 0.517 | 0.870 | 0.319 |
| 1162 | **Rdh19** | Unchanged | G5E8H9 | 13 | 126 | 0 | 318 | 35.9 | 9.01 | 40 | 2.225 | 0.323 | 7.792 | 0.181 | 0.456 | 0.295 | 17.084 | 0.150 |
| 1163 | **Sprr1a** | Unchanged | Q62266 | 3 | 7 | 3 | 144 | 15.8 | 7.85 | 40 | 3.047 | 0.138 | 0.796 | 0.789 | 9.949 | 0.065 | 0.080 | 0.061 |
| 1164 | **Dnaja1** | Unchanged | P63037 | 9 | 117 | 9 | 397 | 44.8 | 7.08 | 40 | 1.123 | 0.603 | 0.752 | 0.239 | 0.540 | 0.068 | 1.394 | 0.227 |
| 1165 | **Acyp1** | Unchanged | P56376 | 3 | 9 | 3 | 99 | 11.2 | 9.04 | 39 | 0.643 | 0.620 | 0.135 | 0.100 | 0.096 | 0.088 | 1.396 | 0.370 |
| 1166 | **Cutc** | UP | Q9D8X1 | 6 | 36 | 6 | 272 | 29 | 7.34 | 39 | 1.862 | 0.062 | 1.726 | 0.258 | 1.837 | 0.039 | 0.939 | 0.871 |
| 1167 | **Urad** | Unchanged | Q283N4 | 6 | 80 | 6 | 178 | 20 | 6.11 | 39 | 0.842 | 0.628 | 1.290 | 0.606 | 0.790 | 0.493 | 1.633 | 0.377 |
| 1168 | **Cluh** | Unchanged | Q5SW19 | 33 | 240 | 33 | 1315 | 148 | 6.02 | 39 | 0.994 | 0.953 | 1.170 | 0.296 | 1.147 | 0.403 | 1.020 | 0.916 |
| 1169 | **Tmed2** | Unchanged | Q9R0Q3 | 6 | 44 | 6 | 201 | 22.7 | 5.17 | 39 | 0.920 | 0.877 | 1.797 | 0.386 | 1.269 | 0.570 | 1.416 | 0.513 |
| 1170 | **Gpam** | Unchanged | Q61586 | 18 | 135 | 17 | 827 | 93.6 | 7.84 | 39 | 0.838 | 0.382 | 0.897 | 0.544 | 0.750 | 0.261 | 1.195 | 0.403 |
| 1171 | **Arpc4** | Unchanged | P59999 | 5 | 58 | 5 | 168 | 19.7 | 8.43 | 39 | 0.524 | 0.420 | 0.205 | 0.167 | 0.538 | 0.402 | 0.381 | 0.001 |
| 1172 | **Anp32a** | Unchanged | O35381 | 10 | 158 | 8 | 247 | 28.5 | 4.07 | 39 | 0.777 | 0.208 | 0.865 | 0.399 | 0.838 | 0.248 | 1.032 | 0.822 |
| 1173 | **Coro1c** | Unchanged | Q9WUM4 | 13 | 92 | 13 | 474 | 53.1 | 7.08 | 39 | 1.321 | 0.492 | 1.811 | 0.064 | 2.067 | 0.068 | 0.876 | 0.615 |
| 1174 | **Eif2s3y** | Down | Q9Z0N2 | 13 | 134 | 2 | 472 | 51.1 | 8.5 | 39 | 0.243 | 0.091 | 0.281 | 0.120 | 0.087 | 0.043 | 3.237 | 0.290 |
| 1175 | **Phyhd1** | Unchanged | Q9DB26 | 8 | 88 | 8 | 291 | 32.5 | 6.34 | 39 | 1.051 | 0.841 | 1.248 | 0.399 | 1.588 | 0.265 | 0.786 | 0.514 |
| 1176 | **Tomm70** | Unchanged | Q9CZW5 | 20 | 123 | 20 | 611 | 67.5 | 7.53 | 39 | 1.276 | 0.301 | 1.441 | 0.215 | 0.752 | 0.298 | 1.915 | 0.035 |
| 1177 | **Lman2** | Unchanged | Q9DBH5 | 11 | 113 | 11 | 358 | 40.4 | 6.95 | 39 | 0.800 | 0.458 | 0.733 | 0.366 | 0.752 | 0.342 | 0.974 | 0.916 |
| 1178 | **Tpd52l2** | Unchanged | Q9CYZ2 | 7 | 46 | 7 | 220 | 24 | 6.15 | 39 | 1.106 | 0.526 | 1.194 | 0.542 | 1.388 | 0.306 | 0.860 | 0.671 |

| | | | | | | | | | | | | | | | | | | |
|---|---|---|---|---|---|---|---|---|---|---|---|---|---|---|---|---|---|---|
| 1179 | **Letm1** | Unchanged | Q9Z2I0 | 18 | 146 | 18 | 738 | 82.9 | 6.52 | 39 | 0.480 | 0.154 | 0.527 | 0.181 | 0.623 | 0.293 | 0.845 | 0.359 |
| 1180 | **Iars** | Unchanged | Q8BU30 | 32 | 162 | 32 | 1262 | 144.2 | 6.55 | 38 | 1.081 | 0.842 | 0.943 | 0.890 | 0.849 | 0.643 | 1.111 | 0.795 |
| 1181 | **Blvra** | UP | Q9CY64 | 8 | 59 | 8 | 295 | 33.5 | 7.02 | 39 | 1.574 | 0.228 | 2.031 | 0.075 | 1.752 | 0.034 | 1.159 | 0.628 |
| 1182 | **Coq9** | Unchanged | Q8K1Z0 | 7 | 112 | 7 | 313 | 35.1 | 5.92 | 39 | 0.605 | 0.356 | 0.249 | 0.065 | 1.217 | 0.822 | 0.205 | 0.294 |
| 1183 | **Myl9** | Unchanged | Q9CQ19 | 5 | 46 | 2 | 172 | 19.8 | 4.92 | 39 | 0.865 | 0.798 | 1.250 | 0.686 | 1.707 | 0.479 | 0.732 | 0.634 |
| 1184 | **Pmpca** | Unchanged | Q9DC61 | 15 | 116 | 15 | 524 | 58.2 | 6.83 | 39 | 0.893 | 0.721 | 1.340 | 0.271 | 1.638 | 0.071 | 0.819 | 0.276 |
| 1185 | **Ndufa6** | Unchanged | Q9CQZ5 | 4 | 56 | 4 | 131 | 15.3 | 10.11 | 39 | 1.156 | 0.552 | 1.908 | 0.142 | 0.605 | 0.145 | 3.154 | 0.040 |
| 1186 | **Vamp3** | Down | P63024 | 3 | 16 | 3 | 103 | 11.5 | 8.5 | 39 | 0.644 | 0.069 | 0.468 | 0.030 | 0.305 | 0.000 | 1.537 | 0.452 |
| 1187 | **Eif2s3x** | Unchanged | Q9Z0N1 | 13 | 140 | 2 | 472 | 51 | 8.4 | 39 | 1.206 | 0.279 | 1.451 | 0.342 | 1.472 | 0.083 | 0.985 | 0.966 |
| 1188 | **Sec31aiso** | Unchanged | Q3UPL0 | 36 | 390 | 25 | 1230 | 133.5 | 6.76 | 39 | 1.261 | 0.117 | 1.720 | 0.001 | 1.699 | 0.001 | 1.013 | 0.908 |
| 1189 | **Drg1** | Unchanged | P32233 | 10 | 55 | 9 | 367 | 40.5 | 8.9 | 39 | 1.924 | 0.068 | 1.677 | 0.307 | 1.726 | 0.230 | 0.972 | 0.952 |
| 1190 | **Chordc1** | Unchanged | Q9D1P4 | 7 | 37 | 7 | 331 | 37.3 | 7.9 | 39 | 1.151 | 0.638 | 1.542 | 0.276 | 0.911 | 0.832 | 1.692 | 0.262 |
| 1191 | **Eef1a2** | Unchanged | P62631 | 14 | 721 | 3 | 463 | 50.4 | 9.03 | 39 | 1.056 | 0.917 | 0.548 | 0.248 | 0.741 | 0.573 | 0.740 | 0.626 |
| 1192 | **Gsdmdc1** | Unchanged | Q9D8T2 | 11 | 60 | 11 | 487 | 53.2 | 5.1 | 39 | 0.735 | 0.466 | 1.002 | 0.997 | 0.501 | 0.086 | 2.002 | 0.281 |
| 1193 | **Ptges2** | Unchanged | Q8BWM0 | 9 | 38 | 9 | 384 | 43.3 | 9 | 39 | 1.058 | 0.730 | 1.094 | 0.823 | 1.025 | 0.853 | 1.067 | 0.863 |
| 1194 | **Cyp4a10** | Unchanged | O88833 | 13 | 59 | 7 | 509 | 58.3 | 8.75 | 39 | 1.655 | 0.166 | 1.228 | 0.563 | 1.719 | 0.133 | 0.714 | 0.257 |
| 1195 | **Prkacb** | Unchanged | P68181 | 8 | 36 | 3 | 351 | 40.7 | 8.56 | 38 | 0.569 | 0.307 | 1.625 | 0.374 | 2.224 | 0.244 | 0.730 | 0.588 |
| 1196 | **Acot3** | UP | Q9QYR7 | 10 | 52 | 5 | 432 | 47.5 | 7.46 | 38 | 3.436 | 0.005 | 15.972 | 0.063 | 4.223 | 0.000 | 3.782 | 0.128 |
| 1197 | **Tmed10** | Unchanged | Q9D1D4 | 8 | 46 | 8 | 219 | 24.9 | 6.7 | 38 | 1.124 | 0.801 | 0.859 | 0.595 | 1.067 | 0.775 | 0.805 | 0.337 |
| 1198 | **Ppif** | Down | Q99KR7 | 7 | 43 | 6 | 206 | 21.7 | 9.16 | 38 | 0.374 | 0.005 | 0.358 | 0.010 | 1.929 | 0.331 | 0.185 | 0.116 |
| 1199 | **Nudt21** | Unchanged | Q9CQF3 | 5 | 30 | 5 | 227 | 26.2 | 8.82 | 38 | 0.418 | 0.514 | 0.168 | 0.332 | 0.074 | 0.283 | 2.270 | 0.167 |
| 1200 | **Ckb** | UP | Q04447 | 7 | 73 | 7 | 381 | 42.7 | 5.67 | 38 | 1.181 | 0.713 | 1.756 | 0.157 | 2.178 | 0.006 | 0.806 | 0.423 |
| 1201 | **1600014C10Rik** | Unchanged | Q8WUR0 | 3 | 10 | 3 | 141 | 15.1 | 7.18 | 32 | 1.608 | 0.314 | 2.298 | 0.095 | 1.831 | 0.253 | 1.255 | 0.621 |
| 1202 | **Cars** | UP | Q9ER72 | 25 | 202 | 25 | 831 | 94.8 | 6.76 | 38 | 1.102 | 0.779 | 1.270 | 0.495 | 1.762 | 0.050 | 0.721 | 0.159 |
| 1203 | **Anxa1** | UP | P10107 | 10 | 75 | 10 | 346 | 38.7 | 7.37 | 38 | 1.787 | 0.162 | 2.506 | 0.022 | 5.559 | 0.007 | 0.451 | 0.053 |
| 1204 | **Ptpa** | Unchanged | P58389 | 7 | 42 | 7 | 323 | 36.7 | 6.39 | 38 | 1.126 | 0.687 | 4.069 | 0.209 | 0.959 | 0.851 | 4.245 | 0.202 |
| 1205 | **Pfkfb1** | Unchanged | P70266 | 13 | 80 | 13 | 471 | 54.8 | 6.76 | 38 | 1.051 | 0.809 | 1.234 | 0.434 | 1.198 | 0.261 | 1.030 | 0.881 |
| 1206 | **Srp72** | Unchanged | F8VQC1 | 14 | 57 | 14 | 671 | 74.6 | 9.23 | 38 | 1.123 | 0.257 | 0.960 | 0.864 | 0.755 | 0.091 | 1.271 | 0.423 |
| 1207 | **Rab10** | Unchanged | P61027 | 7 | 66 | 3 | 200 | 22.5 | 8.38 | 38 | 0.431 | 0.172 | 2.114 | 0.234 | 0.615 | 0.346 | 3.436 | 0.096 |
| 1208 | **Galt** | UP | Q03249 | 7 | 33 | 7 | 379 | 43.2 | 6.77 | 38 | 2.322 | 0.029 | 4.753 | 0.002 | 1.476 | 0.167 | 3.220 | 0.004 |
| 1209 | **Bzw1** | Down | Q9CQC6 | 11 | 66 | 11 | 419 | 48 | 5.92 | 38 | 0.628 | 0.049 | 0.648 | 0.151 | 0.256 | 0.000 | 2.527 | 0.115 |
| 1210 | **Aspg** | Unchanged | A0JNU3 | 14 | 104 | 14 | 564 | 60.6 | 5.88 | 38 | 1.526 | 0.396 | 0.730 | 0.249 | 1.033 | 0.870 | 0.707 | 0.248 |
| 1211 | **Eif3l** | UP | Q8QZY1 | 16 | 155 | 16 | 564 | 66.6 | 6.44 | 38 | 4.009 | 0.052 | 3.333 | 0.139 | 7.042 | 0.005 | 0.473 | 0.118 |
| 1212 | **Egfr** | Unchanged | Q01279 | 31 | 231 | 31 | 1210 | 134.8 | 6.86 | 38 | 0.781 | 0.475 | 0.882 | 0.732 | 0.687 | 0.303 | 1.283 | 0.341 |
| 1213 | **Nnmt** | Unchanged | O55239 | 5 | 27 | 5 | 264 | 29.6 | 5.39 | 38 | 1.389 | 0.589 | 1.719 | 0.425 | 1.384 | 0.548 | 1.242 | 0.720 |
| 1214 | **Psmd9** | Unchanged | Q9CR00 | 7 | 37 | 7 | 222 | 24.7 | 6.43 | 38 | 0.776 | 0.646 | 0.340 | 0.111 | 0.220 | 0.068 | 1.549 | 0.110 |
| 1215 | **Cops7a** | Unchanged | Q9CZ04 | 5 | 50 | 5 | 275 | 30.2 | 7.87 | 38 | 1.122 | 0.746 | 1.721 | 0.050 | 1.537 | 0.177 | 1.120 | 0.566 |
| 1216 | **Qars** | Unchanged | Q8BML9 | 22 | 73 | 22 | 775 | 87.6 | 7.31 | 38 | 0.899 | 0.722 | 2.052 | 0.431 | 1.093 | 0.764 | 1.878 | 0.465 |
| 1217 | **Cmah** | Unchanged | Q61419 | 14 | 69 | 14 | 577 | 66.9 | 6.38 | 38 | 1.060 | 0.856 | 0.855 | 0.714 | 0.609 | 0.177 | 1.404 | 0.454 |
| 1218 | **Sqor** | Unchanged | Q9R112 | 13 | 77 | 13 | 450 | 50.3 | 9.09 | 38 | 0.638 | 0.395 | 1.181 | 0.773 | 0.577 | 0.325 | 2.045 | 0.235 |
| 1219 | **Rpl18** | Down | P35980 | 7 | 145 | 7 | 188 | 21.6 | 11.78 | 38 | 0.954 | 0.837 | 0.584 | 0.082 | 0.630 | 0.030 | 0.927 | 0.788 |

| | | | | | | | | | | | | | | | | | |
|---|---|---|---|---|---|---|---|---|---|---|---|---|---|---|---|---|---|
| 1220 | **Ndufs5** | Unchanged | Q99LY9 | 4 | 31 | 4 | 106 | 12.6 | 8.92 | 38 | 1.560 | 0.231 | 1.932 | 0.090 | 0.882 | 0.718 | 2.191 | 0.035 |
| 1221 | **Rap2c** | Unchanged | Q8BU31 | 4 | 23 | 2 | 183 | 20.7 | 4.94 | 38 | 0.558 | 0.270 | 1.262 | 0.658 | 0.549 | 0.270 | 2.300 | 0.164 |
| 1222 | **Lypla2** | Unchanged | Q9WTL7 | 4 | 28 | 4 | 231 | 24.8 | 7.23 | 38 | 2.292 | 0.135 | 2.987 | 0.144 | 1.996 | 0.105 | 1.496 | 0.411 |
| 1223 | **Akr1b8** | UP | P45377 | 7 | 64 | 4 | 316 | 36.1 | 6.37 | 38 | 1.604 | 0.285 | 5.310 | 0.138 | 2.253 | 0.044 | 2.357 | 0.282 |
| 1224 | **Rps25** | Unchanged | P62852 | 7 | 101 | 7 | 125 | 13.7 | 10.11 | 38 | 1.130 | 0.767 | 0.950 | 0.905 | 1.009 | 0.980 | 0.942 | 0.831 |
| 1225 | **Ap1m1** | UP | P35585 | 10 | 47 | 10 | 423 | 48.5 | 7.3 | 38 | 1.031 | 0.906 | 7.952 | 0.047 | 0.815 | 0.535 | 9.760 | 0.043 |
| 1226 | **Gnai3** | Unchanged | Q9DC51 | 8 | 30 | 6 | 354 | 40.5 | 5.69 | 38 | 0.767 | 0.633 | 0.581 | 0.274 | 0.264 | 0.076 | 2.200 | 0.024 |
| 1227 | **Tmem205** | Unchanged | Q91XE8 | 5 | 63 | 5 | 189 | 21.2 | 9.42 | 38 | 0.690 | 0.379 | 0.505 | 0.158 | 0.375 | 0.073 | 1.347 | 0.337 |
| 1228 | **Plin3** | Unchanged | Q9DBG5 | 9 | 38 | 9 | 437 | 47.2 | 5.62 | 38 | 1.503 | 0.298 | 1.881 | 0.078 | 0.607 | 0.336 | 3.096 | 0.003 |
| 1229 | **Rpl21** | Down | O09167 | 8 | 106 | 8 | 160 | 18.6 | 10.49 | 38 | 0.677 | 0.107 | 0.488 | 0.021 | 0.305 | 0.002 | 1.599 | 0.097 |
| 1230 | **Uqcr10** | Unchanged | Q8R1I1 | 2 | 87 | 2 | 64 | 7.4 | 9.19 | 38 | 0.925 | 0.743 | 1.360 | 0.402 | 0.817 | 0.553 | 1.665 | 0.280 |
| 1231 | **Pygb** | UP | Q8CI94 | 24 | 174 | 17 | 843 | 96.7 | 6.73 | 37 | 1.539 | 0.329 | 2.783 | 0.007 | 2.947 | 0.002 | 0.944 | 0.788 |
| 1232 | **Aldoart1** | Unchanged | A6ZI46 | 11 | 77 | 2 | 419 | 45.3 | 7.05 | 37 | #DIV/0! | #DIV/0! | #DIV/0! | #DIV/0! | #DIV/0! | #DIV/0! | #DIV/0! | #DIV/0! |
| 1233 | **Mrrf** | Down | Q9D6S7 | 6 | 27 | 6 | 262 | 29 | 9.85 | 37 | 0.657 | 0.014 | 0.757 | 0.485 | 0.535 | 0.015 | 1.414 | 0.531 |
| 1234 | **Txn2** | Unchanged | P97493 | 5 | 76 | 5 | 166 | 18.2 | 7.9 | 37 | 0.818 | 0.714 | 0.912 | 0.850 | 0.729 | 0.552 | 1.251 | 0.497 |
| 1235 | **Cp** | Unchanged | Q61147 | 25 | 221 | 25 | 1061 | 121.1 | 5.85 | 37 | 0.849 | 0.664 | 1.544 | 0.412 | 1.019 | 0.948 | 1.515 | 0.442 |
| 1236 | **Sacm1l** | Unchanged | Q9EP69 | 14 | 78 | 14 | 587 | 66.9 | 7.3 | 37 | 0.859 | 0.492 | 1.038 | 0.888 | 0.964 | 0.854 | 1.077 | 0.764 |
| 1237 | **Hist1h2bf** | Unchanged | P10853 | 6 | 98 | 6 | 126 | 13.9 | 10.32 | 37 | 1.241 | 0.688 | 3.004 | 0.343 | 5.177 | 0.230 | 0.580 | 0.579 |
| 1238 | **Ddx5** | Unchanged | Q61656 | 19 | 107 | 14 | 614 | 69.2 | 8.92 | 37 | 1.063 | 0.731 | 0.700 | 0.118 | 0.950 | 0.809 | 0.737 | 0.310 |
| 1239 | **Ugt2b35** | Unchanged | Q8BJL9 | 12 | 85 | 10 | 529 | 60.4 | 8.1 | 37 | 0.729 | 0.116 | 0.935 | 0.763 | 1.066 | 0.696 | 0.877 | 0.469 |
| 1240 | **Ddx39a** | Unchanged | Q8VDW0 | 12 | 80 | 5 | 427 | 49 | 5.68 | 37 | 1.339 | 0.461 | 1.735 | 0.097 | 1.636 | 0.188 | 1.060 | 0.852 |
| 1241 | **Cpped1** | Unchanged | Q8BFS6 | 6 | 29 | 6 | 312 | 35.2 | 5.34 | 37 | 0.912 | 0.781 | 0.686 | 0.100 | 0.821 | 0.557 | 0.836 | 0.654 |
| 1242 | **Dnaja2** | Unchanged | Q9QYJ0 | 9 | 98 | 9 | 412 | 45.7 | 6.48 | 37 | 0.832 | 0.511 | 0.881 | 0.658 | 0.641 | 0.258 | 1.375 | 0.346 |
| 1243 | **Hnrnpc iso** | Unchanged | Q9Z204 | 10 | 100 | 5 | 313 | 34.4 | 5.05 | 37 | 1.449 | 0.312 | 1.538 | 0.127 | 1.411 | 0.083 | 1.090 | 0.734 |
| 1244 | **Prkaca** | Unchanged | P05132 | 9 | 41 | 4 | 351 | 40.5 | 8.79 | 37 | 0.810 | 0.637 | 1.074 | 0.905 | 1.708 | 0.133 | 0.629 | 0.349 |
| 1245 | **Ube2g2** | Unchanged | P60605 | 3 | 3 | 3 | 165 | 18.6 | 4.7 | 37 | 2.339 | 0.077 | 2.182 | 0.203 | 0.501 | 0.253 | 4.353 | 0.061 |
| 1246 | **Dpp7** | Unchanged | Q9ET22 | 10 | 87 | 10 | 506 | 56.2 | 5.39 | 34 | 0.738 | 0.544 | 0.926 | 0.847 | 0.364 | 0.126 | 2.540 | 0.000 |
| 1247 | **Gmps** | Unchanged | Q3THK7 | 15 | 114 | 15 | 693 | 76.7 | 6.73 | 37 | 1.225 | 0.622 | 1.446 | 0.356 | 1.107 | 0.812 | 1.306 | 0.510 |
| 1248 | **Cml1** | Unchanged | Q9JIZ0 | 6 | 28 | 5 | 222 | 25 | 9.55 | 37 | 1.340 | 0.557 | 2.511 | 0.110 | 1.972 | 0.230 | 1.273 | 0.643 |
| 1249 | **Hp** | Down | Q61646 | 15 | 192 | 15 | 347 | 38.7 | 6.29 | 37 | 0.421 | 0.026 | 0.555 | 0.115 | 0.254 | 0.006 | 2.183 | 0.100 |
| 1250 | **Timm13** | Unchanged | P62075 | 3 | 36 | 3 | 95 | 10.5 | 8.18 | 37 | 1.194 | 0.609 | 0.950 | 0.920 | 1.211 | 0.577 | 0.785 | 0.583 |
| 1251 | **Rpe** | Unchanged | Q8VEE0 | 4 | 32 | 3 | 228 | 24.9 | 5.41 | 37 | 1.017 | 0.971 | 0.918 | 0.874 | 2.188 | 0.227 | 0.420 | 0.206 |
| 1252 | **Mogs** | Unchanged | Q80UM7 | 17 | 145 | 17 | 834 | 91.8 | 9 | 37 | 1.009 | 0.969 | 1.497 | 0.089 | 0.976 | 0.917 | 1.534 | 0.091 |
| 1253 | **Adsl** | UP | P54822 | 12 | 96 | 12 | 484 | 54.8 | 7.27 | 37 | 1.328 | 0.248 | 1.721 | 0.034 | 1.625 | 0.036 | 1.059 | 0.723 |
| 1254 | **Ostf1** | Unchanged | Q62422 | 5 | 52 | 5 | 215 | 23.8 | 5.68 | 37 | 1.443 | 0.336 | 1.301 | 0.514 | 1.580 | 0.318 | 0.823 | 0.588 |
| 1255 | **Eif2b1** | Unchanged | Q99LC8 | 7 | 28 | 7 | 305 | 33.8 | 8.32 | 37 | 1.616 | 0.228 | 2.237 | 0.092 | 1.073 | 0.858 | 2.086 | 0.148 |
| 1256 | **Cdv3** | Down | Q4VAA2 | 4 | 30 | 4 | 281 | 29.7 | 6.1 | 37 | 0.179 | 0.021 | 0.126 | 0.016 | 0.452 | 0.116 | 0.278 | 0.045 |
| 1257 | **Npm1** | Unchanged | Q61937 | 9 | 143 | 9 | 292 | 32.5 | 4.77 | 37 | 1.206 | 0.283 | 5.328 | 0.220 | 1.507 | 0.247 | 3.535 | 0.277 |
| 1258 | **Txndc17** | Unchanged | Q9CQM5 | 4 | 34 | 4 | 123 | 14 | 4.77 | 37 | 0.787 | 0.719 | 1.268 | 0.690 | 0.759 | 0.678 | 1.671 | 0.387 |
| 1259 | **Erh** | Down | P84089 | 4 | 50 | 4 | 104 | 12.3 | 5.92 | 37 | 0.973 | 0.885 | 0.471 | 0.038 | 0.620 | 0.118 | 0.760 | 0.488 |
| 1260 | **Ndufv3** | Down | Q8BK30 | 2 | 26 | 1 | 104 | 11.8 | 9.35 | 37 | 0.184 | 0.021 | 0.119 | 0.008 | 0.044 | 0.004 | 2.688 | 0.407 |

| | | | | | | | | | | | | | | | | | | |
|---|---|---|---|---|---|---|---|---|---|---|---|---|---|---|---|---|---|---|
| 1261 | **Rps26** | Down | P62855 | 3 | 54 | 3 | 115 | 13 | 11 | 37 | 0.800 | 0.445 | 0.400 | 0.018 | 0.809 | 0.561 | 0.495 | 0.132 |
| 1262 | **Lactb** | UP | Q9EP89 | 12 | 57 | 12 | 551 | 60.7 | 8.9 | 36 | 1.763 | 0.018 | 1.327 | 0.210 | 1.411 | 0.332 | 0.940 | 0.826 |
| 1263 | **Kpna4** | UP | O35343 | 10 | 100 | 6 | 521 | 57.9 | 4.94 | 36 | 1.293 | 0.204 | 2.535 | 0.032 | 1.693 | 0.106 | 1.497 | 0.231 |
| 1264 | **Atp1a1** | Unchanged | Q8VDN2 | 31 | 482 | 25 | 1023 | 112.9 | 5.45 | 36 | 0.928 | 0.434 | 1.708 | 0.183 | 0.853 | 0.098 | 2.002 | 0.120 |
| 1265 | **Tnnt1** | Unchanged | E9PVN9 | 1 | 1 | 1 | 118 | 14.1 | 4.83 | 36 | 1.070 | 0.942 | 0.104 | 0.280 | 0.420 | 0.490 | 0.247 | 0.174 |
| 1266 | **Endog** | Unchanged | O08600 | 7 | 52 | 7 | 294 | 32.2 | 9.54 | 36 | 0.458 | 0.249 | 0.798 | 0.694 | 0.297 | 0.141 | 2.688 | 0.082 |
| 1267 | **Lonp2** | Unchanged | Q9DBN5 | 22 | 140 | 22 | 852 | 94.5 | 7.23 | 36 | 1.076 | 0.792 | 1.732 | 0.086 | 0.628 | 0.075 | 2.758 | 0.010 |
| 1268 | **Cyp51a1** | Down | Q8K0C4 | 13 | 90 | 13 | 503 | 56.7 | 8.41 | 36 | 0.579 | 0.024 | 1.116 | 0.721 | 0.980 | 0.913 | 1.139 | 0.681 |
| 1269 | **Dpy30** | Unchanged | Q99LT0 | 2 | 22 | 2 | 99 | 11.2 | 4.88 | 36 | 2.141 | 0.149 | 7.849 | 0.133 | 7.476 | 0.117 | 1.050 | 0.948 |
| 1270 | **Sec24a** | Unchanged | Q3U2P1 | 26 | 170 | 1 | 1090 | 118.7 | 7.83 | 36 | 1.113 | 0.761 | 0.488 | 0.076 | 1.115 | 0.747 | 0.438 | 0.117 |
| 1271 | **Tpp2** | Unchanged | Q64514 | 32 | 235 | 32 | 1262 | 139.8 | 6.58 | 36 | 1.024 | 0.888 | 1.061 | 0.789 | 0.868 | 0.502 | 1.222 | 0.323 |
| 1272 | **Kpna3** | Unchanged | O35344 | 11 | 101 | 7 | 521 | 57.7 | 4.94 | 36 | 1.167 | 0.536 | 1.784 | 0.076 | 1.015 | 0.968 | 1.758 | 0.144 |
| 1273 | **Sec24aiso** | Unchanged | A2AA71 | 26 | 96 | 1 | 1089 | 118.6 | 7.83 | 36 | 1.370 | 0.057 | 1.506 | 0.087 | 1.255 | 0.233 | 1.200 | 0.417 |
| 1274 | **Slco1a1** | Unchanged | Q9QXZ6 | 18 | 124 | 18 | 670 | 74.3 | 8.22 | 36 | 1.698 | 0.514 | 0.579 | 0.053 | 0.961 | 0.893 | 0.603 | 0.162 |
| 1275 | **Iars2** | Unchanged | Q8BIJ6 | 22 | 179 | 22 | 1012 | 112.7 | 6.81 | 36 | 1.528 | 0.309 | 1.563 | 0.062 | 0.976 | 0.915 | 1.601 | 0.053 |
| 1276 | **Mrpl10** | Unchanged | Q3TBW2 | 5 | 30 | 5 | 262 | 29.4 | 9.69 | 36 | 2.431 | 0.322 | 4.770 | 0.135 | 2.811 | 0.122 | 1.697 | 0.453 |
| 1277 | **Uso1** | Unchanged | Q9Z1Z0 | 26 | 196 | 26 | 959 | 106.9 | 4.93 | 36 | 0.737 | 0.016 | 1.859 | 0.060 | 1.076 | 0.764 | 1.727 | 0.118 |
| 1278 | **Iyd** | Unchanged | Q9DCX8 | 8 | 104 | 8 | 285 | 32.8 | 6.37 | 36 | 0.346 | 0.251 | 0.398 | 0.285 | 0.666 | 0.644 | 0.598 | 0.592 |
| 1279 | **Snx5** | Down | Q9D8U8 | 9 | 46 | 9 | 404 | 46.8 | 6.62 | 36 | 1.151 | 0.496 | 1.689 | 0.047 | 0.606 | 0.009 | 2.788 | 0.006 |
| 1280 | **Shpk** | Unchanged | Q9D5J6 | 11 | 109 | 11 | 476 | 51.3 | 6.24 | 36 | 0.485 | 0.096 | 0.962 | 0.925 | 0.924 | 0.826 | 1.041 | 0.918 |
| 1281 | **Plpbp** | Unchanged | Q9Z2Y8 | 6 | 96 | 6 | 274 | 30 | 8.27 | 36 | 0.898 | 0.438 | 1.010 | 0.935 | 0.956 | 0.759 | 1.057 | 0.583 |
| 1282 | **Ce9** | Unchanged | P01644 | 2 | 21 | 2 | 108 | 11.9 | 7.97 | 36 | 0.711 | 0.133 | 1.950 | 0.091 | 1.110 | 0.718 | 1.757 | 0.172 |
| 1283 | **Ube2d3** | Unchanged | P61079 | 3 | 41 | 3 | 147 | 16.7 | 7.8 | 36 | 1.415 | 0.527 | 1.447 | 0.556 | 1.112 | 0.882 | 1.301 | 0.714 |
| 1284 | **Acbd5** | Unchanged | Q5XG73 | 14 | 88 | 14 | 508 | 56.6 | 5.41 | 36 | 1.157 | 0.673 | 1.836 | 0.192 | 0.954 | 0.905 | 1.925 | 0.158 |
| 1285 | **Nap1l4** | Unchanged | Q78ZA7 | 7 | 58 | 7 | 375 | 42.7 | 4.67 | 36 | 0.852 | 0.454 | 2.290 | 0.151 | 1.350 | 0.316 | 1.696 | 0.306 |
| 1286 | **Mgll** | UP | O35678 | 9 | 68 | 9 | 303 | 33.4 | 7.15 | 36 | 1.923 | 0.105 | 3.581 | 0.001 | 2.160 | 0.040 | 1.658 | 0.042 |
| 1287 | **Ddb1** | Down | Q3U1J4 | 30 | 154 | 30 | 1140 | 126.8 | 5.26 | 36 | 0.646 | 0.013 | 0.868 | 0.393 | 1.087 | 0.381 | 0.799 | 0.170 |
| 1288 | **Cops4** | UP | O88544 | 11 | 84 | 11 | 406 | 46.3 | 5.83 | 36 | 0.913 | 0.466 | 2.021 | 0.080 | 2.018 | 0.021 | 1.002 | 0.996 |
| 1289 | **Htatip2** | Unchanged | Q9Z2G9 | 6 | 25 | 6 | 242 | 26.9 | 8.59 | 36 | 1.627 | 0.601 | 8.554 | 0.197 | 0.803 | 0.688 | 10.654 | 0.185 |
| 1290 | **Nsfl1c** | Unchanged | Q9CZ44 | 9 | 83 | 9 | 370 | 40.7 | 5.15 | 36 | 0.626 | 0.108 | 0.617 | 0.166 | 0.906 | 0.670 | 0.682 | 0.112 |
| 1291 | **Noct** | Unchanged | A0A0A6YY01 | 1 | 1 | 1 | 117 | 13.3 | 5.49 | 36 | 1.863 | 0.323 | 2.136 | 0.293 | 1.682 | 0.117 | 1.270 | 0.662 |
| 1292 | **Acot4** | Unchanged | Q8BWN8 | 11 | 81 | 7 | 421 | 46.5 | 7.85 | 36 | 1.265 | 0.225 | 2.005 | 0.158 | 2.087 | 0.105 | 0.960 | 0.925 |
| 1293 | **Acsm3** | Unchanged | Q3UNX5 | 17 | 121 | 17 | 580 | 65.6 | 7.46 | 36 | 1.340 | 0.055 | 2.148 | 0.242 | 0.825 | 0.321 | 2.604 | 0.187 |
| 1294 | **Sar1aiso** | Unchanged | P36536 | 5 | 96 | 1 | 198 | 22.4 | 6.93 | 36 | 0.360 | 0.222 | 0.002 | 0.063 | 0.566 | 0.438 | 0.004 | 0.064 |
| 1295 | **Apool** | Unchanged | Q78IK4 | 7 | 58 | 7 | 265 | 29.2 | 9.31 | 36 | 1.088 | 0.675 | 1.409 | 0.105 | 0.876 | 0.640 | 1.608 | 0.132 |
| 1296 | **Eif3m** | UP | Q99JX4 | 7 | 67 | 7 | 374 | 42.5 | 5.74 | 36 | 1.664 | 0.169 | 3.094 | 0.090 | 1.702 | 0.010 | 1.818 | 0.238 |
| 1297 | **Atp2a2** | Unchanged | O55143 | 29 | 308 | 20 | 1044 | 114.8 | 5.34 | 36 | 0.946 | 0.833 | 1.572 | 0.244 | 0.754 | 0.203 | 2.086 | 0.128 |
| 1298 | **Hmgb1** | Unchanged | P63158 | 9 | 53 | 8 | 215 | 24.9 | 5.74 | 36 | 0.927 | 0.888 | 1.420 | 0.599 | 2.386 | 0.198 | 0.595 | 0.384 |
| 1299 | **Umps** | UP | P13439 | 11 | 108 | 11 | 481 | 52.3 | 6.61 | 36 | 1.760 | 0.027 | 2.266 | 0.015 | 2.041 | 0.001 | 1.111 | 0.608 |
| 1300 | **Abce1** | Down | P61222 | 15 | 114 | 15 | 599 | 67.3 | 8.34 | 36 | 0.926 | 0.570 | 1.023 | 0.936 | 0.504 | 0.001 | 2.029 | 0.099 |
| 1301 | **Slirp** | Unchanged | Q9D8T7 | 3 | 14 | 3 | 112 | 12.6 | 9.82 | 36 | 1.227 | 0.606 | 1.728 | 0.206 | 1.111 | 0.778 | 1.556 | 0.177 |

| | | | | | | | | | | | | | | | | | | |
|---|---|---|---|---|---|---|---|---|---|---|---|---|---|---|---|---|---|---|
| 1302 | **Prkcsh** | Unchanged | O08795 | 15 | 140 | 15 | 521 | 58.8 | 4.46 | 36 | 0.951 | 0.777 | 1.512 | 0.419 | 0.897 | 0.747 | 1.685 | 0.366 |
| 1303 | **A1cf** | UP | Q5YD48 | 15 | 81 | 14 | 595 | 65.6 | 8.59 | 36 | 1.169 | 0.599 | 2.208 | 0.044 | 1.179 | 0.586 | 1.873 | 0.112 |
| 1304 | **Ak1** | Unchanged | Q9R0Y5 | 5 | 11 | 5 | 194 | 21.5 | 5.81 | 36 | 0.683 | 0.630 | 1.447 | 0.626 | 0.494 | 0.446 | 2.926 | 0.174 |
| 1305 | **Tmod3** | Unchanged | Q9JHJ0 | 9 | 42 | 9 | 352 | 39.5 | 5.14 | 36 | 0.688 | 0.421 | 1.507 | 0.303 | 1.172 | 0.660 | 1.285 | 0.371 |
| 1306 | **Sec24d** | UP | Q6NXL1 | 21 | 63 | 21 | 1032 | 112.6 | 7.18 | 35 | 1.419 | 0.016 | 1.974 | 0.001 | 1.757 | 0.011 | 1.123 | 0.409 |
| 1307 | **Atad1** | Unchanged | Q9D5T0 | 8 | 43 | 8 | 361 | 40.7 | 6.9 | 35 | 0.897 | 0.577 | 1.644 | 0.267 | 1.197 | 0.636 | 1.374 | 0.502 |
| 1308 | **Mars** | Unchanged | E9QB02 | 21 | 64 | 21 | 910 | 102.3 | 6.92 | 35 | 1.023 | 0.953 | 1.003 | 0.993 | 1.175 | 0.579 | 0.853 | 0.576 |
| 1309 | **B2m** | Unchanged | P01887 | 3 | 13 | 3 | 119 | 13.8 | 8.44 | 35 | 0.958 | 0.941 | 0.299 | 0.231 | 2.014 | 0.199 | 0.149 | 0.016 |
| 1310 | **Txndc12** | Unchanged | Q9CQU0 | 5 | 45 | 5 | 170 | 19 | 5.26 | 35 | 1.032 | 0.943 | 1.319 | 0.498 | 1.868 | 0.163 | 0.706 | 0.392 |
| 1311 | **Gstm5** | UP | P48774 | 7 | 64 | 6 | 224 | 26.6 | 7.21 | 35 | 2.255 | 0.138 | 4.233 | 0.041 | 2.253 | 0.105 | 1.879 | 0.220 |
| 1312 | **Hnmt** | Unchanged | Q91VF2 | 7 | 51 | 7 | 295 | 33.6 | 5.06 | 35 | 0.996 | 0.995 | 1.156 | 0.829 | 0.659 | 0.475 | 1.756 | 0.407 |
| 1313 | **Asgr1** | Unchanged | P34927 | 6 | 80 | 6 | 284 | 32.6 | 6.44 | 35 | 1.007 | 0.975 | 0.755 | 0.302 | 0.632 | 0.052 | 1.194 | 0.501 |
| 1314 | **Ran** | Unchanged | P62827 | 8 | 101 | 8 | 216 | 24.4 | 7.49 | 35 | 0.696 | 0.148 | 2.309 | 0.055 | 2.033 | 0.061 | 1.136 | 0.707 |
| 1315 | **Ap2m1** | Unchanged | P84091 | 11 | 80 | 11 | 435 | 49.6 | 9.54 | 35 | 0.704 | 0.340 | 0.809 | 0.566 | 1.082 | 0.780 | 0.748 | 0.260 |
| 1316 | **Aimp1** | Unchanged | P31230 | 7 | 34 | 7 | 310 | 34 | 8.35 | 35 | 0.877 | 0.656 | 0.872 | 0.724 | 0.737 | 0.327 | 1.184 | 0.704 |
| 1317 | **Carhsp1** | Unchanged | Q9CR86 | 2 | 43 | 2 | 148 | 16.1 | 8.21 | 35 | 1.306 | 0.602 | 1.201 | 0.724 | 1.484 | 0.360 | 0.809 | 0.437 |
| 1318 | **Cops9** | Unchanged | Q3U898 | 1 | 7 | 1 | 57 | 6.2 | 3.83 | 35 | 0.340 | 0.237 | 0.697 | 0.577 | 0.199 | 0.145 | 3.493 | 0.040 |
| 1319 | **H2-Q10** | Unchanged | P01898 | 9 | 48 | 6 | 325 | 37.2 | 5.25 | 35 | 0.805 | 0.487 | 1.315 | 0.357 | 0.687 | 0.246 | 1.915 | 0.052 |
| 1320 | **Mlec** | Unchanged | Q6ZQI3 | 9 | 46 | 9 | 291 | 32.3 | 6.05 | 35 | 1.177 | 0.453 | 0.591 | 0.138 | 1.086 | 0.817 | 0.544 | 0.191 |
| 1321 | **S100a10** | UP | P08207 | 2 | 7 | 2 | 97 | 11.2 | 6.77 | 18 | 0.825 | 0.647 | 1.152 | 0.781 | 3.118 | 0.024 | 0.370 | 0.050 |
| 1322 | **Nudcd2** | Down | Q9CQ48 | 3 | 9 | 3 | 157 | 17.6 | 5.07 | 35 | 0.251 | 0.046 | 0.104 | 0.014 | 0.281 | 0.052 | 0.370 | 0.231 |
| 1323 | **Aldoc** | Unchanged | P05063 | 10 | 53 | 7 | 363 | 39.4 | 7.12 | 35 | 0.544 | 0.211 | 0.830 | 0.639 | 7.367 | 0.150 | 0.113 | 0.140 |
| 1324 | **Idh3a** | Unchanged | Q9D6R2 | 11 | 153 | 11 | 366 | 39.6 | 6.73 | 35 | 1.523 | 0.387 | 0.948 | 0.823 | 1.058 | 0.839 | 0.896 | 0.727 |
| 1325 | **Ttc36** | Unchanged | Q8VBW8 | 7 | 67 | 7 | 186 | 20.1 | 5.29 | 35 | 0.586 | 0.253 | 0.679 | 0.381 | 0.687 | 0.380 | 0.988 | 0.954 |
| 1326 | **Fkbp4** | Unchanged | P30416 | 14 | 68 | 14 | 458 | 51.5 | 5.72 | 35 | 0.445 | 0.085 | 0.444 | 0.062 | 0.700 | 0.314 | 0.634 | 0.175 |
| 1327 | **Kras** | UP | P32883 | 5 | 12 | 2 | 189 | 21.6 | 6.77 | 35 | 1.439 | 0.550 | 0.674 | 0.510 | 3.472 | 0.033 | 0.194 | 0.017 |
| 1328 | **Nono** | Down | Q99K48 | 9 | 91 | 9 | 473 | 54.5 | 8.95 | 35 | 0.148 | 0.000 | 0.131 | 0.000 | 0.160 | 0.000 | 0.819 | 0.444 |
| 1329 | **Glrx5** | Unchanged | Q80Y14 | 4 | 64 | 4 | 152 | 16.3 | 6.55 | 35 | 0.968 | 0.939 | 1.182 | 0.672 | 0.884 | 0.754 | 1.337 | 0.456 |
| 1330 | **Arpc3** | Down | Q9JM76 | 7 | 58 | 7 | 178 | 20.5 | 8.59 | 35 | 0.332 | 0.018 | 1.183 | 0.603 | 0.494 | 0.125 | 2.396 | 0.105 |
| 1331 | **Rpl32** | Unchanged | P62911 | 6 | 41 | 6 | 135 | 15.9 | 11.33 | 35 | 0.933 | 0.857 | 0.817 | 0.612 | 0.487 | 0.141 | 1.679 | 0.265 |
| 1332 | **Hnrnpu** | Unchanged | Q8VEK3 | 21 | 174 | 20 | 800 | 87.9 | 6.24 | 35 | 0.889 | 0.376 | 1.627 | 0.077 | 1.229 | 0.072 | 1.324 | 0.245 |
| 1333 | **Hspa1b** | Down | P17879 | 14 | 190 | 8 | 642 | 70.1 | 5.72 | 35 | 0.420 | 0.017 | 0.495 | 0.063 | 0.421 | 0.019 | 1.177 | 0.669 |
| 1334 | **Eif5** | UP | P59325 | 12 | 72 | 12 | 429 | 48.9 | 5.52 | 35 | 1.165 | 0.631 | 1.774 | 0.043 | 1.030 | 0.923 | 1.723 | 0.045 |
| 1335 | **Cops5** | Unchanged | O35864 | 8 | 45 | 8 | 334 | 37.5 | 6.54 | 35 | 0.710 | 0.279 | 0.718 | 0.325 | 1.223 | 0.454 | 0.587 | 0.058 |
| 1336 | **Nnt** | Unchanged | Q61941 | 31 | 294 | 31 | 1086 | 113.8 | 7.64 | 35 | 0.623 | 0.195 | 0.728 | 0.299 | 0.733 | 0.322 | 0.993 | 0.966 |
| 1337 | **Syncrip** | Unchanged | Q7TMK9 | 16 | 190 | 12 | 623 | 69.6 | 8.59 | 35 | 1.493 | 0.341 | 1.935 | 0.109 | 1.525 | 0.231 | 1.269 | 0.518 |
| 1338 | **Vps29** | Unchanged | Q9QZ88 | 4 | 51 | 4 | 182 | 20.5 | 6.79 | 35 | 0.833 | 0.653 | 0.957 | 0.895 | 1.229 | 0.483 | 0.779 | 0.248 |
| 1339 | **Idh3g** | Unchanged | P70404 | 10 | 110 | 10 | 393 | 42.8 | 9.01 | 35 | 1.052 | 0.857 | 0.734 | 0.269 | 1.092 | 0.668 | 0.672 | 0.159 |
| 1340 | **Aqp1** | Unchanged | Q02013 | 4 | 54 | 4 | 269 | 28.8 | 7.43 | 35 | 1.119 | 0.750 | 2.206 | 0.397 | 1.085 | 0.796 | 2.033 | 0.435 |
| 1341 | **Nags** | Unchanged | Q8R4H7 | 11 | 54 | 11 | 527 | 57.5 | 9 | 35 | 1.300 | 0.555 | 1.684 | 0.402 | 1.017 | 0.970 | 1.656 | 0.340 |
| 1342 | **Aacs** | Unchanged | Q9D2R0 | 16 | 86 | 16 | 672 | 75.2 | 6.71 | 35 | 0.566 | 0.111 | 1.480 | 0.338 | 0.560 | 0.116 | 2.642 | 0.062 |

| | | | | | | | | | | | | | | | | | | |
|---|---|---|---|---|---|---|---|---|---|---|---|---|---|---|---|---|---|---|
| 1343 | **Wars** | Unchanged | P32921 | 11 | 77 | 11 | 481 | 54.3 | 6.89 | 35 | 0.909 | 0.430 | 1.248 | 0.403 | 0.913 | 0.434 | 1.368 | 0.260 |
| 1344 | **Akr1b1** | Unchanged | P45376 | 9 | 74 | 8 | 316 | 35.7 | 7.18 | 34 | 1.511 | 0.516 | 0.844 | 0.746 | 1.388 | 0.628 | 0.608 | 0.478 |
| 1345 | **Sec13** | UP | Q9D1M0 | 7 | 89 | 7 | 322 | 35.5 | 5.38 | 34 | 1.380 | 0.268 | 1.508 | 0.405 | 1.667 | 0.034 | 0.904 | 0.780 |
| 1346 | **Pdcd10** | Unchanged | Q8VE70 | 4 | 7 | 4 | 212 | 24.7 | 8.19 | 34 | 1.245 | 0.480 | 1.904 | 0.213 | 1.768 | 0.251 | 1.077 | 0.884 |
| 1347 | **Ighg2c** | Unchanged | F6TQW2 | 11 | 96 | 9 | 404 | 44.2 | 6.9 | 34 | 0.578 | 0.187 | 0.599 | 0.238 | 0.606 | 0.286 | 0.988 | 0.976 |
| 1348 | **Tmed4** | Unchanged | Q8R1V4 | 6 | 74 | 5 | 227 | 26 | 8.18 | 34 | 1.173 | 0.676 | 1.424 | 0.385 | 0.966 | 0.911 | 1.474 | 0.359 |
| 1349 | **Mapk1** | Unchanged | P63085 | 8 | 39 | 4 | 358 | 41.2 | 6.98 | 34 | 1.363 | 0.522 | 0.615 | 0.420 | 0.989 | 0.979 | 0.621 | 0.316 |
| 1350 | **Eif4a3** | Unchanged | Q91VC3 | 11 | 112 | 8 | 411 | 46.8 | 6.73 | 34 | 0.862 | 0.498 | 0.636 | 0.139 | 0.828 | 0.359 | 0.768 | 0.322 |
| 1351 | **LOC433053** | Unchanged | P01843 | 2 | 23 | 2 | 105 | 11.6 | 6.27 | 34 | 0.324 | 0.133 | 1.732 | 0.428 | 0.876 | 0.811 | 1.976 | 0.359 |
| 1352 | **Atp1b3** | Unchanged | P97370 | 5 | 24 | 5 | 278 | 31.8 | 8.51 | 34 | 0.635 | 0.350 | 1.028 | 0.939 | 1.387 | 0.574 | 0.742 | 0.580 |
| 1353 | **Ndufc2** | Unchanged | Q9CQ54 | 3 | 57 | 3 | 120 | 14.2 | 9.2 | 34 | 1.034 | 0.936 | 0.844 | 0.632 | 0.703 | 0.413 | 1.200 | 0.460 |
| 1354 | **Gorasp2** | Unchanged | Q99JX3 | 9 | 79 | 9 | 451 | 47 | 4.79 | 34 | 0.694 | 0.313 | 1.149 | 0.643 | 0.800 | 0.443 | 1.436 | 0.114 |
| 1355 | **Ier3ip1** | Down | Q9CR20 | 2 | 27 | 2 | 82 | 9 | 8.22 | 34 | 0.686 | 0.194 | 0.728 | 0.282 | 0.369 | 0.028 | 1.974 | 0.045 |
| 1356 | **Ldhb** | Unchanged | P16125 | 10 | 42 | 9 | 334 | 36.5 | 6.05 | 34 | 0.792 | 0.497 | 1.012 | 0.978 | 1.149 | 0.695 | 0.881 | 0.760 |
| 1357 | **Ufc1** | Unchanged | Q9CR09 | 4 | 52 | 4 | 167 | 19.5 | 7.4 | 34 | 1.825 | 0.235 | 1.562 | 0.164 | 2.009 | 0.064 | 0.777 | 0.406 |
| 1358 | **Atg7** | Down | Q9D906 | 15 | 84 | 15 | 698 | 77.5 | 6.4 | 34 | 0.829 | 0.420 | 0.675 | 0.137 | 0.444 | 0.016 | 1.520 | 0.118 |
| 1359 | **Hnrnpa1** | UP | P49312 | 10 | 80 | 8 | 320 | 34.2 | 9.23 | 34 | 1.095 | 0.787 | 1.987 | 0.137 | 3.266 | 0.007 | 0.608 | 0.135 |
| 1360 | **Tpm1** | UP | F8WID5 | 15 | 68 | 5 | 326 | 37.4 | 4.72 | 34 | 1.831 | 0.134 | 3.972 | 0.213 | 3.750 | 0.002 | 1.059 | 0.923 |
| 1361 | **Sdf2l1** | Down | Q9ESP1 | 4 | 37 | 4 | 221 | 23.6 | 7.42 | 34 | 0.660 | 0.067 | 0.614 | 0.021 | 0.673 | 0.178 | 0.912 | 0.760 |
| 1362 | **Tpm4** | Unchanged | Q6IRU2 | 11 | 60 | 10 | 248 | 28.5 | 4.68 | 34 | 0.288 | 0.062 | 0.720 | 0.495 | 1.125 | 0.837 | 0.640 | 0.470 |
| 1363 | **Ighm** | Unchanged | A0A075B5P6 | 12 | 52 | 12 | 455 | 50 | 7.01 | 34 | 1.549 | 0.051 | 1.122 | 0.586 | 1.261 | 0.230 | 0.889 | 0.551 |
| 1364 | **Hsd17b7** | UP | O88736 | 7 | 49 | 7 | 334 | 37.3 | 6.73 | 34 | 1.577 | 0.075 | 2.898 | 0.021 | 1.266 | 0.491 | 2.290 | 0.055 |
| 1365 | **Retsat** | Unchanged | Q64FW2 | 11 | 73 | 11 | 609 | 67.3 | 8.95 | 34 | 0.514 | 0.317 | 0.590 | 0.424 | 0.378 | 0.208 | 1.563 | 0.362 |
| 1366 | **Scfd1** | UP | Q8BRF7 | 11 | 92 | 11 | 639 | 72.3 | 6.38 | 34 | 1.283 | 0.094 | 2.199 | 0.017 | 1.295 | 0.027 | 1.698 | 0.054 |
| 1367 | **Nfs1** | Unchanged | Q9Z1J3 | 12 | 45 | 12 | 459 | 50.5 | 8.16 | 34 | 1.313 | 0.311 | 0.979 | 0.954 | 1.131 | 0.433 | 0.866 | 0.670 |
| 1368 | **Stat3** | Unchanged | P42227 | 16 | 85 | 16 | 770 | 88 | 6.3 | 34 | 0.896 | 0.660 | 1.667 | 0.138 | 1.236 | 0.695 | 1.349 | 0.530 |
| 1369 | **Uchl5** | Unchanged | Q9WUP7 | 9 | 58 | 9 | 329 | 37.6 | 5.33 | 34 | 0.831 | 0.594 | 0.588 | 0.126 | 0.477 | 0.060 | 1.231 | 0.135 |
| 1370 | **Idnk** | Unchanged | Q8R0J8 | 4 | 25 | 4 | 184 | 20 | 6.33 | 34 | 1.105 | 0.836 | 0.980 | 0.965 | 0.735 | 0.384 | 1.334 | 0.529 |
| 1371 | **Csrp1** | UP | P97315 | 4 | 28 | 4 | 193 | 20.6 | 8.57 | 34 | 0.926 | 0.882 | 1.964 | 0.368 | 3.832 | 0.046 | 0.512 | 0.243 |
| 1372 | **Ighv1-43** | Unchanged | A0A075B5V7 | 2 | 7 | 1 | 98 | 10.8 | 8.98 | 34 | #DIV/0! | #DIV/0! | #DIV/0! | #DIV/0! | #DIV/0! | #DIV/0! | #DIV/0! | #DIV/0! |
| 1373 | **Grpel1** | Unchanged | Q99LP6 | 7 | 90 | 7 | 217 | 24.3 | 8.38 | 34 | 1.193 | 0.339 | 1.261 | 0.206 | 0.908 | 0.503 | 1.389 | 0.090 |
| 1374 | **Nomo1** | Unchanged | Q6GQT9 | 24 | 114 | 24 | 1214 | 133.3 | 6.09 | 34 | 0.561 | 0.328 | 0.602 | 0.379 | 0.366 | 0.156 | 1.644 | 0.135 |
| 1375 | **Comtd1** | Unchanged | Q8BIG7 | 5 | 20 | 5 | 262 | 28.9 | 8.32 | 34 | 1.118 | 0.629 | 3.390 | 0.129 | 1.152 | 0.400 | 2.943 | 0.153 |
| 1376 | **Isg15** | Unchanged | Q64339 | 2 | 6 | 2 | 161 | 17.9 | 7.9 | 34 | 3.297 | 0.052 | 7.596 | 0.111 | 4.276 | 0.104 | 1.777 | 0.438 |
| 1377 | **Dlst** | Unchanged | Q9D2G2 | 13 | 280 | 13 | 454 | 49 | 8.95 | 33 | 0.985 | 0.921 | 1.019 | 0.928 | 1.014 | 0.925 | 1.005 | 0.978 |
| 1378 | **Pmm2** | Unchanged | Q9Z2M7 | 8 | 46 | 8 | 242 | 27.6 | 6.42 | 33 | 0.895 | 0.481 | 0.672 | 0.117 | 0.759 | 0.182 | 0.886 | 0.631 |
| 1379 | **C8a** | Unchanged | Q8K182 | 12 | 60 | 12 | 587 | 66 | 6.54 | 33 | 2.562 | 0.421 | 7.686 | 0.315 | 4.136 | 0.100 | 1.858 | 0.596 |
| 1380 | **Sptbn1** | Unchanged | Q62261 | 53 | 303 | 50 | 2363 | 274.1 | 5.58 | 33 | 1.555 | 0.418 | 1.573 | 0.403 | 1.260 | 0.588 | 1.248 | 0.670 |
| 1381 | **Nras** | Unchanged | P08556 | 5 | 6 | 2 | 189 | 21.2 | 5.17 | 33 | 1.236 | 0.658 | 0.938 | 0.866 | 0.469 | 0.183 | 1.999 | 0.002 |
| 1382 | **Dnm2** | UP | P39054 | 20 | 109 | 20 | 870 | 98.1 | 7.43 | 33 | 1.021 | 0.922 | 1.884 | 0.118 | 1.550 | 0.035 | 1.215 | 0.532 |
| 1383 | **Rabif** | Unchanged | Q91X96 | 1 | 1 | 1 | 123 | 13.9 | 5.52 | 33 | 7.702 | 0.390 | 30.860 | 0.156 | 1.507 | 0.311 | 20.475 | 0.162 |

| | | | | | | | | | | | | | | | | | |
|---|---|---|---|---|---|---|---|---|---|---|---|---|---|---|---|---|---|
| 1384 | **Nmt1** | UP | O70310 | 11 | 52 | 11 | 496 | 56.9 | 8 | 33 | 1.277 | 0.400 | 2.057 | 0.006 | 1.242 | 0.388 | 1.656 | 0.046 |
| 1385 | **Ifi35** | Unchanged | Q9D8C4 | 5 | 44 | 5 | 286 | 31.9 | 5.87 | 33 | 1.008 | 0.969 | 1.110 | 0.798 | 0.965 | 0.866 | 1.151 | 0.722 |
| 1386 | **Fadd** | UP | Q61160 | 3 | 16 | 3 | 205 | 22.9 | 6.04 | 33 | 2.920 | 0.193 | 5.113 | 0.028 | 1.084 | 0.876 | 4.715 | 0.030 |
| 1387 | **Ddx6** | Unchanged | P54823 | 10 | 47 | 10 | 483 | 54.2 | 8.66 | 33 | 0.709 | 0.289 | 0.992 | 0.977 | 0.964 | 0.913 | 1.030 | 0.912 |
| 1388 | **Pir** | UP | Q9D711 | 5 | 31 | 5 | 290 | 32 | 7.11 | 33 | 1.586 | 0.203 | 3.496 | 0.147 | 4.818 | 0.000 | 0.726 | 0.429 |
| 1389 | **Asb3** | Unchanged | D6RDR7 | 1 | 1 | 1 | 124 | 13.9 | 9.07 | 33 | #DIV/0! | #DIV/0! | #DIV/0! | #DIV/0! | #DIV/0! | #DIV/0! | #DIV/0! | #DIV/0! |
| 1390 | **Arpc1b** | UP | Q9WV32 | 8 | 96 | 7 | 372 | 41 | 8.35 | 33 | 0.958 | 0.864 | 0.903 | 0.859 | 2.654 | 0.001 | 0.340 | 0.012 |
| 1391 | **U2af2** | UP | P26369 | 9 | 40 | 9 | 475 | 53.5 | 9.09 | 33 | 16.467 | 0.131 | 7.595 | 0.089 | 26.223 | 0.002 | 0.290 | 0.020 |
| 1392 | **Tpd52** | Unchanged | Q62393 | 5 | 18 | 5 | 224 | 24.3 | 4.72 | 33 | 0.704 | 0.460 | 1.278 | 0.461 | 1.050 | 0.899 | 1.217 | 0.430 |
| 1393 | **Irgm1** | Unchanged | Q60766 | 9 | 64 | 9 | 409 | 46.5 | 8.28 | 33 | 0.822 | 0.356 | 0.820 | 0.592 | 0.865 | 0.521 | 0.947 | 0.885 |
| 1394 | **Hnrnpab** | UP | Q99020 | 10 | 117 | 9 | 285 | 30.8 | 7.91 | 33 | 1.607 | 0.190 | 2.633 | 0.141 | 2.177 | 0.039 | 1.209 | 0.688 |
| 1395 | **Lars** | Unchanged | Q8BMJ2 | 28 | 223 | 28 | 1178 | 134.1 | 7.05 | 33 | 1.297 | 0.098 | 1.249 | 0.399 | 0.967 | 0.884 | 1.291 | 0.426 |
| 1396 | **Tpk1** | Down | Q9R0M5 | 4 | 26 | 4 | 243 | 27.1 | 5.76 | 33 | 1.075 | 0.823 | 1.136 | 0.643 | 0.362 | 0.004 | 3.134 | 0.025 |
| 1397 | **Rdh11** | Unchanged | Q9QYF1 | 6 | 36 | 6 | 316 | 35.1 | 8.91 | 33 | 0.573 | 0.327 | 0.778 | 0.600 | 0.577 | 0.321 | 1.348 | 0.431 |
| 1398 | **Cox6c** | Unchanged | Q9CPQ1 | 4 | 78 | 4 | 76 | 8.5 | 10.14 | 17 | 1.193 | 0.603 | 0.472 | 0.086 | 0.882 | 0.768 | 0.535 | 0.314 |
| 1399 | **Pex5** | Unchanged | O09012 | 12 | 54 | 12 | 639 | 70.7 | 4.55 | 33 | 1.059 | 0.906 | 1.219 | 0.596 | 1.154 | 0.698 | 1.056 | 0.831 |
| 1400 | **Agxt** | Unchanged | O35423 | 12 | 143 | 12 | 414 | 45.9 | 8.18 | 33 | 0.837 | 0.712 | 0.826 | 0.676 | 0.623 | 0.425 | 1.327 | 0.575 |
| 1401 | **Reep6** | Unchanged | Q9JM62 | 5 | 104 | 5 | 201 | 22.2 | 7.33 | 33 | 1.072 | 0.645 | 1.071 | 0.802 | 0.965 | 0.803 | 1.110 | 0.733 |
| 1402 | **Rrbp1** | Unchanged | Q99PL5 | 45 | 598 | 45 | 1605 | 172.8 | 9.33 | 33 | 0.930 | 0.792 | 0.788 | 0.387 | 0.770 | 0.398 | 1.022 | 0.953 |
| 1403 | **Igkv1-117** | Unchanged | P01631 | 2 | 13 | 2 | 113 | 12.3 | 8.88 | 33 | 0.186 | 0.206 | 1.788 | 0.608 | 0.527 | 0.456 | 3.394 | 0.383 |
| 1404 | **Oxsm** | Unchanged | Q9D404 | 7 | 65 | 7 | 459 | 48.6 | 7.06 | 33 | 1.410 | 0.242 | 1.734 | 0.130 | 1.282 | 0.416 | 1.352 | 0.396 |
| 1405 | **Rbm3** | Unchanged | O89086 | 2 | 5 | 2 | 153 | 16.6 | 7.5 | 33 | 2.866 | 0.352 | 5.883 | 0.137 | 0.518 | 0.320 | 11.359 | 0.104 |
| 1406 | **Coro1b** | Unchanged | Q9WUM3 | 11 | 82 | 11 | 484 | 53.9 | 5.78 | 33 | 1.033 | 0.895 | 1.647 | 0.103 | 1.866 | 0.166 | 0.882 | 0.740 |
| 1407 | **Gimap4** | UP | D3YTN4 | 7 | 28 | 7 | 328 | 38 | 6.23 | 33 | 1.284 | 0.580 | 2.255 | 0.008 | 1.194 | 0.662 | 1.889 | 0.019 |
| 1408 | **Erap1** | UP | Q9EQH2 | 24 | 193 | 24 | 930 | 106.5 | 6.2 | 33 | 1.455 | 0.171 | 2.245 | 0.000 | 0.764 | 0.165 | 2.938 | 0.000 |
| 1409 | **C030006K11Rik** | Down | Q8VE95 | 5 | 28 | 5 | 218 | 24.3 | 8.7 | 33 | 0.670 | 0.288 | 0.951 | 0.888 | 0.468 | 0.015 | 2.030 | 0.187 |
| 1410 | **Ranbp1** | UP | P34022 | 4 | 48 | 4 | 203 | 23.6 | 5.22 | 33 | 0.987 | 0.951 | 1.273 | 0.454 | 2.159 | 0.036 | 0.590 | 0.140 |
| 1411 | **Sept11** | Unchanged | Q8C1B7 | 9 | 74 | 7 | 431 | 49.7 | 6.68 | 32 | 0.966 | 0.880 | 0.935 | 0.852 | 1.348 | 0.373 | 0.694 | 0.362 |
| 1412 | **Rmdn1** | Unchanged | Q9DCV4 | 7 | 43 | 7 | 305 | 35 | 8.7 | 32 | 1.279 | 0.390 | 0.943 | 0.874 | 1.083 | 0.668 | 0.871 | 0.687 |
| 1413 | **Ruvbl1** | Unchanged | P60122 | 7 | 36 | 7 | 456 | 50.2 | 6.42 | 32 | 1.633 | 0.179 | 0.804 | 0.656 | 0.748 | 0.354 | 1.076 | 0.878 |
| 1414 | **Rpl27a** | Unchanged | P14115 | 7 | 79 | 7 | 148 | 16.6 | 11.12 | 32 | 0.802 | 0.220 | 0.845 | 0.382 | 0.992 | 0.972 | 0.851 | 0.551 |
| 1415 | **Yars2** | Unchanged | Q8BYL4 | 7 | 16 | 7 | 472 | 52.6 | 9.16 | 32 | 0.814 | 0.359 | 1.587 | 0.299 | 1.034 | 0.921 | 1.535 | 0.381 |
| 1416 | **Rpl26** | Unchanged | P61255 | 9 | 67 | 9 | 145 | 17.2 | 10.55 | 32 | 0.554 | 0.239 | 0.931 | 0.891 | 0.770 | 0.560 | 1.209 | 0.726 |
| 1417 | **Apoa2** | Unchanged | P09813 | 2 | 20 | 2 | 102 | 11.3 | 7.18 | 32 | 0.758 | 0.692 | 0.233 | 0.178 | 0.467 | 0.336 | 0.499 | 0.067 |
| 1418 | **Alcam** | UP | Q61490 | 12 | 45 | 12 | 583 | 65.1 | 6.15 | 32 | 1.084 | 0.772 | 1.047 | 0.901 | 1.745 | 0.032 | 0.600 | 0.073 |
| 1419 | **Dlat** | Unchanged | Q8BMF4 | 16 | 214 | 16 | 642 | 67.9 | 8.57 | 32 | 0.930 | 0.855 | 0.819 | 0.664 | 0.817 | 0.592 | 1.002 | 0.996 |
| 1420 | **Tcea1** | Unchanged | P10711 | 7 | 29 | 6 | 301 | 33.9 | 8.38 | 32 | 0.563 | 0.183 | 0.954 | 0.898 | 1.579 | 0.324 | 0.604 | 0.279 |
| 1421 | **Snrpd2** | Unchanged | P62317 | 4 | 62 | 4 | 118 | 13.5 | 9.91 | 32 | 1.342 | 0.235 | 1.797 | 0.151 | 3.051 | 0.081 | 0.589 | 0.300 |
| 1422 | **Opa1** | Unchanged | P58281 | 21 | 130 | 21 | 960 | 111.3 | 7.55 | 32 | 1.153 | 0.443 | 1.378 | 0.206 | 1.330 | 0.175 | 1.036 | 0.860 |
| 1423 | **Emc3** | Unchanged | Q99KI3 | 5 | 21 | 5 | 261 | 30 | 6.81 | 32 | 0.703 | 0.607 | 0.336 | 0.151 | 1.198 | 0.691 | 0.281 | 0.008 |
| 1424 | **Psmd4** | Unchanged | O35226 | 7 | 72 | 7 | 376 | 40.7 | 4.79 | 28 | 1.004 | 0.988 | 0.852 | 0.618 | 0.719 | 0.241 | 1.185 | 0.505 |

| | | | | | | | | | | | | | | | | | | |
|---|---|---|---|---|---|---|---|---|---|---|---|---|---|---|---|---|---|---|
| 1425 | **Mtx1** | Unchanged | P47802 | 7 | 34 | 7 | 317 | 35.6 | 6.18 | 32 | 1.031 | 0.887 | 1.512 | 0.119 | 0.748 | 0.446 | 2.020 | 0.071 |
| 1426 | **Ifitm3** | Unchanged | Q9CQW9 | 3 | 10 | 2 | 137 | 14.9 | 7.4 | 32 | 3.450 | 0.467 | 2.357 | 0.380 | 3.506 | 0.084 | 0.672 | 0.538 |
| 1427 | **Scrn3** | Unchanged | Q3TMH2 | 8 | 89 | 8 | 418 | 47.6 | 5.66 | 32 | 1.053 | 0.885 | 1.182 | 0.580 | 1.021 | 0.948 | 1.158 | 0.101 |
| 1428 | **Ptrh2** | Unchanged | Q8R2Y8 | 4 | 54 | 4 | 181 | 19.5 | 7.42 | 32 | 1.322 | 0.694 | 1.539 | 0.543 | 0.323 | 0.263 | 4.762 | 0.096 |
| 1429 | **Mcatiso** | Unchanged | Q8R3F5 | 6 | 13 | 4 | 381 | 41.9 | 8.1 | 32 | 0.918 | 0.777 | 1.055 | 0.850 | 1.024 | 0.919 | 1.030 | 0.907 |
| 1430 | **Sbds** | Unchanged | P70122 | 7 | 27 | 7 | 250 | 28.8 | 8.76 | 32 | 0.175 | 0.087 | 0.343 | 0.171 | 0.296 | 0.137 | 1.158 | 0.764 |
| 1431 | **Ssr4** | Unchanged | Q62186 | 5 | 82 | 5 | 172 | 18.9 | 5.78 | 32 | 1.506 | 0.308 | 1.103 | 0.721 | 0.989 | 0.973 | 1.115 | 0.764 |
| 1432 | **Anxa7** | UP | Q07076 | 11 | 114 | 11 | 463 | 49.9 | 6.18 | 32 | 1.924 | 0.014 | 1.758 | 0.062 | 1.074 | 0.712 | 1.637 | 0.079 |
| 1433 | **Lasp1** | Unchanged | Q61792 | 8 | 62 | 8 | 263 | 30 | 7.05 | 32 | 2.009 | 0.294 | 0.806 | 0.360 | 1.920 | 0.149 | 0.420 | 0.093 |
| 1434 | **Cdc37** | UP | Q61081 | 9 | 68 | 9 | 379 | 44.6 | 5.34 | 32 | 1.557 | 0.126 | 1.582 | 0.198 | 2.566 | 0.023 | 0.616 | 0.178 |
| 1435 | **Pmpcb** | Unchanged | Q9CXT8 | 12 | 84 | 11 | 489 | 54.6 | 6.99 | 32 | 0.721 | 0.096 | 1.075 | 0.596 | 0.700 | 0.173 | 1.537 | 0.065 |
| 1436 | **Ppid** | Unchanged | Q9CR16 | 10 | 77 | 10 | 370 | 40.7 | 7.43 | 32 | 0.582 | 0.190 | 0.639 | 0.211 | 0.439 | 0.060 | 1.458 | 0.184 |
| 1437 | **Lgals3bp** | Down | Q07797 | 13 | 80 | 12 | 577 | 64.5 | 5.14 | 32 | 0.501 | 0.015 | 1.407 | 0.367 | 0.735 | 0.224 | 1.915 | 0.168 |
| 1438 | **Gart** | UP | Q64737 | 16 | 124 | 16 | 1010 | 107.4 | 6.68 | 32 | 1.697 | 0.030 | 2.159 | 0.043 | 1.612 | 0.005 | 1.340 | 0.286 |
| 1439 | **Rpl24** | Unchanged | Q8BP67 | 6 | 101 | 6 | 157 | 17.8 | 11.25 | 32 | 0.734 | 0.369 | 0.775 | 0.566 | 0.793 | 0.524 | 0.977 | 0.965 |
| 1440 | **Serpinc1** | Unchanged | P32261 | 9 | 58 | 9 | 465 | 52 | 6.46 | 32 | 0.955 | 0.910 | 1.081 | 0.849 | 1.142 | 0.720 | 0.947 | 0.866 |
| 1441 | **Creg1** | UP | O88668 | 5 | 61 | 5 | 220 | 24.4 | 6.42 | 32 | 1.010 | 0.979 | 3.969 | 0.045 | 2.194 | 0.074 | 1.809 | 0.205 |
| 1442 | **Gm5678** | Unchanged | A0A1B0GRG8 | 7 | 76 | 1 | 277 | 30.7 | 6.27 | 32 | 4.584 | 0.273 | 7.713 | 0.302 | 6.505 | 0.110 | 1.186 | 0.863 |
| 1443 | **Rps6** | Unchanged | P62754 | 10 | 189 | 10 | 249 | 28.7 | 10.84 | 32 | 0.943 | 0.698 | 1.052 | 0.823 | 1.008 | 0.962 | 1.043 | 0.862 |
| 1444 | **F2** | Unchanged | P19221 | 13 | 94 | 13 | 618 | 70.2 | 6.43 | 32 | 1.362 | 0.398 | 1.395 | 0.310 | 2.968 | 0.164 | 0.470 | 0.262 |
| 1445 | **Ugt1a10** | UP | E9PXN7 | 12 | 175 | 1 | 530 | 60.3 | 8.35 | 32 | 0.739 | 0.374 | 0.703 | 0.334 | 2.957 | 0.000 | 0.238 | 0.000 |
| 1446 | **Mpi** | Unchanged | Q924M7 | 8 | 47 | 8 | 423 | 46.5 | 5.95 | 32 | 1.448 | 0.420 | 1.791 | 0.346 | 0.841 | 0.730 | 2.130 | 0.220 |
| 1447 | **Eif2a** | Unchanged | Q8BJW6 | 12 | 81 | 12 | 581 | 64.4 | 8.91 | 30 | 0.710 | 0.137 | 1.087 | 0.677 | 0.746 | 0.340 | 1.457 | 0.173 |
| 1448 | **Acss2** | Unchanged | Q9QXG4 | 15 | 123 | 15 | 701 | 78.8 | 6.64 | 32 | 0.879 | 0.684 | 0.959 | 0.885 | 0.679 | 0.320 | 1.412 | 0.330 |
| 1449 | **Clpp** | Unchanged | O88696 | 6 | 75 | 6 | 272 | 29.8 | 7.47 | 32 | 1.480 | 0.534 | 1.023 | 0.964 | 1.341 | 0.648 | 0.763 | 0.703 |
| 1450 | **Tstd1** | Unchanged | E9PY03 | 1 | 5 | 1 | 133 | 14.3 | 8.48 | 32 | 1.199 | 0.777 | 2.167 | 0.278 | 0.498 | 0.269 | 4.348 | 0.107 |
| 1451 | **Tubb6** | Unchanged | Q922F4 | 14 | 241 | 3 | 447 | 50.1 | 4.89 | 32 | 1.249 | 0.692 | 2.258 | 0.342 | 0.985 | 0.981 | 2.292 | 0.357 |
| 1452 | **Rbbp4** | Unchanged | Q60972 | 6 | 23 | 4 | 425 | 47.6 | 4.89 | 32 | 1.103 | 0.807 | 1.154 | 0.726 | 1.923 | 0.286 | 0.600 | 0.375 |
| 1453 | **Pfdn6** | Unchanged | Q03958 | 2 | 10 | 2 | 127 | 14.4 | 8.88 | 31 | 1.052 | 0.906 | 2.157 | 0.279 | 0.376 | 0.063 | 5.733 | 0.107 |
| 1454 | **L2hgdh** | Unchanged | Q91YP0 | 12 | 79 | 12 | 464 | 50.9 | 8.29 | 31 | 2.655 | 0.067 | 1.715 | 0.207 | 1.421 | 0.216 | 1.207 | 0.616 |
| 1455 | **Ppm1a** | Unchanged | P49443 | 8 | 60 | 7 | 382 | 42.4 | 5.36 | 31 | 0.899 | 0.611 | 1.167 | 0.604 | 1.156 | 0.652 | 1.009 | 0.981 |
| 1456 | **Nme3** | Unchanged | Q9WV85 | 3 | 18 | 3 | 169 | 19.1 | 6.68 | 31 | 1.107 | 0.857 | 0.952 | 0.944 | 1.528 | 0.368 | 0.623 | 0.273 |
| 1457 | **Dync1h1** | Unchanged | Q9JHU4 | 98 | 641 | 98 | 4644 | 531.7 | 6.42 | 31 | 1.392 | 0.519 | 1.371 | 0.489 | 1.277 | 0.624 | 1.074 | 0.887 |
| 1458 | **Ap2a1** | Unchanged | P17426 | 20 | 117 | 14 | 977 | 107.6 | 7.03 | 30 | 0.991 | 0.962 | 1.952 | 0.090 | 0.794 | 0.124 | 2.457 | 0.048 |
| 1459 | **Tfg** | Unchanged | Q9Z1A1 | 9 | 26 | 9 | 397 | 43 | 5.1 | 31 | 1.452 | 0.331 | 1.479 | 0.308 | 1.397 | 0.258 | 1.059 | 0.866 |
| 1460 | **Blmh** | Unchanged | Q8R016 | 8 | 84 | 8 | 455 | 52.5 | 6.48 | 31 | 1.217 | 0.513 | 1.121 | 0.682 | 0.845 | 0.599 | 1.327 | 0.205 |
| 1461 | **Nans** | Unchanged | Q99J77 | 7 | 36 | 7 | 359 | 40 | 7.06 | 31 | 1.045 | 0.928 | 0.918 | 0.800 | 0.912 | 0.714 | 1.007 | 0.979 |
| 1462 | **Cops6** | Unchanged | O88545 | 6 | 27 | 6 | 324 | 35.9 | 5.73 | 31 | 1.527 | 0.099 | 1.781 | 0.157 | 0.983 | 0.957 | 1.813 | 0.151 |
| 1463 | **Cbx3iso** | Unchanged | P23198 | 7 | 29 | 1 | 183 | 20.8 | 5.22 | 25 | 1.019 | 0.970 | 1.153 | 0.785 | 1.323 | 0.551 | 0.871 | 0.793 |
| 1464 | **Gusb** | UP | P12265 | 13 | 65 | 13 | 648 | 74.1 | 6.7 | 31 | 2.302 | 0.033 | 3.244 | 0.021 | 2.019 | 0.081 | 1.607 | 0.214 |
| 1465 | **Gars** | UP | Q9CZD3 | 17 | 102 | 17 | 729 | 81.8 | 6.65 | 31 | 1.547 | 0.181 | 2.430 | 0.067 | 2.097 | 0.004 | 1.159 | 0.657 |

| | | | | | | | | | | | | | | | | | | |
|---|---|---|---|---|---|---|---|---|---|---|---|---|---|---|---|---|---|---|
| 1466 | Sept7 | Unchanged | O55131 | 9 | 107 | 8 | 436 | 50.5 | 8.57 | 31 | 1.117 | 0.670 | 1.177 | 0.318 | 1.407 | 0.279 | 0.836 | 0.541 |
| 1467 | Ndufaf2 | Unchanged | Q59J78 | 4 | 29 | 4 | 168 | 19.6 | 8.25 | 31 | 1.320 | 0.490 | 1.248 | 0.603 | 1.088 | 0.810 | 1.147 | 0.744 |
| 1468 | Igh-3 | Unchanged | P01867 | 10 | 77 | 1 | 404 | 44.2 | 6.52 | 31 | 0.094 | 0.076 | 1.104 | 0.894 | 0.034 | 0.061 | 32.436 | 0.117 |
| 1469 | H2-D1 | Unchanged | P01899 | 10 | 69 | 5 | 362 | 40.8 | 6.73 | 31 | 1.277 | 0.630 | 1.970 | 0.117 | 1.669 | 0.188 | 1.180 | 0.593 |
| 1470 | Pex11a | Unchanged | Q9Z211 | 4 | 42 | 4 | 246 | 28.1 | 9.83 | 31 | 1.327 | 0.534 | 1.355 | 0.538 | 1.422 | 0.415 | 0.953 | 0.914 |
| 1471 | Hsph1 | Down | Q61699 | 19 | 106 | 17 | 858 | 96.3 | 5.53 | 31 | 0.247 | 0.027 | 0.337 | 0.047 | 0.355 | 0.051 | 0.951 | 0.825 |
| 1472 | Pura | Unchanged | P42669 | 8 | 121 | 7 | 321 | 34.9 | 6.44 | 31 | 0.518 | 0.174 | 0.290 | 0.061 | 0.431 | 0.114 | 0.674 | 0.118 |
| 1473 | Ctsl | Unchanged | P06797 | 6 | 64 | 6 | 334 | 37.5 | 6.83 | 31 | 0.918 | 0.809 | 1.772 | 0.266 | 1.341 | 0.490 | 1.321 | 0.554 |
| 1474 | Jchain | Unchanged | P01592 | 4 | 16 | 4 | 159 | 18 | 4.89 | 31 | 0.650 | 0.106 | 1.304 | 0.587 | 0.710 | 0.186 | 1.837 | 0.303 |
| 1475 | Atp5f1e | Unchanged | P56382 | 2 | 8 | 2 | 52 | 5.8 | 10.01 | 31 | 0.926 | 0.932 | 0.249 | 0.272 | 0.039 | 0.156 | 6.389 | 0.250 |
| 1476 | Pafah2 | Unchanged | Q8VDG7 | 8 | 75 | 8 | 390 | 43.5 | 7.47 | 31 | 0.724 | 0.020 | 0.968 | 0.812 | 0.878 | 0.552 | 1.103 | 0.665 |
| 1477 | G3bp1 | Unchanged | P97855 | 9 | 67 | 8 | 465 | 51.8 | 5.59 | 28 | 1.416 | 0.186 | 0.922 | 0.751 | 1.089 | 0.246 | 0.846 | 0.484 |
| 1478 | Sec11a | Unchanged | Q9R0P6 | 6 | 27 | 6 | 179 | 20.6 | 9.33 | 31 | 1.327 | 0.460 | 1.493 | 0.404 | 0.687 | 0.390 | 2.175 | 0.164 |
| 1479 | Ndufs8 | Unchanged | Q8K3J1 | 7 | 70 | 7 | 212 | 24 | 6.21 | 31 | 2.706 | 0.170 | 1.771 | 0.170 | 0.672 | 0.429 | 2.635 | 0.023 |
| 1480 | Anpep | UP | P97449 | 19 | 146 | 19 | 966 | 109.6 | 5.9 | 31 | 1.008 | 0.976 | 2.345 | 0.010 | 1.644 | 0.129 | 1.427 | 0.177 |
| 1481 | Taco1 | UP | Q8K0Z7 | 4 | 32 | 4 | 294 | 32.3 | 8.12 | 31 | 1.934 | 0.125 | 2.101 | 0.185 | 2.472 | 0.047 | 0.850 | 0.707 |
| 1482 | Apmap | Unchanged | Q9D7N9 | 7 | 49 | 7 | 415 | 46.4 | 6.32 | 31 | 1.214 | 0.747 | 1.772 | 0.414 | 0.560 | 0.439 | 3.167 | 0.146 |
| 1483 | Psmd8 | Unchanged | Q9CX56 | 7 | 42 | 7 | 353 | 39.9 | 9.58 | 31 | 1.268 | 0.591 | 2.680 | 0.105 | 1.662 | 0.155 | 1.612 | 0.312 |
| 1484 | Eftud2 | Unchanged | O08810 | 19 | 78 | 18 | 971 | 109.3 | 5 | 31 | 1.097 | 0.735 | 0.899 | 0.540 | 1.125 | 0.606 | 0.799 | 0.312 |
| 1485 | Acnat1 | Unchanged | A2AKK5 | 8 | 61 | 5 | 416 | 46 | 8.73 | 31 | 0.853 | 0.632 | 1.257 | 0.675 | 0.871 | 0.612 | 1.444 | 0.541 |
| 1486 | Armc1 | Unchanged | Q9D7A8 | 4 | 17 | 4 | 282 | 31.2 | 5.57 | 30 | 1.507 | 0.596 | 1.476 | 0.574 | 2.411 | 0.220 | 0.612 | 0.496 |
| 1487 | Gba | Unchanged | P17439 | 8 | 39 | 8 | 515 | 57.6 | 7.75 | 30 | 0.676 | 0.354 | 1.333 | 0.463 | 0.823 | 0.575 | 1.619 | 0.179 |
| 1488 | Rpl36 | Down | Q6ZWZ4 | 4 | 36 | 4 | 105 | 12.2 | 11.59 | 30 | 0.505 | 0.028 | 0.387 | 0.010 | 0.449 | 0.004 | 0.862 | 0.670 |
| 1489 | Hnrnpd | Unchanged | Q60668 | 11 | 134 | 9 | 355 | 38.3 | 7.81 | 30 | 0.908 | 0.689 | 0.918 | 0.769 | 0.742 | 0.094 | 1.238 | 0.500 |
| 1490 | Cpne3 | Down | Q8BT60 | 10 | 76 | 10 | 533 | 59.5 | 5.78 | 30 | 0.891 | 0.654 | 0.747 | 0.504 | 0.445 | 0.028 | 1.679 | 0.383 |
| 1491 | Cfh | Unchanged | P06909 | 22 | 109 | 22 | 1234 | 139 | 6.99 | 30 | 0.945 | 0.873 | 1.306 | 0.511 | 0.578 | 0.111 | 2.259 | 0.108 |
| 1492 | Ddx3x | Unchanged | Q62167 | 15 | 146 | 3 | 662 | 73.1 | 7.18 | 30 | 1.208 | 0.515 | 1.816 | 0.186 | 1.320 | 0.472 | 1.376 | 0.448 |
| 1493 | Eloc | Unchanged | P83940 | 2 | 25 | 2 | 112 | 12.5 | 4.78 | 30 | 0.713 | 0.479 | 1.568 | 0.202 | 0.617 | 0.410 | 2.540 | 0.058 |
| 1494 | Impa1 | Unchanged | Q924B0 | 8 | 48 | 8 | 277 | 30.4 | 5.19 | 30 | 1.010 | 0.946 | 1.486 | 0.260 | 0.961 | 0.853 | 1.546 | 0.245 |
| 1495 | Flad1 | Unchanged | Q8R123 | 11 | 72 | 11 | 492 | 54.7 | 6.64 | 30 | 2.819 | 0.084 | 3.365 | 0.250 | 0.272 | 0.088 | 12.383 | 0.137 |
| 1496 | Vwa8 | Unchanged | Q8CC88 | 37 | 185 | 37 | 1905 | 213.3 | 6.6 | 30 | 0.525 | 0.313 | 0.606 | 0.430 | 0.955 | 0.922 | 0.635 | 0.173 |
| 1497 | Tmed9 | Unchanged | Q99KF1 | 8 | 100 | 7 | 235 | 27.1 | 8.41 | 30 | 2.285 | 0.197 | 3.763 | 0.172 | 1.668 | 0.293 | 2.257 | 0.305 |
| 1498 | Rab21 | UP | P35282 | 6 | 36 | 6 | 222 | 24.1 | 7.94 | 30 | 2.124 | 0.075 | 5.656 | 0.088 | 5.454 | 0.001 | 1.037 | 0.938 |
| 1499 | Csnk2a1 | Unchanged | Q60737 | 8 | 28 | 8 | 391 | 45.1 | 7.74 | 30 | 1.081 | 0.800 | 1.391 | 0.392 | 1.720 | 0.185 | 0.809 | 0.613 |
| 1500 | Selenoo | Unchanged | Q9DBC0 | 12 | 80 | 12 | 667 | 74.2 | 5.83 | 30 | 3.258 | 0.329 | 1.065 | 0.905 | 1.277 | 0.552 | 0.834 | 0.727 |
| 1501 | H2afy | Unchanged | Q9QZQ8 | 5 | 13 | 5 | 372 | 39.7 | 9.8 | 30 | 1.935 | 0.252 | 1.831 | 0.236 | 4.131 | 0.072 | 0.443 | 0.194 |
| 1502 | Set | UP | Q9EQU5 | 5 | 62 | 5 | 289 | 33.4 | 4.32 | 30 | 1.208 | 0.523 | 1.868 | 0.038 | 2.193 | 0.101 | 0.852 | 0.647 |
| 1503 | Rps24 | Unchanged | P62849 | 5 | 82 | 5 | 133 | 15.4 | 10.78 | 30 | 1.239 | 0.608 | 1.241 | 0.726 | 1.284 | 0.489 | 0.967 | 0.954 |
| 1504 | Grn | Unchanged | P28798 | 9 | 116 | 9 | 589 | 63.4 | 6.8 | 30 | 0.576 | 0.538 | 0.572 | 0.528 | 0.426 | 0.378 | 1.344 | 0.698 |
| 1505 | Atl2 | Unchanged | Q6PA06 | 10 | 87 | 9 | 583 | 66.2 | 5.43 | 30 | 1.070 | 0.727 | 2.163 | 0.093 | 1.571 | 0.405 | 1.377 | 0.522 |
| 1506 | Gaa | Down | P70699 | 20 | 138 | 20 | 953 | 106.2 | 5.83 | 30 | 1.073 | 0.711 | 0.971 | 0.853 | 0.619 | 0.023 | 1.568 | 0.071 |

| | | | | | | | | | | | | | | | | | | |
|---|---|---|---|---|---|---|---|---|---|---|---|---|---|---|---|---|---|---|
| 1507 | **Igkv1-135** | Down | A0A0B4J1H7 | 3 | 3 | 3 | 120 | 13.3 | 7.12 | 30 | 0.356 | 0.001 | 0.893 | 0.845 | 0.153 | 0.000 | 5.847 | 0.189 |
| 1508 | **Coro7** | Unchanged | Q9D2V7 | 12 | 53 | 12 | 922 | 100.7 | 5.77 | 30 | 1.664 | 0.148 | 1.771 | 0.114 | 0.826 | 0.553 | 2.144 | 0.023 |
| 1509 | **Vtn** | UP | P29788 | 8 | 47 | 8 | 478 | 54.8 | 5.88 | 30 | 1.854 | 0.027 | 3.970 | 0.024 | 3.181 | 0.152 | 1.248 | 0.661 |
| 1510 | **Vnn1** | Unchanged | Q9Z0K8 | 9 | 55 | 9 | 512 | 57.1 | 5.94 | 30 | 0.689 | 0.600 | 0.400 | 0.320 | 0.509 | 0.421 | 0.786 | 0.637 |
| 1511 | **Nudt12** | Unchanged | Q9DCN1 | 10 | 93 | 10 | 462 | 51.5 | 7.12 | 30 | 1.141 | 0.681 | 1.493 | 0.148 | 1.095 | 0.739 | 1.363 | 0.232 |
| 1512 | **Med21** | Unchanged | Q9CQ39 | 1 | 1 | 1 | 144 | 15.6 | 4.53 | 30 | #DIV/0! | #DIV/0! | #DIV/0! | #DIV/0! | #DIV/0! | #DIV/0! | #DIV/0! | #DIV/0! |
| 1513 | **Fkbp11** | Unchanged | Q9D1M7 | 4 | 33 | 4 | 201 | 22.1 | 9.36 | 30 | 0.732 | 0.648 | 1.929 | 0.256 | 1.756 | 0.436 | 1.099 | 0.859 |
| 1514 | **Txnrd2** | UP | Q9JLT4 | 8 | 52 | 7 | 524 | 56.6 | 8.46 | 30 | 1.241 | 0.489 | 2.459 | 0.086 | 1.750 | 0.039 | 1.405 | 0.369 |
| 1515 | **Ifi47** | Unchanged | Q61635 | 9 | 23 | 9 | 420 | 46.8 | 6.74 | 30 | 0.728 | 0.299 | 0.514 | 0.071 | 1.195 | 0.714 | 0.430 | 0.232 |
| 1516 | **Unc5a** | Unchanged | F6TGW0 | 1 | 1 | 1 | 128 | 14.4 | 6.27 | 30 | 0.574 | 0.590 | 0.013 | 0.186 | 1.317 | 0.741 | 0.010 | 0.070 |
| 1517 | **Acot12** | Unchanged | Q9DBK0 | 13 | 95 | 13 | 556 | 61.7 | 7.18 | 30 | 0.983 | 0.946 | 1.163 | 0.584 | 0.839 | 0.529 | 1.386 | 0.176 |
| 1518 | **Cald1** | Unchanged | E9QA16 | 16 | 40 | 1 | 536 | 61.8 | 6.93 | 30 | 0.679 | 0.656 | 0.405 | 0.375 | 1.039 | 0.964 | 0.390 | 0.370 |
| 1519 | **Vapb** | Down | Q9QY76 | 7 | 105 | 6 | 243 | 26.9 | 7.78 | 30 | 0.560 | 0.179 | 1.180 | 0.711 | 0.313 | 0.040 | 3.768 | 0.051 |
| 1520 | **Hnrnph1** | Unchanged | O35737 | 9 | 124 | 3 | 449 | 49.2 | 6.3 | 30 | 0.681 | 0.598 | 1.228 | 0.682 | 1.207 | 0.682 | 1.017 | 0.953 |
| 1521 | **Sdhc** | UP | Q9CZB0 | 5 | 19 | 5 | 169 | 18.4 | 9.94 | 30 | 3.310 | 0.002 | 3.078 | 0.222 | 3.002 | 0.002 | 1.025 | 0.963 |
| 1522 | **Stom** | UP | P54116 | 5 | 43 | 5 | 284 | 31.4 | 6.93 | 30 | 2.089 | 0.140 | 4.707 | 0.001 | 2.615 | 0.001 | 1.800 | 0.026 |
| 1523 | **Wdr61** | Unchanged | Q9ERF3 | 5 | 34 | 5 | 305 | 33.8 | 5.36 | 30 | 1.543 | 0.378 | 1.050 | 0.899 | 1.016 | 0.940 | 1.033 | 0.929 |
| 1524 | **Ncl** | Unchanged | P09405 | 23 | 311 | 23 | 707 | 76.7 | 4.75 | 29 | 1.156 | 0.672 | 1.585 | 0.409 | 1.690 | 0.190 | 0.938 | 0.895 |
| 1525 | **Rps20** | Unchanged | P60867 | 5 | 50 | 5 | 119 | 13.4 | 9.94 | 29 | 0.853 | 0.747 | 0.736 | 0.554 | 0.904 | 0.828 | 0.814 | 0.567 |
| 1526 | **Ubqln1** | Down | Q8R317 | 10 | 84 | 7 | 582 | 61.9 | 4.94 | 29 | 0.986 | 0.929 | 1.575 | 0.191 | 0.659 | 0.019 | 2.390 | 0.047 |
| 1527 | **Pfkl** | UP | P12382 | 13 | 95 | 11 | 780 | 85.3 | 7.17 | 29 | 1.747 | 0.043 | 2.408 | 0.120 | 2.168 | 0.000 | 1.111 | 0.777 |
| 1528 | **Tagln** | Unchanged | P37804 | 6 | 30 | 6 | 201 | 22.6 | 8.81 | 29 | 0.863 | 0.812 | 1.968 | 0.407 | 0.920 | 0.889 | 2.139 | 0.377 |
| 1529 | **Pgp** | Unchanged | Q8CHP8 | 6 | 53 | 6 | 321 | 34.5 | 5.35 | 29 | 0.557 | 0.082 | 1.051 | 0.891 | 1.644 | 0.254 | 0.639 | 0.317 |
| 1530 | **Rpl35** | Unchanged | Q6ZWV7 | 4 | 48 | 4 | 123 | 14.5 | 11.05 | 29 | 1.237 | 0.685 | 0.564 | 0.166 | 1.130 | 0.818 | 0.499 | 0.290 |
| 1531 | **Gspt1** | Down | Q8R050 | 12 | 47 | 12 | 636 | 68.6 | 5.21 | 29 | 0.694 | 0.144 | 0.614 | 0.088 | 0.507 | 0.028 | 1.211 | 0.285 |
| 1532 | **Sec61a1** | UP | P61620 | 9 | 122 | 9 | 476 | 52.2 | 8.06 | 29 | 1.427 | 0.531 | 3.414 | 0.049 | 1.484 | 0.339 | 2.301 | 0.122 |
| 1533 | **Ndufaf3** | Unchanged | Q9JKL4 | 3 | 10 | 3 | 185 | 20.7 | 8.05 | 29 | 0.893 | 0.837 | 0.978 | 0.973 | 0.332 | 0.185 | 2.943 | 0.216 |
| 1534 | **Mrpl21** | UP | Q9D1N9 | 4 | 15 | 4 | 209 | 23.4 | 9.99 | 29 | 1.232 | 0.451 | 1.716 | 0.334 | 1.889 | 0.045 | 0.909 | 0.832 |
| 1535 | **Gm2056** | Unchanged | J3QQ02 | 2 | 13 | 1 | 144 | 16.6 | 5.35 | 29 | 0.540 | 0.050 | 0.542 | 0.109 | 0.718 | 0.276 | 0.755 | 0.506 |
| 1536 | **Mrpl11** | Unchanged | Q9CQF0 | 5 | 21 | 5 | 192 | 20.7 | 9.73 | 29 | 0.845 | 0.801 | 0.291 | 0.224 | 0.780 | 0.696 | 0.373 | 0.059 |
| 1537 | **M6pr** | Down | P24668 | 5 | 32 | 5 | 278 | 31.2 | 5.39 | 29 | 0.765 | 0.179 | 0.858 | 0.607 | 0.520 | 0.020 | 1.649 | 0.234 |
| 1538 | **Dbnl** | Unchanged | Q62418 | 8 | 69 | 8 | 436 | 48.7 | 4.92 | 29 | 0.686 | 0.037 | 1.710 | 0.445 | 1.519 | 0.404 | 1.126 | 0.860 |
| 1539 | **Rpl35a** | Unchanged | O55142 | 5 | 27 | 5 | 110 | 12.5 | 10.89 | 29 | 1.315 | 0.606 | 1.146 | 0.780 | 1.488 | 0.407 | 0.770 | 0.460 |
| 1540 | **Rpl13a** | Unchanged | P19253 | 6 | 39 | 6 | 203 | 23.4 | 11.02 | 24 | 0.744 | 0.572 | 0.489 | 0.203 | 0.467 | 0.187 | 1.048 | 0.919 |
| 1541 | **Cyp4f13** | UP | Q99N19 | 9 | 34 | 7 | 523 | 59.8 | 7.69 | 29 | 1.773 | 0.164 | 3.025 | 0.000 | 2.262 | 0.051 | 1.337 | 0.213 |
| 1542 | **Mrps35** | Unchanged | Q8BJZ4 | 5 | 17 | 5 | 320 | 36 | 8.59 | 29 | 1.255 | 0.521 | 1.032 | 0.946 | 0.834 | 0.629 | 1.237 | 0.685 |
| 1543 | **Srsf1** | Unchanged | Q6PDM2 | 9 | 54 | 9 | 248 | 27.7 | 10.36 | 29 | 0.993 | 0.970 | 1.066 | 0.809 | 1.126 | 0.749 | 0.946 | 0.899 |
| 1544 | **Slc9a3r1** | Unchanged | P70441 | 9 | 82 | 9 | 355 | 38.6 | 5.9 | 29 | 0.958 | 0.860 | 0.861 | 0.673 | 1.184 | 0.505 | 0.727 | 0.339 |
| 1545 | **Rap2a** | UP | Q80ZJ1 | 3 | 15 | 1 | 183 | 20.6 | 4.82 | 29 | 1.897 | 0.370 | 4.273 | 0.022 | 3.055 | 0.022 | 1.399 | 0.378 |
| 1546 | **Rps21** | Down | Q9CQR2 | 2 | 36 | 2 | 83 | 9.1 | 8.51 | 29 | 0.574 | 0.084 | 0.496 | 0.058 | 0.303 | 0.004 | 1.637 | 0.247 |
| 1547 | **U2af1** | Unchanged | Q9D883 | 4 | 23 | 4 | 239 | 27.8 | 8.81 | 29 | 1.644 | 0.482 | 1.707 | 0.371 | 1.541 | 0.517 | 1.108 | 0.863 |

| | | | | | | | | | | | | | | | | | | |
|---|---|---|---|---|---|---|---|---|---|---|---|---|---|---|---|---|---|---|
| 1548 | **Twf1** | Unchanged | Q91YR1 | 8 | 61 | 8 | 350 | 40.1 | 6.67 | 29 | 0.851 | 0.252 | 0.764 | 0.303 | 0.869 | 0.253 | 0.879 | 0.596 |
| 1549 | **Pabpc4** | Unchanged | Q6PHQ9 | 18 | 74 | 10 | 660 | 72.2 | 9.39 | 29 | 1.395 | 0.235 | 1.320 | 0.263 | 1.249 | 0.288 | 1.057 | 0.824 |
| 1550 | **Pitpna** | Down | P53810 | 6 | 43 | 5 | 271 | 31.9 | 6.37 | 29 | 0.821 | 0.201 | 0.619 | 0.027 | 0.995 | 0.977 | 0.623 | 0.028 |
| 1551 | **Atp6v1e1** | UP | P50518 | 6 | 58 | 6 | 226 | 26.1 | 8.43 | 29 | 1.390 | 0.327 | 2.198 | 0.009 | 1.098 | 0.683 | 2.003 | 0.024 |
| 1552 | **Ddc** | Down | O88533 | 8 | 81 | 8 | 480 | 53.8 | 6.6 | 29 | 0.745 | 0.271 | 0.774 | 0.440 | 0.589 | 0.022 | 1.314 | 0.465 |
| 1553 | **Igk-V9;Igkv14-111** | Unchanged | P01638 | 2 | 17 | 2 | 115 | 13 | 7.81 | 29 | 2.692 | 0.208 | 1.134 | 0.883 | 6.015 | 0.052 | 0.189 | 0.067 |
| 1554 | **Eef1d** | Unchanged | F6ZFU0 | 11 | 94 | 1 | 552 | 61.4 | 7.39 | 29 | 0.046 | 0.217 | 1.069 | 0.941 | 0.300 | 0.366 | 3.563 | 0.223 |
| 1555 | **Prps2** | Unchanged | Q9CS42 | 7 | 94 | 2 | 318 | 34.8 | 6.61 | 29 | 1.361 | 0.528 | 1.636 | 0.311 | 0.554 | 0.272 | 2.952 | 0.062 |
| 1556 | **Pyurf** | Unchanged | Q9D1C3 | 1 | 1 | 1 | 112 | 12.5 | 8.34 | 29 | 0.841 | 0.682 | 0.858 | 0.575 | 1.101 | 0.756 | 0.779 | 0.391 |
| 1557 | **Sgta** | Unchanged | Q8BJU0 | 7 | 20 | 7 | 315 | 34.3 | 5.06 | 29 | 1.132 | 0.725 | 2.682 | 0.091 | 0.689 | 0.390 | 3.891 | 0.042 |
| 1558 | **Ctsh** | Unchanged | P49935 | 6 | 82 | 6 | 333 | 37.1 | 8.4 | 29 | 1.118 | 0.806 | 2.273 | 0.101 | 1.257 | 0.660 | 1.808 | 0.218 |
| 1559 | **Ldah** | UP | A0A0F6AIX5 | 6 | 21 | 6 | 326 | 37.3 | 8.28 | 29 | 1.415 | 0.489 | 4.102 | 0.047 | 2.324 | 0.099 | 1.765 | 0.264 |
| 1560 | **Atp11c** | Unchanged | Q9QZW0 | 22 | 197 | 22 | 1129 | 129.2 | 7.3 | 29 | 0.842 | 0.415 | 0.715 | 0.058 | 1.311 | 0.580 | 0.545 | 0.290 |
| 1561 | **Prpsap2** | Unchanged | Q8R574 | 7 | 26 | 6 | 369 | 40.9 | 7.17 | 28 | 1.324 | 0.405 | 1.281 | 0.539 | 1.259 | 0.516 | 1.018 | 0.949 |
| 1562 | **Gypc** | Unchanged | Q78HU7 | 1 | 1 | 1 | 95 | 10.3 | 5.35 | 28 | 9.006 | 0.316 | 17.385 | 0.137 | 4.865 | 0.160 | 3.574 | 0.255 |
| 1563 | **Serping1** | UP | P97290 | 9 | 32 | 9 | 504 | 55.5 | 6.29 | 28 | 1.122 | 0.680 | 1.891 | 0.011 | 1.251 | 0.390 | 1.511 | 0.032 |
| 1564 | **Bche** | Unchanged | Q03311 | 8 | 79 | 8 | 603 | 68.4 | 7.25 | 28 | 1.378 | 0.382 | 1.073 | 0.870 | 0.993 | 0.986 | 1.081 | 0.817 |
| 1565 | **Scpep1** | Unchanged | Q920A5 | 9 | 66 | 9 | 452 | 50.9 | 5.66 | 28 | 0.807 | 0.309 | 0.941 | 0.850 | 1.261 | 0.340 | 0.746 | 0.379 |
| 1566 | **Acacb** | Unchanged | E9Q4Z2 | 41 | 262 | 34 | 2448 | 275.6 | 6.32 | 28 | 0.949 | 0.844 | 1.568 | 0.052 | 1.151 | 0.463 | 1.362 | 0.113 |
| 1567 | **Acss3** | Unchanged | Q14DH7 | 12 | 55 | 12 | 682 | 74.5 | 8.63 | 28 | 0.800 | 0.586 | 1.471 | 0.272 | 0.605 | 0.223 | 2.431 | 0.019 |
| 1568 | **Tpp1** | Unchanged | O89023 | 7 | 98 | 7 | 562 | 61.3 | 6.57 | 28 | 1.321 | 0.442 | 1.390 | 0.383 | 1.283 | 0.544 | 1.084 | 0.832 |
| 1569 | **Erlin2** | Unchanged | Q8BFZ9 | 7 | 32 | 5 | 340 | 37.8 | 5.5 | 28 | 0.749 | 0.373 | 0.771 | 0.408 | 0.976 | 0.924 | 0.790 | 0.410 |
| 1570 | **Ppm1b** | Unchanged | P36993 | 7 | 53 | 6 | 390 | 42.8 | 5.19 | 28 | 1.123 | 0.759 | 2.213 | 0.211 | 1.260 | 0.600 | 1.756 | 0.358 |
| 1571 | **Gabarapl1** | Unchanged | Q8R3R8 | 4 | 17 | 4 | 117 | 14 | 8.73 | 28 | 2.368 | 0.212 | 4.472 | 0.059 | 1.280 | 0.556 | 3.493 | 0.080 |
| 1572 | **Rpl34** | Down | Q9D1R9 | 4 | 37 | 4 | 117 | 13.3 | 11.47 | 28 | 0.326 | 0.003 | 0.871 | 0.709 | 0.211 | 0.000 | 4.131 | 0.077 |
| 1573 | **Mug2** | UP | P28666 | 30 | 271 | 12 | 1451 | 162.3 | 6.74 | 28 | 1.002 | 0.993 | 2.153 | 0.007 | 0.704 | 0.202 | 3.058 | 0.002 |
| 1574 | **Bpgm** | Unchanged | P15327 | 3 | 17 | 3 | 259 | 30 | 7.06 | 28 | 0.960 | 0.912 | 0.558 | 0.181 | 1.589 | 0.472 | 0.351 | 0.206 |
| 1575 | **Acsf3** | Down | Q3URE1 | 9 | 49 | 9 | 583 | 65 | 8.02 | 28 | 0.721 | 0.180 | 2.071 | 0.088 | 0.555 | 0.024 | 3.731 | 0.024 |
| 1576 | **Eif3g** | Unchanged | Q9Z1D1 | 6 | 39 | 6 | 320 | 35.6 | 5.9 | 18 | 0.982 | 0.932 | 1.351 | 0.409 | 1.094 | 0.746 | 1.235 | 0.583 |
| 1577 | **Czib** | Down | Q8BHG2 | 3 | 14 | 3 | 160 | 18 | 5.12 | 28 | 1.125 | 0.719 | 0.996 | 0.993 | 0.255 | 0.012 | 3.909 | 0.057 |
| 1578 | **Chil4** | UP | Q91Z98 | 7 | 13 | 5 | 402 | 44.9 | 6.19 | 28 | 1.996 | 0.495 | 6.592 | 0.064 | 4.947 | 0.025 | 1.332 | 0.588 |
| 1579 | **Serpina1e** | Unchanged | Q00898 | 9 | 143 | 3 | 413 | 45.9 | 5.73 | 28 | 0.614 | 0.113 | 0.929 | 0.825 | 0.887 | 0.644 | 1.048 | 0.891 |
| 1580 | **Naglu** | Unchanged | O88325 | 16 | 59 | 16 | 739 | 82.5 | 6.61 | 28 | 1.126 | 0.459 | 1.245 | 0.236 | 1.251 | 0.291 | 0.996 | 0.985 |
| 1581 | **Acp6** | Unchanged | Q8BP40 | 7 | 46 | 7 | 418 | 47.6 | 7.72 | 28 | 0.891 | 0.796 | 0.872 | 0.775 | 1.072 | 0.882 | 0.814 | 0.664 |
| 1582 | **Prps1** | UP | Q9D7G0 | 7 | 113 | 2 | 318 | 34.8 | 6.98 | 28 | 1.462 | 0.164 | 1.797 | 0.004 | 1.590 | 0.016 | 1.130 | 0.359 |
| 1583 | **Eefsec** | UP | Q9JHW4 | 8 | 61 | 8 | 583 | 63.5 | 8.29 | 28 | 0.834 | 0.641 | 1.756 | 0.033 | 2.594 | 0.003 | 0.677 | 0.104 |
| 1584 | **Cacybp** | Unchanged | Q9CXW3 | 4 | 19 | 4 | 229 | 26.5 | 7.87 | 28 | 1.095 | 0.874 | 1.613 | 0.440 | 1.649 | 0.296 | 0.978 | 0.964 |
| 1585 | **Hdhd5** | UP | Q91WM2 | 7 | 49 | 7 | 419 | 46.3 | 7.88 | 28 | 1.419 | 0.412 | 4.482 | 0.040 | 1.146 | 0.588 | 3.910 | 0.046 |
| 1586 | **Gnaq** | UP | P21279 | 7 | 24 | 4 | 359 | 42.1 | 5.68 | 28 | 0.980 | 0.925 | 1.596 | 0.039 | 1.655 | 0.007 | 0.964 | 0.801 |
| 1587 | **Golph3** | Unchanged | Q9CRA5 | 6 | 41 | 6 | 298 | 33.7 | 6.44 | 28 | 2.303 | 0.345 | 2.873 | 0.053 | 1.047 | 0.936 | 2.745 | 0.086 |
| 1588 | **Snx3** | Down | Q78ZM0 | 6 | 27 | 4 | 162 | 18.8 | 8.66 | 28 | 0.607 | 0.090 | 0.446 | 0.006 | 0.458 | 0.005 | 0.975 | 0.943 |

| | | | | | | | | | | | | | | | | | | |
|---|---|---|---|---|---|---|---|---|---|---|---|---|---|---|---|---|---|---|
| 1589 | **Eif3c** | Unchanged | Q8R1B4 | 19 | 185 | 19 | 911 | 105.5 | 5.78 | 28 | 1.313 | 0.152 | 1.939 | 0.089 | 1.261 | 0.070 | 1.537 | 0.200 |
| 1590 | **Bcap31** | Unchanged | Q61335 | 4 | 85 | 4 | 245 | 27.9 | 8.7 | 28 | 1.180 | 0.703 | 1.005 | 0.992 | 1.304 | 0.531 | 0.771 | 0.633 |
| 1591 | **Snrpd1** | Unchanged | P62315 | 2 | 59 | 2 | 119 | 13.3 | 11.56 | 28 | 0.962 | 0.841 | 1.583 | 0.124 | 1.435 | 0.125 | 1.103 | 0.679 |
| 1592 | **Ptpn6** | Unchanged | P29351 | 9 | 46 | 9 | 595 | 67.5 | 7.81 | 28 | 0.993 | 0.969 | 1.140 | 0.562 | 1.147 | 0.594 | 0.994 | 0.981 |
| 1593 | **Stard5** | UP | Q9EPQ7 | 3 | 20 | 3 | 213 | 23.9 | 6.39 | 28 | 3.320 | 0.040 | 3.015 | 0.093 | 2.494 | 0.146 | 1.209 | 0.712 |
| 1594 | **Steap4** | Down | Q923B6 | 9 | 70 | 9 | 470 | 53 | 9.17 | 28 | 0.643 | 0.228 | 0.715 | 0.399 | 0.342 | 0.028 | 2.094 | 0.168 |
| 1595 | **Ybx1** | Unchanged | P62960 | 7 | 49 | 3 | 322 | 35.7 | 9.88 | 28 | 0.782 | 0.260 | 0.648 | 0.208 | 0.693 | 0.156 | 0.935 | 0.862 |
| 1596 | **Atp6v0d1** | UP | P51863 | 6 | 31 | 6 | 351 | 40.3 | 5 | 28 | 1.293 | 0.468 | 1.765 | 0.045 | 1.070 | 0.766 | 1.650 | 0.056 |
| 1597 | **Atp1b1** | Unchanged | P14094 | 6 | 64 | 6 | 304 | 35.2 | 8.65 | 28 | 1.465 | 0.234 | 1.921 | 0.178 | 1.090 | 0.745 | 1.762 | 0.230 |
| 1598 | **Cyp4a14** | Unchanged | O35728 | 11 | 60 | 7 | 507 | 58.7 | 8.78 | 28 | 0.962 | 0.953 | 1.366 | 0.638 | 2.219 | 0.086 | 0.616 | 0.243 |
| 1599 | **Cd81** | Unchanged | P35762 | 3 | 71 | 3 | 236 | 25.8 | 5.83 | 28 | 1.264 | 0.423 | 1.404 | 0.118 | 1.057 | 0.846 | 1.328 | 0.236 |
| 1600 | **Dync1li1** | Down | Q8R1Q8 | 8 | 21 | 8 | 523 | 56.6 | 6.42 | 28 | 0.734 | 0.314 | 0.940 | 0.835 | 0.272 | 0.004 | 3.460 | 0.014 |
| 1601 | **Ddx3y** | Unchanged | Q62095 | 14 | 128 | 2 | 658 | 73.4 | 7.53 | 28 | 1.858 | 0.142 | 2.547 | 0.151 | 1.147 | 0.764 | 2.220 | 0.185 |
| 1602 | **Acad10** | Unchanged | Q8K370 | 21 | 102 | 12 | 1069 | 118.9 | 8.24 | 28 | 1.660 | 0.241 | 2.587 | 0.281 | 0.955 | 0.898 | 2.710 | 0.269 |
| 1603 | **Mrps17** | Down | Q9CQE3 | 2 | 8 | 2 | 120 | 13.4 | 9.89 | 28 | 1.373 | 0.640 | 1.753 | 0.544 | 0.075 | 0.047 | 23.378 | 0.174 |
| 1604 | **Efhd2** | Unchanged | Q9D8Y0 | 7 | 41 | 7 | 240 | 26.8 | 5.06 | 28 | 0.908 | 0.658 | 1.457 | 0.074 | 1.343 | 0.098 | 1.086 | 0.624 |
| 1605 | **Dnaja3** | Unchanged | Q99M87 | 9 | 58 | 9 | 480 | 52.4 | 9.22 | 28 | 0.873 | 0.736 | 0.841 | 0.706 | 0.616 | 0.220 | 1.367 | 0.517 |
| 1606 | **Mrps25** | Down | Q9D125 | 3 | 12 | 3 | 171 | 19.9 | 9.23 | 27 | 0.717 | 0.439 | 0.578 | 0.342 | 0.188 | 0.004 | 3.067 | 0.326 |
| 1607 | **Chil3** | Unchanged | O35744 | 7 | 32 | 5 | 398 | 44.4 | 5.69 | 27 | 0.469 | 0.216 | 1.591 | 0.471 | 0.437 | 0.149 | 3.639 | 0.146 |
| 1608 | **Dpp4** | Unchanged | P28843 | 16 | 157 | 16 | 760 | 87.4 | 6.42 | 27 | 0.815 | 0.555 | 8.657 | 0.184 | 0.390 | 0.064 | 22.196 | 0.155 |
| 1609 | **Nap1l1** | Unchanged | P28656 | 5 | 36 | 5 | 391 | 45.3 | 4.46 | 27 | 1.047 | 0.920 | 1.069 | 0.865 | 1.160 | 0.685 | 0.921 | 0.796 |
| 1610 | **Ufl1** | Unchanged | Q8CCJ3 | 12 | 87 | 12 | 793 | 89.5 | 6.67 | 27 | 0.887 | 0.725 | 0.901 | 0.765 | 0.694 | 0.329 | 1.299 | 0.258 |
| 1611 | **Srpra** | Unchanged | Q9DBG7 | 11 | 53 | 11 | 636 | 69.6 | 8.95 | 26 | 1.228 | 0.512 | 1.594 | 0.183 | 0.993 | 0.982 | 1.606 | 0.239 |
| 1612 | **Sdsl** | Unchanged | Q8R238 | 3 | 6 | 3 | 329 | 34.7 | 6.43 | 27 | 1.849 | 0.350 | 2.884 | 0.211 | 2.104 | 0.229 | 1.371 | 0.642 |
| 1613 | **Igkv4-57** | Unchanged | A0A0A6YYE7 | 2 | 4 | 2 | 117 | 12.7 | 8.91 | 27 | 1.637 | 0.642 | 0.266 | 0.266 | 0.140 | 0.196 | 1.896 | 0.214 |
| 1614 | **Cd38** | Unchanged | P56528 | 5 | 29 | 5 | 304 | 34.4 | 8.29 | 27 | 1.213 | 0.597 | 2.560 | 0.103 | 0.678 | 0.360 | 3.778 | 0.054 |
| 1615 | **Atp5mf** | Unchanged | P56135 | 3 | 40 | 3 | 88 | 10.3 | 9.95 | 27 | 1.062 | 0.921 | 1.070 | 0.926 | 0.876 | 0.849 | 1.222 | 0.809 |
| 1616 | **Pcyt1a** | Unchanged | P49586 | 6 | 17 | 6 | 367 | 41.6 | 7.03 | 27 | 1.455 | 0.018 | 0.889 | 0.591 | 0.944 | 0.824 | 0.942 | 0.847 |
| 1617 | **Stard4** | Unchanged | Q99JV5 | 3 | 7 | 3 | 224 | 25.6 | 5.67 | 27 | 1.374 | 0.408 | 2.342 | 0.288 | 0.641 | 0.315 | 3.654 | 0.171 |
| 1618 | **Lmna** | Unchanged | P48678 | 16 | 79 | 16 | 665 | 74.2 | 6.98 | 27 | 1.291 | 0.436 | 0.893 | 0.722 | 1.136 | 0.668 | 0.786 | 0.490 |
| 1619 | **Sec24c** | Unchanged | G3X972 | 18 | 67 | 18 | 1096 | 118.5 | 6.84 | 27 | 1.162 | 0.550 | 1.199 | 0.272 | 1.030 | 0.891 | 1.164 | 0.424 |
| 1620 | **Fhl1** | Unchanged | P97447 | 5 | 11 | 5 | 280 | 31.9 | 8.37 | 27 | 0.367 | 0.368 | 0.616 | 0.588 | 0.410 | 0.402 | 1.503 | 0.325 |
| 1621 | **As3mt** | Unchanged | Q91WU5 | 7 | 34 | 7 | 376 | 41.8 | 5.86 | 27 | 1.050 | 0.897 | 2.363 | 0.206 | 2.874 | 0.231 | 0.822 | 0.771 |
| 1622 | **Uncharacterizedprotein** | UP | A0A338P6I9 | 1 | 3 | 1 | 59 | 6.8 | 8.63 | 27 | 1.747 | 0.458 | 2.891 | 0.116 | 2.710 | 0.008 | 1.067 | 0.867 |
| 1623 | **Coro1a** | UP | O89053 | 7 | 58 | 7 | 461 | 51 | 6.48 | 27 | 1.777 | 0.383 | 2.116 | 0.097 | 3.185 | 0.008 | 0.665 | 0.216 |
| 1624 | **Aox1** | UP | G3X8P9 | 26 | 83 | 26 | 1333 | 146.6 | 6.93 | 27 | 0.912 | 0.667 | 1.624 | 0.069 | 1.645 | 0.045 | 0.987 | 0.955 |
| 1625 | **Rab8a** | Down | P55258 | 6 | 39 | 2 | 207 | 23.7 | 9.07 | 27 | 0.693 | 0.599 | 0.106 | 0.013 | 0.146 | 0.016 | 0.727 | 0.455 |
| 1626 | **Cyp8b1** | Unchanged | O88962 | 12 | 75 | 12 | 500 | 57.7 | 8.84 | 27 | 1.184 | 0.534 | 1.509 | 0.253 | 0.697 | 0.164 | 2.167 | 0.071 |
| 1627 | **Banf1** | UP | O54962 | 1 | 3 | 1 | 89 | 10.1 | 6.09 | 27 | 8.621 | 0.124 | 26.601 | 0.268 | 22.087 | 0.025 | 1.204 | 0.848 |
| 1628 | **Cops3** | Unchanged | O88543 | 7 | 37 | 7 | 423 | 47.8 | 6.65 | 27 | 0.830 | 0.687 | 0.711 | 0.452 | 1.248 | 0.733 | 0.570 | 0.422 |
| 1629 | **Arl3** | Unchanged | Q9WUL7 | 3 | 13 | 3 | 182 | 20.5 | 7.24 | 27 | 1.379 | 0.560 | 5.158 | 0.085 | 1.706 | 0.264 | 3.024 | 0.141 |

| | | | | | | | | | | | | | | | | | | |
|---|---|---|---|---|---|---|---|---|---|---|---|---|---|---|---|---|---|---|
| 1630 | **Crip2** | Unchanged | Q9DCT8 | 3 | 26 | 3 | 208 | 22.7 | 8.63 | 27 | 1.109 | 0.899 | 0.622 | 0.564 | 0.349 | 0.293 | 1.780 | 0.462 |
| 1631 | **Sfxn5** | Unchanged | Q925N0 | 4 | 16 | 4 | 342 | 37.3 | 9.44 | 27 | 1.669 | 0.273 | 1.260 | 0.603 | 1.489 | 0.307 | 0.846 | 0.705 |
| 1632 | **Flna** | Unchanged | Q8BTM8 | 40 | 236 | 37 | 2647 | 281 | 6.04 | 27 | 1.292 | 0.540 | 1.815 | 0.116 | 1.780 | 0.105 | 1.020 | 0.956 |
| 1633 | **Enpep** | UP | P16406 | 20 | 137 | 20 | 945 | 107.9 | 5.44 | 27 | 1.301 | 0.228 | 1.798 | 0.069 | 2.227 | 0.037 | 0.808 | 0.480 |
| 1634 | **Cygb** | UP | Q9CX80 | 3 | 19 | 3 | 190 | 21.5 | 6.8 | 27 | 2.205 | 0.249 | 2.830 | 0.066 | 3.711 | 0.027 | 0.763 | 0.504 |
| 1635 | **Tstd3** | UP | Q9D0B5 | 3 | 14 | 3 | 157 | 17.3 | 8.13 | 27 | 3.140 | 0.144 | 3.063 | 0.192 | 4.070 | 0.005 | 0.752 | 0.545 |
| 1636 | **Igkv8-19** | Unchanged | A0A140T8M4 | 2 | 2 | 1 | 101 | 11.1 | 5.25 | 27 | 0.496 | 0.435 | 0.118 | 0.073 | 0.242 | 0.118 | 0.487 | 0.152 |
| 1637 | **Bhmt2** | Unchanged | Q91WS4 | 8 | 116 | 4 | 363 | 39.8 | 6.54 | 27 | 0.986 | 0.965 | 3.817 | 0.228 | 7.010 | 0.147 | 0.544 | 0.480 |
| 1638 | **Tmem97** | Unchanged | Q8VD00 | 3 | 9 | 3 | 176 | 20.8 | 9.28 | 27 | 1.108 | 0.864 | 1.140 | 0.819 | 1.039 | 0.940 | 1.097 | 0.841 |
| 1639 | **Ipo7** | Unchanged | Q9EPL8 | 15 | 83 | 15 | 1038 | 119.4 | 4.82 | 27 | 0.948 | 0.836 | 1.834 | 0.141 | 0.955 | 0.863 | 1.920 | 0.104 |
| 1640 | **Xpnpep3** | Unchanged | B7ZMP1 | 6 | 32 | 6 | 506 | 56.6 | 7.68 | 27 | 0.792 | 0.452 | 1.177 | 0.532 | 0.796 | 0.447 | 1.478 | 0.016 |
| 1641 | **Hexb** | Unchanged | P20060 | 9 | 41 | 9 | 536 | 61.1 | 8.12 | 27 | 1.071 | 0.877 | 1.537 | 0.287 | 1.070 | 0.747 | 1.437 | 0.301 |
| 1642 | **Lgmn** | Unchanged | O89017 | 9 | 100 | 9 | 435 | 49.3 | 6.39 | 27 | 0.652 | 0.235 | 2.484 | 0.231 | 0.456 | 0.061 | 5.444 | 0.108 |
| 1643 | **Cnn3** | Unchanged | Q9DAW9 | 3 | 18 | 3 | 330 | 36.4 | 5.72 | 27 | 1.290 | 0.680 | 2.121 | 0.266 | 1.108 | 0.802 | 1.914 | 0.315 |
| 1644 | **Chchd4** | Unchanged | Q8VEA4 | 2 | 23 | 2 | 139 | 15.5 | 4.32 | 27 | 1.071 | 0.833 | 0.869 | 0.677 | 0.996 | 0.993 | 0.873 | 0.793 |
| 1645 | **Emc2** | Unchanged | Q9CRD2 | 4 | 16 | 4 | 297 | 34.9 | 6.81 | 27 | 1.232 | 0.736 | 2.384 | 0.264 | 1.587 | 0.450 | 1.502 | 0.464 |
| 1646 | **S100a1** | Unchanged | P56565 | 4 | 60 | 4 | 94 | 10.5 | 4.5 | 27 | 1.635 | 0.300 | 1.641 | 0.146 | 1.632 | 0.262 | 1.005 | 0.984 |
| 1647 | **Cse1l** | Unchanged | Q9ERK4 | 16 | 145 | 16 | 971 | 110.4 | 5.77 | 26 | 1.170 | 0.527 | 1.379 | 0.403 | 2.787 | 0.258 | 0.495 | 0.376 |
| 1648 | **Spryd4** | Unchanged | Q91WK1 | 4 | 13 | 4 | 207 | 23.3 | 9.45 | 27 | 0.851 | 0.722 | 1.942 | 0.055 | 1.027 | 0.955 | 1.891 | 0.036 |
| 1649 | **Nmral1** | Unchanged | Q8K2T1 | 5 | 32 | 5 | 309 | 34.4 | 6.86 | 27 | 1.060 | 0.915 | 1.217 | 0.750 | 0.917 | 0.889 | 1.327 | 0.664 |
| 1650 | **Tpmtiso** | Unchanged | A0A286YCA5 | 1 | 1 | 1 | 83 | 9.9 | 10.4 | 27 | 2.989 | 0.144 | 2.817 | 0.206 | 1.391 | 0.642 | 2.026 | 0.362 |
| 1651 | **Tango2** | Unchanged | P54797 | 3 | 19 | 3 | 276 | 30.9 | 5.1 | 26 | 1.217 | 0.677 | 0.924 | 0.854 | 2.104 | 0.221 | 0.439 | 0.192 |
| 1652 | **Usp9x** | Unchanged | G3UY52 | 2 | 3 | 1 | 174 | 20.5 | 6.64 | 26 | 1.204 | 0.696 | 1.241 | 0.758 | 0.302 | 0.144 | 4.116 | 0.175 |
| 1653 | **Fdx2** | Unchanged | Q9CPW2 | 3 | 7 | 3 | 174 | 18.8 | 6.04 | 26 | 0.694 | 0.651 | 0.462 | 0.426 | 2.803 | 0.172 | 0.165 | 0.052 |
| 1654 | **Metap1** | Unchanged | Q8BP48 | 6 | 33 | 6 | 386 | 43.2 | 7.17 | 26 | 1.141 | 0.647 | 1.866 | 0.330 | 1.533 | 0.114 | 1.217 | 0.693 |
| 1655 | **Vps36** | UP | Q91XD6 | 7 | 25 | 7 | 386 | 43.7 | 7.15 | 26 | 2.473 | 0.058 | 3.875 | 0.006 | 1.063 | 0.858 | 3.645 | 0.007 |
| 1656 | **Gps1** | Unchanged | Q99LD4 | 7 | 43 | 7 | 471 | 53.4 | 6.84 | 26 | 0.531 | 0.192 | 0.487 | 0.168 | 0.532 | 0.190 | 0.915 | 0.707 |
| 1657 | **Lzic** | UP | Q8K3C3 | 2 | 7 | 2 | 190 | 21.5 | 4.96 | 26 | 1.599 | 0.046 | 1.768 | 0.100 | 1.757 | 0.061 | 1.006 | 0.983 |
| 1658 | **Snrpg** | Unchanged | P62309 | 2 | 18 | 2 | 76 | 8.5 | 8.88 | 26 | 2.739 | 0.266 | 6.073 | 0.190 | 4.322 | 0.114 | 1.405 | 0.672 |
| 1659 | **Ccs** | Unchanged | Q9WU84 | 3 | 27 | 3 | 274 | 28.9 | 6.1 | 26 | 7.085 | 0.355 | 10.329 | 0.185 | 4.363 | 0.085 | 2.367 | 0.396 |
| 1660 | **Mrpl53** | Unchanged | Q9D1H8 | 2 | 7 | 2 | 118 | 12.7 | 9.52 | 26 | 0.772 | 0.577 | 41.228 | 0.189 | 132.430 | 0.057 | 0.311 | 0.202 |
| 1661 | **Cdk5rap3** | Unchanged | Q99LM2 | 8 | 42 | 8 | 503 | 57 | 4.83 | 26 | 0.858 | 0.754 | 1.037 | 0.935 | 0.884 | 0.775 | 1.173 | 0.689 |
| 1662 | **Ndufa11** | Unchanged | Q9D8B4 | 3 | 29 | 3 | 141 | 15 | 8.35 | 26 | 1.276 | 0.740 | 2.244 | 0.345 | 1.123 | 0.869 | 1.998 | 0.410 |
| 1663 | **Nop56** | Unchanged | Q9D6Z1 | 10 | 46 | 10 | 580 | 64.4 | 9.14 | 26 | 0.544 | 0.160 | 0.516 | 0.219 | 0.396 | 0.078 | 1.303 | 0.621 |
| 1664 | **Eps8l2** | UP | Q99K30 | 11 | 51 | 11 | 729 | 82.2 | 7.18 | 26 | 0.931 | 0.805 | 1.812 | 0.015 | 0.900 | 0.667 | 2.013 | 0.018 |
| 1665 | **Ctsf** | Down | Q9R013 | 7 | 29 | 7 | 462 | 51.6 | 6.55 | 26 | 0.566 | 0.142 | 0.517 | 0.030 | 0.578 | 0.156 | 0.895 | 0.808 |
| 1666 | **Grcc10** | Unchanged | O35127 | 2 | 7 | 2 | 126 | 13.2 | 5.14 | 26 | #DIV/0! | #DIV/0! | #DIV/0! | #DIV/0! | #DIV/0! | #DIV/0! | #DIV/0! | #DIV/0! |
| 1667 | **Tnpo1** | Unchanged | Q8BFY9 | 12 | 63 | 10 | 898 | 102.3 | 4.98 | 26 | 0.934 | 0.750 | 1.007 | 0.981 | 0.937 | 0.781 | 1.075 | 0.809 |
| 1668 | **Deptor** | Down | Q570Y9 | 8 | 31 | 8 | 409 | 46.1 | 7.91 | 26 | 1.582 | 0.284 | 0.530 | 0.038 | 0.838 | 0.488 | 0.632 | 0.202 |
| 1669 | **Eif3k** | Unchanged | Q9DBZ5 | 5 | 36 | 5 | 218 | 25.1 | 4.93 | 26 | 0.676 | 0.273 | 0.711 | 0.443 | 0.981 | 0.938 | 0.725 | 0.405 |
| 1670 | **Sult1b1** | Unchanged | Q9QWG7 | 6 | 64 | 6 | 299 | 34.9 | 6.8 | 26 | 1.600 | 0.151 | 0.659 | 0.347 | 1.985 | 0.180 | 0.332 | 0.065 |

| | | | | | | | | | | | | | | | | | | |
|---|---|---|---|---|---|---|---|---|---|---|---|---|---|---|---|---|---|---|
| 1671 | **Igkv6-23** | Unchanged | A0A140T8N5 | 2 | 2 | 1 | 115 | 12.8 | 5.73 | 26 | 0.330 | 0.203 | 5.530 | 0.415 | 4.153 | 0.053 | 1.332 | 0.806 |
| 1672 | **Lum** | UP | P51885 | 7 | 45 | 7 | 338 | 38.2 | 6.43 | 26 | 0.583 | 0.060 | 1.892 | 0.148 | 1.826 | 0.019 | 1.036 | 0.915 |
| 1673 | **Ufsp2** | Unchanged | Q99K23 | 6 | 11 | 6 | 461 | 52.5 | 6.76 | 26 | 1.113 | 0.793 | 2.860 | 0.114 | 3.288 | 0.291 | 0.870 | 0.854 |
| 1674 | **Mrpl23** | Down | O35972 | 3 | 11 | 3 | 146 | 17.1 | 9.76 | 26 | 0.518 | 0.031 | 0.323 | 0.005 | 0.093 | 0.001 | 3.461 | 0.004 |
| 1675 | **Psap** | Unchanged | Q8BFQ1 | 11 | 85 | 11 | 554 | 61 | 5.24 | 26 | 0.649 | 0.391 | 0.761 | 0.641 | 0.771 | 0.632 | 0.986 | 0.984 |
| 1676 | **Ddx19a** | Unchanged | Q61655 | 7 | 31 | 7 | 478 | 53.9 | 6.67 | 26 | 5.237 | 0.081 | 7.763 | 0.256 | 3.438 | 0.109 | 2.258 | 0.468 |
| 1677 | **Prelp** | Unchanged | Q9JK53 | 8 | 46 | 8 | 378 | 43.3 | 9.54 | 26 | 1.017 | 0.936 | 2.202 | 0.103 | 2.638 | 0.194 | 0.835 | 0.750 |
| 1678 | **Nudc** | Unchanged | O35685 | 5 | 24 | 5 | 332 | 38.3 | 5.26 | 15 | 1.200 | 0.790 | 1.833 | 0.410 | 1.195 | 0.764 | 1.534 | 0.521 |
| 1679 | **Rab9a** | Unchanged | Q9R0M6 | 3 | 11 | 3 | 201 | 22.9 | 5.66 | 26 | 1.155 | 0.722 | 2.558 | 0.393 | 1.269 | 0.691 | 2.016 | 0.487 |
| 1680 | **Ube2v1** | Unchanged | Q9CZY3 | 3 | 23 | 3 | 147 | 16.3 | 7.96 | 26 | 1.100 | 0.797 | 0.731 | 0.425 | 0.925 | 0.881 | 0.791 | 0.622 |
| 1681 | **Sf3b1** | Unchanged | Q99NB9 | 20 | 109 | 20 | 1304 | 145.7 | 7.09 | 26 | 1.058 | 0.808 | 2.107 | 0.155 | 1.301 | 0.596 | 1.619 | 0.387 |
| 1682 | **Eif3a** | Unchanged | P23116 | 30 | 304 | 30 | 1344 | 161.8 | 6.77 | 25 | 1.166 | 0.799 | 0.596 | 0.505 | 0.699 | 0.602 | 0.854 | 0.834 |
| 1683 | **Rpl14** | Down | Q9CR57 | 5 | 74 | 5 | 217 | 23.5 | 11.02 | 26 | 0.523 | 0.282 | 0.084 | 0.043 | 1.096 | 0.811 | 0.077 | 0.000 |
| 1684 | **Tmem170b** | Unchanged | P86050 | 1 | 3 | 1 | 132 | 14.4 | 10.8 | 26 | 1.192 | 0.895 | 0.151 | 0.394 | 0.732 | 0.800 | 0.206 | 0.181 |
| 1685 | **Puf60** | Unchanged | Q3UEB3 | 8 | 39 | 8 | 564 | 60.2 | 5.29 | 26 | 0.753 | 0.406 | 0.943 | 0.893 | 0.634 | 0.229 | 1.487 | 0.370 |
| 1686 | **Macrod1** | Unchanged | Q922B1 | 4 | 25 | 4 | 323 | 35.3 | 8.85 | 26 | 0.659 | 0.430 | 0.920 | 0.843 | 1.446 | 0.264 | 0.637 | 0.101 |
| 1687 | **Anxa3** | Unchanged | O35639 | 6 | 40 | 5 | 323 | 36.4 | 5.76 | 26 | 0.387 | 0.418 | 1.846 | 0.355 | 2.140 | 0.182 | 0.863 | 0.628 |
| 1688 | **Pdlim1** | Down | O70400 | 7 | 25 | 7 | 327 | 35.8 | 6.84 | 26 | 0.851 | 0.684 | 0.440 | 0.012 | 0.325 | 0.002 | 1.352 | 0.257 |
| 1689 | **Ube2a** | UP | Q9Z255 | 1 | 2 | 1 | 152 | 17.3 | 5.15 | 26 | 1.691 | 0.379 | 3.134 | 0.140 | 5.504 | 0.015 | 0.569 | 0.252 |
| 1690 | **Rhog** | Down | P84096 | 3 | 18 | 3 | 191 | 21.3 | 8.12 | 26 | 0.791 | 0.300 | 1.284 | 0.631 | 0.473 | 0.008 | 2.713 | 0.187 |
| 1691 | **Ighv5-17** | UP | A0A075B5R1 | 2 | 3 | 2 | 117 | 12.9 | 8.28 | 26 | 7.431 | 0.165 | 27.956 | 0.004 | 3.298 | 0.256 | 8.478 | 0.007 |
| 1692 | **St13** | Unchanged | Q99L47 | 9 | 92 | 9 | 371 | 41.6 | 5.26 | 26 | 0.741 | 0.259 | 0.597 | 0.130 | 0.977 | 0.918 | 0.612 | 0.043 |
| 1693 | **Itih4** | Unchanged | A6X935 | 17 | 124 | 17 | 942 | 104.6 | 6.4 | 26 | 1.084 | 0.824 | 1.381 | 0.348 | 0.826 | 0.547 | 1.673 | 0.181 |
| 1694 | **Srp9** | Unchanged | P49962 | 2 | 9 | 2 | 86 | 10.2 | 7.99 | 13 | 0.531 | 0.389 | 1.019 | 0.979 | 1.064 | 0.895 | 0.957 | 0.938 |
| 1695 | **Tmem30a** | Unchanged | Q8VEK0 | 7 | 54 | 7 | 364 | 41 | 8.37 | 26 | 2.620 | 0.291 | 1.141 | 0.745 | 1.573 | 0.083 | 0.725 | 0.309 |
| 1696 | **Ssb** | UP | P32067 | 8 | 77 | 8 | 415 | 47.7 | 9.77 | 26 | 1.138 | 0.701 | 1.846 | 0.078 | 2.020 | 0.014 | 0.914 | 0.613 |
| 1697 | **Copg2** | Unchanged | Q9QXK3 | 13 | 70 | 11 | 871 | 97.6 | 5.8 | 25 | 1.152 | 0.647 | 1.051 | 0.907 | 1.139 | 0.607 | 0.922 | 0.841 |
| 1698 | **Gmppa** | Down | Q922H4 | 8 | 27 | 8 | 420 | 46.2 | 7.62 | 25 | 0.442 | 0.018 | 1.062 | 0.880 | 0.627 | 0.441 | 1.693 | 0.456 |
| 1699 | **Gosr2** | Unchanged | O35166 | 4 | 8 | 4 | 212 | 24.7 | 8.21 | 25 | 1.009 | 0.976 | 1.331 | 0.620 | 1.796 | 0.084 | 0.741 | 0.503 |
| 1700 | **Srp14** | Unchanged | P16254 | 3 | 17 | 3 | 110 | 12.5 | 10.17 | 13 | 1.169 | 0.764 | 0.495 | 0.292 | 1.138 | 0.829 | 0.435 | 0.234 |
| 1701 | **Stbd1** | Unchanged | Q8C7E7 | 6 | 45 | 6 | 338 | 36.1 | 5.99 | 25 | 1.141 | 0.809 | 0.472 | 0.235 | 0.402 | 0.177 | 1.174 | 0.649 |
| 1702 | **2210016F16Rik** | Unchanged | G3X8U3 | 6 | 13 | 6 | 338 | 38.6 | 5.64 | 25 | 0.716 | 0.215 | 1.581 | 0.086 | 1.109 | 0.751 | 1.426 | 0.299 |
| 1703 | **Tmpo** | Unchanged | Q61029 | 7 | 77 | 7 | 452 | 50.3 | 9.45 | 23 | 0.701 | 0.316 | 0.969 | 0.947 | 0.780 | 0.474 | 1.242 | 0.681 |
| 1704 | **Ist1** | Unchanged | Q9CX00 | 4 | 10 | 4 | 362 | 39.4 | 5.44 | 25 | 1.111 | 0.799 | 1.476 | 0.491 | 1.155 | 0.711 | 1.278 | 0.609 |
| 1705 | **Crkl** | UP | P47941 | 5 | 18 | 5 | 303 | 33.8 | 6.74 | 25 | 2.295 | 0.066 | 2.312 | 0.031 | 1.849 | 0.213 | 1.250 | 0.546 |
| 1706 | **Kctd12** | Unchanged | A0A0R4J2B2 | 6 | 12 | 6 | 327 | 35.9 | 5.81 | 25 | 0.874 | 0.769 | 7.742 | 0.294 | 1.032 | 0.946 | 7.498 | 0.296 |
| 1707 | **Pepd** | UP | Q11136 | 8 | 83 | 8 | 493 | 55 | 5.78 | 25 | 1.069 | 0.634 | 1.549 | 0.027 | 1.059 | 0.713 | 1.463 | 0.028 |
| 1708 | **Magoh** | Unchanged | P61327 | 3 | 14 | 3 | 146 | 17.2 | 6.11 | 25 | 0.751 | 0.429 | 1.080 | 0.856 | 0.566 | 0.142 | 1.908 | 0.185 |
| 1709 | **Galk2** | UP | Q68FH4 | 5 | 15 | 5 | 458 | 50.5 | 6.9 | 25 | 1.233 | 0.571 | 2.410 | 0.238 | 2.536 | 0.019 | 0.950 | 0.917 |
| 1710 | **Pgam2** | Unchanged | O70250 | 6 | 22 | 3 | 253 | 28.8 | 8.5 | 25 | 0.160 | 0.190 | 0.893 | 0.887 | 1.224 | 0.776 | 0.730 | 0.626 |
| 1711 | **Apoo** | Unchanged | Q9DCZ4 | 3 | 14 | 3 | 198 | 22.6 | 9.25 | 25 | 1.100 | 0.765 | 0.917 | 0.870 | 1.506 | 0.144 | 0.609 | 0.267 |

| | | | | | | | | | | | | | | | | | | |
|---|---|---|---|---|---|---|---|---|---|---|---|---|---|---|---|---|---|---|
| 1712 | **Hnrnpm** | Unchanged | Q9D0E1 | 13 | 108 | 13 | 729 | 77.6 | 8.63 | 25 | 1.481 | 0.369 | 2.452 | 0.129 | 1.311 | 0.249 | 1.870 | 0.224 |
| 1713 | **Hnrnpul2** | Unchanged | Q00PI9 | 15 | 66 | 15 | 745 | 84.9 | 4.89 | 25 | 1.238 | 0.573 | 1.736 | 0.302 | 1.983 | 0.128 | 0.875 | 0.776 |
| 1714 | **Atp6v1h** | Unchanged | A0A0A6YWP6 | 7 | 24 | 1 | 441 | 50.8 | 6.7 | 25 | 0.899 | 0.855 | 0.175 | 0.075 | 0.216 | 0.090 | 0.809 | 0.694 |
| 1715 | **Mrpl46** | Unchanged | Q9EQI8 | 4 | 33 | 4 | 283 | 32.1 | 7.4 | 20 | 1.130 | 0.756 | 1.449 | 0.299 | 0.879 | 0.650 | 1.648 | 0.134 |
| 1716 | **Acp5** | Unchanged | Q05117 | 4 | 6 | 4 | 327 | 36.8 | 8.9 | 25 | 0.749 | 0.382 | 1.569 | 0.131 | 1.181 | 0.640 | 1.329 | 0.422 |
| 1717 | **Aldh16a1** | Unchanged | Q571I9 | 12 | 87 | 12 | 802 | 84.7 | 6.32 | 25 | 0.708 | 0.281 | 1.111 | 0.684 | 0.981 | 0.944 | 1.132 | 0.672 |
| 1718 | **Atp6v1h iso** | Unchanged | Q8BVE3 | 7 | 32 | 1 | 483 | 55.8 | 6.61 | 25 | 0.451 | 0.124 | 0.446 | 0.159 | 0.629 | 0.269 | 0.708 | 0.418 |
| 1719 | **Ssbp1** | Unchanged | Q9CYR0 | 3 | 44 | 3 | 152 | 17.3 | 9.92 | 25 | 0.994 | 0.988 | 1.370 | 0.548 | 1.474 | 0.440 | 0.929 | 0.891 |
| 1720 | **Myl1** | Unchanged | P05977 | 4 | 10 | 3 | 188 | 20.6 | 5.03 | 25 | 0.854 | 0.813 | 0.462 | 0.363 | 0.005 | 0.061 | 99.550 | 0.189 |
| 1721 | **Tpm2** | Unchanged | P58774 | 10 | 77 | 2 | 284 | 32.8 | 4.7 | 25 | 0.699 | 0.404 | 0.803 | 0.569 | 7.968 | 0.127 | 0.101 | 0.117 |
| 1722 | **Fkbp1a** | Unchanged | P26883 | 2 | 37 | 2 | 108 | 11.9 | 8.16 | 13 | 0.822 | 0.377 | 0.691 | 0.180 | 1.067 | 0.771 | 0.648 | 0.120 |
| 1723 | **Ica** | Unchanged | Q9DBD0 | 10 | 45 | 10 | 700 | 76.7 | 7.25 | 25 | 0.870 | 0.736 | 1.446 | 0.350 | 1.189 | 0.622 | 1.216 | 0.459 |
| 1724 | **Faf2** | Unchanged | Q3TDN2 | 6 | 33 | 6 | 445 | 52.4 | 5.47 | 25 | 0.916 | 0.669 | 1.359 | 0.485 | 0.848 | 0.500 | 1.602 | 0.311 |
| 1725 | **Serpinh1** | Unchanged | P19324 | 7 | 54 | 7 | 417 | 46.5 | 8.82 | 25 | 1.422 | 0.427 | 1.742 | 0.158 | 3.817 | 0.055 | 0.456 | 0.158 |
| 1726 | **Hmbs** | Unchanged | P22907 | 7 | 62 | 7 | 361 | 39.3 | 6.87 | 25 | 1.702 | 0.131 | 1.650 | 0.235 | 1.586 | 0.071 | 1.041 | 0.898 |
| 1727 | **Mrps30** | UP | Q9D0G0 | 6 | 18 | 6 | 442 | 49.9 | 9.38 | 25 | 4.007 | 0.086 | 4.128 | 0.007 | 5.711 | 0.045 | 0.723 | 0.483 |
| 1728 | **Rbp4** | UP | Q00724 | 5 | 26 | 5 | 201 | 23.2 | 5.99 | 25 | 6.161 | 0.060 | 2.881 | 0.191 | 4.784 | 0.010 | 0.602 | 0.299 |
| 1729 | **Mat2b** | UP | Q99LB6 | 5 | 35 | 4 | 334 | 37.4 | 6.95 | 25 | 1.037 | 0.947 | 1.318 | 0.684 | 0.599 | 0.273 | 2.202 | 0.334 |
| 1730 | **Rpl28** | Unchanged | P41105 | 6 | 46 | 6 | 137 | 15.7 | 12.02 | 25 | 1.892 | 0.076 | 1.374 | 0.645 | 0.967 | 0.933 | 1.422 | 0.593 |
| 1731 | **Dap3** | Unchanged | Q9ER88 | 6 | 29 | 6 | 391 | 44.7 | 8.94 | 25 | 1.435 | 0.437 | 2.497 | 0.294 | 1.148 | 0.735 | 2.174 | 0.334 |
| 1732 | **Hypk** | Unchanged | Q9CR41 | 2 | 6 | 2 | 129 | 14.7 | 4.93 | 25 | 0.710 | 0.700 | 0.505 | 0.526 | 0.835 | 0.823 | 0.605 | 0.511 |
| 1733 | **Ebp** | UP | P70245 | 3 | 55 | 3 | 230 | 26.2 | 7.56 | 25 | 1.942 | 0.293 | 2.453 | 0.222 | 2.907 | 0.049 | 0.844 | 0.712 |
| 1734 | **Tmem256** | Unchanged | Q5F285 | 1 | 6 | 1 | 113 | 11.6 | 8.94 | 25 | 1.577 | 0.598 | 1.691 | 0.590 | 0.086 | 0.152 | 19.768 | 0.179 |
| 1735 | **S100a9** | Unchanged | P31725 | 4 | 55 | 4 | 113 | 13 | 7.17 | 25 | 0.966 | 0.962 | 2.007 | 0.312 | 5.146 | 0.148 | 0.390 | 0.273 |
| 1736 | **Rab4a** | Unchanged | P56371 | 4 | 25 | 3 | 218 | 24.4 | 6.07 | 25 | 0.564 | 0.503 | 0.027 | 0.093 | 0.039 | 0.097 | 0.693 | 0.387 |
| 1737 | **Ppcs** | Unchanged | Q8VDG5 | 7 | 56 | 7 | 311 | 33.8 | 6.55 | 25 | 1.292 | 0.582 | 0.928 | 0.755 | 1.043 | 0.904 | 0.890 | 0.749 |
| 1738 | **Fbxo6** | UP | Q9QZN4 | 5 | 17 | 5 | 295 | 34.5 | 8.18 | 16 | 3.181 | 0.211 | 4.608 | 0.013 | 3.138 | 0.056 | 1.468 | 0.336 |
| 1739 | **Fga** | Unchanged | E9PV24 | 15 | 177 | 15 | 789 | 87.4 | 6.11 | 25 | 0.574 | 0.208 | 0.559 | 0.230 | 0.640 | 0.292 | 0.873 | 0.621 |
| 1740 | **Alyref** | Unchanged | O08583 | 4 | 11 | 4 | 255 | 26.9 | 11.15 | 25 | 1.538 | 0.416 | 2.893 | 0.200 | 2.285 | 0.170 | 1.266 | 0.709 |
| 1741 | **Plaa** | Unchanged | P27612 | 12 | 34 | 12 | 794 | 87.2 | 6.14 | 25 | 0.892 | 0.645 | 1.095 | 0.734 | 1.067 | 0.872 | 1.026 | 0.952 |
| 1742 | **Pgm2** | UP | Q7TSV4 | 10 | 44 | 10 | 620 | 68.7 | 6.14 | 25 | 1.202 | 0.670 | 2.378 | 0.020 | 0.952 | 0.853 | 2.496 | 0.011 |
| 1743 | **Fnta** | Unchanged | Q61239 | 6 | 65 | 6 | 377 | 44 | 4.93 | 25 | 1.754 | 0.342 | 2.048 | 0.068 | 1.994 | 0.056 | 1.027 | 0.931 |
| 1744 | **Ahnak** | Unchanged | E9Q616 | 19 | 69 | 19 | 5656 | 603.9 | 6.3 | 25 | 1.173 | 0.755 | 1.548 | 0.447 | 0.497 | 0.156 | 3.114 | 0.126 |
| 1745 | **Opa3** | Unchanged | Q505D7 | 3 | 27 | 2 | 179 | 20.1 | 8.34 | 25 | 0.309 | 0.067 | 0.855 | 0.771 | 0.398 | 0.111 | 2.146 | 0.294 |
| 1746 | **Npepps** | Unchanged | Q11011 | 15 | 73 | 15 | 920 | 103.3 | 5.9 | 25 | 0.750 | 0.313 | 1.335 | 0.438 | 1.009 | 0.978 | 1.323 | 0.450 |
| 1747 | **Dap** | Unchanged | Q91XC8 | 1 | 7 | 1 | 102 | 11.1 | 9.36 | 25 | 1.377 | 0.667 | 2.102 | 0.452 | 0.484 | 0.305 | 4.343 | 0.257 |
| 1748 | **Srp68** | UP | Q8BMA6 | 12 | 70 | 12 | 625 | 70.5 | 8.57 | 24 | 1.538 | 0.226 | 2.717 | 0.005 | 2.087 | 0.045 | 1.302 | 0.327 |
| 1749 | **Slc4a1** | Unchanged | P04919 | 13 | 88 | 13 | 929 | 103.1 | 5.45 | 24 | 1.470 | 0.176 | 1.284 | 0.471 | 1.253 | 0.445 | 1.025 | 0.934 |
| 1750 | **Snrpf** | Down | P62307 | 3 | 25 | 3 | 86 | 9.7 | 4.67 | 24 | 0.877 | 0.737 | 0.470 | 0.007 | 0.476 | 0.041 | 0.988 | 0.979 |
| 1751 | **Prkar1a** | Unchanged | Q9DBC7 | 6 | 48 | 6 | 381 | 43.2 | 5.35 | 24 | 1.222 | 0.322 | 1.209 | 0.595 | 0.904 | 0.543 | 1.336 | 0.458 |
| 1752 | **Tmem19** | Unchanged | Q91W52 | 3 | 24 | 3 | 336 | 36.3 | 7.72 | 24 | 0.265 | 0.157 | 0.054 | 0.064 | 0.424 | 0.291 | 0.128 | 0.193 |

| | | | | | | | | | | | | | | | | | | |
|---|---|---|---|---|---|---|---|---|---|---|---|---|---|---|---|---|---|---|
| 1753 | **Gm4450** | Unchanged | E9Q007 | 6 | 32 | 1 | 373 | 41.7 | 8.44 | 24 | 1.456 | 0.471 | 0.907 | 0.872 | 0.893 | 0.844 | 1.015 | 0.971 |
| 1754 | **Eef1diso** | Unchanged | A0A0R4J1E2 | 11 | 108 | 1 | 660 | 72.9 | 6.51 | 24 | 0.846 | 0.353 | 0.899 | 0.701 | 1.079 | 0.749 | 0.833 | 0.561 |
| 1755 | **Fkbp8** | Unchanged | O35465 | 6 | 43 | 6 | 402 | 43.5 | 5.16 | 24 | 0.550 | 0.514 | 0.955 | 0.953 | 0.358 | 0.332 | 2.666 | 0.167 |
| 1756 | **Gpx4** | Unchanged | O70325 | 5 | 24 | 5 | 197 | 22.2 | 8.46 | 24 | 0.557 | 0.367 | 0.750 | 0.603 | 0.434 | 0.214 | 1.728 | 0.209 |
| 1757 | **Mrps9** | Unchanged | Q9D7N3 | 6 | 24 | 6 | 390 | 44.9 | 8.81 | 24 | 0.104 | 0.070 | 0.132 | 0.079 | 0.091 | 0.067 | 1.440 | 0.543 |
| 1758 | **Map4** | Unchanged | P27546 | 15 | 110 | 15 | 1125 | 117.4 | 4.98 | 24 | 0.317 | 0.109 | 0.436 | 0.233 | 0.586 | 0.342 | 0.744 | 0.603 |
| 1759 | **Gm2a** | UP | Q60648 | 3 | 41 | 3 | 193 | 20.8 | 5.9 | 24 | 6.111 | 0.018 | 4.102 | 0.034 | 2.023 | 0.209 | 2.028 | 0.157 |
| 1760 | **Pygm** | Unchanged | Q9WUB3 | 14 | 119 | 6 | 842 | 97.2 | 7.11 | 24 | 1.703 | 0.424 | 0.849 | 0.822 | 1.626 | 0.499 | 0.522 | 0.323 |
| 1761 | **Naga** | UP | Q9QWR8 | 8 | 85 | 8 | 415 | 47.2 | 6.44 | 24 | 0.775 | 0.537 | 2.017 | 0.013 | 0.689 | 0.399 | 2.926 | 0.000 |
| 1762 | **Timm44** | Unchanged | O35857 | 11 | 61 | 11 | 452 | 51.1 | 8.13 | 24 | 1.340 | 0.477 | 0.824 | 0.681 | 1.265 | 0.653 | 0.651 | 0.340 |
| 1763 | **Spcs2** | Unchanged | Q9CYN2 | 5 | 35 | 5 | 226 | 25 | 8.57 | 24 | 0.952 | 0.881 | 1.056 | 0.907 | 0.979 | 0.948 | 1.078 | 0.868 |
| 1764 | **Mapk3** | Unchanged | Q63844 | 6 | 26 | 2 | 380 | 43 | 6.61 | 24 | 1.249 | 0.482 | 2.514 | 0.144 | 2.252 | 0.097 | 1.116 | 0.818 |
| 1765 | **Paip2** | Down | Q9D6V8 | 1 | 5 | 1 | 124 | 14.7 | 4.12 | 24 | 0.805 | 0.186 | 0.687 | 0.412 | 0.419 | 0.012 | 1.641 | 0.516 |
| 1766 | **Kpna1** | Unchanged | Q60960 | 6 | 51 | 5 | 538 | 60.1 | 5.01 | 24 | 1.149 | 0.811 | 6.562 | 0.061 | 1.385 | 0.361 | 4.737 | 0.078 |
| 1767 | **Cd302** | Down | Q9DCG2 | 4 | 34 | 4 | 228 | 25.4 | 4.61 | 24 | 0.609 | 0.104 | 0.494 | 0.035 | 0.616 | 0.105 | 0.802 | 0.244 |
| 1768 | **Ptcd3** | UP | Q14C51 | 10 | 67 | 10 | 685 | 77.7 | 5.88 | 24 | 1.500 | 0.118 | 1.821 | 0.003 | 1.805 | 0.011 | 1.009 | 0.948 |
| 1769 | **Ambp** | Down | Q07456 | 7 | 38 | 7 | 349 | 39 | 6.32 | 24 | 0.475 | 0.019 | 0.861 | 0.523 | 0.528 | 0.035 | 1.631 | 0.043 |
| 1770 | **Ube2m** | Unchanged | P61082 | 4 | 28 | 4 | 183 | 20.9 | 7.69 | 24 | 0.879 | 0.696 | 0.795 | 0.560 | 1.049 | 0.930 | 0.758 | 0.663 |
| 1771 | **Eif5b** | Unchanged | Q05D44 | 16 | 74 | 16 | 1216 | 137.5 | 5.59 | 24 | 1.070 | 0.788 | 1.480 | 0.217 | 1.238 | 0.381 | 1.196 | 0.510 |
| 1772 | **Mtdh** | Unchanged | Q80WJ7 | 10 | 76 | 10 | 579 | 63.8 | 9.33 | 24 | 1.824 | 0.221 | 3.836 | 0.164 | 1.337 | 0.447 | 2.870 | 0.222 |
| 1773 | **Ighv1-81** | Down | A0A075B5Y4 | 2 | 3 | 2 | 117 | 13 | 8.79 | 24 | 0.148 | 0.017 | 0.075 | 0.011 | 1.412 | 0.650 | 0.053 | 0.145 |
| 1774 | **Ighv1-76** | Unchanged | A0A0G2JFE9 | 2 | 2 | 2 | 117 | 13 | 9.07 | 24 | 0.259 | 0.364 | 2.408 | 0.533 | 2.247 | 0.303 | 1.072 | 0.943 |
| 1775 | **Mapre1** | Unchanged | Q61166 | 4 | 9 | 4 | 268 | 30 | 5.22 | 24 | 0.577 | 0.095 | 1.138 | 0.782 | 1.865 | 0.058 | 0.610 | 0.231 |
| 1776 | **Coq5** | UP | Q9CXI0 | 4 | 6 | 4 | 327 | 37.3 | 7.49 | 24 | 9.643 | 0.067 | 15.585 | 0.016 | 9.680 | 0.026 | 1.610 | 0.337 |
| 1777 | **Sgpl1** | Unchanged | Q8R0X7 | 10 | 62 | 10 | 568 | 63.6 | 9.1 | 24 | 0.878 | 0.702 | 1.264 | 0.588 | 0.791 | 0.371 | 1.597 | 0.308 |
| 1778 | **Sigmar1** | Unchanged | O55242 | 3 | 15 | 3 | 223 | 25.2 | 5.91 | 24 | 0.419 | 0.260 | 0.333 | 0.179 | 0.555 | 0.395 | 0.600 | 0.315 |
| 1779 | **Sumo1** | Unchanged | P63166 | 1 | 1 | 1 | 101 | 11.6 | 5.52 | 24 | 0.679 | 0.532 | 0.683 | 0.612 | 0.761 | 0.690 | 0.897 | 0.901 |
| 1780 | **Mrpl15** | Unchanged | Q9CPR5 | 5 | 13 | 5 | 295 | 33.5 | 10.07 | 24 | 2.761 | 0.210 | 6.263 | 0.055 | 0.984 | 0.963 | 6.364 | 0.053 |
| 1781 | **Ndufab1** | UP | Q9CR21 | 4 | 16 | 4 | 156 | 17.4 | 5.21 | 24 | 1.391 | 0.502 | 1.713 | 0.523 | 5.686 | 0.024 | 0.301 | 0.073 |
| 1782 | **Tecr** | Unchanged | Q9CY27 | 10 | 75 | 10 | 308 | 36.1 | 9.55 | 24 | 1.028 | 0.849 | 1.052 | 0.793 | 0.771 | 0.113 | 1.364 | 0.163 |
| 1783 | **Crk** | Unchanged | Q64010 | 5 | 52 | 5 | 304 | 33.8 | 5.55 | 24 | 0.743 | 0.323 | 1.210 | 0.668 | 0.975 | 0.937 | 1.241 | 0.627 |
| 1784 | **Fis1** | Down | Q9CQ92 | 3 | 24 | 3 | 152 | 17 | 8.53 | 24 | 0.253 | 0.034 | 1.049 | 0.941 | 0.831 | 0.673 | 1.263 | 0.743 |
| 1785 | **Ilk** | Unchanged | O55222 | 8 | 53 | 8 | 452 | 51.3 | 8.07 | 24 | 3.182 | 0.318 | 13.521 | 0.224 | 2.058 | 0.282 | 6.569 | 0.264 |
| 1786 | **Pdcd4** | Unchanged | Q61823 | 7 | 25 | 7 | 469 | 51.7 | 5.16 | 24 | 7.649 | 0.078 | 5.560 | 0.143 | 6.537 | 0.091 | 0.851 | 0.814 |
| 1787 | **Mrpl55** | Unchanged | Q9CZ83 | 2 | 5 | 2 | 127 | 15.1 | 10.51 | 24 | 1.029 | 0.938 | 2.159 | 0.062 | 0.921 | 0.816 | 2.345 | 0.069 |
| 1788 | **Pgm3** | UP | Q9CYR6 | 9 | 62 | 9 | 542 | 59.4 | 6.2 | 24 | 1.420 | 0.244 | 2.706 | 0.128 | 1.837 | 0.048 | 1.473 | 0.412 |
| 1789 | **Fam25c** | Unchanged | Q8CF02 | 1 | 1 | 1 | 89 | 9.2 | 6.05 | 24 | 0.783 | 0.573 | 11.530 | 0.068 | 3.217 | 0.160 | 3.584 | 0.147 |
| 1790 | **Serpinb6** | Unchanged | Q60854 | 7 | 45 | 7 | 378 | 42.6 | 5.74 | 24 | 1.144 | 0.715 | 1.243 | 0.617 | 0.513 | 0.051 | 2.424 | 0.134 |
| 1791 | **Snrpa1** | Unchanged | P57784 | 3 | 18 | 3 | 255 | 28.3 | 8.62 | 24 | 0.873 | 0.664 | 0.600 | 0.373 | 0.879 | 0.745 | 0.683 | 0.519 |
| 1792 | **Atad3** | UP | Q925I1 | 9 | 57 | 9 | 591 | 66.7 | 9.29 | 24 | 2.015 | 0.042 | 2.697 | 0.173 | 1.293 | 0.356 | 2.086 | 0.238 |
| 1793 | **Mybbp1a** | Unchanged | Q7TPV4 | 18 | 102 | 18 | 1344 | 151.9 | 8.95 | 24 | 0.780 | 0.612 | 0.860 | 0.693 | 0.831 | 0.672 | 1.035 | 0.920 |

| | | | | | | | | | | | | | | | | | |
|---|---|---|---|---|---|---|---|---|---|---|---|---|---|---|---|---|---|
| 1794 | **Actr10** | Unchanged | Q9QZB7 | 5 | 5 | 5 | 417 | 46.2 | 7.61 | 24 | 1.294 | 0.552 | 4.161 | 0.062 | 0.658 | 0.441 | 6.324 | 0.037 |
| 1795 | **S100a13** | Unchanged | P97352 | 2 | 13 | 2 | 98 | 11.2 | 6.13 | 23 | 1.082 | 0.883 | 1.261 | 0.708 | 1.243 | 0.683 | 1.015 | 0.981 |
| 1796 | **Hrg** | Unchanged | Q9ESB3 | 11 | 102 | 11 | 525 | 59.1 | 7.66 | 23 | 1.078 | 0.774 | 1.054 | 0.868 | 3.854 | 0.058 | 0.273 | 0.060 |
| 1797 | **Ugt3a1** | Unchanged | Q3UP75 | 8 | 77 | 5 | 523 | 59.7 | 7.34 | 23 | 1.182 | 0.650 | 1.403 | 0.303 | 1.652 | 0.242 | 0.849 | 0.587 |
| 1798 | **Ndufv3iso** | Unchanged | Q3U422 | 4 | 6 | 3 | 468 | 50.5 | 8.97 | 23 | 1.795 | 0.340 | 1.652 | 0.300 | 7.945 | 0.157 | 0.208 | 0.192 |
| 1799 | **Nceh1** | Unchanged | Q8BLF1 | 5 | 42 | 5 | 408 | 45.7 | 7.05 | 23 | 1.256 | 0.326 | 1.567 | 0.137 | 1.048 | 0.908 | 1.496 | 0.304 |
| 1800 | **Abcb11** | Unchanged | Q9QY30 | 20 | 193 | 20 | 1321 | 146.7 | 7.66 | 23 | 1.082 | 0.636 | 1.290 | 0.155 | 1.295 | 0.281 | 0.995 | 0.985 |
| 1801 | **Sri** | Unchanged | Q6P069 | 4 | 12 | 4 | 198 | 21.6 | 5.59 | 23 | 1.209 | 0.751 | 1.939 | 0.094 | 0.818 | 0.730 | 2.370 | 0.039 |
| 1802 | **Iscu** | Unchanged | Q9D7P6 | 3 | 5 | 3 | 168 | 18.1 | 9.29 | 23 | 0.327 | 0.427 | 0.277 | 0.397 | 0.028 | 0.248 | 9.924 | 0.263 |
| 1803 | **Higd1a** | UP | Q9JLR9 | 1 | 10 | 1 | 95 | 10.4 | 9.79 | 23 | 1.761 | 0.301 | 3.912 | 0.005 | 2.689 | 0.095 | 1.455 | 0.289 |
| 1804 | **Sept9** | Unchanged | Q80UG5 | 9 | 43 | 9 | 583 | 65.5 | 8.9 | 23 | 0.953 | 0.898 | 1.522 | 0.338 | 1.697 | 0.088 | 0.896 | 0.733 |
| 1805 | **Alg2** | Down | Q9DBE8 | 5 | 20 | 5 | 415 | 47.4 | 7.97 | 23 | 0.752 | 0.534 | 1.194 | 0.751 | 0.281 | 0.043 | 4.250 | 0.112 |
| 1806 | **Mrpl4** | Unchanged | Q9DCU6 | 5 | 18 | 5 | 294 | 33.1 | 9.82 | 23 | 0.849 | 0.593 | 2.917 | 0.063 | 0.571 | 0.067 | 5.108 | 0.032 |
| 1807 | **Tnnc2** | Unchanged | P20801 | 3 | 6 | 3 | 160 | 18.1 | 4.2 | 23 | 0.766 | 0.714 | 2.089 | 0.328 | 1.271 | 0.707 | 1.644 | 0.421 |
| 1808 | **Dhx9** | Unchanged | E9QNN1 | 21 | 101 | 21 | 1384 | 149.6 | 6.83 | 23 | 1.014 | 0.918 | 1.011 | 0.972 | 0.729 | 0.071 | 1.388 | 0.397 |
| 1809 | **Snx2** | Unchanged | Q9CWK8 | 8 | 53 | 7 | 519 | 58.4 | 5.12 | 23 | 0.902 | 0.798 | 0.874 | 0.687 | 1.812 | 0.090 | 0.483 | 0.027 |
| 1810 | **Chp1** | Unchanged | P61022 | 4 | 18 | 4 | 195 | 22.4 | 5.1 | 23 | 1.123 | 0.798 | 1.194 | 0.687 | 1.586 | 0.263 | 0.753 | 0.291 |
| 1811 | **Dhrs11** | Unchanged | Q3U0B3 | 4 | 26 | 4 | 260 | 28.3 | 6.34 | 23 | 1.008 | 0.989 | 1.674 | 0.212 | 0.584 | 0.394 | 2.868 | 0.007 |
| 1812 | **Snrpd3** | UP | P62320 | 2 | 17 | 2 | 126 | 13.9 | 10.32 | 23 | 5.173 | 0.053 | 2.579 | 0.005 | 1.243 | 0.381 | 2.075 | 0.016 |
| 1813 | **Ecsit** | Unchanged | Q9QZH6 | 5 | 19 | 5 | 435 | 49.8 | 6.6 | 23 | 0.658 | 0.207 | 1.062 | 0.820 | 0.564 | 0.103 | 1.883 | 0.034 |
| 1814 | **Gcn1** | Unchanged | E9PVA8 | 39 | 225 | 39 | 2671 | 292.8 | 7.36 | 23 | 1.083 | 0.608 | 2.320 | 0.058 | 0.998 | 0.990 | 2.326 | 0.059 |
| 1815 | **Lyz2** | Unchanged | P08905 | 4 | 27 | 2 | 148 | 16.7 | 8.81 | 23 | 0.884 | 0.834 | 0.779 | 0.631 | 0.519 | 0.283 | 1.502 | 0.355 |
| 1816 | **Nt5c3a** | Unchanged | Q9D020 | 5 | 17 | 5 | 331 | 37.2 | 6.65 | 23 | 0.611 | 0.342 | 0.868 | 0.753 | 0.575 | 0.383 | 1.508 | 0.454 |
| 1817 | **Pin4** | Unchanged | Q9CWW6 | 1 | 2 | 1 | 131 | 13.8 | 9.77 | 23 | 0.139 | 0.086 | 0.713 | 0.581 | 0.095 | 0.073 | 7.484 | 0.036 |
| 1818 | **Sae1** | Unchanged | Q9R1T2 | 5 | 36 | 5 | 350 | 38.6 | 5.36 | 23 | 1.176 | 0.740 | 1.516 | 0.414 | 1.306 | 0.569 | 1.161 | 0.731 |
| 1819 | **Pdk2** | Unchanged | Q9JK42 | 5 | 9 | 5 | 407 | 46 | 6.61 | 23 | 0.835 | 0.682 | 0.746 | 0.531 | 0.724 | 0.476 | 1.031 | 0.941 |
| 1820 | **Uba5** | Unchanged | Q8VE47 | 5 | 30 | 5 | 403 | 44.8 | 4.96 | 23 | 0.644 | 0.176 | 0.512 | 0.107 | 3.346 | 0.218 | 0.153 | 0.143 |
| 1821 | **Asns** | Unchanged | Q61024 | 8 | 37 | 8 | 561 | 64.2 | 6.58 | 23 | 0.596 | 0.204 | 1.075 | 0.876 | 0.782 | 0.429 | 1.375 | 0.508 |
| 1822 | **Mrpl58** | UP | Q8R035 | 3 | 6 | 3 | 206 | 23.5 | 10.18 | 23 | 1.458 | 0.370 | 2.245 | 0.024 | 0.391 | 0.163 | 5.735 | 0.000 |
| 1823 | **Upp2** | Unchanged | Q8CGR7 | 3 | 6 | 3 | 320 | 35.7 | 6.52 | 23 | 2.760 | 0.240 | 1.291 | 0.610 | 0.467 | 0.279 | 2.764 | 0.064 |
| 1824 | **Ddah2** | Unchanged | Q99LD8 | 4 | 17 | 3 | 285 | 29.6 | 6.01 | 23 | 0.732 | 0.631 | 2.876 | 0.100 | 0.756 | 0.640 | 3.804 | 0.049 |
| 1825 | **Rab5b** | Unchanged | P61021 | 4 | 44 | 2 | 215 | 23.7 | 8.13 | 23 | 0.734 | 0.640 | 0.298 | 0.139 | 1.277 | 0.675 | 0.233 | 0.075 |
| 1826 | **Ptms** | Unchanged | Q9D0J8 | 3 | 59 | 3 | 101 | 11.4 | 4.22 | 23 | 1.551 | 0.490 | 0.602 | 0.284 | 1.274 | 0.692 | 0.472 | 0.361 |
| 1827 | **Gltpd2** | Unchanged | Q8K0R6 | 4 | 16 | 4 | 321 | 34.4 | 9.16 | 23 | 0.774 | 0.696 | 1.667 | 0.409 | 0.607 | 0.470 | 2.748 | 0.115 |
| 1828 | **Ada** | Unchanged | P03958 | 4 | 25 | 4 | 352 | 40 | 5.72 | 23 | 1.218 | 0.740 | 0.560 | 0.445 | 2.385 | 0.058 | 0.235 | 0.006 |
| 1829 | **Mavs** | Unchanged | Q8VCF0 | 6 | 71 | 6 | 503 | 53.4 | 6.37 | 23 | 0.618 | 0.572 | 0.605 | 0.517 | 0.400 | 0.309 | 1.512 | 0.459 |
| 1830 | **Flot2** | Unchanged | Q60634 | 7 | 32 | 7 | 428 | 47 | 5.2 | 23 | 0.764 | 0.345 | 1.010 | 0.956 | 1.019 | 0.953 | 0.992 | 0.975 |
| 1831 | **Mindy1** | Unchanged | Q76LS9 | 5 | 28 | 5 | 468 | 51.2 | 4.73 | 23 | 1.211 | 0.801 | 2.107 | 0.257 | 1.421 | 0.496 | 1.483 | 0.450 |
| 1832 | **Samhd1** | UP | Q60710 | 9 | 45 | 9 | 658 | 75.8 | 7.93 | 23 | 1.447 | 0.378 | 2.339 | 0.069 | 1.941 | 0.025 | 1.205 | 0.511 |
| 1833 | **Parp9** | Down | Q8CAS9 | 13 | 63 | 13 | 866 | 96.6 | 7.09 | 23 | 1.230 | 0.429 | 1.574 | 0.253 | 0.519 | 0.009 | 3.032 | 0.049 |
| 1834 | **Dnajb11** | Unchanged | Q99KV1 | 7 | 73 | 7 | 358 | 40.5 | 6.32 | 23 | 1.984 | 0.293 | 1.406 | 0.518 | 1.638 | 0.413 | 0.858 | 0.804 |

| | | | | | | | | | | | | | | | | | |
|---|---|---|---|---|---|---|---|---|---|---|---|---|---|---|---|---|---|
| 1835 | **Lamtor3** | Unchanged | O88653 | 1 | 2 | 1 | 124 | 13.5 | 7.34 | 23 | 0.917 | 0.891 | 1.610 | 0.515 | 0.213 | 0.150 | 7.553 | 0.100 |
| 1836 | **Ctsa** | UP | P16675 | 8 | 92 | 8 | 474 | 53.8 | 5.86 | 23 | 1.629 | 0.029 | 2.606 | 0.000 | 2.742 | 0.000 | 0.950 | 0.663 |
| 1837 | **Poldip2** | Unchanged | Q91VA6 | 5 | 37 | 5 | 368 | 41.8 | 8.63 | 23 | 1.601 | 0.107 | 1.298 | 0.538 | 1.244 | 0.345 | 1.043 | 0.908 |
| 1838 | **Ctsc** | UP | P97821 | 8 | 75 | 8 | 462 | 52.3 | 6.89 | 23 | 1.115 | 0.782 | 1.503 | 0.240 | 2.254 | 0.017 | 0.667 | 0.193 |
| 1839 | **Klhdc4** | Unchanged | G3UZC6 | 1 | 1 | 1 | 40 | 4.6 | 10.89 | 23 | 0.920 | 0.851 | 1.900 | 0.106 | 1.509 | 0.277 | 1.258 | 0.346 |
| 1840 | **Ahsa1** | Unchanged | Q8BK64 | 5 | 87 | 5 | 338 | 38.1 | 5.53 | 22 | 0.978 | 0.923 | 1.311 | 0.220 | 1.167 | 0.621 | 1.124 | 0.654 |
| 1841 | **Naa50** | Unchanged | Q6PGB6 | 3 | 4 | 3 | 169 | 19.4 | 8.81 | 22 | 0.767 | 0.443 | 0.930 | 0.841 | 0.386 | 0.063 | 2.413 | 0.040 |
| 1842 | **Mrpl13** | UP | Q9D1P0 | 3 | 7 | 3 | 178 | 20.7 | 9.35 | 22 | 6.150 | 0.007 | 2.766 | 0.164 | 1.733 | 0.347 | 1.596 | 0.450 |
| 1843 | **Abcf1** | UP | Q6P542 | 8 | 30 | 8 | 837 | 94.9 | 6.51 | 22 | 3.471 | 0.024 | 3.900 | 0.016 | 3.781 | 0.032 | 1.031 | 0.934 |
| 1844 | **Vbp1** | Unchanged | P61759 | 2 | 8 | 2 | 196 | 22.4 | 6.28 | 22 | 2.729 | 0.312 | 5.248 | 0.187 | 1.221 | 0.394 | 4.299 | 0.208 |
| 1845 | **Eif4b** | Unchanged | Q8BGD9 | 10 | 48 | 10 | 611 | 68.8 | 5.67 | 22 | 0.737 | 0.215 | 3.750 | 0.291 | 1.455 | 0.213 | 2.577 | 0.375 |
| 1846 | **Api5** | UP | O35841 | 6 | 65 | 6 | 504 | 56.8 | 5.92 | 22 | 2.203 | 0.109 | 2.783 | 0.009 | 2.546 | 0.013 | 1.093 | 0.649 |
| 1847 | **Emc7** | Unchanged | Q9EP72 | 3 | 14 | 3 | 241 | 26.3 | 9.23 | 22 | 0.697 | 0.611 | 1.219 | 0.766 | 0.754 | 0.655 | 1.616 | 0.370 |
| 1848 | **Ptbp3** | Unchanged | Q8BHD7 | 7 | 37 | 4 | 523 | 56.7 | 9.13 | 22 | 1.543 | 0.334 | 1.375 | 0.494 | 0.438 | 0.054 | 3.139 | 0.085 |
| 1849 | **Sntb1** | Unchanged | Q99L88 | 9 | 50 | 9 | 537 | 58 | 8.31 | 22 | 0.508 | 0.267 | 0.732 | 0.543 | 1.458 | 0.490 | 0.502 | 0.190 |
| 1850 | **Fdx1** | Down | P46656 | 3 | 73 | 3 | 188 | 20.1 | 5.62 | 22 | 0.756 | 0.296 | 0.463 | 0.047 | 0.239 | 0.007 | 1.937 | 0.142 |
| 1851 | **Mocs1** | Unchanged | Q5RKZ7 | 11 | 61 | 11 | 636 | 69.8 | 9.14 | 22 | 0.946 | 0.785 | 0.801 | 0.280 | 0.909 | 0.701 | 0.881 | 0.638 |
| 1852 | **Oxct1** | Unchanged | Q9D0K2 | 6 | 14 | 6 | 520 | 56 | 8.53 | 22 | 0.473 | 0.275 | 1.212 | 0.732 | 1.049 | 0.942 | 1.156 | 0.796 |
| 1853 | **Lgals3bpiso** | Unchanged | F6VRP8 | 2 | 10 | 1 | 139 | 15.4 | 8.47 | 22 | 1.890 | 0.468 | 3.018 | 0.108 | 2.147 | 0.221 | 1.406 | 0.506 |
| 1854 | **Creld2** | Unchanged | Q9CYA0 | 6 | 43 | 6 | 350 | 38.2 | 4.58 | 22 | 1.368 | 0.619 | 1.501 | 0.508 | 1.895 | 0.326 | 0.792 | 0.709 |
| 1855 | **Ap3m1** | UP | Q9JKC8 | 6 | 40 | 6 | 418 | 46.9 | 6.93 | 22 | 1.541 | 0.194 | 2.056 | 0.018 | 1.075 | 0.786 | 1.913 | 0.026 |
| 1856 | **Sec63** | UP | Q8VHE0 | 10 | 63 | 10 | 760 | 87.8 | 5.38 | 22 | 1.546 | 0.049 | 2.628 | 0.081 | 2.382 | 0.002 | 1.103 | 0.776 |
| 1857 | **Orm1** | Unchanged | Q60590 | 4 | 20 | 4 | 207 | 23.9 | 5.85 | 22 | 0.819 | 0.724 | 0.531 | 0.325 | 0.641 | 0.428 | 0.827 | 0.694 |
| 1858 | **Ccdc58** | Unchanged | Q8R3Q6 | 2 | 19 | 2 | 144 | 16.7 | 8.16 | 22 | 0.655 | 0.450 | 0.494 | 0.270 | 0.788 | 0.618 | 0.627 | 0.305 |
| 1859 | **Uhrf1bp1** | Unchanged | A0A3B2W3R7 | 1 | 1 | 1 | 108 | 11.9 | 7.84 | 22 | #DIV/0! | #DIV/0! | #DIV/0! | #DIV/0! | #DIV/0! | #DIV/0! | #DIV/0! | #DIV/0! |
| 1860 | **Acsl4** | UP | Q9QUJ7 | 8 | 40 | 8 | 711 | 79 | 8.28 | 22 | 1.488 | 0.407 | 1.672 | 0.123 | 1.942 | 0.012 | 0.861 | 0.481 |
| 1861 | **Man2a1** | Down | P27046 | 18 | 132 | 18 | 1150 | 131.5 | 8.03 | 22 | 0.872 | 0.676 | 0.794 | 0.454 | 0.341 | 0.034 | 2.328 | 0.000 |
| 1862 | **Sf3a3** | Down | Q9D554 | 7 | 19 | 7 | 501 | 58.8 | 5.34 | 22 | 0.760 | 0.322 | 0.511 | 0.042 | 0.526 | 0.066 | 0.970 | 0.951 |
| 1863 | **Ap1g1** | Down | P22892 | 13 | 89 | 13 | 822 | 91.3 | 6.8 | 22 | 1.072 | 0.735 | 1.485 | 0.368 | 0.578 | 0.033 | 2.569 | 0.109 |
| 1864 | **Eif2b4** | Down | Q61749 | 5 | 18 | 5 | 524 | 57.6 | 9.25 | 22 | 0.963 | 0.942 | 0.192 | 0.044 | 1.610 | 0.530 | 0.119 | 0.142 |
| 1865 | **Cops2** | UP | P61202 | 5 | 21 | 5 | 443 | 51.6 | 5.53 | 22 | 2.590 | 0.033 | 7.103 | 0.074 | 3.759 | 0.012 | 1.890 | 0.309 |
| 1866 | **Pdap1** | Unchanged | Q3UHX2 | 4 | 34 | 4 | 181 | 20.6 | 7.39 | 22 | 1.121 | 0.792 | 1.313 | 0.452 | 0.837 | 0.621 | 1.568 | 0.276 |
| 1867 | **Bola2** | UP | Q8BGS2 | 1 | 3 | 1 | 86 | 10.2 | 6.16 | 22 | 2.769 | 0.183 | 8.551 | 0.038 | 7.286 | 0.008 | 1.174 | 0.727 |
| 1868 | **Uba3** | Unchanged | Q8C878 | 5 | 23 | 5 | 462 | 51.7 | 5.45 | 22 | 1.656 | 0.089 | 3.010 | 0.092 | 1.620 | 0.157 | 1.858 | 0.228 |
| 1869 | **Abcb7** | Unchanged | Q61102 | 10 | 70 | 10 | 752 | 82.5 | 9.32 | 22 | 1.095 | 0.708 | 2.375 | 0.080 | 0.974 | 0.911 | 2.438 | 0.074 |
| 1870 | **Trabd** | UP | Q99JY4 | 6 | 32 | 6 | 376 | 42.2 | 8.41 | 22 | 1.585 | 0.347 | 2.843 | 0.213 | 2.151 | 0.037 | 1.322 | 0.635 |
| 1871 | **Vwa5a** | Unchanged | Q99KC8 | 11 | 96 | 11 | 793 | 87.1 | 6.58 | 22 | 0.990 | 0.915 | 1.189 | 0.524 | 1.101 | 0.140 | 1.080 | 0.769 |
| 1872 | **Preb** | Unchanged | Q9WUQ2 | 6 | 37 | 6 | 417 | 45.4 | 8.76 | 22 | 1.019 | 0.941 | 0.989 | 0.967 | 0.915 | 0.770 | 1.080 | 0.765 |
| 1873 | **Mat2a** | Unchanged | Q3THS6 | 9 | 103 | 5 | 395 | 43.7 | 6.48 | 22 | 1.211 | 0.413 | 0.864 | 0.623 | 0.640 | 0.126 | 1.351 | 0.270 |
| 1874 | **Zbtb8os** | Unchanged | Q505B7 | 2 | 6 | 2 | 168 | 19.7 | 4.56 | 22 | 1.280 | 0.563 | 2.142 | 0.265 | 0.370 | 0.080 | 5.791 | 0.092 |
| 1875 | **Prxl2a** | Unchanged | Q9CYH2 | 6 | 60 | 6 | 218 | 24.4 | 9.17 | 22 | 0.968 | 0.893 | 3.736 | 0.216 | 0.862 | 0.572 | 4.332 | 0.196 |

| | | | | | | | | | | | | | | | | | | |
|---|---|---|---|---|---|---|---|---|---|---|---|---|---|---|---|---|---|---|
| 1876 | **Kif5b** | UP | Q61768 | 13 | 48 | 13 | 963 | 109.5 | 6.44 | 22 | 3.536 | 0.019 | 3.746 | 0.088 | 3.980 | 0.090 | 0.941 | 0.913 |
| 1877 | **Ca8** | Unchanged | P28651 | 4 | 24 | 4 | 291 | 33.1 | 4.78 | 22 | 1.408 | 0.213 | 1.112 | 0.815 | 1.450 | 0.420 | 0.767 | 0.616 |
| 1878 | **Lyrm4** | Unchanged | Q8K215 | 3 | 11 | 3 | 91 | 10.8 | 10.13 | 22 | 1.027 | 0.948 | 0.839 | 0.671 | 1.174 | 0.612 | 0.714 | 0.305 |
| 1879 | **Lamtor5** | Unchanged | Q9D1L9 | 1 | 8 | 1 | 91 | 9.6 | 4.87 | 22 | 0.659 | 0.368 | 0.900 | 0.750 | 0.668 | 0.334 | 1.347 | 0.428 |
| 1880 | **Eif2b3** | Unchanged | B1AUN2 | 5 | 10 | 5 | 452 | 50.5 | 6.74 | 22 | 2.481 | 0.241 | 1.760 | 0.399 | 1.079 | 0.852 | 1.631 | 0.473 |
| 1881 | **Ctnnb1** | Unchanged | Q02248 | 13 | 57 | 11 | 781 | 85.4 | 5.86 | 22 | 0.768 | 0.413 | 1.768 | 0.124 | 0.818 | 0.548 | 2.162 | 0.056 |
| 1882 | **Arpc1a** | Unchanged | Q9R0Q6 | 4 | 23 | 3 | 370 | 41.6 | 8.18 | 22 | 1.026 | 0.953 | 1.061 | 0.925 | 1.636 | 0.236 | 0.648 | 0.418 |
| 1883 | **Anxa11** | Unchanged | P97384 | 9 | 59 | 8 | 503 | 54 | 7.66 | 22 | 0.974 | 0.901 | 1.249 | 0.057 | 0.875 | 0.451 | 1.428 | 0.050 |
| 1884 | **Kif13b** | Down | A0A286YCV9 | 26 | 89 | 25 | 1847 | 204.8 | 5.8 | 22 | 0.614 | 0.015 | 0.812 | 0.130 | 0.613 | 0.014 | 1.325 | 0.079 |
| 1885 | **Pex14** | Unchanged | Q9R0A0 | 5 | 58 | 5 | 376 | 41.2 | 5.11 | 22 | 1.462 | 0.387 | 2.434 | 0.113 | 1.654 | 0.414 | 1.472 | 0.462 |
| 1886 | **Hmox1** | Unchanged | P14901 | 3 | 7 | 3 | 289 | 32.9 | 6.54 | 22 | 0.969 | 0.962 | 1.106 | 0.895 | 0.134 | 0.078 | 8.266 | 0.174 |
| 1887 | **Nucks1** | Unchanged | Q80XU3 | 3 | 33 | 3 | 234 | 26.3 | 5.14 | 22 | 0.680 | 0.464 | 1.425 | 0.567 | 0.622 | 0.409 | 2.292 | 0.257 |
| 1888 | **G3bp2** | Unchanged | P97379 | 6 | 25 | 5 | 482 | 54.1 | 5.62 | 19 | 0.093 | 0.148 | 0.092 | 0.148 | 0.079 | 0.142 | 1.173 | 0.835 |
| 1889 | **Samm50** | UP | Q8BGH2 | 7 | 47 | 7 | 469 | 51.8 | 6.8 | 22 | 1.558 | 0.409 | 5.326 | 0.014 | 0.762 | 0.625 | 6.992 | 0.010 |
| 1890 | **Mydgf** | Unchanged | Q9CPT4 | 3 | 57 | 3 | 166 | 18 | 6.79 | 22 | 0.878 | 0.713 | 1.026 | 0.952 | 1.257 | 0.383 | 0.816 | 0.587 |
| 1891 | **Dguok** | Unchanged | Q9QX60 | 3 | 14 | 3 | 277 | 32.3 | 7.66 | 22 | 0.845 | 0.758 | 0.769 | 0.654 | 1.388 | 0.413 | 0.554 | 0.065 |
| 1892 | **Nsf** | Unchanged | P46460 | 12 | 68 | 12 | 744 | 82.6 | 6.95 | 22 | 0.458 | 0.213 | 1.191 | 0.794 | 0.179 | 0.064 | 6.665 | 0.129 |
| 1893 | **Ighg3** | Unchanged | P03987 | 5 | 25 | 5 | 398 | 43.9 | 7.14 | 22 | 2.101 | 0.147 | 6.004 | 0.168 | 1.131 | 0.812 | 5.311 | 0.176 |
| 1894 | **Pam16** | Unchanged | Q9CQV1 | 1 | 6 | 1 | 125 | 13.8 | 9.64 | 22 | 7.492 | 0.309 | 2.491 | 0.338 | 0.604 | 0.547 | 4.123 | 0.193 |
| 1895 | **Trmt112** | Unchanged | Q9DCG9 | 2 | 13 | 2 | 125 | 14.1 | 5.27 | 22 | 1.644 | 0.267 | 0.955 | 0.926 | 4.087 | 0.077 | 0.234 | 0.067 |
| 1896 | **Lmnb1** | Unchanged | P14733 | 9 | 46 | 9 | 588 | 66.7 | 5.16 | 22 | 1.066 | 0.917 | 1.011 | 0.986 | 1.632 | 0.470 | 0.619 | 0.456 |
| 1897 | **Gch1** | Unchanged | Q05915 | 3 | 22 | 3 | 241 | 27 | 8.27 | 22 | 1.289 | 0.569 | 0.837 | 0.679 | 1.318 | 0.406 | 0.635 | 0.139 |
| 1898 | **Nhp2** | Unchanged | Q9CRB2 | 2 | 14 | 2 | 153 | 17.2 | 8.41 | 22 | 1.477 | 0.525 | 1.762 | 0.391 | 0.350 | 0.087 | 5.029 | 0.113 |
| 1899 | **Hacd3** | Unchanged | Q8K2C9 | 5 | 70 | 5 | 362 | 43.1 | 9.13 | 22 | 1.143 | 0.806 | 1.051 | 0.919 | 0.873 | 0.797 | 1.203 | 0.518 |
| 1900 | **Pea15** | Unchanged | Q62048 | 2 | 2 | 2 | 130 | 15 | 5.02 | 22 | 0.924 | 0.908 | 2.157 | 0.196 | 0.667 | 0.497 | 3.234 | 0.067 |
| 1901 | **Timm50** | Unchanged | Q9D880 | 6 | 28 | 6 | 353 | 39.8 | 8.13 | 22 | 1.506 | 0.135 | 1.564 | 0.132 | 1.435 | 0.007 | 1.090 | 0.705 |
| 1902 | **Khsrp** | Unchanged | Q3U0V1 | 11 | 68 | 11 | 748 | 76.7 | 7.33 | 22 | 0.897 | 0.765 | 1.516 | 0.393 | 1.183 | 0.557 | 1.281 | 0.583 |
| 1903 | **Timm17b** | UP | Q9Z0V7 | 2 | 12 | 2 | 172 | 18.3 | 9.03 | 22 | 1.647 | 0.312 | 2.589 | 0.034 | 1.049 | 0.915 | 2.469 | 0.009 |
| 1904 | **Eif4g1** | UP | Q6NZJ6 | 27 | 188 | 25 | 1600 | 176 | 5.4 | 22 | 1.524 | 0.044 | 2.402 | 0.000 | 1.589 | 0.078 | 1.511 | 0.021 |
| 1905 | **Myo1b** | Unchanged | P46735 | 15 | 90 | 15 | 1107 | 128.5 | 9.26 | 21 | 0.995 | 0.987 | 0.927 | 0.792 | 0.672 | 0.218 | 1.380 | 0.114 |
| 1906 | **Ppm1f** | Unchanged | Q8CGA0 | 4 | 14 | 4 | 452 | 49.6 | 5.3 | 21 | 0.857 | 0.790 | 1.038 | 0.946 | 1.047 | 0.914 | 0.992 | 0.983 |
| 1907 | **Eci3** | Unchanged | Q78JN3 | 5 | 88 | 1 | 317 | 35.2 | 9.16 | 21 | 1.730 | 0.378 | 3.548 | 0.124 | 1.732 | 0.455 | 2.048 | 0.301 |
| 1908 | **Uqcr11** | Unchanged | Q9CPX8 | 1 | 7 | 1 | 56 | 6.5 | 9.82 | 21 | 2.408 | 0.259 | 2.958 | 0.257 | 2.950 | 0.165 | 1.003 | 0.997 |
| 1909 | **Gon7** | Unchanged | P0C8B4 | 1 | 4 | 1 | 98 | 10.3 | 4.09 | 21 | 0.269 | 0.294 | 3.344 | 0.169 | 0.681 | 0.719 | 4.914 | 0.118 |
| 1910 | **Coq6** | Unchanged | Q8R1S0 | 6 | 30 | 6 | 476 | 51.4 | 7.17 | 21 | 0.787 | 0.739 | 3.234 | 0.274 | 0.295 | 0.119 | 10.972 | 0.153 |
| 1911 | **Pfdn5** | Unchanged | Q9WU28 | 2 | 16 | 2 | 154 | 17.3 | 6.33 | 21 | 0.837 | 0.621 | 3.353 | 0.126 | 0.833 | 0.129 | 4.023 | 0.105 |
| 1912 | **Sdhd** | Unchanged | Q9CXV1 | 3 | 17 | 3 | 159 | 17 | 9.1 | 21 | 1.511 | 0.478 | 2.770 | 0.211 | 1.851 | 0.406 | 1.497 | 0.563 |
| 1913 | **Kng1** | Unchanged | O08677 | 10 | 67 | 10 | 661 | 73.1 | 6.54 | 21 | 0.973 | 0.958 | 0.827 | 0.733 | 0.723 | 0.569 | 1.144 | 0.585 |
| 1914 | **2810459M11Rik** | Unchanged | Q9CYS6 | 4 | 21 | 4 | 286 | 30.3 | 6.05 | 21 | 1.450 | 0.588 | 2.260 | 0.330 | 1.213 | 0.630 | 1.863 | 0.432 |
| 1915 | **Myg1** | Unchanged | Q9JK81 | 6 | 18 | 6 | 380 | 42.7 | 7.02 | 21 | 0.542 | 0.159 | 0.466 | 0.078 | 0.561 | 0.219 | 0.830 | 0.687 |
| 1916 | **Gfm1** | Unchanged | Q8K0D5 | 12 | 78 | 12 | 751 | 83.5 | 6.92 | 21 | 1.155 | 0.597 | 1.092 | 0.635 | 0.671 | 0.125 | 1.628 | 0.000 |

| | | | | | | | | | | | | | | | | | | |
|---|---|---|---|---|---|---|---|---|---|---|---|---|---|---|---|---|---|---|
| 1917 | **Dffb** | Unchanged | A0A1W2P7L3 | 1 | 1 | 1 | 160 | 18 | 5.78 | 21 | #DIV/0! | #DIV/0! | #DIV/0! | #DIV/0! | #DIV/0! | #DIV/0! | #DIV/0! | #DIV/0! |
| 1918 | **Impdh2** | Unchanged | P24547 | 7 | 14 | 7 | 514 | 55.8 | 7.28 | 21 | 0.809 | 0.586 | 1.202 | 0.723 | 0.925 | 0.844 | 1.300 | 0.625 |
| 1919 | **Ppp2r5e** | Unchanged | Q61151 | 5 | 18 | 5 | 467 | 54.7 | 6.95 | 21 | 1.336 | 0.542 | 3.601 | 0.200 | 1.106 | 0.602 | 3.257 | 0.217 |
| 1920 | **Surf4** | Unchanged | Q64310 | 4 | 38 | 4 | 269 | 30.4 | 7.78 | 14 | 1.186 | 0.776 | 1.227 | 0.729 | 0.895 | 0.849 | 1.371 | 0.503 |
| 1921 | **Rprd1b** | Unchanged | Q9CSU0 | 4 | 13 | 4 | 326 | 36.9 | 5.97 | 21 | 0.997 | 0.993 | 1.499 | 0.425 | 0.511 | 0.191 | 2.931 | 0.100 |
| 1922 | **Rab6a** | Unchanged | P35279 | 4 | 28 | 3 | 208 | 23.6 | 5.54 | 21 | 1.608 | 0.365 | 2.774 | 0.065 | 1.671 | 0.236 | 1.660 | 0.147 |
| 1923 | **Rbm39** | Unchanged | Q8VH51 | 7 | 31 | 7 | 530 | 59.4 | 10.1 | 21 | 1.714 | 0.309 | 5.542 | 0.064 | 1.520 | 0.371 | 3.646 | 0.099 |
| 1924 | **Mrpl1** | Down | Q99N96 | 4 | 13 | 4 | 336 | 37.6 | 8.72 | 21 | 0.340 | 0.161 | 0.144 | 0.053 | 0.111 | 0.045 | 1.296 | 0.534 |
| 1925 | **Pcna** | Unchanged | P17918 | 3 | 23 | 3 | 261 | 28.8 | 4.77 | 21 | 0.964 | 0.929 | 0.978 | 0.968 | 1.111 | 0.840 | 0.881 | 0.842 |
| 1926 | **Auh** | Down | Q9JLZ3 | 5 | 37 | 5 | 314 | 33.4 | 9.51 | 21 | 0.028 | 0.045 | 0.065 | 0.052 | 0.023 | 0.044 | 2.879 | 0.096 |
| 1927 | **Ap2m1 iso** | Unchanged | A0A338P6T4 | 1 | 6 | 1 | 119 | 13.7 | 5.2 | 21 | 0.974 | 0.968 | 0.425 | 0.352 | 1.278 | 0.724 | 0.333 | 0.137 |
| 1928 | **Dhx15** | UP | O35286 | 11 | 60 | 11 | 795 | 90.9 | 7.46 | 21 | 1.289 | 0.286 | 1.304 | 0.336 | 1.906 | 0.001 | 0.684 | 0.092 |
| 1929 | **Psmb9** | Down | A0A0R4J256 | 4 | 4 | 4 | 219 | 23.3 | 5.01 | 21 | 0.698 | 0.433 | 0.327 | 0.045 | 0.432 | 0.081 | 0.759 | 0.569 |
| 1930 | **Nipsnap2** | Down | O55126 | 4 | 14 | 4 | 281 | 32.9 | 9.26 | 21 | 0.374 | 0.006 | 0.595 | 0.048 | 0.310 | 0.003 | 1.917 | 0.031 |
| 1931 | **Ssr1** | Unchanged | Q9CY50 | 4 | 58 | 4 | 286 | 32 | 4.45 | 21 | 1.251 | 0.551 | 1.528 | 0.534 | 1.321 | 0.439 | 1.157 | 0.816 |
| 1932 | **Clpx** | Unchanged | Q9JHS4 | 8 | 26 | 8 | 634 | 69.2 | 7.71 | 21 | 1.076 | 0.825 | 1.199 | 0.570 | 0.619 | 0.191 | 1.937 | 0.113 |
| 1933 | **Mrps28** | Unchanged | Q9CY16 | 2 | 31 | 2 | 186 | 20.5 | 8.92 | 21 | 1.214 | 0.634 | 1.607 | 0.183 | 0.919 | 0.858 | 1.749 | 0.077 |
| 1934 | **Jup** | Unchanged | Q02257 | 10 | 29 | 8 | 745 | 81.7 | 6.14 | 21 | 1.396 | 0.314 | 2.281 | 0.079 | 1.216 | 0.376 | 1.876 | 0.130 |
| 1935 | **Iglv1** | Unchanged | P01725 | 1 | 2 | 1 | 110 | 11.6 | 8 | 21 | 0.317 | 0.183 | 0.581 | 0.389 | 0.482 | 0.289 | 1.206 | 0.458 |
| 1936 | **Pvalb** | Unchanged | P32848 | 2 | 3 | 2 | 110 | 11.9 | 5.19 | 21 | 0.198 | 0.095 | 0.412 | 0.202 | 0.341 | 0.162 | 1.209 | 0.569 |
| 1937 | **Cyp3a25** | UP | O09158 | 8 | 34 | 7 | 503 | 58.1 | 8.81 | 21 | 1.647 | 0.141 | 1.932 | 0.076 | 4.859 | 0.007 | 0.397 | 0.032 |
| 1938 | **Azgp1** | UP | Q64726 | 6 | 50 | 6 | 307 | 35.3 | 6.18 | 21 | 1.077 | 0.644 | 2.223 | 0.014 | 0.984 | 0.939 | 2.260 | 0.022 |
| 1939 | **Cald1 iso** | Unchanged | E9QA15 | 16 | 37 | 1 | 768 | 89.2 | 5.5 | 21 | 0.843 | 0.728 | 1.323 | 0.662 | 1.044 | 0.929 | 1.267 | 0.725 |
| 1940 | **Clu** | Unchanged | Q06890 | 7 | 59 | 7 | 448 | 51.6 | 5.67 | 21 | 0.991 | 0.974 | 1.392 | 0.402 | 1.166 | 0.559 | 1.194 | 0.613 |
| 1941 | **Sfpq** | Unchanged | Q8VIJ6 | 10 | 57 | 10 | 699 | 75.4 | 9.44 | 21 | 2.705 | 0.116 | 1.470 | 0.281 | 0.688 | 0.432 | 2.136 | 0.004 |
| 1942 | **Rbks** | Unchanged | Q8R1Q9 | 2 | 11 | 2 | 323 | 34.1 | 5.47 | 21 | 1.967 | 0.066 | 4.204 | 0.159 | 0.969 | 0.815 | 4.336 | 0.155 |
| 1943 | **Dhcr24** | UP | Q8VCH6 | 8 | 43 | 8 | 516 | 60.1 | 8.16 | 21 | 3.958 | 0.246 | 5.269 | 0.078 | 3.701 | 0.002 | 1.424 | 0.495 |
| 1944 | **Acad12** | Unchanged | D3Z7X0 | 10 | 36 | 1 | 555 | 61.4 | 9.36 | 21 | 0.677 | 0.569 | 0.465 | 0.276 | 0.210 | 0.113 | 2.213 | 0.081 |
| 1945 | **Mrpl44** | Unchanged | Q9CY73 | 4 | 19 | 4 | 333 | 37.5 | 8.51 | 21 | 1.614 | 0.246 | 2.487 | 0.096 | 2.083 | 0.140 | 1.194 | 0.690 |
| 1946 | **Acnat2** | Down | Q8BGG9 | 5 | 13 | 2 | 420 | 46.6 | 7.97 | 16 | 0.701 | 0.451 | 0.550 | 0.205 | 0.268 | 0.029 | 2.056 | 0.183 |
| 1947 | **Hax1** | Unchanged | O35387 | 3 | 17 | 3 | 280 | 31.6 | 4.91 | 21 | 0.338 | 0.180 | 0.196 | 0.084 | 0.083 | 0.053 | 2.358 | 0.119 |
| 1948 | **Ipo5** | Unchanged | Q8BKC5 | 15 | 51 | 15 | 1097 | 123.5 | 4.93 | 21 | 0.917 | 0.802 | 0.654 | 0.140 | 1.344 | 0.146 | 0.487 | 0.016 |
| 1949 | **Gm4951** | Unchanged | Q3UED7 | 6 | 26 | 3 | 406 | 47.2 | 5.71 | 21 | 1.626 | 0.537 | 1.757 | 0.483 | 1.655 | 0.428 | 1.062 | 0.928 |
| 1950 | **Dynlt3** | Unchanged | P56387 | 1 | 6 | 1 | 116 | 12.9 | 5.12 | 21 | 0.844 | 0.804 | 2.879 | 0.236 | 0.265 | 0.170 | 10.881 | 0.097 |
| 1951 | **Dazap1** | Unchanged | Q9JII5 | 5 | 41 | 5 | 406 | 43.2 | 8.56 | 21 | 0.938 | 0.698 | 1.180 | 0.528 | 0.978 | 0.935 | 1.206 | 0.586 |
| 1952 | **Serbp1** | Unchanged | Q9CY58 | 8 | 41 | 8 | 407 | 44.7 | 8.54 | 21 | 0.882 | 0.662 | 1.084 | 0.833 | 1.469 | 0.420 | 0.738 | 0.557 |
| 1953 | **Cpq** | Unchanged | Q9WVJ3 | 6 | 63 | 6 | 470 | 51.8 | 6.21 | 21 | 0.942 | 0.779 | 1.006 | 0.980 | 1.004 | 0.981 | 1.002 | 0.992 |
| 1954 | **Rab32** | Unchanged | Q9CZE3 | 3 | 9 | 3 | 223 | 25.1 | 6.68 | 21 | 0.448 | 0.298 | 0.978 | 0.973 | 0.466 | 0.299 | 2.099 | 0.288 |
| 1955 | **Snrnp200** | Unchanged | Q6P4T2 | 27 | 120 | 27 | 2136 | 244.4 | 6.06 | 21 | 1.057 | 0.850 | 1.276 | 0.413 | 1.218 | 0.402 | 1.047 | 0.834 |
| 1956 | **Matr3** | Unchanged | Q8K310 | 11 | 65 | 11 | 846 | 94.6 | 6.25 | 21 | 1.712 | 0.177 | 1.951 | 0.295 | 0.741 | 0.554 | 2.633 | 0.150 |
| 1957 | **Eif4h** | Unchanged | Q9WUK2 | 4 | 28 | 4 | 248 | 27.3 | 7.23 | 21 | 1.260 | 0.659 | 1.691 | 0.225 | 1.690 | 0.226 | 1.001 | 0.999 |

| | | | | | | | | | | | | | | | | | | |
|---|---|---|---|---|---|---|---|---|---|---|---|---|---|---|---|---|---|---|
| 1958 | **Chmp5** | Unchanged | Q9D7S9 | 2 | 13 | 2 | 219 | 24.6 | 4.79 | 21 | 0.839 | 0.727 | 1.428 | 0.290 | 1.828 | 0.304 | 0.781 | 0.614 |
| 1959 | **Itih1** | Unchanged | Q61702 | 8 | 60 | 8 | 907 | 101 | 6.96 | 21 | 1.714 | 0.125 | 1.350 | 0.341 | 1.206 | 0.590 | 1.120 | 0.729 |
| 1960 | **Mvk** | UP | Q9R008 | 3 | 8 | 3 | 395 | 41.9 | 6.68 | 21 | 5.758 | 0.195 | 5.004 | 0.045 | 2.292 | 0.241 | 2.184 | 0.173 |
| 1961 | **Ppil1** | Unchanged | Q9D0W5 | 2 | 5 | 2 | 166 | 18.2 | 7.99 | 20 | 1.086 | 0.918 | 1.280 | 0.772 | 2.015 | 0.274 | 0.635 | 0.472 |
| 1962 | **Apoc1** | Down | P34928 | 2 | 12 | 2 | 88 | 9.7 | 9.09 | 20 | 0.256 | 0.001 | 0.363 | 0.001 | 0.496 | 0.056 | 0.732 | 0.564 |
| 1963 | **Brox** | Unchanged | Q8K2Q7 | 4 | 19 | 4 | 411 | 46.2 | 7.69 | 20 | 1.170 | 0.712 | 1.181 | 0.791 | 1.570 | 0.270 | 0.753 | 0.534 |
| 1964 | **Sptbn2** | Unchanged | Q68FG2 | 31 | 79 | 28 | 2388 | 270.8 | 5.86 | 20 | 0.851 | 0.495 | 1.339 | 0.467 | 0.523 | 0.055 | 2.562 | 0.080 |
| 1965 | **Sh3bgrl3** | Unchanged | Q91VW3 | 1 | 12 | 1 | 93 | 10.5 | 5.14 | 20 | 1.036 | 0.966 | 2.056 | 0.479 | 0.250 | 0.255 | 8.228 | 0.198 |
| 1966 | **Asph** | Unchanged | Q8BSY0 | 10 | 105 | 10 | 741 | 83 | 5.08 | 20 | 0.859 | 0.675 | 1.126 | 0.715 | 0.877 | 0.665 | 1.283 | 0.398 |
| 1967 | **Fau** | Unchanged | P62862 | 3 | 24 | 3 | 59 | 6.6 | 12.15 | 20 | 0.843 | 0.743 | 0.800 | 0.684 | 0.605 | 0.384 | 1.323 | 0.649 |
| 1968 | **Hdac1** | Down | O09106 | 5 | 21 | 3 | 482 | 55 | 5.48 | 20 | 0.309 | 0.014 | 0.183 | 0.005 | 0.151 | 0.003 | 1.216 | 0.531 |
| 1969 | **Slc26a1** | Unchanged | P58735 | 8 | 61 | 8 | 704 | 75.7 | 8.09 | 20 | 1.047 | 0.895 | 0.547 | 0.156 | 0.547 | 0.112 | 1.001 | 0.998 |
| 1970 | **Fam136a** | Down | Q9CR98 | 2 | 13 | 2 | 138 | 15.7 | 7.61 | 20 | 1.153 | 0.673 | 0.309 | 0.008 | 3.503 | 0.151 | 0.088 | 0.076 |
| 1971 | **Ncln** | Unchanged | Q8VCM8 | 8 | 35 | 1 | 563 | 62.9 | 6.49 | 20 | 1.383 | 0.552 | 1.840 | 0.250 | 0.992 | 0.989 | 1.854 | 0.274 |
| 1972 | **Nudt3** | Unchanged | Q9JI46 | 2 | 4 | 2 | 168 | 19 | 6.34 | 20 | #DIV/0! | #DIV/0! | #DIV/0! | #DIV/0! | #DIV/0! | #DIV/0! | #DIV/0! | #DIV/0! |
| 1973 | **Dnajc11** | UP | Q5U458 | 6 | 31 | 6 | 559 | 63.2 | 8.32 | 20 | 1.752 | 0.125 | 1.748 | 0.119 | 1.950 | 0.042 | 0.896 | 0.732 |
| 1974 | **Gid8** | Unchanged | Q9D7M1 | 2 | 3 | 2 | 228 | 26.8 | 4.97 | 20 | 1.962 | 0.147 | 4.573 | 0.069 | 1.626 | 0.255 | 2.812 | 0.134 |
| 1975 | **Tspan4** | Unchanged | Q9DCK3 | 2 | 6 | 2 | 238 | 26 | 5.48 | 20 | 1.995 | 0.423 | 4.115 | 0.212 | 0.909 | 0.779 | 4.526 | 0.200 |
| 1976 | **Lrrc47** | Unchanged | Q505F5 | 8 | 37 | 8 | 581 | 63.6 | 8.1 | 20 | 0.723 | 0.466 | 0.954 | 0.930 | 0.465 | 0.102 | 2.052 | 0.308 |
| 1977 | **Cd47** | Unchanged | Q61735 | 3 | 17 | 3 | 303 | 33.1 | 8.63 | 20 | 2.488 | 0.066 | 3.057 | 0.175 | 0.722 | 0.496 | 4.234 | 0.125 |
| 1978 | **Mrpl50** | UP | Q8VDT9 | 3 | 9 | 3 | 159 | 18.2 | 9.33 | 20 | 5.984 | 0.047 | 4.293 | 0.092 | 8.029 | 0.030 | 0.535 | 0.269 |
| 1979 | **Nclniso** | Unchanged | D3YU17 | 8 | 20 | 1 | 562 | 62.7 | 6.49 | 20 | 0.998 | 0.996 | 0.714 | 0.565 | 0.739 | 0.497 | 0.966 | 0.956 |
| 1980 | **Ndufb5** | Unchanged | Q9CQH3 | 5 | 50 | 5 | 189 | 21.7 | 9.41 | 20 | 0.932 | 0.820 | 1.796 | 0.073 | 0.895 | 0.687 | 2.006 | 0.019 |
| 1981 | **Tmed5** | Unchanged | Q9CXE7 | 4 | 17 | 4 | 229 | 26.2 | 4.93 | 20 | 0.911 | 0.912 | 2.450 | 0.209 | 1.473 | 0.522 | 1.664 | 0.292 |
| 1982 | **Sec23b** | Unchanged | Q9D662 | 12 | 113 | 8 | 767 | 86.4 | 6.96 | 20 | 1.859 | 0.068 | 1.712 | 0.164 | 1.473 | 0.066 | 1.162 | 0.629 |
| 1983 | **Capn1** | Unchanged | O35350 | 10 | 38 | 10 | 713 | 82.1 | 5.87 | 20 | 1.576 | 0.079 | 2.102 | 0.181 | 1.700 | 0.202 | 1.236 | 0.661 |
| 1984 | **Gna11** | Unchanged | P21278 | 6 | 43 | 3 | 359 | 42 | 5.97 | 20 | 0.712 | 0.486 | 0.862 | 0.794 | 0.668 | 0.480 | 1.291 | 0.746 |
| 1985 | **Pef1** | Unchanged | Q8BFY6 | 3 | 18 | 3 | 275 | 29.2 | 6.3 | 20 | 0.474 | 0.215 | 1.715 | 0.483 | 0.683 | 0.427 | 2.512 | 0.299 |
| 1986 | **Lsm2** | Unchanged | O35900 | 1 | 34 | 1 | 95 | 10.8 | 6.52 | 20 | 1.476 | 0.266 | 1.310 | 0.658 | 1.306 | 0.484 | 1.003 | 0.996 |
| 1987 | **Hbq1a** | Unchanged | Q8BYM1 | 1 | 2 | 1 | 145 | 15.9 | 6.61 | 20 | 0.083 | 0.129 | 0.157 | 0.161 | 0.431 | 0.349 | 0.363 | 0.189 |
| 1988 | **Btd** | UP | Q8CIF4 | 6 | 29 | 6 | 520 | 58.1 | 5.8 | 20 | 0.957 | 0.906 | 3.499 | 0.015 | 1.367 | 0.519 | 2.559 | 0.052 |
| 1989 | **Rpl29** | Unchanged | P47915 | 3 | 28 | 3 | 160 | 17.6 | 11.84 | 20 | 1.109 | 0.897 | 0.679 | 0.593 | 0.418 | 0.274 | 1.623 | 0.585 |
| 1990 | **Prkag1** | UP | O54950 | 3 | 56 | 3 | 330 | 37.5 | 7.14 | 20 | 1.400 | 0.418 | 2.534 | 0.036 | 1.325 | 0.461 | 1.912 | 0.056 |
| 1991 | **Uap1l1** | Unchanged | Q3TW96 | 7 | 57 | 7 | 507 | 56.6 | 5.43 | 20 | 0.735 | 0.140 | 1.239 | 0.628 | 1.237 | 0.349 | 1.002 | 0.997 |
| 1992 | **Trim28** | Unchanged | Q62318 | 7 | 28 | 7 | 834 | 88.8 | 5.77 | 20 | 0.465 | 0.119 | 0.803 | 0.736 | 0.546 | 0.169 | 1.472 | 0.614 |
| 1993 | **Rala** | Unchanged | P63321 | 3 | 24 | 3 | 206 | 23.5 | 7.11 | 20 | 3.270 | 0.169 | 3.431 | 0.187 | 3.676 | 0.082 | 0.933 | 0.909 |
| 1994 | **Rab35** | Unchanged | Q6PHN9 | 3 | 41 | 1 | 201 | 23 | 8.29 | 20 | 0.910 | 0.818 | 2.547 | 0.111 | 1.846 | 0.079 | 1.380 | 0.423 |
| 1995 | **Eml2** | Unchanged | Q7TNG5 | 9 | 23 | 9 | 649 | 70.7 | 6.28 | 19 | 1.468 | 0.259 | 1.710 | 0.303 | 1.475 | 0.411 | 1.159 | 0.775 |
| 1996 | **Sfxn2** | UP | Q925N2 | 4 | 29 | 4 | 322 | 36.1 | 8.82 | 20 | 1.976 | 0.130 | 2.737 | 0.190 | 2.435 | 0.016 | 1.124 | 0.812 |
| 1997 | **Etnppl** | Unchanged | Q8BWU8 | 7 | 48 | 7 | 499 | 55.5 | 6.93 | 20 | 5.001 | 0.066 | 3.313 | 0.083 | 18.975 | 0.157 | 0.175 | 0.212 |
| 1998 | **Ppp5c** | Unchanged | Q60676 | 7 | 31 | 7 | 499 | 56.8 | 6.2 | 20 | 1.713 | 0.139 | 4.308 | 0.120 | 1.298 | 0.457 | 3.318 | 0.156 |

| | | | | | | | | | | | | | | | | | | |
|---|---|---|---|---|---|---|---|---|---|---|---|---|---|---|---|---|---|---|
| 1999 | **Higd2a** | UP | Q9CQJ1 | 1 | 6 | 1 | 106 | 11.4 | 9.57 | 20 | 1.300 | 0.472 | 2.889 | 0.009 | 3.768 | 0.002 | 0.767 | 0.221 |
| 2000 | **Hspa4l** | Unchanged | P48722 | 11 | 37 | 8 | 838 | 94.3 | 5.74 | 20 | 0.970 | 0.937 | 1.628 | 0.170 | 1.233 | 0.522 | 1.321 | 0.255 |
| 2001 | **Rbpms** | Unchanged | Q9WVB0 | 3 | 11 | 2 | 197 | 21.8 | 8.07 | 20 | 0.935 | 0.908 | 4.516 | 0.172 | 3.220 | 0.245 | 1.403 | 0.661 |
| 2002 | **Tapbp** | UP | Q9R233 | 5 | 23 | 5 | 465 | 49.7 | 8.5 | 20 | 0.972 | 0.947 | 3.843 | 0.048 | 1.447 | 0.524 | 2.655 | 0.115 |
| 2003 | **Camp** | Down | P51437 | 2 | 25 | 2 | 172 | 19.4 | 8.68 | 20 | 0.642 | 0.193 | 0.495 | 0.052 | 0.329 | 0.000 | 1.506 | 0.487 |
| 2004 | **Man2b1** | Unchanged | O09159 | 14 | 81 | 14 | 1013 | 114.6 | 8.13 | 20 | 1.135 | 0.669 | 1.859 | 0.205 | 1.572 | 0.220 | 1.183 | 0.697 |
| 2005 | **Ap2s1** | Down | P62743 | 3 | 10 | 3 | 142 | 17 | 6.18 | 20 | 0.871 | 0.765 | 0.585 | 0.324 | 0.151 | 0.014 | 3.876 | 0.174 |
| 2006 | **Fabp2** | Unchanged | P55050 | 2 | 47 | 2 | 132 | 15.1 | 7.24 | 20 | 1.295 | 0.535 | 0.459 | 0.127 | 0.392 | 0.073 | 1.171 | 0.689 |
| 2007 | **Gm11214** | Unchanged | V9GX06 | 3 | 59 | 1 | 183 | 19.9 | 7.42 | 20 | #DIV/0! | #DIV/0! | #DIV/0! | #DIV/0! | #DIV/0! | #DIV/0! | #DIV/0! | #DIV/0! |
| 2008 | **Rpl22l1** | Unchanged | Q9D7S7 | 1 | 8 | 1 | 122 | 14.5 | 9.45 | 20 | 1.397 | 0.782 | 1.384 | 0.764 | 1.920 | 0.317 | 0.721 | 0.584 |
| 2009 | **Mt2** | Unchanged | P02798 | 1 | 2 | 1 | 61 | 6.1 | 7.83 | 20 | 0.061 | 0.334 | 0.053 | 0.330 | 0.060 | 0.334 | 0.876 | 0.775 |
| 2010 | **Mt1** | Unchanged | P02802 | 1 | 2 | 1 | 61 | 6 | 7.96 | 20 | #DIV/0! | #DIV/0! | #DIV/0! | #DIV/0! | #DIV/0! | #DIV/0! | #DIV/0! | #DIV/0! |
| 2011 | **Cox20** | Unchanged | Q9D7J4 | 1 | 7 | 1 | 117 | 13.2 | 9.01 | 20 | 2.597 | 0.391 | 7.450 | 0.134 | 5.043 | 0.183 | 1.477 | 0.626 |
| 2012 | **Sf3b3** | Unchanged | Q921M3 | 15 | 107 | 15 | 1217 | 135.5 | 5.26 | 20 | 1.120 | 0.632 | 1.582 | 0.216 | 0.865 | 0.594 | 1.829 | 0.142 |
| 2013 | **Abcc2** | UP | Q8VI47 | 16 | 92 | 16 | 1543 | 173.6 | 8.65 | 20 | 1.610 | 0.009 | 1.972 | 0.013 | 1.945 | 0.008 | 1.014 | 0.941 |
| 2014 | **Lsm3** | Unchanged | P62311 | 2 | 17 | 2 | 102 | 11.8 | 4.7 | 20 | 0.816 | 0.459 | 0.808 | 0.670 | 1.221 | 0.385 | 0.661 | 0.352 |
| 2015 | **Rpl39** | Unchanged | P62892 | 1 | 4 | 1 | 51 | 6.4 | 12.56 | 20 | 1.384 | 0.698 | 0.698 | 0.705 | 0.606 | 0.535 | 1.152 | 0.881 |
| 2016 | **G6pc** | UP | P35576 | 4 | 30 | 4 | 357 | 40.4 | 9.01 | 20 | 3.347 | 0.079 | 7.212 | 0.185 | 3.478 | 0.040 | 2.073 | 0.419 |
| 2017 | **Pitrm1** | Unchanged | Q8K411 | 12 | 63 | 12 | 1036 | 117.3 | 7.2 | 20 | 0.875 | 0.709 | 0.793 | 0.566 | 0.786 | 0.495 | 1.009 | 0.979 |
| 2018 | **Rbbp7** | UP | Q60973 | 5 | 16 | 3 | 425 | 47.8 | 5.05 | 20 | 2.280 | 0.121 | 2.826 | 0.012 | 3.270 | 0.017 | 0.864 | 0.629 |
| 2019 | **Cul4a** | Unchanged | Q3TCH7 | 10 | 28 | 5 | 759 | 87.7 | 8.35 | 19 | 1.543 | 0.469 | 0.612 | 0.355 | 0.969 | 0.968 | 0.632 | 0.617 |
| 2020 | **Dad1** | Unchanged | P61804 | 2 | 15 | 2 | 113 | 12.5 | 7.08 | 19 | 0.798 | 0.399 | 0.856 | 0.745 | 1.034 | 0.899 | 0.828 | 0.721 |
| 2021 | **Gpx3** | Unchanged | P46412 | 3 | 11 | 3 | 226 | 25.4 | 8.22 | 19 | 1.250 | 0.404 | 0.612 | 0.091 | 1.354 | 0.412 | 0.452 | 0.117 |
| 2022 | **Hnrnpr** | Unchanged | Q8VHM5 | 10 | 35 | 6 | 632 | 70.8 | 8.13 | 19 | 2.410 | 0.241 | 1.466 | 0.392 | 1.530 | 0.302 | 0.958 | 0.899 |
| 2023 | **Apoh** | Unchanged | Q01339 | 5 | 28 | 5 | 345 | 38.6 | 8.22 | 19 | 0.935 | 0.869 | 1.861 | 0.334 | 2.205 | 0.125 | 0.844 | 0.749 |
| 2024 | **Mmab** | Unchanged | Q9D273 | 3 | 19 | 3 | 237 | 26.3 | 9.2 | 19 | 0.595 | 0.401 | 1.074 | 0.904 | 1.013 | 0.980 | 1.061 | 0.903 |
| 2025 | **Gm16971;Ighv5-4** | Unchanged | P18528 | 1 | 11 | 1 | 98 | 11 | 7.93 | 19 | 0.435 | 0.053 | 1.779 | 0.062 | 0.679 | 0.333 | 2.621 | 0.019 |
| 2026 | **Ighv1-5** | Unchanged | A0A075B5T5 | 1 | 2 | 1 | 98 | 10.8 | 9.07 | 19 | 0.878 | 0.888 | 0.682 | 0.706 | 0.393 | 0.448 | 1.736 | 0.361 |
| 2027 | **Mtfr1l** | UP | Q9CWE0 | 3 | 5 | 3 | 289 | 31.7 | 6.1 | 19 | 1.930 | 0.178 | 2.344 | 0.021 | 1.442 | 0.279 | 1.625 | 0.088 |
| 2028 | **Acot8** | Unchanged | P58137 | 4 | 36 | 4 | 320 | 35.8 | 7.64 | 19 | 0.633 | 0.236 | 2.188 | 0.512 | 0.512 | 0.089 | 4.270 | 0.359 |
| 2029 | **Hmox2** | Unchanged | O70252 | 4 | 14 | 4 | 315 | 35.7 | 5.87 | 19 | 1.610 | 0.146 | 0.865 | 0.654 | 1.211 | 0.631 | 0.714 | 0.307 |
| 2030 | **Gimd1** | Unchanged | E9PW74 | 2 | 6 | 2 | 217 | 24.2 | 6.61 | 19 | 0.835 | 0.553 | 1.548 | 0.301 | 1.223 | 0.469 | 1.266 | 0.515 |
| 2031 | **Clec4f** | Unchanged | P70194 | 8 | 34 | 8 | 548 | 61.2 | 7.15 | 19 | 0.996 | 0.992 | 1.164 | 0.788 | 0.881 | 0.787 | 1.320 | 0.664 |
| 2032 | **Lyn** | UP | P25911 | 7 | 25 | 6 | 512 | 58.8 | 7.15 | 19 | 1.295 | 0.482 | 1.595 | 0.021 | 1.141 | 0.743 | 1.398 | 0.312 |
| 2033 | **Fxn** | Unchanged | O35943 | 3 | 12 | 3 | 207 | 22.9 | 8.02 | 19 | 1.200 | 0.613 | 1.041 | 0.935 | 0.328 | 0.070 | 3.171 | 0.104 |
| 2034 | **Upf1** | UP | Q9EPU0 | 15 | 105 | 15 | 1124 | 123.9 | 6.61 | 19 | 1.693 | 0.165 | 1.986 | 0.033 | 2.472 | 0.071 | 0.804 | 0.560 |
| 2035 | **2310039H08Rik** | Unchanged | Q9D727 | 1 | 1 | 1 | 114 | 12 | 6.29 | 19 | #DIV/0! | #DIV/0! | #DIV/0! | #DIV/0! | #DIV/0! | #DIV/0! | #DIV/0! | #DIV/0! |
| 2036 | **Ndrg1** | UP | Q62433 | 4 | 11 | 4 | 394 | 43 | 6.1 | 19 | 2.882 | 0.194 | 2.791 | 0.152 | 3.120 | 0.027 | 0.895 | 0.811 |
| 2037 | **Diaph1** | UP | O08808 | 14 | 64 | 14 | 1255 | 139.3 | 5.57 | 19 | 0.949 | 0.865 | 1.957 | 0.041 | 1.262 | 0.429 | 1.550 | 0.093 |
| 2038 | **Dync1i2** | Unchanged | O88487 | 5 | 29 | 5 | 612 | 68.4 | 5.29 | 19 | 2.229 | 0.195 | 1.005 | 0.991 | 1.017 | 0.954 | 0.989 | 0.983 |
| 2039 | **Mrps27** | Down | Q8BK72 | 6 | 24 | 6 | 415 | 47.7 | 5.5 | 19 | 0.733 | 0.457 | 1.110 | 0.670 | 0.388 | 0.035 | 2.858 | 0.008 |

| | | | | | | | | | | | | | | | | | | |
|---|---|---|---|---|---|---|---|---|---|---|---|---|---|---|---|---|---|---|
| 2040 | **Mrpl49** | Down | Q9CQ40 | 3 | 25 | 3 | 166 | 19.1 | 9.5 | 19 | 0.755 | 0.269 | 0.583 | 0.082 | 0.518 | 0.028 | 1.125 | 0.699 |
| 2041 | **Mrpl54** | Unchanged | Q9CPW3 | 1 | 2 | 1 | 135 | 15.4 | 10.04 | 19 | 0.941 | 0.709 | 1.490 | 0.355 | 1.669 | 0.577 | 0.893 | 0.889 |
| 2042 | **Vkorc1** | Unchanged | Q9CRC0 | 2 | 18 | 2 | 161 | 17.8 | 9.14 | 19 | 0.791 | 0.669 | 0.873 | 0.800 | 0.303 | 0.156 | 2.881 | 0.110 |
| 2043 | **Denr** | Unchanged | Q9CQJ6 | 2 | 34 | 2 | 198 | 22.2 | 5.3 | 19 | 0.906 | 0.795 | 0.679 | 0.381 | 0.764 | 0.537 | 0.888 | 0.794 |
| 2044 | **Tysnd1** | Unchanged | Q9DBA6 | 5 | 11 | 5 | 568 | 59 | 7.23 | 19 | 1.012 | 0.963 | 1.882 | 0.211 | 0.963 | 0.838 | 1.954 | 0.191 |
| 2045 | **Lpgat1** | Unchanged | Q91YX5 | 6 | 9 | 6 | 370 | 43.1 | 8.79 | 19 | 0.434 | 0.152 | 0.654 | 0.447 | 0.459 | 0.194 | 1.427 | 0.556 |
| 2046 | **Ckmt1** | Unchanged | P30275 | 4 | 7 | 2 | 418 | 47 | 8.16 | 19 | 1.081 | 0.645 | 1.412 | 0.419 | 0.650 | 0.055 | 2.174 | 0.165 |
| 2047 | **Snapin** | UP | Q9Z266 | 1 | 1 | 1 | 136 | 14.9 | 9.31 | 19 | 1.306 | 0.728 | 2.339 | 0.047 | 1.353 | 0.289 | 1.729 | 0.132 |
| 2048 | **Tom1** | Unchanged | O88746 | 4 | 29 | 4 | 492 | 54.3 | 4.94 | 19 | 1.002 | 0.996 | 1.316 | 0.493 | 1.572 | 0.351 | 0.837 | 0.628 |
| 2049 | **Pacsin3** | Down | Q99JB8 | 5 | 47 | 5 | 424 | 48.6 | 6.1 | 19 | 0.717 | 0.110 | 0.616 | 0.251 | 0.431 | 0.011 | 1.431 | 0.553 |
| 2050 | **Eppk1** | Unchanged | Q8R0W0 | 13 | 56 | 12 | 6548 | 724.2 | 6.07 | 17 | 1.135 | 0.603 | 1.397 | 0.413 | 1.876 | 0.106 | 0.745 | 0.451 |
| 2051 | **Enoph1** | Unchanged | Q8BGB7 | 2 | 10 | 2 | 257 | 28.6 | 4.92 | 9 | 2.453 | 0.193 | 2.058 | 0.164 | 1.294 | 0.681 | 1.591 | 0.256 |
| 2052 | **Itih2** | Unchanged | Q61703 | 9 | 33 | 9 | 946 | 105.9 | 7.27 | 19 | 1.238 | 0.513 | 1.715 | 0.283 | 1.972 | 0.075 | 0.870 | 0.724 |
| 2053 | **Elavl1** | Unchanged | P70372 | 5 | 63 | 5 | 326 | 36.1 | 9.04 | 19 | 0.903 | 0.687 | 0.875 | 0.620 | 0.833 | 0.467 | 1.051 | 0.865 |
| 2054 | **Rpa3** | Unchanged | Q9CQ71 | 1 | 1 | 1 | 121 | 13.6 | 4.84 | 19 | 0.741 | 0.438 | 0.910 | 0.755 | 2.884 | 0.084 | 0.315 | 0.072 |
| 2055 | **Fdft1** | Unchanged | P53798 | 4 | 10 | 4 | 416 | 48.1 | 6.32 | 19 | 1.625 | 0.488 | 0.265 | 0.133 | 0.697 | 0.646 | 0.381 | 0.379 |
| 2056 | **Ero1a** | Unchanged | Q8R180 | 5 | 24 | 5 | 464 | 54.1 | 6.54 | 19 | 0.867 | 0.509 | 0.869 | 0.696 | 0.715 | 0.405 | 1.215 | 0.731 |
| 2057 | **Map2k2** | Down | Q63932 | 5 | 26 | 2 | 401 | 44.4 | 7.05 | 19 | 0.605 | 0.189 | 1.238 | 0.662 | 0.454 | 0.020 | 2.729 | 0.151 |
| 2058 | **Pcmt1** | Unchanged | P23506 | 3 | 17 | 3 | 227 | 24.6 | 7.65 | 19 | 0.652 | 0.109 | 0.553 | 0.155 | 0.860 | 0.682 | 0.643 | 0.409 |
| 2059 | **Ebpl** | Unchanged | Q9D0P0 | 2 | 6 | 2 | 206 | 23.3 | 6.89 | 4 | 1.638 | 0.270 | 4.715 | 0.176 | 1.555 | 0.486 | 3.032 | 0.252 |
| 2060 | **Pon2** | Unchanged | Q62086 | 5 | 31 | 5 | 354 | 39.6 | 5.83 | 19 | 0.724 | 0.343 | 1.282 | 0.560 | 0.673 | 0.308 | 1.905 | 0.196 |
| 2061 | **Tomm6** | Unchanged | Q9CQN3 | 1 | 9 | 1 | 74 | 7.9 | 4.89 | 19 | 1.780 | 0.324 | 1.922 | 0.207 | 1.376 | 0.439 | 1.397 | 0.513 |
| 2062 | **Sub1** | Unchanged | P11031 | 2 | 4 | 2 | 127 | 14.4 | 9.6 | 19 | 3.912 | 0.129 | 4.361 | 0.257 | 14.230 | 0.062 | 0.306 | 0.179 |
| 2063 | **Grb2** | Unchanged | Q60631 | 2 | 26 | 2 | 217 | 25.2 | 6.32 | 19 | 0.543 | 0.144 | 0.549 | 0.167 | 0.743 | 0.330 | 0.739 | 0.401 |
| 2064 | **Bloc1s2** | Down | Q9CWG9 | 1 | 4 | 1 | 143 | 16.3 | 4.88 | 19 | 0.397 | 0.027 | 1.361 | 0.233 | 0.404 | 0.027 | 3.368 | 0.011 |
| 2065 | **Snrpc** | Unchanged | Q62241 | 2 | 15 | 2 | 159 | 17.4 | 9.67 | 19 | 1.249 | 0.689 | 2.125 | 0.310 | 0.263 | 0.150 | 8.070 | 0.082 |
| 2066 | **Xpo1** | Unchanged | Q6P5F9 | 14 | 50 | 14 | 1071 | 123 | 6.07 | 19 | 1.269 | 0.615 | 3.275 | 0.126 | 1.224 | 0.668 | 2.676 | 0.166 |
| 2067 | **Ighv1-72** | UP | P01753 | 2 | 11 | 2 | 117 | 12.9 | 9.13 | 19 | 1.129 | 0.853 | 4.816 | 0.039 | 0.563 | 0.470 | 8.553 | 0.018 |
| 2068 | **Itgb1** | Down | P09055 | 11 | 73 | 11 | 798 | 88.2 | 5.94 | 19 | 0.639 | 0.154 | 1.124 | 0.666 | 0.463 | 0.037 | 2.429 | 0.027 |
| 2069 | **Igkv4-81** | Unchanged | A0A075B5L6 | 1 | 1 | 1 | 96 | 10.2 | 5.1 | 19 | 0.431 | 0.181 | 0.357 | 0.132 | 0.497 | 0.229 | 0.717 | 0.270 |
| 2070 | **Stxbp3** | Unchanged | Q60770 | 7 | 32 | 7 | 592 | 67.9 | 8.02 | 19 | 1.261 | 0.420 | 3.722 | 0.129 | 1.530 | 0.127 | 2.432 | 0.210 |
| 2071 | **Khdrbs1** | Unchanged | Q60749 | 5 | 15 | 5 | 443 | 48.3 | 8.72 | 19 | 0.870 | 0.807 | 0.830 | 0.744 | 1.081 | 0.867 | 0.768 | 0.369 |
| 2072 | **Arl6ip1** | Unchanged | Q9JKW0 | 2 | 12 | 2 | 203 | 23.4 | 9.32 | 19 | 2.033 | 0.170 | 1.155 | 0.806 | 6.547 | 0.061 | 0.176 | 0.068 |
| 2073 | **Mat2biso** | Unchanged | E0CYU5 | 2 | 3 | 1 | 102 | 11.7 | 7.47 | 19 | 3.470 | 0.038 | 3.781 | 0.069 | 4.272 | 0.084 | 0.885 | 0.819 |
| 2074 | **Fbxw11** | Unchanged | F7BGU3 | 1 | 2 | 1 | 43 | 5 | 8.31 | 19 | 0.809 | 0.731 | 1.884 | 0.392 | 0.800 | 0.729 | 2.354 | 0.278 |
| 2075 | **Slc35d1** | Unchanged | A2AKQ0 | 3 | 13 | 3 | 355 | 39.2 | 8.84 | 19 | 1.139 | 0.843 | 1.375 | 0.617 | 1.758 | 0.276 | 0.782 | 0.468 |
| 2076 | **Stt3a** | Unchanged | P46978 | 13 | 87 | 12 | 705 | 80.5 | 8.1 | 17 | 0.998 | 0.995 | 0.823 | 0.615 | 0.742 | 0.432 | 1.109 | 0.709 |
| 2077 | **Pabpn1** | Unchanged | Q8CCS6 | 2 | 11 | 2 | 302 | 32.3 | 5.17 | 19 | 1.150 | 0.719 | 1.269 | 0.717 | 1.689 | 0.191 | 0.751 | 0.581 |
| 2078 | **Tat** | Down | Q8QZR1 | 5 | 9 | 5 | 454 | 50.5 | 5.53 | 19 | 1.386 | 0.553 | 0.765 | 0.590 | 0.273 | 0.014 | 2.799 | 0.205 |
| 2079 | **Acox3** | Unchanged | Q9EPL9 | 8 | 32 | 8 | 700 | 78.4 | 7.12 | 18 | 0.778 | 0.292 | 1.121 | 0.671 | 0.862 | 0.422 | 1.301 | 0.356 |
| 2080 | **Tfr2** | Unchanged | Q9JKX3 | 10 | 37 | 10 | 798 | 88.3 | 6.13 | 18 | 0.656 | 0.301 | 2.671 | 0.253 | 2.194 | 0.072 | 1.217 | 0.745 |

| | | | | | | | | | | | | | | | | | | |
|---|---|---|---|---|---|---|---|---|---|---|---|---|---|---|---|---|---|---|
| 2081 | **Mrps6** | Down | P58064 | 3 | 7 | 3 | 125 | 14.3 | 9.5 | 17 | 0.250 | 0.001 | 1.558 | 0.635 | 1.766 | 0.213 | 0.882 | 0.872 |
| 2082 | **Mtnd4l** | Unchanged | P03903 | 1 | 2 | 1 | 98 | 10.6 | 5.71 | 18 | 1.574 | 0.614 | 0.852 | 0.792 | 0.596 | 0.364 | 1.428 | 0.583 |
| 2083 | **Rab8b** | Down | P61028 | 4 | 36 | 1 | 207 | 23.6 | 9.07 | 18 | 0.280 | 0.005 | 0.375 | 0.018 | 0.618 | 0.078 | 0.606 | 0.238 |
| 2084 | **Srsf7** | Unchanged | Q8BL97 | 5 | 49 | 4 | 267 | 30.8 | 11.9 | 18 | 0.683 | 0.259 | 1.094 | 0.836 | 0.580 | 0.191 | 1.885 | 0.223 |
| 2085 | **Nif3l1** | UP | Q9EQ80 | 4 | 34 | 4 | 376 | 41.7 | 6.76 | 18 | 1.595 | 0.275 | 1.605 | 0.345 | 3.046 | 0.023 | 0.527 | 0.158 |
| 2086 | **Prorsd1** | Unchanged | Q9D820 | 2 | 8 | 2 | 169 | 18.9 | 6.14 | 18 | 0.590 | 0.083 | 0.812 | 0.691 | 0.642 | 0.166 | 1.265 | 0.702 |
| 2087 | **Defa6** | Unchanged | P50704 | 2 | 3 | 2 | 93 | 10.4 | 5.01 | 18 | 13.324 | 0.163 | 19.981 | 0.334 | 28.003 | 0.093 | 0.714 | 0.739 |
| 2088 | **Pdcd5** | Unchanged | P56812 | 2 | 12 | 2 | 126 | 14.3 | 5.68 | 18 | 1.369 | 0.416 | 1.342 | 0.464 | 0.986 | 0.977 | 1.361 | 0.410 |
| 2089 | **Osbpl1a** | UP | Q91XL9 | 8 | 32 | 8 | 950 | 107.7 | 6.44 | 18 | 1.931 | 0.147 | 3.185 | 0.042 | 1.878 | 0.089 | 1.696 | 0.161 |
| 2090 | **Alg5** | Down | Q9DB25 | 4 | 10 | 4 | 324 | 36.8 | 8.6 | 18 | 1.251 | 0.576 | 1.658 | 0.523 | 0.146 | 0.023 | 11.357 | 0.145 |
| 2091 | **Gm4756** | Unchanged | A0A140LIS8 | 3 | 10 | 3 | 335 | 37.8 | 7.69 | 18 | 0.497 | 0.269 | 0.328 | 0.161 | 0.279 | 0.113 | 1.179 | 0.768 |
| 2092 | **Sf3a1** | Unchanged | Q8K4Z5 | 9 | 28 | 9 | 791 | 88.5 | 5.22 | 18 | 0.771 | 0.533 | 0.756 | 0.591 | 2.410 | 0.329 | 0.314 | 0.264 |
| 2093 | **Mvd** | Unchanged | Q99JF5 | 4 | 23 | 4 | 401 | 44 | 6.3 | 18 | 0.763 | 0.496 | 1.591 | 0.486 | 0.925 | 0.830 | 1.720 | 0.428 |
| 2094 | **Pdk1** | UP | Q8BFP9 | 5 | 18 | 5 | 434 | 49 | 8.19 | 18 | 1.573 | 0.029 | 2.611 | 0.000 | 1.421 | 0.054 | 1.837 | 0.002 |
| 2095 | **Scarb2** | Unchanged | O35114 | 6 | 75 | 6 | 478 | 54 | 5.1 | 18 | 0.635 | 0.226 | 0.620 | 0.287 | 0.467 | 0.084 | 1.327 | 0.585 |
| 2096 | **Bola3** | Unchanged | Q8CEI1 | 1 | 8 | 1 | 110 | 12.2 | 8.97 | 18 | 0.793 | 0.731 | 0.508 | 0.443 | 0.157 | 0.131 | 3.225 | 0.371 |
| 2097 | **Ces1b** | UP | D3Z5G7 | 8 | 40 | 2 | 567 | 62.2 | 5.22 | 18 | 11.200 | 0.108 | 12.011 | 0.050 | 11.505 | 0.047 | 1.044 | 0.940 |
| 2098 | **Mrpl19** | Unchanged | Q9D338 | 3 | 18 | 3 | 292 | 33.6 | 9.44 | 18 | 0.953 | 0.917 | 0.956 | 0.929 | 0.486 | 0.106 | 1.968 | 0.326 |
| 2099 | **Sfn** | Unchanged | O70456 | 4 | 62 | 1 | 248 | 27.7 | 4.78 | 18 | 1.722 | 0.584 | 0.616 | 0.542 | 0.386 | 0.234 | 1.595 | 0.628 |
| 2100 | **Cnih2** | Unchanged | O35089 | 1 | 1 | 1 | 160 | 18.9 | 7.25 | 18 | 1.450 | 0.656 | 0.387 | 0.431 | 1.269 | 0.764 | 0.305 | 0.098 |
| 2101 | **Cryzl1** | UP | Q921W4 | 3 | 3 | 3 | 348 | 38.7 | 5.97 | 18 | 1.896 | 0.450 | 1.453 | 0.582 | 2.779 | 0.006 | 0.523 | 0.123 |
| 2102 | **Msrb1** | Unchanged | Q9JLC3 | 1 | 1 | 1 | 116 | 12.8 | 8.27 | 18 | 0.317 | 0.141 | 1.721 | 0.505 | 2.635 | 0.062 | 0.653 | 0.445 |
| 2103 | **Pemt** | Unchanged | Q61907 | 2 | 8 | 2 | 199 | 22.6 | 8.6 | 18 | 1.809 | 0.629 | 3.873 | 0.226 | 0.811 | 0.821 | 4.777 | 0.185 |
| 2104 | **Ezr** | Unchanged | P26040 | 11 | 133 | 2 | 586 | 69.4 | 6.1 | 18 | 1.149 | 0.766 | 1.280 | 0.609 | 1.110 | 0.819 | 1.153 | 0.752 |
| 2105 | **Cuta** | Unchanged | Q9CQ89 | 2 | 15 | 2 | 177 | 18.9 | 6.77 | 18 | 1.215 | 0.721 | 1.704 | 0.431 | 1.654 | 0.403 | 1.030 | 0.962 |
| 2106 | **Gsn** | Unchanged | P13020 | 7 | 28 | 7 | 780 | 85.9 | 6.18 | 18 | 0.656 | 0.055 | 0.758 | 0.360 | 1.420 | 0.029 | 0.534 | 0.046 |
| 2107 | **Sirt5** | UP | Q8K2C6 | 4 | 35 | 4 | 310 | 34.1 | 8.53 | 18 | 3.588 | 0.010 | 4.547 | 0.025 | 3.460 | 0.068 | 1.314 | 0.540 |
| 2108 | **Ifitm2** | Unchanged | Q99J93 | 2 | 5 | 1 | 144 | 15.7 | 7.3 | 18 | 4.745 | 0.405 | 7.217 | 0.365 | 4.689 | 0.180 | 1.539 | 0.725 |
| 2109 | **Dhrs7** | Down | Q9CXR1 | 3 | 15 | 3 | 338 | 38.1 | 8.32 | 18 | 0.577 | 0.168 | 0.497 | 0.105 | 0.313 | 0.043 | 1.589 | 0.245 |
| 2110 | **Fgf7** | Unchanged | P36363 | 1 | 1 | 1 | 194 | 22.3 | 9.32 | 18 | #DIV/0! | #DIV/0! | #DIV/0! | #DIV/0! | #DIV/0! | #DIV/0! | #DIV/0! | #DIV/0! |
| 2111 | **Csrp3** | Down | P50462 | 2 | 8 | 2 | 194 | 20.9 | 8.54 | 18 | 0.945 | 0.913 | 0.152 | 0.041 | 0.395 | 0.150 | 0.386 | 0.203 |
| 2112 | **Ilvbl** | Unchanged | Q8BU33 | 6 | 32 | 6 | 632 | 68.1 | 8.69 | 18 | 1.449 | 0.352 | 2.521 | 0.239 | 0.824 | 0.510 | 3.059 | 0.199 |
| 2113 | **Saa1** | Unchanged | P05366 | 2 | 10 | 2 | 122 | 13.8 | 7.03 | 18 | 0.034 | 0.137 | 0.043 | 0.140 | 0.177 | 0.211 | 0.242 | 0.425 |
| 2114 | **Vps4b** | Unchanged | P46467 | 4 | 16 | 4 | 444 | 49.4 | 7.11 | 18 | 1.323 | 0.533 | 1.552 | 0.386 | 1.678 | 0.342 | 0.925 | 0.872 |
| 2115 | **Pla2g2c** | Unchanged | A2APR1 | 1 | 1 | 1 | 150 | 17 | 8.28 | 18 | 0.770 | 0.777 | 0.317 | 0.336 | 0.519 | 0.493 | 0.610 | 0.275 |
| 2116 | **Fus** | Unchanged | P56959 | 7 | 51 | 5 | 518 | 52.6 | 9.36 | 18 | 0.843 | 0.626 | 0.842 | 0.727 | 1.190 | 0.698 | 0.707 | 0.544 |
| 2117 | **Ighv5-4** | UP | A0A075B5P9 | 1 | 1 | 1 | 117 | 13 | 8.6 | 18 | 13.555 | 0.097 | 16.128 | 0.045 | 14.060 | 0.096 | 1.147 | 0.832 |
| 2118 | **Des** | Down | P31001 | 5 | 12 | 3 | 469 | 53.5 | 5.27 | 18 | 0.546 | 0.326 | 0.906 | 0.888 | 0.072 | 0.017 | 12.613 | 0.181 |
| 2119 | **Rad23a** | Unchanged | P54726 | 4 | 13 | 3 | 363 | 39.7 | 4.58 | 18 | 1.563 | 0.400 | 0.753 | 0.692 | 0.594 | 0.465 | 1.267 | 0.774 |
| 2120 | **Cltb** | Unchanged | Q6IRU5 | 6 | 26 | 6 | 229 | 25.2 | 4.63 | 18 | 0.965 | 0.907 | 0.535 | 0.212 | 0.748 | 0.601 | 0.715 | 0.703 |
| 2121 | **Clta** | Unchanged | Q6PFA2 | 3 | 35 | 2 | 218 | 23.6 | 4.5 | 18 | 0.843 | 0.633 | 0.766 | 0.555 | 0.685 | 0.372 | 1.118 | 0.831 |

| | | | | | | | | | | | | | | | | | | |
|---|---|---|---|---|---|---|---|---|---|---|---|---|---|---|---|---|---|---|
| 2122 | **Fn1** | Unchanged | P11276 | 25 | 202 | 25 | 2477 | 272.4 | 5.59 | 18 | 1.010 | 0.972 | 0.841 | 0.513 | 0.894 | 0.679 | 0.941 | 0.818 |
| 2123 | **Lgals8** | Unchanged | Q9JL15 | 4 | 6 | 4 | 316 | 36.1 | 8.95 | 18 | 1.145 | 0.843 | 0.964 | 0.969 | 0.166 | 0.230 | 5.811 | 0.233 |
| 2124 | **Mtco1** | UP | P00397 | 3 | 6 | 3 | 514 | 56.9 | 6.7 | 18 | 20.108 | 0.045 | 30.718 | 0.052 | 31.166 | 0.013 | 0.986 | 0.979 |
| 2125 | **Timm23** | Unchanged | Q9WTQ8 | 2 | 6 | 2 | 209 | 22 | 9.13 | 18 | 0.607 | 0.257 | 0.559 | 0.246 | 0.538 | 0.161 | 1.038 | 0.953 |
| 2126 | **Ykt6** | Unchanged | Q9CQW1 | 2 | 11 | 2 | 198 | 22.3 | 6.35 | 18 | 0.940 | 0.932 | 0.383 | 0.293 | 1.363 | 0.591 | 0.281 | 0.035 |
| 2127 | **Srxn1** | Unchanged | Q9D975 | 1 | 8 | 1 | 136 | 14.1 | 8.02 | 18 | 0.281 | 0.318 | 3.327 | 0.380 | 0.865 | 0.853 | 3.844 | 0.339 |
| 2128 | **Anp32b** | Unchanged | Q9EST5 | 4 | 37 | 2 | 272 | 31.1 | 4.01 | 18 | 0.834 | 0.837 | 1.114 | 0.910 | 0.300 | 0.280 | 3.713 | 0.324 |
| 2129 | **Epb41** | Unchanged | F6S4K9 | 5 | 14 | 1 | 375 | 40.3 | 4.64 | 18 | 0.408 | 0.208 | 0.080 | 0.051 | 0.495 | 0.286 | 0.161 | 0.057 |
| 2130 | **Lamp2** | Unchanged | P17047 | 7 | 55 | 7 | 415 | 45.7 | 7.39 | 18 | 1.179 | 0.620 | 1.527 | 0.220 | 1.384 | 0.338 | 1.103 | 0.760 |
| 2131 | **Ado** | Unchanged | Q6PDY2 | 2 | 5 | 2 | 256 | 28.4 | 5.97 | 18 | 0.884 | 0.874 | 0.317 | 0.264 | 0.238 | 0.207 | 1.331 | 0.669 |
| 2132 | **Snx12** | Unchanged | O70493 | 3 | 22 | 1 | 165 | 19.1 | 7.34 | 18 | 1.351 | 0.497 | 0.704 | 0.565 | 3.061 | 0.150 | 0.230 | 0.107 |
| 2133 | **Map2k1** | Unchanged | P31938 | 4 | 20 | 1 | 393 | 43.4 | 6.7 | 18 | 1.718 | 0.284 | 3.613 | 0.223 | 3.327 | 0.126 | 1.086 | 0.903 |
| 2134 | **Rpia** | Unchanged | P47968 | 3 | 6 | 3 | 303 | 32.4 | 8.02 | 17 | 1.013 | 0.969 | 1.593 | 0.504 | 0.942 | 0.869 | 1.691 | 0.480 |
| 2135 | **Crygn** | Unchanged | Q8VHL5 | 2 | 5 | 2 | 183 | 21.4 | 6.71 | 17 | 1.061 | 0.947 | 1.237 | 0.837 | 0.771 | 0.759 | 1.604 | 0.614 |
| 2136 | **Ciapin1** | Unchanged | F8WIK0 | 2 | 8 | 2 | 309 | 33.4 | 5.2 | 17 | 1.258 | 0.748 | 1.405 | 0.527 | 1.615 | 0.323 | 0.870 | 0.676 |
| 2137 | **Mrpl40** | Unchanged | Q9Z2Q5 | 3 | 19 | 3 | 206 | 24.3 | 9.47 | 17 | 1.846 | 0.335 | 2.585 | 0.179 | 1.565 | 0.464 | 1.652 | 0.400 |
| 2138 | **Sf3b5** | Unchanged | Q923D4 | 1 | 4 | 1 | 86 | 10.1 | 6.35 | 17 | #DIV/0! | #DIV/0! | #DIV/0! | #DIV/0! | #DIV/0! | #DIV/0! | #DIV/0! | #DIV/0! |
| 2139 | **Arfip1** | Unchanged | G5E8V9 | 4 | 11 | 4 | 373 | 41.5 | 6.76 | 17 | 0.805 | 0.734 | 0.969 | 0.963 | 1.026 | 0.968 | 0.944 | 0.936 |
| 2140 | **Tgm1** | Unchanged | A0A0R4J293 | 10 | 34 | 10 | 815 | 89.7 | 6.58 | 17 | 0.844 | 0.614 | 1.042 | 0.888 | 0.548 | 0.058 | 1.900 | 0.091 |
| 2141 | **Fbxo22** | Unchanged | Q78JE5 | 5 | 13 | 5 | 402 | 44.2 | 7.61 | 17 | 5.480 | 0.079 | 10.520 | 0.076 | 3.304 | 0.173 | 3.184 | 0.178 |
| 2142 | **Mpdu1** | Unchanged | Q9R0Q9 | 3 | 39 | 3 | 247 | 26.5 | 8.75 | 17 | 0.893 | 0.771 | 0.847 | 0.607 | 0.863 | 0.684 | 0.982 | 0.947 |
| 2143 | **Ptp4a2** | Unchanged | O70274 | 1 | 2 | 1 | 167 | 19.1 | 8.37 | 17 | 1.672 | 0.069 | 2.248 | 0.272 | 0.894 | 0.772 | 2.516 | 0.253 |
| 2144 | **Cyp2c38** | Unchanged | P56655 | 8 | 109 | 1 | 490 | 56.2 | 8.29 | 17 | #DIV/0! | #DIV/0! | #DIV/0! | #DIV/0! | #DIV/0! | #DIV/0! | #DIV/0! | #DIV/0! |
| 2145 | **Csk** | Unchanged | P41241 | 6 | 16 | 6 | 450 | 50.7 | 7.06 | 17 | 1.519 | 0.478 | 0.912 | 0.821 | 0.743 | 0.392 | 1.226 | 0.549 |
| 2146 | **Casp3** | Unchanged | P70677 | 4 | 33 | 4 | 277 | 31.5 | 6.92 | 17 | 0.936 | 0.867 | 1.098 | 0.762 | 0.810 | 0.563 | 1.356 | 0.259 |
| 2147 | **Slc25a42** | Unchanged | Q8R0Y8 | 3 | 47 | 3 | 318 | 35.2 | 10.05 | 17 | 1.336 | 0.648 | 1.530 | 0.564 | 0.723 | 0.548 | 2.116 | 0.402 |
| 2148 | **Ucp1** | Unchanged | P12242 | 3 | 7 | 3 | 307 | 33.2 | 9.14 | 17 | 0.474 | 0.209 | 0.953 | 0.939 | 0.685 | 0.520 | 1.392 | 0.666 |
| 2149 | **Usp4** | UP | P35123 | 10 | 51 | 10 | 962 | 108.3 | 5.64 | 17 | 2.847 | 0.002 | 2.593 | 0.043 | 2.240 | 0.102 | 1.158 | 0.700 |
| 2150 | **Crat** | Unchanged | P47934 | 7 | 18 | 7 | 626 | 70.8 | 8.44 | 17 | 0.781 | 0.453 | 1.605 | 0.211 | 0.513 | 0.100 | 3.128 | 0.025 |
| 2151 | **Cryl1** | UP | Q99KP3 | 3 | 22 | 3 | 319 | 35.2 | 5.86 | 17 | 1.474 | 0.502 | 2.236 | 0.225 | 2.749 | 0.032 | 0.813 | 0.617 |
| 2152 | **Mbl2** | Down | P41317 | 3 | 28 | 3 | 244 | 25.9 | 5.06 | 17 | 0.319 | 0.015 | 0.209 | 0.007 | 0.212 | 0.007 | 0.989 | 0.972 |
| 2153 | **Bax** | Unchanged | Q07813 | 2 | 24 | 2 | 192 | 21.4 | 4.98 | 17 | 0.491 | 0.379 | 0.761 | 0.710 | 0.699 | 0.620 | 1.090 | 0.899 |
| 2154 | **Chchd3** | Down | Q9CRB9 | 3 | 25 | 3 | 227 | 26.3 | 8.37 | 17 | 1.095 | 0.829 | 1.161 | 0.766 | 0.317 | 0.006 | 3.659 | 0.140 |
| 2155 | **Fam3c** | Down | Q91VU0 | 3 | 7 | 3 | 227 | 24.7 | 8.29 | 17 | 0.969 | 0.928 | 0.790 | 0.563 | 0.293 | 0.030 | 2.700 | 0.056 |
| 2156 | **Ahcyl1** | Unchanged | Q80SW1 | 8 | 70 | 1 | 530 | 58.9 | 6.89 | 17 | 3.662 | 0.111 | 5.673 | 0.139 | 4.264 | 0.094 | 1.331 | 0.678 |
| 2157 | **Kpna6** | UP | O35345 | 5 | 35 | 4 | 536 | 59.9 | 4.94 | 17 | 3.220 | 0.084 | 4.884 | 0.115 | 7.413 | 0.007 | 0.659 | 0.396 |
| 2158 | **Derl2** | Unchanged | Q8BNI4 | 2 | 15 | 2 | 239 | 27.6 | 7.28 | 17 | 0.815 | 0.510 | 0.874 | 0.770 | 0.637 | 0.263 | 1.372 | 0.568 |
| 2159 | **Stx12** | Unchanged | Q9ER00 | 3 | 16 | 3 | 274 | 31.2 | 5.44 | 17 | 0.528 | 0.118 | 1.794 | 0.185 | 0.922 | 0.805 | 1.946 | 0.159 |
| 2160 | **Ptpmt1** | Down | Q66GT5 | 3 | 18 | 3 | 193 | 21.9 | 9.72 | 17 | 0.943 | 0.847 | 0.984 | 0.965 | 0.361 | 0.024 | 2.723 | 0.084 |
| 2161 | **Hpcal1** | Down | P62748 | 3 | 11 | 3 | 193 | 22.3 | 5.5 | 17 | 0.715 | 0.159 | 0.966 | 0.887 | 0.331 | 0.004 | 2.915 | 0.014 |
| 2162 | **Ighv1-39** | Unchanged | A0A075B5V5 | 1 | 1 | 1 | 117 | 12.9 | 8.41 | 17 | 1.154 | 0.842 | 1.051 | 0.956 | 0.597 | 0.466 | 1.760 | 0.591 |

| | | | | | | | | | | | | | | | | | | |
|---|---|---|---|---|---|---|---|---|---|---|---|---|---|---|---|---|---|---|
| 2163 | Gpd1l | UP | Q3ULJ0 | 3 | 39 | 2 | 351 | 38.2 | 6.77 | 17 | 18.734 | 0.000 | 16.906 | 0.132 | 11.787 | 0.054 | 1.434 | 0.643 |
| 2164 | Vps25 | Unchanged | Q9CQ80 | 3 | 15 | 3 | 176 | 20.7 | 6.34 | 17 | 2.073 | 0.145 | 1.591 | 0.377 | 7.963 | 0.070 | 0.200 | 0.095 |
| 2165 | Rnf114 | Unchanged | Q9ET26 | 3 | 5 | 3 | 229 | 25.7 | 7.03 | 17 | 0.793 | 0.737 | 1.410 | 0.645 | 0.276 | 0.131 | 5.100 | 0.164 |
| 2166 | Fetub | Unchanged | Q9QXC1 | 3 | 10 | 3 | 388 | 42.7 | 6.61 | 17 | 0.877 | 0.794 | 4.367 | 0.199 | 0.876 | 0.803 | 4.982 | 0.181 |
| 2167 | Mbnl1 | Unchanged | Q9JKP5 | 2 | 4 | 2 | 341 | 37 | 8.6 | 17 | 0.614 | 0.524 | 0.759 | 0.483 | 0.711 | 0.437 | 1.067 | 0.853 |
| 2168 | Plbd2 | Unchanged | Q3TCN2 | 7 | 35 | 7 | 594 | 66.2 | 6.13 | 17 | 0.873 | 0.671 | 2.283 | 0.108 | 0.752 | 0.318 | 3.035 | 0.063 |
| 2169 | Ube2l6 | Unchanged | Q9QZU9 | 1 | 2 | 1 | 153 | 17.8 | 8.09 | 17 | 3.138 | 0.362 | 6.959 | 0.157 | 4.788 | 0.333 | 1.454 | 0.693 |
| 2170 | Rbpms2 | Unchanged | Q8VC52 | 3 | 21 | 2 | 206 | 22.4 | 8.41 | 17 | 4.760 | 0.062 | 1.945 | 0.278 | 1.575 | 0.360 | 1.234 | 0.716 |
| 2171 | Sugct | Unchanged | Q7TNE1 | 4 | 27 | 4 | 436 | 47.7 | 8.68 | 17 | 2.030 | 0.113 | 1.755 | 0.353 | 1.871 | 0.322 | 0.938 | 0.919 |
| 2172 | Pfkm | Unchanged | P47857 | 8 | 32 | 7 | 780 | 85.2 | 8 | 17 | 2.179 | 0.316 | 0.268 | 0.135 | 0.122 | 0.081 | 2.190 | 0.090 |
| 2173 | Lrp1 | Unchanged | Q91ZX7 | 48 | 240 | 48 | 4545 | 504.4 | 5.36 | 17 | 1.021 | 0.939 | 0.929 | 0.771 | 0.769 | 0.148 | 1.208 | 0.467 |
| 2174 | Luc7l3 | Unchanged | Q5SUF2 | 5 | 19 | 5 | 432 | 51.4 | 9.77 | 17 | 0.892 | 0.848 | 1.402 | 0.479 | 0.886 | 0.813 | 1.582 | 0.269 |
| 2175 | Hist1h1e | Unchanged | P43274 | 4 | 42 | 1 | 219 | 22 | 11.11 | 17 | 0.580 | 0.246 | 1.005 | 0.994 | 0.908 | 0.886 | 1.106 | 0.907 |
| 2176 | Lyz1 | Unchanged | P17897 | 3 | 11 | 1 | 148 | 16.8 | 9.41 | 17 | #DIV/0! | #DIV/0! | #DIV/0! | #DIV/0! | #DIV/0! | #DIV/0! | #DIV/0! | #DIV/0! |
| 2177 | Pfdn2 | Unchanged | O70591 | 2 | 14 | 2 | 154 | 16.5 | 6.58 | 17 | 0.912 | 0.900 | 0.980 | 0.977 | 0.224 | 0.166 | 4.367 | 0.128 |
| 2178 | Cnbp | Unchanged | P53996 | 2 | 6 | 2 | 178 | 19.6 | 7.58 | 17 | 0.804 | 0.811 | 0.168 | 0.169 | 0.082 | 0.134 | 2.045 | 0.177 |
| 2179 | Eny2 | UP | Q9JIX0 | 1 | 3 | 1 | 101 | 11.5 | 9.33 | 17 | 1.283 | 0.686 | 9.872 | 0.013 | 7.264 | 0.005 | 1.359 | 0.436 |
| 2180 | Igkv4-86 | Unchanged | P01675 | 1 | 2 | 1 | 107 | 11.6 | 8.44 | 17 | #DIV/0! | #DIV/0! | #DIV/0! | #DIV/0! | #DIV/0! | #DIV/0! | #DIV/0! | #DIV/0! |
| 2181 | Gfpt1 | Unchanged | P47856 | 8 | 44 | 8 | 697 | 78.5 | 6.84 | 17 | 1.695 | 0.413 | 1.385 | 0.678 | 0.545 | 0.323 | 2.544 | 0.340 |
| 2182 | Elovl2 | Unchanged | Q9JLJ4 | 2 | 22 | 2 | 292 | 34.2 | 9.26 | 17 | 3.494 | 0.328 | 9.550 | 0.174 | 2.991 | 0.265 | 3.192 | 0.301 |
| 2183 | Rbm47 | Unchanged | Q91WT8 | 6 | 28 | 5 | 590 | 64 | 8.19 | 17 | 0.806 | 0.723 | 0.819 | 0.740 | 0.341 | 0.124 | 2.404 | 0.230 |
| 2184 | Ptp4a1 | Unchanged | Q63739 | 1 | 3 | 1 | 173 | 19.8 | 8.97 | 17 | 1.385 | 0.610 | 0.986 | 0.987 | 1.129 | 0.869 | 0.873 | 0.852 |
| 2185 | Hist1h1d | Unchanged | P43277 | 4 | 25 | 1 | 221 | 22.1 | 11.03 | 17 | 0.192 | 0.095 | 0.652 | 0.516 | 0.392 | 0.211 | 1.664 | 0.444 |
| 2186 | Ap3b1 | Unchanged | Q9Z1T1 | 12 | 40 | 12 | 1105 | 122.7 | 5.66 | 17 | 1.091 | 0.829 | 1.828 | 0.129 | 0.895 | 0.782 | 2.043 | 0.030 |
| 2187 | Gns | Down | Q8BFR4 | 6 | 16 | 6 | 544 | 61.1 | 8.24 | 17 | 0.534 | 0.038 | 0.744 | 0.303 | 1.251 | 0.417 | 0.595 | 0.157 |
| 2188 | Hs1bp3 | Unchanged | Q3TC93 | 3 | 7 | 3 | 395 | 43.7 | 4.98 | 17 | 1.031 | 0.927 | 1.402 | 0.230 | 1.322 | 0.525 | 1.060 | 0.888 |
| 2189 | Fermt3 | Unchanged | Q8K1B8 | 6 | 18 | 6 | 665 | 75.6 | 7.05 | 17 | 0.762 | 0.411 | 0.925 | 0.847 | 3.486 | 0.245 | 0.265 | 0.236 |
| 2190 | Commd9 | Unchanged | Q8K2Q0 | 2 | 8 | 2 | 198 | 21.8 | 5.82 | 17 | 1.805 | 0.329 | 0.803 | 0.687 | 1.519 | 0.382 | 0.528 | 0.260 |
| 2191 | Lsm8 | UP | Q6ZWM4 | 1 | 18 | 1 | 96 | 10.4 | 4.48 | 17 | 2.671 | 0.082 | 4.042 | 0.023 | 5.566 | 0.005 | 0.726 | 0.345 |
| 2192 | Rac2 | Unchanged | Q05144 | 3 | 9 | 1 | 192 | 21.4 | 7.61 | 17 | 1.898 | 0.485 | 5.461 | 0.065 | 1.392 | 0.538 | 3.923 | 0.091 |
| 2193 | Chchd10 | Unchanged | Q7TNL9 | 1 | 2 | 1 | 138 | 13.8 | 8.22 | 17 | 0.575 | 0.514 | 0.364 | 0.308 | 0.283 | 0.247 | 1.286 | 0.690 |
| 2194 | Sepsecs | Unchanged | Q6P6M7 | 5 | 27 | 5 | 504 | 55.3 | 8.06 | 17 | 1.236 | 0.556 | 2.292 | 0.090 | 0.608 | 0.116 | 3.769 | 0.039 |
| 2195 | L3hypdh | Unchanged | Q9CXA2 | 4 | 28 | 4 | 354 | 37.8 | 6.7 | 17 | 1.045 | 0.914 | 1.118 | 0.820 | 0.440 | 0.084 | 2.540 | 0.146 |
| 2196 | Cenpv | UP | Q9CXS4 | 2 | 3 | 2 | 252 | 27.5 | 9.79 | 17 | 3.077 | 0.085 | 2.229 | 0.373 | 13.877 | 0.040 | 0.161 | 0.064 |
| 2197 | Fads2 | Unchanged | Q9Z0R9 | 7 | 49 | 7 | 444 | 52.4 | 8.82 | 17 | 0.659 | 0.294 | 0.520 | 0.151 | 0.711 | 0.422 | 0.732 | 0.470 |
| 2198 | Creld1 | UP | Q91XD7 | 5 | 12 | 5 | 420 | 45.7 | 5.02 | 17 | 3.905 | 0.126 | 4.370 | 0.021 | 3.695 | 0.055 | 1.183 | 0.698 |
| 2199 | Ybx3 | Unchanged | Q9JKB3 | 5 | 32 | 1 | 361 | 38.8 | 9.69 | 17 | 0.931 | 0.912 | 1.255 | 0.839 | 1.171 | 0.775 | 1.072 | 0.945 |
| 2200 | Htra2 | Unchanged | Q9JIY5 | 6 | 21 | 6 | 458 | 49.3 | 9.6 | 17 | 1.381 | 0.629 | 1.650 | 0.552 | 0.270 | 0.149 | 6.113 | 0.182 |
| 2201 | Eif4e | Unchanged | P63073 | 3 | 19 | 3 | 217 | 25 | 6.15 | 17 | 1.700 | 0.247 | 1.449 | 0.365 | 1.470 | 0.399 | 0.986 | 0.976 |
| 2202 | Tsnax | Unchanged | Q9QZE7 | 4 | 26 | 4 | 290 | 32.9 | 6.55 | 17 | 0.943 | 0.901 | 1.672 | 0.244 | 1.499 | 0.259 | 1.115 | 0.790 |
| 2203 | Glb1 | Unchanged | P23780 | 9 | 40 | 9 | 647 | 73.1 | 7.47 | 17 | 2.607 | 0.276 | 2.138 | 0.387 | 1.828 | 0.258 | 1.169 | 0.830 |

| | | | | | | | | | | | | | | | | | | |
|---|---|---|---|---|---|---|---|---|---|---|---|---|---|---|---|---|---|---|
| 2204 | **Marcks** | Down | P26645 | 3 | 13 | 3 | 309 | 29.6 | 4.34 | 17 | 0.347 | 0.131 | 0.752 | 0.660 | 0.045 | 0.022 | 16.698 | 0.136 |
| 2205 | **Arl6ip5** | Unchanged | Q8R5J9 | 2 | 5 | 2 | 188 | 21.5 | 9.61 | 16 | 0.821 | 0.775 | 1.642 | 0.444 | 1.222 | 0.722 | 1.344 | 0.547 |
| 2206 | **Irf3** | Unchanged | P70671 | 4 | 26 | 4 | 419 | 46.8 | 5.43 | 16 | 1.205 | 0.642 | 2.796 | 0.228 | 0.975 | 0.948 | 2.867 | 0.223 |
| 2207 | **Rhot2** | Unchanged | Q8JZN7 | 6 | 25 | 5 | 620 | 69 | 6 | 16 | 0.618 | 0.224 | 0.947 | 0.872 | 0.539 | 0.090 | 1.756 | 0.179 |
| 2208 | **Picalm** | Unchanged | Q7M6Y3 | 9 | 82 | 4 | 660 | 71.5 | 7.91 | 16 | 0.760 | 0.598 | 1.214 | 0.710 | 0.572 | 0.291 | 2.124 | 0.192 |
| 2209 | **Aspscr1** | Unchanged | Q8VBT9 | 5 | 28 | 5 | 550 | 59.8 | 6.96 | 16 | 0.725 | 0.375 | 1.334 | 0.565 | 0.606 | 0.105 | 2.201 | 0.204 |
| 2210 | **Arpc5l** | Unchanged | Q9D898 | 2 | 11 | 2 | 153 | 17 | 6.8 | 16 | 0.743 | 0.309 | 1.178 | 0.576 | 0.846 | 0.603 | 1.392 | 0.278 |
| 2211 | **Isyna1** | Unchanged | Q9JHU9 | 6 | 20 | 6 | 557 | 60.9 | 6.42 | 16 | 0.804 | 0.790 | 0.037 | 0.112 | 0.599 | 0.520 | 0.062 | 0.057 |
| 2212 | **Ppp2r2a** | UP | Q6P1F6 | 5 | 35 | 5 | 447 | 51.7 | 6.2 | 16 | 2.472 | 0.073 | 3.379 | 0.034 | 1.137 | 0.669 | 2.973 | 0.051 |
| 2213 | **Ahcyl2** | Unchanged | Q68FL4 | 9 | 58 | 2 | 613 | 66.9 | 7.36 | 16 | 1.771 | 0.170 | 1.226 | 0.517 | 2.466 | 0.254 | 0.497 | 0.320 |
| 2214 | **Dtx3l** | Unchanged | Q3UIR3 | 7 | 20 | 7 | 748 | 83 | 8.09 | 16 | 1.900 | 0.216 | 1.533 | 0.343 | 1.975 | 0.105 | 0.776 | 0.550 |
| 2215 | **Prcp** | Unchanged | Q7TMR0 | 5 | 25 | 5 | 491 | 55 | 6.79 | 16 | 1.236 | 0.609 | 1.753 | 0.186 | 1.134 | 0.684 | 1.546 | 0.291 |
| 2216 | **Csde1** | Unchanged | Q91W50 | 9 | 51 | 9 | 798 | 88.7 | 6.37 | 16 | 0.713 | 0.425 | 0.906 | 0.785 | 0.440 | 0.115 | 2.061 | 0.041 |
| 2217 | **Stat1** | Unchanged | P42225 | 8 | 44 | 8 | 749 | 87.1 | 5.58 | 16 | 1.071 | 0.786 | 1.178 | 0.644 | 0.717 | 0.345 | 1.643 | 0.309 |
| 2218 | **Ears2** | Unchanged | Q9CXJ1 | 4 | 9 | 4 | 523 | 58.3 | 8.75 | 16 | 0.914 | 0.821 | 1.638 | 0.409 | 0.750 | 0.412 | 2.183 | 0.237 |
| 2219 | **Cox7a1** | Unchanged | P56392 | 1 | 2 | 1 | 80 | 9 | 9.79 | 16 | 0.363 | 0.477 | 0.155 | 0.349 | 0.278 | 0.425 | 0.559 | 0.405 |
| 2220 | **Atg3** | Unchanged | Q9CPX6 | 4 | 16 | 4 | 314 | 35.8 | 4.72 | 16 | 0.295 | 0.158 | 0.297 | 0.148 | 0.311 | 0.170 | 0.954 | 0.942 |
| 2221 | **Ighv5-6** | Down | A0A075B5Q0 | 1 | 1 | 1 | 117 | 12.9 | 8.6 | 16 | 2.314 | 0.214 | 0.074 | 0.043 | 3.003 | 0.228 | 0.025 | 0.084 |
| 2222 | **Sirt3** | Unchanged | Q8R104 | 5 | 29 | 5 | 334 | 36.6 | 7.44 | 16 | 1.223 | 0.257 | 1.450 | 0.200 | 0.898 | 0.546 | 1.614 | 0.111 |
| 2223 | **Dscr3** | Unchanged | O35075 | 3 | 12 | 3 | 297 | 32.9 | 7.68 | 16 | 0.898 | 0.823 | 1.465 | 0.466 | 0.332 | 0.120 | 4.417 | 0.048 |
| 2224 | **Ppp2r1b** | Unchanged | Q7TNP2 | 9 | 111 | 3 | 601 | 65.9 | 5.1 | 16 | 0.226 | 0.059 | 0.901 | 0.889 | 0.823 | 0.642 | 1.096 | 0.907 |
| 2225 | **Ifi30** | Unchanged | Q9ESY9 | 5 | 21 | 5 | 248 | 27.8 | 5.01 | 16 | 1.049 | 0.920 | 1.438 | 0.378 | 0.708 | 0.443 | 2.030 | 0.051 |
| 2226 | **Mapk14** | Unchanged | P47811 | 4 | 27 | 4 | 360 | 41.3 | 5.88 | 16 | 1.503 | 0.335 | 3.704 | 0.111 | 1.534 | 0.151 | 2.415 | 0.191 |
| 2227 | **Clns1a** | Unchanged | Q61189 | 2 | 5 | 2 | 236 | 26 | 4.12 | 16 | 1.178 | 0.204 | 2.043 | 0.411 | 1.839 | 0.215 | 1.111 | 0.884 |
| 2228 | **Mrpl3** | Unchanged | Q99N95 | 2 | 3 | 2 | 348 | 39.1 | 9.55 | 16 | 1.541 | 0.204 | 1.070 | 0.827 | 1.095 | 0.839 | 0.977 | 0.960 |
| 2229 | **Col14a1** | Unchanged | Q80X19 | 16 | 47 | 16 | 1797 | 192.9 | 5.1 | 16 | 1.156 | 0.611 | 1.359 | 0.367 | 1.350 | 0.209 | 1.007 | 0.982 |
| 2230 | **Tollip** | Unchanged | Q9QZ06 | 4 | 28 | 4 | 274 | 30.3 | 5.17 | 16 | 1.283 | 0.699 | 0.385 | 0.276 | 0.124 | 0.129 | 3.096 | 0.026 |
| 2231 | **Lsm4** | Unchanged | Q9QXA5 | 3 | 10 | 3 | 137 | 15.1 | 10.05 | 11 | 0.872 | 0.741 | 1.544 | 0.514 | 0.916 | 0.771 | 1.685 | 0.451 |
| 2232 | **Abracl** | Unchanged | Q4KML4 | 1 | 2 | 1 | 81 | 9 | 5.8 | 16 | 2.586 | 0.380 | 5.565 | 0.060 | 0.610 | 0.266 | 9.122 | 0.044 |
| 2233 | **4932414N04Rik** | Unchanged | A2AQL4 | 1 | 1 | 1 | 162 | 18.1 | 5.4 | 16 | 1.652 | 0.598 | 1.973 | 0.540 | 1.470 | 0.653 | 1.342 | 0.747 |
| 2234 | **Picalmiso** | Unchanged | A0A140LHQ8 | 6 | 30 | 1 | 531 | 57 | 7.59 | 16 | 0.772 | 0.644 | 0.824 | 0.779 | 0.675 | 0.482 | 1.221 | 0.785 |
| 2235 | **Rbmxl1** | Unchanged | Q91VM5 | 5 | 22 | 1 | 388 | 42.1 | 9.99 | 16 | 1.003 | 0.997 | 0.572 | 0.426 | 6.596 | 0.191 | 0.087 | 0.161 |
| 2236 | **Adssl1** | Unchanged | P28650 | 5 | 39 | 5 | 457 | 50.2 | 8.43 | 16 | 1.183 | 0.551 | 0.770 | 0.507 | 1.133 | 0.744 | 0.680 | 0.394 |
| 2237 | **Asgr2** | Unchanged | P24721 | 3 | 14 | 3 | 301 | 34.9 | 5.21 | 16 | 1.268 | 0.469 | 1.053 | 0.912 | 0.924 | 0.884 | 1.139 | 0.837 |
| 2238 | **Casp6** | Down | O08738 | 4 | 12 | 4 | 276 | 31.6 | 6.89 | 16 | 0.318 | 0.030 | 1.134 | 0.659 | 0.806 | 0.473 | 1.407 | 0.144 |
| 2239 | **Emc8** | Unchanged | O70378 | 2 | 10 | 2 | 207 | 23.3 | 6.15 | 16 | 1.956 | 0.271 | 4.225 | 0.070 | 1.705 | 0.307 | 2.479 | 0.151 |
| 2240 | **Srr** | Unchanged | Q9QZX7 | 3 | 43 | 3 | 339 | 36.3 | 6.02 | 16 | 1.560 | 0.232 | 1.687 | 0.106 | 1.360 | 0.274 | 1.240 | 0.383 |
| 2241 | **Ecpas** | Unchanged | Q6PDI5 | 16 | 97 | 16 | 1840 | 203.6 | 7.06 | 16 | 1.031 | 0.896 | 1.485 | 0.243 | 1.141 | 0.502 | 1.301 | 0.388 |
| 2242 | **Capg** | Unchanged | P24452 | 3 | 4 | 3 | 352 | 39.2 | 7.2 | 16 | 0.824 | 0.397 | 1.010 | 0.958 | 4.619 | 0.395 | 0.219 | 0.397 |
| 2243 | **Metap2** | Unchanged | O08663 | 5 | 14 | 5 | 478 | 52.9 | 5.82 | 16 | 1.076 | 0.848 | 0.495 | 0.101 | 1.332 | 0.451 | 0.371 | 0.050 |
| 2244 | **Aga** | Unchanged | Q64191 | 3 | 9 | 3 | 346 | 37 | 6.44 | 16 | 2.369 | 0.100 | 3.737 | 0.222 | 2.543 | 0.089 | 1.469 | 0.600 |

| | | | | | | | | | | | | | | | | | | |
|---|---|---|---|---|---|---|---|---|---|---|---|---|---|---|---|---|---|---|
| 2245 | **Fmo2** | UP | Q8K2I3 | 4 | 10 | 4 | 535 | 60.9 | 8.47 | 16 | 1.186 | 0.398 | 1.674 | 0.031 | 0.889 | 0.642 | 1.882 | 0.041 |
| 2246 | **Rbmx** | Unchanged | Q9WV02 | 5 | 20 | 1 | 391 | 42.3 | 10.05 | 16 | 0.795 | 0.678 | 1.314 | 0.718 | 1.160 | 0.771 | 1.133 | 0.848 |
| 2247 | **Lrpap1** | Unchanged | P55302 | 4 | 32 | 4 | 360 | 42.2 | 7.87 | 16 | 1.246 | 0.751 | 1.130 | 0.867 | 1.046 | 0.931 | 1.081 | 0.910 |
| 2248 | **Lpp** | Unchanged | Q8BFW7 | 6 | 30 | 6 | 613 | 65.8 | 7.37 | 16 | 0.794 | 0.692 | 0.611 | 0.389 | 1.087 | 0.908 | 0.562 | 0.507 |
| 2249 | **Ap1s1** | Unchanged | P61967 | 2 | 40 | 2 | 158 | 18.7 | 5.73 | 16 | 0.716 | 0.706 | 0.437 | 0.404 | 0.094 | 0.170 | 4.670 | 0.159 |
| 2250 | **Csnk2b** | Unchanged | P67871 | 2 | 6 | 2 | 215 | 24.9 | 5.55 | 16 | 1.425 | 0.633 | 1.781 | 0.496 | 2.260 | 0.249 | 0.788 | 0.724 |
| 2251 | **Apom** | Unchanged | Q9Z1R3 | 2 | 6 | 2 | 190 | 21.3 | 6.52 | 16 | 0.884 | 0.840 | 1.560 | 0.545 | 2.047 | 0.131 | 0.762 | 0.610 |
| 2252 | **Nedd4** | Unchanged | P46935 | 9 | 64 | 6 | 887 | 102.6 | 5.26 | 16 | 0.997 | 0.994 | 1.311 | 0.368 | 0.727 | 0.356 | 1.802 | 0.042 |
| 2253 | **Cfb** | Unchanged | P04186 | 8 | 69 | 8 | 761 | 85 | 7.37 | 16 | 0.964 | 0.939 | 1.690 | 0.122 | 1.583 | 0.203 | 1.067 | 0.599 |
| 2254 | **Atp6v1c1** | Unchanged | Q9Z1G3 | 4 | 7 | 4 | 382 | 43.9 | 7.46 | 16 | 1.146 | 0.805 | 1.100 | 0.858 | 0.902 | 0.830 | 1.220 | 0.705 |
| 2255 | **Chchd2** | Down | Q9D1L0 | 1 | 15 | 1 | 153 | 15.7 | 9.66 | 16 | 0.503 | 0.076 | 0.400 | 0.053 | 0.200 | 0.012 | 2.006 | 0.321 |
| 2256 | **Bsg** | Unchanged | P18572 | 5 | 77 | 5 | 389 | 42.4 | 5.85 | 16 | 1.019 | 0.960 | 1.326 | 0.536 | 1.100 | 0.693 | 1.205 | 0.675 |
| 2257 | **Slc2a2** | Unchanged | P14246 | 7 | 38 | 7 | 523 | 57.1 | 6.73 | 16 | 1.129 | 0.887 | 0.949 | 0.948 | 0.936 | 0.939 | 1.014 | 0.985 |
| 2258 | **Ppp1r14a** | Unchanged | Q91VC7 | 1 | 2 | 1 | 147 | 16.6 | 7.34 | 16 | 2.340 | 0.368 | 4.513 | 0.153 | 1.432 | 0.329 | 3.152 | 0.199 |
| 2259 | **Bub3** | Unchanged | Q9WVA3 | 4 | 17 | 4 | 326 | 36.9 | 6.84 | 16 | 1.333 | 0.684 | 3.740 | 0.159 | 1.646 | 0.302 | 2.273 | 0.286 |
| 2260 | **Abca6** | Unchanged | Q8K441 | 15 | 78 | 15 | 1624 | 183.2 | 6.93 | 16 | 1.107 | 0.743 | 1.474 | 0.125 | 0.988 | 0.966 | 1.492 | 0.055 |
| 2261 | **Pacsin2** | Unchanged | Q9WVE8 | 6 | 40 | 6 | 486 | 55.8 | 5.2 | 16 | 0.862 | 0.649 | 1.539 | 0.326 | 1.388 | 0.394 | 1.108 | 0.791 |
| 2262 | **Tma7** | Unchanged | Q8K003 | 1 | 6 | 1 | 64 | 7.1 | 9.99 | 16 | 0.483 | 0.118 | 0.757 | 0.501 | 0.796 | 0.688 | 0.951 | 0.943 |
| 2263 | **Hnrnpdl** | Unchanged | Q9Z130 | 4 | 22 | 2 | 301 | 33.5 | 7.31 | 16 | 1.165 | 0.807 | 1.537 | 0.468 | 1.942 | 0.225 | 0.791 | 0.615 |
| 2264 | **Tmem208** | Unchanged | Q9CR96 | 1 | 1 | 1 | 173 | 19.6 | 9.23 | 16 | #DIV/0! | #DIV/0! | #DIV/0! | #DIV/0! | #DIV/0! | #DIV/0! | #DIV/0! | #DIV/0! |
| 2265 | **Xpnpep1** | Unchanged | Q6P1B1 | 8 | 84 | 8 | 623 | 69.5 | 5.54 | 16 | 1.221 | 0.037 | 1.486 | 0.174 | 1.457 | 0.012 | 1.020 | 0.933 |
| 2266 | **Pdhx** | Unchanged | Q8BKZ9 | 6 | 49 | 6 | 501 | 54 | 7.75 | 16 | 0.821 | 0.169 | 0.732 | 0.052 | 0.946 | 0.845 | 0.774 | 0.432 |
| 2267 | **Map2k3** | Unchanged | O09110 | 3 | 15 | 3 | 347 | 39.3 | 7.43 | 16 | 2.811 | 0.244 | 4.338 | 0.125 | 0.983 | 0.962 | 4.415 | 0.121 |
| 2268 | **Timm10** | Unchanged | P62073 | 2 | 22 | 2 | 90 | 10.3 | 6.29 | 16 | 0.288 | 0.325 | 1.644 | 0.619 | 0.296 | 0.333 | 5.562 | 0.238 |
| 2269 | **Mrpl22** | Unchanged | Q8BU88 | 3 | 11 | 3 | 206 | 23.8 | 9.86 | 16 | 1.475 | 0.262 | 1.981 | 0.242 | 2.484 | 0.169 | 0.798 | 0.694 |
| 2270 | **Scd1** | Unchanged | P13516 | 4 | 59 | 4 | 355 | 41 | 9.07 | 15 | 1.026 | 0.967 | 0.912 | 0.893 | 1.184 | 0.767 | 0.770 | 0.730 |
| 2271 | **Sept10** | Unchanged | Q8C650 | 4 | 46 | 3 | 452 | 52.4 | 6.6 | 15 | 1.432 | 0.166 | 1.922 | 0.333 | 0.606 | 0.240 | 3.174 | 0.186 |
| 2272 | **Tspan9** | Unchanged | Q8BJU2 | 2 | 4 | 2 | 239 | 26.7 | 7.42 | 15 | 1.113 | 0.855 | 25.971 | 0.179 | 0.447 | 0.339 | 58.166 | 0.170 |
| 2273 | **Abhd6** | Unchanged | Q8R2Y0 | 3 | 19 | 3 | 336 | 38.2 | 8.47 | 15 | 1.838 | 0.311 | 2.380 | 0.167 | 1.546 | 0.467 | 1.540 | 0.417 |
| 2274 | **Sec23ip** | Unchanged | Q6NZC7 | 8 | 20 | 7 | 998 | 110.7 | 5.94 | 15 | 1.255 | 0.540 | 1.817 | 0.205 | 1.728 | 0.149 | 1.051 | 0.894 |
| 2275 | **Usp10** | Unchanged | P52479 | 8 | 57 | 8 | 792 | 87 | 5.17 | 15 | 0.886 | 0.598 | 1.113 | 0.645 | 0.880 | 0.521 | 1.265 | 0.371 |
| 2276 | **Maoa** | UP | Q64133 | 6 | 30 | 5 | 526 | 59.6 | 7.81 | 15 | 1.512 | 0.181 | 2.000 | 0.076 | 1.942 | 0.000 | 1.030 | 0.911 |
| 2277 | **Acsbg1** | Down | D3YZ56 | 1 | 1 | 1 | 65 | 7.2 | 10.18 | 15 | 0.810 | 0.698 | 1.122 | 0.900 | 0.050 | 0.048 | 22.601 | 0.243 |
| 2278 | **4931414P19Rik** | Unchanged | A0A2I3BS37 | 1 | 1 | 1 | 39 | 4.6 | 11.59 | 15 | 0.754 | 0.661 | 0.023 | 0.034 | 0.265 | 0.095 | 0.088 | 0.005 |
| 2279 | **Nln** | UP | Q91YP2 | 8 | 33 | 8 | 704 | 80.4 | 6.44 | 15 | 1.798 | 0.107 | 1.602 | 0.049 | 1.646 | 0.116 | 0.974 | 0.924 |
| 2280 | **Ciao1** | UP | Q99KN2 | 3 | 11 | 3 | 339 | 37.6 | 4.88 | 15 | 4.400 | 0.032 | 4.704 | 0.228 | 4.866 | 0.032 | 0.967 | 0.960 |
| 2281 | **Mia3** | Unchanged | Q8BI84 | 21 | 167 | 21 | 1930 | 213.5 | 4.75 | 15 | 0.933 | 0.683 | 0.917 | 0.777 | 0.776 | 0.531 | 1.182 | 0.739 |
| 2282 | **Esyt1** | Unchanged | Q3U7R1 | 11 | 51 | 11 | 1092 | 121.5 | 5.95 | 15 | 1.056 | 0.857 | 2.066 | 0.106 | 1.183 | 0.569 | 1.747 | 0.191 |
| 2283 | **Tmem167a** | Unchanged | Q9CR64 | 1 | 1 | 1 | 72 | 8.1 | 8.95 | 15 | 0.708 | 0.343 | 0.734 | 0.443 | 0.826 | 0.524 | 0.889 | 0.770 |
| 2284 | **Ppp1r11** | Unchanged | Q8K1L5 | 1 | 4 | 1 | 131 | 14.5 | 6.09 | 15 | #DIV/0! | #DIV/0! | #DIV/0! | #DIV/0! | #DIV/0! | #DIV/0! | #DIV/0! | #DIV/0! |
| 2285 | **Rab43** | Unchanged | Q8CG50 | 3 | 25 | 2 | 210 | 23.2 | 5.86 | 15 | 1.092 | 0.789 | 0.567 | 0.194 | 0.486 | 0.083 | 1.166 | 0.772 |

| | | | | | | | | | | | | | | | | | | |
|---|---|---|---|---|---|---|---|---|---|---|---|---|---|---|---|---|---|---|
| 2286 | **Prune1** | Down | Q8BIW1 | 4 | 5 | 4 | 454 | 50.2 | 5.11 | 15 | 0.599 | 0.232 | 0.570 | 0.201 | 0.339 | 0.035 | 1.683 | 0.342 |
| 2287 | **Syndig1l** | Unchanged | Q3USQ7 | 1 | 1 | 1 | 237 | 25.5 | 4.77 | 15 | 1.422 | 0.618 | 0.204 | 0.187 | 1.247 | 0.761 | 0.164 | 0.131 |
| 2288 | **Clint1** | Unchanged | Q5SUH6 | 7 | 16 | 7 | 641 | 69.7 | 6.65 | 15 | 0.545 | 0.445 | 3.361 | 0.326 | 0.866 | 0.841 | 3.884 | 0.296 |
| 2289 | **Oxnad1** | Unchanged | Q8VE38 | 3 | 13 | 3 | 311 | 34.7 | 8.13 | 15 | 1.384 | 0.379 | 1.469 | 0.364 | 1.252 | 0.632 | 1.173 | 0.735 |
| 2290 | **Mrc1** | Unchanged | Q61830 | 16 | 50 | 16 | 1456 | 164.9 | 6.83 | 15 | 0.970 | 0.896 | 0.943 | 0.726 | 1.036 | 0.892 | 0.910 | 0.709 |
| 2291 | **Cfi** | Unchanged | Q61129 | 5 | 12 | 5 | 603 | 67.2 | 7.46 | 15 | 0.818 | 0.570 | 0.543 | 0.174 | 0.598 | 0.219 | 0.908 | 0.832 |
| 2292 | **Tsg101** | Unchanged | Q61187 | 4 | 14 | 4 | 391 | 44.1 | 6.71 | 15 | 0.760 | 0.372 | 1.233 | 0.693 | 0.555 | 0.094 | 2.222 | 0.235 |
| 2293 | **Stk24** | Unchanged | Q99KH8 | 4 | 16 | 4 | 431 | 47.9 | 5.43 | 15 | 1.251 | 0.582 | 1.908 | 0.177 | 0.804 | 0.568 | 2.373 | 0.090 |
| 2294 | **Colgalt1** | Unchanged | Q8K297 | 8 | 11 | 8 | 617 | 71 | 7.28 | 15 | 0.456 | 0.236 | 0.776 | 0.681 | 1.021 | 0.966 | 0.760 | 0.583 |
| 2295 | **Cmas** | Unchanged | Q99KK2 | 6 | 27 | 6 | 432 | 48 | 8.1 | 15 | 0.702 | 0.470 | 0.983 | 0.978 | 1.343 | 0.489 | 0.731 | 0.558 |
| 2296 | **Rraga** | Unchanged | Q80X95 | 3 | 8 | 3 | 313 | 36.5 | 7.72 | 15 | 1.641 | 0.420 | 3.679 | 0.209 | 1.674 | 0.297 | 2.198 | 0.354 |
| 2297 | **Tor1a** | Unchanged | Q9ER39 | 4 | 11 | 4 | 333 | 37.8 | 7.17 | 15 | 0.414 | 0.090 | 0.787 | 0.561 | 0.449 | 0.117 | 1.750 | 0.170 |
| 2298 | **Casp8** | Unchanged | O89110 | 5 | 16 | 5 | 480 | 55.3 | 5.25 | 15 | 0.399 | 0.125 | 0.239 | 0.060 | 0.222 | 0.051 | 1.075 | 0.861 |
| 2299 | **Hspa1l** | Unchanged | P16627 | 7 | 131 | 1 | 641 | 70.6 | 6.24 | 15 | 0.373 | 0.202 | 0.263 | 0.140 | 0.572 | 0.377 | 0.460 | 0.220 |
| 2300 | **Pfas** | Unchanged | Q5SUR0 | 9 | 41 | 9 | 1337 | 144.5 | 5.67 | 15 | 0.849 | 0.239 | 1.228 | 0.397 | 1.367 | 0.055 | 0.898 | 0.630 |
| 2301 | **Timm22** | Unchanged | Q9CQ85 | 1 | 1 | 1 | 194 | 20.1 | 8.22 | 15 | 1.464 | 0.428 | 1.850 | 0.202 | 1.829 | 0.260 | 1.012 | 0.975 |
| 2302 | **Mrps21** | Unchanged | P58059 | 1 | 8 | 1 | 87 | 10.6 | 10.32 | 15 | 1.122 | 0.867 | 1.295 | 0.719 | 0.089 | 0.135 | 14.607 | 0.069 |
| 2303 | **Commd6** | Unchanged | Q3V4B5 | 1 | 2 | 1 | 87 | 9.8 | 5.38 | 15 | 1.030 | 0.953 | 4.711 | 0.358 | 0.876 | 0.818 | 5.376 | 0.345 |
| 2304 | **Rangap1** | Unchanged | P46061 | 6 | 23 | 6 | 589 | 63.5 | 4.68 | 15 | 1.348 | 0.610 | 1.118 | 0.886 | 1.303 | 0.664 | 0.858 | 0.842 |
| 2305 | **Pex11g** | Unchanged | Q6P6M5 | 2 | 13 | 2 | 241 | 27.1 | 9.61 | 15 | 0.519 | 0.444 | 0.951 | 0.958 | 0.490 | 0.352 | 1.941 | 0.598 |
| 2306 | **FAM120A** | Unchanged | Q6A0A9 | 11 | 46 | 11 | 1112 | 121.6 | 8.92 | 14 | 1.283 | 0.417 | 0.856 | 0.660 | 1.213 | 0.610 | 0.706 | 0.474 |
| 2307 | **Nop58** | Unchanged | Q6DFW4 | 5 | 26 | 5 | 536 | 60.3 | 8.34 | 15 | 1.624 | 0.643 | 0.666 | 0.704 | 0.606 | 0.652 | 1.098 | 0.865 |
| 2308 | **Nfu1** | Unchanged | Q9QZ23 | 3 | 26 | 3 | 255 | 28.5 | 5.03 | 15 | 3.193 | 0.244 | 4.815 | 0.156 | 4.316 | 0.119 | 1.115 | 0.874 |
| 2309 | **Scamp3** | Unchanged | O35609 | 3 | 20 | 3 | 349 | 38.4 | 7.64 | 15 | 0.633 | 0.463 | 1.412 | 0.628 | 0.555 | 0.368 | 2.543 | 0.293 |
| 2310 | **Ptgr1** | Unchanged | Q91YR9 | 3 | 12 | 3 | 329 | 35.5 | 7.96 | 15 | 0.481 | 0.214 | 2.178 | 0.133 | 2.583 | 0.264 | 0.843 | 0.779 |
| 2311 | **Gfm2** | Unchanged | Q8R2Q4 | 6 | 39 | 6 | 779 | 86.1 | 6.43 | 15 | 1.041 | 0.938 | 1.402 | 0.584 | 1.160 | 0.712 | 1.209 | 0.737 |
| 2312 | **Ttll12** | Unchanged | Q3UDE2 | 5 | 6 | 5 | 639 | 74 | 5.63 | 15 | 0.885 | 0.690 | 1.229 | 0.547 | 0.774 | 0.453 | 1.589 | 0.187 |
| 2313 | **Cobll1** | Unchanged | B1AZ15 | 10 | 20 | 10 | 1240 | 133.2 | 7.36 | 15 | 1.141 | 0.766 | 0.847 | 0.737 | 0.814 | 0.686 | 1.041 | 0.942 |
| 2314 | **Ergic1** | Down | Q9DC16 | 3 | 11 | 3 | 290 | 32.5 | 7.06 | 15 | 1.098 | 0.723 | 0.391 | 0.019 | 0.046 | 0.000 | 8.504 | 0.083 |
| 2315 | **Rer1** | Unchanged | Q9CQU3 | 2 | 38 | 2 | 196 | 23 | 9.51 | 15 | 1.322 | 0.442 | 1.052 | 0.923 | 0.982 | 0.954 | 1.072 | 0.890 |
| 2316 | **Mrps22** | UP | Q9CXW2 | 4 | 10 | 4 | 359 | 41.2 | 8.56 | 15 | 5.580 | 0.194 | 22.612 | 0.160 | 3.868 | 0.031 | 5.846 | 0.218 |
| 2317 | **Gstp3** | Unchanged | F6RWR5 | 2 | 21 | 1 | 210 | 24.1 | 7.88 | 15 | 1.043 | 0.911 | 3.570 | 0.155 | 0.910 | 0.833 | 3.925 | 0.146 |
| 2318 | **Snap23** | Unchanged | O09044 | 2 | 18 | 2 | 210 | 23.2 | 4.98 | 15 | 1.681 | 0.458 | 3.232 | 0.196 | 0.966 | 0.894 | 3.346 | 0.188 |
| 2319 | **Mov10** | Unchanged | P23249 | 9 | 27 | 9 | 1004 | 113.5 | 9.01 | 15 | 0.692 | 0.558 | 0.317 | 0.165 | 0.715 | 0.605 | 0.444 | 0.217 |
| 2320 | **Cartpt** | Unchanged | P56388 | 1 | 4 | 1 | 129 | 14.3 | 8.87 | 15 | 1.763 | 0.469 | 4.045 | 0.167 | 2.133 | 0.356 | 1.896 | 0.413 |
| 2321 | **Eps15** | Unchanged | P42567 | 10 | 36 | 10 | 897 | 98.4 | 4.6 | 15 | 1.406 | 0.358 | 0.909 | 0.688 | 1.470 | 0.416 | 0.619 | 0.325 |
| 2322 | **Ctnnd1** | Unchanged | P30999 | 8 | 45 | 8 | 938 | 104.9 | 6.87 | 15 | 0.225 | 0.080 | 0.203 | 0.073 | 0.455 | 0.217 | 0.447 | 0.111 |
| 2323 | **Tbl2** | Unchanged | Q9R099 | 5 | 35 | 5 | 442 | 49.6 | 9.04 | 15 | 1.072 | 0.917 | 1.221 | 0.730 | 0.544 | 0.426 | 2.245 | 0.070 |
| 2324 | **Abcc6** | Unchanged | Q9R1S7 | 13 | 82 | 13 | 1498 | 164.8 | 8.21 | 15 | 1.001 | 0.997 | 1.214 | 0.397 | 0.673 | 0.124 | 1.803 | 0.031 |
| 2325 | **Rdh14** | Unchanged | Q9ERI6 | 3 | 15 | 3 | 334 | 36.3 | 8.18 | 15 | 1.930 | 0.477 | 5.435 | 0.110 | 1.048 | 0.918 | 5.188 | 0.112 |
| 2326 | **Nhlrc2** | UP | Q8BZW8 | 5 | 14 | 5 | 725 | 78.4 | 5.54 | 15 | 0.976 | 0.951 | 2.339 | 0.296 | 2.727 | 0.007 | 0.857 | 0.759 |

| | | | | | | | | | | | | | | | | | | |
|---|---|---|---|---|---|---|---|---|---|---|---|---|---|---|---|---|---|---|
| 2327 | **Ceacam1** | Unchanged | P31809 | 4 | 27 | 4 | 521 | 57 | 5.58 | 15 | 0.757 | 0.650 | 0.419 | 0.234 | 0.424 | 0.233 | 0.988 | 0.978 |
| 2328 | **Pex16** | Unchanged | Q91XC9 | 3 | 10 | 3 | 336 | 38.7 | 9.72 | 15 | 1.274 | 0.685 | 2.256 | 0.308 | 1.489 | 0.362 | 1.515 | 0.496 |
| 2329 | **Sugt1** | Unchanged | Q9CX34 | 3 | 22 | 3 | 336 | 38.1 | 5.45 | 15 | 0.550 | 0.458 | 0.284 | 0.217 | 0.163 | 0.151 | 1.747 | 0.417 |
| 2330 | **Zc3h15** | Unchanged | Q3TIV5 | 4 | 27 | 4 | 426 | 48.3 | 5.3 | 15 | 1.048 | 0.919 | 1.092 | 0.810 | 0.770 | 0.404 | 1.418 | 0.330 |
| 2331 | **Cpne1** | Unchanged | Q8C166 | 5 | 32 | 5 | 536 | 58.8 | 5.66 | 15 | 1.295 | 0.222 | 1.974 | 0.136 | 1.014 | 0.950 | 1.946 | 0.156 |
| 2332 | **Eif2b2** | UP | Q99LD9 | 2 | 28 | 2 | 351 | 38.9 | 6.24 | 15 | 4.306 | 0.100 | 7.205 | 0.000 | 4.143 | 0.004 | 1.739 | 0.024 |
| 2333 | **Commd7** | Unchanged | Q8BG94 | 1 | 5 | 1 | 200 | 22.6 | 5.94 | 15 | 0.656 | 0.630 | 1.187 | 0.810 | 1.964 | 0.447 | 0.604 | 0.531 |
| 2334 | **Mipep** | Unchanged | A6H611 | 5 | 10 | 5 | 711 | 80.8 | 7.15 | 14 | 0.586 | 0.240 | 0.936 | 0.859 | 0.568 | 0.148 | 1.649 | 0.259 |
| 2335 | **Atp8b2** | Unchanged | F6WT01 | 1 | 5 | 1 | 145 | 16 | 6.15 | 14 | 1.054 | 0.935 | 1.066 | 0.914 | 0.671 | 0.514 | 1.589 | 0.426 |
| 2336 | **Ddx17** | Unchanged | Q501J6 | 8 | 75 | 3 | 650 | 72.4 | 8.59 | 14 | 0.866 | 0.700 | 2.104 | 0.226 | 1.762 | 0.081 | 1.194 | 0.696 |
| 2337 | **Psmd4iso** | Unchanged | F7AI87 | 2 | 8 | 2 | 194 | 21.3 | 4.45 | 14 | 0.825 | 0.716 | 0.572 | 0.278 | 0.516 | 0.252 | 1.110 | 0.891 |
| 2338 | **Cdk6** | Unchanged | Q64261 | 3 | 7 | 3 | 326 | 37 | 6.62 | 14 | 12.197 | 0.103 | 3.762 | 0.080 | 1.610 | 0.228 | 2.337 | 0.172 |
| 2339 | **Serpinf1** | Unchanged | P97298 | 3 | 9 | 3 | 417 | 46.2 | 6.98 | 14 | 1.539 | 0.551 | 6.895 | 0.164 | 0.776 | 0.592 | 8.886 | 0.149 |
| 2340 | **Cast** | Unchanged | P51125 | 5 | 40 | 5 | 788 | 84.9 | 5.52 | 14 | 0.421 | 0.094 | 0.783 | 0.516 | 0.495 | 0.154 | 1.581 | 0.144 |
| 2341 | **Afg3l2** | UP | Q8JZQ2 | 8 | 28 | 7 | 802 | 89.5 | 8.6 | 14 | 0.891 | 0.782 | 1.478 | 0.452 | 1.858 | 0.023 | 0.795 | 0.506 |
| 2342 | **Ppp6r3** | Unchanged | Q922D4 | 8 | 38 | 7 | 844 | 94.6 | 4.58 | 14 | 1.200 | 0.747 | 1.710 | 0.414 | 1.414 | 0.555 | 1.210 | 0.752 |
| 2343 | **Agt** | Unchanged | Q3UTR7 | 3 | 8 | 3 | 482 | 52.6 | 5.54 | 14 | 1.110 | 0.844 | 0.879 | 0.847 | 0.794 | 0.687 | 1.108 | 0.835 |
| 2344 | **Nrdc** | Unchanged | Q8BHG1 | 11 | 42 | 11 | 1161 | 132.8 | 4.87 | 14 | 0.888 | 0.718 | 0.985 | 0.957 | 0.567 | 0.110 | 1.738 | 0.027 |
| 2345 | **Pex11b** | Unchanged | Q9Z210 | 3 | 14 | 3 | 259 | 28.7 | 9.95 | 14 | 0.273 | 0.110 | 0.431 | 0.215 | 0.184 | 0.076 | 2.342 | 0.115 |
| 2346 | **Ndufa1** | Unchanged | O35683 | 1 | 3 | 1 | 70 | 8.1 | 9.48 | 14 | 0.126 | 0.154 | 0.225 | 0.201 | 0.655 | 0.601 | 0.344 | 0.210 |
| 2347 | **Cox7c** | Unchanged | P17665 | 1 | 16 | 1 | 63 | 7.3 | 11 | 14 | 0.607 | 0.270 | 0.793 | 0.565 | 1.966 | 0.293 | 0.403 | 0.182 |
| 2348 | **Pex3** | Unchanged | Q9QXY9 | 3 | 16 | 3 | 372 | 42.2 | 7.77 | 14 | 0.806 | 0.658 | 0.997 | 0.996 | 0.825 | 0.658 | 1.209 | 0.757 |
| 2349 | **Sf1** | Unchanged | Q64213 | 6 | 33 | 6 | 653 | 70.4 | 8.98 | 14 | 0.478 | 0.191 | 0.739 | 0.544 | 1.185 | 0.684 | 0.623 | 0.243 |
| 2350 | **Akr1b10** | Unchanged | G5E895 | 4 | 15 | 1 | 316 | 35.8 | 7.28 | 14 | 1.419 | 0.167 | 1.296 | 0.299 | 1.541 | 0.211 | 0.841 | 0.528 |
| 2351 | **Akr1b7** | Unchanged | P21300 | 4 | 28 | 1 | 316 | 36 | 7.25 | 14 | 4.331 | 0.272 | 19.721 | 0.331 | 1.025 | 0.919 | 19.244 | 0.332 |
| 2352 | **Ptpn11** | Unchanged | P35235 | 5 | 23 | 5 | 597 | 68.4 | 7.3 | 14 | 1.681 | 0.203 | 1.500 | 0.532 | 0.963 | 0.904 | 1.558 | 0.481 |
| 2353 | **Naa15** | Unchanged | Q80UM3 | 8 | 31 | 7 | 865 | 100.9 | 7.62 | 14 | 7.259 | 0.285 | 3.192 | 0.235 | 1.673 | 0.341 | 1.908 | 0.430 |
| 2354 | **Cers2** | Unchanged | Q924Z4 | 3 | 50 | 3 | 380 | 45 | 8.75 | 14 | 0.944 | 0.910 | 0.590 | 0.309 | 0.924 | 0.870 | 0.638 | 0.411 |
| 2355 | **Psmg1** | Unchanged | Q9JK23 | 3 | 7 | 3 | 289 | 33.1 | 6.37 | 14 | 0.855 | 0.749 | 1.114 | 0.843 | 0.716 | 0.461 | 1.556 | 0.451 |
| 2356 | **Nploc4** | UP | P60670 | 5 | 18 | 5 | 608 | 68 | 6.46 | 14 | 1.721 | 0.137 | 1.910 | 0.008 | 0.997 | 0.992 | 1.915 | 0.003 |
| 2357 | **Guk1** | Unchanged | Q64520 | 2 | 14 | 2 | 198 | 21.9 | 6.55 | 14 | 1.303 | 0.688 | 4.108 | 0.086 | 1.953 | 0.398 | 2.103 | 0.272 |
| 2358 | **Uba7** | Unchanged | Q9DBK7 | 5 | 9 | 5 | 977 | 108.6 | 5.97 | 14 | 1.568 | 0.280 | 2.947 | 0.092 | 0.812 | 0.396 | 3.627 | 0.071 |
| 2359 | **Cul4b** | Unchanged | A2A432 | 10 | 28 | 5 | 970 | 110.6 | 8.37 | 14 | 0.462 | 0.402 | 0.845 | 0.821 | 0.181 | 0.182 | 4.661 | 0.101 |
| 2360 | **Mrpl20** | Unchanged | Q9CQL4 | 2 | 4 | 2 | 149 | 17.6 | 11.11 | 14 | 1.365 | 0.324 | 1.988 | 0.216 | 1.782 | 0.109 | 1.116 | 0.800 |
| 2361 | **Bag1** | Unchanged | Q60739 | 2 | 4 | 2 | 355 | 39.7 | 8.46 | 14 | 0.595 | 0.080 | 1.471 | 0.505 | 0.630 | 0.084 | 2.333 | 0.242 |
| 2362 | **Lars2** | Unchanged | Q8VDC0 | 5 | 11 | 5 | 902 | 101.4 | 8.19 | 14 | 1.390 | 0.344 | 1.943 | 0.096 | 1.226 | 0.520 | 1.585 | 0.264 |
| 2363 | **Surf1** | Unchanged | P09925 | 2 | 11 | 2 | 306 | 34.8 | 9.72 | 14 | 0.691 | 0.393 | 0.581 | 0.254 | 0.773 | 0.538 | 0.751 | 0.568 |
| 2364 | **Mrpl2** | Unchanged | Q9D773 | 2 | 4 | 2 | 306 | 33.3 | 11.06 | 14 | 1.075 | 0.907 | 1.532 | 0.408 | 0.474 | 0.364 | 3.233 | 0.009 |
| 2365 | **Mrpl28** | Unchanged | Q9D1B9 | 2 | 22 | 2 | 257 | 30.2 | 9.29 | 14 | 2.398 | 0.142 | 3.007 | 0.067 | 2.724 | 0.062 | 1.104 | 0.791 |
| 2366 | **Igtp** | Unchanged | Q9DCE9 | 5 | 18 | 5 | 423 | 48.5 | 7.77 | 14 | 0.923 | 0.907 | 0.748 | 0.686 | 0.682 | 0.590 | 1.095 | 0.893 |
| 2367 | **Bin1** | Unchanged | O08539 | 4 | 13 | 4 | 588 | 64.4 | 5.03 | 14 | 0.763 | 0.192 | 2.761 | 0.153 | 2.925 | 0.060 | 0.944 | 0.910 |

| | | | | | | | | | | | | | | | | | | |
|---|---|---|---|---|---|---|---|---|---|---|---|---|---|---|---|---|---|---|
| 2368 | **Sts** | Unchanged | P50427 | 6 | 30 | 6 | 624 | 66.5 | 8.5 | 14 | 0.883 | 0.533 | 1.668 | 0.095 | 1.475 | 0.062 | 1.131 | 0.592 |
| 2369 | **Slco1b2** | Unchanged | Q9JJL3 | 6 | 84 | 6 | 689 | 76.7 | 8.68 | 14 | 1.044 | 0.931 | 0.827 | 0.720 | 1.204 | 0.685 | 0.687 | 0.454 |
| 2370 | **Igkv19-93** | Down | A0A0G2JFZ3 | 1 | 1 | 1 | 115 | 12.8 | 8.29 | 14 | 0.114 | 0.046 | 2.335 | 0.313 | 1.205 | 0.757 | 1.937 | 0.407 |
| 2371 | **Serpina3n** | Unchanged | Q91WP6 | 5 | 74 | 3 | 418 | 46.7 | 5.82 | 14 | 0.918 | 0.883 | 1.047 | 0.950 | 1.523 | 0.427 | 0.687 | 0.592 |
| 2372 | **Ndufaf4** | Down | Q9D1H6 | 2 | 5 | 2 | 173 | 20.1 | 9.39 | 14 | 1.205 | 0.725 | 1.131 | 0.785 | 0.347 | 0.013 | 3.264 | 0.104 |
| 2373 | **Tradd** | Unchanged | Q3U0V2 | 2 | 18 | 2 | 310 | 34.6 | 5.2 | 14 | 0.967 | 0.927 | 1.405 | 0.261 | 0.636 | 0.150 | 2.209 | 0.039 |
| 2374 | **Arfgap2** | Unchanged | Q99K28 | 5 | 23 | 5 | 520 | 56.6 | 8.18 | 14 | 1.540 | 0.455 | 1.437 | 0.490 | 1.314 | 0.647 | 1.094 | 0.872 |
| 2375 | **Mesd** | Down | Q9ERE7 | 3 | 7 | 3 | 224 | 25.2 | 6.34 | 14 | 0.280 | 0.047 | 0.556 | 0.316 | 0.441 | 0.127 | 1.262 | 0.719 |
| 2376 | **Ggct** | UP | Q9D7X8 | 2 | 4 | 2 | 188 | 21.2 | 5.67 | 14 | 3.171 | 0.004 | 3.203 | 0.032 | 5.174 | 0.120 | 0.619 | 0.450 |
| 2377 | **Tomm20** | Unchanged | Q9DCC8 | 1 | 15 | 1 | 145 | 16.3 | 8.6 | 14 | 1.030 | 0.967 | 2.027 | 0.309 | 1.025 | 0.970 | 1.977 | 0.307 |
| 2378 | **Vti1b** | Unchanged | O88384 | 2 | 3 | 2 | 232 | 26.7 | 8.79 | 14 | 0.972 | 0.964 | 1.362 | 0.648 | 0.297 | 0.170 | 4.591 | 0.120 |
| 2379 | **Mp68** | Down | P56379 | 2 | 19 | 2 | 58 | 6.7 | 9.99 | 14 | 2.135 | 0.110 | 0.376 | 0.044 | 0.476 | 0.081 | 0.790 | 0.472 |
| 2380 | **Mrpl14** | Unchanged | Q9D1I6 | 2 | 6 | 2 | 145 | 15.9 | 10.99 | 14 | 1.123 | 0.851 | 1.354 | 0.669 | 0.439 | 0.296 | 3.085 | 0.217 |
| 2381 | **Olfr1298** | Unchanged | Q8VGE6 | 2 | 2 | 2 | 312 | 35.3 | 8.22 | 14 | 0.180 | 0.225 | 11.755 | 0.276 | 0.212 | 0.243 | 55.356 | 0.244 |
| 2382 | **Raly** | Down | Q64012 | 3 | 10 | 3 | 312 | 33.2 | 8.84 | 14 | 0.506 | 0.054 | 0.336 | 0.012 | 0.583 | 0.182 | 0.576 | 0.356 |
| 2383 | **Psme3** | Unchanged | P61290 | 3 | 12 | 3 | 254 | 29.5 | 5.95 | 14 | 1.598 | 0.357 | 1.466 | 0.325 | 0.951 | 0.934 | 1.541 | 0.432 |
| 2384 | **Rpl19** | Unchanged | P84099 | 3 | 52 | 3 | 196 | 23.5 | 11.47 | 14 | 0.886 | 0.718 | 0.705 | 0.480 | 0.992 | 0.983 | 0.711 | 0.560 |
| 2385 | **Nt5c2** | Unchanged | Q3V1L4 | 4 | 6 | 4 | 560 | 64.8 | 6.21 | 14 | 1.289 | 0.695 | 0.283 | 0.184 | 0.255 | 0.165 | 1.110 | 0.827 |
| 2386 | **Mrpl30** | Down | Q9D7N6 | 1 | 3 | 1 | 160 | 18.3 | 10.15 | 14 | 0.812 | 0.716 | 0.559 | 0.285 | 0.138 | 0.018 | 4.052 | 0.141 |
| 2387 | **Tm9sf2** | Unchanged | P58021 | 6 | 25 | 6 | 662 | 75.3 | 7.43 | 14 | 1.502 | 0.437 | 2.620 | 0.096 | 1.252 | 0.674 | 2.092 | 0.134 |
| 2388 | **Stau1** | Unchanged | A2A5S3 | 5 | 13 | 5 | 495 | 54.8 | 9.52 | 14 | 0.808 | 0.673 | 0.891 | 0.837 | 1.578 | 0.310 | 0.564 | 0.176 |
| 2389 | **Rras2** | Unchanged | P62071 | 2 | 8 | 2 | 204 | 23.4 | 6.01 | 14 | 0.793 | 0.682 | 1.505 | 0.521 | 0.215 | 0.073 | 6.999 | 0.084 |
| 2390 | **Tax1bp3** | Unchanged | Q9DBG9 | 1 | 10 | 1 | 124 | 13.7 | 8.48 | 14 | 0.649 | 0.361 | 1.304 | 0.500 | 1.150 | 0.685 | 1.135 | 0.699 |
| 2391 | **Cda** | Unchanged | P56389 | 1 | 2 | 1 | 146 | 16.1 | 5.58 | 14 | 2.309 | 0.204 | 2.921 | 0.122 | 0.820 | 0.749 | 3.561 | 0.076 |
| 2392 | **Nmi** | Unchanged | O35309 | 2 | 27 | 2 | 314 | 35.2 | 5.05 | 14 | 1.146 | 0.528 | 1.070 | 0.858 | 0.915 | 0.788 | 1.169 | 0.726 |
| 2393 | **Mrpl38** | Unchanged | Q8K2M0 | 3 | 15 | 3 | 380 | 45 | 8.1 | 14 | 0.718 | 0.213 | 2.226 | 0.390 | 1.960 | 0.249 | 1.135 | 0.867 |
| 2394 | **Igkv17-127** | Down | A0A0G2JDN5 | 1 | 4 | 1 | 117 | 12.8 | 4.36 | 14 | 0.430 | 0.120 | 2.829 | 0.442 | 0.341 | 0.034 | 8.288 | 0.302 |
| 2395 | **Hars2** | Unchanged | Q99KK9 | 6 | 29 | 3 | 505 | 56.9 | 8.32 | 14 | 1.193 | 0.670 | 1.995 | 0.282 | 0.659 | 0.213 | 3.029 | 0.145 |
| 2396 | **Ube2h** | Unchanged | P62257 | 2 | 6 | 2 | 183 | 20.6 | 4.67 | 14 | 2.715 | 0.098 | 2.052 | 0.359 | 1.167 | 0.814 | 1.758 | 0.423 |
| 2397 | **Dnajb4** | Unchanged | Q9D832 | 3 | 9 | 2 | 337 | 37.8 | 8.59 | 14 | 1.654 | 0.276 | 1.563 | 0.341 | 2.015 | 0.123 | 0.776 | 0.437 |
| 2398 | **Odr4** | Unchanged | Q4PJX1 | 3 | 9 | 3 | 447 | 50 | 6.14 | 14 | 0.903 | 0.847 | 1.032 | 0.959 | 0.301 | 0.113 | 3.425 | 0.166 |
| 2399 | **Dhrs7b** | Unchanged | Q99J47 | 4 | 7 | 4 | 323 | 35 | 9.63 | 14 | 0.472 | 0.376 | 0.488 | 0.392 | 0.194 | 0.189 | 2.513 | 0.028 |
| 2400 | **Hist1h1a** | Unchanged | P43275 | 2 | 28 | 1 | 213 | 21.8 | 10.93 | 14 | 2.224 | 0.506 | 2.690 | 0.390 | 51.557 | 0.053 | 0.052 | 0.060 |
| 2401 | **Icam1** | Down | P13597 | 5 | 15 | 5 | 537 | 58.8 | 6.13 | 14 | 0.595 | 0.075 | 0.849 | 0.483 | 0.475 | 0.039 | 1.788 | 0.050 |
| 2402 | **Hspg2** | Unchanged | E9PZ16 | 32 | 110 | 32 | 4383 | 469.5 | 6.47 | 14 | 1.494 | 0.202 | 2.869 | 0.119 | 2.404 | 0.075 | 1.193 | 0.718 |
| 2403 | **Rab27a** | Unchanged | Q9ERI2 | 1 | 1 | 1 | 221 | 25 | 5.36 | 14 | 2.250 | 0.077 | 149.636 | 0.242 | 301.682 | 0.068 | 0.496 | 0.435 |
| 2404 | **Rfk** | Unchanged | Q8CFV9 | 1 | 6 | 1 | 155 | 17.4 | 7.55 | 14 | 3.535 | 0.355 | 9.608 | 0.142 | 0.715 | 0.266 | 13.444 | 0.130 |
| 2405 | **Mrps31** | Unchanged | Q61733 | 3 | 44 | 2 | 384 | 43.9 | 8.51 | 14 | 1.458 | 0.496 | 0.988 | 0.982 | 0.806 | 0.633 | 1.225 | 0.750 |
| 2406 | **Ptma** | Unchanged | P26350 | 3 | 62 | 3 | 111 | 12.2 | 3.79 | 14 | 0.282 | 0.364 | 13.276 | 0.352 | 1.030 | 0.978 | 12.894 | 0.353 |
| 2407 | **Dpysl3** | Unchanged | Q62188 | 4 | 35 | 3 | 570 | 61.9 | 6.49 | 14 | 1.910 | 0.322 | 0.995 | 0.989 | 0.667 | 0.463 | 1.493 | 0.496 |
| 2408 | **Serhl** | Unchanged | Q9EPB5 | 2 | 3 | 2 | 311 | 35.3 | 8.1 | 14 | 1.138 | 0.746 | 7.311 | 0.072 | 1.156 | 0.727 | 6.322 | 0.080 |

| | | | | | | | | | | | | | | | | | | |
|---|---|---|---|---|---|---|---|---|---|---|---|---|---|---|---|---|---|---|
| 2409 | **Pycr3** | Unchanged | Q9DCC4 | 2 | 17 | 2 | 274 | 28.7 | 7.27 | 14 | 0.846 | 0.438 | 1.356 | 0.650 | 0.619 | 0.086 | 2.191 | 0.352 |
| 2410 | **Diablo** | Unchanged | Q9JIQ3 | 1 | 1 | 1 | 237 | 26.8 | 6.37 | 14 | 2.543 | 0.253 | 1.729 | 0.470 | 1.300 | 0.630 | 1.330 | 0.700 |
| 2411 | **Tha1** | Down | Q6XPS7 | 3 | 4 | 3 | 400 | 43.5 | 7.2 | 14 | 1.345 | 0.362 | 2.321 | 0.244 | 0.459 | 0.024 | 5.058 | 0.111 |
| 2412 | **Pgghg** | Unchanged | Q8BP56 | 5 | 23 | 5 | 690 | 76.4 | 5.33 | 13 | 0.740 | 0.186 | 1.831 | 0.071 | 0.638 | 0.073 | 2.871 | 0.018 |
| 2413 | **Mocos** | Unchanged | Q14CH1 | 7 | 23 | 7 | 862 | 95 | 6.62 | 13 | 0.984 | 0.973 | 1.426 | 0.470 | 0.655 | 0.295 | 2.179 | 0.150 |
| 2414 | **Fbl** | Unchanged | P35550 | 2 | 6 | 2 | 327 | 34.3 | 10.24 | 13 | 0.946 | 0.941 | 1.128 | 0.873 | 0.944 | 0.944 | 1.195 | 0.752 |
| 2415 | **Ighv8-12** | Unchanged | A0A0G2JDE1 | 1 | 1 | 1 | 119 | 13.3 | 9.17 | 13 | 1.278 | 0.829 | 9.116 | 0.277 | 0.343 | 0.383 | 26.599 | 0.241 |
| 2416 | **Myo6** | Unchanged | E9PVU0 | 10 | 13 | 10 | 1266 | 146.2 | 8.47 | 13 | 0.643 | 0.468 | 0.926 | 0.878 | 0.341 | 0.153 | 2.714 | 0.029 |
| 2417 | **Hmgn2** | Unchanged | F6W687 | 1 | 1 | 1 | 112 | 11.4 | 9.83 | 13 | #DIV/0! | #DIV/0! | #DIV/0! | #DIV/0! | #DIV/0! | #DIV/0! | #DIV/0! | #DIV/0! |
| 2418 | **Adss** | Unchanged | P46664 | 4 | 10 | 4 | 456 | 50 | 6.38 | 13 | 0.609 | 0.225 | 1.239 | 0.635 | 1.772 | 0.337 | 0.699 | 0.535 |
| 2419 | **Bcl2l13** | Unchanged | P59017 | 4 | 39 | 4 | 434 | 46.7 | 4.59 | 13 | 0.942 | 0.893 | 4.513 | 0.105 | 1.913 | 0.320 | 2.359 | 0.241 |
| 2420 | **Dctn1** | Unchanged | O08788 | 10 | 28 | 10 | 1281 | 141.6 | 5.9 | 13 | 0.625 | 0.054 | 1.032 | 0.918 | 0.677 | 0.110 | 1.524 | 0.230 |
| 2421 | **Iqgap1** | Unchanged | Q9JKF1 | 13 | 65 | 10 | 1657 | 188.6 | 6.48 | 13 | 1.446 | 0.528 | 1.126 | 0.766 | 1.357 | 0.500 | 0.830 | 0.622 |
| 2422 | **Paqr9** | Unchanged | Q6TCG2 | 3 | 23 | 3 | 375 | 42.7 | 8.76 | 13 | 1.018 | 0.982 | 0.461 | 0.328 | 0.430 | 0.298 | 1.071 | 0.811 |
| 2423 | **Prkab1** | UP | Q9R078 | 2 | 3 | 2 | 270 | 30.3 | 6.23 | 13 | 0.555 | 0.384 | 2.823 | 0.420 | 6.307 | 0.047 | 0.448 | 0.286 |
| 2424 | **Flot1** | Down | O08917 | 4 | 12 | 4 | 428 | 47.5 | 7.15 | 13 | 0.577 | 0.140 | 0.250 | 0.024 | 0.169 | 0.009 | 1.477 | 0.564 |
| 2425 | **Igh-1a** | Unchanged | P01865 | 4 | 45 | 2 | 398 | 43.9 | 6.29 | 13 | 0.557 | 0.160 | 13.893 | 0.234 | 0.575 | 0.158 | 24.149 | 0.220 |
| 2426 | **Tmx1** | Down | Q8VBT0 | 3 | 11 | 3 | 278 | 31.4 | 5.29 | 13 | 1.315 | 0.489 | 1.497 | 0.354 | 0.305 | 0.041 | 4.914 | 0.025 |
| 2427 | **Pck2** | UP | Q8BH04 | 6 | 12 | 6 | 640 | 70.5 | 7.28 | 13 | 2.635 | 0.166 | 7.462 | 0.036 | 1.976 | 0.106 | 3.775 | 0.068 |
| 2428 | **Col6a3** | Unchanged | E9PWQ3 | 27 | 92 | 27 | 3284 | 353.7 | 6.93 | 13 | 1.120 | 0.811 | 1.158 | 0.703 | 1.070 | 0.871 | 1.082 | 0.787 |
| 2429 | **Fmc1** | Down | Q9CR13 | 1 | 2 | 1 | 113 | 12.7 | 10.08 | 13 | 0.849 | 0.737 | 0.573 | 0.345 | 0.223 | 0.041 | 2.576 | 0.328 |
| 2430 | **Crybb3** | Unchanged | Q9JJU9 | 2 | 4 | 2 | 211 | 24.3 | 7.23 | 13 | 0.311 | 0.180 | 0.395 | 0.272 | 0.267 | 0.154 | 1.481 | 0.593 |
| 2431 | **Hspb8** | Unchanged | Q9JK92 | 1 | 11 | 1 | 196 | 21.5 | 5.02 | 13 | 2.054 | 0.299 | 1.830 | 0.312 | 1.722 | 0.474 | 1.063 | 0.905 |
| 2432 | **Lmnb2** | Unchanged | P21619 | 5 | 22 | 5 | 596 | 67.3 | 5.5 | 13 | 0.847 | 0.711 | 0.922 | 0.883 | 0.660 | 0.443 | 1.397 | 0.520 |
| 2433 | **Timm8b** | Unchanged | P62077 | 1 | 10 | 1 | 83 | 9.3 | 5.12 | 13 | 2.256 | 0.317 | 2.661 | 0.412 | 2.019 | 0.462 | 1.318 | 0.756 |
| 2434 | **G6pdx** | Unchanged | Q00612 | 5 | 19 | 5 | 515 | 59.2 | 6.49 | 13 | 0.931 | 0.894 | 0.614 | 0.377 | 1.584 | 0.436 | 0.387 | 0.148 |
| 2435 | **Akap1** | Unchanged | O08715 | 6 | 27 | 6 | 857 | 92.1 | 5.02 | 13 | 0.735 | 0.278 | 0.703 | 0.244 | 0.602 | 0.184 | 1.167 | 0.718 |
| 2436 | **Rwdd1** | Unchanged | Q9CQK7 | 1 | 1 | 1 | 243 | 27.8 | 4.26 | 13 | #DIV/0! | #DIV/0! | #DIV/0! | #DIV/0! | #DIV/0! | #DIV/0! | #DIV/0! | #DIV/0! |
| 2437 | **Zglp1** | Unchanged | Q1WG82 | 1 | 1 | 1 | 266 | 29.1 | 9.11 | 13 | 1.622 | 0.268 | 1.551 | 0.627 | 1.727 | 0.107 | 0.898 | 0.882 |
| 2438 | **Ninj1** | Unchanged | O70131 | 1 | 30 | 1 | 152 | 16.5 | 6.55 | 13 | 0.964 | 0.931 | 0.888 | 0.849 | 1.229 | 0.594 | 0.722 | 0.510 |
| 2439 | **Fam210b** | Down | Q9D8B6 | 2 | 3 | 2 | 190 | 20.3 | 10.51 | 13 | 0.473 | 0.170 | 1.431 | 0.409 | 0.208 | 0.041 | 6.885 | 0.011 |
| 2440 | **Caprin1** | Down | Q60865 | 7 | 98 | 7 | 707 | 78.1 | 5.25 | 13 | 0.306 | 0.022 | 0.360 | 0.032 | 0.224 | 0.012 | 1.607 | 0.050 |
| 2441 | **Usp15** | Unchanged | Q8R5H1 | 7 | 29 | 7 | 981 | 112.3 | 5.17 | 13 | 1.262 | 0.510 | 2.046 | 0.333 | 1.673 | 0.200 | 1.222 | 0.735 |
| 2442 | **Olfr412** | Unchanged | Q7TRW7 | 1 | 1 | 1 | 312 | 35.2 | 8.72 | 13 | 11.693 | 0.145 | 1.671 | 0.320 | 7.566 | 0.178 | 0.221 | 0.225 |
| 2443 | **Osgep** | Unchanged | A0A0R4J1Y3 | 3 | 22 | 3 | 335 | 36.3 | 6.24 | 13 | 1.133 | 0.642 | 0.882 | 0.701 | 1.455 | 0.206 | 0.606 | 0.199 |
| 2444 | **Dcun1d1** | Unchanged | Q9QZ73 | 2 | 9 | 2 | 259 | 30.1 | 5.34 | 13 | 0.874 | 0.807 | 1.758 | 0.486 | 1.043 | 0.941 | 1.685 | 0.526 |
| 2445 | **Ptgs1** | Unchanged | P22437 | 4 | 22 | 4 | 602 | 69 | 6.83 | 13 | 0.509 | 0.129 | 1.047 | 0.916 | 0.483 | 0.112 | 2.166 | 0.195 |
| 2446 | **Stk16** | Unchanged | O88697 | 2 | 7 | 2 | 305 | 34.4 | 6.46 | 13 | 3.225 | 0.261 | 6.599 | 0.160 | 3.712 | 0.112 | 1.778 | 0.477 |
| 2447 | **Dtd2** | Unchanged | Q8BHA3 | 1 | 19 | 1 | 168 | 18.2 | 7.91 | 13 | 1.153 | 0.768 | 1.341 | 0.459 | 1.487 | 0.309 | 0.902 | 0.567 |
| 2448 | **Fxyd1** | Unchanged | Q9Z239 | 1 | 6 | 1 | 92 | 10.3 | 8.54 | 13 | 0.362 | 0.229 | 0.228 | 0.138 | 3.520 | 0.451 | 0.065 | 0.328 |
| 2449 | **Mtnd3** | Unchanged | P03899 | 1 | 8 | 1 | 115 | 13.2 | 4.64 | 13 | 0.986 | 0.986 | 0.370 | 0.304 | 0.707 | 0.676 | 0.523 | 0.463 |

| | | | | | | | | | | | | | | | | | |
|---|---|---|---|---|---|---|---|---|---|---|---|---|---|---|---|---|---|
| 2450 | **Them4** | Unchanged | Q3UUI3 | 2 | 8 | 2 | 230 | 26 | 9.64 | 13 | 1.821 | 0.211 | 2.247 | 0.054 | 0.889 | 0.805 | 2.528 | 0.068 |
| 2451 | **Serpinf2** | Unchanged | Q61247 | 4 | 29 | 4 | 491 | 54.9 | 6.3 | 13 | 0.785 | 0.279 | 0.718 | 0.347 | 1.147 | 0.611 | 0.626 | 0.229 |
| 2452 | **Gyg** | Unchanged | K3W4S6 | 2 | 7 | 2 | 377 | 41.9 | 6.02 | 13 | 1.293 | 0.728 | 0.543 | 0.495 | 0.950 | 0.952 | 0.571 | 0.530 |
| 2453 | **Cbr3** | Unchanged | Q8K354 | 2 | 18 | 1 | 277 | 30.9 | 6.57 | 13 | 0.544 | 0.085 | 2.741 | 0.357 | 0.974 | 0.930 | 2.813 | 0.352 |
| 2454 | **Acot9** | Down | Q9R0X4 | 4 | 19 | 4 | 439 | 50.5 | 8.59 | 13 | 1.321 | 0.207 | 1.682 | 0.237 | 0.491 | 0.017 | 3.426 | 0.048 |
| 2455 | **Gm9970** | Unchanged | F6ZED4 | 1 | 1 | 1 | 185 | 19.1 | 11.37 | 13 | 0.253 | 0.071 | 0.864 | 0.831 | 0.195 | 0.053 | 4.443 | 0.223 |
| 2456 | **Rbx1** | Unchanged | P62878 | 2 | 9 | 2 | 108 | 12.3 | 6.96 | 13 | 1.183 | 0.598 | 1.208 | 0.568 | 0.769 | 0.383 | 1.570 | 0.153 |
| 2457 | **Cyp4b1** | Unchanged | Q64462 | 6 | 15 | 6 | 511 | 58.9 | 8.44 | 13 | 1.926 | 0.146 | 2.162 | 0.111 | 1.355 | 0.580 | 1.595 | 0.368 |
| 2458 | **Drg2** | Unchanged | Q9QXB9 | 4 | 15 | 3 | 364 | 40.7 | 8.88 | 13 | 2.649 | 0.349 | 2.566 | 0.150 | 1.244 | 0.651 | 2.062 | 0.239 |
| 2459 | **Fam162a** | Unchanged | Q9D6U8 | 2 | 25 | 2 | 155 | 17.7 | 9.88 | 13 | 1.352 | 0.627 | 0.625 | 0.403 | 0.994 | 0.991 | 0.629 | 0.411 |
| 2460 | **Parva** | Unchanged | Q9EPC1 | 4 | 22 | 3 | 372 | 42.3 | 5.95 | 13 | 5.997 | 0.334 | 1.482 | 0.296 | 1.762 | 0.070 | 0.841 | 0.622 |
| 2461 | **Olfr222** | UP | Q7TS30 | 1 | 1 | 1 | 318 | 35.7 | 8.57 | 13 | 0.898 | 0.724 | 2.732 | 0.381 | 2.096 | 0.045 | 1.303 | 0.745 |
| 2462 | **Ccdc90b** | UP | Q8C3X2 | 2 | 2 | 2 | 256 | 29.6 | 8.53 | 13 | 4.575 | 0.008 | 5.715 | 0.035 | 4.033 | 0.083 | 1.417 | 0.504 |
| 2463 | **Bcas2** | Down | Q9D287 | 2 | 3 | 2 | 225 | 26.1 | 5.66 | 13 | 0.528 | 0.218 | 0.467 | 0.137 | 0.186 | 0.018 | 2.511 | 0.151 |
| 2464 | **Cul1** | Unchanged | Q9WTX6 | 7 | 23 | 7 | 776 | 89.6 | 8 | 13 | 0.814 | 0.658 | 1.703 | 0.114 | 0.707 | 0.414 | 2.411 | 0.005 |
| 2465 | **Ciao2b** | Unchanged | Q9D187 | 1 | 1 | 1 | 163 | 17.7 | 4.98 | 13 | 0.832 | 0.778 | 1.275 | 0.687 | 0.244 | 0.125 | 5.216 | 0.096 |
| 2466 | **Tatdn1** | Down | Q6P8M1 | 3 | 4 | 3 | 295 | 33.3 | 6.18 | 13 | 0.974 | 0.949 | 0.183 | 0.027 | 0.606 | 0.472 | 0.302 | 0.351 |
| 2467 | **Fcgrt** | Unchanged | Q61559 | 3 | 15 | 3 | 365 | 40.1 | 5.29 | 13 | 0.542 | 0.282 | 0.868 | 0.729 | 0.345 | 0.145 | 2.515 | 0.062 |
| 2468 | **Abcb10** | Unchanged | Q9JI39 | 6 | 28 | 6 | 715 | 77.1 | 9.6 | 13 | 1.629 | 0.373 | 1.317 | 0.443 | 2.224 | 0.078 | 0.592 | 0.241 |
| 2469 | **Lrg1** | Unchanged | Q91XL1 | 3 | 11 | 3 | 342 | 37.4 | 6.61 | 13 | 0.447 | 0.319 | 0.571 | 0.459 | 0.076 | 0.108 | 7.520 | 0.045 |
| 2470 | **Mt-Cyb** | Unchanged | P00158 | 2 | 19 | 2 | 381 | 43.2 | 7.97 | 13 | 2.969 | 0.244 | 0.937 | 0.885 | 0.463 | 0.141 | 2.023 | 0.237 |
| 2471 | **Sirt2** | Unchanged | Q8VDQ8 | 3 | 16 | 3 | 389 | 43.2 | 5.35 | 13 | 1.037 | 0.959 | 0.772 | 0.697 | 1.165 | 0.808 | 0.663 | 0.498 |
| 2472 | **Casq1** | Unchanged | O09165 | 3 | 6 | 3 | 405 | 46.3 | 4.12 | 13 | 0.557 | 0.543 | 0.515 | 0.527 | 0.518 | 0.501 | 0.995 | 0.994 |
| 2473 | **Tcirg1** | Unchanged | Q9JHF5 | 7 | 15 | 7 | 834 | 93.4 | 6.38 | 13 | 1.983 | 0.228 | 2.265 | 0.070 | 1.629 | 0.243 | 1.390 | 0.365 |
| 2474 | **Fam98b** | Unchanged | Q80VD1 | 4 | 13 | 4 | 429 | 45.3 | 8.5 | 13 | 0.916 | 0.853 | 0.932 | 0.910 | 0.670 | 0.488 | 1.391 | 0.630 |
| 2475 | **Ighv1-18** | Unchanged | A0A0A6YXN4 | 1 | 1 | 1 | 117 | 12.9 | 7.94 | 13 | #DIV/0! | #DIV/0! | #DIV/0! | #DIV/0! | #DIV/0! | #DIV/0! | #DIV/0! | #DIV/0! |
| 2476 | **Fam210a** | Unchanged | Q8BGY7 | 2 | 22 | 2 | 273 | 31.6 | 9.82 | 13 | 1.109 | 0.896 | 0.867 | 0.870 | 1.733 | 0.462 | 0.500 | 0.339 |
| 2477 | **Mrps7** | Unchanged | Q80X85 | 2 | 16 | 2 | 242 | 28 | 9.94 | 13 | 1.019 | 0.975 | 2.098 | 0.290 | 1.553 | 0.579 | 1.351 | 0.652 |
| 2478 | **Srsf3** | Unchanged | P84104 | 2 | 38 | 1 | 164 | 19.3 | 11.65 | 13 | 1.902 | 0.327 | 2.042 | 0.505 | 2.987 | 0.066 | 0.684 | 0.584 |
| 2479 | **Srrt** | Unchanged | Q99MR6 | 7 | 25 | 7 | 875 | 100.4 | 5.97 | 13 | 1.757 | 0.391 | 1.311 | 0.666 | 0.151 | 0.117 | 8.655 | 0.049 |
| 2480 | **Snf8** | Unchanged | Q9CZ28 | 2 | 12 | 2 | 258 | 28.9 | 6.65 | 13 | 1.020 | 0.977 | 0.172 | 0.157 | 1.793 | 0.324 | 0.096 | 0.017 |
| 2481 | **Eps15l1** | Unchanged | Q60902 | 8 | 30 | 8 | 907 | 99.2 | 5.02 | 13 | 2.230 | 0.309 | 2.447 | 0.185 | 2.285 | 0.259 | 1.071 | 0.906 |
| 2482 | **Emc1** | Unchanged | Q8C7X2 | 7 | 33 | 7 | 997 | 111.5 | 7.43 | 13 | 1.370 | 0.277 | 1.227 | 0.461 | 1.274 | 0.119 | 0.963 | 0.879 |
| 2483 | **Pxmp4** | Unchanged | Q9JJW0 | 2 | 9 | 2 | 212 | 24.2 | 9.83 | 6 | 1.587 | 0.499 | 2.384 | 0.369 | 0.835 | 0.802 | 2.857 | 0.315 |
| 2484 | **Slc25a30** | Unchanged | Q9CR58 | 1 | 1 | 1 | 291 | 32.3 | 9.38 | 13 | 0.964 | 0.925 | 1.134 | 0.751 | 3.069 | 0.090 | 0.370 | 0.113 |
| 2485 | **Pafah1b1** | Unchanged | P63005 | 3 | 31 | 3 | 410 | 46.6 | 7.37 | 13 | 2.104 | 0.363 | 2.425 | 0.334 | 1.681 | 0.486 | 1.443 | 0.643 |
| 2486 | **Lancl2** | Unchanged | Q9JJK2 | 2 | 8 | 2 | 450 | 50.7 | 7.28 | 13 | 1.099 | 0.677 | 2.181 | 0.287 | 1.050 | 0.862 | 2.078 | 0.318 |
| 2487 | **Fam114a1** | Unchanged | Q9D281 | 3 | 3 | 3 | 569 | 61 | 4.56 | 10 | 1.046 | 0.811 | 4.343 | 0.069 | 0.810 | 0.342 | 5.363 | 0.058 |
| 2488 | **Mtfp1** | Unchanged | Q9CRB8 | 1 | 2 | 1 | 166 | 18.3 | 8.68 | 13 | 1.578 | 0.524 | 2.469 | 0.256 | 0.704 | 0.425 | 3.508 | 0.174 |
| 2489 | **Acsl3** | Down | Q9CZW4 | 5 | 10 | 5 | 720 | 80.4 | 8.54 | 13 | 0.475 | 0.015 | 0.398 | 0.005 | 1.007 | 0.981 | 0.395 | 0.073 |
| 2490 | **Cxadr** | Unchanged | P97792 | 3 | 12 | 3 | 365 | 39.9 | 6.96 | 13 | 0.321 | 0.086 | 0.766 | 0.592 | 1.195 | 0.703 | 0.641 | 0.371 |

| | | | | | | | | | | | | | | | | | | |
|---|---|---|---|---|---|---|---|---|---|---|---|---|---|---|---|---|---|---|
| 2491 | **Map2k4** | Unchanged | P47809 | 3 | 15 | 3 | 397 | 44.1 | 8.07 | 13 | 0.536 | 0.061 | 0.620 | 0.206 | 0.757 | 0.493 | 0.819 | 0.680 |
| 2492 | **Tln2** | Unchanged | Q71LX4 | 16 | 85 | 10 | 2375 | 253.5 | 5.8 | 13 | 1.119 | 0.489 | 1.131 | 0.661 | 0.855 | 0.612 | 1.323 | 0.471 |
| 2493 | **Pqbp1** | Unchanged | Q91VJ5 | 1 | 1 | 1 | 263 | 30.6 | 6.23 | 13 | 1.438 | 0.473 | 2.793 | 0.176 | 2.953 | 0.143 | 0.946 | 0.922 |
| 2494 | **Lmf2** | Unchanged | Q8C3X8 | 4 | 24 | 4 | 702 | 79.9 | 9.99 | 13 | 0.228 | 0.297 | 0.654 | 0.639 | 0.512 | 0.503 | 1.278 | 0.483 |
| 2495 | **Cmpk2** | Unchanged | Q3U5Q7 | 2 | 4 | 2 | 447 | 50 | 7.27 | 13 | 0.542 | 0.213 | 2.285 | 0.338 | 0.634 | 0.260 | 3.603 | 0.216 |
| 2496 | **C6** | Unchanged | E9Q6D8 | 5 | 10 | 5 | 934 | 104.6 | 6.24 | 13 | 0.557 | 0.193 | 0.801 | 0.679 | 0.601 | 0.234 | 1.334 | 0.606 |
| 2497 | **Snrpn** | Unchanged | P63163 | 3 | 24 | 3 | 240 | 24.6 | 11.19 | 13 | 1.480 | 0.336 | 1.773 | 0.332 | 1.877 | 0.130 | 0.945 | 0.912 |
| 2498 | **Krtcap2** | Unchanged | Q5RL79 | 1 | 12 | 1 | 136 | 14.7 | 9.61 | 13 | 0.210 | 0.102 | 0.427 | 0.291 | 0.693 | 0.529 | 0.616 | 0.464 |
| 2499 | **Majin** | UP | Q9D992 | 1 | 1 | 1 | 256 | 29.2 | 6.9 | 13 | 1.934 | 0.029 | 1.084 | 0.842 | 1.631 | 0.093 | 0.665 | 0.174 |
| 2500 | **Hbs1l** | Down | Q69ZS7 | 5 | 7 | 5 | 682 | 75.1 | 6.46 | 12 | 0.932 | 0.830 | 0.482 | 0.046 | 0.148 | 0.000 | 3.259 | 0.083 |
| 2501 | **Cul5** | Unchanged | Q9D5V5 | 6 | 35 | 6 | 780 | 90.9 | 7.81 | 12 | 0.870 | 0.761 | 0.849 | 0.762 | 0.398 | 0.157 | 2.134 | 0.184 |
| 2502 | **Erlin1** | Unchanged | Q91X78 | 3 | 19 | 1 | 346 | 38.9 | 7.21 | 12 | 0.462 | 0.207 | 0.761 | 0.669 | 0.249 | 0.090 | 3.060 | 0.214 |
| 2503 | **Sf3b2** | Unchanged | Q3UJB0 | 7 | 20 | 7 | 878 | 98.1 | 5.62 | 12 | 2.410 | 0.140 | 3.953 | 0.291 | 2.583 | 0.093 | 1.531 | 0.629 |
| 2504 | **Nudt8** | Unchanged | Q9CR24 | 2 | 9 | 2 | 210 | 23.2 | 6.52 | 12 | 1.078 | 0.897 | 1.727 | 0.420 | 1.290 | 0.636 | 1.339 | 0.630 |
| 2505 | **Ufd1** | Unchanged | P70362 | 3 | 23 | 3 | 307 | 34.5 | 6.7 | 12 | 2.884 | 0.331 | 5.663 | 0.122 | 0.610 | 0.403 | 9.284 | 0.097 |
| 2506 | **Sult5a1** | Unchanged | Q91X36 | 3 | 3 | 3 | 291 | 33.7 | 6.68 | 12 | 2.834 | 0.257 | 8.800 | 0.089 | 6.544 | 0.107 | 1.345 | 0.666 |
| 2507 | **Atg5** | Unchanged | Q99J83 | 2 | 7 | 2 | 275 | 32.4 | 5.91 | 12 | 0.754 | 0.631 | 4.411 | 0.200 | 4.039 | 0.088 | 1.092 | 0.899 |
| 2508 | **Hnrnpll** | Unchanged | Q921F4 | 4 | 18 | 4 | 591 | 64.1 | 5.85 | 12 | 1.347 | 0.515 | 1.194 | 0.774 | 0.766 | 0.581 | 1.558 | 0.518 |
| 2509 | **Cyp39a1** | UP | Q9JKJ9 | 3 | 14 | 3 | 470 | 53.5 | 8.84 | 12 | 0.845 | 0.701 | 2.887 | 0.109 | 1.866 | 0.047 | 1.547 | 0.347 |
| 2510 | **Osbp** | Unchanged | Q3B7Z2 | 6 | 33 | 6 | 805 | 88.7 | 7.2 | 12 | 0.783 | 0.370 | 0.873 | 0.723 | 1.017 | 0.947 | 0.858 | 0.723 |
| 2511 | **Psmg3** | Unchanged | Q9CZH3 | 1 | 2 | 1 | 122 | 13.3 | 8.66 | 12 | 0.894 | 0.903 | 0.794 | 0.832 | 0.385 | 0.469 | 2.063 | 0.484 |
| 2512 | **Ccdc22** | Down | Q9JIG7 | 4 | 14 | 4 | 627 | 70.8 | 6.01 | 12 | 0.839 | 0.444 | 2.231 | 0.188 | 0.476 | 0.006 | 4.691 | 0.072 |
| 2513 | **Ear6** | Unchanged | Q923L7 | 1 | 1 | 1 | 155 | 17.3 | 8.43 | 12 | 0.885 | 0.551 | 2.067 | 0.280 | 0.574 | 0.100 | 3.601 | 0.143 |
| 2514 | **Ctbp1** | Unchanged | O88712 | 3 | 6 | 3 | 441 | 47.7 | 6.77 | 12 | 0.528 | 0.247 | 0.694 | 0.442 | 0.322 | 0.084 | 2.157 | 0.088 |
| 2515 | **Tmem214** | Unchanged | Q8BM55 | 7 | 35 | 7 | 687 | 76.4 | 9.29 | 12 | 1.036 | 0.894 | 1.605 | 0.143 | 1.450 | 0.268 | 1.107 | 0.723 |
| 2516 | **Prpf8** | Unchanged | Q99PV0 | 17 | 96 | 17 | 2335 | 273.4 | 8.84 | 12 | 0.978 | 0.933 | 1.085 | 0.761 | 0.672 | 0.130 | 1.614 | 0.117 |
| 2517 | **Rhot1** | Unchanged | Q8BG51 | 6 | 54 | 5 | 631 | 72.2 | 6.49 | 12 | 2.265 | 0.341 | 2.809 | 0.287 | 2.183 | 0.334 | 1.287 | 0.743 |
| 2518 | **Ear2** | Unchanged | P97425 | 1 | 3 | 1 | 156 | 17.6 | 9.35 | 12 | 1.114 | 0.885 | 1.640 | 0.555 | 1.910 | 0.192 | 0.859 | 0.825 |
| 2519 | **Hm13** | Unchanged | Q9D8V0 | 4 | 38 | 4 | 378 | 41.7 | 6.04 | 12 | 0.543 | 0.182 | 0.312 | 0.054 | 0.653 | 0.457 | 0.478 | 0.348 |
| 2520 | **Mif4gd** | Down | Q3UBZ5 | 2 | 7 | 2 | 222 | 25.5 | 5.12 | 12 | 0.846 | 0.573 | 0.663 | 0.346 | 0.460 | 0.003 | 1.440 | 0.566 |
| 2521 | **Celf1** | Unchanged | P28659 | 3 | 13 | 3 | 486 | 52.1 | 8.46 | 12 | 0.486 | 0.089 | 0.914 | 0.787 | 0.531 | 0.135 | 1.721 | 0.147 |
| 2522 | **Prkaa2** | Unchanged | Q8BRK8 | 4 | 38 | 2 | 552 | 62 | 7.87 | 12 | 1.826 | 0.384 | 1.676 | 0.201 | 1.245 | 0.173 | 1.347 | 0.384 |
| 2523 | **Sept8** | Unchanged | Q8CHH9 | 3 | 39 | 1 | 429 | 49.8 | 6 | 12 | #DIV/0! | #DIV/0! | #DIV/0! | #DIV/0! | #DIV/0! | #DIV/0! | #DIV/0! | #DIV/0! |
| 2524 | **Lgals3** | UP | P16110 | 3 | 13 | 3 | 264 | 27.5 | 8.38 | 12 | 0.350 | 0.169 | 5.694 | 0.180 | 12.167 | 0.019 | 0.468 | 0.226 |
| 2525 | **Apob** | Unchanged | E9Q414 | 36 | 186 | 36 | 4505 | 509.1 | 6.81 | 12 | 0.916 | 0.548 | 1.046 | 0.890 | 0.774 | 0.123 | 1.351 | 0.413 |
| 2526 | **Klc4** | Unchanged | Q9DBS5 | 5 | 21 | 4 | 619 | 68.6 | 6.09 | 12 | 1.231 | 0.760 | 0.968 | 0.956 | 1.210 | 0.752 | 0.800 | 0.716 |
| 2527 | **Phpt1** | Unchanged | Q9DAK9 | 1 | 7 | 1 | 124 | 14 | 5.53 | 12 | 0.838 | 0.820 | 1.154 | 0.830 | 1.036 | 0.959 | 1.115 | 0.854 |
| 2528 | **Ruvbl2** | Unchanged | Q9WTM5 | 4 | 19 | 4 | 463 | 51.1 | 5.64 | 12 | 1.208 | 0.518 | 1.054 | 0.876 | 0.678 | 0.161 | 1.555 | 0.268 |
| 2529 | **Igbp1** | Unchanged | Q61249 | 3 | 38 | 3 | 340 | 38.9 | 6.18 | 12 | 1.024 | 0.927 | 1.086 | 0.754 | 1.272 | 0.589 | 0.854 | 0.735 |
| 2530 | **Mmaa** | UP | Q8C7H1 | 3 | 16 | 3 | 415 | 45.9 | 9.32 | 12 | 5.981 | 0.035 | 4.256 | 0.024 | 7.455 | 0.031 | 0.571 | 0.268 |
| 2531 | **Clptm1** | Unchanged | Q8VBZ3 | 5 | 17 | 5 | 664 | 75.2 | 6.3 | 12 | 1.109 | 0.734 | 1.058 | 0.901 | 1.056 | 0.897 | 1.003 | 0.996 |

| | | | | | | | | | | | | | | | | | | |
|---|---|---|---|---|---|---|---|---|---|---|---|---|---|---|---|---|---|---|
| 2532 | **Pdlim5** | Down | Q8CI51 | 5 | 27 | 5 | 591 | 63.3 | 8.25 | 12 | 0.617 | 0.185 | 0.515 | 0.082 | 0.426 | 0.043 | 1.210 | 0.458 |
| 2533 | **Ipo9** | Unchanged | Q91YE6 | 5 | 14 | 5 | 1041 | 116 | 4.81 | 12 | 0.880 | 0.568 | 1.764 | 0.231 | 1.212 | 0.545 | 1.455 | 0.416 |
| 2534 | **Tmem38b** | Unchanged | Q9DAV9 | 2 | 5 | 2 | 292 | 32.6 | 9.11 | 12 | 1.388 | 0.442 | 2.159 | 0.291 | 1.842 | 0.132 | 1.172 | 0.759 |
| 2535 | **Tubb1** | Unchanged | A2AQ07 | 6 | 55 | 1 | 451 | 50.4 | 5.07 | 12 | 1.188 | 0.750 | 1.305 | 0.445 | 1.030 | 0.914 | 1.267 | 0.418 |
| 2536 | **Slc39a4** | Unchanged | Q78IQ7 | 3 | 10 | 3 | 660 | 71 | 5.57 | 12 | 0.869 | 0.765 | 1.493 | 0.481 | 1.900 | 0.280 | 0.786 | 0.684 |
| 2537 | **Rnaset2a** | UP | C0HKG5 | 3 | 13 | 3 | 259 | 29.6 | 6.37 | 12 | 1.396 | 0.357 | 2.188 | 0.003 | 1.738 | 0.072 | 1.259 | 0.161 |
| 2538 | **Asrgl1** | Down | Q8C0M9 | 3 | 14 | 3 | 326 | 33.9 | 7.65 | 12 | 0.615 | 0.265 | 0.204 | 0.024 | 0.811 | 0.608 | 0.252 | 0.023 |
| 2539 | **Atg4b** | Unchanged | A0A0R4J065 | 3 | 6 | 3 | 393 | 44.4 | 5.07 | 12 | 0.963 | 0.928 | 0.900 | 0.813 | 0.567 | 0.199 | 1.588 | 0.298 |
| 2540 | **Oxsr1** | Unchanged | Q6P9R2 | 4 | 32 | 4 | 527 | 58.2 | 6.43 | 12 | 1.375 | 0.448 | 1.494 | 0.385 | 1.100 | 0.818 | 1.359 | 0.501 |
| 2541 | **Manba** | Unchanged | A0A0R4J092 | 8 | 15 | 8 | 879 | 100.8 | 7.08 | 12 | 2.297 | 0.271 | 14.280 | 0.347 | 0.523 | 0.146 | 27.299 | 0.331 |
| 2542 | **Tmem109** | Unchanged | Q3UBX0 | 2 | 3 | 2 | 243 | 26.3 | 9.89 | 12 | 1.344 | 0.471 | 1.463 | 0.463 | 0.872 | 0.777 | 1.678 | 0.270 |
| 2543 | **Ppp2r5d** | UP | Q7TNL5 | 4 | 7 | 4 | 595 | 69.1 | 7.96 | 12 | 0.812 | 0.377 | 1.711 | 0.015 | 0.982 | 0.972 | 1.742 | 0.185 |
| 2544 | **Smpdl3a** | Unchanged | P70158 | 2 | 3 | 2 | 445 | 49.8 | 6.46 | 12 | 2.066 | 0.338 | 3.859 | 0.282 | 0.750 | 0.618 | 5.147 | 0.241 |
| 2545 | **Olfr800** | Unchanged | Q8VFH7 | 1 | 1 | 1 | 311 | 35.4 | 8.54 | 12 | 1.828 | 0.120 | 1.489 | 0.391 | 5.262 | 0.091 | 0.283 | 0.132 |
| 2546 | **Tigar** | Unchanged | Q8BZA9 | 2 | 12 | 2 | 269 | 29.2 | 8.18 | 12 | 0.564 | 0.155 | 1.770 | 0.274 | 0.973 | 0.950 | 1.818 | 0.286 |
| 2547 | **Vamp8** | Unchanged | A0A0R4J0R1 | 1 | 2 | 1 | 101 | 11.4 | 8.19 | 12 | 2.613 | 0.305 | 2.109 | 0.487 | 1.629 | 0.674 | 1.294 | 0.811 |
| 2548 | **Myl10** | Unchanged | Q62082 | 1 | 1 | 1 | 202 | 22.5 | 4.98 | 12 | 1.603 | 0.573 | 1.125 | 0.882 | 2.227 | 0.290 | 0.505 | 0.342 |
| 2549 | **Vma21** | Unchanged | Q78T54 | 1 | 2 | 1 | 101 | 11.4 | 7.24 | 12 | 1.939 | 0.162 | 4.975 | 0.271 | 0.888 | 0.834 | 5.602 | 0.258 |
| 2550 | **Uap1** | UP | Q91YN5 | 4 | 21 | 4 | 522 | 58.6 | 6.49 | 12 | 4.273 | 0.153 | 3.697 | 0.015 | 1.025 | 0.950 | 3.605 | 0.014 |
| 2551 | **Xpo7** | Down | Q9EPK7 | 8 | 47 | 8 | 1087 | 123.7 | 6.38 | 12 | 0.540 | 0.132 | 0.138 | 0.002 | 0.079 | 0.001 | 1.740 | 0.455 |
| 2552 | **Tmx2** | UP | Q9D710 | 3 | 3 | 3 | 295 | 33.9 | 8.75 | 7 | 4.676 | 0.001 | 8.034 | 0.043 | 2.419 | 0.198 | 3.321 | 0.105 |
| 2553 | **Mrps23** | Unchanged | Q8VE22 | 3 | 25 | 3 | 177 | 20.3 | 8.59 | 12 | 0.590 | 0.444 | 0.630 | 0.490 | 0.347 | 0.203 | 1.816 | 0.301 |
| 2554 | **Prmt1** | Unchanged | Q9JIF0 | 3 | 12 | 3 | 371 | 42.4 | 5.43 | 12 | 0.704 | 0.327 | 0.808 | 0.560 | 0.731 | 0.303 | 1.105 | 0.737 |
| 2555 | **Pxn** | Unchanged | Q8VI36 | 3 | 15 | 3 | 591 | 64.4 | 6.02 | 12 | 1.148 | 0.641 | 1.779 | 0.197 | 1.152 | 0.674 | 1.544 | 0.339 |
| 2556 | **Hnrnpa0** | Unchanged | Q9CX86 | 2 | 4 | 2 | 305 | 30.5 | 9.31 | 12 | 1.854 | 0.144 | 2.017 | 0.151 | 2.049 | 0.142 | 0.984 | 0.965 |
| 2557 | **Cav1** | Unchanged | P49817 | 1 | 1 | 1 | 178 | 20.5 | 6.02 | 12 | 1.087 | 0.936 | 3.069 | 0.488 | 0.241 | 0.165 | 12.720 | 0.343 |
| 2558 | **Epb41iso** | Unchanged | P48193 | 8 | 38 | 3 | 858 | 95.9 | 5.6 | 12 | 0.918 | 0.831 | 1.226 | 0.737 | 0.512 | 0.185 | 2.396 | 0.265 |
| 2559 | **Dnajb1** | UP | Q9QYJ3 | 3 | 4 | 2 | 340 | 38.1 | 8.63 | 12 | 3.445 | 0.043 | 3.767 | 0.032 | 2.618 | 0.129 | 1.439 | 0.433 |
| 2560 | **Ube2r2** | Unchanged | Q6ZWZ2 | 1 | 2 | 1 | 238 | 27.1 | 4.42 | 12 | 0.928 | 0.855 | 4.495 | 0.346 | 1.128 | 0.784 | 3.986 | 0.362 |
| 2561 | **Dctpp1** | Down | Q9QY93 | 1 | 9 | 1 | 170 | 18.8 | 5.03 | 12 | 0.852 | 0.637 | 0.526 | 0.032 | 0.911 | 0.663 | 0.577 | 0.014 |
| 2562 | **Mpp6** | UP | Q9JLB0 | 5 | 18 | 5 | 553 | 62.6 | 6.4 | 12 | 0.866 | 0.590 | 2.254 | 0.001 | 0.830 | 0.465 | 2.716 | 0.000 |
| 2563 | **Rab4b** | Unchanged | Q91ZR1 | 2 | 18 | 1 | 213 | 23.6 | 6.06 | 12 | 0.118 | 0.157 | 0.420 | 0.360 | 0.589 | 0.509 | 0.712 | 0.555 |
| 2564 | **Mms19** | Unchanged | Q9D071 | 6 | 27 | 6 | 1031 | 113 | 6.21 | 12 | 1.284 | 0.369 | 2.029 | 0.059 | 0.971 | 0.848 | 2.089 | 0.058 |
| 2565 | **Mrps14** | UP | Q9CR88 | 1 | 3 | 1 | 128 | 14.9 | 11.39 | 12 | 6.488 | 0.051 | 7.943 | 0.156 | 11.929 | 0.002 | 0.666 | 0.448 |
| 2566 | **Bccip** | Down | Q9CWI3 | 2 | 7 | 2 | 316 | 35.9 | 4.45 | 12 | 0.698 | 0.448 | 0.856 | 0.773 | 0.337 | 0.029 | 2.541 | 0.248 |
| 2567 | **Smarcb1** | Unchanged | Q9Z0H3 | 2 | 4 | 2 | 385 | 44.1 | 6.23 | 12 | 1.290 | 0.732 | 0.411 | 0.339 | 1.974 | 0.331 | 0.208 | 0.083 |
| 2568 | **Lin7c** | Unchanged | O88952 | 2 | 9 | 1 | 197 | 21.8 | 8.43 | 12 | 1.274 | 0.646 | 1.029 | 0.969 | 0.220 | 0.200 | 4.669 | 0.148 |
| 2569 | **Cd82** | Unchanged | P40237 | 3 | 13 | 3 | 266 | 29.6 | 5.02 | 12 | 0.761 | 0.715 | 0.308 | 0.180 | 0.075 | 0.067 | 4.077 | 0.235 |
| 2570 | **Yes1** | Unchanged | Q04736 | 4 | 12 | 3 | 541 | 60.6 | 6.64 | 12 | 0.848 | 0.462 | 10.314 | 0.295 | 1.056 | 0.786 | 9.771 | 0.298 |
| 2571 | **Nsun2** | Unchanged | Q1HFZ0 | 7 | 26 | 7 | 757 | 85.4 | 6.58 | 12 | 0.475 | 0.271 | 0.447 | 0.252 | 0.552 | 0.363 | 0.810 | 0.728 |
| 2572 | **Psat1** | Unchanged | Q99K85 | 4 | 7 | 4 | 370 | 40.4 | 8.03 | 12 | 1.304 | 0.344 | 1.872 | 0.153 | 2.281 | 0.084 | 0.821 | 0.640 |

| | | | | | | | | | | | | | | | | | | |
|---|---|---|---|---|---|---|---|---|---|---|---|---|---|---|---|---|---|---|
| 2573 | **Atp6v0c** | Unchanged | P63082 | 1 | 1 | 1 | 155 | 15.8 | 9.13 | 12 | 1.356 | 0.543 | 1.804 | 0.270 | 1.419 | 0.372 | 1.271 | 0.620 |
| 2574 | **Lss** | Unchanged | Q8BLN5 | 6 | 24 | 6 | 733 | 83.1 | 6.4 | 12 | 0.802 | 0.685 | 0.960 | 0.945 | 0.674 | 0.463 | 1.426 | 0.509 |
| 2575 | **Scarb1** | Unchanged | Q61009 | 2 | 8 | 2 | 509 | 56.7 | 8.09 | 12 | 3.782 | 0.340 | 12.802 | 0.143 | 7.448 | 0.177 | 1.719 | 0.545 |
| 2576 | **Defa20** | Unchanged | Q45VN2 | 1 | 2 | 1 | 95 | 10.6 | 5.81 | 12 | #DIV/0! | #DIV/0! | #DIV/0! | #DIV/0! | #DIV/0! | #DIV/0! | #DIV/0! | #DIV/0! |
| 2577 | **Slc22a18** | Unchanged | Q78KK3 | 3 | 12 | 3 | 406 | 43 | 9.07 | 12 | 1.007 | 0.989 | 1.448 | 0.540 | 1.787 | 0.427 | 0.810 | 0.756 |
| 2578 | **Abcb9** | Unchanged | D6RFD9 | 2 | 3 | 1 | 268 | 29.5 | 7.78 | 12 | 3.837 | 0.336 | 0.501 | 0.440 | 1.670 | 0.438 | 0.300 | 0.071 |
| 2579 | **Nubp1** | Unchanged | Q9R060 | 2 | 2 | 2 | 320 | 34.1 | 5.9 | 12 | 5.563 | 0.113 | 33.876 | 0.337 | 7.069 | 0.177 | 4.792 | 0.433 |
| 2580 | **Mrpl47** | Unchanged | Q8K2Y7 | 3 | 9 | 3 | 252 | 29.7 | 10.21 | 12 | 5.568 | 0.381 | 0.378 | 0.166 | 0.431 | 0.217 | 0.877 | 0.832 |
| 2581 | **Prss38** | Unchanged | Q3UKY7 | 1 | 1 | 1 | 322 | 35.7 | 7.84 | 11 | 1.485 | 0.573 | 1.164 | 0.833 | 1.980 | 0.449 | 0.588 | 0.547 |
| 2582 | **Rnase4** | Unchanged | Q8C7E4 | 1 | 3 | 1 | 148 | 17 | 8.85 | 11 | #DIV/0! | #DIV/0! | #DIV/0! | #DIV/0! | #DIV/0! | #DIV/0! | #DIV/0! | #DIV/0! |
| 2583 | **Eea1** | Unchanged | Q8BL66 | 11 | 44 | 11 | 1411 | 160.8 | 5.77 | 11 | 0.765 | 0.448 | 0.834 | 0.563 | 0.968 | 0.918 | 0.861 | 0.400 |
| 2584 | **Dnajc8** | UP | Q6NZB0 | 1 | 1 | 1 | 253 | 29.8 | 9.06 | 11 | 0.635 | 0.188 | 7.331 | 0.025 | 3.863 | 0.104 | 1.898 | 0.245 |
| 2585 | **Man1a1** | Unchanged | P45700 | 6 | 24 | 6 | 655 | 73.2 | 6.81 | 11 | 1.426 | 0.265 | 1.554 | 0.443 | 1.116 | 0.653 | 1.392 | 0.533 |
| 2586 | **Hmgb2** | Unchanged | P30681 | 2 | 7 | 1 | 210 | 24.1 | 7.31 | 11 | #DIV/0! | #DIV/0! | #DIV/0! | #DIV/0! | #DIV/0! | #DIV/0! | #DIV/0! | #DIV/0! |
| 2587 | **Cst3** | Unchanged | P21460 | 1 | 4 | 1 | 140 | 15.5 | 9 | 11 | 0.534 | 0.456 | 0.152 | 0.056 | 1.145 | 0.805 | 0.133 | 0.047 |
| 2588 | **Pla2g15** | Unchanged | Q8VEB4 | 3 | 29 | 3 | 412 | 47.3 | 6.47 | 11 | 1.122 | 0.815 | 1.836 | 0.329 | 1.189 | 0.676 | 1.544 | 0.427 |
| 2589 | **Dusp13** | Unchanged | Z4YLI2 | 1 | 2 | 1 | 316 | 35.4 | 9.74 | 11 | 1.683 | 0.081 | 0.835 | 0.678 | 1.259 | 0.541 | 0.663 | 0.288 |
| 2590 | **Armt1** | Unchanged | A6H630 | 2 | 10 | 2 | 439 | 50.5 | 5.92 | 11 | 0.624 | 0.460 | 1.536 | 0.502 | 1.368 | 0.500 | 1.123 | 0.811 |
| 2591 | **C5** | Unchanged | P06684 | 13 | 45 | 13 | 1680 | 188.8 | 6.81 | 11 | 0.772 | 0.389 | 1.306 | 0.200 | 0.756 | 0.261 | 1.727 | 0.000 |
| 2592 | **Cdh2** | Unchanged | P15116 | 5 | 31 | 5 | 906 | 99.7 | 4.78 | 11 | 0.918 | 0.828 | 0.882 | 0.790 | 0.892 | 0.798 | 0.988 | 0.977 |
| 2593 | **Mapk9** | Unchanged | Q9WTU6 | 3 | 11 | 3 | 423 | 48.2 | 5.97 | 11 | 0.487 | 0.273 | 5.037 | 0.212 | 0.940 | 0.947 | 5.361 | 0.216 |
| 2594 | **Mrps33** | Unchanged | Q9D2R8 | 1 | 6 | 1 | 106 | 12.5 | 10.26 | 11 | 1.506 | 0.389 | 3.147 | 0.169 | 0.497 | 0.281 | 6.332 | 0.094 |
| 2595 | **Adrm1** | Unchanged | Q9JKV1 | 5 | 43 | 5 | 407 | 42 | 5.07 | 11 | 1.401 | 0.222 | 1.866 | 0.222 | 0.862 | 0.652 | 2.164 | 0.163 |
| 2596 | **Esrp2** | Unchanged | Q8K0G8 | 4 | 8 | 4 | 717 | 77.3 | 6.38 | 11 | 0.873 | 0.703 | 0.860 | 0.682 | 0.652 | 0.297 | 1.319 | 0.289 |
| 2597 | **Ppie** | Unchanged | Q9QZH3 | 2 | 8 | 1 | 301 | 33.4 | 5.6 | 11 | 1.258 | 0.515 | 2.055 | 0.262 | 0.931 | 0.825 | 2.206 | 0.222 |
| 2598 | **Adpgk** | Unchanged | Q8VDL4 | 3 | 11 | 3 | 496 | 53.9 | 5.62 | 11 | 1.378 | 0.375 | 2.472 | 0.170 | 1.138 | 0.761 | 2.171 | 0.208 |
| 2599 | **Nlrx1** | UP | Q3TL44 | 4 | 19 | 4 | 975 | 107.8 | 7.37 | 11 | 1.946 | 0.465 | 0.486 | 0.423 | 3.812 | 0.039 | 0.128 | 0.014 |
| 2600 | **Sdr39u1** | Unchanged | Q5M8N4 | 2 | 7 | 2 | 293 | 31.4 | 9.31 | 11 | 1.043 | 0.918 | 1.428 | 0.435 | 1.069 | 0.860 | 1.336 | 0.520 |
| 2601 | **Neu1** | Unchanged | O35657 | 2 | 2 | 2 | 409 | 44.6 | 6.02 | 11 | 1.086 | 0.763 | 1.058 | 0.822 | 2.242 | 0.127 | 0.472 | 0.156 |
| 2602 | **Acbd3** | Unchanged | Q8BMP6 | 3 | 7 | 3 | 525 | 60.1 | 5.11 | 11 | 0.933 | 0.901 | 2.298 | 0.341 | 3.381 | 0.143 | 0.680 | 0.573 |
| 2603 | **Uqcc3** | Unchanged | Q8K2T4 | 1 | 1 | 1 | 89 | 9.6 | 9.52 | 11 | 1.511 | 0.511 | 1.467 | 0.625 | 0.502 | 0.289 | 2.923 | 0.297 |
| 2604 | **Cnpy3** | Unchanged | Q9DAU1 | 2 | 22 | 2 | 276 | 30.5 | 5.62 | 11 | 2.581 | 0.081 | 1.374 | 0.659 | 1.855 | 0.227 | 0.741 | 0.640 |
| 2605 | **Mrps18a** | Unchanged | Q99N85 | 1 | 2 | 1 | 196 | 22.3 | 10.18 | 11 | 2.935 | 0.114 | 6.421 | 0.143 | 1.802 | 0.135 | 3.562 | 0.204 |
| 2606 | **Cavin1** | Unchanged | O54724 | 3 | 9 | 3 | 392 | 43.9 | 5.52 | 11 | 1.083 | 0.915 | 2.083 | 0.526 | 0.983 | 0.979 | 2.119 | 0.502 |
| 2607 | **Olfr481** | Unchanged | Q8VGI5 | 1 | 1 | 1 | 312 | 34.3 | 7.58 | 11 | 1.112 | 0.891 | 0.681 | 0.630 | 0.804 | 0.748 | 0.848 | 0.641 |
| 2608 | **Abhd3** | Unchanged | Q91ZH7 | 2 | 11 | 2 | 411 | 46.2 | 7.49 | 11 | 0.989 | 0.984 | 2.106 | 0.237 | 1.102 | 0.840 | 1.912 | 0.290 |
| 2609 | **Fads1** | Unchanged | Q920L1 | 3 | 12 | 3 | 447 | 52.3 | 9.29 | 8 | 1.833 | 0.357 | 3.448 | 0.134 | 1.296 | 0.647 | 2.660 | 0.202 |
| 2610 | **Commd1** | UP | Q8K4M5 | 1 | 5 | 1 | 188 | 21 | 7.59 | 11 | 1.751 | 0.374 | 2.846 | 0.032 | 3.788 | 0.010 | 0.751 | 0.395 |
| 2611 | **Snrnp70** | Unchanged | Q62376 | 4 | 16 | 4 | 448 | 52 | 9.94 | 11 | 0.847 | 0.710 | 3.324 | 0.317 | 2.269 | 0.184 | 1.465 | 0.659 |
| 2612 | **Chmp4b** | UP | Q9D8B3 | 2 | 3 | 2 | 224 | 24.9 | 4.82 | 11 | 26.873 | 0.047 | 9.390 | 0.156 | 132.237 | 0.222 | 0.071 | 0.251 |
| 2613 | **Tor1aip2** | Unchanged | Q8BYU6 | 4 | 21 | 4 | 502 | 54.5 | 4.86 | 11 | 0.591 | 0.080 | 0.727 | 0.427 | 0.850 | 0.664 | 0.855 | 0.778 |

| | | | | | | | | | | | | | | | | | | |
|---|---|---|---|---|---|---|---|---|---|---|---|---|---|---|---|---|---|---|
| 2614 | **Uba2** | Unchanged | Q9Z1F9 | 3 | 13 | 3 | 638 | 70.5 | 5.24 | 11 | 2.333 | 0.330 | 1.993 | 0.331 | 1.652 | 0.438 | 1.206 | 0.730 |
| 2615 | **Gnas** | Down | Q6R0H7 | 8 | 54 | 8 | 1133 | 121.4 | 4.81 | 11 | 0.512 | 0.037 | 0.657 | 0.243 | 0.313 | 0.001 | 2.100 | 0.202 |
| 2616 | **Mrpl24** | UP | Q9CQ06 | 1 | 8 | 1 | 216 | 24.9 | 9.5 | 11 | 3.679 | 0.024 | 1.517 | 0.567 | 1.980 | 0.232 | 0.766 | 0.611 |
| 2617 | **Bgn** | Unchanged | P28653 | 3 | 21 | 3 | 369 | 41.6 | 7.27 | 11 | 4.394 | 0.239 | 6.698 | 0.107 | 3.845 | 0.185 | 1.742 | 0.454 |
| 2618 | **Mrpl41** | Unchanged | Q9CQN7 | 1 | 2 | 1 | 135 | 15.3 | 9.82 | 11 | 0.995 | 0.993 | 0.482 | 0.277 | 0.414 | 0.203 | 1.164 | 0.824 |
| 2619 | **Babam1** | Unchanged | Q3U143 | 1 | 1 | 1 | 333 | 36.8 | 4.55 | 11 | #DIV/0! | #DIV/0! | #DIV/0! | #DIV/0! | #DIV/0! | #DIV/0! | #DIV/0! | #DIV/0! |
| 2620 | **Ctnna2** | Unchanged | E0CXB9 | 7 | 23 | 3 | 966 | 106.7 | 5.69 | 11 | 1.442 | 0.609 | 2.611 | 0.275 | 1.292 | 0.740 | 2.021 | 0.377 |
| 2621 | **Ppp4c** | Unchanged | P97470 | 2 | 3 | 1 | 307 | 35.1 | 5.06 | 11 | 0.778 | 0.658 | 0.540 | 0.266 | 0.830 | 0.723 | 0.651 | 0.436 |
| 2622 | **Cd9** | UP | P40240 | 1 | 4 | 1 | 226 | 25.2 | 7.23 | 11 | 2.416 | 0.008 | 2.142 | 0.029 | 7.497 | 0.128 | 0.286 | 0.198 |
| 2623 | **Usp7** | UP | Q6A4J8 | 7 | 19 | 7 | 1103 | 128.4 | 5.5 | 11 | 1.797 | 0.039 | 3.235 | 0.146 | 0.976 | 0.910 | 3.314 | 0.143 |
| 2624 | **Olfr128** | Unchanged | Q7TRJ1 | 1 | 1 | 1 | 308 | 34.3 | 8.22 | 11 | 0.804 | 0.787 | 0.158 | 0.224 | 0.435 | 0.410 | 0.364 | 0.282 |
| 2625 | **Cd36** | Unchanged | Q08857 | 3 | 23 | 3 | 472 | 52.7 | 8.35 | 11 | 0.684 | 0.283 | 1.024 | 0.953 | 0.390 | 0.083 | 2.628 | 0.112 |
| 2626 | **Pgpep1** | Unchanged | Q9ESW8 | 1 | 2 | 1 | 209 | 22.9 | 5.39 | 11 | 1.016 | 0.983 | 1.618 | 0.564 | 2.363 | 0.276 | 0.685 | 0.588 |
| 2627 | **Sars2** | Unchanged | Q9JJL8 | 3 | 7 | 3 | 518 | 58.3 | 7.9 | 11 | 0.991 | 0.972 | 1.351 | 0.343 | 1.656 | 0.328 | 0.815 | 0.648 |
| 2628 | **Brcc3** | Unchanged | P46737 | 2 | 12 | 2 | 291 | 33.3 | 5.83 | 11 | 0.204 | 0.265 | 0.778 | 0.747 | 0.247 | 0.292 | 3.146 | 0.001 |
| 2629 | **Aifm2** | Unchanged | Q8BUE4 | 2 | 4 | 2 | 373 | 40.6 | 8.98 | 11 | 0.993 | 0.987 | 1.492 | 0.395 | 2.009 | 0.193 | 0.742 | 0.539 |
| 2630 | **H2-Aa** | Unchanged | P14434 | 1 | 1 | 1 | 256 | 28.1 | 4.78 | 11 | #DIV/0! | #DIV/0! | #DIV/0! | #DIV/0! | #DIV/0! | #DIV/0! | #DIV/0! | #DIV/0! |
| 2631 | **Rbm8a** | Unchanged | Q9CWZ3 | 1 | 1 | 1 | 174 | 19.9 | 5.72 | 11 | #DIV/0! | #DIV/0! | #DIV/0! | #DIV/0! | #DIV/0! | #DIV/0! | #DIV/0! | #DIV/0! |
| 2632 | **Sumo3** | Unchanged | Q9Z172 | 1 | 4 | 1 | 110 | 12.4 | 5.96 | 11 | 0.058 | 0.072 | 0.050 | 0.070 | 0.022 | 0.063 | 2.201 | 0.484 |
| 2633 | **Plbd1** | Unchanged | Q8VCI0 | 4 | 8 | 4 | 550 | 63 | 8.87 | 11 | 1.552 | 0.597 | 1.929 | 0.432 | 1.360 | 0.630 | 1.419 | 0.662 |
| 2634 | **Usp9xiso** | Unchanged | P70398 | 16 | 88 | 14 | 2559 | 290.5 | 5.87 | 11 | 1.231 | 0.551 | 1.432 | 0.446 | 1.769 | 0.200 | 0.810 | 0.641 |
| 2635 | **Pigr** | Unchanged | O70570 | 8 | 36 | 8 | 771 | 84.9 | 5.4 | 11 | 1.055 | 0.804 | 0.958 | 0.837 | 1.234 | 0.230 | 0.776 | 0.297 |
| 2636 | **Zpr1** | Unchanged | Q62384 | 3 | 7 | 3 | 459 | 50.7 | 4.78 | 11 | 12.516 | 0.126 | 7.092 | 0.098 | 8.049 | 0.059 | 0.881 | 0.839 |
| 2637 | **Trim25** | Unchanged | Q61510 | 4 | 13 | 4 | 634 | 71.7 | 8.28 | 11 | 0.921 | 0.781 | 1.007 | 0.989 | 1.109 | 0.671 | 0.908 | 0.833 |
| 2638 | **Ppme1** | Unchanged | Q8BVQ5 | 2 | 5 | 2 | 386 | 42.2 | 5.97 | 11 | 0.144 | 0.343 | 0.230 | 0.393 | 0.363 | 0.493 | 0.633 | 0.641 |
| 2639 | **Grsf1** | UP | Q8C5Q4 | 3 | 8 | 3 | 479 | 53 | 6.67 | 11 | 6.408 | 0.146 | 4.124 | 0.003 | 5.148 | 0.029 | 0.801 | 0.525 |
| 2640 | **Sh3bgrl2** | Unchanged | A0A087WS96 | 1 | 1 | 1 | 120 | 13.5 | 8.43 | 11 | 1.031 | 0.960 | 0.708 | 0.673 | 1.302 | 0.602 | 0.543 | 0.380 |
| 2641 | **9430069I07Rik** | Unchanged | Q9CX21 | 1 | 4 | 1 | 111 | 12.7 | 9.83 | 11 | 1.161 | 0.821 | 0.793 | 0.735 | 1.560 | 0.531 | 0.508 | 0.391 |
| 2642 | **Atad2b** | Unchanged | A0A1W2P7J4 | 1 | 1 | 1 | 259 | 28.8 | 7.61 | 11 | 0.824 | 0.580 | 0.856 | 0.718 | 0.730 | 0.371 | 1.172 | 0.704 |
| 2643 | **Selenot** | Unchanged | P62342 | 1 | 1 | 1 | 195 | 22.3 | 8.6 | 11 | #DIV/0! | #DIV/0! | #DIV/0! | #DIV/0! | #DIV/0! | #DIV/0! | #DIV/0! | #DIV/0! |
| 2644 | **Commd3** | UP | Q63829 | 1 | 8 | 1 | 195 | 22 | 5.59 | 11 | 1.676 | 0.169 | 3.357 | 0.034 | 0.572 | 0.267 | 5.868 | 0.014 |
| 2645 | **Gm5571** | Unchanged | P01639 | 1 | 2 | 1 | 130 | 14.3 | 5.48 | 11 | 0.492 | 0.361 | 2.044 | 0.605 | 2.990 | 0.485 | 0.684 | 0.780 |
| 2646 | **Clmn** | UP | Q8C5W0 | 6 | 23 | 6 | 1052 | 117.2 | 5.03 | 11 | 1.804 | 0.031 | 1.417 | 0.203 | 1.480 | 0.309 | 0.957 | 0.903 |
| 2647 | **Vmn1r224** | Unchanged | E9PWK2 | 1 | 1 | 1 | 298 | 34.1 | 9.72 | 11 | 2.201 | 0.276 | 3.326 | 0.103 | 0.605 | 0.204 | 5.497 | 0.065 |
| 2648 | **Stx7** | Unchanged | O70439 | 2 | 7 | 2 | 261 | 29.8 | 5.78 | 11 | 0.975 | 0.953 | 1.378 | 0.626 | 1.182 | 0.672 | 1.166 | 0.816 |
| 2649 | **Grem2** | Unchanged | O88273 | 1 | 1 | 1 | 168 | 19.3 | 9.17 | 11 | 1.717 | 0.503 | 0.512 | 0.469 | 1.650 | 0.519 | 0.310 | 0.257 |
| 2650 | **Nucb2** | Unchanged | P81117 | 3 | 8 | 3 | 420 | 50.3 | 5.15 | 11 | 0.709 | 0.593 | 0.917 | 0.863 | 0.451 | 0.226 | 2.035 | 0.188 |
| 2651 | **Psmf1** | Unchanged | Q8BHL8 | 2 | 11 | 2 | 271 | 29.6 | 5.25 | 11 | 1.046 | 0.900 | 0.792 | 0.561 | 1.039 | 0.927 | 0.762 | 0.551 |
| 2652 | **Mycbp** | Unchanged | Q9EQS3 | 1 | 1 | 1 | 103 | 12 | 5.91 | 11 | #DIV/0! | #DIV/0! | #DIV/0! | #DIV/0! | #DIV/0! | #DIV/0! | #DIV/0! | #DIV/0! |
| 2653 | **Abcb4** | Unchanged | P21440 | 8 | 18 | 8 | 1276 | 140.3 | 8.78 | 11 | 1.576 | 0.446 | 3.039 | 0.118 | 0.380 | 0.070 | 7.989 | 0.046 |
| 2654 | **Hdac2** | Unchanged | P70288 | 4 | 13 | 2 | 488 | 55.3 | 5.91 | 11 | 1.107 | 0.878 | 2.906 | 0.367 | 0.337 | 0.229 | 8.630 | 0.220 |

| | | | | | | | | | | | | | | | | | | |
|---|---|---|---|---|---|---|---|---|---|---|---|---|---|---|---|---|---|---|
| 2655 | **Maip1** | Unchanged | Q8BHE8 | 3 | 5 | 3 | 291 | 33 | 9.14 | 8 | 2.477 | 0.258 | 0.049 | 0.066 | 0.077 | 0.073 | 0.629 | 0.540 |
| 2656 | **Tbrg4** | Unchanged | Q91YM4 | 4 | 12 | 4 | 630 | 71.5 | 8.34 | 11 | 1.445 | 0.267 | 1.826 | 0.057 | 1.154 | 0.706 | 1.583 | 0.056 |
| 2657 | **Sqstm1** | Unchanged | Q64337 | 2 | 2 | 2 | 442 | 48.1 | 5.21 | 11 | #DIV/0! | #DIV/0! | #DIV/0! | #DIV/0! | #DIV/0! | #DIV/0! | #DIV/0! | #DIV/0! |
| 2658 | **Gm34653** | Unchanged | A0A1L1SR01 | 1 | 1 | 1 | 368 | 41.7 | 6.05 | 11 | #DIV/0! | #DIV/0! | #DIV/0! | #DIV/0! | #DIV/0! | #DIV/0! | #DIV/0! | #DIV/0! |
| 2659 | **Ctss** | Unchanged | O70370 | 2 | 13 | 2 | 340 | 38.4 | 6.96 | 11 | 0.530 | 0.208 | 2.247 | 0.415 | 0.767 | 0.520 | 2.930 | 0.332 |
| 2660 | **Ube2g1** | Unchanged | P62254 | 1 | 2 | 1 | 170 | 19.5 | 5.3 | 11 | #DIV/0! | #DIV/0! | #DIV/0! | #DIV/0! | #DIV/0! | #DIV/0! | #DIV/0! | #DIV/0! |
| 2661 | **Ppp3r1** | UP | Q63810 | 1 | 1 | 1 | 170 | 19.3 | 4.81 | 11 | 2.300 | 0.018 | 2.675 | 0.142 | 1.971 | 0.227 | 1.357 | 0.569 |
| 2662 | **Reep5** | UP | G3X8R0 | 2 | 3 | 2 | 189 | 21.4 | 8.12 | 11 | 1.846 | 0.045 | 0.790 | 0.598 | 3.592 | 0.093 | 0.220 | 0.070 |
| 2663 | **Ndufb3** | Down | Q9CQZ6 | 1 | 6 | 1 | 104 | 11.7 | 9.04 | 11 | 0.215 | 0.002 | 1.242 | 0.596 | 0.106 | 0.001 | 11.762 | 0.024 |
| 2664 | **Tm9sf4** | Unchanged | Q8BH24 | 5 | 17 | 5 | 643 | 74.6 | 7.23 | 11 | 0.893 | 0.836 | 1.238 | 0.750 | 0.873 | 0.808 | 1.417 | 0.626 |
| 2665 | **Exoc4** | Unchanged | O35382 | 5 | 12 | 5 | 975 | 110.5 | 6.49 | 11 | 1.266 | 0.225 | 1.763 | 0.080 | 1.399 | 0.142 | 1.260 | 0.398 |
| 2666 | **Cab39** | Unchanged | Q06138 | 3 | 19 | 3 | 341 | 39.8 | 6.89 | 11 | 1.420 | 0.489 | 4.145 | 0.159 | 1.468 | 0.392 | 2.824 | 0.225 |
| 2667 | **Selenop** | Unchanged | P70274 | 2 | 10 | 2 | 380 | 42.7 | 7.09 | 11 | 2.769 | 0.186 | 4.211 | 0.092 | 2.850 | 0.151 | 1.477 | 0.493 |
| 2668 | **Fabp3** | Unchanged | P11404 | 1 | 6 | 1 | 133 | 14.8 | 6.57 | 11 | 1.347 | 0.814 | 0.129 | 0.314 | 0.024 | 0.260 | 5.387 | 0.303 |
| 2669 | **C8b** | Down | Q8BH35 | 4 | 9 | 4 | 589 | 66.2 | 7.77 | 11 | 0.680 | 0.312 | 0.482 | 0.061 | 0.424 | 0.013 | 1.137 | 0.792 |
| 2670 | **Tmem273** | Unchanged | E9PVZ2 | 1 | 1 | 1 | 95 | 10.2 | 8.79 | 11 | 0.380 | 0.306 | 0.049 | 0.115 | 0.310 | 0.245 | 0.158 | 0.080 |
| 2671 | **Gas2** | Unchanged | P11862 | 2 | 8 | 2 | 314 | 34.9 | 8.57 | 11 | 0.359 | 0.235 | 0.441 | 0.276 | 0.838 | 0.770 | 0.527 | 0.173 |
| 2672 | **Slc22a28** | Unchanged | B2RT89 | 5 | 30 | 5 | 552 | 62.1 | 8.24 | 11 | 0.972 | 0.932 | 0.798 | 0.566 | 0.653 | 0.364 | 1.221 | 0.687 |
| 2673 | **Lcn2** | Unchanged | P11672 | 1 | 2 | 1 | 200 | 22.9 | 8.81 | 11 | 2.816 | 0.128 | 0.708 | 0.630 | 0.301 | 0.217 | 2.351 | 0.342 |
| 2674 | **Ddrgk1** | Unchanged | Q80WW9 | 3 | 15 | 3 | 315 | 36 | 5.35 | 10 | 2.831 | 0.243 | 3.645 | 0.055 | 1.610 | 0.296 | 2.264 | 0.136 |
| 2675 | **Mrps11** | Unchanged | Q9DCA2 | 1 | 4 | 1 | 191 | 20.2 | 10.77 | 10 | 0.749 | 0.758 | 1.638 | 0.496 | 0.341 | 0.360 | 4.807 | 0.071 |
| 2676 | **Sap18** | Unchanged | O55128 | 1 | 2 | 1 | 153 | 17.6 | 9.35 | 10 | 0.713 | 0.469 | 5.803 | 0.338 | 0.684 | 0.414 | 8.486 | 0.309 |
| 2677 | **Ldlr** | Unchanged | P35951 | 5 | 15 | 5 | 862 | 94.9 | 5.02 | 10 | 1.115 | 0.697 | 0.940 | 0.823 | 1.102 | 0.825 | 0.852 | 0.742 |
| 2678 | **Mid1ip1** | Down | Q9CQ20 | 1 | 2 | 1 | 182 | 20.3 | 5.5 | 10 | 0.156 | 0.049 | 1.590 | 0.711 | 0.645 | 0.437 | 2.464 | 0.551 |
| 2679 | **Tmem126b** | Unchanged | Q9D1R1 | 2 | 6 | 2 | 230 | 25.4 | 9.51 | 10 | 1.066 | 0.938 | 1.756 | 0.303 | 1.382 | 0.636 | 1.271 | 0.703 |
| 2680 | **Tubg2** | Unchanged | Q8VCK3 | 2 | 4 | 2 | 451 | 51.1 | 6.21 | 10 | 1.775 | 0.504 | 4.406 | 0.173 | 0.710 | 0.447 | 6.204 | 0.142 |
| 2681 | **Cd1d1** | Unchanged | P11609 | 2 | 5 | 2 | 336 | 38.5 | 7.14 | 10 | 1.855 | 0.278 | 6.908 | 0.154 | 1.551 | 0.108 | 4.454 | 0.190 |
| 2682 | **Bag3** | Unchanged | Q9JLV1 | 6 | 23 | 6 | 577 | 61.8 | 7.27 | 10 | 1.079 | 0.897 | 1.298 | 0.671 | 1.211 | 0.687 | 1.072 | 0.865 |
| 2683 | **Kpna2** | Unchanged | P52293 | 3 | 13 | 3 | 529 | 57.9 | 5.68 | 10 | 2.060 | 0.396 | 0.875 | 0.797 | 1.152 | 0.852 | 0.760 | 0.725 |
| 2684 | **Ncald** | Unchanged | Q91X97 | 2 | 3 | 2 | 193 | 22.2 | 5.35 | 10 | 1.016 | 0.974 | 2.323 | 0.256 | 1.344 | 0.392 | 1.728 | 0.396 |
| 2685 | **Cds2** | Unchanged | Q99L43 | 3 | 39 | 3 | 444 | 51.3 | 7.05 | 10 | 0.926 | 0.822 | 2.080 | 0.293 | 1.082 | 0.858 | 1.922 | 0.357 |
| 2686 | **Ankrd7** | Unchanged | A0A0R4J1L1 | 1 | 1 | 1 | 280 | 31.3 | 9.07 | 10 | 0.255 | 0.349 | 0.828 | 0.851 | 0.709 | 0.735 | 1.167 | 0.850 |
| 2687 | **Krt5** | Unchanged | Q922U2 | 7 | 92 | 1 | 580 | 61.7 | 7.75 | 10 | 5.421 | 0.084 | 2.726 | 0.083 | 1.776 | 0.198 | 1.536 | 0.367 |
| 2688 | **Atp13a1** | Unchanged | Q9EPE9 | 7 | 24 | 7 | 1200 | 132.3 | 8.03 | 10 | 1.156 | 0.799 | 2.528 | 0.181 | 0.948 | 0.922 | 2.665 | 0.140 |
| 2689 | **F11r** | Down | O88792 | 2 | 2 | 2 | 300 | 32.4 | 6.77 | 10 | 0.048 | 0.022 | 1.065 | 0.954 | 0.626 | 0.462 | 1.702 | 0.697 |
| 2690 | **Mrps2** | Unchanged | Q924T2 | 2 | 11 | 2 | 291 | 32.3 | 9.14 | 10 | 1.726 | 0.403 | 3.488 | 0.163 | 3.822 | 0.154 | 0.913 | 0.893 |
| 2691 | **Atxn7l3b** | Unchanged | Q3UD01 | 1 | 2 | 1 | 97 | 10.7 | 4.32 | 10 | 0.210 | 0.077 | 0.435 | 0.248 | 0.625 | 0.374 | 0.697 | 0.582 |
| 2692 | **Epb41isob** | Unchanged | A0A1D5RLV1 | 4 | 15 | 1 | 476 | 54 | 9.09 | 10 | 0.962 | 0.808 | 1.522 | 0.635 | 0.922 | 0.866 | 1.651 | 0.613 |
| 2693 | **Tprkb** | Unchanged | Q8QZZ7 | 2 | 16 | 2 | 175 | 19.5 | 7.39 | 10 | 2.069 | 0.339 | 4.841 | 0.151 | 3.008 | 0.234 | 1.609 | 0.538 |
| 2694 | **Cdk5** | Unchanged | P49615 | 2 | 8 | 2 | 292 | 33.3 | 7.66 | 10 | 1.558 | 0.501 | 1.496 | 0.554 | 0.446 | 0.172 | 3.353 | 0.179 |
| 2695 | **Tmem176b** | Unchanged | Q9R1Q6 | 2 | 3 | 2 | 263 | 28.3 | 7.84 | 10 | 1.079 | 0.863 | 1.825 | 0.127 | 0.612 | 0.390 | 2.984 | 0.008 |

| | | | | | | | | | | | | | | | | | | |
|---|---|---|---|---|---|---|---|---|---|---|---|---|---|---|---|---|---|---|
| 2696 | **Napsa** | Unchanged | O09043 | 1 | 9 | 1 | 419 | 45.5 | 7.5 | 10 | 1.085 | 0.907 | 0.720 | 0.701 | 1.100 | 0.874 | 0.654 | 0.495 |
| 2697 | **Tbcb** | Unchanged | Q9D1E6 | 1 | 2 | 1 | 244 | 27.4 | 5.24 | 10 | 0.881 | 0.750 | 0.578 | 0.139 | 0.479 | 0.067 | 1.209 | 0.586 |
| 2698 | **Fcgr2b** | Unchanged | A0A0B4J1G1 | 2 | 4 | 2 | 293 | 33 | 7.23 | 10 | 1.103 | 0.868 | 0.466 | 0.271 | 1.036 | 0.958 | 0.450 | 0.319 |
| 2699 | **Prkra** | Unchanged | Q9WTX2 | 2 | 4 | 2 | 313 | 34.3 | 8.43 | 10 | 0.383 | 0.201 | 0.585 | 0.459 | 0.477 | 0.277 | 1.226 | 0.744 |
| 2700 | **Msi1** | Unchanged | Q61474 | 1 | 1 | 1 | 362 | 39.1 | 7.87 | 10 | 1.082 | 0.840 | 0.749 | 0.592 | 7.768 | 0.073 | 0.096 | 0.066 |
| 2701 | **C2cd2** | Unchanged | E9Q3C1 | 4 | 6 | 4 | 696 | 76.5 | 7.72 | 10 | 1.103 | 0.852 | 1.676 | 0.261 | 0.363 | 0.160 | 4.615 | 0.009 |
| 2702 | **Mrpl9** | Unchanged | Q99N94 | 2 | 4 | 2 | 265 | 30.2 | 10.08 | 10 | 0.545 | 0.402 | 0.497 | 0.360 | 0.224 | 0.139 | 2.219 | 0.301 |
| 2703 | **Zc3hc1** | Unchanged | Q80YV2 | 3 | 4 | 3 | 501 | 55.2 | 5.38 | 10 | 1.266 | 0.694 | 0.589 | 0.505 | 1.068 | 0.906 | 0.552 | 0.452 |
| 2704 | **Dcn** | UP | P28654 | 3 | 10 | 3 | 354 | 39.8 | 8.68 | 10 | 0.674 | 0.512 | 7.604 | 0.127 | 4.648 | 0.034 | 1.636 | 0.492 |
| 2705 | **Otol1** | Unchanged | Q4ZJM7 | 1 | 1 | 1 | 482 | 49.6 | 8.81 | 10 | 1.276 | 0.712 | 1.153 | 0.815 | 2.897 | 0.307 | 0.398 | 0.357 |
| 2706 | **Pid1** | Unchanged | Q3UBG2 | 1 | 1 | 1 | 217 | 24.8 | 6.99 | 10 | 1.428 | 0.640 | 1.684 | 0.476 | 0.247 | 0.213 | 6.826 | 0.093 |
| 2707 | **Pnkd** | Unchanged | Q69ZP3 | 2 | 7 | 2 | 385 | 43 | 8.94 | 10 | 1.361 | 0.638 | 1.084 | 0.890 | 1.063 | 0.920 | 1.020 | 0.975 |
| 2708 | **Tmem258** | Unchanged | P61166 | 1 | 1 | 1 | 79 | 9.1 | 5.83 | 10 | 0.405 | 0.260 | 0.692 | 0.564 | 0.090 | 0.071 | 7.711 | 0.052 |
| 2709 | **Chac2** | Unchanged | Q9CQG1 | 1 | 6 | 1 | 178 | 20.1 | 5.68 | 10 | 1.691 | 0.438 | 2.450 | 0.335 | 2.049 | 0.523 | 1.195 | 0.846 |
| 2710 | **Snrpa** | Unchanged | Q62189 | 1 | 1 | 1 | 287 | 31.8 | 9.8 | 10 | #DIV/0! | #DIV/0! | #DIV/0! | #DIV/0! | #DIV/0! | #DIV/0! | #DIV/0! | #DIV/0! |
| 2711 | **S1pr4** | Unchanged | Q9Z0L1 | 1 | 1 | 1 | 386 | 42.2 | 9.77 | 10 | 0.376 | 0.349 | 0.286 | 0.286 | 0.196 | 0.232 | 1.459 | 0.306 |
| 2712 | **Ubxn4** | Down | A0A0R4J078 | 3 | 15 | 3 | 506 | 56.4 | 6.61 | 10 | 0.560 | 0.100 | 0.544 | 0.148 | 0.376 | 0.014 | 1.447 | 0.498 |
| 2713 | **Napb** | Unchanged | P28663 | 2 | 11 | 1 | 298 | 33.5 | 5.47 | 10 | 1.048 | 0.926 | 1.049 | 0.941 | 1.262 | 0.627 | 0.831 | 0.773 |
| 2714 | **Slc25a21** | Unchanged | Q8BZ09 | 3 | 19 | 3 | 298 | 33.2 | 9.72 | 10 | 0.898 | 0.885 | 0.276 | 0.190 | 0.521 | 0.411 | 0.529 | 0.316 |
| 2715 | **Commd4** | UP | Q9CQ02 | 1 | 9 | 1 | 199 | 21.8 | 6.67 | 10 | 2.606 | 0.400 | 3.797 | 0.138 | 4.580 | 0.032 | 0.829 | 0.721 |
| 2716 | **Il1b** | Unchanged | P10749 | 1 | 2 | 1 | 269 | 30.9 | 4.7 | 10 | 1.163 | 0.855 | 0.816 | 0.821 | 0.991 | 0.991 | 0.824 | 0.750 |
| 2717 | **Srsf6** | Unchanged | Q3TWW8 | 3 | 5 | 3 | 339 | 39 | 11.46 | 10 | 1.358 | 0.561 | 1.592 | 0.616 | 1.361 | 0.579 | 1.170 | 0.856 |
| 2718 | **Man2c1** | Unchanged | Q91W89 | 6 | 19 | 6 | 1039 | 115.6 | 6.51 | 10 | 0.626 | 0.368 | 1.185 | 0.732 | 0.853 | 0.829 | 1.390 | 0.631 |
| 2719 | **Dcaf8** | Unchanged | Q8N7N5 | 4 | 23 | 4 | 591 | 66 | 5.87 | 7 | 1.791 | 0.332 | 12.619 | 0.254 | 3.177 | 0.326 | 3.972 | 0.357 |
| 2720 | **Wbp2** | Unchanged | P97765 | 2 | 2 | 2 | 261 | 28 | 6.33 | 10 | 0.951 | 0.931 | 1.743 | 0.242 | 0.869 | 0.790 | 2.005 | 0.132 |
| 2721 | **Rcan1** | Unchanged | Q9JHG6 | 2 | 3 | 2 | 251 | 28.1 | 5.41 | 10 | 1.117 | 0.846 | 1.193 | 0.767 | 1.873 | 0.378 | 0.637 | 0.522 |
| 2722 | **Mia2** | Down | Q91ZV0 | 12 | 35 | 12 | 1396 | 156.4 | 4.55 | 10 | 0.677 | 0.190 | 0.815 | 0.482 | 0.515 | 0.043 | 1.582 | 0.131 |
| 2723 | **Lrat** | Unchanged | Q9JI60 | 1 | 1 | 1 | 231 | 25.8 | 8.7 | 10 | 0.821 | 0.784 | 2.243 | 0.454 | 1.270 | 0.724 | 1.765 | 0.537 |
| 2724 | **Setd3** | Unchanged | Q91WC0 | 4 | 8 | 4 | 594 | 67.1 | 5.6 | 10 | 0.623 | 0.191 | 0.791 | 0.581 | 0.436 | 0.088 | 1.816 | 0.308 |
| 2725 | **Prss54** | Unchanged | A6H685 | 1 | 1 | 1 | 383 | 42.8 | 8.35 | 10 | 0.569 | 0.210 | 0.652 | 0.149 | 0.857 | 0.619 | 0.760 | 0.348 |
| 2726 | **Gtf2i** | Unchanged | Q9ESZ8 | 6 | 13 | 6 | 998 | 112.2 | 6.55 | 10 | 1.638 | 0.388 | 0.970 | 0.961 | 1.767 | 0.431 | 0.549 | 0.438 |
| 2727 | **Commd10** | Unchanged | Q8JZY2 | 1 | 1 | 1 | 202 | 22.8 | 6.65 | 10 | 0.725 | 0.458 | 1.440 | 0.297 | 1.703 | 0.118 | 0.846 | 0.389 |
| 2728 | **Slc22a1** | Unchanged | O08966 | 4 | 51 | 4 | 556 | 61.5 | 6.67 | 7 | 1.425 | 0.416 | 0.832 | 0.673 | 1.458 | 0.404 | 0.570 | 0.319 |
| 2729 | **Arhgef1** | Unchanged | Q61210 | 5 | 8 | 5 | 920 | 102.7 | 5.6 | 10 | 0.941 | 0.888 | 1.084 | 0.779 | 1.236 | 0.408 | 0.877 | 0.574 |
| 2730 | **Lin7a** | Down | Q8JZS0 | 2 | 3 | 1 | 233 | 26 | 8.72 | 10 | 2.964 | 0.138 | 0.226 | 0.004 | 1.653 | 0.386 | 0.136 | 0.076 |
| 2731 | **Ccar2** | Unchanged | Q8VDP4 | 4 | 13 | 4 | 922 | 102.9 | 5.25 | 10 | 1.140 | 0.852 | 1.020 | 0.978 | 0.680 | 0.571 | 1.499 | 0.536 |
| 2732 | **Hist1h1b** | Unchanged | P43276 | 2 | 8 | 2 | 223 | 22.6 | 10.92 | 10 | 0.996 | 0.992 | 1.891 | 0.330 | 4.623 | 0.199 | 0.409 | 0.340 |
| 2733 | **Hexa** | Unchanged | P29416 | 3 | 5 | 3 | 528 | 60.6 | 6.54 | 10 | 1.871 | 0.575 | 1.573 | 0.398 | 1.220 | 0.773 | 1.290 | 0.603 |
| 2734 | **Dnph1** | Unchanged | Q80VJ3 | 1 | 8 | 1 | 173 | 19 | 5.38 | 10 | 0.629 | 0.679 | 1.677 | 0.589 | 1.205 | 0.820 | 1.392 | 0.649 |
| 2735 | **Necap1** | Unchanged | Q9CR95 | 1 | 12 | 1 | 275 | 29.6 | 6.38 | 10 | 0.879 | 0.847 | 0.388 | 0.243 | 0.673 | 0.552 | 0.576 | 0.444 |
| 2736 | **Tmem106b** | Unchanged | Q80X71 | 1 | 1 | 1 | 275 | 31.2 | 6.68 | 10 | 0.716 | 0.731 | 0.302 | 0.392 | 0.659 | 0.675 | 0.459 | 0.389 |

| | | | | | | | | | | | | | | | | | | |
|---|---|---|---|---|---|---|---|---|---|---|---|---|---|---|---|---|---|---|
| 2737 | **Rragc** | Unchanged | Q99K70 | 3 | 8 | 3 | 398 | 44.1 | 5.1 | 10 | 1.387 | 0.601 | 0.945 | 0.932 | 0.235 | 0.107 | 4.014 | 0.177 |
| 2738 | **P2rx4** | Unchanged | Q9JJX6 | 2 | 3 | 2 | 388 | 43.4 | 7.99 | 10 | 0.626 | 0.232 | 0.501 | 0.111 | 0.966 | 0.916 | 0.519 | 0.079 |
| 2739 | **Ca14** | Down | Q9WVT6 | 2 | 9 | 2 | 337 | 37.5 | 6.35 | 10 | 1.894 | 0.100 | 1.046 | 0.888 | 0.445 | 0.045 | 2.351 | 0.082 |
| 2740 | **Mob4** | Unchanged | Q6PEB6 | 1 | 3 | 1 | 225 | 26 | 5.78 | 10 | 0.800 | 0.658 | 1.509 | 0.510 | 0.635 | 0.423 | 2.377 | 0.200 |
| 2741 | **Slc37a4** | Unchanged | A0A1L1SUI3 | 2 | 8 | 2 | 450 | 48.4 | 8.32 | 10 | 0.743 | 0.399 | 0.443 | 0.059 | 0.396 | 0.092 | 1.120 | 0.856 |
| 2742 | **Pdrg1** | Unchanged | P59048 | 1 | 1 | 1 | 133 | 15.4 | 8.1 | 10 | 0.161 | 0.284 | 1.650 | 0.569 | 0.218 | 0.316 | 7.565 | 0.118 |
| 2743 | **Sel1l** | Unchanged | Q9Z2G6 | 4 | 21 | 4 | 790 | 88.3 | 5.57 | 10 | 1.065 | 0.901 | 0.759 | 0.591 | 0.420 | 0.123 | 1.807 | 0.362 |
| 2744 | **Nmrk1** | Unchanged | Q91W63 | 1 | 1 | 1 | 195 | 22.3 | 4.89 | 10 | #DIV/0! | #DIV/0! | #DIV/0! | #DIV/0! | #DIV/0! | #DIV/0! | #DIV/0! | #DIV/0! |
| 2745 | **Bid** | Unchanged | P70444 | 1 | 8 | 1 | 195 | 21.9 | 4.81 | 10 | 0.895 | 0.881 | 1.393 | 0.550 | 0.581 | 0.499 | 2.397 | 0.110 |
| 2746 | **Ppbp** | Unchanged | Q9EQI5 | 1 | 1 | 1 | 113 | 12.2 | 8.72 | 10 | #DIV/0! | #DIV/0! | #DIV/0! | #DIV/0! | #DIV/0! | #DIV/0! | #DIV/0! | #DIV/0! |
| 2747 | **Dio1** | Unchanged | Q61153 | 1 | 2 | 1 | 257 | 29.6 | 9.29 | 10 | #DIV/0! | #DIV/0! | #DIV/0! | #DIV/0! | #DIV/0! | #DIV/0! | #DIV/0! | #DIV/0! |
| 2748 | **Neil2** | Unchanged | Q6R2P8 | 1 | 1 | 1 | 329 | 36.8 | 8.5 | 10 | 8.580 | 0.195 | 8.271 | 0.366 | 0.616 | 0.263 | 13.430 | 0.343 |
| 2749 | **Fuk** | Unchanged | Q7TMC8 | 5 | 22 | 5 | 1090 | 119.2 | 6.6 | 10 | 1.010 | 0.978 | 4.071 | 0.123 | 0.890 | 0.786 | 4.577 | 0.110 |
| 2750 | **Dlg1** | Unchanged | Q811D0 | 5 | 8 | 5 | 905 | 100.1 | 5.8 | 10 | 1.168 | 0.705 | 1.153 | 0.779 | 2.019 | 0.189 | 0.571 | 0.288 |
| 2751 | **Mob3c** | Unchanged | Q8BJG4 | 1 | 1 | 1 | 216 | 25.5 | 8.44 | 10 | #DIV/0! | #DIV/0! | #DIV/0! | #DIV/0! | #DIV/0! | #DIV/0! | #DIV/0! | #DIV/0! |
| 2752 | **Lima1** | Unchanged | Q9ERG0 | 5 | 20 | 5 | 753 | 84 | 6.6 | 10 | 0.812 | 0.513 | 1.046 | 0.886 | 0.785 | 0.517 | 1.331 | 0.413 |
| 2753 | **Mrpl37** | Unchanged | Q921S7 | 3 | 32 | 3 | 423 | 48.3 | 8.84 | 10 | 1.441 | 0.234 | 1.544 | 0.212 | 1.018 | 0.953 | 1.517 | 0.290 |
| 2754 | **Amy2** | Unchanged | P00688 | 2 | 2 | 2 | 508 | 57.3 | 7.28 | 10 | 3.623 | 0.403 | 0.748 | 0.692 | 1.860 | 0.437 | 0.402 | 0.262 |
| 2755 | **Ggcx** | Unchanged | Q9QYC7 | 5 | 57 | 5 | 757 | 87.1 | 8.02 | 10 | 0.952 | 0.898 | 2.221 | 0.225 | 1.356 | 0.592 | 1.638 | 0.463 |
| 2756 | **Napg** | UP | Q9CWZ7 | 2 | 6 | 2 | 312 | 34.7 | 5.41 | 10 | 1.561 | 0.180 | 1.103 | 0.840 | 2.997 | 0.007 | 0.368 | 0.013 |
| 2757 | **Dhrs3** | Unchanged | O88876 | 2 | 3 | 2 | 302 | 33.6 | 8.69 | 10 | 2.601 | 0.130 | 7.654 | 0.115 | 2.979 | 0.101 | 2.569 | 0.266 |
| 2758 | **Enpp1** | Unchanged | P06802 | 5 | 27 | 5 | 906 | 103.1 | 6.6 | 10 | 1.232 | 0.420 | 1.910 | 0.112 | 1.050 | 0.870 | 1.820 | 0.140 |
| 2759 | **Sf3b6** | Unchanged | P59708 | 1 | 8 | 1 | 125 | 14.6 | 9.38 | 10 | 1.084 | 0.780 | 1.048 | 0.851 | 1.194 | 0.575 | 0.877 | 0.678 |
| 2760 | **Mocs3** | Unchanged | A2BDX3 | 2 | 18 | 2 | 460 | 49.3 | 7.49 | 10 | 0.904 | 0.848 | 0.865 | 0.631 | 0.696 | 0.308 | 1.243 | 0.452 |
| 2761 | **Mien1** | Unchanged | Q9CQ86 | 1 | 6 | 1 | 115 | 12.3 | 4.51 | 10 | 5.974 | 0.398 | 0.197 | 0.109 | 0.239 | 0.113 | 0.822 | 0.830 |
| 2762 | **Ldb3** | Unchanged | Q9JKS4 | 3 | 6 | 3 | 723 | 76.4 | 7.75 | 10 | 0.404 | 0.220 | 0.739 | 0.645 | 0.593 | 0.458 | 1.247 | 0.734 |
| 2763 | **Pak2** | Unchanged | Q8CIN4 | 3 | 14 | 3 | 524 | 57.9 | 5.77 | 10 | 1.266 | 0.647 | 1.325 | 0.591 | 2.586 | 0.115 | 0.512 | 0.179 |
| 2764 | **Cttn** | Unchanged | Q60598 | 4 | 26 | 4 | 546 | 61.2 | 5.4 | 10 | 0.713 | 0.622 | 0.335 | 0.191 | 0.204 | 0.119 | 1.642 | 0.221 |
| 2765 | **Shtn1** | Down | Q8K2Q9 | 5 | 12 | 5 | 631 | 71.3 | 5.44 | 8 | 0.208 | 0.004 | 0.814 | 0.522 | 0.801 | 0.610 | 1.017 | 0.973 |
| 2766 | **Cul3** | Unchanged | Q9JLV5 | 5 | 13 | 5 | 768 | 88.9 | 8.46 | 10 | 0.651 | 0.236 | 1.498 | 0.188 | 0.846 | 0.615 | 1.770 | 0.046 |
| 2767 | **Ece1** | Unchanged | Q4PZA2 | 4 | 11 | 4 | 769 | 87 | 5.91 | 9 | 1.114 | 0.768 | 2.942 | 0.113 | 4.762 | 0.070 | 0.618 | 0.406 |
| 2768 | **Spdye4c** | Unchanged | I6XKQ3 | 1 | 1 | 1 | 339 | 38.2 | 9.79 | 9 | 0.083 | 0.164 | 0.136 | 0.187 | 0.143 | 0.190 | 0.954 | 0.758 |
| 2769 | **Uros** | Unchanged | P51163 | 1 | 3 | 1 | 265 | 28.5 | 6.54 | 9 | 0.762 | 0.556 | 1.029 | 0.954 | 0.946 | 0.902 | 1.087 | 0.882 |
| 2770 | **Kdelr2** | Unchanged | Q9CQM2 | 2 | 5 | 2 | 212 | 24.4 | 8.72 | 9 | 1.369 | 0.640 | 1.776 | 0.405 | 1.471 | 0.577 | 1.208 | 0.750 |
| 2771 | **Lmf1** | Unchanged | Q3U3R4 | 4 | 7 | 4 | 574 | 65.8 | 9.29 | 9 | 1.961 | 0.338 | 2.927 | 0.253 | 1.561 | 0.486 | 1.875 | 0.415 |
| 2772 | **Creb3** | Unchanged | Q61817 | 1 | 1 | 1 | 404 | 45.1 | 5.01 | 9 | 1.307 | 0.397 | 1.135 | 0.741 | 1.295 | 0.496 | 0.877 | 0.758 |
| 2773 | **Ubqln2** | UP | Q9QZM0 | 4 | 41 | 1 | 638 | 67.3 | 5.22 | 9 | 1.034 | 0.968 | 1.339 | 0.750 | 28.884 | 0.006 | 0.046 | 0.006 |
| 2774 | **Faf1** | UP | P54731 | 4 | 6 | 4 | 649 | 73.8 | 4.86 | 9 | 2.303 | 0.041 | 3.362 | 0.156 | 1.336 | 0.400 | 2.517 | 0.222 |
| 2775 | **Oasl1** | Unchanged | Q8VI94 | 3 | 8 | 3 | 511 | 59.1 | 7.14 | 9 | 0.609 | 0.116 | 1.019 | 0.968 | 2.658 | 0.318 | 0.383 | 0.333 |
| 2776 | **Col25a1** | Unchanged | E9Q5L6 | 2 | 3 | 2 | 640 | 62.6 | 8.84 | 9 | 1.208 | 0.854 | 0.380 | 0.548 | 1.275 | 0.815 | 0.298 | 0.175 |
| 2777 | **Otulin** | Unchanged | Q3UCV8 | 3 | 5 | 3 | 352 | 40.3 | 5.68 | 9 | 1.155 | 0.767 | 1.395 | 0.576 | 2.175 | 0.149 | 0.641 | 0.371 |

| | | | | | | | | | | | | | | | | | | |
|---|---|---|---|---|---|---|---|---|---|---|---|---|---|---|---|---|---|---|
| 2778 | **Myo1c** | Unchanged | Q9WTI7 | 7 | 48 | 7 | 1063 | 121.9 | 9.35 | 9 | 1.282 | 0.020 | 1.349 | 0.222 | 0.801 | 0.500 | 1.685 | 0.174 |
| 2779 | **Erg28** | Unchanged | Q9ERY9 | 1 | 2 | 1 | 140 | 15.8 | 9.91 | 9 | 0.680 | 0.414 | 1.127 | 0.799 | 0.803 | 0.594 | 1.403 | 0.543 |
| 2780 | **Cnp** | Unchanged | P16330 | 2 | 8 | 2 | 420 | 47.1 | 8.97 | 9 | 1.807 | 0.275 | 2.380 | 0.199 | 1.508 | 0.304 | 1.578 | 0.398 |
| 2781 | **Copz2** | Unchanged | Q9JHH9 | 1 | 2 | 1 | 205 | 22.9 | 5.17 | 9 | 0.521 | 0.182 | 0.348 | 0.059 | 1.247 | 0.782 | 0.279 | 0.302 |
| 2782 | **Zmpste24** | Unchanged | Q80W54 | 3 | 29 | 3 | 475 | 54.7 | 6.95 | 9 | 1.336 | 0.582 | 1.895 | 0.137 | 1.919 | 0.130 | 0.988 | 0.948 |
| 2783 | **Lsr** | Unchanged | Q99KG5 | 4 | 6 | 4 | 594 | 66.1 | 6.54 | 9 | 1.824 | 0.420 | 0.560 | 0.323 | 0.712 | 0.460 | 0.788 | 0.624 |
| 2784 | **Apoa5** | Unchanged | Q8C7G5 | 2 | 21 | 2 | 368 | 41.2 | 6.01 | 9 | 2.528 | 0.334 | 3.182 | 0.169 | 1.042 | 0.832 | 3.054 | 0.178 |
| 2785 | **H2-K1iso** | Unchanged | P01902 | 3 | 21 | 1 | 368 | 41.5 | 6.62 | 9 | #DIV/0! | #DIV/0! | #DIV/0! | #DIV/0! | #DIV/0! | #DIV/0! | #DIV/0! | #DIV/0! |
| 2786 | **Rae1** | Down | Q8C570 | 2 | 7 | 2 | 368 | 40.9 | 7.83 | 9 | 1.581 | 0.275 | 0.240 | 0.002 | 2.539 | 0.239 | 0.094 | 0.092 |
| 2787 | **Krt17** | Unchanged | Q9QWL7 | 3 | 5 | 2 | 433 | 48.1 | 5.06 | 9 | 4.356 | 0.221 | 0.976 | 0.955 | 0.907 | 0.802 | 1.076 | 0.871 |
| 2788 | **Yme1l1** | UP | O88967 | 4 | 7 | 4 | 715 | 80 | 8.97 | 9 | 1.982 | 0.286 | 4.626 | 0.041 | 1.919 | 0.150 | 2.410 | 0.117 |
| 2789 | **Eif2b5** | Unchanged | Q8CHW4 | 3 | 8 | 3 | 717 | 80 | 5.07 | 9 | 0.374 | 0.297 | 0.430 | 0.358 | 0.592 | 0.485 | 0.727 | 0.603 |
| 2790 | **Clec2d** | Unchanged | Q91V08 | 1 | 1 | 1 | 207 | 23.6 | 7.62 | 9 | #DIV/0! | #DIV/0! | #DIV/0! | #DIV/0! | #DIV/0! | #DIV/0! | #DIV/0! | #DIV/0! |
| 2791 | **Exoc1** | Unchanged | Q8R3S6 | 4 | 17 | 4 | 894 | 101.8 | 6.52 | 9 | 1.188 | 0.737 | 1.959 | 0.151 | 1.763 | 0.144 | 1.111 | 0.764 |
| 2792 | **Gm5478** | Down | A0A2R8VHP3 | 5 | 34 | 2 | 535 | 57.9 | 6.2 | 9 | 0.569 | 0.109 | 0.855 | 0.761 | 0.273 | 0.006 | 3.130 | 0.222 |
| 2793 | **Mrpl42** | Unchanged | Q9CPV3 | 1 | 1 | 1 | 142 | 16.5 | 9.19 | 9 | #DIV/0! | #DIV/0! | #DIV/0! | #DIV/0! | #DIV/0! | #DIV/0! | #DIV/0! | #DIV/0! |
| 2794 | **Snx30** | Unchanged | Q8CE50 | 1 | 1 | 1 | 437 | 49.5 | 5.35 | 9 | 1.202 | 0.483 | 0.750 | 0.513 | 0.632 | 0.256 | 1.187 | 0.716 |
| 2795 | **Dhcr7** | Unchanged | O88455 | 3 | 33 | 3 | 471 | 53.9 | 8.35 | 9 | 0.526 | 0.078 | 0.646 | 0.259 | 0.640 | 0.298 | 1.009 | 0.988 |
| 2796 | **Myh11** | UP | A0A338P6K2 | 13 | 56 | 3 | 1979 | 227.7 | 5.48 | 9 | 1.789 | 0.132 | 1.658 | 0.387 | 5.053 | 0.022 | 0.328 | 0.062 |
| 2797 | **Pin1** | Unchanged | Q9QUR7 | 1 | 2 | 1 | 165 | 18.4 | 8.79 | 9 | 1.040 | 0.948 | 0.658 | 0.537 | 0.284 | 0.122 | 2.318 | 0.303 |
| 2798 | **Arsa** | Unchanged | P50428 | 2 | 26 | 2 | 506 | 53.7 | 5.87 | 9 | 0.903 | 0.855 | 1.289 | 0.654 | 4.987 | 0.054 | 0.259 | 0.073 |
| 2799 | **Adh7** | Unchanged | Q64437 | 3 | 25 | 1 | 374 | 39.9 | 7.85 | 9 | 1.256 | 0.508 | 5.889 | 0.301 | 1.554 | 0.188 | 3.789 | 0.356 |
| 2800 | **Dnajc12** | Unchanged | Q9R022 | 1 | 1 | 1 | 198 | 22.8 | 6.15 | 9 | #DIV/0! | #DIV/0! | #DIV/0! | #DIV/0! | #DIV/0! | #DIV/0! | #DIV/0! | #DIV/0! |
| 2801 | **Hint3** | Unchanged | Q9CPS6 | 1 | 2 | 1 | 165 | 18.8 | 6.33 | 9 | 1.282 | 0.621 | 1.382 | 0.700 | 0.933 | 0.901 | 1.480 | 0.637 |
| 2802 | **Fli1** | Unchanged | P26323 | 1 | 1 | 1 | 452 | 51 | 7.06 | 9 | #DIV/0! | #DIV/0! | #DIV/0! | #DIV/0! | #DIV/0! | #DIV/0! | #DIV/0! | #DIV/0! |
| 2803 | **Farp1** | Unchanged | F8VPU2 | 5 | 16 | 5 | 1048 | 118.8 | 7.88 | 9 | 0.835 | 0.452 | 1.206 | 0.501 | 1.206 | 0.626 | 1.000 | 1.000 |
| 2804 | **Stx4** | Unchanged | P70452 | 2 | 4 | 2 | 298 | 34.1 | 6.14 | 9 | 0.895 | 0.787 | 0.263 | 0.056 | 0.939 | 0.887 | 0.279 | 0.030 |
| 2805 | **Eif4g2** | Unchanged | Q62448 | 7 | 51 | 7 | 906 | 102 | 7.14 | 9 | 0.962 | 0.907 | 1.077 | 0.849 | 0.823 | 0.607 | 1.309 | 0.454 |
| 2806 | **Pls1** | Unchanged | Q3V0K9 | 5 | 44 | 1 | 630 | 70.4 | 5.38 | 9 | 3.778 | 0.177 | 8.432 | 0.181 | 5.021 | 0.296 | 1.679 | 0.598 |
| 2807 | **Amelx** | Unchanged | P63277 | 1 | 3 | 1 | 210 | 23.5 | 7.25 | 9 | 2.404 | 0.300 | 1.425 | 0.688 | 2.492 | 0.254 | 0.572 | 0.494 |
| 2808 | **Xrn2** | Unchanged | Q9DBR1 | 5 | 21 | 5 | 951 | 108.6 | 7.59 | 9 | 0.605 | 0.523 | 0.855 | 0.809 | 0.426 | 0.355 | 2.006 | 0.006 |
| 2809 | **Sfxn3** | Unchanged | Q91V61 | 2 | 12 | 1 | 321 | 35.4 | 9.51 | 9 | 0.153 | 0.230 | 0.006 | 0.162 | 0.004 | 0.162 | 1.324 | 0.601 |
| 2810 | **Tcf25** | Unchanged | Q8R3L2 | 2 | 3 | 2 | 676 | 76.6 | 6.51 | 9 | 1.405 | 0.355 | 1.597 | 0.295 | 0.945 | 0.911 | 1.691 | 0.240 |
| 2811 | **Mrps26** | Unchanged | Q80ZS3 | 1 | 14 | 1 | 200 | 23.4 | 9.96 | 9 | 0.272 | 0.267 | 1.736 | 0.616 | 0.586 | 0.540 | 2.964 | 0.400 |
| 2812 | **Gbp2** | Unchanged | Q9Z0E6 | 3 | 6 | 3 | 589 | 66.7 | 5.71 | 9 | 0.995 | 0.990 | 1.898 | 0.428 | 1.070 | 0.853 | 1.774 | 0.465 |
| 2813 | **Pcnp** | Unchanged | Q6P8I4 | 1 | 3 | 1 | 178 | 19 | 7.49 | 9 | 0.153 | 0.105 | 1.576 | 0.669 | 3.015 | 0.259 | 0.523 | 0.494 |
| 2814 | **Snap47** | Unchanged | Q8R570 | 3 | 9 | 3 | 413 | 46.5 | 5.76 | 9 | 1.272 | 0.575 | 1.741 | 0.392 | 0.818 | 0.543 | 2.128 | 0.279 |
| 2815 | **Pfdn4** | Unchanged | Q3UWL8 | 1 | 1 | 1 | 134 | 15.2 | 4.61 | 9 | 0.614 | 0.491 | 0.214 | 0.085 | 0.416 | 0.201 | 0.515 | 0.256 |
| 2816 | **Miga2** | Unchanged | Q8BK03 | 2 | 5 | 2 | 593 | 65.5 | 5.41 | 9 | 0.702 | 0.308 | 0.834 | 0.568 | 0.810 | 0.457 | 1.030 | 0.910 |
| 2817 | **Fam98a** | Down | Q3TJZ6 | 3 | 20 | 3 | 515 | 55 | 8.95 | 9 | 0.744 | 0.477 | 0.556 | 0.276 | 0.194 | 0.029 | 2.865 | 0.182 |
| 2818 | **Commd5** | Unchanged | A0A0R4J0U7 | 1 | 2 | 1 | 224 | 24.5 | 8.06 | 9 | #DIV/0! | #DIV/0! | #DIV/0! | #DIV/0! | #DIV/0! | #DIV/0! | #DIV/0! | #DIV/0! |

| | | | | | | | | | | | | | | | | | | |
|---|---|---|---|---|---|---|---|---|---|---|---|---|---|---|---|---|---|---|
| 2819 | **Apcs** | Unchanged | P12246 | 1 | 12 | 1 | 224 | 26.2 | 6.35 | 9 | 0.319 | 0.099 | 0.362 | 0.140 | 0.700 | 0.532 | 0.517 | 0.436 |
| 2820 | **Ap3d1** | Unchanged | O54774 | 6 | 17 | 6 | 1199 | 135 | 7.37 | 9 | 3.847 | 0.215 | 4.725 | 0.051 | 4.774 | 0.280 | 0.990 | 0.990 |
| 2821 | **Coq3** | Down | Q8BMS4 | 2 | 19 | 2 | 370 | 40.9 | 7.96 | 9 | 0.182 | 0.026 | 0.217 | 0.036 | 0.173 | 0.023 | 1.251 | 0.740 |
| 2822 | **Fubp1** | Unchanged | Q91WJ8 | 4 | 14 | 4 | 651 | 68.5 | 7.93 | 9 | 0.662 | 0.472 | 1.010 | 0.987 | 0.640 | 0.461 | 1.580 | 0.499 |
| 2823 | **Mapre3** | Unchanged | Q6PER3 | 1 | 2 | 1 | 281 | 31.9 | 5.54 | 9 | 1.638 | 0.518 | 1.572 | 0.593 | 1.705 | 0.452 | 0.922 | 0.907 |
| 2824 | **Rab17** | Unchanged | P35292 | 1 | 8 | 1 | 214 | 23.6 | 5.44 | 9 | 0.807 | 0.666 | 3.520 | 0.133 | 1.889 | 0.204 | 1.863 | 0.321 |
| 2825 | **Tbc1d7** | UP | Q9D0K0 | 1 | 1 | 1 | 293 | 33.8 | 7.2 | 9 | 8.829 | 0.066 | 1.447 | 0.636 | 17.011 | 0.047 | 0.085 | 0.053 |
| 2826 | **Apoc4** | Unchanged | Q61268 | 1 | 4 | 1 | 124 | 14.3 | 9.52 | 9 | 0.727 | 0.521 | 0.515 | 0.194 | 0.997 | 0.996 | 0.517 | 0.340 |
| 2827 | **Trim14** | UP | Q8BVW3 | 3 | 7 | 3 | 440 | 49.6 | 7.93 | 7 | 6.595 | 0.040 | 0.551 | 0.217 | 2.391 | 0.102 | 0.230 | 0.034 |
| 2828 | **Rnps1** | Unchanged | Q99M28 | 2 | 23 | 2 | 305 | 34.2 | 11.84 | 9 | 1.062 | 0.847 | 1.029 | 0.952 | 0.775 | 0.499 | 1.328 | 0.600 |
| 2829 | **Rabggtb** | Unchanged | P53612 | 2 | 3 | 2 | 339 | 37.8 | 5.16 | 9 | 0.967 | 0.936 | 3.029 | 0.111 | 0.792 | 0.473 | 3.822 | 0.080 |
| 2830 | **Dpm1** | Unchanged | O70152 | 2 | 23 | 2 | 260 | 29.2 | 9.51 | 9 | 1.786 | 0.493 | 2.310 | 0.170 | 0.372 | 0.060 | 6.208 | 0.047 |
| 2831 | **Slc17a3** | Unchanged | G3UWD9 | 2 | 2 | 2 | 498 | 54.5 | 8.6 | 9 | 0.849 | 0.828 | 0.396 | 0.406 | 0.314 | 0.332 | 1.259 | 0.741 |
| 2832 | **Nadsyn1** | Down | Q711T7 | 4 | 12 | 4 | 725 | 81.6 | 6.49 | 9 | 1.062 | 0.829 | 0.416 | 0.013 | 0.796 | 0.518 | 0.523 | 0.186 |
| 2833 | **Gbf1** | Unchanged | Q6DFZ1 | 10 | 15 | 10 | 1861 | 206.7 | 5.8 | 9 | 2.249 | 0.056 | 3.195 | 0.262 | 9.254 | 0.268 | 0.345 | 0.422 |
| 2834 | **Hjv** | UP | Q7TQ32 | 1 | 1 | 1 | 420 | 44.8 | 7.84 | 9 | 1.607 | 0.339 | 3.108 | 0.058 | 2.435 | 0.013 | 1.276 | 0.494 |
| 2835 | **Pycard** | Unchanged | Q9EPB4 | 2 | 4 | 2 | 193 | 21.4 | 5.43 | 9 | 0.693 | 0.596 | 1.671 | 0.518 | 0.799 | 0.663 | 2.093 | 0.361 |
| 2836 | **Pex7** | Unchanged | P97865 | 2 | 10 | 2 | 318 | 35.5 | 5.58 | 9 | 2.222 | 0.315 | 0.946 | 0.884 | 0.620 | 0.216 | 1.526 | 0.389 |
| 2837 | **Tmlhe** | Unchanged | Q91ZE0 | 3 | 16 | 3 | 421 | 49.6 | 8.25 | 9 | 0.942 | 0.884 | 2.253 | 0.209 | 1.269 | 0.476 | 1.775 | 0.306 |
| 2838 | **Irf6** | Unchanged | P97431 | 2 | 7 | 2 | 467 | 53.1 | 5.33 | 9 | 0.947 | 0.905 | 1.308 | 0.518 | 0.811 | 0.624 | 1.612 | 0.187 |
| 2839 | **Klkb1** | Down | H9H9R5 | 4 | 14 | 4 | 638 | 71.3 | 8.02 | 9 | 0.861 | 0.612 | 0.394 | 0.012 | 0.457 | 0.022 | 0.863 | 0.621 |
| 2840 | **Nenf** | Unchanged | Q9CQ45 | 1 | 1 | 1 | 171 | 18.9 | 5.27 | 9 | 1.017 | 0.970 | 1.485 | 0.511 | 0.793 | 0.649 | 1.872 | 0.292 |
| 2841 | **Timmdc1** | UP | Q8BUY5 | 1 | 4 | 1 | 285 | 31.8 | 9.29 | 9 | 1.529 | 0.446 | 2.159 | 0.192 | 3.565 | 0.003 | 0.606 | 0.098 |
| 2842 | **Apon** | Unchanged | G3X9D6 | 1 | 4 | 1 | 251 | 27.9 | 5.69 | 9 | 0.789 | 0.373 | 0.621 | 0.164 | 0.646 | 0.232 | 0.960 | 0.919 |
| 2843 | **Nudcd1** | Unchanged | Q6PIP5 | 3 | 10 | 3 | 582 | 66.7 | 5.3 | 9 | 2.265 | 0.206 | 0.699 | 0.418 | 0.617 | 0.305 | 1.134 | 0.824 |
| 2844 | **C9** | Unchanged | P06683 | 3 | 14 | 3 | 548 | 62 | 5.78 | 9 | 1.225 | 0.527 | 1.462 | 0.583 | 0.948 | 0.862 | 1.542 | 0.530 |
| 2845 | **Map1lc3b** | Unchanged | M0QWC2 | 1 | 1 | 1 | 160 | 18.2 | 9.86 | 9 | 1.163 | 0.723 | 0.512 | 0.138 | 0.620 | 0.228 | 0.825 | 0.649 |
| 2846 | **Gatm** | Unchanged | Q9D964 | 2 | 9 | 2 | 423 | 48.3 | 7.88 | 9 | 1.486 | 0.331 | 1.636 | 0.328 | 0.835 | 0.514 | 1.960 | 0.203 |
| 2847 | **Nagk** | Unchanged | Q9QZ08 | 1 | 4 | 1 | 343 | 37.2 | 5.73 | 9 | #DIV/0! | #DIV/0! | #DIV/0! | #DIV/0! | #DIV/0! | #DIV/0! | #DIV/0! | #DIV/0! |
| 2848 | **Dcaf11** | Unchanged | Q91VU6 | 3 | 8 | 3 | 549 | 62 | 6.44 | 9 | 1.236 | 0.770 | 1.403 | 0.622 | 0.255 | 0.266 | 5.502 | 0.048 |
| 2849 | **Emc4** | Unchanged | Q9CZX9 | 1 | 17 | 1 | 183 | 20.1 | 8.62 | 9 | 1.238 | 0.518 | 0.820 | 0.578 | 1.026 | 0.949 | 0.799 | 0.647 |
| 2850 | **Thop1** | Down | Q8C1A5 | 5 | 23 | 5 | 687 | 78 | 6.06 | 9 | 2.514 | 0.137 | 2.766 | 0.125 | 0.413 | 0.016 | 6.698 | 0.050 |
| 2851 | **D6Wsu163e** | Unchanged | Q91YN0 | 2 | 3 | 2 | 552 | 63.6 | 6.18 | 9 | #DIV/0! | #DIV/0! | #DIV/0! | #DIV/0! | #DIV/0! | #DIV/0! | #DIV/0! | #DIV/0! |
| 2852 | **Spryd7** | Unchanged | Q3TFQ1 | 1 | 1 | 1 | 196 | 21.7 | 6.7 | 9 | 126.465 | 0.340 | 0.632 | 0.232 | 0.929 | 0.812 | 0.681 | 0.405 |
| 2853 | **Krt79** | Unchanged | Q8VED5 | 5 | 58 | 2 | 531 | 57.5 | 7.69 | 9 | 11.960 | 0.107 | 3.866 | 0.059 | 2.840 | 0.102 | 1.361 | 0.522 |
| 2854 | **Nol3** | Unchanged | Q9D1X0 | 1 | 1 | 1 | 220 | 24.6 | 4.07 | 9 | 0.153 | 0.084 | 0.180 | 0.091 | 1.882 | 0.218 | 0.095 | 0.010 |
| 2855 | **Tor1b** | Unchanged | Q9ER41 | 2 | 3 | 2 | 336 | 37.8 | 8.02 | 9 | 1.126 | 0.762 | 3.358 | 0.065 | 1.578 | 0.072 | 2.128 | 0.145 |
| 2856 | **Wdr82** | Unchanged | Q8BFQ4 | 2 | 8 | 2 | 313 | 35.1 | 7.69 | 9 | 0.727 | 0.398 | 0.798 | 0.654 | 0.886 | 0.682 | 0.901 | 0.831 |
| 2857 | **Grip2** | Unchanged | E0CYS1 | 1 | 1 | 1 | 174 | 19.1 | 10.26 | 9 | 1.417 | 0.661 | 3.235 | 0.220 | 0.768 | 0.682 | 4.211 | 0.171 |
| 2858 | **Rcn1** | Unchanged | Q05186 | 2 | 3 | 2 | 325 | 38.1 | 4.84 | 9 | 1.838 | 0.368 | 2.347 | 0.369 | 0.892 | 0.852 | 2.630 | 0.325 |
| 2859 | **Trp53rkb** | Unchanged | Q543M9 | 1 | 1 | 1 | 244 | 27.4 | 9.23 | 9 | 0.706 | 0.440 | 0.880 | 0.823 | 0.362 | 0.057 | 2.428 | 0.304 |

| | | | | | | | | | | | | | | | | | | |
|---|---|---|---|---|---|---|---|---|---|---|---|---|---|---|---|---|---|---|
| 2860 | **Smad9** | Unchanged | Q9JIW5 | 2 | 2 | 2 | 430 | 48.6 | 8.16 | 9 | 0.279 | 0.110 | 0.625 | 0.508 | 0.641 | 0.473 | 0.976 | 0.973 |
| 2861 | **Scyl2** | UP | Q8CFE4 | 4 | 18 | 4 | 930 | 103.3 | 8.02 | 9 | 0.859 | 0.639 | 1.358 | 0.456 | 2.903 | 0.001 | 0.468 | 0.010 |
| 2862 | **Dctn3** | Unchanged | Q9Z0Y1 | 1 | 4 | 1 | 186 | 21 | 6.06 | 9 | 2.078 | 0.516 | 0.489 | 0.292 | 1.055 | 0.932 | 0.464 | 0.225 |
| 2863 | **Abcf2** | Down | Q99LE6 | 4 | 16 | 4 | 628 | 71.7 | 7.05 | 9 | 0.769 | 0.278 | 0.548 | 0.144 | 0.232 | 0.007 | 2.364 | 0.183 |
| 2864 | **Fndc3a** | Unchanged | Q8BX90 | 5 | 10 | 5 | 1198 | 131.9 | 6.64 | 9 | 1.050 | 0.897 | 0.860 | 0.583 | 0.502 | 0.057 | 1.713 | 0.112 |
| 2865 | **Eng** | Unchanged | Q63961 | 3 | 11 | 3 | 653 | 70 | 6.2 | 9 | 1.192 | 0.705 | 1.094 | 0.875 | 0.699 | 0.457 | 1.566 | 0.415 |
| 2866 | **Ilf3** | Unchanged | Q9Z1X4 | 5 | 31 | 5 | 898 | 96 | 8.76 | 9 | 0.793 | 0.362 | 2.522 | 0.112 | 0.885 | 0.541 | 2.848 | 0.088 |
| 2867 | **Olfr715** | Unchanged | Q8VG49 | 1 | 1 | 1 | 315 | 35.3 | 9.32 | 9 | #DIV/0! | #DIV/0! | #DIV/0! | #DIV/0! | #DIV/0! | #DIV/0! | #DIV/0! | #DIV/0! |
| 2868 | **Ero1b** | Unchanged | Q8R2E9 | 3 | 26 | 3 | 467 | 53.5 | 8.09 | 9 | 2.957 | 0.114 | 2.450 | 0.115 | 3.449 | 0.103 | 0.710 | 0.536 |
| 2869 | **Gpld1** | Down | Q8VCU2 | 3 | 5 | 3 | 842 | 93.6 | 6.8 | 9 | 0.794 | 0.623 | 0.701 | 0.447 | 0.132 | 0.036 | 5.314 | 0.007 |
| 2870 | **Cradd** | UP | O88843 | 1 | 1 | 1 | 199 | 22.6 | 6.21 | 9 | 5.479 | 0.130 | 5.223 | 0.001 | 4.844 | 0.062 | 1.078 | 0.848 |
| 2871 | **Trnt1** | Unchanged | Q8K1J6 | 3 | 19 | 3 | 434 | 49.9 | 8.56 | 9 | 1.458 | 0.586 | 1.068 | 0.884 | 0.713 | 0.366 | 1.499 | 0.384 |
| 2872 | **Gorasp1** | Down | Q91X51 | 1 | 2 | 1 | 446 | 46.9 | 4.65 | 9 | 0.635 | 0.141 | 0.626 | 0.113 | 0.579 | 0.048 | 1.081 | 0.769 |
| 2873 | **Rpa2** | Down | Q62193 | 1 | 6 | 1 | 270 | 29.7 | 6.15 | 9 | 0.584 | 0.045 | 2.318 | 0.239 | 0.599 | 0.005 | 3.867 | 0.137 |
| 2874 | **Dut** | Unchanged | Q8VCG1 | 1 | 1 | 1 | 200 | 21.2 | 9.58 | 9 | #DIV/0! | #DIV/0! | #DIV/0! | #DIV/0! | #DIV/0! | #DIV/0! | #DIV/0! | #DIV/0! |
| 2875 | **Mtnd4** | Unchanged | P03911 | 3 | 39 | 3 | 459 | 51.8 | 9.38 | 8 | 1.602 | 0.516 | 5.976 | 0.073 | 3.023 | 0.126 | 1.977 | 0.296 |
| 2876 | **Cmc1** | Unchanged | Q9CPZ8 | 1 | 6 | 1 | 106 | 12.5 | 8.24 | 8 | 1.213 | 0.627 | 2.378 | 0.172 | 1.029 | 0.952 | 2.310 | 0.191 |
| 2877 | **ATP5IF1** | Unchanged | O35143 | 1 | 5 | 1 | 106 | 12.2 | 9.64 | 8 | 0.705 | 0.608 | 1.439 | 0.626 | 0.815 | 0.734 | 1.765 | 0.473 |
| 2878 | **Pggt1b** | Unchanged | Q8BUY9 | 1 | 2 | 1 | 377 | 42.3 | 6.83 | 8 | 0.562 | 0.648 | 2.513 | 0.411 | 2.378 | 0.289 | 1.057 | 0.939 |
| 2879 | **Mmtag2** | Unchanged | Q99LX5 | 2 | 2 | 2 | 260 | 29.3 | 9.86 | 8 | 127.885 | 0.075 | 3.410 | 0.381 | 124.115 | 0.071 | 0.027 | 0.076 |
| 2880 | **Srrm1** | Down | Q52KI8 | 4 | 15 | 4 | 946 | 106.8 | 11.87 | 8 | 0.932 | 0.870 | 1.974 | 0.137 | 0.191 | 0.000 | 10.340 | 0.016 |
| 2881 | **Snrpb2** | Unchanged | Q9CQI7 | 1 | 9 | 1 | 225 | 25.3 | 9.72 | 8 | 1.963 | 0.509 | 4.854 | 0.156 | 3.604 | 0.089 | 1.347 | 0.667 |
| 2882 | **Plec** | Unchanged | Q9QXS1 | 24 | 95 | 23 | 4691 | 533.9 | 5.96 | 8 | 1.067 | 0.662 | 1.268 | 0.478 | 1.133 | 0.274 | 1.120 | 0.722 |
| 2883 | **Arl13b** | Unchanged | Q640N2 | 1 | 1 | 1 | 427 | 48.1 | 6.32 | 8 | 0.484 | 0.459 | 0.576 | 0.542 | 1.585 | 0.491 | 0.363 | 0.084 |
| 2884 | **Selenos** | Unchanged | Q9BCZ4 | 1 | 2 | 1 | 190 | 21.5 | 9.39 | 8 | 1.815 | 0.386 | 1.500 | 0.727 | 0.780 | 0.737 | 1.923 | 0.609 |
| 2885 | **Tmem11** | Unchanged | Q8BK08 | 1 | 4 | 1 | 190 | 21.3 | 7.36 | 8 | 1.389 | 0.651 | 2.036 | 0.348 | 0.213 | 0.138 | 9.554 | 0.085 |
| 2886 | **Fn3k** | Unchanged | Q9ER35 | 2 | 7 | 2 | 309 | 35 | 8.4 | 8 | 0.262 | 0.126 | 0.174 | 0.086 | 0.155 | 0.075 | 1.127 | 0.887 |
| 2887 | **Gosr1** | Unchanged | O88630 | 2 | 13 | 2 | 250 | 28.5 | 9.29 | 8 | 2.862 | 0.192 | 2.988 | 0.154 | 3.937 | 0.151 | 0.759 | 0.677 |
| 2888 | **Atpaf2** | Unchanged | A0A0R4J1C5 | 1 | 1 | 1 | 298 | 34.3 | 6.8 | 8 | 0.962 | 0.952 | 5.521 | 0.090 | 1.682 | 0.266 | 3.282 | 0.142 |
| 2889 | **Ogn** | Unchanged | Q62000 | 2 | 8 | 2 | 298 | 34 | 5.74 | 8 | 0.268 | 0.084 | 0.535 | 0.247 | 0.960 | 0.943 | 0.557 | 0.322 |
| 2890 | **Olfr444** | Unchanged | Q8VFS6 | 1 | 1 | 1 | 310 | 35 | 8.43 | 8 | 1.054 | 0.921 | 0.872 | 0.734 | 0.532 | 0.201 | 1.639 | 0.192 |
| 2891 | **Tm6sf1** | Unchanged | P58749 | 1 | 1 | 1 | 370 | 41.6 | 7.64 | 8 | 1.048 | 0.884 | 0.498 | 0.132 | 1.682 | 0.151 | 0.296 | 0.027 |
| 2892 | **Tbc1d24** | Unchanged | Q3UUG6 | 3 | 5 | 3 | 561 | 63.2 | 7.24 | 8 | 0.830 | 0.758 | 0.920 | 0.875 | 0.458 | 0.285 | 2.009 | 0.053 |
| 2893 | **Tsr2** | UP | Q8C8T8 | 1 | 23 | 1 | 191 | 20.9 | 4.23 | 8 | 1.989 | 0.270 | 1.267 | 0.678 | 3.147 | 0.048 | 0.403 | 0.096 |
| 2894 | **Cavin2** | Unchanged | Q63918 | 2 | 4 | 2 | 418 | 46.7 | 5.21 | 8 | 1.040 | 0.910 | 0.589 | 0.275 | 0.559 | 0.195 | 1.054 | 0.882 |
| 2895 | **Dalrd3** | Down | Q6PJN8 | 2 | 3 | 2 | 538 | 58.7 | 8.15 | 8 | 0.514 | 0.106 | 0.561 | 0.189 | 0.300 | 0.026 | 1.868 | 0.206 |
| 2896 | **Gsdma2** | Unchanged | Q32M21 | 2 | 4 | 2 | 443 | 49.8 | 5.9 | 7 | 1.088 | 0.904 | 0.350 | 0.267 | 0.074 | 0.116 | 4.735 | 0.112 |
| 2897 | **Pmvk** | Unchanged | Q9D1G2 | 1 | 2 | 1 | 192 | 21.9 | 5.99 | 8 | 0.579 | 0.264 | 0.395 | 0.100 | 0.551 | 0.214 | 0.717 | 0.219 |
| 2898 | **Atp1a4** | Unchanged | Q9WV27 | 7 | 117 | 1 | 1032 | 114.8 | 5.71 | 8 | 0.828 | 0.779 | 0.285 | 0.168 | 0.237 | 0.131 | 1.202 | 0.756 |
| 2899 | **Cdkn2a** | Unchanged | P51480 | 1 | 1 | 1 | 168 | 17.9 | 6.52 | 8 | #DIV/0! | #DIV/0! | #DIV/0! | #DIV/0! | #DIV/0! | #DIV/0! | #DIV/0! | #DIV/0! |
| 2900 | **Krt7** | Unchanged | Q9DCV7 | 4 | 20 | 2 | 457 | 50.7 | 5.87 | 8 | 2.893 | 0.274 | 2.465 | 0.219 | 1.691 | 0.144 | 1.458 | 0.495 |

| | | | | | | | | | | | | | | | | | | |
|---|---|---|---|---|---|---|---|---|---|---|---|---|---|---|---|---|---|---|
| 2901 | **Pip4k2c** | Unchanged | Q91XU3 | 1 | 2 | 1 | 421 | 47.3 | 6.89 | 8 | 0.882 | 0.603 | 0.841 | 0.406 | 1.043 | 0.864 | 0.807 | 0.402 |
| 2902 | **Gnpda1** | Unchanged | O88958 | 2 | 4 | 2 | 289 | 32.5 | 6.6 | 8 | 66.566 | 0.141 | 57.171 | 0.172 | 138.395 | 0.056 | 0.413 | 0.292 |
| 2903 | **C1sa** | Unchanged | Q8CG14 | 3 | 4 | 3 | 688 | 76.8 | 5.08 | 8 | 0.739 | 0.381 | 2.341 | 0.310 | 7.232 | 0.194 | 0.324 | 0.314 |
| 2904 | **Zfp839** | Unchanged | A0A1Y7VL21 | 1 | 1 | 1 | 145 | 15.7 | 10.1 | 8 | 4.309 | 0.188 | 3.419 | 0.319 | 2.879 | 0.260 | 1.188 | 0.845 |
| 2905 | **Synj2bp** | Unchanged | Q9D6K5 | 1 | 6 | 1 | 145 | 15.8 | 6.3 | 8 | #DIV/0! | #DIV/0! | #DIV/0! | #DIV/0! | #DIV/0! | #DIV/0! | #DIV/0! | #DIV/0! |
| 2906 | **Pex6** | Unchanged | Q99LC9 | 4 | 12 | 4 | 981 | 104.5 | 7.34 | 8 | 1.192 | 0.539 | 4.438 | 0.119 | 2.215 | 0.148 | 2.003 | 0.319 |
| 2907 | **Ttc37** | Unchanged | F8VPK0 | 7 | 12 | 7 | 1563 | 173.8 | 7.12 | 8 | 0.658 | 0.501 | 0.838 | 0.732 | 0.518 | 0.307 | 1.619 | 0.124 |
| 2908 | **Ppp3ca** | Unchanged | P63328 | 2 | 7 | 2 | 521 | 58.6 | 5.86 | 8 | 0.864 | 0.777 | 0.754 | 0.643 | 2.023 | 0.335 | 0.373 | 0.229 |
| 2909 | **Dkc1** | Unchanged | Q9ESX5 | 3 | 23 | 3 | 509 | 57.4 | 9.28 | 8 | 1.566 | 0.444 | 1.866 | 0.287 | 1.554 | 0.360 | 1.201 | 0.731 |
| 2910 | **Sec16a** | Down | E9QAT4 | 10 | 30 | 10 | 2357 | 254 | 5.81 | 8 | 0.319 | 0.003 | 2.084 | 0.336 | 0.344 | 0.006 | 6.062 | 0.137 |
| 2911 | **Cyp20a1** | Unchanged | Q8BKE6 | 2 | 2 | 2 | 462 | 52.1 | 6.95 | 8 | 2.617 | 0.264 | 5.131 | 0.111 | 2.308 | 0.364 | 2.223 | 0.315 |
| 2912 | **Sh3glb1** | UP | Q9JK48 | 1 | 3 | 1 | 365 | 40.8 | 6.04 | 8 | 1.492 | 0.212 | 2.587 | 0.043 | 0.667 | 0.309 | 3.880 | 0.019 |
| 2913 | **Rrm1** | Unchanged | P07742 | 3 | 9 | 3 | 792 | 90.2 | 6.7 | 8 | 1.287 | 0.652 | 1.280 | 0.722 | 1.612 | 0.315 | 0.794 | 0.587 |
| 2914 | **Cnbpiso** | Unchanged | A0A0N4SVS6 | 1 | 6 | 1 | 171 | 18.8 | 7.71 | 8 | 0.698 | 0.649 | 0.295 | 0.253 | 0.600 | 0.518 | 0.491 | 0.222 |
| 2915 | **Arhgef3** | Unchanged | A0A286YDE6 | 2 | 2 | 2 | 551 | 63 | 6.8 | 8 | 2.011 | 0.085 | 1.374 | 0.517 | 0.583 | 0.140 | 2.356 | 0.169 |
| 2916 | **Tmem120a** | Unchanged | Q8C1E7 | 2 | 6 | 2 | 343 | 40.7 | 8.88 | 8 | 0.256 | 0.121 | 0.326 | 0.213 | 0.446 | 0.228 | 0.732 | 0.716 |
| 2917 | **Olfr178** | Unchanged | E9Q9T3 | 1 | 2 | 1 | 319 | 36.5 | 8.34 | 8 | 0.431 | 0.400 | 0.089 | 0.166 | 0.633 | 0.579 | 0.140 | 0.037 |
| 2918 | **Stx5a** | Down | I1E4X1 | 2 | 2 | 2 | 442 | 49.1 | 8.98 | 8 | 0.860 | 0.717 | 2.576 | 0.253 | 0.197 | 0.005 | 13.082 | 0.097 |
| 2919 | **Kcmf1** | Unchanged | Q80UY2 | 2 | 6 | 2 | 381 | 41.8 | 5.76 | 8 | 1.125 | 0.695 | 2.144 | 0.097 | 0.639 | 0.154 | 3.357 | 0.031 |
| 2920 | **Ldhc** | Unchanged | P00342 | 3 | 39 | 1 | 332 | 35.9 | 8.21 | 8 | 0.855 | 0.783 | 0.543 | 0.369 | 0.166 | 0.075 | 3.261 | 0.197 |
| 2921 | **Try5** | Unchanged | Q9QUK9 | 1 | 5 | 1 | 246 | 26.3 | 5.3 | 8 | 3.829 | 0.412 | 0.711 | 0.654 | 1.086 | 0.911 | 0.655 | 0.611 |
| 2922 | **Prss1** | Unchanged | Q9Z1R9 | 1 | 21 | 1 | 246 | 26.1 | 4.94 | 8 | 0.747 | 0.557 | 0.805 | 0.657 | 0.866 | 0.762 | 0.929 | 0.869 |
| 2923 | **Try10** | Unchanged | Q792Z1 | 2 | 12 | 2 | 246 | 26.2 | 5.83 | 8 | 0.807 | 0.732 | 0.884 | 0.843 | 0.952 | 0.930 | 0.928 | 0.803 |
| 2924 | **Spg21** | Unchanged | Q9CQC8 | 1 | 1 | 1 | 308 | 34.9 | 6.28 | 8 | 3.265 | 0.288 | 6.375 | 0.156 | 0.834 | 0.602 | 7.642 | 0.145 |
| 2925 | **Pawr** | Unchanged | Q925B0 | 2 | 5 | 2 | 333 | 35.9 | 5.92 | 8 | 1.271 | 0.649 | 0.653 | 0.519 | 0.756 | 0.510 | 0.864 | 0.866 |
| 2926 | **Rabac1** | Unchanged | Q9Z0S9 | 1 | 2 | 1 | 185 | 20.6 | 7.9 | 8 | 1.136 | 0.832 | 1.266 | 0.736 | 2.247 | 0.571 | 0.563 | 0.665 |
| 2927 | **C4bpa** | Unchanged | P08607 | 3 | 15 | 3 | 469 | 51.5 | 7.15 | 8 | 0.411 | 0.151 | 0.263 | 0.079 | 0.316 | 0.091 | 0.833 | 0.594 |
| 2928 | **Ptpn1** | Down | P35821 | 2 | 4 | 2 | 432 | 49.6 | 6.16 | 8 | 0.577 | 0.297 | 0.197 | 0.023 | 0.019 | 0.006 | 10.363 | 0.146 |
| 2929 | **Polr2e** | UP | Q80UW8 | 1 | 2 | 1 | 210 | 24.6 | 5.95 | 8 | 6.742 | 0.017 | 10.282 | 0.015 | 2.983 | 0.373 | 3.446 | 0.082 |
| 2930 | **Fn3krp** | Down | Q8K274 | 2 | 15 | 2 | 309 | 34.4 | 7.87 | 8 | 0.458 | 0.082 | 0.496 | 0.121 | 0.225 | 0.002 | 2.202 | 0.349 |
| 2931 | **Gm8909** | Unchanged | G3UXE9 | 3 | 11 | 1 | 396 | 44.6 | 6.29 | 8 | 0.407 | 0.354 | 0.596 | 0.536 | 27.499 | 0.112 | 0.022 | 0.108 |
| 2932 | **Ccdc127** | Unchanged | Q3TC33 | 1 | 1 | 1 | 260 | 30.5 | 9.13 | 8 | 2.720 | 0.132 | 1.038 | 0.935 | 1.783 | 0.326 | 0.582 | 0.384 |
| 2933 | **Spcs1** | Unchanged | Q9D958 | 1 | 8 | 1 | 161 | 18.2 | 10.01 | 8 | 0.813 | 0.786 | 0.627 | 0.584 | 0.060 | 0.109 | 10.387 | 0.190 |
| 2934 | **Clpb** | UP | E9PY58 | 3 | 4 | 3 | 707 | 79.5 | 8.56 | 8 | 5.397 | 0.015 | 11.221 | 0.023 | 4.145 | 0.283 | 2.707 | 0.158 |
| 2935 | **Cldn34b4** | Unchanged | Q3V0X2 | 1 | 1 | 1 | 211 | 23.8 | 8.48 | 8 | 1.372 | 0.145 | 1.511 | 0.091 | 6.898 | 0.127 | 0.219 | 0.159 |
| 2936 | **Ostc** | Unchanged | Q78XF5 | 1 | 14 | 1 | 149 | 16.8 | 9.13 | 8 | 1.397 | 0.659 | 1.875 | 0.476 | 1.374 | 0.711 | 1.364 | 0.705 |
| 2937 | **Aup1** | Down | P70295 | 2 | 7 | 2 | 410 | 46.1 | 8.4 | 8 | 0.612 | 0.190 | 0.452 | 0.044 | 0.430 | 0.033 | 1.052 | 0.850 |
| 2938 | **Scfd2** | Unchanged | Q8BTY8 | 3 | 8 | 3 | 684 | 74.7 | 6.81 | 8 | 0.755 | 0.563 | 1.022 | 0.967 | 0.376 | 0.109 | 2.721 | 0.138 |
| 2939 | **Sptlc1** | Unchanged | O35704 | 1 | 1 | 1 | 473 | 52.5 | 6.4 | 8 | 0.508 | 0.102 | 1.419 | 0.529 | 0.826 | 0.469 | 1.718 | 0.386 |
| 2940 | **Gga1** | Unchanged | Q8R0H9 | 2 | 11 | 2 | 635 | 69.9 | 5.27 | 8 | 1.040 | 0.870 | 1.129 | 0.802 | 0.906 | 0.646 | 1.246 | 0.670 |
| 2941 | **Cisd3** | Unchanged | B1AR13 | 1 | 3 | 1 | 137 | 15.7 | 9.79 | 8 | 1.020 | 0.981 | 0.771 | 0.774 | 0.107 | 0.172 | 7.230 | 0.217 |

| | | | | | | | | | | | | | | | | | | |
|---|---|---|---|---|---|---|---|---|---|---|---|---|---|---|---|---|---|---|
| 2942 | **Kdm1a** | Unchanged | A3KG93 | 2 | 2 | 2 | 873 | 94.6 | 6.25 | 8 | 1.306 | 0.796 | 0.525 | 0.367 | 0.681 | 0.571 | 0.771 | 0.672 |
| 2943 | **Vmn1r60** | Unchanged | G3UY47 | 1 | 1 | 1 | 300 | 34.2 | 8.78 | 8 | 0.554 | 0.585 | 0.657 | 0.682 | 0.296 | 0.384 | 2.218 | 0.262 |
| 2944 | **Scamp1** | Unchanged | Q8K021 | 2 | 3 | 2 | 338 | 38 | 7.71 | 8 | 1.107 | 0.880 | 0.679 | 0.614 | 0.584 | 0.509 | 1.164 | 0.543 |
| 2945 | **Tra2b** | Unchanged | P62996 | 2 | 12 | 2 | 288 | 33.6 | 11.25 | 8 | 1.910 | 0.237 | 0.685 | 0.438 | 2.062 | 0.559 | 0.332 | 0.453 |
| 2946 | **Tmco1** | Unchanged | Q921L3 | 1 | 8 | 1 | 188 | 21.2 | 9.74 | 8 | 2.136 | 0.384 | 4.344 | 0.278 | 0.549 | 0.414 | 7.916 | 0.216 |
| 2947 | **Arfgap1** | Unchanged | Q9EPJ9 | 2 | 5 | 2 | 414 | 45.3 | 5.57 | 8 | 0.963 | 0.944 | 0.572 | 0.389 | 0.488 | 0.301 | 1.173 | 0.674 |
| 2948 | **Gak** | Unchanged | Q99KY4 | 6 | 7 | 6 | 1305 | 143.6 | 5.73 | 8 | 2.656 | 0.061 | 0.924 | 0.788 | 1.459 | 0.550 | 0.633 | 0.495 |
| 2949 | **Mrpl17** | Unchanged | Q9D8P4 | 1 | 1 | 1 | 176 | 20.2 | 9.82 | 8 | 1.573 | 0.469 | 1.935 | 0.221 | 0.966 | 0.892 | 2.004 | 0.196 |
| 2950 | **Mier2** | Unchanged | Q9CZI4 | 1 | 5 | 1 | 264 | 27.9 | 9.5 | 8 | 1.129 | 0.830 | 1.304 | 0.645 | 0.336 | 0.211 | 3.887 | 0.066 |
| 2951 | **Herc4** | Unchanged | Q6PAV2 | 4 | 7 | 4 | 1057 | 118.3 | 6.29 | 8 | 1.566 | 0.601 | 0.408 | 0.187 | 0.566 | 0.345 | 0.720 | 0.356 |
| 2952 | **Sphk2** | Unchanged | Q9JIA7 | 3 | 8 | 3 | 617 | 65.6 | 6.57 | 8 | 1.177 | 0.730 | 0.999 | 0.999 | 1.060 | 0.921 | 0.943 | 0.915 |
| 2953 | **Rdh10** | Unchanged | Q8VCH7 | 2 | 3 | 2 | 341 | 38.1 | 7.59 | 8 | 1.640 | 0.290 | 2.430 | 0.214 | 0.948 | 0.883 | 2.564 | 0.180 |
| 2954 | **Tial1** | Unchanged | P70318 | 2 | 11 | 2 | 392 | 43.4 | 7.99 | 8 | 0.439 | 0.412 | 2.136 | 0.292 | 0.286 | 0.293 | 7.481 | 0.047 |
| 2955 | **Nccrp1** | Unchanged | A0A0A0MQ97 | 1 | 1 | 1 | 291 | 33 | 6.89 | 8 | 1.450 | 0.307 | 1.325 | 0.479 | 0.953 | 0.928 | 1.391 | 0.410 |
| 2956 | **Fkbp5** | Unchanged | Q64378 | 2 | 7 | 2 | 456 | 50.9 | 7.8 | 8 | 1.249 | 0.755 | 3.240 | 0.147 | 0.057 | 0.148 | 56.714 | 0.036 |
| 2957 | **Phospho2** | Unchanged | Q9D9M5 | 1 | 1 | 1 | 241 | 27.5 | 6.54 | 8 | #DIV/0! | #DIV/0! | #DIV/0! | #DIV/0! | #DIV/0! | #DIV/0! | #DIV/0! | #DIV/0! |
| 2958 | **Smpd2** | Unchanged | O70572 | 3 | 17 | 3 | 419 | 47.4 | 6.44 | 8 | 1.081 | 0.793 | 1.271 | 0.525 | 0.924 | 0.824 | 1.375 | 0.443 |
| 2959 | **Prkaa1** | Unchanged | Q5EG47 | 3 | 31 | 1 | 559 | 63.9 | 8.12 | 8 | 1.201 | 0.793 | 2.572 | 0.342 | 0.335 | 0.281 | 7.687 | 0.162 |
| 2960 | **Ube2z** | Unchanged | Q3UE37 | 2 | 3 | 2 | 356 | 38.3 | 5.62 | 8 | 1.078 | 0.889 | 1.560 | 0.480 | 0.995 | 0.993 | 1.567 | 0.530 |
| 2961 | **Ythdf3** | Unchanged | Q8BYK6 | 4 | 10 | 4 | 585 | 63.9 | 9.04 | 8 | 0.436 | 0.252 | 0.578 | 0.382 | 0.223 | 0.084 | 2.595 | 0.198 |
| 2962 | **Msr1** | Down | P30204 | 2 | 5 | 2 | 458 | 50.1 | 6.54 | 8 | 0.206 | 0.030 | 0.050 | 0.012 | 0.015 | 0.010 | 3.333 | 0.263 |
| 2963 | **Ahsa2** | Unchanged | Q8N9S3 | 1 | 1 | 1 | 331 | 37.6 | 6.44 | 8 | 0.845 | 0.641 | 1.512 | 0.192 | 0.965 | 0.904 | 1.567 | 0.147 |
| 2964 | **Fam98c** | Down | E9PYD1 | 2 | 2 | 2 | 344 | 37.2 | 8.62 | 8 | 0.358 | 0.004 | 1.166 | 0.804 | 1.130 | 0.780 | 1.032 | 0.963 |
| 2965 | **Fam114a2** | UP | Q8VE88 | 2 | 8 | 2 | 497 | 54 | 4.93 | 8 | 4.468 | 0.128 | 7.928 | 0.035 | 2.491 | 0.146 | 3.183 | 0.096 |
| 2966 | **Nudt4** | Unchanged | Q8R2U6 | 1 | 8 | 1 | 179 | 20.1 | 6.35 | 8 | 0.839 | 0.745 | 1.177 | 0.766 | 0.591 | 0.323 | 1.990 | 0.277 |
| 2967 | **Ddx23** | Unchanged | D3Z0M9 | 5 | 9 | 5 | 819 | 95.4 | 9.58 | 8 | 0.497 | 0.235 | 1.181 | 0.735 | 0.683 | 0.452 | 1.730 | 0.223 |
| 2968 | **Dpep1** | Unchanged | P31428 | 1 | 1 | 1 | 410 | 45.7 | 6.37 | 8 | 0.659 | 0.309 | 1.305 | 0.551 | 4.433 | 0.131 | 0.294 | 0.169 |
| 2969 | **Gar1** | Unchanged | Q9CY66 | 1 | 1 | 1 | 231 | 23.5 | 11.02 | 8 | 0.852 | 0.473 | 0.658 | 0.200 | 4.293 | 0.064 | 0.153 | 0.046 |
| 2970 | **Zcchc7** | Unchanged | F7CLJ4 | 1 | 4 | 1 | 77 | 9.1 | 9.19 | 8 | 0.695 | 0.605 | 0.170 | 0.120 | 0.657 | 0.538 | 0.260 | 0.157 |
| 2971 | **Isca2** | Unchanged | Q9DCB8 | 1 | 3 | 1 | 154 | 16.7 | 5.6 | 8 | 1.255 | 0.694 | 2.764 | 0.278 | 0.779 | 0.595 | 3.548 | 0.228 |
| 2972 | **Ap3s1** | Down | Q9DCR2 | 1 | 8 | 1 | 193 | 21.7 | 5.39 | 8 | 0.630 | 0.363 | 0.240 | 0.029 | 0.315 | 0.060 | 0.762 | 0.713 |
| 2973 | **Sra1** | Unchanged | Q80VJ2 | 1 | 1 | 1 | 232 | 25.5 | 6.43 | 8 | 0.467 | 0.409 | 0.288 | 0.261 | 1.624 | 0.556 | 0.177 | 0.158 |
| 2974 | **Skint2** | Unchanged | A7XUX6 | 1 | 1 | 1 | 400 | 45.6 | 6.93 | 8 | 1.656 | 0.560 | 3.537 | 0.114 | 1.323 | 0.418 | 2.674 | 0.164 |
| 2975 | **Tiprl** | Unchanged | Q8BH58 | 1 | 5 | 1 | 271 | 31.2 | 5.63 | 8 | 1.155 | 0.664 | 2.355 | 0.309 | 2.126 | 0.188 | 1.108 | 0.875 |
| 2976 | **Zbtb25** | Unchanged | G3UW50 | 1 | 1 | 1 | 439 | 49.2 | 6.74 | 8 | #DIV/0! | #DIV/0! | #DIV/0! | #DIV/0! | #DIV/0! | #DIV/0! | #DIV/0! | #DIV/0! |
| 2977 | **Fgf1** | Unchanged | P61148 | 1 | 6 | 1 | 155 | 17.4 | 7.02 | 8 | 1.126 | 0.709 | 4.190 | 0.246 | 1.693 | 0.130 | 2.475 | 0.357 |
| 2978 | **Mrpl39** | Unchanged | Q9JKF7 | 2 | 5 | 2 | 336 | 38.5 | 7.94 | 8 | 0.400 | 0.354 | 2.007 | 0.519 | 1.157 | 0.853 | 1.735 | 0.583 |
| 2979 | **Afg1l** | Unchanged | Q3V384 | 2 | 3 | 2 | 480 | 54.3 | 8.03 | 8 | 0.943 | 0.825 | 1.656 | 0.192 | 1.080 | 0.735 | 1.534 | 0.233 |
| 2980 | **Ankfy1** | Unchanged | Q810B6 | 5 | 14 | 5 | 1169 | 128.6 | 5.91 | 8 | 1.252 | 0.636 | 1.340 | 0.521 | 1.139 | 0.760 | 1.176 | 0.735 |
| 2981 | **Rex1bd** | Unchanged | Q9CYZ6 | 1 | 2 | 1 | 169 | 18.7 | 5.81 | 8 | 1.291 | 0.562 | 1.950 | 0.314 | 1.961 | 0.189 | 0.994 | 0.992 |
| 2982 | **Srsf2** | Unchanged | Q62093 | 1 | 44 | 1 | 221 | 25.5 | 11.85 | 8 | 1.183 | 0.720 | 1.268 | 0.710 | 0.851 | 0.682 | 1.490 | 0.547 |

| | | | | | | | | | | | | | | | | | | |
|---|---|---|---|---|---|---|---|---|---|---|---|---|---|---|---|---|---|---|
| 2983 | **Gm6970** | Unchanged | F7DCP6 | 1 | 1 | 1 | 221 | 22.8 | 10.59 | 8 | 3.874 | 0.252 | 4.950 | 0.216 | 7.138 | 0.105 | 0.693 | 0.631 |
| 2984 | **Phgdh** | Unchanged | Q61753 | 1 | 1 | 1 | 533 | 56.5 | 6.54 | 8 | #DIV/0! | #DIV/0! | #DIV/0! | #DIV/0! | #DIV/0! | #DIV/0! | #DIV/0! | #DIV/0! |
| 2985 | **Spg7** | Unchanged | Q3ULF4 | 3 | 15 | 3 | 781 | 85.9 | 9.04 | 8 | 1.331 | 0.568 | 1.765 | 0.281 | 1.215 | 0.724 | 1.453 | 0.487 |
| 2986 | **P2ry10b** | Unchanged | Q8BY68 | 1 | 1 | 1 | 352 | 40.4 | 8.46 | 8 | #DIV/0! | #DIV/0! | #DIV/0! | #DIV/0! | #DIV/0! | #DIV/0! | #DIV/0! | #DIV/0! |
| 2987 | **Dtd1** | Unchanged | Q9DD18 | 1 | 4 | 1 | 209 | 23.4 | 7.87 | 8 | #DIV/0! | #DIV/0! | #DIV/0! | #DIV/0! | #DIV/0! | #DIV/0! | #DIV/0! | #DIV/0! |
| 2988 | **Gnrhr** | Unchanged | Q01776 | 1 | 1 | 1 | 327 | 37.7 | 9.55 | 8 | #DIV/0! | #DIV/0! | #DIV/0! | #DIV/0! | #DIV/0! | #DIV/0! | #DIV/0! | #DIV/0! |
| 2989 | **Hpn** | Unchanged | E9Q5P0 | 2 | 2 | 2 | 445 | 47.7 | 7.28 | 8 | 0.504 | 0.340 | 0.066 | 0.078 | 0.351 | 0.212 | 0.188 | 0.060 |
| 2990 | **Phkg2** | Unchanged | Q9DB30 | 2 | 3 | 2 | 406 | 46.5 | 6.48 | 8 | 1.015 | 0.974 | 1.584 | 0.412 | 0.951 | 0.931 | 1.667 | 0.414 |
| 2991 | **Fam107b** | Unchanged | Q3TGF2 | 1 | 2 | 1 | 131 | 15.6 | 8.31 | 8 | 1.904 | 0.340 | 0.840 | 0.484 | 1.612 | 0.427 | 0.521 | 0.324 |
| 2992 | **Ppt2** | UP | O35448 | 2 | 11 | 2 | 302 | 34.3 | 6.39 | 8 | 2.691 | 0.140 | 5.292 | 0.053 | 5.626 | 0.007 | 0.941 | 0.886 |
| 2993 | **Camk2d** | Unchanged | Q6PHZ2 | 2 | 7 | 2 | 499 | 56.3 | 7.25 | 8 | 1.610 | 0.280 | 2.698 | 0.121 | 0.734 | 0.345 | 3.673 | 0.077 |
| 2994 | **Rexo2** | UP | Q9D8S4 | 1 | 9 | 1 | 237 | 26.7 | 7.15 | 8 | 1.352 | 0.494 | 5.278 | 0.059 | 5.803 | 0.015 | 0.910 | 0.836 |
| 2995 | **Ptgis** | Down | O35074 | 3 | 8 | 3 | 501 | 57 | 6.73 | 8 | 0.996 | 0.995 | 0.972 | 0.929 | 0.415 | 0.018 | 2.342 | 0.063 |
| 2996 | **Amdhd2** | Unchanged | Q8JZV7 | 1 | 2 | 1 | 409 | 43.5 | 6.23 | 8 | 0.538 | 0.368 | 0.752 | 0.612 | 0.394 | 0.172 | 1.909 | 0.336 |
| 2997 | **Ssr3** | Unchanged | Q9DCF9 | 1 | 40 | 1 | 185 | 21.1 | 9.61 | 8 | 0.753 | 0.377 | 0.683 | 0.285 | 0.751 | 0.410 | 0.910 | 0.680 |
| 2998 | **Fam45a** | Unchanged | Q9D8N2 | 1 | 3 | 1 | 357 | 40.4 | 6.47 | 8 | 1.322 | 0.651 | 1.591 | 0.617 | 0.536 | 0.135 | 2.968 | 0.375 |
| 2999 | **Rmnd1** | Unchanged | Q8CI78 | 2 | 6 | 2 | 450 | 51.8 | 7.99 | 8 | 0.663 | 0.287 | 0.950 | 0.852 | 0.663 | 0.118 | 1.433 | 0.222 |
| 3000 | **Eif2d** | Down | Q61211 | 2 | 3 | 2 | 570 | 62.8 | 8.24 | 8 | 1.013 | 0.974 | 1.273 | 0.695 | 0.343 | 0.050 | 3.707 | 0.168 |
| 3001 | **Mtnd2** | UP | P03893 | 1 | 14 | 1 | 345 | 38.7 | 9.92 | 8 | 6.858 | 0.043 | 18.156 | 0.006 | 13.430 | 0.001 | 1.352 | 0.403 |
| 3002 | **Fscn1** | Unchanged | Q61553 | 2 | 7 | 2 | 493 | 54.5 | 6.89 | 8 | 1.932 | 0.482 | 2.335 | 0.190 | 1.694 | 0.228 | 1.379 | 0.547 |
| 3003 | **Epn1** | Unchanged | Q80VP1 | 3 | 13 | 3 | 575 | 60.2 | 4.81 | 7 | 1.414 | 0.175 | 1.629 | 0.069 | 0.970 | 0.916 | 1.679 | 0.124 |
| 3004 | **Aldh1l2** | Unchanged | Q8K009 | 7 | 100 | 1 | 923 | 101.5 | 6.29 | 7 | 1.895 | 0.630 | 15.631 | 0.145 | 0.449 | 0.541 | 34.831 | 0.131 |
| 3005 | **Mea1** | Unchanged | Q64327 | 1 | 1 | 1 | 174 | 18.6 | 4.08 | 7 | #DIV/0! | #DIV/0! | #DIV/0! | #DIV/0! | #DIV/0! | #DIV/0! | #DIV/0! | #DIV/0! |
| 3006 | **Fam129a** | Unchanged | Q3UW53 | 3 | 8 | 3 | 926 | 102.6 | 4.78 | 7 | 1.872 | 0.578 | 2.316 | 0.428 | 3.548 | 0.197 | 0.653 | 0.547 |
| 3007 | **Cpsf6** | Unchanged | Q6NVF9 | 2 | 35 | 2 | 551 | 59.1 | 7.15 | 7 | 0.445 | 0.171 | 2.886 | 0.211 | 0.366 | 0.120 | 7.881 | 0.097 |
| 3008 | **Qrsl1** | Unchanged | Q9CZN8 | 1 | 2 | 1 | 525 | 56.7 | 5.95 | 7 | 0.850 | 0.627 | 1.209 | 0.613 | 1.210 | 0.510 | 0.999 | 0.998 |
| 3009 | **Slc35a3** | Unchanged | E9Q3V2 | 1 | 2 | 1 | 202 | 21.7 | 11.47 | 7 | 2.173 | 0.402 | 1.289 | 0.767 | 0.325 | 0.262 | 3.962 | 0.250 |
| 3010 | **Abhd10** | Unchanged | Q6PE15 | 1 | 2 | 1 | 297 | 33 | 8.79 | 7 | 1.545 | 0.461 | 0.969 | 0.964 | 1.068 | 0.898 | 0.907 | 0.882 |
| 3011 | **Parvb** | Unchanged | Q9ES46 | 2 | 6 | 1 | 365 | 41.6 | 6.33 | 7 | 3.838 | 0.101 | 5.933 | 0.063 | 3.537 | 0.143 | 1.677 | 0.407 |
| 3012 | **Vmp1** | Unchanged | Q99KU0 | 1 | 24 | 1 | 406 | 45.9 | 6.95 | 7 | 1.430 | 0.642 | 0.916 | 0.921 | 1.822 | 0.358 | 0.503 | 0.139 |
| 3013 | **Ccdc51** | Unchanged | Q3URS9 | 2 | 8 | 2 | 406 | 45.1 | 8.09 | 7 | 0.208 | 0.099 | 2.107 | 0.488 | 0.561 | 0.408 | 3.757 | 0.330 |
| 3014 | **Stat5b** | Unchanged | P42232 | 4 | 15 | 4 | 786 | 89.9 | 6.05 | 7 | 0.673 | 0.306 | 1.650 | 0.137 | 0.711 | 0.513 | 2.319 | 0.119 |
| 3015 | **Saa2** | Unchanged | P05367 | 1 | 4 | 1 | 122 | 13.6 | 6.93 | 7 | 0.083 | 0.316 | 0.094 | 0.322 | 0.086 | 0.318 | 1.094 | 0.794 |
| 3016 | **Ube4a** | Unchanged | G3X9Y5 | 4 | 8 | 4 | 1085 | 124.4 | 5.19 | 7 | 0.646 | 0.395 | 0.991 | 0.984 | 0.776 | 0.664 | 1.277 | 0.637 |
| 3017 | **Olfr845** | Unchanged | Q7TRG2 | 1 | 1 | 1 | 312 | 34.6 | 6.95 | 7 | 0.687 | 0.332 | 0.437 | 0.164 | 0.724 | 0.478 | 0.604 | 0.497 |
| 3018 | **Tnni2** | Down | A2A6J8 | 1 | 1 | 1 | 190 | 22.2 | 8.95 | 7 | 1.099 | 0.894 | 0.634 | 0.460 | 0.067 | 0.023 | 9.473 | 0.131 |
| 3019 | **Ncstn** | UP | P57716 | 3 | 15 | 3 | 708 | 78.4 | 6.09 | 7 | 5.370 | 0.049 | 10.367 | 0.088 | 6.121 | 0.010 | 1.694 | 0.425 |
| 3020 | **Tppp** | Unchanged | Q7TQD2 | 1 | 2 | 1 | 218 | 23.6 | 9.42 | 7 | #DIV/0! | #DIV/0! | #DIV/0! | #DIV/0! | #DIV/0! | #DIV/0! | #DIV/0! | #DIV/0! |
| 3021 | **Bco2** | Unchanged | Q99NF1 | 2 | 7 | 2 | 532 | 60.1 | 6.54 | 7 | 0.903 | 0.716 | 1.406 | 0.335 | 0.843 | 0.643 | 1.667 | 0.172 |
| 3022 | **Psmb10** | Unchanged | O35955 | 1 | 12 | 1 | 273 | 29 | 6.87 | 7 | 3.769 | 0.203 | 10.890 | 0.095 | 2.007 | 0.337 | 5.426 | 0.132 |
| 3023 | **Serpind1** | Unchanged | P49182 | 3 | 13 | 3 | 478 | 54.5 | 7.34 | 7 | 0.841 | 0.799 | 1.243 | 0.768 | 0.312 | 0.186 | 3.988 | 0.185 |

| | | | | | | | | | | | | | | | | | | |
|---|---|---|---|---|---|---|---|---|---|---|---|---|---|---|---|---|---|---|
| 3024 | **B4galnt1** | Unchanged | Q09200 | 2 | 3 | 2 | 533 | 59.2 | 8.59 | 7 | 0.551 | 0.349 | 1.445 | 0.673 | 0.485 | 0.273 | 2.977 | 0.337 |
| 3025 | **Cpn2** | Unchanged | Q9DBB9 | 2 | 9 | 2 | 547 | 60.4 | 5.88 | 7 | 0.916 | 0.889 | 0.436 | 0.303 | 0.531 | 0.367 | 0.822 | 0.791 |
| 3026 | **Itih3** | Unchanged | Q61704 | 3 | 6 | 3 | 889 | 99.3 | 6.05 | 7 | 0.980 | 0.971 | 1.741 | 0.473 | 4.254 | 0.172 | 0.409 | 0.312 |
| 3027 | **Babam2** | Unchanged | Q8K3W0 | 2 | 22 | 2 | 383 | 43.5 | 5.94 | 7 | 1.323 | 0.548 | 1.509 | 0.550 | 1.343 | 0.546 | 1.124 | 0.859 |
| 3028 | **Anp32e** | Unchanged | P97822 | 1 | 10 | 1 | 260 | 29.6 | 3.88 | 7 | 1.177 | 0.798 | 1.296 | 0.660 | 2.053 | 0.215 | 0.631 | 0.385 |
| 3029 | **Arhgap1** | Unchanged | A2AH25 | 2 | 2 | 2 | 479 | 54.4 | 5.87 | 7 | 0.802 | 0.704 | 5.514 | 0.366 | 0.386 | 0.192 | 14.290 | 0.306 |
| 3030 | **Ccz1** | Unchanged | Q8C1Y8 | 3 | 4 | 3 | 480 | 55.5 | 6.02 | 7 | 1.062 | 0.883 | 0.646 | 0.414 | 0.327 | 0.081 | 1.971 | 0.223 |
| 3031 | **Vac14** | Unchanged | Q80WQ2 | 4 | 17 | 4 | 782 | 88 | 6.13 | 7 | 1.841 | 0.234 | 3.111 | 0.099 | 1.428 | 0.388 | 2.178 | 0.189 |
| 3032 | **Minpp1** | Unchanged | Q9Z2L6 | 2 | 2 | 2 | 481 | 54.5 | 7.49 | 7 | 1.393 | 0.588 | 1.621 | 0.323 | 0.964 | 0.928 | 1.681 | 0.284 |
| 3033 | **Dedd2** | Unchanged | Q8QZV0 | 1 | 1 | 1 | 330 | 36.8 | 8.57 | 7 | 0.271 | 0.258 | 0.731 | 0.696 | 2.276 | 0.305 | 0.321 | 0.177 |
| 3034 | **Usp39** | Down | Q3TIX9 | 3 | 3 | 3 | 564 | 65.1 | 8.9 | 7 | 0.136 | 0.043 | 0.444 | 0.234 | 0.158 | 0.048 | 2.813 | 0.270 |
| 3035 | **Man2b2** | Unchanged | O54782 | 5 | 13 | 5 | 1018 | 115.5 | 7.39 | 7 | 0.290 | 0.141 | 0.326 | 0.156 | 0.342 | 0.169 | 0.953 | 0.821 |
| 3036 | **Cyfip1** | Unchanged | Q7TMB8 | 7 | 12 | 7 | 1253 | 145.1 | 6.9 | 7 | 2.032 | 0.350 | 0.567 | 0.081 | 1.726 | 0.100 | 0.328 | 0.020 |
| 3037 | **Paip1** | Unchanged | Q8VE62 | 2 | 15 | 2 | 400 | 45.7 | 4.55 | 7 | 2.222 | 0.152 | 2.025 | 0.414 | 2.912 | 0.232 | 0.696 | 0.628 |
| 3038 | **Abraxas2** | Down | Q3TCJ1 | 2 | 11 | 2 | 415 | 46.9 | 6.18 | 7 | 0.911 | 0.847 | 0.423 | 0.143 | 0.135 | 0.002 | 3.139 | 0.367 |
| 3039 | **Slc3a2** | Unchanged | P10852 | 2 | 6 | 2 | 526 | 58.3 | 5.91 | 7 | 0.719 | 0.178 | 0.927 | 0.670 | 1.735 | 0.425 | 0.534 | 0.382 |
| 3040 | **Trappc3** | UP | O55013 | 1 | 2 | 1 | 180 | 20.3 | 4.96 | 7 | 1.765 | 0.206 | 3.546 | 0.040 | 0.715 | 0.581 | 4.961 | 0.020 |
| 3041 | **Mrpl18** | Unchanged | Q9CQL5 | 1 | 2 | 1 | 180 | 20.7 | 9.28 | 7 | 0.856 | 0.828 | 1.762 | 0.563 | 0.962 | 0.965 | 1.833 | 0.574 |
| 3042 | **Wbp11** | Unchanged | A0A0N4SWF7 | 1 | 1 | 1 | 194 | 22.7 | 10.56 | 7 | 1.156 | 0.810 | 0.983 | 0.981 | 0.677 | 0.558 | 1.451 | 0.640 |
| 3043 | **Vmn1r69** | Unchanged | Q8VIC1 | 1 | 1 | 1 | 319 | 36.5 | 9.69 | 7 | 1.500 | 0.382 | 0.597 | 0.214 | 1.914 | 0.180 | 0.312 | 0.079 |
| 3044 | **Sept14** | Unchanged | Q9DA97 | 2 | 37 | 1 | 430 | 49.8 | 7.27 | 7 | 1.877 | 0.051 | 10.086 | 0.075 | 1.162 | 0.500 | 8.677 | 0.080 |
| 3045 | **Ppp2r5a** | Unchanged | Q6PD03 | 2 | 13 | 2 | 486 | 56.3 | 7.03 | 7 | 1.157 | 0.714 | 0.439 | 0.129 | 0.782 | 0.573 | 0.562 | 0.241 |
| 3046 | **Lamtor2** | Down | Q9JHS3 | 1 | 10 | 1 | 125 | 13.5 | 5.4 | 7 | 3.261 | 0.282 | 3.319 | 0.152 | 0.209 | 0.032 | 15.903 | 0.062 |
| 3047 | **Nacc1** | Unchanged | Q7TSZ8 | 1 | 2 | 1 | 514 | 56.5 | 5.73 | 7 | 1.194 | 0.815 | 1.691 | 0.526 | 0.986 | 0.985 | 1.716 | 0.508 |
| 3048 | **Bcs1l** | Unchanged | Q9CZP5 | 2 | 2 | 2 | 418 | 47.4 | 7.93 | 7 | 0.620 | 0.324 | 1.535 | 0.545 | 0.582 | 0.295 | 2.635 | 0.267 |
| 3049 | **Slc6a12** | Unchanged | P31651 | 3 | 7 | 2 | 614 | 69.6 | 7.17 | 7 | 0.923 | 0.852 | 1.167 | 0.740 | 0.365 | 0.131 | 3.198 | 0.071 |
| 3050 | **Rbm17** | Unchanged | Q8JZX4 | 2 | 2 | 2 | 405 | 45.3 | 5.82 | 7 | 0.617 | 0.365 | 0.566 | 0.217 | 0.370 | 0.181 | 1.530 | 0.587 |
| 3051 | **Cdkn2c** | Unchanged | Q60772 | 1 | 1 | 1 | 168 | 18.1 | 6 | 7 | #DIV/0! | #DIV/0! | #DIV/0! | #DIV/0! | #DIV/0! | #DIV/0! | #DIV/0! | #DIV/0! |
| 3052 | **Slc39a7** | Unchanged | Q31125 | 2 | 3 | 2 | 476 | 50.6 | 6.87 | 7 | 1.039 | 0.965 | 1.418 | 0.645 | 0.695 | 0.632 | 2.042 | 0.359 |
| 3053 | **Nhej1** | Unchanged | Q3KNJ2 | 1 | 2 | 1 | 295 | 32.7 | 6.74 | 7 | 1.400 | 0.645 | 2.151 | 0.230 | 0.745 | 0.677 | 2.890 | 0.076 |
| 3054 | **Stam** | Unchanged | P70297 | 2 | 3 | 2 | 548 | 59.7 | 4.84 | 7 | 1.401 | 0.492 | 0.634 | 0.455 | 0.499 | 0.222 | 1.270 | 0.708 |
| 3055 | **Cldn13** | Unchanged | Q9Z0S4 | 1 | 1 | 1 | 211 | 23.5 | 5.86 | 7 | 3.084 | 0.125 | 2.405 | 0.288 | 0.200 | 0.219 | 12.047 | 0.078 |
| 3056 | **Dek** | Unchanged | Q7TNV0 | 2 | 9 | 2 | 380 | 43.1 | 6.86 | 7 | 0.523 | 0.283 | 0.927 | 0.905 | 1.942 | 0.071 | 0.477 | 0.088 |
| 3057 | **Grb7** | Unchanged | Q03160 | 3 | 14 | 3 | 535 | 59.9 | 8.19 | 7 | 1.344 | 0.526 | 3.409 | 0.210 | 1.048 | 0.881 | 3.254 | 0.213 |
| 3058 | **Naa25** | Unchanged | Q8BWZ3 | 4 | 14 | 4 | 972 | 111.6 | 6.52 | 7 | 1.072 | 0.908 | 0.897 | 0.868 | 0.939 | 0.927 | 0.955 | 0.926 |
| 3059 | **Phkb** | Unchanged | Q7TSH2 | 4 | 20 | 4 | 1085 | 123.8 | 6.83 | 7 | 1.009 | 0.987 | 1.904 | 0.128 | 0.562 | 0.218 | 3.391 | 0.014 |
| 3060 | **Glra3** | Unchanged | G5E811 | 1 | 1 | 1 | 480 | 55.6 | 8.72 | 7 | 1.467 | 0.542 | 0.477 | 0.251 | 0.541 | 0.311 | 0.882 | 0.748 |
| 3061 | **Arhgap22** | Unchanged | D3Z1Y5 | 1 | 1 | 1 | 537 | 58.9 | 7.18 | 7 | 1.673 | 0.090 | 14.784 | 0.126 | 18.838 | 0.052 | 0.785 | 0.728 |
| 3062 | **Inpp1** | Unchanged | P49442 | 2 | 29 | 2 | 396 | 43.3 | 5.01 | 7 | 1.630 | 0.262 | 3.034 | 0.088 | 1.235 | 0.611 | 2.456 | 0.141 |
| 3063 | **Eva1a** | Unchanged | Q91WM6 | 1 | 2 | 1 | 156 | 17.8 | 6.55 | 7 | #DIV/0! | #DIV/0! | #DIV/0! | #DIV/0! | #DIV/0! | #DIV/0! | #DIV/0! | #DIV/0! |
| 3064 | **Arih1** | Unchanged | Q9Z1K5 | 2 | 7 | 2 | 555 | 64 | 5.08 | 7 | 0.777 | 0.571 | 0.373 | 0.066 | 0.464 | 0.103 | 0.804 | 0.735 |

| | | | | | | | | | | | | | | | | | | |
|---|---|---|---|---|---|---|---|---|---|---|---|---|---|---|---|---|---|---|
| 3065 | **Ipo4** | UP | Q8VI75 | 5 | 17 | 5 | 1082 | 119.2 | 5.03 | 7 | 3.795 | 0.046 | 4.460 | 0.018 | 1.011 | 0.979 | 4.413 | 0.016 |
| 3066 | **Fads6** | Unchanged | Q80UG1 | 1 | 1 | 1 | 342 | 38.8 | 8.46 | 7 | 1.259 | 0.666 | 1.954 | 0.311 | 2.958 | 0.317 | 0.661 | 0.618 |
| 3067 | **Sf3a2** | Unchanged | G3UVU2 | 2 | 5 | 2 | 485 | 51.2 | 9.64 | 7 | 1.954 | 0.405 | 0.990 | 0.966 | 0.873 | 0.642 | 1.134 | 0.700 |
| 3068 | **Ugt2a2** | Unchanged | Q6PDD0 | 3 | 25 | 1 | 528 | 59.9 | 8.35 | 7 | 1.764 | 0.542 | 2.478 | 0.381 | 0.393 | 0.347 | 6.300 | 0.198 |
| 3069 | **Mfap4** | UP | Q9D1H9 | 1 | 4 | 1 | 257 | 28.9 | 5.44 | 7 | 4.451 | 0.315 | 6.766 | 0.057 | 7.685 | 0.004 | 0.880 | 0.771 |
| 3070 | **Cul2** | Unchanged | Q9D4H8 | 5 | 25 | 5 | 745 | 86.8 | 7.01 | 7 | 2.048 | 0.146 | 1.177 | 0.750 | 0.512 | 0.271 | 2.297 | 0.109 |
| 3071 | **Snx7** | Unchanged | Q9CY18 | 1 | 4 | 1 | 387 | 45 | 5.07 | 7 | 17.383 | 0.345 | 5.654 | 0.087 | 1.151 | 0.713 | 4.911 | 0.099 |
| 3072 | **Ppih** | Unchanged | A2BGI8 | 1 | 1 | 1 | 158 | 17.2 | 8.34 | 7 | 1.076 | 0.874 | 0.896 | 0.858 | 0.457 | 0.279 | 1.961 | 0.439 |
| 3073 | **Nup93** | Unchanged | Q8BJ71 | 3 | 5 | 3 | 819 | 93.2 | 5.72 | 7 | 0.532 | 0.117 | 1.735 | 0.174 | 0.870 | 0.693 | 1.995 | 0.110 |
| 3074 | **Rab3ip** | Unchanged | A0A1W2P7K6 | 2 | 2 | 2 | 460 | 50.9 | 5.94 | 7 | 3.331 | 0.303 | 1.810 | 0.249 | 1.923 | 0.415 | 0.941 | 0.924 |
| 3075 | **Fubp3** | Down | Q3TIX6 | 2 | 2 | 2 | 575 | 61.9 | 8.38 | 7 | 0.569 | 0.257 | 0.431 | 0.121 | 0.241 | 0.038 | 1.785 | 0.255 |
| 3076 | **Krt19** | Unchanged | P19001 | 2 | 18 | 1 | 403 | 44.5 | 5.39 | 7 | 0.622 | 0.342 | 0.990 | 0.974 | 1.776 | 0.484 | 0.558 | 0.482 |
| 3077 | **Bud31** | UP | Q6PGH1 | 1 | 1 | 1 | 144 | 17 | 8.82 | 7 | 3.514 | 0.172 | 2.029 | 0.035 | 4.061 | 0.152 | 0.500 | 0.326 |
| 3078 | **Gfy** | Unchanged | J3KML8 | 1 | 1 | 1 | 504 | 53.8 | 6.2 | 7 | #DIV/0! | #DIV/0! | #DIV/0! | #DIV/0! | #DIV/0! | #DIV/0! | #DIV/0! | #DIV/0! |
| 3079 | **Acot11** | Unchanged | Q8VHQ9 | 2 | 4 | 2 | 594 | 67.3 | 6.8 | 7 | 1.156 | 0.789 | 1.144 | 0.863 | 0.486 | 0.173 | 2.353 | 0.406 |
| 3080 | **Snx1** | Unchanged | Q9WV80 | 3 | 17 | 2 | 522 | 58.9 | 5.22 | 7 | 1.154 | 0.854 | 1.436 | 0.632 | 0.808 | 0.787 | 1.778 | 0.432 |
| 3081 | **Itfg1** | Down | Q99KW9 | 1 | 1 | 1 | 610 | 67.4 | 5.73 | 7 | 0.622 | 0.149 | 2.168 | 0.368 | 0.450 | 0.004 | 4.819 | 0.200 |
| 3082 | **Myh10** | Unchanged | Q5SV64 | 12 | 50 | 1 | 2007 | 232.3 | 5.54 | 7 | 0.408 | 0.233 | 0.657 | 0.633 | 0.088 | 0.065 | 7.496 | 0.330 |
| 3083 | **Rcn2** | Unchanged | Q8BP92 | 2 | 6 | 1 | 320 | 37.2 | 4.42 | 7 | 2.584 | 0.174 | 1.866 | 0.618 | 1.486 | 0.619 | 1.256 | 0.834 |
| 3084 | **Mcrip2** | Unchanged | Q9CQB2 | 1 | 1 | 1 | 160 | 17.9 | 7.3 | 7 | 0.623 | 0.313 | 1.592 | 0.653 | 3.403 | 0.301 | 0.468 | 0.487 |
| 3085 | **Slc25a23** | Unchanged | Q6GQS1 | 2 | 5 | 2 | 467 | 52.5 | 7.59 | 7 | 1.386 | 0.457 | 6.250 | 0.317 | 0.906 | 0.810 | 6.898 | 0.309 |
| 3086 | **Hgs** | Unchanged | Q99LI8 | 3 | 6 | 3 | 775 | 86 | 6.16 | 7 | 0.886 | 0.702 | 2.431 | 0.274 | 1.303 | 0.474 | 1.866 | 0.395 |
| 3087 | **Rufy3** | Unchanged | Q9D394 | 2 | 5 | 2 | 469 | 53 | 5.49 | 7 | 1.028 | 0.930 | 2.347 | 0.167 | 0.863 | 0.565 | 2.718 | 0.127 |
| 3088 | **Baiap2l1** | Unchanged | Q9DBJ3 | 2 | 8 | 2 | 514 | 57.2 | 8.75 | 7 | 0.648 | 0.488 | 0.830 | 0.797 | 0.251 | 0.089 | 3.303 | 0.313 |
| 3089 | **Slc47a2** | Unchanged | Q3V050 | 1 | 1 | 1 | 573 | 61.7 | 7.83 | 7 | 0.025 | 0.066 | 0.036 | 0.069 | 0.281 | 0.176 | 0.130 | 0.154 |
| 3090 | **Ergic3** | Unchanged | Q9CQE7 | 1 | 1 | 1 | 383 | 43.2 | 6.47 | 7 | 2.126 | 0.476 | 5.935 | 0.212 | 0.063 | 0.222 | 94.268 | 0.139 |
| 3091 | **Vps28** | Unchanged | Q9D1C8 | 1 | 3 | 1 | 221 | 25.4 | 5.54 | 7 | 0.976 | 0.922 | 1.075 | 0.805 | 1.312 | 0.213 | 0.819 | 0.423 |
| 3092 | **MINDY3** | Unchanged | Q9CV28 | 2 | 6 | 2 | 444 | 49.6 | 4.78 | 7 | 0.940 | 0.859 | 0.934 | 0.860 | 0.643 | 0.377 | 1.452 | 0.504 |
| 3093 | **9030624J02Rik** | Unchanged | Q8BWQ6 | 5 | 10 | 5 | 963 | 109 | 7.31 | 7 | 0.954 | 0.909 | 1.887 | 0.133 | 0.486 | 0.134 | 3.885 | 0.014 |
| 3094 | **Mfn2** | Unchanged | Q80U63 | 3 | 11 | 3 | 757 | 86.1 | 6.77 | 7 | 0.953 | 0.933 | 0.883 | 0.827 | 1.355 | 0.615 | 0.652 | 0.480 |
| 3095 | **Ppox** | Unchanged | P51175 | 2 | 4 | 2 | 477 | 50.8 | 8.84 | 7 | 2.383 | 0.102 | 3.246 | 0.389 | 2.695 | 0.242 | 1.204 | 0.847 |
| 3096 | **Aipl1** | Unchanged | Q924K1 | 1 | 2 | 1 | 328 | 38.2 | 5.95 | 7 | 0.831 | 0.769 | 0.026 | 0.063 | 0.059 | 0.071 | 0.435 | 0.079 |
| 3097 | **H1f0** | Unchanged | P10922 | 1 | 7 | 1 | 194 | 20.8 | 10.9 | 7 | 0.166 | 0.055 | 0.941 | 0.926 | 13.753 | 0.154 | 0.068 | 0.153 |
| 3098 | **Slc16a1** | Unchanged | P53986 | 2 | 35 | 2 | 493 | 53.2 | 7.47 | 7 | 0.553 | 0.142 | 1.007 | 0.990 | 0.788 | 0.604 | 1.277 | 0.729 |
| 3099 | **Castor2** | Unchanged | Q8CAB8 | 1 | 1 | 1 | 329 | 36.1 | 5.45 | 7 | 1.125 | 0.642 | 0.752 | 0.312 | 1.173 | 0.629 | 0.641 | 0.305 |
| 3100 | **Spcs3** | Unchanged | Q6ZWQ7 | 1 | 8 | 1 | 180 | 20.3 | 8.62 | 7 | 0.636 | 0.391 | 0.655 | 0.476 | 0.741 | 0.570 | 0.885 | 0.824 |
| 3101 | **Nup62** | Unchanged | Q63850 | 2 | 2 | 2 | 526 | 53.2 | 5.31 | 7 | 2.259 | 0.267 | 4.115 | 0.102 | 1.147 | 0.754 | 3.586 | 0.119 |
| 3102 | **Retreg3** | Unchanged | Q9CQV4 | 1 | 1 | 1 | 466 | 51.6 | 4.97 | 7 | 1.232 | 0.663 | 1.225 | 0.583 | 1.208 | 0.638 | 1.014 | 0.973 |
| 3103 | **Tapbpl** | Unchanged | Q8VD31 | 2 | 2 | 2 | 451 | 48.7 | 6.46 | 7 | 0.553 | 0.331 | 0.152 | 0.056 | 0.466 | 0.202 | 0.325 | 0.018 |
| 3104 | **Mlph** | Unchanged | A0A0R4J075 | 1 | 1 | 1 | 590 | 65 | 6.07 | 7 | #DIV/0! | #DIV/0! | #DIV/0! | #DIV/0! | #DIV/0! | #DIV/0! | #DIV/0! | #DIV/0! |
| 3105 | **Cdc5l** | UP | Q6A068 | 3 | 5 | 3 | 802 | 92.1 | 8.02 | 7 | 9.434 | 0.091 | 3.763 | 0.097 | 10.345 | 0.025 | 0.364 | 0.106 |

| | | | | | | | | | | | | | | | | | | |
|---|---|---|---|---|---|---|---|---|---|---|---|---|---|---|---|---|---|---|
| 3106 | **Olfr154** | Unchanged | Q9QY00 | 1 | 1 | 1 | 318 | 35.9 | 8.66 | 7 | 1.226 | 0.552 | 0.845 | 0.549 | 0.848 | 0.542 | 0.996 | 0.991 |
| 3107 | **Emc10** | Unchanged | Q3TAS6 | 1 | 4 | 1 | 258 | 27 | 5.91 | 7 | 1.081 | 0.889 | 1.241 | 0.757 | 0.414 | 0.202 | 3.000 | 0.238 |
| 3108 | **Jagn1** | UP | Q5XKN4 | 1 | 2 | 1 | 183 | 21.1 | 9.8 | 7 | 0.346 | 0.112 | 1.803 | 0.317 | 9.983 | 0.041 | 0.181 | 0.060 |
| 3109 | **Ripk1** | Unchanged | Q60855 | 2 | 2 | 2 | 656 | 74.8 | 6.47 | 7 | 7.584 | 0.207 | 4.880 | 0.071 | 1.674 | 0.328 | 2.915 | 0.120 |
| 3110 | **Mfsd3** | Unchanged | Q5U419 | 1 | 8 | 1 | 412 | 43.1 | 9 | 7 | 1.473 | 0.201 | 1.526 | 0.221 | 1.422 | 0.286 | 1.073 | 0.791 |
| 3111 | **Slc33a1** | Unchanged | Q99J27 | 2 | 27 | 2 | 550 | 61 | 8.16 | 7 | 0.926 | 0.917 | 0.319 | 0.170 | 0.508 | 0.322 | 0.628 | 0.408 |
| 3112 | **Ubr4** | Unchanged | A2AN08 | 18 | 64 | 18 | 5180 | 571.9 | 6.06 | 7 | 0.879 | 0.859 | 0.511 | 0.402 | 0.128 | 0.139 | 3.982 | 0.040 |
| 3113 | **Ascc2** | Unchanged | Q91WR3 | 2 | 5 | 2 | 749 | 85.6 | 5.03 | 7 | 1.033 | 0.919 | 1.424 | 0.372 | 1.067 | 0.864 | 1.335 | 0.449 |
| 3114 | **Mrpl45** | Unchanged | Q9D0Q7 | 1 | 2 | 1 | 306 | 35.4 | 9.23 | 7 | 1.682 | 0.526 | 7.321 | 0.178 | 1.118 | 0.836 | 6.549 | 0.187 |
| 3115 | **Marcksl1** | Unchanged | P28667 | 1 | 2 | 1 | 200 | 20.2 | 4.61 | 7 | #DIV/0! | #DIV/0! | #DIV/0! | #DIV/0! | #DIV/0! | #DIV/0! | #DIV/0! | #DIV/0! |
| 3116 | **Fitm2** | Unchanged | P59266 | 1 | 2 | 1 | 262 | 30 | 9 | 6 | 1.113 | 0.778 | 1.942 | 0.316 | 0.746 | 0.536 | 2.605 | 0.192 |
| 3117 | **Ddx58** | Unchanged | Q6Q899 | 3 | 14 | 3 | 926 | 105.9 | 6.65 | 6 | 0.874 | 0.727 | 1.410 | 0.556 | 0.921 | 0.722 | 1.532 | 0.468 |
| 3118 | **Atp6v1d** | Unchanged | P57746 | 1 | 3 | 1 | 247 | 28.4 | 9.45 | 6 | 0.728 | 0.604 | 24.427 | 0.088 | 67.746 | 0.050 | 0.361 | 0.205 |
| 3119 | **Vta1** | Unchanged | Q9CR26 | 1 | 2 | 1 | 309 | 33.9 | 6.13 | 6 | 1.034 | 0.950 | 1.486 | 0.561 | 0.390 | 0.259 | 3.805 | 0.132 |
| 3120 | **Stk38** | UP | Q91VJ4 | 1 | 1 | 1 | 465 | 54.1 | 7.34 | 6 | 1.334 | 0.313 | 1.777 | 0.036 | 1.004 | 0.994 | 1.771 | 0.102 |
| 3121 | **Col6a1** | UP | Q04857 | 4 | 44 | 4 | 1025 | 108.4 | 5.36 | 6 | 3.877 | 0.103 | 5.649 | 0.022 | 3.070 | 0.036 | 1.840 | 0.168 |
| 3122 | **Bnip3** | Unchanged | O55003 | 1 | 3 | 1 | 187 | 21 | 7.14 | 6 | 0.472 | 0.261 | 1.324 | 0.623 | 0.529 | 0.306 | 2.505 | 0.167 |
| 3123 | **Prpf31** | Unchanged | Q8CCF0 | 2 | 3 | 2 | 499 | 55.4 | 5.67 | 6 | 1.346 | 0.347 | 1.470 | 0.381 | 0.963 | 0.901 | 1.526 | 0.314 |
| 3124 | **Pym1** | Unchanged | Q8CHP5 | 1 | 22 | 1 | 203 | 22.7 | 9.5 | 6 | 0.567 | 0.222 | 0.746 | 0.494 | 0.674 | 0.220 | 1.107 | 0.824 |
| 3125 | **Lamp1** | Unchanged | P11438 | 2 | 6 | 2 | 406 | 43.8 | 8.4 | 6 | 0.669 | 0.636 | 0.711 | 0.718 | 0.211 | 0.214 | 3.376 | 0.349 |
| 3126 | **Bst2** | Unchanged | Q8R2Q8 | 1 | 1 | 1 | 172 | 19.1 | 7.34 | 6 | #DIV/0! | #DIV/0! | #DIV/0! | #DIV/0! | #DIV/0! | #DIV/0! | #DIV/0! | #DIV/0! |
| 3127 | **Lgalsl** | Unchanged | Q8VED9 | 1 | 2 | 1 | 172 | 18.9 | 5.35 | 6 | 0.628 | 0.538 | 0.503 | 0.405 | 0.134 | 0.132 | 3.756 | 0.158 |
| 3128 | **Sdf2** | Unchanged | A0A0R4IZW9 | 1 | 3 | 1 | 219 | 24 | 7.61 | 6 | #DIV/0! | #DIV/0! | #DIV/0! | #DIV/0! | #DIV/0! | #DIV/0! | #DIV/0! | #DIV/0! |
| 3129 | **Nckap1** | UP | P28660 | 4 | 11 | 4 | 1128 | 128.7 | 6.62 | 6 | 1.931 | 0.036 | 2.250 | 0.217 | 1.076 | 0.782 | 2.090 | 0.234 |
| 3130 | **Mrpl16** | Unchanged | Q99N93 | 1 | 3 | 1 | 251 | 28.8 | 10.27 | 6 | 0.925 | 0.804 | 1.541 | 0.540 | 2.110 | 0.233 | 0.730 | 0.633 |
| 3131 | **Sod3** | Unchanged | O09164 | 1 | 4 | 1 | 251 | 27.4 | 6.84 | 6 | 1.086 | 0.909 | 2.441 | 0.095 | 0.455 | 0.281 | 5.369 | 0.011 |
| 3132 | **Appl1** | Unchanged | Q8K3H0 | 2 | 8 | 2 | 707 | 79.3 | 5.41 | 6 | 0.886 | 0.638 | 1.567 | 0.402 | 1.300 | 0.329 | 1.205 | 0.682 |
| 3133 | **Stx8** | Unchanged | O88983 | 1 | 4 | 1 | 236 | 26.9 | 5.01 | 6 | #DIV/0! | #DIV/0! | #DIV/0! | #DIV/0! | #DIV/0! | #DIV/0! | #DIV/0! | #DIV/0! |
| 3134 | **Galns** | Unchanged | Q571E4 | 2 | 10 | 2 | 520 | 57.6 | 6.52 | 6 | 1.923 | 0.306 | 5.015 | 0.201 | 1.452 | 0.483 | 3.454 | 0.254 |
| 3135 | **Slc35c1** | Unchanged | Q8BLX4 | 1 | 1 | 1 | 363 | 39.9 | 8.62 | 6 | 2.476 | 0.366 | 1.847 | 0.311 | 0.968 | 0.943 | 1.909 | 0.313 |
| 3136 | **Mgat2** | Unchanged | Q921V5 | 1 | 1 | 1 | 442 | 51 | 8.59 | 6 | 1.164 | 0.680 | 1.287 | 0.450 | 1.248 | 0.610 | 1.031 | 0.937 |
| 3137 | **Ogfod2** | Unchanged | Q9CQ04 | 1 | 2 | 1 | 349 | 39.2 | 6.39 | 6 | #DIV/0! | #DIV/0! | #DIV/0! | #DIV/0! | #DIV/0! | #DIV/0! | #DIV/0! | #DIV/0! |
| 3138 | **Tm9sf3** | Unchanged | Q9ET30 | 2 | 21 | 2 | 587 | 67.5 | 7.21 | 6 | 0.882 | 0.848 | 1.841 | 0.214 | 0.796 | 0.664 | 2.313 | 0.080 |
| 3139 | **Actl6a** | Unchanged | Q9Z2N8 | 2 | 7 | 2 | 429 | 47.4 | 5.6 | 6 | 1.494 | 0.401 | 70.476 | 0.141 | 0.830 | 0.715 | 84.884 | 0.140 |
| 3140 | **Emilin1** | Unchanged | Q99K41 | 3 | 10 | 3 | 1017 | 107.5 | 5.3 | 6 | 2.462 | 0.289 | 1.802 | 0.347 | 1.033 | 0.949 | 1.745 | 0.413 |
| 3141 | **Dohh** | UP | Q99LN9 | 1 | 1 | 1 | 302 | 32.9 | 4.84 | 6 | 0.914 | 0.808 | 2.442 | 0.093 | 2.032 | 0.017 | 1.202 | 0.619 |
| 3142 | **Mtnd1** | Unchanged | P03888 | 1 | 4 | 1 | 318 | 36 | 6.93 | 6 | 1.578 | 0.390 | 2.578 | 0.325 | 1.546 | 0.513 | 1.667 | 0.534 |
| 3143 | **Hspa14** | Unchanged | Q99M31 | 1 | 3 | 1 | 509 | 54.6 | 5.92 | 6 | 1.571 | 0.270 | 1.757 | 0.234 | 1.755 | 0.356 | 1.001 | 0.998 |
| 3144 | **Hnrnpul1** | Unchanged | Q8VDM6 | 4 | 18 | 3 | 859 | 95.9 | 6.58 | 6 | 0.974 | 0.948 | 4.237 | 0.205 | 1.815 | 0.352 | 2.334 | 0.353 |
| 3145 | **Cryab** | Unchanged | P23927 | 1 | 1 | 1 | 175 | 20.1 | 7.33 | 6 | 0.274 | 0.221 | 0.285 | 0.229 | 3.985 | 0.366 | 0.071 | 0.262 |
| 3146 | **Fmo4** | Unchanged | Q8VHG0 | 2 | 3 | 1 | 560 | 63.8 | 8.94 | 6 | 1.422 | 0.324 | 1.328 | 0.428 | 0.604 | 0.330 | 2.199 | 0.079 |

| | | | | | | | | | | | | | | | | | | |
|---|---|---|---|---|---|---|---|---|---|---|---|---|---|---|---|---|---|---|
| 3147 | **Usp47** | Unchanged | Q8BY87 | 6 | 30 | 6 | 1376 | 157.4 | 5.11 | 6 | 0.945 | 0.896 | 0.889 | 0.807 | 1.044 | 0.930 | 0.851 | 0.697 |
| 3148 | **Ccdc25** | Unchanged | Q78PG9 | 1 | 2 | 1 | 208 | 24.5 | 6.95 | 6 | #DIV/0! | #DIV/0! | #DIV/0! | #DIV/0! | #DIV/0! | #DIV/0! | #DIV/0! | #DIV/0! |
| 3149 | **Ccdc85a** | Unchanged | E0CX32 | 1 | 1 | 1 | 545 | 59.6 | 8.68 | 6 | #DIV/0! | #DIV/0! | #DIV/0! | #DIV/0! | #DIV/0! | #DIV/0! | #DIV/0! | #DIV/0! |
| 3150 | **Smu1** | Unchanged | Q3UKJ7 | 3 | 7 | 3 | 513 | 57.5 | 7.18 | 6 | 0.410 | 0.145 | 1.117 | 0.844 | 1.388 | 0.543 | 0.805 | 0.699 |
| 3151 | **Krt76** | Unchanged | Q3UV17 | 4 | 13 | 1 | 594 | 62.8 | 8.43 | 6 | 0.677 | 0.217 | 0.669 | 0.455 | 0.545 | 0.114 | 1.226 | 0.759 |
| 3152 | **Cdca8** | Unchanged | Q8BHX3 | 1 | 2 | 1 | 289 | 32.2 | 10.13 | 6 | 0.410 | 0.374 | 0.624 | 0.569 | 0.986 | 0.985 | 0.633 | 0.476 |
| 3153 | **Pde2a** | Unchanged | Q922S4 | 3 | 6 | 3 | 916 | 103.2 | 5.38 | | 0.787 | 0.782 | 1.212 | 0.845 | 0.238 | 0.295 | 5.090 | 0.258 |
| 3154 | **Gbp6** | Unchanged | A0A0G2JDV3 | 2 | 4 | 2 | 611 | 70.4 | 6.11 | 6 | 1.117 | 0.720 | 2.941 | 0.089 | 1.397 | 0.237 | 2.105 | 0.165 |
| 3155 | **Adprhl2** | Unchanged | Q8CG72 | 2 | 3 | 2 | 370 | 39.4 | 4.96 | 6 | 0.600 | 0.432 | 0.871 | 0.842 | 0.210 | 0.093 | 4.151 | 0.206 |
| 3156 | **Fxr1** | Unchanged | Q61584 | 3 | 29 | 3 | 677 | 76.2 | 6.98 | 6 | 0.927 | 0.639 | 0.906 | 0.769 | 0.957 | 0.760 | 0.948 | 0.882 |
| 3157 | **Vnn3** | Unchanged | Q9QZ25 | 1 | 2 | 1 | 500 | 56.3 | 6.29 | 6 | 1.167 | 0.758 | 1.765 | 0.312 | 0.851 | 0.767 | 2.075 | 0.246 |
| 3158 | **Krt14** | Unchanged | Q61781 | 2 | 4 | 1 | 484 | 52.8 | 5.17 | 6 | 6.737 | 0.319 | 9.506 | 0.220 | 9.971 | 0.092 | 0.953 | 0.955 |
| 3159 | **Parp3** | UP | Q3ULW8 | 2 | 2 | 2 | 533 | 59.9 | 7.12 | 6 | 5.731 | 0.049 | 5.707 | 0.174 | 2.577 | 0.007 | 2.215 | 0.353 |
| 3160 | **Washc3** | Unchanged | Q9CR27 | 1 | 3 | 1 | 194 | 21.1 | 4.46 | 6 | 1.155 | 0.777 | 1.025 | 0.966 | 0.428 | 0.253 | 2.394 | 0.149 |
| 3161 | **Snx6** | Down | Q6P8X1 | 2 | 8 | 2 | 406 | 46.6 | 6.16 | 6 | 1.461 | 0.497 | 2.814 | 0.222 | 0.397 | 0.039 | 7.082 | 0.112 |
| 3162 | **Tmx3** | Unchanged | Q8BXZ1 | 1 | 7 | 1 | 456 | 51.8 | 5.16 | 6 | 1.294 | 0.691 | 0.817 | 0.754 | 2.097 | 0.217 | 0.390 | 0.106 |
| 3163 | **Slc38a3** | Unchanged | Q9DCP2 | 2 | 46 | 2 | 505 | 55.6 | 7.09 | 6 | 1.333 | 0.496 | 1.220 | 0.404 | 1.534 | 0.128 | 0.796 | 0.410 |
| 3164 | **Bin2** | Unchanged | S4R270 | 2 | 2 | 2 | 505 | 54.7 | 4.81 | 6 | #DIV/0! | #DIV/0! | #DIV/0! | #DIV/0! | #DIV/0! | #DIV/0! | #DIV/0! | #DIV/0! |
| 3165 | **Slc39a14** | Unchanged | Q75N73 | 1 | 6 | 1 | 489 | 53.9 | 5.27 | 6 | 0.638 | 0.586 | 0.864 | 0.833 | 0.202 | 0.142 | 4.269 | 0.142 |
| 3166 | **Ppic** | Unchanged | P30412 | 1 | 3 | 1 | 212 | 22.8 | 7.5 | 6 | 1.319 | 0.586 | 2.043 | 0.072 | 0.787 | 0.506 | 2.595 | 0.037 |
| 3167 | **Krt1** | Unchanged | P04104 | 5 | 56 | 3 | 637 | 65.6 | 8.15 | 6 | 2.975 | 0.201 | 1.166 | 0.809 | 1.974 | 0.323 | 0.591 | 0.430 |
| 3168 | **U2surp** | Unchanged | Q6NV83 | 3 | 6 | 3 | 1029 | 118.2 | 8.47 | 5 | 0.789 | 0.371 | 0.970 | 0.873 | 0.828 | 0.379 | 1.172 | 0.354 |
| 3169 | **Seh1l** | Unchanged | Q8R2U0 | 1 | 2 | 1 | 360 | 39.7 | 8.05 | 6 | 1.510 | 0.567 | 1.763 | 0.420 | 0.286 | 0.101 | 6.163 | 0.109 |
| 3170 | **Srsf10** | UP | Q9R0U0 | 1 | 9 | 1 | 262 | 31.3 | 11.27 | 6 | 4.456 | 0.166 | 12.550 | 0.129 | 7.121 | 0.012 | 1.763 | 0.464 |
| 3171 | **Slc10a1** | Unchanged | O08705 | 2 | 13 | 2 | 362 | 39.4 | 9.03 | 6 | 1.680 | 0.276 | 0.398 | 0.211 | 0.643 | 0.484 | 0.618 | 0.334 |
| 3172 | **Cdc42bpb** | Unchanged | Q7TT50 | 5 | 14 | 5 | 1713 | 194.6 | 6.46 | 6 | 1.305 | 0.654 | 1.268 | 0.627 | 0.711 | 0.541 | 1.783 | 0.267 |
| 3173 | **Ces2f** | Unchanged | Q08ED5 | 3 | 20 | 1 | 561 | 62.7 | 7.47 | 6 | 0.548 | 0.249 | 0.380 | 0.059 | 0.451 | 0.089 | 0.841 | 0.813 |
| 3174 | **Tcea2** | Unchanged | F6XMY4 | 2 | 2 | 1 | 297 | 33.1 | 8.68 | 6 | 0.645 | 0.624 | 0.613 | 0.540 | 10.523 | 0.181 | 0.058 | 0.165 |
| 3175 | **Nhlrc3** | Unchanged | Q8CCH2 | 2 | 7 | 2 | 347 | 38.2 | 6.23 | 6 | 0.255 | 0.314 | 0.228 | 0.296 | 0.107 | 0.232 | 2.128 | 0.013 |
| 3176 | **Srl** | Unchanged | Q7TQ48 | 4 | 6 | 4 | 910 | 99.1 | 4.46 | 6 | 0.405 | 0.357 | 0.349 | 0.316 | 0.307 | 0.288 | 1.138 | 0.532 |
| 3177 | **Usp24** | Unchanged | B1AY13 | 8 | 19 | 8 | 2617 | 293.8 | 6.19 | 6 | 0.779 | 0.349 | 1.277 | 0.414 | 1.575 | 0.056 | 0.811 | 0.389 |
| 3178 | **Hddc2** | Unchanged | Q3SXD3 | 1 | 1 | 1 | 199 | 22.7 | 5.02 | 6 | 0.226 | 0.181 | 1.316 | 0.734 | 0.473 | 0.361 | 2.784 | 0.289 |
| 3179 | **Pank1** | Unchanged | Q8K4K6 | 2 | 10 | 2 | 548 | 60.1 | 8.03 | 6 | 4.901 | 0.318 | 1.066 | 0.944 | 1.390 | 0.691 | 0.767 | 0.727 |
| 3180 | **Smyd5** | Unchanged | Q3TYX3 | 1 | 4 | 1 | 416 | 47.1 | 5.21 | 6 | 3.452 | 0.222 | 4.498 | 0.131 | 2.069 | 0.097 | 2.174 | 0.276 |
| 3181 | **Thumpd1** | UP | Q99J36 | 2 | 7 | 2 | 350 | 38.9 | 6.07 | 6 | 0.951 | 0.897 | 5.688 | 0.020 | 1.109 | 0.725 | 5.127 | 0.021 |
| 3182 | **Smc3** | Down | Q9CW03 | 5 | 7 | 5 | 1217 | 141.5 | 7.18 | 6 | 1.958 | 0.317 | 0.616 | 0.214 | 0.446 | 0.047 | 1.382 | 0.402 |
| 3183 | **Thrap3** | Unchanged | Q569Z6 | 4 | 9 | 4 | 951 | 108.1 | 10.17 | 6 | 2.289 | 0.221 | 5.240 | 0.141 | 3.800 | 0.150 | 1.379 | 0.653 |
| 3184 | **Arsb** | Down | P50429 | 3 | 16 | 3 | 534 | 59.6 | 7.24 | 6 | 0.629 | 0.152 | 0.489 | 0.122 | 0.274 | 0.016 | 1.784 | 0.321 |
| 3185 | **Ccdc124** | Unchanged | Q9D8X2 | 1 | 4 | 1 | 217 | 25.3 | 9.64 | 6 | 1.311 | 0.704 | 1.384 | 0.721 | 1.004 | 0.995 | 1.378 | 0.718 |
| 3186 | **Rdh13** | UP | Q8CEE7 | 2 | 3 | 2 | 334 | 36.4 | 8.85 | 6 | 0.424 | 0.406 | 0.377 | 0.372 | 5.582 | 0.035 | 0.068 | 0.015 |
| 3187 | **Mospd2** | Unchanged | Q9CWP6 | 2 | 5 | 2 | 518 | 59.8 | 5.67 | 6 | 0.968 | 0.944 | 1.324 | 0.362 | 1.739 | 0.214 | 0.761 | 0.534 |

| | | | | | | | | | | | | | | | | | | |
|---|---|---|---|---|---|---|---|---|---|---|---|---|---|---|---|---|---|---|
| 3188 | **Fcamr** | Unchanged | Q2TB54 | 1 | 1 | 1 | 535 | 57.8 | 9.22 | 6 | 1.169 | 0.733 | 0.809 | 0.716 | 1.032 | 0.938 | 0.784 | 0.641 |
| 3189 | **Slc6a13** | Down | P31649 | 3 | 15 | 2 | 602 | 68.2 | 6.25 | 6 | 1.472 | 0.168 | 0.327 | 0.048 | 0.867 | 0.792 | 0.377 | 0.228 |
| 3190 | **Rheb** | Unchanged | Q921J2 | 1 | 2 | 1 | 184 | 20.4 | 5.92 | 6 | 6.164 | 0.201 | 5.694 | 0.090 | 0.619 | 0.071 | 9.205 | 0.071 |
| 3191 | **Krt16** | Unchanged | Q9Z2K1 | 2 | 7 | 2 | 469 | 51.6 | 5.2 | 6 | 1.750 | 0.420 | 1.740 | 0.431 | 1.718 | 0.287 | 1.013 | 0.981 |
| 3192 | **Nudt5** | Unchanged | Q9JKX6 | 1 | 4 | 1 | 218 | 24 | 5.48 | 6 | 0.119 | 0.134 | 0.136 | 0.144 | 0.769 | 0.718 | 0.177 | 0.095 |
| 3193 | **Dnajc2** | Unchanged | P54103 | 3 | 8 | 3 | 621 | 71.7 | 8.7 | 6 | 0.850 | 0.790 | 0.504 | 0.261 | 0.480 | 0.238 | 1.049 | 0.916 |
| 3194 | **Tap1** | UP | P21958 | 2 | 4 | 2 | 724 | 78.8 | 8.79 | 6 | 2.308 | 0.266 | 4.003 | 0.008 | 1.145 | 0.728 | 3.495 | 0.011 |
| 3195 | **Mylpf** | Unchanged | P97457 | 1 | 1 | 1 | 169 | 18.9 | 4.92 | 6 | 0.250 | 0.369 | 0.571 | 0.618 | 0.575 | 0.624 | 0.993 | 0.992 |
| 3196 | **Mrps18b** | Unchanged | Q99N84 | 1 | 2 | 1 | 254 | 28.7 | 8.35 | 6 | #DIV/0! | #DIV/0! | #DIV/0! | #DIV/0! | #DIV/0! | #DIV/0! | #DIV/0! | #DIV/0! |
| 3197 | **Tjp1** | Unchanged | P39447 | 6 | 18 | 6 | 1745 | 194.6 | 6.64 | 6 | 0.910 | 0.801 | 1.011 | 0.981 | 0.870 | 0.640 | 1.162 | 0.730 |
| 3198 | **Fbp2** | Unchanged | P70695 | 1 | 1 | 1 | 339 | 36.9 | 6.16 | 6 | 0.467 | 0.502 | 0.590 | 0.620 | 3.191 | 0.405 | 0.185 | 0.312 |
| 3199 | **Pcbd2** | Unchanged | Q9CZL5 | 1 | 4 | 1 | 136 | 14.8 | 9.16 | 6 | 0.598 | 0.512 | 0.875 | 0.851 | 0.080 | 0.138 | 10.881 | 0.037 |
| 3200 | **Large2** | Unchanged | A2AHG6 | 1 | 1 | 1 | 715 | 82.3 | 7.2 | 6 | #DIV/0! | #DIV/0! | #DIV/0! | #DIV/0! | #DIV/0! | #DIV/0! | #DIV/0! | #DIV/0! |
| 3201 | **Skap2** | Unchanged | Q3UND0 | 1 | 2 | 1 | 358 | 40.7 | 4.7 | 6 | #DIV/0! | #DIV/0! | #DIV/0! | #DIV/0! | #DIV/0! | #DIV/0! | #DIV/0! | #DIV/0! |
| 3202 | **Glyr1** | Unchanged | Q922P9 | 2 | 17 | 2 | 546 | 59.7 | 9.22 | 6 | 1.184 | 0.675 | 2.538 | 0.213 | 1.297 | 0.583 | 1.957 | 0.331 |
| 3203 | **Mbl1** | Down | P39039 | 1 | 17 | 1 | 239 | 25.4 | 7.61 | 6 | 0.320 | 0.113 | 0.346 | 0.145 | 0.120 | 0.034 | 2.875 | 0.330 |
| 3204 | **Agfg2** | Unchanged | Q80WC7 | 2 | 11 | 2 | 479 | 48.9 | 9.1 | 6 | 1.413 | 0.428 | 5.231 | 0.064 | 1.102 | 0.821 | 4.748 | 0.074 |
| 3205 | **Trim30d** | Unchanged | E9PWL0 | 1 | 2 | 1 | 497 | 57.5 | 5.94 | 6 | 2.836 | 0.089 | 1.359 | 0.661 | 2.527 | 0.151 | 0.538 | 0.291 |
| 3206 | **Acin1** | Unchanged | Q9JIX8 | 5 | 27 | 5 | 1338 | 150.6 | 5.91 | 6 | 0.982 | 0.938 | 1.461 | 0.115 | 0.821 | 0.489 | 1.781 | 0.018 |
| 3207 | **Afdn** | Unchanged | Q9QZQ1 | 6 | 20 | 6 | 1820 | 206.4 | 6.32 | 6 | 0.885 | 0.786 | 0.612 | 0.365 | 0.332 | 0.112 | 1.843 | 0.211 |
| 3208 | **Clcn7** | Unchanged | F6SUM2 | 2 | 4 | 2 | 860 | 94.8 | 7.24 | 6 | 2.242 | 0.322 | 0.946 | 0.881 | 1.018 | 0.969 | 0.929 | 0.851 |
| 3209 | **Pik3r1** | Unchanged | P26450 | 2 | 5 | 2 | 724 | 83.5 | 6.28 | 6 | 0.819 | 0.678 | 0.569 | 0.255 | 0.863 | 0.724 | 0.660 | 0.230 |
| 3210 | **Krt10** | Unchanged | P02535 | 3 | 21 | 3 | 570 | 57.7 | 5.11 | 6 | 1.914 | 0.375 | 0.802 | 0.659 | 0.442 | 0.125 | 1.813 | 0.292 |
| 3211 | **Ghitm** | Unchanged | Q91VC9 | 1 | 9 | 1 | 346 | 37.3 | 9.8 | 6 | 1.290 | 0.641 | 0.852 | 0.747 | 1.173 | 0.733 | 0.727 | 0.368 |
| 3212 | **Acot7** | Unchanged | Q91V12 | 2 | 27 | 2 | 381 | 42.5 | 8.68 | 6 | 1.555 | 0.582 | 2.388 | 0.364 | 1.708 | 0.513 | 1.398 | 0.681 |
| 3213 | **Pld4** | Down | Q8BG07 | 2 | 5 | 2 | 503 | 56.1 | 7.31 | 6 | 0.302 | 0.001 | 0.283 | 0.001 | 0.143 | 0.000 | 1.976 | 0.178 |
| 3214 | **Sema6b** | Down | O54951 | 2 | 2 | 2 | 886 | 95.4 | 8.29 | 6 | 2.133 | 0.295 | 0.966 | 0.945 | 0.149 | 0.011 | 6.480 | 0.082 |
| 3215 | **Snx27** | Unchanged | Q3UHD6 | 2 | 4 | 2 | 539 | 61 | 6.4 | 6 | 1.734 | 0.219 | 3.056 | 0.345 | 0.746 | 0.427 | 4.098 | 0.288 |
| 3216 | **Arrb1** | Unchanged | Q8BWG8 | 1 | 16 | 1 | 418 | 46.9 | 6.28 | 6 | 1.497 | 0.594 | 1.471 | 0.615 | 1.728 | 0.300 | 0.851 | 0.780 |
| 3217 | **Ncf2** | Unchanged | O70145 | 2 | 2 | 2 | 525 | 59.4 | 6.61 | 6 | 0.859 | 0.763 | 1.046 | 0.909 | 1.565 | 0.211 | 0.668 | 0.201 |
| 3218 | **Tspan31** | Unchanged | Q9CQ88 | 1 | 1 | 1 | 210 | 22.7 | 8.44 | 6 | #DIV/0! | #DIV/0! | #DIV/0! | #DIV/0! | #DIV/0! | #DIV/0! | #DIV/0! | #DIV/0! |
| 3219 | **Gm5795** | Unchanged | E9PZN8 | 2 | 6 | 2 | 263 | 30.9 | 8.16 | 6 | 1.833 | 0.381 | 3.168 | 0.143 | 1.552 | 0.445 | 2.041 | 0.275 |
| 3220 | **Ctrb1** | Unchanged | Q9CR35 | 1 | 1 | 1 | 263 | 27.8 | 5.07 | 6 | 1.000 | 1.000 | 0.677 | 0.334 | 1.788 | 0.168 | 0.378 | 0.053 |
| 3221 | **Olfr1150-ps1** | Unchanged | A0A1L1STL1 | 1 | 2 | 1 | 316 | 35.5 | 8.62 | 6 | 0.618 | 0.537 | 0.617 | 0.518 | 0.460 | 0.363 | 1.342 | 0.415 |
| 3222 | **Gm1587** | Unchanged | Q3UT80 | 1 | 3 | 1 | 123 | 14.3 | 8.51 | 6 | 1.500 | 0.589 | 2.424 | 0.232 | 2.394 | 0.229 | 1.013 | 0.983 |
| 3223 | **Atxn10** | Unchanged | P28658 | 2 | 3 | 2 | 475 | 53.7 | 5.25 | 6 | 1.830 | 0.397 | 3.493 | 0.184 | 0.922 | 0.788 | 3.789 | 0.176 |
| 3224 | **Rint1** | Unchanged | Q8BZ36 | 2 | 3 | 2 | 792 | 90 | 5.14 | 6 | 0.711 | 0.126 | 1.121 | 0.611 | 1.018 | 0.943 | 1.102 | 0.741 |
| 3225 | **Dnajc7** | Unchanged | Q9QYI3 | 2 | 3 | 2 | 494 | 56.4 | 6.49 | 6 | 1.103 | 0.877 | 1.706 | 0.415 | 0.843 | 0.789 | 2.025 | 0.268 |
| 3226 | **Tjp2** | Unchanged | Q9Z0U1 | 5 | 7 | 4 | 1167 | 131.2 | 6.79 | 4 | 0.430 | 0.087 | 0.320 | 0.050 | 0.397 | 0.083 | 0.806 | 0.497 |
| 3227 | **Optn** | Unchanged | Q8K3K8 | 2 | 8 | 2 | 584 | 67 | 5.26 | 6 | 4.455 | 0.150 | 2.645 | 0.171 | 1.015 | 0.973 | 2.605 | 0.193 |
| 3228 | **Slc17a2** | Unchanged | Q5SZA1 | 1 | 1 | 1 | 478 | 51.9 | 8.63 | 6 | 0.810 | 0.791 | 0.764 | 0.744 | 1.498 | 0.525 | 0.510 | 0.195 |

| | | | | | | | | | | | | | | | | | | |
|---|---|---|---|---|---|---|---|---|---|---|---|---|---|---|---|---|---|---|
| 3229 | **Armc8** | Unchanged | Q9DBR3 | 2 | 8 | 2 | 673 | 75.3 | 6.73 | 6 | 1.627 | 0.162 | 2.126 | 0.128 | 1.015 | 0.968 | 2.095 | 0.125 |
| 3230 | **Nudt16** | Unchanged | Q6P3D0 | 1 | 6 | 1 | 195 | 21.8 | 7.12 | 6 | 0.984 | 0.979 | 2.591 | 0.216 | 0.249 | 0.083 | 10.421 | 0.071 |
| 3231 | **Sco1** | Down | Q5SUC9 | 1 | 2 | 1 | 284 | 31.6 | 8.47 | 6 | 0.854 | 0.706 | 1.260 | 0.547 | 0.136 | 0.007 | 9.252 | 0.013 |
| 3232 | **Tnpo3** | Unchanged | Q6P2B1 | 3 | 8 | 3 | 923 | 104.1 | 5.57 | 6 | 5.452 | 0.342 | 2.338 | 0.114 | 1.526 | 0.415 | 1.532 | 0.362 |
| 3233 | **Dcakd** | Unchanged | Q8BHC4 | 1 | 8 | 1 | 231 | 26.5 | 9.58 | 6 | 0.950 | 0.939 | 3.368 | 0.179 | 2.193 | 0.173 | 1.535 | 0.529 |
| 3234 | **Ddx46** | Unchanged | Q569Z5 | 4 | 12 | 4 | 1032 | 117.4 | 9.26 | 6 | 1.655 | 0.523 | 0.714 | 0.517 | 1.204 | 0.712 | 0.593 | 0.312 |
| 3235 | **Tcf7l2** | UP | D3Z002 | 1 | 1 | 1 | 178 | 19.7 | 9.33 | 6 | 2.393 | 0.311 | 7.350 | 0.016 | 1.363 | 0.573 | 5.394 | 0.024 |
| 3236 | **Nae1** | Down | Q8VBW6 | 2 | 14 | 2 | 534 | 60.2 | 5.52 | 6 | 0.451 | 0.033 | 0.828 | 0.719 | 0.352 | 0.020 | 2.355 | 0.308 |
| 3237 | **Cdk4** | Unchanged | P30285 | 1 | 1 | 1 | 303 | 33.7 | 6.62 | 6 | #DIV/0! | #DIV/0! | #DIV/0! | #DIV/0! | #DIV/0! | #DIV/0! | #DIV/0! | #DIV/0! |
| 3238 | **Casp7** | Unchanged | P97864 | 1 | 1 | 1 | 303 | 34 | 6.33 | 6 | 1.269 | 0.579 | 0.859 | 0.800 | 2.170 | 0.227 | 0.396 | 0.157 |
| 3239 | **Gpatch2l** | Unchanged | Q6PE65 | 1 | 9 | 1 | 482 | 54 | 8.25 | 6 | 1.594 | 0.541 | 1.397 | 0.707 | 1.924 | 0.407 | 0.726 | 0.658 |
| 3240 | **Mbp** | UP | P04370 | 1 | 2 | 1 | 250 | 27.2 | 9.58 | 6 | 17.778 | 0.145 | 1.798 | 0.278 | 3.034 | 0.013 | 0.593 | 0.142 |
| 3241 | **Xrcc6** | Unchanged | P23475 | 2 | 4 | 2 | 608 | 69.4 | 6.79 | 6 | 1.614 | 0.259 | 2.527 | 0.112 | 1.803 | 0.131 | 1.401 | 0.470 |
| 3242 | **Stt3b** | Unchanged | Q3TDQ1 | 5 | 30 | 4 | 823 | 93.2 | 8.95 | 6 | 0.759 | 0.431 | 3.438 | 0.106 | 0.513 | 0.085 | 6.696 | 0.060 |
| 3243 | **Cnn2** | Unchanged | Q08093 | 1 | 1 | 1 | 305 | 33.1 | 7.62 | 6 | 1.591 | 0.522 | 0.309 | 0.165 | 0.733 | 0.621 | 0.421 | 0.139 |
| 3244 | **Taf15** | UP | Q8BQ46 | 3 | 17 | 1 | 557 | 58.6 | 8.4 | 6 | 14.834 | 0.052 | 3.401 | 0.011 | 26.326 | 0.019 | 0.129 | 0.029 |
| 3245 | **Larp4** | UP | Q8BWW4 | 2 | 8 | 2 | 719 | 79.7 | 6.39 | 6 | 5.511 | 0.077 | 6.309 | 0.142 | 3.362 | 0.016 | 1.877 | 0.404 |
| 3246 | **Gtpbp1** | Unchanged | O08582 | 2 | 5 | 2 | 668 | 72.3 | 8.29 | 6 | 1.375 | 0.477 | 1.913 | 0.350 | 0.676 | 0.432 | 2.831 | 0.188 |
| 3247 | **Nup155** | Unchanged | Q99P88 | 4 | 28 | 4 | 1391 | 155 | 6.15 | 4 | 0.702 | 0.384 | 1.913 | 0.263 | 1.204 | 0.710 | 1.589 | 0.409 |
| 3248 | **Mvb12a** | Unchanged | Q78HU3 | 1 | 3 | 1 | 271 | 28.7 | 9.11 | 6 | #DIV/0! | #DIV/0! | #DIV/0! | #DIV/0! | #DIV/0! | #DIV/0! | #DIV/0! | #DIV/0! |
| 3249 | **Stx16** | Unchanged | Q8BVI5 | 1 | 4 | 1 | 326 | 37.1 | 5.86 | 6 | 0.796 | 0.633 | 1.779 | 0.382 | 1.484 | 0.496 | 1.199 | 0.764 |
| 3250 | **Hspa13** | Unchanged | Q8BM72 | 2 | 13 | 2 | 471 | 51.7 | 5.63 | 6 | 0.788 | 0.711 | 0.923 | 0.915 | 0.891 | 0.863 | 1.036 | 0.955 |
| 3251 | **Sumf2** | UP | Q8BPG6 | 1 | 1 | 1 | 308 | 34.7 | 7.11 | 6 | 4.411 | 0.135 | 8.635 | 0.003 | 0.992 | 0.984 | 8.707 | 0.003 |
| 3252 | **Plcb3** | Unchanged | P51432 | 3 | 8 | 3 | 1234 | 139.4 | 5.94 | 6 | 0.943 | 0.924 | 0.813 | 0.777 | 0.926 | 0.912 | 0.878 | 0.822 |
| 3253 | **Uba6** | Unchanged | Q8C7R4 | 4 | 13 | 4 | 1053 | 117.9 | 6.11 | 6 | 1.451 | 0.558 | 1.568 | 0.697 | 1.667 | 0.454 | 0.940 | 0.950 |
| 3254 | **Aasdhppt** | Unchanged | Q9CQF6 | 1 | 4 | 1 | 309 | 35.7 | 7.14 | 6 | 0.945 | 0.925 | 3.834 | 0.285 | 0.826 | 0.730 | 4.642 | 0.255 |
| 3255 | **Vps51** | Unchanged | Q3UVL4 | 2 | 10 | 2 | 782 | 86.1 | 6.24 | 5 | 1.301 | 0.300 | 1.908 | 0.087 | 1.379 | 0.102 | 1.383 | 0.318 |
| 3256 | **Myo5b** | Down | H3BLK6 | 1 | 2 | 1 | 128 | 14.9 | 5.01 | 5 | 0.246 | 0.038 | 0.657 | 0.450 | 0.550 | 0.252 | 1.194 | 0.790 |
| 3257 | **Ago3** | Unchanged | Q8CJF9 | 3 | 7 | 3 | 860 | 97.2 | 9.1 | 5 | 1.942 | 0.399 | 1.955 | 0.205 | 2.419 | 0.220 | 0.808 | 0.705 |
| 3258 | **Etnk2** | Unchanged | A7MCT6 | 1 | 4 | 1 | 385 | 44.5 | 7.21 | 5 | 1.158 | 0.864 | 0.433 | 0.393 | 0.872 | 0.842 | 0.497 | 0.294 |
| 3259 | **Sf3b4** | UP | Q8QZY9 | 1 | 7 | 1 | 424 | 44.3 | 8.56 | 5 | 4.900 | 0.289 | 17.013 | 0.043 | 8.336 | 0.153 | 2.041 | 0.316 |
| 3260 | **Slc25a16** | Unchanged | Q8C0K5 | 2 | 3 | 2 | 332 | 36.2 | 9.88 | 5 | 14.415 | 0.127 | 25.440 | 0.089 | 22.437 | 0.193 | 1.134 | 0.882 |
| 3261 | **Rab24** | Down | P35290 | 1 | 1 | 1 | 203 | 23.1 | 6.23 | 5 | 0.009 | 0.047 | 0.037 | 0.053 | 0.086 | 0.064 | 0.431 | 0.287 |
| 3262 | **Pdcl3** | Unchanged | Q8BVF2 | 1 | 1 | 1 | 240 | 27.6 | 4.72 | 5 | #DIV/0! | #DIV/0! | #DIV/0! | #DIV/0! | #DIV/0! | #DIV/0! | #DIV/0! | #DIV/0! |
| 3263 | **1700021F05Rik** | Unchanged | D3Z316 | 1 | 6 | 1 | 240 | 27.8 | 9.36 | 5 | 1.205 | 0.805 | 1.889 | 0.394 | 7.671 | 0.008 | 0.246 | 0.019 |
| 3264 | **Lipe** | Unchanged | E9Q4M2 | 3 | 5 | 3 | 1072 | 117.2 | 6.83 | 5 | 0.872 | 0.821 | 1.022 | 0.978 | 1.160 | 0.823 | 0.881 | 0.873 |
| 3265 | **Sart3** | Unchanged | Q9JLI8 | 3 | 7 | 3 | 962 | 109.6 | 5.24 | 5 | 1.133 | 0.776 | 1.461 | 0.488 | 1.000 | 1.000 | 1.461 | 0.486 |
| 3266 | **Trmt5** | Down | Q9D0C4 | 1 | 2 | 1 | 501 | 56.8 | 7.91 | 5 | 0.709 | 0.108 | 0.860 | 0.406 | 0.613 | 0.007 | 1.403 | 0.170 |
| 3267 | **Moxd1** | Unchanged | Q9CXI3 | 2 | 4 | 2 | 613 | 69.6 | 6.67 | 5 | 0.402 | 0.429 | 3.394 | 0.424 | 0.897 | 0.896 | 3.783 | 0.394 |
| 3268 | **Cyp2j12** | Unchanged | G3UXT0 | 2 | 9 | 1 | 502 | 57.7 | 7.4 | 5 | 0.243 | 0.168 | 0.023 | 0.059 | 0.059 | 0.068 | 0.381 | 0.148 |
| 3269 | **Rhox12** | Unchanged | Q4TU81 | 1 | 2 | 1 | 186 | 20.9 | 6.07 | 5 | 1.916 | 0.507 | 6.282 | 0.154 | 0.507 | 0.536 | 12.402 | 0.121 |

| | | | | | | | | | | | | | | | | | |
|---|---|---|---|---|---|---|---|---|---|---|---|---|---|---|---|---|---|
| 3270 | **Snta1** | Unchanged | Q61234 | 2 | 3 | 2 | 503 | 53.6 | 6.86 | 5 | 1.249 | 0.627 | 0.997 | 0.995 | 0.437 | 0.184 | 2.282 | 0.162 |
| 3271 | **Fam160b1** | Unchanged | Q8CDM8 | 2 | 3 | 2 | 764 | 86 | 5.25 | 5 | 0.979 | 0.956 | 0.734 | 0.455 | 0.440 | 0.080 | 1.666 | 0.188 |
| 3272 | **Drap1** | Unchanged | Q9D6N5 | 1 | 2 | 1 | 205 | 22.3 | 5.26 | 5 | 1.856 | 0.337 | 0.932 | 0.867 | 0.935 | 0.877 | 0.996 | 0.991 |
| 3273 | **mt-Co3** | Unchanged | P00416 | 1 | 25 | 1 | 261 | 29.9 | 7.3 | 5 | 0.809 | 0.757 | 0.378 | 0.257 | 0.390 | 0.295 | 0.969 | 0.976 |
| 3274 | **Serpina10** | UP | Q8R121 | 1 | 2 | 1 | 448 | 51.8 | 5.67 | 5 | 1.709 | 0.234 | 3.278 | 0.016 | 1.801 | 0.168 | 1.821 | 0.128 |
| 3275 | **Erc1** | Unchanged | Q99MI1 | 5 | 23 | 5 | 1120 | 128.3 | 5.87 | 5 | 1.283 | 0.463 | 1.182 | 0.700 | 0.991 | 0.987 | 1.193 | 0.781 |
| 3276 | **Vps26b** | Unchanged | Q8C0E2 | 1 | 4 | 1 | 336 | 39.1 | 7.37 | 5 | 0.169 | 0.382 | 0.113 | 0.351 | 0.217 | 0.408 | 0.520 | 0.057 |
| 3277 | **Cybc1** | Unchanged | Q3TYS2 | 1 | 2 | 1 | 187 | 20.9 | 8.59 | 5 | 1.293 | 0.211 | 1.828 | 0.231 | 1.449 | 0.259 | 1.262 | 0.592 |
| 3278 | **Itgb6** | Unchanged | Q9Z0T9 | 2 | 2 | 2 | 787 | 86 | 5.47 | 5 | 0.797 | 0.782 | 0.365 | 0.296 | 0.192 | 0.178 | 1.905 | 0.292 |
| 3279 | **Snx4** | Unchanged | Q91YJ2 | 2 | 9 | 2 | 450 | 51.7 | 5.8 | 5 | 0.872 | 0.804 | 1.702 | 0.201 | 0.620 | 0.463 | 2.745 | 0.004 |
| 3280 | **Esyt2** | Unchanged | Q3TZZ7 | 3 | 12 | 3 | 845 | 94.1 | 7.75 | 5 | 1.106 | 0.891 | 0.792 | 0.702 | 0.195 | 0.115 | 4.063 | 0.055 |
| 3281 | **Elp6** | Unchanged | A0A0H2UH13 | 1 | 1 | 1 | 263 | 28.7 | 6.98 | 5 | 0.851 | 0.806 | 0.335 | 0.246 | 0.366 | 0.270 | 0.916 | 0.845 |
| 3282 | **Cpsf7** | Unchanged | Q8BTV2 | 1 | 4 | 1 | 471 | 52 | 8 | 5 | 1.465 | 0.346 | 1.378 | 0.548 | 1.225 | 0.619 | 1.125 | 0.789 |
| 3283 | **Trappc11** | Unchanged | B2RXC1 | 3 | 5 | 3 | 1133 | 128.3 | 7.58 | 5 | 1.525 | 0.477 | 1.386 | 0.242 | 1.061 | 0.831 | 1.306 | 0.283 |
| 3284 | **Hfe** | Unchanged | P70387 | 1 | 1 | 1 | 359 | 40.5 | 6.24 | 5 | 1.830 | 0.425 | 4.596 | 0.111 | 1.188 | 0.585 | 3.869 | 0.127 |
| 3285 | **Letmd1** | Unchanged | Q924L1 | 1 | 2 | 1 | 360 | 41.7 | 10.54 | 5 | 1.114 | 0.759 | 0.712 | 0.468 | 0.445 | 0.055 | 1.600 | 0.523 |
| 3286 | **Gnl1** | Unchanged | P36916 | 2 | 4 | 2 | 607 | 68.7 | 5.68 | 5 | 3.592 | 0.078 | 1.080 | 0.869 | 2.356 | 0.401 | 0.459 | 0.440 |
| 3287 | **Afm** | Unchanged | O89020 | 2 | 5 | 2 | 608 | 69.3 | 5.78 | 5 | 0.392 | 0.098 | 0.535 | 0.216 | 0.411 | 0.104 | 1.302 | 0.555 |
| 3288 | **Apcdd1** | Unchanged | Q3U128 | 1 | 1 | 1 | 514 | 58.6 | 8.31 | 5 | 1.265 | 0.302 | 0.763 | 0.530 | 0.857 | 0.540 | 0.890 | 0.772 |
| 3289 | **Gm3854** | Unchanged | A0A1B0GSS6 | 1 | 1 | 1 | 495 | 57 | 9.25 | 5 | 3.002 | 0.321 | 1.645 | 0.566 | 0.279 | 0.238 | 5.894 | 0.176 |
| 3290 | **Colq** | Unchanged | O35348 | 1 | 1 | 1 | 457 | 47.7 | 8.51 | 5 | #DIV/0! | #DIV/0! | #DIV/0! | #DIV/0! | #DIV/0! | #DIV/0! | #DIV/0! | #DIV/0! |
| 3291 | **Krt2** | Unchanged | Q3TTY5 | 6 | 18 | 4 | 707 | 70.9 | 8.06 | 5 | 2.315 | 0.465 | 2.163 | 0.217 | 3.192 | 0.067 | 0.678 | 0.454 |
| 3292 | **Txndc15** | Down | Q6P6J9 | 1 | 4 | 1 | 344 | 38.1 | 4.77 | 5 | 1.189 | 0.615 | 3.436 | 0.238 | 0.404 | 0.028 | 8.499 | 0.151 |
| 3293 | **Armc10** | Unchanged | Q9D0L7 | 1 | 2 | 1 | 306 | 33.3 | 7.99 | 5 | #DIV/0! | #DIV/0! | #DIV/0! | #DIV/0! | #DIV/0! | #DIV/0! | #DIV/0! | #DIV/0! |
| 3294 | **Mrpl48** | Unchanged | E9QPQ8 | 1 | 1 | 1 | 211 | 24 | 9.52 | 5 | #DIV/0! | #DIV/0! | #DIV/0! | #DIV/0! | #DIV/0! | #DIV/0! | #DIV/0! | #DIV/0! |
| 3295 | **Galnt3** | Unchanged | P70419 | 1 | 4 | 1 | 633 | 72.9 | 8.25 | 5 | 2.683 | 0.134 | 0.638 | 0.471 | 4.463 | 0.180 | 0.143 | 0.138 |
| 3296 | **Huwe1** | Unchanged | Q7TMY8 | 11 | 34 | 11 | 4377 | 482.3 | 5.22 | 5 | 1.519 | 0.229 | 1.737 | 0.261 | 1.483 | 0.069 | 1.171 | 0.703 |
| 3297 | **Aars2** | Unchanged | Q14CH7 | 2 | 3 | 2 | 980 | 106.7 | 6.18 | 5 | #DIV/0! | #DIV/0! | #DIV/0! | #DIV/0! | #DIV/0! | #DIV/0! | #DIV/0! | #DIV/0! |
| 3298 | **Ktn1** | Unchanged | Q61595 | 4 | 9 | 4 | 1327 | 152.5 | 5.86 | 5 | 1.129 | 0.680 | 4.567 | 0.382 | 1.903 | 0.063 | 2.400 | 0.510 |
| 3299 | **Rnf121** | Unchanged | Q8R1Z9 | 1 | 2 | 1 | 327 | 38 | 7.06 | 5 | 0.575 | 0.224 | 0.550 | 0.306 | 0.562 | 0.204 | 0.978 | 0.966 |
| 3300 | **Hyal1** | Unchanged | Q91ZJ9 | 1 | 4 | 1 | 462 | 52.1 | 6.4 | 5 | 0.919 | 0.838 | 3.850 | 0.110 | 0.892 | 0.775 | 4.317 | 0.099 |
| 3301 | **Cog3** | Unchanged | E9QL65 | 1 | 4 | 1 | 828 | 94.1 | 5.72 | 5 | 0.682 | 0.448 | 0.386 | 0.139 | 0.617 | 0.341 | 0.625 | 0.388 |
| 3302 | **Ctbs** | Unchanged | Q8R242 | 1 | 2 | 1 | 366 | 41.3 | 5.6 | 5 | #DIV/0! | #DIV/0! | #DIV/0! | #DIV/0! | #DIV/0! | #DIV/0! | #DIV/0! | #DIV/0! |
| 3303 | **Tarsl2** | Unchanged | Q8BLY2 | 3 | 9 | 1 | 790 | 91.3 | 7.53 | 5 | #DIV/0! | #DIV/0! | #DIV/0! | #DIV/0! | #DIV/0! | #DIV/0! | #DIV/0! | #DIV/0! |
| 3304 | **Nedd4l** | Unchanged | Q8CFI0 | 4 | 7 | 1 | 1004 | 115.3 | 6.15 | 5 | 1.948 | 0.374 | 0.799 | 0.802 | 1.472 | 0.490 | 0.543 | 0.440 |
| 3305 | **Ccdc137** | Unchanged | Q8R0K4 | 1 | 1 | 1 | 290 | 32.9 | 10.68 | 5 | 0.685 | 0.693 | 0.100 | 0.136 | 3.912 | 0.143 | 0.026 | 0.057 |
| 3306 | **Wipi2** | Unchanged | Q80W47 | 1 | 1 | 1 | 445 | 48.4 | 5.86 | 5 | 0.704 | 0.614 | 1.123 | 0.862 | 0.327 | 0.218 | 3.438 | 0.131 |
| 3307 | **Ddx25** | Unchanged | Q9QY15 | 1 | 5 | 1 | 484 | 54.8 | 5.99 | 5 | 6.841 | 0.054 | 3.968 | 0.148 | 7.627 | 0.054 | 0.520 | 0.307 |
| 3308 | **Nup62cl** | Unchanged | A2AG10 | 1 | 1 | 1 | 272 | 31.2 | 4.68 | 5 | 0.590 | 0.115 | 3.482 | 0.185 | 2.175 | 0.128 | 1.601 | 0.497 |
| 3309 | **Gp1bb** | Unchanged | B2RR03 | 1 | 1 | 1 | 214 | 22.5 | 10.02 | 5 | 0.292 | 0.277 | 1.152 | 0.843 | 2.097 | 0.354 | 0.549 | 0.387 |
| 3310 | **Mfn1** | Unchanged | Q811U4 | 3 | 4 | 3 | 741 | 83.7 | 6.51 | 5 | 3.863 | 0.125 | 5.367 | 0.167 | 4.921 | 0.106 | 1.091 | 0.904 |

| | | | | | | | | | | | | | | | | | |
|---|---|---|---|---|---|---|---|---|---|---|---|---|---|---|---|---|---|
| 3311 | **Exosc2** | Unchanged | Q8VBV3 | 1 | 2 | 1 | 293 | 32.6 | 7.5 | 5 | 1.057 | 0.910 | 0.544 | 0.382 | 0.415 | 0.196 | 1.312 | 0.680 |
| 3312 | **Clip3** | Down | B9EHT4 | 1 | 1 | 1 | 547 | 59.5 | 7.94 | 5 | 0.424 | 0.049 | 0.572 | 0.112 | 0.320 | 0.012 | 1.788 | 0.097 |
| 3313 | **Sarg** | Unchanged | Q8BI29 | 1 | 1 | 1 | 606 | 65.1 | 9.14 | 5 | 1.851 | 0.203 | 2.699 | 0.202 | 1.044 | 0.869 | 2.584 | 0.211 |
| 3314 | **Abcg4** | Unchanged | Q91WA9 | 1 | 1 | 1 | 646 | 72.1 | 8.19 | 5 | 2.169 | 0.480 | 3.257 | 0.413 | 0.792 | 0.380 | 4.112 | 0.374 |
| 3315 | **Samsn1** | Unchanged | P57725 | 1 | 1 | 1 | 372 | 41.6 | 5.25 | 5 | 0.907 | 0.872 | 0.761 | 0.718 | 1.294 | 0.708 | 0.589 | 0.523 |
| 3316 | **Pde12** | UP | Q3TIU4 | 2 | 12 | 2 | 608 | 67.5 | 6.92 | 5 | 1.070 | 0.847 | 6.022 | 0.015 | 2.789 | 0.094 | 2.159 | 0.118 |
| 3317 | **Fcn1** | Unchanged | O70165 | 1 | 1 | 1 | 334 | 36.3 | 6.52 | 5 | #DIV/0! | #DIV/0! | #DIV/0! | #DIV/0! | #DIV/0! | #DIV/0! | #DIV/0! | #DIV/0! |
| 3318 | **Ndst2** | Unchanged | Q2TBA8 | 1 | 1 | 1 | 513 | 58.5 | 8.47 | 5 | 1.330 | 0.622 | 0.363 | 0.202 | 0.497 | 0.310 | 0.730 | 0.606 |
| 3319 | **Myd88** | UP | P22366 | 1 | 12 | 1 | 296 | 33.7 | 5.52 | 5 | 8.058 | 0.061 | 12.830 | 0.060 | 5.738 | 0.035 | 2.236 | 0.248 |
| 3320 | **Rrp1** | Unchanged | P56183 | 1 | 2 | 1 | 494 | 54.7 | 5.17 | 5 | 0.610 | 0.400 | 1.032 | 0.964 | 1.064 | 0.906 | 0.970 | 0.961 |
| 3321 | **Cpt1b** | Unchanged | Q924X2 | 3 | 4 | 2 | 772 | 88.2 | 8.38 | 5 | 22.935 | 0.062 | 21.582 | 0.085 | 0.891 | 0.862 | 24.224 | 0.084 |
| 3322 | **Anxa13** | Unchanged | Q99JG3 | 1 | 1 | 1 | 317 | 35.9 | 5.99 | 5 | #DIV/0! | #DIV/0! | #DIV/0! | #DIV/0! | #DIV/0! | #DIV/0! | #DIV/0! | #DIV/0! |
| 3323 | **Paf1** | Down | Q8K2T8 | 2 | 4 | 2 | 535 | 60.5 | 4.65 | 5 | 0.675 | 0.182 | 1.080 | 0.853 | 0.419 | 0.045 | 2.579 | 0.127 |
| 3324 | **Palld** | Unchanged | Q9ET54 | 3 | 5 | 3 | 1408 | 152 | 6.25 | 5 | 0.296 | 0.323 | 0.752 | 0.784 | 0.330 | 0.335 | 2.277 | 0.504 |
| 3325 | **Pcmtd1** | Unchanged | P59913 | 1 | 11 | 1 | 357 | 40.7 | 5.66 | 5 | 2.420 | 0.359 | 2.676 | 0.218 | 1.962 | 0.359 | 1.364 | 0.633 |
| 3326 | **Ggt6** | Unchanged | Q6PDE7 | 1 | 1 | 1 | 497 | 51.4 | 6.24 | 5 | 1.376 | 0.351 | 0.597 | 0.279 | 1.585 | 0.092 | 0.376 | 0.007 |
| 3327 | **Abcb8** | Unchanged | Q9CXJ4 | 2 | 12 | 2 | 717 | 78 | 9.07 | 5 | 1.879 | 0.292 | 4.204 | 0.175 | 2.808 | 0.077 | 1.497 | 0.552 |
| 3328 | **Lztfl1** | Unchanged | Q9JHQ5 | 1 | 1 | 1 | 299 | 34.8 | 5.17 | 5 | 2.612 | 0.390 | 3.028 | 0.460 | 4.894 | 0.157 | 0.619 | 0.613 |
| 3329 | **B4galt1** | Unchanged | P15535 | 2 | 6 | 2 | 399 | 44.4 | 9.38 | 5 | 0.609 | 0.324 | 0.956 | 0.930 | 0.994 | 0.992 | 0.962 | 0.943 |
| 3330 | **Smap1** | Unchanged | Q91VZ6 | 1 | 2 | 1 | 440 | 47.6 | 8.51 | 5 | 1.083 | 0.860 | 0.442 | 0.148 | 0.336 | 0.095 | 1.316 | 0.553 |
| 3331 | **Cyp2b10** | Unchanged | P12791 | 2 | 3 | 2 | 500 | 56.7 | 7.59 | 5 | 0.985 | 0.988 | 0.203 | 0.218 | 0.291 | 0.268 | 0.698 | 0.376 |
| 3332 | **Prpf4** | Unchanged | Q9DAW6 | 2 | 4 | 2 | 521 | 58.3 | 7.28 | 5 | 0.829 | 0.544 | 1.110 | 0.895 | 0.844 | 0.595 | 1.314 | 0.760 |
| 3333 | **Tap2** | Unchanged | P36371 | 1 | 1 | 1 | 702 | 77.4 | 7.24 | 5 | #DIV/0! | #DIV/0! | #DIV/0! | #DIV/0! | #DIV/0! | #DIV/0! | #DIV/0! | #DIV/0! |
| 3334 | **Qki** | Unchanged | Q9QYS9 | 1 | 1 | 1 | 341 | 37.6 | 8.56 | 5 | 0.724 | 0.691 | 3.082 | 0.314 | 4.888 | 0.053 | 0.631 | 0.483 |
| 3335 | **Appl2** | Unchanged | Q8K3G9 | 2 | 4 | 2 | 662 | 73.8 | 5.03 | 4 | 181.140 | 0.157 | 36.186 | 0.343 | 246.814 | 0.153 | 0.147 | 0.223 |
| 3336 | **Tra2a** | Unchanged | Q6PFR5 | 1 | 2 | 1 | 281 | 32.3 | 11.28 | 5 | 0.780 | 0.656 | 0.527 | 0.311 | 0.573 | 0.344 | 0.920 | 0.751 |
| 3337 | **Tars2** | Unchanged | Q3UQ84 | 3 | 9 | 3 | 723 | 81.6 | 7.87 | 5 | 1.069 | 0.893 | 2.099 | 0.336 | 0.602 | 0.368 | 3.484 | 0.199 |
| 3338 | **Atp2c2** | Unchanged | A7L9Z8 | 1 | 1 | 1 | 944 | 102.5 | 6.44 | 5 | 1.059 | 0.961 | 0.144 | 0.305 | 0.140 | 0.302 | 1.026 | 0.952 |
| 3339 | **Pfkp** | Unchanged | Q9WUA3 | 2 | 14 | 1 | 784 | 85.4 | 7.11 | 5 | 1.273 | 0.723 | 1.538 | 0.522 | 2.130 | 0.290 | 0.722 | 0.611 |
| 3340 | **Cyp7a1** | Unchanged | Q64505 | 2 | 7 | 2 | 503 | 57.2 | 8.78 | 5 | 0.717 | 0.635 | 0.930 | 0.916 | 1.565 | 0.433 | 0.594 | 0.371 |
| 3341 | **Papss1** | Unchanged | Q60967 | 2 | 13 | 1 | 624 | 70.7 | 6.77 | 5 | #DIV/0! | #DIV/0! | #DIV/0! | #DIV/0! | #DIV/0! | #DIV/0! | #DIV/0! | #DIV/0! |
| 3342 | **Nek7** | UP | Q9ES74 | 1 | 1 | 1 | 302 | 34.5 | 8.25 | 5 | 4.290 | 0.042 | 0.387 | 0.282 | 0.935 | 0.932 | 0.415 | 0.332 |
| 3343 | **Tnpo2** | Unchanged | Q99LG2 | 3 | 9 | 1 | 887 | 100.4 | 4.98 | 5 | 0.324 | 0.266 | 0.418 | 0.354 | 0.116 | 0.137 | 3.607 | 0.277 |
| 3344 | **Pds5a** | Unchanged | Q6A026 | 4 | 7 | 3 | 1332 | 150.2 | 7.85 | 5 | 0.640 | 0.118 | 1.092 | 0.877 | 0.722 | 0.260 | 1.511 | 0.521 |
| 3345 | **Degs1** | Unchanged | O09005 | 1 | 2 | 1 | 323 | 38.2 | 7.77 | 5 | 2.044 | 0.517 | 0.381 | 0.273 | 4.002 | 0.246 | 0.095 | 0.161 |
| 3346 | **Soat2** | Unchanged | O88908 | 2 | 2 | 2 | 525 | 60.6 | 8.81 | 5 | 0.307 | 0.130 | 0.258 | 0.112 | 2.018 | 0.211 | 0.128 | 0.024 |
| 3347 | **Ifi27l2b** | Unchanged | Q8VC49 | 1 | 1 | 1 | 283 | 27.8 | 4.7 | 5 | #DIV/0! | #DIV/0! | #DIV/0! | #DIV/0! | #DIV/0! | #DIV/0! | #DIV/0! | #DIV/0! |
| 3348 | **Gjb1** | UP | P28230 | 1 | 1 | 1 | 283 | 32 | 8.95 | 5 | 0.755 | 0.429 | 1.059 | 0.834 | 5.447 | 0.037 | 0.195 | 0.040 |
| 3349 | **Hook1** | Unchanged | Q8BIL5 | 2 | 3 | 2 | 728 | 84.4 | 5.22 | 5 | 0.949 | 0.862 | 12.950 | 0.268 | 1.128 | 0.669 | 11.478 | 0.273 |
| 3350 | **Ube3a** | Unchanged | O08759 | 3 | 7 | 3 | 870 | 99.8 | 5.08 | 5 | 2.998 | 0.254 | 2.082 | 0.376 | 0.602 | 0.436 | 3.457 | 0.220 |
| 3351 | **Vwf** | UP | S4R195 | 1 | 2 | 1 | 385 | 41.9 | 7.59 | 5 | 1.409 | 0.500 | 0.965 | 0.951 | 2.675 | 0.017 | 0.361 | 0.001 |

| | | | | | | | | | | | | | | | | | | |
|---|---|---|---|---|---|---|---|---|---|---|---|---|---|---|---|---|---|---|
| 3352 | **Rbm14** | Unchanged | Q8C2Q3 | 1 | 3 | 1 | 669 | 69.4 | 9.67 | 5 | 1.663 | 0.303 | 0.805 | 0.641 | 1.304 | 0.413 | 0.618 | 0.196 |
| 3353 | **Ppp1r3b** | Unchanged | Q8C767 | 1 | 1 | 1 | 284 | 32.4 | 5.87 | 5 | 0.312 | 0.281 | 2.484 | 0.326 | 0.397 | 0.346 | 6.260 | 0.148 |
| 3354 | **Zc3hav1** | Unchanged | D3Z5I1 | 3 | 8 | 3 | 996 | 112.5 | 8.07 | 5 | 1.738 | 0.489 | 0.587 | 0.323 | 0.884 | 0.757 | 0.665 | 0.340 |
| 3355 | **Ppp1r32** | Unchanged | A0A0R4J0F3 | 1 | 1 | 1 | 427 | 47.8 | 8.84 | 5 | 1.040 | 0.960 | 1.081 | 0.941 | 0.938 | 0.924 | 1.153 | 0.899 |
| 3356 | **Vps45** | Unchanged | P97390 | 2 | 4 | 2 | 570 | 65 | 8.25 | 5 | 0.933 | 0.734 | 1.941 | 0.290 | 1.038 | 0.810 | 1.870 | 0.305 |
| 3357 | **Cars2** | Unchanged | Q8BYM8 | 2 | 3 | 2 | 551 | 61.2 | 8.18 | 5 | #DIV/0! | #DIV/0! | #DIV/0! | #DIV/0! | #DIV/0! | #DIV/0! | #DIV/0! | #DIV/0! |
| 3358 | **Add1** | Unchanged | Q9QYC0 | 2 | 20 | 2 | 735 | 80.6 | 5.9 | 5 | 2.860 | 0.129 | 3.462 | 0.143 | 3.018 | 0.060 | 1.147 | 0.799 |
| 3359 | **Tifa** | Unchanged | Q793I8 | 1 | 1 | 1 | 184 | 21.5 | 5 | 5 | #DIV/0! | #DIV/0! | #DIV/0! | #DIV/0! | #DIV/0! | #DIV/0! | #DIV/0! | #DIV/0! |
| 3360 | **Il18rap** | Unchanged | Q9Z2B1 | 1 | 1 | 1 | 614 | 70 | 8.28 | 5 | 1.420 | 0.360 | 9.219 | 0.268 | 1.400 | 0.429 | 6.587 | 0.290 |
| 3361 | **Zw10** | Down | O54692 | 2 | 6 | 2 | 779 | 88 | 5.92 | 5 | 1.184 | 0.422 | 1.472 | 0.428 | 0.571 | 0.019 | 2.576 | 0.140 |
| 3362 | **Prmt5** | Unchanged | A0A0R4J049 | 1 | 1 | 1 | 637 | 72.7 | 6.34 | 5 | 1.193 | 0.616 | 3.833 | 0.156 | 1.111 | 0.767 | 3.450 | 0.170 |
| 3363 | **Tmem33** | Unchanged | Q9CR67 | 1 | 8 | 1 | 247 | 28 | 9.66 | 5 | 0.598 | 0.359 | 10.077 | 0.323 | 0.670 | 0.527 | 15.031 | 0.307 |
| 3364 | **2210010C04Rik** | Unchanged | Q9CPN9 | 1 | 16 | 1 | 247 | 26.4 | 7.93 | 5 | 0.716 | 0.261 | 1.144 | 0.711 | 0.567 | 0.043 | 2.016 | 0.130 |
| 3365 | **Uhrf2** | Unchanged | Q7TMI3 | 1 | 1 | 1 | 803 | 90 | 8.03 | 5 | 0.069 | 0.205 | 0.061 | 0.201 | 0.059 | 0.200 | 1.034 | 0.895 |
| 3366 | **Rbm42** | Unchanged | Q91V81 | 1 | 1 | 1 | 474 | 49.8 | 9.63 | 5 | 0.675 | 0.328 | 3.163 | 0.066 | 0.917 | 0.804 | 3.450 | 0.054 |
| 3367 | **Col18a1** | Down | P39061 | 4 | 14 | 4 | 1774 | 182.1 | 5.62 | 5 | 0.814 | 0.589 | 0.981 | 0.965 | 0.264 | 0.009 | 3.710 | 0.084 |
| 3368 | **Tbc1d8b** | Unchanged | A3KGB4 | 4 | 28 | 3 | 1114 | 127.8 | 5.49 | 5 | 0.466 | 0.130 | 0.422 | 0.184 | 0.352 | 0.078 | 1.199 | 0.850 |
| 3369 | **Mbd3l1** | Unchanged | Q9D9H3 | 1 | 9 | 1 | 186 | 20.7 | 8.06 | 5 | 1.605 | 0.505 | 1.579 | 0.587 | 1.162 | 0.806 | 1.359 | 0.691 |
| 3370 | **Scrib** | Unchanged | Q80U72 | 4 | 6 | 4 | 1612 | 174 | 5.12 | 5 | 0.924 | 0.822 | 1.273 | 0.474 | 1.082 | 0.819 | 1.176 | 0.696 |
| 3371 | **Hpgd** | UP | Q8VCC1 | 1 | 6 | 1 | 269 | 29.2 | 6.54 | 5 | 6.066 | 0.241 | 3.523 | 0.128 | 4.369 | 0.005 | 0.806 | 0.593 |
| 3372 | **Vps37c** | Unchanged | Q8R105 | 1 | 2 | 1 | 352 | 38.4 | 5.31 | 5 | 2.680 | 0.417 | 0.630 | 0.372 | 0.487 | 0.198 | 1.292 | 0.619 |
| 3373 | **Insl5** | Unchanged | A2AIY3 | 1 | 2 | 1 | 145 | 16.7 | 9.1 | 5 | 1.135 | 0.815 | 1.027 | 0.959 | 0.450 | 0.191 | 2.285 | 0.165 |
| 3374 | **Tpte** | Unchanged | G5E8H5 | 1 | 1 | 1 | 664 | 76.7 | 8.68 | 5 | 0.138 | 0.074 | 0.247 | 0.110 | 1.463 | 0.558 | 0.169 | 0.091 |
| 3375 | **Zfp345** | Unchanged | A2AQA1 | 1 | 1 | 1 | 561 | 66 | 9.28 | 5 | 3.005 | 0.361 | 6.654 | 0.088 | 1.787 | 0.152 | 3.724 | 0.136 |
| 3376 | **Sept4** | Unchanged | P28661 | 2 | 8 | 2 | 478 | 54.9 | 5.87 | 5 | 0.984 | 0.959 | 1.810 | 0.270 | 0.955 | 0.880 | 1.897 | 0.255 |
| 3377 | **Sorbs1** | Unchanged | Q62417 | 4 | 9 | 4 | 1290 | 143 | 8.25 | 5 | 0.862 | 0.735 | 0.634 | 0.296 | 0.566 | 0.240 | 1.121 | 0.738 |
| 3378 | **Irgq** | Unchanged | Q8VIM9 | 1 | 1 | 1 | 583 | 59.3 | 4.83 | 5 | #DIV/0! | #DIV/0! | #DIV/0! | #DIV/0! | #DIV/0! | #DIV/0! | #DIV/0! | #DIV/0! |
| 3379 | **Tmem150a** | Unchanged | Q91WN2 | 1 | 4 | 1 | 271 | 29 | 6.04 | 5 | 0.653 | 0.333 | 0.377 | 0.069 | 0.649 | 0.394 | 0.581 | 0.385 |
| 3380 | **Mcam** | Unchanged | D6RFP0 | 1 | 1 | 1 | 146 | 15.4 | 11.44 | 5 | 0.307 | 0.297 | 0.680 | 0.667 | 0.098 | 0.177 | 6.939 | 0.160 |
| 3381 | **Vash2** | Unchanged | Q8C5G2 | 1 | 5 | 1 | 355 | 40.5 | 9.98 | 5 | 21.907 | 0.055 | 0.145 | 0.207 | 26.091 | 0.054 | 0.006 | 0.047 |
| 3382 | **Nasp** | UP | B1AU75 | 2 | 3 | 2 | 773 | 84 | 4.37 | 5 | 0.549 | 0.212 | 1.021 | 0.951 | 2.960 | 0.044 | 0.345 | 0.047 |
| 3383 | **Tm7sf2** | Down | Q71KT5 | 2 | 21 | 2 | 418 | 46.5 | 8.28 | 5 | 0.475 | 0.079 | 0.281 | 0.021 | 0.239 | 0.008 | 1.179 | 0.756 |
| 3384 | **Proc** | Unchanged | P33587 | 2 | 2 | 2 | 460 | 51.8 | 6.35 | 5 | 1.241 | 0.715 | 0.659 | 0.411 | 0.376 | 0.090 | 1.755 | 0.358 |
| 3385 | **Erlec1** | Unchanged | Q8VEH8 | 1 | 1 | 1 | 483 | 54.9 | 6.25 | 5 | 0.357 | 0.089 | 0.453 | 0.135 | 0.728 | 0.427 | 0.623 | 0.094 |
| 3386 | **AW549877** | UP | Q8BR90 | 1 | 2 | 1 | 294 | 33.5 | 5.19 | 5 | 1.764 | 0.173 | 4.876 | 0.006 | 0.580 | 0.205 | 8.402 | 0.003 |
| 3387 | **Sarnp** | Unchanged | Q9D1J3 | 1 | 5 | 1 | 210 | 23.5 | 6.65 | 5 | 0.316 | 0.137 | 0.183 | 0.083 | 0.546 | 0.329 | 0.335 | 0.168 |
| 3388 | **Nqo1** | Unchanged | Q64669 | 1 | 1 | 1 | 274 | 30.9 | 8.72 | 5 | 1.821 | 0.464 | 4.540 | 0.172 | 2.342 | 0.206 | 1.938 | 0.399 |
| 3389 | **Hrc** | Unchanged | G5E8J6 | 1 | 1 | 1 | 738 | 85.1 | 4.63 | 5 | 1.136 | 0.611 | 0.850 | 0.658 | 0.378 | 0.050 | 2.249 | 0.099 |
| 3390 | **Klhl4** | UP | B1AZR0 | 1 | 1 | 1 | 717 | 80 | 7.05 | 5 | 740.167 | 0.045 | 86.167 | 0.108 | 839.333 | 0.072 | 0.103 | 0.102 |
| 3391 | **Ogfr** | Unchanged | Q99PG2 | 1 | 1 | 1 | 633 | 70.6 | 4.77 | 5 | 0.538 | 0.159 | 0.410 | 0.079 | 0.417 | 0.093 | 0.984 | 0.973 |
| 3392 | **Rela** | UP | Q04207 | 2 | 5 | 2 | 549 | 60.2 | 5.76 | 5 | 0.438 | 0.298 | 1.993 | 0.365 | 3.074 | 0.018 | 0.648 | 0.332 |

| | | | | | | | | | | | | | | | | | | |
|---|---|---|---|---|---|---|---|---|---|---|---|---|---|---|---|---|---|---|
| 3393 | **Nup50** | Unchanged | Q9JIH2 | 1 | 3 | 1 | 466 | 49.5 | 6.24 | 5 | 1.713 | 0.286 | 3.221 | 0.141 | 1.409 | 0.385 | 2.286 | 0.206 |
| 3394 | **Sqle** | Unchanged | P52019 | 2 | 6 | 2 | 572 | 63.7 | 8.47 | 5 | 0.623 | 0.593 | 0.886 | 0.873 | 0.440 | 0.406 | 2.014 | 0.283 |
| 3395 | **Bag5** | Unchanged | Q8CI32 | 1 | 2 | 1 | 447 | 50.9 | 6.05 | 5 | #DIV/0! | #DIV/0! | #DIV/0! | #DIV/0! | #DIV/0! | #DIV/0! | #DIV/0! | #DIV/0! |
| 3396 | **Naaa** | Unchanged | Q9D7V9 | 1 | 3 | 1 | 362 | 40.1 | 6.46 | 5 | #DIV/0! | #DIV/0! | #DIV/0! | #DIV/0! | #DIV/0! | #DIV/0! | #DIV/0! | #DIV/0! |
| 3397 | **Elac2** | Unchanged | Q80Y81 | 3 | 14 | 3 | 831 | 92.7 | 7.34 | 5 | 3.199 | 0.176 | 4.060 | 0.100 | 2.785 | 0.205 | 1.458 | 0.540 |
| 3398 | **Thoc6** | Unchanged | Q5U4D9 | 1 | 3 | 1 | 341 | 37.3 | 7.12 | 5 | 1.002 | 0.994 | 1.847 | 0.119 | 0.813 | 0.308 | 2.272 | 0.066 |
| 3399 | **Slc25a25** | Unchanged | A2ASZ8 | 1 | 2 | 1 | 469 | 52.6 | 8.54 | 5 | 1.014 | 0.978 | 1.987 | 0.382 | 0.761 | 0.377 | 2.611 | 0.277 |
| 3400 | **Tgfbi** | Unchanged | P82198 | 2 | 4 | 2 | 683 | 74.6 | 7.06 | 5 | #DIV/0! | #DIV/0! | #DIV/0! | #DIV/0! | #DIV/0! | #DIV/0! | #DIV/0! | #DIV/0! |
| 3401 | **Ncbp1** | Down | Q3UYV9 | 3 | 6 | 3 | 790 | 91.9 | 6.64 | 5 | 0.279 | 0.032 | 0.993 | 0.985 | 0.247 | 0.021 | 4.020 | 0.023 |
| 3402 | **Saraf** | Unchanged | A0A0R4J0D1 | 1 | 2 | 1 | 363 | 38.7 | 8.57 | 5 | #DIV/0! | #DIV/0! | #DIV/0! | #DIV/0! | #DIV/0! | #DIV/0! | #DIV/0! | #DIV/0! |
| 3403 | **Prpf4b** | Unchanged | Q61136 | 3 | 6 | 3 | 1007 | 116.9 | 10.23 | 5 | 2.604 | 0.226 | 2.127 | 0.422 | 3.491 | 0.194 | 0.609 | 0.530 |
| 3404 | **Amfr** | Unchanged | Q9R049 | 3 | 3 | 3 | 643 | 73.1 | 6.46 | 5 | 0.693 | 0.633 | 0.342 | 0.269 | 0.139 | 0.154 | 2.453 | 0.128 |
| 3405 | **Larp1** | UP | Q6ZQ58 | 4 | 21 | 4 | 1072 | 121.1 | 8.79 | 5 | 1.882 | 0.266 | 2.804 | 0.059 | 2.285 | 0.050 | 1.227 | 0.595 |
| 3406 | **Rpusd3** | Unchanged | Q14AI6 | 1 | 1 | 1 | 344 | 38.1 | 9.52 | 5 | 0.859 | 0.727 | 1.384 | 0.632 | 0.785 | 0.562 | 1.764 | 0.444 |
| 3407 | **Znf319** | Unchanged | Q9ERR8 | 1 | 1 | 1 | 581 | 65.6 | 8.37 | 5 | 1.473 | 0.414 | 3.422 | 0.203 | 0.947 | 0.905 | 3.615 | 0.182 |
| 3408 | **Krt42** | Down | Q6IFX2 | 2 | 10 | 2 | 452 | 50.1 | 5.16 | 5 | 0.559 | 0.231 | 0.530 | 0.212 | 0.234 | 0.040 | 2.268 | 0.119 |
| 3409 | **Pdss1** | Unchanged | Q33DR2 | 1 | 2 | 1 | 409 | 45.9 | 8.88 | 5 | 0.871 | 0.644 | 0.483 | 0.078 | 0.868 | 0.605 | 0.557 | 0.230 |
| 3410 | **Tdrkh** | Unchanged | Q80VL1 | 1 | 1 | 1 | 560 | 62.1 | 4.94 | 5 | 0.199 | 0.328 | 0.226 | 0.348 | 0.150 | 0.301 | 1.507 | 0.512 |
| 3411 | **Ppat** | Unchanged | Q8CIH9 | 2 | 6 | 2 | 517 | 57.4 | 6.96 | 5 | 0.772 | 0.614 | 0.856 | 0.824 | 0.816 | 0.694 | 1.050 | 0.949 |
| 3412 | **Erich6** | Unchanged | D3Z6S9 | 1 | 1 | 1 | 713 | 80.9 | 5.03 | 5 | 0.002 | 0.172 | 0.002 | 0.172 | 0.003 | 0.173 | 0.627 | 0.186 |
| 3413 | **Mmp28** | Unchanged | Q8CGV8 | 1 | 1 | 1 | 520 | 58.6 | 9.5 | 5 | #DIV/0! | #DIV/0! | #DIV/0! | #DIV/0! | #DIV/0! | #DIV/0! | #DIV/0! | #DIV/0! |
| 3414 | **Nup35** | Unchanged | Q8R4R6 | 1 | 2 | 1 | 325 | 34.8 | 9.25 | 5 | 0.149 | 0.065 | 0.480 | 0.321 | 0.167 | 0.069 | 2.877 | 0.314 |
| 3415 | **Ppp1r21** | Unchanged | Q3TDD9 | 2 | 3 | 2 | 780 | 88.3 | 6.9 | 5 | 0.821 | 0.562 | 1.212 | 0.318 | 0.813 | 0.308 | 1.491 | 0.014 |
| 3416 | **Fbxw11iso** | Unchanged | Q5SRY7 | 1 | 3 | 1 | 542 | 62 | 7.12 | 5 | 1.249 | 0.675 | 1.227 | 0.739 | 1.803 | 0.126 | 0.681 | 0.376 |
| 3417 | **Stab2** | Unchanged | Q8R4U0 | 6 | 11 | 6 | 2559 | 277.3 | 6.93 | 5 | 0.699 | 0.665 | 0.882 | 0.889 | 1.157 | 0.843 | 0.763 | 0.721 |
| 3418 | **Scyl1** | Unchanged | Q9EQC5 | 1 | 2 | 1 | 806 | 89.1 | 6.44 | 5 | 3.942 | 0.244 | 2.003 | 0.276 | 1.761 | 0.063 | 1.137 | 0.777 |
| 3419 | **Gm6569** | Unchanged | D3YTM9 | 1 | 2 | 1 | 175 | 18.6 | 8.19 | 5 | 2.166 | 0.464 | 2.504 | 0.334 | 0.287 | 0.242 | 8.712 | 0.140 |
| 3420 | **Tmem115** | Unchanged | Q9WUH1 | 1 | 6 | 1 | 350 | 38.1 | 7.5 | 5 | 1.316 | 0.672 | 1.332 | 0.625 | 0.926 | 0.897 | 1.438 | 0.557 |
| 3421 | **Abhd5** | Unchanged | Q9DBL9 | 1 | 3 | 1 | 351 | 39.1 | 6.79 | 5 | 1.244 | 0.563 | 1.257 | 0.562 | 0.705 | 0.387 | 1.783 | 0.165 |
| 3422 | **Snx18** | Unchanged | Q8C788 | 1 | 1 | 1 | 615 | 67.7 | 6.46 | 5 | #DIV/0! | #DIV/0! | #DIV/0! | #DIV/0! | #DIV/0! | #DIV/0! | #DIV/0! | #DIV/0! |
| 3423 | **Cd2ap** | Down | Q9JLQ0 | 2 | 7 | 2 | 637 | 70.4 | 6.38 | 5 | 0.536 | 0.153 | 1.183 | 0.616 | 0.091 | 0.007 | 12.994 | 0.002 |
| 3424 | **Utrn** | Unchanged | E9Q6R7 | 11 | 17 | 11 | 3430 | 392.5 | 5.33 | 5 | 1.127 | 0.658 | 3.166 | 0.263 | 0.734 | 0.354 | 4.315 | 0.214 |
| 3425 | **Serpina7** | Unchanged | P61939 | 1 | 2 | 1 | 418 | 47 | 6.54 | 5 | 0.576 | 0.556 | 1.489 | 0.710 | 0.221 | 0.234 | 6.738 | 0.289 |
| 3426 | **Lkaaear1** | Unchanged | Q8BIG2 | 1 | 1 | 1 | 198 | 22.1 | 10.29 | 5 | 1.522 | 0.430 | 1.721 | 0.311 | 1.184 | 0.627 | 1.453 | 0.377 |
| 3427 | **Gatd1** | Unchanged | Q8BFQ8 | 1 | 15 | 1 | 220 | 23.3 | 7.05 | 5 | 1.230 | 0.754 | 3.147 | 0.311 | 1.548 | 0.551 | 2.032 | 0.466 |
| 3428 | **Mcmbp** | Unchanged | H9KV04 | 1 | 1 | 1 | 660 | 74.5 | 5.47 | 5 | 1.185 | 0.856 | 2.305 | 0.186 | 0.560 | 0.440 | 4.117 | 0.048 |
| 3429 | **Safb2** | Unchanged | Q80YR5 | 2 | 15 | 2 | 991 | 111.8 | 6.38 | 5 | 0.706 | 0.557 | 0.817 | 0.711 | 0.719 | 0.551 | 1.136 | 0.820 |
| 3430 | **Tor1aip1** | Unchanged | Q921T2 | 2 | 6 | 2 | 595 | 66.7 | 7.05 | 5 | 0.889 | 0.845 | 1.303 | 0.723 | 1.091 | 0.888 | 1.194 | 0.816 |
| 3431 | **Dnajc13** | UP | D4AFX7 | 7 | 12 | 7 | 2248 | 254.9 | 6.77 | 5 | 1.201 | 0.417 | 1.838 | 0.035 | 1.290 | 0.053 | 1.425 | 0.146 |
| 3432 | **Rnf14** | Unchanged | Q9JI90 | 1 | 9 | 1 | 485 | 54.9 | 4.7 | 5 | 0.884 | 0.796 | 0.973 | 0.960 | 1.125 | 0.760 | 0.865 | 0.677 |
| 3433 | **Ppip5k2** | Unchanged | Q6ZQB6 | 2 | 9 | 2 | 1129 | 128.3 | 7.94 | 5 | 2.153 | 0.085 | 3.387 | 0.055 | 1.846 | 0.175 | 1.835 | 0.204 |

| | | | | | | | | | | | | | | | | | | |
|---|---|---|---|---|---|---|---|---|---|---|---|---|---|---|---|---|---|---|
| 3434 | **Lrrc42** | Unchanged | Q8R2U7 | 1 | 1 | 1 | 421 | 47.9 | 7.4 | 5 | 1.179 | 0.670 | 1.823 | 0.105 | 0.899 | 0.768 | 2.027 | 0.088 |
| 3435 | **Arhgap17** | Unchanged | Q3UIA2 | 2 | 3 | 2 | 846 | 92.1 | 7.56 | 4 | 0.721 | 0.411 | 0.942 | 0.892 | 1.418 | 0.451 | 0.665 | 0.413 |
| 3436 | **Exosc4** | Unchanged | Q921I9 | 1 | 6 | 1 | 245 | 26.2 | 6.15 | 4 | 0.747 | 0.498 | 1.830 | 0.375 | 0.738 | 0.137 | 2.480 | 0.254 |
| 3437 | **Tekt3** | UP | Q6X6Z7 | 1 | 1 | 1 | 490 | 56.6 | 7.05 | 4 | 2.275 | 0.221 | 2.001 | 0.029 | 0.714 | 0.326 | 2.803 | 0.005 |
| 3438 | **Hspbp1** | Unchanged | Q99P31 | 1 | 2 | 1 | 357 | 39.1 | 5.36 | 4 | #DIV/0! | #DIV/0! | #DIV/0! | #DIV/0! | #DIV/0! | #DIV/0! | #DIV/0! | #DIV/0! |
| 3439 | **Adck5** | UP | Q80V03 | 2 | 3 | 2 | 582 | 66.6 | 8.76 | 4 | 1.494 | 0.355 | 2.961 | 0.032 | 1.101 | 0.803 | 2.690 | 0.041 |
| 3440 | **St6gal1** | UP | Q64685 | 1 | 2 | 1 | 403 | 46.6 | 8.87 | 4 | 0.728 | 0.372 | 0.691 | 0.339 | 3.063 | 0.049 | 0.225 | 0.031 |
| 3441 | **Npc1** | Unchanged | O35604 | 4 | 22 | 4 | 1277 | 142.8 | 5.71 | 4 | 1.337 | 0.595 | 1.526 | 0.391 | 1.188 | 0.733 | 1.285 | 0.614 |
| 3442 | **Adap2** | Unchanged | Q8R2V5 | 1 | 3 | 1 | 381 | 44 | 9.14 | 4 | 1.685 | 0.294 | 3.225 | 0.140 | 1.456 | 0.373 | 2.216 | 0.225 |
| 3443 | **Ccdc7b** | UP | E9Q9Y3 | 1 | 1 | 1 | 381 | 44.2 | 9.72 | 4 | 43.719 | 0.149 | 25.514 | 0.129 | 66.753 | 0.036 | 0.382 | 0.205 |
| 3444 | **Srsf5** | Unchanged | O35326 | 1 | 7 | 1 | 269 | 30.9 | 11.56 | 4 | 1.283 | 0.641 | 1.300 | 0.762 | 1.078 | 0.887 | 1.206 | 0.834 |
| 3445 | **Mtnd5** | Unchanged | P03921 | 2 | 9 | 2 | 607 | 68.4 | 9.03 | 4 | 1.569 | 0.404 | 4.140 | 0.196 | 2.320 | 0.344 | 1.785 | 0.490 |
| 3446 | **Tomm40** | Unchanged | Q9QYA2 | 1 | 8 | 1 | 361 | 37.9 | 7.74 | 4 | 0.781 | 0.731 | 0.186 | 0.146 | 0.202 | 0.152 | 0.916 | 0.760 |
| 3447 | **Trappc6b** | Unchanged | Q9D289 | 1 | 10 | 1 | 158 | 17.9 | 8.68 | 4 | 0.868 | 0.818 | 0.606 | 0.477 | 0.674 | 0.524 | 0.899 | 0.881 |
| 3448 | **Ewsr1** | Unchanged | Q61545 | 2 | 12 | 2 | 655 | 68.4 | 9.33 | 4 | 1.034 | 0.965 | 0.956 | 0.948 | 1.064 | 0.919 | 0.898 | 0.883 |
| 3449 | **Rbm25** | Unchanged | B2RY56 | 2 | 11 | 2 | 838 | 99.5 | 6.32 | 4 | 1.474 | 0.377 | 0.570 | 0.293 | 0.654 | 0.303 | 0.873 | 0.796 |
| 3450 | **Gdpd5** | Unchanged | A0A1L1STK0 | 1 | 1 | 1 | 612 | 69.5 | 8.02 | 4 | 0.534 | 0.571 | 0.861 | 0.876 | 0.819 | 0.814 | 1.050 | 0.937 |
| 3451 | **Gabpa** | Unchanged | Q00422 | 2 | 3 | 2 | 454 | 51.3 | 5 | 4 | 1.111 | 0.810 | 1.434 | 0.192 | 0.957 | 0.857 | 1.498 | 0.128 |
| 3452 | **Ttc4** | Unchanged | Q8R3H9 | 1 | 1 | 1 | 386 | 44.3 | 5.22 | 4 | #DIV/0! | #DIV/0! | #DIV/0! | #DIV/0! | #DIV/0! | #DIV/0! | #DIV/0! | #DIV/0! |
| 3453 | **Ublcp1** | UP | Q8BGR9 | 1 | 4 | 1 | 318 | 36.8 | 6.46 | 4 | 0.523 | 0.264 | 5.149 | 0.018 | 3.794 | 0.180 | 1.357 | 0.575 |
| 3454 | **Ccny** | Unchanged | Q8BGU5 | 1 | 1 | 1 | 341 | 39.4 | 7.2 | 4 | 1.515 | 0.157 | 4.305 | 0.173 | 1.464 | 0.594 | 2.941 | 0.260 |
| 3455 | **Pspc1** | Down | Q8R326 | 1 | 4 | 1 | 523 | 58.7 | 6.67 | 4 | 1.645 | 0.204 | 1.898 | 0.286 | 0.423 | 0.032 | 4.491 | 0.090 |
| 3456 | **Lifr** | Unchanged | P42703 | 3 | 5 | 2 | 1092 | 122.5 | 6.04 | 4 | 0.905 | 0.826 | 0.334 | 0.094 | 0.386 | 0.139 | 0.866 | 0.785 |
| 3457 | **Golim4** | Down | D3YVW2 | 2 | 6 | 2 | 683 | 79.9 | 4.86 | 4 | 0.330 | 0.035 | 2.316 | 0.226 | 0.658 | 0.388 | 3.517 | 0.140 |
| 3458 | **Ltf** | Unchanged | P08071 | 2 | 5 | 2 | 707 | 77.8 | 8.53 | 4 | 0.917 | 0.772 | 1.603 | 0.343 | 11.679 | 0.117 | 0.137 | 0.137 |
| 3459 | **Cbl** | Unchanged | P22682 | 1 | 1 | 1 | 913 | 100.5 | 6.67 | 4 | 0.814 | 0.754 | 0.355 | 0.233 | 1.098 | 0.894 | 0.324 | 0.190 |
| 3460 | **Cnn1** | Unchanged | Q08091 | 1 | 1 | 1 | 297 | 33.3 | 8.97 | 4 | #DIV/0! | #DIV/0! | #DIV/0! | #DIV/0! | #DIV/0! | #DIV/0! | #DIV/0! | #DIV/0! |
| 3461 | **Pdcd2** | Unchanged | A0A0R4J0N5 | 1 | 1 | 1 | 343 | 38.3 | 5.38 | 4 | 1.368 | 0.448 | 2.300 | 0.164 | 1.501 | 0.310 | 1.532 | 0.393 |
| 3462 | **Cox15** | Unchanged | Q8BJ03 | 1 | 1 | 1 | 413 | 45.8 | 9.79 | 4 | 0.950 | 0.945 | 0.814 | 0.821 | 0.219 | 0.291 | 3.709 | 0.182 |
| 3463 | **Rock1** | Unchanged | P70335 | 3 | 6 | 2 | 1354 | 158.1 | 5.86 | 4 | 0.726 | 0.496 | 1.457 | 0.238 | 1.497 | 0.190 | 0.973 | 0.887 |
| 3464 | **Rspry1** | UP | Q8BVR6 | 1 | 1 | 1 | 576 | 64.3 | 5.5 | 4 | 0.815 | 0.690 | 1.354 | 0.396 | 2.026 | 0.008 | 0.668 | 0.042 |
| 3465 | **Fbln5** | Unchanged | A0A1Y7VJW9 | 2 | 3 | 2 | 461 | 51.7 | 4.78 | 4 | 0.999 | 0.997 | 0.623 | 0.101 | 1.675 | 0.190 | 0.372 | 0.051 |
| 3466 | **Cdk7** | Unchanged | Q03147 | 1 | 1 | 1 | 346 | 38.9 | 8.47 | 4 | #DIV/0! | #DIV/0! | #DIV/0! | #DIV/0! | #DIV/0! | #DIV/0! | #DIV/0! | #DIV/0! |
| 3467 | **Dcaf15** | UP | Q6PFH3 | 1 | 4 | 1 | 600 | 66.6 | 6.58 | 4 | 2.932 | 0.391 | 2.360 | 0.048 | 7.379 | 0.179 | 0.320 | 0.282 |
| 3468 | **Col4a3bp** | UP | Q9EQG9 | 2 | 4 | 2 | 624 | 71.1 | 5.44 | 4 | 3.300 | 0.067 | 1.461 | 0.389 | 9.802 | 0.024 | 0.149 | 0.031 |
| 3469 | **Ppp4r2** | Unchanged | A0A0R4J0U2 | 1 | 4 | 1 | 417 | 46.4 | 4.56 | 4 | 0.944 | 0.899 | 8.403 | 0.059 | 5.463 | 0.154 | 1.538 | 0.522 |
| 3470 | **Tdrd3** | Unchanged | Q91W18 | 1 | 1 | 1 | 743 | 82.2 | 9.11 | 4 | 0.458 | 0.437 | 0.386 | 0.378 | 4.828 | 0.178 | 0.080 | 0.115 |
| 3471 | **Prim1** | Unchanged | J3QN19 | 1 | 1 | 1 | 418 | 49.4 | 8.6 | 4 | 0.843 | 0.785 | 2.108 | 0.521 | 1.127 | 0.846 | 1.870 | 0.572 |
| 3472 | **Abhd16a** | Unchanged | Q9Z1Q2 | 2 | 2 | 2 | 558 | 63 | 8.25 | 4 | 0.628 | 0.693 | 0.654 | 0.706 | 0.040 | 0.294 | 16.184 | 0.030 |
| 3473 | **Mmrn1** | Unchanged | B2RPV6 | 2 | 2 | 2 | 1210 | 136 | 7.71 | 4 | 3.545 | 0.268 | 9.362 | 0.283 | 4.559 | 0.209 | 2.054 | 0.550 |
| 3474 | **Lamc2** | Unchanged | G5E874 | 2 | 4 | 2 | 1193 | 130.3 | 6.47 | 4 | 1.256 | 0.709 | 0.437 | 0.355 | 1.017 | 0.979 | 0.430 | 0.382 |

| | | | | | | | | | | | | | | | | | |
|---|---|---|---|---|---|---|---|---|---|---|---|---|---|---|---|---|---|
| 3475 | **Nectin1** | Unchanged | Q9JKF6 | 1 | 1 | 1 | 515 | 57 | 6.35 | 4 | 6.348 | 0.187 | 9.092 | 0.170 | 0.844 | 0.831 | 10.768 | 0.161 |
| 3476 | **Ppp4r3a** | Down | Q6P2K6 | 2 | 7 | 2 | 820 | 93.8 | 4.89 | 4 | 1.489 | 0.180 | 0.115 | 0.001 | 0.467 | 0.092 | 0.247 | 0.147 |
| 3477 | **Igfbp2** | Unchanged | P47877 | 1 | 2 | 1 | 305 | 32.8 | 7.64 | 4 | 1.766 | 0.484 | 3.057 | 0.352 | 0.199 | 0.204 | 15.335 | 0.191 |
| 3478 | **Tbc1d13** | Unchanged | Q8R3D1 | 1 | 1 | 1 | 400 | 46.4 | 5.36 | 4 | #DIV/0! | #DIV/0! | #DIV/0! | #DIV/0! | #DIV/0! | #DIV/0! | #DIV/0! | #DIV/0! |
| 3479 | **Lhx3** | Unchanged | P50481 | 1 | 1 | 1 | 400 | 44 | 7.71 | 4 | 1.113 | 0.772 | 1.614 | 0.233 | 0.628 | 0.200 | 2.569 | 0.039 |
| 3480 | **Slc7a6os** | Unchanged | Q7TPE5 | 1 | 3 | 1 | 306 | 35 | 4.58 | 4 | #DIV/0! | #DIV/0! | #DIV/0! | #DIV/0! | #DIV/0! | #DIV/0! | #DIV/0! | #DIV/0! |
| 3481 | **Gm17949** | Unchanged | A0A1B0GRV7 | 1 | 7 | 1 | 212 | 24.2 | 8.37 | 4 | 0.746 | 0.711 | 2.084 | 0.387 | 2.126 | 0.248 | 0.980 | 0.973 |
| 3482 | **Unc45a** | Unchanged | Q99KD5 | 2 | 21 | 2 | 944 | 103.4 | 6.34 | 4 | 0.626 | 0.168 | 1.017 | 0.974 | 0.928 | 0.808 | 1.096 | 0.854 |
| 3483 | **Trub2** | Unchanged | Q91WG3 | 1 | 1 | 1 | 331 | 36.8 | 8.72 | 4 | #DIV/0! | #DIV/0! | #DIV/0! | #DIV/0! | #DIV/0! | #DIV/0! | #DIV/0! | #DIV/0! |
| 3484 | **Rtn4** | Unchanged | Q99P72 | 4 | 11 | 4 | 1162 | 126.5 | 4.54 | 4 | 38.483 | 0.083 | 42.355 | 0.091 | 0.802 | 0.795 | 52.835 | 0.090 |
| 3485 | **Zfpl1** | Unchanged | Q9DB43 | 1 | 2 | 1 | 310 | 34.1 | 8.19 | 4 | #DIV/0! | #DIV/0! | #DIV/0! | #DIV/0! | #DIV/0! | #DIV/0! | #DIV/0! | #DIV/0! |
| 3486 | **Strip1** | Unchanged | Q8C079 | 1 | 1 | 1 | 837 | 95.5 | 6.25 | 4 | 1.757 | 0.200 | 1.769 | 0.244 | 1.517 | 0.375 | 1.166 | 0.728 |
| 3487 | **Gip** | Unchanged | P48756 | 1 | 1 | 1 | 144 | 16.4 | 7.3 | 4 | 0.008 | 0.070 | 0.106 | 0.098 | 0.009 | 0.070 | 12.155 | 0.129 |
| 3488 | **Dag1** | Unchanged | Q62165 | 2 | 2 | 2 | 893 | 96.8 | 8.44 | 4 | 0.772 | 0.647 | 0.511 | 0.262 | 0.477 | 0.202 | 1.072 | 0.867 |
| 3489 | **Ptprd** | Unchanged | Q64487 | 4 | 8 | 3 | 1912 | 214.3 | 6.55 | 4 | 0.421 | 0.242 | 0.240 | 0.133 | 0.819 | 0.701 | 0.293 | 0.009 |
| 3490 | **Bckdk** | Unchanged | O55028 | 1 | 8 | 1 | 412 | 46.6 | 8.91 | 4 | 1.154 | 0.656 | 2.021 | 0.210 | 1.973 | 0.064 | 1.025 | 0.956 |
| 3491 | **Hip1r** | Down | Q9JKY5 | 3 | 6 | 3 | 1068 | 119.4 | 6.52 | 4 | 0.505 | 0.047 | 0.442 | 0.016 | 0.375 | 0.010 | 1.178 | 0.692 |
| 3492 | **Nup85** | Unchanged | Q8R480 | 2 | 6 | 2 | 656 | 74.7 | 5.57 | 3 | 0.764 | 0.450 | 1.402 | 0.139 | 1.085 | 0.814 | 1.292 | 0.388 |
| 3493 | **Cideb** | Down | O70303 | 1 | 1 | 1 | 219 | 24.8 | 8.6 | 4 | 0.720 | 0.289 | 1.206 | 0.547 | 0.362 | 0.006 | 3.335 | 0.025 |
| 3494 | **Kiaa0391** | Unchanged | Q8JZY4 | 1 | 3 | 1 | 584 | 66.8 | 8.79 | 4 | 1.040 | 0.942 | 2.211 | 0.067 | 1.563 | 0.389 | 1.415 | 0.238 |
| 3495 | **Fgf8** | Unchanged | P37237 | 1 | 1 | 1 | 268 | 30.4 | 10.39 | 4 | 1.040 | 0.922 | 2.974 | 0.148 | 0.806 | 0.586 | 3.691 | 0.113 |
| 3496 | **Gopc** | Unchanged | K3W4Q9 | 1 | 1 | 1 | 463 | 50.6 | 6.35 | 4 | 1.106 | 0.871 | 2.331 | 0.188 | 2.387 | 0.126 | 0.976 | 0.961 |
| 3497 | **Herpud1** | Unchanged | Q9JJK5 | 1 | 1 | 1 | 391 | 43.9 | 5.53 | 4 | 1.362 | 0.401 | 0.880 | 0.791 | 0.962 | 0.928 | 0.915 | 0.884 |
| 3498 | **Pip4k2b** | Unchanged | Q80XI4 | 1 | 2 | 1 | 416 | 47.3 | 7.33 | 4 | #DIV/0! | #DIV/0! | #DIV/0! | #DIV/0! | #DIV/0! | #DIV/0! | #DIV/0! | #DIV/0! |
| 3499 | **Gm12845** | Unchanged | A0A0N4SUM5 | 1 | 1 | 1 | 588 | 65.8 | 8.09 | 4 | #DIV/0! | #DIV/0! | #DIV/0! | #DIV/0! | #DIV/0! | #DIV/0! | #DIV/0! | #DIV/0! |
| 3500 | **Slc4a4** | Unchanged | O88343 | 2 | 3 | 2 | 1079 | 121.4 | 6.84 | 4 | 0.783 | 0.575 | 1.511 | 0.580 | 3.138 | 0.153 | 0.482 | 0.328 |
| 3501 | **Pofut1** | Unchanged | Q91ZW2 | 1 | 1 | 1 | 393 | 44.7 | 8.41 | 4 | 0.875 | 0.777 | 0.441 | 0.185 | 0.291 | 0.074 | 1.519 | 0.650 |
| 3502 | **Sgcb** | Unchanged | P82349 | 1 | 1 | 1 | 320 | 34.9 | 8.6 | 4 | 1.104 | 0.828 | 1.076 | 0.869 | 2.066 | 0.467 | 0.521 | 0.510 |
| 3503 | **Sdr42e1** | Unchanged | Q9D665 | 1 | 2 | 1 | 394 | 43.9 | 6.8 | 4 | #DIV/0! | #DIV/0! | #DIV/0! | #DIV/0! | #DIV/0! | #DIV/0! | #DIV/0! | #DIV/0! |
| 3504 | **Arglu1** | UP | Q3UL36 | 1 | 1 | 1 | 271 | 32.9 | 10.36 | 4 | 1.907 | 0.250 | 1.134 | 0.854 | 202.144 | 0.048 | 0.006 | 0.048 |
| 3505 | **Sorbs2** | Unchanged | A0A0R4J1T1 | 3 | 3 | 3 | 1134 | 127.7 | 8.12 | 4 | 2.487 | 0.058 | 1.237 | 0.634 | 1.128 | 0.776 | 1.097 | 0.837 |
| 3506 | **Slc22a15** | Unchanged | Q504N2 | 1 | 1 | 1 | 544 | 60.2 | 6.19 | 4 | 0.811 | 0.637 | 1.020 | 0.957 | 1.105 | 0.784 | 0.924 | 0.817 |
| 3507 | **Tcea3** | Unchanged | P23881 | 1 | 2 | 1 | 347 | 38.8 | 9.33 | 4 | 1.042 | 0.952 | 0.703 | 0.704 | 1.810 | 0.292 | 0.389 | 0.191 |
| 3508 | **Sgk3** | Unchanged | Q9ERE3 | 1 | 1 | 1 | 496 | 57.1 | 7.15 | 4 | 2.368 | 0.359 | 0.204 | 0.125 | 2.063 | 0.313 | 0.099 | 0.066 |
| 3509 | **Il20ra** | Unchanged | Q6PHB0 | 1 | 1 | 1 | 546 | 61.9 | 5.02 | 4 | 0.742 | 0.737 | 0.818 | 0.774 | 0.286 | 0.148 | 2.856 | 0.326 |
| 3510 | **Ccar1** | Unchanged | Q8CH18 | 3 | 16 | 3 | 1146 | 132 | 5.76 | 4 | 1.493 | 0.340 | 2.062 | 0.190 | 1.343 | 0.596 | 1.536 | 0.460 |
| 3511 | **Tbcd** | Unchanged | Q8BYA0 | 2 | 3 | 2 | 1196 | 133.2 | 6.51 | 4 | 0.913 | 0.758 | 1.033 | 0.882 | 0.937 | 0.842 | 1.102 | 0.732 |
| 3512 | **Fam49b** | Unchanged | Q921M7 | 1 | 7 | 1 | 324 | 36.8 | 6.06 | 4 | 2.360 | 0.374 | 6.043 | 0.223 | 1.263 | 0.485 | 4.783 | 0.244 |
| 3513 | **Ckap4** | Unchanged | Q8BMK4 | 2 | 4 | 2 | 575 | 63.7 | 5.64 | 4 | 1.292 | 0.700 | 0.538 | 0.365 | 0.547 | 0.407 | 0.983 | 0.982 |
| 3514 | **Folr2** | Unchanged | Q05685 | 1 | 1 | 1 | 251 | 28.8 | 7.55 | 4 | #DIV/0! | #DIV/0! | #DIV/0! | #DIV/0! | #DIV/0! | #DIV/0! | #DIV/0! | #DIV/0! |
| 3515 | **Ribc2** | Unchanged | G3X8W0 | 1 | 1 | 1 | 377 | 45.2 | 9.28 | 4 | 2.444 | 0.292 | 7.746 | 0.068 | 5.448 | 0.108 | 1.422 | 0.584 |

| | | | | | | | | | | | | | | | | | | |
|---|---|---|---|---|---|---|---|---|---|---|---|---|---|---|---|---|---|---|
| 3516 | **Cog2** | Unchanged | Q921L5 | 1 | 4 | 1 | 731 | 82 | 6.21 | 4 | 0.503 | 0.069 | 4.262 | 0.059 | 1.426 | 0.364 | 2.989 | 0.099 |
| 3517 | **Trappc8** | Unchanged | E9PWG2 | 4 | 7 | 4 | 1437 | 160.8 | 7.01 | 4 | 1.468 | 0.268 | 2.284 | 0.201 | 0.933 | 0.782 | 2.447 | 0.173 |
| 3518 | **Mcm8** | Unchanged | Q9CWV1 | 1 | 1 | 1 | 833 | 92.3 | 7.62 | 4 | 2.255 | 0.306 | 1.944 | 0.635 | 2.212 | 0.358 | 0.879 | 0.907 |
| 3519 | **March5** | Unchanged | Q3KNM2 | 1 | 3 | 1 | 278 | 31.2 | 8.7 | | 0.875 | 0.715 | 0.954 | 0.925 | 1.909 | 0.141 | 0.500 | 0.181 |
| 3520 | **Ttyh2** | Unchanged | Q3TH73 | 1 | 2 | 1 | 532 | 59 | 6.07 | 4 | 0.673 | 0.304 | 1.295 | 0.618 | 1.943 | 0.298 | 0.667 | 0.528 |
| 3521 | **Bhlhe22** | Unchanged | A0A0R4J056 | 1 | 1 | 1 | 355 | 35.1 | 7.39 | 4 | 0.437 | 0.336 | 0.181 | 0.098 | 0.227 | 0.111 | 0.797 | 0.569 |
| 3522 | **Flii** | Unchanged | Q9JJ28 | 3 | 9 | 3 | 1271 | 144.7 | 6.06 | 4 | 1.008 | 0.981 | 1.365 | 0.481 | 0.933 | 0.779 | 1.463 | 0.396 |
| 3523 | **Impad1** | Unchanged | Q80V26 | 1 | 1 | 1 | 356 | 38.6 | 6.47 | 4 | 0.872 | 0.821 | 0.637 | 0.625 | 1.244 | 0.725 | 0.512 | 0.470 |
| 3524 | **Baiap2** | Unchanged | Q8BKX1 | 2 | 4 | 2 | 535 | 59.2 | 9.04 | 4 | 0.899 | 0.688 | 0.946 | 0.795 | 1.503 | 0.230 | 0.629 | 0.169 |
| 3525 | **H2-Ea** | Unchanged | P01904 | 1 | 2 | 1 | 255 | 29.1 | 5.12 | 4 | 0.368 | 0.286 | 0.051 | 0.113 | 0.695 | 0.626 | 0.073 | 0.051 |
| 3526 | **Vav2** | UP | Q60992 | 2 | 10 | 2 | 868 | 99.9 | 6.84 | 4 | 1.071 | 0.809 | 1.795 | 0.048 | 1.283 | 0.200 | 1.400 | 0.196 |
| 3527 | **Ndufaf6** | Unchanged | A2AIL4 | 1 | 1 | 1 | 333 | 38.3 | 9.44 | 4 | 1.005 | 0.996 | 1.147 | 0.860 | 2.190 | 0.202 | 0.523 | 0.100 |
| 3528 | **Wdfy1** | Unchanged | E9Q4P1 | 1 | 1 | 1 | 410 | 46.2 | 7.34 | 4 | 11.888 | 0.353 | 1.467 | 0.559 | 0.625 | 0.255 | 2.349 | 0.302 |
| 3529 | **Pafah1b3** | Unchanged | Q61205 | 1 | 4 | 1 | 232 | 25.8 | 6.93 | 4 | 1.184 | 0.670 | 2.778 | 0.200 | 2.531 | 0.200 | 1.097 | 0.883 |
| 3530 | **Nadk** | Unchanged | P58058 | 1 | 5 | 1 | 439 | 48.6 | 6.52 | 4 | 0.855 | 0.659 | 0.765 | 0.615 | 0.480 | 0.077 | 1.593 | 0.484 |
| 3531 | **Stxbp2** | Unchanged | F8WGM5 | 1 | 1 | 1 | 594 | 66.4 | 6.74 | 4 | 0.992 | 0.984 | 1.161 | 0.724 | 1.027 | 0.958 | 1.131 | 0.811 |
| 3532 | **Cybb** | Unchanged | Q61093 | 1 | 2 | 1 | 570 | 65.3 | 7.72 | 4 | 1.815 | 0.462 | 2.731 | 0.124 | 0.589 | 0.573 | 4.639 | 0.022 |
| 3533 | **Alkbh3** | Unchanged | A2AKV6 | 1 | 1 | 1 | 287 | 33.3 | 8.29 | 4 | #DIV/0! | #DIV/0! | #DIV/0! | #DIV/0! | #DIV/0! | #DIV/0! | #DIV/0! | #DIV/0! |
| 3534 | **Ikbkb** | Unchanged | O88351 | 2 | 6 | 2 | 757 | 86.6 | 6.4 | 4 | 0.807 | 0.729 | 1.197 | 0.724 | 1.243 | 0.723 | 0.963 | 0.937 |
| 3535 | **Luc7l2** | Unchanged | Q7TNC4 | 1 | 1 | 1 | 392 | 46.6 | 10.1 | 4 | #DIV/0! | #DIV/0! | #DIV/0! | #DIV/0! | #DIV/0! | #DIV/0! | #DIV/0! | #DIV/0! |
| 3536 | **Lrrc58** | UP | Q3UGP9 | 1 | 1 | 1 | 366 | 40.1 | 6.62 | 4 | 1.734 | 0.261 | 3.653 | 0.013 | 1.790 | 0.288 | 2.040 | 0.063 |
| 3537 | **Sgk2** | Unchanged | Q9QZS5 | 1 | 6 | 1 | 367 | 41.3 | 6.47 | 4 | 1.499 | 0.572 | 2.485 | 0.283 | 0.625 | 0.466 | 3.973 | 0.162 |
| 3538 | **Pdxdc1** | Unchanged | Q99K01 | 2 | 6 | 2 | 787 | 87.3 | 5.48 | 4 | 2.055 | 0.352 | 2.276 | 0.258 | 1.500 | 0.528 | 1.517 | 0.529 |
| 3539 | **Smad4** | Unchanged | P97471 | 1 | 3 | 1 | 551 | 60.3 | 6.99 | 4 | 1.755 | 0.085 | 1.824 | 0.179 | 0.663 | 0.420 | 2.750 | 0.080 |
| 3540 | **Pds5b** | Down | Q4VA53 | 4 | 21 | 2 | 1446 | 164.3 | 8.5 | 4 | 0.813 | 0.513 | 3.825 | 0.151 | 0.385 | 0.001 | 9.940 | 0.089 |
| 3541 | **Arfgef2** | Unchanged | A2A5R2 | 5 | 15 | 5 | 1792 | 202.1 | 6.55 | 4 | 1.086 | 0.766 | 2.216 | 0.158 | 0.738 | 0.199 | 3.003 | 0.093 |
| 3542 | **Fto** | Unchanged | Q8BGW1 | 1 | 1 | 1 | 502 | 58 | 5.12 | 4 | #DIV/0! | #DIV/0! | #DIV/0! | #DIV/0! | #DIV/0! | #DIV/0! | #DIV/0! | #DIV/0! |
| 3543 | **Pigs** | Unchanged | Q6PD26 | 1 | 2 | 1 | 555 | 61.7 | 6.93 | 4 | #DIV/0! | #DIV/0! | #DIV/0! | #DIV/0! | #DIV/0! | #DIV/0! | #DIV/0! | #DIV/0! |
| 3544 | **Zbtb6** | Unchanged | Q8K088 | 1 | 1 | 1 | 423 | 48.2 | 6.62 | 4 | #DIV/0! | #DIV/0! | #DIV/0! | #DIV/0! | #DIV/0! | #DIV/0! | #DIV/0! | #DIV/0! |
| 3545 | **Serpina6** | Down | Q06770 | 1 | 4 | 1 | 397 | 44.7 | 5.24 | 4 | 1.382 | 0.692 | 0.263 | 0.105 | 0.055 | 0.037 | 4.744 | 0.184 |
| 3546 | **Sv2a** | Unchanged | Q9JIS5 | 1 | 1 | 1 | 742 | 82.6 | 5.57 | 4 | 0.466 | 0.380 | 0.323 | 0.290 | 0.699 | 0.665 | 0.462 | 0.473 |
| 3547 | **Chmp7** | Unchanged | Q8R1T1 | 1 | 4 | 1 | 451 | 50.6 | 5.22 | 4 | 0.708 | 0.427 | 2.600 | 0.254 | 0.880 | 0.683 | 2.956 | 0.223 |
| 3548 | **Itm2b** | Unchanged | O89051 | 1 | 4 | 1 | 266 | 30.2 | 5.3 | 4 | 0.683 | 0.424 | 0.651 | 0.193 | 1.650 | 0.294 | 0.394 | 0.126 |
| 3549 | **Myo18a** | Unchanged | Q9JMH9 | 5 | 14 | 5 | 2050 | 232.6 | 6.28 | 4 | 1.089 | 0.858 | 0.651 | 0.279 | 0.971 | 0.938 | 0.671 | 0.347 |
| 3550 | **Pdcl2** | Unchanged | Q78Y63 | 1 | 1 | 1 | 240 | 27.8 | 4.89 | 4 | 1.707 | 0.498 | 2.385 | 0.372 | 0.204 | 0.075 | 11.714 | 0.162 |
| 3551 | **Akt2** | Unchanged | Q60823 | 2 | 7 | 2 | 481 | 55.7 | 6.37 | 4 | 0.753 | 0.508 | 1.513 | 0.513 | 0.721 | 0.532 | 2.098 | 0.314 |
| 3552 | **Smap2** | Unchanged | Q7TN29 | 1 | 1 | 1 | 428 | 46.5 | 8.87 | 4 | 1.014 | 0.984 | 0.365 | 0.130 | 0.159 | 0.056 | 2.291 | 0.041 |
| 3553 | **Exoc7** | UP | O35250 | 1 | 3 | 1 | 697 | 79.9 | 6.98 | 4 | 1.657 | 0.350 | 3.111 | 0.003 | 1.769 | 0.206 | 1.758 | 0.046 |
| 3554 | **Wdr4** | Unchanged | E9Q156 | 1 | 1 | 1 | 456 | 50.4 | 7.96 | 4 | 0.411 | 0.313 | 0.941 | 0.933 | 1.703 | 0.248 | 0.553 | 0.199 |
| 3555 | **Pde3b** | Unchanged | Q61409 | 2 | 2 | 2 | 1100 | 122.1 | 5.71 | 4 | 1.069 | 0.855 | 1.266 | 0.417 | 0.481 | 0.143 | 2.634 | 0.002 |
| 3556 | **Zfr** | Unchanged | O88532 | 3 | 4 | 3 | 1074 | 116.8 | 9.04 | 4 | 0.211 | 0.158 | 0.639 | 0.522 | 0.240 | 0.175 | 2.663 | 0.120 |

| | | | | | | | | | | | | | | | | | | |
|---|---|---|---|---|---|---|---|---|---|---|---|---|---|---|---|---|---|---|
| 3557 | **Agpat3** | Unchanged | Q9D517 | 1 | 7 | 1 | 376 | 43.3 | 8.51 | 4 | 1.070 | 0.924 | 2.681 | 0.256 | 1.296 | 0.659 | 2.069 | 0.314 |
| 3558 | **Dpp9** | Unchanged | Q8BVG4 | 2 | 16 | 2 | 862 | 97.9 | 6.65 | 4 | 3.425 | 0.159 | 4.482 | 0.174 | 3.616 | 0.115 | 1.240 | 0.758 |
| 3559 | **Tctn1** | Unchanged | Q8BZ64 | 1 | 1 | 1 | 593 | 63.4 | 7.28 | 4 | 2.797 | 0.256 | 4.726 | 0.099 | 1.877 | 0.299 | 2.517 | 0.212 |
| 3560 | **Gucy1b1** | Unchanged | O54865 | 2 | 4 | 2 | 620 | 70.6 | 5.31 | 4 | 1.531 | 0.134 | 1.218 | 0.701 | 0.839 | 0.569 | 1.451 | 0.532 |
| 3561 | **Flnc** | Unchanged | Q8VHX6 | 6 | 40 | 1 | 2726 | 290.9 | 5.95 | 4 | 0.565 | 0.556 | 0.248 | 0.243 | 6.049 | 0.167 | 0.041 | 0.114 |
| 3562 | **Ranbp2** | Unchanged | Q9ERU9 | 7 | 12 | 7 | 3053 | 340.9 | 6.18 | 4 | 1.080 | 0.896 | 1.060 | 0.924 | 1.007 | 0.990 | 1.053 | 0.900 |
| 3563 | **Atp6v0a1** | Unchanged | Q9Z1G4 | 2 | 5 | 2 | 839 | 96.4 | 6.76 | 4 | 0.546 | 0.305 | 1.998 | 0.342 | 0.687 | 0.496 | 2.908 | 0.198 |
| 3564 | **Yap1** | Unchanged | P46938 | 1 | 1 | 1 | 488 | 52.4 | 5.11 | 4 | 0.668 | 0.449 | 0.905 | 0.803 | 0.890 | 0.819 | 1.017 | 0.962 |
| 3565 | **Brd7** | Unchanged | O88665 | 1 | 1 | 1 | 651 | 74 | 6.38 | 4 | 0.770 | 0.778 | 0.407 | 0.377 | 0.981 | 0.982 | 0.415 | 0.333 |
| 3566 | **Cad** | Unchanged | B2RQC6 | 7 | 57 | 4 | 2225 | 243.1 | 6.43 | 4 | 0.990 | 0.983 | 1.298 | 0.605 | 1.135 | 0.806 | 1.144 | 0.795 |
| 3567 | **Nrip2** | Unchanged | A0A0N4SUN5 | 1 | 6 | 1 | 190 | 21 | 8.21 | 4 | 1.184 | 0.833 | 1.340 | 0.740 | 1.054 | 0.946 | 1.271 | 0.770 |
| 3568 | **Ehd2** | Unchanged | Q8BH64 | 2 | 3 | 1 | 543 | 61.1 | 6.51 | 4 | 2.628 | 0.411 | 2.346 | 0.170 | 3.595 | 0.338 | 0.653 | 0.651 |
| 3569 | **Col6a2** | Unchanged | Q02788 | 2 | 3 | 2 | 1034 | 110.3 | 6.42 | 4 | 1.566 | 0.418 | 7.073 | 0.076 | 1.975 | 0.157 | 3.581 | 0.132 |
| 3570 | **Abcb9 iso** | Unchanged | Q9JJ59 | 2 | 5 | 1 | 762 | 83.9 | 7.62 | 4 | 2.014 | 0.357 | 3.181 | 0.259 | 1.052 | 0.875 | 3.024 | 0.273 |
| 3571 | **Mrps34** | Unchanged | Q9JIK9 | 1 | 7 | 1 | 218 | 25.8 | 10.43 | 4 | 1.323 | 0.736 | 2.036 | 0.439 | 0.243 | 0.270 | 8.362 | 0.144 |
| 3572 | **Rab3gap1** | Unchanged | Q80UJ7 | 2 | 8 | 2 | 981 | 110.1 | 5.73 | 4 | 1.429 | 0.355 | 1.138 | 0.763 | 0.482 | 0.129 | 2.362 | 0.110 |
| 3573 | **Sigirr** | Unchanged | Q9JLZ8 | 1 | 2 | 1 | 409 | 46.1 | 5.74 | 4 | #DIV/0! | #DIV/0! | #DIV/0! | #DIV/0! | #DIV/0! | #DIV/0! | #DIV/0! | #DIV/0! |
| 3574 | **Ints11** | Unchanged | Q9CWS4 | 1 | 1 | 1 | 600 | 67.8 | 7.94 | 4 | 1.596 | 0.491 | 0.769 | 0.662 | 6.454 | 0.213 | 0.119 | 0.195 |
| 3575 | **Stim1** | Unchanged | P70302 | 2 | 3 | 2 | 685 | 77.5 | 6.54 | 4 | 0.792 | 0.645 | 2.819 | 0.238 | 0.701 | 0.487 | 4.019 | 0.163 |
| 3576 | **Prss50** | Unchanged | Q8BLH5 | 1 | 2 | 1 | 439 | 48.6 | 9.66 | 4 | 0.722 | 0.589 | 1.120 | 0.826 | 0.768 | 0.661 | 1.458 | 0.526 |
| 3577 | **Cstf2t** | Unchanged | Q8C7E9 | 1 | 2 | 1 | 632 | 65.8 | 7.25 | 4 | 0.886 | 0.800 | 0.923 | 0.853 | 1.408 | 0.324 | 0.656 | 0.188 |
| 3578 | **Cd300lg** | Unchanged | Q1ERP8 | 1 | 2 | 1 | 331 | 36.7 | 5.85 | 4 | 0.151 | 0.082 | 0.133 | 0.075 | 0.747 | 0.700 | 0.178 | 0.227 |
| 3579 | **Trmt10c** | Unchanged | Q3UFY8 | 1 | 1 | 1 | 414 | 48.4 | 9.38 | 4 | #DIV/0! | #DIV/0! | #DIV/0! | #DIV/0! | #DIV/0! | #DIV/0! | #DIV/0! | #DIV/0! |
| 3580 | **Wasf2** | Unchanged | Q8BH43 | 2 | 4 | 2 | 497 | 54 | 5.53 | 4 | 1.074 | 0.894 | 1.217 | 0.739 | 0.335 | 0.166 | 3.634 | 0.098 |
| 3581 | **Elp3** | Unchanged | A0A286YDB8 | 1 | 1 | 1 | 553 | 63 | 8 | 4 | 1.251 | 0.457 | 2.211 | 0.291 | 0.968 | 0.927 | 2.285 | 0.279 |
| 3582 | **Pitpnc1** | Unchanged | Q8K4R4 | 1 | 2 | 1 | 332 | 38.4 | 6.32 | 4 | 0.165 | 0.098 | 0.222 | 0.138 | 0.155 | 0.102 | 1.431 | 0.758 |
| 3583 | **Ptpn12** | Unchanged | P35831 | 2 | 3 | 2 | 775 | 86.5 | 5.99 | 4 | 0.890 | 0.584 | 0.784 | 0.160 | 0.558 | 0.067 | 1.406 | 0.340 |
| 3584 | **Tmem74** | Unchanged | Q8BQU7 | 1 | 1 | 1 | 305 | 33.4 | 4.77 | 4 | #DIV/0! | #DIV/0! | #DIV/0! | #DIV/0! | #DIV/0! | #DIV/0! | #DIV/0! | #DIV/0! |
| 3585 | **Kctd1** | Unchanged | E9Q6T9 | 1 | 1 | 1 | 861 | 96.1 | 5.81 | 4 | 0.978 | 0.975 | 0.456 | 0.428 | 0.517 | 0.478 | 0.882 | 0.753 |
| 3586 | **Sec24b** | Unchanged | Q80ZX0 | 3 | 6 | 3 | 1251 | 135.5 | 6.96 | 4 | 1.144 | 0.760 | 1.766 | 0.164 | 0.858 | 0.333 | 2.058 | 0.100 |
| 3587 | **Hk1** | Unchanged | P17710 | 3 | 13 | 3 | 974 | 108.2 | 6.8 | 4 | 0.965 | 0.944 | 1.341 | 0.521 | 1.295 | 0.477 | 1.035 | 0.917 |
| 3588 | **Mtpap** | Unchanged | Q9D0D3 | 2 | 5 | 2 | 585 | 65.2 | 8.88 | 4 | 0.810 | 0.418 | 1.048 | 0.904 | 1.061 | 0.750 | 0.988 | 0.973 |
| 3589 | **Tbx18** | Unchanged | G3X919 | 1 | 1 | 1 | 613 | 65.4 | 8.43 | 4 | 0.686 | 0.556 | 3.850 | 0.170 | 0.647 | 0.458 | 5.954 | 0.125 |
| 3590 | **Grwd1** | Unchanged | Q810D6 | 1 | 1 | 1 | 446 | 49.2 | 4.81 | 4 | 2.950 | 0.198 | 4.675 | 0.218 | 0.174 | 0.121 | 26.901 | 0.135 |
| 3591 | **Psme4** | Unchanged | Q5SSW2 | 3 | 5 | 3 | 1843 | 211.1 | 7.01 | 4 | 1.187 | 0.577 | 1.605 | 0.205 | 0.971 | 0.914 | 1.653 | 0.170 |
| 3592 | **Tpr** | Down | F6ZDS4 | 5 | 19 | 5 | 2431 | 273.8 | 5.03 | 4 | 0.786 | 0.418 | 0.342 | 0.030 | 1.290 | 0.302 | 0.265 | 0.000 |
| 3593 | **Pnpt1** | Unchanged | Q8K1R3 | 2 | 5 | 2 | 783 | 85.6 | 8.03 | 4 | 0.918 | 0.916 | 1.383 | 0.712 | 0.291 | 0.262 | 4.752 | 0.218 |
| 3594 | **Sptlc2** | UP | P97363 | 1 | 3 | 1 | 560 | 62.9 | 8.18 | 4 | 0.950 | 0.888 | 2.949 | 0.036 | 2.328 | 0.119 | 1.267 | 0.561 |
| 3595 | **Grpel2** | Unchanged | O88396 | 1 | 8 | 1 | 224 | 25 | 7.72 | 4 | 0.055 | 0.092 | 0.089 | 0.103 | 1.147 | 0.836 | 0.078 | 0.060 |
| 3596 | **Cep162** | Unchanged | Q6ZQ06 | 2 | 3 | 2 | 1403 | 160.8 | 5.53 | 4 | 1.577 | 0.352 | 1.204 | 0.734 | 0.338 | 0.129 | 3.560 | 0.119 |
| 3597 | **Rcc1** | Unchanged | Q8VE37 | 1 | 2 | 1 | 421 | 44.9 | 8.1 | 4 | 1.098 | 0.864 | 1.719 | 0.543 | 1.038 | 0.954 | 1.656 | 0.597 |

| | | | | | | | | | | | | | | | | | | |
|---|---|---|---|---|---|---|---|---|---|---|---|---|---|---|---|---|---|---|
| 3598 | **Cbwd1** | Unchanged | Q8VEH6 | 1 | 1 | 1 | 393 | 43.7 | 4.89 | 4 | #DIV/0! | #DIV/0! | #DIV/0! | #DIV/0! | #DIV/0! | #DIV/0! | #DIV/0! | #DIV/0! |
| 3599 | **Chid1** | UP | Q922Q9 | 1 | 3 | 1 | 393 | 44.9 | 7.97 | 4 | 8.777 | 0.044 | 0.365 | 0.014 | 1.111 | 0.774 | 0.329 | 0.069 |
| 3600 | **Syap1** | UP | Q9D5V6 | 2 | 2 | 2 | 365 | 41.3 | 4.54 | 4 | 9.183 | 0.027 | 6.897 | 0.175 | 8.048 | 0.034 | 0.857 | 0.817 |
| 3601 | **Atxn2l** | Unchanged | A0A0U1RPL0 | 2 | 2 | 2 | 1096 | 115.2 | 8.66 | 4 | 0.239 | 0.167 | 1.290 | 0.786 | 0.843 | 0.826 | 1.531 | 0.674 |
| 3602 | **Chkb** | Down | O55229 | 1 | 1 | 1 | 394 | 45.1 | 5.41 | 4 | 0.958 | 0.933 | 0.106 | 0.031 | 0.973 | 0.940 | 0.109 | 0.000 |
| 3603 | **Ctnnbl1** | Unchanged | Q9CWL8 | 2 | 5 | 2 | 563 | 64.9 | 5.07 | 4 | 0.747 | 0.617 | 1.676 | 0.344 | 0.266 | 0.160 | 6.291 | 0.021 |
| 3604 | **Cdkn2aip** | Unchanged | Q8BI72 | 1 | 8 | 1 | 563 | 59.7 | 9.16 | 4 | 1.851 | 0.607 | 3.911 | 0.190 | 0.559 | 0.530 | 7.001 | 0.128 |
| 3605 | **Xrcc5** | Unchanged | P27641 | 1 | 7 | 1 | 732 | 83 | 5.16 | 4 | 0.518 | 0.421 | 1.276 | 0.683 | 1.767 | 0.252 | 0.722 | 0.335 |
| 3606 | **Fbxo3** | Unchanged | Q9DC63 | 1 | 8 | 1 | 480 | 55.2 | 5.02 | 4 | 0.864 | 0.841 | 1.034 | 0.964 | 1.419 | 0.670 | 0.728 | 0.694 |
| 3607 | **Dhx58** | Unchanged | Q99J87 | 1 | 1 | 1 | 678 | 76.7 | 8.19 | 4 | #DIV/0! | #DIV/0! | #DIV/0! | #DIV/0! | #DIV/0! | #DIV/0! | #DIV/0! | #DIV/0! |
| 3608 | **Bcam** | Unchanged | Q9R069 | 1 | 1 | 1 | 622 | 67.6 | 6.25 | 4 | 0.407 | 0.082 | 0.817 | 0.605 | 0.729 | 0.274 | 1.121 | 0.802 |
| 3609 | **Dhx30** | Unchanged | Q99PU8 | 2 | 3 | 2 | 1217 | 136.6 | 8.75 | 4 | 0.992 | 0.969 | 1.440 | 0.294 | 0.861 | 0.548 | 1.673 | 0.158 |
| 3610 | **Gm18336** | Unchanged | A0A140LIU8 | 1 | 1 | 1 | 765 | 78.7 | 4.25 | 4 | 0.837 | 0.780 | 1.643 | 0.554 | 0.444 | 0.257 | 3.698 | 0.250 |
| 3611 | **Plxnb2** | Unchanged | B2RXS4 | 4 | 12 | 4 | 1842 | 206.1 | 5.87 | 4 | 1.000 | 1.000 | 0.853 | 0.742 | 0.376 | 0.134 | 2.269 | 0.059 |
| 3612 | **Pcdh1** | Unchanged | F7BJK1 | 3 | 3 | 3 | 1077 | 116.4 | 5.08 | 4 | 0.566 | 0.500 | 0.300 | 0.223 | 0.199 | 0.167 | 1.513 | 0.242 |
| 3613 | **Abhd12** | Unchanged | Q99LR1 | 1 | 1 | 1 | 398 | 45.2 | 8.72 | 4 | 6.679 | 0.252 | 3.970 | 0.080 | 1.279 | 0.505 | 3.105 | 0.112 |
| 3614 | **Ptk2** | Down | P34152 | 2 | 12 | 1 | 1052 | 119.2 | 6.62 | 4 | 0.628 | 0.041 | 0.895 | 0.430 | 0.541 | 0.038 | 1.655 | 0.123 |
| 3615 | **Lig4** | Unchanged | Q8BTF7 | 1 | 1 | 1 | 911 | 104.1 | 8.16 | 4 | 0.162 | 0.095 | 0.595 | 0.505 | 1.042 | 0.945 | 0.571 | 0.439 |
| 3616 | **Gtpbp8** | Unchanged | Q9CY28 | 1 | 2 | 1 | 285 | 31.9 | 9.28 | 4 | 1.148 | 0.672 | 0.435 | 0.123 | 1.195 | 0.644 | 0.364 | 0.046 |
| 3617 | **Apobr** | Unchanged | Q8VBT6 | 1 | 1 | 1 | 942 | 102.6 | 4.36 | 4 | 1.397 | 0.538 | 0.211 | 0.086 | 1.188 | 0.801 | 0.178 | 0.142 |
| 3618 | **Ppp1r12a** | Unchanged | Q9DBR7 | 2 | 8 | 2 | 1029 | 114.9 | 5.49 | 3 | 1.865 | 0.199 | 2.478 | 0.245 | 1.826 | 0.336 | 1.357 | 0.636 |
| 3619 | **Mtor** | Unchanged | Q9JLN9 | 5 | 8 | 5 | 2549 | 288.6 | 7.17 | 3 | 0.944 | 0.862 | 1.226 | 0.562 | 0.843 | 0.619 | 1.453 | 0.226 |
| 3620 | **Gpnmb** | UP | Q99P91 | 2 | 20 | 2 | 574 | 63.6 | 7.61 | 3 | 2.357 | 0.271 | 4.181 | 0.003 | 6.579 | 0.001 | 0.635 | 0.059 |
| 3621 | **1700029J07Rik** | Unchanged | Q3U1D9 | 1 | 4 | 1 | 316 | 35.2 | 9.73 | 3 | 1.406 | 0.654 | 0.641 | 0.672 | 1.959 | 0.362 | 0.327 | 0.250 |
| 3622 | **Rbms3** | Unchanged | Q8BWL5 | 1 | 4 | 1 | 431 | 46.9 | 7.99 | 3 | 0.783 | 0.660 | 0.559 | 0.376 | 0.283 | 0.086 | 1.979 | 0.426 |
| 3623 | **Mtus1** | Unchanged | Q5HZI1 | 3 | 9 | 3 | 1210 | 134.3 | 6.86 | 3 | 1.125 | 0.805 | 0.679 | 0.615 | 0.926 | 0.890 | 0.734 | 0.725 |
| 3624 | **Klhl26** | Unchanged | Q8BGY4 | 1 | 1 | 1 | 606 | 67 | 7.01 | 3 | 1.394 | 0.431 | 9.563 | 0.126 | 2.619 | 0.118 | 3.652 | 0.209 |
| 3625 | **Strn** | Unchanged | O55106 | 2 | 4 | 2 | 780 | 85.9 | 5.27 | 3 | 2.473 | 0.078 | 2.079 | 0.350 | 0.807 | 0.716 | 2.578 | 0.267 |
| 3626 | **Pip4k2a** | Unchanged | O70172 | 1 | 1 | 1 | 405 | 46.1 | 6.99 | 3 | #DIV/0! | #DIV/0! | #DIV/0! | #DIV/0! | #DIV/0! | #DIV/0! | #DIV/0! | #DIV/0! |
| 3627 | **Bend7** | Unchanged | Q8BSV3 | 1 | 1 | 1 | 434 | 48.4 | 9.41 | 3 | 1.040 | 0.954 | 2.525 | 0.192 | 2.707 | 0.212 | 0.933 | 0.911 |
| 3628 | **Arih2** | Unchanged | Q9Z1K6 | 1 | 4 | 1 | 492 | 57.7 | 5.69 | 3 | 0.284 | 0.273 | 1.763 | 0.530 | 1.316 | 0.819 | 1.340 | 0.780 |
| 3629 | **Nup205** | Unchanged | A0A0J9YUD5 | 3 | 3 | 3 | 2061 | 233 | 6.62 | 3 | 1.116 | 0.812 | 1.298 | 0.586 | 0.935 | 0.839 | 1.388 | 0.440 |
| 3630 | **Tns1** | Unchanged | E9Q0S6 | 4 | 6 | 3 | 1888 | 201.3 | 7.75 | 3 | 0.608 | 0.071 | 0.691 | 0.450 | 0.859 | 0.547 | 0.804 | 0.661 |
| 3631 | **Vps13c** | Unchanged | Q8BX70 | 8 | 13 | 8 | 3748 | 419.8 | 6.81 | 3 | 0.678 | 0.176 | 0.908 | 0.760 | 0.609 | 0.098 | 1.491 | 0.189 |
| 3632 | **Mcm9** | Unchanged | Q2KHI9 | 2 | 2 | 2 | 1134 | 125.7 | 7.37 | 3 | 0.598 | 0.257 | 0.465 | 0.161 | 0.745 | 0.468 | 0.625 | 0.273 |
| 3633 | **Elmo1** | Down | Q8BPU7 | 2 | 3 | 2 | 727 | 83.9 | 6.28 | 2 | 0.282 | 0.020 | 0.365 | 0.066 | 0.205 | 0.004 | 1.781 | 0.523 |
| 3634 | **Hectd1** | Down | Q69ZR2 | 6 | 19 | 6 | 2618 | 289.9 | 5.41 | 3 | 0.775 | 0.332 | 1.196 | 0.609 | 0.571 | 0.034 | 2.095 | 0.094 |
| 3635 | **Ca13** | Unchanged | Q9D6N1 | 1 | 4 | 1 | 262 | 29.5 | 6.65 | 3 | 0.873 | 0.825 | 0.362 | 0.213 | 0.731 | 0.613 | 0.495 | 0.354 |
| 3636 | **Naca** | Unchanged | P70670 | 7 | 144 | 7 | 2187 | 220.4 | 9.35 | 3 | 0.999 | 0.997 | 1.081 | 0.780 | 1.144 | 0.590 | 0.944 | 0.828 |
| 3637 | **Cabp1** | Unchanged | D3Z1M4 | 1 | 2 | 1 | 350 | 37.9 | 8.28 | 3 | 0.698 | 0.521 | 0.239 | 0.115 | 0.550 | 0.403 | 0.435 | 0.392 |
| 3638 | **Klhdc9** | Unchanged | Q3USL1 | 1 | 6 | 1 | 350 | 38.2 | 8.5 | 3 | 0.938 | 0.920 | 0.622 | 0.503 | 2.018 | 0.116 | 0.308 | 0.027 |

| | | | | | | | | | | | | | | | | | | |
|---|---|---|---|---|---|---|---|---|---|---|---|---|---|---|---|---|---|---|
| 3639 | **Atxn2** | Unchanged | O70305 | 3 | 8 | 3 | 1285 | 136.4 | 9.55 | 3 | 1.069 | 0.916 | 1.585 | 0.595 | 0.379 | 0.174 | 4.180 | 0.251 |
| 3640 | **Fam234a** | UP | Q8C0Z1 | 2 | 4 | 2 | 555 | 60.5 | 5.68 | 3 | 0.832 | 0.726 | 3.344 | 0.291 | 14.846 | 0.033 | 0.225 | 0.080 |
| 3641 | **Kti12** | Unchanged | Q9D1R2 | 1 | 2 | 1 | 351 | 38.4 | 6.92 | 3 | 0.139 | 0.199 | 0.157 | 0.209 | 0.535 | 0.499 | 0.293 | 0.178 |
| 3642 | **Rgs18** | Unchanged | Q99PG4 | 1 | 2 | 1 | 235 | 27.6 | 8.48 | 3 | 1.920 | 0.333 | 11.623 | 0.254 | 1.206 | 0.785 | 9.641 | 0.263 |
| 3643 | **Ric8a** | Unchanged | Q3TIR3 | 1 | 1 | 1 | 530 | 59.8 | 5.68 | 3 | 0.821 | 0.842 | 0.172 | 0.304 | 0.354 | 0.422 | 0.486 | 0.339 |
| 3644 | **Ubxn6** | Unchanged | Q99PL6 | 1 | 2 | 1 | 442 | 49.8 | 8.54 | 3 | 0.346 | 0.353 | 0.616 | 0.577 | 1.011 | 0.990 | 0.609 | 0.457 |
| 3645 | **Nmd3** | Unchanged | Q99L48 | 1 | 3 | 1 | 503 | 57.6 | 7.17 | 3 | 1.607 | 0.549 | 0.327 | 0.210 | 2.677 | 0.230 | 0.122 | 0.097 |
| 3646 | **Ngly1** | Unchanged | Q9JI78 | 1 | 1 | 1 | 651 | 74.2 | 6.89 | 3 | 0.930 | 0.903 | 2.786 | 0.122 | 0.561 | 0.389 | 4.968 | 0.044 |
| 3647 | **Elp1** | Unchanged | Q7TT37 | 3 | 30 | 3 | 1333 | 149.5 | 6 | 3 | 1.170 | 0.697 | 1.191 | 0.694 | 0.385 | 0.107 | 3.096 | 0.038 |
| 3648 | **Saal1** | Unchanged | Q9D2C2 | 1 | 1 | 1 | 474 | 52.7 | 4.53 | 3 | 0.612 | 0.262 | 0.934 | 0.859 | 0.575 | 0.243 | 1.624 | 0.251 |
| 3649 | **Tmbim6** | Unchanged | Q9D2C7 | 1 | 16 | 1 | 237 | 26.5 | 8.73 | 3 | 1.011 | 0.975 | 0.634 | 0.420 | 2.403 | 0.144 | 0.264 | 0.092 |
| 3650 | **Washc2** | Unchanged | Q6PGL7 | 2 | 3 | 2 | 1334 | 145.2 | 4.77 | 3 | 0.461 | 0.381 | 0.929 | 0.929 | 1.160 | 0.789 | 0.801 | 0.676 |
| 3651 | **Slc25a24** | Unchanged | Q8BMD8 | 1 | 4 | 1 | 475 | 52.9 | 7.43 | 3 | 4.472 | 0.212 | 12.888 | 0.056 | 4.762 | 0.060 | 2.706 | 0.184 |
| 3652 | **Hectd3** | Unchanged | Q3U487 | 2 | 8 | 2 | 861 | 97.3 | 5.47 | 3 | 1.376 | 0.398 | 2.440 | 0.114 | 0.672 | 0.270 | 3.634 | 0.057 |
| 3653 | **Trim72** | Unchanged | Q1XH17 | 1 | 1 | 1 | 477 | 52.8 | 6.42 | 3 | 0.329 | 0.170 | 6.749 | 0.137 | 0.505 | 0.294 | 13.372 | 0.108 |
| 3654 | **Dgat2** | Unchanged | Q9DCV3 | 1 | 2 | 1 | 388 | 43.7 | 9.48 | 3 | 1.961 | 0.291 | 2.143 | 0.297 | 0.560 | 0.350 | 3.828 | 0.160 |
| 3655 | **Hspb1** | Down | P14602 | 1 | 1 | 1 | 209 | 23 | 6.55 | 3 | 1.145 | 0.823 | 1.360 | 0.509 | 0.283 | 0.013 | 4.805 | 0.064 |
| 3656 | **Nek5** | Unchanged | Q7TSC3 | 1 | 1 | 1 | 627 | 71.7 | 5.54 | 3 | 0.236 | 0.113 | 0.580 | 0.405 | 0.514 | 0.445 | 1.128 | 0.898 |
| 3657 | **Asnsd1** | Unchanged | Q8BFS9 | 1 | 1 | 1 | 627 | 69.7 | 6.87 | 3 | #DIV/0! | #DIV/0! | #DIV/0! | #DIV/0! | #DIV/0! | #DIV/0! | #DIV/0! | #DIV/0! |
| 3658 | **Abcg2** | Unchanged | Q7TMS5 | 2 | 16 | 2 | 657 | 72.9 | 8.54 | 3 | 1.117 | 0.867 | 1.955 | 0.344 | 1.197 | 0.809 | 1.633 | 0.489 |
| 3659 | **Aip** | Down | O08915 | 1 | 1 | 1 | 330 | 37.6 | 6.4 | 3 | 0.232 | 0.141 | 0.184 | 0.091 | 0.006 | 0.043 | 29.443 | 0.087 |
| 3660 | **Nup54** | Down | Q8BTS4 | 1 | 5 | 1 | 510 | 55.7 | 7.02 | 3 | 0.424 | 0.078 | 0.348 | 0.032 | 0.233 | 0.016 | 1.496 | 0.189 |
| 3661 | **Thbs2** | Down | Q03350 | 1 | 1 | 1 | 1172 | 129.8 | 4.82 | 3 | 0.836 | 0.301 | 0.341 | 0.000 | 1.573 | 0.319 | 0.217 | 0.052 |
| 3662 | **Srsf11** | Unchanged | E9Q6E5 | 1 | 3 | 1 | 511 | 56.7 | 10.7 | 3 | 1.022 | 0.972 | 1.000 | 1.000 | 0.334 | 0.149 | 2.996 | 0.287 |
| 3663 | **Itga5** | UP | P11688 | 3 | 6 | 3 | 1053 | 115 | 5.95 | 3 | 3.455 | 0.174 | 5.136 | 0.054 | 3.473 | 0.007 | 1.479 | 0.409 |
| 3664 | **Kank2** | Unchanged | Q8BX02 | 2 | 5 | 2 | 843 | 90.2 | 5.55 | 3 | 1.218 | 0.484 | 1.480 | 0.153 | 1.198 | 0.630 | 1.236 | 0.405 |
| 3665 | **Lrrfip1** | Unchanged | A0A087WPK3 | 2 | 4 | 2 | 663 | 75 | 5.85 | 3 | 0.370 | 0.243 | 0.134 | 0.124 | 1.217 | 0.807 | 0.110 | 0.165 |
| 3666 | **Araf** | Unchanged | P04627 | 2 | 4 | 2 | 604 | 67.5 | 9.11 | 3 | 1.838 | 0.406 | 2.046 | 0.323 | 1.149 | 0.568 | 1.781 | 0.393 |
| 3667 | **Acp2** | Unchanged | P24638 | 1 | 6 | 1 | 423 | 48.5 | 7.02 | 3 | 5.300 | 0.087 | 5.672 | 0.064 | 0.998 | 0.997 | 5.683 | 0.064 |
| 3668 | **Clcc1** | Unchanged | A2AEM2 | 1 | 1 | 1 | 544 | 61.2 | 5.77 | 3 | 0.322 | 0.185 | 0.824 | 0.771 | 0.570 | 0.466 | 1.446 | 0.617 |
| 3669 | **Dtymk** | Unchanged | P97930 | 1 | 1 | 1 | 212 | 23.9 | 7.9 | 3 | 1.131 | 0.682 | 0.509 | 0.222 | 0.999 | 0.998 | 0.509 | 0.417 |
| 3670 | **Smarcc2** | Unchanged | Q6PDG5 | 2 | 14 | 2 | 1213 | 132.5 | 5.59 | 3 | 1.309 | 0.555 | 0.671 | 0.335 | 1.582 | 0.520 | 0.424 | 0.329 |
| 3671 | **Afg3l1** | Down | Q920A7 | 2 | 2 | 1 | 789 | 87 | 8.85 | 3 | 1.244 | 0.629 | 1.270 | 0.651 | 0.295 | 0.026 | 4.309 | 0.113 |
| 3672 | **Enpp4** | Unchanged | Q8BTJ4 | 2 | 4 | 2 | 456 | 51.6 | 6.73 | 3 | 0.969 | 0.936 | 1.021 | 0.914 | 0.947 | 0.653 | 1.078 | 0.685 |
| 3673 | **Cpn1** | Unchanged | Q9JJN5 | 1 | 2 | 1 | 457 | 51.8 | 8.28 | 3 | 0.973 | 0.921 | 2.063 | 0.236 | 1.221 | 0.455 | 1.690 | 0.337 |
| 3674 | **Oas1a** | Unchanged | P11928 | 1 | 1 | 1 | 367 | 42.4 | 8.09 | 3 | 2.159 | 0.384 | 0.576 | 0.452 | 3.104 | 0.249 | 0.186 | 0.172 |
| 3675 | **Pik3c3** | Unchanged | Q6PF93 | 2 | 2 | 2 | 887 | 101.4 | 6.73 | 3 | #DIV/0! | #DIV/0! | #DIV/0! | #DIV/0! | #DIV/0! | #DIV/0! | #DIV/0! | #DIV/0! |
| 3676 | **Rpa1** | Unchanged | Q5SWN2 | 1 | 1 | 1 | 644 | 71.4 | 8.18 | 3 | 1.051 | 0.951 | 0.237 | 0.051 | 0.368 | 0.125 | 0.643 | 0.591 |
| 3677 | **Arhgef37** | Unchanged | A1IGU4 | 1 | 6 | 1 | 676 | 76.5 | 5.9 | 3 | 0.821 | 0.813 | 0.627 | 0.592 | 4.458 | 0.066 | 0.141 | 0.045 |
| 3678 | **Grk6** | Unchanged | A0A286YDA5 | 1 | 1 | 1 | 584 | 66.6 | 8.73 | 3 | 2.495 | 0.299 | 1.811 | 0.604 | 3.667 | 0.235 | 0.494 | 0.478 |
| 3679 | **Ppil4** | Unchanged | Q9CXG3 | 1 | 1 | 1 | 492 | 57.2 | 6.11 | 3 | #DIV/0! | #DIV/0! | #DIV/0! | #DIV/0! | #DIV/0! | #DIV/0! | #DIV/0! | #DIV/0! |

| | | | | | | | | | | | | | | | | | | |
|---|---|---|---|---|---|---|---|---|---|---|---|---|---|---|---|---|---|---|
| 3680 | **Dync1li2** | Unchanged | Q6PDL0 | 1 | 1 | 1 | 492 | 54.2 | 6.28 | 3 | #DIV/0! | #DIV/0! | #DIV/0! | #DIV/0! | #DIV/0! | #DIV/0! | #DIV/0! | #DIV/0! |
| 3681 | **Naa16** | Unchanged | Q9DBB4 | 3 | 12 | 2 | 864 | 101.2 | 8.35 | 2 | 2.236 | 0.239 | 1.919 | 0.172 | 1.092 | 0.897 | 1.757 | 0.370 |
| 3682 | **Mrps5** | Unchanged | Q99N87 | 1 | 4 | 1 | 432 | 48.2 | 10.14 | 3 | 2.499 | 0.270 | 5.101 | 0.187 | 0.631 | 0.486 | 8.086 | 0.150 |
| 3683 | **Mob1b** | Unchanged | Q8BPB0 | 1 | 6 | 1 | 216 | 25.1 | 6.73 | 3 | 1.725 | 0.211 | 3.730 | 0.133 | 0.729 | 0.523 | 5.115 | 0.106 |
| 3684 | **Ifi204** | Unchanged | P0DOV2 | 2 | 3 | 2 | 619 | 69.4 | 8.69 | 2 | #DIV/0! | #DIV/0! | #DIV/0! | #DIV/0! | #DIV/0! | #DIV/0! | #DIV/0! | #DIV/0! |
| 3685 | **Ifit3** | Unchanged | Q64345 | 1 | 2 | 1 | 403 | 47.2 | 5.64 | 3 | 0.927 | 0.847 | 13.115 | 0.146 | 4.337 | 0.407 | 3.024 | 0.328 |
| 3686 | **Nbas** | Unchanged | E9Q411 | 3 | 4 | 3 | 2356 | 265.6 | 5.95 | 3 | 1.354 | 0.661 | 1.955 | 0.384 | 1.023 | 0.971 | 1.911 | 0.385 |
| 3687 | **Zyg11b** | Unchanged | Q3UFS0 | 2 | 2 | 2 | 744 | 83.9 | 6.87 | 3 | 10.839 | 0.341 | 1.231 | 0.451 | 1.105 | 0.592 | 1.114 | 0.682 |
| 3688 | **Ank1** | Unchanged | Q02357 | 4 | 10 | 4 | 1862 | 204.1 | 6.55 | 3 | 0.716 | 0.458 | 0.838 | 0.702 | 1.911 | 0.091 | 0.439 | 0.035 |
| 3689 | **Itpr2** | UP | Q9Z329 | 7 | 16 | 4 | 2701 | 307.3 | 6.44 | 3 | 1.789 | 0.125 | 2.971 | 0.030 | 2.699 | 0.001 | 1.101 | 0.734 |
| 3690 | **Mpp1** | Unchanged | P70290 | 1 | 11 | 1 | 466 | 52.2 | 7.2 | 3 | 0.833 | 0.712 | 2.877 | 0.394 | 0.815 | 0.666 | 3.528 | 0.349 |
| 3691 | **Vsig4** | Unchanged | F6TUL9 | 1 | 1 | 1 | 280 | 31.4 | 7.42 | 3 | 0.062 | 0.119 | 0.058 | 0.118 | 1.771 | 0.534 | 0.033 | 0.144 |
| 3692 | **Cstf3** | Unchanged | Q99LI7 | 1 | 1 | 1 | 717 | 82.8 | 8.12 | 3 | #DIV/0! | #DIV/0! | #DIV/0! | #DIV/0! | #DIV/0! | #DIV/0! | #DIV/0! | #DIV/0! |
| 3693 | **Pisd** | UP | Q8BSF4 | 1 | 2 | 1 | 406 | 45.9 | 9.57 | 3 | 1.478 | 0.291 | 3.100 | 0.290 | 5.890 | 0.025 | 0.526 | 0.306 |
| 3694 | **Ptprn2** | Unchanged | P80560 | 1 | 1 | 1 | 1001 | 111.4 | 5.95 | 3 | 0.644 | 0.257 | 0.667 | 0.278 | 1.315 | 0.240 | 0.508 | 0.020 |
| 3695 | **Washc5** | Unchanged | Q8C2E7 | 2 | 3 | 2 | 1159 | 134 | 7.12 | 3 | 1.233 | 0.586 | 1.508 | 0.466 | 1.230 | 0.539 | 1.226 | 0.699 |
| 3696 | **Sgsh** | UP | Q9EQ08 | 1 | 1 | 1 | 502 | 56.7 | 6.37 | 3 | 1.550 | 0.048 | 4.427 | 0.230 | 1.036 | 0.922 | 4.273 | 0.238 |
| 3697 | **Adgre5** | Unchanged | E9QJS7 | 1 | 1 | 1 | 818 | 90.4 | 7.3 | 3 | 1.497 | 0.463 | 1.038 | 0.890 | 0.762 | 0.524 | 1.362 | 0.464 |
| 3698 | **1700102P08Rik** | Unchanged | Q9D9C7 | 1 | 2 | 1 | 252 | 28.5 | 5.44 | 3 | 2.078 | 0.403 | 1.289 | 0.708 | 1.712 | 0.554 | 0.753 | 0.737 |
| 3699 | **Gria2** | Down | E9QKC0 | 2 | 2 | 2 | 883 | 98.8 | 7.75 | 3 | 0.027 | 0.048 | 0.070 | 0.056 | 0.061 | 0.055 | 1.140 | 0.860 |
| 3700 | **Nr1h5** | Unchanged | E9Q5A6 | 1 | 1 | 1 | 505 | 57.1 | 6.89 | 3 | 0.672 | 0.641 | 0.500 | 0.472 | 0.235 | 0.243 | 2.124 | 0.360 |
| 3701 | **Eif4g3** | Unchanged | Q80XI3 | 4 | 17 | 2 | 1579 | 174.8 | 5.53 | 3 | 1.952 | 0.259 | 2.127 | 0.195 | 1.226 | 0.552 | 1.735 | 0.291 |
| 3702 | **Galnt2** | Unchanged | Q6PB93 | 1 | 1 | 1 | 570 | 64.5 | 8.53 | 3 | #DIV/0! | #DIV/0! | #DIV/0! | #DIV/0! | #DIV/0! | #DIV/0! | #DIV/0! | #DIV/0! |
| 3703 | **Igfals** | UP | P70389 | 1 | 4 | 1 | 603 | 66.9 | 6.6 | 3 | 2.360 | 0.309 | 2.829 | 0.184 | 2.885 | 0.017 | 0.981 | 0.968 |
| 3704 | **Neurl3** | Unchanged | Q8CJC5 | 1 | 1 | 1 | 254 | 28 | 6.9 | 3 | 5.277 | 0.357 | 2.331 | 0.415 | 1.687 | 0.491 | 1.382 | 0.707 |
| 3705 | **Gm7168** | Unchanged | A0A0R4J163 | 1 | 1 | 1 | 508 | 57 | 8.22 | 3 | 1.227 | 0.594 | 17.621 | 0.138 | 21.819 | 0.112 | 0.808 | 0.792 |
| 3706 | **Epb41l5** | Unchanged | Q8BGS1 | 2 | 11 | 2 | 731 | 81.6 | 6.39 | 3 | 1.085 | 0.891 | 2.037 | 0.407 | 1.191 | 0.739 | 1.710 | 0.492 |
| 3707 | **Mbd2** | Unchanged | Q9Z2E1 | 1 | 1 | 1 | 414 | 43.5 | 10.04 | 3 | 1.046 | 0.925 | 0.737 | 0.611 | 1.931 | 0.116 | 0.382 | 0.009 |
| 3708 | **Srd5a1** | Unchanged | Q68FF9 | 1 | 2 | 1 | 255 | 29.3 | 8.63 | | 5.463 | 0.197 | 7.769 | 0.070 | 2.055 | 0.433 | 3.781 | 0.133 |
| 3709 | **Ggta1l1** | Unchanged | A2AUQ7 | 1 | 1 | 1 | 319 | 37.9 | 9.29 | 3 | 1.402 | 0.617 | 1.502 | 0.525 | 1.900 | 0.393 | 0.791 | 0.732 |
| 3710 | **Zfx** | Unchanged | P17012 | 1 | 1 | 1 | 799 | 90 | 6.19 | 3 | 1.095 | 0.862 | 4.006 | 0.173 | 1.345 | 0.346 | 2.977 | 0.219 |
| 3711 | **Pgam5** | Unchanged | Q8BX10 | 1 | 2 | 1 | 288 | 32 | 9.04 | 3 | 0.917 | 0.887 | 0.098 | 0.091 | 0.770 | 0.724 | 0.127 | 0.152 |
| 3712 | **Aox2** | Unchanged | Q5SGK3 | 3 | 7 | 3 | 1345 | 147.8 | 6.86 | 3 | 0.799 | 0.776 | 0.028 | 0.140 | 0.035 | 0.143 | 0.823 | 0.519 |
| 3713 | **Ncoa5** | Unchanged | Q91W39 | 1 | 2 | 1 | 579 | 65.3 | 9.82 | 3 | 0.731 | 0.403 | 1.520 | 0.161 | 1.305 | 0.174 | 1.165 | 0.538 |
| 3714 | **Mapkapk2** | Unchanged | P49138 | 1 | 1 | 1 | 386 | 44 | 8.57 | 3 | 0.940 | 0.923 | 0.364 | 0.185 | 0.327 | 0.159 | 1.113 | 0.812 |
| 3715 | **Rsl1d1** | Unchanged | Q8BVY0 | 1 | 2 | 1 | 452 | 50.4 | 9.98 | 3 | 0.601 | 0.497 | 1.703 | 0.678 | 0.152 | 0.086 | 11.172 | 0.353 |
| 3716 | **Ints2** | Unchanged | Q80UK8 | 2 | 2 | 2 | 1198 | 133.4 | 6.11 | 3 | 1.728 | 0.216 | 5.056 | 0.165 | 1.065 | 0.806 | 4.745 | 0.170 |
| 3717 | **Tubd1** | Unchanged | Q8CDD3 | 1 | 1 | 1 | 486 | 54.4 | 7.11 | 3 | 2.141 | 0.417 | 0.159 | 0.066 | 1.356 | 0.655 | 0.117 | 0.106 |
| 3718 | **Mtch1** | UP | Q791T5 | 1 | 1 | 1 | 389 | 41.5 | 9.32 | 3 | 2.041 | 0.256 | 6.083 | 0.040 | 1.442 | 0.471 | 4.220 | 0.059 |
| 3719 | **Sec1** | Unchanged | Q8CDC9 | 1 | 1 | 1 | 389 | 44.1 | 10.02 | 3 | 4.573 | 0.185 | 0.230 | 0.122 | 2.576 | 0.204 | 0.089 | 0.056 |
| 3720 | **Klc1** | Unchanged | Q8CD76 | 2 | 2 | 1 | 616 | 69.5 | 5.96 | 3 | 1.885 | 0.352 | 2.297 | 0.483 | 1.855 | 0.401 | 1.238 | 0.825 |

| | | | | | | | | | | | | | | | | | |
|---|---|---|---|---|---|---|---|---|---|---|---|---|---|---|---|---|---|
| 3721 | **Cml5** | Down | Q9QXS8 | 1 | 6 | 1 | 227 | 25.7 | 9.19 | 3 | 0.371 | 0.053 | 0.333 | 0.030 | 0.164 | 0.005 | 2.033 | 0.267 |
| 3722 | **Map1s** | Unchanged | Q8C052 | 3 | 6 | 3 | 973 | 102.9 | 7.02 | 3 | 0.540 | 0.130 | 0.780 | 0.464 | 1.936 | 0.058 | 0.403 | 0.033 |
| 3723 | **Ttc23** | Unchanged | Q8CHY7 | 1 | 1 | 1 | 488 | 54.6 | 7.75 | 3 | 1.243 | 0.758 | 1.325 | 0.746 | 1.335 | 0.600 | 0.992 | 0.992 |
| 3724 | **Ptprf** | Unchanged | A2A8L5 | 3 | 4 | 2 | 1898 | 211.4 | 6.65 | 3 | 0.116 | 0.256 | 2.280 | 0.322 | 2.991 | 0.266 | 0.762 | 0.701 |
| 3725 | **Ddx21** | Unchanged | Q9JIK5 | 2 | 7 | 2 | 851 | 93.5 | 9.11 | 3 | 1.143 | 0.782 | 1.269 | 0.692 | 0.440 | 0.202 | 2.882 | 0.151 |
| 3726 | **Ranbp3** | Unchanged | Q9CT10 | 1 | 2 | 1 | 491 | 52.5 | 5.12 | 3 | 1.404 | 0.717 | 0.445 | 0.508 | 0.207 | 0.319 | 2.152 | 0.442 |
| 3727 | **Gpc4** | Unchanged | P51655 | 1 | 2 | 1 | 557 | 62.5 | 6.33 | 3 | 0.786 | 0.510 | 2.028 | 0.287 | 0.592 | 0.128 | 3.424 | 0.140 |
| 3728 | **Sult1e1** | Unchanged | P49891 | 1 | 1 | 1 | 295 | 35.6 | 7.01 | 3 | 1.513 | 0.611 | 1.371 | 0.703 | 1.351 | 0.715 | 1.015 | 0.987 |
| 3729 | **Ate1** | UP | A0A1L1SQ41 | 1 | 1 | 1 | 559 | 64.1 | 7.8 | 3 | 21.530 | 0.092 | 29.663 | 0.030 | 28.129 | 0.064 | 1.055 | 0.929 |
| 3730 | **Phyhip** | Unchanged | Q8K0S0 | 1 | 1 | 1 | 330 | 37.5 | 7.01 | 3 | #DIV/0! | #DIV/0! | #DIV/0! | #DIV/0! | #DIV/0! | #DIV/0! | #DIV/0! | #DIV/0! |
| 3731 | **Zc3h11a** | Unchanged | Q6NZF1 | 1 | 3 | 1 | 792 | 86.4 | 8.13 | 3 | 0.065 | 0.170 | 0.026 | 0.154 | 0.081 | 0.177 | 0.324 | 0.292 |
| 3732 | **Hnf1a** | Unchanged | P22361 | 1 | 4 | 1 | 628 | 67.2 | 6.16 | 3 | 0.563 | 0.128 | 1.178 | 0.564 | 0.682 | 0.237 | 1.727 | 0.164 |
| 3733 | **Rapgef1** | Unchanged | Q3UHC1 | 2 | 3 | 2 | 1224 | 136.4 | 6.24 | 3 | 1.800 | 0.422 | 1.570 | 0.525 | 1.631 | 0.538 | 0.963 | 0.949 |
| 3734 | **Plod1** | Unchanged | Q9R0E2 | 1 | 1 | 1 | 728 | 83.5 | 6.54 | 3 | #DIV/0! | #DIV/0! | #DIV/0! | #DIV/0! | #DIV/0! | #DIV/0! | #DIV/0! | #DIV/0! |
| 3735 | **Ccdc93** | Unchanged | Q7TQK5 | 2 | 5 | 2 | 629 | 72.6 | 8.29 | 3 | 9.305 | 0.145 | 10.701 | 0.154 | 8.190 | 0.093 | 1.307 | 0.737 |
| 3736 | **Sec62** | Unchanged | Q8BU14 | 1 | 1 | 1 | 398 | 45.6 | 7.31 | 3 | #DIV/0! | #DIV/0! | #DIV/0! | #DIV/0! | #DIV/0! | #DIV/0! | #DIV/0! | #DIV/0! |
| 3737 | **Oxr1** | UP | Q4KMM3 | 1 | 3 | 1 | 866 | 95.9 | 5.33 | 3 | 2.308 | 0.264 | 2.861 | 0.043 | 2.771 | 0.099 | 1.032 | 0.932 |
| 3738 | **Ctif** | Unchanged | Q6PEE2 | 1 | 4 | 1 | 600 | 67.8 | 6.65 | 3 | 2.568 | 0.143 | 15.091 | 0.127 | 2.195 | 0.270 | 6.875 | 0.160 |
| 3739 | **Slc47a1** | Unchanged | Q8K0H1 | 1 | 1 | 1 | 567 | 61.6 | 7.52 | 3 | 0.068 | 0.140 | 0.053 | 0.135 | 0.089 | 0.148 | 0.603 | 0.204 |
| 3740 | **Plcg1** | Unchanged | Q62077 | 2 | 2 | 2 | 1302 | 149.6 | 5.8 | 3 | 1.306 | 0.656 | 0.882 | 0.900 | 1.522 | 0.600 | 0.580 | 0.604 |
| 3741 | **Chn2** | Unchanged | Q3V2R3 | 1 | 1 | 1 | 468 | 53.8 | 7.4 | 3 | 1.148 | 0.746 | 4.120 | 0.292 | 0.855 | 0.624 | 4.820 | 0.271 |
| 3742 | **Tjp3** | Unchanged | Q921G9 | 2 | 3 | 1 | 904 | 99.1 | 6.76 | 3 | 0.332 | 0.073 | 0.660 | 0.550 | 0.887 | 0.843 | 0.743 | 0.733 |
| 3743 | **Magt1** | Unchanged | Q9CQY5 | 1 | 2 | 1 | 335 | 37.9 | 9.76 | 3 | 1.666 | 0.327 | 2.091 | 0.291 | 0.131 | 0.106 | 15.965 | 0.048 |
| 3744 | **Foxr1** | Unchanged | Q3UTB7 | 1 | 1 | 1 | 268 | 30.6 | 9.1 | 3 | 0.415 | 0.248 | 0.760 | 0.682 | 0.576 | 0.398 | 1.319 | 0.673 |
| 3745 | **Pxdn** | Unchanged | Q3UQ28 | 1 | 1 | 1 | 1475 | 165 | 7.14 | 3 | #DIV/0! | #DIV/0! | #DIV/0! | #DIV/0! | #DIV/0! | #DIV/0! | #DIV/0! | #DIV/0! |
| 3746 | **Spart** | Unchanged | Q8R1X6 | 1 | 1 | 1 | 671 | 72.6 | 5.86 | 3 | 0.519 | 0.348 | 0.472 | 0.239 | 0.355 | 0.153 | 1.330 | 0.298 |
| 3747 | **Clk3** | Unchanged | O35492 | 1 | 1 | 1 | 638 | 73.8 | 9.94 | 3 | 0.476 | 0.456 | 0.525 | 0.492 | 0.268 | 0.301 | 1.959 | 0.017 |
| 3748 | **Os9** | Unchanged | Q8K2C7 | 1 | 1 | 1 | 672 | 76.1 | 4.84 | 3 | 1.530 | 0.206 | 3.286 | 0.061 | 1.404 | 0.208 | 2.340 | 0.108 |
| 3749 | **Snx32** | Unchanged | Q80ZJ7 | 1 | 1 | 1 | 404 | 46.6 | 6.96 | 3 | 1.039 | 0.937 | 1.049 | 0.920 | 6.672 | 0.126 | 0.157 | 0.129 |
| 3750 | **Rapgef4** | Unchanged | Q9EQZ6 | 2 | 5 | 2 | 1011 | 115.4 | 6.92 | 3 | 1.670 | 0.230 | 2.368 | 0.125 | 1.878 | 0.074 | 1.261 | 0.556 |
| 3751 | **Rabl3** | Unchanged | Q9D4V7 | 1 | 2 | 1 | 236 | 26.3 | 7.15 | 3 | 1.884 | 0.372 | 3.373 | 0.379 | 0.037 | 0.205 | 90.067 | 0.210 |
| 3752 | **Pnpla8** | Unchanged | Q8K1N1 | 2 | 4 | 2 | 776 | 87.3 | 9.23 | 3 | 1.229 | 0.629 | 1.351 | 0.399 | 0.348 | 0.108 | 3.879 | 0.000 |
| 3753 | **Ubp1** | Unchanged | Q811S7 | 1 | 2 | 1 | 540 | 60.2 | 6.27 | 3 | 1.894 | 0.338 | 1.741 | 0.191 | 0.639 | 0.215 | 2.725 | 0.083 |
| 3754 | **Panx2** | UP | Q6IMP4 | 1 | 3 | 1 | 677 | 74.6 | 7.52 | 3 | 4.099 | 0.069 | 1.580 | 0.355 | 6.539 | 0.041 | 0.242 | 0.055 |
| 3755 | **Rars2** | Unchanged | Q3U186 | 1 | 4 | 1 | 578 | 65.3 | 8.02 | 3 | 0.555 | 0.286 | 0.687 | 0.485 | 1.539 | 0.430 | 0.446 | 0.167 |
| 3756 | **Ppp1r10** | Unchanged | Q80W00 | 1 | 2 | 1 | 888 | 94.3 | 9.16 | 3 | 1.592 | 0.197 | 2.264 | 0.144 | 0.680 | 0.369 | 3.328 | 0.067 |
| 3757 | **Gga3** | Unchanged | Q8BMI3 | 1 | 1 | 1 | 718 | 77.9 | 5.66 | 3 | 0.275 | 0.197 | 0.745 | 0.704 | 1.757 | 0.369 | 0.424 | 0.202 |
| 3758 | **Smarce1** | Unchanged | O54941 | 1 | 1 | 1 | 411 | 46.6 | 4.88 | 3 | #DIV/0! | #DIV/0! | #DIV/0! | #DIV/0! | #DIV/0! | #DIV/0! | #DIV/0! | #DIV/0! |
| 3759 | **Pde6a** | Unchanged | Q8K0A8 | 1 | 3 | 1 | 860 | 99.5 | 5.55 | 3 | #DIV/0! | #DIV/0! | #DIV/0! | #DIV/0! | #DIV/0! | #DIV/0! | #DIV/0! | #DIV/0! |
| 3760 | **Txlna** | UP | Q6PAM1 | 1 | 2 | 1 | 554 | 62.3 | 6.74 | 3 | 0.640 | 0.474 | 1.143 | 0.879 | 3.657 | 0.050 | 0.312 | 0.107 |
| 3761 | **Yipf3** | Unchanged | Q3UDR8 | 1 | 3 | 1 | 347 | 38 | 5.94 | 3 | 0.369 | 0.197 | 0.402 | 0.245 | 0.384 | 0.208 | 1.045 | 0.955 |

| | | | | | | | | | | | | | | | | | | |
|---|---|---|---|---|---|---|---|---|---|---|---|---|---|---|---|---|---|---|
| 3762 | **Rrp36** | Unchanged | A0A3B2WB30 | 1 | 2 | 1 | 244 | 28.5 | 10.45 | 3 | 1.750 | 0.411 | 2.948 | 0.440 | 2.698 | 0.278 | 1.093 | 0.929 |
| 3763 | **Atp2b1** | Unchanged | G5E829 | 2 | 4 | 2 | 1220 | 134.7 | 5.91 | 3 | 0.468 | 0.074 | 1.034 | 0.950 | 0.460 | 0.102 | 2.248 | 0.285 |
| 3764 | **Ryk** | Unchanged | Q01887 | 1 | 1 | 1 | 594 | 66.2 | 7.14 | 3 | 0.809 | 0.754 | 1.171 | 0.822 | 0.221 | 0.128 | 5.307 | 0.141 |
| 3765 | **Cog7** | Unchanged | Q3UM29 | 1 | 2 | 1 | 770 | 86 | 5.38 | 3 | 1.644 | 0.276 | 1.431 | 0.504 | 0.842 | 0.815 | 1.699 | 0.359 |
| 3766 | **Nup133** | UP | Q8R0G9 | 2 | 2 | 2 | 1155 | 128.5 | 5.2 | 3 | 2.442 | 0.335 | 4.139 | 0.047 | 0.784 | 0.451 | 5.281 | 0.036 |
| 3767 | **Trim2** | Unchanged | E9QKC6 | 1 | 1 | 1 | 770 | 84.3 | 7.47 | 3 | 0.931 | 0.838 | 0.928 | 0.818 | 1.102 | 0.763 | 0.842 | 0.578 |
| 3768 | **Tmem69** | UP | Q3KQJ0 | 1 | 2 | 1 | 245 | 27.3 | 9.86 | 3 | 20.872 | 0.043 | 2.958 | 0.216 | 27.392 | 0.041 | 0.108 | 0.056 |
| 3769 | **Macf1** | Unchanged | Q9QXZ0 | 11 | 18 | 11 | 7354 | 831.4 | 5.43 | 3 | 1.339 | 0.676 | 1.101 | 0.876 | 1.977 | 0.370 | 0.557 | 0.411 |
| 3770 | **Myh14** | Down | Q6URW6 | 5 | 42 | 1 | 2000 | 228.4 | 5.55 | 3 | 0.664 | 0.427 | 0.409 | 0.152 | 0.228 | 0.042 | 1.794 | 0.410 |
| 3771 | **Ambn** | Unchanged | Q5M8P3 | 1 | 4 | 1 | 422 | 45.3 | 6.47 | 3 | 0.779 | 0.633 | 0.241 | 0.079 | 0.678 | 0.563 | 0.355 | 0.303 |
| 3772 | **Abhd15** | Down | Q5F2F2 | 1 | 3 | 1 | 459 | 51.1 | 7.99 | 3 | 0.898 | 0.624 | 0.475 | 0.113 | 0.071 | 0.001 | 6.656 | 0.114 |
| 3773 | **Ptar1** | Unchanged | E9QAB6 | 1 | 12 | 1 | 424 | 48.3 | 6.98 | 3 | 0.898 | 0.866 | 0.776 | 0.718 | 0.777 | 0.702 | 0.998 | 0.997 |
| 3774 | **Slc29a1** | UP | Q9JIM1 | 1 | 1 | 1 | 460 | 50.2 | 8.31 | 3 | 2.310 | 0.156 | 2.862 | 0.071 | 3.773 | 0.021 | 0.759 | 0.248 |
| 3775 | **Cdc73** | Unchanged | Q8JZM7 | 2 | 2 | 2 | 531 | 60.5 | 9.61 | 3 | 2.189 | 0.188 | 10.146 | 0.109 | 1.339 | 0.637 | 7.578 | 0.122 |
| 3776 | **Rb1** | Down | P13405 | 1 | 1 | 1 | 921 | 105.3 | 7.93 | 3 | 0.504 | 0.056 | 5.737 | 0.035 | 0.501 | 0.047 | 11.443 | 0.023 |
| 3777 | **Nanp** | Unchanged | Q9CPT3 | 1 | 1 | 1 | 248 | 27.8 | 6.07 | 3 | 2.338 | 0.271 | 2.618 | 0.212 | 0.373 | 0.238 | 7.016 | 0.075 |
| 3778 | **Rbm12** | Unchanged | Q8R4X3 | 2 | 3 | 2 | 992 | 102.7 | 8.32 | 3 | 1.002 | 0.996 | 4.479 | 0.212 | 0.562 | 0.251 | 7.976 | 0.163 |
| 3779 | **Cnot1** | Unchanged | Q6ZQ08 | 4 | 21 | 4 | 2375 | 266.6 | 7.11 | 3 | 1.990 | 0.396 | 1.494 | 0.617 | 1.863 | 0.422 | 0.802 | 0.720 |
| 3780 | **Pdp2** | Unchanged | Q504M2 | 1 | 2 | 1 | 532 | 59.9 | 6.32 | 3 | 0.689 | 0.616 | 0.990 | 0.991 | 0.465 | 0.361 | 2.129 | 0.470 |
| 3781 | **Mtm1** | Unchanged | Q9Z2C5 | 1 | 5 | 1 | 603 | 69.5 | 7.75 | 3 | 2.112 | 0.289 | 3.723 | 0.180 | 2.302 | 0.191 | 1.618 | 0.485 |
| 3782 | **Env** | Unchanged | P10404 | 1 | 1 | 1 | 641 | 69.6 | 7.96 | 3 | #DIV/0! | #DIV/0! | #DIV/0! | #DIV/0! | #DIV/0! | #DIV/0! | #DIV/0! | #DIV/0! |
| 3783 | **Ckap5** | Unchanged | A2AGT5 | 4 | 6 | 4 | 2032 | 225.5 | 7.96 | 3 | 6.974 | 0.366 | 0.684 | 0.132 | 0.757 | 0.342 | 0.904 | 0.673 |
| 3784 | **Ppp6r1** | Unchanged | Q7TSI3 | 3 | 5 | 2 | 856 | 94.5 | 4.64 | 3 | 0.822 | 0.731 | 1.574 | 0.503 | 1.069 | 0.915 | 1.473 | 0.578 |
| 3785 | **Rap1gds1** | Unchanged | E9Q912 | 1 | 2 | 1 | 607 | 66 | 5.35 | 3 | 0.763 | 0.731 | 3.521 | 0.216 | 0.668 | 0.578 | 5.267 | 0.156 |
| 3786 | **Btg4** | Unchanged | O70552 | 1 | 1 | 1 | 250 | 28.6 | 8.73 | 3 | 1.305 | 0.688 | 0.179 | 0.115 | 0.067 | 0.066 | 2.669 | 0.520 |
| 3787 | **5730455P16Rik** | Unchanged | J3KMQ6 | 1 | 1 | 1 | 393 | 44.4 | 5.05 | 3 | 0.918 | 0.822 | 1.361 | 0.483 | 0.918 | 0.818 | 1.484 | 0.389 |
| 3788 | **Snw1** | Unchanged | A0A0B4J1E2 | 1 | 1 | 1 | 536 | 61.4 | 9.48 | 3 | 0.209 | 0.185 | 0.512 | 0.430 | 1.244 | 0.797 | 0.412 | 0.373 |
| 3789 | **Tinagl1** | Unchanged | Q99JR5 | 1 | 2 | 1 | 466 | 52.6 | 6.77 | 3 | #DIV/0! | #DIV/0! | #DIV/0! | #DIV/0! | #DIV/0! | #DIV/0! | #DIV/0! | #DIV/0! |
| 3790 | **Prl7c1** | Unchanged | Q9CRB5 | 1 | 5 | 1 | 251 | 29.2 | 5.45 | 3 | 0.937 | 0.933 | 1.119 | 0.889 | 0.165 | 0.185 | 6.792 | 0.150 |
| 3791 | **1700074P13Rik** | Unchanged | Q9D9G7 | 1 | 3 | 1 | 251 | 28.4 | 7.69 | 3 | 2.359 | 0.332 | 0.856 | 0.841 | 2.564 | 0.117 | 0.334 | 0.087 |
| 3792 | **Slc2a9** | Unchanged | Q3T9X0 | 1 | 4 | 1 | 538 | 58.6 | 8.07 | 3 | 0.820 | 0.739 | 1.193 | 0.801 | 0.372 | 0.175 | 3.209 | 0.229 |
| 3793 | **Spta1** | UP | P08032 | 4 | 10 | 4 | 2415 | 279.7 | 5.03 | 3 | 2.750 | 0.042 | 3.850 | 0.063 | 1.536 | 0.234 | 2.506 | 0.130 |
| 3794 | **Supt5h** | Unchanged | O55201 | 2 | 3 | 2 | 1082 | 120.6 | 5.05 | 3 | 72.710 | 0.075 | 2.198 | 0.281 | 76.771 | 0.076 | 0.029 | 0.080 |
| 3795 | **Carm1** | Unchanged | D3YUP1 | 1 | 1 | 1 | 651 | 70.3 | 7.18 | 3 | #DIV/0! | #DIV/0! | #DIV/0! | #DIV/0! | #DIV/0! | #DIV/0! | #DIV/0! | #DIV/0! |
| 3796 | **Shc1** | Unchanged | P98083 | 1 | 3 | 1 | 579 | 62.6 | 6.54 | 3 | 0.893 | 0.877 | 0.625 | 0.562 | 0.124 | 0.105 | 5.038 | 0.244 |
| 3797 | **Cpd** | Unchanged | O89001 | 1 | 1 | 1 | 1377 | 152.6 | 6.18 | 3 | 1.654 | 0.330 | 1.938 | 0.093 | 1.124 | 0.603 | 1.724 | 0.161 |
| 3798 | **Slk** | Unchanged | O54988 | 2 | 5 | 2 | 1233 | 141.4 | 5.14 | 3 | 0.598 | 0.577 | 0.430 | 0.394 | 0.114 | 0.184 | 3.761 | 0.100 |
| 3799 | **Lancl1** | Unchanged | O89112 | 1 | 1 | 1 | 399 | 45.3 | 7.77 | 3 | 0.740 | 0.755 | 0.018 | 0.138 | 0.010 | 0.135 | 1.865 | 0.202 |
| 3800 | **Epb42** | Unchanged | P49222 | 1 | 2 | 1 | 691 | 76.7 | 7.31 | 3 | #DIV/0! | #DIV/0! | #DIV/0! | #DIV/0! | #DIV/0! | #DIV/0! | #DIV/0! | #DIV/0! |
| 3801 | **Vps16** | Unchanged | Q920Q4 | 2 | 3 | 2 | 839 | 94.9 | 7.01 | 3 | 1.201 | 0.714 | 0.350 | 0.063 | 0.786 | 0.676 | 0.444 | 0.373 |
| 3802 | **Ciao3** | Unchanged | Q7TMW6 | 1 | 2 | 1 | 476 | 53.1 | 6.48 | 3 | 0.655 | 0.384 | 2.559 | 0.220 | 0.395 | 0.123 | 6.483 | 0.090 |

| | | | | | | | | | | | | | | | | | | |
|---|---|---|---|---|---|---|---|---|---|---|---|---|---|---|---|---|---|---|
| 3803 | **Anapc4** | Unchanged | Q91W96 | 1 | 1 | 1 | 807 | 91.6 | 5.4 | 3 | 0.827 | 0.832 | 0.212 | 0.213 | 0.052 | 0.138 | 4.099 | 0.094 |
| 3804 | **Krt222** | Down | Q8CCX5 | 1 | 4 | 1 | 294 | 34.2 | 5.78 | 3 | 0.838 | 0.727 | 0.497 | 0.128 | 0.022 | 0.003 | 22.545 | 0.027 |
| 3805 | **Eif2ak2** | Unchanged | Q03963 | 1 | 1 | 1 | 515 | 58.2 | 8.57 | 3 | #DIV/0! | #DIV/0! | #DIV/0! | #DIV/0! | #DIV/0! | #DIV/0! | #DIV/0! | #DIV/0! |
| 3806 | **Ednrb** | Down | P48302 | 1 | 1 | 1 | 442 | 49.5 | 9.28 | 3 | 1.046 | 0.832 | 1.004 | 0.991 | 0.471 | 0.019 | 2.133 | 0.120 |
| 3807 | **Arhgap18** | Down | Q8K0Q5 | 1 | 1 | 1 | 663 | 74.9 | 6.73 | 3 | 0.851 | 0.533 | 1.021 | 0.913 | 0.350 | 0.009 | 2.919 | 0.001 |
| 3808 | **Nudt14** | Unchanged | Q9D142 | 1 | 4 | 1 | 222 | 24.4 | 5.14 | 3 | 2.055 | 0.077 | 1.651 | 0.346 | 1.008 | 0.988 | 1.637 | 0.362 |
| 3809 | **Dhx36** | Unchanged | Q8VHK9 | 2 | 12 | 2 | 1001 | 113.8 | 8.29 | 3 | 1.263 | 0.606 | 2.639 | 0.145 | 0.858 | 0.599 | 3.074 | 0.118 |
| 3810 | **Rtn3** | Unchanged | Q9ES97 | 3 | 10 | 3 | 964 | 103.8 | 4.92 | 3 | 1.704 | 0.640 | 2.036 | 0.355 | 1.412 | 0.438 | 1.443 | 0.556 |
| 3811 | **Ncoa1** | Unchanged | P70365 | 1 | 4 | 1 | 1447 | 156.9 | 6.14 | 3 | 1.418 | 0.658 | 0.433 | 0.378 | 3.788 | 0.255 | 0.114 | 0.165 |
| 3812 | **Mta2** | Unchanged | Q9R190 | 1 | 2 | 1 | 668 | 75 | 9.67 | 3 | 0.121 | 0.059 | 0.185 | 0.076 | 0.707 | 0.552 | 0.262 | 0.071 |
| 3813 | **Serpini1** | Unchanged | O35684 | 1 | 1 | 1 | 410 | 46.3 | 4.72 | 3 | 0.716 | 0.694 | 0.892 | 0.902 | 0.138 | 0.178 | 6.459 | 0.260 |
| 3814 | **Sppl2a** | Unchanged | Q9JJF9 | 1 | 1 | 1 | 523 | 58.1 | 7.06 | 3 | 1.220 | 0.516 | 1.385 | 0.404 | 0.796 | 0.613 | 1.741 | 0.222 |
| 3815 | **Cd163** | Unchanged | Q2VLH6 | 2 | 4 | 2 | 1121 | 120.8 | 6.24 | 3 | 0.344 | 0.360 | 0.071 | 0.181 | 0.200 | 0.248 | 0.357 | 0.278 |
| 3816 | **Lpcat3** | Unchanged | Q91V01 | 1 | 7 | 1 | 487 | 56.1 | 8.56 | 3 | 0.754 | 0.558 | 0.759 | 0.572 | 0.351 | 0.153 | 2.160 | 0.220 |
| 3817 | **Trim24** | Unchanged | Q64127 | 1 | 1 | 1 | 1051 | 116.6 | 7.06 | 3 | 1.419 | 0.239 | 2.104 | 0.185 | 0.773 | 0.357 | 2.720 | 0.124 |
| 3818 | **Cyb5d2** | Unchanged | Q5SSH8 | 1 | 2 | 1 | 263 | 29.1 | 7.42 | 3 | 0.837 | 0.720 | 0.386 | 0.147 | 0.279 | 0.086 | 1.384 | 0.445 |
| 3819 | **Srsf4** | Unchanged | Q8VE97 | 1 | 10 | 1 | 489 | 55.9 | 11.4 | 3 | 2.224 | 0.456 | 2.143 | 0.313 | 2.088 | 0.364 | 1.027 | 0.970 |
| 3820 | **Lrrc40** | Unchanged | Q9CRC8 | 1 | 2 | 1 | 602 | 68 | 6.89 | 3 | 1.803 | 0.140 | 2.364 | 0.076 | 1.869 | 0.183 | 1.265 | 0.555 |
| 3821 | **Pdk3** | Unchanged | Q922H2 | 1 | 2 | 1 | 415 | 47.9 | 8.82 | 3 | 0.536 | 0.465 | 0.372 | 0.331 | 0.423 | 0.366 | 0.879 | 0.862 |
| 3822 | **Cept1** | Unchanged | Q8BGS7 | 1 | 6 | 1 | 416 | 46.4 | 8.1 | 3 | 0.888 | 0.872 | 0.951 | 0.948 | 0.768 | 0.734 | 1.239 | 0.787 |
| 3823 | **Epha3** | Unchanged | Q8BRB1 | 1 | 1 | 1 | 984 | 110 | 6.93 | 3 | 0.908 | 0.688 | 0.561 | 0.099 | 0.890 | 0.740 | 0.630 | 0.288 |
| 3824 | **Trpm1** | Unchanged | A0A0U1RQ32 | 2 | 3 | 2 | 1628 | 184.2 | 8.05 | 3 | 1.062 | 0.800 | 1.017 | 0.956 | 1.178 | 0.426 | 0.863 | 0.539 |
| 3825 | **Tldc1** | UP | Q8K0P3 | 1 | 2 | 1 | 455 | 50.8 | 6.52 | 3 | 2.062 | 0.291 | 4.792 | 0.029 | 1.348 | 0.480 | 3.554 | 0.043 |
| 3826 | **Slco2b1** | Unchanged | Q8BXB6 | 1 | 5 | 1 | 683 | 74.5 | 8.31 | 3 | 1.245 | 0.675 | 2.502 | 0.246 | 0.911 | 0.829 | 2.745 | 0.206 |
| 3827 | **Bnip1** | Down | Q6QD59 | 1 | 1 | 1 | 228 | 26.2 | 8.75 | 3 | 0.645 | 0.366 | 0.825 | 0.628 | 0.213 | 0.027 | 3.870 | 0.019 |
| 3828 | **Ghdc** | Unchanged | Q99J23 | 1 | 1 | 1 | 532 | 58.5 | 7.56 | 3 | 0.539 | 0.380 | 0.486 | 0.148 | 0.541 | 0.178 | 0.897 | 0.610 |
| 3829 | **Sptb** | Unchanged | P15508 | 4 | 12 | 3 | 2128 | 245.1 | 5.33 | 3 | 0.825 | 0.724 | 0.538 | 0.342 | 0.411 | 0.223 | 1.309 | 0.306 |
| 3830 | **Slc3a1** | Unchanged | Q91WV7 | 1 | 1 | 1 | 685 | 78.1 | 5.66 | 3 | 1.466 | 0.273 | 2.184 | 0.135 | 0.870 | 0.785 | 2.509 | 0.083 |
| 3831 | **Numa1** | Unchanged | E9Q7G0 | 4 | 5 | 4 | 2094 | 235.5 | 5.87 | 3 | 0.759 | 0.499 | 0.578 | 0.256 | 0.740 | 0.431 | 0.781 | 0.602 |
| 3832 | **Slc1a2** | Unchanged | P43006 | 1 | 2 | 1 | 572 | 62 | 6.68 | 3 | #DIV/0! | #DIV/0! | #DIV/0! | #DIV/0! | #DIV/0! | #DIV/0! | #DIV/0! | #DIV/0! |
| 3833 | **Ddx31** | Unchanged | Q6NZQ2 | 1 | 1 | 1 | 687 | 76.9 | 9.74 | 3 | 1.501 | 0.323 | 20.140 | 0.337 | 1.339 | 0.527 | 15.038 | 0.345 |
| 3834 | **Epb41l1** | Unchanged | A2AUK5 | 1 | 1 | 1 | 879 | 98.3 | 5.66 | 3 | 1.029 | 0.959 | 1.050 | 0.933 | 2.332 | 0.385 | 0.450 | 0.407 |
| 3835 | **Cnksr2** | Unchanged | Q80YA9 | 1 | 1 | 1 | 1032 | 117.3 | 6.79 | 3 | 0.898 | 0.890 | 0.665 | 0.617 | 1.058 | 0.931 | 0.628 | 0.480 |
| 3836 | **Abcb6** | Unchanged | Q9DC29 | 1 | 3 | 1 | 842 | 93.7 | 8.05 | 3 | 0.073 | 0.140 | 0.216 | 0.211 | 0.070 | 0.139 | 3.064 | 0.257 |
| 3837 | **Armc4** | Unchanged | B2RY50 | 1 | 1 | 1 | 1037 | 115.2 | 6.09 | 3 | 1.119 | 0.806 | 2.102 | 0.427 | 1.500 | 0.667 | 1.401 | 0.728 |
| 3838 | **Stat2** | Unchanged | Q9WVL2 | 2 | 4 | 2 | 923 | 105.4 | 5.27 | 3 | 1.283 | 0.622 | 2.698 | 0.096 | 0.638 | 0.286 | 4.230 | 0.042 |
| 3839 | **Zzef1** | Unchanged | Q5SSH7 | 4 | 6 | 4 | 2924 | 328.1 | 6.11 | 3 | 1.348 | 0.535 | 2.350 | 0.176 | 1.126 | 0.637 | 2.087 | 0.223 |
| 3840 | **Abca8b** | Unchanged | Q8K440 | 3 | 6 | 3 | 1620 | 182.9 | 7.18 | 3 | 0.302 | 0.277 | 0.201 | 0.212 | 0.328 | 0.298 | 0.612 | 0.448 |
| 3841 | **Cand2** | Down | Q6ZQ73 | 3 | 8 | 1 | 1235 | 135.5 | 5.58 | 3 | 0.902 | 0.688 | 0.261 | 0.000 | 1.204 | 0.760 | 0.217 | 0.186 |
| 3842 | **Olfr525** | Unchanged | Q8VGL5 | 1 | 1 | 1 | 309 | 34.1 | 8.37 | 3 | 0.893 | 0.734 | 0.859 | 0.726 | 1.124 | 0.617 | 0.764 | 0.524 |
| 3843 | **Klhdc7a** | Unchanged | A2APT9 | 1 | 1 | 1 | 773 | 83.5 | 6.33 | 3 | 1.087 | 0.795 | 1.065 | 0.848 | 1.234 | 0.615 | 0.863 | 0.695 |

| | | | | | | | | | | | | | | | | | | |
|---|---|---|---|---|---|---|---|---|---|---|---|---|---|---|---|---|---|---|
| 3844 | **Acsm4** | Unchanged | Q80W40 | 2 | 5 | 1 | 580 | 65.4 | 7.72 | 3 | 0.588 | 0.431 | 0.811 | 0.759 | 0.789 | 0.699 | 1.028 | 0.967 |
| 3845 | **Pigt** | UP | Q8BXQ2 | 1 | 6 | 1 | 582 | 65.7 | 8.4 | 3 | 1.956 | 0.013 | 2.143 | 0.209 | 2.202 | 0.015 | 0.973 | 0.948 |
| 3846 | **Nrxn1** | Unchanged | Q9CS84 | 1 | 1 | 1 | 1514 | 166.1 | 6.09 | 3 | 2.910 | 0.398 | 1.688 | 0.182 | 1.512 | 0.322 | 1.116 | 0.761 |
| 3847 | **Spag9** | Unchanged | Q58A65 | 2 | 6 | 2 | 1321 | 146.1 | 5.15 | 3 | 0.990 | 0.975 | 1.614 | 0.417 | 0.865 | 0.762 | 1.867 | 0.361 |
| 3848 | **Lats1** | Unchanged | Q8BYR2 | 1 | 1 | 1 | 1129 | 126.2 | 8.73 | 3 | #DIV/0! | #DIV/0! | #DIV/0! | #DIV/0! | #DIV/0! | #DIV/0! | #DIV/0! | #DIV/0! |
| 3849 | **Styk1** | Down | Q6J9G1 | 1 | 1 | 1 | 429 | 48.1 | 8.29 | 3 | 0.072 | 0.033 | 3.193 | 0.128 | 2.302 | 0.199 | 1.387 | 0.570 |
| 3850 | **Kmt5c** | Unchanged | Q6Q783 | 1 | 5 | 1 | 468 | 53.1 | 9 | 3 | 9.448 | 0.152 | 11.392 | 0.090 | 4.440 | 0.089 | 2.566 | 0.253 |
| 3851 | **Abcc3** | UP | B2RX12 | 2 | 8 | 2 | 1523 | 169 | 7.06 | 3 | 1.624 | 0.049 | 2.195 | 0.225 | 1.551 | 0.304 | 1.415 | 0.533 |
| 3852 | **Gtf2f1** | UP | Q3THK3 | 1 | 1 | 1 | 508 | 57.2 | 7.01 | 3 | 33.038 | 0.029 | 72.992 | 0.057 | 44.492 | 0.029 | 1.641 | 0.455 |
| 3853 | **Tnks1bp1** | Unchanged | P58871 | 3 | 4 | 3 | 1720 | 181.7 | 4.88 | 3 | 0.841 | 0.790 | 0.938 | 0.934 | 1.712 | 0.180 | 0.548 | 0.283 |
| 3854 | **Add3** | Unchanged | Q9QYB5 | 1 | 1 | 1 | 706 | 78.7 | 5.95 | 3 | 0.921 | 0.834 | 3.334 | 0.143 | 0.960 | 0.913 | 3.474 | 0.136 |
| 3855 | **Relch** | Unchanged | G3X9J4 | 2 | 2 | 2 | 1216 | 134.5 | 5.34 | 3 | 3.024 | 0.249 | 1.243 | 0.718 | 0.992 | 0.990 | 1.253 | 0.735 |
| 3856 | **Carmil2** | Unchanged | Q3V3V9 | 1 | 1 | 1 | 1296 | 141.3 | 7.18 | 3 | 2.586 | 0.253 | 1.711 | 0.494 | 2.376 | 0.059 | 0.720 | 0.492 |
| 3857 | **Lemd2** | Unchanged | Q6DVA0 | 1 | 2 | 1 | 511 | 57.5 | 9.01 | 3 | 2.661 | 0.387 | 8.922 | 0.167 | 1.845 | 0.596 | 4.836 | 0.229 |
| 3858 | **Cmss1** | Unchanged | Q9CZT6 | 1 | 6 | 1 | 276 | 31.6 | 8.46 | 3 | 2.404 | 0.412 | 1.566 | 0.516 | 2.076 | 0.420 | 0.754 | 0.725 |
| 3859 | **Farp2** | Unchanged | Q91VS8 | 2 | 7 | 2 | 1065 | 121.2 | 8.44 | 3 | 1.891 | 0.177 | 0.562 | 0.222 | 0.549 | 0.415 | 1.023 | 0.978 |
| 3860 | **Slc25a32** | Unchanged | Q8BMG8 | 1 | 1 | 1 | 316 | 35 | 9.47 | 3 | 0.253 | 0.151 | 0.218 | 0.138 | 0.029 | 0.071 | 7.573 | 0.068 |
| 3861 | **Rufy1** | Unchanged | Q8BIJ7 | 2 | 3 | 2 | 712 | 80.3 | 5.68 | 2 | 1.487 | 0.469 | 0.742 | 0.706 | 1.082 | 0.897 | 0.686 | 0.499 |
| 3862 | **Olfr686** | Unchanged | Q8VGX3 | 1 | 2 | 1 | 317 | 34.5 | 8.46 | 3 | 0.176 | 0.344 | 0.406 | 0.504 | 0.014 | 0.260 | 28.903 | 0.143 |
| 3863 | **Olfr559** | Unchanged | Q8VH14 | 1 | 1 | 1 | 318 | 35.4 | 9.45 | 3 | #DIV/0! | #DIV/0! | #DIV/0! | #DIV/0! | #DIV/0! | #DIV/0! | #DIV/0! | #DIV/0! |
| 3864 | **Lcat** | UP | P16301 | 1 | 1 | 1 | 438 | 49.7 | 6.43 | 3 | 4.484 | 0.416 | 16.189 | 0.041 | 0.524 | 0.285 | 30.903 | 0.037 |
| 3865 | **Mark3** | Unchanged | A0A1Y7VNZ6 | 1 | 1 | 1 | 797 | 88.8 | 9.38 | 3 | 29.538 | 0.306 | 0.791 | 0.706 | 2.856 | 0.232 | 0.277 | 0.167 |
| 3866 | **Ecm1** | Unchanged | Q61508 | 1 | 3 | 1 | 559 | 62.8 | 6.8 | 3 | 0.606 | 0.468 | 0.537 | 0.404 | 0.045 | 0.069 | 11.865 | 0.099 |
| 3867 | **Sap25** | Unchanged | H3BLC5 | 1 | 3 | 1 | 280 | 29.8 | 4.81 | 3 | 0.380 | 0.374 | 2.260 | 0.329 | 1.177 | 0.813 | 1.920 | 0.367 |
| 3868 | **Olfr509** | Unchanged | Q8VF20 | 1 | 2 | 1 | 321 | 36.3 | 9.35 | 2 | 1.156 | 0.822 | 3.924 | 0.253 | 0.159 | 0.126 | 24.713 | 0.144 |
| 3869 | **Xpot** | Unchanged | Q9CRT8 | 2 | 8 | 2 | 963 | 109.7 | 5.25 | 2 | 1.268 | 0.605 | 0.757 | 0.451 | 0.927 | 0.880 | 0.816 | 0.683 |
| 3870 | **Pnkp** | Unchanged | Q9JLV6 | 1 | 4 | 1 | 522 | 57.2 | 7.9 | 2 | 0.222 | 0.083 | 18.916 | 0.193 | 1.162 | 0.772 | 16.280 | 0.196 |
| 3871 | **Cpsf3** | Down | Q9QXK7 | 1 | 3 | 1 | 684 | 77.5 | 5.6 | 2 | 0.503 | 0.251 | 0.532 | 0.243 | 0.188 | 0.042 | 2.831 | 0.079 |
| 3872 | **Rnf213** | Unchanged | E9Q555 | 7 | 28 | 7 | 5152 | 584.4 | 6.8 | 2 | 0.904 | 0.763 | 1.429 | 0.499 | 0.615 | 0.162 | 2.323 | 0.183 |
| 3873 | **Supt16h** | UP | Q920B9 | 1 | 5 | 1 | 1047 | 119.7 | 5.66 | 2 | 5.988 | 0.213 | 6.810 | 0.077 | 10.300 | 0.004 | 0.661 | 0.368 |
| 3874 | **Insr** | Unchanged | P15208 | 2 | 7 | 2 | 1372 | 155.5 | 5.95 | 2 | 0.835 | 0.735 | 1.430 | 0.556 | 0.960 | 0.938 | 1.490 | 0.444 |
| 3875 | **Haghl** | Unchanged | Q9DB32 | 1 | 3 | 1 | 283 | 31.5 | 5.55 | | #DIV/0! | #DIV/0! | #DIV/0! | #DIV/0! | #DIV/0! | #DIV/0! | #DIV/0! | #DIV/0! |
| 3876 | **Phlda1** | Down | Q62392 | 2 | 12 | 2 | 405 | 45.6 | 9.7 | 2 | 0.887 | 0.827 | 0.546 | 0.247 | 0.369 | 0.040 | 1.481 | 0.527 |
| 3877 | **Ttc22** | Unchanged | Q8C159 | 1 | 1 | 1 | 568 | 63.1 | 5.1 | 2 | 0.854 | 0.860 | 0.505 | 0.487 | 0.010 | 0.141 | 50.750 | 0.146 |
| 3878 | **Sel1l3** | Unchanged | Q80TS8 | 1 | 1 | 1 | 1137 | 128.6 | 6.64 | 2 | 0.662 | 0.347 | 0.921 | 0.826 | 3.158 | 0.178 | 0.292 | 0.162 |
| 3879 | **Sept1** | Unchanged | P42209 | 1 | 1 | 1 | 366 | 42 | 5.82 | 2 | 1.301 | 0.759 | 0.555 | 0.507 | 0.096 | 0.146 | 5.782 | 0.178 |
| 3880 | **Vsig1** | Unchanged | A0A0B4J1E9 | 1 | 3 | 1 | 407 | 44 | 4.55 | 2 | 1.726 | 0.451 | 1.621 | 0.611 | 1.394 | 0.623 | 1.162 | 0.860 |
| 3881 | **Cep83** | Unchanged | Q9D5R3 | 2 | 2 | 2 | 692 | 81.9 | 6.61 | 1 | 0.959 | 0.922 | 0.167 | 0.058 | 0.513 | 0.276 | 0.326 | 0.151 |
| 3882 | **Bcl9** | UP | Q9D219 | 1 | 1 | 1 | 1425 | 148.9 | 8.91 | 2 | 2.177 | 0.344 | 0.778 | 0.631 | 4.775 | 0.048 | 0.163 | 0.037 |
| 3883 | **Kif1c** | Unchanged | O35071 | 2 | 12 | 1 | 1100 | 122.4 | 7.03 | 2 | #DIV/0! | #DIV/0! | #DIV/0! | #DIV/0! | #DIV/0! | #DIV/0! | #DIV/0! | #DIV/0! |
| 3884 | **Cyp2b9** | Unchanged | P12790 | 1 | 1 | 1 | 491 | 55.7 | 7.66 | 2 | #DIV/0! | #DIV/0! | #DIV/0! | #DIV/0! | #DIV/0! | #DIV/0! | #DIV/0! | #DIV/0! |

| | | | | | | | | | | | | | | | | | | |
|---|---|---|---|---|---|---|---|---|---|---|---|---|---|---|---|---|---|---|
| 3885 | Poli | Unchanged | E9QJU6 | 1 | 1 | 1 | 737 | 81.6 | 6.47 | 2 | 2.079 | 0.495 | 8.700 | 0.200 | 0.705 | 0.390 | 12.345 | 0.185 |
| 3886 | Parp4 | Unchanged | E9PYK3 | 3 | 4 | 3 | 1969 | 216 | 6.07 | 2 | 1.278 | 0.330 | 1.576 | 0.355 | 1.511 | 0.246 | 1.043 | 0.922 |
| 3887 | Jak1 | Unchanged | B1ASP2 | 2 | 3 | 2 | 1153 | 133.3 | 7.53 | 2 | 1.499 | 0.327 | 2.330 | 0.151 | 2.873 | 0.111 | 0.811 | 0.677 |
| 3888 | Socs7 | Unchanged | Q8VHQ2 | 1 | 1 | 1 | 579 | 62.7 | 8.18 | 2 | #DIV/0! | #DIV/0! | #DIV/0! | #DIV/0! | #DIV/0! | #DIV/0! | #DIV/0! | #DIV/0! |
| 3889 | Stkld1 | Unchanged | Q80YS9 | 2 | 3 | 2 | 662 | 74.8 | 4.97 | 2 | 0.660 | 0.565 | 1.460 | 0.495 | 1.269 | 0.712 | 1.150 | 0.774 |
| 3890 | Tut1 | Unchanged | Q8R3F9 | 1 | 8 | 1 | 869 | 94.5 | 5.81 | 2 | 1.076 | 0.823 | 0.961 | 0.909 | 1.161 | 0.610 | 0.828 | 0.501 |
| 3891 | Fbxo16 | Unchanged | Q9QZM9 | 1 | 17 | 1 | 334 | 39 | 9.85 | 2 | 0.479 | 0.372 | 0.498 | 0.392 | 1.176 | 0.783 | 0.424 | 0.135 |
| 3892 | Ctsr | Unchanged | Q9JIA9 | 1 | 12 | 1 | 334 | 37.4 | 8.28 | 2 | 1.472 | 0.579 | 0.920 | 0.893 | 1.209 | 0.774 | 0.761 | 0.682 |
| 3893 | Thbs1 | Unchanged | P35441 | 2 | 5 | 2 | 1170 | 129.6 | 4.96 | 2 | 0.036 | 0.144 | 0.109 | 0.174 | 0.103 | 0.171 | 1.061 | 0.910 |
| 3894 | Parp12 | Unchanged | Q8BZ20 | 1 | 2 | 1 | 711 | 79.9 | 8.54 | 2 | 1.375 | 0.306 | 2.041 | 0.330 | 1.944 | 0.128 | 1.050 | 0.929 |
| 3895 | Ogt | Unchanged | Q8CGY8 | 2 | 6 | 2 | 1046 | 116.9 | 6.7 | 2 | 1.704 | 0.512 | 0.150 | 0.081 | 0.755 | 0.669 | 0.199 | 0.130 |
| 3896 | Dyrk3 | Unchanged | Q922Y0 | 1 | 1 | 1 | 586 | 65.5 | 9.55 | 2 | 0.882 | 0.619 | 3.227 | 0.382 | 1.314 | 0.530 | 2.455 | 0.456 |
| 3897 | Hk3 | Unchanged | Q3TRM8 | 1 | 6 | 1 | 922 | 100 | 5.94 | 2 | 1.493 | 0.562 | 0.596 | 0.456 | 2.323 | 0.107 | 0.257 | 0.018 |
| 3898 | Trmt44 | Unchanged | Q9D2Q2 | 1 | 1 | 1 | 713 | 79.8 | 7.25 | 2 | #DIV/0! | #DIV/0! | #DIV/0! | #DIV/0! | #DIV/0! | #DIV/0! | #DIV/0! | #DIV/0! |
| 3899 | Me2 | Unchanged | Q99KE1 | 1 | 2 | 1 | 589 | 65.8 | 7.61 | 2 | #DIV/0! | #DIV/0! | #DIV/0! | #DIV/0! | #DIV/0! | #DIV/0! | #DIV/0! | #DIV/0! |
| 3900 | Calcrl | UP | Q9R1W5 | 1 | 1 | 1 | 463 | 53.2 | 6.71 | 2 | 37.062 | 0.057 | 34.171 | 0.042 | 31.394 | 0.082 | 1.088 | 0.896 |
| 3901 | E130308A19Rik | Unchanged | Q8C4P0 | 1 | 1 | 1 | 716 | 79.1 | 6.93 | 2 | 0.688 | 0.377 | 1.080 | 0.852 | 6.470 | 0.060 | 0.167 | 0.063 |
| 3902 | Krt23 | Unchanged | Q99PS0 | 1 | 1 | 1 | 422 | 48 | 6.23 | 2 | 17.222 | 0.192 | 14.094 | 0.363 | 12.638 | 0.152 | 1.115 | 0.927 |
| 3903 | Tmem164 | Unchanged | Q6PHN7 | 1 | 12 | 1 | 296 | 33.4 | 8.1 | 2 | 1.074 | 0.883 | 2.603 | 0.337 | 1.842 | 0.449 | 1.414 | 0.682 |
| 3904 | Tcerg1 | Unchanged | Q8CGF7 | 2 | 2 | 2 | 1100 | 123.7 | 8.65 | 2 | 0.879 | 0.670 | 4.229 | 0.384 | 0.850 | 0.668 | 4.975 | 0.365 |
| 3905 | Cacnb1 | Unchanged | A0A0R4J194 | 1 | 2 | 1 | 677 | 74.1 | 5.99 | 2 | 0.511 | 0.482 | 0.094 | 0.129 | 1.507 | 0.586 | 0.063 | 0.085 |
| 3906 | Itga8 | Unchanged | A2ARA8 | 1 | 1 | 1 | 1062 | 117.5 | 6.16 | 2 | 0.874 | 0.864 | 0.294 | 0.300 | 4.005 | 0.087 | 0.073 | 0.029 |
| 3907 | Csf2ra | Unchanged | Q00941 | 1 | 1 | 1 | 388 | 41.7 | 7.78 | 2 | 1.003 | 0.996 | 0.727 | 0.686 | 0.787 | 0.720 | 0.925 | 0.915 |
| 3908 | Itga9 | Unchanged | B8JK39 | 1 | 1 | 1 | 1036 | 114.3 | 6.07 | 2 | 1.043 | 0.924 | 3.017 | 0.146 | 1.239 | 0.593 | 2.434 | 0.201 |
| 3909 | Lgr5 | Unchanged | Q9Z1P4 | 1 | 1 | 1 | 907 | 99.6 | 6.61 | 2 | 1.688 | 0.505 | 2.622 | 0.270 | 1.258 | 0.768 | 2.084 | 0.337 |
| 3910 | Mlxipl | Unchanged | Q99MZ3 | 1 | 4 | 1 | 864 | 94.8 | 7.85 | 2 | 1.238 | 0.735 | 4.119 | 0.076 | 0.677 | 0.299 | 6.080 | 0.052 |
| 3911 | Abcd1 | Unchanged | P48410 | 1 | 1 | 1 | 736 | 81.8 | 8.7 | 2 | #DIV/0! | #DIV/0! | #DIV/0! | #DIV/0! | #DIV/0! | #DIV/0! | #DIV/0! | #DIV/0! |
| 3912 | Mybpc1 | Unchanged | D3YU50 | 2 | 2 | 2 | 1127 | 126.5 | 5.88 | 2 | 0.346 | 0.343 | 0.197 | 0.249 | 0.331 | 0.333 | 0.594 | 0.279 |
| 3913 | Nars2 | Unchanged | Q8BGV0 | 1 | 1 | 1 | 477 | 53.9 | 7.36 | 2 | #DIV/0! | #DIV/0! | #DIV/0! | #DIV/0! | #DIV/0! | #DIV/0! | #DIV/0! | #DIV/0! |
| 3914 | Tapt1 | Unchanged | Q4VBD2 | 1 | 1 | 1 | 564 | 63.9 | 8.22 | 2 | 1.216 | 0.700 | 1.732 | 0.573 | 0.428 | 0.273 | 4.049 | 0.331 |
| 3915 | Fkbp15 | Unchanged | Q6P9Q6 | 2 | 6 | 2 | 1216 | 132.9 | 5.07 | 2 | 1.001 | 0.996 | 1.686 | 0.092 | 0.507 | 0.083 | 3.324 | 0.008 |
| 3916 | Polr2b | Unchanged | Q8CFI7 | 2 | 3 | 2 | 1174 | 133.8 | 6.87 | 2 | 1.093 | 0.770 | 4.485 | 0.189 | 0.550 | 0.069 | 8.156 | 0.142 |
| 3917 | Zc3h10 | Unchanged | Q8R205 | 1 | 1 | 1 | 435 | 46.1 | 7.62 | 2 | 0.548 | 0.583 | 0.102 | 0.207 | 0.093 | 0.203 | 1.104 | 0.672 |
| 3918 | Gba2 | Unchanged | Q69ZF3 | 1 | 4 | 1 | 918 | 103.2 | 5.59 | 2 | 0.190 | 0.173 | 0.314 | 0.241 | 0.231 | 0.191 | 1.364 | 0.445 |
| 3919 | Hormad2 | Unchanged | Q5SQP1 | 1 | 2 | 1 | 306 | 34.8 | 7.49 | 2 | 0.752 | 0.491 | 0.931 | 0.896 | 0.702 | 0.444 | 1.327 | 0.636 |
| 3920 | F10 | Unchanged | O88947 | 1 | 2 | 1 | 481 | 54 | 5.66 | 2 | 0.987 | 0.984 | 1.161 | 0.842 | 0.625 | 0.522 | 1.856 | 0.373 |
| 3921 | Mthfd2 | Unchanged | P18155 | 1 | 1 | 1 | 350 | 37.8 | 8.88 | | #DIV/0! | #DIV/0! | #DIV/0! | #DIV/0! | #DIV/0! | #DIV/0! | #DIV/0! | #DIV/0! |
| 3922 | Nup210 | Unchanged | Q9QY81 | 2 | 2 | 2 | 1886 | 204 | 6.65 | 2 | 1.199 | 0.807 | 1.049 | 0.955 | 2.561 | 0.150 | 0.409 | 0.169 |
| 3923 | Birc6 | Unchanged | O88738 | 5 | 16 | 5 | 4882 | 531.8 | 6.07 | 2 | 0.676 | 0.403 | 0.965 | 0.929 | 0.772 | 0.534 | 1.250 | 0.260 |
| 3924 | Fam129b | Unchanged | Q8R1F1 | 1 | 1 | 1 | 749 | 84.8 | 5.94 | 2 | 6.301 | 0.368 | 7.397 | 0.350 | 12.514 | 0.058 | 0.591 | 0.554 |
| 3925 | Parp1 | Unchanged | Q921K2 | 2 | 2 | 2 | 1014 | 112.7 | 8.95 | 2 | 0.121 | 0.079 | 0.522 | 0.480 | 1.052 | 0.924 | 0.496 | 0.378 |

| | | | | | | | | | | | | | | | | | | |
|---|---|---|---|---|---|---|---|---|---|---|---|---|---|---|---|---|---|---|
| 3926 | **Nptn** | Unchanged | P97300 | 1 | 1 | 1 | 397 | 44.3 | 7.74 | 2 | 1.228 | 0.770 | 0.722 | 0.666 | 0.208 | 0.191 | 3.463 | 0.114 |
| 3927 | **Mme** | Unchanged | Q61391 | 2 | 3 | 2 | 750 | 85.6 | 5.81 | 2 | 2.201 | 0.137 | 1.618 | 0.318 | 1.010 | 0.985 | 1.602 | 0.395 |
| 3928 | **Nfkb1** | Unchanged | P25799 | 2 | 2 | 2 | 971 | 105.5 | 5.39 | 2 | 3.544 | 0.114 | 0.635 | 0.692 | 2.802 | 0.337 | 0.227 | 0.220 |
| 3929 | **C1ra** | Unchanged | Q8CG16 | 1 | 2 | 1 | 707 | 80 | 5.66 | 2 | 0.982 | 0.981 | 0.702 | 0.603 | 1.391 | 0.757 | 0.505 | 0.610 |
| 3930 | **Itpr1** | Unchanged | P11881 | 5 | 10 | 2 | 2749 | 313 | 6.04 | 2 | 1.186 | 0.581 | 2.880 | 0.387 | 6.389 | 0.137 | 0.451 | 0.387 |
| 3931 | **Asah2** | Unchanged | Q9JHE3 | 1 | 2 | 1 | 756 | 83.5 | 6.86 | 2 | #DIV/0! | #DIV/0! | #DIV/0! | #DIV/0! | #DIV/0! | #DIV/0! | #DIV/0! | #DIV/0! |
| 3932 | **Homer3** | Unchanged | Q99JP6 | 1 | 1 | 1 | 356 | 39.7 | 5.36 | 2 | #DIV/0! | #DIV/0! | #DIV/0! | #DIV/0! | #DIV/0! | #DIV/0! | #DIV/0! | #DIV/0! |
| 3933 | **Zfp383** | Unchanged | A0A087WRR7 | 1 | 1 | 1 | 490 | 56.1 | 8.21 | 2 | 0.034 | 0.064 | 0.046 | 0.066 | 0.088 | 0.077 | 0.524 | 0.224 |
| 3934 | **Serpine1** | Unchanged | G5E899 | 1 | 2 | 1 | 402 | 45 | 6.47 | 2 | 0.847 | 0.784 | 0.619 | 0.511 | 0.720 | 0.593 | 0.860 | 0.836 |
| 3935 | **Shoc2** | Unchanged | O88520 | 1 | 2 | 1 | 582 | 64.9 | 8.46 | 2 | 1.021 | 0.963 | 0.713 | 0.480 | 0.592 | 0.313 | 1.205 | 0.703 |
| 3936 | **Alox5** | Unchanged | P48999 | 1 | 1 | 1 | 674 | 77.9 | 6.18 | 2 | 1.517 | 0.501 | 1.416 | 0.422 | 0.173 | 0.068 | 8.169 | 0.005 |
| 3937 | **Adgrf5** | Unchanged | G5E8Q8 | 1 | 1 | 1 | 1348 | 149.3 | 7.02 | 2 | 2.759 | 0.321 | 16.337 | 0.258 | 3.079 | 0.102 | 5.306 | 0.325 |
| 3938 | **Skil** | Unchanged | Q60665 | 1 | 1 | 1 | 675 | 76.3 | 6.58 | 2 | 0.818 | 0.805 | 0.106 | 0.151 | 1.051 | 0.942 | 0.101 | 0.047 |
| 3939 | **Srrm2** | Unchanged | Q8BTI8 | 4 | 22 | 4 | 2703 | 294.7 | 12.03 | 2 | 0.461 | 0.317 | 1.063 | 0.911 | 1.071 | 0.902 | 0.992 | 0.982 |
| 3940 | **Col11a1** | Unchanged | Q61245 | 2 | 3 | 2 | 1804 | 180.9 | 5.21 | 2 | 0.338 | 0.247 | 0.187 | 0.159 | 0.662 | 0.586 | 0.283 | 0.175 |
| 3941 | **Tns2** | Unchanged | Q8CGB6 | 3 | 7 | 2 | 1400 | 151.9 | 8.48 | 2 | 0.374 | 0.138 | 0.234 | 0.064 | 0.283 | 0.081 | 0.825 | 0.500 |
| 3942 | **Olfr622** | UP | Q8VGY7 | 1 | 2 | 1 | 317 | 35.3 | 8.95 | 2 | 0.484 | 0.353 | 4.743 | 0.394 | 7.011 | 0.012 | 0.676 | 0.627 |
| 3943 | **Glt8d2** | Unchanged | E9PWN8 | 1 | 2 | 1 | 363 | 40.7 | 8.47 | 2 | 0.511 | 0.353 | 0.556 | 0.391 | 0.478 | 0.285 | 1.163 | 0.787 |
| 3944 | **Grm8** | Unchanged | P47743 | 1 | 3 | 1 | 908 | 101.8 | 8.15 | 2 | 0.812 | 0.779 | 0.374 | 0.121 | 0.375 | 0.139 | 0.998 | 0.996 |
| 3945 | **Cwc22** | Unchanged | Q8C5N3 | 1 | 2 | 1 | 908 | 104.7 | 8.09 | 2 | 0.690 | 0.240 | 1.120 | 0.643 | 0.911 | 0.799 | 1.229 | 0.503 |
| 3946 | **Casp9** | Unchanged | Q8C3Q9 | 1 | 2 | 1 | 454 | 49.9 | 6.83 | 2 | 0.632 | 0.641 | 0.448 | 0.493 | 0.127 | 0.270 | 3.515 | 0.208 |
| 3947 | **Ube2j1** | Unchanged | Q9JJZ4 | 1 | 8 | 1 | 318 | 35 | 6.99 | 2 | 0.876 | 0.858 | 2.123 | 0.339 | 0.645 | 0.571 | 3.293 | 0.168 |
| 3948 | **Tefm** | Unchanged | Q5SSK3 | 1 | 1 | 1 | 364 | 41.8 | 9.51 | 2 | 1.726 | 0.527 | 1.536 | 0.566 | 0.349 | 0.261 | 4.401 | 0.145 |
| 3949 | **Gpat4** | Unchanged | Q8K2C8 | 1 | 1 | 1 | 456 | 52.1 | 9.35 | 2 | 0.715 | 0.577 | 0.464 | 0.267 | 0.578 | 0.403 | 0.802 | 0.739 |
| 3950 | **Llgl2** | Unchanged | J3QJU5 | 2 | 4 | 2 | 1051 | 116.9 | 7.33 | 2 | 1.059 | 0.862 | 1.051 | 0.898 | 0.644 | 0.107 | 1.631 | 0.279 |
| 3951 | **Rgl2** | Down | Q61193 | 1 | 1 | 1 | 778 | 83.8 | 6.73 | 2 | 0.844 | 0.713 | 0.069 | 0.031 | 0.027 | 0.026 | 2.570 | 0.082 |
| 3952 | **Snx9** | Unchanged | Q91VH2 | 1 | 1 | 1 | 595 | 66.5 | 5.52 | 2 | 0.708 | 0.493 | 0.839 | 0.667 | 0.404 | 0.116 | 2.075 | 0.056 |
| 3953 | **Ampd2** | Unchanged | A2AE27 | 1 | 2 | 1 | 824 | 94.6 | 6.51 | 2 | 0.873 | 0.749 | 0.697 | 0.349 | 0.505 | 0.153 | 1.382 | 0.256 |
| 3954 | **Thumpd3** | Unchanged | P97770 | 1 | 2 | 1 | 505 | 56.4 | 7.43 | 2 | 1.218 | 0.565 | 1.223 | 0.728 | 0.609 | 0.286 | 2.008 | 0.311 |
| 3955 | **Cdk11b** | Unchanged | P24788 | 1 | 5 | 1 | 784 | 91.5 | 5.39 | 2 | 0.841 | 0.716 | 0.580 | 0.317 | 0.620 | 0.313 | 0.934 | 0.852 |
| 3956 | **Bag6** | Unchanged | Q9Z1R2 | 2 | 7 | 2 | 1154 | 121 | 5.71 | 2 | 4.595 | 0.199 | 0.337 | 0.310 | 6.546 | 0.143 | 0.051 | 0.103 |
| 3957 | **Gm4788** | Unchanged | E9Q8B5 | 1 | 1 | 1 | 879 | 99.4 | 7.44 | 2 | #DIV/0! | #DIV/0! | #DIV/0! | #DIV/0! | #DIV/0! | #DIV/0! | #DIV/0! | #DIV/0! |
| 3958 | **Anxa10** | Unchanged | Q9QZ10 | 1 | 1 | 1 | 324 | 37.3 | 5.4 | 2 | 2.155 | 0.282 | 7.000 | 0.302 | 3.423 | 0.078 | 2.045 | 0.536 |
| 3959 | **Ranbp10** | Unchanged | A0A0R4J0G4 | 1 | 1 | 1 | 648 | 70 | 6.68 | 2 | #DIV/0! | #DIV/0! | #DIV/0! | #DIV/0! | #DIV/0! | #DIV/0! | #DIV/0! | #DIV/0! |
| 3960 | **Larp4b** | Unchanged | Q6A0A2 | 1 | 2 | 1 | 741 | 81.6 | 7.49 | 2 | 1.845 | 0.332 | 5.126 | 0.204 | 1.144 | 0.549 | 4.480 | 0.218 |
| 3961 | **Slc34a2** | Unchanged | Q9DBP0 | 1 | 1 | 1 | 697 | 76.2 | 7.68 | 2 | 1.296 | 0.661 | 1.358 | 0.660 | 0.579 | 0.476 | 2.345 | 0.247 |
| 3962 | **Stat6** | Unchanged | P52633 | 1 | 1 | 1 | 837 | 93.4 | 6.23 | 2 | 1.817 | 0.085 | 1.675 | 0.157 | 1.600 | 0.213 | 1.047 | 0.851 |
| 3963 | **Olfr1115** | Unchanged | Q7TR53 | 1 | 1 | 1 | 326 | 37 | 8.78 | 2 | 0.254 | 0.203 | 0.504 | 0.416 | 0.148 | 0.148 | 3.417 | 0.157 |
| 3964 | **Etnk1** | Down | A0A0N4SVC4 | 1 | 1 | 1 | 373 | 42.5 | 5.78 | 2 | 0.606 | 0.032 | 0.676 | 0.236 | 0.536 | 0.081 | 1.261 | 0.631 |
| 3965 | **Osbpl8** | Unchanged | B9EJ86 | 1 | 2 | 1 | 889 | 101.2 | 6.96 | 2 | #DIV/0! | #DIV/0! | #DIV/0! | #DIV/0! | #DIV/0! | #DIV/0! | #DIV/0! | #DIV/0! |
| 3966 | **Daam1** | Unchanged | Q8BPM0 | 2 | 5 | 2 | 1077 | 123.3 | 7.46 | 2 | 0.842 | 0.797 | 0.635 | 0.504 | 0.860 | 0.788 | 0.739 | 0.537 |

| | | | | | | | | | | | | | | | | | | |
|---|---|---|---|---|---|---|---|---|---|---|---|---|---|---|---|---|---|---|
| 3967 | **Thoc2** | Unchanged | B1AZI6 | 3 | 6 | 3 | 1594 | 182.7 | 8.44 | 2 | 0.810 | 0.750 | 1.242 | 0.792 | 0.643 | 0.489 | 1.932 | 0.485 |
| 3968 | **Masp1** | Unchanged | P98064 | 1 | 2 | 1 | 704 | 79.9 | 5.55 | 2 | 1.303 | 0.554 | 0.474 | 0.272 | 0.694 | 0.515 | 0.683 | 0.388 |
| 3969 | **Son** | Unchanged | Q9QX47 | 3 | 7 | 3 | 2444 | 265.5 | 5.6 | 2 | 0.923 | 0.924 | 0.180 | 0.209 | 1.060 | 0.942 | 0.169 | 0.143 |
| 3970 | **Dhx29** | Unchanged | Q6PGC1 | 2 | 2 | 2 | 1365 | 153.9 | 7.94 | 2 | 0.559 | 0.123 | 1.171 | 0.723 | 1.636 | 0.486 | 0.716 | 0.632 |
| 3971 | **Inf2** | UP | Q0GNC1 | 2 | 3 | 2 | 1273 | 138.5 | 5.21 | 2 | 2.225 | 0.298 | 3.770 | 0.020 | 2.091 | 0.162 | 1.803 | 0.159 |
| 3972 | **Mylk** | Down | Q6PDN3 | 3 | 6 | 3 | 1941 | 212.8 | 6.25 | 2 | 1.140 | 0.826 | 0.051 | 0.043 | 0.018 | 0.038 | 2.790 | 0.120 |
| 3973 | **Pla2g6** | Unchanged | P97819 | 1 | 2 | 1 | 807 | 89.5 | 7.24 | 2 | 1.468 | 0.434 | 2.668 | 0.253 | 1.078 | 0.894 | 2.476 | 0.279 |
| 3974 | **Pkd2l1** | Unchanged | A2A259 | 1 | 1 | 1 | 760 | 87.2 | 5.31 | 2 | 27.022 | 0.075 | 116.152 | 0.302 | 0.754 | 0.285 | 154.125 | 0.301 |
| 3975 | **Hcfc1** | Unchanged | Q61191 | 2 | 9 | 2 | 2045 | 210.3 | 7.18 | 2 | 1.452 | 0.340 | 3.697 | 0.156 | 2.251 | 0.081 | 1.643 | 0.449 |
| 3976 | **Golgb1** | Unchanged | E9PVZ8 | 5 | 14 | 5 | 3238 | 369.9 | 5.05 | 2 | 1.199 | 0.784 | 0.914 | 0.896 | 1.172 | 0.808 | 0.779 | 0.744 |
| 3977 | **Ackr1** | Unchanged | A0A0R4J0J4 | 1 | 1 | 1 | 334 | 36.6 | 7.77 | 2 | 0.018 | 0.075 | 0.032 | 0.079 | 0.023 | 0.076 | 1.397 | 0.645 |
| 3978 | **Kcnab2** | Unchanged | E0CXZ9 | 1 | 2 | 1 | 382 | 42.7 | 8.98 | 2 | 0.596 | 0.423 | 0.241 | 0.133 | 0.575 | 0.389 | 0.420 | 0.254 |
| 3979 | **Ppfia3** | Unchanged | P60469 | 1 | 1 | 1 | 1194 | 133.3 | 5.64 | 2 | 3.658 | 0.366 | 6.082 | 0.147 | 3.356 | 0.201 | 1.812 | 0.463 |
| 3980 | **Gpaa1** | Unchanged | Q9WTK3 | 1 | 1 | 1 | 621 | 67.9 | 8.4 | 2 | #DIV/0! | #DIV/0! | #DIV/0! | #DIV/0! | #DIV/0! | #DIV/0! | #DIV/0! | #DIV/0! |
| 3981 | **Pi4ka** | Unchanged | E9Q3L2 | 4 | 7 | 4 | 2105 | 236.9 | 7.06 | 2 | 0.557 | 0.227 | 0.347 | 0.116 | 0.513 | 0.230 | 0.676 | 0.562 |
| 3982 | **Usp33** | Unchanged | Q8R5K2 | 1 | 1 | 1 | 909 | 102.7 | 5.73 | 2 | #DIV/0! | #DIV/0! | #DIV/0! | #DIV/0! | #DIV/0! | #DIV/0! | #DIV/0! | #DIV/0! |
| 3983 | **Cdc37l1** | Unchanged | Q9CZP7 | 1 | 1 | 1 | 335 | 38.4 | 5.39 | 2 | 1.298 | 0.849 | 0.067 | 0.339 | 2.651 | 0.270 | 0.025 | 0.039 |
| 3984 | **Cstf1** | Unchanged | Q99LC2 | 1 | 1 | 1 | 431 | 48.4 | 6.58 | 2 | #DIV/0! | #DIV/0! | #DIV/0! | #DIV/0! | #DIV/0! | #DIV/0! | #DIV/0! | #DIV/0! |
| 3985 | **N4bp2l2** | Unchanged | Q8JZS6 | 1 | 1 | 1 | 575 | 66.2 | 5.22 | 2 | 1.438 | 0.574 | 0.038 | 0.078 | 0.085 | 0.091 | 0.449 | 0.109 |
| 3986 | **Myo10** | Unchanged | F8VQB6 | 1 | 1 | 1 | 2062 | 237.2 | 6.25 | 2 | 0.092 | 0.269 | 0.074 | 0.261 | 0.077 | 0.262 | 0.966 | 0.914 |
| 3987 | **Iqgap3** | Unchanged | F8VQ29 | 3 | 4 | 2 | 1632 | 185.2 | 7.52 | 2 | 1.590 | 0.460 | 1.416 | 0.693 | 2.768 | 0.196 | 0.511 | 0.402 |
| 3988 | **Ubap2l** | Unchanged | Q80X50 | 2 | 11 | 2 | 1107 | 116.7 | 7.11 | 2 | 0.824 | 0.559 | 0.826 | 0.579 | 1.148 | 0.662 | 0.720 | 0.318 |
| 3989 | **Lrrc39** | Unchanged | Q8BGI7 | 1 | 2 | 1 | 337 | 39 | 6.11 | 2 | 0.515 | 0.491 | 0.302 | 0.305 | 0.322 | 0.313 | 0.937 | 0.943 |
| 3990 | **Coq8b** | Unchanged | E9QLB8 | 1 | 1 | 1 | 533 | 59.1 | 7.75 | 2 | 0.947 | 0.928 | 9.556 | 0.079 | 1.735 | 0.474 | 5.509 | 0.107 |
| 3991 | **Iqca1l** | Unchanged | A6H690 | 2 | 33 | 2 | 825 | 95.9 | 9.51 | 2 | 1.049 | 0.932 | 0.960 | 0.938 | 0.944 | 0.907 | 1.017 | 0.975 |
| 3992 | **Als2** | Unchanged | Q920R0 | 2 | 4 | 2 | 1651 | 182.5 | 6.3 | 2 | 2.115 | 0.337 | 2.322 | 0.292 | 3.048 | 0.105 | 0.762 | 0.645 |
| 3993 | **Btnl9** | Unchanged | Q8BJE2 | 1 | 2 | 1 | 536 | 60.5 | 6.42 | 2 | 620.969 | 0.144 | 1.750 | 0.223 | 1.313 | 0.626 | 1.333 | 0.486 |
| 3994 | **Ctps2** | Unchanged | P70303 | 1 | 2 | 1 | 586 | 65.5 | 6.49 | 2 | 0.501 | 0.276 | 1.044 | 0.940 | 0.349 | 0.110 | 2.991 | 0.166 |
| 3995 | **Mthfd1l** | UP | Q3V3R1 | 1 | 1 | 1 | 977 | 105.7 | 7.02 | 2 | 1.936 | 0.007 | 1.258 | 0.470 | 1.766 | 0.088 | 0.713 | 0.274 |
| 3996 | **1810055G02Rik** | Unchanged | Q9D8N1 | 1 | 1 | 1 | 392 | 41.1 | 8.15 | 2 | 0.026 | 0.039 | 3.230 | 0.324 | 0.884 | 0.811 | 3.653 | 0.297 |
| 3997 | **Dus3l** | Unchanged | A0A0R4IZY9 | 1 | 1 | 1 | 637 | 71.1 | 7.9 | 2 | #DIV/0! | #DIV/0! | #DIV/0! | #DIV/0! | #DIV/0! | #DIV/0! | #DIV/0! | #DIV/0! |
| 3998 | **Unc79** | Unchanged | E5CYJ9 | 2 | 2 | 2 | 2654 | 297.1 | 6.39 | 2 | 0.167 | 0.099 | 0.952 | 0.940 | 0.097 | 0.078 | 9.806 | 0.087 |
| 3999 | **Pml** | UP | Q60953 | 1 | 1 | 1 | 885 | 98.2 | 5.63 | 2 | 1.846 | 0.282 | 3.057 | 0.010 | 1.545 | 0.187 | 1.979 | 0.043 |
| 4000 | **Ubac2** | Down | Q8R1K1 | 1 | 2 | 1 | 345 | 39 | 9.44 | 2 | 1.945 | 0.244 | 1.984 | 0.235 | 0.099 | 0.003 | 20.079 | 0.034 |
| 4001 | **St5** | Unchanged | Q924W7 | 1 | 1 | 1 | 1134 | 126.8 | 9.11 | 2 | 1.009 | 0.995 | 0.102 | 0.313 | 0.087 | 0.306 | 1.166 | 0.438 |
| 4002 | **Ptpn23** | Unchanged | Q6PB44 | 2 | 4 | 2 | 1692 | 185.1 | 6.8 | 2 | 1.009 | 0.981 | 1.662 | 0.454 | 1.285 | 0.585 | 1.293 | 0.686 |
| 4003 | **Srpk1** | Unchanged | O70551 | 1 | 2 | 1 | 648 | 73 | 6.19 | 2 | 1.975 | 0.102 | 3.030 | 0.173 | 1.002 | 0.997 | 3.023 | 0.178 |
| 4004 | **Tmprss9** | UP | D3YTR8 | 1 | 2 | 1 | 1099 | 118.2 | 7.74 | 2 | 5.370 | 0.029 | 3.440 | 0.121 | 16.929 | 0.040 | 0.203 | 0.077 |
| 4005 | **Ccdc62** | Unchanged | E9PVD1 | 1 | 15 | 1 | 701 | 79.3 | 6.23 | 2 | 0.888 | 0.844 | 0.598 | 0.431 | 0.863 | 0.796 | 0.692 | 0.484 |
| 4006 | **Kif1b** | Unchanged | Q60575 | 3 | 5 | 3 | 1816 | 204 | 5.62 | 2 | 0.592 | 0.497 | 0.945 | 0.928 | 0.780 | 0.716 | 1.210 | 0.768 |
| 4007 | **Tm9sf1** | Unchanged | Q9DBU0 | 1 | 3 | 1 | 606 | 68.9 | 7.09 | 2 | 0.533 | 0.233 | 2.152 | 0.088 | 2.720 | 0.068 | 0.791 | 0.543 |

| | | | | | | | | | | | | | | | | | | |
|---|---|---|---|---|---|---|---|---|---|---|---|---|---|---|---|---|---|---|
| 4008 | **Mon1a** | Unchanged | Q6PDG8 | 1 | 2 | 1 | 556 | 62.1 | 6.1 | 2 | 0.252 | 0.111 | 0.359 | 0.165 | 0.286 | 0.125 | 1.254 | 0.472 |
| 4009 | **Atp8b2 iso** | Unchanged | D3YXQ5 | 1 | 1 | 1 | 1214 | 137.3 | 7.11 | 2 | 0.848 | 0.691 | 0.875 | 0.719 | 0.899 | 0.824 | 0.973 | 0.951 |
| 4010 | **Kcnk13** | UP | Q8R1P5 | 1 | 1 | 1 | 405 | 44.9 | 8.18 | 2 | 9.994 | 0.020 | 14.582 | 0.100 | 14.026 | 0.000 | 1.040 | 0.943 |
| 4011 | **Tmprss6** | Unchanged | Q9DBI0 | 1 | 2 | 1 | 811 | 90.9 | 6.34 | 2 | 0.609 | 0.475 | 1.261 | 0.751 | 0.480 | 0.302 | 2.624 | 0.261 |
| 4012 | **Pik3ap1** | Unchanged | Q9EQ32 | 1 | 6 | 1 | 811 | 90.9 | 5.17 | 2 | 0.554 | 0.362 | 1.253 | 0.809 | 0.270 | 0.120 | 4.645 | 0.322 |
| 4013 | **Arhgef7** | Unchanged | Q9ES28 | 1 | 2 | 1 | 862 | 97 | 6.8 | 2 | #DIV/0! | #DIV/0! | #DIV/0! | #DIV/0! | #DIV/0! | #DIV/0! | #DIV/0! | #DIV/0! |
| 4014 | **Mcm4** | Unchanged | P49717 | 1 | 2 | 1 | 862 | 96.7 | 7.2 | 2 | #DIV/0! | #DIV/0! | #DIV/0! | #DIV/0! | #DIV/0! | #DIV/0! | #DIV/0! | #DIV/0! |
| 4015 | **Tgds** | Unchanged | Q8VDR7 | 1 | 4 | 1 | 355 | 40.5 | 6.33 | 2 | 0.230 | 0.219 | 0.178 | 0.183 | 0.131 | 0.161 | 1.357 | 0.706 |
| 4016 | **Tmem135** | UP | Q9CYV5 | 1 | 7 | 1 | 458 | 52.3 | 9.5 | 2 | 2.459 | 0.188 | 6.954 | 0.033 | 3.852 | 0.231 | 1.805 | 0.352 |
| 4017 | **Ptpn13** | Unchanged | Q64512 | 2 | 2 | 2 | 2453 | 270.2 | 6.37 | 2 | 0.595 | 0.364 | 1.043 | 0.959 | 1.345 | 0.791 | 0.775 | 0.836 |
| 4018 | **Akap11** | Unchanged | E9Q774 | 1 | 2 | 1 | 1895 | 208.7 | 5.36 | 2 | 1.363 | 0.745 | 0.098 | 0.322 | 1.061 | 0.950 | 0.092 | 0.042 |
| 4019 | **Zfp871** | Unchanged | G5E905 | 1 | 1 | 1 | 615 | 71.5 | 8.28 | 2 | #DIV/0! | #DIV/0! | #DIV/0! | #DIV/0! | #DIV/0! | #DIV/0! | #DIV/0! | #DIV/0! |
| 4020 | **Grtp1** | Unchanged | Q9D3N8 | 1 | 8 | 1 | 359 | 40.6 | 8.47 | 2 | 1.113 | 0.889 | 1.892 | 0.493 | 0.249 | 0.216 | 7.602 | 0.179 |
| 4021 | **Rnf146** | UP | Q9CZW6 | 1 | 3 | 1 | 359 | 38.9 | 5.2 | 2 | 9.726 | 0.023 | 17.794 | 0.109 | 0.464 | 0.279 | 38.369 | 0.100 |
| 4022 | **Rab3gap2** | Unchanged | E9QKE4 | 1 | 2 | 1 | 1387 | 154.5 | 6.07 | 2 | 0.817 | 0.623 | 1.265 | 0.558 | 1.151 | 0.700 | 1.099 | 0.753 |
| 4023 | **Elmo3** | Unchanged | Q8BYZ7 | 1 | 1 | 1 | 720 | 81.7 | 6.47 | 2 | 2.141 | 0.308 | 1.497 | 0.219 | 1.039 | 0.912 | 1.440 | 0.246 |
| 4024 | **Heatr6** | Unchanged | Q6P1G0 | 1 | 3 | 1 | 1184 | 128.8 | 7.24 | 2 | 1.371 | 0.488 | 0.835 | 0.724 | 1.160 | 0.733 | 0.719 | 0.364 |
| 4025 | **Nxf1** | Unchanged | Q99JX7 | 1 | 1 | 1 | 618 | 70.3 | 8.73 | 2 | 0.817 | 0.621 | 2.931 | 0.148 | 0.684 | 0.274 | 4.285 | 0.100 |
| 4026 | **Slc30a9** | Unchanged | Q5IRJ6 | 1 | 3 | 1 | 567 | 62.8 | 8.69 | 2 | #DIV/0! | #DIV/0! | #DIV/0! | #DIV/0! | #DIV/0! | #DIV/0! | #DIV/0! | #DIV/0! |
| 4027 | **Gvin1** | Unchanged | Q80SU7 | 4 | 7 | 4 | 2427 | 280.6 | 6.57 | 2 | 0.937 | 0.888 | 1.423 | 0.407 | 0.700 | 0.266 | 2.034 | 0.152 |
| 4028 | **Gigyf2** | Unchanged | Q6Y7W8 | 1 | 2 | 1 | 1291 | 149.1 | 5.57 | 2 | 0.928 | 0.801 | 1.694 | 0.135 | 1.130 | 0.726 | 1.500 | 0.305 |
| 4029 | **Ptprc** | Unchanged | P06800 | 1 | 1 | 1 | 1293 | 144.7 | 5.94 | 2 | 0.787 | 0.681 | 1.703 | 0.308 | 0.624 | 0.506 | 2.730 | 0.049 |
| 4030 | **Nfib** | Unchanged | P97863 | 1 | 2 | 1 | 570 | 63.5 | 8.66 | 2 | 0.950 | 0.953 | 1.587 | 0.665 | 0.014 | 0.126 | 116.091 | 0.216 |
| 4031 | **Med27** | Unchanged | Q9DB40 | 1 | 3 | 1 | 311 | 35.3 | 9.31 | 2 | 0.122 | 0.054 | 0.721 | 0.580 | 0.268 | 0.101 | 2.688 | 0.173 |
| 4032 | **Col1a1** | Unchanged | P11087 | 1 | 2 | 1 | 1453 | 137.9 | 5.85 | 2 | 2.137 | 0.074 | 2.170 | 0.159 | 2.751 | 0.052 | 0.789 | 0.581 |
| 4033 | **Hspa12a** | Unchanged | Q8K0U4 | 1 | 1 | 1 | 675 | 74.8 | 6.77 | 2 | #DIV/0! | #DIV/0! | #DIV/0! | #DIV/0! | #DIV/0! | #DIV/0! | #DIV/0! | #DIV/0! |
| 4034 | **Nup160** | Unchanged | Q9Z0W3 | 1 | 1 | 1 | 1402 | 158.1 | 5.52 | 2 | #DIV/0! | #DIV/0! | #DIV/0! | #DIV/0! | #DIV/0! | #DIV/0! | #DIV/0! | #DIV/0! |
| 4035 | **Gnpat** | Unchanged | P98192 | 1 | 4 | 1 | 678 | 76.8 | 8.13 | 2 | 0.587 | 0.243 | 2.303 | 0.193 | 1.001 | 0.998 | 2.300 | 0.190 |
| 4036 | **Lbr** | Unchanged | Q3U9G9 | 1 | 1 | 1 | 626 | 71.4 | 9.36 | 2 | 0.789 | 0.715 | 1.512 | 0.659 | 0.030 | 0.052 | 50.160 | 0.190 |
| 4037 | **Sh3gl1** | Unchanged | Q62419 | 1 | 1 | 1 | 368 | 41.5 | 5.72 | | 1.928 | 0.288 | 0.486 | 0.509 | 0.602 | 0.591 | 0.808 | 0.786 |
| 4038 | **Pou4f1** | Down | P17208 | 1 | 1 | 1 | 421 | 42.7 | 9.01 | 2 | 0.195 | 0.026 | 2.994 | 0.328 | 2.538 | 0.113 | 1.180 | 0.830 |
| 4039 | **Zfp397** | Unchanged | Q7TNK4 | 1 | 1 | 1 | 527 | 60.5 | 8.15 | 2 | 3.746 | 0.152 | 0.728 | 0.722 | 9.496 | 0.132 | 0.077 | 0.123 |
| 4040 | **Gemin4** | Unchanged | Q6P6L6 | 1 | 2 | 1 | 1058 | 120.1 | 5.15 | 2 | #DIV/0! | #DIV/0! | #DIV/0! | #DIV/0! | #DIV/0! | #DIV/0! | #DIV/0! | #DIV/0! |
| 4041 | **Tex13c2** | Unchanged | A0A0G2JGA3 | 1 | 1 | 1 | 530 | 58.8 | 5.24 | 2 | 1.336 | 0.685 | 1.005 | 0.995 | 0.337 | 0.200 | 2.982 | 0.198 |
| 4042 | **Vcpip1** | Unchanged | Q8CDG3 | 2 | 6 | 2 | 1220 | 134.4 | 7.17 | 2 | 0.278 | 0.149 | 0.333 | 0.195 | 0.095 | 0.076 | 3.514 | 0.176 |
| 4043 | **Scnn1b** | Unchanged | Q9WU38 | 1 | 2 | 1 | 638 | 72.1 | 6.09 | 2 | 1.447 | 0.570 | 1.573 | 0.539 | 1.705 | 0.352 | 0.923 | 0.886 |
| 4044 | **Nub1** | Down | A0A0G2JGQ4 | 1 | 1 | 1 | 638 | 73.1 | 6.47 | 2 | 0.644 | 0.103 | 0.581 | 0.281 | 0.016 | 0.000 | 36.759 | 0.124 |
| 4045 | **Wdr11** | Unchanged | Q8K1X1 | 2 | 4 | 2 | 1223 | 135.9 | 7.11 | 2 | 1.250 | 0.372 | 1.893 | 0.255 | 0.704 | 0.437 | 2.690 | 0.153 |
| 4046 | **Pex1** | Unchanged | Q5BL07 | 2 | 4 | 2 | 1284 | 141.3 | 6.35 | 2 | 0.380 | 0.082 | 2.547 | 0.152 | 0.943 | 0.889 | 2.702 | 0.138 |
| 4047 | **Skint4** | Unchanged | E9QLU2 | 1 | 1 | 1 | 482 | 54.8 | 6.35 | 2 | #DIV/0! | #DIV/0! | #DIV/0! | #DIV/0! | #DIV/0! | #DIV/0! | #DIV/0! | #DIV/0! |
| 4048 | **Nr1i2** | Unchanged | O54915 | 1 | 1 | 1 | 431 | 49.5 | 7.66 | 2 | 0.997 | 0.993 | 1.357 | 0.639 | 0.602 | 0.311 | 2.254 | 0.337 |

| | | | | | | | | | | | | | | | | | | |
|---|---|---|---|---|---|---|---|---|---|---|---|---|---|---|---|---|---|---|
| 4049 | **Dnaaf2** | Unchanged | Q8BPI1 | 1 | 1 | 1 | 814 | 88.3 | 4.96 | 2 | 0.198 | 0.123 | 0.332 | 0.185 | 0.431 | 0.275 | 0.770 | 0.595 |
| 4050 | **Bysl** | Unchanged | O54825 | 1 | 1 | 1 | 436 | 49.8 | 7.78 | 2 | 1.448 | 0.444 | 2.375 | 0.254 | 4.276 | 0.109 | 0.555 | 0.392 |
| 4051 | **Ptgs2** | Unchanged | Q05769 | 1 | 3 | 1 | 604 | 69 | 7.37 | 2 | 2.695 | 0.235 | 3.943 | 0.230 | 2.252 | 0.291 | 1.751 | 0.512 |
| 4052 | **Ddx18** | Unchanged | Q8K363 | 1 | 2 | 1 | 660 | 74.1 | 9.52 | 2 | 12.504 | 0.145 | 5.006 | 0.072 | 22.642 | 0.063 | 0.221 | 0.123 |
| 4053 | **Rpap3** | UP | Q9D706 | 1 | 1 | 1 | 660 | 74.1 | 7.99 | 2 | 2.492 | 0.006 | 2.182 | 0.088 | 2.556 | 0.061 | 0.854 | 0.681 |
| 4054 | **Mmp2** | Unchanged | P33434 | 1 | 1 | 1 | 662 | 74.1 | 5.53 | 2 | #DIV/0! | #DIV/0! | #DIV/0! | #DIV/0! | #DIV/0! | #DIV/0! | #DIV/0! | #DIV/0! |
| 4055 | **Hook3** | Unchanged | Q8BUK6 | 1 | 2 | 1 | 718 | 83.2 | 5.19 | 2 | 0.802 | 0.727 | 0.458 | 0.263 | 0.615 | 0.386 | 0.745 | 0.464 |
| 4056 | **Aff4** | Unchanged | Q9ESC8 | 1 | 1 | 1 | 1160 | 126.6 | 9.36 | 2 | 2.459 | 0.395 | 7.988 | 0.130 | 2.798 | 0.192 | 2.855 | 0.260 |
| 4057 | **Rin1** | Unchanged | A0A2C9F2A2 | 2 | 4 | 2 | 774 | 84.3 | 8.81 | 2 | 1.842 | 0.429 | 5.138 | 0.206 | 3.052 | 0.270 | 1.684 | 0.560 |
| 4058 | **Taar1** | Unchanged | Q923Y8 | 1 | 3 | 1 | 332 | 37.6 | 8.91 | 2 | 1.359 | 0.736 | 0.530 | 0.539 | 0.028 | 0.153 | 19.137 | 0.241 |
| 4059 | **Vps53** | Unchanged | Q8CCB4 | 1 | 1 | 1 | 832 | 94.4 | 6.61 | 2 | 0.989 | 0.971 | 1.488 | 0.435 | 23.589 | 0.141 | 0.063 | 0.149 |
| 4060 | **Map3k21** | Unchanged | Q8VDG6 | 1 | 1 | 1 | 1002 | 110 | 8.19 | 2 | 0.378 | 0.128 | 1.892 | 0.364 | 0.717 | 0.544 | 2.640 | 0.232 |
| 4061 | **Edem3** | Unchanged | A0A087WR24 | 1 | 1 | 1 | 947 | 105.8 | 5.11 | 2 | 1.112 | 0.838 | 10.363 | 0.333 | 25.331 | 0.055 | 0.409 | 0.320 |
| 4062 | **Anapc2** | Unchanged | Q8BZQ7 | 1 | 4 | 1 | 837 | 95.2 | 5.31 | 2 | 1.097 | 0.889 | 1.698 | 0.507 | 1.125 | 0.862 | 1.509 | 0.605 |
| 4063 | **D11Wsu47e** | Unchanged | Q6PIX9 | 1 | 1 | 1 | 558 | 60.2 | 9.5 | 2 | 5.437 | 0.373 | 16.537 | 0.172 | 1.094 | 0.853 | 15.118 | 0.174 |
| 4064 | **Fam91a1** | Down | Q3UVG3 | 1 | 2 | 1 | 837 | 93.4 | 6.35 | 2 | 1.420 | 0.495 | 0.447 | 0.089 | 0.204 | 0.022 | 2.191 | 0.009 |
| 4065 | **Gm49324** | Unchanged | A0A1L1SQF0 | 1 | 1 | 1 | 893 | 99.1 | 8.12 | 2 | 19.037 | 0.125 | 1.378 | 0.248 | 15.620 | 0.078 | 0.088 | 0.085 |
| 4066 | **Slc6a17** | Unchanged | Q8BJI1 | 1 | 3 | 1 | 727 | 81 | 6.23 | 2 | #DIV/0! | #DIV/0! | #DIV/0! | #DIV/0! | #DIV/0! | #DIV/0! | #DIV/0! | #DIV/0! |
| 4067 | **Kifc5b** | UP | E9PUA5 | 1 | 1 | 1 | 672 | 73.6 | 8.81 | 2 | 2.295 | 0.035 | 2.642 | 0.124 | 1.249 | 0.663 | 2.115 | 0.190 |
| 4068 | **Prpf40a** | Unchanged | Q9R1C7 | 1 | 2 | 1 | 953 | 108.4 | 7.69 | 2 | #DIV/0! | #DIV/0! | #DIV/0! | #DIV/0! | #DIV/0! | #DIV/0! | #DIV/0! | #DIV/0! |
| 4069 | **Golga5** | Unchanged | Q9QYE6 | 1 | 1 | 1 | 729 | 82.3 | 6.23 | 2 | #DIV/0! | #DIV/0! | #DIV/0! | #DIV/0! | #DIV/0! | #DIV/0! | #DIV/0! | #DIV/0! |
| 4070 | **Ipo8** | Unchanged | Q7TMY7 | 1 | 1 | 1 | 1010 | 117 | 5.16 | 2 | 2.048 | 0.251 | 1.382 | 0.660 | 1.733 | 0.188 | 0.797 | 0.716 |
| 4071 | **Gpcpd1** | Unchanged | Q8C0L9 | 1 | 5 | 1 | 675 | 76.5 | 5.58 | 2 | 1.324 | 0.567 | 0.762 | 0.623 | 0.661 | 0.394 | 1.153 | 0.736 |
| 4072 | **Jakmip3** | Unchanged | Q5DTN8 | 1 | 1 | 1 | 844 | 98.7 | 5.74 | 2 | 2.492 | 0.287 | 2.539 | 0.243 | 4.279 | 0.130 | 0.593 | 0.451 |
| 4073 | **Cdk18** | Unchanged | Q04899 | 1 | 2 | 1 | 451 | 51.8 | 8.51 | 2 | 2.090 | 0.236 | 0.797 | 0.711 | 1.839 | 0.273 | 0.433 | 0.191 |
| 4074 | **Dhodh** | Unchanged | O35435 | 1 | 1 | 1 | 395 | 42.7 | 9.55 | | #DIV/0! | #DIV/0! | #DIV/0! | #DIV/0! | #DIV/0! | #DIV/0! | #DIV/0! | #DIV/0! |
| 4075 | **Ell3** | Unchanged | Q80VR2 | 1 | 1 | 1 | 395 | 44.7 | 5.27 | 2 | 0.903 | 0.875 | 0.403 | 0.229 | 0.786 | 0.715 | 0.513 | 0.395 |
| 4076 | **Dlg5** | Unchanged | E9Q9R9 | 2 | 7 | 2 | 1921 | 214.3 | 7.4 | 2 | 0.570 | 0.606 | 0.364 | 0.455 | 0.744 | 0.762 | 0.488 | 0.438 |
| 4077 | **Ryr1** | Unchanged | E9PZQ0 | 6 | 14 | 5 | 5035 | 564.7 | 5.27 | 2 | 1.096 | 0.653 | 1.016 | 0.982 | 26.828 | 0.146 | 0.038 | 0.146 |
| 4078 | **Kifap3** | Unchanged | P70188 | 1 | 1 | 1 | 793 | 91.2 | 5.11 | 2 | 1.219 | 0.581 | 1.155 | 0.642 | 0.974 | 0.934 | 1.186 | 0.663 |
| 4079 | **Cfap57** | Unchanged | Q9D180 | 2 | 6 | 2 | 1249 | 144.8 | 5.74 | 1 | 0.855 | 0.683 | 0.986 | 0.983 | 0.389 | 0.088 | 2.531 | 0.395 |
| 4080 | **Prpf3** | UP | Q922U1 | 1 | 1 | 1 | 683 | 77.4 | 9.5 | 2 | 2841.000 | 0.065 | 3951.000 | 0.072 | 3205.500 | 0.046 | 1.233 | 0.759 |
| 4081 | **Abcd2** | Unchanged | Q61285 | 1 | 2 | 1 | 741 | 83.4 | 9.09 | 2 | 1.311 | 0.689 | 1.816 | 0.590 | 0.351 | 0.174 | 5.176 | 0.323 |
| 4082 | **Alx4** | Unchanged | O35137 | 1 | 1 | 1 | 399 | 42.7 | 8.32 | | 52.039 | 0.198 | 6.535 | 0.371 | 69.465 | 0.229 | 0.094 | 0.268 |
| 4083 | **Irf7** | Unchanged | P70434 | 1 | 1 | 1 | 457 | 51.2 | 6 | 2 | 66.350 | 0.151 | 93.756 | 0.217 | 1.512 | 0.490 | 62.000 | 0.219 |
| 4084 | **Kif6** | Unchanged | E9PX57 | 1 | 1 | 1 | 802 | 90.6 | 7.03 | 2 | 2.359 | 0.248 | 1.518 | 0.674 | 0.673 | 0.602 | 2.255 | 0.475 |
| 4085 | **Krt82** | Unchanged | Q99M74 | 1 | 1 | 1 | 516 | 57.1 | 6.25 | 2 | 1.354 | 0.663 | 0.117 | 0.076 | 0.621 | 0.409 | 0.188 | 0.000 |
| 4086 | **H2-M1** | Unchanged | F7CXU4 | 1 | 3 | 1 | 344 | 40.2 | 8.98 | 2 | 1.099 | 0.851 | 0.865 | 0.891 | 0.082 | 0.058 | 10.536 | 0.387 |
| 4087 | **Ehmt2** | Unchanged | Q9Z148 | 1 | 2 | 1 | 1263 | 138 | 5.83 | 2 | 0.811 | 0.757 | 0.868 | 0.882 | 0.990 | 0.988 | 0.877 | 0.888 |
| 4088 | **Qsox1** | Unchanged | Q8BND5 | 1 | 1 | 1 | 748 | 82.7 | 7.17 | 2 | #DIV/0! | #DIV/0! | #DIV/0! | #DIV/0! | #DIV/0! | #DIV/0! | #DIV/0! | #DIV/0! |
| 4089 | **Rock2** | Unchanged | A0A1Y7VMN0 | 2 | 4 | 1 | 1444 | 166.7 | 5.9 | 2 | 0.282 | 0.250 | 2.536 | 0.244 | 0.430 | 0.331 | 5.895 | 0.093 |

| | | | | | | | | | | | | | | | | | | |
|---|---|---|---|---|---|---|---|---|---|---|---|---|---|---|---|---|---|---|
| 4090 | **Sntb2** | Unchanged | Q61235 | 1 | 2 | 1 | 520 | 56.3 | 8.69 | 2 | 1.197 | 0.716 | 1.040 | 0.942 | 2.992 | 0.113 | 0.348 | 0.111 |
| 4091 | **Fcho2** | Unchanged | Q3UQN2 | 1 | 4 | 1 | 809 | 88.7 | 6.89 | 2 | #DIV/0! | #DIV/0! | #DIV/0! | #DIV/0! | #DIV/0! | #DIV/0! | #DIV/0! | #DIV/0! |
| 4092 | **Nelfb** | Unchanged | Q8C4Y3 | 1 | 4 | 1 | 580 | 65.6 | 6.02 | 2 | 0.907 | 0.876 | 3.693 | 0.224 | 0.398 | 0.199 | 9.287 | 0.140 |
| 4093 | **Pcdhgb2** | Unchanged | Q91XX7 | 1 | 3 | 1 | 928 | 100.8 | 4.97 | 2 | 1.014 | 0.980 | 1.885 | 0.263 | 0.623 | 0.268 | 3.027 | 0.113 |
| 4094 | **Lpin2** | Unchanged | E9PWN0 | 1 | 1 | 1 | 931 | 103.7 | 5.4 | 2 | 0.729 | 0.478 | 1.043 | 0.915 | 1.122 | 0.761 | 0.929 | 0.846 |
| 4095 | **Pcdhga9** | Unchanged | Q91XX9 | 1 | 2 | 1 | 932 | 101.1 | 5.12 | 2 | 2.776 | 0.274 | 4.588 | 0.158 | 2.578 | 0.266 | 1.780 | 0.458 |
| 4096 | **Siglec1** | Unchanged | Q62230 | 2 | 4 | 2 | 1695 | 182.9 | 6.67 | 2 | 1.486 | 0.485 | 1.365 | 0.350 | 1.985 | 0.415 | 0.688 | 0.611 |
| 4097 | **Cherp** | Unchanged | Q8CGZ0 | 1 | 2 | 1 | 936 | 106.1 | 9.14 | 2 | 1.031 | 0.948 | 1.063 | 0.891 | 1.318 | 0.422 | 0.806 | 0.590 |
| 4098 | **Myh1** | Unchanged | Q5SX40 | 2 | 3 | 2 | 1942 | 223.2 | 5.76 | 2 | 0.892 | 0.770 | 0.831 | 0.661 | 0.323 | 0.071 | 2.573 | 0.066 |
| 4099 | **Kat6a** | Unchanged | G3X940 | 1 | 1 | 1 | 2003 | 224.7 | 5.58 | 2 | 0.767 | 0.676 | 0.556 | 0.427 | 0.431 | 0.260 | 1.289 | 0.683 |
| 4100 | **Mlkl** | Unchanged | Q9D2Y4 | 1 | 1 | 1 | 472 | 54.3 | 8.25 | 2 | 0.910 | 0.909 | 0.242 | 0.313 | 0.945 | 0.948 | 0.256 | 0.137 |
| 4101 | **Allc** | Unchanged | Q9JHX6 | 1 | 1 | 1 | 414 | 46.3 | 6.25 | 2 | #DIV/0! | #DIV/0! | #DIV/0! | #DIV/0! | #DIV/0! | #DIV/0! | #DIV/0! | #DIV/0! |
| 4102 | **Heatr5b** | Unchanged | Q8C547 | 1 | 1 | 1 | 2070 | 224.2 | 7.14 | 2 | #DIV/0! | #DIV/0! | #DIV/0! | #DIV/0! | #DIV/0! | #DIV/0! | #DIV/0! | #DIV/0! |
| 4103 | **Grb14** | Unchanged | Q9JLM9 | 1 | 1 | 1 | 538 | 60.5 | 8.44 | 2 | 0.161 | 0.057 | 1.274 | 0.729 | 1.157 | 0.696 | 1.101 | 0.867 |
| 4104 | **Gripap1** | Unchanged | A2AEW8 | 1 | 1 | 1 | 837 | 95.9 | 5.17 | 2 | 1.283 | 0.731 | 0.542 | 0.423 | 4.450 | 0.077 | 0.122 | 0.043 |
| 4105 | **Ganc** | Unchanged | Q8BVW0 | 1 | 4 | 1 | 898 | 101.9 | 6.32 | 2 | 0.590 | 0.379 | 0.352 | 0.135 | 0.328 | 0.127 | 1.073 | 0.844 |
| 4106 | **Akt1** | Unchanged | P31750 | 1 | 2 | 1 | 480 | 55.7 | 5.9 | 2 | 1.159 | 0.740 | 3.021 | 0.171 | 1.245 | 0.616 | 2.427 | 0.219 |
| 4107 | **Fut10** | Down | Q5F2L2 | 1 | 8 | 1 | 481 | 55.7 | 7.33 | | 1.018 | 0.976 | 1.041 | 0.939 | 0.132 | 0.037 | 7.873 | 0.051 |
| 4108 | **Cpsf2** | Unchanged | O35218 | 1 | 6 | 1 | 782 | 88.3 | 5.11 | 2 | 1.116 | 0.791 | 2.264 | 0.244 | 1.083 | 0.808 | 2.091 | 0.275 |
| 4109 | **Btbd18** | Unchanged | A0A0A6YY25 | 1 | 4 | 1 | 723 | 79.4 | 4.97 | 2 | 0.696 | 0.589 | 0.005 | 0.053 | 0.323 | 0.184 | 0.015 | 0.073 |
| 4110 | **Ppfibp1** | Down | Q8C8U0 | 1 | 2 | 1 | 969 | 108.5 | 5.49 | 2 | 0.744 | 0.236 | 1.066 | 0.783 | 0.568 | 0.043 | 1.878 | 0.100 |
| 4111 | **Mtif2** | Unchanged | Q91YJ5 | 1 | 1 | 1 | 727 | 81.2 | 7.24 | 2 | 1.371 | 0.708 | 1.531 | 0.689 | 0.083 | 0.315 | 18.383 | 0.168 |
| 4112 | **Ints3** | Down | Q7TPD0 | 1 | 4 | 1 | 1041 | 117.9 | 5.8 | 2 | 2.321 | 0.476 | 2.174 | 0.295 | 0.444 | 0.012 | 4.901 | 0.137 |
| 4113 | **Heatr3** | Unchanged | Q8BQM4 | 1 | 1 | 1 | 679 | 74.3 | 5.01 | 2 | 1.512 | 0.409 | 2.401 | 0.223 | 0.630 | 0.331 | 3.810 | 0.141 |
| 4114 | **Ube4b** | Unchanged | Q9ES00 | 1 | 1 | 1 | 1173 | 133.2 | 6.07 | 2 | 1.349 | 0.654 | 1.787 | 0.576 | 1.525 | 0.603 | 1.172 | 0.863 |
| 4115 | **Eml4** | Unchanged | Q3UMY5 | 1 | 2 | 1 | 988 | 110 | 6.54 | 2 | 0.415 | 0.178 | 0.595 | 0.347 | 0.434 | 0.197 | 1.372 | 0.520 |
| 4116 | **Chd4** | Unchanged | Q6PDQ2 | 2 | 4 | 2 | 1915 | 217.6 | 5.81 | 2 | 0.899 | 0.848 | 0.478 | 0.228 | 0.234 | 0.066 | 2.043 | 0.271 |
| 4117 | **Dhx32** | Unchanged | Q8BZS9 | 1 | 6 | 1 | 744 | 83.9 | 4.96 | 2 | 2.126 | 0.160 | 3.286 | 0.054 | 1.692 | 0.227 | 1.942 | 0.161 |
| 4118 | **Itga1** | Unchanged | Q3V3R4 | 2 | 14 | 2 | 1179 | 130.7 | 6.2 | 2 | 0.613 | 0.420 | 1.591 | 0.485 | 0.529 | 0.288 | 3.007 | 0.180 |
| 4119 | **Pias2** | Unchanged | Q8C5D8 | 1 | 1 | 1 | 621 | 68.4 | 7.65 | 2 | 0.638 | 0.663 | 1.842 | 0.544 | 0.404 | 0.449 | 4.553 | 0.229 |
| 4120 | **Wasl** | Down | Q91YD9 | 1 | 1 | 1 | 501 | 54.2 | 7.93 | 2 | 0.142 | 0.002 | 0.564 | 0.371 | 0.007 | 0.000 | 85.494 | 0.229 |
| 4121 | **Helz2** | Unchanged | E9QAM5 | 3 | 4 | 3 | 2947 | 331.4 | 8.07 | 2 | 1.511 | 0.450 | 2.209 | 0.170 | 1.163 | 0.689 | 1.900 | 0.234 |
| 4122 | **Crnkl1** | Unchanged | P63154 | 1 | 9 | 1 | 690 | 83.4 | 6.93 | 2 | 1.086 | 0.899 | 1.066 | 0.946 | 1.594 | 0.519 | 0.669 | 0.649 |
| 4123 | **Gtf2e1** | Unchanged | Q9D0D5 | 1 | 2 | 1 | 440 | 49.6 | 4.86 | 2 | #DIV/0! | #DIV/0! | #DIV/0! | #DIV/0! | #DIV/0! | #DIV/0! | #DIV/0! | #DIV/0! |
| 4124 | **Dnajb12** | Down | Q8C4C9 | 1 | 8 | 1 | 378 | 42.4 | 8.5 | 2 | 0.860 | 0.706 | 0.268 | 0.046 | 0.584 | 0.207 | 0.460 | 0.136 |
| 4125 | **Mroh1** | Unchanged | E0CZ22 | 2 | 2 | 2 | 1640 | 181.7 | 6.39 | 2 | 0.723 | 0.597 | 0.527 | 0.355 | 0.197 | 0.102 | 2.671 | 0.157 |
| 4126 | **Bicdl2** | Unchanged | Q8CHW5 | 1 | 2 | 1 | 507 | 57.5 | 4.97 | 2 | 1.115 | 0.878 | 0.874 | 0.837 | 1.100 | 0.886 | 0.794 | 0.750 |
| 4127 | **Itsn1** | Unchanged | Q9Z0R4 | 2 | 3 | 2 | 1714 | 194.2 | 7.91 | 2 | 0.704 | 0.663 | 0.364 | 0.209 | 0.816 | 0.745 | 0.446 | 0.200 |
| 4128 | **Ube3c** | Unchanged | Q80U95 | 1 | 1 | 1 | 1083 | 123.9 | 6.39 | 2 | 0.616 | 0.252 | 0.554 | 0.204 | 1.008 | 0.983 | 0.550 | 0.141 |
| 4129 | **Agmo** | Unchanged | Q8BS35 | 1 | 8 | 1 | 447 | 51.7 | 7.27 | 2 | 1.054 | 0.875 | 1.066 | 0.881 | 0.840 | 0.661 | 1.268 | 0.512 |
| 4130 | **Aak1** | UP | Q3UHJ0 | 1 | 1 | 1 | 959 | 103.3 | 6.7 | 2 | 28.041 | 0.137 | 45.089 | 0.240 | 88.463 | 0.022 | 0.510 | 0.377 |

| | | | | | | | | | | | | | | | | | | |
|---|---|---|---|---|---|---|---|---|---|---|---|---|---|---|---|---|---|---|
| 4131 | **Sidt2** | Unchanged | Q8CIF6 | 1 | 1 | 1 | 832 | 94.4 | 7.21 | 2 | 0.315 | 0.072 | 3.390 | 0.068 | 1.828 | 0.584 | 1.854 | 0.407 |
| 4132 | **A2m** | Unchanged | Q6GQT1 | 2 | 11 | 1 | 1474 | 164.2 | 6.61 | 2 | #DIV/0! | #DIV/0! | #DIV/0! | #DIV/0! | #DIV/0! | #DIV/0! | #DIV/0! | #DIV/0! |
| 4133 | **Acvr2a** | Unchanged | P27038 | 1 | 1 | 1 | 513 | 57.9 | 5.92 | 2 | 0.562 | 0.510 | 0.674 | 0.645 | 0.126 | 0.151 | 5.341 | 0.210 |
| 4134 | **Def6** | Down | A0A3B2W417 | 1 | 1 | 1 | 385 | 45 | 6.67 | 2 | 0.460 | 0.226 | 0.167 | 0.069 | 0.081 | 0.048 | 2.048 | 0.271 |
| 4135 | **Zfp292** | Down | Q9Z2U2 | 1 | 1 | 1 | 2698 | 300.9 | 7.53 | 2 | 0.474 | 0.016 | 0.916 | 0.681 | 0.744 | 0.276 | 1.231 | 0.524 |
| 4136 | **Hip1** | Unchanged | Q8VD75 | 1 | 1 | 1 | 1029 | 115.1 | 5.43 | 2 | 0.917 | 0.851 | 1.368 | 0.403 | 1.256 | 0.528 | 1.089 | 0.735 |
| 4137 | **Far2** | Down | Q7TNT2 | 1 | 2 | 1 | 515 | 59.1 | 9.19 | | 0.840 | 0.546 | 0.841 | 0.610 | 0.487 | 0.015 | 1.726 | 0.248 |
| 4138 | **Itga2b** | Unchanged | Q9QUM0 | 1 | 2 | 1 | 1033 | 112.6 | 5.76 | 2 | 0.625 | 0.401 | 0.308 | 0.134 | 0.277 | 0.112 | 1.113 | 0.898 |
| 4139 | **Tfap2d** | Unchanged | Q91ZK0 | 1 | 1 | 1 | 452 | 49.5 | 8.16 | | #DIV/0! | #DIV/0! | #DIV/0! | #DIV/0! | #DIV/0! | #DIV/0! | #DIV/0! | #DIV/0! |
| 4140 | **Atf7ip2** | Unchanged | Q3UL97 | 1 | 2 | 1 | 452 | 50.9 | 9.45 | | 3.257 | 0.268 | 3.312 | 0.336 | 4.552 | 0.176 | 0.728 | 0.687 |
| 4141 | **Cd44** | UP | P15379 | 1 | 3 | 1 | 778 | 85.6 | 4.96 | 2 | 0.787 | 0.606 | 1.246 | 0.725 | 4.367 | 0.009 | 0.285 | 0.021 |
| 4142 | **Blnk** | Unchanged | Q9QUN3 | 1 | 3 | 1 | 457 | 50.6 | 8.25 | | 1.099 | 0.898 | 1.759 | 0.538 | 1.279 | 0.682 | 1.376 | 0.677 |
| 4143 | **Ddx11** | Unchanged | A0A286YDY6 | 1 | 5 | 1 | 393 | 45.2 | 5.64 | 2 | 1.232 | 0.793 | 0.879 | 0.872 | 0.137 | 0.137 | 6.420 | 0.188 |
| 4144 | **Ppp3cb** | Unchanged | P48453 | 1 | 1 | 1 | 525 | 59.1 | 5.91 | 2 | 1.331 | 0.705 | 1.474 | 0.675 | 0.249 | 0.129 | 5.926 | 0.250 |
| 4145 | **Rmc1** | Unchanged | Q8VC42 | 1 | 1 | 1 | 657 | 74.9 | 7.83 | 2 | 0.566 | 0.290 | 1.834 | 0.375 | 0.556 | 0.264 | 3.295 | 0.159 |
| 4146 | **Itga3** | Unchanged | Q62470 | 1 | 1 | 1 | 1053 | 116.7 | 6.57 | 2 | #DIV/0! | #DIV/0! | #DIV/0! | #DIV/0! | #DIV/0! | #DIV/0! | #DIV/0! | #DIV/0! |
| 4147 | **Slc39a8** | Unchanged | Q91W10 | 1 | 1 | 1 | 462 | 50.1 | 5.86 | 2 | 13.908 | 0.068 | 15.255 | 0.145 | 19.347 | 0.065 | 0.789 | 0.748 |
| 4148 | **Vars2** | Unchanged | Q3U2A8 | 1 | 8 | 1 | 1060 | 118.7 | 7.14 | 2 | 1.516 | 0.330 | 2.580 | 0.186 | 1.613 | 0.137 | 1.599 | 0.390 |
| 4149 | **Gapvd1** | Unchanged | Q6PAR5 | 2 | 2 | 2 | 1458 | 162.3 | 5.19 | 2 | 0.363 | 0.256 | 0.136 | 0.098 | 0.572 | 0.420 | 0.238 | 0.068 |
| 4150 | **Vps33a** | Unchanged | Q9D2N9 | 1 | 4 | 1 | 598 | 67.5 | 7.08 | 2 | 1.094 | 0.817 | 1.202 | 0.628 | 0.571 | 0.106 | 2.105 | 0.097 |
| 4151 | **Bbof1** | Unchanged | Q3V079 | 1 | 1 | 1 | 533 | 62.5 | 9.19 | 2 | 1.453 | 0.547 | 1.268 | 0.691 | 1.124 | 0.840 | 1.128 | 0.843 |
| 4152 | **Zfp948** | Unchanged | Q6DFU8 | 1 | 2 | 1 | 667 | 78.1 | 8.81 | 1 | 2.109 | 0.339 | 1.487 | 0.685 | 3.141 | 0.194 | 0.473 | 0.385 |
| 4153 | **Dym** | Unchanged | Q8CHY3 | 1 | 2 | 1 | 669 | 75.8 | 5.77 | 1 | 0.658 | 0.260 | 0.954 | 0.890 | 0.684 | 0.261 | 1.395 | 0.337 |
| 4154 | **Gm4847** | Unchanged | G3X946 | 1 | 1 | 1 | 537 | 61.3 | 9.16 | 1 | #DIV/0! | #DIV/0! | #DIV/0! | #DIV/0! | #DIV/0! | #DIV/0! | #DIV/0! | #DIV/0! |
| 4155 | **Traf4** | Down | Q61382 | 1 | 1 | 1 | 470 | 53.5 | 8.15 | 1 | 0.615 | 0.317 | 1.672 | 0.289 | 0.059 | 0.001 | 28.389 | 0.021 |
| 4156 | **Efcab12** | Unchanged | V9GXH0 | 1 | 1 | 1 | 673 | 77.7 | 9.82 | 1 | 0.939 | 0.935 | 0.639 | 0.539 | 2.001 | 0.308 | 0.319 | 0.143 |
| 4157 | **Abcg8** | Unchanged | Q9DBM0 | 1 | 3 | 1 | 673 | 75.9 | 7.33 | 1 | 1.233 | 0.527 | 1.119 | 0.734 | 0.591 | 0.124 | 1.893 | 0.136 |
| 4158 | **Ptpn5** | Unchanged | P54830 | 1 | 8 | 1 | 541 | 60.8 | 5.2 | 1 | 1.664 | 0.419 | 1.163 | 0.813 | 1.151 | 0.804 | 1.010 | 0.988 |
| 4159 | **Terf2** | Unchanged | O35144 | 1 | 1 | 1 | 541 | 60.2 | 9.5 | 1 | #DIV/0! | #DIV/0! | #DIV/0! | #DIV/0! | #DIV/0! | #DIV/0! | #DIV/0! | #DIV/0! |
| 4160 | **Msh6** | Down | P54276 | 2 | 3 | 2 | 1358 | 151 | 6.73 | 1 | 0.822 | 0.773 | 0.565 | 0.381 | 0.003 | 0.014 | 196.917 | 0.141 |
| 4161 | **Mfsd12** | Unchanged | Q3U481 | 1 | 22 | 1 | 476 | 51.5 | 7.81 | 1 | 0.349 | 0.178 | 0.166 | 0.080 | 0.213 | 0.096 | 0.778 | 0.722 |
| 4162 | **Wdr91** | Unchanged | Q7TMQ7 | 1 | 3 | 1 | 748 | 83.4 | 6.64 | 1 | #DIV/0! | #DIV/0! | #DIV/0! | #DIV/0! | #DIV/0! | #DIV/0! | #DIV/0! | #DIV/0! |
| 4163 | **Serpina3i** | Unchanged | D3Z450 | 1 | 4 | 1 | 408 | 45.7 | 6.95 | 1 | 6.344 | 0.321 | 0.024 | 0.235 | 2.499 | 0.420 | 0.010 | 0.159 |
| 4164 | **Aldh3b2** | Unchanged | E9Q3E1 | 1 | 6 | 1 | 479 | 52.9 | 6.09 | 1 | 2.431 | 0.235 | 0.822 | 0.791 | 0.174 | 0.133 | 4.736 | 0.166 |
| 4165 | **Rnf157** | Unchanged | Q3TEL6 | 1 | 1 | 1 | 685 | 74.5 | 4.81 | 1 | 2.615 | 0.225 | 2.707 | 0.274 | 0.500 | 0.243 | 5.416 | 0.156 |
| 4166 | **Coro2b** | Unchanged | Q8BH44 | 1 | 1 | 1 | 480 | 54.9 | 8.27 | 1 | 0.885 | 0.802 | 1.461 | 0.553 | 1.526 | 0.401 | 0.958 | 0.935 |
| 4167 | **Gm14226** | Unchanged | Q3TZL0 | 1 | 1 | 1 | 686 | 76.3 | 8.06 | 1 | 0.055 | 0.071 | 0.096 | 0.082 | 0.124 | 0.092 | 0.775 | 0.699 |
| 4168 | **Phka2** | Unchanged | Q8BWJ3 | 1 | 2 | 1 | 1235 | 138.4 | 6.37 | 1 | 2.057 | 0.392 | 2.085 | 0.393 | 0.550 | 0.054 | 3.793 | 0.238 |
| 4169 | **Myb** | UP | A0A087WPA7 | 1 | 1 | 1 | 755 | 84.1 | 7.3 | 1 | 2.212 | 0.010 | 1.273 | 0.681 | 601.636 | 0.044 | 0.002 | 0.044 |
| 4170 | **Abi1** | Unchanged | Q8CBW3 | 1 | 5 | 1 | 481 | 52.3 | 7.64 | 1 | 1.207 | 0.658 | 0.658 | 0.432 | 0.928 | 0.879 | 0.709 | 0.625 |
| 4171 | **Slc22a26** | Unchanged | Q91WJ2 | 1 | 5 | 1 | 552 | 61.3 | 8.27 | 1 | 1.378 | 0.630 | 2.273 | 0.257 | 1.023 | 0.969 | 2.222 | 0.266 |

| | | | | | | | | | | | | | | | | | | |
|---|---|---|---|---|---|---|---|---|---|---|---|---|---|---|---|---|---|---|
| 4172 | **Nxf3** | Unchanged | Q4ZGD9 | 1 | 4 | 1 | 553 | 63.6 | 8.13 | 1 | 1.408 | 0.704 | 0.805 | 0.820 | 1.145 | 0.877 | 0.703 | 0.700 |
| 4173 | **Nt5dc2** | Unchanged | A0A2I3BR81 | 1 | 1 | 1 | 553 | 63.8 | 7.59 | 1 | 1.202 | 0.842 | 1.485 | 0.676 | 0.032 | 0.161 | 45.885 | 0.156 |
| 4174 | **Nup214** | Unchanged | Q80U93 | 1 | 3 | 1 | 2085 | 212.8 | 7.08 | 1 | 0.633 | 0.284 | 1.512 | 0.093 | 0.863 | 0.706 | 1.752 | 0.093 |
| 4175 | **Lrba** | Down | E9Q3Y4 | 3 | 5 | 3 | 2854 | 316.6 | 5.54 | 1 | 0.366 | 0.089 | 0.254 | 0.033 | 0.452 | 0.131 | 0.562 | 0.301 |
| 4176 | **Fmn1** | Unchanged | Q05860 | 2 | 2 | 2 | 1466 | 163.5 | 8.59 | 1 | 0.777 | 0.686 | 1.368 | 0.649 | 1.600 | 0.300 | 0.855 | 0.788 |
| 4177 | **Rai14** | Unchanged | Q9EP71 | 1 | 6 | 1 | 979 | 108.8 | 6.27 | 1 | 0.938 | 0.909 | 1.090 | 0.864 | 1.014 | 0.982 | 1.075 | 0.887 |
| 4178 | **Nek9** | Unchanged | Q8K1R7 | 1 | 2 | 1 | 984 | 107.1 | 5.63 | 1 | 0.912 | 0.868 | 1.089 | 0.886 | 0.522 | 0.325 | 2.089 | 0.319 |
| 4179 | **She** | Unchanged | Q8BSD5 | 1 | 1 | 1 | 492 | 53.3 | 9.17 | 1 | 0.835 | 0.636 | 0.422 | 0.189 | 1.088 | 0.896 | 0.388 | 0.312 |
| 4180 | **Map2** | Unchanged | P20357 | 1 | 1 | 1 | 1828 | 199 | 4.91 | 1 | #DIV/0! | #DIV/0! | #DIV/0! | #DIV/0! | #DIV/0! | #DIV/0! | #DIV/0! | #DIV/0! |
| 4181 | **C2cd2l** | Unchanged | Q80X80 | 1 | 1 | 1 | 706 | 76.3 | 7.21 | 1 | 1.657 | 0.415 | 3.085 | 0.174 | 0.676 | 0.321 | 4.562 | 0.125 |
| 4182 | **Map1a** | Unchanged | A2ARP8 | 1 | 1 | 1 | 3014 | 325.7 | 5.01 | 1 | #DIV/0! | #DIV/0! | #DIV/0! | #DIV/0! | #DIV/0! | #DIV/0! | #DIV/0! | #DIV/0! |
| 4183 | **Rsrc2** | Unchanged | S4R2L4 | 1 | 1 | 1 | 434 | 50.6 | 11.36 | 1 | 0.845 | 0.845 | 0.361 | 0.332 | 0.393 | 0.400 | 0.918 | 0.938 |
| 4184 | **Arntl2** | Unchanged | Q2VPD4 | 1 | 2 | 1 | 579 | 64.4 | 8.09 | 1 | 0.868 | 0.854 | 0.006 | 0.067 | 0.110 | 0.096 | 0.058 | 0.044 |
| 4185 | **Ppm1j** | Unchanged | Q149T7 | 1 | 2 | 1 | 507 | 55.5 | 7.84 | 1 | 1.702 | 0.228 | 1.733 | 0.192 | 1.002 | 0.997 | 1.729 | 0.190 |
| 4186 | **Scel** | Unchanged | Q9EQG3 | 1 | 8 | 1 | 652 | 72.9 | 9.47 | 1 | 1.547 | 0.526 | 1.413 | 0.749 | 1.794 | 0.438 | 0.788 | 0.802 |
| 4187 | **Gm49368** | Unchanged | Q3U1U4 | 1 | 1 | 1 | 1232 | 135.9 | 8.21 | 1 | 0.835 | 0.776 | 0.928 | 0.902 | 0.490 | 0.273 | 1.893 | 0.288 |
| 4188 | **Dis3l2** | Unchanged | Q8CI75 | 1 | 1 | 1 | 870 | 97.7 | 5.9 | 1 | #DIV/0! | #DIV/0! | #DIV/0! | #DIV/0! | #DIV/0! | #DIV/0! | #DIV/0! | #DIV/0! |
| 4189 | **Sass6** | Unchanged | Q80UK7 | 1 | 3 | 1 | 654 | 74 | 7.37 | 1 | 0.839 | 0.774 | 0.790 | 0.725 | 0.861 | 0.809 | 0.917 | 0.902 |
| 4190 | **Nepn** | Unchanged | Q9CQ76 | 1 | 8 | 1 | 512 | 57.7 | 8.24 | | 1.023 | 0.972 | 0.511 | 0.402 | 0.771 | 0.721 | 0.663 | 0.617 |
| 4191 | **Asic2** | Unchanged | Q925H0 | 1 | 3 | 1 | 512 | 57.7 | 5.19 | | 0.952 | 0.956 | 0.411 | 0.402 | 0.020 | 0.141 | 20.152 | 0.219 |
| 4192 | **Myom2** | Unchanged | Q14BI5 | 1 | 1 | 1 | 1463 | 164.6 | 5.77 | 1 | 0.438 | 0.249 | 0.319 | 0.173 | 0.383 | 0.206 | 0.834 | 0.677 |
| 4193 | **Zfhx4** | Unchanged | H3BLK8 | 2 | 4 | 2 | 3606 | 397.8 | 6.37 | 1 | 0.855 | 0.857 | 0.338 | 0.330 | 0.051 | 0.154 | 6.612 | 0.220 |
| 4194 | **Osbpl9** | Unchanged | A2A8Z1 | 1 | 2 | 1 | 736 | 83.1 | 6.07 | 1 | 1.412 | 0.645 | 1.282 | 0.775 | 0.339 | 0.279 | 3.787 | 0.263 |
| 4195 | **Prox1** | Unchanged | P48437 | 1 | 4 | 1 | 737 | 83.1 | 7.08 | 1 | 19.244 | 0.303 | 2.469 | 0.080 | 3.361 | 0.089 | 0.735 | 0.536 |
| 4196 | **Vcam1** | Unchanged | P29533 | 1 | 1 | 1 | 739 | 81.3 | 5.3 | 1 | 1.732 | 0.316 | 2.100 | 0.250 | 2.219 | 0.123 | 0.947 | 0.912 |
| 4197 | **Cercam** | Unchanged | A3KGW5 | 1 | 1 | 1 | 592 | 67.6 | 6.76 | 1 | #DIV/0! | #DIV/0! | #DIV/0! | #DIV/0! | #DIV/0! | #DIV/0! | #DIV/0! | #DIV/0! |
| 4198 | **Frrs1** | Unchanged | A0A0G2JFP4 | 1 | 2 | 1 | 592 | 66 | 8.32 | 1 | 0.715 | 0.699 | 0.960 | 0.956 | 0.102 | 0.169 | 9.448 | 0.045 |
| 4199 | **Ints1** | Unchanged | A0A0G2JH17 | 1 | 1 | 1 | 2222 | 247.8 | 6.34 | 1 | 0.731 | 0.408 | 0.478 | 0.074 | 0.677 | 0.331 | 0.707 | 0.408 |
| 4200 | **Mtf2** | Down | Q02395 | 1 | 1 | 1 | 593 | 66.9 | 8.68 | 1 | 0.884 | 0.829 | 0.314 | 0.189 | 0.009 | 0.046 | 36.000 | 0.222 |
| 4201 | **Mtrex** | Down | Q9CZU3 | 1 | 2 | 1 | 1040 | 117.6 | 6.4 | 1 | 0.317 | 0.078 | 0.240 | 0.045 | 0.427 | 0.137 | 0.561 | 0.268 |
| 4202 | **Tbc1d9b** | Down | Q5SVR0 | 2 | 3 | 1 | 1263 | 141.7 | 5.29 | 1 | 0.995 | 0.982 | 1.079 | 0.853 | 0.209 | 0.007 | 5.157 | 0.042 |
| 4203 | **Pank4** | Unchanged | Q80YV4 | 1 | 2 | 1 | 820 | 91.5 | 6.71 | 1 | 1.656 | 0.216 | 22.366 | 0.277 | 4.473 | 0.310 | 5.000 | 0.364 |
| 4204 | **Susd2** | Down | Q9DBX3 | 1 | 1 | 1 | 820 | 90.6 | 6.62 | 1 | 0.122 | 0.008 | 1.051 | 0.934 | 1.043 | 0.952 | 1.007 | 0.993 |
| 4205 | **Zbtb18** | Unchanged | Q9WUK6 | 1 | 2 | 1 | 522 | 58.3 | 5.69 | 1 | 0.241 | 0.230 | 0.471 | 0.417 | 1.669 | 0.504 | 0.282 | 0.185 |
| 4206 | **Lzts1** | Unchanged | P60853 | 1 | 2 | 1 | 599 | 67.2 | 7.56 | 1 | 0.643 | 0.553 | 0.581 | 0.482 | 0.841 | 0.804 | 0.691 | 0.572 |
| 4207 | **Cenpb** | Unchanged | P27790 | 1 | 13 | 1 | 599 | 65.3 | 4.53 | 1 | 2.255 | 0.343 | 1.866 | 0.365 | 1.717 | 0.470 | 1.087 | 0.899 |
| 4208 | **D430041D05Rik** | Unchanged | A0A2R8VKG2 | 1 | 1 | 1 | 2099 | 225.8 | 9.14 | 1 | #DIV/0! | #DIV/0! | #DIV/0! | #DIV/0! | #DIV/0! | #DIV/0! | #DIV/0! | #DIV/0! |
| 4209 | **Efl1** | Unchanged | Q8C0D5 | 1 | 4 | 1 | 1127 | 125.7 | 6.16 | 1 | 0.865 | 0.809 | 0.524 | 0.352 | 0.346 | 0.212 | 1.512 | 0.407 |
| 4210 | **Epha2** | Unchanged | Q03145 | 1 | 1 | 1 | 977 | 108.8 | 6.23 | 1 | 2.035 | 0.630 | 0.085 | 0.262 | 0.200 | 0.330 | 0.427 | 0.437 |
| 4211 | **Inava** | Unchanged | G3X9Z8 | 1 | 1 | 1 | 677 | 74.1 | 8.82 | 1 | 1.194 | 0.750 | 3.622 | 0.324 | 1.898 | 0.392 | 1.908 | 0.528 |
| 4212 | **Pole2** | Unchanged | O54956 | 1 | 1 | 1 | 527 | 59.3 | 6.25 | | 2.234 | 0.302 | 4.094 | 0.211 | 3.476 | 0.193 | 1.178 | 0.827 |

| | | | | | | | | | | | | | | | | | | |
|---|---|---|---|---|---|---|---|---|---|---|---|---|---|---|---|---|---|---|
| 4213 | **Traf6** | Down | P70196 | 1 | 1 | 1 | 530 | 60 | 6.55 | | 0.370 | 0.146 | 0.223 | 0.059 | 0.081 | 0.019 | 2.753 | 0.417 |
| 4214 | **Slc29a2** | Unchanged | Q61672 | 1 | 2 | 1 | 456 | 50.2 | 6.51 | 1 | 1.155 | 0.558 | 0.989 | 0.975 | 0.913 | 0.756 | 1.083 | 0.829 |
| 4215 | **Kntc1** | Unchanged | Q8C3Y4 | 1 | 1 | 1 | 2207 | 250.2 | 6 | 1 | 0.494 | 0.507 | 0.524 | 0.532 | 2.046 | 0.377 | 0.256 | 0.165 |
| 4216 | **Fam184a** | Unchanged | E9PW83 | 1 | 1 | 1 | 1143 | 132.9 | 5.96 | 1 | #DIV/0! | #DIV/0! | #DIV/0! | #DIV/0! | #DIV/0! | #DIV/0! | #DIV/0! | #DIV/0! |
| 4217 | **Krt78** | Unchanged | E9Q0F0 | 2 | 19 | 2 | 1068 | 112.2 | 7.97 | 1 | 3.386 | 0.217 | 1.836 | 0.394 | 1.745 | 0.343 | 1.052 | 0.933 |
| 4218 | **Elf1** | Unchanged | Q60775 | 1 | 1 | 1 | 612 | 66.2 | 4.97 | 1 | 2.005 | 0.364 | 1.043 | 0.969 | 1.989 | 0.362 | 0.524 | 0.488 |
| 4219 | **Fgd4** | Unchanged | Q91ZT5 | 1 | 2 | 1 | 766 | 86.5 | 5.95 | 1 | 7.627 | 0.212 | 10.509 | 0.160 | 7.819 | 0.085 | 1.344 | 0.711 |
| 4220 | **Top1** | Unchanged | Q04750 | 1 | 1 | 1 | 767 | 90.8 | 9.33 | 1 | 0.841 | 0.778 | 3.913 | 0.121 | 1.723 | 0.348 | 2.271 | 0.250 |
| 4221 | **Slmap** | Down | Q3URD3 | 1 | 3 | 1 | 845 | 96.9 | 5.33 | 1 | 0.208 | 0.011 | 0.429 | 0.086 | 0.115 | 0.004 | 3.735 | 0.139 |
| 4222 | **Gnl3** | Unchanged | Q8CI11 | 1 | 4 | 1 | 538 | 60.7 | 9.11 | 1 | 0.739 | 0.344 | 1.534 | 0.439 | 0.784 | 0.359 | 1.958 | 0.272 |
| 4223 | **Rundc1** | Unchanged | Q0VDN7 | 1 | 1 | 1 | 615 | 67.8 | 5.78 | | 0.094 | 0.246 | 0.018 | 0.211 | 0.186 | 0.295 | 0.095 | 0.048 |
| 4224 | **Taok1** | Unchanged | Q5F2E8 | 1 | 5 | 1 | 1001 | 116 | 7.55 | 1 | 0.930 | 0.909 | 0.883 | 0.845 | 0.594 | 0.443 | 1.486 | 0.560 |
| 4225 | **Ttc39b** | Unchanged | Q8BYY4 | 1 | 2 | 1 | 617 | 70.2 | 6.64 | 1 | #DIV/0! | #DIV/0! | #DIV/0! | #DIV/0! | #DIV/0! | #DIV/0! | #DIV/0! | #DIV/0! |
| 4226 | **Dnah1** | Unchanged | E9Q8T7 | 2 | 2 | 2 | 4250 | 486.8 | 5.57 | 1 | 1.043 | 0.945 | 3.738 | 0.294 | 0.980 | 0.974 | 3.812 | 0.282 |
| 4227 | **Pdcd11** | Unchanged | Q6NS46 | 1 | 1 | 1 | 1862 | 207.6 | 8.75 | 1 | 0.713 | 0.413 | 2.464 | 0.320 | 0.681 | 0.309 | 3.619 | 0.224 |
| 4228 | **Lrrc29** | Unchanged | A0A1B0GQY9 | 1 | 7 | 1 | 621 | 66.9 | 6.57 | 1 | 0.905 | 0.873 | 0.458 | 0.365 | 1.406 | 0.404 | 0.326 | 0.064 |
| 4229 | **Tnks** | Unchanged | Q6PFX9 | 1 | 1 | 1 | 1320 | 140.9 | 7.05 | 1 | 4.151 | 0.311 | 6.562 | 0.106 | 1.132 | 0.825 | 5.795 | 0.109 |
| 4230 | **Nt5dc1** | Unchanged | Q8C5P5 | 1 | 1 | 1 | 467 | 53.1 | 6.33 | 1 | 0.493 | 0.223 | 3.121 | 0.342 | 0.263 | 0.084 | 11.847 | 0.204 |
| 4231 | **9930021J03Rik** | Unchanged | H3BKP8 | 1 | 2 | 1 | 2027 | 218.8 | 9.04 | 1 | 0.607 | 0.665 | 0.233 | 0.385 | 1.116 | 0.911 | 0.209 | 0.154 |
| 4232 | **Keap1** | Unchanged | Q9Z2X8 | 1 | 3 | 1 | 624 | 69.5 | 6.44 | 1 | 0.962 | 0.947 | 0.655 | 0.558 | 0.329 | 0.112 | 1.992 | 0.481 |
| 4233 | **Inpp5k** | Unchanged | Q8C5L6 | 1 | 3 | 1 | 468 | 54.1 | 5.88 | 1 | 0.532 | 0.591 | 0.893 | 0.927 | 0.689 | 0.730 | 1.296 | 0.815 |
| 4234 | **Glg1** | UP | Q61543 | 1 | 6 | 1 | 1175 | 133.6 | 6.84 | 1 | 52.394 | 0.307 | 31.234 | 0.131 | 7.119 | 0.009 | 4.387 | 0.218 |
| 4235 | **Erf** | Unchanged | P70459 | 1 | 1 | 1 | 551 | 59 | 7.28 | | #DIV/0! | #DIV/0! | #DIV/0! | #DIV/0! | #DIV/0! | #DIV/0! | #DIV/0! | #DIV/0! |
| 4236 | **Poln** | Down | A0A0R4J0E3 | 1 | 1 | 1 | 866 | 96.9 | 7.84 | 1 | 1.930 | 0.199 | 0.291 | 0.145 | 0.045 | 0.048 | 6.531 | 0.156 |
| 4237 | **Cadm1** | Unchanged | E9PYN1 | 1 | 1 | 1 | 474 | 51.7 | 4.87 | 1 | 0.384 | 0.166 | 3.536 | 0.296 | 1.407 | 0.424 | 2.514 | 0.376 |
| 4238 | **Tbx2** | Unchanged | Q60707 | 1 | 3 | 1 | 711 | 75 | 9.14 | 1 | 1.122 | 0.867 | 0.383 | 0.294 | 0.981 | 0.977 | 0.390 | 0.283 |
| 4239 | **Zfp251** | Unchanged | Q6PCX8 | 1 | 1 | 1 | 632 | 70.9 | 8.4 | 1 | 672.100 | 0.091 | 694.700 | 0.054 | 632.200 | 0.086 | 1.099 | 0.892 |
| 4240 | **Gpr179** | Down | E9PY61 | 1 | 1 | 1 | 2293 | 248.6 | 5.55 | 1 | 0.541 | 0.046 | 1.445 | 0.699 | 0.516 | 0.056 | 2.798 | 0.424 |
| 4241 | **Mageb4** | Unchanged | A2A9R3 | 1 | 1 | 1 | 556 | 60.9 | 9.16 | 1 | 0.418 | 0.349 | 0.676 | 0.597 | 0.341 | 0.290 | 1.984 | 0.138 |
| 4242 | **Casd1** | UP | Q7TN73 | 1 | 1 | 1 | 797 | 91.5 | 8.76 | 1 | 2.260 | 0.043 | 3.802 | 0.363 | 4.842 | 0.007 | 0.785 | 0.746 |
| 4243 | **Ubr2** | Unchanged | Q6WKZ8 | 1 | 1 | 1 | 1755 | 199 | 6.33 | 1 | 3.274 | 0.199 | 1.876 | 0.297 | 2.401 | 0.176 | 0.781 | 0.578 |
| 4244 | **Col15a1** | Unchanged | O35206 | 2 | 6 | 2 | 1367 | 140.4 | 4.89 | 1 | 0.864 | 0.801 | 0.958 | 0.946 | 0.473 | 0.275 | 2.024 | 0.311 |
| 4245 | **Cr1l** | Unchanged | Q64735 | 1 | 3 | 1 | 483 | 53.7 | 6.65 | 1 | 0.845 | 0.835 | 1.762 | 0.441 | 1.049 | 0.954 | 1.680 | 0.472 |
| 4246 | **Map3k6** | Unchanged | A0A0A0MQ82 | 1 | 2 | 1 | 1291 | 142.9 | 6.62 | 1 | 1.031 | 0.962 | 1.588 | 0.055 | 0.615 | 0.218 | 2.582 | 0.002 |
| 4247 | **Ripk3** | Unchanged | Q9QZL0 | 1 | 1 | 1 | 486 | 53.3 | 7.53 | 1 | #DIV/0! | #DIV/0! | #DIV/0! | #DIV/0! | #DIV/0! | #DIV/0! | #DIV/0! | #DIV/0! |
| 4248 | **Heatr9** | Down | Q5QNV8 | 1 | 4 | 1 | 569 | 65.8 | 9.7 | 1 | 0.938 | 0.851 | 1.040 | 0.863 | 0.401 | 0.032 | 2.596 | 0.009 |
| 4249 | **Npr1** | Unchanged | P18293 | 1 | 3 | 1 | 1057 | 119 | 6.89 | 1 | 2.004 | 0.341 | 0.985 | 0.987 | 2.276 | 0.298 | 0.433 | 0.362 |
| 4250 | **Zc3h4** | Unchanged | Q6ZPZ3 | 1 | 1 | 1 | 1304 | 140.9 | 6.27 | 1 | 1.366 | 0.490 | 2.702 | 0.212 | 1.158 | 0.732 | 2.334 | 0.247 |
| 4251 | **Focad** | Unchanged | A2AKG8 | 1 | 6 | 1 | 1798 | 198.8 | 6.74 | 1 | 1.838 | 0.138 | 1.493 | 0.407 | 1.314 | 0.569 | 1.135 | 0.766 |
| 4252 | **Clip1** | Down | Q922J3 | 1 | 2 | 1 | 1391 | 155.7 | 5.24 | 1 | 0.916 | 0.780 | 0.974 | 0.958 | 0.312 | 0.038 | 3.122 | 0.176 |
| 4253 | **Wdfy3** | Unchanged | G3UYW1 | 2 | 2 | 2 | 3526 | 394 | 6.77 | 1 | 0.716 | 0.450 | 0.900 | 0.820 | 1.377 | 0.322 | 0.654 | 0.260 |

| | | | | | | | | | | | | | | | | | | |
|---|---|---|---|---|---|---|---|---|---|---|---|---|---|---|---|---|---|---|
| 4254 | **Clca3a1** | Unchanged | Q9QX15 | 1 | 1 | 1 | 902 | 100 | 6.86 | 1 | 1.481 | 0.330 | 2.692 | 0.094 | 1.331 | 0.262 | 2.023 | 0.170 |
| 4255 | **Luzp1** | Unchanged | Q8R4U7 | 1 | 2 | 1 | 1068 | 119.2 | 7.99 | 1 | 0.787 | 0.761 | 0.271 | 0.170 | 0.245 | 0.154 | 1.109 | 0.737 |
| 4256 | **Strc** | Unchanged | Q8VIM6 | 1 | 1 | 1 | 1809 | 196.2 | 5.4 | 1 | 70.277 | 0.134 | 320.872 | 0.074 | 33.362 | 0.230 | 9.618 | 0.106 |
| 4257 | **Phka1** | Unchanged | P18826 | 1 | 2 | 1 | 1241 | 138.7 | 5.86 | 1 | 3.989 | 0.154 | 2.521 | 0.160 | 0.957 | 0.870 | 2.633 | 0.154 |
| 4258 | **Chuk** | Unchanged | Q60680 | 1 | 2 | 1 | 745 | 84.7 | 6.77 | 1 | 0.881 | 0.655 | 1.328 | 0.560 | 0.613 | 0.191 | 2.166 | 0.208 |
| 4259 | **Th** | Unchanged | P24529 | 1 | 2 | 1 | 498 | 56 | 6.15 | 1 | 2.464 | 0.342 | 0.584 | 0.482 | 0.100 | 0.105 | 5.867 | 0.120 |
| 4260 | **Atg2a** | Unchanged | Q6P4T0 | 1 | 1 | 1 | 1914 | 210.8 | 6.06 | 1 | #DIV/0! | #DIV/0! | #DIV/0! | #DIV/0! | #DIV/0! | #DIV/0! | #DIV/0! | #DIV/0! |
| 4261 | **Ppl** | Unchanged | Q9R269 | 2 | 6 | 2 | 1755 | 203.9 | 5.54 | 1 | 1.474 | 0.143 | 1.176 | 0.712 | 1.528 | 0.209 | 0.769 | 0.545 |
| 4262 | **Nexn** | Unchanged | A0A0G2JEX1 | 1 | 1 | 1 | 671 | 80 | 5.22 | 1 | 1.071 | 0.926 | 1.244 | 0.751 | 0.925 | 0.909 | 1.344 | 0.602 |
| 4263 | **Ptch1** | Unchanged | Q61115 | 1 | 1 | 1 | 1434 | 159.2 | 6.77 | 1 | 1.190 | 0.841 | 3.016 | 0.259 | 1.770 | 0.441 | 1.704 | 0.514 |
| 4264 | **Kmt2a** | Unchanged | P55200 | 1 | 1 | 1 | 3966 | 429.4 | 9.16 | 1 | #DIV/0! | #DIV/0! | #DIV/0! | #DIV/0! | #DIV/0! | #DIV/0! | #DIV/0! | #DIV/0! |
| 4265 | **Epor** | Unchanged | P14753 | 1 | 2 | 1 | 507 | 55.2 | 5 | 1 | 1.565 | 0.478 | 0.675 | 0.619 | 7.382 | 0.111 | 0.091 | 0.094 |
| 4266 | **Ptpn9** | Unchanged | O35239 | 1 | 1 | 1 | 593 | 67.9 | 8.1 | 1 | #DIV/0! | #DIV/0! | #DIV/0! | #DIV/0! | #DIV/0! | #DIV/0! | #DIV/0! | #DIV/0! |
| 4267 | **Zfp334** | Unchanged | A2A4U6 | 1 | 5 | 1 | 679 | 78.2 | 9.45 | 1 | 0.662 | 0.453 | 0.556 | 0.317 | 0.308 | 0.105 | 1.804 | 0.210 |
| 4268 | **Ifih1** | Unchanged | Q8R5F7 | 1 | 2 | 1 | 1025 | 115.9 | 6.25 | 1 | 0.893 | 0.675 | 2.108 | 0.192 | 1.206 | 0.344 | 1.747 | 0.276 |
| 4269 | **Usp49** | Unchanged | Q6P9L4 | 1 | 1 | 1 | 685 | 78.2 | 9.23 | 1 | #DIV/0! | #DIV/0! | #DIV/0! | #DIV/0! | #DIV/0! | #DIV/0! | #DIV/0! | #DIV/0! |
| 4270 | **Gimap8** | Unchanged | Q75N62 | 1 | 1 | 1 | 688 | 76.8 | 7.27 | 1 | 1.106 | 0.836 | 0.530 | 0.409 | 1.308 | 0.668 | 0.405 | 0.283 |
| 4271 | **Pkn1** | Unchanged | P70268 | 1 | 1 | 1 | 946 | 104.3 | 6.27 | 1 | 1.356 | 0.704 | 0.075 | 0.145 | 0.021 | 0.124 | 3.511 | 0.424 |
| 4272 | **Gigyf1** | Unchanged | Q99MR1 | 1 | 1 | 1 | 1044 | 116.2 | 5.39 | 1 | #DIV/0! | #DIV/0! | #DIV/0! | #DIV/0! | #DIV/0! | #DIV/0! | #DIV/0! | #DIV/0! |
| 4273 | **Msh4** | Unchanged | Q99MT2 | 1 | 4 | 1 | 958 | 106.9 | 8.18 | 1 | 1.141 | 0.788 | 0.033 | 0.076 | 1.592 | 0.496 | 0.021 | 0.051 |
| 4274 | **Dis3** | Unchanged | Q9CSH3 | 1 | 2 | 1 | 958 | 108.8 | 7.53 | 1 | 1.026 | 0.959 | 0.443 | 0.266 | 0.165 | 0.071 | 2.678 | 0.279 |
| 4275 | **Pfpl** | Unchanged | Q5RKV8 | 1 | 2 | 1 | 702 | 78 | 6.73 | 1 | #DIV/0! | #DIV/0! | #DIV/0! | #DIV/0! | #DIV/0! | #DIV/0! | #DIV/0! | #DIV/0! |
| 4276 | **Ccdc187** | Unchanged | A0A2I3BPI0 | 1 | 1 | 1 | 2202 | 239.3 | 6.61 | 1 | 0.914 | 0.879 | 0.323 | 0.143 | 0.277 | 0.096 | 1.168 | 0.851 |
| 4277 | **Setx** | Unchanged | A2AKX3 | 1 | 1 | 1 | 2646 | 297.4 | 6.83 | 1 | 1.230 | 0.604 | 1.800 | 0.198 | 1.086 | 0.800 | 1.658 | 0.222 |
| 4278 | **Fam234b** | Unchanged | Q8BYI8 | 1 | 1 | 1 | 624 | 67 | 5.03 | 1 | 44.000 | 0.108 | 43.356 | 0.311 | 63.144 | 0.090 | 0.687 | 0.706 |
| 4279 | **Srgap2** | Unchanged | Q91Z67 | 1 | 2 | 1 | 1071 | 120.7 | 6.64 | 1 | #DIV/0! | #DIV/0! | #DIV/0! | #DIV/0! | #DIV/0! | #DIV/0! | #DIV/0! | #DIV/0! |
| 4280 | **Tbc1d14** | Unchanged | G3UVU5 | 1 | 2 | 1 | 714 | 80.6 | 6.65 | 1 | 1.112 | 0.687 | 12.442 | 0.131 | 12.730 | 0.087 | 0.977 | 0.975 |
| 4281 | **Heatr1** | Unchanged | G3X9B1 | 1 | 1 | 1 | 2143 | 241.9 | 6.98 | 1 | 0.688 | 0.343 | 1.743 | 0.332 | 0.496 | 0.136 | 3.511 | 0.116 |
| 4282 | **Uvssa** | Unchanged | Q9D479 | 1 | 1 | 1 | 717 | 81.7 | 6.61 | 1 | #DIV/0! | #DIV/0! | #DIV/0! | #DIV/0! | #DIV/0! | #DIV/0! | #DIV/0! | #DIV/0! |
| 4283 | **Tsc22d1** | Unchanged | P62500 | 1 | 14 | 1 | 1077 | 109.7 | 5.83 | 1 | 0.996 | 0.995 | 1.147 | 0.792 | 0.572 | 0.379 | 2.004 | 0.200 |
| 4284 | **Mcm7** | Down | Q61881 | 1 | 1 | 1 | 719 | 81.2 | 6.37 | 1 | 1.433 | 0.728 | 0.071 | 0.056 | 0.010 | 0.043 | 7.127 | 0.273 |
| 4285 | **Zswim2** | Unchanged | Q9D9X6 | 1 | 6 | 1 | 631 | 71.7 | 9.03 | 1 | 1.852 | 0.397 | 1.487 | 0.540 | 1.897 | 0.377 | 0.784 | 0.708 |
| 4286 | **Plekha6** | Unchanged | Q7TQG1 | 1 | 1 | 1 | 1173 | 131.3 | 8.97 | 1 | 0.015 | 0.055 | 0.229 | 0.145 | 0.174 | 0.111 | 1.318 | 0.821 |
| 4287 | **Capn7** | Down | Q9R1S8 | 1 | 1 | 1 | 813 | 92.5 | 7.97 | 1 | 0.507 | 0.027 | 1.879 | 0.366 | 0.484 | 0.058 | 3.879 | 0.167 |
| 4288 | **Cd177** | Unchanged | Q8R2S8 | 1 | 1 | 1 | 817 | 87 | 5.67 | 1 | 1.392 | 0.655 | 1.152 | 0.886 | 1.449 | 0.595 | 0.795 | 0.805 |
| 4289 | **Slc34a1** | UP | Q60825 | 1 | 1 | 1 | 637 | 68.7 | 8.69 | 1 | 0.227 | 0.007 | 49.630 | 0.020 | 0.166 | 0.004 | 299.308 | 0.019 |
| 4290 | **Nt5dc3** | Unchanged | Q3UHB1 | 1 | 3 | 1 | 546 | 63.1 | 8.56 | 1 | 0.729 | 0.744 | 0.189 | 0.272 | 0.010 | 0.173 | 19.206 | 0.356 |
| 4291 | **Ddx51** | Unchanged | Q6P9R1 | 1 | 1 | 1 | 639 | 70.3 | 9.47 | 1 | 1.255 | 0.743 | 0.758 | 0.689 | 1.220 | 0.768 | 0.621 | 0.454 |
| 4292 | **Ptk2b** | Unchanged | Q9QVP9 | 1 | 1 | 1 | 1009 | 115.7 | 6.35 | 1 | #DIV/0! | #DIV/0! | #DIV/0! | #DIV/0! | #DIV/0! | #DIV/0! | #DIV/0! | #DIV/0! |
| 4293 | **Mus81** | Down | Q91ZJ0 | 1 | 1 | 1 | 551 | 61.5 | 9.41 | 1 | 0.946 | 0.802 | 0.337 | 0.053 | 0.026 | 0.000 | 13.226 | 0.228 |
| 4294 | **Cdc42bpa** | Unchanged | A0A1D5RLQ9 | 1 | 1 | 1 | 1748 | 198.5 | 6.51 | 1 | 1.366 | 0.687 | 0.531 | 0.542 | 2.122 | 0.322 | 0.250 | 0.126 |

| | | | | | | | | | | | | | | | | | | |
|---|---|---|---|---|---|---|---|---|---|---|---|---|---|---|---|---|---|---|
| 4295 | **Tmtc3** | Unchanged | G5E8C4 | 1 | 1 | 1 | 920 | 104.1 | 8.53 | 1 | 1.248 | 0.548 | 2.546 | 0.060 | 1.743 | 0.075 | 1.461 | 0.293 |
| 4296 | **Dysf** | Unchanged | A0A0N4SV63 | 1 | 1 | 1 | 2120 | 241 | 5.67 | 1 | #DIV/0! | #DIV/0! | #DIV/0! | #DIV/0! | #DIV/0! | #DIV/0! | #DIV/0! | #DIV/0! |
| 4297 | **Dip2b** | Unchanged | Q3UH60 | 1 | 1 | 1 | 1574 | 171 | 8.09 | 1 | 0.798 | 0.536 | 1.070 | 0.833 | 0.779 | 0.504 | 1.373 | 0.364 |
| 4298 | **Cask** | UP | O70589 | 1 | 1 | 1 | 926 | 105 | 6.43 | 1 | 2.692 | 0.265 | 3.663 | 0.212 | 4.977 | 0.004 | 0.736 | 0.559 |
| 4299 | **Cntnap5a** | Unchanged | Q0V8T9 | 1 | 1 | 1 | 1304 | 145.6 | 6.28 | 1 | 2.286 | 0.424 | 5.896 | 0.225 | 1.159 | 0.882 | 5.086 | 0.247 |
| 4300 | **Lnpep** | Unchanged | Q8C129 | 1 | 1 | 1 | 1025 | 117.2 | 5.96 | 1 | #DIV/0! | #DIV/0! | #DIV/0! | #DIV/0! | #DIV/0! | #DIV/0! | #DIV/0! | #DIV/0! |
| 4301 | **Kif20b** | Unchanged | Q80WE4 | 2 | 3 | 2 | 1774 | 203.4 | 5.78 | 1 | 1.620 | 0.321 | 0.864 | 0.839 | 1.693 | 0.449 | 0.510 | 0.446 |
| 4302 | **Prpf6** | Unchanged | Q91YR7 | 1 | 2 | 1 | 941 | 106.7 | 8.09 | 1 | 0.933 | 0.834 | 1.202 | 0.747 | 0.637 | 0.135 | 1.886 | 0.382 |
| 4303 | **Pde4d** | Unchanged | F6QFD1 | 1 | 6 | 1 | 754 | 85.5 | 5.25 | 1 | 1.354 | 0.633 | 0.809 | 0.742 | 0.767 | 0.674 | 1.054 | 0.935 |
| 4304 | **Plxna2** | Unchanged | P70207 | 1 | 2 | 1 | 1894 | 211.4 | 6.54 | 1 | 3.037 | 0.331 | 15.529 | 0.120 | 4.824 | 0.053 | 3.219 | 0.243 |
| 4305 | **Nom1** | Unchanged | Q3UFM5 | 1 | 2 | 1 | 854 | 95.9 | 8.87 | 1 | 2.983 | 0.310 | 1.588 | 0.634 | 0.446 | 0.503 | 3.557 | 0.265 |
| 4306 | **Cntnap2** | UP | E9QNF7 | 1 | 1 | 1 | 1332 | 148.2 | 6.76 | 1 | 1.506 | 0.571 | 31.353 | 0.000 | 1.290 | 0.385 | 24.305 | 0.001 |
| 4307 | **Rptor** | Unchanged | A2ACM0 | 1 | 1 | 1 | 1335 | 149.4 | 6.87 | 1 | 0.210 | 0.290 | 0.274 | 0.330 | 0.154 | 0.257 | 1.779 | 0.371 |
| 4308 | **Rnf214** | Unchanged | Q8BFU3 | 1 | 2 | 1 | 668 | 73.6 | 6.23 | 1 | 5.858 | 0.235 | 5.379 | 0.283 | 5.464 | 0.154 | 0.984 | 0.986 |
| 4309 | **Troap** | Unchanged | B7ZNG4 | 1 | 5 | 1 | 668 | 71.9 | 8.7 | 1 | 1.004 | 0.995 | 0.396 | 0.254 | 0.473 | 0.308 | 0.836 | 0.774 |
| 4310 | **Sec16b** | Unchanged | Q91XT4 | 1 | 2 | 1 | 1051 | 115.4 | 6.05 | 1 | #DIV/0! | #DIV/0! | #DIV/0! | #DIV/0! | #DIV/0! | #DIV/0! | #DIV/0! | #DIV/0! |
| 4311 | **Pdzd8** | Unchanged | B9EJ80 | 1 | 1 | 1 | 1147 | 127.7 | 6.04 | 1 | 0.950 | 0.809 | 1.030 | 0.930 | 0.914 | 0.750 | 1.127 | 0.729 |
| 4312 | **Skiv2l** | Unchanged | Q6NZR5 | 1 | 1 | 1 | 1244 | 137.4 | 6.28 | 1 | 0.871 | 0.534 | 1.086 | 0.740 | 2.032 | 0.179 | 0.534 | 0.237 |
| 4313 | **Mgam** | Unchanged | B5THE2 | 1 | 1 | 1 | 1827 | 208.4 | 5.95 | 1 | #DIV/0! | #DIV/0! | #DIV/0! | #DIV/0! | #DIV/0! | #DIV/0! | #DIV/0! | #DIV/0! |
| 4314 | **Ints4** | Unchanged | Q8CIM8 | 1 | 2 | 1 | 964 | 108.1 | 6.42 | 1 | 0.834 | 0.569 | 1.088 | 0.804 | 0.919 | 0.692 | 1.184 | 0.611 |
| 4315 | **Trip12** | Unchanged | G5E870 | 1 | 3 | 1 | 2025 | 224 | 8.35 | 1 | 1.420 | 0.339 | 1.419 | 0.293 | 0.798 | 0.559 | 1.777 | 0.082 |
| 4316 | **Wdr81** | Unchanged | Q5ND34 | 1 | 4 | 1 | 1934 | 211.8 | 5.77 | 1 | 0.907 | 0.879 | 0.853 | 0.771 | 1.531 | 0.417 | 0.557 | 0.243 |
| 4317 | **Lrit3** | Unchanged | W8DXL4 | 1 | 1 | 1 | 681 | 74.8 | 6.74 | | 2.810 | 0.330 | 15.076 | 0.079 | 1.076 | 0.896 | 14.013 | 0.081 |
| 4318 | **Usp34** | UP | F6WJB7 | 1 | 1 | 1 | 3602 | 409.7 | 5.69 | 1 | 4.485 | 0.119 | 12.848 | 0.115 | 3.127 | 0.032 | 4.109 | 0.187 |
| 4319 | **Mast4** | Unchanged | E9QPR4 | 1 | 1 | 1 | 2441 | 265.9 | 8.65 | 1 | #DIV/0! | #DIV/0! | #DIV/0! | #DIV/0! | #DIV/0! | #DIV/0! | #DIV/0! | #DIV/0! |
| 4320 | **Kif21a** | Unchanged | E9Q0J5 | 1 | 1 | 1 | 1672 | 186.5 | 6.47 | 1 | 0.187 | 0.082 | 2.471 | 0.303 | 1.232 | 0.784 | 2.005 | 0.420 |
| 4321 | **Capn9** | Unchanged | Q9D805 | 1 | 3 | 1 | 690 | 78.9 | 5.36 | 1 | 16.004 | 0.345 | 5.108 | 0.214 | 3.077 | 0.351 | 1.660 | 0.596 |
| 4322 | **Calcoco1** | Unchanged | Q8CGU1 | 1 | 1 | 1 | 691 | 77.2 | 4.82 | 1 | #DIV/0! | #DIV/0! | #DIV/0! | #DIV/0! | #DIV/0! | #DIV/0! | #DIV/0! | #DIV/0! |
| 4323 | **Ppfia4** | Unchanged | B8QI36 | 1 | 9 | 1 | 1187 | 133.6 | 6.27 | 1 | 1.211 | 0.770 | 1.836 | 0.422 | 1.794 | 0.386 | 1.023 | 0.972 |
| 4324 | **Epg5** | Unchanged | Q80TA9 | 1 | 1 | 1 | 2572 | 290.6 | 6.37 | 1 | 1.077 | 0.935 | 0.695 | 0.753 | 2.856 | 0.239 | 0.243 | 0.131 |
| 4325 | **Ahdc1** | Down | Q6PAL7 | 1 | 1 | 1 | 1594 | 168 | 8.97 | 1 | 0.886 | 0.770 | 1.397 | 0.376 | 0.441 | 0.020 | 3.168 | 0.044 |
| 4326 | **Chd1l** | Unchanged | Q9CXF7 | 1 | 1 | 1 | 900 | 101.4 | 6.64 | | 0.981 | 0.952 | 6.881 | 0.058 | 1.170 | 0.633 | 5.883 | 0.065 |
| 4327 | **E2f7** | Unchanged | Q6S7F2 | 1 | 1 | 1 | 904 | 99.5 | 6.77 | 1 | 0.195 | 0.181 | 1.302 | 0.803 | 2.049 | 0.201 | 0.635 | 0.539 |
| 4328 | **Kiaa1211** | Unchanged | Q5PR69 | 1 | 1 | 1 | 1207 | 132.2 | 5.4 | 1 | 0.671 | 0.502 | 1.714 | 0.616 | 3.041 | 0.266 | 0.564 | 0.550 |
| 4329 | **Dicer1** | Unchanged | Q8R418 | 1 | 1 | 1 | 1916 | 216.7 | 5.82 | 1 | 0.779 | 0.748 | 2.437 | 0.502 | 2.281 | 0.326 | 1.068 | 0.947 |
| 4330 | **Scmh1** | Unchanged | Q8K214 | 1 | 1 | 1 | 706 | 78.6 | 9.31 | 1 | 0.029 | 0.082 | 0.020 | 0.079 | 0.176 | 0.135 | 0.114 | 0.139 |
| 4331 | **Uaca** | Unchanged | A0A0R4J0S7 | 1 | 1 | 1 | 1413 | 160.8 | 7.2 | 1 | #DIV/0! | #DIV/0! | #DIV/0! | #DIV/0! | #DIV/0! | #DIV/0! | #DIV/0! | #DIV/0! |
| 4332 | **Lrrc4b** | Unchanged | P0C192 | 1 | 1 | 1 | 709 | 76.1 | 7.24 | 1 | 0.733 | 0.728 | 0.571 | 0.451 | 0.438 | 0.306 | 1.303 | 0.683 |
| 4333 | **Cnnm3** | Unchanged | Q32NY4 | 1 | 5 | 1 | 713 | 76.2 | 5.58 | 1 | #DIV/0! | #DIV/0! | #DIV/0! | #DIV/0! | #DIV/0! | #DIV/0! | #DIV/0! | #DIV/0! |
| 4334 | **Top2a** | Unchanged | Q01320 | 1 | 1 | 1 | 1528 | 172.7 | 8.6 | 1 | 1.363 | 0.295 | 0.989 | 0.976 | 1.218 | 0.645 | 0.812 | 0.683 |
| 4335 | **Dnah11** | Unchanged | E9Q7N9 | 2 | 2 | 2 | 4488 | 516 | 6.14 | 1 | 0.984 | 0.981 | 0.215 | 0.096 | 0.148 | 0.074 | 1.452 | 0.286 |

| | | | | | | | | | | | | | | | | | | |
|---|---|---|---|---|---|---|---|---|---|---|---|---|---|---|---|---|---|---|
| 4336 | **Tbc1d2** | Unchanged | B1AVH7 | 1 | 2 | 1 | 922 | 104.2 | 6.43 | 1 | 0.156 | 0.132 | 0.060 | 0.095 | 2.775 | 0.449 | 0.022 | 0.247 |
| 4337 | **Exoc2** | Unchanged | Q9D4H1 | 1 | 2 | 1 | 924 | 103.9 | 7.18 | 1 | 0.821 | 0.669 | 1.049 | 0.921 | 0.857 | 0.768 | 1.225 | 0.685 |
| 4338 | **Sarm1** | Unchanged | Q6PDS3 | 1 | 2 | 1 | 724 | 79.6 | 6.27 | 1 | 2.243 | 0.102 | 1.474 | 0.659 | 2.807 | 0.107 | 0.525 | 0.367 |
| 4339 | **Uimc1** | Unchanged | A0A0R4J073 | 1 | 2 | 1 | 727 | 81.4 | 6.51 | 1 | 0.576 | 0.523 | 0.304 | 0.302 | 0.004 | 0.143 | 75.093 | 0.074 |
| 4340 | **Arhgap28** | Unchanged | Q8BN58 | 1 | 3 | 1 | 729 | 81.7 | 7.78 | | 1.327 | 0.581 | 1.745 | 0.176 | 32.092 | 0.151 | 0.054 | 0.160 |
| 4341 | **Ttc3** | Unchanged | G5E8T2 | 2 | 2 | 2 | 1979 | 223.8 | 6.25 | 1 | 2.971 | 0.271 | 2.328 | 0.158 | 1.186 | 0.732 | 1.963 | 0.229 |
| 4342 | **Ankrd13b** | Unchanged | Q5F259 | 1 | 3 | 1 | 626 | 70.3 | 7.21 | 1 | 0.400 | 0.274 | 0.229 | 0.168 | 0.043 | 0.090 | 5.373 | 0.130 |
| 4343 | **Itgav** | Unchanged | P43406 | 1 | 2 | 1 | 1044 | 115.3 | 5.63 | 1 | 0.735 | 0.526 | 1.303 | 0.630 | 0.413 | 0.186 | 3.155 | 0.096 |
| 4344 | **Pik3r4** | Unchanged | Q8VD65 | 1 | 1 | 1 | 1358 | 152.5 | 7.12 | 1 | 1.565 | 0.534 | 1.256 | 0.576 | 3.071 | 0.124 | 0.409 | 0.174 |
| 4345 | **Smg1** | Down | Q8BKX6 | 3 | 3 | 3 | 3658 | 409.5 | 6.4 | 1 | 1.404 | 0.596 | 0.897 | 0.735 | 0.228 | 0.004 | 3.930 | 0.030 |
| 4346 | **Bcorl1** | Unchanged | A2AQH4 | 1 | 2 | 1 | 1781 | 190.4 | 7.78 | 1 | 0.435 | 0.259 | 0.251 | 0.115 | 0.492 | 0.282 | 0.510 | 0.346 |
| 4347 | **Parp6** | Down | G5E856 | 1 | 2 | 1 | 630 | 71.1 | 8.29 | 1 | 1.311 | 0.528 | 1.411 | 0.600 | 0.223 | 0.002 | 6.331 | 0.150 |
| 4348 | **Frem3** | Unchanged | F8WHZ0 | 1 | 2 | 1 | 2127 | 235.3 | 5.16 | 1 | 1.264 | 0.727 | 0.420 | 0.266 | 0.475 | 0.402 | 0.885 | 0.915 |
| 4349 | **Thsd1** | Unchanged | Q9JM61 | 1 | 4 | 1 | 851 | 94.6 | 8.19 | | 1.097 | 0.784 | 1.197 | 0.817 | 1.133 | 0.582 | 1.057 | 0.941 |
| 4350 | **Tut7** | Unchanged | Q5BLK4 | 1 | 1 | 1 | 1491 | 169 | 6.65 | 1 | 0.821 | 0.512 | 0.624 | 0.248 | 1.094 | 0.734 | 0.571 | 0.180 |
| 4351 | **Ext1** | Unchanged | P97464 | 1 | 7 | 1 | 746 | 86.3 | 9.04 | 1 | 0.895 | 0.800 | 0.503 | 0.289 | 0.979 | 0.974 | 0.514 | 0.428 |
| 4352 | **Hecw1** | Unchanged | Q8K4P8 | 1 | 1 | 1 | 1604 | 179.4 | 5.39 | 1 | 0.717 | 0.664 | 0.528 | 0.431 | 0.117 | 0.061 | 4.500 | 0.336 |
| 4353 | **Ankle2** | Unchanged | Q6P1H6 | 1 | 1 | 1 | 964 | 106.1 | 7.37 | 1 | #DIV/0! | #DIV/0! | #DIV/0! | #DIV/0! | #DIV/0! | #DIV/0! | #DIV/0! | #DIV/0! |
| 4354 | **Mphosph8** | Unchanged | Q3TYA6 | 1 | 2 | 1 | 858 | 97.4 | 5.85 | | 0.604 | 0.604 | 0.222 | 0.265 | 0.913 | 0.902 | 0.243 | 0.017 |
| 4355 | **Lrrc49** | Unchanged | E9QM16 | 1 | 8 | 1 | 752 | 85.6 | 8.34 | 1 | 2.901 | 0.223 | 1.411 | 0.743 | 2.324 | 0.286 | 0.607 | 0.547 |
| 4356 | **Golga1** | Unchanged | Q9CW79 | 1 | 6 | 1 | 758 | 87.3 | 5.52 | 1 | 0.816 | 0.755 | 0.721 | 0.618 | 0.120 | 0.093 | 6.023 | 0.067 |
| 4357 | **Cep152** | Unchanged | A2AUM9 | 2 | 12 | 2 | 1736 | 196.4 | 5.66 | 1 | 1.347 | 0.533 | 0.752 | 0.507 | 1.000 | 1.000 | 0.752 | 0.527 |
| 4358 | **Gpr155** | Unchanged | A2AWR3 | 1 | 12 | 1 | 868 | 96.3 | 6.87 | 1 | 1.089 | 0.890 | 1.103 | 0.881 | 1.033 | 0.958 | 1.067 | 0.868 |
| 4359 | **Ercc2** | Unchanged | O08811 | 1 | 6 | 1 | 760 | 86.8 | 6.92 | 1 | 0.777 | 0.707 | 0.012 | 0.060 | 0.745 | 0.660 | 0.016 | 0.057 |
| 4360 | **Phtf1** | Unchanged | Q9QZ09 | 1 | 1 | 1 | 761 | 86.7 | 9.63 | 1 | 0.769 | 0.406 | 23.181 | 0.077 | 10.830 | 0.087 | 2.140 | 0.335 |
| 4361 | **Cfap58** | Down | B2RW38 | 1 | 1 | 1 | 873 | 103.5 | 8.16 | 1 | 0.414 | 0.130 | 0.118 | 0.031 | 0.040 | 0.021 | 2.963 | 0.108 |
| 4362 | **Igfn1** | Unchanged | Q3KNY0 | 1 | 1 | 1 | 2849 | 303.5 | 6.58 | 1 | 0.481 | 0.350 | 0.635 | 0.521 | 1.003 | 0.995 | 0.633 | 0.288 |
| 4363 | **Gcfc2** | Unchanged | Q8BKT3 | 1 | 1 | 1 | 769 | 87.4 | 5.55 | 1 | 0.249 | 0.127 | 0.233 | 0.123 | 0.277 | 0.140 | 0.841 | 0.549 |
| 4364 | **Chl1** | Unchanged | P70232 | 1 | 1 | 1 | 1209 | 135 | 5.57 | 1 | 0.819 | 0.777 | 1.335 | 0.639 | 1.168 | 0.890 | 1.143 | 0.895 |
| 4365 | **Clasp1** | Unchanged | A0A087WQ31 | 1 | 1 | 1 | 1552 | 170.9 | 9.03 | 1 | #DIV/0! | #DIV/0! | #DIV/0! | #DIV/0! | #DIV/0! | #DIV/0! | #DIV/0! | #DIV/0! |
| 4366 | **Ccdc150** | Unchanged | Q8CDI7 | 1 | 1 | 1 | 1110 | 128.4 | 6.93 | 1 | 3.409 | 0.443 | 0.881 | 0.849 | 1.248 | 0.732 | 0.706 | 0.681 |
| 4367 | **Arap1** | Down | Q4LDD4 | 1 | 2 | 1 | 1452 | 162.2 | 6.18 | 1 | 1.092 | 0.901 | 0.220 | 0.046 | 0.177 | 0.031 | 1.240 | 0.733 |
| 4368 | **Ubr5** | Unchanged | A0A2I3BQS6 | 1 | 1 | 1 | 2798 | 308.8 | 5.83 | 1 | #DIV/0! | #DIV/0! | #DIV/0! | #DIV/0! | #DIV/0! | #DIV/0! | #DIV/0! | #DIV/0! |
| 4369 | **Dock7** | Unchanged | Q8R1A4 | 1 | 2 | 1 | 2130 | 241.3 | 6.71 | 1 | 1.124 | 0.830 | 1.374 | 0.360 | 1.376 | 0.507 | 0.998 | 0.997 |
| 4370 | **Cfap43** | Unchanged | E9Q7R9 | 1 | 2 | 1 | 1682 | 193.3 | 5.62 | 1 | 1.572 | 0.491 | 1.753 | 0.387 | 1.807 | 0.392 | 0.970 | 0.961 |
| 4371 | **Gon4l** | Unchanged | K4DI71 | 1 | 1 | 1 | 2243 | 246.7 | 4.86 | 1 | 0.754 | 0.749 | 0.540 | 0.476 | 0.731 | 0.690 | 0.738 | 0.551 |
| 4372 | **Pik3c2a** | Unchanged | F8VPL2 | 1 | 1 | 1 | 1686 | 190.6 | 8.02 | 1 | #DIV/0! | #DIV/0! | #DIV/0! | #DIV/0! | #DIV/0! | #DIV/0! | #DIV/0! | #DIV/0! |
| 4373 | **Srebf2** | Unchanged | Q3U1N2 | 1 | 4 | 1 | 1130 | 122.8 | 8.44 | 1 | 2.431 | 0.274 | 1.184 | 0.780 | 2.131 | 0.320 | 0.556 | 0.412 |
| 4374 | **Swt1** | Unchanged | Q9DBQ9 | 1 | 6 | 1 | 907 | 103.1 | 8.22 | 1 | 1.802 | 0.285 | 2.063 | 0.324 | 1.385 | 0.631 | 1.490 | 0.549 |
| 4375 | **Cip2a** | Unchanged | D3Z7B5 | 1 | 1 | 1 | 907 | 102.1 | 6.39 | 1 | #DIV/0! | #DIV/0! | #DIV/0! | #DIV/0! | #DIV/0! | #DIV/0! | #DIV/0! | #DIV/0! |
| 4376 | **Tbc1d10b** | Unchanged | Q8BHL3 | 1 | 8 | 1 | 798 | 87.2 | 8.9 | 1 | 1.327 | 0.581 | 3.242 | 0.275 | 2.173 | 0.385 | 1.492 | 0.643 |

| | | | | | | | | | | | | | | | | | | |
|---|---|---|---|---|---|---|---|---|---|---|---|---|---|---|---|---|---|---|
| 4377 | **Taf5** | Unchanged | Q8C092 | 1 | 22 | 1 | 801 | 87 | 5.64 | 1 | 0.243 | 0.344 | 0.388 | 0.449 | 0.251 | 0.347 | 1.550 | 0.533 |
| 4378 | **Rasa4** | Unchanged | Q6PFQ7 | 1 | 13 | 1 | 802 | 90 | 8 | 1 | 0.935 | 0.925 | 1.516 | 0.565 | 0.905 | 0.866 | 1.675 | 0.488 |
| 4379 | **Marf1** | Unchanged | Q8BJ34 | 2 | 19 | 2 | 1730 | 191.9 | 8.22 | 1 | 0.866 | 0.878 | 0.648 | 0.680 | 0.015 | 0.163 | 43.778 | 0.257 |
| 4380 | **Lrrc8a** | UP | Q80WG5 | 1 | 1 | 1 | 810 | 94.1 | 7.94 | 1 | 2.472 | 0.229 | 4.273 | 0.406 | 11.108 | 0.001 | 0.385 | 0.141 |
| 4381 | **Ddx42** | Unchanged | Q810A7 | 1 | 1 | 1 | 929 | 101.9 | 6.98 | 1 | 1.068 | 0.857 | 1.421 | 0.240 | 0.591 | 0.190 | 2.403 | 0.016 |
| 4382 | **Galnt5** | Unchanged | Q8C102 | 1 | 1 | 1 | 930 | 105.7 | 9.09 | 1 | 1.411 | 0.549 | 0.966 | 0.952 | 0.715 | 0.494 | 1.351 | 0.688 |
| 4383 | **Tax1bp1** | Unchanged | Q3UKC1 | 1 | 2 | 1 | 814 | 93.6 | 5.33 | 1 | 0.158 | 0.200 | 0.200 | 0.230 | 0.275 | 0.266 | 0.727 | 0.672 |
| 4384 | **Ddhd2** | Unchanged | Q80Y98 | 1 | 1 | 1 | 699 | 79.5 | 5.31 | 1 | 2.521 | 0.175 | 3.488 | 0.371 | 0.682 | 0.626 | 5.113 | 0.324 |
| 4385 | **Gtf3c4** | Unchanged | Q8BMQ2 | 1 | 2 | 1 | 817 | 91.6 | 6.73 | | #DIV/0! | #DIV/0! | #DIV/0! | #DIV/0! | #DIV/0! | #DIV/0! | #DIV/0! | #DIV/0! |
| 4386 | **Smarca5** | Unchanged | Q91ZW3 | 1 | 2 | 1 | 1051 | 121.6 | 8.15 | 1 | 1.474 | 0.304 | 1.302 | 0.588 | 1.342 | 0.392 | 0.970 | 0.942 |
| 4387 | **Ttll6** | Unchanged | A4Q9E8 | 1 | 2 | 1 | 822 | 94.4 | 9.17 | 1 | 0.814 | 0.788 | 1.038 | 0.958 | 0.412 | 0.290 | 2.521 | 0.240 |
| 4388 | **Slc12a8** | Unchanged | Q8VI23 | 1 | 9 | 1 | 705 | 77 | 6.21 | 1 | 1.708 | 0.455 | 1.690 | 0.448 | 1.905 | 0.345 | 0.887 | 0.851 |
| 4389 | **Elfn1** | Unchanged | Q8C8T7 | 1 | 1 | 1 | 828 | 90.8 | 8.18 | 1 | 1.100 | 0.896 | 1.594 | 0.598 | 0.190 | 0.122 | 8.374 | 0.190 |
| 4390 | **Arhgef12** | Unchanged | F8VQN6 | 1 | 1 | 1 | 1544 | 172.4 | 5.74 | 1 | 0.940 | 0.935 | 0.550 | 0.558 | 1.079 | 0.918 | 0.509 | 0.409 |
| 4391 | **Cgn** | Unchanged | P59242 | 1 | 9 | 1 | 1191 | 136.4 | 5.91 | 1 | 1.467 | 0.613 | 1.283 | 0.725 | 0.648 | 0.402 | 1.979 | 0.392 |
| 4392 | **Sec14l1** | Unchanged | A8Y5H7 | 1 | 3 | 1 | 715 | 81.2 | 6.34 | 1 | 1.664 | 0.486 | 3.578 | 0.134 | 1.922 | 0.356 | 1.862 | 0.364 |
| 4393 | **Kidins220** | Unchanged | E9Q9B7 | 1 | 1 | 1 | 1793 | 199.1 | 6.67 | 1 | #DIV/0! | #DIV/0! | #DIV/0! | #DIV/0! | #DIV/0! | #DIV/0! | #DIV/0! | #DIV/0! |
| 4394 | **Cmtr1** | Unchanged | Q9DBC3 | 1 | 3 | 1 | 837 | 95.6 | 7.27 | 1 | 0.701 | 0.087 | 0.670 | 0.201 | 1.954 | 0.208 | 0.343 | 0.112 |
| 4395 | **Heatr5a** | Unchanged | Q5PRF0 | 1 | 1 | 1 | 2038 | 219.8 | 6.76 | 1 | 1.493 | 0.385 | 2.263 | 0.415 | 0.978 | 0.954 | 2.313 | 0.397 |
| 4396 | **Nlrc3** | Unchanged | A0A2R8VHN0 | 1 | 3 | 1 | 1199 | 128.7 | 7.5 | 1 | 0.378 | 0.217 | 0.194 | 0.098 | 0.340 | 0.195 | 0.570 | 0.477 |
| 4397 | **Mast2** | Unchanged | E9Q1Q1 | 1 | 1 | 1 | 1800 | 197.9 | 7.91 | 1 | 0.974 | 0.981 | 5.224 | 0.357 | 1.765 | 0.386 | 2.960 | 0.449 |
| 4398 | **Xpo5** | Unchanged | Q924C1 | 1 | 1 | 1 | 1204 | 136.9 | 5.82 | 1 | #DIV/0! | #DIV/0! | #DIV/0! | #DIV/0! | #DIV/0! | #DIV/0! | #DIV/0! | #DIV/0! |
| 4399 | **Parp14** | Unchanged | Q2EMV9 | 1 | 3 | 1 | 1817 | 203.7 | 6.79 | 1 | #DIV/0! | #DIV/0! | #DIV/0! | #DIV/0! | #DIV/0! | #DIV/0! | #DIV/0! | #DIV/0! |
| 4400 | **Osbpl3** | Unchanged | Q9DBS9 | 1 | 2 | 1 | 855 | 96.9 | 6.51 | 1 | 0.766 | 0.769 | 0.611 | 0.611 | 0.481 | 0.510 | 1.271 | 0.733 |
| 4401 | **Szt2** | Unchanged | A2A9C3 | 1 | 2 | 1 | 3431 | 377.4 | 6.19 | 1 | 1.835 | 0.071 | 1.350 | 0.585 | 1.638 | 0.239 | 0.824 | 0.695 |
| 4402 | **Ano8** | Unchanged | A0A1L1SR96 | 1 | 2 | 1 | 1104 | 123.9 | 6.01 | 1 | 0.961 | 0.937 | 0.933 | 0.936 | 1.296 | 0.673 | 0.720 | 0.691 |
| 4403 | **Zmym1** | Down | Q3TJB1 | 1 | 2 | 1 | 983 | 110.3 | 6.42 | 1 | 0.322 | 0.030 | 1.945 | 0.557 | 1.617 | 0.473 | 1.203 | 0.853 |
| 4404 | **Mcf2** | Unchanged | A0A2I3BR92 | 1 | 1 | 1 | 1111 | 127.1 | 7.14 | 1 | 1.223 | 0.767 | 1.775 | 0.431 | 0.985 | 0.982 | 1.802 | 0.415 |
| 4405 | **Daam2** | Unchanged | Q80U19 | 1 | 2 | 1 | 1115 | 128.3 | 6.92 | 1 | 0.043 | 0.071 | 0.074 | 0.079 | 0.047 | 0.071 | 1.585 | 0.498 |
| 4406 | **Smarca4** | Unchanged | A0A0R4J170 | 1 | 1 | 1 | 1617 | 181.7 | 8.07 | 1 | 0.960 | 0.935 | 1.347 | 0.723 | 0.535 | 0.381 | 2.517 | 0.418 |
| 4407 | **Tert** | Unchanged | O70372 | 1 | 1 | 1 | 1122 | 127.9 | 9.91 | 1 | 0.004 | 0.060 | 0.004 | 0.060 | 0.052 | 0.071 | 0.082 | 0.147 |
| 4408 | **Arhgap35** | Unchanged | Q91YM2 | 1 | 1 | 1 | 1499 | 170.3 | 6.61 | 1 | 1.368 | 0.573 | 0.715 | 0.629 | 0.125 | 0.103 | 5.698 | 0.101 |
| 4409 | **Asap2** | Unchanged | E9PX52 | 1 | 5 | 1 | 1000 | 111.1 | 6.8 | 1 | 4.716 | 0.217 | 0.782 | 0.746 | 0.211 | 0.256 | 3.698 | 0.029 |
| 4410 | **Ppp4r4** | Unchanged | Q8C0Y0 | 1 | 18 | 1 | 875 | 99.4 | 7.71 | 1 | 0.437 | 0.237 | 0.486 | 0.290 | 0.479 | 0.264 | 1.015 | 0.980 |
| 4411 | **Nlrp9a** | Down | D3Z7F5 | 1 | 1 | 1 | 1004 | 115.9 | 6.61 | 1 | 0.635 | 0.481 | 0.560 | 0.315 | 0.243 | 0.030 | 2.302 | 0.337 |
| 4412 | **Fam135a** | Unchanged | Q6NS59 | 1 | 1 | 1 | 1506 | 166.8 | 5.47 | 1 | #DIV/0! | #DIV/0! | #DIV/0! | #DIV/0! | #DIV/0! | #DIV/0! | #DIV/0! | #DIV/0! |
| 4413 | **Incenp** | Unchanged | Q9WU62 | 1 | 1 | 1 | 880 | 101.1 | 9.55 | 1 | #DIV/0! | #DIV/0! | #DIV/0! | #DIV/0! | #DIV/0! | #DIV/0! | #DIV/0! | #DIV/0! |
| 4414 | **Lrrk2** | Unchanged | Q5S006 | 2 | 2 | 2 | 2527 | 284.5 | 6.83 | 1 | 1.189 | 0.641 | 1.107 | 0.798 | 0.597 | 0.303 | 1.856 | 0.238 |
| 4415 | **Rpgrip1l** | Down | Q8CG73 | 1 | 1 | 1 | 1264 | 144.9 | 5.43 | 1 | 0.462 | 0.155 | 0.009 | 0.004 | 0.011 | 0.004 | 0.778 | 0.638 |
| 4416 | **Axl** | Unchanged | Q00993 | 1 | 3 | 1 | 888 | 98.1 | 5.74 | 1 | 0.573 | 0.407 | 0.792 | 0.720 | 0.618 | 0.421 | 1.283 | 0.660 |
| 4417 | **Mroh9** | Down | G5E8L9 | 1 | 1 | 1 | 891 | 101.6 | 7.44 | 1 | 0.664 | 0.243 | 0.756 | 0.536 | 0.289 | 0.022 | 2.615 | 0.160 |

| | | | | | | | | | | | | | | | | | | |
|---|---|---|---|---|---|---|---|---|---|---|---|---|---|---|---|---|---|---|
| 4418 | **Atp9b** | Uncha nged | D3YV00 | 1 | 2 | 1 | 114 6 | 129 | 7.7 1 | 1 | 0.920 | 0.88 2 | 1.189 | 0.75 1 | 0.318 | 0.08 5 | 3.741 | 0.107 |
| 4419 | **Ston2** | Uncha nged | E9PXP7 | 1 | 1 | 1 | 898 | 99. 8 | 5.4 8 | 1 | 0.889 | 0.89 0 | 0.156 | 0.22 7 | 1.351 | 0.70 5 | 0.115 | 0.091 |
| 4420 | **Vinac1** | Uncha nged | A2AP89 | 1 | 6 | 1 | 141 3 | 155 .7 | 6.4 8 | 1 | 1.744 | 0.49 5 | 0.101 | 0.16 6 | 0.267 | 0.26 8 | 0.380 | 0.397 |
| 4421 | **Trpm5** | Uncha nged | Q9JJH7 | 1 | 1 | 1 | 115 8 | 130 .8 | 6.4 | 1 | 1.637 | 0.56 8 | 0.045 | 0.29 8 | 0.423 | 0.52 9 | 0.106 | 0.076 |
| 4422 | **Zc3h12c** | Uncha nged | E9Q1I3 | 1 | 1 | 1 | 903 | 101 | 7.4 4 | 1 | #DIV/0! | #DIV /0! | #DIV/0! | #DIV /0! | #DIV/0! | #DIV /0! | #DIV/0 ! | #DIV/0! |
| 4423 | **Nrip1** | Uncha nged | Q8CBD1 | 1 | 1 | 1 | 116 1 | 126 .3 | 7.9 7 | 1 | 0.373 | 0.22 9 | 0.347 | 0.20 8 | 0.429 | 0.27 7 | 0.808 | 0.730 |
| 4424 | **Crybg1** | Uncha nged | A0A0G2 JG52 | 1 | 1 | 1 | 206 5 | 223 .3 | 5.7 2 | 1 | 1.009 | 0.98 2 | 1.948 | 0.34 2 | 0.725 | 0.48 0 | 2.686 | 0.222 |
| 4425 | **Cdh4** | Uncha nged | P39038 | 1 | 2 | 1 | 913 | 100 | 4.8 1 | 1 | 1.602 | 0.46 7 | 0.384 | 0.16 8 | 1.849 | 0.29 3 | 0.207 | 0.054 |
| 4426 | **Adprhl1** | Uncha nged | A0A0N4 SVK8 | 1 | 2 | 1 | 169 7 | 184 .1 | 7.6 6 | 1 | 37.128 | 0.15 3 | 51.819 | 0.25 0 | 44.289 | 0.07 2 | 1.170 | 0.874 |
| 4427 | **Rapgef3** | Down | Q8VCC8 | 1 | 1 | 1 | 918 | 103 .5 | 7.8 1 | 1 | 0.523 | 0.20 1 | 0.433 | 0.11 5 | 0.272 | 0.04 1 | 1.589 | 0.271 |
| 4428 | **Col4a2** | Uncha nged | P08122 | 1 | 4 | 1 | 170 7 | 167 .2 | 8.4 8 | 1 | 2.211 | 0.40 5 | 3.406 | 0.26 1 | 2.294 | 0.26 1 | 1.485 | 0.594 |
| 4429 | **Cfap46** | Uncha nged | E9Q2C0 | 1 | 1 | 1 | 264 1 | 302 .4 | 6.8 4 | 1 | 1.574 | 0.65 7 | 1.576 | 0.74 8 | 0.110 | 0.31 6 | 14.282 | 0.364 |
| 4430 | **Nisch** | Uncha nged | Q80TM9 | 1 | 1 | 1 | 159 3 | 174 .9 | 5.1 6 | 1 | 0.572 | 0.41 2 | 2.499 | 0.41 8 | 0.331 | 0.18 6 | 7.560 | 0.239 |
| 4431 | **Heg1** | Uncha nged | E9Q7X6 | 1 | 1 | 1 | 133 7 | 141 .8 | 6.6 7 | 1 | 0.734 | 0.70 2 | 0.509 | 0.45 9 | 0.089 | 0.16 5 | 5.695 | 0.088 |
| 4432 | **Zmiz1** | Uncha nged | Q6P1E1 | 1 | 1 | 1 | 107 2 | 115 .8 | 7.4 4 | 1 | 1.142 | 0.73 7 | 1.750 | 0.41 8 | 0.622 | 0.39 4 | 2.814 | 0.217 |
| 4433 | **Svil** | Uncha nged | Q8K4L3 | 1 | 5 | 1 | 217 0 | 243 | 6.8 7 | 1 | #DIV/0! | #DIV /0! | #DIV/0! | #DIV /0! | #DIV/0! | #DIV /0! | #DIV/0 ! | #DIV/0! |
| 4434 | **Adamtsl2** | Uncha nged | Q7TSK7 | 1 | 6 | 1 | 957 | 105 .6 | 6.8 3 | 1 | 1.297 | 0.51 2 | 1.399 | 0.54 9 | 0.972 | 0.94 5 | 1.440 | 0.553 |
| 4435 | **Diaph2** | Uncha nged | E9Q4U7 | 1 | 1 | 1 | 110 2 | 125 .3 | 6.7 7 | 1 | #DIV/0! | #DIV /0! | #DIV/0! | #DIV /0! | #DIV/0! | #DIV /0! | #DIV/0 ! | #DIV/0! |
| 4436 | **Pkd2** | Uncha nged | O35245 | 1 | 1 | 1 | 966 | 108 .9 | 5.6 6 | | 1.463 | 0.64 9 | 0.970 | 0.96 7 | 3.225 | 0.17 1 | 0.301 | 0.166 |
| 4437 | **Dnah6** | Uncha nged | E9Q0B6 | 1 | 1 | 1 | 414 4 | 474 .5 | 5.6 3 | 1 | 0.880 | 0.68 2 | 1.322 | 0.46 9 | 0.757 | 0.49 7 | 1.746 | 0.223 |
| 4438 | **Nup98** | Uncha nged | Q6PFD9 | 1 | 2 | 1 | 181 6 | 197 .1 | 6.1 8 | 1 | 1.007 | 0.98 6 | 0.794 | 0.57 2 | 1.548 | 0.50 5 | 0.513 | 0.344 |
| 4439 | **Fbrs** | Uncha nged | A0A1N9 PTV1 | 1 | 1 | 1 | 980 | 103 .9 | 9.6 1 | 1 | 0.122 | 0.13 2 | 0.015 | 0.09 4 | 0.015 | 0.09 4 | 1.047 | 0.913 |
| 4440 | **Sycp2l** | Uncha nged | A0A0M3 U1B0 | 1 | 1 | 1 | 842 | 96 | 6.0 5 | 1 | 0.878 | 0.82 9 | 1.195 | 0.81 8 | 1.128 | 0.83 1 | 1.060 | 0.941 |
| 4441 | **Nr3c2** | Uncha nged | E9Q8M8 | 1 | 2 | 1 | 984 | 107 | 7.8 7 | 1 | 4.018 | 0.13 6 | 4.544 | 0.12 8 | 3.559 | 0.12 2 | 1.277 | 0.705 |
| 4442 | **Plcl2** | Down | Q8K394 | 1 | 1 | 1 | 112 8 | 125 .7 | 6.9 2 | 1 | 0.019 | 0.04 8 | 0.019 | 0.04 8 | 0.015 | 0.04 7 | 1.261 | 0.616 |
| 4443 | **Dlgap4** | Uncha nged | B1AZP2 | 1 | 1 | 1 | 992 | 108 | 7.0 8 | 1 | #DIV/0! | #DIV /0! | #DIV/0! | #DIV /0! | #DIV/0! | #DIV /0! | #DIV/0 ! | #DIV/0! |
| 4444 | **Prdm16** | Uncha nged | A2A933 | 1 | 16 | 1 | 127 6 | 140 .9 | 6.1 | 1 | 1.012 | 0.96 2 | 0.736 | 0.34 4 | 0.985 | 0.93 8 | 0.747 | 0.407 |
| 4445 | **Trrap** | Uncha nged | A0A1D5 RLL4 | 1 | 2 | 1 | 385 9 | 437 .1 | 8.1 8 | 1 | 1.506 | 0.39 9 | 1.075 | 0.89 8 | 1.469 | 0.39 4 | 0.732 | 0.535 |
| 4446 | **Ttf1** | Uncha nged | Q62187 | 1 | 1 | 1 | 859 | 97. 7 | 9.3 8 | 1 | 0.149 | 0.18 9 | 0.131 | 0.17 7 | 0.137 | 0.18 5 | 0.958 | 0.965 |
| 4447 | **Nalcn** | Uncha nged | E9QLE4 | 1 | 1 | 1 | 173 9 | 200 .4 | 8.6 8 | 1 | 0.125 | 0.08 7 | 0.289 | 0.17 8 | 1.661 | 0.34 8 | 0.174 | 0.029 |
| 4448 | **Vcan** | Uncha nged | Q62059 | 1 | 2 | 1 | 335 7 | 366 .6 | 4.6 4 | 1 | 5.067 | 0.10 0 | 5.549 | 0.12 4 | 8.127 | 0.06 7 | 0.683 | 0.560 |
| 4449 | **Morc2b** | Uncha nged | Q8C5W 4 | 1 | 1 | 1 | 102 2 | 117 .1 | 7.2 7 | | 1.245 | 0.73 0 | 0.642 | 0.55 1 | 1.189 | 0.77 1 | 0.540 | 0.412 |
| 4450 | **Tnk2** | Down | O54967 | 1 | 1 | 1 | 105 5 | 116 .9 | 7.4 3 | | 0.888 | 0.71 6 | 0.293 | 0.03 4 | 0.615 | 0.34 3 | 0.476 | 0.339 |
| 4451 | **Atp13a3** | Uncha nged | Q5XF89 | 1 | 10 | 1 | 121 9 | 137 .4 | 6.6 7 | 1 | 1.168 | 0.78 7 | 1.062 | 0.92 0 | 0.941 | 0.91 4 | 1.129 | 0.844 |
| 4452 | **Specc1** | UP | Q5SXY1 | 1 | 1 | 1 | 106 7 | 118 | 6.6 4 | 1 | 1.793 | 0.17 4 | 59.346 | 0.07 6 | 2.586 | 0.00 4 | 22.951 | 0.082 |
| 4453 | **Sugp2** | Uncha nged | Q8CH09 | 1 | 2 | 1 | 106 7 | 118 | 8.3 1 | | 1.256 | 0.76 4 | 1.189 | 0.84 0 | 0.488 | 0.34 2 | 2.436 | 0.395 |
| 4454 | **Abcc1** | Uncha nged | O35379 | 1 | 3 | 1 | 152 8 | 171 .1 | 7.3 6 | 1 | 1.109 | 0.86 0 | 0.812 | 0.73 7 | 1.174 | 0.79 2 | 0.692 | 0.579 |
| 4455 | **Tango6** | Down | Q8C3S2 | 1 | 4 | 1 | 107 9 | 118 .7 | 6.3 2 | 1 | 1.254 | 0.70 6 | 0.176 | 0.02 6 | 1.002 | 0.99 6 | 0.176 | 0.033 |
| 4456 | **Pdzph1** | Uncha nged | Q8BGR1 | 1 | 1 | 1 | 123 8 | 138 .5 | 6.7 6 | 1 | 1.289 | 0.70 0 | 1.354 | 0.66 1 | 0.280 | 0.27 2 | 4.842 | 0.092 |
| 4457 | **Trio** | Uncha nged | Q0KL02 | 1 | 1 | 1 | 310 2 | 347 .6 | 6.3 5 | 1 | 2.538 | 0.40 0 | 0.730 | 0.53 7 | 4.104 | 0.12 4 | 0.178 | 0.092 |
| 4458 | **Kcnq5** | Uncha nged | E9Q9F1 | 1 | 1 | 1 | 952 | 104 .6 | 9.5 8 | 1 | 0.976 | 0.97 2 | 1.588 | 0.48 2 | 1.136 | 0.80 4 | 1.398 | 0.564 |

| | | | | | | | | | | | | | | | | | | |
|---|---|---|---|---|---|---|---|---|---|---|---|---|---|---|---|---|---|---|
| 4459 | **Zeb1** | Unchanged | Q64318 | 1 | 1 | 1 | 1117 | 122.4 | 4.88 | | 3.600 | 0.260 | 3.419 | 0.372 | 3.683 | 0.193 | 0.928 | 0.935 |
| 4460 | **Pcnt** | Unchanged | F8VPV0 | 2 | 3 | 2 | 2916 | 331.1 | 5.38 | 1 | 1.125 | 0.770 | 1.096 | 0.831 | 1.064 | 0.875 | 1.030 | 0.952 |
| 4461 | **Rfx7** | Unchanged | F8VPJ6 | 1 | 1 | 1 | 1459 | 157.5 | 6.42 | 1 | 0.008 | 0.118 | 1.277 | 0.739 | 0.002 | 0.116 | 620.333 | 0.055 |
| 4462 | **Cog1** | Unchanged | Q9Z160 | 1 | 1 | 1 | 980 | 109 | 7.18 | 1 | #DIV/0! | #DIV/0! | #DIV/0! | #DIV/0! | #DIV/0! | #DIV/0! | #DIV/0! | #DIV/0! |
| 4463 | **Syne2** | Unchanged | Q6ZWQ0 | 2 | 11 | 2 | 6874 | 782.2 | 5.33 | 1 | 0.948 | 0.891 | 0.429 | 0.104 | 0.940 | 0.875 | 0.456 | 0.080 |
| 4464 | **Col12a1** | Unchanged | Q60847 | 2 | 2 | 2 | 3120 | 340 | 5.64 | 1 | 3.716 | 0.247 | 4.249 | 0.217 | 5.250 | 0.136 | 0.809 | 0.780 |
| 4465 | **Pdzd2** | Unchanged | E9Q1M1 | 1 | 1 | 1 | 2796 | 296.5 | 7.65 | 1 | #DIV/0! | #DIV/0! | #DIV/0! | #DIV/0! | #DIV/0! | #DIV/0! | #DIV/0! | #DIV/0! |
| 4466 | **Stk36** | Unchanged | Q69ZM6 | 1 | 2 | 1 | 1316 | 144.1 | 5.92 | 1 | 0.446 | 0.190 | 0.493 | 0.234 | 0.506 | 0.220 | 0.973 | 0.958 |
| 4467 | **Ccdc88c** | Unchanged | Q6VGS5 | 1 | 1 | 1 | 2009 | 226.4 | 6.15 | 1 | #DIV/0! | #DIV/0! | #DIV/0! | #DIV/0! | #DIV/0! | #DIV/0! | #DIV/0! | #DIV/0! |
| 4468 | **Slc9c1** | Unchanged | Q6UJY2 | 1 | 1 | 1 | 1175 | 135.5 | 6.74 | 1 | 1.185 | 0.603 | 1.108 | 0.830 | 2.064 | 0.404 | 0.537 | 0.463 |
| 4469 | **Nbea** | Unchanged | Q9EPN1 | 1 | 2 | 1 | 2936 | 326.5 | 6.2 | 1 | 1.722 | 0.460 | 1.426 | 0.671 | 1.467 | 0.510 | 0.972 | 0.970 |
| 4470 | **Dennd4c** | Unchanged | A6H8H2 | 1 | 3 | 1 | 1906 | 211.3 | 6.4 | 1 | 1.138 | 0.746 | 0.769 | 0.503 | 0.709 | 0.346 | 1.085 | 0.832 |
| 4471 | **Ms4a14** | Unchanged | A0A087WSD2 | 1 | 1 | 1 | 1232 | 137.5 | 4.69 | 1 | #DIV/0! | #DIV/0! | #DIV/0! | #DIV/0! | #DIV/0! | #DIV/0! | #DIV/0! | #DIV/0! |
| 4472 | **Srcin1** | Unchanged | Q9QWI6 | 1 | 3 | 1 | 1250 | 134.8 | 9.32 | | 0.737 | 0.541 | 2.181 | 0.311 | 0.871 | 0.778 | 2.505 | 0.271 |
| 4473 | **Top2b** | Unchanged | Q64511 | 1 | 2 | 1 | 1612 | 181.8 | 8.29 | 1 | 1.143 | 0.812 | 1.950 | 0.352 | 0.924 | 0.862 | 2.111 | 0.272 |
| 4474 | **Fyco1** | Unchanged | Q8VDC1 | 1 | 1 | 1 | 1437 | 162.2 | 4.98 | 1 | 0.048 | 0.140 | 0.019 | 0.130 | 0.043 | 0.138 | 0.452 | 0.295 |
| 4475 | **Anks1b** | Unchanged | Q8BIZ1 | 1 | 1 | 1 | 1259 | 139 | 6.29 | | 0.561 | 0.570 | 0.348 | 0.387 | 1.092 | 0.920 | 0.319 | 0.254 |
| 4476 | **Tacc2** | Unchanged | E9Q8T1 | 1 | 1 | 1 | 2879 | 305 | 4.79 | 1 | 0.252 | 0.240 | 0.162 | 0.184 | 0.028 | 0.127 | 5.725 | 0.128 |
| 4477 | **Plcg2** | Unchanged | Q8CIH5 | 1 | 2 | 1 | 1265 | 147.5 | 6.77 | 1 | 1.368 | 0.566 | 0.723 | 0.622 | 0.376 | 0.247 | 1.924 | 0.223 |
| 4478 | **Obscn** | Unchanged | A2AAJ9 | 2 | 2 | 2 | 8891 | 965.8 | 5.95 | 1 | 3.654 | 0.273 | 2.913 | 0.374 | 3.530 | 0.213 | 0.825 | 0.824 |
| 4479 | **Nos1** | Unchanged | F8WGF2 | 1 | 9 | 1 | 1463 | 164.2 | 7.24 | 1 | 1.279 | 0.656 | 1.119 | 0.820 | 1.618 | 0.455 | 0.691 | 0.573 |
| 4480 | **Per1** | Unchanged | O35973 | 1 | 1 | 1 | 1291 | 136.3 | 6.13 | 1 | 0.941 | 0.905 | 0.886 | 0.760 | 0.440 | 0.086 | 2.013 | 0.081 |
| 4481 | **Sbf1** | UP | Q6ZPE2 | 1 | 2 | 1 | 1867 | 208.6 | 7.12 | 1 | 1.615 | 0.087 | 1.445 | 0.265 | 1.813 | 0.004 | 0.797 | 0.356 |
| 4482 | **Nutm1** | Unchanged | Q8BHP2 | 1 | 1 | 1 | 1126 | 120.6 | 5.45 | 1 | 0.450 | 0.396 | 0.035 | 0.062 | 0.035 | 0.062 | 1.002 | 0.995 |
| 4483 | **Ccdc171** | Unchanged | E9Q1U1 | 1 | 7 | 1 | 1324 | 152.2 | 6.76 | 1 | 1.125 | 0.852 | 0.311 | 0.209 | 1.120 | 0.861 | 0.278 | 0.189 |
| 4484 | **Atp8b3** | Unchanged | Q6UQ17 | 1 | 2 | 1 | 1335 | 151.9 | 8.02 | 1 | 1.214 | 0.749 | 0.655 | 0.549 | 1.457 | 0.520 | 0.450 | 0.249 |
| 4485 | **Wdr19** | Unchanged | Q3UGF1 | 1 | 1 | 1 | 1341 | 151.4 | 6.57 | | 1.235 | 0.817 | 0.607 | 0.632 | 0.004 | 0.141 | 147.000 | 0.267 |
| 4486 | **Nol6** | Unchanged | Q8R5K4 | 1 | 1 | 1 | 1152 | 129.1 | 6.79 | 1 | 2.283 | 0.339 | 3.246 | 0.254 | 2.529 | 0.272 | 1.283 | 0.740 |
| 4487 | **Pcnx4** | Down | E9QN69 | 1 | 6 | 1 | 1177 | 131.4 | 6.54 | 1 | 1.798 | 0.447 | 0.112 | 0.038 | 0.009 | 0.024 | 12.127 | 0.015 |
| 4488 | **Arhgef5** | UP | E9Q7D5 | 1 | 2 | 1 | 1581 | 176.6 | 5.83 | 1 | 14.329 | 0.266 | 4.335 | 0.311 | 6.310 | 0.017 | 0.687 | 0.590 |
| 4489 | **Dst** | Unchanged | Q91ZU6 | 3 | 19 | 3 | 7393 | 833.7 | 5.31 | 1 | 0.683 | 0.572 | 1.214 | 0.796 | 0.876 | 0.835 | 1.385 | 0.675 |
| 4490 | **Muc5b** | Unchanged | E9Q5I3 | 1 | 1 | 1 | 4800 | 515.1 | 5.73 | 1 | #DIV/0! | #DIV/0! | #DIV/0! | #DIV/0! | #DIV/0! | #DIV/0! | #DIV/0! | #DIV/0! |
| 4491 | **D430041D05Rikiso** | Unchanged | A0A2R8VHF1 | 1 | 1 | 1 | 1415 | 153.4 | 9.39 | 0 | 0.671 | 0.494 | 1.084 | 0.923 | 0.004 | 0.053 | 302.154 | 0.169 |
| 4492 | **Dmxl2** | Unchanged | B0V2P5 | 1 | 1 | 1 | 3054 | 340.5 | 6.4 | 0 | #DIV/0! | #DIV/0! | #DIV/0! | #DIV/0! | #DIV/0! | #DIV/0! | #DIV/0! | #DIV/0! |
| 4493 | **Rnf17** | Unchanged | Q99MV7 | 1 | 3 | 1 | 1640 | 185.5 | 5.48 | 0 | 0.851 | 0.454 | 0.835 | 0.663 | 0.599 | 0.087 | 1.393 | 0.550 |
| 4494 | **Ccdc7a** | Unchanged | A0A1L1SS82 | 1 | 2 | 1 | 1642 | 184.8 | 9.13 | 0 | 4.050 | 0.080 | 1.658 | 0.462 | 4.793 | 0.082 | 0.346 | 0.164 |
| 4495 | **Myom3** | UP | A2ABU4 | 1 | 1 | 1 | 1439 | 161.7 | 5.72 | 0 | 1.126 | 0.875 | 4.235 | 0.024 | 0.519 | 0.433 | 8.152 | 0.009 |
| 4496 | **Zfhx3** | Unchanged | Q61329 | 1 | 2 | 1 | 3726 | 406.3 | 6.24 | 0 | 2.663 | 0.360 | 4.372 | 0.224 | 0.115 | 0.160 | 37.912 | 0.127 |
| 4497 | **Dock5** | Unchanged | B2RY04 | 1 | 3 | 1 | 1868 | 214.3 | 7.55 | 0 | 33.321 | 0.079 | 1.046 | 0.920 | 40.958 | 0.082 | 0.026 | 0.082 |
| 4498 | **Nktr** | Unchanged | P30415 | 1 | 14 | 1 | 1453 | 163.4 | 10.05 | 0 | 1.642 | 0.503 | 2.086 | 0.335 | 1.925 | 0.369 | 1.084 | 0.901 |
| 4499 | **Htt** | Down | G3X9H5 | 1 | 1 | 1 | 3120 | 344.6 | 6.28 | 0 | 0.865 | 0.336 | 0.915 | 0.751 | 0.546 | 0.019 | 1.677 | 0.220 |

| | | | | | | | | | | | | | | | | | | |
|---|---|---|---|---|---|---|---|---|---|---|---|---|---|---|---|---|---|---|
| 4500 | **Plxna4** | Unchanged | Q80UG2 | 1 | 8 | 1 | 1893 | 212.4 | 6.83 | 0 | 0.990 | 0.989 | 1.115 | 0.888 | 1.045 | 0.941 | 1.067 | 0.933 |
| 4501 | **Plxna1** | Unchanged | P70206 | 1 | 3 | 1 | 1894 | 211 | 6.9 | 0 | 1.378 | 0.547 | 1.777 | 0.429 | 1.082 | 0.871 | 1.643 | 0.474 |
| 4502 | **Arhgap29** | Unchanged | Q8CGF1 | 1 | 1 | 1 | 1266 | 142.3 | 7.17 | 0 | 0.732 | 0.699 | 0.584 | 0.540 | 0.261 | 0.229 | 2.240 | 0.360 |
| 4503 | **Mki67** | Unchanged | E9PVX6 | 1 | 1 | 1 | 3177 | 350.7 | 9.72 | 0 | 1.053 | 0.927 | 2.099 | 0.336 | 0.997 | 0.996 | 2.105 | 0.336 |
| 4504 | **Gemin5** | Unchanged | Q8BX17 | 1 | 2 | 1 | 1502 | 166.5 | 6.71 | | #DIV/0! | #DIV/0! | #DIV/0! | #DIV/0! | #DIV/0! | #DIV/0! | #DIV/0! | #DIV/0! |
| 4505 | **Tep1** | Down | P97499 | 1 | 1 | 1 | 2629 | 291.3 | 7.24 | 0 | 0.240 | 0.059 | 0.112 | 0.033 | 0.239 | 0.065 | 0.468 | 0.226 |
| 4506 | **Vps13d** | Unchanged | B1ART2 | 1 | 1 | 1 | 4390 | 490.7 | 6.55 | 0 | 1.136 | 0.844 | 0.324 | 0.211 | 0.595 | 0.430 | 0.544 | 0.313 |
| 4507 | **Eml5** | Unchanged | Q8BQM8 | 1 | 1 | 1 | 1977 | 220 | 7.8 | 0 | 0.853 | 0.810 | 0.796 | 0.776 | 0.178 | 0.100 | 4.478 | 0.286 |
| 4508 | **Ryr3** | Unchanged | A2AGL3 | 2 | 10 | 1 | 4863 | 551 | 5.78 | 0 | 1.144 | 0.864 | 1.788 | 0.531 | 0.747 | 0.772 | 2.394 | 0.426 |
| 4509 | **Kdm5c** | Unchanged | P41230 | 1 | 7 | 1 | 1554 | 175.2 | 5.44 | 0 | 1.069 | 0.920 | 1.518 | 0.669 | 1.705 | 0.485 | 0.890 | 0.892 |
| 4510 | **Synm** | Unchanged | Q70IV5 | 1 | 6 | 1 | 1561 | 173.1 | 5.12 | 0 | 1.708 | 0.528 | 1.167 | 0.821 | 1.122 | 0.884 | 1.040 | 0.957 |
| 4511 | **Gm7173** | Down | A0A0G2JEB6 | 1 | 1 | 1 | 3132 | 354.5 | 7.99 | 0 | 1.584 | 0.482 | 0.012 | 0.035 | 2.180 | 0.318 | 0.005 | 0.070 |
| 4512 | **Zfp831** | Unchanged | A2ADM8 | 1 | 3 | 1 | 1628 | 174 | 7.9 | 0 | 1.257 | 0.795 | 1.449 | 0.715 | 0.119 | 0.240 | 12.141 | 0.208 |
| 4513 | **Kat6b** | Unchanged | Q8BRB7 | 1 | 4 | 1 | 1872 | 208.4 | 5.97 | | 0.461 | 0.362 | 0.301 | 0.236 | 0.494 | 0.395 | 0.610 | 0.485 |
| 4514 | **Dchs1** | Unchanged | E9PVD3 | 1 | 1 | 1 | 3291 | 346.2 | 5 | 0 | 5.793 | 0.181 | 0.965 | 0.961 | 8.347 | 0.080 | 0.116 | 0.081 |
| 4515 | **Muc16** | Unchanged | A0A140LJ72 | 1 | 1 | 1 | 8478 | 908.2 | 6.04 | 0 | 1.809 | 0.527 | 1.742 | 0.541 | 0.289 | 0.319 | 6.018 | 0.166 |
| 4516 | **Ryr2** | Unchanged | E9Q401 | 1 | 3 | 1 | 4966 | 564.5 | 6.09 | 0 | 5.661 | 0.100 | 14.006 | 0.215 | 3.780 | 0.050 | 3.705 | 0.323 |
| 4517 | **Shprh** | Unchanged | Q7TPQ3 | 1 | 1 | 1 | 1674 | 191.4 | 7.46 | 0 | 1.702 | 0.566 | 0.625 | 0.610 | 0.074 | 0.146 | 8.444 | 0.220 |
| 4518 | **Plch1** | Unchanged | Q4KWH5 | 1 | 2 | 1 | 1682 | 187.6 | 7.33 | 0 | #DIV/0! | #DIV/0! | #DIV/0! | #DIV/0! | #DIV/0! | #DIV/0! | #DIV/0! | #DIV/0! |
| 4519 | **Ythdc2** | Unchanged | B2RR83 | 1 | 1 | 1 | 1445 | 161 | 8.51 | 0 | #DIV/0! | #DIV/0! | #DIV/0! | #DIV/0! | #DIV/0! | #DIV/0! | #DIV/0! | #DIV/0! |
| 4520 | **Mtcl1** | Unchanged | Q3UHU5 | 1 | 1 | 1 | 1945 | 213.7 | 6.14 | 0 | 1.310 | 0.739 | 0.829 | 0.846 | 1.729 | 0.521 | 0.479 | 0.413 |
| 4521 | **Nbeal1** | UP | E9PYP2 | 1 | 1 | 1 | 2688 | 306 | 6.39 | 0 | 1.011 | 0.976 | 44.729 | 0.018 | 23.676 | 0.031 | 1.889 | 0.254 |
| 4522 | **Chd6** | Unchanged | A3KFM7 | 1 | 6 | 1 | 2711 | 305.2 | 6.38 | 0 | 1.472 | 0.613 | 2.158 | 0.414 | 2.041 | 0.059 | 1.058 | 0.931 |
| 4523 | **Shoc1** | Unchanged | A2ALV5 | 1 | 6 | 1 | 1481 | 168 | 5.91 | 0 | 0.395 | 0.264 | 0.087 | 0.093 | 2.497 | 0.279 | 0.035 | 0.079 |
| 4524 | **Brca1** | Unchanged | P48754 | 1 | 1 | 1 | 1812 | 198.7 | 5.76 | 0 | 0.777 | 0.546 | 0.506 | 0.207 | 0.632 | 0.295 | 0.800 | 0.520 |
| 4525 | **Tro** | Unchanged | Q6DIC6 | 1 | 1 | 1 | 2087 | 205.9 | 9.31 | 0 | 0.741 | 0.713 | 0.322 | 0.221 | 0.231 | 0.136 | 1.395 | 0.806 |
| 4526 | **Dmd** | Unchanged | P11531 | 1 | 1 | 1 | 3678 | 425.6 | 5.94 | 0 | #DIV/0! | #DIV/0! | #DIV/0! | #DIV/0! | #DIV/0! | #DIV/0! | #DIV/0! | #DIV/0! |
| 4527 | **Adgrl3** | Unchanged | D3Z5M6 | 1 | 1 | 1 | 1580 | 175.6 | 6.74 | 0 | 4.529 | 0.376 | 3.712 | 0.424 | 4.049 | 0.115 | 0.917 | 0.927 |
| 4528 | **Syne1** | Unchanged | A0A1L1STC6 | 3 | 3 | 3 | 8799 | 1009 | 5.58 | 0 | 1.171 | 0.812 | 1.548 | 0.431 | 0.589 | 0.470 | 2.627 | 0.076 |
| 4529 | **Synj1** | Unchanged | D3Z656 | 1 | 1 | 1 | 1607 | 176.2 | 7.72 | 0 | 2.047 | 0.114 | 0.941 | 0.901 | 1.944 | 0.295 | 0.484 | 0.298 |
| 4530 | **Herc1** | Unchanged | E9PZP8 | 1 | 1 | 1 | 4859 | 532.3 | 6.04 | 0 | #DIV/0! | #DIV/0! | #DIV/0! | #DIV/0! | #DIV/0! | #DIV/0! | #DIV/0! | #DIV/0! |
| 4531 | **Kiaa1549** | Unchanged | Q68FD9 | 1 | 3 | 1 | 1940 | 209.1 | 6.1 | 0 | 2.381 | 0.173 | 0.087 | 0.186 | 0.104 | 0.192 | 0.843 | 0.861 |
| 4532 | **Dnah17** | Unchanged | K3W4R0 | 1 | 1 | 1 | 4481 | 511.1 | 5.63 | 0 | 0.867 | 0.813 | 1.219 | 0.793 | 0.064 | 0.059 | 18.919 | 0.132 |
| 4533 | **Polr2a** | UP | P08775 | 1 | 1 | 1 | 1970 | 217 | 7.37 | 0 | 0.935 | 0.949 | 3.346 | 0.023 | 1.480 | 0.667 | 2.260 | 0.151 |
| 4534 | **Zfc3h1** | Down | B2RT41 | 1 | 4 | 1 | 1992 | 224.9 | 8.15 | 0 | 1.088 | 0.741 | 0.934 | 0.889 | 0.030 | 0.002 | 30.749 | 0.058 |
| 4535 | **Fcgbp** | Unchanged | E9Q0B5 | 1 | 7 | 1 | 2583 | 275 | 5.03 | 0 | 2.970 | 0.223 | 2.325 | 0.338 | 1.300 | 0.661 | 1.788 | 0.490 |
| 4536 | **Pcm1** | Unchanged | Q9R0L6 | 1 | 3 | 1 | 2025 | 228.7 | 5.01 | 0 | 2.390 | 0.275 | 2.289 | 0.422 | 2.593 | 0.136 | 0.883 | 0.862 |
| 4537 | **Spg11** | Unchanged | Q3UHA3 | 1 | 1 | 1 | 2430 | 273.8 | 5.94 | 0 | 1.504 | 0.505 | 1.078 | 0.933 | 1.637 | 0.495 | 0.658 | 0.634 |
| 4538 | **Igf2r** | Unchanged | Q07113 | 1 | 2 | 1 | 2483 | 273.6 | 5.71 | 0 | #DIV/0! | #DIV/0! | #DIV/0! | #DIV/0! | #DIV/0! | #DIV/0! | #DIV/0! | #DIV/0! |
| 4539 | **Myo15a** | Unchanged | Q9QZZ4 | 1 | 1 | 1 | 3511 | 395.4 | 9.16 | 0 | 0.497 | 0.116 | 1.264 | 0.535 | 3.027 | 0.287 | 0.417 | 0.352 |
| 4540 | **Bod1l** | Unchanged | E9Q6J5 | 1 | 1 | 1 | 3032 | 327.3 | 5.33 | 0 | 0.663 | 0.532 | 1.280 | 0.719 | 0.602 | 0.387 | 2.126 | 0.370 |

| | | | | | | | | | | | | | | | | | | |
|---|---|---|---|---|---|---|---|---|---|---|---|---|---|---|---|---|---|---|
| 4541 | **Ncor1** | Unchanged | Q5RIM6 | 1 | 1 | 1 | 2454 | 270.6 | 6.93 | 0 | 1.822 | 0.120 | 6.349 | 0.350 | 16.674 | 0.081 | 0.381 | 0.309 |
| 4542 | **Ncor2** | Unchanged | F8VQL9 | 1 | 1 | 1 | 2507 | 273.2 | 7.44 | 0 | 5.175 | 0.309 | 0.190 | 0.253 | 2.462 | 0.461 | 0.077 | 0.236 |
| 4543 | **Gm11639** | Unchanged | A0A1D5RLM8 | 2 | 4 | 2 | 5808 | 653.8 | 5.4 | 0 | #DIV/0! | #DIV/0! | #DIV/0! | #DIV/0! | #DIV/0! | #DIV/0! | #DIV/0! | #DIV/0! |
| 4544 | **Frem1** | Down | Q684R7 | 1 | 1 | 1 | 2191 | 244.4 | 6.39 | 0 | 1.477 | 0.591 | 1.794 | 0.261 | 0.158 | 0.042 | 11.378 | 0.019 |
| 4545 | **Myo18b** | Unchanged | E9PV66 | 1 | 1 | 1 | 2605 | 288.7 | 6.67 | 0 | #DIV/0! | #DIV/0! | #DIV/0! | #DIV/0! | #DIV/0! | #DIV/0! | #DIV/0! | #DIV/0! |
| 4546 | **Trank1** | Unchanged | Q8BV79 | 1 | 3 | 1 | 2999 | 343.1 | 7.81 | 0 | 1.406 | 0.517 | 0.879 | 0.821 | 0.840 | 0.724 | 1.047 | 0.942 |
| 4547 | **Aspm** | Unchanged | Q8CJ27 | 1 | 2 | 1 | 3122 | 364 | 10.62 | 0 | 33.806 | 0.165 | 124.903 | 0.118 | 1.573 | 0.092 | 79.426 | 0.120 |
| 4548 | **Nf1** | Unchanged | Q04690 | 1 | 2 | 1 | 2841 | 319.4 | 7.39 | 0 | 2.251 | 0.316 | 2.594 | 0.310 | 0.457 | 0.173 | 5.682 | 0.175 |
| 4549 | **Fbn1** | Unchanged | Q61554 | 1 | 1 | 1 | 2873 | 312.1 | 4.92 | 0 | 0.914 | 0.881 | 0.643 | 0.512 | 0.560 | 0.419 | 1.148 | 0.676 |
| 4550 | **Fat3** | Unchanged | Q8BNA6 | 1 | 1 | 1 | 4555 | 501.7 | 4.87 | 0 | #DIV/0! | #DIV/0! | #DIV/0! | #DIV/0! | #DIV/0! | #DIV/0! | #DIV/0! | #DIV/0! |
| 4551 | **Ttn** | Unchanged | A2ASS6 | 6 | 14 | 6 | 35213 | 3904 | 6.2 | 0 | 0.889 | 0.777 | 0.612 | 0.276 | 0.694 | 0.259 | 0.882 | 0.742 |
| 4552 | **Abca12** | Unchanged | E9Q876 | 1 | 6 | 1 | 2595 | 292.4 | 7.93 | 0 | 0.594 | 0.373 | 0.418 | 0.192 | 0.546 | 0.327 | 0.765 | 0.583 |
| 4553 | **Ep400** | Unchanged | Q8CHI8 | 1 | 4 | 1 | 3072 | 337 | 9.14 | 0 | 0.829 | 0.792 | 0.297 | 0.132 | 1.053 | 0.916 | 0.282 | 0.032 |
| 4554 | **Dnah8** | Unchanged | Q91XQ0 | 1 | 3 | 1 | 4731 | 540.9 | 6.18 | 0 | 0.734 | 0.612 | 0.511 | 0.352 | 0.561 | 0.375 | 0.911 | 0.895 |
| 4555 | **Pkd1** | Unchanged | O08852 | 1 | 8 | 1 | 4293 | 466.3 | 5.87 | 0 | 0.998 | 0.997 | 1.196 | 0.792 | 0.488 | 0.283 | 2.453 | 0.254 |
| 4556 | **Dync2h1** | Unchanged | Q45VK7 | 1 | 1 | 1 | 4306 | 492 | 6.6 | 0 | 2.024 | 0.120 | 5.376 | 0.091 | 1.178 | 0.723 | 4.563 | 0.100 |
| 4557 | **Hmcn1** | UP | D3YXG0 | 1 | 2 | 1 | 5634 | 611.2 | 6.64 | 0 | 4.544 | 0.157 | 13.514 | 0.093 | 11.244 | 0.009 | 1.202 | 0.760 |
| 4558 | **Neb** | Unchanged | E9Q1W3 | 1 | 1 | 1 | 7152 | 828.1 | 9.07 | 0 | 0.375 | 0.208 | 0.446 | 0.281 | 0.440 | 0.273 | 1.014 | 0.984 |

**Supplementary Table 4: Identified proteins association with OMIM database for liver fibrosis**

Supplementary Table 4. Identified proteins association with OMIM database for liver fibrosis.

| Classification | Proteins | ASA | CCl4 | ASA-CCl4 | Control |
|---|---|---|---|---|---|
| Fibrosis | Mpv17 | 7.43 | 11.57 | 8.10 | 1.00 |
| | Ryr2 | 5.66 | 14.01 | 3.78 | 1.00 |
| | Nr3c2 | 4.02 | 4.54 | 3.56 | 1.00 |
| | Nfkb1 | 3.54 | -1.57 | 2.80 | 1.00 |
| | Hjv | 1.61 | 3.11 | 2.44 | 1.00 |
| | Dnajb11 | 1.98 | 1.41 | 1.64 | 1.00 |
| | Tardbp | 2.05 | 3.70 | 1.64 | 1.00 |
| | Urod | 1.49 | 2.13 | 1.56 | 1.00 |
| | Sec61a1 | 1.43 | 3.41 | 1.48 | 1.00 |
| | Ppa2 | 1.22 | 2.39 | 1.48 | 1.00 |
| | Abcd3 | 1.11 | 1.61 | 1.41 | 1.00 |
| | Dbt | 1.21 | 1.63 | 1.38 | 1.00 |
| | Btd | -1.04 | 3.50 | 1.37 | 1.00 |
| | Acox2 | 1.17 | 2.67 | 1.30 | 1.00 |
| | Gbe1 | 1.23 | 1.17 | 1.30 | 1.00 |
| | Col25a1 | 1.21 | -2.63 | 1.27 | 1.00 |
| | Lpp | -1.26 | -1.64 | 1.09 | 1.00 |
| | Trappc11 | 1.53 | 1.39 | 1.06 | 1.00 |
| | Vps45 | -1.07 | 1.94 | 1.04 | 1.00 |
| | Ap3b1 | 1.09 | 1.83 | -1.12 | 1.00 |
| | Ctnnb1 | -1.30 | 1.77 | -1.22 | 1.00 |
| | Nup133 | 2.44 | 4.14 | -1.28 | 1.00 |
| | Ttn | -1.13 | -1.63 | -1.44 | 1.00 |
| | Ptprc | -1.27 | 1.70 | -1.60 | 1.00 |
| | Agxt | -1.19 | -1.21 | -1.61 | 1.00 |
| | Cnbp | -1.43 | -3.39 | -1.67 | 1.00 |
| | Bcs1l | -1.61 | 1.53 | -1.72 | 1.00 |
| | Slc34a2 | 1.30 | 1.36 | -1.73 | 1.00 |
| | Tert | -28.55 | -23.89 | -1.91 | 1.00 |
| | Ttc37 | -1.52 | -1.19 | -1.93 | 1.00 |
| | Dag1 | -1.30 | -1.96 | -2.10 | 1.00 |
| | Apoa2 | -1.32 | -4.29 | -2.14 | 1.00 |
| | Neb | -2.67 | -2.24 | -2.27 | 1.00 |
| | Casp8 | -2.50 | -4.18 | -4.50 | 1.00 |
| | Itgb6 | -1.25 | -2.74 | -5.22 | 1.00 |
| | Des | -1.83 | -1.10 | -13.92 | 1.00 |

| | | | | | |
|---|---|---|---|---|---|
| | Myl1 | -1.17 | -2.16 | -25.45 | 1.00 |
| | Wdr19 | 1.24 | -1.65 | -24.31 | 1.00 |
| **Liver and Fibrosis** | Ece1 | -0.90 | 2.94 | 4.76 | 1.00 |
| | Dhcr24 | 3.96 | 5.27 | 3.70 | 1.00 |
| | Ptprf | -8.60 | 2.28 | 2.99 | 1.00 |
| | Gnb1 | 1.95 | 2.67 | 1.93 | 1.00 |
| | Slc35d1 | 1.14 | 1.37 | 1.76 | 1.00 |
| | Szt2 | 1.83 | 1.35 | 1.64 | 1.00 |
| | Faah | 1.17 | 2.10 | 1.50 | 1.00 |
| | Ppt1 | 1.48 | 1.72 | 1.41 | 1.00 |
| | H6pd | 1.16 | 2.44 | 1.13 | 1.00 |
| | Ddost | -1.04 | 1.48 | -1.11 | 1.00 |
| | Kif1b | -1.69 | -1.06 | -1.28 | 1.00 |
| | Gale | 1.62 | 1.13 | -1.29 | 1.00 |
| | Yars | -1.30 | -1.76 | -1.98 | 1.00 |
| | Epb41 | -2.45 | -12.53 | -2.02 | 1.00 |
| | Adprhl2 | -1.67 | -1.15 | -4.77 | 1.00 |
| | Epha2 | 2.03 | -11.70 | -5.00 | 1.00 |

**Red= >1.5**

**Green = <0.65**

### Supplementary Table 5. List of identified metabolites across all mice liver samples and their quantification.

| Supplementary Table 5. List of identified metabolites across all mice liver samples and their quantification. | | | | | | | | | | | | | | | | | | | |
|---|---|---|---|---|---|---|---|---|---|---|---|---|---|---|---|---|---|---|---|
| Sr. no | Samples | Attribution | Mode of ionisation | Identification | column Hilic/ C18 | Molecular Weight | RT | KEGG ALL | HMDB | PubChem | KEGG Pathways | ASA-control | pvalue ASA/control | CCl4/control | pvalue CCl4/control | CCL4ASA/ control | p-value CCl4ASA/ control | CCL4/ccl4-ASA | Pvalue CCl4/ccl4ASA |
| 1 | Glycine | [M+H]+1 | POS | Mass, Spectral match | Hilic significant | 75.033 | 6.630 | C00037 | HMDB0000123 | 750 | Metabolic pathways;Biosynthesis of secondary metabolites;Microbial metabolism in diverse environments;Biosynthesis of antibiotics;Biosynthesis of amino acids;Biosynthesis of plant secondary metabolites;ABC transporters;Protein digestion and absorption;Cyanoamino acid metabolism;Aminoacyl-tRNA biosynthesis;Central carbon metabolism in cancer;Glycine, serine and threonine metabolism;Lysine degradation;Other | 0.68 | 0.063 | 0.92 | 0.695 | 1.08 | 0.256 | 0.85 | 0.482201 |
| 2 | Pyridine | [M+H]+1 | POS | Mass, Spectral match | Hilic | 79.043 | 2.727 | C00747 | HMDB0000926 | 1049 | | 1.31 | 0.182 | 0.86 | 0.484 | 0.89 | 0.561 | 0.96 | 0.717244 |
| 3 | Pyrimidine | [M+H]+1 | POS | Mass, Spectral match | C18 | 80.038 | 16.441 | C00396 | HMDB0003361 | 9260 | | 0.70 | 0.468 | 1.05 | 0.875 | 1.15 | 0.156 | 0.91 | 0.538843 |
| 4 | Pyrimidine | [M+H]+1 | POS | Mass, Spectral match | Hilic | 80.038 | 4.463 | C00396 | HMDB0003361 | 9260 | | 1.23 | 0.696 | 0.63 | 0.427 | 0.62 | 0.453 | 1.01 | 0.813424 |
| 5 | Cobalt | [M+H]+1 | POS | Mass, Spectral match | Hilic | 81.923 | 1.142 | C00175 | HMDB0000608 | 104729 | | 1.98 | 0.287 | 4.77 | 0.009 | 5.54 | 0.086 | 0.86 | 0.934204 |
| 6 | Dimethylamine | [M+H]+1 | POS | Mass, Spectral match | C18 | 84.021 | 0.554 | C00543 | HMDB0000087 | 674 | Other | 0.95 | 0.879 | 2.28 | 0.431 | 2.44 | 0.436 | 0.93 | 0.871852 |
| 7 | 4_Hydroxy_2_butynal | [M+H]+1 | POS | Mass, Spectral match | Hilic | 84.021 | 3.211 | C04486 | HMDB0034453 | 440357 | Other | 1.24 | 0.234 | 1.12 | 0.562 | 0.99 | 0.856 | 1.13 | 0.481923 |
| 8 | Acetonecyanohydrin | [M+H]+1 | POS | Mass, Spectral match | Hilic | 85.053 | 1.239 | C02659 | MA3TEM059 | 5632 | Biosynthesis of secondary metabolites;Cyanoamino acid metabolism | 1.43 | 0.349 | 1.18 | 0.295 | 1.03 | 0.434 | 1.15 | 0.408128 |
| 9 | Piperidine | [M+H]+1 | POS | Mass, Spectral match | Hilic | 85.089 | 0.947 | C01746 | HMDB0034301 | 8082 | Protein digestion and absorption;Other | 1.04 | 0.228 | 1.03 | 0.265 | 1.07 | 0.369 | 0.96 | 0.564573 |
| 10 | Piperidine | [M+H]+1 | POS | Mass, Spectral match | C18 | 85.090 | 0.740 | C01746 | HMDB0034301 | 8082 | Protein digestion and absorption;Tropane, piperidine and pyridine alkaloid biosynthesis | 1.10 | 0.563 | 0.88 | 0.513 | 1.04 | 0.624 | 0.85 | 0.379426 |
| 11 | Crotonicacid | [M+H]+1 | POS | Mass, Spectral match | Hilic | 86.036 | 6.713 | C01771 | HMDB0010720 | 637090 | | 1.13 | 0.577 | 0.93 | 0.540 | 1.01 | 0.378 | 0.92 | 0.875156 |
| 12 | _AminobutyricacidGABA | [M+H]+1 | POS | Mass, Spectral | Hilic | 86.037 | 1.159 | C00334 | HMDB00112 | 119 | Microbial metabolism in diverse environments;Other | 0.93 | 0.438 | 0.94 | 0.509 | 0.91 | 0.773 | 1.03 | 0.762855 |

| | | | | | | | | | | | | | | | | | | | |
|---|---|---|---|---|---|---|---|---|---|---|---|---|---|---|---|---|---|---|---|
| | | | | match | | | | | | | | | | | | | | | |
| 13 | 3_Aminopyrrolidine | [M+H]+1 | POS | Mass, Spectral match | Hilic | 86.085 | 5.235 | C10164 | HMDB0002243 | 1018 | | 1.89 | 0.354 | 1.20 | 0.612 | 0.95 | 0.846 | 1.26 | 0.609776 |
| 14 | 2_Aminoacrylic_acid | [M+H]+1 | POS | Mass, Spectral match | Hilic | 87.032 | 8.537 | C02218 | HMDB0003609 | 123991 | Metabolic pathways;Cysteine and methionine metabolism;Pyrimidine metabolism | 1.03 | 0.861 | 1.27 | 0.472 | 7.20 | 0.354 | 0.18 | 0.411372 |
| 15 | 2_Aminoacrylic_acid | [M+H]+1 | POS | Mass, Spectral match | C18 | 87.032 | 0.511 | C02218 | HMDB0003609 | 123991 | Metabolic pathways;Other | 1.01 | 0.915 | 0.70 | 0.005 | 0.86 | 0.946 | 0.81 | 0.089208 |
| 16 | 4_Aminobutyraldehyde | [M+H]+1 | POS | Mass, Spectral match | Hilic | 87.068 | 0.799 | C00555 | HMDB0001080 | 118 | Metabolic pathways;Biosynthesis of secondary metabolites;Cyanoamino acid metabolism;Arginine and proline metabolism;2-Oxocarboxylic acid metabolism;Other | 0.97 | 0.874 | 1.42 | 0.113 | 0.69 | 0.528 | 2.06 | 0.004104 |
| 20 | D_Lactic_acid | [M+H]+1 | POS | Mass, Spectral match | Hilic significant | 90.032 | 9.375 | C00256 | HMDB0001311 | 61503 | Metabolic pathways;Biosynthesis of secondary metabolites;Microbial metabolism in diverse environments;Biosynthesis of antibiotics;Central carbon metabolism in cancer;Pyrimidine metabolism;Other | 1.01 | 0.983 | 0.53 | 0.023 | 0.65 | 0.527 | 0.81 | 0.340093 |
| 18 | L_Alanine | [M+H]+1 | POS | Mass, Spectral match | C18 | 89.048 | 0.491 | C00041 | HMDB0000161 | 5950 | Metabolic pathways;Biosynthesis of secondary metabolites;Microbial metabolism in diverse environments;Biosynthesis of plant secondary metabolites;Biosynthesis of amino acids;ABC transporters;Protein digestion and absorption;Arginine and proline metabolism;Central carbon metabolism in cancer;Aminoacyl-tRNA biosynthesis;Neuroactive ligand-receptor interaction;Other | 0.92 | 0.484 | 1.00 | 0.999 | 0.95 | 0.542 | 1.05 | 0.657148 |
| 19 | _Alanine | [M+H]+1 | POS | Mass, Spectral match | Hilic | 89.048 | 4.707 | C00041 | HMDB0000161 | 5950 | Metabolic pathways;Biosynthesis of secondary metabolites;Microbial metabolism in diverse environments;Biosynthesis of amino acids;Biosynthesis of plant secondary metabolites;ABC transporters;Protein digestion and absorption;Cyanoamino acid metabolism;Arginine and proline metabolism;Aminoacyl-tRNA biosynthesis;Central carbon metabolism in cancer;Glycine, | 0.65 | 0.325 | 0.93 | 0.819 | 0.95 | 0.499 | 0.98 | 0.907996 |

| | | | | | | | | | | | | | | | | | | | |
|---|---|---|---|---|---|---|---|---|---|---|---|---|---|---|---|---|---|---|---|
| | | | | | | | | | | | serine and threonine metabolism;Cysteine and methionine metabolism;Pyrimidine metabolism;Other | | | | | | | | |
| 46 | 3_Hydroxybutyricacid | [M+H]+1 | POS | Mass, Spectral match | Hilic | 104.046 | 6.413 | C01089 | HMDB0000357 | 441 | | 0.82 | 0.300 | 0.72 | 0.189 | 0.54 | 0.377 | 1.34 | 0.360542 |
| 21 | Toluene | [M+H]+1 | POS | Mass, Spectral match | C18 | 92.063 | 5.239 | C01455 | HMDB0034168 | 1140 | Metabolic pathways;Microbial metabolism in diverse environments;Degradation of aromatic compounds;Other | 0.98 | 0.842 | 0.84 | 0.161 | 0.89 | 0.298 | 0.93 | 0.453641 |
| 22 | Aniline | [M+H]+1 | POS | Mass,RT, Spectral match | Hilic | 93.058 | 2.642 | C00292 | HMDB0003012 | 6115 | Microbial metabolism in diverse environments;Other | 0.87 | 0.565 | 0.96 | 0.854 | 0.80 | 0.663 | 1.21 | 0.525041 |
| 23 | Aniline | [M+H]+1 | POS | Mass, RT, Spectral match | C18 | 93.058 | 18.119 | C00292 | HMDB0003012 | 6115 | Microbial metabolism in diverse environments;Other | 0.91 | 0.232 | 0.95 | 0.445 | 0.99 | 0.544 | 0.96 | 0.496791 |
| 24 | Phenol | [M+H]+1 | POS | Mass,RT, Spectral match | Hilic | 94.042 | 4.321 | C00146 | HMDB0000228 | 996 | Microbial metabolism in diverse environments;Protein digestion and absorption;Degradation of aromatic compounds;Other | 1.22 | 0.494 | 0.84 | 0.547 | 0.92 | 0.798 | 0.92 | 0.983664 |
| 25 | Ethanolamine | [M+H]+1 | POS | Mass, RT, Spectral match | C18 | 96.021 | 0.528 | C00189 | HMDB0000149 | 700 | Microbial metabolism in diverse environments;Other | 0.54 | 0.029 | 0.61 | 0.042 | 0.51 | 0.344 | 1.20 | 0.079052 |
| 26 | Ethylene_oxide | [M+H]+1 | POS | Mass, Spectral match | C18 | 97.006 | 18.620 | C01380 | HMDB0032262 | 174 | | 0.91 | 0.585 | 0.74 | 0.054 | 0.98 | 0.526 | 0.75 | 0.026979 |
| 47 | 4_Hydroxybutyric_acid | [M+H]+1 | POS | Mass, Spectral match | Hilic | 104.047 | 6.442 | C00989 | HMDB0000710 | 10413 | Metabolic pathways;Microbial metabolism in diverse environments;Other | 0.84 | 0.472 | 1.47 | 0.400 | 1.13 | 0.506 | 1.29 | 0.574653 |
| 28 | Phosphate | [M+H]+1 | POS | Mass, Spectral match | C18 | 97.977 | 0.547 | C00009 | HMDB0001429 | 1061 | | 0.93 | 0.721 | 0.92 | 0.617 | 0.76 | 0.748 | 1.21 | 0.035078 |
| 29 | Hydrogen_carbonate | [M-H]-1 | NEG | Mass, Spectral match | C18 | 99.955 | 0.572 | C00288 | HMDB0000595 | 769 | | 0.34 | 0.299 | 0.53 | 0.465 | 3.16 | 0.469 | 0.17 | 0.599873 |
| 30 | Hydantoin | [M+H]+1 | POS | Mass, Spectral match | Hilic | 100.027 | 2.396 | C01551 | HMDB0000462 | 204 | | 1.30 | 0.366 | 1.27 | 0.511 | 1.11 | 0.278 | 1.15 | 0.959232 |
| 31 | 1_Aminocyclopropane_1_carboxylate | [M+H]+1 | POS | Mass, Spectral match | C18 | 101.047 | 0.523 | C01234 | HMDB36458 | 535 | Metabolic pathways;Biosynthesis of secondary metabolites;Biosynthesis of plant secondary metabolites;Other | 1.11 | 0.569 | 0.88 | 0.440 | 0.74 | 0.752 | 1.18 | 0.328783 |
| 32 | 1_Aminocyclopropane_1_carboxylate | [M+H]+1 | POS | Mass, Spectral match | Hilic | 101.048 | 7.191 | C01234 | HMDB36458 | 535 | Metabolic pathways;Biosynthesis of secondary metabolites;Biosynthesis of plant secondary metabolites;Cysteine and methionine metabolism;Other | 1.09 | 0.599 | 0.95 | 0.776 | 0.96 | 0.866 | 0.99 | 0.996623 |
| 33 | Betaine_aldehyde | [M+H]+1 | POS | Mass, RT, Spectral match | C18 | 101.084 | 0.509 | C00576 | HMDB0001252 | 249 | Metabolic pathways;Biosynthesis of secondary metabolites;Microbial metabolism in diverse environments;2-Oxocarboxylic acid metabolism;Tropane, | 1.20 | 0.514 | 1.65 | 0.155 | 0.55 | 0.292 | 3.02 | 0.009127 |

| | | | | | | | | | | | | | | | | | | | |
|---|---|---|---|---|---|---|---|---|---|---|---|---|---|---|---|---|---|---|---|
| | | | | | | | | | | | piperidine and pyridine alkaloid biosynthesis;Other | | | | | | | | |
| 34 | Cadaverine | [M+H]+1 | POS | Mass, RT, Spectral match | C18 | 101.108 | 0.603 | C01672 | HMDB0002322 | 273 | | 1.55 | 0.532 | 1.62 | 0.429 | 0.68 | 0.600 | 2.39 | 0.277788 |
| 35 | Hexylamine | [M+H]+1 | POS | Mass, RT, Spectral match | C18 | 101.121 | 18.113 | C08306 | HMDB0032323 | 8102 | | 0.87 | 0.040 | 0.96 | 0.530 | 0.89 | 0.241 | 1.08 | 0.41037 |
| 36 | Hexylamine | [M+H]+1 | POS | Mass, Spectral match | Hilic | 101.121 | 0.748 | C08306 | HMDB0032323 | 8102 | | 1.05 | 0.738 | 1.07 | 0.574 | 1.08 | 0.360 | 1.00 | 0.924167 |
| 292 | Monodehydroascorbate | [M+H]+1 | POS | Mass, Spectral match | Hilic | 174.016 | 2.617 | C01041 | METPA0114 | NA | | 8.84 | 0.126 | 2.11 | 0.425 | 7.09 | 0.243 | 0.30 | 0.324403 |
| 38 | 2_Ketobutyric_acid | [M-H]-1 | NEG | Mass, Spectral match | Hilic | 102.032 | 9.831 | C00109 | HMDB0000005 | 58 | Metabolic pathways;Biosynthesis of secondary metabolites;Microbial metabolism in diverse environments;Biosynthesis of antibiotics;Biosynthesis of amino acids;2-Oxocarboxylic acid metabolism;Glycine, serine and threonine metabolism;Cysteine and methionine metabolism;Lysine degradation;Other | 0.95 | 0.751 | 1.32 | 0.077 | 1.27 | 0.327 | 1.04 | 0.644951 |
| 39 | L_cycloserine | [M+H]+1 | POS | Mass, Spectral match | Hilic | 102.043 | 7.071 | C08057 | HMDB0014405 | 6234 | Biosynthesis of antibiotics;Other | 0.86 | 0.422 | 0.99 | 0.967 | 0.87 | 0.874 | 1.14 | 0.485688 |
| 40 | NN_Dimethylglycine | [M+H]+1 | POS | Mass, Spectral match | Hilic | 103.064 | 3.890 | C01026 | HMDB0000092 | 673 | Metabolic pathways;Microbial metabolism in diverse environments;Arginine and proline metabolism;Glycine, serine and threonine metabolism;Cysteine and methionine metabolism;Pyrimidine metabolism;Other | 1.01 | 0.984 | 1.07 | 0.856 | 1.60 | 0.141 | 0.66 | 0.421272 |
| 41 | Gamma_Aminobutyric_acid | [M+H]+1 | POS | Mass, Spectral match | C18 | 103.064 | 0.492 | C00334 | HMDB0000112 | 119 | Metabolic pathways;Microbial metabolism in diverse environments;Arginine and proline metabolism;Neuroactive ligand-receptor interaction;Other | 0.86 | 0.145 | 0.91 | 0.319 | 0.90 | 0.718 | 1.01 | 0.97976 |
| 42 | DL_3_Aminoisobutyric_acid | [M-H]-1 | NEG | Mass, Spectral match | Hilic significant | 103.064 | 3.329 | C05145 | HMDB0003911 | 64956 | Metabolic pathways;Microbial metabolism in diverse environments;Arginine and proline metabolism;Glycine, serine and threonine metabolism;Cysteine and methionine metabolism;Pyrimidine metabolism;Other | 1.13 | 0.798 | 0.99 | 0.959 | 2.92 | 0.020 | 0.34 | 0.014619 |
| 43 | Choline | [M+H]+1 | POS | Mass, Spectral match | Hilic | 103.100 | 0.736 | C00114 | HMDB0000097 | 305 | | 1.08 | 0.464 | 1.04 | 0.718 | 0.92 | 0.761 | 1.12 | 0.388503 |
| 44 | Choline | [M+H]+1 | POS | Mass, RT, Spectral match | C18 | 103.100 | 0.492 | C00114 | HMDB0000097 | 305 | | 1.15 | 0.243 | 0.95 | 0.563 | 0.85 | 0.893 | 1.11 | 0.355431 |

| | | | | | | | | | | | | | | | | | | | |
|---|---|---|---|---|---|---|---|---|---|---|---|---|---|---|---|---|---|---|---|
| 45 | Diethylenetriamine | [M+H]+1 | POS | Mass, Spectral match | Hilic | 103.111 | 6.919 | | HMDB0031413 | 8111 | | 2.24 | 0.092 | 3.14 | 0.011 | 3.37 | 0.045 | 0.93 | 0.933116 |
| 407 | Sedoheptulose | [M+H]+1 | POS | Mass, Spectral match | Hilic | 210.073 | 3.106 | C02076 | HMDB0003219 | 102926 | Other | 0.71 | 0.095 | 0.72 | 0.245 | 0.81 | 0.847 | 0.89 | 0.597117 |
| 439 | N_Acetyl_D_glucosamine | [M+H]+1 | POS | Mass, Spectral match | Hilic significant | 221.090 | 1.943 | C00140 | HMDB0000215 | 439174 | Metabolic pathways;ABC transporters;Other | 0.54 | 0.061 | 0.63 | 0.129 | 0.44 | 0.166 | 1.44 | 0.146235 |
| 48 | D_Serine | [M+H]+1 | POS | Mass, Spectral match | Hilic | 105.043 | 8.522 | C00065 | HMDB0000187 | 5951 | Metabolic pathways;Biosynthesis of secondary metabolites;Microbial metabolism in diverse environments;Biosynthesis of antibiotics;Biosynthesis of amino acids;Biosynthesis of plant secondary metabolites;ABC transporters;Protein digestion and absorption;Cyanoamino acid metabolism;Aminoacyl-tRNA biosynthesis;Central carbon metabolism in cancer;Glycine, serine and threonine metabolism;Cysteine and methionine metabolism;Other | 1.03 | 0.854 | 0.97 | 0.848 | 0.92 | 0.945 | 1.04 | 0.807171 |
| 49 | Serine | [M+H]+1 | POS | Mass, RT, Spectral match | C18 significant | 105.043 | 0.500 | C00065 | HMDB0000187 | 5951 | Metabolic pathways;Biosynthesis of secondary metabolites;Biosynthesis of antibiotics;Microbial metabolism in diverse environments;Biosynthesis of plant secondary metabolites;Biosynthesis of amino acids;ABC transporters;Protein digestion and absorption;Central carbon metabolism in cancer;Aminoacyl-tRNA biosynthesis;Other | 1.34 | 0.043 | 1.09 | 0.589 | 1.10 | 0.397 | 0.98 | 0.943353 |
| 442 | Ethyl__D_glucuronide | [M+H]+1 | POS | Mass, Spectral match | Hilic | 222.073 | 9.510 | | HMDB0010316 | 83944 | | 0.90 | 0.694 | 1.61 | 0.060 | 1.14 | 0.292 | 1.41 | 0.084775 |
| 51 | Benzaldehyde | [M+H]+1 | POS | Mass, Spectral match | Hilic | 106.041 | 2.608 | C00193 | HMDB0006115 | 240 | Microbial metabolism in diverse environments;Degradation of aromatic compounds;Other | 1.06 | 0.829 | 1.14 | 0.505 | 1.06 | 0.599 | 1.07 | 0.689475 |
| 52 | Cyclopentanone | [M-H]-1 | NEG | Mass, RT, Spectral match | C18 | 107.047 | 18.057 | C00557 | HMDB0031407 | NA | | 1.05 | 0.785 | 1.32 | 0.210 | 1.14 | 0.742 | 1.15 | 0.523641 |
| 53 | Cyclopentanone | [M-H]-1 | NEG | Mass, Spectral match | Hilic | 107.047 | 5.740 | C00557 | HMDB0031407 | NA | | 0.74 | 0.383 | 0.72 | 0.370 | 0.56 | 0.079 | 1.28 | 0.480849 |
| 54 | o_Toluidine | [M+H]+1 | POS | Mass, RT, Spectral match | C18 | 107.074 | 18.113 | C14403 | HMDB41965 | 7242 | | 0.91 | 0.112 | 1.00 | 0.966 | 0.90 | 0.085 | 1.12 | 0.246552 |
| 55 | Hypotaurine | [M+H]+1 | POS | Mass, Spectral match | Hilic | 109.020 | 9.202 | C00519 | HMDB0000965 | 107812 | Metabolic pathways;Other | 0.95 | 0.890 | 0.85 | 0.720 | 1.01 | 0.347 | 0.85 | 0.803454 |

| | | | | | | | | | | | | | | | | | | | |
|---|---|---|---|---|---|---|---|---|---|---|---|---|---|---|---|---|---|---|---|
| 56 | Quinone | [M+H]+1 | POS | Mass, Spectral match | Hilic | 109.028 | 2.089 | C00472 | HMDB0003364 | 4650 | | 2.81 | 0.304 | 2.07 | 0.327 | 2.49 | 0.155 | 0.83 | 0.800166 |
| 57 | Prenol | [2M+H]+1 | POS | Mass, Spectral match | Hilic | 109.063 | 1.940 | C01390 | HMDB0030124 | 11173 | | 1.18 | 0.642 | 1.20 | 0.621 | 0.75 | 0.827 | 1.58 | 0.331495 |
| 58 | Benzyl_alcohol | [M+H]+1 | POS | Mass, Spectral match | C18 | 109.064 | 17.931 | C00556 | HMDB0003119 | 244 | | 0.56 | 0.140 | 0.60 | 0.082 | 0.77 | 0.790 | 0.78 | 0.718176 |
| 59 | Benzyl_alcohol | [M+H]+1 | POS | Mass, Spectral match | Hilic | 109.064 | 1.822 | C01279 | METPA0166 | - | | 1.40 | 0.232 | 1.09 | 0.723 | 0.95 | 0.775 | 1.15 | 0.772563 |
| 60 | Pyrocatechol | [M+H]+1 | POS | Mass, Spectral match | C18 | 111.043 | 17.687 | C00090 | HMDB0000957 | 289 | Metabolic pathways;Other | 1.71 | 0.391 | 0.81 | 0.764 | 1.40 | 0.505 | 0.58 | 0.584504 |
| 61 | Cytosine | [M+H]+1 | POS | Mass, Spectral match | Hilic | 111.043 | 1.942 | C00380 | HMDB0000630 | 597 | Metabolic pathways;Pyrimidine metabolism | 0.86 | 0.510 | 0.71 | 0.023 | 0.46 | 0.217 | 1.53 | 0.047645 |
| 62 | histamine | [M+H]+1 | POS | Mass,RT, Spectral match | Hilic significant | 111.080 | 4.817 | C00388 | HMDB0000870 | 774 | Metabolic pathways;Biosynthesis of secondary metabolites;Protein digestion and absorption;Histidine metabolism;Other | 1.33 | 0.207 | 1.74 | 0.036 | 1.32 | 0.949 | 1.32 | 0.256821 |
| 63 | Histamine | [M+H]+1 | POS | Mass, Spectral match | C18 significant | 111.080 | 16.850 | C00388 | HMDB0000870 | 774 | Metabolic pathways;Biosynthesis of secondary metabolites;Protein digestion and absorption;Neuroactive ligand-receptor interaction;Other | 0.51 | 0.067 | 0.46 | 0.033 | 0.86 | 0.883 | 0.54 | 0.025824 |
| 64 | Uracil | [M+H]+1 | POS | Mass,RT, Spectral match | Hilic | 112.027 | 2.036 | C00106 | HMDB0000300 | 1174 | Metabolic pathways;Pyrimidine metabolism;Other | 1.30 | 0.193 | 1.09 | 0.492 | 1.38 | 0.123 | 0.79 | 0.329217 |
| 65 | Uracil | [M+H]+1 | POS | Mass, Spectral match | C18 | 112.027 | 0.579 | C00106 | HMDB0000300 | 1174 | Metabolic pathways;Other | 0.82 | 0.368 | 0.92 | 0.575 | 0.90 | 0.938 | 1.02 | 0.718414 |
| 66 | Arenediol | [M+H]+1 | POS | Mass, Spectral match | C18 | 112.053 | 8.829 | C01066 | #N/A | 10434 | Microbial metabolism in diverse environments;Degradation of aromatic compounds;Other | 0.95 | 0.890 | 0.71 | 0.380 | 0.66 | 0.389 | 1.09 | 0.559562 |
| 67 | 13_Diaminopropane | [M+H]+1 | POS | Mass, Spectral match | C18 | 113.048 | 0.556 | C00986 | HMDB0000002 | 428 | Metabolic pathways;Biosynthesis of secondary metabolites;Biosynthesis of antibiotics;Biosynthesis of amino acids;Arginine and proline metabolism;Other | 0.63 | 0.040 | 0.91 | 0.654 | 0.73 | 0.526 | 1.25 | 0.256735 |
| 68 | S_1_Pyrroline_5_carboxylate | [M+H]+1 | POS | Mass, Spectral match | Hilic | 113.048 | 3.400 | C04322 | HMDB0001301 | 1196 | Metabolic pathways;Biosynthesis of secondary metabolites;Biosynthesis of antibiotics;Biosynthesis of amino acids;Arginine and proline metabolism;Other | 0.40 | 0.022 | 0.90 | 0.779 | 0.81 | 0.827 | 1.12 | 0.872492 |
| 69 | Creatinine | [M+H]+1 | POS | Mass, Spectral match | Hilic significant | 113.059 | 1.203 | C00791 | HMDB0000562 | 588 | Metabolic pathways;Arginine and proline metabolism | 1.30 | 0.506 | 2.10 | 0.111 | 2.59 | 0.005 | 0.81 | 0.404539 |
| 70 | Trimethylamine_N_oxide | [M-H]-1 | NEG | Mass, Spectral match | C18 | 114.030 | 0.527 | C01104 | HMDB0000925 | 1145 | | 1.49 | 0.022 | 1.49 | 0.132 | 1.66 | 0.056 | 0.89 | 0.604317 |

| | | | | | | | | | | | | | | | | | | | |
|---|---|---|---|---|---|---|---|---|---|---|---|---|---|---|---|---|---|---|---|
| 71 | Dihydrouracil | [M+H]+1 | POS | Mass,RT, Spectral match | Hilic | 114.043 | 3.974 | C00429 | HMDB0000076 | 649 | Metabolic pathways;Biosynthesis of secondary metabolites;Cyanoamino acid metabolism;Arginine and proline metabolism;Pyrimidine metabolism;Other | 0.87 | 0.762 | 0.97 | 0.932 | 1.18 | 0.192 | 0.82 | 0.294897 |
| 72 | 3_Hydroxycyclohexanone | [M+H]+1 | POS | Mass, Spectral match | Hilic | 114.068 | 2.268 | C01147 | #N/A | 10785 | Microbial metabolism in diverse environments;Degradation of aromatic compounds;Other | 0.97 | 0.854 | 1.23 | 0.554 | 1.14 | 0.364 | 1.08 | 0.924979 |
| 73 | Prolinamide | [M+H]+1 | POS | Mass, Spectral match | Hilic | 114.079 | 2.784 | NA | HMDB0005777 | 119303 | | 0.85 | 0.732 | 1.29 | 0.484 | 1.14 | 0.403 | 1.13 | 0.633937 |
| 74 | Prolinamide | [M+H]+1 | POS | Mass, Spectral match | C18 | 114.079 | 0.446 | C19781 | #N/A | 111306 | | 0.97 | 0.834 | 0.87 | 0.469 | 0.99 | 0.804 | 0.88 | 0.455554 |
| 75 | Maleamate | [M+H]+1 | POS | Mass, Spectral match | Hilic | 115.027 | 8.479 | C01596 | METPA0190 | NA | Microbial metabolism in diverse environments;Other | 1.32 | 0.173 | 0.98 | 0.941 | 1.19 | 0.469 | 0.83 | 0.304358 |
| 76 | Glycerol | [M-H]-1 | NEG | Mass, Spectral match | Hilic | 115.038 | 2.439 | C00116 | HMDB0000131 | 753 | | 0.94 | 0.877 | 1.10 | 0.796 | 0.80 | 0.456 | 1.38 | 0.411534 |
| 77 | 2_Hydroxy_24_pentadienoate | [M+H]+1 | POS | Mass, Spectral match | Hilic | 115.038 | 2.451 | C00596 | METPA0060 | NA | | 0.92 | 0.766 | 1.08 | 0.769 | 0.89 | 0.656 | 1.22 | 0.214513 |
| 78 | L_Proline | [M+H]+1 | POS | Mass, Spectral match | C18 | 115.063 | 0.515 | C00148 | HMDB0000162 | 145742 | Metabolic pathways;Biosynthesis of secondary metabolites;Biosynthesis of antibiotics;Biosynthesis of amino acids;ABC transporters;Protein digestion and absorption;Arginine and proline metabolism;Central carbon metabolism in cancer;Aminoacyl-tRNA biosynthesis;Other | 1.04 | 0.836 | 0.83 | 0.353 | 0.88 | 0.858 | 0.95 | 0.673894 |
| 79 | Proline | [M+H]+1 | POS | Mass,RT, Spectral match | Hilic | 115.063 | 4.386 | C00148 | HMDB0000162 | 145742 | Metabolic pathways;Biosynthesis of secondary metabolites;Biosynthesis of antibiotics;Biosynthesis of amino acids;ABC transporters;Protein digestion and absorption;Arginine and proline metabolism;Aminoacyl-tRNA biosynthesis;Central carbon metabolism in cancer;Other | 0.98 | 0.916 | 0.95 | 0.806 | 0.90 | 0.928 | 1.05 | 0.738751 |
| 80 | Fumaric_acid | [M-H]-1 | NEG | Mass,RT, Spectral match | Hilic | 116.010 | 8.532 | C00122 | HMDB0000134 | 444972 | Metabolic pathways;Biosynthesis of secondary metabolites;Microbial metabolism in diverse environments;Biosynthesis of antibiotics;Biosynthesis of plant secondary metabolites;Central carbon metabolism in cancer;Degradation of aromatic compounds;Other | 1.81 | 0.108 | 2.65 | 0.091 | 4.64 | 0.066 | 0.57 | 0.418509 |

| | | | | | | | | | | | | | | | | | | | |
|---|---|---|---|---|---|---|---|---|---|---|---|---|---|---|---|---|---|---|---|
| 453 | Deoxyuridine | [M+H]+1 | POS | Mass, Spectral match | Hilic | 228.074 | 1.558 | C00526 | HMDB0000012 | 13712 | Pyrimidine metabolism | 0.69 | 0.480 | 0.93 | 0.879 | 0.53 | 0.154 | 1.77 | 0.202882 |
| 82 | Guanidineaceticacid | [M+H]+1 | POS | Mass, Spectral match | Hilic | 117.054 | 2.337 | C00581 | HMDB0000128 | 763 | Metabolic pathways;Arginine and proline metabolism;Glycine, serine and threonine metabolism | 1.30 | 0.317 | 1.25 | 0.536 | 1.07 | 0.350 | 1.16 | 0.915241 |
| 83 | Indole | [M+H]+1 | POS | Mass, Spectral match | C18 | 117.058 | 1.651 | C00463 | HMDB0000738 | 798 | Metabolic pathways;Biosynthesis of secondary metabolites;Microbial metabolism in diverse environments;Protein digestion and absorption;Other | 1.34 | 0.235 | 1.09 | 0.719 | 1.12 | 0.737 | 0.97 | 0.911773 |
| 84 | Valine | [M-H]-1 | NEG | Mass, Spectral match | C18 significant | 117.079 | 18.510 | C00183 | HMDB0000883 | 6287 | Metabolic pathways;Biosynthesis of secondary metabolites;Biosynthesis of antibiotics;Microbial metabolism in diverse environments;Biosynthesis of plant secondary metabolites;Biosynthesis of amino acids;ABC transporters;Protein digestion and absorption;Arginine and proline metabolism;Central carbon metabolism in cancer;2-Oxocarboxylic acid metabolism;Biosynthesis of alkaloids derived from shikimate pathway;Aminoacyl-tRNA biosynthesis;Other | 0.81 | 0.177 | 0.78 | 0.192 | 0.76 | 0.135 | 1.03 | 0.946353 |
| 85 | Betaine | [M+H]+1 | POS | Mass,RT, Spectral match | Hilic | 117.079 | 2.912 | C00183 | HMDB0000883 | 6287 | Metabolic pathways;Biosynthesis of secondary metabolites;Microbial metabolism in diverse environments;Biosynthesis of antibiotics;Biosynthesis of amino acids;Biosynthesis of plant secondary metabolites;ABC transporters;Protein digestion and absorption;Cyanoamino acid metabolism;Arginine and proline metabolism;2-Oxocarboxylic acid metabolism;Aminoacyl-tRNA biosynthesis;Central carbon metabolism in cancer;Glycine, serine and threonine metabolism;Lysine degradation;Other | 1.68 | 0.229 | 1.28 | 0.640 | 2.89 | 0.068 | 0.44 | 0.158031 |
| 86 | L_Valine | [M+H]+1 | POS | Mass, Spectral match | C18 | 117.079 | 0.536 | C00183 | HMDB0000883 | 6287 | Metabolic pathways;Biosynthesis of secondary metabolites;Biosynthesis of antibiotics;Microbial metabolism in diverse environments;Biosynthesis of | 0.86 | 0.337 | 0.94 | 0.645 | 1.55 | 0.044 | 0.60 | 0.011902 |

| | | | | | | | | | | | | | | | | | | | |
|---|---|---|---|---|---|---|---|---|---|---|---|---|---|---|---|---|---|---|---|
| | | | | | | | | | | | plant secondary metabolites;Biosynthesis of amino acids;ABC transporters;Protein digestion and absorption;Arginine and proline metabolism;Central carbon metabolism in cancer;2-Oxocarboxylic acid metabolism;Biosynthesis of alkaloids derived from shikimate pathway;Aminoacyl-tRNA biosynthesis;Other | | | | | | | | |
| 87 | Succinic_acid | [M-H]-1 | NEG | Mass, RT, Spectral match | C18 | 118.025 | 0.672 | C00042 | HMDB0000254 | 1110 | | 3.66 | 0.231 | 1.67 | 0.270 | 3.22 | 0.268 | 0.52 | 0.755879 |
| 88 | Guanidoacetic_acid | [M-H]-1 | NEG | Mass, RT, Spectral match | C18 | 118.062 | 1.692 | C00581 | HMDB0000128 | 763 | | 1.72 | 0.178 | 1.10 | 0.649 | 1.35 | 0.228 | 0.82 | 0.69052 |
| 89 | L_Threonine | [M+H]+1 | POS | Mass, Spectral match | Hilic | 119.058 | 7.231 | C00188 | HMDB0000167 | 6288 | Metabolic pathways;Biosynthesis of secondary metabolites;Microbial metabolism in diverse environments;Biosynthesis of antibiotics;Biosynthesis of amino acids;Biosynthesis of plant secondary metabolites;ABC transporters;Protein digestion and absorption;Aminoacyl-tRNA biosynthesis;Glycine, serine and threonine metabolism;Cysteine and methionine metabolism;Other | 1.09 | 0.631 | 0.95 | 0.794 | 0.89 | 0.959 | 1.07 | 0.745758 |
| 90 | L_Threonine | [M+H]+1 | POS | Mass, RT, Spectral match | C18 | 119.058 | 0.509 | C00188 | HMDB0000167 | 6288 | Metabolic pathways;Biosynthesis of secondary metabolites;Biosynthesis of antibiotics;Microbial metabolism in diverse environments;Biosynthesis of plant secondary metabolites;Biosynthesis of amino acids;ABC transporters;Protein digestion and absorption;Aminoacyl-tRNA biosynthesis;Other | 1.02 | 0.849 | 0.86 | 0.302 | 1.07 | 0.341 | 0.81 | 0.226778 |
| 456 | Amiloride | [M+H]+1 | POS | Mass,RT, Spectral match | Hilic | 229.047 | 0.690 | C06821 | HMDB0014732 | 16231 | | 1.03 | 0.840 | 0.98 | 0.855 | 1.06 | 0.908 | 0.93 | 0.623178 |
| 92 | Phenylacetaldehyde | [M+H]+1 | POS | Mass, Spectral match | Hilic | 120.057 | 0.708 | C00601 | HMDB0006236 | 998 | Metabolic pathways;Microbial metabolism in diverse environments;Degradation of aromatic compounds;Other | 0.84 | 0.327 | 0.98 | 0.909 | 0.98 | 0.275 | 0.99 | 0.760553 |
| 93 | Phenylacetaldehyde | [M+H]+1 | POS | Mass, RT, Spectral match | C18 | 120.058 | 0.554 | C00601 | HMDB0006236 | 998 | Metabolic pathways;Microbial metabolism in diverse environments;Degradation of aromatic compounds;Other | 1.51 | 0.320 | 1.24 | 0.538 | 1.38 | 0.101 | 0.90 | 0.812038 |
| 94 | Acetophenone | [M+H]+1 | POS | Mass, RT, Spectral match | C18 | 120.058 | 11.949 | C07113 | HMDB0033910 | 7410 | Metabolic pathways;Microbial metabolism in diverse environments;Degradation of aromatic compounds;Other | 0.97 | 0.532 | 0.94 | 0.407 | 0.88 | 0.777 | 1.07 | 0.441525 |

| | | | | | | | | | | | | | | | | | | | |
|---|---|---|---|---|---|---|---|---|---|---|---|---|---|---|---|---|---|---|---|
| 95 | Benzoate | [M+H]+1 | POS | Mass, Spectral match | Hilic | 121.030 | 1.148 | C00180 | HMDB01870 | 243 | | 0.56 | 0.080 | 0.59 | 0.106 | 0.79 | 0.699 | 0.74 | 0.780411 |
| 96 | Cyclohexanone | [M+H]+1 | POS | Mass, Spectral match | Hilic | 121.063 | 7.268 | C00414 | HMDB0003315 | 7967 | | 1.14 | 0.491 | 0.98 | 0.919 | 1.00 | 0.805 | 0.98 | 0.963027 |
| 97 | NN_Dimethylaniline | [M+H]+1 | POS | Mass, Spectral match | Hilic | 121.089 | 6.992 | C02846 | HMDB0001020 | 949 | Other | 0.50 | 0.187 | 0.47 | 0.167 | 0.43 | 0.312 | 1.09 | 0.12045 |
| 98 | NN_Dimethylaniline | [M+H]+1 | POS | Mass, RT, Spectral match | C18 significant | 121.090 | 17.830 | C02846 | HMDB0001020 | 949 | Other | 0.70 | 0.338 | 0.35 | 0.041 | 0.77 | 0.707 | 0.45 | 0.085677 |
| 99 | 4_Hydroxybenzaldehyde | [M+H]+1 | POS | Mass,RT, Spectral match | Hilic significant | 122.037 | 3.984 | C00633 | HMDB0011718 | 126 | Metabolic pathways;Biosynthesis of secondary metabolites;Microbial metabolism in diverse environments;Degradation of aromatic compounds;Other | 0.10 | 0.036 | 0.24 | 0.098 | 0.07 | 0.098 | 3.64 | 0.867842 |
| 100 | Nicotinamide | [M+H]+1 | POS | Mass,RT, Spectral match | Hilic | 122.048 | 1.164 | C00153 | HMDB0001406 | 936 | Metabolic pathways;Other | 1.45 | 0.222 | 0.92 | 0.703 | 1.32 | 0.138 | 0.69 | 0.345 |
| 101 | Niacinamide | [M+H]+1 | POS | Mass, RT, Spectral match | C18 | 122.048 | 0.602 | C00153 | HMDB0001406 | 936 | Metabolic pathways;Other | 1.49 | 0.230 | 1.31 | 0.320 | 1.44 | 0.100 | 0.91 | 0.690975 |
| 102 | Nicotinicacid | [M+H]+1 | POS | Mass, Spectral match | Hilic | 123.032 | 10.112 | C00253 | HMDB0001488 | 938 | Metabolic pathways;Biosynthesis of secondary metabolites;Microbial metabolism in diverse environments;Biosynthesis of plant secondary metabolites;Degradation of aromatic compounds;Other | 1.17 | 0.318 | 1.48 | 0.167 | 1.20 | 0.145 | 1.24 | 0.478559 |
| 103 | 1_Hexanol | [M+H]+1 | POS | Mass, Spectral match | Hilic | 123.078 | 1.183 | C00854 | HMDB0012971 | 8103 | | 0.91 | 0.286 | 0.93 | 0.441 | 0.89 | 0.791 | 1.04 | 0.758297 |
| 104 | 1_Hexanol | [M+H]+1 | POS | Mass, Spectral match | C18 | 123.078 | 0.494 | C00854 | HMDB0012971 | 8103 | | 1.13 | 0.290 | 1.07 | 0.606 | 1.00 | 0.666 | 1.07 | 0.613954 |
| 105 | 5_Hydroxypyrazinamide | [M-H]-1 | NEG | Mass, Spectral match | Hilic | 124.051 | 1.086 | C01956 | HMDB0001978 | 151823 | | 0.98 | 0.694 | 0.95 | 0.474 | 0.95 | 0.430 | 1.00 | 0.975824 |
| 106 | Methylimidazole_acetaldehyde | [M+H]+1 | POS | Mass, RT, Spectral match | C18 | 124.064 | 16.176 | C05827 | HMDB0004181 | 193545 | Other | 0.80 | 0.527 | 0.32 | 0.059 | 0.69 | 0.320 | 0.46 | 0.201628 |
| 107 | Taurine | [M+H]+1 | POS | Mass, Spectral match | Hilic | 125.015 | 8.144 | C00245 | HMDB0000251 | 1123 | Metabolic pathways;ABC transporters;Other | 1.01 | 0.967 | 0.75 | 0.155 | 0.64 | 0.737 | 1.16 | 0.750414 |
| 108 | Taurine | [M+H]+1 | POS | Mass, Spectral match | C18 significant | 125.015 | 0.500 | C00245 | HMDB0000251 | 1123 | Metabolic pathways;ABC transporters;Neuroactive ligand-receptor interaction;Other | 1.07 | 0.809 | 0.55 | 0.030 | 0.41 | 0.270 | 1.33 | 0.703422 |
| 109 | 1_methylhistamine | [M+H]+1 | POS | Mass, Spectral match | Hilic significant | 125.096 | 0.394 | C05127 | HMDB0000898 | 3614 | Histidine metabolism;Other | 0.37 | 0.104 | 0.20 | 0.045 | 0.30 | 0.217 | 0.68 | 0.119262 |
| 110 | 1_methylhistamine | [M+H]+1 | POS | Mass, Spectral match | C18 significant | 125.096 | 16.850 | C05127 | HMDB0000898 | 3614 | Other | 0.59 | 0.199 | 0.66 | 0.198 | 1.11 | 0.612 | 0.59 | 0.034964 |
| 111 | Sulcatone | [M+H]+1 | POS | Mass, Spectral match | C18 | 126.104 | 4.938 | C07287 | HMDB0035915 | 9862 | Biosynthesis of antibiotics;Other | 0.80 | 0.619 | 0.97 | 0.949 | 0.95 | 0.555 | 1.03 | 0.872077 |

| | | | | | | | | | | | | | | | | | | | |
|---|---|---|---|---|---|---|---|---|---|---|---|---|---|---|---|---|---|---|---|
| 112 | Urea_1_carboxylate | [M+H]+1 | POS | Mass, Spectral match | C18 | 127.012 | 0.434 | C01010 | METPA0111 | NA | | 0.98 | 0.817 | 0.93 | 0.571 | 1.28 | 0.131 | 0.73 | 0.041207 |
| 113 | S_2_Propylpiperidine | [M+H]+1 | POS | Mass, Spectral match | C18 | 127.136 | 18.166 | C06523 | HMDB0030285 | 9985 | Biosynthesis of plant secondary metabolites;Other | 0.90 | 0.332 | 0.84 | 0.177 | 0.96 | 0.844 | 0.88 | 0.274915 |
| 114 | 4E_2_Oxohexenoicacid | [M+H]+1 | POS | Mass, Spectral match | Hilic | 128.047 | 5.943 | C01125 | #N/A | 39 | Metabolic pathways;Microbial metabolism in diverse environments;Degradation of aromatic compounds;Other | 1.46 | 0.257 | 1.36 | 0.403 | 1.37 | 0.291 | 0.99 | 0.993436 |
| 115 | Dihydrothymine | [M-H]-1 | NEG | Mass, Spectral match | Hilic | 128.057 | 7.837 | C00906 | HMDB0000079 | 93556 | | 0.83 | 0.187 | 1.06 | 0.554 | 0.99 | 0.495 | 1.07 | 0.586134 |
| 116 | 4_Chlorophenol | [M+H]+1 | NEG | Mass, Spectral match | C18 | 129.009 | 0.438 | C02124 | METPA0251 | NA | | 0.98 | 0.835 | 0.94 | 0.610 | 1.25 | 0.138 | 0.75 | 0.053751 |
| 117 | 4_Oxoproline | [M+H]+1 | POS | Mass, Spectral match | Hilic | 129.042 | 7.933 | C01877 | METPA0228 | NA | Metabolic pathways;Arginine and proline metabolism;Other | 1.20 | 0.283 | 1.16 | 0.437 | 1.12 | 0.430 | 1.04 | 0.849458 |
| 118 | L_Pyroglutamic_acid_oxoproline | [M+H]+1 | NEG | Mass, Spectral match | C18 significant | 129.043 | 0.510 | C01877 | METPA0228 | NA | Metabolic pathways;Arginine and proline metabolism;Other | 1.35 | 0.037 | 1.14 | 0.474 | 1.19 | 0.260 | 0.96 | 0.792897 |
| 119 | Pipecolic_acid | [M+H]+1 | POS | Mass, Spectral match | C18 | 129.079 | 0.462 | C00408 | HMDB0000070 | 849 | Metabolic pathways;Biosynthesis of secondary metabolites;Biosynthesis of plant secondary metabolites;Arginine and proline metabolism;Tropane, piperidine and pyridine alkaloid biosynthesis;Biosynthesis of alkaloids derived from ornithine, lysine and nicotinic acid;Other | 0.91 | 0.598 | 0.83 | 0.377 | 0.89 | 0.908 | 0.93 | 0.616951 |
| 120 | Pipecolinic_acid_ | [M+H]+1 | POS | Mass, Spectral match | Hilic significant | 129.079 | 1.250 | C00408 | HMDB00070 | 439227 | Metabolic pathways;Biosynthesis of secondary metabolites;Biosynthesis of plant secondary metabolites;Arginine and proline metabolism;Lysine degradation;Other | 1.63 | 0.049 | 1.30 | 0.096 | 1.77 | 0.016 | 0.73 | 0.115254 |
| 121 | Octanal | [M+H]+1 | POS | Mass, Spectral match | Hilic | 129.126 | 9.776 | C01545 | HMDB0001140 | 454 | | 1.00 | 0.991 | 0.89 | 0.629 | 0.99 | 0.744 | 0.89 | 0.521299 |
| 122 | 2_Oxoglutarate_semialdehyde | [M-H]-1 | NEG | Mass, Spectral match | Hilic | 130.025 | 2.633 | C00433 | #N/A | 523 | | 7.73 | 0.128 | 1.68 | 0.487 | 5.04 | 0.283 | 0.33 | 0.537822 |
| 123 | N_Acetylputrescine_ | [M+H]+1 | POS | Mass, Spectral match | C18 significant | 130.111 | 18.210 | C02714 | HMDB0002064 | 122356 | Metabolic pathways;Arginine and proline metabolism | 0.93 | 0.353 | 0.92 | 0.325 | 0.84 | 0.126 | 1.09 | 0.318005 |
| 485 | Cytidine | [M+H]+1 | POS | Mass,RT, Spectral match | Hilic significant | 243.086 | 2.002 | C00475 | HMDB0000089 | 6175 | Metabolic pathways;Cyanoamino acid metabolism;Pyrimidine metabolism | 78.54 | 0.017 | 58.36 | 0.008 | 35.63 | 0.167 | 1.64 | 0.53688 |
| 125 | 5_Aminolevulinic_acid | [M+H]+1 | POS | Mass,RT, Spectral match | Hilic | 131.058 | 7.388 | C00430 | HMDB0001149 | 137 | Metabolic pathways;Biosynthesis of secondary metabolites;Microbial metabolism in diverse environments;Biosynthesis of | 0.92 | 0.822 | 1.00 | 0.996 | 1.22 | 0.176 | 0.82 | 0.649305 |

| | | | | | | | | | | | | | | | | | | | |
|---|---|---|---|---|---|---|---|---|---|---|---|---|---|---|---|---|---|---|---|
| | | | | | | | | | | | antibiotics;Biosynthesis of amino acids;Biosynthesis of plant secondary metabolites;ABC transporters;Arginine and proline metabolism;Glycine, serine and threonine metabolism;Other | | | | | | | | |
| 126 | cis_4_hydroxy_D_proline | [M+H]+1 | POS | Mass,RT, Spectral match | Hilic significant | 131.059 | 1.685 | C01015 | HMDB0006055 | 69248 | Metabolic pathways;Biosynthesis of secondary metabolites;Microbial metabolism in diverse environments;Biosynthesis of antibiotics;Biosynthesis of amino acids;Biosynthesis of plant secondary metabolites;ABC transporters;Arginine and proline metabolism;Glycine, serine and threonine metabolism;Other | 0.55 | 0.018 | 0.84 | 0.521 | 0.79 | 0.883 | 1.05 | 0.878639 |
| 127 | Ketoleucine | [M+H]+1 | POS | Mass, Spectral match | C18 | 131.069 | 0.507 | C00233 | HMDB0000695 | 70 | Metabolic pathways;Arginine and proline metabolism;Other | 1.42 | 0.297 | 1.47 | 0.231 | 1.38 | 0.121 | 1.06 | 0.748404 |
| 128 | Creatine | [M+H]+1 | POS | Mass,RT, Spectral match | Hilic | 131.069 | 1.675 | C00300 | HMDB0000064 | 586 | Metabolic pathways;Arginine and proline metabolism;Glycine, serine and threonine metabolism | 0.57 | 0.215 | 1.15 | 0.754 | 1.08 | 0.321 | 1.07 | 0.976605 |
| 129 | 3_Methylindole | [M+H]+1 | POS | Mass, Spectral match | Hilic | 131.073 | 2.554 | C08313 | HMDB0000466 | 6736 | | 1.14 | 0.521 | 1.17 | 0.473 | 1.22 | 0.482 | 0.95 | 0.909681 |
| 130 | DL_Leucine | [M-H]-1 | NEG | Mass, Spectral match | Hilic | 131.093 | 2.272 | C00123 | HMDB0000687 | 6106 | | 0.83 | 0.496 | 0.94 | 0.689 | 0.94 | 0.887 | 0.99 | 0.981567 |
| 131 | L_Norleucine | [M+H]+1 | POS | Mass,RT, Spectral match | Hilic | 131.095 | 2.280 | C01933 | HMDB0001645 | 21236 | Metabolic pathways;Biosynthesis of secondary metabolites;Microbial metabolism in diverse environments;Biosynthesis of antibiotics;Biosynthesis of amino acids;Biosynthesis of plant secondary metabolites;ABC transporters;Protein digestion and absorption;Cyanoamino acid metabolism;2-Oxocarboxylic acid metabolism;Aminoacyl-tRNA biosynthesis;Central carbon metabolism in cancer;Other | 0.87 | 0.180 | 1.27 | 0.517 | 1.22 | 0.251 | 1.04 | 0.840966 |
| 132 | L_Leucine | [M+H]+1 | POS | Mass, RT, Spectral match | C18 | 131.095 | 0.745 | C00123 | HMDB0000687 | 6106 | Metabolic pathways;Biosynthesis of secondary metabolites;Biosynthesis of antibiotics;Microbial metabolism in diverse environments;Biosynthesis of plant secondary metabolites;Biosynthesis of amino acids;ABC transporters;Protein digestion | 1.07 | 0.653 | 0.91 | 0.536 | 0.97 | 0.770 | 0.93 | 0.688524 |

| | | | | | | | | | | | | | | | | | | | |
|---|---|---|---|---|---|---|---|---|---|---|---|---|---|---|---|---|---|---|---|
| | | | | | | | | | | | and absorption;Central carbon metabolism in cancer;2-Oxocarboxylic acid metabolism;Aminoacyl-tRNA biosynthesis;Tropane, piperidine and pyridine alkaloid biosynthesis;Biosynthesis of alkaloids derived from ornithine, lysine and nicotinic acid;Other | | | | | | | | |
| 133 | N_ACETYLputrescine | [M+H]+1 | POS | Mass, Spectral match | Hilic | 131.106 | 6.918 | C02714 | HMDB0002064 | 122356 | Metabolic pathways;Arginine and proline metabolism | 0.96 | 0.908 | 1.72 | 0.256 | 1.30 | 0.545 | 1.32 | 0.568659 |
| 134 | Glutaric_acid | [M+H]+1 | POS | Mass, RT, Spectral match | C18 | 132.041 | 0.827 | C00489 | HMDB0000661 | 743 | | 2.17 | 0.253 | 1.73 | 0.079 | 2.13 | 0.059 | 0.81 | 0.561008 |
| 135 | Ureidoglycine | [M+H]+1 | POS | Mass, Spectral match | Hilic | 132.042 | 9.846 | C02091 | METPA0247 | NA | Metabolic pathways;Biosynthesis of secondary metabolites;Microbial metabolism in diverse environments;Biosynthesis of antibiotics;Biosynthesis of amino acids;2-Oxocarboxylic acid metabolism;Degradation of aromatic compounds;Lysine degradation;Other | 1.39 | 0.444 | 1.12 | 0.802 | 1.24 | 0.147 | 0.90 | 0.723802 |
| 136 | Glycylglycine | [M+Cl]-1 | POS | Mass, Spectral match | Hilic | 132.052 | 8.483 | C02037 | HMDB0011733 | 11163 | | 1.15 | 0.616 | 0.81 | 0.563 | 0.75 | 0.552 | 1.08 | 0.689864 |
| 137 | Gly_Gly | [M+Cl]-1 | POS | Mass, Spectral match | Hilic significant | 132.053 | 2.910 | C02037 | HMDB0011733 | 11163 | | 15.17 | 0.008 | 4.75 | 0.433 | 11.61 | 0.096 | 0.41 | 0.132147 |
| 138 | L_Asparagine | [M+H]+1 | POS | Mass, Spectral match | C18 | 132.053 | 0.539 | C00152 | HMDB0000168 | 6267 | Metabolic pathways;Biosynthesis of secondary metabolites;Biosynthesis of plant secondary metabolites;Biosynthesis of amino acids;Protein digestion and absorption;Arginine and proline metabolism;Central carbon metabolism in cancer;Aminoacyl-tRNA biosynthesis;Other | 1.13 | 0.568 | 0.81 | 0.426 | 1.07 | 0.241 | 0.76 | 0.151879 |
| 139 | Asparagine | [M+H]+1 | POS | Mass,RT, Spectral match | Hilic | 132.053 | 8.479 | C00152 | HMDB0000168 | 6267 | Metabolic pathways;Biosynthesis of secondary metabolites;Biosynthesis of amino acids;Biosynthesis of plant secondary metabolites;Protein digestion and absorption;Cyanoamino acid metabolism;Arginine and proline metabolism;Aminoacyl-tRNA biosynthesis;Central carbon metabolism in cancer;Pyrimidine metabolism;Other | 1.32 | 0.176 | 0.98 | 0.932 | 1.18 | 0.480 | 0.83 | 0.30872 |

| | | | | | | | | | | | | | | | | | | | |
|---|---|---|---|---|---|---|---|---|---|---|---|---|---|---|---|---|---|---|---|
| 140 | D_Asparagine | [M+H]+1 | POS | Mass,RT, Spectral match | Hilic | 132.053 | 7.211 | C01905 | HMDB0033780 | 439600 | Metabolic pathways;Biosynthesis of secondary metabolites;Biosynthesis of amino acids;Biosynthesis of plant secondary metabolites;Protein digestion and absorption;Cyanoamino acid metabolism;Arginine and proline metabolism;Aminoacyl-tRNA biosynthesis;Central carbon metabolism in cancer;Pyrimidine metabolism;Other | 0.90 | 0.571 | 0.74 | 0.182 | 0.90 | 0.935 | 0.83 | 0.198209 |
| 141 | 2_Hydroxycaproicacid | [M-H]-1 | NEG | Mass, Spectral match | Hilic | 132.077 | 3.329 | C06103 | HMDB0001624 | 99824 | | 1.64 | 0.218 | 1.40 | 0.346 | 1.25 | 0.111 | 1.12 | 0.858031 |
| 142 | Ornithine | [M+H]+1 | POS | Mass, Spectral match | C18 | 132.090 | 0.447 | C00077 | HMDB0000214 | 6262 | Metabolic pathways;Biosynthesis of secondary metabolites;Biosynthesis of antibiotics;Biosynthesis of plant secondary metabolites;Biosynthesis of amino acids;ABC transporters;Arginine and proline metabolism;2-Oxocarboxylic acid metabolism;Biosynthesis of alkaloids derived from ornithine, lysine and nicotinic acid;Other | 0.72 | 0.352 | 0.62 | 0.142 | 0.83 | 0.898 | 0.74 | 0.476038 |
| 143 | 2R_4S_2_4_Diaminopentanoate | [M+H]+1 | POS | Mass,RT, Spectral match | Hilic | 132.090 | 1.238 | C03943 | METPA0448 | NA | Metabolic pathways;Biosynthesis of secondary metabolites;Biosynthesis of antibiotics;Biosynthesis of amino acids;Biosynthesis of plant secondary metabolites;ABC transporters;Arginine and proline metabolism;2-Oxocarboxylic acid metabolism;Other | 1.36 | 0.383 | 1.05 | 0.918 | 5.15 | 0.339 | 0.20 | 0.371078 |
| 144 | Ornithine | [M+H]+1 | POS | Mass,RT, Spectral match | Hilic significant | 132.090 | 6.077 | C00077 | HMDB0000214 | | Metabolic pathways;Biosynthesis of secondary metabolites;Biosynthesis of antibiotics;Biosynthesis of amino acids;Biosynthesis of plant secondary metabolites;ABC transporters;Arginine and proline metabolism;2-Oxocarboxylic acid metabolism;Other | 1.33 | 0.690 | 3.39 | 0.098 | 0.85 | 0.617 | 3.97 | 0.018484 |
| 145 | L_Aspartic_acid | [M+H]+1 | POS | Mass, Spectral match | C18 | 133.037 | 0.508 | C00049 | HMDB0000191 | 5960 | Metabolic pathways;Biosynthesis of secondary metabolites;Biosynthesis of antibiotics;Microbial metabolism in diverse | 0.85 | 0.701 | 0.60 | 0.140 | 0.63 | 0.307 | 0.96 | 0.859757 |

| | | | | | | | | | | | | | | | | | | | |
|---|---|---|---|---|---|---|---|---|---|---|---|---|---|---|---|---|---|---|---|
| | | | | | | | | | | | environments;Biosynthesis of plant secondary metabolites;Biosynthesis of amino acids;ABC transporters;Protein digestion and absorption;Central carbon metabolism in cancer;2-Oxocarboxylic acid metabolism;Aminoacyl-tRNA biosynthesis;Biosynthesis of alkaloids derived from ornithine, lysine and nicotinic acid;Neuroactive ligand-receptor interaction;Other | | | | | | | | |
| 146 | L_Asparticacid | [M+H]+1 | POS | Mass,RT, Spectral match | Hilic | 133.037 | 21.515 | C01042 | HMDB0000812 | 65065 | Metabolic pathways;Biosynthesis of secondary metabolites;Microbial metabolism in diverse environments;Biosynthesis of antibiotics;Biosynthesis of amino acids;Biosynthesis of plant secondary metabolites;ABC transporters;Protein digestion and absorption;Cyanoamino acid metabolism;2-Oxocarboxylic acid metabolism;Aminoacyl-tRNA biosynthesis;Central carbon metabolism in cancer;Glycine, serine and threonine metabolism;Cysteine and methionine metabolism;Histidine metabolism;Other | 0.90 | 0.604 | 0.87 | 0.490 | 0.78 | 0.618 | 1.11 | 0.476514 |
| 147 | D_Aspartate | [M+H]+1 | POS | Mass,RT, Spectral match | Hilic | 133.038 | 6.966 | C00402 | HMDB0006483 | 83887 | Metabolic pathways;Biosynthesis of secondary metabolites;Microbial metabolism in diverse environments;Biosynthesis of antibiotics;Biosynthesis of amino acids;Biosynthesis of plant secondary metabolites;ABC transporters;Protein digestion and absorption;Cyanoamino acid metabolism;2-Oxocarboxylic acid metabolism;Aminoacyl-tRNA biosynthesis;Central carbon metabolism in cancer;Glycine, serine and threonine metabolism;Cysteine and methionine metabolism;Histidine metabolism;Other | 1.04 | 0.898 | 1.34 | 0.395 | 1.18 | 0.565 | 1.14 | 0.712407 |
| 148 | L_Malic_acid | [M-H]-1 | NEG | Mass, Spectral match | C18 | 134.020 | 0.536 | C00149 | HMDB0000156 | 222656 | | 1.10 | 0.507 | 1.25 | 0.272 | 1.19 | 0.167 | 1.05 | 0.9129 |
| 149 | chavicol | [M+H]+1 | POS | Mass, Spectral | C18 | 134.073 | 13.880 | C01710 | HMDB59601 | 22819 | Biosynthesis of secondary metabolites;Biosynthesis of | 0.66 | 0.317 | 0.92 | 0.810 | 1.04 | 0.341 | 0.88 | 0.680736 |

| | | | | | | | | | | | | | | | | | | | |
|---|---|---|---|---|---|---|---|---|---|---|---|---|---|---|---|---|---|---|---|
| | | | | match | | | | | | | plant secondary metabolites;Other | | | | | | | | |
| 150 | Adenine | [M+H]+1 | POS | Mass, Spectral match | Hilic significant | 135.055 | 1.106 | C00147 | HMDB0000034 | 190 | Metabolic pathways;Other | 0.60 | 0.007 | 0.94 | 0.835 | 1.30 | 0.180 | 0.73 | 0.257069 |
| 151 | O_Acetylaniline | [M+H]+1 | POS | Mass, Spectral match | Hilic | 135.068 | 4.272 | | HMDB0032630 | 11086 | Biosynthesis of secondary metabolites;Microbial metabolism in diverse environments;Cyanoamino acid metabolism;2-Oxocarboxylic acid metabolism;Other | 1.14 | 0.533 | 0.99 | 0.968 | 0.99 | 0.767 | 1.00 | 0.776861 |
| 152 | Acetanilide | [M+H]+1 | POS | Mass, Spectral match | Hilic | 135.068 | 4.031 | C07565 | HMDB0001250 | 904 | Biosynthesis of secondary metabolites;Microbial metabolism in diverse environments;Cyanoamino acid metabolism;2-Oxocarboxylic acid metabolism;Other | 0.13 | 0.078 | 0.21 | 0.106 | 0.40 | 0.617 | 0.52 | 0.693201 |
| 153 | N_Benzylformamide | [M+H]+1 | POS | Mass, Spectral match | Hilic | 135.068 | 4.681 | C04744 | HMDB0011657 | 127546 | Biosynthesis of secondary metabolites;Microbial metabolism in diverse environments;Cyanoamino acid metabolism;2-Oxocarboxylic acid metabolism;Other | 1.45 | 0.094 | 1.46 | 0.212 | 1.26 | 0.402 | 1.15 | 0.511694 |
| 486 | Pseudouridine | [M+H]+1 | POS | Mass,RT, Spectral match | Hilic | 244.069 | 3.417 | C02067 | HMDB0000767 | 15047 | Metabolic pathways;Pyrimidine metabolism | 0.85 | 0.492 | 0.82 | 0.405 | 0.56 | 0.219 | 1.46 | 0.02866 |
| 155 | 4_Deoxythreonic_acid | [M-H]-1 | NEG | Mass, Spectral match | Hilic | 136.037 | 3.199 | | HMDB0002453 | 10964471 | Other | 2.90 | 0.319 | 0.94 | 0.936 | 1.30 | 0.176 | 0.72 | 0.586038 |
| 156 | Hypoxanthine | [M+H]+1 | POS | Mass, Spectral match | Hilic | 136.038 | 2.054 | C00262 | HMDB0000157 | 790 | Metabolic pathways;Other | 5.66 | 0.222 | 1.69 | 0.529 | 4.17 | 0.141 | 0.40 | 0.439746 |
| 157 | allopurinol | [M+H]+1 | POS | Mass, Spectral match | Hilic | 136.038 | 3.163 | | HMDB0014581 | 2094 | Metabolic pathways;Other | 0.43 | 0.159 | 0.49 | 0.156 | 0.88 | 0.828 | 0.56 | 0.540881 |
| 158 | Hypoxanthine | [M+H]+1 | POS | Mass, Spectral match | C18 | 136.038 | 0.570 | C00262 | HMDB0000157 | 790 | Metabolic pathways;Other | 4.15 | 0.217 | 1.49 | 0.525 | 3.17 | 0.192 | 0.47 | 0.542417 |
| 159 | 2_Methylnicotinamide | [M+H]+1 | POS | Mass,RT, Spectral match | Hilic | 136.064 | 0.774 | C02918 | HMDB0000699 | 457 | | 2.40 | 0.085 | 3.27 | 0.114 | 2.28 | 0.084 | 1.43 | 0.709787 |
| 160 | 2_Hydroxy_24_pentadienoate | [M+H]+1 | POS | Mass, RT, Spectral match | C18 | 137.020 | 18.609 | C00596 | METPA0060 | NA | | 0.81 | 0.184 | 1.06 | 0.695 | 1.01 | 0.845 | 1.04 | 0.732325 |
| 161 | 3_Pyridylaceticacid | [M+H]+1 | POS | Mass,RT, Spectral match | Hilic | 137.047 | 2.938 | | HMDB0001538 | 108 | Metabolic pathways;Biosynthesis of secondary metabolites;Microbial metabolism in diverse environments;Biosynthesis of antibiotics;Biosynthesis of amino acids;Biosynthesis of plant secondary metabolites;Degradation of aromatic compounds;Other | 0.55 | 0.108 | 0.53 | 0.100 | 0.44 | 0.293 | 1.21 | 0.80166 |
| 162 | 2_Aminobenzoic_acid | [M+H]+1 | POS | Mass, RT, Spectral match | C18 significant | 137.048 | 0.520 | C00108 | HMDB0001123 | 227 | Metabolic pathways;Biosynthesis of secondary metabolites;Biosynthesis of | 0.55 | 0.055 | 0.49 | 0.034 | 0.44 | 0.278 | 1.10 | 0.923022 |

| | | | | | | | | | | | | | | | | | | | |
|---|---|---|---|---|---|---|---|---|---|---|---|---|---|---|---|---|---|---|---|
| | | | | | | | | | | | antibiotics;Microbial metabolism in diverse environments;Biosynthesis of plant secondary metabolites;Biosynthesis of amino acids;Degradation of aromatic compounds;Biosynthesis of alkaloids derived from shikimate pathway;Other | | | | | | | | |
| 163 | 4_Hydroxybenzoic_acid | [M-H]-1 | NEG | Mass, Spectral match | C18 | 138.031 | 5.890 | C00156 | HMDB0000500 | 135 | | 0.65 | 0.076 | 0.71 | 0.088 | 4.28 | 0.013 | 0.17 | 0.001272 |
| 164 | 4_Nitro_aniline | [M+H]+1 | POS | Mass,RT, Spectral match | Hilic | 138.043 | 2.093 | NA | #N/A | #N/A | Metabolic pathways;Histidine metabolism | 0.01 | 0.025 | 0.01 | 0.025 | 0.01 | 0.199 | 1.22 | 0.947425 |
| 165 | Urocanicacid | [M+H]+1 | POS | Mass,RT, Spectral match | Hilic | 138.043 | 2.310 | C00785 | HMDB0000301 | 736715 | Metabolic pathways;Histidine metabolism | 0.02 | 0.072 | 0.60 | 0.470 | 0.56 | 0.299 | 1.08 | 0.830557 |
| 166 | Hydroxypyruvic_acid | [M+H]+1 | POS | Mass, Spectral match | C18 | 138.980 | 18.565 | C00168 | HMDB0001352 | 964 | | 0.97 | 0.678 | 0.99 | 0.857 | 0.85 | 0.015 | 1.16 | 0.12201 |
| 167 | Tropinone | [M+H]+1 | POS | Mass, Spectral match | C18 | 139.100 | 0.521 | C00783 | METPA0078 | NA | Metabolic pathways;Biosynthesis of secondary metabolites;Tropane, piperidine and pyridine alkaloid biosynthesis;Biosynthesis of alkaloids derived from ornithine, lysine and nicotinic acid | 1.00 | 0.992 | 0.88 | 0.373 | 0.88 | 0.150 | 1.00 | 0.820806 |
| 168 | Acetylphosphate | [M+H]+1 | POS | Mass, Spectral match | C18 | 139.988 | 18.574 | C00227 | HMDB0001494 | 186 | Metabolic pathways;Microbial metabolism in diverse environments;Other | 0.97 | 0.755 | 1.00 | 0.995 | 0.94 | 0.264 | 1.07 | 0.565379 |
| 169 | Methylimidazoleacetic_acid | [M+H]+1 | POS | Mass, Spectral match | C18 | 140.058 | 0.523 | C05828 | HMDB0002820 | 75810 | Other | 1.58 | 0.047 | 1.36 | 0.327 | 1.54 | 0.229 | 0.88 | 0.659595 |
| 170 | L_Histidinol | [M+H]+1 | POS | Mass, Spectral match | Hilic | 140.084 | 27.572 | C00860 | HMDB0003431 | 165271 | Metabolic pathways;Microbial metabolism in diverse environments;Degradation of aromatic compounds;Other | 1.05 | 0.188 | 0.95 | 0.062 | 0.84 | 0.140 | 1.13 | 0.075824 |
| 171 | O_Phosphorylethanolamine | [2M-H]-1 | NEG | Mass, Spectral match | Hilic | 141.019 | 15.550 | C00682 | METPA0068 | NA | Metabolic pathways;Other | 1.31 | 0.373 | 0.74 | 0.423 | 0.98 | 0.478 | 0.75 | 0.370887 |
| 172 | Dimetridazole | [M+H]+1 | POS | Mass, Spectral match | C18 | 141.054 | 1.720 | D07855 | #N/A | 3090 | | 1.09 | 0.714 | 1.13 | 0.503 | 0.86 | 0.956 | 1.31 | 0.206593 |
| 173 | Crotono_betaine | [M+H]+1 | POS | Mass, Spectral match | Hilic | 141.115 | 0.760 | C00719 | HMDB0000043 | 247 | Metabolic pathways;Biosynthesis of secondary metabolites;Other | 1.06 | 0.440 | 1.07 | 0.343 | 1.08 | 0.633 | 0.99 | 0.848269 |
| 174 | Tropine | [M+H]+1 | NEG | Mass, Spectral match | C18 | 141.115 | 0.525 | C00729 | METPA0074 | NA | Metabolic pathways;Biosynthesis of secondary metabolites;Tropane, piperidine and pyridine alkaloid biosynthesis;Biosynthesis of alkaloids derived from ornithine, lysine and nicotinic acid | 1.04 | 0.523 | 1.04 | 0.517 | 0.97 | 0.259 | 1.08 | 0.258883 |

| | | | | | | | | | | | | | | | | | | | |
|---|---|---|---|---|---|---|---|---|---|---|---|---|---|---|---|---|---|---|---|
| 175 | Ectoine | [M+H]+1 | POS | Mass, Spectral match | Hilic | 142.074 | 1.915 | C06231 | METPA0797 | NA | Metabolic pathways;Microbial metabolism in diverse environments;Glycine, serine and threonine metabolism | 0.36 | 0.093 | 0.43 | 0.130 | 0.33 | 0.407 | 1.30 | 0.749322 |
| 176 | Proline_betaine | [M+H]+1 | POS | Mass, Spectral match | C18 | 143.094 | 0.523 | C10172 | HMDB0004827 | 115244 | ABC transporters | 1.12 | 0.614 | 0.92 | 0.607 | 1.39 | 0.075 | 0.66 | 0.153304 |
| 177 | Stachydrine | [M+H]+1 | POS | Mass,RT, Spectral match | Hilic significant | 143.095 | 2.513 | C10172 | HMDB0029230 | 375481 | ABC transporters | 1.41 | 0.208 | 1.19 | 0.335 | 2.13 | 0.027 | 0.56 | 0.059077 |
| 178 | 2_3_Dimethylmaleate | [M+H]+1 | POS | Mass, Spectral match | C18 | 144.042 | 0.522 | C00922 | HMDB36232 | 3032309 | Microbial metabolism in diverse environments;Other | 1.08 | 0.457 | 1.74 | 0.006 | 1.81 | 0.083 | 0.96 | 0.875366 |
| 179 | 4_Guanidinobutanamide | [M+H]+1 | POS | Mass, Spectral match | Hilic | 144.101 | 0.844 | C03078 | METPA0359 | NA | Arginine and proline metabolism | 1.20 | 0.752 | 1.26 | 0.694 | 1.44 | 0.759 | 0.88 | 0.996064 |
| 180 | 12H_Isoquinolinone | [M+H]+1 | POS | Mass, Spectral match | Hilic | 145.053 | 2.613 | C13650 | HMDB0006468 | 107669 | | 1.37 | 0.225 | 1.26 | 0.370 | 1.19 | 0.671 | 1.06 | 0.59721 |
| 181 | 4_Trimethylammoniobutanoic_acid | [M+H]+1 | POS | Mass, RT, Spectral match | C18 | 145.110 | 0.501 | C01181 | HMDB0001161 | 134 | | 1.34 | 0.332 | 0.94 | 0.734 | 1.05 | 0.379 | 0.90 | 0.837041 |
| 182 | Acetylcholine | [M+Na]+1 | POS | Mass, Spectral match | Hilic | 145.110 | 1.206 | C01996 | HMDB0000895 | 187 | | 2.19 | 0.430 | 1.22 | 0.544 | 1.80 | 0.093 | 0.68 | 0.389802 |
| 183 | Spermidine | [M+H]+1 | POS | Mass,RT, Spectral match | Hilic | 145.158 | 11.288 | C00315 | HMDB0001257 | 1102 | Metabolic pathways;ABC transporters;Arginine and proline metabolism;Other | 1.03 | 0.931 | 1.02 | 0.953 | 0.75 | 0.459 | 1.37 | 0.499335 |
| 184 | Spermidine | [M+H]+1 | POS | Mass, RT, Spectral match | C18 | 145.158 | 0.487 | C00315 | HMDB0001257 | 1102 | Metabolic pathways;Bile secretion;ABC transporters;Arginine and proline metabolism;Other | 0.88 | 0.577 | 1.11 | 0.669 | 0.98 | 0.579 | 1.13 | 0.794153 |
| 185 | Oxoglutaric_acid | [M-H]-1 | NEG | Mass, Spectral match | C18 | 146.021 | 0.576 | C00026 | HMDB0000208 | 51 | | 8.95 | 0.133 | 0.77 | 0.466 | 1.64 | 0.753 | 0.47 | 0.018239 |
| 186 | 3_Methylglutaric_acid | [M-H]-1 | NEG | Mass, RT, Spectral match | C18 | 146.057 | 0.786 | C00659 | #N/A | 21 | Metabolic pathways;Biosynthesis of secondary metabolites;Biosynthesis of antibiotics;Microbial metabolism in diverse environments;Biosynthesis of amino acids;Degradation of aromatic compounds;2-Oxocarboxylic acid metabolism;Other | 0.84 | 0.699 | 1.02 | 0.968 | 0.66 | 0.222 | 1.54 | 0.345775 |
| 187 | Adipic_acid_ | [M+H]+1 | POS | Mass,RT, Spectral match | Hilic significant | 146.058 | 5.925 | C06104 | HMDB0000448 | 196 | Metabolic pathways;Biosynthesis of secondary metabolites;Microbial metabolism in diverse environments;Biosynthesis of antibiotics;Biosynthesis of amino acids;2-Oxocarboxylic acid metabolism;Degradation of aromatic compounds;Other | 1.52 | 0.052 | 1.42 | 0.262 | 1.49 | 0.218 | 0.95 | 0.909194 |
| 188 | L_Glutamine | [M+H]+1 | POS | Mass, Spectral match | C18 | 146.069 | 0.503 | C00064 | HMDB0000641 | 5961 | Metabolic pathways;Microbial metabolism in diverse environments;Biosynthesis of plant secondary metabolites;Biosynthesis of amino acids;ABC | 1.34 | 0.051 | 1.14 | 0.480 | 1.21 | 0.230 | 0.94 | 0.71882 |

| | | | | | | | | | | | | | | | | | | | |
|---|---|---|---|---|---|---|---|---|---|---|---|---|---|---|---|---|---|---|---|
| | | | | | | | | | | | transporters;Protein digestion and absorption;Central carbon metabolism in cancer;Aminoacyl-tRNA biosynthesis;Other | | | | | | | | |
| 189 | 3_Ureidoisobutyrate | [M+H]+1 | POS | Mass,RT, Spectral match | Hilic | 146.069 | 1.908 | C05100 | HMDB0002031 | 160663 | Metabolic pathways;Microbial metabolism in diverse environments;Biosynthesis of amino acids;Biosynthesis of plant secondary metabolites;ABC transporters;Protein digestion and absorption;Aminoacyl-tRNA biosynthesis;Central carbon metabolism in cancer;Pyrimidine metabolism;Histidine metabolism;Other | 1.51 | 0.193 | 1.27 | 0.315 | 1.25 | 0.462 | 1.02 | 0.944151 |
| 190 | D___Glutamine | [M+H]+1 | POS | Mass,RT, Spectral match | Hilic | 146.069 | 7.936 | C00064 | HMDB0000641 | 5961 | Metabolic pathways;Microbial metabolism in diverse environments;Biosynthesis of amino acids;Biosynthesis of plant secondary metabolites;ABC transporters;Protein digestion and absorption;Aminoacyl-tRNA biosynthesis;Central carbon metabolism in cancer;Pyrimidine metabolism;Histidine metabolism;Other | 1.19 | 0.284 | 1.16 | 0.428 | 1.12 | 0.421 | 1.04 | 0.85315 |
| 191 | 4_Guanidinobutanoic_acid | [M-H]-1 | NEG | Mass, RT, Spectral match | C18 | 146.093 | 6.377 | C01035 | HMDB0003464 | 500 | | 0.95 | 0.901 | 1.08 | 0.789 | 3.42 | 0.359 | 0.32 | 0.551205 |
| 192 | L_Lysine | [M+H]+1 | POS | Mass, RT, Spectral match | C18 | 146.105 | 0.455 | C00047 | HMDB0000182 | 5962 | Metabolic pathways;Biosynthesis of secondary metabolites;Biosynthesis of antibiotics;Microbial metabolism in diverse environments;Biosynthesis of plant secondary metabolites;Biosynthesis of amino acids;ABC transporters;Protein digestion and absorption;2-Oxocarboxylic acid metabolism;Aminoacyl-tRNA biosynthesis;Tropane, piperidine and pyridine alkaloid biosynthesis;Biosynthesis of alkaloids derived from ornithine, lysine and nicotinic acid;Other | 0.98 | 0.911 | 0.83 | 0.308 | 1.07 | 0.561 | 0.78 | 0.163513 |
| 193 | Lysine | [M+H]+1 | POS | Mass, Spectral match | Hilic significant | 146.106 | 1.337 | C00047 | HMDB0000182 | 5962 | Metabolic pathways;Biosynthesis of secondary metabolites;Microbial metabolism in diverse environments;Biosynthesis of antibiotics;Biosynthesis of amino acids;Biosynthesis of | 0.27 | 0.065 | 0.51 | 0.189 | 0.98 | 0.936 | 0.52 | 0.184057 |

| | | | | | | | | | | | | | | | | | | | |
|---|---|---|---|---|---|---|---|---|---|---|---|---|---|---|---|---|---|---|---|
| | | | | | | | | | | | plant secondary metabolites;ABC transporters;Protein digestion and absorption;2-Oxocarboxylic acid metabolism;Aminoacyl-tRNA biosynthesis;Lysine degradation;Other | | | | | | | | |
| 194 | Cycloalliin | [M+H]+1 | POS | Mass, Spectral match | Hilic | 147.035 | 1.803 | | HMDB0029417 | 12305351 | | 1.00 | 0.948 | 1.06 | 0.459 | 24.22 | 0.364 | 0.04 | 0.300616 |
| 195 | L_Glutamicacid | [M+H]+1 | POS | Mass, Spectral match | Hilic | 147.053 | 15.653 | C00025 | HMDB0000148 | 33032 | Metabolic pathways;Biosynthesis of secondary metabolites;Microbial metabolism in diverse environments;Biosynthesis of antibiotics;Biosynthesis of amino acids;Biosynthesis of plant secondary metabolites;ABC transporters;Protein digestion and absorption;Cyanoamino acid metabolism;Arginine and proline metabolism;2-Oxocarboxylic acid metabolism;Aminoacyl-tRNA biosynthesis;Central carbon metabolism in cancer;Cysteine and methionine metabolism;Histidine metabolism;Other | 1.11 | 0.604 | 0.95 | 0.808 | 1.18 | 0.364 | 0.81 | 0.259356 |
| 196 | L_Glutamic_acid | [M+H]+1 | POS | Mass, RT, Spectral match | C18 | 147.053 | 0.513 | C00025 | HMDB0000148 | 33032 | Metabolic pathways;Biosynthesis of secondary metabolites;Biosynthesis of antibiotics;Microbial metabolism in diverse environments;Biosynthesis of plant secondary metabolites;Biosynthesis of amino acids;ABC transporters;Protein digestion and absorption;Arginine and proline metabolism;Central carbon metabolism in cancer;2-Oxocarboxylic acid metabolism;Aminoacyl-tRNA biosynthesis;Biosynthesis of alkaloids derived from ornithine, lysine and nicotinic acid;Neuroactive ligand-receptor interaction;Other | 1.27 | 0.207 | 1.15 | 0.502 | 1.38 | 0.114 | 0.83 | 0.369175 |
| 197 | 3_Methyloxindole | [M+H]+1 | POS | Mass, Spectral match | Hilic | 147.068 | 2.606 | C05834 | HMDB0004186 | 151066 | | 1.28 | 0.287 | 1.22 | 0.385 | 1.20 | 0.504 | 1.01 | 0.890134 |
| 198 | 2_Aminophenol | [M+H]+1 | POS | Mass, RT, Spectral match | C18 | 148.016 | 11.394 | C01987 | METPA0236 | NA | | 0.90 | 0.084 | 0.92 | 0.193 | 0.94 | 0.278 | 0.97 | 0.660272 |
| 488 | Uridine | [M+H]+1 | POS | Mass,RT, Spectral match | Hilic significant | 244.070 | 3.417 | C00299 | HMDB0000296 | 6029 | Metabolic pathways;Pyrimidine metabolism | 0.85 | 0.492 | 0.82 | 0.405 | 0.56 | 0.219 | 1.46 | 0.02866 |

| | | | | | | | | | | | | | | | | | |
|---|---|---|---|---|---|---|---|---|---|---|---|---|---|---|---|---|---|
| 200 | 1_phenylpropane_1_2_dione | [M+H]+1 | POS | Mass,RT, Spectral match | Hilic | 148.052 | 2.600 | C17268 | HMDB0035243 | 11363 | Metabolic pathways;Biosynthesis of secondary metabolites;Microbial metabolism in diverse environments;Biosynthesis of plant secondary metabolites;Degradation of aromatic compounds;Other | 1.13 | 0.481 | 0.98 | 0.905 | 1.07 | 0.605 | 0.92 | 0.697651 |
| 201 | trans_Cinnamic_acid | [M+H]+1 | POS | Mass, Spectral match | C18 | 148.052 | 0.997 | C00423 | HMDB0000930 | 444539 | Metabolic pathways;Biosynthesis of secondary metabolites;Microbial metabolism in diverse environments;Biosynthesis of plant secondary metabolites;Degradation of aromatic compounds;Biosynthesis of alkaloids derived from shikimate pathway;Other | 1.19 | 0.374 | 1.02 | 0.939 | 1.03 | 0.706 | 0.98 | 0.989003 |
| 202 | Cinnamic_acid | [M+H]+1 | POS | Mass,RT, Spectral match | Hilic significant | 148.053 | 2.377 | C10438 | HMDB0000567 | 5372954 | Metabolic pathways;Biosynthesis of secondary metabolites;Microbial metabolism in diverse environments;Biosynthesis of plant secondary metabolites;Degradation of aromatic compounds;Other | 0.01 | 0.048 | 0.54 | 0.426 | 0.29 | 0.674 | 1.84 | 0.513158 |
| 203 | L_Methionine | [M+H]+1 | POS | Mass, Spectral match | C18 | 149.051 | 0.611 | C00073 | HMDB0000696 | 6137 | Metabolic pathways;Biosynthesis of secondary metabolites;Biosynthesis of plant secondary metabolites;Biosynthesis of amino acids;ABC transporters;Protein digestion and absorption;Central carbon metabolism in cancer;2-Oxocarboxylic acid metabolism;Aminoacyl-tRNA biosynthesis;Other | 1.01 | 0.969 | 1.01 | 0.971 | 0.92 | 0.968 | 1.10 | 0.41532 |
| 204 | Methionine | [M+H]+1 | POS | Mass, Spectral match | Hilic | 149.051 | 3.219 | C00073 | HMDB0000696 | 6137 | Metabolic pathways;Biosynthesis of secondary metabolites;Biosynthesis of amino acids;Biosynthesis of plant secondary metabolites;ABC transporters;Protein digestion and absorption;2-Oxocarboxylic acid metabolism;Aminoacyl-tRNA biosynthesis;Central carbon metabolism in cancer;Cysteine and methionine metabolism;Other | 1.20 | 0.295 | 1.07 | 0.739 | 0.95 | 0.932 | 1.12 | 0.468022 |
| 205 | Methamphetamine | [M+H]+1 | POS | Mass, Spectral match | Hilic | 149.120 | 0.352 | | HMDB0015517 | 10836 | | 0.76 | 0.554 | 0.27 | 0.073 | 0.74 | 0.325 | 0.37 | 0.250105 |
| 206 | Phentermine | [M+H]+1 | POS | Mass, Spectral | C18 | 149.120 | 18.128 | C07438 | HMDB0014337 | 4771 | | 0.98 | 0.763 | 0.98 | 0.807 | 1.02 | 0.928 | 0.96 | 0.558568 |

| | | | | | | | | | | | | | | | | | | | |
|---|---|---|---|---|---|---|---|---|---|---|---|---|---|---|---|---|---|---|---|
| | | | | match | | | | | | | | | | | | | | | |
| 207 | Phenylglyoxylic_acid | [M+H]+1 | POS | Mass, Spectral match | C18 | 150.031 | 0.544 | C02137 | HMDB0001587 | 11915 | Microbial metabolism in diverse environments;Other | 1.03 | 0.817 | 1.05 | 0.679 | 0.84 | 0.568 | 1.25 | 0.052022 |
| 515 | 1_Ribosylimidazole_4_acetate | [M+H]+1 | POS | Mass, Spectral match | Hilic | 258.085 | 3.487 | C05131 | HMDB0002331 | 440569 | Histidine metabolism | 0.04 | 0.000 | 0.75 | 0.451 | 0.16 | 0.048 | 4.85 | 0.202438 |
| 209 | Hypotaurocyamine | [M+H]+1 | POS | Mass, Spectral match | Hilic | 151.041 | 9.182 | C00642 | HMDB0000020 | 127 | | 1.03 | 0.950 | 0.74 | 0.433 | 0.55 | 0.529 | 1.35 | 0.73435 |
| 210 | Guanine | [M+H]+1 | POS | Mass, RT, Spectral match | C18 | 151.049 | 0.621 | C00242 | HMDB0000132 | 764 | | 0.20 | 0.098 | 0.66 | 0.611 | 0.46 | 0.920 | 1.43 | 0.764726 |
| 211 | Guanine | [M+K]+1 | POS | Mass, Spectral match | Hilic significant | 151.049 | 9.352 | C00242 | HMDB0000132 | 764 | | 1.10 | 0.801 | 0.58 | 0.036 | 0.67 | 0.835 | 0.87 | 0.286708 |
| 212 | Xanthine | [M-H]-1 | NEG | Mass, RT, Spectral match | C18 | 152.033 | 0.578 | C00385 | HMDB0000292 | 1188 | | 2.75 | 0.325 | 1.91 | 0.404 | 2.38 | 0.109 | 0.80 | 0.593851 |
| 213 | Xanthine | [2M-H]-1 | NEG | Mass,RT, Spectral match | Hilic | 152.033 | 2.086 | C00385 | HMDB0000292 | 1188 | Metabolic pathways;Biosynthesis of secondary metabolites;Microbial metabolism in diverse environments;Biosynthesis of plant secondary metabolites;Other | 3.43 | 0.291 | 2.57 | 0.306 | 3.07 | 0.124 | 0.84 | 0.602587 |
| 214 | N1_Methyl_2_pyridone_5_carboxamide | [M+H]+1 | POS | Mass, Spectral match | Hilic | 152.058 | 1.247 | C05842 | HMDB0004193 | 69698 | Metabolic pathways;Microbial metabolism in diverse environments;Other | 1.03 | 0.928 | 0.69 | 0.103 | 0.78 | 0.816 | 0.89 | 0.367061 |
| 215 | N1_Methyl_2_pyridone_5_carboxamide | [M+Na]+1 | POS | Mass, RT, Spectral match | C18 | 152.058 | 0.663 | C05842 | HMDB0004193 | 69698 | Metabolic pathways;Microbial metabolism in diverse environments;Other | 0.87 | 0.536 | 0.57 | 0.016 | 0.47 | 0.207 | 1.21 | 0.376327 |
| 216 | L_Arabitol | [M+Cl]-1 | POS | Mass, Spectral match | Hilic | 152.067 | 4.025 | C00532 | HMDB0001851 | 439255 | | 1.06 | 0.893 | 0.91 | 0.823 | 1.47 | 0.044 | 0.62 | 0.210852 |
| 217 | D_Arabitol | [M+Cl]-1 | POS | Mass, Spectral match | Hilic | 152.067 | 3.883 | C01904 | HMDB0000568 | 827 | | 1.50 | 0.114 | 1.45 | 0.172 | 1.63 | 0.049 | 0.89 | 0.602848 |
| 218 | 2_Pyrocatechuic_acid | [M+H]+1 | POS | Mass, RT, Spectral match | C18 | 153.018 | 0.573 | C00196 | HMDB0000397 | 19 | | 1.47 | 0.377 | 1.39 | 0.477 | 1.23 | 0.314 | 1.13 | 0.944171 |
| 219 | Imidazol_5_yl_pyruvate | [M+H]+1 | POS | Mass, Spectral match | Hilic | 154.038 | 6.796 | C03277 | METPA0379 | NA | Histidine metabolism | 1.55 | 0.278 | 1.06 | 0.899 | 1.17 | 0.621 | 0.91 | 0.668582 |
| 220 | D_1_Aminopropan_2_ol O_phosphate | [M+H]+1 | POS | Mass, Spectral match | Hilic | 155.035 | 2.886 | C04122 | METPA0470 | NA | Other | 1.13 | 0.522 | 1.23 | 0.039 | 1.54 | 0.062 | 0.80 | 0.274243 |
| 221 | L_Histidine | [M+H]+1 | POS | Mass, Spectral match | C18 | 155.069 | 0.464 | C00135 | HMDB0000177 | 6274 | Metabolic pathways;Biosynthesis of secondary metabolites;Biosynthesis of plant secondary metabolites;Biosynthesis of amino acids;ABC transporters;Protein digestion and absorption;Central carbon metabolism in | 1.45 | 0.203 | 1.28 | 0.230 | 1.30 | 0.211 | 0.98 | 0.950515 |

| | | | | | | | | | | | | | | | | | | | |
|---|---|---|---|---|---|---|---|---|---|---|---|---|---|---|---|---|---|---|---|
| | | | | | | | | | | | cancer;Aminoacyl-tRNA biosynthesis;Other | | | | | | | | |
| 222 | Histidine | [2M-H]-1 | NEG | Mass, Spectral match | Hilic significant | 155.070 | 2.662 | C00135 | HMDB0000177 | 6274 | Metabolic pathways;Biosynthesis of secondary metabolites;Biosynthesis of amino acids;Biosynthesis of plant secondary metabolites;ABC transporters;Protein digestion and absorption;Aminoacyl-tRNA biosynthesis;Central carbon metabolism in cancer;Histidine metabolism;Other | 3.34 | 0.047 | 2.08 | 0.003 | 1.97 | 0.366 | 1.06 | 0.364345 |
| 223 | Phosphoglycolic_acid | [M+H]+1 | POS | Mass, Spectral match | C18 | 155.983 | 18.580 | C00988 | HMDB0000816 | 529 | Metabolic pathways;Biosynthesis of secondary metabolites;Biosynthesis of antibiotics;Other | 1.05 | 0.640 | 0.96 | 0.560 | 0.90 | 0.162 | 1.06 | 0.442673 |
| 224 | 4_Imidazolone_5_propanoate | [M+H]+1 | POS | Mass, Spectral match | Hilic | 156.053 | 2.353 | C03680 | HMDB0001014 | 128 | Metabolic pathways;Histidine metabolism | 0.37 | 0.075 | 0.71 | 0.526 | 0.22 | 0.208 | 3.21 | 0.249658 |
| 225 | Imidazolelactic_acid | [M+H]+1 | POS | Mass, Spectral match | Hilic significant | 156.054 | 2.353 | C05132 | HMDB0002320 | 459122 | Metabolic pathways;Histidine metabolism | 0.37 | 0.075 | 0.71 | 0.526 | 0.22 | 0.208 | 3.21 | 0.249658 |
| 226 | 3_Methylcrotonylglycine | [M+H]+1 | POS | Mass,RT, Spectral match | Hilic | 157.074 | 1.255 | | HMDB0000459 | 169485 | Biosynthesis of antibiotics | 1.32 | 0.565 | 0.77 | 0.324 | 2.56 | 0.143 | 0.30 | 0.145348 |
| 227 | Adenine | [M-H]-1 | NEG | Mass, RT, Spectral match | C18 | 158.043 | 0.506 | C00147 | HMDB0000034 | 190 | Metabolic pathways;Microbial metabolism in diverse environments;Other | 0.88 | 0.628 | 0.88 | 0.670 | 0.73 | 0.426 | 1.21 | 0.448395 |
| 228 | R__Allantoin | [2M-H]-1 | NEG | Mass,RT, Spectral match | Hilic | 158.044 | 2.455 | C02348 | METPA0275 | NA | Metabolic pathways;Microbial metabolism in diverse environments;Other | 0.94 | 0.817 | 1.10 | 0.712 | 0.90 | 0.703 | 1.21 | 0.198389 |
| 229 | L_2_Amino_4_methylenepentanedioic_acid | [M+H]+1 | POS | Mass,RT, Spectral match | Hilic | 158.047 | 9.356 | C00651 | HMDB0029433 | 96407 | | 1.11 | 0.712 | 0.60 | 0.127 | 0.71 | 0.007 | 0.85 | 0.334206 |
| 230 | Oxoadipic_acid | [M+H]+1 | POS | Mass, Spectral match | Hilic | 159.029 | 2.742 | C00322 | HMDB0000225 | 71 | | 0.11 | 0.194 | 0.64 | 0.598 | 0.07 | 0.392 | 8.82 | 0.019379 |
| 231 | N_Isovalerylglycine | [M-H]-1 | NEG | Mass, Spectral match | Hilic | 159.088 | 6.386 | | HMDB0000678 | 546304 | | 0.76 | 0.489 | 0.77 | 0.485 | 0.90 | 0.447 | 0.86 | 0.719219 |
| 232 | 3_Dehydrocarnitine | [M+H]+1 | POS | Mass, Spectral match | Hilic | 159.089 | 2.367 | C02636 | HMDB0012154 | 6991982 | Lysine degradation;Other | 1.52 | 0.237 | 1.45 | 0.338 | 2.06 | 0.222 | 0.71 | 0.724761 |
| 233 | 3_Dehydrocarnitine | [M+H]+1 | POS | Mass, RT, Spectral match | C18 | 159.089 | 0.562 | C02636 | HMDB0012154 | 6991982 | Tropane, piperidine and pyridine alkaloid biosynthesis;Other | 0.53 | 0.038 | 0.67 | 0.250 | 0.74 | 0.934 | 0.90 | 0.456826 |
| 234 | Acetyl_methylcholine | [M+H]+1 | POS | Mass, Spectral match | Hilic | 159.126 | 1.144 | C07471 | HMDB0015654 | 1993 | | 1.87 | 0.355 | 1.06 | 0.875 | 1.54 | 0.224 | 0.69 | 0.38286 |
| 235 | Methacholine | [M+H]+1 | POS | Mass, RT, Spectral match | C18 | 159.126 | 0.530 | C07471 | HMDB0015654 | 1993 | | 1.15 | 0.245 | 0.98 | 0.811 | 1.01 | 0.435 | 0.98 | 0.422758 |
| 236 | Aminopropylcadaverine | [M+H]+1 | POS | Mass, Spectral match | Hilic | 159.173 | 21.495 | C16565 | HMDB0012189 | 65523 | Biosynthesis of secondary metabolites;Biosynthesis of plant secondary metabolites;Other | 0.86 | 0.604 | 0.98 | 0.935 | 0.91 | 0.293 | 1.08 | 0.57698 |

| | | | | | | | | | | | | | | | | | | | |
|---|---|---|---|---|---|---|---|---|---|---|---|---|---|---|---|---|---|---|---|
| 237 | D_Ala_D_Ala | [M+Na]+1 | POS | Mass,RT, Spectral match | Hilic | 160.084 | 1.420 | C00993 | HMDB0003459 | 5460362 | Metabolic pathways;Microbial metabolism in diverse environments;Glycine, serine and threonine metabolism;Other | 1.65 | 0.488 | 1.14 | 0.834 | 2.30 | 0.293 | 0.50 | 0.496895 |
| 238 | D_Alanyl_D_alanine | [M+H]+1 | POS | Mass, RT, Spectral match | C18 | 160.085 | 0.522 | C00993 | HMDB0003459 | 5460362 | Metabolic pathways;Microbial metabolism in diverse environments;Other | 1.82 | 0.217 | 0.96 | 0.835 | 1.33 | 0.203 | 0.73 | 0.48161 |
| 239 | D_Ala_D_Ala | [M+Na]+1 | POS | Mass,RT, Spectral match | Hilic significant | 160.085 | 1.640 | C00993 | HMDB0003459 | 5460362 | Metabolic pathways;Microbial metabolism in diverse environments;Glycine, serine and threonine metabolism;Other | 0.68 | 0.341 | 1.34 | 0.499 | 10.19 | 0.053 | 0.13 | 0.050111 |
| 240 | L_Carnitinamide | [M+H]+1 | POS | Mass,RT, Spectral match | Hilic | 160.121 | 1.492 | C00318 | HMDB0000062 | 2724480 | | 1.06 | 0.869 | 1.24 | 0.514 | 0.89 | 0.891 | 1.39 | 0.442151 |
| 241 | allylcysteine | [M+H]+1 | POS | Mass, Spectral match | Hilic | 161.051 | 3.468 | C16759 | HMDB0034323 | 98280 | | 1.13 | 0.658 | 1.08 | 0.838 | 0.69 | 0.352 | 1.56 | 0.313385 |
| 242 | 2_Aminoadipicacid | [M+H]+1 | POS | Mass, Spectral match | Hilic | 161.068 | 12.187 | C00956 | HMDB0000510 | 469 | Metabolic pathways;Biosynthesis of secondary metabolites;Microbial metabolism in diverse environments;Biosynthesis of antibiotics;Biosynthesis of amino acids;2-Oxocarboxylic acid metabolism;Cysteine and methionine metabolism;Lysine degradation;Other | 1.17 | 0.653 | 1.72 | 0.191 | 2.63 | 0.068 | 0.65 | 0.425596 |
| 243 | Alpha_D_Aminoadipic_acid | [M+H]+1 | POS | Mass, RT, Spectral match | C18 | 161.069 | 0.516 | C00956 | HMDB0000510 | 469 | Metabolic pathways;Biosynthesis of secondary metabolites;Biosynthesis of antibiotics;Microbial metabolism in diverse environments;Biosynthesis of amino acids;2-Oxocarboxylic acid metabolism;Other | 1.02 | 0.887 | 1.66 | 0.005 | 1.77 | 0.034 | 0.94 | 0.992773 |
| 244 | Alpha_D_Aminoadipic_acid | [M-H]-1 | NEG | Mass, Spectral match | Hilic significant | 161.069 | 12.187 | C00956 | HMDB0000510 | 469 | Metabolic pathways;Biosynthesis of secondary metabolites;Microbial metabolism in diverse environments;Biosynthesis of antibiotics;Biosynthesis of amino acids;2-Oxocarboxylic acid metabolism;Cysteine and methionine metabolism;Lysine degradation;Other | 1.17 | 0.653 | 1.72 | 0.191 | 2.63 | 0.068 | 0.65 | 0.425596 |
| 245 | L_Carnitine | [M+Na]+1 | POS | Mass, Spectral match | Hilic | 161.105 | 1.258 | C00487 | #N/A | 85 | Other | 1.18 | 0.599 | 0.73 | 0.435 | 1.48 | 0.047 | 0.50 | 0.123973 |
| 246 | DL_Carnitine | [M+H]+1 | POS | Mass, Spectral match | C18 | 161.105 | 0.501 | C00318 | HMDB0000062 | 2724480 | Bile secretion;Other | 1.49 | 0.057 | 1.07 | 0.676 | 1.38 | 0.038 | 0.77 | 0.143493 |
| 247 | 5_Hydroxypyrazinamide | [M-H]-1 | NEG | Mass, Spectral match | C18 | 162.005 | 0.846 | C01956 | HMDB0001978 | 151823 | | 0.94 | 0.863 | 0.77 | 0.510 | 0.76 | 0.665 | 1.01 | 0.934428 |

| | | | | | | | | | | | | | | | | | | | |
|---|---|---|---|---|---|---|---|---|---|---|---|---|---|---|---|---|---|---|---|
| 248 | 2R_3S_2_3_Dimethylmalate | [M+H]+1 | POS | Mass, Spectral match | Hilic | 162.052 | 12.424 | C03652 | METPA0421 | NA | Microbial metabolism in diverse environments;Biosynthesis of antibiotics;Other | 1.15 | 0.800 | 1.66 | 0.205 | 2.31 | 0.010 | 0.72 | 0.24368 |
| 249 | 3_Hydroxy_3_methylglutaric_acid_ | [M+H]+1 | POS | Mass, Spectral match | Hilic significant | 162.053 | 9.804 | C03761 | HMDB0000355 | 1662 | Microbial metabolism in diverse environments;Biosynthesis of antibiotics;Other | 1.37 | 0.381 | 2.65 | 0.008 | 2.18 | 0.050 | 1.22 | 0.416806 |
| 250 | 3_Hydroxy_L_glutamate | [M+Na]+1 | POS | Mass, Spectral match | Hilic | 163.049 | 4.938 | C03079 | HMDB0002273 | 5460078 | Metabolic pathways;Arginine and proline metabolism | 16.36 | 0.153 | 1.81 | 0.321 | 11.06 | 0.254 | 0.16 | 0.332465 |
| 251 | Pterin | [M+Na]+1 | POS | Mass, Spectral match | Hilic | 163.049 | 2.459 | C00715 | HMDB0000802 | 73000 | Metabolic pathways;Arginine and proline metabolism | 1.84 | 0.477 | 1.89 | 0.404 | 1.92 | 0.071 | 0.98 | 0.710137 |
| 252 | O_ureido_l_serine | [M+H]+1 | POS | Mass, Spectral match | C18 | 163.059 | 0.179 | | HMDB0012271 | 21126175 | Biosynthesis of antibiotics;Other | 0.97 | 0.102 | 0.97 | 0.165 | 0.97 | 0.997 | 1.00 | 0.888621 |
| 253 | Rhamnose | [M-H]-1 | NEG | Mass, Spectral match | Hilic | 163.061 | 7.040 | C00507 | HMDB0000849 | 25310 | | 0.91 | 0.719 | 0.99 | 0.982 | 1.05 | 0.732 | 0.95 | 0.71338 |
| 254 | 1_Deoxynojirimycin | [M+H]+1 | POS | Mass, Spectral match | C18 | 163.084 | 0.532 | C16843 | HMDB0035359 | 1374 | Biosynthesis of secondary metabolites;Biosynthesis of antibiotics;Other | 0.83 | 0.240 | 1.07 | 0.732 | 0.97 | 0.812 | 1.11 | 0.647514 |
| 255 | 1_deoxynojirimycin | [M+H]+1 | POS | Mass, Spectral match | Hilic | 163.084 | 6.642 | C16843 | HMDB0035359 | 1374 | Biosynthesis of secondary metabolites;Biosynthesis of antibiotics;Cyanoamino acid metabolism;Other | 1.93 | 0.324 | 1.13 | 0.459 | 11.45 | 0.348 | 0.10 | 0.262645 |
| 256 | 2_Hydroxycinnamicacid | [M+H]+1 | POS | Mass,RT, Spectral match | Hilic | 164.047 | 4.309 | C01772 | HMDB0002641 | 637540 | Metabolic pathways;Biosynthesis of secondary metabolites;Microbial metabolism in diverse environments;Biosynthesis of antibiotics;Biosynthesis of amino acids;Biosynthesis of plant secondary metabolites;2-Oxocarboxylic acid metabolism;Degradation of aromatic compounds;Other | 0.97 | 0.903 | 0.89 | 0.597 | 0.83 | 0.750 | 1.08 | 0.572717 |
| 257 | Phenylpyruvic_acid | [M+H]+1 | POS | Mass, Spectral match | C18 | 164.047 | 0.621 | C00166 | HMDB0000205 | 997 | Metabolic pathways;Biosynthesis of secondary metabolites;Biosynthesis of antibiotics;Microbial metabolism in diverse environments;Biosynthesis of plant secondary metabolites;Biosynthesis of amino acids;Degradation of aromatic compounds;2-Oxocarboxylic acid metabolism;Tropane, piperidine and pyridine alkaloid biosynthesis;Other | 1.11 | 0.493 | 0.96 | 0.808 | 0.94 | 0.754 | 1.02 | 0.99204 |
| 258 | p_Coumaric_acid_ | [M+H]+1 | POS | Mass,RT, Spectral match | Hilic significant | 164.048 | 4.015 | C00811 | HMDB0002035 | 637542 | Metabolic pathways;Biosynthesis of secondary metabolites;Microbial metabolism in diverse environments;Biosynthesis of antibiotics;Biosynthesis of amino acids;Biosynthesis of plant secondary | 0.06 | 0.010 | 0.15 | 0.020 | 0.04 | 0.084 | 3.63 | 0.355215 |

| | | | | | | | | | | | | | | | | | | | |
|---|---|---|---|---|---|---|---|---|---|---|---|---|---|---|---|---|---|---|---|
| | | | | | | | | | | | metabolites;2-Oxocarboxylic acid metabolism;Degradation of aromatic compounds;Other | | | | | | | | |
| 259 | 1_5_Anhydro_D_mannitol | [M+H]+1 | POS | Mass, Spectral match | Hilic | 164.068 | 5.905 | C07326 | HMDB0002712 | 64960 | Metabolic pathways;Microbial metabolism in diverse environments;ABC transporters;Other | 1.61 | 0.033 | 1.61 | 0.164 | 1.57 | 0.203 | 1.02 | 0.980507 |
| 260 | Beta_D_fucose | [M+H]+1 | POS | Mass,RT, Spectral match | Hilic significant | 164.069 | 5.905 | C02095 | HMDB0003081 | 439650 | Metabolic pathways;Microbial metabolism in diverse environments;ABC transporters;Other | 1.61 | 0.033 | 1.61 | 0.164 | 1.57 | 0.203 | 1.02 | 0.980507 |
| 261 | L_MethionineS_oxide | [M+H]+1 | POS | Mass,RT, Spectral match | Hilic | 165.046 | 3.233 | C02989 | METPA0344 | NA | Cysteine and methionine metabolism | 0.09 | 0.000 | 0.30 | 0.024 | 0.20 | 0.087 | 1.50 | 0.867473 |
| 262 | Methionine_sulfoxide | [M+H]+1 | POS | Mass,RT, Spectral match | Hilic significant | 165.046 | 3.003 | | HMDB0002005 | 847 | Cysteine and methionine metabolism | 0.09 | 0.000 | 0.30 | 0.024 | 0.20 | 0.087 | 1.50 | 0.867473 |
| 263 | L_Phenylalanine | [M-H]-1 | NEG | Mass, Spectral match | C18 | 165.078 | 1.003 | C00079 | HMDB0000159 | 6140 | Metabolic pathways;Biosynthesis of secondary metabolites;Biosynthesis of antibiotics;Biosynthesis of plant secondary metabolites;Biosynthesis of amino acids;ABC transporters;Protein digestion and absorption;Central carbon metabolism in cancer;2-Oxocarboxylic acid metabolism;Biosynthesis of alkaloids derived from shikimate pathway;Aminoacyl-tRNA biosynthesis;Tropane, piperidine and pyridine alkaloid biosynthesis;Biosynthesis of alkaloids derived from ornithine, lysine and nicotinic acid;Other | 1.09 | 0.673 | 0.99 | 0.967 | 1.10 | 0.553 | 0.90 | 0.859867 |
| 264 | L_Phenylalanine | [M+H]+1 | POS | Mass,RT, Spectral match | Hilic | 165.079 | 2.481 | C00079 | HMDB0000159 | 6140 | Metabolic pathways;Biosynthesis of secondary metabolites;Biosynthesis of antibiotics;Biosynthesis of amino acids;Biosynthesis of plant secondary metabolites;ABC transporters;Protein digestion and absorption;Cyanoamino acid metabolism;2-Oxocarboxylic acid metabolism;Aminoacyl-tRNA biosynthesis;Central carbon metabolism in cancer;Other | 0.01 | 0.041 | 0.54 | 0.402 | 0.30 | 0.648 | 1.80 | 0.544865 |
| 265 | Phthalic_acid | [M+H]+1 | POS | Mass, RT, Spectral match | C18 | 166.026 | 10.573 | C01606 | HMDB0002107 | 1017 | Metabolic pathways;Biosynthesis of antibiotics;Microbial metabolism in diverse environments;ABC transporters;Degradation of aromatic compounds;Other | 0.77 | 0.099 | 0.70 | 0.059 | 0.76 | 0.943 | 0.93 | 0.585989 |

| | | | | | | | | | | | | | | | | | | | |
|---|---|---|---|---|---|---|---|---|---|---|---|---|---|---|---|---|---|---|---|
| 266 | D_Xylonate | [M+H]+1 | POS | Mass, RT, Spectral match | C18 | 166.047 | 0.523 | C00502 | METPA0050 | NA | Metabolic pathways;Other | 1.30 | 0.087 | 1.31 | 0.230 | 1.22 | 0.346 | 1.07 | 0.879746 |
| 267 | D+_Phenyllacticacid | [M-H]-1 | NEG | Mass, Spectral match | Hilic | 166.062 | 5.126 | C05607 | HMDB00563 | 643327 | Metabolic pathways;Biosynthesis of secondary metabolites;Microbial metabolism in diverse environments;Degradation of aromatic compounds;Other | 1.38 | 0.498 | 1.11 | 0.727 | 1.57 | 0.358 | 0.71 | 0.911511 |
| 268 | 3_2_Hydroxyphenylpropanoic_acid | [M-H]-1 | NEG | Mass, Spectral match | C18 | 166.062 | 4.749 | C01198 | HMDB0033752 | 873 | Metabolic pathways;Biosynthesis of secondary metabolites;Microbial metabolism in diverse environments;Degradation of aromatic compounds;Tropane, piperidine and pyridine alkaloid biosynthesis;Biosynthesis of alkaloids derived from ornithine, lysine and nicotinic acid;Other | 1.01 | 0.533 | 0.98 | 0.421 | 0.98 | 0.918 | 1.00 | 0.901884 |
| 269 | N_Carbamoylputrescine | [M+H]+1 | POS | Mass, Spectral match | Hilic | 166.076 | 2.676 | C00436 | HMDB0033458 | 502 | | 1.43 | 0.251 | 1.21 | 0.471 | 1.39 | 0.381 | 0.87 | 0.50627 |
| 270 | Homogentisic_acid | [2M+H]+1 | POS | Mass, Spectral match | Hilic | 167.036 | 7.872 | C00544 | HMDB0000130 | 780 | Biosynthesis of antibiotics;Other | 0.95 | 0.873 | 0.56 | 0.042 | 0.89 | 0.811 | 0.63 | 0.097524 |
| 271 | Pyridoxal | [M+H]+1 | POS | Mass, Spectral match | C18 | 167.058 | 0.543 | C00250 | HMDB0001545 | 1050 | Metabolic pathways;Biosynthesis of secondary metabolites;Biosynthesis of antibiotics;Microbial metabolism in diverse environments;Other | 0.92 | 0.574 | 1.29 | 0.202 | 0.85 | 0.513 | 1.52 | 0.025313 |
| 272 | Pyridoxamine | [M+H]+1 | POS | Mass, Spectral match | Hilic | 167.083 | 2.608 | C00534 | HMDB0001431 | 1052 | | 1.04 | 0.875 | 1.12 | 0.568 | 1.12 | 0.543 | 1.00 | 0.934979 |
| 635 | 6_alpha_D_Glucosaminyl_1D_myo_inositol | [M+H]+1 | POS | Mass, Spectral match | Hilic | 341.132 | 10.483 | C15658 | HMDB0011668 | NA | | 1.31 | 0.572 | 1.03 | 0.952 | 1.03 | 0.616 | 1.00 | 0.966649 |
| 274 | Uric_acid | [M+Cl]-1 | NEG | Mass, RT, Spectral match | C18 | 168.027 | 0.528 | C00366 | HMDB0000289 | 1175 | | 1.01 | 0.948 | 0.84 | 0.583 | 0.70 | 0.871 | 1.21 | 0.894381 |
| 275 | Uricacid | [M+H]+1 | POS | Mass, Spectral match | Hilic | 168.028 | 9.250 | C00366 | HMDB0000289 | 1175 | Metabolic pathways;Microbial metabolism in diverse environments;Other | 0.98 | 0.928 | 0.92 | 0.811 | 0.77 | 0.663 | 1.20 | 0.823104 |
| 637 | Lactulose | [M+H]+1 | POS | Mass, Spectral match | Hilic significant | 342.117 | 9.774 | C07064 | HMDB0000740 | 11333 | Metabolic pathways;Biosynthesis of secondary metabolites;ABC transporters;Other | 1.00 | 0.990 | 2.09 | 0.039 | 1.95 | 0.092 | 1.07 | 0.77764 |
| 277 | Pyridoxine | [M+H]+1 | POS | Mass, Spectral match | C18 | 169.074 | 0.584 | C00314 | HMDB0000239 | 1054 | Metabolic pathways;Microbial metabolism in diverse environments;Neuroactive ligand-receptor interaction;Other | 1.33 | 0.262 | 1.19 | 0.549 | 0.97 | 0.972 | 1.23 | 0.56557 |
| 278 | 1_Methylhistidine | [M+H]+1 | POS | Mass,RT, Spectral match | Hilic | 169.085 | 1.559 | C01152 | HMDB0000001 | 92105 | Histidine metabolism | 1.99 | 0.094 | 1.44 | 0.196 | 1.52 | 0.243 | 0.95 | 0.956392 |

| | | | | | | | | | | | | | | | | | | | |
|---|---|---|---|---|---|---|---|---|---|---|---|---|---|---|---|---|---|---|---|
| 279 | Dihydroxyacetone_phosphate | [M+H]+1 | POS | Mass, Spectral match | C18 | 169.998 | 18.683 | C00111 | HMDB0001473 | 668 | Metabolic pathways;Biosynthesis of secondary metabolites;Biosynthesis of antibiotics;Microbial metabolism in diverse environments;Biosynthesis of plant secondary metabolites;Biosynthesis of amino acids;Biosynthesis of alkaloids derived from shikimate pathway;Biosynthesis of alkaloids derived from ornithine, lysine and nicotinic acid;Other | 1.01 | 0.877 | 0.89 | 0.153 | 0.94 | 0.538 | 0.95 | 0.553138 |
| 280 | 2_Aminobenzoic_acid | [M+H]+1 | POS | Mass, Spectral match | Hilic | 172.015 | 9.185 | C00108 | HMDB0001123 | 227 | | 0.92 | 0.806 | 0.67 | 0.270 | 0.75 | 0.581 | 0.89 | 0.804507 |
| 281 | Hydantoin_5_propionate | [M+H]+1 | POS | Mass, Spectral match | Hilic | 172.048 | 6.569 | C05565 | HMDB0001212 | 782 | Histidine metabolism | 1.25 | 0.374 | 0.95 | 0.844 | 0.70 | 0.391 | 1.35 | 0.331278 |
| 282 | Tyramine | [M+H]+1 | POS | Mass, Spectral match | Hilic | 172.053 | 7.072 | C00483 | HMDB0000306 | 5610 | Metabolic pathways;Microbial metabolism in diverse environments;Degradation of aromatic compounds;Other | 1.30 | 0.347 | 1.38 | 0.315 | 1.39 | 0.210 | 0.99 | 0.791913 |
| 283 | Gly_Pro | [M+H]+1 | POS | Mass,RT, Spectral match | Hilic | 172.084 | 2.026 | | HMDB0000721 | 79101 | | 2.79 | 0.036 | 2.23 | 0.031 | 3.10 | 0.026 | 0.72 | 0.273588 |
| 284 | Prolylglycine | [M+H]+1 | POS | Mass,RT, Spectral match | Hilic | 172.085 | 1.499 | | HMDB0011178 | 98206 | | 1.53 | 0.100 | 1.23 | 0.382 | 1.39 | 0.343 | 0.88 | 0.533672 |
| 285 | Gly_Pro | [M+H]+1 | POS | Mass,RT, Spectral match | Hilic significant | 172.085 | 2.026 | | HMDB0000721 | 79101 | | 2.79 | 0.036 | 2.23 | 0.031 | 3.10 | 0.026 | 0.72 | 0.273588 |
| 286 | L_Arginine | [M+H]+1 | POS | Mass, Spectral match | Hilic | 173.104 | 4.245 | C00062 | HMDB0000517 | 6322 | | 0.93 | 0.808 | 1.14 | 0.735 | 1.11 | 0.806 | 1.03 | 0.90318 |
| 287 | L_Arginine | [M+H]+1 | POS | Mass, Spectral match | C18 | 173.105 | 0.820 | C00062 | HMDB0000517 | 6322 | Tropane, piperidine and pyridine alkaloid biosynthesis;Biosynthesis of alkaloids derived from ornithine, lysine and nicotinic acid | 0.70 | 0.484 | 0.95 | 0.912 | 0.86 | 0.720 | 1.10 | 0.933956 |
| 288 | L_Indospicine | [M+H]+1 | POS | Mass, Spectral match | Hilic | 173.116 | 6.948 | NA | #N/A | #N/A | | 1.12 | 0.499 | 0.98 | 0.909 | 0.90 | 0.976 | 1.09 | 0.703933 |
| 289 | 4_Nitrophenol | [M+H]+1 | POS | Mass, Spectral match | Hilic | 173.994 | 2.159 | C00870 | HMDB0001232 | 980 | | 3.36 | 0.264 | 1.52 | 0.594 | 2.31 | 0.116 | 0.66 | 0.49265 |
| 290 | Homocysteine | [M-H]-1 | NEG | Mass, Spectral match | C18 | 173.998 | 1.687 | C00155 | HMDB0000742 | 778 | | 0.51 | 0.107 | 0.36 | 0.039 | 1.59 | 0.044 | 0.23 | 0.002023 |
| 291 | cis_Aconitic_acid | [M-H]-1 | NEG | Mass, Spectral match | C18 | 174.015 | 0.586 | C00417 | HMDB0000072 | 643757 | | 1.19 | 0.772 | 1.00 | 0.998 | 1.19 | 0.176 | 0.84 | 0.557296 |
| 695 | Cholic_acid | [M+H]+1 | POS | Mass,RT, Spectral match | Hilic significant | 408.288 | 1.183 | C00695 | HMDB0000619 | 221493 | Metabolic pathways;Other | 1.91 | 0.239 | 2.48 | 0.035 | 4.02 | 0.041 | 0.62 | 0.444355 |
| 293 | formyl_isoglutamine | [M+H]+1 | POS | Mass,RT, Spectral match | Hilic | 174.064 | 12.163 | C16674 | METPA1416 | NA | Metabolic pathways;Histidine metabolism | 0.79 | 0.746 | 0.23 | 0.181 | 0.30 | 0.137 | 0.77 | 0.937187 |

| | | | | | | | | | | | | | | | | | | | |
|---|---|---|---|---|---|---|---|---|---|---|---|---|---|---|---|---|---|---|---|
| 294 | Citrulline | [M+H]+1 | POS | Mass, Spectral match | C18 | 174.090 | 15.630 | C00327 | HMDB0000904 | 9750 | Other | 1.11 | 0.530 | 0.74 | 0.155 | 1.13 | 0.563 | 0.65 | 0.478263 |
| 295 | N_Acetylornithine | [M+H]+1 | POS | Mass, Spectral match | C18 | 174.100 | 0.607 | C00437 | HMDB0003357 | 439232 | Metabolic pathways;Biosynthesis of secondary metabolites;Biosynthesis of antibiotics;Biosynthesis of amino acids;2-Oxocarboxylic acid metabolism;Other | 2.16 | 0.165 | 1.52 | 0.214 | 2.10 | 0.057 | 0.72 | 0.367387 |
| 296 | N_Acetyl_ornithine_ | [M+H]+1 | POS | Mass, Spectral match | Hilic significant | 174.101 | 6.935 | C00437 | HMDB0003357 | 439232 | Metabolic pathways;Biosynthesis of secondary metabolites;Biosynthesis of antibiotics;Biosynthesis of amino acids;2-Oxocarboxylic acid metabolism;Other | 1.04 | 0.888 | 1.35 | 0.215 | 0.61 | 0.966 | 2.23 | 0.004115 |
| 297 | N_Acetyl_ornithine_ | [M+H]+1 | POS | Mass, Spectral match | C18 significant | 174.101 | 0.610 | C00437 | HMDB0003357 | 439232 | Metabolic pathways;Biosynthesis of secondary metabolites;Biosynthesis of antibiotics;Biosynthesis of amino acids;2-Oxocarboxylic acid metabolism;Other | 2.16 | 0.165 | 1.52 | 0.214 | 2.10 | 0.057 | 0.72 | 0.367387 |
| 298 | Allantoic_acid | [M-H]-1 | NEG | Mass, Spectral match | C18 | 175.047 | 0.579 | C00499 | HMDB0001209 | 203 | | 0.80 | 0.608 | 0.53 | 0.219 | 0.85 | 0.629 | 0.63 | 0.66633 |
| 299 | Allantoic_acid | [M-H]-1 | NEG | Mass, Spectral match | Hilic | 175.048 | 6.002 | C00499 | HMDB0001209 | 203 | Metabolic pathways;Histidine metabolism;Other | 0.52 | 0.087 | 0.57 | 0.172 | 0.47 | 0.782 | 1.19 | 0.971781 |
| 300 | N_Acetyl_L_aspartic_acid | [M-H]-1 | NEG | Mass, Spectral match | C18 | 175.049 | 0.583 | C01042 | HMDB0000812 | 65065 | Metabolic pathways;Other | 3.40 | 0.213 | 2.23 | 0.249 | 3.96 | 0.194 | 0.56 | 0.686546 |
| 301 | N_Amidino_L_aspartate | [M-H]-1 | NEG | Mass, Spectral match | Hilic | 175.059 | 14.850 | C03139 | HMDB0003157 | 439918 | | 3.04 | 0.185 | 2.94 | 0.119 | 2.13 | 0.130 | 1.38 | 0.895648 |
| 302 | L_Citrulline | [M+H]+1 | POS | Mass, Spectral match | Hilic | 175.095 | 8.550 | C00327 | HMDB0000904 | 9750 | Metabolic pathways;Biosynthesis of secondary metabolites;Biosynthesis of antibiotics;Biosynthesis of amino acids;Other | 1.19 | 0.469 | 1.14 | 0.620 | 1.23 | 0.411 | 0.93 | 0.794676 |
| 303 | Monodehydroascorbate | [M-H]-1 | NEG | Mass, Spectral match | C18 | 176.031 | 0.533 | C01041 | METPA0114 | NA | | 3.08 | 0.315 | 0.29 | 0.079 | 0.23 | 0.073 | 1.26 | 0.883531 |
| 304 | Tyramine | [M+H]+1 | POS | Mass, Spectral match | C18 | 176.047 | 8.894 | C00483 | HMDB0000306 | 5610 | | 1.01 | 0.908 | 1.01 | 0.946 | 0.96 | 0.463 | 1.05 | 0.813063 |
| 305 | Canavanine | [M-H]-1 | NEG | Mass, RT, Spectral match | C18 | 177.100 | 0.532 | C00308 | HMDB0002706 | 439202 | | 1.28 | 0.402 | 1.00 | 0.984 | 1.01 | 0.343 | 0.99 | 0.776487 |
| 306 | __Menthone | [M+H]+1 | POS | Mass, RT, Spectral match | C18 | 177.126 | 1.972 | C00843 | HMDB0035162 | 26447 | | 0.82 | 0.828 | 0.58 | 0.595 | 1.07 | 0.984 | 0.55 | 0.629717 |
| 307 | Gluconolactone | [M+Cl]-1 | NEG | Mass, Spectral match | C18 | 178.047 | 0.511 | C00198 | HMDB0000150 | 7027 | Metabolic pathways;Biosynthesis of secondary metabolites;Biosynthesis of antibiotics;Microbial metabolism in diverse environments;Other | 1.18 | 0.484 | 1.02 | 0.954 | 1.58 | 0.136 | 0.64 | 0.227321 |

| | | | | | | | | | | | | | | | | | | | |
|---|---|---|---|---|---|---|---|---|---|---|---|---|---|---|---|---|---|---|---|
| 308 | Isoxanthopterin | [M+H]+1 | POS | Mass, Spectral match | Hilic | 179.044 | 3.711 | C03975 | HMDB0000704 | 10729 | | 0.90 | 0.619 | 1.26 | 0.388 | 0.92 | 0.980 | 1.38 | 0.369209 |
| 309 | Scytalone | [M+H]+1 | POS | Mass,RT, Spectral match | Hilic | 179.094 | 2.279 | C09642 | HMDB0005199 | 54456 | Biosynthesis of secondary metabolites;2-Oxocarboxylic acid metabolism;Other | 1.05 | 0.612 | 1.03 | 0.756 | 0.99 | 0.768 | 1.03 | 0.323693 |
| 310 | O_Phosphoethanolamine | [M+H]+1 | POS | Mass, Spectral match | C18 | 179.982 | 18.744 | C00346 | HMDB0000224 | 1015 | | 0.91 | 0.491 | 0.75 | 0.020 | 0.66 | 0.003 | 1.14 | 0.034463 |
| 311 | 2_Keto_glutaramic_acid | [M+H]+1 | POS | Mass, RT, Spectral match | C18 | 180.007 | 18.561 | C00940 | HMDB0001552 | 48 | | 1.10 | 0.383 | 1.06 | 0.565 | 0.89 | 0.198 | 1.19 | 0.119621 |
| 312 | L_Histidinol | [M+H]+1 | POS | Mass, RT, Spectral match | C18 | 180.052 | 5.168 | C00860 | HMDB0003431 | 165271 | | 1.68 | 0.305 | 0.91 | 0.583 | 1.03 | 0.451 | 0.88 | 0.640862 |
| 757 | Diacetoxyscirpenol | [M+H]+1 | POS | Mass,RT, Spectral match | Hilic | 744.410 | 1.100 | C09662 | HMDB0035104 | 422111 | | 0.89 | 0.294 | 1.01 | 0.955 | 0.90 | 0.867 | 1.12 | 0.396542 |
| 314 | 1D_chiro_inositol | [M+Cl]-1 | POS | Mass,RT, Spectral match | Hilic | 180.063 | 6.049 | C19891 | METPA1881 | NA | Metabolic pathways;Biosynthesis of secondary metabolites;Microbial metabolism in diverse environments;Biosynthesis of antibiotics;Biosynthesis of plant secondary metabolites;ABC transporters;Central carbon metabolism in cancer;Other | 1.28 | 0.002 | 1.39 | 0.047 | 1.34 | 0.092 | 1.04 | 0.877846 |
| 315 | L_Threo_3_Phenylserine | [M+H]+1 | POS | Mass,RT, Spectral match | Hilic significant | 181.074 | 4.633 | | HMDB0002184 | 12314153 | Metabolic pathways;Biosynthesis of secondary metabolites;Biosynthesis of antibiotics;Biosynthesis of amino acids;Biosynthesis of plant secondary metabolites;Protein digestion and absorption;Cyanoamino acid metabolism;2-Oxocarboxylic acid metabolism;Aminoacyl-tRNA biosynthesis;Central carbon metabolism in cancer;Other | 1.69 | 0.066 | 1.75 | 0.082 | 1.92 | 0.062 | 0.91 | 0.737235 |
| 316 | 9_Methyluricacid | [M-H]-1 | NEG | Mass, Spectral match | Hilic | 182.043 | 3.903 | | HMDB0001973 | 108714 | Other | 3.79 | 0.237 | 4.82 | 0.170 | 1.61 | 0.754 | 3.00 | 0.397985 |
| 317 | DL_4_Hydroxyphenyllacticacid | [M-H]-1 | NEG | Mass,RT, Spectral match | Hilic | 182.057 | 9.043 | C03672 | HMDB0000755 | 9378 | Metabolic pathways;Biosynthesis of secondary metabolites;Microbial metabolism in diverse environments;Biosynthesis of antibiotics;Degradation of aromatic compounds;Other | 0.99 | 0.979 | 0.79 | 0.209 | 0.94 | 0.556 | 0.84 | 0.718642 |
| 318 | Indoleacetaldehyde | [M-H]-1 | NEG | Mass, Spectral match | C18 | 182.057 | 1.585 | C00637 | HMDB0001190 | 800 | Metabolic pathways;Biosynthesis of secondary metabolites;Biosynthesis of antibiotics;Microbial metabolism in diverse environments;Degradation of aromatic compounds;Other | 0.98 | 0.947 | 0.86 | 0.462 | 1.01 | 0.471 | 0.85 | 0.813945 |

| | | | | | | | | | | | | | | | | | | | |
|---|---|---|---|---|---|---|---|---|---|---|---|---|---|---|---|---|---|---|---|
| 319 | Diethylphosphate | [M+H]+1 | POS | Mass, Spectral match | C18 | 182.071 | 6.570 | C06608 | HMDB0012209 | 654 | | 0.87 | 0.067 | 0.90 | 0.215 | 0.83 | 0.358 | 1.08 | 0.337107 |
| 320 | Mannitol | [M-H]-1 | NEG | Mass, Spectral match | C18 | 182.078 | 0.494 | C00392 | HMDB0000765 | 6251 | | 1.51 | 0.041 | 1.03 | 0.852 | 1.20 | 0.100 | 0.87 | 0.415925 |
| 321 | Dulcitol | [M+H]+1 | POS | Mass,RT, Spectral match | Hilic | 182.078 | 5.888 | C01697 | HMDB0000107 | 11850 | | 1.42 | 0.050 | 1.29 | 0.277 | 1.45 | 0.074 | 0.89 | 0.565497 |
| 322 | 3_5_Dihydroxy_phenylglycine | [M+H]+1 | POS | Mass, Spectral match | Hilic | 183.052 | 2.957 | C06689 | HMDB0014653 | 14969 | Metabolic pathways;Microbial metabolism in diverse environments;Biosynthesis of antibiotics;Other | 0.43 | 0.052 | 0.45 | 0.061 | 0.42 | 0.239 | 1.07 | 0.986724 |
| 323 | 4_Pyridoxic_acid | [M-H]-1 | NEG | Mass, RT, Spectral match | C18 | 183.052 | 0.666 | C00847 | HMDB0000017 | 6723 | Metabolic pathways;Biosynthesis of antibiotics;Microbial metabolism in diverse environments;Other | 1.21 | 0.386 | 0.82 | 0.351 | 1.13 | 0.570 | 0.73 | 0.617836 |
| 324 | 4_Pyridoxicacid | [M+H]+1 | POS | Mass, Spectral match | Hilic | 183.052 | 9.290 | C00847 | HMDB0000017 | 6723 | Metabolic pathways;Microbial metabolism in diverse environments;Biosynthesis of antibiotics;Other | 1.12 | 0.657 | 0.67 | 0.103 | 0.80 | 0.854 | 0.84 | 0.657572 |
| 325 | Phosphorylcholine | [M+H]+1 | POS | Mass, RT, Spectral match | C18 | 183.066 | 10.817 | C00588 | HMDB0001565 | 1014 | | 1.05 | 0.816 | 0.85 | 0.504 | 1.06 | 0.642 | 0.81 | 0.878722 |
| 326 | Phosphorylcholine | [M+H]+1 | POS | Mass,RT, Spectral match | Hilic | 183.066 | 14.806 | C00588 | HMDB0001565 | 1014 | | 1.04 | 0.850 | 0.93 | 0.780 | 0.93 | 0.678 | 1.00 | 0.919256 |
| 327 | Ecgonine | [M+H]+1 | POS | Mass, Spectral match | Hilic | 185.105 | 9.119 | C10858 | HMDB0006548 | 443003 | Other | 1.96 | 0.252 | 1.77 | 0.198 | 1.73 | 0.337 | 1.02 | 0.813054 |
| 328 | D___3_Phosphoglyceric_acid_disodium_salt | [M-H]-1 | NEG | Mass, Spectral match | C18 | 185.992 | 0.522 | C00197 | HMDB60180 | 439183 | Metabolic pathways;Biosynthesis of secondary metabolites;Biosynthesis of antibiotics;Microbial metabolism in diverse environments;Biosynthesis of plant secondary metabolites;Biosynthesis of amino acids;Central carbon metabolism in cancer;Biosynthesis of alkaloids derived from shikimate pathway;Biosynthesis of alkaloids derived from ornithine, lysine and nicotinic acid;Other | 1.24 | 0.611 | 1.00 | 0.998 | 0.77 | 0.445 | 1.30 | 0.789197 |
| 329 | 3_Phosphoglycericacid | [M-H]-1 | NEG | Mass, Spectral match | Hilic | 185.994 | 6.796 | C00597 | HMDB0000807 | 724 | | 0.75 | 0.701 | 0.69 | 0.643 | 1.86 | 0.152 | 0.37 | 0.194375 |
| 330 | Phosphoserine | [M+H]+1 | POS | Mass, Spectral match | C18 | 186.015 | 4.188 | C01005 | HMDB0000272 | 68841 | | 0.62 | 0.595 | 1.31 | 0.745 | 0.64 | 0.370 | 2.04 | 0.792056 |
| 331 | Ala_Pro | [M+H]+1 | POS | Mass, Spectral match | Hilic | 186.100 | 1.205 | | HMDB0028695 | NA | | 1.72 | 0.103 | 1.08 | 0.725 | 1.63 | 0.255 | 0.66 | 0.300923 |
| 332 | 3_Isopropylcatechol | [M-H]-1 | NEG | Mass, Spectral match | Hilic | 187.052 | 9.167 | C01014 | #N/A | 16498 | | 1.43 | 0.377 | 1.17 | 0.621 | 1.01 | 0.581 | 1.15 | 0.71643 |

| | | | | | | | | | | | | | | | | | | | |
|---|---|---|---|---|---|---|---|---|---|---|---|---|---|---|---|---|---|---|---|
| 333 | Indole_3_acrylicacid | [M-H]-1 | NEG | Mass, Spectral match | Hilic | 187.063 | 2.562 | | HMDB0000734 | 15030923 | Microbial metabolism in diverse environments;Other | 1.24 | 0.199 | 1.14 | 0.402 | 1.15 | 0.507 | 0.99 | 0.819929 |
| 334 | Indole_3_acrylic_acid | [M+H]+1 | POS | Mass,RT, Spectral match | Hilic | 187.063 | 2.333 | | HMDB0000734 | 15030923 | Microbial metabolism in diverse environments;Other | 0.03 | 0.047 | 0.40 | 0.313 | 0.24 | 0.603 | 1.65 | 0.91541 |
| 335 | trans_3_Indoleacrylicacid | [M+H]+1 | POS | Mass, Spectral match | C18 | 187.063 | 1.651 | | HMDB0000734 | 15030923 | Microbial metabolism in diverse environments;Other | 1.22 | 0.316 | 1.01 | 0.943 | 1.04 | 0.704 | 0.97 | 0.905937 |
| 336 | Homo_L_arginine | [M-H]-1 | NEG | Mass,RT, Spectral match | Hilic | 187.120 | 4.558 | C01924 | HMDB0000670 | 9085 | Metabolic pathways;Other | 1.30 | 0.705 | 1.55 | 0.492 | 2.13 | 0.075 | 0.73 | 0.456055 |
| 337 | Homo_L_arginine | [M+H]+1 | POS | Mass, Spectral match | C18 | 187.121 | 0.630 | C01924 | HMDB0000670 | 9085 | Metabolic pathways;Other | 1.36 | 0.327 | 1.24 | 0.369 | 1.09 | 0.109 | 1.14 | 0.51879 |
| 338 | N8_Acetylspermidine | [M+H]+1 | POS | Mass, Spectral match | Hilic | 187.168 | 1.288 | C01029 | HMDB0002189 | 123689 | | 4.39 | 0.277 | 0.54 | 0.196 | 1.05 | 0.872 | 0.51 | 0.24464 |
| 339 | p_Cresol_sulfate | [M-H]-1 | NEG | Mass, Spectral match | Hilic | 188.013 | 1.226 | | HMDB0011635 | 4615423 | Microbial metabolism in diverse environments;Degradation of aromatic compounds;Other | 0.73 | 0.626 | 1.31 | 0.734 | 5.84 | 0.312 | 0.22 | 0.494161 |
| 340 | 6_Hydroxy_2_naphthoic acid | [M-H]-1 | NEG | Mass,RT, Spectral match | Hilic | 188.048 | 9.222 | C03203 | METPA0374 | NA | Metabolic pathways;Microbial metabolism in diverse environments;Degradation of aromatic compounds;Other | 1.99 | 0.163 | 1.64 | 0.155 | 1.34 | 0.413 | 1.23 | 0.580926 |
| 341 | 2_oxosuberate | [M-H]-1 | NEG | Mass,RT, Spectral match | Hilic | 188.068 | 3.853 | | HMDB0000325 | 22328017 | Microbial metabolism in diverse environments;2-Oxocarboxylic acid metabolism;Other | 0.96 | 0.878 | 1.83 | 0.202 | 1.90 | 0.576 | 0.96 | 0.432138 |
| 342 | N_Acetyl_L_glutamine | [M+H]+1 | POS | Mass, Spectral match | Hilic | 188.079 | 17.434 | | HMDB0006029 | 25561 | Biosynthesis of antibiotics;Other | 1.15 | 0.581 | 1.38 | 0.427 | 1.83 | 0.075 | 0.76 | 0.327302 |
| 343 | N_Acetylglutamine | [M+H]+1 | POS | Mass, Spectral match | C18 | 188.079 | 0.548 | | HMDB0006029 | 25561 | Biosynthesis of antibiotics;Other | 0.91 | 0.606 | 1.25 | 0.555 | 1.48 | 0.106 | 0.84 | 0.412943 |
| 344 | N_Acetylglutamine | [M+H]+1 | POS | Mass, Spectral match | Hilic | 188.079 | 17.611 | | HMDB0006029 | 25561 | Biosynthesis of antibiotics;Other | 1.44 | 0.199 | 1.20 | 0.710 | 1.88 | 0.347 | 0.64 | 0.996999 |
| 345 | Ephedrine | [M-H]-1 | NEG | Mass, Spectral match | C18 | 188.104 | 6.312 | C01575 | HMDB0015451 | 9294 | | 0.88 | 0.295 | 1.05 | 0.719 | 0.89 | 0.274 | 1.17 | 0.149815 |
| 346 | 6_Acetamido_3_aminohexanoate | [M+H]+1 | POS | Mass, Spectral match | Hilic | 188.115 | 1.267 | C05548 | HMDB0012150 | 194080 | Lysine degradation | 1.57 | 0.037 | 1.09 | 0.629 | 1.49 | 0.171 | 0.74 | 0.121493 |
| 347 | Glycylleucine | [M+H]+1 | POS | Mass, Spectral match | C18 | 188.116 | 1.438 | C02155 | HMDB0000759 | 92843 | Other | 1.49 | 0.176 | 1.02 | 0.935 | 1.21 | 0.595 | 0.84 | 0.341656 |
| 348 | N6_Acetyl_L_lysine | [M+H]+1 | POS | Mass, Spectral match | Hilic significant | 188.116 | 1.267 | C02727 | HMDB0000206 | 92832 | Lysine degradation | 1.57 | 0.037 | 1.09 | 0.629 | 1.49 | 0.171 | 0.74 | 0.121493 |
| 349 | N6_Acetyl_L_lysine | [M+H]+1 | POS | Mass, Spectral match | C18 significant | 188.116 | 0.760 | C02727 | HMDB0000206 | 92832 | Other | 1.75 | 0.154 | 0.94 | 0.898 | 2.22 | 0.068 | 0.43 | 0.042149 |
| 350 | N6N6N6_Trimethyl_L_lysine | [M+H]+1 | POS | Mass,RT, Spectral match | Hilic | 188.152 | 1.450 | C03793 | HMDB0001325 | 440120 | Metabolic pathways;Lysine degradation;Other | 1.09 | 0.693 | 1.14 | 0.667 | 0.73 | 0.562 | 1.56 | 0.292147 |
| 351 | Kynurenic_acid | [M-H]-1 | NEG | Mass,RT, Spectral match | Hilic | 189.048 | #N/A | C01717 | HMDB0000715 | 3845 | Other | 1.02 | 0.912 | 1.05 | 0.767 | 0.89 | 0.665 | 1.18 | 0.260177 |
| 352 | 3_Indolepropionicacid | [M+H]+1 | POS | Mass, Spectral | Hilic | 189.079 | 2.658 | | HMDB0002302 | 3744 | | 1.28 | 0.240 | 1.11 | 0.470 | 1.13 | 0.500 | 0.98 | 0.922655 |

| | | | | | | | | | | | | | | | | | | | |
|---|---|---|---|---|---|---|---|---|---|---|---|---|---|---|---|---|---|---|---|
| | | | | match | | | | | | | | | | | | | | | |
| 353 | L_Homocitrulline | [M+H]+1 | POS | Mass, Spectral match | Hilic | 189.111 | 7.855 | C02427 | HMDB0000679 | 65072 | | 1.09 | 0.756 | 0.95 | 0.820 | 1.05 | 0.540 | 0.90 | 0.998673 |
| 354 | 8_Methylthiooctanaldoxime | [M+H]+1 | POS | Mass, Spectral match | C18 | 189.118 | 10.977 | C17251 | METPA1776 | NA | 2-Oxocarboxylic acid metabolism;Other | 1.10 | 0.665 | 0.81 | 0.346 | 0.86 | 0.622 | 0.94 | 0.684648 |
| 355 | 4_Nitrocatechol | [M-H]-1 | NEG | Mass, RT, Spectral match | C18 | 189.992 | 2.651 | C02235 | HMDB0002916 | 3505109 | | 1.91 | 0.366 | 1.73 | 0.221 | 0.99 | 0.518 | 1.75 | 0.267786 |
| 356 | 34_dihydroxybenzenesulfonicacid | [M-H]-1 | NEG | Mass, Spectral match | Hilic | 189.993 | 0.946 | C01161 | HMDB0001336 | 547 | Microbial metabolism in diverse environments;Other | 0.84 | 0.084 | 1.01 | 0.890 | 0.90 | 0.399 | 1.13 | 0.22458 |
| 357 | 2_6_Diaminoheptanedioicacid | [M+H]+1 | POS | Mass, Spectral match | Hilic | 190.095 | 1.935 | C04421 | HMDB0012267 | 25202447 | Metabolic pathways;Biosynthesis of secondary metabolites;Microbial metabolism in diverse environments;Biosynthesis of antibiotics;Biosynthesis of amino acids;Other | 3.09 | 0.080 | 1.78 | 0.304 | 2.98 | 0.166 | 0.60 | 0.24329 |
| 358 | Quinic_acid | [M+H]+1 | POS | Mass, Spectral match | Hilic | 191.058 | 2.649 | C00296 | #N/A | 6508 | Metabolic pathways;Other | 1.12 | 0.553 | 0.78 | 0.351 | 0.52 | 0.236 | 1.50 | 0.373643 |
| 359 | 5_Hydroxyindoleacetic_acid | [M+H]+1 | POS | Mass, RT, Spectral match | C18 significant | 191.059 | 0.950 | C05635 | HMDB0000763 | 1826 | Metabolic pathways;Other | 0.98 | 0.897 | 0.68 | 0.229 | 0.41 | 0.125 | 1.68 | 0.330097 |
| 360 | N_Acetyl_L_methionine | [M-H]-1 | NEG | Mass, Spectral match | Hilic | 191.061 | 7.309 | C02712 | HMDB0011745 | 6180 | | 1.20 | 0.648 | 0.80 | 0.290 | 0.88 | 0.916 | 0.91 | 0.408242 |
| 361 | N_Acetyl_L_methionine | [M-H]-1 | NEG | Mass, RT, Spectral match | C18 | 191.061 | 2.078 | C02712 | HMDB0011745 | 6180 | | 1.16 | 0.642 | 1.04 | 0.861 | 1.05 | 0.707 | 0.99 | 0.436786 |
| 362 | Pyridoxamine | [M+H]+1 | POS | Mass, RT, Spectral match | C18 | 191.079 | 0.513 | C00534 | HMDB0001431 | 1052 | Biosynthesis of secondary metabolites;Biosynthesis of antibiotics;Other | 0.80 | 0.363 | 0.80 | 0.217 | 0.75 | 0.815 | 1.08 | 0.563512 |
| 363 | Citric_acid | [M+Cl]-1 | NEG | Mass, RT, Spectral match | C18 | 192.026 | 0.520 | C00158 | HMDB0000094 | 311 | Metabolic pathways;Biosynthesis of secondary metabolites;Biosynthesis of antibiotics;Microbial metabolism in diverse environments;Biosynthesis of plant secondary metabolites;Biosynthesis of amino acids;Central carbon metabolism in cancer;2-Oxocarboxylic acid metabolism;Biosynthesis of alkaloids derived from shikimate pathway;Biosynthesis of alkaloids derived from ornithine, lysine and nicotinic acid;Other | 1.37 | 0.526 | 0.69 | 0.395 | 1.09 | 0.475 | 0.63 | 0.423502 |
| 364 | Citric_acid | [M-H]-1 | NEG | Mass, Spectral match | Hilic significant | 192.027 | 2.633 | C00158 | HMDB0000094 | 311 | Metabolic pathways;Biosynthesis of secondary metabolites;Microbial metabolism in diverse environments;Biosynthesis of | 13.09 | 0.073 | 4.36 | 0.240 | 13.45 | 0.216 | 0.32 | 0.4352 |

| | | | | | | | | | | | | | | | | | | | |
|---|---|---|---|---|---|---|---|---|---|---|---|---|---|---|---|---|---|---|---|
| | | | | | | | | | | | antibiotics;Biosynthesis of amino acids;Biosynthesis of plant secondary metabolites;2-Oxocarboxylic acid metabolism;Central carbon metabolism in cancer;Other | | | | | | | | |
| 365 | 45_Dihydroorotic_acid | [M-H]-1 | NEG | Mass, Spectral match | Hilic | 193.001 | 8.132 | C00337 | HMDB0000528 | 648 | | 0.95 | 0.950 | 3.38 | 0.216 | 3.89 | 0.202 | 0.87 | 0.834797 |
| 366 | 2_Methylhippuricacid | [M-H]-1 | NEG | Mass, Spectral match | Hilic | 193.073 | 6.511 | C01586 | HMDB0011723 | 91637 | Other | 1.05 | 0.907 | 0.75 | 0.536 | 1.14 | 0.199 | 0.65 | 0.208307 |
| 367 | Hippuric_acid | [M-H]-1 | NEG | Mass, RT, Spectral match | C18 | 193.073 | 4.251 | C01586 | HMDB0000714 | 464 | Other | 1.11 | 0.795 | 0.88 | 0.797 | 1.43 | 0.157 | 0.61 | 0.178682 |
| 368 | Indoleacetaldehyde | [M+H]+1 | POS | Mass, Spectral match | Hilic | 194.036 | 10.216 | C00637 | HMDB0001190 | 800 | | 1.27 | 0.390 | 0.92 | 0.783 | 1.57 | 0.021 | 0.59 | 0.087314 |
| 369 | 2_Dehydro_D_gluconate | [M-H]-1 | NEG | Mass, Spectral match | Hilic | 194.042 | 1.598 | C15673 | HMDB0011732 | 50 | Metabolic pathways;Microbial metabolism in diverse environments;ABC transporters;Other | 2.05 | 0.143 | 1.47 | 0.324 | 1.91 | 0.208 | 0.77 | 0.640681 |
| 370 | D_Glucuronic_acid | [M-H]-1 | NEG | Mass, RT, Spectral match | C18 | 194.042 | 0.509 | C00191 | HMDB0000127 | 444791 | Metabolic pathways;Microbial metabolism in diverse environments;ABC transporters;Other | 2.60 | 0.312 | 0.38 | 0.027 | 0.35 | 0.115 | 1.07 | 0.927031 |
| 371 | Galacturonic_acid | [M-H]-1 | NEG | Mass, Spectral match | Hilic significant | 194.043 | 6.840 | C08348 | HMDB0002545 | 84740 | Metabolic pathways;Microbial metabolism in diverse environments;ABC transporters;Other | 0.92 | 0.640 | 0.63 | 0.027 | 0.84 | 0.508 | 0.76 | 0.137334 |
| 372 | Butylparaben | [M+H]+1 | POS | Mass, RT, Spectral match | C18 | 194.094 | 8.470 | D01420 | #N/A | 7184 | | 1.04 | 0.688 | 1.03 | 0.717 | 1.01 | 0.337 | 1.02 | 0.842086 |
| 373 | 6_Hydroxypseudooxynicotine | [M-H]-1 | NEG | Mass, RT, Spectral match | C18 | 194.106 | 7.705 | C01297 | METPA0168 | NA | Microbial metabolism in diverse environments;Other | 1.03 | 0.880 | 1.06 | 0.716 | 1.06 | 0.894 | 1.00 | 0.845683 |
| 374 | Gluconic_acid | [M-H]-1 | NEG | Mass, RT, Spectral match | C18 | 196.057 | 0.504 | C00257 | HMDB0000625 | 10690 | Metabolic pathways;Biosynthesis of secondary metabolites;Biosynthesis of antibiotics;Microbial metabolism in diverse environments;Other | 0.89 | 0.621 | 0.44 | 0.004 | 0.80 | 0.915 | 0.55 | 0.019175 |
| 375 | 2_Carboxy_D_arabinitol | [M-H]-1 | NEG | Mass, Spectral match | Hilic | 196.058 | 2.339 | | HMDB0031201 | 14865845 | Metabolic pathways;Biosynthesis of secondary metabolites;Microbial metabolism in diverse environments;Biosynthesis of antibiotics;Other | 1.58 | 0.150 | 1.22 | 0.341 | 1.36 | 0.158 | 0.90 | 0.552428 |
| 376 | Gluconicacid | [M-H]-1 | NEG | Mass, Spectral match | Hilic | 196.058 | 24.414 | C00257 | HMDB0000625 | 10690 | Metabolic pathways;Biosynthesis of secondary metabolites;Microbial metabolism in diverse environments;Biosynthesis of antibiotics;Other | 1.04 | 0.920 | 0.49 | 0.007 | 0.98 | 0.327 | 0.50 | 0.056712 |
| 377 | Galactonic_acid | [M-H]-1 | NEG | Mass, RT, Spectral match | C18 significant | 196.058 | 0.510 | C00880 | HMDB0000565 | 128869 | Metabolic pathways;Biosynthesis of secondary metabolites;Biosynthesis of | 0.89 | 0.621 | 0.44 | 0.004 | 0.80 | 0.915 | 0.55 | 0.019175 |

| | | | | | | | | | | | | | | | | | | | |
|---|---|---|---|---|---|---|---|---|---|---|---|---|---|---|---|---|---|---|---|
| | | | | | | | | | | | antibiotics;Microbial metabolism in diverse environments;Other | | | | | | | | |
| 378 | Anisotropine_Methylbromide | [M+H]+1 | POS | Mass, Spectral match | Hilic | 197.035 | 8.811 | C06830 | HMDB0014658 | 21867154 | | 2.90 | 0.250 | 0.82 | 0.589 | 1.96 | 0.432 | 0.42 | 0.609318 |
| 379 | N_Acetylhistidine | [M+H]+1 | POS | Mass, Spectral match | Hilic | 197.079 | 3.002 | C02997 | HMDB0032055 | 273260 | | 1.07 | 0.876 | 1.28 | 0.574 | 1.70 | 0.079 | 0.75 | 0.348043 |
| 380 | Tacrine | [M+H]+1 | POS | Mass, RT, Spectral match | C18 | 197.107 | 10.965 | C01453 | HMDB0014526 | 1935 | | 1.59 | 0.202 | 1.17 | 0.707 | 1.35 | 0.032 | 0.86 | 0.423077 |
| 381 | __Menthylacetate | [M+H]+1 | POS | Mass, Spectral match | Hilic | 198.162 | 3.938 | C09870 | HMDB41264 | 220674 | Other | 1.30 | 0.443 | 1.34 | 0.331 | 1.12 | 0.393 | 1.20 | 0.593895 |
| 382 | MCPA | [M-H]-1 | NEG | Mass, Spectral match | Hilic | 200.023 | 4.190 | NA | #N/A | #N/A | | 0.90 | 0.800 | 1.68 | 0.286 | 1.26 | 0.148 | 1.33 | 0.684 |
| 383 | O_Phosphohomoserine | [M-H]-1 | NEG | Mass, Spectral match | C18 | 200.030 | 8.302 | C01102 | HMDB0003484 | 151187 | | 0.93 | 0.876 | 0.50 | 0.221 | 0.39 | 0.139 | 1.28 | 0.725252 |
| 384 | 3_2_Hydroxyphenylpropanoic_acid | [M+H]+1 | POS | Mass, Spectral match | Hilic | 201.030 | 8.354 | C01198 | HMDB0033752 | 873 | | 0.87 | 0.769 | 0.67 | 0.319 | 30.92 | 0.372 | 0.02 | 0.404955 |
| 385 | S_6_Hydroxynicotine | [M+H]+1 | POS | Mass, Spectral match | C18 | 201.100 | 0.638 | C01056 | METPA0118 | NA | Biosynthesis of antibiotics;Other | 2.04 | 0.015 | 1.83 | 0.145 | 1.84 | 0.054 | 1.00 | 0.760137 |
| 386 | Spermine | [M+H]+1 | POS | Mass, Spectral match | C18 | 201.209 | 8.115 | C00750 | HMDB0001256 | 1103 | | 0.96 | 0.383 | 1.04 | 0.498 | 1.08 | 0.100 | 0.97 | 0.531022 |
| 387 | 4_Sulfobenzoic_acid | [M-H]-1 | NEG | Mass, Spectral match | C18 | 201.993 | 0.589 | C02236 | #N/A | 69469 | Microbial metabolism in diverse environments;Degradation of aromatic compounds;Other | 0.54 | 0.233 | 1.10 | 0.893 | 2.14 | 0.437 | 0.51 | 0.401231 |
| 388 | Sebacicacid | [M-H]-1 | NEG | Mass, Spectral match | Hilic | 202.120 | 2.614 | | HMDB0000350 | 3017884 | | 0.99 | 0.978 | 1.18 | 0.545 | 1.75 | 0.309 | 0.67 | 0.636738 |
| 389 | N3N4_Dimethyl_L_arginine | [M+Na]+1 | POS | Mass, Spectral match | Hilic | 202.143 | 1.436 | C03626 | HMDB0001539 | 123831 | | 0.66 | 0.414 | 0.79 | 0.602 | 0.23 | 0.038 | 3.42 | 0.02215 |
| 390 | Symmetric_dimethylarginine | [M+Na]+1 | POS | Mass, Spectral match | Hilic | 202.143 | 1.227 | | HMDB0003334 | 169148 | | 0.94 | 0.932 | 1.36 | 0.665 | 1.98 | 0.018 | 0.69 | 0.279998 |
| 391 | N2_Acetyl_L_aminoadipate | [M+H]+1 | POS | Mass, Spectral match | C18 | 203.079 | 0.519 | C12986 | METPA1105 | NA | | 0.95 | 0.714 | 1.06 | 0.690 | 1.02 | 0.562 | 1.04 | 0.766582 |
| 392 | N2_Acetyl_L_aminoadipate | [M+H]+1 | POS | Mass, Spectral match | Hilic | 203.079 | 4.398 | C12986 | METPA1105 | NA | | 4.03 | 0.045 | 3.19 | 0.081 | 4.23 | 0.040 | 0.75 | 0.603724 |
| 393 | L_Acetylcarnitine | [M+H]+1 | POS | Mass, Spectral match | C18 | 203.115 | 0.574 | C02571 | HMDB0000201 | 1 | | 0.82 | 0.261 | 1.16 | 0.538 | 0.84 | 0.701 | 1.38 | 0.252226 |
| 394 | L_Acetylcarnitine | [M+H]+1 | POS | Mass, Spectral match | Hilic significant | 203.116 | 6.431 | C02571 | HMDB0000201 | 1 | | 1.11 | 0.664 | 0.63 | 0.075 | 0.74 | 0.691 | 0.84 | 0.734389 |
| 395 | Indolelactic_acid | [M+H]+1 | POS | Mass, Spectral match | Hilic | 204.066 | 20.601 | C02043 | HMDB0000671 | 92904 | | 1.11 | 0.840 | 1.39 | 0.634 | 3.01 | 0.227 | 0.46 | 0.684733 |
| 396 | DL_Tryptophan | [M+H]+1 | POS | Mass, Spectral match | C18 | 204.090 | 1.649 | C01365 | #N/A | #N/A | Metabolic pathways;Biosynthesis of secondary metabolites;Biosynthesis of | 1.22 | 0.316 | 1.01 | 0.942 | 1.04 | 0.704 | 0.97 | 0.906537 |

| | | | | | | | | | | | | | | | | | | | |
|---|---|---|---|---|---|---|---|---|---|---|---|---|---|---|---|---|---|---|---|
| | | | | | | | | | | | antibiotics;Biosynthesis of plant secondary metabolites;Biosynthesis of amino acids;Protein digestion and absorption;Central carbon metabolism in cancer;2-Oxocarboxylic acid metabolism;Biosynthesis of alkaloids derived from shikimate pathway;Aminoacyl-tRNA biosynthesis;Other | | | | | | | | |
| 397 | DL_Tryptophan | [M+H]+1 | POS | Mass,RT, Spectral match | Hilic significant | 204.090 | 2.332 | C00525 | HMDB0013609 | 9060 | Metabolic pathways;Biosynthesis of secondary metabolites;Biosynthesis of antibiotics;Biosynthesis of amino acids;Biosynthesis of plant secondary metabolites;Protein digestion and absorption;2-Oxocarboxylic acid metabolism;Aminoacyl-tRNA biosynthesis;Central carbon metabolism in cancer;Glycine, serine and threonine metabolism;Other | 0.04 | 0.047 | 0.40 | 0.311 | 0.24 | 0.598 | 1.64 | 0.989104 |
| 398 | 3_Hydroxy_N6N6N6_trimethyl_L_lysine | [M+H]+1 | POS | Mass, Spectral match | Hilic | 204.147 | 0.725 | C01259 | HMDB0001422 | 439460 | | 0.79 | 0.309 | 1.39 | 0.128 | 0.32 | 0.044 | 4.28 | 0.000738 |
| 399 | Gallic_acid | [M-H]-1 | NEG | Mass, Spectral match | Hilic | 204.992 | 4.889 | C01424 | HMDB0005807 | 370 | | 0.72 | 0.314 | 1.04 | 0.932 | 2.58 | 0.401 | 0.40 | 0.939835 |
| 400 | 3_Oxalomalate | [M-H]-1 | NEG | Mass, Spectral match | Hilic | 204.999 | 15.656 | C01990 | METPA0238 | NA | | 2.01 | 0.146 | 2.29 | 0.021 | 3.03 | 0.021 | 0.76 | 0.480871 |
| 401 | N_Acetyl_L_phenylalanine | [M-H]-1 | NEG | Mass, Spectral match | Hilic | 207.089 | 6.108 | C03519 | HMDB0000512 | 74839 | Other | 0.90 | 0.796 | 0.75 | 0.460 | 0.80 | 0.719 | 0.94 | 0.856219 |
| 402 | 1_Methylhistidine | [M+H]+1 | POS | Mass, Spectral match | C18 | 208.047 | 5.248 | C01152 | HMDB0000001 | 92105 | | 0.59 | 0.594 | 2.93 | 0.324 | 0.47 | 0.327 | 6.22 | 0.365609 |
| 403 | Dihydrolipoate | [M-H]-1 | NEG | Mass, RT, Spectral match | C18 | 208.058 | 0.538 | C02147 | HMDB0012210 | 421 | | 1.57 | 0.099 | 2.97 | 0.023 | 2.40 | 0.127 | 1.24 | 0.479069 |
| 404 | L_Kynurenine | [M+H]+1 | POS | Mass, Spectral match | C18 | 208.084 | 0.949 | C00328 | HMDB0000684 | 161166 | Metabolic pathways;Other | 0.97 | 0.865 | 0.68 | 0.234 | 0.40 | 0.125 | 1.69 | 0.332089 |
| 405 | Pyocyanine | [M+H]+1 | POS | Mass, Spectral match | Hilic | 209.072 | 3.712 | C01748 | #N/A | 439562 | | 0.88 | 0.731 | 0.98 | 0.971 | 0.56 | 0.153 | 1.75 | 0.905252 |
| 406 | tolonidine | [M-H]-1 | NEG | Mass, RT, Spectral match | C18 | 209.072 | 0.504 | C01748 | #N/A | 439562 | | 0.77 | 0.232 | 1.20 | 0.550 | 0.63 | 0.474 | 1.91 | 0.091084 |
| 17 | Pyruvic_acid | [M-H]-1 | NEG | Mass, Spectral match | C18 | 88.015 | 0.520 | C00022 | HMDB0000243 | 1060 | | 3.63 | 0.301 | 0.22 | 0.099 | 0.08 | 0.043 | 2.89 | 0.388501 |
| 408 | Sedoheptulose | [M-H]-1 | NEG | Mass, Spectral match | C18 | 210.074 | 0.489 | C02076 | HMDB0003219 | 102926 | Other | 1.08 | 0.249 | 1.05 | 0.448 | 1.09 | 0.234 | 0.96 | 0.614985 |

| | | | | | | | | | | | | | | | | | | | |
|---|---|---|---|---|---|---|---|---|---|---|---|---|---|---|---|---|---|---|---|
| 409 | Canavanine | [M-H]-1 | NEG | Mass, Spectral match | Hilic | 211.060 | 2.374 | C00308 | HMDB0002706 | 439202 | | 2.16 | 0.089 | 5.85 | 0.040 | 4.05 | 0.073 | 1.45 | 0.432841 |
| 410 | Cysteinylglycine | [M-H]-1 | NEG | Mass, RT, Spectral match | C18 | 213.009 | 2.051 | C01419 | HMDB0000078 | 439498 | | 0.92 | 0.857 | 1.69 | 0.519 | 1.82 | 0.101 | 0.93 | 0.961749 |
| 411 | Dethiobiotin | [M+H]+1 | POS | Mass, RT, Spectral match | C18 | 214.131 | 0.872 | C01909 | HMDB0003581 | 445027 | Metabolic pathways;Other | 1.50 | 0.200 | 1.23 | 0.543 | 1.30 | 0.663 | 0.94 | 0.748638 |
| 412 | Coniferyl_alcohol | [M+H]+1 | POS | Mass, Spectral match | C18 | 215.047 | 0.744 | C00590 | HMDB0012915 | 1549095 | | 1.10 | 0.507 | 0.96 | 0.746 | 0.99 | 0.620 | 0.97 | 0.918541 |
| 413 | sn_glycero_3_Phosphoethanolamine | [2M-H]-1 | NEG | Mass, Spectral match | Hilic | 215.055 | 11.294 | C04438 | METPA0497 | NA | Other | 1.60 | 0.226 | 1.22 | 0.594 | 3.42 | 0.108 | 0.36 | 0.090009 |
| 414 | gamma_Glutamyl_gamma_aminobutyraldehyde | [M+H]+1 | POS | Mass, Spectral match | Hilic | 216.110 | 1.467 | C15700 | METPA1197 | NA | Metabolic pathways;Arginine and proline metabolism | 1.18 | 0.676 | 0.85 | 0.696 | 1.00 | 0.758 | 0.85 | 0.666938 |
| 415 | undecanedioic_acid | [M-H]-1 | NEG | Mass, Spectral match | C18 | 216.135 | 7.626 | | HMDB0000888 | 15816 | | 0.89 | 0.405 | 1.00 | 0.991 | 0.92 | 0.544 | 1.08 | 0.522938 |
| 416 | Xanthan | [M-H]-1 | NEG | Mass, Spectral match | C18 | 217.043 | 0.492 | C01464 | HMDB0029923 | NA | | 1.09 | 0.219 | 1.05 | 0.632 | 0.99 | 0.366 | 1.06 | 0.669474 |
| 417 | Xanthan | [M-H]-1 | NEG | Mass, Spectral match | Hilic | 217.043 | 8.508 | C01464 | HMDB0029923 | NA | | 4.63 | 0.032 | 8.05 | 0.045 | 6.55 | 0.080 | 1.23 | 0.765126 |
| 418 | N_a_Acetylcitrulline | [M+Na]+1 | POS | Mass, Spectral match | Hilic | 217.106 | 3.280 | C15532 | HMDB0000856 | 11506771 | Metabolic pathways;Biosynthesis of amino acids;Other | 2.09 | 0.126 | 1.52 | 0.274 | 1.82 | 0.076 | 0.83 | 0.435141 |
| 419 | gamma_L_Glutamylputrescine | [M+H]+1 | POS | Mass, Spectral match | Hilic | 217.142 | 1.610 | C15699 | HMDB0012230 | 20841830 | Metabolic pathways;Arginine and proline metabolism;Other | 4.42 | 0.180 | 3.03 | 0.240 | 4.73 | 0.294 | 0.64 | 0.709139 |
| 420 | Choline_sulfate | [2M-H]-1 | NEG | Mass, Spectral match | Hilic | 218.026 | 0.387 | C00919 | #N/A | 485 | | 0.65 | 0.535 | 0.33 | 0.200 | 0.97 | 0.703 | 0.34 | 0.175028 |
| 421 | N2_D_1_Carboxyethyl_L_lysine | [M+H]+1 | POS | Mass, Spectral match | Hilic | 218.126 | 1.476 | C04020 | METPA0456 | NA | Lysine degradation | 2.66 | 0.078 | 1.93 | 0.120 | 2.09 | 0.207 | 0.92 | 0.793959 |
| 422 | N2_D_1_Carboxyethyl_L_lysine | [M+H]+1 | POS | Mass, RT, Spectral match | C18 | 218.126 | 1.259 | C04020 | METPA0456 | NA | Other | 2.81 | 0.129 | 1.94 | 0.137 | 2.57 | 0.157 | 0.75 | 0.644137 |
| 423 | Imidazole_acetol_phosphate | [M-H]-1 | NEG | Mass, Spectral match | Hilic | 219.017 | 10.452 | C01267 | HMDB0012236 | 792 | | 0.73 | 0.504 | 0.27 | 0.012 | 0.27 | 0.170 | 1.01 | 0.855567 |
| 424 | Carboxybenzalpyruvate | [M-H]-1 | NEG | Mass, RT, Spectral match | C18 | 219.029 | 0.072 | C01275 | #N/A | 5280421 | | 0.30 | 0.177 | 1.52 | 0.226 | 0.70 | 0.582 | 2.18 | 0.048626 |
| 425 | trans_2_Carboxybenzalpyruvate | [M-H]-1 | NEG | Mass, Spectral match | Hilic | 219.029 | 0.266 | C01275 | #N/A | 5280421 | | 1.23 | 0.684 | 1.41 | 0.375 | 0.87 | 0.096 | 1.62 | 0.145865 |
| 426 | Isoflurophate | [M-H]-1 | NEG | Mass, Spectral match | Hilic | 219.036 | 3.342 | C00202 | HMDB0014815 | 5936 | | 0.14 | 0.089 | 0.80 | 0.799 | 0.14 | 0.273 | 5.65 | 0.303999 |
| 427 | O_Succinyl_L_homoserine | [M-H]-1 | NEG | Mass, RT, Spectral match | C18 | 219.074 | 0.650 | C01118 | METPA0134 | NA | Metabolic pathways;Biosynthesis of secondary metabolites;Biosynthesis of amino acids;Other | 1.55 | 0.180 | 1.55 | 0.249 | 1.73 | 0.092 | 0.89 | 0.614977 |
| 428 | O_Succinyl_L_homoserin | [M+H]+1 | POS | Mass, Spectral | Hilic | 219.074 | 6.024 | C01118 | METPA0134 | NA | Metabolic pathways;Biosynthesis of | 0.96 | 0.897 | 0.82 | 0.558 | 0.69 | 0.678 | 1.19 | 0.708976 |

| | | | | | | | | | | | | | | | | | | | |
|---|---|---|---|---|---|---|---|---|---|---|---|---|---|---|---|---|---|---|---|
| | e | | | match | | | | | | | secondary metabolites;Biosynthesis of amino acids;Cysteine and methionine metabolism;Other | | | | | | | | |
| 429 | N_Phenyl_1_naphthylamine | [M+H]+1 | POS | Mass, RT, Spectral match | C18 | 219.104 | 8.320 | C14405 | #N/A | 7013 | | 1.00 | 0.999 | 1.03 | 0.308 | 0.90 | 0.999 | 1.14 | 0.258478 |
| 430 | Pantothenic_acid | [2M-H]-1 | NEG | Mass, Spectral match | Hilic | 219.110 | 4.045 | C00864 | HMDB0000210 | 988 | Metabolic pathways;Biosynthesis of secondary metabolites;Other | 1.31 | 0.216 | 1.45 | 0.066 | 1.13 | 0.482 | 1.28 | 0.262736 |
| 431 | Cis_zeatin | [M+Cl]-1 | NEG | Mass, RT, Spectral match | C18 | 219.110 | 1.132 | C00371 | HMDB0012204 | 449093 | Metabolic pathways;Biosynthesis of secondary metabolites;Other | 1.28 | 0.299 | 1.28 | 0.255 | 0.96 | 0.763 | 1.33 | 0.243003 |
| 432 | Pantothenic_acid | [M-H]-1 | NEG | Mass, Spectral match | Hilic significant | 219.111 | 4.045 | C00864 | HMDB0000210 | 988 | Metabolic pathways;Biosynthesis of secondary metabolites;Other | 1.31 | 0.216 | 1.45 | 0.066 | 1.13 | 0.482 | 1.28 | 0.262736 |
| 433 | Imidazole_acetol_phosphate | [M-H]-1 | NEG | Mass, RT, Spectral match | C18 | 220.026 | 18.561 | C01267 | HMDB0012236 | 792 | | 0.97 | 0.059 | 1.01 | 0.855 | 0.93 | 0.023 | 1.08 | 0.304465 |
| 434 | L_Histidinol_phosphate | [M+H]+1 | POS | Mass, Spectral match | Hilic | 220.051 | 20.536 | C01100 | METPA0128 | NA | | 1.73 | 0.358 | 1.28 | 0.574 | 1.58 | 0.084 | 0.81 | 0.410688 |
| 435 | Oxitriptan | [M+H]+1 | POS | Mass, Spectral match | C18 | 220.084 | 1.039 | C00643 | HMDB0015571 | 439280 | Metabolic pathways;Biosynthesis of alkaloids derived from shikimate pathway;Other | 1.12 | 0.599 | 0.89 | 0.678 | 3.38 | 0.347 | 0.26 | 0.368447 |
| 436 | 5_Hydroxy_DL_tryptophan | [M+H]+1 | POS | Mass, Spectral match | Hilic | 220.084 | 6.023 | C01017 | HMDB0000472 | 144 | Metabolic pathways;Other | 1.20 | 0.389 | 1.54 | 0.131 | 3.31 | 0.255 | 0.47 | 0.445005 |
| 437 | N_Acetyl_beta_D_glucosaminylamine | [M+H]+1 | POS | Mass, Spectral match | Hilic | 220.105 | 2.984 | C01239 | HMDB0001104 | 439454 | | 2.55 | 0.026 | 1.74 | 0.100 | 2.08 | 0.138 | 0.83 | 0.557383 |
| 438 | Allocystathionine | [M+H]+1 | POS | Mass, Spectral match | C18 | 221.060 | 4.271 | C00542 | HMDB0000455 | 10104953 | | 3.55 | 0.268 | 0.32 | 0.156 | 0.43 | 0.846 | 0.73 | 0.507464 |
| 27 | Haloacetate | [M-H]-1 | NEG | Mass, RT, Spectral match | C18 | 97.976 | 0.540 | C01812 | #N/A | #N/A | | 1.06 | 0.476 | 1.02 | 0.814 | 0.94 | 0.545 | 1.09 | 0.453691 |
| 440 | 2_Acetamidomethylenesuccinate | [M-H]-1 | NEG | Mass, Spectral match | Hilic | 222.019 | 9.261 | C01215 | METPA0147 | NA | | 0.85 | 0.598 | 0.60 | 0.217 | 0.47 | 0.611 | 1.28 | 0.680557 |
| 441 | Cystathionine | [M+H]+1 | POS | Mass, Spectral match | Hilic | 222.067 | 12.085 | C02291 | HMDB0000099 | 439258 | Metabolic pathways;Biosynthesis of secondary metabolites;Biosynthesis of antibiotics;Biosynthesis of amino acids;Glycine, serine and threonine metabolism;Cysteine and methionine metabolism | 1.16 | 0.722 | 0.64 | 0.374 | 1.33 | 0.239 | 0.48 | 0.19971 |
| 37 | Pyridine | [M-H]-1 | NEG | Mass, RT, Spectral match | C18 | 102.031 | 0.519 | C00747 | HMDB0000926 | 1049 | | 1.32 | 0.356 | 1.44 | 0.325 | 1.18 | 0.651 | 1.22 | 0.788492 |
| 443 | N_Acetyl_D_glucosamine | [M-H]-1 | NEG | Mass, RT, Spectral match | C18 | 222.074 | 0.503 | C00140 | HMDB0000215 | 439174 | | 1.04 | 0.788 | 0.93 | 0.702 | 1.35 | 0.316 | 0.69 | 0.33472 |
| 444 | Monobutyl_phthalate | [M+H]+1 | POS | Mass, Spectral match | C18 | 222.089 | 8.253 | | HMDB0013247 | 8575 | Metabolic pathways;Biosynthesis of secondary metabolites;Other | 1.43 | 0.217 | 2.01 | 0.034 | 2.38 | 0.359 | 0.84 | 0.606617 |
| 445 | Sinapic_acid | [M+H]+1 | POS | Mass, Spectral match | C18 | 223.063 | 7.952 | C00482 | HMDB0032616 | 637775 | | 1.00 | 0.983 | 1.30 | 0.387 | 1.33 | 0.210 | 0.97 | 0.727087 |

| | | | | | | | | | | | | | | | | | | | |
|---|---|---|---|---|---|---|---|---|---|---|---|---|---|---|---|---|---|---|---|
| 446 | 78_Diaminononanoate | [M-H]-1 | NEG | Mass, Spectral match | Hilic | 223.120 | 3.725 | C01037 | METPA0113 | NA | Other | 1.67 | 0.411 | 0.97 | 0.945 | 0.96 | 0.541 | 1.01 | 0.975785 |
| 447 | Kynurenic_acid | [M-H]-1 | NEG | Mass, Spectral match | C18 | 224.011 | 2.597 | C01717 | HMDB0000715 | 3845 | | 1.82 | 0.449 | 1.38 | 0.517 | 1.18 | 0.297 | 1.17 | 0.49149 |
| 448 | Naphthazarin | [M-H]-1 | NEG | Mass, Spectral match | Hilic | 224.995 | 7.862 | C01938 | #N/A | 10965344 | | 1.73 | 0.157 | 1.76 | 0.188 | 3.11 | 0.040 | 0.57 | 0.276801 |
| 449 | Diaminopimelic_acid | [2M+H]+1 | POS | Mass, Spectral match | Hilic | 225.063 | 10.584 | C00666 | HMDB0001370 | 439283 | Biosynthesis of antibiotics;Other | 2.46 | 0.120 | 1.39 | 0.411 | 2.19 | 0.062 | 0.63 | 0.211332 |
| 450 | Tetradecenoic_acid | [M+H]+1 | POS | Mass, Spectral match | Hilic | 226.193 | 3.916 | C08322 | HMDB0002000 | 5281119 | | 1.21 | 0.546 | 0.92 | 0.794 | 0.89 | 0.453 | 1.02 | 0.963195 |
| 451 | Deoxycytidine | [M+H]+1 | POS | Mass, RT, Spectral match | C18 | 227.089 | 4.661 | C00881 | HMDB0000014 | 13711 | | 1.72 | 0.322 | 0.84 | 0.410 | 1.06 | 0.344 | 0.79 | 0.470622 |
| 452 | Deoxycytidine | [M+H]+1 | POS | Mass,RT, Spectral match | Hilic | 227.090 | 4.390 | C00881 | HMDB0000014 | 13711 | Pyrimidine metabolism | 4.73 | 0.156 | 5.75 | 0.258 | 6.92 | 0.305 | 0.83 | 0.715215 |
| 50 | Glyceric_acid | [M-H]-1 | NEG | Mass, RT, Spectral match | C18 | 106.026 | 0.506 | C00258 | HMDB0000139 | 439194 | | 1.13 | 0.608 | 0.82 | 0.552 | 0.73 | 0.402 | 1.13 | 0.953354 |
| 454 | Leu_Pro | [M-H]-1 | NEG | Mass, Spectral match | Hilic significant | 228.147 | 1.294 | | HMDB11175 | 44369311 | | 1.64 | 0.031 | 1.22 | 0.378 | 1.53 | 0.213 | 0.80 | 0.358344 |
| 455 | Leucylproline | [M+H]+1 | POS | Mass, RT, Spectral match | C18 | 228.147 | 1.797 | | HMDB0011175 | 44369311 | | 1.70 | 0.130 | 1.18 | 0.549 | 1.55 | 0.350 | 0.76 | 0.357188 |
| 81 | Fumaric_acid | [M-H]-1 | NEG | Mass, Spectral match | C18 | 116.010 | 0.529 | C00122 | HMDB0000134 | 444972 | Metabolic pathways;Biosynthesis of secondary metabolites;Biosynthesis of antibiotics;Microbial metabolism in diverse environments;Biosynthesis of plant secondary metabolites;Degradation of aromatic compounds;Central carbon metabolism in cancer;Biosynthesis of alkaloids derived from shikimate pathway;Biosynthesis of alkaloids derived from ornithine, lysine and nicotinic acid;Other | 3.01 | 0.197 | 1.33 | 0.111 | 1.24 | 0.329 | 1.07 | 0.784215 |
| 457 | Mevalonic_acid_5P | [M+Cl]-1 | NEG | Mass, Spectral match | C18 | 229.048 | 1.067 | C01107 | HMDB0001343 | 439400 | | 1.03 | 0.160 | 1.01 | 0.328 | 1.01 | 0.958 | 1.00 | 0.864228 |
| 458 | Naproxen | [M+H]+1 | POS | Mass,RT, Spectral match | Hilic | 229.088 | 8.049 | C01517 | HMDB0001923 | 156391 | | 1.08 | 0.844 | 0.66 | 0.191 | 0.63 | 0.670 | 1.04 | 0.751928 |
| 459 | Naproxen | [M+H]+1 | POS | Mass, RT, Spectral match | C18 | 229.089 | 7.837 | C01517 | HMDB0001923 | 156391 | | 1.65 | 0.452 | 1.62 | 0.409 | 3.80 | 0.222 | 0.43 | 0.637678 |
| 460 | D_Ribose_5_phosphate | [M-H]-1 | NEG | Mass, Spectral match | C18 | 230.019 | 0.530 | C00117 | HMDB0001548 | 439167 | Metabolic pathways;Biosynthesis of secondary metabolites;Biosynthesis of antibiotics;Microbial | 0.98 | 0.917 | 0.72 | 0.254 | 0.66 | 0.765 | 1.08 | 0.59551 |

| | | | | | | | | | | | | | | | | | | | |
|---|---|---|---|---|---|---|---|---|---|---|---|---|---|---|---|---|---|---|---|
| | | | | | | | | | | | metabolism in diverse environments;Biosynthesis of amino acids;Other | | | | | | | | |
| 461 | L_Formylkynurenine | [M-H]-1 | NEG | Mass, Spectral match | Hilic | 236.079 | 5.763 | C05648 | HMDB0004086 | 440744 | Metabolic pathways;Other | 1.52 | 0.112 | 1.40 | 0.372 | 16.20 | 0.366 | 0.09 | 0.451294 |
| 462 | 4_Sulfobenzoic_acid. | [M-H]-1 | NEG | Mass, Spectral match | Hilic | 236.960 | 15.676 | C02236 | #N/A | 69469 | | 0.29 | 0.010 | 0.34 | 0.073 | 0.34 | 0.340 | 1.00 | 0.872373 |
| 463 | 3_Hydroxyflavone | [M+H]+1 | POS | Mass, Spectral match | C18 | 237.055 | 2.804 | C01495 | HMDB0031816 | 11349 | | 10.04 | 0.264 | 0.39 | 0.300 | 0.52 | 0.581 | 0.75 | 0.555732 |
| 464 | N_Acetyl_D_glucosamine | [M+Na]+1 | POS | Mass, Spectral match | Hilic | 237.084 | 10.365 | C00140 | HMDB0000215 | 439174 | | 0.82 | 0.308 | 0.74 | 0.208 | 0.79 | 0.917 | 0.95 | 0.88959 |
| 465 | Biopterin | [M+H]+1 | POS | Mass, Spectral match | Hilic | 237.085 | 6.018 | C06313 | HMDB0000468 | 445040 | Metabolic pathways;Other | 1.14 | 0.642 | 0.96 | 0.897 | 1.07 | 0.573 | 0.90 | 0.905344 |
| 466 | 6_Pyruvoyltetrahydropterin | [M+H]+1 | POS | Mass, Spectral match | Hilic | 237.086 | 4.087 | C03684 | HMDB0001195 | 128973 | Metabolic pathways;Other | 1.03 | 0.944 | 1.17 | 0.590 | 1.68 | 0.250 | 0.70 | 0.740783 |
| 467 | Ketamine | [M+H]+1 | POS | Mass, Spectral match | C18 | 237.092 | 5.331 | C07525 | HMDB0015352 | 3821 | | 1.89 | 0.164 | 1.47 | 0.147 | 1.29 | 0.514 | 1.14 | 0.605768 |
| 468 | 4a_Carbinolamine_tetrahydrobiopterin | [M-H]-1 | NEG | Mass, Spectral match | Hilic | 238.095 | 0.699 | C00268 | HMDB0002215 | 133246 | Other | 2.93 | 0.108 | 2.57 | 0.086 | 2.34 | 0.302 | 1.10 | 0.600528 |
| 469 | Sepiapterin | [M+H]+1 | POS | Mass, Spectral match | C18 | 238.095 | 5.320 | C00835 | HMDB0000238 | 65253 | Other | 1.58 | 0.122 | 1.42 | 0.245 | 1.21 | 0.537 | 1.17 | 0.551789 |
| 470 | Muscone | [M+H]+1 | POS | Mass, Spectral match | C18 | 238.229 | 13.039 | | HMDB0034181 | 10947 | | 1.24 | 0.472 | 0.90 | 0.595 | 1.23 | 0.281 | 0.73 | 0.631509 |
| 471 | Clomazone | [M+H]+1 | POS | Mass, Spectral match | C18 | 239.071 | 6.841 | C11095 | #N/A | 54778 | | 2.93 | 0.204 | 0.36 | 0.068 | 1.66 | 0.129 | 0.22 | 0.096294 |
| 472 | 4a_Carbinolamine_tetrahydrobiopterin | [M+Na]+1 | POS | Mass, RT, Spectral match | C18 | 239.102 | 0.583 | C00268 | HMDB0002215 | 133246 | Metabolic pathways;Other | 1.48 | 0.284 | 1.04 | 0.918 | 1.02 | 0.584 | 1.01 | 0.802701 |
| 473 | L_Cystine | [M+H]+1 | POS | Mass, Spectral match | Hilic | 240.023 | 12.622 | C00491 | HMDB0000192 | 67678 | Metabolic pathways;ABC transporters;Protein digestion and absorption;Cysteine and methionine metabolism;Other | 1.35 | 0.337 | 1.58 | 0.093 | 1.31 | 0.393 | 1.20 | 0.462912 |
| 474 | Dinoseb | [M-H]-1 | NEG | Mass, Spectral match | Hilic | 240.074 | 1.294 | C14302 | HMDB0032559 | 6950 | | 1.54 | 0.338 | 2.66 | 0.089 | 1.49 | 0.909 | 1.79 | 0.561921 |
| 475 | Anserine | [M+H]+1 | POS | Mass, Spectral match | Hilic | 240.122 | 6.774 | C01262 | HMDB0000194 | 112072 | Metabolic pathways;Arginine and proline metabolism;Histidine metabolism;Other | 1.52 | 0.202 | 1.94 | 0.115 | 2.00 | 0.124 | 0.97 | 0.874844 |
| 476 | R_lipoic_acid | [M+H]+1 | POS | Mass, Spectral match | Hilic | 241.015 | 7.822 | C00725 | HMDB0001451 | 6112 | | 0.79 | 0.017 | 0.90 | 0.450 | 0.94 | 0.540 | 0.96 | 0.878866 |
| 477 | Lumichrome | [M+H]+1 | POS | Mass, Spectral match | Hilic | 241.073 | 1.037 | C01727 | METPA0216 | NA | Biosynthesis of secondary metabolites;Other | 1.04 | 0.469 | 0.86 | 0.006 | 1.02 | 0.376 | 0.84 | 0.017836 |
| 478 | Lumichrome | [M+H]+1 | POS | Mass, RT, Spectral match | C18 | 241.073 | 8.317 | C01727 | METPA0216 | NA | Biosynthesis of secondary metabolites;Biosynthesis of alkaloids derived from shikimate pathway;Other | 0.92 | 0.243 | 0.85 | 0.025 | 0.76 | 0.212 | 1.12 | 0.226072 |

| | | | | | | | | | | | | | | | | | | | |
|---|---|---|---|---|---|---|---|---|---|---|---|---|---|---|---|---|---|---|---|
| 479 | Bis2_ethylhexylamine | [M+H]+1 | POS | Mass, Spectral match | Hilic | 241.276 | 27.866 | C02455 | HMDB0002017 | 7408 | | 0.96 | 0.695 | 0.94 | 0.575 | 1.13 | 0.180 | 0.84 | 0.048255 |
| 480 | L_Kynurenine | [M+H]+1 | POS | Mass, Spectral match | Hilic | 243.054 | 6.890 | C00328 | HMDB0000684 | 161166 | | 1.43 | 0.660 | 1.55 | 0.472 | 1.71 | 0.250 | 0.91 | 0.645756 |
| 481 | Chalcone | [M-H]-1 | NEG | Mass, Spectral match | Hilic | 243.059 | 8.474 | C01484 | HMDB0003066 | 637760 | | 1.02 | 0.901 | 1.23 | 0.305 | 1.43 | 0.342 | 0.86 | 0.998786 |
| 482 | Chalcone | [M-H]-1 | NEG | Mass, RT, Spectral match | C18 | 243.059 | 0.732 | C01484 | HMDB0003066 | 637760 | | 1.52 | 0.023 | 1.37 | 0.131 | 1.49 | 0.262 | 0.92 | 0.878856 |
| 483 | Biotin | [M-H]-1 | NEG | Mass, RT, Spectral match | C18 | 243.080 | 18.685 | C00120 | HMDB0000030 | 171548 | | 0.95 | 0.200 | 1.01 | 0.853 | 0.93 | 0.736 | 1.09 | 0.116582 |
| 484 | Biotin | [M+H]+1 | POS | Mass, Spectral match | Hilic | 243.080 | 10.991 | C00120 | HMDB0000030 | 171548 | | 0.83 | 0.417 | 1.00 | 0.996 | 0.87 | 0.836 | 1.16 | 0.298449 |
| 91 | D___Erythrose | [M-H]-1 | NEG | Mass, RT, Spectral match | C18 | 120.041 | 0.522 | C01796 | HMDB0002649 | 439574 | | 1.32 | 0.074 | 1.32 | 0.156 | 1.33 | 0.188 | 0.99 | 0.93412 |
| 124 | 5_Aminolevulinic_acid | [M-H]-1 | NEG | Mass, RT, Spectral match | C18 | 131.057 | 0.690 | C00430 | HMDB0001149 | 137 | | 1.38 | 0.208 | 1.01 | 0.960 | 1.21 | 0.390 | 0.83 | 0.507954 |
| 154 | Threonic_acid | [M-H]-1 | NEG | Mass, Spectral match | C18 | 136.036 | 0.500 | C01620 | HMDB0000943 | 151152 | | 0.89 | 0.617 | 0.44 | 0.003 | 0.80 | 0.903 | 0.55 | 0.017082 |
| 199 | D_?_Hydroxyglutaric_acid | [M-H]-1 | NEG | Mass, Spectral match | C18 | 148.036 | 0.595 | C00652 | #N/A | 17723 | | 1.22 | 0.409 | 0.82 | 0.351 | 1.05 | 0.512 | 0.79 | 0.464683 |
| 489 | Cytidine | [M+H]+1 | NEG | Mass, Spectral match | C18 | 244.092 | 2.492 | C00475 | HMDB0000089 | 6175 | | 1.15 | 0.620 | 0.55 | 0.062 | 1.43 | 0.288 | 0.39 | 0.045938 |
| 490 | Aminocaproic_acid | [M-H]-1 | NEG | Mass, Spectral match | Hilic | 244.178 | 1.236 | C02378 | HMDB0001901 | 564 | Microbial metabolism in diverse environments;Other | 2.16 | 0.244 | 1.62 | 0.296 | 2.18 | 0.160 | 0.74 | 0.743697 |
| 491 | N_6_Aminohexanoyl_6_aminohexanoate | [M-H]-1 | NEG | Mass, Spectral match | C18 | 244.178 | 4.833 | C01255 | #N/A | 895 | Microbial metabolism in diverse environments;Other | 3.45 | 0.179 | 2.29 | 0.163 | 3.37 | 0.182 | 0.68 | 0.751817 |
| 492 | Glucaric_acid | [M-H]-1 | NEG | Mass, Spectral match | Hilic | 245.006 | 10.139 | C00818 | HMDB0000663 | 33037 | | 1.87 | 0.010 | 2.27 | 0.018 | 2.02 | 0.133 | 1.12 | 0.671835 |
| 493 | Pyridoxal_5_phosphate | [M-H]-1 | NEG | Mass, Spectral match | C18 | 246.020 | 2.811 | C00018 | HMDB0001491 | 1051 | | 1.58 | 0.608 | 1.00 | 0.994 | 0.44 | 0.426 | 2.30 | 0.421661 |
| 494 | Benzoin | [M+H]+1 | POS | Mass, Spectral match | Hilic | 247.054 | 5.350 | C01408 | HMDB0032039 | 8400 | Other | 1.02 | 0.947 | 0.85 | 0.540 | 0.87 | 0.712 | 0.98 | 0.676236 |
| 495 | Lansoprazole | [M+H]+1 | POS | Mass, RT, Spectral match | C18 | 247.105 | 0.540 | C01594 | HMDB0005008 | 3883 | Biosynthesis of secondary metabolites;Other | 0.73 | 0.099 | 0.92 | 0.827 | 1.11 | 0.483 | 0.82 | 0.574706 |
| 496 | Linamarin | [M+H]+1 | POS | Mass, Spectral match | Hilic | 247.105 | 8.806 | C01594 | HMDB0033699 | 11128 | Biosynthesis of secondary metabolites;Cyanoamino acid metabolism | 4.01 | 0.120 | 12.80 | 0.041 | 11.99 | 0.075 | 1.07 | 0.962611 |
| 497 | Pyridoxamine5_phosphate | [M+H]+1 | POS | Mass, Spectral match | Hilic | 248.056 | 13.896 | C00627 | HMDB0001319 | 1055 | Metabolic pathways;Other | 2.11 | 0.181 | 1.65 | 0.220 | 1.47 | 0.414 | 1.13 | 0.584268 |
| 498 | Pyridoxine_5_phosphate | [M-H]-1 | NEG | Mass, RT, Spectral match | C18 | 250.045 | 1.584 | C00627 | HMDB0001319 | 1055 | | 0.99 | 0.976 | 0.86 | 0.497 | 1.06 | 0.390 | 0.81 | 0.641745 |

| | | | | | | | | | | | | | | | | | | | |
|---|---|---|---|---|---|---|---|---|---|---|---|---|---|---|---|---|---|---|---|
| 499 | L_Arogenate | [M+H]+1 | POS | Mass, RT, Spectral match | C18 | 250.070 | 0.894 | C00826 | METPA0086 | NA | | 1.40 | 0.338 | 0.94 | 0.692 | 0.78 | 0.848 | 1.21 | 0.323267 |
| 500 | Deoxyadenosine | [M+H]+1 | POS | Mass, RT, Spectral match | C18 | 251.100 | 0.517 | C00559 | HMDB0000101 | 13730 | | 0.64 | 0.246 | 0.78 | 0.439 | 0.90 | 0.801 | 0.87 | 0.95137 |
| 501 | Muramicacid | [M+H]+1 | POS | Mass,RT, Spectral match | Hilic | 251.100 | 3.306 | C06470 | HMDB0003254 | 22833605 | | 0.73 | 0.451 | 0.76 | 0.529 | 0.74 | 0.853 | 1.03 | 0.878724 |
| 502 | 2_Deoxyadenosine | [M+Cl]-1 | POS | Mass, Spectral match | Hilic | 251.100 | 5.097 | C00559 | HMDB0000101 | 13730 | | 0.67 | 0.131 | 0.62 | 0.092 | 0.57 | 0.325 | 1.08 | 0.584173 |
| 503 | Ala_Tyr | [M-H]-1 | NEG | Mass, Spectral match | Hilic | 252.111 | 1.613 | NA | HMDB0028699 | NA | | 2.20 | 0.123 | 1.85 | 0.069 | 2.56 | 0.162 | 0.72 | 0.62917 |
| 504 | 2_Hydroxy_6_oxo_6_phenylhexa_24_dienoate | [M+H]+1 | POS | Mass, RT, Spectral match | C18 | 253.025 | 1.339 | C01273 | METPA0162 | NA | | 0.79 | 0.327 | 1.15 | 0.430 | 0.84 | 0.283 | 1.38 | 0.00804 |
| 505 | Ketoprofen | [M+H]+1 | POS | Mass,RT, Spectral match | Hilic | 253.086 | 0.722 | C01716 | HMDB0015144 | 3825 | | 1.63 | 0.306 | 0.91 | 0.592 | 0.83 | 0.642 | 1.09 | 0.44412 |
| 506 | N_Ribosylnicotinamide | [M-H]-1 | NEG | Mass, Spectral match | Hilic | 254.090 | 4.311 | C03150 | HMDB0000855 | 439924 | Other | 0.84 | 0.745 | 1.19 | 0.702 | 1.59 | 0.024 | 0.75 | 0.361585 |
| 507 | 2_beta_D_Glucosyl_sn_glycerol | [M-H]-1 | NEG | Mass, Spectral match | Hilic | 254.100 | 7.128 | C04046 | METPA0461 | NA | Other | 1.35 | 0.180 | 2.49 | 0.067 | 2.49 | 0.140 | 1.00 | 0.972472 |
| 508 | Hexadecanamide | [M+H]+1 | POS | Mass, RT, Spectral match | C18 | 255.256 | 12.796 | | HMDB0012273 | 69421 | | 1.05 | 0.645 | 1.01 | 0.957 | 1.02 | 0.211 | 0.98 | 0.868419 |
| 509 | Hexadecanamide | [M+H]+1 | POS | Mass,RT, Spectral match | Hilic | 255.256 | 4.120 | | HMDB0012273 | 69421 | | 0.90 | 0.572 | 0.86 | 0.410 | 0.88 | 0.301 | 0.98 | 0.950454 |
| 510 | Dihydrozeatin | [M-H]-1 | NEG | Mass, Spectral match | Hilic | 256.096 | 2.398 | C02029 | HMDB0012215 | 439631 | Other | 1.01 | 0.961 | 1.26 | 0.264 | 0.67 | 0.630 | 1.89 | 0.030457 |
| 208 | L_Arabinose | [M-H]-1 | NEG | Mass, Spectral match | C18 | 150.052 | 0.505 | C00121 | HMDB0000283 | 5779 | Metabolic pathways;ABC transporters;Other | 0.89 | 0.627 | 0.45 | 0.004 | 0.81 | 0.902 | 0.56 | 0.022466 |
| 512 | Glycerophosphocholine | [M+H]+1 | POS | Mass, RT, Spectral match | C18 | 257.102 | 0.492 | C00670 | HMDB0000086 | 71920 | | 1.86 | 0.045 | 1.73 | 0.311 | 4.64 | 0.119 | 0.37 | 0.163639 |
| 513 | sn_glycero_3_Phosphocholine | [M-H]-1 | NEG | Mass, Spectral match | Hilic | 257.103 | 7.679 | C00670 | HMDB0000086 | 71920 | | 0.55 | 0.201 | 1.00 | 0.993 | 0.53 | 0.936 | 1.88 | 0.345082 |
| 514 | 78_Dihydro_78_dihydroxykynurenate | [M+H]+1 | POS | Mass, Spectral match | Hilic | 258.016 | 21.512 | C01249 | METPA0157 | NA | | 1.10 | 0.671 | 0.95 | 0.798 | 0.99 | 0.935 | 0.96 | 0.775761 |
| 273 | Phosphoenolpyruvic_acid | [M-H]-1 | NEG | Mass, RT, Spectral match | C18 | 167.982 | 0.507 | C00074 | HMDB0000263 | 1005 | Metabolic pathways;Biosynthesis of secondary metabolites;Biosynthesis of antibiotics;Microbial metabolism in diverse environments;Biosynthesis of plant secondary metabolites;Biosynthesis of amino acids;Central carbon metabolism in cancer;Biosynthesis of alkaloids derived from | 3.26 | 0.218 | 1.13 | 0.779 | 1.26 | 0.412 | 0.89 | 0.950543 |

| | | | | | | | | | | | | | | | | | | | |
|---|---|---|---|---|---|---|---|---|---|---|---|---|---|---|---|---|---|---|---|
| | | | | | | | | | | | shikimate pathway;Other | | | | | | | | |
| 516 | Norbutorphanol | [M+H]+1 | POS | Mass, Spectral match | Hilic | 259.157 | 0.729 | C06863 | HMDB0014749 | 5361092 | Biosynthesis of antibiotics;Other | 1.08 | 0.316 | 1.09 | 0.218 | 1.11 | 0.530 | 0.98 | 0.754164 |
| 517 | Propranolol | [M+H]+1 | POS | Mass, RT, Spectral match | C18 | 259.157 | 6.152 | C07407 | HMDB0001849 | 4946 | Biosynthesis of antibiotics;Other | 1.03 | 0.462 | 1.03 | 0.328 | 1.02 | 0.891 | 1.01 | 0.504227 |
| 518 | Dihydrozeatin | [M-H]-1 | NEG | Mass, RT, Spectral match | C18 | 260.090 | 8.229 | C02029 | HMDB0012215 | 439631 | | 0.30 | 0.034 | 0.21 | 0.012 | 0.59 | 0.857 | 0.37 | 0.077271 |
| 519 | Carnosine | [M+H]+1 | POS | Mass, Spectral match | Hilic | 261.078 | 8.067 | C00386 | HMDB0000033 | 439224 | | 1.21 | 0.498 | 0.93 | 0.806 | 0.89 | 0.760 | 1.05 | 0.752579 |
| 520 | Lotaustralin | [M+H]+1 | POS | Mass, Spectral match | Hilic | 261.121 | 8.211 | C08334 | HMDB0033865 | 4626626 | Biosynthesis of secondary metabolites;Cyanoamino acid metabolism | 0.69 | 0.140 | 0.82 | 0.590 | 0.96 | 0.750 | 0.85 | 0.844038 |
| 521 | Lotaustralin | [M+H]+1 | POS | Mass, RT, Spectral match | C18 | 261.121 | 0.632 | C08334 | HMDB0033865 | 4626626 | Biosynthesis of secondary metabolites;Other | 0.84 | 0.325 | 0.99 | 0.982 | 0.89 | 0.624 | 1.12 | 0.896036 |
| 522 | Albendazole | [M-H]-1 | NEG | Mass, Spectral match | Hilic | 264.081 | 2.484 | C01779 | HMDB0014659 | 2082 | | 0.93 | 0.541 | 0.97 | 0.663 | 0.96 | 0.335 | 1.02 | 0.798167 |
| 523 | Thiamine | [M+H]+1 | POS | Mass, RT, Spectral match | C18 | 264.104 | 0.474 | C00378 | HMDB0000235 | 1130 | 2-Oxocarboxylic acid metabolism;Other | 1.56 | 0.033 | 1.47 | 0.118 | 1.23 | 0.292 | 1.20 | 0.473275 |
| 524 | Thiamine | [M-H]-1 | NEG | Mass, Spectral match | Hilic | 264.104 | 0.809 | C00378 | HMDB0000235 | 1130 | 2-Oxocarboxylic acid metabolism;Other | 1.26 | 0.338 | 1.01 | 0.958 | 1.05 | 0.463 | 0.97 | 0.97681 |
| 525 | Adenosine | [M-H]-1 | NEG | Mass, RT, Spectral match | C18 | 266.090 | 0.857 | C00212 | HMDB0000050 | 60961 | | 1.55 | 0.352 | 1.22 | 0.541 | 0.97 | 0.664 | 1.26 | 0.535298 |
| 526 | D_Ribitol_5_phosphate | [M+H]+1 | POS | Mass, Spectral match | Hilic | 267.003 | 0.726 | C01068 | METPA0121 | NA | | 1.00 | 0.982 | 1.01 | 0.919 | 0.95 | 0.355 | 1.07 | 0.701525 |
| 527 | L_Fuculose_1_phosphate | [M-H]-1 | NEG | Mass, Spectral match | C18 | 267.022 | 1.059 | C01099 | METPA0127 | NA | | 1.00 | 0.925 | 0.99 | 0.755 | 0.93 | 0.971 | 1.06 | 0.336717 |
| 528 | Neuraminicacid | [M+H]+1 | POS | Mass, Spectral match | Hilic | 267.095 | 11.031 | C06469 | HMDB0000830 | 513472 | | 0.80 | 0.410 | 0.89 | 0.761 | 0.97 | 0.731 | 0.92 | 0.87855 |
| 529 | N_Acetylgalactosamine | [M+K]+1 | POS | Mass, Spectral match | Hilic | 267.095 | 4.258 | C01074 | HMDB0000212 | 84265 | Metabolic pathways;Other | 1.11 | 0.749 | 0.69 | 0.153 | 0.78 | 0.760 | 0.88 | 0.613956 |
| 530 | Adenosine | [M+Cl]-1 | POS | Mass,RT, Spectral match | Hilic | 267.096 | 2.379 | C00212 | HMDB0000050 | 60961 | Metabolic pathways;Other | 0.45 | 0.444 | 0.08 | 0.195 | 0.06 | 0.180 | 1.35 | 0.534056 |
| 276 | Homogentisic_acid | [M-H]-1 | NEG | Mass, Spectral match | C18 | 169.050 | 0.496 | C00544 | HMDB0000130 | 780 | Other | 1.33 | 0.052 | 1.00 | 0.993 | 2.25 | 0.124 | 0.45 | 0.10321 |
| 532 | Inosine | [2M-H]-1 | NEG | Mass, Spectral match | Hilic significant | 268.081 | 1.840 | C00294 | HMDB0000195 | 6021 | Metabolic pathways;Other | 1.61 | 0.044 | 2.08 | 0.200 | 1.59 | 0.299 | 1.31 | 0.937646 |
| 533 | Morphine | [M+H]+1 | POS | Mass, RT, Spectral match | C18 | 271.157 | 6.743 | C01516 | HMDB0014440 | 5288826 | | 0.96 | 0.241 | 0.99 | 0.721 | 0.95 | 0.108 | 1.04 | 0.294914 |
| 534 | S_Acetyldihydrolipoamide | [M-H]-1 | NEG | Mass, Spectral match | C18 | 272.077 | 1.651 | C01136 | HMDB0001526 | 1076 | | 1.17 | 0.376 | 0.98 | 0.918 | 1.02 | 0.737 | 0.96 | 0.916527 |
| 535 | 16_Hydroxyhexadecanoicacid | [M-H]-1 | NEG | Mass, Spectral match | Hilic | 272.235 | 1.205 | C08317 | HMDB0002059 | 79034 | Metabolic pathways;Other | 0.71 | 0.396 | 0.84 | 0.675 | 0.89 | 0.163 | 0.95 | 0.780418 |

| | | | | | | | | | | | | | | | | | | | |
|---|---|---|---|---|---|---|---|---|---|---|---|---|---|---|---|---|---|---|---|
| 536 | N2_Succinyl_L_arginine | [M-H]-1 | NEG | Mass, Spectral match | Hilic | 274.127 | 5.513 | C03296 | METPA0384 | NA | Metabolic pathways;Arginine and proline metabolism | 3.14 | 0.196 | 2.37 | 0.087 | 4.25 | 0.070 | 0.56 | 0.213552 |
| 537 | Mefenamic_acid | [M+H]+1 | POS | Mass, Spectral match | Hilic | 276.082 | 6.458 | C02168 | HMDB0014922 | 4044 | | 1.16 | 0.437 | 1.98 | 0.147 | 1.72 | 0.229 | 1.15 | 0.786223 |
| 538 | Ggamma_L_Glutamyl_L_glutamicacid | [M+H]+1 | POS | Mass, Spectral match | Hilic | 276.096 | 19.966 | C05282 | HMDB0011737 | 92865 | Biosynthesis of antibiotics;Other | 1.26 | 0.425 | 1.39 | 0.224 | 1.37 | 0.374 | 1.01 | 0.60437 |
| 539 | Queuine | [M+H]+1 | POS | Mass, Spectral match | Hilic | 276.108 | 2.757 | C01449 | HMDB0001495 | 53477731 | Metabolic pathways;Biosynthesis of secondary metabolites;Microbial metabolism in diverse environments;Other | 0.09 | 0.015 | 0.29 | 0.060 | 0.04 | 0.064 | 8.02 | 0.203833 |
| 540 | L_Saccharopine | [M+H]+1 | POS | Mass, Spectral match | Hilic | 276.132 | 17.853 | C00449 | HMDB0000279 | 160556 | Metabolic pathways;Biosynthesis of secondary metabolites;Biosynthesis of antibiotics;Biosynthesis of amino acids;Lysine degradation;Other | 0.90 | 0.782 | 1.42 | 0.328 | 2.06 | 0.121 | 0.69 | 0.53026 |
| 541 | Pantetheine | [M+H]+1 | POS | Mass, RT, Spectral match | C18 | 278.129 | 10.911 | C00831 | HMDB0003426 | 479 | Biosynthesis of antibiotics;Other | 0.62 | 0.106 | 0.63 | 0.160 | 0.57 | 0.236 | 1.11 | 0.948418 |
| 542 | Dibutylphthalate | [M-H]-1 | NEG | Mass, Spectral match | Hilic | 278.151 | 27.836 | C14214 | HMDB0033244 | 3026 | | 0.99 | 0.800 | 1.03 | 0.394 | 1.01 | 0.134 | 1.02 | 0.322779 |
| 543 | Dibutyl_phthalate | [M-H]-1 | NEG | Mass, RT, Spectral match | C18 | 278.152 | 10.574 | C14214 | HMDB0033244 | 3026 | | 0.80 | 0.166 | 0.72 | 0.088 | 0.78 | 0.891 | 0.92 | 0.514005 |
| 544 | Aspartyl_Phenylalanine | [M+H]+1 | POS | Mass, Spectral match | C18 | 280.105 | 1.975 | | HMDB0000706 | 93078 | | 3.34 | 0.204 | 2.97 | 0.145 | 3.25 | 0.251 | 0.91 | 0.967142 |
| 545 | Aspartylphenylalanine | [M-H]-1 | NEG | Mass, Spectral match | Hilic | 280.106 | 8.069 | | HMDB0000706 | 93078 | | 1.71 | 0.081 | 1.49 | 0.139 | 1.88 | 0.090 | 0.79 | 0.276186 |
| 546 | Imipramine | [M+H]+1 | POS | Mass, Spectral match | C18 | 280.193 | 7.871 | C07049 | HMDB0001848 | 3696 | | 1.03 | 0.449 | 0.98 | 0.480 | 1.00 | 0.917 | 0.98 | 0.41199 |
| 547 | Linoleic_acid | [M+H]+1 | POS | Mass, Spectral match | C18 | 280.240 | 13.087 | C01595 | HMDB0000673 | 5280450 | Metabolic pathways;Biosynthesis of plant secondary metabolites;Other | 1.04 | 0.865 | 1.09 | 0.662 | 1.19 | 0.697 | 0.92 | 0.674097 |
| 548 | Alverine | [M+H]+1 | POS | Mass, Spectral match | C18 | 281.214 | 13.304 | D07440 | #N/A | 3678 | | 0.85 | 0.198 | 0.80 | 0.041 | 0.81 | 0.817 | 0.98 | 0.823199 |
| 549 | Oleamide | [M+H]+1 | POS | Mass, Spectral match | C18 | 281.271 | 13.005 | C19670 | HMDB0002117 | 5283387 | | 0.87 | 0.050 | 0.91 | 0.103 | 0.82 | 0.808 | 1.12 | 0.229401 |
| 550 | Oleamide | [M+H]+1 | POS | Mass, Spectral match | Hilic | 281.271 | 28.191 | C19670 | HMDB0002117 | 5283387 | | 1.10 | 0.795 | 1.04 | 0.903 | 1.06 | 0.487 | 0.98 | 0.960783 |
| 551 | Glucosamine_6_phosphate | [M-H]-1 | NEG | Mass, RT, Spectral match | C18 | 282.033 | 0.788 | C00352 | HMDB0001254 | 439217 | | 0.81 | 0.516 | 1.08 | 0.791 | 0.79 | 0.453 | 1.36 | 0.254947 |
| 552 | 6Z_Octadecenoicacid | [M+H]+1 | POS | Mass, Spectral match | Hilic | 282.255 | 3.984 | | HMDB40900 | 5283009 | Biosynthesis of plant secondary metabolites;Other | 0.84 | 0.105 | 0.98 | 0.796 | 1.15 | 0.458 | 0.85 | 0.251086 |
| 313 | D_Glucose | [M-H]-1 | NEG | Mass, Spectral match | C18 | 180.063 | 0.721 | C00031 | HMDB0000122 | 5793 | | 2.36 | 0.039 | 1.94 | 0.185 | 2.85 | 0.048 | 0.68 | 0.282087 |

| | | | | | | | | | | | | | | | | | | | |
|---|---|---|---|---|---|---|---|---|---|---|---|---|---|---|---|---|---|---|---|
| 554 | Guanosine | [M-H]-1 | NEG | Mass, Spectral match | Hilic | 283.091 | 5.756 | C00387 | HMDB0000133 | 6802 | Metabolic pathways;Other | 0.18 | 0.085 | 0.43 | 0.258 | 0.39 | 0.790 | 1.12 | 0.908282 |
| 555 | Octadecanamide | [M+H]+1 | POS | Mass, RT, Spectral match | C18 | 283.287 | 13.701 | C13846 | HMDB0034146 | 31292 | | 0.88 | 0.484 | 0.99 | 0.943 | 1.00 | 0.569 | 0.99 | 0.858195 |
| 556 | Xanthosine | [M-H]-1 | NEG | Mass,RT, Spectral match | Hilic | 284.075 | 6.876 | C01762 | HMDB0000299 | 64959 | Metabolic pathways;Biosynthesis of secondary metabolites;Other | 1.98 | 0.281 | 1.27 | 0.649 | 1.34 | 0.158 | 0.95 | 0.815586 |
| 557 | Xanthosine | [M+Cl]-1 | NEG | Mass, RT, Spectral match | C18 | 284.075 | 0.646 | C01762 | HMDB0000299 | 64959 | Metabolic pathways;Biosynthesis of secondary metabolites;Other | 1.48 | 0.308 | 1.02 | 0.958 | 1.04 | 0.235 | 0.98 | 0.82869 |
| 558 | Retinal | [M+H]+1 | POS | Mass, RT, Spectral match | C18 | 284.213 | 11.489 | C00376 | HMDB0001358 | 638015 | Metabolic pathways;Other | 0.72 | 0.356 | 1.44 | 0.435 | 0.77 | 0.429 | 1.88 | 0.466246 |
| 559 | Stearic_acid | [M-H]-1 | NEG | Mass, Spectral match | C18 | 284.272 | 14.260 | C01530 | HMDB0000827 | 5281 | Biosynthesis of plant secondary metabolites;Other | 1.32 | 0.558 | 1.22 | 0.459 | 1.54 | 0.361 | 0.80 | 0.927093 |
| 560 | Diflunisal | [M-H]-1 | NEG | Mass, Spectral match | Hilic | 285.010 | 7.011 | C01691 | HMDB0014999 | 3059 | | 0.97 | 0.837 | 0.85 | 0.232 | 0.65 | 0.657 | 1.30 | 0.159994 |
| 561 | Vitamin_A | [M+H]+1 | POS | Mass, Spectral match | C18 | 285.222 | 12.914 | C00473 | HMDB0000305 | 445354 | | 0.85 | 0.666 | 0.74 | 0.349 | 0.66 | 0.421 | 1.12 | 0.409049 |
| 562 | N_Acetyl_L_tyrosine | [M+H]+1 | POS | Mass, Spectral match | Hilic | 286.083 | 6.910 | C01657 | HMDB0000866 | 68310 | Biosynthesis of secondary metabolites;Biosynthesis of antibiotics;Other | 0.70 | 0.360 | 0.86 | 0.723 | 1.35 | 0.450 | 0.64 | 0.314806 |
| 563 | Cycasin | [M-H]-1 | NEG | Mass, Spectral match | Hilic | 287.067 | 10.447 | C01418 | HMDB38850 | 5459896 | | 0.91 | 0.841 | 0.52 | 0.104 | 0.58 | 0.870 | 0.89 | 0.854409 |
| 564 | alpha_Hexachlorocyclohexane | [M-H]-1 | NEG | Mass, Spectral match | Hilic | 287.861 | 0.784 | C07075 | HMDB0014575 | NA | Metabolic pathways;Microbial metabolism in diverse environments;Other | 4.29 | 0.028 | 5.07 | 0.054 | 7.86 | 0.132 | 0.64 | 0.676124 |
| 565 | Atropine | [M+Na]+1 | POS | Mass, Spectral match | Hilic | 289.167 | 0.696 | C01479 | HMDB0014712 | NA | Metabolic pathways;Biosynthesis of secondary metabolites;Other | 1.01 | 0.913 | 1.01 | 0.863 | 1.07 | 0.818 | 0.94 | 0.213318 |
| 566 | Phenyllactic_acid | [M+H]+1 | POS | Mass, Spectral match | C18 | 289.167 | 5.236 | C01479 | HMDB0000779 | 3848 | Metabolic pathways;Biosynthesis of secondary metabolites;Tropane, piperidine and pyridine alkaloid biosynthesis;Biosynthesis of alkaloids derived from ornithine, lysine and nicotinic acid | 0.97 | 0.312 | 0.95 | 0.148 | 0.99 | 0.964 | 0.95 | 0.202993 |
| 567 | Androsterone | [M+H]+1 | POS | Mass, Spectral match | C18 | 290.224 | 11.929 | C00523 | HMDB0000031 | 12306765 | Microbial metabolism in diverse environments;Other | 0.92 | 0.390 | 0.97 | 0.781 | 0.91 | 0.736 | 1.06 | 0.525566 |
| 568 | Aspartame | [M-H]-1 | NEG | Mass, Spectral match | Hilic | 294.121 | 5.347 | C11045 | HMDB0001894 | 134601 | Other | 1.99 | 0.075 | 1.28 | 0.558 | 1.36 | 0.623 | 0.94 | 0.949978 |
| 569 | Gingerol | [M+Cl]-1 | NEG | Mass, Spectral match | C18 | 294.183 | 9.196 | C10462 | HMDB0005783 | 3473 | Biosynthesis of secondary metabolites;Other | 0.97 | 0.364 | 1.02 | 0.536 | 1.03 | 0.267 | 0.99 | 0.814656 |
| 570 | Trimipramine | [M+H]+1 | POS | Mass, Spectral match | C18 | 294.209 | 7.870 | C14029 | #N/A | 5282318 | | 1.07 | 0.934 | 1.41 | 0.649 | 0.39 | 0.836 | 3.61 | 0.190174 |
| 571 | Monosaccharide_1_phosphate | [M-H]-1 | NEG | Mass, Spectral match | Hilic | 297.059 | 14.799 | C00994 | #N/A | 439361 | | 0.62 | 0.170 | 0.29 | 0.004 | 0.32 | 0.172 | 0.92 | 0.827833 |

| | | | | | | | | | | | | | | | | | | | |
|---|---|---|---|---|---|---|---|---|---|---|---|---|---|---|---|---|---|---|---|
| 572 | N2_Methylguanosine | [M+H]+1 | POS | Mass, Spectral match | Hilic | 297.107 | 2.615 | C04153 | HMDB0006040 | 24828 | | 1.27 | 0.499 | 1.00 | 0.995 | 0.70 | 0.254 | 1.42 | 0.028007 |
| 573 | 2_Methylguanosine | [M+H]+1 | POS | Mass, Spectral match | C18 | 297.107 | 0.658 | C00837 | #N/A | 439323 | | 0.92 | 0.843 | 0.71 | 0.403 | 0.50 | 0.174 | 1.41 | 0.150155 |
| 574 | 5_Methylthioadenosine | [M+H]+1 | POS | Mass, Spectral match | C18 | 298.098 | 1.753 | C00170 | HMDB0001173 | 439176 | | 0.41 | 0.171 | 1.15 | 0.810 | 0.41 | 0.845 | 2.83 | 0.567601 |
| 575 | 2_Oxooctadecanoicacid | [M+H]+1 | POS | Mass, Spectral match | Hilic | 298.250 | 4.030 | | HMDB40901 | 5283010 | Other | 1.04 | 0.895 | 1.66 | 0.019 | 1.19 | 0.360 | 1.40 | 0.152166 |
| 576 | 2_Oxooctadecanoicacid | [M-H]-1 | NEG | Mass, Spectral match | C18 | 298.251 | 12.471 | C00869 | #N/A | 439332 | Other | 1.61 | 0.328 | 1.21 | 0.390 | 1.11 | 0.494 | 1.09 | 0.609872 |
| 577 | D_4_Phosphopantothenate | [M-H]-1 | NEG | Mass, RT, Spectral match | C18 | 299.077 | 0.641 | C00994 | #N/A | 439361 | Metabolic pathways;Other | 1.17 | 0.423 | 1.05 | 0.833 | 0.91 | 0.538 | 1.15 | 0.732302 |
| 578 | Sphingosine | [M+H]+1 | POS | Mass, RT, Spectral match | C18 | 299.282 | 10.598 | C00319 | HMDB0000252 | 5353955 | Metabolic pathways;Neuroactive ligand-receptor interaction;Other | 2.25 | 0.297 | 1.85 | 0.081 | 1.28 | 0.531 | 1.44 | 0.329434 |
| 579 | beta_D_3_Ribofuranosyluric_acid | [M-H]-1 | NEG | Mass, Spectral match | Hilic | 300.070 | 5.412 | C05513 | HMDB0029920 | NA | Other | 1.23 | 0.607 | 0.84 | 0.571 | 0.60 | 0.757 | 1.39 | 0.374031 |
| 580 | 2_Methoxyestrone | [M-H]-1 | NEG | Mass, Spectral match | Hilic | 300.172 | 1.050 | C05299 | HMDB0000010 | 440624 | Metabolic pathways;Microbial metabolism in diverse environments;Other | 1.04 | 0.504 | 0.96 | 0.595 | 1.03 | 0.861 | 0.94 | 0.247577 |
| 581 | Testolactone | [M-H]-1 | NEG | Mass, RT, Spectral match | C18 | 300.172 | 7.160 | C02197 | HMDB0015031 | 13769 | Metabolic pathways;Microbial metabolism in diverse environments;Other | 1.00 | 0.940 | 0.97 | 0.327 | 0.96 | 0.120 | 1.01 | 0.778688 |
| 582 | N_Acetyl_D_Glucosamine_6_Phosphate | [M-H]-1 | NEG | Mass, Spectral match | C18 | 301.057 | 0.531 | C00357 | HMDB0001062 | 440996 | Metabolic pathways;Biosynthesis of antibiotics;Other | 0.89 | 0.635 | 1.20 | 0.607 | 0.95 | 0.807 | 1.26 | 0.606816 |
| 583 | Sphinganine | [M+H]+1 | POS | Mass, Spectral match | Hilic | 301.297 | 0.871 | C00836 | HMDB0000269 | 91486 | Metabolic pathways;Other | 2.73 | 0.123 | 1.07 | 0.777 | 0.89 | 0.712 | 1.20 | 0.372988 |
| 584 | Sphinganine | [M+H]+1 | POS | Mass, RT, Spectral match | C18 | 301.297 | 10.837 | C00836 | HMDB0000269 | 91486 | Metabolic pathways;Other | 2.09 | 0.299 | 1.26 | 0.544 | 1.24 | 0.337 | 1.02 | 0.695044 |
| 585 | Hesperetin | [M-H]-1 | NEG | Mass, Spectral match | C18 | 302.078 | 18.553 | C01709 | HMDB0005782 | 72281 | Biosynthesis of antibiotics;Other | 1.09 | 0.180 | 0.93 | 0.478 | 0.94 | 0.974 | 0.99 | 0.811322 |
| 586 | Abietic_acid | [M+H]+1 | POS | Mass, Spectral match | Hilic | 302.224 | 4.098 | NA | #N/A | #N/A | Metabolic pathways;Biosynthesis of secondary metabolites;Other | 0.87 | 0.093 | 0.85 | 0.026 | 0.83 | 0.109 | 1.03 | 0.729745 |
| 587 | Taxifolin | [M-H]-1 | NEG | Mass, Spectral match | Hilic | 303.053 | 4.896 | C01617 | METPA0191 | NA | | 0.33 | 0.208 | 1.10 | 0.883 | 0.41 | 0.175 | 2.69 | 0.134233 |
| 588 | 312_Dihydroxy_13_methoxypodocarpa_81113_trien_7_one | [M+H]+1 | POS | Mass, Spectral match | C18 | 304.164 | 9.357 | C01804 | #N/A | 194139 | Biosynthesis of secondary metabolites;Other | 0.89 | 0.294 | 0.96 | 0.690 | 0.85 | 0.984 | 1.12 | 0.30116 |
| 589 | Arachidonic_acid | [M-H]-1 | NEG | Mass, Spectral match | C18 | 304.240 | 12.995 | C00219 | HMDB0001043 | 444899 | Metabolic pathways;Biosynthesis of secondary metabolites;Other | 1.03 | 0.813 | 1.14 | 0.457 | 1.27 | 0.551 | 0.90 | 0.59947 |
| 590 | Arachidonicacid | [M-H]-1 | NEG | Mass, Spectral match | Hilic | 304.240 | 0.945 | C00219 | HMDB0001043 | 444899 | Metabolic pathways;Biosynthesis of secondary metabolites;Other | 0.89 | 0.622 | 1.11 | 0.677 | 1.18 | 0.640 | 0.94 | 0.72513 |
| 591 | Apigenin | [M-H]-1 | NEG | Mass, Spectral | C18 | 305.019 | 6.478 | C01477 | HMDB0002124 | 5280443 | | 1.96 | 0.090 | 1.43 | 0.503 | 1.37 | 0.586 | 1.04 | 0.639956 |

| | | | | | | | | | | | | | | | | | | | |
|---|---|---|---|---|---|---|---|---|---|---|---|---|---|---|---|---|---|---|---|
| | | | | match | | | | | | | | | | | | | | | |
| 592 | Apigenin | [M-H]-1 | NEG | Mass, Spectral match | Hilic | 305.019 | 2.911 | C01477 | HMDB0002124 | 5280443 | | 1.82 | 0.329 | 1.30 | 0.678 | 1.24 | 0.942 | 1.05 | 0.580955 |
| 593 | D_Lombricine | [M-H]-1 | NEG | Mass, Spectral match | Hilic | 305.041 | 23.332 | C01726 | METPA0215 | NA | Pyrimidine metabolism | 0.71 | 0.407 | 1.09 | 0.848 | 0.54 | 0.364 | 2.02 | 0.219248 |
| 594 | 6S_Hydroxyhyoscyamine | [M+H]+1 | POS | Mass, RT, Spectral match | C18 | 305.162 | 5.578 | C01564 | #N/A | 2680 | Metabolic pathways;Biosynthesis of secondary metabolites;Tropane, piperidine and pyridine alkaloid biosynthesis | 0.90 | 0.129 | 0.92 | 0.252 | 1.00 | 0.681 | 0.92 | 0.161204 |
| 595 | N_Butyl_N_2_phenoxyphenylurea | [M+H]+1 | POS | Mass, Spectral match | Hilic | 306.140 | 5.736 | C19415 | HMDB0032565 | 9013 | | 0.57 | 0.267 | 1.09 | 0.819 | 0.99 | 0.177 | 1.10 | 0.791944 |
| 596 | Glutathione_ | [M-H]-1 | NEG | Mass, RT, Spectral match | C18 | 307.084 | 0.553 | C00051 | HMDB0000125 | 124886 | Metabolic pathways;Bile secretion;ABC transporters;Other | 2.43 | 0.077 | 1.53 | 0.103 | 2.42 | 0.139 | 0.63 | 0.823919 |
| 597 | Hydroxycarteolol | [M-H]-1 | NEG | Mass, Spectral match | C18 | 308.173 | 6.213 | C00732 | #N/A | 439299 | | 1.04 | 0.892 | 1.25 | 0.426 | 1.01 | 0.159 | 1.23 | 0.537036 |
| 598 | N_Acetylneuraminicacid | [M-H]-1 | NEG | Mass,RT, Spectral match | Hilic | 309.106 | 18.883 | C19910 | HMDB0000230 | 445063 | Metabolic pathways;Other | 1.15 | 0.631 | 1.17 | 0.710 | 1.40 | 0.294 | 0.84 | 0.668862 |
| 599 | N_Acetylneuraminicacid NeuAc | [M-H]-1 | NEG | Mass, RT, Spectral match | C18 | 309.106 | 0.543 | C00270 | #N/A | 439197 | Metabolic pathways;Other | 1.09 | 0.736 | 1.39 | 0.424 | 1.17 | 0.403 | 1.19 | 0.811681 |
| 600 | Taxiphyllin | [M+H]+1 | POS | Mass, RT, Spectral match | C18 | 311.103 | 0.628 | C01855 | HMDB0030704 | 4475719 | | 0.68 | 0.276 | 0.74 | 0.501 | 0.93 | 0.837 | 0.80 | 0.904709 |
| 601 | Thebaine | [M+H]+1 | POS | Mass, RT, Spectral match | C18 | 311.152 | 10.973 | | HMDB0029378 | 408120 | Metabolic pathways;Biosynthesis of secondary metabolites;Biosynthesis of alkaloids derived from shikimate pathway;Other | 0.74 | 0.220 | 0.79 | 0.345 | 0.62 | 0.354 | 1.27 | 0.38493 |
| 602 | 7S_8S_DiHODE | [M-H]-1 | NEG | Mass, Spectral match | Hilic | 312.230 | 1.008 | C07354 | METPA0927 | NA | Metabolic pathways;Biosynthesis of secondary metabolites;Other | 0.77 | 0.040 | 1.31 | 0.065 | 7.28 | 0.366 | 0.18 | 0.644025 |
| 603 | 7S8S_DiHODE | [M-H]-1 | NEG | Mass, RT, Spectral match | C18 | 312.230 | 9.708 | C07354 | METPA0927 | NA | Metabolic pathways;Biosynthesis of secondary metabolites;Other | 0.79 | 0.047 | 0.88 | 0.460 | 4.92 | 0.378 | 0.18 | 0.407483 |
| 604 | Geranyl_PP | [M-H]-1 | NEG | Mass, Spectral match | Hilic | 313.060 | 9.343 | C00341 | HMDB0001285 | 445995 | | 0.83 | 0.685 | 0.59 | 0.324 | 1.21 | 0.506 | 0.49 | 0.748544 |
| 605 | 78_Dihydropteroic_acid | [M+H]+1 | POS | Mass, Spectral match | Hilic | 313.105 | 7.881 | C00921 | HMDB0001412 | 170 | | 1.14 | 0.409 | 0.79 | 0.237 | 0.99 | 0.661 | 0.81 | 0.401629 |
| 606 | octadecenoicacid | [M-H]-1 | NEG | Mass, Spectral match | C18 | 314.246 | 13.508 | | HMDB40900 | 5283009 | Other | 0.80 | 0.646 | 0.93 | 0.864 | 1.31 | 0.260 | 0.71 | 0.252446 |
| 607 | Phytosphingosine | [M+H]+1 | POS | Mass, Spectral match | C18 | 317.292 | 9.282 | C12144 | HMDB0004610 | 122121 | Metabolic pathways;Other | 0.88 | 0.161 | 0.99 | 0.865 | 1.06 | 0.116 | 0.93 | 0.378516 |
| 608 | Leukotriene_A4 | [M-H]-1 | NEG | Mass, Spectral match | C18 | 318.219 | 10.769 | C00909 | HMDB0001337 | 5280383 | Metabolic pathways;Biosynthesis of secondary metabolites;Other | 0.56 | 0.105 | 1.13 | 0.834 | 0.35 | 0.075 | 3.24 | 0.095618 |
| 609 | Morphine | [M-H]-1 | NEG | Mass, Spectral match | Hilic | 320.107 | 1.246 | C01516 | HMDB0014440 | 5288826 | | 1.81 | 0.089 | 3.25 | 0.051 | 4.00 | 0.074 | 0.81 | 0.770684 |

| | | | | | | | | | | | | | | | | | | | |
|---|---|---|---|---|---|---|---|---|---|---|---|---|---|---|---|---|---|---|---|
| 610 | Mycophenolic_acid | [M+H]+1 | POS | Mass, Spectral match | Hilic | 320.123 | 1.207 | | HMDB0015159 | 446541 | | 0.99 | 0.933 | 0.62 | 0.018 | 0.84 | 0.609 | 0.74 | 0.089886 |
| 611 | Testolactone | [M+H]+1 | POS | Mass, Spectral match | Hilic | 320.198 | 4.122 | C02197 | HMDB0015031 | 13769 | | 0.81 | 0.024 | 0.84 | 0.028 | 0.84 | 0.262 | 1.01 | 0.844263 |
| 612 | 20_Hydroxyeicosatetraenoic_acid | [M-H]-1 | NEG | Mass, Spectral match | C18 | 320.235 | 11.060 | C14748 | HMDB0005998 | 5283157 | Metabolic pathways;Other | 0.74 | 0.329 | 1.23 | 0.480 | 0.73 | 0.440 | 1.68 | 0.144027 |
| 613 | 1112_Epoxyeicosatrienoic_acid | [M-H]-1 | NEG | Mass, Spectral match | C18 | 320.235 | 11.280 | C14770 | HMDB0004673 | 5353269 | Metabolic pathways;Other | 0.60 | 0.131 | 1.20 | 0.701 | 0.59 | 0.217 | 2.03 | 0.116674 |
| 614 | Digallic_acid | [M-H]-1 | NEG | Mass, Spectral match | Hilic | 321.026 | 10.126 | C01572 | #N/A | 341 | | 0.59 | 0.030 | 0.41 | 0.001 | 0.32 | 0.139 | 1.27 | 0.315168 |
| 615 | 5_Thymidylic_acid | [M-H]-1 | NEG | Mass, Spectral match | Hilic | 321.047 | 1.177 | C00364 | HMDB0001227 | 9700 | | 0.73 | 0.554 | 0.25 | 0.047 | 0.20 | 0.040 | 1.26 | 0.484817 |
| 616 | Salicin | [M-H]-1 | NEG | Mass, Spectral match | Hilic | 321.076 | 14.797 | C01451 | HMDB0003546 | 439503 | | 0.99 | 0.973 | 0.68 | 0.200 | 0.71 | 0.849 | 0.97 | 0.933953 |
| 617 | 5_O_1_Carboxyvinyl_3_phosphoshikimate | [M-H]-1 | NEG | Mass, Spectral match | Hilic | 323.020 | 7.312 | C01269 | METPA0160 | NA | | 0.65 | 0.028 | 0.48 | 0.001 | 0.35 | 0.113 | 1.37 | 0.23629 |
| 618 | Cytidine_monophosphate | [M+H]+1 | POS | Mass, Spectral match | C18 | 323.052 | 0.512 | C00055 | HMDB0000095 | 6131 | Metabolic pathways;Other | 0.98 | 0.885 | 1.33 | 0.041 | 0.94 | 0.484 | 1.41 | 0.010298 |
| 619 | Cytidine_phosphate | [M-H]-1 | NEG | Mass, Spectral match | Hilic significant | 323.052 | 24.124 | C00239 | HMDB0001202 | 13945 | Metabolic pathways;Pyrimidine metabolism | 1.56 | 0.184 | 1.70 | 0.160 | 2.75 | 0.030 | 0.62 | 0.236025 |
| 620 | Cytidine_PO4_ | [M+H]+1 | POS | Mass, RT, Spectral match | C18 significant | 323.052 | 0.520 | C00239 | HMDB0001202 | 13945 | Metabolic pathways;Other | 0.98 | 0.885 | 1.33 | 0.041 | 0.94 | 0.484 | 1.41 | 0.010298 |
| 621 | N_Glycolylneuraminic_acid | [M-H]-1 | NEG | Mass, Spectral match | C18 | 325.102 | 0.518 | C03410 | HMDB0000833 | 123802 | Other | 0.63 | 0.112 | 0.71 | 0.231 | 0.55 | 0.326 | 1.30 | 0.3846 |
| 622 | Imidocarb | [M+H]+1 | POS | Mass, Spectral match | Hilic | 326.193 | 1.158 | | HMDB0031801 | 5372582 | | 0.88 | 0.136 | 0.85 | 0.023 | 0.92 | 0.560 | 0.93 | 0.655368 |
| 623 | DOCEBENONE | [M+H]+1 | POS | Mass, Spectral match | C18 | 327.198 | 10.946 | C01349 | #N/A | 1967 | | 1.04 | 0.937 | 1.33 | 0.599 | 1.47 | 0.160 | 0.90 | 0.87327 |
| 624 | NN_Dimethylsphingosine | [M+H]+1 | POS | Mass, Spectral match | Hilic | 327.313 | 4.227 | C13914 | HMDB0013645 | 5282309 | | 0.89 | 0.238 | 0.85 | 0.116 | 0.85 | 0.051 | 1.01 | 0.892503 |
| 625 | Desulfoglucotropeolin | [M+H]+1 | POS | Mass, Spectral match | Hilic | 328.087 | 6.905 | C01069 | METPA1778 | NA | | 1.42 | 0.408 | 1.62 | 0.193 | 1.42 | 0.201 | 1.14 | 0.90829 |
| 626 | 7_Hydroxy_6_methyl_8_ribityllumazine | [M-H]-1 | NEG | Mass, Spectral match | Hilic | 328.100 | 9.778 | C05995 | HMDB0004256 | NA | Other | 0.79 | 0.567 | 1.64 | 0.160 | 1.64 | 0.195 | 1.00 | 0.889785 |
| 627 | 17_Hydroxyprogesterone | [M+H]+1 | POS | Mass, Spectral match | C18 | 330.216 | 0.463 | C01176 | HMDB0000374 | 6238 | | 1.57 | 0.074 | 1.59 | 0.072 | 1.70 | 0.122 | 0.94 | 0.89957 |
| 628 | 1_palmitoylglycerol | [M+Na]+1 | POS | Mass, Spectral match | Hilic | 330.276 | 4.181 | | HMDB0031074 | 14900 | | 0.81 | 0.186 | 0.82 | 0.005 | 0.81 | 0.281 | 1.02 | 0.769795 |
| 629 | Deoxyadenosine_monophosphate | [M-H]-1 | NEG | Mass, RT, Spectral match | C18 | 331.067 | 0.504 | C00360 | HMDB0000905 | 12599 | | 0.56 | 0.051 | 0.53 | 0.062 | 0.59 | 0.900 | 0.90 | 0.508785 |

| | | | | | | | | | | | | | | | | | | | |
|---|---|---|---|---|---|---|---|---|---|---|---|---|---|---|---|---|---|---|---|
| 630 | Tos_Lys_CH2Cl | [M+H]+1 | POS | Mass,RT, Spectral match | Hilic | 331.088 | 3.170 | C00896 | #N/A | 73094 | | 1.74 | 0.572 | 0.60 | 0.623 | 0.61 | 0.273 | 0.99 | 0.703195 |
| 631 | 2_deoxyinosine5_monophosphatedIMP | [M+Na]+1 | POS | Mass, Spectral match | Hilic | 332.053 | 15.660 | C06196 | HMDB0006555 | 91531 | Other | 1.17 | 0.546 | 0.97 | 0.895 | 1.22 | 0.367 | 0.79 | 0.233102 |
| 632 | ?_Nicotinamidemononucleotide | [M-H]-1 | NEG | Mass, Spectral match | Hilic | 334.056 | 14.531 | C01185 | HMDB0001132 | 53477721 | Metabolic pathways;Other | 1.88 | 0.340 | 1.72 | 0.264 | 1.16 | 0.264 | 1.48 | 0.623821 |
| 633 | Dicumarol | [M-H]-1 | NEG | Mass, Spectral match | Hilic | 335.055 | 11.286 | C00796 | HMDB0014411 | 54676038 | | 1.77 | 0.304 | 0.70 | 0.255 | 1.44 | 0.109 | 0.48 | 0.083208 |
| 634 | Prostaglandin_B1 | [M+H]+1 | POS | Mass, Spectral match | C18 | 336.227 | 10.014 | C00959 | HMDB0002982 | 5280388 | | 1.03 | 0.869 | 1.19 | 0.277 | 1.12 | 0.516 | 1.07 | 0.553741 |
| 487 | Uridine | [M-H]-1 | NEG | Mass, Spectral match | C18 | 244.069 | 0.612 | C00299 | HMDB0000296 | 6029 | Metabolic pathways;Other | 1.26 | 0.345 | 0.98 | 0.901 | 1.18 | 0.344 | 0.82 | 0.435506 |
| 636 | dCMP | [M-H]-1 | NEG | Mass, Spectral match | Hilic | 342.022 | 2.095 | C00239 | HMDB0001202 | 13945 | | 4.47 | 0.250 | 2.42 | 0.352 | 1.78 | 0.183 | 1.36 | 0.854545 |
| 511 | Palmitic_acid | [M-H]-1 | NEG | Mass, RT, Spectral match | C18 | 256.240 | 13.520 | C00249 | HMDB0000220 | 985 | Metabolic pathways;Biosynthesis of plant secondary metabolites;Other | 1.19 | 0.608 | 1.21 | 0.520 | 1.72 | 0.313 | 0.70 | 0.809204 |
| 531 | Inosine | [M-H]-1 | NEG | Mass, Spectral match | C18 | 268.080 | 0.533 | C00294 | HMDB0000195 | 6021 | Metabolic pathways;Other | 0.14 | 0.103 | 0.30 | 0.190 | 0.42 | 0.648 | 0.72 | 0.848565 |
| 639 | Thiamine_monophosphate | [M+H]+1 | POS | Mass, Spectral match | Hilic | 344.070 | 8.797 | C01081 | HMDB0002666 | 3382778 | | 1.83 | 0.373 | 1.70 | 0.247 | 1.38 | 0.227 | 1.23 | 0.650453 |
| 640 | Adenosine_monophosphate | [M+Na]+1 | POS | Mass, Spectral match | Hilic | 346.057 | 7.871 | C00020 | HMDB0000045 | 6083 | | 0.97 | 0.866 | 1.33 | 0.127 | 1.37 | 0.136 | 0.97 | 0.995748 |
| 641 | Gibberellin_A3 | [M-H]-1 | NEG | Mass, RT, Spectral match | C18 | 346.139 | 10.574 | C01699 | HMDB0003559 | 9819600 | | 0.87 | 0.334 | 0.78 | 0.163 | 0.82 | 0.903 | 0.94 | 0.583377 |
| 642 | Adenosine_monophosphate | [M-H]-1 | NEG | Mass, RT, Spectral match | C18 | 347.063 | 0.547 | C00020 | HMDB0000045 | 6083 | Metabolic pathways;Biosynthesis of secondary metabolites;Biosynthesis of antibiotics;Biosynthesis of plant secondary metabolites;Other | 1.10 | 0.712 | 1.50 | 0.393 | 1.25 | 0.359 | 1.20 | 0.875397 |
| 643 | Gibberellin_A1 | [M+H]+1 | POS | Mass, Spectral match | Hilic | 347.148 | 6.939 | C00859 | METPA0089 | NA | | 0.44 | 0.019 | 0.46 | 0.029 | 0.34 | 0.110 | 1.38 | 0.601641 |
| 644 | Pentyl_2_furyl_ketone | [M+H]+1 | POS | Mass, Spectral match | Hilic | 350.063 | 10.459 | C02088 | HMDB0032462 | 61738 | | 0.92 | 0.841 | 0.44 | 0.046 | 0.49 | 0.619 | 0.90 | 0.921465 |
| 645 | 25_Diamino_6_5_phosphoribosylamino_4_pyrimidineone | [M-H]-1 | NEG | Mass, Spectral match | Hilic | 352.063 | 6.868 | C01304 | METPA0171 | NA | Other | 1.25 | 0.564 | 0.81 | 0.594 | 0.84 | 0.246 | 0.97 | 0.832433 |
| 646 | 25_Diamino_6_5_phosphoribosylamino_4_pyrimidineone | [M-H]-1 | NEG | Mass, RT, Spectral match | C18 | 352.064 | 0.595 | C01304 | METPA0171 | NA | Other | 23.96 | 0.101 | 1.61 | 0.490 | 17.86 | 0.052 | 0.09 | 0.052851 |

| | | | | | | | | | | | | | | | | | | | |
|---|---|---|---|---|---|---|---|---|---|---|---|---|---|---|---|---|---|---|---|
| 647 | S_Reticuline | [M+H]+1 | POS | Mass, RT, Spectral match | C18 | 352.149 | 8.856 | C02105 | HMDB0003601 | 439653 | | 0.68 | 0.355 | 0.65 | 0.314 | 0.64 | 0.803 | 1.02 | 0.892427 |
| 648 | Prostaglandin_H2 | [M-H]-1 | NEG | Mass, RT, Spectral match | C18 | 352.224 | 7.376 | C00427 | HMDB0001381 | 445049 | Metabolic pathways;Biosynthesis of antibiotics;Bile secretion;Neuroactive ligand-receptor interaction;Other | 0.54 | 0.314 | 0.93 | 0.865 | 0.24 | 0.032 | 3.86 | 0.006188 |
| 649 | 5_Amino_6_5_phosphoribosylaminouracil | [M-H]-1 | NEG | Mass, Spectral match | Hilic | 353.048 | 6.015 | C01268 | METPA0159 | NA | | 1.09 | 0.801 | 1.80 | 0.193 | 1.62 | 0.153 | 1.11 | 0.87539 |
| 650 | Chlorogenic_acid | [M+H]+1 | POS | Mass, Spectral match | Hilic | 353.091 | 6.932 | C00852 | HMDB0003164 | 1794427 | | 0.43 | 0.013 | 0.57 | 0.051 | 0.71 | 0.813 | 0.80 | 0.843915 |
| 651 | Prostaglandin_F2a | [M-H]-1 | NEG | Mass, RT, Spectral match | C18 | 354.241 | 8.199 | C00639 | HMDB0001139 | 5283078 | Metabolic pathways;Bile secretion;Neuroactive ligand-receptor interaction;Other | 0.48 | 0.221 | 0.75 | 0.503 | 0.21 | 0.031 | 3.63 | 0.00375 |
| 652 | S_Adenosylmethionine | [M-H]-1 | NEG | Mass, RT, Spectral match | C18 | 356.165 | 13.520 | C01137 | HMDB0000988 | 439415 | | 0.84 | 0.536 | 0.85 | 0.560 | 0.87 | 0.860 | 0.98 | 0.700379 |
| 653 | Docosahexaenoic_acid_ethyl_ester | [M+H]+1 | POS | Mass, Spectral match | C18 | 356.271 | 9.167 | C16185 | #N/A | 9831416 | | 2.74 | 0.227 | 10.91 | 0.127 | 1.25 | 0.893 | 8.73 | 0.092539 |
| 654 | E_64 | [M-H]-1 | NEG | Mass, Spectral match | C18 | 357.201 | 3.625 | C01341 | #N/A | 123985 | | 1.02 | 0.934 | 1.19 | 0.500 | 1.03 | 0.134 | 1.15 | 0.682802 |
| 655 | Pantetheine_4_phosphate | [M-H]-1 | NEG | Mass, Spectral match | C18 | 358.097 | 1.032 | C01134 | HMDB0001416 | 987 | Metabolic pathways;Biosynthesis of antibiotics;Other | 3.34 | 0.114 | 2.21 | 0.028 | 2.36 | 0.189 | 0.94 | 0.645724 |
| 656 | Prednisone | [M+Cl]-1 | NEG | Mass, Spectral match | C18 | 358.178 | 7.159 | C07370 | HMDB0014773 | 5865 | | 1.00 | 0.979 | 0.99 | 0.568 | 0.98 | 0.608 | 1.00 | 0.932953 |
| 657 | Prednisone | [M+Cl]-1 | POS | Mass, Spectral match | Hilic | 358.178 | 1.056 | C07370 | HMDB0014773 | 5865 | | 1.08 | 0.065 | 1.03 | 0.442 | 1.08 | 0.439 | 0.95 | 0.196103 |
| 658 | 1_Stearoylglycerol | [M+Na]+1 | POS | Mass, Spectral match | C18 | 358.307 | 13.866 | D01947 | #N/A | 24699 | | 1.37 | 0.474 | 0.96 | 0.893 | 1.44 | 0.282 | 0.67 | 0.652947 |
| 659 | Stearyl_monoglyceridyl_citrate | [M+H]+1 | POS | Mass, Spectral match | Hilic | 358.308 | 4.357 | | HMDB0032522 | 62110 | | 1.13 | 0.398 | 1.02 | 0.903 | 0.93 | 0.246 | 1.09 | 0.494436 |
| 660 | Rosmarinic_acid | [M-H]-1 | NEG | Mass, Spectral match | Hilic | 359.077 | 5.952 | C01850 | HMDB0003572 | 5281792 | | 1.30 | 0.145 | 1.19 | 0.466 | 1.20 | 0.235 | 0.99 | 0.825433 |
| 661 | _Lactose | [M+H]+1 | POS | Mass, Spectral match | Hilic | 359.142 | 9.495 | C00243 | HMDB0000186 | 84571 | | 1.38 | 0.441 | 2.64 | 0.008 | 2.78 | 0.079 | 0.95 | 0.995853 |
| 662 | Cortisone | [M+H]+1 | POS | Mass, Spectral match | C18 | 360.190 | 10.179 | C00762 | HMDB0002802 | 222786 | | 1.08 | 0.123 | 1.01 | 0.722 | 1.04 | 0.836 | 0.98 | 0.546633 |
| 663 | Cortisol | [M+H]+1 | POS | Mass, Spectral match | C18 | 363.219 | 11.812 | C00735 | HMDB0000063 | 657311 | | 0.95 | 0.511 | 0.95 | 0.711 | 0.98 | 0.346 | 0.97 | 0.653191 |
| 664 | D_+_Maltose | [M+H]+1 | POS | Mass, Spectral match | Hilic | 364.097 | 9.492 | C00208 | HMDB0000163 | 10991489 | | 1.51 | 0.431 | 1.78 | 0.002 | 1.71 | 0.094 | 1.04 | 0.670446 |
| 665 | 37_Dimethylquercetin | [M-H]-1 | NEG | Mass, Spectral match | Hilic | 365.046 | 14.794 | C01265 | HMDB0029263 | 5280417 | | 0.48 | 0.086 | 0.19 | 0.006 | 0.16 | 0.039 | 1.16 | 0.984522 |
| 666 | Deoxyadenosine_monop | [M+H]+1 | POS | Mass, Spectral | Hilic | 366.040 | 10.492 | C00360 | HMDB0000905 | 12599 | | 0.85 | 0.739 | 0.44 | 0.081 | 0.49 | 0.675 | 0.89 | 0.903066 |

| | | | | | | | | | | | | | | | | | | | |
|---|---|---|---|---|---|---|---|---|---|---|---|---|---|---|---|---|---|---|---|
| | hosphate | | | match | | | | | | | | | | | | | | | |
| 667 | Adipic_acid | [M+H]+1 | POS | Mass, Spectral match | Hilic | 370.308 | 28.558 | C06104 | HMDB0000448 | 196 | Other | 1.17 | 0.044 | 1.09 | 0.119 | 1.13 | 0.752 | 0.97 | 0.576653 |
| 668 | Berberine | [M-H]-1 | NEG | Mass, Spectral match | Hilic | 371.096 | 10.238 | C00757 | HMDB0003409 | 2353 | | 0.98 | 0.940 | 0.67 | 0.225 | 1.25 | 0.064 | 0.54 | 0.081368 |
| 669 | Haloperidol | [M-H]-1 | NEG | Mass, RT, Spectral match | C18 | 375.139 | 13.514 | C01814 | HMDB0014645 | 3559 | | 0.88 | 0.763 | 0.95 | 0.916 | 1.21 | 0.538 | 0.79 | 0.921275 |
| 670 | Chlorzoxazone | [M+Na]+1 | POS | Mass, Spectral match | Hilic | 376.138 | 2.409 | C07931 | HMDB0014500 | 2733 | Metabolic pathways;Biosynthesis of secondary metabolites;Other | 1.20 | 0.266 | 1.88 | 0.042 | 1.36 | 0.138 | 1.39 | 0.276567 |
| 671 | Riboflavin | [M-H]-1 | NEG | Mass, RT, Spectral match | C18 | 376.138 | 4.934 | C00255 | HMDB0000244 | 493570 | Metabolic pathways;Biosynthesis of secondary metabolites;Other | 1.15 | 0.641 | 1.26 | 0.426 | 0.87 | 0.594 | 1.45 | 0.293263 |
| 672 | Sucrose | [M+Cl]-1 | POS | Mass, Spectral match | Hilic | 377.088 | 0.923 | C00089 | HMDB0000258 | 5988 | | 1.23 | 0.567 | 1.04 | 0.904 | 0.87 | 0.721 | 1.19 | 0.378015 |
| 673 | Sucrose | [M-H]-1 | NEG | Mass, Spectral match | C18 | 377.089 | 12.054 | C00089 | HMDB0000258 | 5988 | | 0.81 | 0.785 | 0.52 | 0.440 | 1.11 | 0.973 | 0.47 | 0.794694 |
| 674 | Coniferin | [M+H]+1 | POS | Mass, Spectral match | Hilic | 377.102 | 10.221 | C00761 | HMDB0013682 | 5280372 | | 1.37 | 0.354 | 0.88 | 0.752 | 2.00 | 0.023 | 0.44 | 0.047504 |
| 675 | Gibberellin_A3 | [M-H]-1 | NEG | Mass, Spectral match | Hilic | 381.109 | 9.339 | C01699 | HMDB0003559 | 9819600 | | 1.09 | 0.855 | 0.69 | 0.286 | 1.63 | 0.327 | 0.42 | 0.450741 |
| 676 | Sphinganine_1_phosphate | [M+H]+1 | POS | Mass, Spectral match | C18 | 382.271 | 11.446 | C01120 | HMDB0001383 | 644260 | | 0.92 | 0.317 | 0.89 | 0.267 | 0.89 | 0.793 | 1.00 | 0.993631 |
| 677 | S_Adenosylhomocysteine | [M+H]+1 | POS | Mass, Spectral match | Hilic | 384.121 | 8.202 | C00021 | HMDB0000939 | 439155 | Metabolic pathways;Biosynthesis of amino acids;Cysteine and methionine metabolism | 2.21 | 0.265 | 2.34 | 0.203 | 3.05 | 0.115 | 0.77 | 0.556684 |
| 678 | Cholestenone | [M+H]+1 | POS | Mass, Spectral match | C18 | 384.338 | 15.807 | C00599 | HMDB0000921 | 91477 | Metabolic pathways;Biosynthesis of antibiotics;Microbial metabolism in diverse environments;Other | 0.87 | 0.468 | 0.85 | 0.472 | 0.93 | 0.561 | 0.92 | 0.991234 |
| 679 | S_Inosyl_L_homocysteine | [M+H]+1 | POS | Mass, Spectral match | Hilic | 385.105 | 9.488 | C03539 | METPA0405 | NA | Biosynthesis of antibiotics;Other | 1.53 | 0.295 | 1.26 | 0.479 | 0.38 | 0.098 | 3.31 | 0.044564 |
| 680 | Vomilenine | [M+H]+1 | POS | Mass, Spectral match | Hilic | 385.128 | 1.280 | C01761 | #N/A | 11953806 | Biosynthesis of antibiotics;Other | 1.89 | 0.150 | 0.83 | 0.380 | 0.74 | 0.724 | 1.13 | 0.21719 |
| 681 | S_Adenosylhomocysteine | [M+H]+1 | POS | Mass, RT, Spectral match | C18 | 385.128 | 5.175 | C00021 | HMDB0000939 | 439155 | Biosynthesis of antibiotics;Other | 1.06 | 0.911 | 0.92 | 0.841 | 1.58 | 0.284 | 0.58 | 0.560932 |
| 682 | S_Adenosylmethioninamine | [M-H]-1 | NEG | Mass, Spectral match | Hilic | 390.126 | 9.817 | C01137 | HMDB0000988 | 439415 | | 0.82 | 0.680 | 1.22 | 0.536 | 1.24 | 0.699 | 0.99 | 0.705012 |
| 683 | Bis2_ethylhexylphthalate | [M+H]+1 | POS | Mass, Spectral match | Hilic | 390.276 | 1.141 | C03343 | HMDB0013248 | 20393 | | 0.74 | 0.012 | 0.79 | 0.104 | 1.16 | 0.194 | 0.67 | 0.093926 |
| 684 | Monoethylhexyl_phthalic_acid | [M+H]+1 | POS | Mass, Spectral match | C18 | 390.276 | 14.441 | C03343 | HMDB0013248 | 20393 | | 0.87 | 0.427 | 0.97 | 0.854 | 0.83 | 0.725 | 1.17 | 0.429051 |
| 685 | Monoethylhexyl_phthalic_acid | [M+H]+1 | POS | Mass, Spectral match | Hilic | 390.276 | 28.655 | C03343 | HMDB0013248 | 20393 | | 0.98 | 0.739 | 1.13 | 0.055 | 1.15 | 0.924 | 0.98 | 0.810243 |

| | | | | | | | | | | | | | | | | | | | |
|---|---|---|---|---|---|---|---|---|---|---|---|---|---|---|---|---|---|---|---|
| 686 | Sulindac | [M-H]-1 | NEG | Mass, Spectral match | Hilic | 391.056 | 9.302 | C01531 | HMDB0014743 | 1548887 | | 0.76 | 0.499 | 0.96 | 0.932 | 0.69 | 0.460 | 1.39 | 0.919703 |
| 687 | S_Adenosyl methionine | [M+H]+1 | POS | Mass, Spectral match | C18 | 398.141 | 9.380 | C00019 | HMDB0001185 | 16757548 | | 0.84 | 0.287 | 1.25 | 0.253 | 1.16 | 0.352 | 1.08 | 0.767733 |
| 688 | Xanthylic_acid | [M+H]+1 | POS | Mass, Spectral match | C18 | 399.012 | 1.333 | C00655 | HMDB0001554 | 73323 | | 0.68 | 0.326 | 1.41 | 0.216 | 0.69 | 0.530 | 2.04 | 0.014336 |
| 689 | Colchicine | [M+H]+1 | POS | Mass, RT, Spectral match | C18 | 399.167 | 6.980 | C07592 | HMDB0015466 | 2833 | Biosynthesis of secondary metabolites;Biosynthesis of alkaloids derived from shikimate pathway;Other | 1.05 | 0.042 | 1.05 | 0.076 | 1.03 | 0.429 | 1.02 | 0.374309 |
| 690 | Calcidiol | [M+H]+1 | POS | Mass, RT, Spectral match | C18 | 400.333 | 14.119 | C01561 | HMDB0003550 | 5283731 | Metabolic pathways;Other | 1.75 | 0.380 | 2.19 | 0.125 | 6.01 | 0.349 | 0.36 | 0.87193 |
| 691 | Guanosine_monophosphate | [M-H]-1 | NEG | Mass, RT, Spectral match | C18 | 402.018 | 2.596 | C00144 | HMDB0001397 | 6804 | | 5.22 | 0.255 | 2.00 | 0.372 | 1.99 | 0.308 | 1.00 | 0.836619 |
| 692 | Isoquinoline_alkaloids | [M-H]-1 | NEG | Mass, Spectral match | Hilic | 404.128 | 10.482 | C02134 | #N/A | 98570 | | 1.09 | 0.853 | 0.53 | 0.194 | 0.55 | 0.720 | 0.96 | 0.948652 |
| 693 | 3alpha_12alpha_Dihydroxy_7_oxo_5beta_cholanate | [M-H]-1 | NEG | Mass, RT, Spectral match | C18 | 406.272 | 8.438 | C01292 | #N/A | 439472 | | 4.18 | 0.010 | 30.13 | 0.223 | 120.18 | 0.245 | 0.25 | 0.276345 |
| 694 | Syringin | [M-H]-1 | NEG | Mass, Spectral match | Hilic | 407.109 | 9.699 | C01533 | METPA1704 | NA | | 0.82 | 0.103 | 0.78 | 0.114 | 0.65 | 0.672 | 1.20 | 0.376115 |
| 553 | Oleic_acid | [M-H]-1 | NEG | Mass, Spectral match | C18 | 282.255 | 13.675 | C00712 | HMDB0000207 | 445639 | Biosynthesis of plant secondary metabolites;Other | 1.16 | 0.540 | 1.64 | 0.118 | 1.77 | 0.385 | 0.93 | 0.716235 |
| 638 | Lactulose | [M-H]-1 | NEG | Mass, RT, Spectral match | C18 significant | 342.117 | 0.500 | C07064 | HMDB0000740 | 11333 | Metabolic pathways;Biosynthesis of secondary metabolites;ABC transporters;Other | 1.15 | 0.690 | 1.88 | 0.003 | 1.76 | 0.035 | 1.06 | 0.595512 |
| 697 | Riboflavin | [M-H]-1 | NEG | Mass,RT, Spectral match | Hilic | 411.107 | 10.146 | C00255 | HMDB0000244 | 493570 | | 0.92 | 0.752 | 0.59 | 0.115 | 0.55 | 0.286 | 1.08 | 0.756789 |
| 698 | Riboflavin_reduced | [M-H]-1 | NEG | Mass,RT, Spectral match | Hilic | 413.123 | 6.890 | C01007 | HMDB0001557 | 22833571 | | 0.95 | 0.911 | 0.67 | 0.365 | 0.85 | 0.258 | 0.79 | 0.571075 |
| 699 | Miconazole | [M+H]+1 | POS | Mass,RT, Spectral match | Hilic | 413.985 | 29.042 | C08070 | HMDB0015242 | 4189 | | 1.08 | 0.348 | 1.07 | 0.252 | 1.18 | 0.286 | 0.91 | 0.480867 |
| 700 | Eplerenone | [M+Na]+1 | NEG | Mass, RT, Spectral match | C18 | 414.203 | 9.823 | C12512 | HMDB0014838 | 443872 | | 1.05 | 0.498 | 0.98 | 0.714 | 1.04 | 0.293 | 0.94 | 0.576474 |
| 701 | 1_O_Sinapoyl_beta_D_glucose | [M+Na]+1 | POS | Mass, Spectral match | Hilic | 421.089 | 9.764 | C01175 | METPA1696 | NA | | 1.10 | 0.559 | 0.87 | 0.473 | 1.05 | 0.152 | 0.83 | 0.475822 |
| 702 | Blasticidin_S | [M+H]+1 | POS | Mass, Spectral match | Hilic | 421.191 | 1.606 | C02010 | HMDB0030452 | 258 | | 1.00 | 0.991 | 1.23 | 0.503 | 1.01 | 0.694 | 1.21 | 0.568505 |
| 703 | Blasticidin_S | [M+H]+1 | NEG | Mass, RT, Spectral match | C18 | 421.192 | 3.591 | C02010 | HMDB0030452 | 258 | | 0.86 | 0.264 | 0.92 | 0.515 | 0.81 | 0.846 | 1.14 | 0.263928 |
| 704 | Secologanin | [M-H]-1 | NEG | Mass, Spectral match | Hilic | 423.102 | 1.240 | C01852 | METPA0226 | NA | | 2.38 | 0.035 | 3.41 | 0.019 | 4.94 | 0.025 | 0.69 | 0.670501 |

| | | | | | | | | | | | | | | | | | | | |
|---|---|---|---|---|---|---|---|---|---|---|---|---|---|---|---|---|---|---|---|
| 705 | S_glutathionyl_L_cysteine | [M-H]-1 | NEG | Mass, Spectral match | Hilic | 426.087 | 21.516 | C05526 | METPA0607 | NA | Cysteine and methionine metabolism | 1.17 | 0.401 | 1.16 | 0.580 | 1.38 | 0.132 | 0.84 | 0.452693 |
| 706 | Vicianin | [M-H]-1 | NEG | Mass, Spectral match | Hilic | 426.137 | 11.370 | C01870 | #N/A | 656493 | | 0.84 | 0.662 | 1.26 | 0.311 | 1.42 | 0.099 | 0.89 | 0.571252 |
| 696 | Cholic_acid | [M-H]-1 | NEG | Mass, RT, Spectral match | C18 significant | 408.288 | 9.020 | C00695 | HMDB0000619 | 221493 | Metabolic pathways;Bile secretion;Secondary bile acid biosynthesis;Other | 2.30 | 0.057 | 9.22 | 0.007 | 18.22 | 0.036 | 0.51 | 0.299499 |
| 708 | Secologanin | [M+H]+1 | NEG | Mass, Spectral match | C18 | 427.103 | 9.034 | C01852 | METPA0226 | NA | | 0.61 | 0.330 | 0.75 | 0.568 | 0.50 | 0.213 | 1.50 | 0.41427 |
| 709 | 3_Phenylpropyl_2_methylpropanoate | [M-H]-1 | NEG | Mass, RT, Spectral match | C18 | 432.191 | 9.803 | C02008 | HMDB0034472 | 7662 | | 0.71 | 0.130 | 0.75 | 0.192 | 0.79 | 0.850 | 0.95 | 0.319879 |
| 710 | Lithocholic_acid_glycine_conjugate | [M+H]+1 | NEG | Mass, RT, Spectral match | C18 | 433.318 | 11.009 | C15557 | HMDB0000698 | 115245 | | 0.80 | 0.286 | 1.45 | 0.346 | 7.29 | 0.381 | 0.20 | 0.6893 |
| 711 | 35_Diiodo_L_tyrosine | [M-H]-1 | NEG | Mass, Spectral match | C18 | 433.873 | 18.195 | C01060 | HMDB0003474 | 9305 | | 0.71 | 0.482 | 0.58 | 0.252 | 0.61 | 0.381 | 0.95 | 0.935603 |
| 712 | Flurandrenolide | [M+H]+1 | NEG | Mass, Spectral match | C18 | 436.227 | 5.522 | D00328 | #N/A | 15209 | | 0.76 | 0.051 | 0.76 | 0.038 | 0.77 | 0.938 | 1.00 | 0.963393 |
| 713 | Flurandrenolide | [M+H]+1 | POS | Mass, Spectral match | Hilic | 436.227 | 1.139 | | HMDB0014984 | 15209 | | 0.86 | 0.086 | 0.87 | 0.049 | 0.92 | 0.656 | 0.95 | 0.914068 |
| 714 | Campesterol | [M+H]+1 | NEG | Mass, Spectral match | C18 | 439.329 | 10.379 | C01789 | HMDB0002869 | 5283637 | | 1.44 | 0.358 | 0.85 | 0.689 | 1.57 | 0.344 | 0.54 | 0.280697 |
| 715 | Tetrahydrofolic_acid | [M-H]-1 | NEG | Mass, Spectral match | Hilic | 444.160 | 3.256 | C00101 | HMDB0001846 | 91443 | | 12.92 | 0.167 | 1.22 | 0.813 | 2.49 | 0.382 | 0.49 | 0.533028 |
| 716 | beta_estradiol_17beta_glucuronide | [M-H]-1 | NEG | Mass, Spectral match | C18 significant | 448.210 | 7.160 | NA | HMDB0010317 | 66424 | Bile secretion;Other | 1.29 | 0.000 | 1.33 | 0.000 | 1.41 | 0.008 | 0.94 | 0.370553 |
| 717 | Fluocinolone acetonide | [M+H]+1 | POS | Mass,RT, Spectral match | Hilic | 452.201 | 1.196 | NA | HMDB0014729 | 6215 | | 0.78 | 0.198 | 0.56 | 0.014 | 0.87 | 0.477 | 0.64 | 0.053677 |
| 718 | Calcitriol | [M-H]-1 | NEG | Mass, RT, Spectral match | C18 | 455.296 | 8.257 | C01673 | HMDB0001903 | 5280453 | | 1.13 | 0.812 | 2.96 | 0.019 | 1.97 | 0.232 | 1.50 | 0.247025 |
| 719 | FMNH2 | [M-H]-1 | NEG | Mass,RT, Spectral match | Hilic | 457.112 | 9.765 | C01847 | HMDB0001142 | 445395 | | 0.45 | 0.001 | 0.42 | 0.004 | 0.27 | 0.081 | 1.54 | 0.371927 |
| 720 | FMN | [M+H]+1 | POS | Mass,RT, Spectral match | Hilic | 458.119 | 12.655 | C00061 | HMDB0001520 | 643976 | Metabolic pathways;Other | 0.55 | 0.155 | 0.96 | 0.920 | 0.98 | 0.323 | 0.98 | 0.91058 |
| 721 | Chitobiose | [M+H]+1 | POS | Mass,RT, Spectral match | Hilic | 459.135 | 10.235 | C01674 | HMDB0003556 | 22833609 | | 1.70 | 0.340 | 1.75 | 0.353 | 2.46 | 0.070 | 0.71 | 0.380088 |
| 722 | Ecdysone | [M-H]-1 | NEG | Mass, Spectral match | C18 | 463.306 | 11.754 | C00477 | METPA1782 | NA | | 1.02 | 0.944 | 2.28 | 0.050 | 4.96 | 0.350 | 0.46 | 0.810781 |
| 723 | Glycocholic_acid_ | [M-H]-1 | NEG | Mass, Spectral match | C18 significant | 465.310 | 8.320 | C01921 | HMDB0000138 | 23617285 | Metabolic pathways;Bile secretion;Secondary bile acid biosynthesis;Other | 10.16 | 0.016 | 11.03 | 0.028 | 12.52 | 0.125 | 0.88 | 0.906554 |
| 724 | 3a7a12a_Trihydroxy_5b_cholestan_26_al | [M+H]+1 | NEG | Mass, RT, Spectral match | C18 | 469.308 | 11.490 | C01301 | HMDB0003533 | 439479 | | 0.94 | 0.359 | 0.99 | 0.888 | 0.93 | 0.649 | 1.07 | 0.426214 |

| | | | | | | | | | | | | | | | | | | | |
|---|---|---|---|---|---|---|---|---|---|---|---|---|---|---|---|---|---|---|---|
| 725 | Kanamycin | [M+H]+1 | POS | Mass, Spectral match | Hilic | 483.229 | 1.085 | C01822 | HMDB0015303 | 6032 | | 0.91 | 0.536 | 1.04 | 0.822 | 0.86 | 0.967 | 1.21 | 0.165335 |
| 726 | Methotrexate | [M+H]+1 | POS | Mass, Spectral match | Hilic | 489.137 | 1.076 | C01937 | HMDB0014703 | 126941 | | 1.04 | 0.408 | 0.96 | 0.257 | 0.99 | 0.338 | 0.97 | 0.447058 |
| 727 | Imatinib | [M+H]+1 | POS | Mass, Spectral match | Hilic | 493.261 | 1.119 | | HMDB0014757 | 5291 | | 1.02 | 0.843 | 0.99 | 0.923 | 0.91 | 0.863 | 1.09 | 0.635323 |
| 728 | 1_Palmitoylglycerophosphocholine | [M+K]+1 | POS | Mass, Spectral match | Hilic | 495.331 | 1.626 | C04517 | METPA0506 | NA | | 1.15 | 0.229 | 1.04 | 0.742 | 1.22 | 0.260 | 0.86 | 0.534697 |
| 729 | 510_Methylene_THF | [M-H]-1 | NEG | Mass, Spectral match | C18 | 496.137 | 4.935 | C00143 | HMDB0001533 | 439175 | | 1.17 | 0.626 | 1.34 | 0.295 | 0.90 | 0.554 | 1.49 | 0.233629 |
| 730 | Taurochenodesoxycholic_acid | [M-H]-1 | NEG | Mass, Spectral match | C18 | 499.297 | 8.524 | C05465 | HMDB0000951 | 387316 | Bile secretion;Secondary bile acid biosynthesis;Other | 4.58 | 0.201 | 7.18 | 0.084 | 0.96 | 0.922 | 7.49 | 0.107257 |
| 731 | Taurodeoxycholic_acid | [M-H]-1 | NEG | Mass, Spectral match | C18 significant | 499.297 | 7.620 | C05463 | HMDB0000896 | 2733768 | Bile secretion;Secondary bile acid biosynthesis;Other | 8.08 | 0.010 | 25.44 | 0.078 | 4.74 | 0.478 | 5.37 | 0.186532 |
| 732 | D_Raffinose | [M-H]-1 | NEG | Mass, Spectral match | Hilic | 504.169 | 11.360 | C00492 | HMDB0003213 | 10542 | ABC transporters;Other | 0.96 | 0.905 | 1.73 | 0.068 | 1.83 | 0.018 | 0.95 | 0.649242 |
| 733 | Taurocholic_acid | [M-H]-1 | NEG | Mass, Spectral match | C18 | 515.291 | 7.000 | C05122 | HMDB0000036 | 440567 | Metabolic pathways;Bile secretion;Secondary bile acid biosynthesis;Other | 12.51 | 0.066 | 13.03 | 0.049 | 6.68 | 0.125 | 1.95 | 0.487385 |
| 734 | Kanamycin | [M+H]+1 | NEG | Mass, Spectral match | C18 | 519.203 | 7.161 | C01822 | HMDB0015303 | 6032 | | 0.97 | 0.488 | 0.96 | 0.304 | 0.93 | 0.151 | 1.03 | 0.408199 |
| 735 | 1_Linoleoylglycerophosphocholine | [M+Na]+1 | POS | Mass, Spectral match | Hilic | 519.332 | 1.427 | C04230 | HMDB0010386 | 11005824 | | 0.76 | 0.265 | 0.78 | 0.370 | 0.78 | 0.697 | 1.00 | 0.96227 |
| 736 | L_Cysteinylglycine | [M+Na]+1 | POS | Mass, Spectral match | Hilic | 521.112 | 8.000 | C01315 | #N/A | 439484 | | 1.68 | 0.354 | 1.13 | 0.829 | 1.08 | 0.856 | 1.05 | 0.735432 |
| 737 | LysoPC18:19Z | [M+Na]+1 | NEG | Mass, Spectral match | C18 | 521.347 | 11.418 | C04230 | HMDB0002815 | 16081932 | | 0.93 | 0.720 | 1.02 | 0.932 | 1.37 | 0.343 | 0.74 | 0.546349 |
| 738 | UDP_D_Xylose | [M-H]-1 | NEG | Mass, Spectral match | C18 | 537.049 | 10.093 | C00190 | HMDB0001018 | 439179 | | 0.90 | 0.868 | 1.20 | 0.832 | 1.10 | 0.981 | 1.09 | 0.45094 |
| 739 | NN_Dimethylsphingosine | [M+H]+1 | NEG | Mass, RT, Spectral match | C18 | 537.511 | 16.236 | C13914 | HMDB0013645 | 5282309 | | 1.54 | 0.570 | 2.45 | 0.107 | 1.12 | 0.461 | 2.18 | 0.142123 |
| 740 | Aliskiren | [M+H]+1 | NEG | Mass, Spectral match | C18 | 551.394 | 13.266 | D03208 | #N/A | 5493444 | | 0.90 | 0.834 | 0.54 | 0.273 | 0.46 | 0.662 | 1.16 | 0.658311 |
| 741 | Adenosine_diphosphate_ribose | [M+H]+1 | NEG | Mass, Spectral match | C18 | 559.071 | 12.848 | C00301 | HMDB0001178 | 192 | Metabolic pathways;Biosynthesis of secondary metabolites;Biosynthesis of amino acids;Other | 1.36 | 0.670 | 0.85 | 0.835 | 0.99 | 0.856 | 0.85 | 0.567556 |
| 742 | CDP_glucose | [M+H]+1 | POS | Mass, Spectral match | Hilic | 564.063 | 9.270 | C00501 | HMDB0003369 | 439244 | | 1.00 | 0.996 | 1.31 | 0.455 | 0.95 | 0.766 | 1.38 | 0.74799 |
| 743 | CDP_glucose | [M-H]-1 | NEG | Mass, RT, Spectral match | C18 | 564.068 | 10.092 | C00501 | HMDB0003369 | 439244 | | 0.80 | 0.721 | 1.02 | 0.978 | 0.95 | 0.830 | 1.07 | 0.484282 |
| 744 | Daunorubicin | [M-H]-1 | NEG | Mass, RT, Spectral match | C18 | 566.146 | 13.062 | C01907 | HMDB0014832 | 30323 | | 0.69 | 0.358 | 1.49 | 0.366 | 0.65 | 0.571 | 2.29 | 0.221589 |

| | | | | | | | | | | | | | | | | | | | |
|---|---|---|---|---|---|---|---|---|---|---|---|---|---|---|---|---|---|---|---|
| 745 | B_Carotene | [M+H]+1 | NEG | Mass, RT, Spectral match | C18 | 575.403 | 11.481 | C02094 | HMDB0000561 | 5280489 | | 0.93 | 0.655 | 0.98 | 0.928 | 0.92 | 0.592 | 1.07 | 0.767855 |
| 746 | Doxorubicin | [M+H]+1 | POS | Mass, Spectral match | C18 | 582.140 | 13.442 | C01661 | HMDB0015132 | 31703 | | 1.02 | 0.973 | 1.41 | 0.614 | 0.90 | 0.698 | 1.56 | 0.388878 |
| 747 | All_trans_hexaprenyl_diphosphate | [M+H]+1 | POS | Mass, Spectral match | C18 | 585.308 | 7.979 | C01230 | HMDB0012188 | 5280413 | | 42.49 | 0.345 | 48.71 | 0.298 | 78.91 | 0.126 | 0.62 | 0.347465 |
| 748 | Oxidized_glutathione | [M-H]-1 | NEG | Mass, Spectral match | Hilic | 611.150 | 9.394 | C00127 | HMDB0003337 | 975 | | 0.62 | 0.156 | 1.12 | 0.753 | 1.59 | 0.367 | 0.70 | 0.972107 |
| 749 | Oxidized_glutathione | [M+H]+1 | POS | Mass, Spectral match | C18 | 612.151 | 0.532 | C00127 | HMDB0003337 | 975 | Other | 0.96 | 0.762 | 1.01 | 0.958 | 1.25 | 0.084 | 0.81 | 0.372522 |
| 750 | Heme | [M+H]+1 | POS | Mass, Spectral match | C18 | 615.168 | 9.306 | C00032 | HMDB0003178 | NA | | 0.27 | 0.218 | 1.19 | 0.842 | 0.18 | 0.071 | 6.65 | 0.34184 |
| 751 | Paromomycin | [M+H]+1 | POS | Mass, RT, Spectral match | C18 | 616.303 | 13.191 | C00832 | HMDB0015490 | 165580 | | 0.68 | 0.224 | 1.44 | 0.342 | 0.93 | 0.905 | 1.55 | 0.335 |
| 752 | Ouabain | [M+H]+1 | POS | Mass, Spectral match | C18 | 619.254 | 14.970 | C01443 | HMDB0015224 | 439501 | | 0.80 | 0.682 | 1.01 | 0.981 | 0.87 | 0.710 | 1.16 | 0.380753 |
| 753 | Chlorophyllide | [M+H]+1 | POS | Mass, Spectral match | Hilic | 649.212 | 12.416 | C02139 | METPA0252 | NA | | 0.90 | 0.856 | 1.70 | 0.295 | 2.93 | 0.007 | 0.58 | 0.134596 |
| 754 | Stachyose | [M-H]-1 | NEG | Mass, Spectral match | Hilic significant | 666.222 | 12.435 | C01613 | HMDB0003553 | 439531 | Other | 0.89 | 0.824 | 1.80 | 0.172 | 2.51 | 0.012 | 0.71 | 0.268142 |
| 755 | Maltotetraose | [M+H]+1 | POS | Mass, Spectral match | Hilic | 688.202 | 12.412 | C02052 | HMDB0001296 | 439639 | | 0.91 | 0.831 | 1.93 | 0.076 | 2.04 | 0.007 | 0.94 | 0.645041 |
| 756 | Erythromycin | [M+Na]+1 | POS | Mass, Spectral match | C18 | 734.468 | 7.060 | C01912 | HMDB0014344 | 12560 | | 1.01 | 0.986 | 1.20 | 0.758 | 0.96 | 0.167 | 1.24 | 0.816687 |
| 707 | Leupeptin | [M-H]-1 | NEG | Mass, Spectral match | C18 | 426.295 | 5.937 | C01591 | #N/A | 5284408 | | 0.96 | 0.879 | 1.19 | 0.570 | 0.99 | 0.177 | 1.21 | 0.717418 |
| 758 | FAD | [M+H]+1 | POS | Mass,RT, Spectral match | Hilic | 784.155 | 10.131 | C00016 | HMDB0001248 | 643975 | | 2.00 | 0.335 | 0.89 | 0.828 | 0.64 | 0.665 | 1.39 | 0.631976 |
| 759 | FAD | [M-H]-1 | NEG | Mass, RT, Spectral match | C18 | 785.157 | 2.451 | C00016 | HMDB0001248 | 643975 | Metabolic pathways;Biosynthesis of secondary metabolites;Other | 1.77 | 0.288 | 1.32 | 0.460 | 1.22 | 0.507 | 1.09 | 0.871501 |
| 760 | Thyroxine | [M+H]+1 | POS | Mass, Spectral match | Hilic | 811.652 | 1.592 | C01829 | HMDB0000248 | 5819 | | 1.03 | 0.947 | 0.90 | 0.772 | 2.61 | 0.115 | 0.34 | 0.148522 |
| 761 | Thyroxine | [M+H]+1 | POS | Mass, Spectral match | C18 | 811.652 | 16.810 | C01829 | HMDB0000248 | 5819 | | 0.61 | 0.287 | 0.73 | 0.477 | 1.62 | 0.489 | 0.45 | 0.260069 |
| 762 | Propionyl_CoA | [M+H]+1 | POS | Mass, Spectral match | Hilic | 822.129 | 12.439 | C00100 | HMDB0001275 | 439164 | | 6.90 | 0.170 | 26.48 | 0.040 | 11.09 | 0.113 | 2.39 | 0.528358 |
| 763 | Coenzyme_Q9 | [M+H]+1 | POS | Mass, Spectral match | Hilic | 829.583 | 1.846 | C01967 | HMDB0006707 | 5280473 | | 1.98 | 0.181 | 1.30 | 0.561 | 1.88 | 0.065 | 0.69 | 0.268598 |
| 764 | Maltopentaose | [M+H]+1 | POS | Mass, Spectral match | Hilic | 850.255 | 13.338 | | HMDB0012254 | 13489094 | | 0.88 | 0.761 | 2.02 | 0.044 | 2.36 | 0.029 | 0.85 | 0.712012 |
| 765 | Cinnamoyl_CoA | [M+H]+1 | POS | Mass, Spectral | Hilic | 896.155 | 8.024 | C00540 | METPA0055 | NA | | 2.80 | 0.311 | 1.42 | 0.688 | 1.82 | 0.664 | 0.78 | 0.515312 |

| | | | | | | | | | | | | | | | | | | | |
|---|---|---|---|---|---|---|---|---|---|---|---|---|---|---|---|---|---|---|---|
| | | | | match | | | | | | | | | | | | | | | |
| 766 | 3_Hydroxy_3_methylglutaryl_CoA | [M+H]+1 | POS | Mass, Spectral match | Hilic | 946.120 | 8.137 | C00356 | HMDB0001375 | 439218 | | 1.16 | 0.772 | 0.43 | 0.051 | 0.56 | 0.714 | 0.78 | 0.896709 |
| 767 | Cyclodextrin_HI | [M+H]+1 | POS | Mass, Spectral match | Hilic | 972.315 | 13.841 | | HMDB0029927 | 86575 | | 1.52 | 0.425 | 2.99 | 0.004 | 2.87 | 0.075 | 1.04 | 0.68402 |
| 768 | Bradykinin_C18 | [M+H]+1 | POS | Mass, Spectral match | C18 | 1058.562 | 10.792 | C00306 | HMDB0004246 | 439201 | | 0.64 | 0.114 | 0.65 | 0.103 | 0.54 | 0.351 | 1.20 | 0.614265 |

Supplementary Table 6: Blood transcription module (BTM) enrichment analysis

| Gene | Sr no | ID | Module title | UP ASA vs CCl4 | Down ASA vs CCl4 | Module size | UP %age | Down %age |
|---|---|---|---|---|---|---|---|---|
| S100a8 | 1 | M37.0 | immune activation - generic cluster | 16 | 18 | 347 | 4.6 | 5.2 |
| Hk3 | | | | | | | | |
| Cyfip1 | | | | | | | | |
| Dtymk | | | | | | | | |
| Ranbp1 | | | | | | | | |
| Myb | | | | | | | | |
| Dock5 | | | | | | | | |
| Lin7a | | | | | | | | |
| Ltf | | | | | | | | |
| Cd9 | | | | | | | | |
| Tm6sf1 | | | | | | | | |
| Lgals3 | | | | | | | | |
| Cse1l | | | | | | | | |
| Iqgap3 | | | | | | | | |
| Gsn | | | | | | | | |
| Ogn | | | | | | | | |
| Adap2 | | | | | | | | |
| Rab32 | | | | | | | | |
| Cda | | | | | | | | |
| Smpdl3a | | | | | | | | |
| Ptk2 | | | | | | | | |
| Eng | | | | | | | | |
| Jup | | | | | | | | |
| Ptgs2 | | | | | | | | |
| Lrrk2 | | | | | | | | |
| Steap4 | | | | | | | | |
| Hp | | | | | | | | |
| Lcn2 | | | | | | | | |
| Ctss | | | | | | | | |
| Tmem176b | | | | | | | | |
| Il18rap | | | | | | | | |
| Pid1 | | | | | | | | |
| Mcm7 | | | | | | | | |
| Lrg1 | | | | | | | | |

| | | | | | | | | |
|---|---|---|---|---|---|---|---|---|
| | 2 | M4.0 | cell cycle and transcription | 9 | 16 | 335 | 2.69 | 4.78 |
| Cst3 | | | | | | | | |
| Kntc1 | | | | | | | | |
| Cd163 | | | | | | | | |
| S100a9 | | | | | | | | |
| Kif20b | | | | | | | | |
| Lmnb1 | | | | | | | | |
| Cdca8 | | | | | | | | |
| E2f7 | | | | | | | | |
| Wdfy3 | | | | | | | | |
| Plxnb2 | | | | | | | | |
| Hmox1 | | | | | | | | |
| Tymp | | | | | | | | |
| Hnmt | | | | | | | | |
| Ctsh | | | | | | | | |
| Prim1 | | | | | | | | |
| Ifi30 | | | | | | | | |
| Mki67 | | | | | | | | |
| Rbm47 | | | | | | | | |
| Fcgrt | | | | | | | | |
| Cd36 | | | | | | | | |
| Cybb | | | | | | | | |
| Kctd12 | | | | | | | | |
| Alox5 | | | | | | | | |
| Marcks | | | | | | | | |
| Aspm | | | | | | | | |
| | 3 | M4.1 | cell cycle (I) | 6 | 3 | 145 | 4.14 | 2.07 |
| Kntc1 | | | | | | | | |
| Kif20b | | | | | | | | |
| Cdca8 | | | | | | | | |
| E2f7 | | | | | | | | |
| Dtymk | | | | | | | | |
| Ranbp1 | | | | | | | | |
| Prim1 | | | | | | | | |
| Mki67 | | | | | | | | |
| Aspm | | | | | | | | |
| M118.0 | 4 | M118.0 | enriched in monocytes (IV) | 2 | 7 | 17 | 11.76 | 41.18 |
| Dock5 | | | | | | | | |
| Lgals3 | | | | | | | | |

| | | | | | | | | |
|---|---|---|---|---|---|---|---|---|
| Steap4 | | | | | | | | |
| Ptgs2 | | | | | | | | |
| Lrrk2 | | | | | | | | |
| Ctss | | | | | | | | |
| Tmem176b | | | | | | | | |
| Pid1 | | | | | | | | |
| Mpp1 | | | | | | | | |
| M11.0 | 5 | M11.0 | enriched in monocytes (II) | 6 | 16 | 189 | 3.17 | 8.47 |
| Cst3 | | | | | | | | |
| Cd163 | | | | | | | | |
| S100a9 | | | | | | | | |
| S100a8 | | | | | | | | |
| Hk3 | | | | | | | | |
| Cyfip1 | | | | | | | | |
| Ctsh | | | | | | | | |
| Plxnb2 | | | | | | | | |
| Tymp | | | | | | | | |
| Hnmt | | | | | | | | |
| Ifi30 | | | | | | | | |
| Rbm47 | | | | | | | | |
| Fcgrt | | | | | | | | |
| Cd36 | | | | | | | | |
| Cybb | | | | | | | | |
| Kctd12 | | | | | | | | |
| Alox5 | | | | | | | | |
| Adap2 | | | | | | | | |
| Vnn3 | | | | | | | | |
| Rab32 | | | | | | | | |
| Cda | | | | | | | | |
| Smpdl3a | | | | | | | | |
| S5 | 6 | S5 | DC surface signature | 3 | 5 | 82 | 3.66 | 6.10 |
| Cd9 | | | | | | | | |
| Alcam | | | | | | | | |
| Gpnmb | | | | | | | | |
| Atp1b1 | | | | | | | | |
| Agpat3 | | | | | | | | |
| Slco2b1 | | | | | | | | |
| Abhd12 | | | | | | | | |
| Msr1 | | | | | | | | |

| | | | | | | | | |
|---|---|---|---|---|---|---|---|---|
| M37.1 | 7 | M37.1 | enriched in neutrophils (I) | 1.92 | 11.54 | 1 | 6 | 52 |
| Lin7a | | | | | | | | |
| Cda | | | | | | | | |
| Steap4 | | | | | | | | |
| Lrg1 | | | | | | | | |
| Vnn3 | | | | | | | | |
| Mme | | | | | | | | |
| Dgat2 | | | | | | | | |
| M5.0 | 8 | M5.0 | regulation of antigen presentation and immune response | 1 | 5 | 81 | 1.23 | 6.17 |
| Plcg1 | | | | | | | | |
| Ctsh | | | | | | | | |
| Chuk | | | | | | | | |
| Myd88 | | | | | | | | |
| Mapk14 | | | | | | | | |
| Ptprc | | | | | | | | |
| M127 | 9 | M127 | type I interferon response | 0 | 5 | 12 | 0.00 | 41.67 |
| Stat1 | | | | | | | | |
| Ifih1 | | | | | | | | |
| Parp9 | | | | | | | | |
| Tap1 | | | | | | | | |
| Irf7 | | | | | | | | |
| M32.1 | 10 | M32.0 | platelet activation (II) | 0 | 5 | 23 | 0.00 | 21.74 |
| Hgs | | | | | | | | |
| Akt1 | | | | | | | | |
| Gnai2 | | | | | | | | |
| Map2k2 | | | | | | | | |
| Pkn1 | | | | | | | | |

| | | | | | | | | |
|---|---|---|---|---|---|---|---|---|
| **M87** | **11** | M87 | **transmembrane transport (I)** | **2** | **5** | **24** | **8.33** | **20.83** |
| **Hk3** | | | | | | | | |
| **Slc25a21** | | | | | | | | |
| **Cd36** | | | | | | | | |
| **Hmox1** | | | | | | | | |
| **Abcg2** | | | | | | | | |
| **Aqp1** | | | | | | | | |
| **Abcb6** | | | | | | | | |
| **M106.0** | **12** | M106.0 | **nuclear pore complex** | **1** | **6** | **17** | **5.88** | **35.29** |
| **Ranbp1** | | | | | | | | |
| **Nup155** | | | | | | | | |
| **Xpo1** | | | | | | | | |
| **Nup35** | | | | | | | | |
| **Nup62** | | | | | | | | |
| **Nup133** | | | | | | | | |
| **Seh1l** | | | | | | | | |
| **M238** | **13** | M238 | **respiratory electron transport chain (mitochondrion)** | **1** | **5** | **17** | **5.88** | **29.41** |
| **Ndufb2** | | | | | | | | |
| **Ndufa4** | | | | | | | | |
| **Uqcrq** | | | | | | | | |
| **Ndufa11** | | | | | | | | |
| **Ndufs8** | | | | | | | | |
| **Ndufa6** | | | | | | | | |
| **M32.6** | **14** | M32.6 | **TBA** | **1** | **5** | **21** | **4.76** | **23.81** |
| **Cdc37** | | | | | | | | |
| **Hgs** | | | | | | | | |

| | | | | | | | | |
|---|---|---|---|---|---|---|---|---|
| Akt1 | | | | | | | | |
| Pkn1 | | | | | | | | |
| Cpne1 | | | | | | | | |
| Psmd3 | | | | | | | | |
| M185 | 15 | M185 | TBA | 1 | 6 | 14 | 7.14 | 42.86 |
| Synj1 | | | | | | | | |
| Ptpn11 | | | | | | | | |
| Fndc3a | | | | | | | | |
| Hprt1 | | | | | | | | |
| Nptn | | | | | | | | |
| Hdac2 | | | | | | | | |
| Rb1 | | | | | | | | |

| Supplementary Table 7. AUROC analysis | | | | | |
|---|---|---|---|---|---|
| | **Markers** | **AUC** | **pvalue** | **FC** | **clusters** |
| **Parameters** | ASTU/L | 1 | 1.27E-07 | 0.7734004 | 1 |
| | TBmg/dl | 1 | 1.686E-07 | 0.3737176 | 1 |
| | Albg/dl | 1 | 4.99E-07 | -0.406857 | 4 |
| | ALTU/L | 1 | 5.43E-07 | 0.6933634 | 1 |
| | Sirius_red | 0.96 | 0.0022053 | 0.671731 | 3 |
| | TGFbeta | 0.96 | 0.0014077 | 0.2563084 | 1 |
| **Validation** | Alox5 | 0.8 | 0.0585509 | 0.1907794 | 1 |
| | Kmo | 1 | 0.0003454 | 0.5491214 | 1 |
| | Arg1 | 1 | 0.0051017 | 0.5813376 | 1 |
| **Proteins** | Rundc1 | 0.9 | 0.0126755 | -0.216607 | 2 |
| | Ppp1r21 | 0.9 | 0.0126869 | 0.0426149 | 3 |
| | Chdh | 0.9 | 0.0168649 | 0.0471785 | 3 |
| | C01727 | 0.9 | 0.0173472 | -0.02092 | 4 |
| | Aifm1 | 0.9 | 0.0198707 | 0.0306008 | 1 |
| | Cyp3a25 | 0.9 | 0.0269685 | -0.079754 | 4 |
| | Fbln5 | 0.9 | 0.0365343 | -0.10426 | 2 |
| | Prps2 | 0.9 | 0.0421509 | 0.1249175 | 1 |
| | Ctrb1 | 0.9 | 0.0632334 | -0.132063 | 2 |
| | Magt1 | 0.92 | 0.0053409 | 0.2760374 | 3 |
| | Ptprd | 0.92 | 0.0055972 | -0.147137 | 2 |
| | Ubac2 | 0.92 | 0.0075021 | 0.2917633 | 1 |
| | Uap1 | 0.92 | 0.0082013 | 0.1221454 | 3 |
| | Psmc3 | 0.92 | 0.0094203 | 0.0243957 | 3 |
| | Lrg1 | 0.92 | 0.0101395 | 0.176595 | 3 |
| | Vps36 | 0.92 | 0.0103102 | 0.1252505 | 3 |
| | Epb41 | 0.92 | 0.0110549 | -0.184191 | 4 |
| **Metabolites** | C00576 | 0.92 | 0.0089975 | 0.1163232 | 3 |
| | C00322 | 0.88 | 0.0192577 | 0.215247 | 1 |
| | C00257a | 0.88 | 0.019452 | -0.054758 | 2 |
| | C00880 | 0.88 | 0.019452 | -0.054758 | 2 |
| | C00388 | 0.88 | 0.0256904 | -0.086562 | 2 |
| | Capn2 | 0.88 | 0.0258243 | -0.078421 | 2 |
| | Gnai3 | 0.88 | 0.025932 | 0.0978863 | 3 |
| | C04153 | 0.88 | 0.0262805 | 0.0370728 | 3 |
| | Mrpl11 | 0.88 | 0.0266667 | -0.117813 | 4 |
| | Lamtor2 | 0.88 | 0.0268749 | 0.2351532 | 3 |

| | | | | |
|---|---|---|---|---|
| Serping1 | 0.88 | 0.0269693 | 0.0395238 | 3 |
| Ap3b1 | 0.88 | 0.0271383 | 0.0575707 | 1 |
| Cyp27a1 | 0.88 | 0.0271738 | 0.0406184 | 3 |
| C01380 | 0.88 | 0.0272242 | -0.036914 | 4 |
| C00299 | 0.88 | 0.0283359 | 0.0395954 | 3 |
| C02067 | 0.88 | 0.0283359 | 0.0395954 | 3 |
| C02846 | 0.84 | 0.0858417 | -0.112753 | 4 |
| C01748 | 0.84 | 0.0884852 | 0.0828892 | 1 |
| C11095 | 0.84 | 0.0959276 | -0.131359 | 2 |
| C00909 | 0.84 | 0.0963517 | 0.1481367 | 1 |
| C00544a | 0.84 | 0.0975317 | -0.049431 | 2 |
| C14770 | 0.84 | 0.117328 | 0.0700761 | 1 |
| C00487 | 0.84 | 0.124024 | -0.217619 | 2 |
| C02037 | 0.84 | 0.1330552 | -0.26056 | 2 |
| C02139 | 0.84 | 0.1343708 | -0.081668 | 2 |
| C01829a | 0.84 | 0.1483727 | -0.120984 | 4 |
| C03680 | 0.84 | 0.2488262 | 0.0874263 | 1 |
| C05132 | 0.84 | 0.2488262 | 0.0874263 | 1 |
| C01938 | 0.84 | 0.2770973 | -0.071596 | 2 |
| Myb | 0.82 | 0.0401152 | -0.435922 | 4 |
| Ddost | 0.82 | 0.0446915 | 0.0401092 | 3 |
| Gripap1 | 0.82 | 0.0487229 | -0.1841 | 2 |
| C02455 | 0.82 | 0.049035 | -0.021835 | 2 |
| C02137 | 0.82 | 0.0497491 | 0.0240744 | 1 |
| Uqcrc1 | 0.82 | 0.0567394 | -0.022174 | 4 |
| Copg1 | 0.82 | 0.0642335 | 0.0308142 | 1 |
| C16185 | 0.82 | 0.0923032 | 0.2949958 | 3 |
| C01586 | 0.82 | 0.1780754 | -0.095314 | 2 |
| Lrrc29 | 0.8 | 0.0249957 | -0.27815 | 4 |
| Tollip | 0.8 | 0.0273222 | 0.0876796 | 3 |
| Frem1 | 0.8 | 0.0326115 | 0.2046837 | 3 |
| S100a8 | 0.8 | 0.035657 | -0.232874 | 4 |
| Cst3 | 0.8 | 0.0357779 | -0.166376 | 4 |
| Nipsnap2 | 0.8 | 0.0388428 | 0.0660442 | 3 |
| Ftl1 | 0.8 | 0.0428682 | -0.077019 | 2 |
| Eloc | 0.8 | 0.0470763 | 0.1694761 | 3 |
| Acot9 | 0.8 | 0.0483218 | 0.0930747 | 3 |
| C01304 | 0.8 | 0.0525851 | -0.293227 | 2 |
| Tbrg4 | 0.8 | 0.0526638 | 0.0487533 | 1 |

| | | | | |
|---|---|---|---|---|
| E130308A19Rik | 0.8 | 0.0557028 | -0.169216 | 2 |
| C10172 | 0.8 | 0.0580989 | -0.049462 | 2 |
| Pdcd6 | 0.8 | 0.0614403 | 0.0872686 | 3 |
| Scfd1 | 0.8 | 0.0626534 | 0.0391436 | 3 |
| Prdx5 | 0.8 | 0.063159 | 0.0151796 | 3 |
| Ces1e | 0.8 | 0.0638373 | 0.0493614 | 3 |
| Metap2 | 0.8 | 0.0640142 | -0.096735 | 4 |
| Trmt112 | 0.8 | 0.0642298 | -0.131285 | 4 |
| Idh3b | 0.8 | 0.0673748 | -0.036617 | 4 |
| Sult1b1 | 0.8 | 0.0698799 | -0.083209 | 4 |
| Stat2 | 0.8 | 0.0699757 | 0.1191923 | 3 |
| Sepsecs | 0.8 | 0.0701941 | 0.1148603 | 3 |
| Cbr4 | 0.8 | 0.0741966 | 0.0406751 | 3 |
| Gpnmb | 0.8 | 0.0748122 | -0.038185 | 4 |
| Nipsnap3b | 0.8 | 0.0755885 | 0.0214171 | 1 |
| C02029 | 0.8 | 0.076832 | -0.146888 | 2 |
| Mrpl55 | 0.8 | 0.0838798 | 0.1078265 | 1 |
| Tial1 | 0.8 | 0.0861593 | 0.1494908 | 3 |
| C00637 | 0.8 | 0.0877364 | -0.073167 | 4 |
| Mup3 | 0.8 | 0.0880359 | 0.0412902 | 3 |
| Cybb | 0.8 | 0.0911218 | 0.12986 | 3 |
| Tgm2 | 0.8 | 0.098086 | -0.025973 | 4 |
| Srrt | 0.8 | 0.1006874 | 0.136053 | 3 |
| C05465 | 0.8 | 0.1074001 | 0.1794914 | 3 |
| Gjb1 | 0.8 | 0.1085727 | -0.133812 | 2 |
| Hist1h1a | 0.8 | 0.1108326 | -0.261823 | 4 |
| Rb1 | 0.8 | 0.1117305 | 0.1817039 | 3 |
| Hectd3 | 0.8 | 0.11924 | 0.1237361 | 3 |
| Ppp2r2a | 0.8 | 0.1280451 | 0.086064 | 3 |
| Tbc1d9b | 0.8 | 0.1332282 | 0.1488189 | 3 |
| C13914 | 0.8 | 0.1410194 | 0.0889518 | 3 |
| Panx2 | 0.8 | 0.143356 | -0.091557 | 4 |
| C14748 | 0.8 | 0.1458268 | 0.0750813 | 3 |
| Nalcn | 0.8 | 0.147401 | -0.171054 | 2 |
| C00183a | 0.8 | 0.1576062 | -0.053418 | 2 |
| C00919 | 0.8 | 0.1747732 | -0.186011 | 2 |
| Ifi30 | 0.8 | 0.194617 | 0.0495526 | 3 |
| Soat2 | 0.8 | 0.2015116 | -0.148946 | 2 |
| C00526 | 0.8 | 0.202214 | 0.0633527 | 1 |

| | | | | |
|---|---|---|---|---|
| Chid1 | 0.8 | 0.2347977 | -0.115163 | 2 |
| Tbl2 | 0.8 | 0.2873241 | 0.0522051 | 3 |
| Pea15 | 0.8 | 0.4546828 | 0.0632306 | 3 |
| Anxa11 | 0.78 | 0.0463322 | 0.0341261 | 3 |
| C01275a | 0.78 | 0.049178 | 0.2323468 | 1 |
| C07354a | 0.78 | 0.646412 | -0.039258 | 4 |
| Gar1 | 0.76 | 0.0388656 | -0.201546 | 2 |
| Thop1 | 0.76 | 0.0493071 | 0.1302681 | 1 |
| C00993 | 0.76 | 0.0499664 | -0.183995 | 4 |
| Grpel2 | 0.76 | 0.0726966 | -0.212309 | 4 |
| Myh11 | 0.76 | 0.0807702 | -0.108873 | 4 |
| Hrg | 0.76 | 0.0814628 | -0.082511 | 4 |
| C04438 | 0.76 | 0.0893406 | -0.083722 | 2 |
| Jagn1 | 0.76 | 0.0980296 | -0.152724 | 4 |
| Sephs2 | 0.76 | 0.0993338 | -0.060221 | 2 |
| Sgk3 | 0.76 | 0.0993927 | -0.170754 | 4 |
| C00544 | 0.76 | 0.1015494 | -0.087615 | 2 |
| Acp2 | 0.76 | 0.115295 | 0.1946427 | 3 |
| C01617 | 0.76 | 0.1349733 | 0.1228759 | 1 |
| C00670 | 0.76 | 0.1633142 | -0.092395 | 2 |
| C00047 | 0.76 | 0.184329 | -0.201092 | 2 |
| C05131 | 0.76 | 0.202125 | 0.1808246 | 1 |
| C01449 | 0.76 | 0.2032033 | 0.1759413 | 1 |
| C01907 | 0.76 | 0.2216864 | 0.1110999 | 1 |
| C03793 | 0.76 | 0.2921246 | 0.0457189 | 3 |
| C16759 | 0.76 | 0.3144799 | 0.0439389 | 1 |
| Apoe | 0.76 | 0.366456 | 0.1130329 | 1 |
| C00376 | 0.76 | 0.4658455 | 0.0406791 | 3 |
| C03079 | 0.74 | 0.3328836 | -0.098191 | 2 |
| Ercc2 | 0.72 | 0.0694804 | -0.258621 | 4 |
| Arntl2 | 0.72 | 0.0820476 | -0.196512 | 4 |
| C00796 | 0.72 | 0.0830918 | -0.079776 | 2 |
| Vapb | 0.72 | 0.0867695 | 0.076843 | 1 |
| Chd1l | 0.72 | 0.0981369 | 0.1400457 | 3 |
| Tmem69 | 0.72 | 0.1277693 | -0.187585 | 4 |
| C05463 | 0.72 | 0.1864071 | 0.1368181 | 3 |
| C14029 | 0.72 | 0.1907347 | 0.124998 | 3 |
| C02291 | 0.72 | 0.2007012 | -0.082194 | 2 |
| C05827 | 0.72 | 0.2015405 | -0.137747 | 2 |

| | | | | |
|---|---|---|---|---|
| C00532 | 0.72 | 0.2111842 | -0.090252 | 2 |
| C00666 | 0.72 | 0.2120997 | -0.078847 | 2 |
| C01726 | 0.72 | 0.2188038 | 0.0631788 | 3 |
| C00198 | 0.72 | 0.2278846 | -0.058596 | 2 |
| C00239a | 0.72 | 0.2354224 | -0.055751 | 2 |
| C01829 | 0.72 | 0.2631688 | -0.065455 | 2 |
| C02235 | 0.72 | 0.2670772 | 0.1342746 | 3 |
| C01789 | 0.72 | 0.2792641 | -0.101016 | 2 |
| C01041 | 0.72 | 0.324593 | -0.147734 | 2 |
| C00832 | 0.72 | 0.3360615 | 0.0892122 | 1 |
| C00670a | 0.72 | 0.3457452 | 0.0561023 | 3 |
| C01847 | 0.72 | 0.3736622 | 0.0399636 | 1 |
| HMDB0001973 | 0.72 | 0.3975801 | 0.1441718 | 3 |
| C00695a | 0.72 | 0.4467702 | -0.036962 | 2 |
| C00100 | 0.72 | 0.5294332 | 0.0712702 | 3 |
| C14302 | 0.72 | 0.5629324 | 0.0292471 | 1 |
| Hnrnpl | 0.68 | 0.0722798 | 0.1119216 | 1 |
| Fdx2 | 0.68 | 0.0913661 | -0.136191 | 4 |
| C01275 | 0.68 | 0.1459183 | 0.1951822 | 1 |
| HMDB0000459 | 0.68 | 0.1461361 | -0.084121 | 4 |
| C00597 | 0.68 | 0.1937064 | -0.131625 | 2 |
| C01029 | 0.68 | 0.2449565 | -0.077975 | 2 |
| C01673 | 0.68 | 0.2476872 | 0.099672 | 3 |
| HMDB0015517 | 0.68 | 0.249626 | -0.204055 | 4 |
| C01292 | 0.68 | 0.2779721 | -0.150085 | 2 |
| C00695 | 0.68 | 0.2990435 | -0.080535 | 2 |
| C01657 | 0.68 | 0.3115947 | -0.057376 | 2 |
| C01390 | 0.68 | 0.3323346 | 0.0494161 | 3 |
| C01230 | 0.68 | 0.3474967 | -0.146599 | 4 |
| C00659 | 0.68 | 0.3477542 | 0.0339626 | 1 |
| C03943 | 0.68 | 0.3707136 | -0.066478 | 2 |
| C01661 | 0.68 | 0.3857715 | 0.1326917 | 1 |
| C02236 | 0.68 | 0.40124 | -0.077197 | 4 |
| C00122 | 0.68 | 0.4176573 | -0.055115 | 2 |
| C00018 | 0.68 | 0.4207778 | 0.0853365 | 3 |
| C00158 | 0.68 | 0.4357701 | -0.112258 | 2 |
| C00262a | 0.68 | 0.4403827 | -0.118204 | 2 |
| C00101 | 0.68 | 0.5331591 | -0.099303 | 2 |
| C00475a | 0.68 | 0.5369849 | 0.1010149 | 3 |

| | | | | |
|---|---|---|---|---|
| C00843 | 0.68 | 0.6276734 | -0.038675 | 4 |
| C00477 | 0.68 | 0.8105134 | 0.0354984 | 3 |
| C02989 | 0.66 | 0.8659081 | -0.014603 | 4 |
| HMDB0002005 | 0.66 | 0.8659081 | -0.014603 | 4 |
| Sec1 | 0.64 | 0.1393016 | -0.153453 | 4 |
| C03296 | 0.64 | 0.2136449 | -0.073975 | 2 |
| C16843 | 0.64 | 0.2637007 | -0.105393 | 2 |
| C00202 | 0.64 | 0.3073355 | 0.1075495 | 1 |
| C00811 | 0.64 | 0.3546638 | 0.0641988 | 3 |
| C00643 | 0.64 | 0.3667584 | -0.076073 | 4 |
| C00022 | 0.64 | 0.3866548 | 0.0830433 | 3 |
| C01852 | 0.64 | 0.4130405 | 0.1264391 | 1 |
| C00158a | 0.64 | 0.4207253 | -0.045684 | 2 |
| C05122 | 0.64 | 0.486162 | 0.0404573 | 3 |
| C00993a | 0.64 | 0.4971141 | -0.092667 | 2 |
| C00433 | 0.64 | 0.5375035 | -0.072026 | 2 |
| C00262 | 0.64 | 0.5415141 | -0.052735 | 2 |
| C00090 | 0.64 | 0.5842699 | -0.039197 | 2 |
| C01042 | 0.64 | 0.688195 | -0.043323 | 2 |
| C00051 | 0.64 | 0.8190424 | -0.010007 | 2 |
| C01672 | 0.6 | 0.2792027 | 0.058374 | 3 |
| C00328 | 0.6 | 0.3270114 | 0.0399082 | 3 |
| C05635 | 0.6 | 0.3344376 | 0.0394673 | 3 |
| C02218 | 0.6 | 0.4091426 | -0.059117 | 4 |
| C01017 | 0.6 | 0.4450468 | -0.045059 | 4 |
| HMDB0011635 | 0.6 | 0.4931659 | -0.098795 | 4 |
| C10438 | 0.6 | 0.5130118 | 0.1143212 | 1 |
| HMDB0014581 | 0.6 | 0.5420141 | -0.046299 | 2 |
| C00021 | 0.6 | 0.5596228 | -0.037717 | 2 |
| C00499 | 0.6 | 0.6653451 | -0.034691 | 2 |
| C07075 | 0.6 | 0.6743459 | -0.046082 | 2 |
| C02043 | 0.6 | 0.6841239 | -0.046692 | 2 |
| C15557 | 0.6 | 0.688676 | -0.038909 | 2 |
| C15699 | 0.6 | 0.710624 | -0.043078 | 2 |
| C01005 | 0.6 | 0.7918338 | 0.0423278 | 3 |
| HMDB0000734 | 0.6 | 0.9172223 | -0.022629 | 4 |
| C00032 | 0.58 | 0.3414331 | 0.1128379 | 1 |
| C01198 | 0.56 | 0.4049942 | -0.107592 | 4 |
| C07354 | 0.56 | 0.4060211 | -0.088512 | 4 |

| | | | | |
|---|---|---|---|---|
| C01699 | 0.56 | 0.4492469 | -0.051985 | 4 |
| C00079 | 0.56 | 0.5458072 | 0.1022969 | 1 |
| C00170 | 0.56 | 0.5700104 | 0.0583193 | 3 |
| C00288 | 0.56 | 0.5996705 | -0.049645 | 4 |
| C06830 | 0.56 | 0.6087825 | -0.052232 | 2 |
| C00341 | 0.56 | 0.7489153 | -0.026685 | 4 |
| C01561 | 0.56 | 0.8700032 | -0.012526 | 4 |
| C01152 | 0.52 | 0.3649306 | 0.0985073 | 3 |
| C05648 | 0.52 | 0.4516997 | -0.088602 | 4 |
| C01035 | 0.52 | 0.5490499 | -0.056416 | 4 |
| C01517 | 0.52 | 0.6364717 | -0.038439 | 2 |
| C07565 | 0.52 | 0.6933024 | -0.022241 | 2 |
| C00042 | 0.52 | 0.7538119 | -0.022058 | 2 |
| C00089 | 0.52 | 0.7952498 | -0.030474 | 4 |
| C00633 | 0.52 | 0.8645646 | 0.006609 | 3 |
| C01748a | 0.52 | 0.9044739 | 0.0051984 | 1 |
| C01424 | 0.52 | 0.9390707 | 0.0211314 | 4 |
| C00525 | 0.52 | 0.9892668 | -0.005498 | 4 |
| HMDB0006029 | 0.52 | 0.9963119 | 0.0083813 | 2 |

**Supplementary Figure-8: Metaproteome analysis: List of identified peptide sequence specific to hepatic microbiome across all mice liver samples and their quantification.**

| Sr no. | Phylum: Order: LCA: Sequence : EC | Average Healthy | Average CCl4 | Average CCl4-ASA | FC CCL4/ Con | FC CCl4+ ASA/ Con | FC CCL4 ASA/ CCl4 | log2FC CCl4/ Con | log2FC CCl4+ASA/ Control | log2FC CCl4+ASA/ CCl4 | p val CCl4 /con | p val CCl4+ ASA/ Con | p value CCL4 ASA/ CCl4 |
|---|---|---|---|---|---|---|---|---|---|---|---|---|---|
| 1 | __Bacteria_AEGLSWR_2.5.1.75 | 1402570.31 | 1.00 | 1.00 | 0.00 | 0.00 | 1.00 | -20.42 | -20.42 | 0.00 | 0.07 | 1.00 | 1.00 |
| 2 | Actinobacteria__Actinobacteria_AGAGVLLR_2.4.1.- | 23266542.22 | 1.00 | 1.00 | 0.00 | 0.00 | 1.00 | -24.47 | -24.47 | 0.00 | 0.15 | 1.00 | 1.00 |
| 3 | __Bacteria_ASQELEITLTSR_ | 209145.28 | 1.00 | 1.00 | 0.00 | 0.00 | 1.00 | -17.67 | -17.67 | 0.00 | 0.00 | 1.00 | 1.00 |
| 4 | Actinobacteria_Kineosporiaceae_Kineococcus radiotolerans_ATGLDDLR_1.2.1.70 | 1.00 | 1.00 | 1.00 | 1.00 | 1.00 | 1.00 | 0.00 | 0.00 | 0.00 | 1.00 | 1.00 | 1.00 |
| 5 | Proteobacteria__Proteobacteria_CVIELDIAK_5.99.1.2 | 1.00 | 1.00 | 1.00 | 1.00 | 1.00 | 1.00 | 0.00 | 0.00 | 0.00 | 1.00 | 1.00 | 1.00 |
| 6 | __Bacteria_DEILIAR_ | 1.00 | 1.00 | 1.00 | 1.00 | 1.00 | 1.00 | 0.00 | 0.00 | 0.00 | 1.00 | 1.00 | 1.00 |
| 7 | __Bacteria_DGLNIWR_6.3.5.4 | 8626625.24 | 1.00 | 1.00 | 0.00 | 0.00 | 1.00 | -23.04 | -23.04 | 0.00 | 0.18 | 1.00 | 1.00 |
| 8 | __Bacteria_DGLVHISK_2.7.7.8 | 1.00 | 1.00 | 1.00 | 1.00 | 1.00 | 1.00 | 0.00 | 0.00 | 0.00 | 1.00 | 1.00 | 1.00 |
| 9 | __Bacteria_DLLLEYNR_ | 1.00 | 1.00 | 1.00 | 1.00 | 1.00 | 1.00 | 0.00 | 0.00 | 0.00 | 1.00 | 1.00 | 1.00 |
| 10 | Spirochaetes_Leptospiraceae_Leptospira_DLLSYLGK_6.1.1.19 | 1.00 | 1.00 | 1.00 | 1.00 | 1.00 | 1.00 | 0.00 | 0.00 | 0.00 | 1.00 | 1.00 | 1.00 |
| 11 | __Bacteria_DVESQDVLLFIDNIFR_3.6.3.14 | 1.00 | 1.00 | 1.00 | 1.00 | 1.00 | 1.00 | 0.00 | 0.00 | 0.00 | 1.00 | 1.00 | 1.00 |
| 12 | Firmicutes__Bacilli_EDGEALTQQK_3.4.11.6 | 11430527 4.40 | 1.00 | 1.00 | 0.00 | 0.00 | 1.00 | -26.77 | -26.77 | 0.00 | 0.28 | 1.00 | 1.00 |
| 13 | Proteobacteria_Enterobacteriaceae_Salmonella enterica_EDLIITAR_2.7.7.7 | 1.00 | 1.00 | 1.00 | 1.00 | 1.00 | 1.00 | 0.00 | 0.00 | 0.00 | 1.00 | 1.00 | 1.00 |
| 14 | __Bacteria_EDPIVEK_1.14.14.18 | 1624386.01 | 1.00 | 1.00 | 0.00 | 0.00 | 1.00 | -20.63 | -20.63 | 0.00 | 0.30 | 1.00 | 1.00 |
| 15 | Thermotogae_Thermotogaceae_Thermotoga neapolitana_EEVISFLEK_2.4.2.17 | 1.00 | 1.00 | 1.00 | 1.00 | 1.00 | 1.00 | 0.00 | 0.00 | 0.00 | 1.00 | 1.00 | 1.00 |
| 16 | Tenericutes_Mycoplasmataceae_Mycoplasma mobile_EISNILDNYEK_ | 124514.99 | 1.00 | 1.00 | 0.00 | 0.00 | 1.00 | -16.93 | -16.93 | 0.00 | 0.00 | 1.00 | 1.00 |
| 17 | Actinobacteria_Streptomycetaceae_Streptomycetaceae_ELGGPTAVTR_3.5.2.6 | 11336426.59 | 1.00 | 1.00 | 0.00 | 0.00 | 1.00 | -23.43 | -23.43 | 0.00 | 0.00 | 1.00 | 1.00 |
| 18 | __Bacteria_ELLAPAIR_3.1.26.4 | 1.00 | 1.00 | 1.00 | 1.00 | 1.00 | 1.00 | 0.00 | 0.00 | 0.00 | 1.00 | 1.00 | 1.00 |
| 19 | __Bacteria_ENTVVIK_ | 12562769.84 | 1.00 | 1.00 | 0.00 | 0.00 | 1.00 | -23.58 | -23.58 | 0.00 | 0.18 | 1.00 | 1.00 |
| 20 | Tenericutes_Mycoplasmataceae_Mycoplasma pneumoniae_ESLVVDNDSFK_5.4.2.8 | 1706497.45 | 1.00 | 1.00 | 0.00 | 0.00 | 1.00 | -20.70 | -20.70 | 0.00 | 0.09 | 1.00 | 1.00 |

| | | | | | | | | | | | | |
|---|---|---|---|---|---|---|---|---|---|---|---|---|
| 21 | Firmicutes__Bacilli_FAAVIFNNVR_2.3.2.3 | 1.00 | 1.00 | 1.00 | 1.00 | 1.00 | 1.00 | 0.00 | 0.00 | 0.00 | 1.00 | 1.00 | 1.00 |
| 22 | Proteobacteria_Desulfovibrionaceae_Desulfovibrio vulgaris_GAVLSILDELK_ | 882271.03 | 1.00 | 1.00 | 0.00 | 0.00 | 1.00 | -19.75 | -19.75 | 0.00 | 0.22 | 1.00 | 1.00 |
| 23 | Proteobacteria_Rickettsiaceae_Rickettsia_GLEVLAEVK_2.5.1.55 | 89129780.51 | 1.00 | 1.00 | 0.00 | 0.00 | 1.00 | -26.41 | -26.41 | 0.00 | 0.05 | 1.00 | 1.00 |
| 24 | Firmicutes_Streptococcaceae_Streptococcus pneumoniae_IAELWGTTDDQVR_6.3.5.5 | 1.00 | 1.00 | 1.00 | 1.00 | 1.00 | 1.00 | 0.00 | 0.00 | 0.00 | 1.00 | 1.00 | 1.00 |
| 25 | __Bacteria_IDEILLVGGSTR_ | 1.00 | 1.00 | 1.00 | 1.00 | 1.00 | 1.00 | 0.00 | 0.00 | 0.00 | 1.00 | 1.00 | 1.00 |
| 26 | __Bacteria_IDILIAK_2.8.4.4 | 1.00 | 1.00 | 1.00 | 1.00 | 1.00 | 1.00 | 0.00 | 0.00 | 0.00 | 1.00 | 1.00 | 1.00 |
| 27 | Actinobacteria_Streptomycetaceae_Streptomycetaceae_IGGVDVLAR_2.2.1.7 | 3971907.06 | 1.00 | 1.00 | 0.00 | 0.00 | 1.00 | -21.92 | -21.92 | 0.00 | 0.02 | 1.00 | 1.00 |
| 28 | Firmicutes__Bacilli_ILVSLSENEIL_ | 1.00 | 1.00 | 1.00 | 1.00 | 1.00 | 1.00 | 0.00 | 0.00 | 0.00 | 1.00 | 1.00 | 1.00 |
| 29 | Proteobacteria_Enterobacteriaceae_Salmonella_IVWPLLK_ | 2961265.48 | 1.00 | 1.00 | 0.00 | 0.00 | 1.00 | -21.50 | -21.50 | 0.00 | 0.14 | 1.00 | 1.00 |
| 30 | Proteobacteria__Gammaproteobacteria_LAQAQVAK_6.1.1.9 | 1.00 | 1.00 | 1.00 | 1.00 | 1.00 | 1.00 | 0.00 | 0.00 | 0.00 | 1.00 | 1.00 | 1.00 |
| 31 | Actinobacteria_Mycobacteriaceae_Mycobacterium leprae_LGLLKPGSR_2.7.8.8 | 1.00 | 1.00 | 1.00 | 1.00 | 1.00 | 1.00 | 0.00 | 0.00 | 0.00 | 1.00 | 1.00 | 1.00 |
| 32 | Firmicutes__Bacilli_LGVTVTLR_ | 1.00 | 1.00 | 1.00 | 1.00 | 1.00 | 1.00 | 0.00 | 0.00 | 0.00 | 1.00 | 1.00 | 1.00 |
| 33 | Proteobacteria__Proteobacteria_LISEEIELSR_2.7.3.9 | 1.00 | 1.00 | 1.00 | 1.00 | 1.00 | 1.00 | 0.00 | 0.00 | 0.00 | 1.00 | 1.00 | 1.00 |
| 34 | __Bacteria_LLVLEDLK_ | 1.00 | 1.00 | 1.00 | 1.00 | 1.00 | 1.00 | 0.00 | 0.00 | 0.00 | 1.00 | 1.00 | 1.00 |
| 35 | __Bacteria_LPPLLNK_2.8.3.16 | 31217602.27 | 1.00 | 1.00 | 0.00 | 0.00 | 1.00 | -24.90 | -24.90 | 0.00 | 0.08 | 1.00 | 1.00 |
| 36 | __Bacteria_MLQEAVDALVDNGR_2.7.7.6 | 1.00 | 1.00 | 1.00 | 1.00 | 1.00 | 1.00 | 0.00 | 0.00 | 0.00 | 1.00 | 1.00 | 1.00 |
| 37 | Proteobacteria_Bdellovibrionaceae_Bdellovibrio bacteriovorus_MMGAADEEK_ | 4210003.18 | 1.00 | 1.00 | 0.00 | 0.00 | 1.00 | -22.01 | -22.01 | 0.00 | 0.03 | 1.00 | 1.00 |
| 38 | Firmicutes_Peptococcaceae_Desulfotomaculum reducens_NILGSVTGR_ | 1.00 | 1.00 | 1.00 | 1.00 | 1.00 | 1.00 | 0.00 | 0.00 | 0.00 | 1.00 | 1.00 | 1.00 |
| 39 | Tenericutes_Mycoplasmataceae_Mycoplasma penetrans_NILGYLQR_2.7.1.11 | 1.00 | 1.00 | 1.00 | 1.00 | 1.00 | 1.00 | 0.00 | 0.00 | 0.00 | 1.00 | 1.00 | 1.00 |
| 40 | __Bacteria_NPIIVDK_2.1.1.- | 1.00 | 1.00 | 1.00 | 1.00 | 1.00 | 1.00 | 0.00 | 0.00 | 0.00 | 1.00 | 1.00 | 1.00 |
| 41 | __Bacteria_NPIVLDK_2.8.3.10 | 169947.00 | 1.00 | 1.00 | 0.00 | 0.00 | 1.00 | -17.37 | -17.37 | 0.00 | 0.07 | 1.00 | 1.00 |
| 42 | Tenericutes_Mycoplasmataceae_Mycoplasma pulmonis_NQEEIFESVLK_2.7.4.9 | 1.00 | 1.00 | 1.00 | 1.00 | 1.00 | 1.00 | 0.00 | 0.00 | 0.00 | 1.00 | 1.00 | 1.00 |
| 43 | Actinobacteria_Corynebacteriaceae_Corynebacterium_QAASVGSDELDR_1.2.1.70 | 1.00 | 1.00 | 1.00 | 1.00 | 1.00 | 1.00 | 0.00 | 0.00 | 0.00 | 1.00 | 1.00 | 1.00 |
| 44 | __Bacteria_QDGTYDK_5.4.3.8 | 42587055 | 1.00 | 1.00 | 0.00 | 0.00 | 1.00 | -25.34 | -25.34 | 0.00 | 0.05 | 1.00 | 1.00 |

| | | | | | | | | | | | |
|---|---|---|---|---|---|---|---|---|---|---|---|
| | | .73 | | | | | | | | | |
| 45 | Firmicutes_Clostridiaceae_Clostridium botulinum_QETIIISIAK_1.1.1.94 | 1.00 | 1.00 | 1.00 | 1.00 | 1.00 | 1.00 | 0.00 | 0.00 | 0.00 | 1.00 | 1.00 | 1.00 |
| 46 | __Bacteria_QMLQDLGR_ | 1.00 | 1.00 | 1.00 | 1.00 | 1.00 | 1.00 | 0.00 | 0.00 | 0.00 | 1.00 | 1.00 | 1.00 |
| 47 | Proteobacteria__Gammaproteobacteria_QPLLIGIAK_ | 1.00 | 1.00 | 1.00 | 1.00 | 1.00 | 1.00 | 0.00 | 0.00 | 0.00 | 1.00 | 1.00 | 1.00 |
| 48 | Thermotogae__Petrotoga_QSELGLYPAVDPLDSTSK_3.6.3.14 | 14491366.41 | 1.00 | 1.00 | 0.00 | 0.00 | 1.00 | -23.79 | -23.79 | 0.00 | 0.05 | 1.00 | 1.00 |
| 49 | __Bacteria_QVAELGIYPAVDPLDSTSR_3.6.3.14 | 745152.63 | 1.00 | 1.00 | 0.00 | 0.00 | 1.00 | -19.51 | -19.51 | 0.00 | 0.11 | 1.00 | 1.00 |
| 50 | Firmicutes__Bacilli_QVLSSVIK_6.1.1.20 | 1.00 | 1.00 | 1.00 | 1.00 | 1.00 | 1.00 | 0.00 | 0.00 | 0.00 | 1.00 | 1.00 | 1.00 |
| 51 | Cyanobacteria_Prochloraceae_Prochlorococcus marinus subsp. pastoris_SAEILVDAIR_4.2.1.9 | 46675733.38 | 1.00 | 1.00 | 0.00 | 0.00 | 1.00 | -25.48 | -25.48 | 0.00 | 0.07 | 1.00 | 1.00 |
| 52 | __Bacteria_SGIEVER_6.1.1.20 | 1.00 | 1.00 | 1.00 | 1.00 | 1.00 | 1.00 | 0.00 | 0.00 | 0.00 | 1.00 | 1.00 | 1.00 |
| 53 | Actinobacteria_Micrococcaceae_Renibacterium salmoninarum_SIDSGCGPADPAVR_6.1.1.4 | 3714765.60 | 1.00 | 1.00 | 0.00 | 0.00 | 1.00 | -21.82 | -21.82 | 0.00 | 0.05 | 1.00 | 1.00 |
| 54 | Tenericutes_Acholeplasmataceae_Candidatus Phytoplasma asteris_SSLISAISK_3.6.5.- | 8001348.74 | 1.00 | 1.00 | 0.00 | 0.00 | 1.00 | -22.93 | -22.93 | 0.00 | 0.06 | 1.00 | 1.00 |
| 55 | __Bacteria_SSLTGILSK_2.7.1.113 | 1.00 | 1.00 | 1.00 | 1.00 | 1.00 | 1.00 | 0.00 | 0.00 | 0.00 | 1.00 | 1.00 | 1.00 |
| 56 | Actinobacteria_Streptomycetaceae_Streptomyces_TAESVVLGAR_3.6.3.- | 13471005.21 | 1.00 | 1.00 | 0.00 | 0.00 | 1.00 | -23.68 | -23.68 | 0.00 | 0.04 | 1.00 | 1.00 |
| 57 | Firmicutes_Bacillaceae_Bacillus subtilis_TATSEMTDVSR_1.-.-.- | 91119176.55 | 1.00 | 1.00 | 0.00 | 0.00 | 1.00 | -26.44 | -26.44 | 0.00 | 0.03 | 1.00 | 1.00 |
| 58 | __Bacteria_TELSALLR_ | 1.00 | 1.00 | 1.00 | 1.00 | 1.00 | 1.00 | 0.00 | 0.00 | 0.00 | 1.00 | 1.00 | 1.00 |
| 59 | Actinobacteria__Actinobacteria_TPDIDEVGK_6.1.1.7 | 9850516.42 | 1.00 | 1.00 | 0.00 | 0.00 | 1.00 | -23.23 | -23.23 | 0.00 | 0.00 | 1.00 | 1.00 |
| 60 | Actinobacteria_Mycobacteriaceae_Mycobacteriaceae_TYHNEDDALVAAK_5.3.1.1 | 9972579.64 | 1.00 | 1.00 | 0.00 | 0.00 | 1.00 | -23.25 | -23.25 | 0.00 | 0.05 | 1.00 | 1.00 |
| 61 | __Bacteria_VAALAELLR_6.5.1.1 | 1.00 | 1.00 | 1.00 | 1.00 | 1.00 | 1.00 | 0.00 | 0.00 | 0.00 | 1.00 | 1.00 | 1.00 |
| 62 | Proteobacteria_Vibrionaceae_Vibrionaceae_VAGFVLGK_3.1.1.29 | 34906674.51 | 1.00 | 1.00 | 0.00 | 0.00 | 1.00 | -25.06 | -25.06 | 0.00 | 0.06 | 1.00 | 1.00 |
| 63 | __Bacteria_VDFISTK_ | 1.00 | 1.00 | 1.00 | 1.00 | 1.00 | 1.00 | 0.00 | 0.00 | 0.00 | 1.00 | 1.00 | 1.00 |
| 64 | __Bacteria_VGLVAGAVVK_ | 1.00 | 1.00 | 1.00 | 1.00 | 1.00 | 1.00 | 0.00 | 0.00 | 0.00 | 1.00 | 1.00 | 1.00 |
| 65 | Firmicutes__Bacilli_VLEEETGHQSLNCEK_2.7.13.3 | 1.00 | 1.00 | 1.00 | 1.00 | 1.00 | 1.00 | 0.00 | 0.00 | 0.00 | 1.00 | 1.00 | 1.00 |
| 66 | Actinobacteria_Frankiaceae_Frankia alni_VLLIGAGAVGSLAAQTAR_1.2.1.70 | 362351.91 | 1.00 | 1.00 | 0.00 | 0.00 | 1.00 | -18.47 | -18.47 | 0.00 | 0.01 | 1.00 | 1.00 |
| 67 | __Bacteria_VLVLGLAR_6.3.2.9 | 3309293. | 1.00 | 1.00 | 0.00 | 0.00 | 1.00 | -21.66 | -21.66 | 0.00 | 0.05 | 1.00 | 1.00 |

| | | | | | | | | | | | | |
|---|---|---|---|---|---|---|---|---|---|---|---|---|
| | | 02 | | | | | | | | | | |
| 68 | Firmicutes_Clostridiaceae_Alkaliphilus oremlandii_VSAIVEGGLER_2.7.7.60 | 13254871.51 | 1.00 | 1.00 | 0.00 | 0.00 | 1.00 | -23.66 | -23.66 | 0.00 | 0.02 | 1.00 | 1.00 |
| 69 | __Bacteria_WGADAIR_2.4.1.211 | 4643641.37 | 1.00 | 1.00 | 0.00 | 0.00 | 1.00 | -22.15 | -22.15 | 0.00 | 0.12 | 1.00 | 1.00 |
| 70 | __Bacteria_YDIDLLGR_ | 292415.39 | 1.00 | 1.00 | 0.00 | 0.00 | 1.00 | -18.16 | -18.16 | 0.00 | 0.01 | 1.00 | 1.00 |
| 71 | __Bacteria_YGGNAMVDDDLK_2.7.2.8 | 4671044.29 | 1.00 | 1.00 | 0.00 | 0.00 | 1.00 | -22.16 | -22.16 | 0.00 | 0.04 | 1.00 | 1.00 |
| 72 | Actinobacteria__Actinobacteria_YVVTNADDPGAAELACR_6.3.2.8 | 1.00 | 1.00 | 1.00 | 1.00 | 1.00 | 1.00 | 0.00 | 0.00 | 0.00 | 1.00 | 1.00 | 1.00 |
| 73 | Thermotogae__Petrotoga_DTNDINFLK_6.3.4.4 | 40944574.32 | 10930751.11 | 1.00 | 0.27 | 0.00 | 0.00 | -1.91 | -25.29 | -23.38 | 0.24 | 0.01 | 0.01 |
| 74 | __Bacteria_EEALAEFR_4.2.1.20 | 1.00 | 66878333.77 | 1.00 | 66878333.77 | 1.00 | 0.00 | 26.00 | 0.00 | -26.00 | 0.06 | 0.06 | 0.06 |
| 75 | Proteobacteria_Desulfovibrionaceae_Desulfovibrio vulgaris_GDHALIIYDDLSK_3.6.3.14 | 13205087.12 | 1250608.08 | 1.00 | 0.09 | 0.00 | 0.00 | -3.40 | -23.65 | -20.25 | 0.06 | 0.15 | 0.15 |
| 76 | Proteobacteria__Proteobacteria_GHMIADVVAIIGTQDIVFGEIDR_1.6.5.11 | 1823307.93 | 690090.38 | 1.00 | 0.38 | 0.00 | 0.00 | -1.40 | -20.80 | -19.40 | 0.54 | 0.25 | 0.25 |
| 77 | __Bacteria_HVLVVYDDLSK_3.6.3.14 | 1.00 | 1.00 | 1.00 | 1.00 | 1.00 | 1.00 | 0.00 | 0.00 | 0.00 | 1.00 | 1.00 | 1.00 |
| 80 | Firmicutes_Bacillaceae_Bacillus_ILQSLVAK_ | 1.00 | 1.00 | 1.00 | 1.00 | 1.00 | 1.00 | 0.00 | 0.00 | 0.00 | 1.00 | 1.00 | 1.00 |
| 81 | __Bacteria_INLIAVGK_2.1.1.177 | 1017286.92 | 1.00 | 1.00 | 0.00 | 0.00 | 1.00 | -19.96 | -19.96 | 0.00 | 0.10 | 1.00 | 1.00 |
| 82 | Cyanobacteria_Synechococcaceae_Synechococcus_LPESILLEMEGLAAR_2.8.4.3 | 797219.36 | 4033790.36 | 1.00 | 5.06 | 0.00 | 0.00 | 2.34 | -19.60 | -21.94 | 0.12 | 0.06 | 0.06 |
| 83 | Proteobacteria_Bradyrhizobiaceae_Bradyrhizobiaceae_LVVIAIGR_2.1.1.177 | 1.00 | 1.00 | 1.00 | 1.00 | 1.00 | 1.00 | 0.00 | 0.00 | 0.00 | 1.00 | 1.00 | 1.00 |
| 84 | Firmicutes_Streptococcaceae_Streptococcus_MTLEEQEFNLLK_3.1.-.- | 1.00 | 1.00 | 1.00 | 1.00 | 1.00 | 1.00 | 0.00 | 0.00 | 0.00 | 1.00 | 1.00 | 1.00 |
| 85 | Firmicutes__Bacilli_SANVIAAHVDHR_6.3.4.19 | 5281081.22 | 14749799.11 | 1.00 | 2.79 | 0.00 | 0.00 | 1.48 | -22.33 | -23.81 | 0.20 | 0.05 | 0.05 |
| 86 | Proteobacteria_Rickettsiaceae_Orientia_TVLIMEIINNVAK_3.6.3.14 | 296519.42 | 1328665.99 | 1.00 | 4.48 | 0.00 | 0.00 | 2.16 | -18.18 | -20.34 | 0.27 | 0.17 | 0.17 |
| 87 | Proteobacteria_Enterobacteriaceae_Klebsiella pneumoniae_VDIIVLPIK_ | 746846.25 | 701182.47 | 1.00 | 0.94 | 0.00 | 0.00 | -0.09 | -19.51 | -19.42 | 0.93 | 0.07 | 0.07 |
| 88 | Proteobacteria__Betaproteobacteria_VGLLLAEGR_1.1.1.94 | 210625.13 | 342235.27 | 1.00 | 1.62 | 0.00 | 0.00 | 0.70 | -17.68 | -18.38 | 0.43 | 0.05 | 0.05 |
| 90 | Tenericutes_Mycoplasmataceae_Mycoplasma penetrans_YSIDLIIK_2.3.1.2 | 2640395.28 | 512027.26 | 1.00 | 0.19 | 0.00 | 0.00 | -2.37 | -21.33 | -18.97 | 0.05 | 0.09 | 0.09 |

| | | | | | | | | | | | | | |
|---|---|---|---|---|---|---|---|---|---|---|---|---|---|
| | 34 | | | | | | | | | | | | |
| 91 | Proteobacteria_Enterobacteriaceae_Enterobacteriaceae_AANIIIHCEGK_5.3.1.13 | 1.00 | 1.00 | 1.00 | 1.00 | 1.00 | 1.00 | 0.00 | 0.00 | 0.00 | 1.00 | 1.00 | 1.00 |
| 93 | __Bacteria_AGDVIEVR_2.7.1.23 | 1.00 | 1.00 | 1.00 | 1.00 | 1.00 | 1.00 | 0.00 | 0.00 | 0.00 | 1.00 | 1.00 | 1.00 |
| 94 | Actinobacteria_Microbacteriaceae_Leifsonia xyli_AGDWAAYGAADSR_ | 1.00 | 1.00 | 1.00 | 1.00 | 1.00 | 1.00 | 0.00 | 0.00 | 0.00 | 1.00 | 1.00 | 1.00 |
| 96 | __Bacteria_ALLNLLAK_5.4.99.5 | 1.00 | 451765.85 | 1.00 | 451765.85 | 1.00 | 0.00 | 18.79 | 0.00 | -18.79 | 0.01 | 0.01 | 0.01 |
| 97 | __Bacteria_AVDGIVLFDK_2.7.13.3 | 1.00 | 2616257.47 | 1.00 | 2616257.47 | 1.00 | 0.00 | 21.32 | 0.00 | -21.32 | 0.12 | 0.12 | 0.12 |
| 99 | Actinobacteria_Nocardiaceae_Rhodococcus_AVLIGPPGAGK_2.7.1.71 | 1.00 | 1.00 | 1.00 | 1.00 | 1.00 | 1.00 | 0.00 | 0.00 | 0.00 | 1.00 | 1.00 | 1.00 |
| 100 | Proteobacteria_Pasteurellaceae_Haemophilus_CVIVSADFK_4.1.1.65 | 1.00 | 1.00 | 1.00 | 1.00 | 1.00 | 1.00 | 0.00 | 0.00 | 0.00 | 1.00 | 1.00 | 1.00 |
| 101 | Proteobacteria__Proteobacteria_DEMGLDVLMFIDNIFR_3.6.3.14 | 1.00 | 27333431.46 | 1.00 | 27333431.46 | 1.00 | 0.00 | 24.70 | 0.00 | -24.70 | 0.29 | 0.29 | 0.29 |
| 104 | __Bacteria_ELPIIIK_ | 1.00 | 5600283.91 | 1.00 | 5600283.91 | 1.00 | 0.00 | 22.42 | 0.00 | -22.42 | 0.07 | 0.07 | 0.07 |
| 105 | Cyanobacteria_Prochloraceae_Prochlorococcus marinus subsp. pastoris_ENLLEIIEK_2.5.1.3 | 1.00 | 1.00 | 1.00 | 1.00 | 1.00 | 1.00 | 0.00 | 0.00 | 0.00 | 1.00 | 1.00 | 1.00 |
| 108 | Tenericutes_Mycoplasmataceae_Mycoplasma pneumoniae_GAANALQAIVK_ | 1.00 | 2093006.67 | 1.00 | 2093006.67 | 1.00 | 0.00 | 21.00 | 0.00 | -21.00 | 0.00 | 0.00 | 0.00 |
| 110 | __Bacteria_GRPGLAER_ | 1.00 | 330055.03 | 1.00 | 330055.03 | 1.00 | 0.00 | 18.33 | 0.00 | -18.33 | 0.01 | 0.01 | 0.01 |
| 112 | Proteobacteria_Erwiniaceae_Erwinia_IAVQLIGR_ | 1.00 | 1.00 | 1.00 | 1.00 | 1.00 | 1.00 | 0.00 | 0.00 | 0.00 | 1.00 | 1.00 | 1.00 |
| 113 | __Bacteria_IGNDVIIK_2.3.1.129 | 1.00 | 1494066.36 | 1.00 | 1494066.36 | 1.00 | 0.00 | 20.51 | 0.00 | -20.51 | 0.10 | 0.10 | 0.10 |
| 114 | Proteobacteria_Erwiniaceae_Buchnera aphidicola_IIGGILGR_2.1.2.3 | 1.00 | 1.00 | 1.00 | 1.00 | 1.00 | 1.00 | 0.00 | 0.00 | 0.00 | 1.00 | 1.00 | 1.00 |
| 115 | Firmicutes_Clostridiaceae_Clostridium botulinum_IIVLDGGIGVGR_2.1.1.195 | 1.00 | 2969223.51 | 1.00 | 2969223.51 | 1.00 | 0.00 | 21.50 | 0.00 | -21.50 | 0.03 | 0.03 | 0.03 |
| 116 | Firmicutes_Bacillaceae_Lysinibacillus_ILDSVILK_2.1.1.74 | 1.00 | 1.00 | 1.00 | 1.00 | 1.00 | 1.00 | 0.00 | 0.00 | 0.00 | 1.00 | 1.00 | 1.00 |
| 117 | Firmicutes_Streptococcaceae_Streptococcus pneumoniae_ILLNMDK_ | 1.00 | 1.00 | 1.00 | 1.00 | 1.00 | 1.00 | 0.00 | 0.00 | 0.00 | 1.00 | 1.00 | 1.00 |
| 119 | Actinobacteria_Mycobacteriaceae_Mycobacteriaceae_LAGAVLIGR_6.3.2.9 | 1.00 | 1.00 | 1.00 | 1.00 | 1.00 | 1.00 | 0.00 | 0.00 | 0.00 | 1.00 | 1.00 | 1.00 |
| 120 | __Bacteria_LEGLPPK_3.2.2.21 | 1.00 | 217263852. | 1.00 | 21726385 2.13 | 1.00 | 0.00 | 27.69 | 0.00 | -27.69 | 0.17 | 0.17 | 0.17 |

| | | | | | | | | | | | | | |
|---|---|---|---|---|---|---|---|---|---|---|---|---|---|
| | | | 13 | | | | | | | | | | |
| 121 | __Bacteria_LELLEPLR_6.3.5.2 | 1.00 | 254321.82 | 1.00 | 254321.82 | 1.00 | 0.00 | 17.96 | 0.00 | -17.96 | 0.04 | 0.04 | 0.04 |
| 122 | Actinobacteria_Mycobacteriaceae_Mycobacterium tuberculosis complex_LEVGPVAR_ | 1.00 | 1.00 | 1.00 | 1.00 | 1.00 | 1.00 | 0.00 | 0.00 | 0.00 | 1.00 | 1.00 | 1.00 |
| 124 | __Bacteria_LLTEIIIK_1.17.1.5 | 1.00 | 1.00 | 1.00 | 1.00 | 1.00 | 1.00 | 0.00 | 0.00 | 0.00 | 1.00 | 1.00 | 1.00 |
| 125 | Firmicutes__Bacilli_LNDLPIIPNK_5.99.1.2 | 1.00 | 1.00 | 1.00 | 1.00 | 1.00 | 1.00 | 0.00 | 0.00 | 0.00 | 1.00 | 1.00 | 1.00 |
| 127 | Firmicutes__Bacilli_LTISLGALDK_2.7.13.3 | 1.00 | 27329221.26 | 1.00 | 27329221.26 | 1.00 | 0.00 | 24.70 | 0.00 | -24.70 | 0.01 | 0.01 | 0.01 |
| 128 | Firmicutes_Streptococcaceae_Streptococcus_MDEEEEQAFK_ | 1.00 | 1.00 | 1.00 | 1.00 | 1.00 | 1.00 | 0.00 | 0.00 | 0.00 | 1.00 | 1.00 | 1.00 |
| 130 | Proteobacteria_Aeromonadaceae_Aeromonas_QILEFGADK_4.1.3.- | 1.00 | 1.00 | 1.00 | 1.00 | 1.00 | 1.00 | 0.00 | 0.00 | 0.00 | 1.00 | 1.00 | 1.00 |
| 131 | __Bacteria_QIVPTLEK_2.7.2.3 | 1.00 | 436537.40 | 1.00 | 436537.40 | 1.00 | 0.00 | 18.74 | 0.00 | -18.74 | 0.07 | 0.07 | 0.07 |
| 133 | Firmicutes__Exiguobacterium_QSAASAELDLR_3.1.-.- | 1.00 | 13550109.04 | 1.00 | 13550109.04 | 1.00 | 0.00 | 23.69 | 0.00 | -23.69 | 0.05 | 0.05 | 0.05 |
| 134 | Actinobacteria_Mycobacteriaceae_Mycobacterium_QVAVVQYPTLSTSK_ | 1.00 | 1.00 | 1.00 | 1.00 | 1.00 | 1.00 | 0.00 | 0.00 | 0.00 | 1.00 | 1.00 | 1.00 |
| 135 | __Bacteria_SALALLGK_3.6.4.12 | 1.00 | 274671.95 | 1.00 | 274671.95 | 1.00 | 0.00 | 18.07 | 0.00 | -18.07 | 0.06 | 0.06 | 0.06 |
| 137 | __Bacteria_SDLNNPR_ | 1.00 | 1.00 | 1.00 | 1.00 | 1.00 | 1.00 | 0.00 | 0.00 | 0.00 | 1.00 | 1.00 | 1.00 |
| 140 | Actinobacteria_Corynebacteriaceae_Corynebacterium efficiens_TLAEGDHVTVGTR_2.7.7.6 | 1.00 | 7105433.68 | 1.00 | 7105433.68 | 1.00 | 0.00 | 22.76 | 0.00 | -22.76 | 0.09 | 0.09 | 0.09 |
| 143 | Firmicutes_Lactobacillaceae_Lactobacillus casei group_VIADHPINIQVLGIGR_3.5.99.6 | 1.00 | 1.00 | 1.00 | 1.00 | 1.00 | 1.00 | 0.00 | 0.00 | 0.00 | 1.00 | 1.00 | 1.00 |
| 144 | Firmicutes__Bacilli_VIAIGEIR_3.2.2.21 | 1.00 | 826644.28 | 1.00 | 826644.28 | 1.00 | 0.00 | 19.66 | 0.00 | -19.66 | 0.07 | 0.07 | 0.07 |
| 145 | __Bacteria_VLQALGLTR_ | 1.00 | 1.00 | 1.00 | 1.00 | 1.00 | 1.00 | 0.00 | 0.00 | 0.00 | 1.00 | 1.00 | 1.00 |
| 147 | __Bacteria_VVAGLGLR_ | 1.00 | 1.00 | 1.00 | 1.00 | 1.00 | 1.00 | 0.00 | 0.00 | 0.00 | 1.00 | 1.00 | 1.00 |
| 148 | __Bacteria_ACGDIDGLER_2.8.4.3 | 1.00 | 1.00 | 1.00 | 1.00 | 1.00 | 1.00 | 0.00 | 0.00 | 0.00 | 1.00 | 1.00 | 1.00 |
| 149 | Actinobacteria__Actinobacteria_AMFDLLVK_2.3.3.13 | 29841971.66 | 1.00 | 28341312.72 | 0.00 | 0.95 | 28341312.72 | -24.83 | -0.07 | 24.76 | 0.17 | 0.05 | 0.05 |
| 150 | Proteobacteria__Proteobacteria_DLDLPVGR_ | 21759890.09 | 1.00 | 50330981.53 | 0.00 | 2.31 | 50330981.53 | -24.38 | 1.21 | 25.58 | 0.04 | 0.04 | 0.04 |
| 151 | Firmicutes__Bacillales_EINLAPSAR_1.-.-.- | 1.00 | 1.00 | 1.00 | 1.00 | 1.00 | 1.00 | 0.00 | 0.00 | 0.00 | 1.00 | 1.00 | 1.00 |
| 152 | Firmicutes__Firmicutes_FELDNLAR_5.3.1.5 | 4310586.99 | 1.00 | 1.00 | 0.00 | 0.00 | 1.00 | -22.04 | -22.04 | 0.00 | 0.15 | 1.00 | 1.00 |

| | | | | | | | | | | | |
|---|---|---|---|---|---|---|---|---|---|---|---|
| 153 | __Bacteria_FIAPSQFK_3.6.3.24 | 1.00 | 1.00 | 1.00 | 1.00 | 1.00 | 1.00 | 0.00 | 0.00 | 0.00 | 1.00 | 1.00 | 1.00 |
| 154 | __Bacteria_GLTFDSGGISIK_3.4.11.10 | 19654863.73 | 1.00 | 21980803.43 | 0.00 | 1.12 | 21980803.43 | -24.23 | 0.16 | 24.39 | 0.13 | 0.00 | 0.00 |
| 155 | Proteobacteria_Desulfobacteraceae_Desulfococcus oleovorans_IAAIIPSR_2.7.7.38 | 1.00 | 1.00 | 1.00 | 1.00 | 1.00 | 1.00 | 0.00 | 0.00 | 0.00 | 1.00 | 1.00 | 1.00 |
| 156 | Firmicutes_Streptococcaceae_Streptococcus uberis_IAGLPGSLLSR_ | 1.00 | 1.00 | 1.00 | 1.00 | 1.00 | 1.00 | 0.00 | 0.00 | 0.00 | 1.00 | 1.00 | 1.00 |
| 157 | __Bacteria_IDIDADGR_6.3.4.3 | 60582954.21 | 1.00 | 96280954.50 | 0.00 | 1.59 | 96280954.50 | -25.85 | 0.67 | 26.52 | 0.01 | 0.07 | 0.07 |
| 158 | Proteobacteria__Proteobacteria_LDITALR_2.3.1.28 | 55387470.49 | 1.00 | 20243664.91 | 0.00 | 0.37 | 20243664.91 | -25.72 | -1.45 | 24.27 | 0.04 | 0.04 | 0.04 |
| 159 | Tenericutes_Mycoplasmataceae_Mycoplasma mycoides group_LLNKPGIK_5.3.1.9 | 6130613.83 | 1.00 | 9989423.32 | 0.00 | 1.63 | 9989423.32 | -22.55 | 0.70 | 23.25 | 0.14 | 0.14 | 0.14 |
| 160 | Firmicutes_Lachnospiraceae_Lachnoclostridium phytofermentans_LVLEDTGK_3.1.-.- | 58564591.66 | 1.00 | 30442498.14 | 0.00 | 0.52 | 30442498.14 | -25.80 | -0.94 | 24.86 | 0.00 | 0.09 | 0.09 |
| 161 | Tenericutes_Mycoplasmataceae_Mycoplasma mycoides group_QGDVVAILK_2.7.1.- | 1.00 | 1.00 | 1.00 | 1.00 | 1.00 | 1.00 | 0.00 | 0.00 | 0.00 | 1.00 | 1.00 | 1.00 |
| 162 | Proteobacteria_Erwiniaceae_Buchnera aphidicola_STLLMLIQR_3.6.3.44 | 7845978.74 | 1.00 | 5402838.86 | 0.00 | 0.69 | 5402838.86 | -22.90 | -0.54 | 22.37 | 0.16 | 0.13 | 0.13 |
| 163 | __Bacteria_TELMNIIK_6.1.1.19 | 25623971.24 | 1.00 | 45715111.02 | 0.00 | 1.78 | 45715111.02 | -24.61 | 0.84 | 25.45 | 0.06 | 0.03 | 0.03 |
| 164 | Proteobacteria_Brucellaceae_Ochrobactrum_TGLPISTYFSGLK_2.7.1.30 | 1.00 | 1.00 | 1.00 | 1.00 | 1.00 | 1.00 | 0.00 | 0.00 | 0.00 | 1.00 | 1.00 | 1.00 |
| 165 | Proteobacteria_Alteromonadaceae_Marinobacter hydrocarbonoclasticus_VADIGLTGGTPK_1.17.7.3 | 1.00 | 1.00 | 1.00 | 1.00 | 1.00 | 1.00 | 0.00 | 0.00 | 0.00 | 1.00 | 1.00 | 1.00 |
| 166 | __Bacteria_VAIVGIPNVGK_ | 1.00 | 1.00 | 1.00 | 1.00 | 1.00 | 1.00 | 0.00 | 0.00 | 0.00 | 1.00 | 1.00 | 1.00 |
| 167 | Actinobacteria_Streptomycetaceae_Streptomycetaceae_VGAIQYAR_6.3.4.2 | 1.00 | 1.00 | 1.00 | 1.00 | 1.00 | 1.00 | 0.00 | 0.00 | 0.00 | 1.00 | 1.00 | 1.00 |
| 168 | __Bacteria_VPVNIIR_2.7.7.85 | 1268595.67 | 1.00 | 1742787.66 | 0.00 | 1.37 | 1742787.66 | -20.27 | 0.46 | 20.73 | 0.12 | 0.09 | 0.09 |
| 169 | __Bacteria_VQEIAALR_6.5.1.2 | 1.00 | 1.00 | 1.00 | 1.00 | 1.00 | 1.00 | 0.00 | 0.00 | 0.00 | 1.00 | 1.00 | 1.00 |
| 170 | Actinobacteria_Corynebacteriaceae_Corynebacterium_AAMIIEGK_ | 3350252.79 | 2069416.73 | 6360831.09 | 0.62 | 1.90 | 3.07 | -0.70 | 0.92 | 1.62 | 0.59 | 0.19 | 0.19 |
| 171 | Actinobacteria_Streptomycetaceae_Streptomycetaceae_AEPGDTVLVAGK_6.3.2.13 | 16231007.64 | 2300007.45 | 42097184.65 | 0.14 | 2.59 | 18.30 | -2.82 | 1.37 | 4.19 | 0.20 | 0.05 | 0.05 |
| 172 | Tenericutes_Mycoplasmataceae_Mycoplasma pneumoniae_AFEVLTNK_ | 1.00 | 1.00 | 1.00 | 1.00 | 1.00 | 1.00 | 0.00 | 0.00 | 0.00 | 1.00 | 1.00 | 1.00 |
| 173 | __Bacteria_ALDIALAYAK_1.1.1.86 | 1.00 | 824530 | 1.00 | 8245300.4 | 1.00 | 0.00 | 22.98 | 0.00 | -22.98 | 0.13 | 0.13 | 0.13 |

| | | | | | | | | | | | | | |
|---|---|---|---|---|---|---|---|---|---|---|---|---|---|
| | | | 0.40 | | 0 | | | | | | | | |
| 174 | Actinobacteria__Actinobacteria_AVKPLVVR_6.2.1.5 | 1.00 | 1.00 | 1.00 | 1.00 | 1.00 | 1.00 | 0.00 | 0.00 | 0.00 | 1.00 | 1.00 | 1.00 |
| 175 | Firmicutes_Clostridiaceae_Clostridium perfringens_DFTPAEIR_2.1.3.3 | 30214019.32 | 3728857.40 | 46325408.29 | 0.12 | 1.53 | 12.42 | -3.02 | 0.62 | 3.63 | 0.19 | 0.11 | 0.11 |
| 177 | Firmicutes_Streptococcaceae_Streptococcus_DVDAILFVVPTK_1.1.1.94 | 2822388.71 | 1.00 | 4371014.20 | 0.00 | 1.55 | 4371014.20 | -21.43 | 0.63 | 22.06 | 0.11 | 0.11 | 0.11 |
| 178 | __Bacteria_DYLLVAK_3.1.-.- | 2807346.67 | 66605629.73 | 4136622.54 | 23.73 | 1.47 | 0.06 | 4.57 | 0.56 | -4.01 | 0.14 | 0.15 | 0.15 |
| 179 | Proteobacteria__Proteobacteria_EEIIAVIAK_ | 1.00 | 1.00 | 1.00 | 1.00 | 1.00 | 1.00 | 0.00 | 0.00 | 0.00 | 1.00 | 1.00 | 1.00 |
| 180 | __Bacteria_EELAALPR_6.1.1.17 | 74015707.59 | 1.00 | 76948256.74 | 0.00 | 1.04 | 76948256.74 | -26.14 | 0.06 | 26.20 | 0.05 | 0.02 | 0.02 |
| 181 | Proteobacteria_Cellvibrionaceae_Teredinibacter turnerae_ELDNIEITR_5.2.1.8 | 1.00 | 1873660.37 | 1.00 | 1873660.37 | 1.00 | 0.00 | 20.84 | 0.00 | -20.84 | 0.01 | 0.01 | 0.01 |
| 183 | __Bacteria_FDFIFPR_ | 339646.34 | 4187699.28 | 769699.86 | 12.33 | 2.27 | 0.18 | 3.62 | 1.18 | -2.44 | 0.03 | 0.06 | 0.06 |
| 184 | Proteobacteria__Alphaproteobacteria_FLAEAVQK_ | 76175150.00 | 2743187.48 | 1.00 | 0.04 | 0.00 | 0.00 | -4.80 | -26.18 | -21.39 | 0.17 | 0.13 | 0.13 |
| 185 | __Bacteria_FQTILQK_2.7.13.3 | 13926648.87 | 1.00 | 2976844.43 | 0.00 | 0.21 | 2976844.43 | -23.73 | -2.23 | 21.51 | 0.00 | 0.10 | 0.10 |
| 186 | __Bacteria_GDLALEVK_3.6.3.17 | 1.00 | 1.00 | 1.00 | 1.00 | 1.00 | 1.00 | 0.00 | 0.00 | 0.00 | 1.00 | 1.00 | 1.00 |
| 187 | Proteobacteria__Gammaproteobacteria_GIIPIPELK_4.1.3.- | 6267262.75 | 1.00 | 6788284.37 | 0.00 | 1.08 | 6788284.37 | -22.58 | 0.12 | 22.69 | 0.01 | 0.08 | 0.08 |
| 188 | Proteobacteria__Proteobacteria_GIPLVILAGK_4.2.1.3 | 8484220.59 | 674489.98 | 29804046.29 | 0.08 | 3.51 | 44.19 | -3.65 | 1.81 | 5.47 | 0.06 | 0.09 | 0.09 |
| 190 | Tenericutes_Mycoplasmataceae_Mycoplasma_IINEPTAAALSFGLDK_ | 1.00 | 1.00 | 1.00 | 1.00 | 1.00 | 1.00 | 0.00 | 0.00 | 0.00 | 1.00 | 1.00 | 1.00 |
| 191 | __Bacteria_IVEVIFK_2.1.3.- | 752701.66 | 1.00 | 979964.21 | 0.00 | 1.30 | 979964.21 | -19.52 | 0.38 | 19.90 | 0.05 | 0.02 | 0.02 |
| 192 | Proteobacteria__Gammaproteobacteria_LAQLGAEIR_2.5.1.7 | 4537450.03 | 2278264.45 | 5201779.03 | 0.50 | 1.15 | 2.28 | -0.99 | 0.20 | 1.19 | 0.24 | 0.18 | 0.18 |
| 193 | Actinobacteria__Actinobacteria_LEVELVVER_2.7.7.6 | 1.00 | 1372618.84 | 11042047.44 | 1372618.84 | 11042047.44 | 8.04 | 20.39 | 23.40 | 3.01 | 0.15 | 0.15 | 0.15 |
| 194 | __Bacteria_LIIGLGNPDPK_3.1.1.29 | 1.00 | 1.00 | 1.00 | 1.00 | 1.00 | 1.00 | 0.00 | 0.00 | 0.00 | 1.00 | 1.00 | 1.00 |
| 195 | __Bacteria_LLIDADIK_ | 25789307.39 | 1.00 | 1075409064.12 | 0.00 | 4.17 | 1075409064.12 | -24.62 | 2.06 | 26.68 | 0.06 | 0.06 | 0.06 |
| 196 | __Bacteria_LTLVEYGFR_ | 1.00 | 1.00 | 1.00 | 1.00 | 1.00 | 1.00 | 0.00 | 0.00 | 0.00 | 1.00 | 1.00 | 1.00 |
| 197 | Deinococcus-Thermus_Deinococcaceae_Deinococcus radiodurans_LVLGLPLR_3.1.-.- | 3214174.58 | 47416149.00 | 1775612.13 | 14.75 | 0.55 | 0.04 | 3.88 | -0.86 | -4.74 | 0.15 | 0.14 | 0.14 |
| 198 | __Bacteria_LVSGGTDNHLMLIDLR_2.1.2.1 | 1.00 | 844240. | 1.00 | 844240.81 | 1.00 | 0.00 | 19.69 | 0.00 | -19.69 | 0.08 | 0.08 | 0.08 |

| | | | | | | | | | | | | | |
|---|---|---|---|---|---|---|---|---|---|---|---|---|---|
| | | | 81 | | | | | | | | | | |
| 199 | Proteobacteria_Hyphomonadaceae_Hyphomonas_MEGALASLASEFAGLR_ | 49450279.85 | 1.00 | 127928139.17 | 0.00 | 2.59 | 127928139.17 | -25.56 | 1.37 | 26.93 | 0.05 | 0.00 | 0.00 |
| 200 | __Bacteria_NESLEDALR_ | 115416358.09 | 61815493.45 | 68994474.50 | 0.54 | 0.60 | 1.12 | -0.90 | -0.74 | 0.16 | 0.15 | 0.80 | 0.80 |
| 201 | Proteobacteria__Proteobacteria_QLPLTDEQK_3.1.11.5 | 110677980.38 | 25725562.80 | 157772171.67 | 0.23 | 1.43 | 6.13 | -2.11 | 0.51 | 2.62 | 0.01 | 0.10 | 0.10 |
| 202 | Proteobacteria_Erwiniaceae_Wigglesworthia glossinidia_SFIIIILK_2.4.2.53 | 1.00 | 126892461.72 | 1.00 | 126892461.72 | 1.00 | 0.00 | 26.92 | 0.00 | -26.92 | 0.03 | 0.03 | 0.03 |
| 203 | Proteobacteria_Vibrionaceae_Vibrio harveyi group_SGYGLTLQDEIK_3.5.2.7 | 1.00 | 1.00 | 1.00 | 1.00 | 1.00 | 1.00 | 0.00 | 0.00 | 0.00 | 1.00 | 1.00 | 1.00 |
| 204 | __Bacteria_SIGIFYR_ | 1.00 | 1.00 | 1.00 | 1.00 | 1.00 | 1.00 | 0.00 | 0.00 | 0.00 | 1.00 | 1.00 | 1.00 |
| 205 | Proteobacteria_Pasteurellaceae_Actinobacillus pleuropneumoniae_TFTTQETMANALTAR_5.3.1.9 | 1.00 | 4440671.74 | 1.00 | 4440671.74 | 1.00 | 0.00 | 22.08 | 0.00 | -22.08 | 0.13 | 0.13 | 0.13 |
| 207 | Proteobacteria__Proteobacteria_TMYVIPFSMGPLGSPIAK_4.1.1.32 | 678706.26 | 1.00 | 7184428.54 | 0.00 | 10.59 | 7184428.54 | -19.37 | 3.40 | 22.78 | 0.12 | 0.16 | 0.16 |
| 208 | Firmicutes_Bacillaceae_Bacillus_TPEQHLESLR_3.1.-.- | 1.00 | 3095352.54 | 1.00 | 3095352.54 | 1.00 | 0.00 | 21.56 | 0.00 | -21.56 | 0.12 | 0.12 | 0.12 |
| 209 | Proteobacteria__Alphaproteobacteria_TTPSVVAFTENER_ | 1.00 | 1.00 | 1.00 | 1.00 | 1.00 | 1.00 | 0.00 | 0.00 | 0.00 | 1.00 | 1.00 | 1.00 |
| 210 | __Bacteria_TTVAIDTIINQK_3.6.3.14 | 1.00 | 1.00 | 1.00 | 1.00 | 1.00 | 1.00 | 0.00 | 0.00 | 0.00 | 1.00 | 1.00 | 1.00 |
| 211 | Proteobacteria_Anaplasmataceae_Anaplasma_TVLIMELISNVAR_3.6.3.14 | 17953389.57 | 1.00 | 40608181.24 | 0.00 | 2.26 | 40608181.24 | -24.10 | 1.18 | 25.28 | 0.03 | 0.01 | 0.01 |
| 212 | __Bacteria_VAIDLGLK_ | 1.00 | 27754437.76 | 145385544.05 | 27754437.76 | 145385544.05 | 5.24 | 24.73 | 27.12 | 2.39 | 0.05 | 0.09 | 0.09 |
| 213 | Proteobacteria__Proteobacteria_VILAGEIR_2.5.1.6 | 1.00 | 22162933.12 | 1.00 | 22162933.12 | 1.00 | 0.00 | 24.40 | 0.00 | -24.40 | 0.14 | 0.14 | 0.14 |
| 214 | Tenericutes_Mycoplasmataceae_Mycoplasma pulmonis_VILPNLAK_2.7.4.9 | 1.00 | 1.00 | 1.00 | 1.00 | 1.00 | 1.00 | 0.00 | 0.00 | 0.00 | 1.00 | 1.00 | 1.00 |
| 215 | Actinobacteria__Corynebacteriales_VIVVTGAGGGLGR_1.1.1.- | 1.00 | 1.00 | 1.00 | 1.00 | 1.00 | 1.00 | 0.00 | 0.00 | 0.00 | 1.00 | 1.00 | 1.00 |
| 216 | Firmicutes_Clostridiaceae_Clostridium perfringens_VMILAYK_ | 4207185.04 | 1.00 | 10650659.37 | 0.00 | 2.53 | 10650659.37 | -22.00 | 1.34 | 23.34 | 0.00 | 0.00 | 0.00 |
| 217 | Firmicutes_Symbiobacteriaceae_Symbiobacterium thermophilum_VTTSPDYGK_6.1.1.16 | 1.00 | 1.00 | 1.00 | 1.00 | 1.00 | 1.00 | 0.00 | 0.00 | 0.00 | 1.00 | 1.00 | 1.00 |
| 218 | Actinobacteria_Streptomycetaceae_Streptomyces_YAPELNTR_3.5.4.4 | 1.00 | 1.00 | 1.00 | 1.00 | 1.00 | 1.00 | 0.00 | 0.00 | 0.00 | 1.00 | 1.00 | 1.00 |

| | | | | | | | | | | | | | |
|---|---|---|---|---|---|---|---|---|---|---|---|---|---|
| 219 | __Bacteria_YVVVSAITPTPLGEGK_6.3.4.3 | 1.00 | 1.00 | 1.00 | 1.00 | 1.00 | 1.00 | 0.00 | 0.00 | 0.00 | 1.00 | 1.00 | 1.00 |
| 220 | __Bacteria_AVIEALVK_3.6.4.12 | 1.00 | 5333038.38 | 50831597.05 | 5333038.38 | 50831597.05 | 9.53 | 22.35 | 25.60 | 3.25 | 0.21 | 0.04 | 0.04 |
| 221 | __Bacteria_DAAVQSAAER_6.1.1.20 | 1.00 | 1.00 | 1.00 | 1.00 | 1.00 | 1.00 | 0.00 | 0.00 | 0.00 | 1.00 | 1.00 | 1.00 |
| 222 | Actinobacteria_Micrococcaceae_Renibacterium salmoninarum_DDLPAGSR_2.4.2.7 | 1.00 | 2169342.14 | 20725197.98 | 2169342.14 | 20725197.98 | 9.55 | 21.05 | 24.30 | 3.26 | 0.14 | 0.09 | 0.09 |
| 223 | __Bacteria_DGVIVFK_ | 1.00 | 40950759.62 | 129069744.41 | 40950759.62 | 129069744.41 | 3.15 | 25.29 | 26.94 | 1.66 | 0.06 | 0.01 | 0.01 |
| 225 | __Bacteria_IAIAQVLK_ | 1.00 | 261021.37 | 1395655.78 | 261021.37 | 1395655.78 | 5.35 | 17.99 | 20.41 | 2.42 | 0.02 | 0.24 | 0.24 |
| 226 | __Bacteria_IGIITIK_2.7.11.1 | 1.00 | 1.00 | 1.00 | 1.00 | 1.00 | 1.00 | 0.00 | 0.00 | 0.00 | 1.00 | 1.00 | 1.00 |
| 227 | __Bacteria_IGQLLLTR_2.7.2.11 | 1.00 | 1.00 | 10567157.75 | 1.00 | 10567157.75 | 10567157.75 | 0.00 | 23.33 | 23.33 | 1.00 | 0.12 | 0.12 |
| 228 | __Bacteria_ILILESLK_ | 1.00 | 1.00 | 1.00 | 1.00 | 1.00 | 1.00 | 0.00 | 0.00 | 0.00 | 1.00 | 1.00 | 1.00 |
| 229 | __Bacteria_ILTTPIK_4.2.1.32 | 1.00 | 3240534.50 | 4923674.48 | 3240534.50 | 4923674.48 | 1.52 | 21.63 | 22.23 | 0.60 | 0.05 | 0.56 | 0.56 |
| 230 | __Bacteria_IMGSLHMTIQTAVLIETLVALGAEVR_3.3.1.1 | 1.00 | 1008313.22 | 1992412.45 | 1008313.22 | 1992412.45 | 1.98 | 19.94 | 20.93 | 0.98 | 0.10 | 0.45 | 0.45 |
| 231 | __Bacteria_LAGGIIER_1.1.1.85 | 1.00 | 1.00 | 1.00 | 1.00 | 1.00 | 1.00 | 0.00 | 0.00 | 0.00 | 1.00 | 1.00 | 1.00 |
| 232 | Actinobacteria__Actinobacteria_LGDALPVLAK_2.1.1.14 | 1.00 | 1.00 | 1.00 | 1.00 | 1.00 | 1.00 | 0.00 | 0.00 | 0.00 | 1.00 | 1.00 | 1.00 |
| 233 | Proteobacteria_Bartonellaceae_Bartonella_LLNPEFSHK_2.3.1.- | 1.00 | 1131960.31 | 1274604.53 | 1131960.31 | 1274604.53 | 1.13 | 20.11 | 20.28 | 0.17 | 0.07 | 0.88 | 0.88 |
| 234 | __Bacteria_LPEEVLQK_2.7.1.15 | 1.00 | 1.00 | 1.00 | 1.00 | 1.00 | 1.00 | 0.00 | 0.00 | 0.00 | 1.00 | 1.00 | 1.00 |
| 236 | __Bacteria_MEEVEVAEVK_3.6.4.- | 1.00 | 7689775.90 | 16333807.62 | 7689775.90 | 16333807.62 | 2.12 | 22.87 | 23.96 | 1.09 | 0.18 | 0.41 | 0.41 |
| 238 | __Bacteria_VQIVGDDLTVTNPK_4.2.1.11 | 1.00 | 3834340.80 | 3050361.56 | 3834340.80 | 3050361.56 | 0.80 | 21.87 | 21.54 | -0.33 | 0.09 | 0.77 | 0.77 |
| 239 | Proteobacteria_Anaplasmataceae_Anaplasma_VSIAVAVVMISK_6.1.1.20 | 1.00 | 1.00 | 1.00 | 1.00 | 1.00 | 1.00 | 0.00 | 0.00 | 0.00 | 1.00 | 1.00 | 1.00 |
| 240 | __Bacteria_WIADLLR_3.1.1.- | 1.00 | 1.00 | 1.00 | 1.00 | 1.00 | 1.00 | 0.00 | 0.00 | 0.00 | 1.00 | 1.00 | 1.00 |
| 241 | Proteobacteria_Erwiniaceae_Buchnera aphidicola_YGLLIIIK_ | 1.00 | 2621874.54 | 2595628.48 | 2621874.54 | 2595628.48 | 0.99 | 21.32 | 21.31 | -0.01 | 0.13 | 0.99 | 0.99 |
| 242 | Actinobacteria_Streptomycetaceae_Streptomyces_ADVMGIDEAK_6.1.1.7 | 1.00 | 1.00 | 1675885.81 | 1.00 | 1675885.81 | 1675885.81 | 0.00 | 20.68 | 20.68 | 1.00 | 0.06 | 0.06 |
| 243 | Thermotogae_Thermotogaceae_Thermotoga_AFIQDIILLK_2.7.2.8 | 1.00 | 1.00 | 1077842.30 | 1.00 | 1077842.30 | 1077842.30 | 0.00 | 20.04 | 20.04 | 1.00 | 0.03 | 0.03 |
| 244 | Actinobacteria__Actinobacteria_AGVGTGGMVTK_2.7.2.11 | 1.00 | 1.00 | 1.00 | 1.00 | 1.00 | 1.00 | 0.00 | 0.00 | 0.00 | 1.00 | 1.00 | 1.00 |
| 245 | Chlorobi_Chlorobiaceae_Chlorobium phaeobacteroides_AISIAGG | 1.00 | 1.00 | 1.00 | 1.00 | 1.00 | 1.00 | 0.00 | 0.00 | 0.00 | 1.00 | 1.00 | 1.00 |

| | | | | | | | | | | | | |
|---|---|---|---|---|---|---|---|---|---|---|---|---|
| | VSANSALR_2.3.1.234 | | | | | | | | | | | |
| 246 | __Bacteria_ALAPQIAR_2.1.1.198 | 1.00 | 1.00 | 1.00 | 1.00 | 1.00 | 1.00 | 0.00 | 0.00 | 0.00 | 1.00 | 1.00 | 1.00 |
| 247 | __Bacteria_ALDIIISLR_ | 1.00 | 1.00 | 1.00 | 1.00 | 1.00 | 1.00 | 0.00 | 0.00 | 0.00 | 1.00 | 1.00 | 1.00 |
| 248 | __Bacteria_AQATEDLR_1.14.14.- | 1.00 | 1.00 | 1.00 | 1.00 | 1.00 | 1.00 | 0.00 | 0.00 | 0.00 | 1.00 | 1.00 | 1.00 |
| 249 | Proteobacteria_Helicobacteraceae_Wolinella succinogenes_AVEAIAYAK_2.3.3.13 | 1.00 | 1.00 | 16327776.87 | 1.00 | 16327776.87 | 16327776.87 | 0.00 | 23.96 | 23.96 | 1.00 | 0.03 | 0.03 |
| 250 | Actinobacteria_Mycobacteriaceae_Mycobacterium tuberculosis complex_AVEGVSAVSELAGR_2.3.1.20 | 1.00 | 1.00 | 1.00 | 1.00 | 1.00 | 1.00 | 0.00 | 0.00 | 0.00 | 1.00 | 1.00 | 1.00 |
| 251 | Tenericutes_Mycoplasmataceae_Mycoplasma genitalium_AVLSGDIK_2.7.7.7 | 1.00 | 1.00 | 1.00 | 1.00 | 1.00 | 1.00 | 0.00 | 0.00 | 0.00 | 1.00 | 1.00 | 1.00 |
| 252 | Actinobacteria__Corynebacteriales_DGIEAVVAR_ | 1.00 | 1.00 | 2842861.36 | 1.00 | 2842861.36 | 2842861.36 | 0.00 | 21.44 | 21.44 | 1.00 | 0.28 | 0.28 |
| 253 | Firmicutes__Firmicutes_DGLENILR_2.7.7.85 | 1.00 | 1.00 | 1.00 | 1.00 | 1.00 | 1.00 | 0.00 | 0.00 | 0.00 | 1.00 | 1.00 | 1.00 |
| 254 | __Bacteria_DGLSELAK_5.1.1.15 | 1.00 | 1.00 | 20819417.35 | 1.00 | 20819417.35 | 20819417.35 | 0.00 | 24.31 | 24.31 | 1.00 | 0.18 | 0.18 |
| 255 | Proteobacteria_Rhizobiaceae_Agrobacterium tumefaciens_DIDLDGYVLPK_1.14.-.- | 1.00 | 1.00 | 34875733.56 | 1.00 | 34875733.56 | 34875733.56 | 0.00 | 25.06 | 25.06 | 1.00 | 0.00 | 0.00 |
| 256 | Tenericutes_Mycoplasmataceae_Mycoplasma gallisepticum_DLSDEELALIR_ | 1.00 | 1.00 | 1.00 | 1.00 | 1.00 | 1.00 | 0.00 | 0.00 | 0.00 | 1.00 | 1.00 | 1.00 |
| 257 | Firmicutes__Bacillales_DLVVQDVLK_2.7.4.9 | 1.00 | 1.00 | 10986205.29 | 1.00 | 10986205.29 | 10986205.29 | 0.00 | 23.39 | 23.39 | 1.00 | 0.04 | 0.04 |
| 258 | Actinobacteria_Nocardiaceae_Rhodococcus jostii_DNGIGMSHDEVVDLIGTLAK_ | 1.00 | 1.00 | 1.00 | 1.00 | 1.00 | 1.00 | 0.00 | 0.00 | 0.00 | 1.00 | 1.00 | 1.00 |
| 259 | Proteobacteria_Vibrionaceae_Vibrio_DVLSQNGK_4.1.1.3 | 1.00 | 1.00 | 485720929.25 | 1.00 | 485720929.25 | 485720929.25 | 0.00 | 28.86 | 28.86 | 1.00 | 0.11 | 0.11 |
| 260 | __Bacteria_EALQDEFK_2.1.2.1 | 1.00 | 1.00 | 1.00 | 1.00 | 1.00 | 1.00 | 0.00 | 0.00 | 0.00 | 1.00 | 1.00 | 1.00 |
| 261 | Tenericutes_Mycoplasmataceae_Mycoplasma penetrans_EENNGGNQ_ | 1.00 | 1.00 | 46266464.39 | 1.00 | 46266464.39 | 46266464.39 | 0.00 | 25.46 | 25.46 | 1.00 | 0.07 | 0.07 |
| 262 | Spirochaetes_Leptospiraceae_Leptospira_EESLLISR_6.3.2.1 | 1.00 | 1.00 | 1.00 | 1.00 | 1.00 | 1.00 | 0.00 | 0.00 | 0.00 | 1.00 | 1.00 | 1.00 |
| 263 | Firmicutes_Ruminococcaceae_Ruminiclostridium thermocellum_EGDVIIATIR_ | 1.00 | 1.00 | 47257345.65 | 1.00 | 47257345.65 | 47257345.65 | 0.00 | 25.49 | 25.49 | 1.00 | 0.11 | 0.11 |
| 264 | Firmicutes__Bacilli_EQLPDIGEMITTANGPAK_ | 1.00 | 1.00 | 1.00 | 1.00 | 1.00 | 1.00 | 0.00 | 0.00 | 0.00 | 1.00 | 1.00 | 1.00 |
| 265 | Firmicutes_Streptococcaceae_Lactococcus lactis_ETDLENIPDLLVK_1.3.1.98 | 1.00 | 1.00 | 1.00 | 1.00 | 1.00 | 1.00 | 0.00 | 0.00 | 0.00 | 1.00 | 1.00 | 1.00 |
| 266 | __Bacteria_FEIEMLK_3.1.21.2 | 1.00 | 1.00 | 1199444.6 | 1.00 | 1199444.63 | 1199444.6 | 0.00 | 20.19 | 20.19 | 1.00 | 0.23 | 0.23 |

| | | | | | | | | | | | | | |
|---|---|---|---|---|---|---|---|---|---|---|---|---|---|
| | | | | 3 | | | 3 | | | | | | |
| 267 | Actinobacteria_Frankiaceae_Frankia_FEVQTPTR_ | 1.00 | 1.00 | 6738499.75 | 1.00 | 6738499.75 | 6738499.75 | 0.00 | 22.68 | 22.68 | 1.00 | 0.03 | 0.03 |
| 268 | Chlamydiae_Parachlamydiaceae_Candidatus Protochlamydia amoebophila_FGLTIIIITHQLETVK_3.6.3.- | 1.00 | 1.00 | 1.00 | 1.00 | 1.00 | 1.00 | 0.00 | 0.00 | 0.00 | 1.00 | 1.00 | 1.00 |
| 269 | Proteobacteria_Neisseriaceae_Neisseria_FVAFLAGK_ | 1.00 | 1.00 | 1.00 | 1.00 | 1.00 | 1.00 | 0.00 | 0.00 | 0.00 | 1.00 | 1.00 | 1.00 |
| 270 | Proteobacteria__Betaproteobacteria_GGILGINIGK_1.3.5.2 | 1.00 | 1.00 | 1.00 | 1.00 | 1.00 | 1.00 | 0.00 | 0.00 | 0.00 | 1.00 | 1.00 | 1.00 |
| 271 | __Bacteria_GIVLPVGGIK_3.4.21.53 | 1.00 | 1.00 | 1638006.78 | 1.00 | 1638006.78 | 1638006.78 | 0.00 | 20.64 | 20.64 | 1.00 | 0.02 | 0.02 |
| 272 | Actinobacteria_Mycobacteriaceae_Mycobacterium_GLIFVIQVYR_ | 1.00 | 1.00 | 4939876.36 | 1.00 | 4939876.36 | 4939876.36 | 0.00 | 22.24 | 22.24 | 1.00 | 0.04 | 0.04 |
| 273 | __Bacteria_HEVNIAIVGK_6.3.4.2 | 1.00 | 1.00 | 1.00 | 1.00 | 1.00 | 1.00 | 0.00 | 0.00 | 0.00 | 1.00 | 1.00 | 1.00 |
| 274 | __Bacteria_IAASLILK_3.1.-.- | 1.00 | 1.00 | 1.00 | 1.00 | 1.00 | 1.00 | 0.00 | 0.00 | 0.00 | 1.00 | 1.00 | 1.00 |
| 275 | Tenericutes_Mycoplasmataceae_Mycoplasma pulmonis_IAASMALTK_ | 1.00 | 1.00 | 1.00 | 1.00 | 1.00 | 1.00 | 0.00 | 0.00 | 0.00 | 1.00 | 1.00 | 1.00 |
| 276 | __Bacteria_IDIDIER_ | 1.00 | 1.00 | 1.00 | 1.00 | 1.00 | 1.00 | 0.00 | 0.00 | 0.00 | 1.00 | 1.00 | 1.00 |
| 277 | __Bacteria_IDLALVK_5.4.99.62 | 1.00 | 1.00 | 1.00 | 1.00 | 1.00 | 1.00 | 0.00 | 0.00 | 0.00 | 1.00 | 1.00 | 1.00 |
| 278 | Aquificae_Aquificaceae_Aquifex aeolicus_IENYPGVPGPLSGYELLK_1.8.1.9 | 1.00 | 1.00 | 2121589.00 | 1.00 | 2121589.00 | 2121589.00 | 0.00 | 21.02 | 21.02 | 1.00 | 0.01 | 0.01 |
| 279 | Firmicutes_Clostridiaceae_Alkaliphilus metalliredigens_IFIDNFLR_ | 1.00 | 1.00 | 28515879.53 | 1.00 | 28515879.53 | 28515879.53 | 0.00 | 24.77 | 24.77 | 1.00 | 0.09 | 0.09 |
| 280 | __Bacteria_IGDAVIIK_6.5.1.2 | 1.00 | 1.00 | 4735759.28 | 1.00 | 4735759.28 | 4735759.28 | 0.00 | 22.18 | 22.18 | 1.00 | 0.05 | 0.05 |
| 281 | __Bacteria_IGIFGGAGVGK_3.6.3.14 | 1.00 | 1.00 | 60768236.73 | 1.00 | 60768236.73 | 60768236.73 | 0.00 | 25.86 | 25.86 | 1.00 | 0.19 | 0.19 |
| 282 | Firmicutes__Firmicutes_IGLLIELK_3.1.4.46 | 1.00 | 1.00 | 1.00 | 1.00 | 1.00 | 1.00 | 0.00 | 0.00 | 0.00 | 1.00 | 1.00 | 1.00 |
| 283 | Actinobacteria_Frankiaceae_Frankia casuarinae_IIDGLLAAGR_2.5.1.75 | 1.00 | 1.00 | 4479080.93 | 1.00 | 4479080.93 | 4479080.93 | 0.00 | 22.09 | 22.09 | 1.00 | 0.15 | 0.15 |
| 284 | Firmicutes_Staphylococcaceae_Staphylococcus_IILLDEVK_ | 1.00 | 1.00 | 1.00 | 1.00 | 1.00 | 1.00 | 0.00 | 0.00 | 0.00 | 1.00 | 1.00 | 1.00 |
| 285 | __Bacteria_IINEPTAAALAYGIDR_ | 1.00 | 1.00 | 2141780.82 | 1.00 | 2141780.82 | 2141780.82 | 0.00 | 21.03 | 21.03 | 1.00 | 0.11 | 0.11 |
| 286 | Proteobacteria__Proteobacteria_ILLDELVK_3.1.3.25 | 1.00 | 1.00 | 1.00 | 1.00 | 1.00 | 1.00 | 0.00 | 0.00 | 0.00 | 1.00 | 1.00 | 1.00 |
| 287 | Firmicutes_Clostridiaceae_Clostridium_IVNVIINIK_ | 1.00 | 1.00 | 1.00 | 1.00 | 1.00 | 1.00 | 0.00 | 0.00 | 0.00 | 1.00 | 1.00 | 1.00 |
| 288 | Firmicutes_Staphylococcaceae_Staphylococcus haemolyticus_LAEVLEGAQRPGHFDGVVTVLNK_6.3.2.1 | 1.00 | 1.00 | 2119542.89 | 1.00 | 2119542.89 | 2119542.89 | 0.00 | 21.02 | 21.02 | 1.00 | 0.05 | 0.05 |

| | | | | | | | | | | | | |
|---|---|---|---|---|---|---|---|---|---|---|---|---|
| 289 | Actinobacteria_Frankiaceae_Frankia casuarinae_LLGLTEAPRPADAADALALALCHLWR_3.1.22.4 | 1.00 | 1.00 | 1.00 | 1.00 | 1.00 | 1.00 | 0.00 | 0.00 | 0.00 | 1.00 | 1.00 | 1.00 |
| 290 | Spirochaetes_Borreliaceae_Borreliaceae_LNIPTGIPLVYELDK_5.4.2.11 | 1.00 | 1.00 | 2631201.52 | 1.00 | 2631201.52 | 2631201.52 | 0.00 | 21.33 | 21.33 | 1.00 | 0.20 | 0.20 |
| 291 | __Bacteria_LSNEGFMK_6.1.1.9 | 1.00 | 1.00 | 1.00 | 1.00 | 1.00 | 1.00 | 0.00 | 0.00 | 0.00 | 1.00 | 1.00 | 1.00 |
| 292 | Firmicutes_Peptoniphilaceae_Finegoldia magna_LVLDDEIPR_3.4.21.53 | 1.00 | 1.00 | 1419378.51 | 1.00 | 1419378.51 | 1419378.51 | 0.00 | 20.44 | 20.44 | 1.00 | 0.04 | 0.04 |
| 293 | Firmicutes__Bacillales_MDSDEVFK_2.6.1.9 | 1.00 | 1.00 | 1.00 | 1.00 | 1.00 | 1.00 | 0.00 | 0.00 | 0.00 | 1.00 | 1.00 | 1.00 |
| 294 | Tenericutes_Mycoplasmataceae_Mycoplasma gallisepticum_MQKPNIIIK_2.7.4.22 | 1.00 | 1.00 | 12094808.86 | 1.00 | 12094808.86 | 12094808.86 | 0.00 | 23.53 | 23.53 | 1.00 | 0.07 | 0.07 |
| 295 | Actinobacteria__Actinobacteria_MVTLAVLR_2.7.4.25 | 1.00 | 1.00 | 582613.81 | 1.00 | 582613.81 | 582613.81 | 0.00 | 19.15 | 19.15 | 1.00 | 0.03 | 0.03 |
| 296 | Tenericutes_Mycoplasmataceae_Mycoplasma pulmonis_NDMSIEEMESLIEK_ | 1.00 | 1.00 | 2200744.21 | 1.00 | 2200744.21 | 2200744.21 | 0.00 | 21.07 | 21.07 | 1.00 | 0.05 | 0.05 |
| 297 | Actinobacteria_Streptomycetaceae_Streptomyces_NFVAEALR_ | 1.00 | 1.00 | 14535017.11 | 1.00 | 14535017.11 | 14535017.11 | 0.00 | 23.79 | 23.79 | 1.00 | 0.11 | 0.11 |
| 298 | Proteobacteria_Cellvibrionaceae_Teredinibacter turnerae_NIIPAIIK_2.7.7.27 | 1.00 | 1.00 | 14208246.72 | 1.00 | 14208246.72 | 14208246.72 | 0.00 | 23.76 | 23.76 | 1.00 | 0.06 | 0.06 |
| 299 | Tenericutes_Mycoplasmataceae_Mycoplasma gallisepticum_NNDLNGLLR_3.6.4.12 | 1.00 | 1.00 | 1.00 | 1.00 | 1.00 | 1.00 | 0.00 | 0.00 | 0.00 | 1.00 | 1.00 | 1.00 |
| 300 | __Bacteria_NPDIVAGVAALK_4.1.1.36 | 1.00 | 1.00 | 1.00 | 1.00 | 1.00 | 1.00 | 0.00 | 0.00 | 0.00 | 1.00 | 1.00 | 1.00 |
| 301 | Actinobacteria_Nocardiaceae_Nocardia farcinica_QIGAAVGLR_3.4.21.88 | 1.00 | 1.00 | 1.00 | 1.00 | 1.00 | 1.00 | 0.00 | 0.00 | 0.00 | 1.00 | 1.00 | 1.00 |
| 302 | Proteobacteria_Vibrionaceae_Vibrio_QLIDIILR_2.7.13.3 | 1.00 | 1.00 | 546944.23 | 1.00 | 546944.23 | 546944.23 | 0.00 | 19.06 | 19.06 | 1.00 | 0.00 | 0.00 |
| 303 | Tenericutes_Mycoplasmataceae_Mycoplasma mobile_QTGSIGDIAGNK_ | 1.00 | 1.00 | 909033.52 | 1.00 | 909033.52 | 909033.52 | 0.00 | 19.79 | 19.79 | 1.00 | 0.00 | 0.00 |
| 304 | __Bacteria_QVFGIPGAK_2.2.1.6 | 1.00 | 1.00 | 1.00 | 1.00 | 1.00 | 1.00 | 0.00 | 0.00 | 0.00 | 1.00 | 1.00 | 1.00 |
| 305 | Firmicutes__Bacilli_QVGAEPLLK_3.6.3.- | 1.00 | 1.00 | 14134275.09 | 1.00 | 14134275.09 | 14134275.09 | 0.00 | 23.75 | 23.75 | 1.00 | 0.06 | 0.06 |
| 306 | __Bacteria_SANIALISYIDGEK_ | 1.00 | 1.00 | 1.00 | 1.00 | 1.00 | 1.00 | 0.00 | 0.00 | 0.00 | 1.00 | 1.00 | 1.00 |
| 307 | Tenericutes_Mycoplasmataceae_Mycoplasma pneumoniae_SANPNSTVVEDAGR_ | 1.00 | 1.00 | 1.00 | 1.00 | 1.00 | 1.00 | 0.00 | 0.00 | 0.00 | 1.00 | 1.00 | 1.00 |
| 308 | Firmicutes__Bacilli_SEQAYENMIPNGR_2.3.1.- | 1.00 | 1.00 | 4018791.83 | 1.00 | 4018791.83 | 4018791.83 | 0.00 | 21.94 | 21.94 | 1.00 | 0.02 | 0.02 |
| 309 | __Bacteria_TGADFYK_ | 1.00 | 1.00 | 52518202.27 | 1.00 | 52518202.27 | 52518202.27 | 0.00 | 25.65 | 25.65 | 1.00 | 0.19 | 0.19 |

| | | | | | | | | | | | | | |
|---|---|---|---|---|---|---|---|---|---|---|---|---|---|
| 310 | Actinobacteria_Mycobacteriaceae_Mycobacteriaceae_TIGRPTPPR_ | 1.00 | 1.00 | 16966432.93 | 1.00 | 16966432.93 | 16966432.93 | 0.00 | 24.02 | 24.02 | 1.00 | 0.00 | 0.00 |
| 311 | Fusobacteria_Leptotrichiaceae_Leptotrichia buccalis_TVEAEVHDSNLK_3.4.24.- | 1.00 | 1.00 | 1.00 | 1.00 | 1.00 | 1.00 | 0.00 | 0.00 | 0.00 | 1.00 | 1.00 | 1.00 |
| 312 | Actinobacteria_Mycobacteriaceae_Mycobacterium leprae_TVGEFPQVLK_6.1.1.19 | 1.00 | 1.00 | 215660.35 | 1.00 | 215660.35 | 215660.35 | 0.00 | 17.72 | 17.72 | 1.00 | 0.01 | 0.01 |
| 313 | Firmicutes__Bacilli_VAEGLGLPIK_1.11.1.6 | 1.00 | 1.00 | 1.00 | 1.00 | 1.00 | 1.00 | 0.00 | 0.00 | 0.00 | 1.00 | 1.00 | 1.00 |
| 314 | __Bacteria_VAGIAEVR_5.99.1.3 | 1.00 | 1.00 | 78461127.48 | 1.00 | 78461127.48 | 78461127.48 | 0.00 | 26.23 | 26.23 | 1.00 | 0.07 | 0.07 |
| 315 | Proteobacteria_Rickettsiaceae_spotted fever group_VIASGLEVIGVEPMDR_6.1.1.19 | 1.00 | 1.00 | 1.00 | 1.00 | 1.00 | 1.00 | 0.00 | 0.00 | 0.00 | 1.00 | 1.00 | 1.00 |
| 316 | Proteobacteria_Enterobacteriaceae_Enterobacteriaceae_VLGVITLK_3.6.3.12 | 1.00 | 1.00 | 14083782.66 | 1.00 | 14083782.66 | 14083782.66 | 0.00 | 23.75 | 23.75 | 1.00 | 0.09 | 0.09 |
| 317 | Proteobacteria__Proteobacteria_VLLEIADLK_3.6.3.- | 1.00 | 1.00 | 5908535.20 | 1.00 | 5908535.20 | 5908535.20 | 0.00 | 22.49 | 22.49 | 1.00 | 0.16 | 0.16 |
| 318 | __Bacteria_VLVFAVGER_ | 1.00 | 1.00 | 1897657.27 | 1.00 | 1897657.27 | 1897657.27 | 0.00 | 20.86 | 20.86 | 1.00 | 0.09 | 0.09 |
| 319 | Firmicutes__Clostridia_VVEVPVGEELLGR_3.6.3.14 | 1.00 | 1.00 | 61030542.67 | 1.00 | 61030542.67 | 61030542.67 | 0.00 | 25.86 | 25.86 | 1.00 | 0.07 | 0.07 |
| 320 | __Bacteria_VVTEIINLK_3.1.3.11 | 1.00 | 1.00 | 13432797.15 | 1.00 | 13432797.15 | 13432797.15 | 0.00 | 23.68 | 23.68 | 1.00 | 0.03 | 0.03 |
| 321 | Proteobacteria_Brucellaceae_Brucellaceae_WGGALLPK_2.7.1.23 | 1.00 | 1.00 | 1.00 | 1.00 | 1.00 | 1.00 | 0.00 | 0.00 | 0.00 | 1.00 | 1.00 | 1.00 |
| 322 | __Bacteria_YDLLLSDR_ | 1.00 | 1.00 | 940308.21 | 1.00 | 940308.21 | 940308.21 | 0.00 | 19.84 | 19.84 | 1.00 | 0.05 | 0.05 |
| 323 | __Bacteria_YLINLVK_2.7.4.25 | 1.00 | 1.00 | 613748.34 | 1.00 | 613748.34 | 613748.34 | 0.00 | 19.23 | 19.23 | 1.00 | 0.20 | 0.20 |
